# Neptune: a toolbox for spinal cord functional MRI data processing and quality assurance

**DOI:** 10.64898/2026.03.03.709443

**Authors:** D Rangaprakash, Robert L Barry

## Abstract

Over the past two decades, open-source research software such as SPM, AFNI and FSL formed the substrate for advancements in the brain functional magnetic resonance imaging (fMRI) field. The spinal cord fMRI field has matured substantially over the past decade, yet there is limited research software tailored for processing cord fMRI data that has distinct noise sources, unique challenges, niche processing requirements and special needs. Spinal cord fMRI data analysis is a ‘different beast’, involving specialized pre- and post-processing steps due to the cord’s unique anatomy and higher distortions/physiological noise, thus requiring extensive and careful quality assessment. Building upon 10+ years of research and development, we present Neptune – a user-interface-based MATLAB toolbox. With 30,000+ lines of in-house code, it is designed to be easy to use and does not require programming knowledge. Neptune builds on our previously published 15-step pre-processing pipeline (Barry et al., 2016) and presents a 19-step pipeline with new processing steps, and enhancements to existing steps. Neptune has a 4-step post-processing pipeline aimed at fMRI connectivity modeling. It generates extensive and novel quality control visuals to enable a thorough assessment of data quality, and displays them in an elegant webpage format. We demonstrate the utility of Neptune on our 7T data. Certain features of the popular Spinal Cord Toolbox (SCT) are integrated into Neptune, and users can import/export between Neptune and other software such as FSL and SPM. The availability of this open-source, easy-to-use software will benefit the spinal cord fMRI community, and also tip the cost-benefit balance for brain fMRI researchers to invest in learning new software to conduct important neuroscientific and clinical research using spinal cord fMRI.

## 1. Introduction

Functional magnetic resonance imaging (fMRI) has revolutionized our understanding of brain function over the past two decades. This quest has been facilitated by open-source research software for robustly and reliably processing brain fMRI data (e.g., SPM [1], AFNI [2], FSL [3]). The brain and spinal cord together comprise the central nervous system (CNS), and the cord is an integral and vital part of neural processing that is also affected in several neurological diseases. Despite advances in spinal cord fMRI [4], an open-source research software tailored for spinal cord fMRI data processing and quality assurance has been elusive. Here, we present a suite of tools, which we call “Neptune”, that combines specialized fMRI data processing steps with helpful quality assurance plots.

Spinal cord fMRI has revealed functional impairments in neurological diseases such as multiple sclerosis [5] and pain [6]. It is an emerging field that has made both fundamental and clinical advancements in the past decade (e.g., [7] [8] [9] [10] [11] [12] [13]). The cord differs from the brain in both anatomy and fMRI data quality. It is a curved, narrow, cylindrical structure surrounded by bone and moving organs (tongue, throat, heart, lungs). Cord fMRI data has considerably higher physiological noise and distortions arising from dynamic B_0_ field inhomogeneities [14]. Spinal cord fMRI data processing is thus highly challenging, requiring extensive data quality assessment at every stage of processing, with a unique blend of pre- and post-processing steps. Despite these requirements, most researchers have used brain fMRI tools to analyze cord data because there is a paucity of publicly available specialized processing software for cord fMRI.

Building upon a decade of software development, Neptune is a MATLAB- and AFNI-based [2] open-source research toolbox. Neptune was necessitated for two reasons: (1) the spinal cord fMRI processing pipeline is different from brain fMRI processing (especially at ultra-high fields), and (2) detailed checking of intermediate outputs is necessary in the form of informative visuals not provided by most brain-based software. The foundation for Neptune was laid in 2012 [8] to process the first 7T resting state (rs) spinal cord fMRI data by the senior author (R.L.B.). The pipeline was further refined as a 15-step procedure in 2016 [10]. The first author (R.D.) built Neptune on this foundation (starting during the COVID-19 pandemic lockdown in 2020) by both updating existing code and adding new processing steps, in addition to introducing a host of parameter choices, quality control (QC), and user interface functionalities. Neptune was first made available to the public at the first Martinos Spinal Cord Workshop in November 2021, and later presented as an oral talk at the 30th ISMRM Annual Meeting in May 2022 [15]. Neptune has undergone further development since then to add new features, fix minor bugs, and improve batch processing, which came about while utilizing Neptune to process newly acquired data. We wrote this article after completing these activities and determining that no further significant additions or modifications are planned. So far, six incremental versions of Neptune have been made public (Nov 2021, Dec 2021, April 2022, May 2022, Nov 2023, March 2026), with the latest release being referenced in this article. Neptune is available on MATLAB Central [16] and GitHub [17]; the current version has also been provided with this article. Sample data is freely and openly available on Mendeley Data [18] for learning and testing the software.

Neptune comprises over 30,000 lines of in-house MATLAB code. It is fully user-interface-based and does not require MATLAB/programming knowledge (command-line options are available). Neptune generates an extensive array of QC outputs and presents them to the user in an elegant interface, which is important for spinal cord fMRI studies. The Spinal Cord Toolbox (SCT) [19], primarily aimed at processing anatomical spinal cord images, is a popular open-source tool in the spinal cord field, and some functionalities of SCT have been integrated into Neptune. A user could choose to perform only part of the steps in Neptune while importing/exporting between other software such as SCT, FSL and SPM. The Neptune pipeline is tailored for processing rs-fMRI data, although it can also be used in part or whole for processing task fMRI data. Neptune is a new, open-source option for spinal cord fMRI researchers to reliably, robustly and efficiently process their data. Here, we present the toolbox and provide validation through standard functional connectivity (FC) analysis of our 7T rs-fMRI data.

## 2. Methods

We have included a supplementary document with this manuscript, which gives complete all-encompassing information about using Neptune from a practical standpoint, replete with 134 figures, screenshots and illustrations. It has four sections – (i) installation and file organization (5 figures), (ii) pre-processing in Neptune (42 figures), (iii) post-processing (20 figures), and (iv) understanding the outputs (67 figures). Here, in the main text, we focus on Neptune at the conceptual level, elaborate on its novelty and utility, and demonstrate it using 7T data. These aims are achieved using a limited set of figures suitable for the main text, and we direct users to the Supplement if further details are desired.

### 2.1. Fundamentals of Neptune

The foundational steps for processing spinal cord rs-fMRI data in our lab, which form the substrate of Neptune, were revised and published in 2016 [10]. We will thus not elaborate on them herein and instead explain how Neptune builds upon the published pipeline in the following six ways.

#### (1) Graphical user interface (GUI)

The source code from 2016 was written for a specific dataset as a single lengthy script that called certain functions and was supposed to be run section by section on the command line (to facilitate manual troubleshooting, code development, and QC). Neptune is a significant upgrade from that prototype. Neptune is user-interface-based and easy to use – the overarching idea was to have a hypothetical new lab member get comfortable using the software within their first week. This functionality has been enabled with 30,000+ lines of in-house code aimed at maximizing functionality, retaining rigor, and minimizing the difficulty of use. Neptune was built on the idea of mimicking a conversation between the user and the software, wherein the software asks the user to make certain GUI choices one after the other, which shape successive conversations. This architecture differs from other programs that use a cohesive settings interface with a host of checkboxes and buttons to elect. Both approaches work well and are merely different styles of programming. **Figure 1** shows Neptune’s interface, examples of choices, and selections made by the user through the GUI. Section 2 of the Supplement shows all GUI choices presented by Neptune to the user (over 30 in a typical execution).

**Figure 1.**
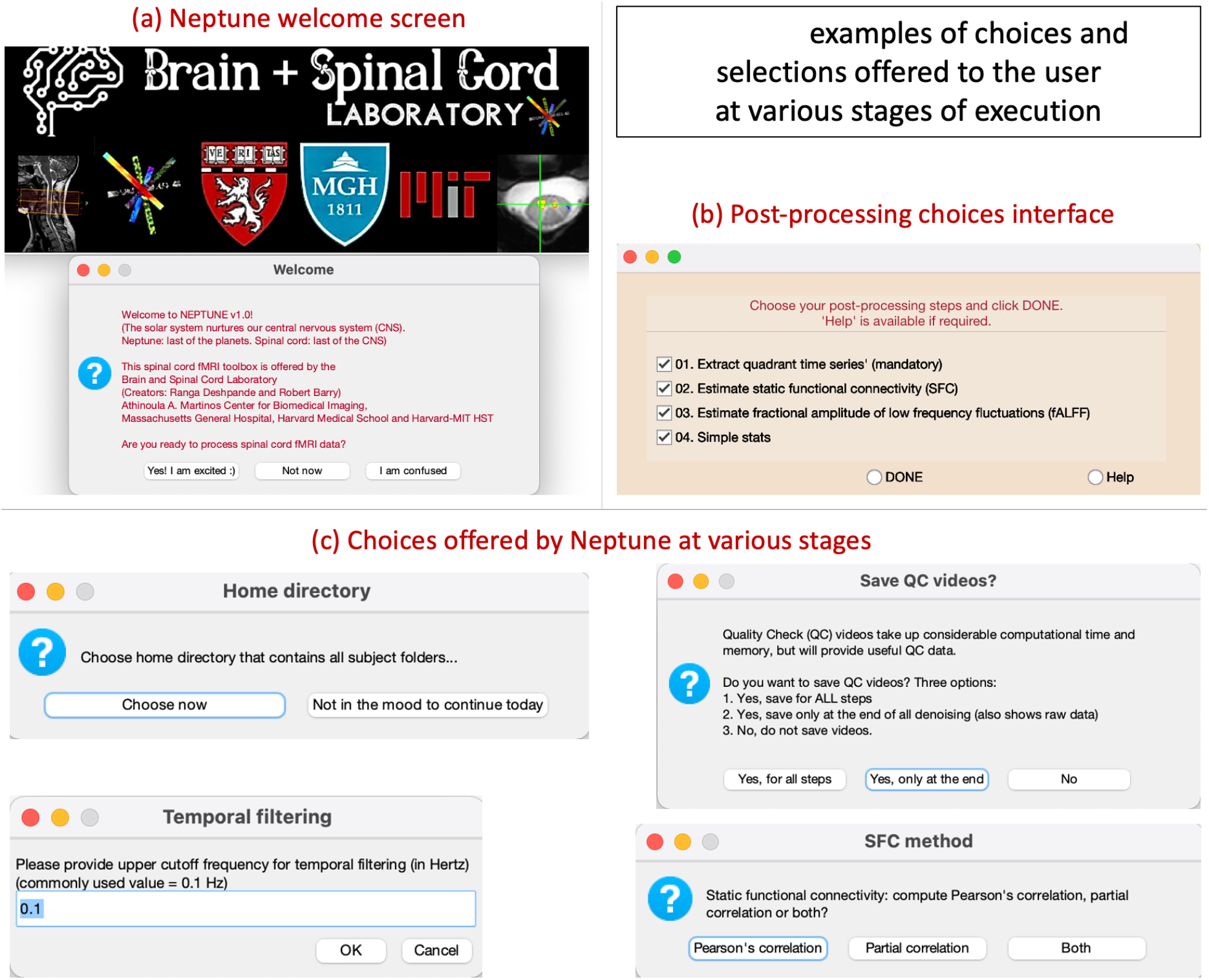
Examples of choices offered to the users at various stages of execution. **(a)** Welcome screen. **(b)** Post-processing choices. **(c)** As an example: choosing the home directory, upper cutoff frequency for temporal filtering, functional connectivity estimation method, and whether to save QC videos.

#### (2) New and improved pre-processing steps

Neptune has a 19-step pre-processing pipeline that includes new steps (compared to the 2016 code) as well as improvements to existing steps. The new steps include options for converting images from DICOM (or Philips PAR/REC) to NIfTI, NORDIC denoising [20], slice-timing correction, regressing out additional covariates (such as motion parameters and their derivatives, scrubbing variables and user-defined covariates), deconvolution, temporal signal-to-noise ratio (tSNR) computation across pre-processing steps and slices, co-registration to standard space, file management (file naming and organization), and finally file compression to reduce disk space usage. Improvements to existing steps include matching functional and anatomical images by reslicing, automated estimation of Gaussian cord mask for motion correction, semi-automated anatomical and mean functional image segmentation, and more efficient temporal filtering. We briefly describe some of these key innovations next. **Figure 2a** shows the list of these 19 steps.

**Figure 2.**
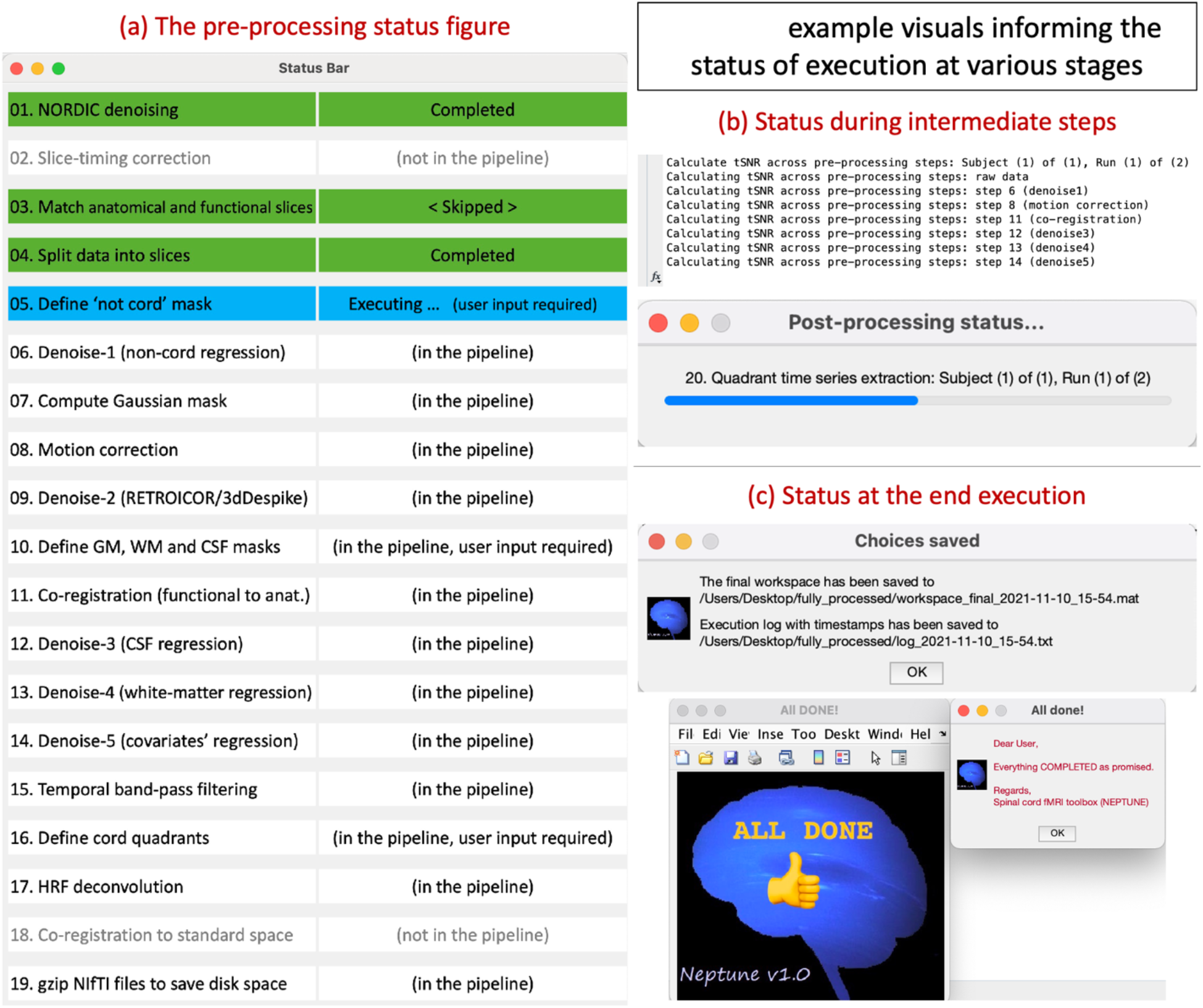
Visuals informing the user of the status of execution. **(a)** The pre-processing status figure that stays open during the entire course of pre-processing. **(b)** Examples of status in the MATLAB command line and in a status bar during intermediate steps. **(c)** Status at the end of execution showing that the workspace and execution log have been saved to specific files.

NORDIC is a recently published technique for minimizing thermal noise in fMRI data [20]. It has gained popularity in the brain fMRI field, and we recently showed that it significantly improves tSNR of highly accelerated acquisitions in the cord, enabling us to achieve sub-second and sub-millimeter fMRI of the cervical cord at 7 Tesla [21]. We thus incorporated NORDIC as step-1 of Neptune by tailoring the source code provided by the authors. It does not perform NORDIC on raw complex K-space data from each receive channel, but instead works directly on NIfTI data using procedures stipulated by the NORDIC authors.

Additional covariates can now be regressed out (for denoising) as part of step-14. Motion parameters and scrubbing variables can be selected, in addition to user-defined covariates input as text, CSV or Excel files. Step-17 provides the option for performing hemodynamic response function (HRF) deconvolution [22]. FMRI is an indirect measure of neural activity, and neurovascular coupling (i.e., the HRF) is the interface between the two. The HRF is variable across individuals and brain regions [23]. HRF variability is generally considered a confound in fMRI studies [24], and ignoring this variability results in erroneous FC estimation [25]. The HRF is altered even in pathological conditions, and ignoring it can result in faulty FC group differences [26] [27]. We recently demonstrated HRF variability in the spinal cord [28]. Hence, performing HRF deconvolution on rs-fMRI data and using the resulting latent neural time series for further processing is a wise choice under most circumstances. This motivated us to incorporate it into Neptune.

Another beneficial feature of Neptune is its integration with SCT. Users can now co-register their images to the standard PAM50 space in step-18. All processing is done within Neptune in MATLAB by calling SCT functions through the ‘*unix*’ command, and its inputs/outputs are integrated into the Neptune pipeline. SCT [19] was also used to improve segmentation steps. Previously [10], the not-cord mask (step-5) and GM/WM/CSF anatomical masks (step-10) were derived by manually drawing boundaries in a tedious and lengthy process. Neptune now estimates WM and CSF boundaries using SCT’s *propseg* function, and GM boundaries using *deepseg*. There is also the option to transform the PAM50 template to the subject’s anatomical space to define the GM boundary. The user is then presented with these boundaries in the user interface, with the freedom to move boundary points to best suit their visual assessment of the tissue boundaries (hence semi-automated). The not-cord mask is estimated from the mean functional image in a similar manner (step-5), which is then used to automatically derive the Gaussian mask (step-7) that is utilized in the motion correction step. (In the original prototype code, the Gaussian mask was manually defined by three points.)

File management was enhanced as well. Although Neptune does the processing slice-by-slice after splitting a 3D volume into 2D slices, the prototype code left the slice-wise data as is, leading to several dozen files per subject. Neptune removes slice-wise files after reconstructing the 3D volume, compresses the NIfTI images to reduce file sizes, and makes the subject folders tractable with only a handful of essential files. Neptune saves information about the amount of space saved and tracks execution time to display the time taken for each step and each subject. Neptune computes tSNR across all slices, as well as across each pre-processing step, which helps assess data quality and detect problematic scans.

#### (3) Status updates

Neptune has a host of continuous visual displays that inform the user about the status of execution (examples in **Figure 2**). Users will benefit from these details. This is not merely a visual aid; all the status updates from the beginning to the end of execution are pooled together and logged, and this informative log is saved. It records all the choices made, the starting and ending times of each step along with the time taken, steps that were completed successfully and those that were skipped, and so on. We tried not to omit any detail to ensure that every relevant piece of information was recorded. As such, this log serves as a permanent record of data pre-processing and analysis that could be reviewed years later, thus supporting transparency and reproducible research. Neptune’s status bar, command-line outputs and status dialog boxes keep the user informed during every execution step.

#### (4) A new post-processing pipeline

Neptune has a new 4-step post-processing pipeline (**Figure 1b**) for obtaining time series from cord quadrants, estimating Pearson’s or partial correlation-based FC within and between vertebral levels, estimating the fractional amplitude of low-frequency fluctuations (fALFF) [29] across the cord, and simple statistical analysis. We also present a useful way to compute and view between-slice FC. The comprehensive within- and between-slice spinal cord connectome is in a format comparable to the brain connectome, allowing for brain network modeling tools to potentially be applied to spinal cord data. This is a distinct way to view the spinal cord connectome because FC between quadrants of the same slice (within-slice FC) has been the preeminent approach in the field [9] [30] [31] [8]. Between-slice FC has been measured to be markedly lower than within-slice FC [9] [32] [33] [34]. Neptune also provides the option to compute fALFF. Finally, simple statistics can be performed in Neptune (without nuisance covariates, multivariate analysis, multiple comparisons correction, or non-parametric statistics). It includes one-way, two-way and paired t-tests. Neptune outputs can be easily fed into other open-source comprehensive statistical analysis packages for further sophisticated statistical analyses.

#### (5) Extensive quality control outputs

Neptune generates QC output visuals at every stage of processing (examples in **Figure 3**). Section 4 of the Supplement gives a detailed overview of all types of QC outputs, which educates the user about the nature of their own data and helps identify problematic runs. Thorough quality assessment is vital for processing spinal cord fMRI data because it is quite common to see noisy runs, useless slices, insufficient motion correction, failed co-registration, inefficient denoising, and a host of other issues. Spinal cord fMRI researchers routinely discard scans and deal with distorted images. Yet, no previous software provides users with QC visuals tailored for spinal cord fMRI data. Addressing this unmet need was one of the driving factors for investing considerable effort into Neptune’s development.

**Figure 3.**
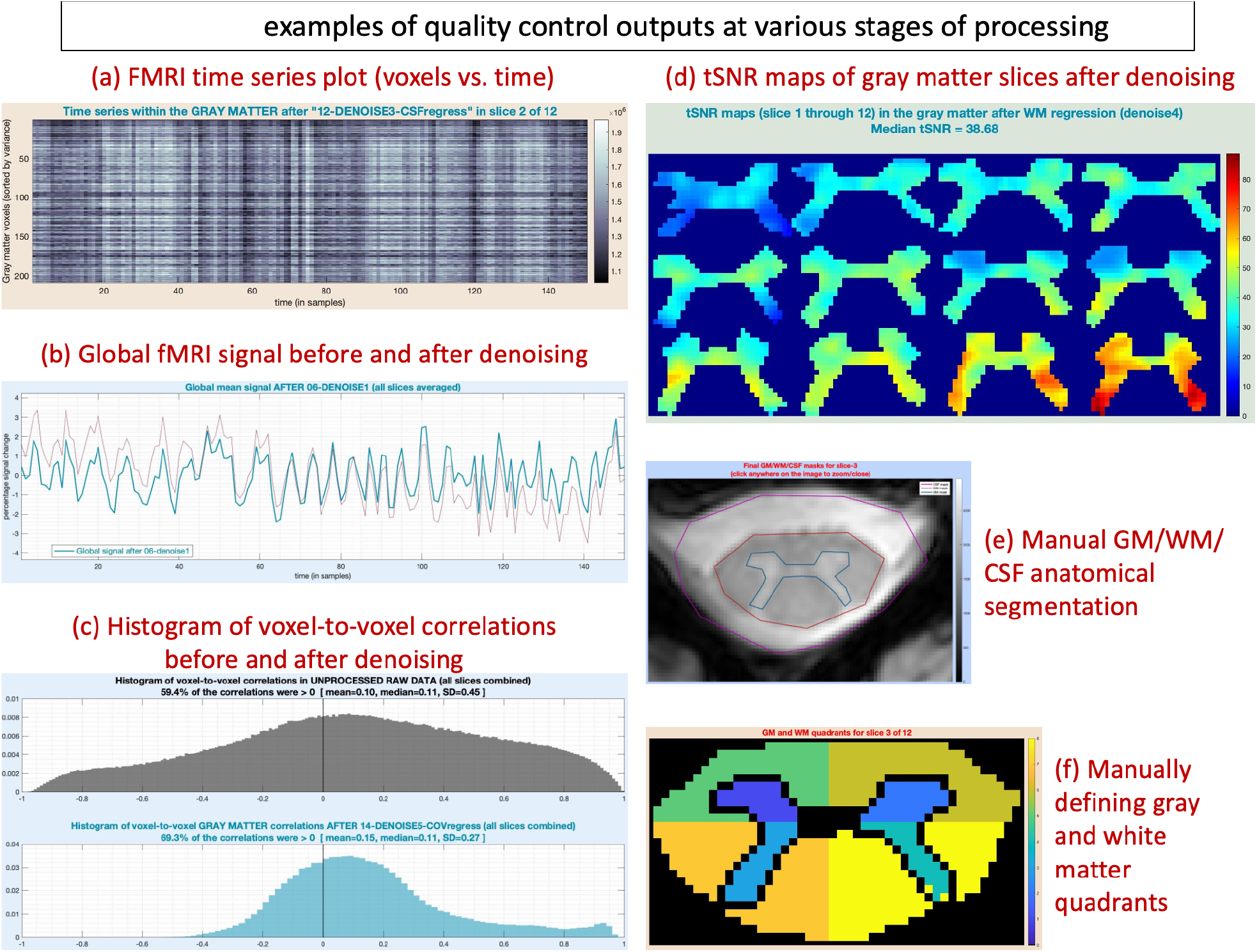
Examples of quality control (QC) outputs. **(a)** A plot of all within-slice gray matter time series, stacked one above the other (to spot systematic noise). Time is along the x-axis, and voxels are along the y-axis. **(b)** Global mean signal before (red) and after (blue) denoising. **(c)** Histogram of voxel-to-voxel correlations before (above, gray) and after (below, blue) denoising, showing a healthy narrowing of the distribution after denoising. **(d)** Gray matter temporal signal-to-noise ratio (tSNR) maps after denoising (most inferior slice in the top left and most superior slice in the bottom right corner). **(e)** Semi-automated segmentation of the anatomical image (shown here is slice-3). **(f)** Defining cord quadrants (left/right dorsal/ventral horns) in the gray and white matter (of slice-3) for post-processing.

#### (6) Viewing results in a novel webpage format

Lastly, Neptune generates a webpage with a novel design at the end of processing (**Figure 4**). The user can comfortably browse through all QC outputs in one place. Neptune generates QC visuals for each slice and additionally aggregates across all slices – both are presented on this webpage.

**Figure 4.**
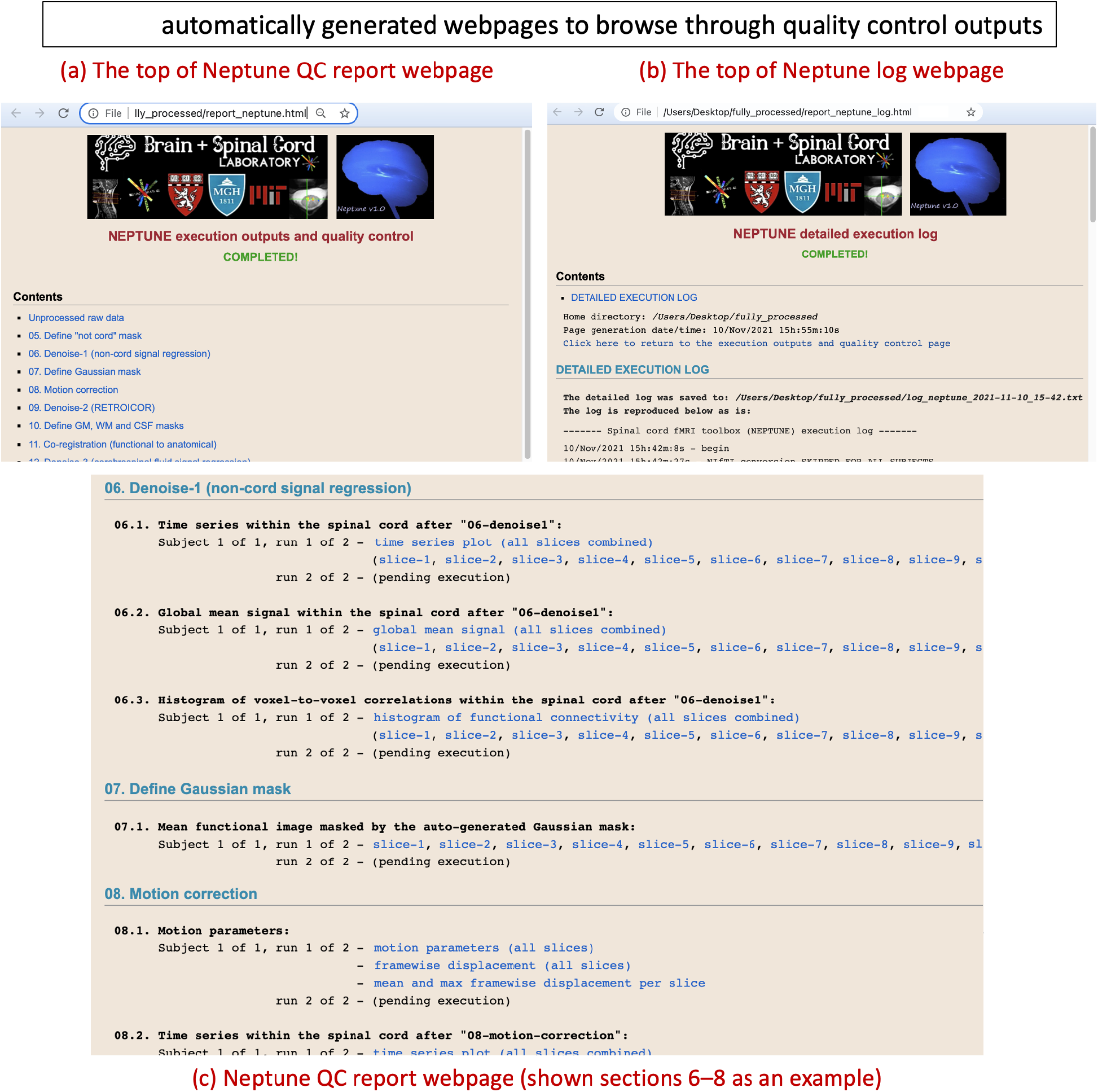
Neptune automatically generates an elegant webpage, which the user can browse to view the generated quality control outputs. **(a)** and **(c)** show the top of the webpage and one of the middle sections, respectively. Neptune also generates a webpage of the detailed execution log that contains (almost) all the information the user may need regarding the execution – the top of this page is shown in **(b)**.

### 2.2. Demonstrating the software using 7T data and an established processing pipeline

The core principles of Neptune have remained consistent over the years of its development: robust motion correction, reliable co-registration of functional images to the anatomical space, and effective denoising. Neptune incorporates the same principles, and the methods used to perform these steps have been the same since 2014. Neptune uses well-validated AFNI functions [2] to perform these tasks. Dozens of 3T and 7T spinal cord fMRI scans have been processed using the same pipeline, and the results have been published [8] [10] [35] [5]. The new functionalities adopted by Neptune have also been well-validated in literature, including NORDIC denoising [20] [36] [37], HRF deconvolution [38] [22] [25] [39] [40], and SCT [19].

Therefore, herein we used existing data with an established processing pipeline to demonstrate the utility of Neptune and test whether we obtain the expected connectivity output. Specifically, we utilized the 7T data that was previously analyzed to characterize the reproducibility of spinal cord networks [10]. Cervical cord rs-fMRI data were acquired in a Philips Achieva 7T scanner (N=23, healthy, 11M/12F, age=26±4.5y). All participants provided informed consent; data acquisition was approved by the Vanderbilt University Institutional Review Board (IRB), and data analysis was approved by the Mass General Brigham IRB. Two sequential rs-fMRI runs were performed using a 3D multi-shot gradient-echo sequence. Acquisition parameters were as follows: volume acquisition time (VAT) = 3.34 s, repetition time (TR) = 17 ms, echo time (TE) = 8.0 ms, flip angle (FA) = 15°, in-plane voxel size = 0.91×0.91 mm^2^, slice thickness = 4 mm, # slices = 12, covering C2–C5 and centered on the C3/C4 junction, field of view (FOV) = 160×160 mm^2^, acceleration (SENSE) = 1.56, phase encoding direction = anterior-posterior (A-P), # volumes = 150, scan duration = 8 min 21 s (excluding 10 initial volumes approaching steady-state magnetization). Physiological data (cardiac/respiratory signals) were acquired using a pulse oximeter and respiratory belt. T2* weighted anatomical images with identical placement were acquired as follows: TR = 303 ms, TE = 8.2 ms, FA = 25°, in-plane voxel size = 0.6×0.6 mm^2^, slice thickness = 4 mm, # slices = 12, FOV = 160×160 mm^2^, SENSE = 2.0, phase encoding direction = A-P, scan duration = 5 min 22 s.

Data were pre-processed in Neptune (steps 3–16) with the standardized pipeline used in our previous publications. Additional denoising covariates for step-14 were motion parameters and their derivatives, and scrubbing variables with aggregate motion greater than 2 mm. A 0.01–0.1 Hz passband was used for bandpass filtering. HRF deconvolution was not chosen due to poor temporal resolution (the rule-of-thumb is that deconvolution requires a VAT of at most 2 sec). Each fMRI slice was divided into four gray matter quadrants as in previous studies [8] [10]: left(L)/right(R) ventral(V)/dorsal(D) horns, and within-slice FC was computed between all pairs of quadrants [35]. For each connection, a one-sample t-test was performed by pooling across all slices, runs and participants (p<0.05, Bonferroni corrected). T-statistics are reported with the expectation that we will replicate the most robust finding in spinal cord rs-fMRI [8] [9] [10] [30], which are higher LV–RV and LD–RD FC (with ventral connectivity being stronger than dorsal connectivity), and lower FC with rest of the connections (LV–LD, RV–RD, LV–RD and LD–RV).

## 3. Results

We examined Neptune’s QC outputs (examples in Figure 3) and verified acceptable fMRI data quality across participants. A complete array of QC outputs from one participant is presented in Section 4 of the Supplement (over 60 figures) to demonstrate the data insights that may be obtained from Neptune. We confirmed robust motion correction and successful co-registration in all subjects. There were no adverse observations during the denoising steps. The de-identified data used here have been made public on Harvard Dataverse, which also contains fully processed data, FC outputs and QC plots for all 23 subjects [41]. Users can download the data and review QC outputs in all subjects to evaluate the utility of automatically generating data quality and outcome measures.

Upon fully processing all 23 subjects and performing statistical analysis, we found the expected pattern of higher T-statistics for LV–RV (*T*=43.97) and LD–RD (*T*=28.75) connections and lower for the rest of the connections (T-value between 13.13 and 18.32) (**Figure 5**). Ventral connectivity was also higher than dorsal connectivity, as predicted.

**Figure 5.**
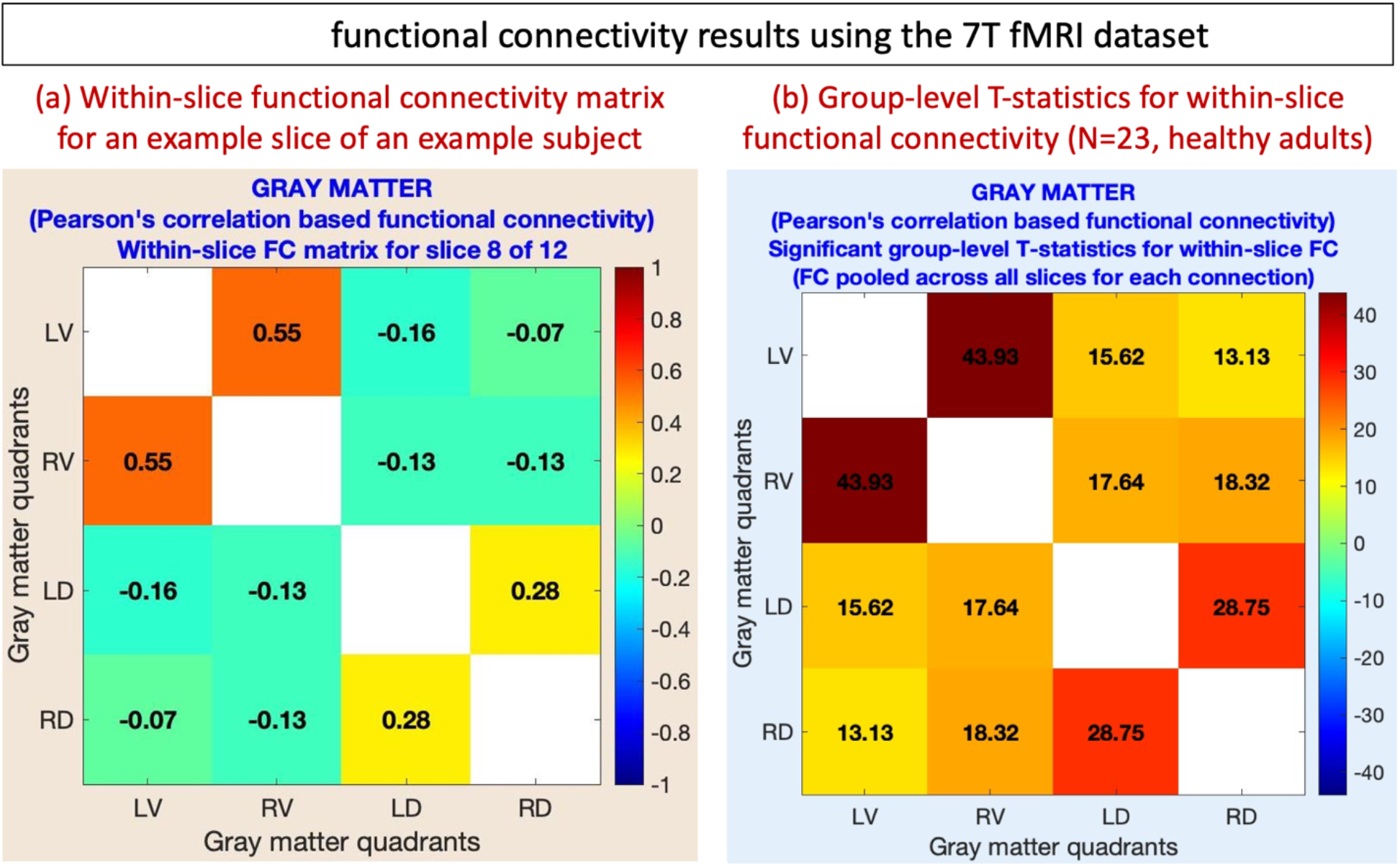
Within-slice functional connectivity (FC) results obtained after running Neptune on our 7T fMRI data (N=23, healthy adults). **(a)** FC of an example slice of a random subject showing high LV-RV and LD-RD FC, but lower FC in the other connections. **(b)** The final result showing the T-statistics, again with high LV-RV and LD-RD FC, replicating robust findings from prior rs-fMRI studies in the cord. L=left, R=right, V=ventral horn, D=dorsal horn.

## 4. Discussion

Convenient and accessible software tools with a quick learning curve significantly boost research in a field, as evident from the breakthroughs facilitated by SPM, AFNI and FSL in the fMRI field. In comparison to the brain, the spinal cord fMRI field is still small, yet it has immense clinical and neuroscientific potential. Tools tailored for the processing and analysis of spinal cord fMRI data to advance the field further have been lacking. Built upon 10+ years of research and development, we presented Neptune: a user-friendly and technically sound specialized toolbox for reliable and informed spinal cord fMRI data analysis and quality assurance.

Notably, a user can run some processing steps outside Neptune (using custom software or an established fMRI toolbox) and then import the semi-processed data into Neptune by keeping the naming convention consistent. Each successive step of Neptune expects input filenames to conform to a predetermined structure. Section 4 of the Supplement explains Neptune’s output file structure; by learning it, a user will be able to run steps outside Neptune, rename the files accordingly (possibly with an automated script), and then run only part of Neptune’s steps on the resultant files. As an example, it is currently common to use the combination of SCT and FSL for task fMRI data processing. Users could run specific anatomical processing steps in SCT, import them to Neptune to run motion correction and denoising, and then use the outputs in FSL to perform activation analyses. These are some of the ways in which users can tailor Neptune to their needs.

In this article, we articulated the need for Neptune and the advancements that it brings to our previously published pipeline. We elaborated on the novel GUI features, status figures, QC outputs and webpage, and explained Neptune’s new and improved processing steps. We demonstrated the utility of Neptune on our previously published 7T data using a standard pipeline. We showed that one can effectively perform quality control and obtain expected patterns of within-slice FC in the cervical cord. In summary, it is our hope that Neptune will play a role in advancing the field of spinal cord fMRI in the following decades by simultaneously simplifying data processing for spinal cord fMRI researchers and serving as a stepping-stone for brain fMRI researchers to study function throughout the central nervous system.

## Supporting information

Neptune user documentation

Neptune software

## Data and Code Availability

Neptune is available to download for free on MATLAB Central [16] and GitHub [17]. The current version (*neptune_v1.260301*) is available with the manuscript as a supplemental attachment. The de-identified data used in this article have been made public and are available to download for free from Harvard Dataverse (it also contains fully processed data, connectivity outputs and QC plots) [41]. The sample data used in the Supplement are available on Mendeley Data [18].

## CRediT (Contributor Roles Taxonomy) – Author Contributions

**D Rangaprakash**: Conceptualization, Methodology, Software, Data Analysis, Investigation, Visualization, Writing - Original Draft, Reviewing and Editing. **Robert L Barry**: Funding Acquisition, Resources, Conceptualization, Methodology, Software, Data Acquisition, Investigation, Writing - Reviewing and Editing, Supervision.

## Supplementary Material

Attached: Neptune user documentation

Attached: current version of the Neptune software (*neptune_v1.260301_setup.zip*)

