## Supplementary material for "Neptune: a toolbox for spinal cord functional MRI data processing and quality assurance": Neptune software: AFNI.txt

Please install AFNI before setting up Neptune:
https://afni.nimh.nih.gov/

AFNI must preferably be installed after installing MATLAB. After setting up, install the AFNI plugin for MATLAB:
https://afni.nimh.nih.gov/pub/dist/doc/OLD/afni_matlab.shtml

In the end, unzip the file 'AFNI_version_issue.zip' provided in this directory and follow the instructions in the readme file.
