## Supplementary material for "Neptune: a toolbox for spinal cord functional MRI data processing and quality assurance": Neptune software: MATLAB.txt

Neptune is a MATLAB-based toolbox. You can get MATLAB here:
https://www.mathworks.com/products/matlab.html

Neptune depends on the following inbuilt MATLAB toolboxes:
 Signal Processing Toolbox
 Image Processing Toolbox
 Statistics and Machine Learning Toolbox
 Parallel Computing Toolbox (optional - it speeds up step-17 HRF deconvolution - but can run without it)

Installing SPM for MATLAB is required only if step-2 (slice timing correction) is chosen:
https://www.fil.ion.ucl.ac.uk/spm/software/download/
