## Supplementary material for "Neptune: a toolbox for spinal cord functional MRI data processing and quality assurance": Neptune software: SCT.txt

Installing the Spinal Cord Toolbox (SCT) is optional if users do not plan to run step-18 (co-registration to PAM50 standard space; it is necessary if this step is chosen). Nevertheless, having it significantly speeds up manual segmentation of tissue boundaries during steps 5 and 10:
https://spinalcordtoolbox.com/
