## Supplementary material for "Neptune: a toolbox for spinal cord functional MRI data processing and quality assurance": Neptune software: how_to_install.txt

1. Neptune has certain dependencies that need to be installed first. This includes MATLAB, AFNI and NIfTI Toolbox. SCT is optional but will expedite manual segmentation steps. The directory '/dependencies' provides links for downloading and installing these software.
   In terms of hardware, Neptune has been tested on MacBook Pro and MacBook Air laptops (2019 models). However, we do not know the minimum processor requirements. The RAM requirement depends on your data size; with fMRI, it is always preferable to have comfortable RAM; Neptune was extensively tested on bulky data in a MacBook Pro with 64 GB RAM without any issues.

2. After completing the above step, the rest of the setup is a breeze. Unzip the downloaded Neptune folder (anywhere you want). In MATLAB, go to the unzipped Neptune folder.

3. Drag and drop the "setup_neptune.m" script to the command window (i.e., run the script). If you open the script in the editor, it tells what it does.

4. Check the command window for potential errors. If there are no errors, then Neptune is installed and ready to use.

5. Type "neptune" in the command window (without the quotes) and hit enter to run Neptune.

This is a work in progress. If you have trouble installing or using the software, or if you want to report bugs, please email Ranga Deshpande or Robert Barry.


Note: in the directory "older_versions", you can find previous public releases, which could help you compare different versions if needed.


For questions, comments, concerns and suggestions, please email:
