## Supplementary material for "Neptune: a toolbox for spinal cord functional MRI data processing and quality assurance": Neptune software: Neptune_user_manual.pdf

---

This is a free-flowing user documentation to help readers understand the practical usage of the Neptune software. This document is divided into four sections – (i) installation and file organization (pages 2–6), (ii) pre-processing in Neptune (pages 6–23), (iii) post-processing (pages 24–30), and (iv) understanding the outputs (pages 31–74). The data used for this demo is the '*fully_processed_data*' made available on Mendeley Data (doi.org/10.17632/75nn8v6znt.1). The quality control figures shown in Section 4 correspond to the first of the two subjects of this sample data. This data is itself part of the complete cohort presented in the main manuscript; hence, the procedures and data acquisition parameters are the same. We next describe the usage of Neptune.

### 1. Installation, file organization, and coding structure

Before setting up Neptune, its dependencies need to be installed first, which include AFNI to process fMRI images (https://afni.nimh.nih.gov) and the NIfTI toolbox to read/write NIfTI files (https://www.mathworks.com/matlabcentral/fileexchange/8797-tools-for-nifti-and-analyze-image). Installing the Spinal Cord Toolbox (SCT) (https://spinalcordtoolbox.com) is optional if users do not plan to run step-18 (co-registration to PAM50 standard space; it is necessary if this step is chosen). Nevertheless, having it significantly speeds up the manual segmentation of tissue boundaries during steps 5 and 10. We assume the user already has MATLAB installed (Neptune has been tested on versions R2019b through R2022a). The Neptune toolbox zip file can be downloaded from MATLAB Central ( ) or GitHub ( ). Here is how the downloaded Neptune folder appears:

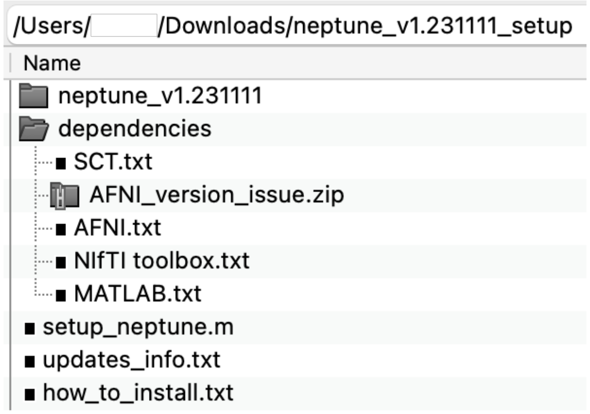


The first step is to read '*how_to_install.txt*'. A history of all significant changes to Neptune across different versions is available in '*updates_info.txt*'. The '*dependencies*' directory provides links to download software before installing Neptune (using '*setup_neptune.m*'). We will next look at the main directory '*neptune_v1.231111*' (here, *v1* refers to version 1, and *23/11/11* is the release date in year-month-day format). Although newer versions of Neptune might be published by the time the reader sees this, the broad contours, folder structures, and file names remain consistent with the descriptions provided here.

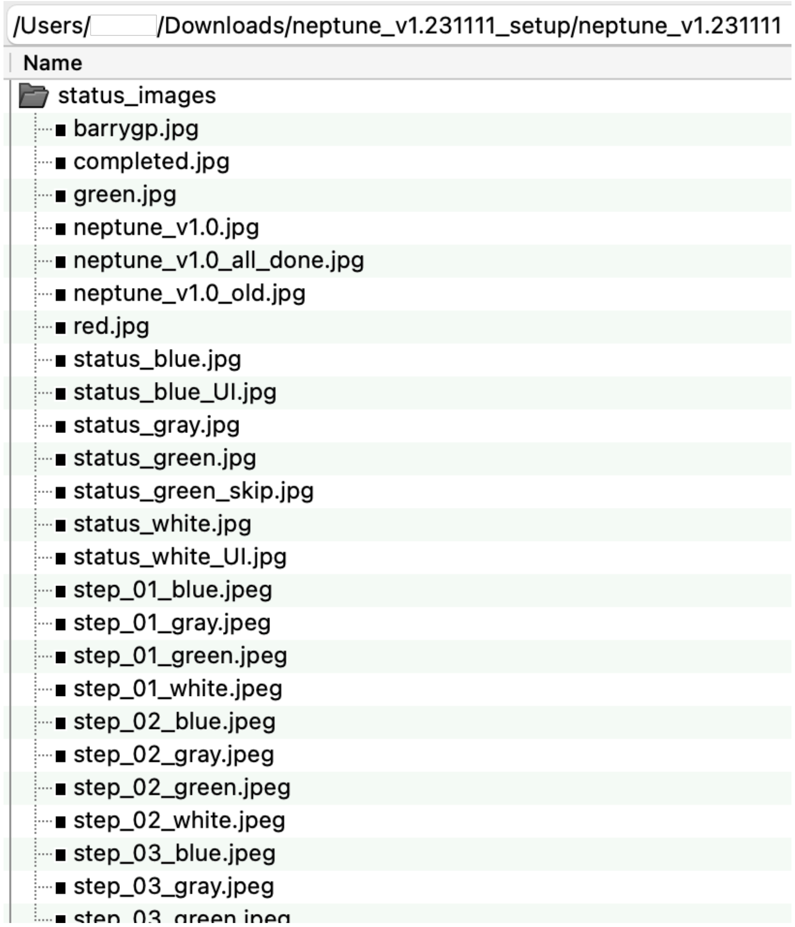


The above directory contains images used for displaying the processing status during execution.

### 2. Pre-processing of spinal cord fMRI data in Neptune

The fundamentals of some of Neptune's core pre-processing steps are described in the 2016 paper (https://pubmed.ncbi.nlm.nih.gov/26924285) (refer to the 15-step pipeline described under Methods → Data processing). Neptune has been built on the idea of mimicking a conversation between the user and the software, wherein the software asks the user to make certain choices one after the other (through GUI dialog boxes), with successive conversations dependent upon the choices made by the user at each stage. This is a different architecture from what most other research software in the field employ, with a single cohesive settings interface having a host of buttons and checkboxes to resolve before hitting the 'Go' button. They are simply different styles of programming, and both work just fine. We next bring out this 'conversation' and explain the choices that pop up when the user types '*neptune*' and hits enter in the MATLAB command window. The description of each figure is *below* the displayed figure.

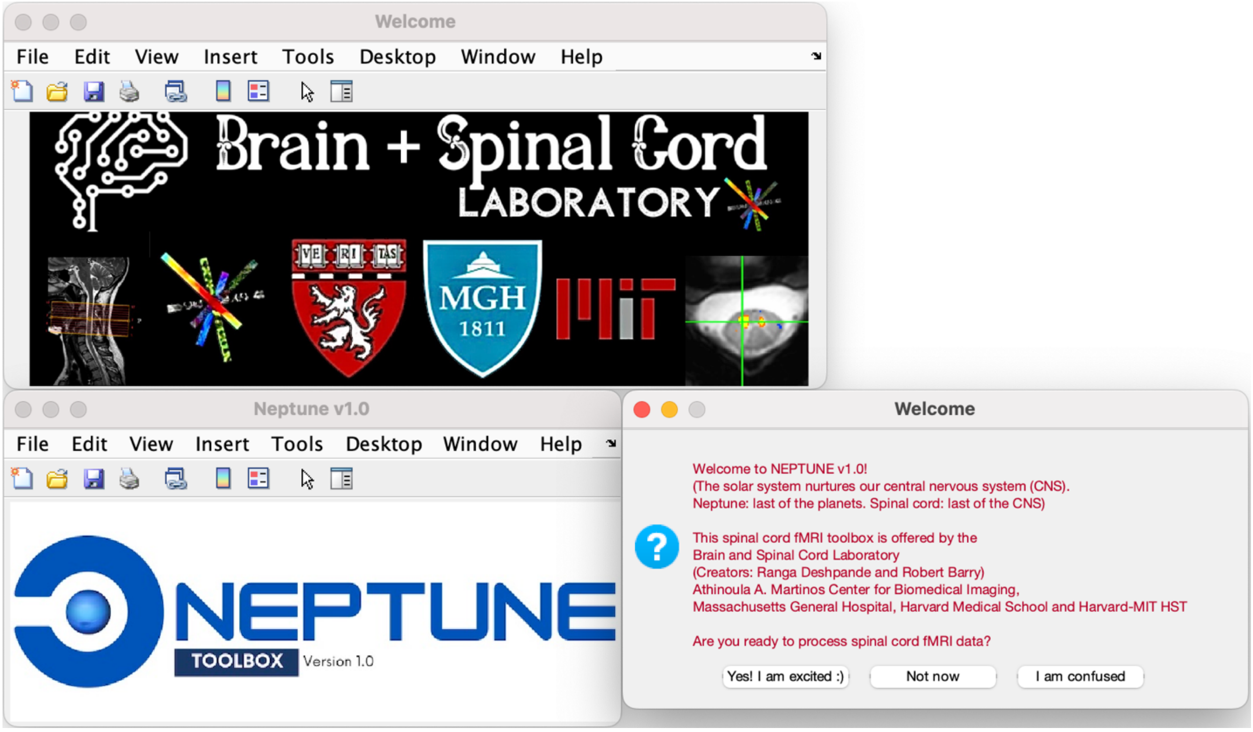


The welcome screen explains why we chose to call our software Neptune. Choosing '*Not now*' will stop the execution, and '*I am confused*' will lead the user to the Neptune manual.

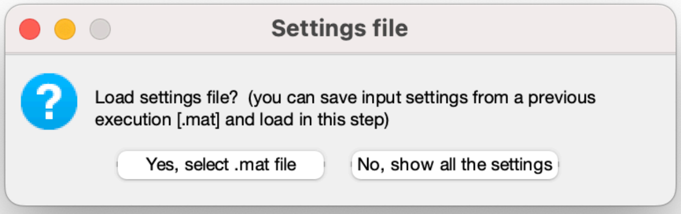


Next, one could choose to load a settings file that was automatically saved if Neptune was run previously. This saves time and effort in reselecting subjects and files and redoing certain choices. Users could also modify an old settings file by loading it into the MATLAB workspace, changing parameter values, and saving it for future use in Neptune. This, however, requires knowledge of different parameter variables used in Neptune, which can be learned by reading the main Neptune functions in MATLAB as they are well-commented. In the above dialog box, one could also choose '*No, show all the settings*', revealing all Neptune choices as described next.

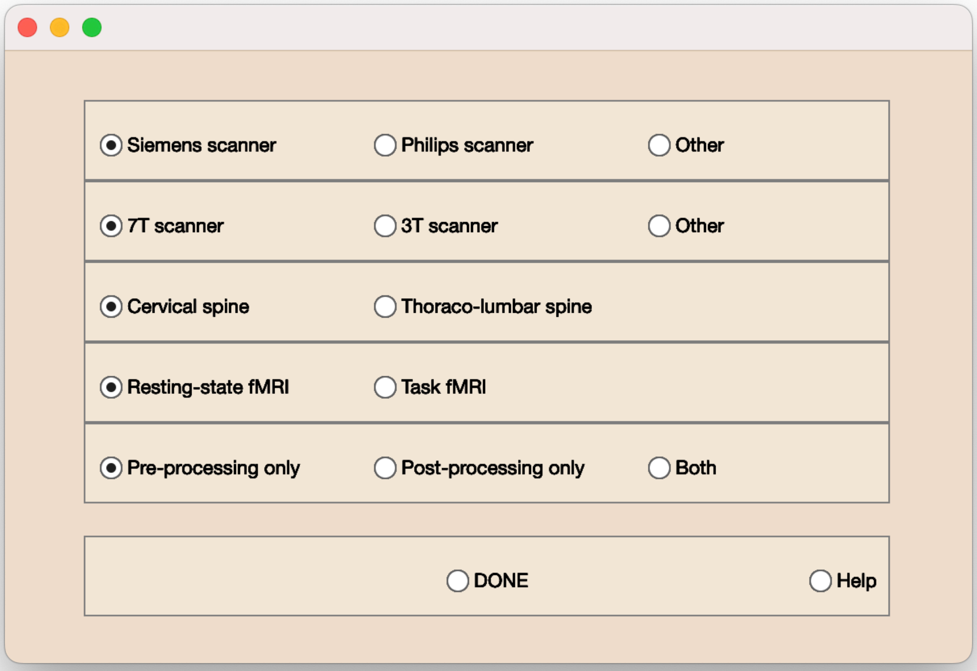


The first four choices here are, perhaps surprisingly, unimportant for most steps. However, they are still included to log what type of data is used, which matters for spinal cord imaging. One could choose to perform pre- or post-processing only (can separately do one after the other), or both.

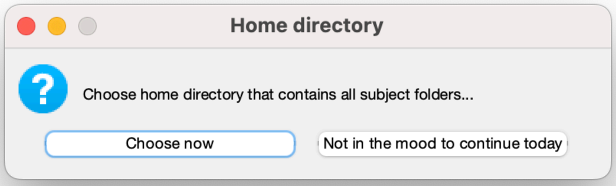


Next, choose the home directory in which all subject folders reside. Users can stop execution by choosing '*Not in the mood…*'; Neptune will save the choices made thus far and quit.

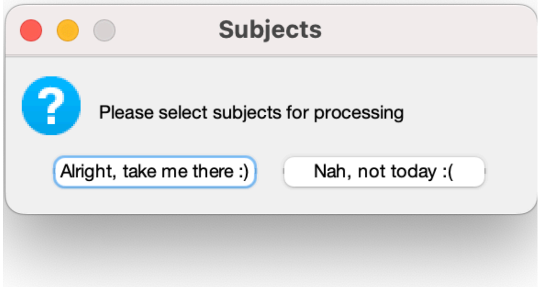


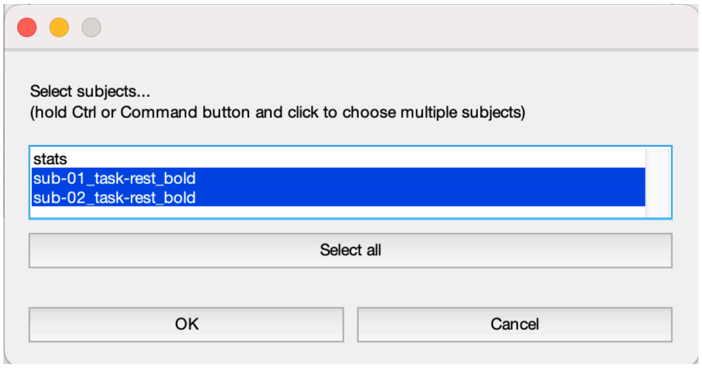


Next, select the subject folders to be processed.

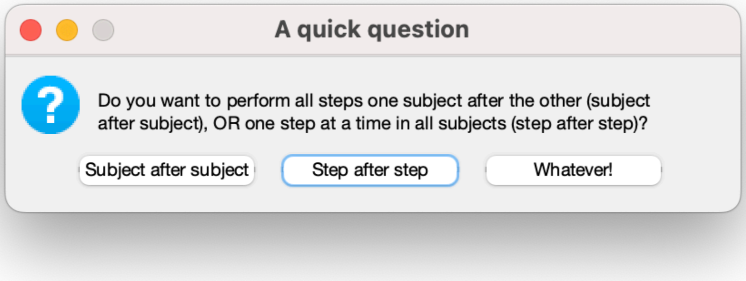


Step-after-step (the default option chosen when 'Whatever!' is selected) will run one step at a time across subjects (e.g., step-1: sub-01 → sub-02, … then step-2: sub-01 → sub-02, … and so on). Subject-after-subject will run all steps in sub-01, then move on to sub-02, and so on. Both are functionally equivalent and will produce the same outputs. The former is more suitable for limited data, while the latter is handy when running a large number of subjects across multiple days, in which case one could stop execution after one night at, say, sub-03 and resume the following night.

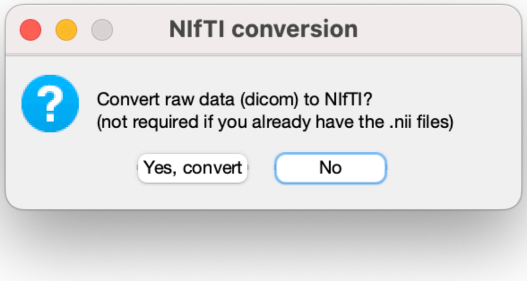


Although this functionality is available, we recommend users perform NIfTI conversion outside Neptune first, because we do not constantly update the DICOM (or PAR/REC) to NIfTI conversion code.

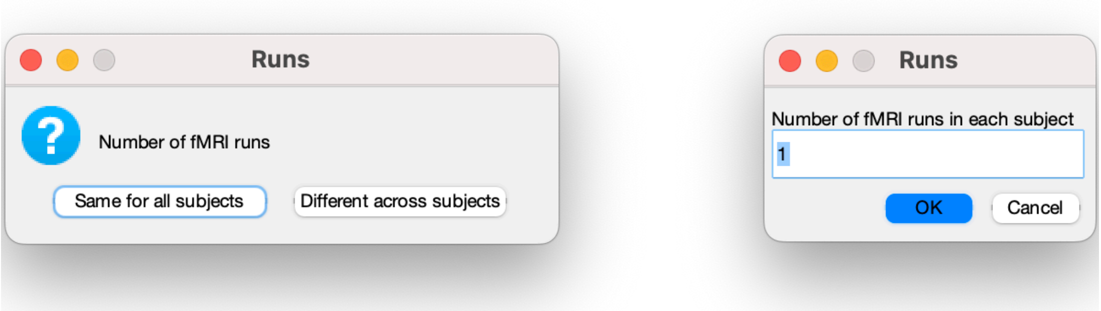


Choose the number of functional runs for each subject.

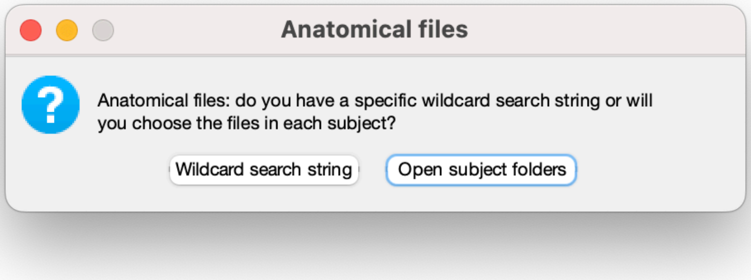


Choose the anatomical NIfTI images (either *.nii* or *.nii.gz*). If '*Open subject folders*' is selected:

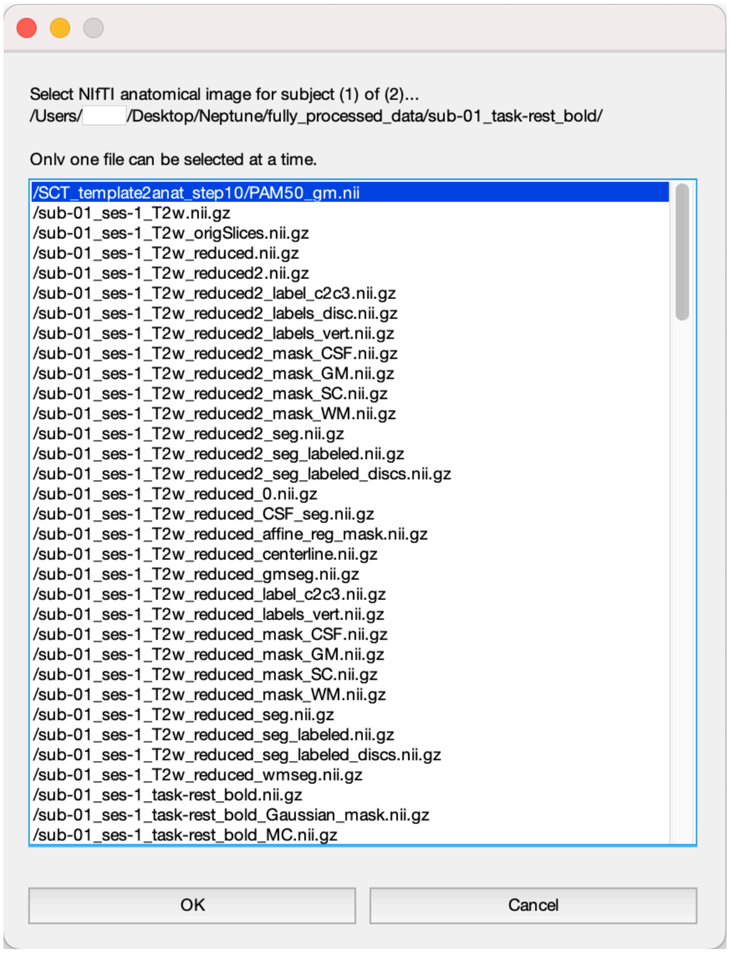

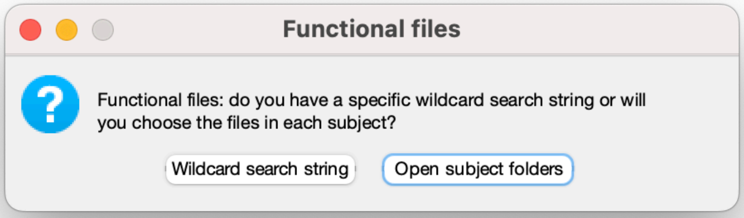


Likewise, choose functional files. If '*Wildcard search string*' is selected:

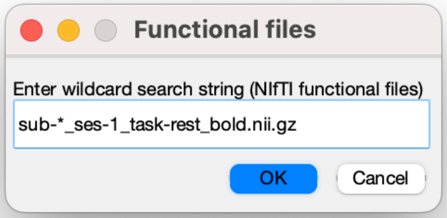


If the files have a regular naming convention (such as BIDS), it is quicker to go the wildcard route.

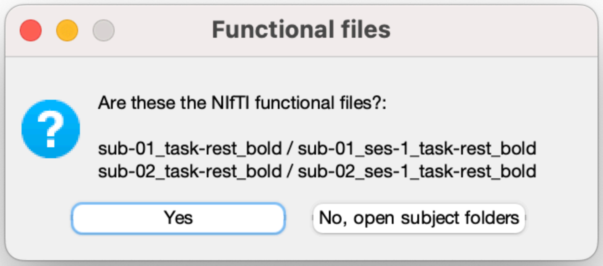


Neptune displays the identified files to make sure that they are what the user intended them to be.

Once these selections are done, Neptune displays the status bar, execution log and status dialogs, and proceeds with seeking pre-processing choices from the user.

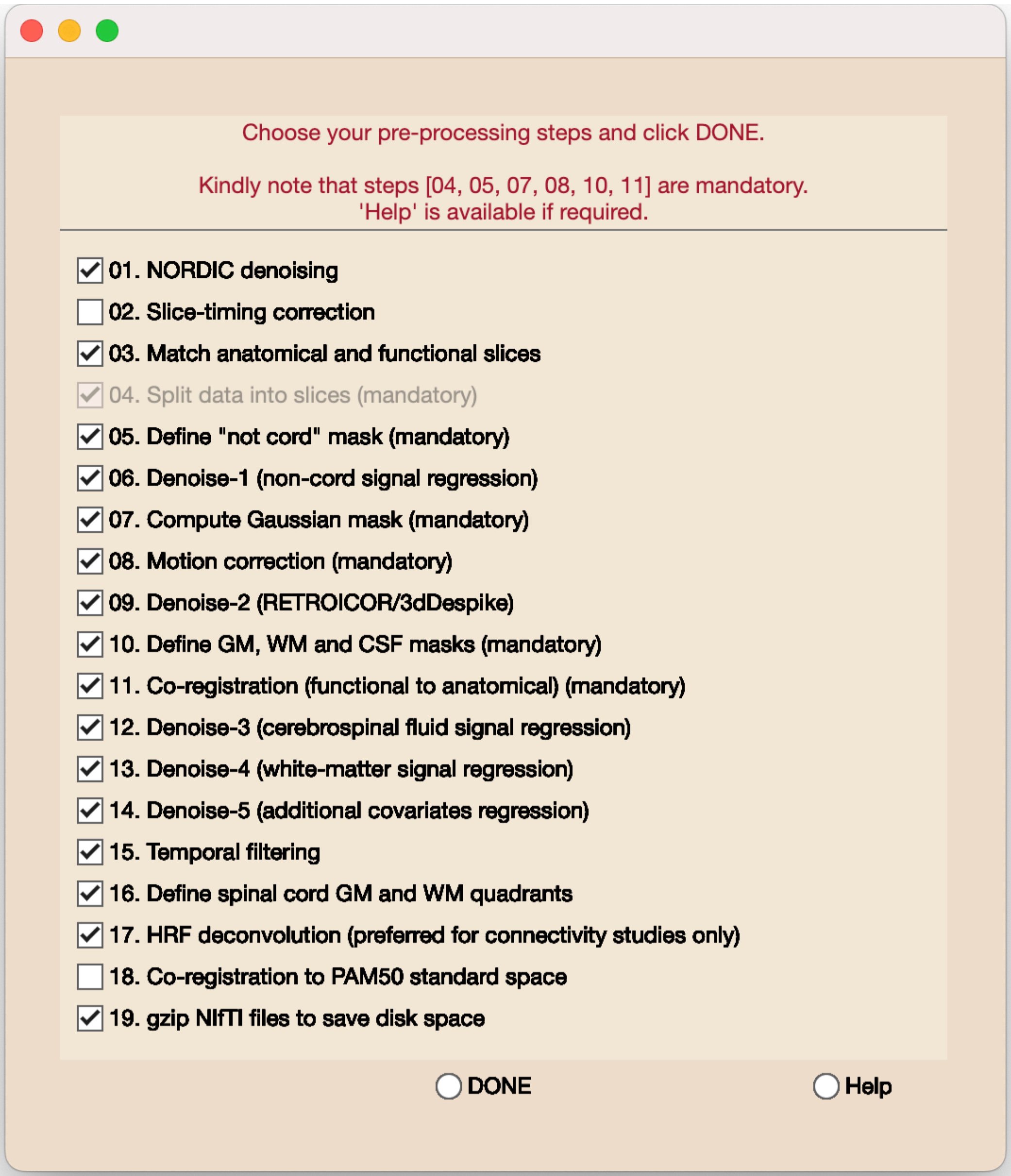


This is the box for selecting pre-processing steps. Clicking '*Help*' will redirect the user to the Neptune manual. Step-4 is run by default; the not-cord mask (step-5) and Gaussian mask (step-7) are required for mandatory motion correction (step-8). Anatomical masks (step-10) and co-registration (step-11) are also mandatory. These steps are mandatory only if further steps are being executed; that is, co-registration is compulsory for doing temporal filtering but not for just performing motion correction, and if motion correction is the endpoint, then further steps need not be chosen. If only NORDIC denoising (step-1) is selected, then none of the other steps need to be selected. Nevertheless, if the user decides to perform a unique combination of steps and is unsure, then they can go ahead and check if Neptune throws an error.

It is possible to run some steps outside Neptune and import, but the naming convention must be kept consistent. Each successive step of Neptune expects its input filenames to be within a few possible options. Although the first part of the filenames is the same as that of the raw NIfTI image, the suffixes will vary. In section 4, we will explain all Neptune output files and corresponding suffixes; the user will be able to use this information to run steps outside Neptune, name the files accordingly, and then run only part of Neptune steps on them.

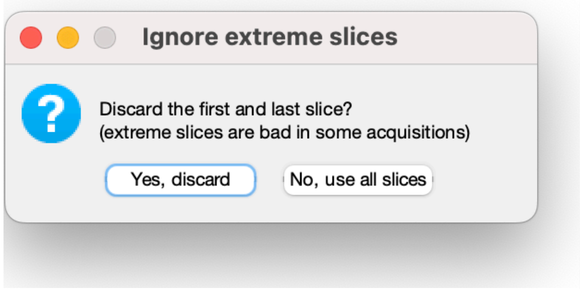


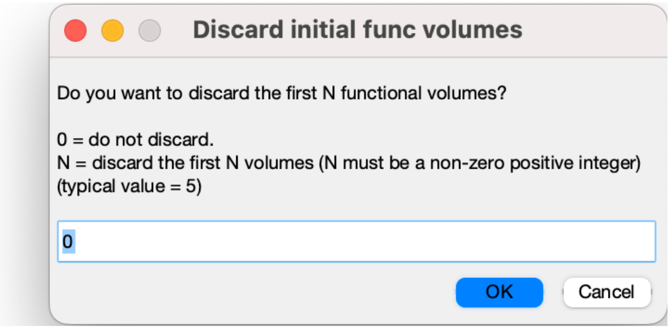


After the selections are complete, Neptune asks the user if they want to discard the top and bottom slices because these slices are of poorer quality in some acquisitions. Neptune also asks if the user wishes to discard the first few functional volumes (to account for the approach to steady-state magnetization).

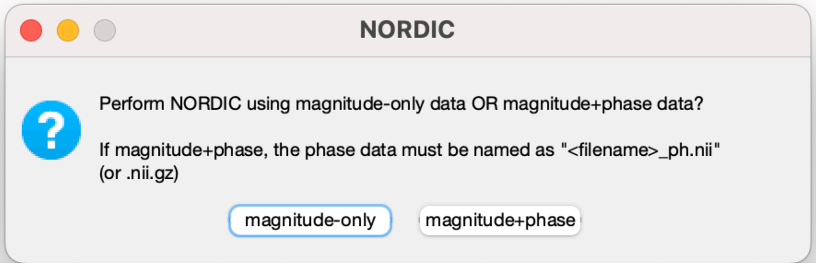


If NORDIC denoising is selected, then Neptune can run NORDIC on just reconstructed magnitude NIfTI data, or, if available, using both magnitude and phase images (which is better). Most fMRI sequences have an option on the scanner to reconstruct both (which we recommend). The NORDIC source code from the original authors was used by us, with some minor modifications of our own. Notably, this implementation does not perform NORDIC on raw complex K-space data from each receive channel (which would be ideal).

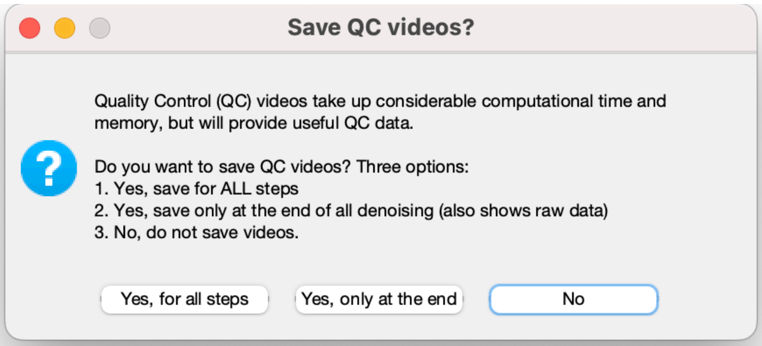


QC videos are motion pictures of fMRI time series saved as *.mp4* files. They can help detect problematic scans and identify issues with the data that are often easy to miss. Generating these videos, however, comes with considerable cost. The videos take up space (sometimes as bulky as the fMRI data itself), and generating these videos takes time (sometimes the lengthiest part of the whole execution). For these reasons, the default option is not to save the videos, but users could choose to save them after each processing step, or only at the end of the last denoising step (compares raw vs. denoised data).

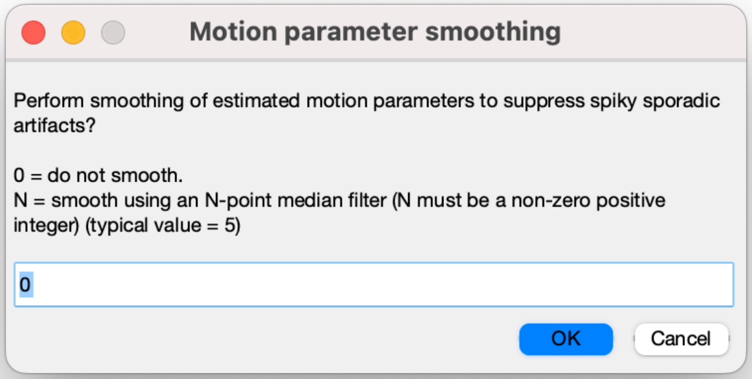


The spinal cord is susceptible to sudden short-lived motion events often arising from nearby organs such as the throat (swallowing). This can affect fMRI data and lead to large motion spikes. Motion parameter smoothing applies an N-point median filter on estimated motion parameters to suppress the spikes, which can sometimes be helpful for effective and efficient motion correction. If '0' is entered, then motion parameter smoothing is not performed.

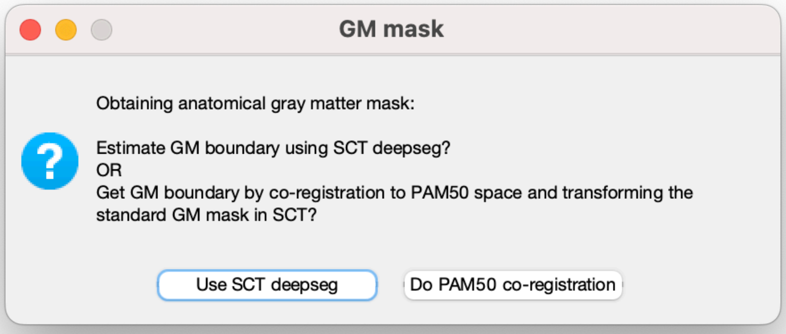


For step-10 (anatomical segmentations), Neptune follows a semi-automated procedure wherein the user finalizes the tissue boundaries, but this process is expedited by using SCT automated segmentations. Neptune presents the SCT-estimated boundaries in the same interface as predefined segmentation, and the user will be able to move around the points to the desired visual accuracy. If SCT exists, the user can either choose to determine the gray matter (GM) boundary in a data-driven way using *deepseg*, or co-register the PAM50 template onto the anatomical space for the boundaries. White matter (WM) and cerebrospinal fluid (CSF) boundaries are always determined using *propseg*, which is generally reliable.

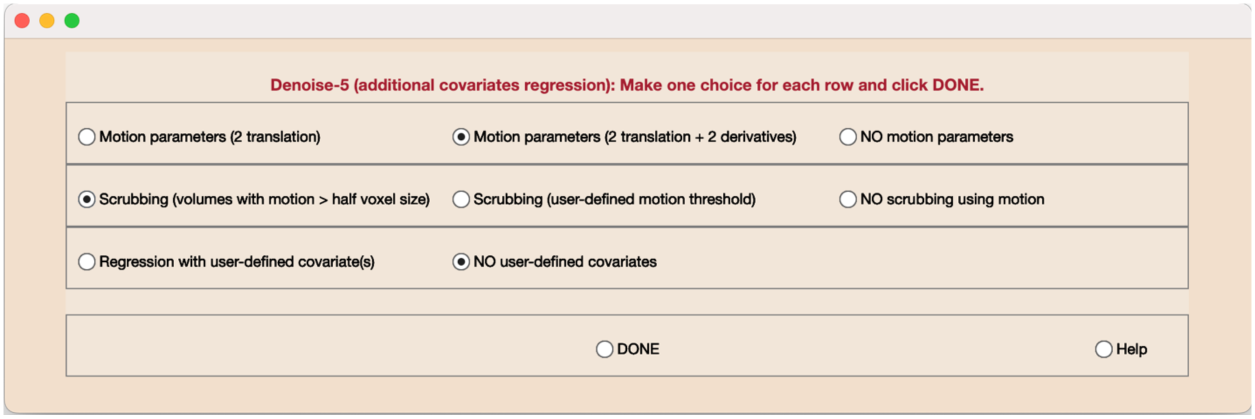


If step-14 (additional covariates regression) is chosen, then the user can pick motion covariates, scrubbing covariates, as well as user-defined covariates. Since motion correction is done slice-wise, there is no *z*-component, and only *x* and *y* translations/rotations are possible. Furthermore, we assume that rotation is negligible because the spinal cord is roughly circular and early testing showed rotation estimates to be unreliable. Scrubbing uses a binary covariate, with each time point being 0 or 1. The threshold can either be automatically picked (over half the in-plane voxel size) or manually entered. User-defined covariates can be *.txt*, *.csv*, *.xls*, or *.xlsx*, with each row corresponding to one time point. The number of rows must equal the number of time points.

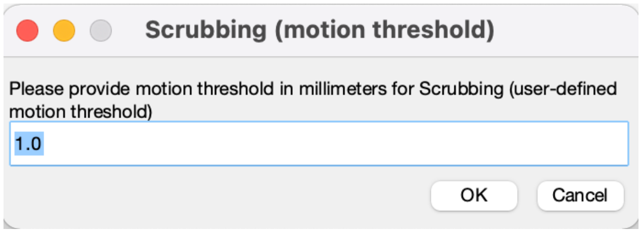


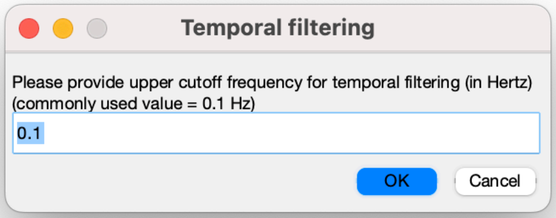


If step-15 (temporal filtering) is selected, then the user can provide the upper cut-off frequency for bandpass filtering (in Hertz). The lower cut-off is always 0.01 Hz (which is also the cut-off for high-pass filtering). It is possible to easily modify this within the code (*scfMRItb_15_filterData.m*).

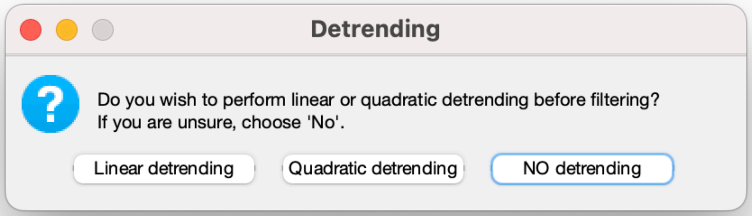


The detrending option is provided to account for linear/quadratic trends introduced in the fMRI time series because of gradual scanner drifts. Detrending, if chosen, is done before filtering.

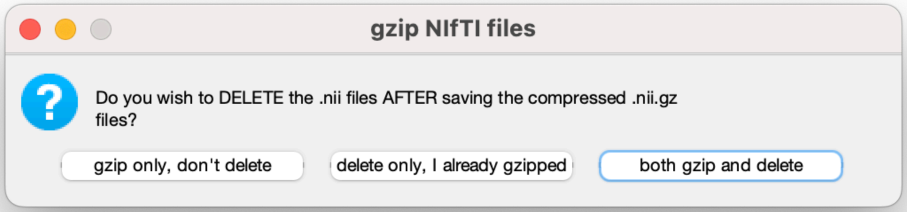


If step-19 (compress NIfTI files) is chosen, then the default (and recommended) option is to both gzip and delete, which will first compress *.nii* files and then delete the *.nii* files only after verifying that the corresponding *.nii.gz* file exists.

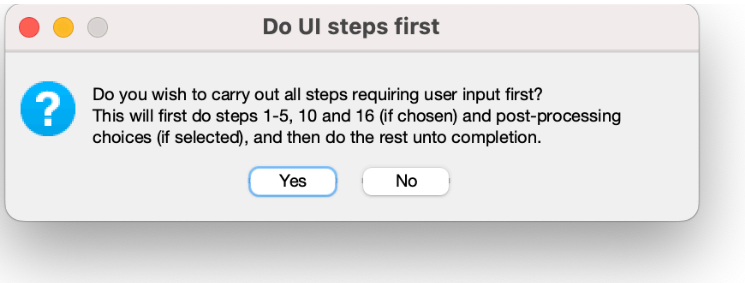


After all these selections are made, Neptune is ready to begin processing. However, before that, it asks the user if they want to first perform steps requiring user input (not-cord mask, anatomical mask, cord quadrants) before doing the rest of the steps. This is helpful because the user can finish all tasks requiring their input (e.g., drawing tissue boundaries) and then let the computer run until the processing is complete. If this is not chosen, then processing will stop midway through the first subject until the user attends to the step that requires interaction.

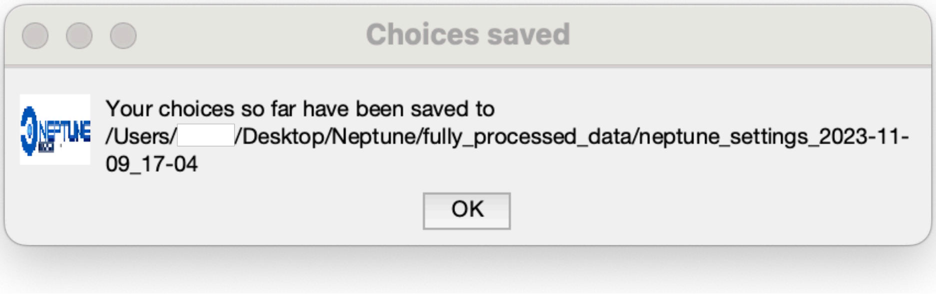


Neptune informs the user that all selections have been saved as a workspace variable in the given location and then commences processing. If the execution were to terminate midway for any reason, then the user can re-run Neptune, load this settings file, and resume processing without needing to go through these selection options again.

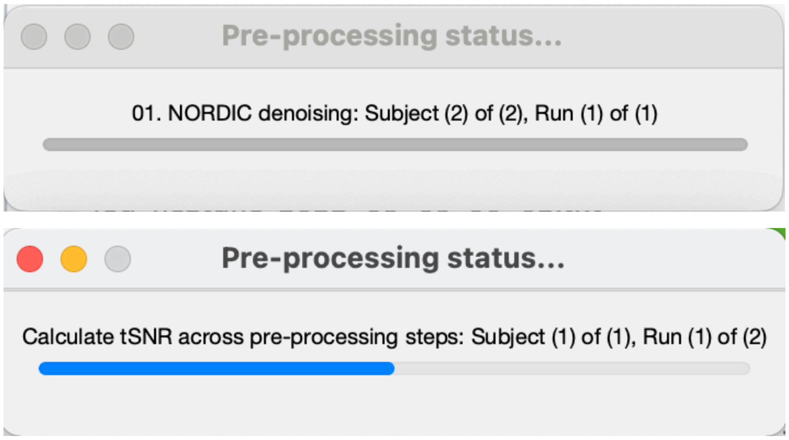


Neptune shows the latest processing status in a dialog box.

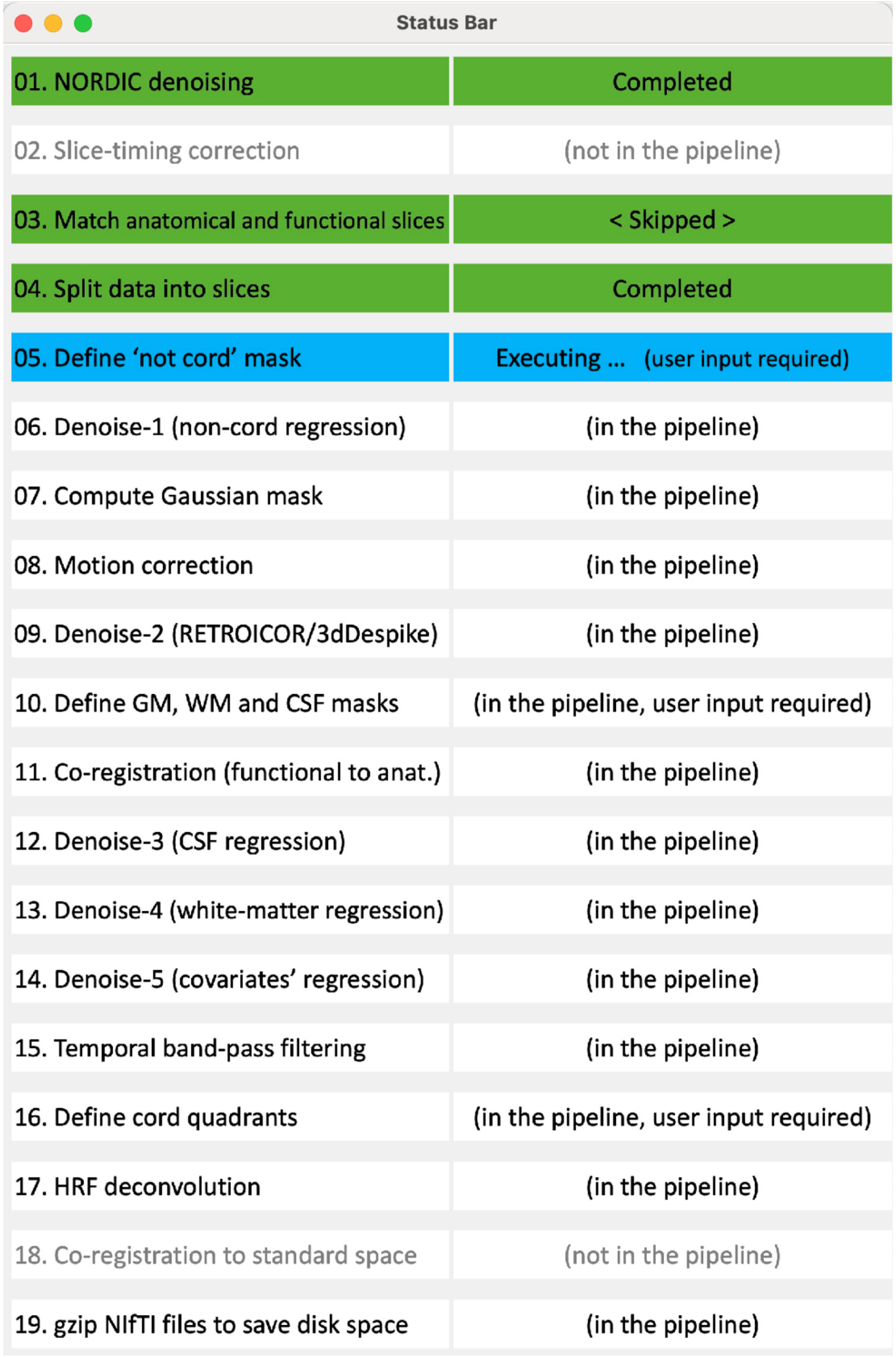

Status Bar

| | |
|---|---|
| 01. NORDIC denoising | Completed |
| 02. Slice-timing correction | (not in the pipeline) |
| 03. Match anatomical and functional slices | < Skipped > |
| 04. Split data into slices | Completed |
| 05. Define 'not cord' mask | Executing ... (user input required) |
| 06. Denoise-1 (non-cord regression) | (in the pipeline) |
| 07. Compute Gaussian mask | (in the pipeline) |
| 08. Motion correction | (in the pipeline) |
| 09. Denoise-2 (RETROICOR/3dDespike) | (in the pipeline) |
| 10. Define GM, WM and CSF masks | (in the pipeline, user input required) |
| 11. Co-registration (functional to anat.) | (in the pipeline) |
| 12. Denoise-3 (CSF regression) | (in the pipeline) |
| 13. Denoise-4 (white-matter regression) | (in the pipeline) |
| 14. Denoise-5 (covariates' regression) | (in the pipeline) |
| 15. Temporal band-pass filtering | (in the pipeline) |
| 16. Define cord quadrants | (in the pipeline, user input required) |
| 17. HRF deconvolution | (in the pipeline) |
| 18. Co-registration to standard space | (not in the pipeline) |
| 19. gzip NIfTI files to save disk space | (in the pipeline) |

After processing starts, three steps require user input – not-cord mask, GM/WM/CSF anatomical masks, and defining cord quadrants. We briefly demonstrate each next.

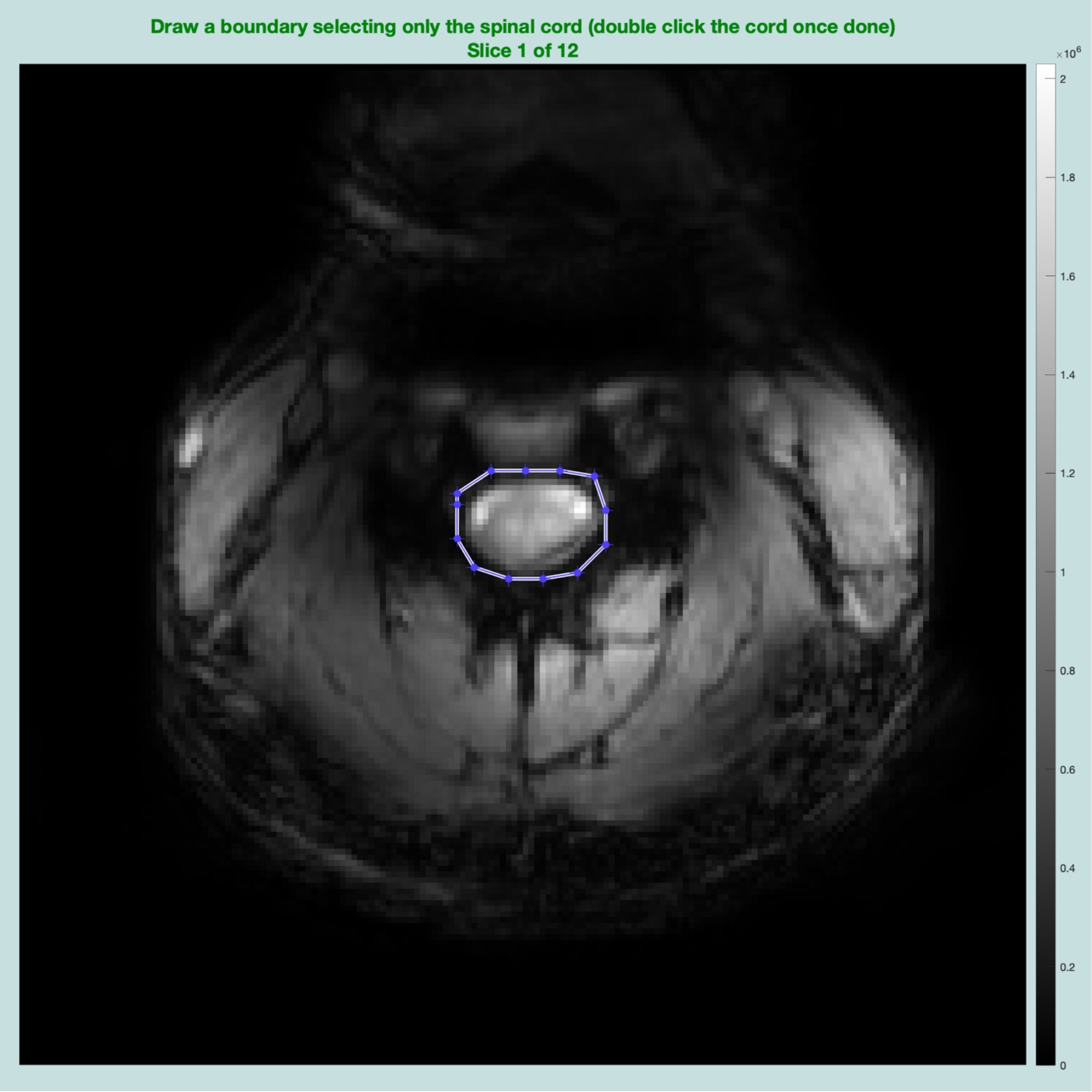


For step-5 (not-cord mask), Neptune runs SCT *propseg* in the background on the mean functional image (if SCT is installed) and displays an estimated boundary for each slice. The user can modify the points suitably and double-click within the chosen volume to make the selection. If the boundary is dissatisfactory, the user can right-click to delete it and then draw their own boundary point-by-point until the shape is closed. The resulting binary mask is used in step-6 (denoise1: not-cord signal regression) to denoise signals from outside the cord. This step is unique to spinal cord fMRI as a comparable step is generally not done when pre-processing brain data. The binary mask is also used to automatically generate a Gaussian mask in step-7, which is then used in step-8 (motion correction) to estimate motion only in the cord. This mask thus does not need to be extremely precise, as an error of a couple of voxels is unlikely to make a significant difference.

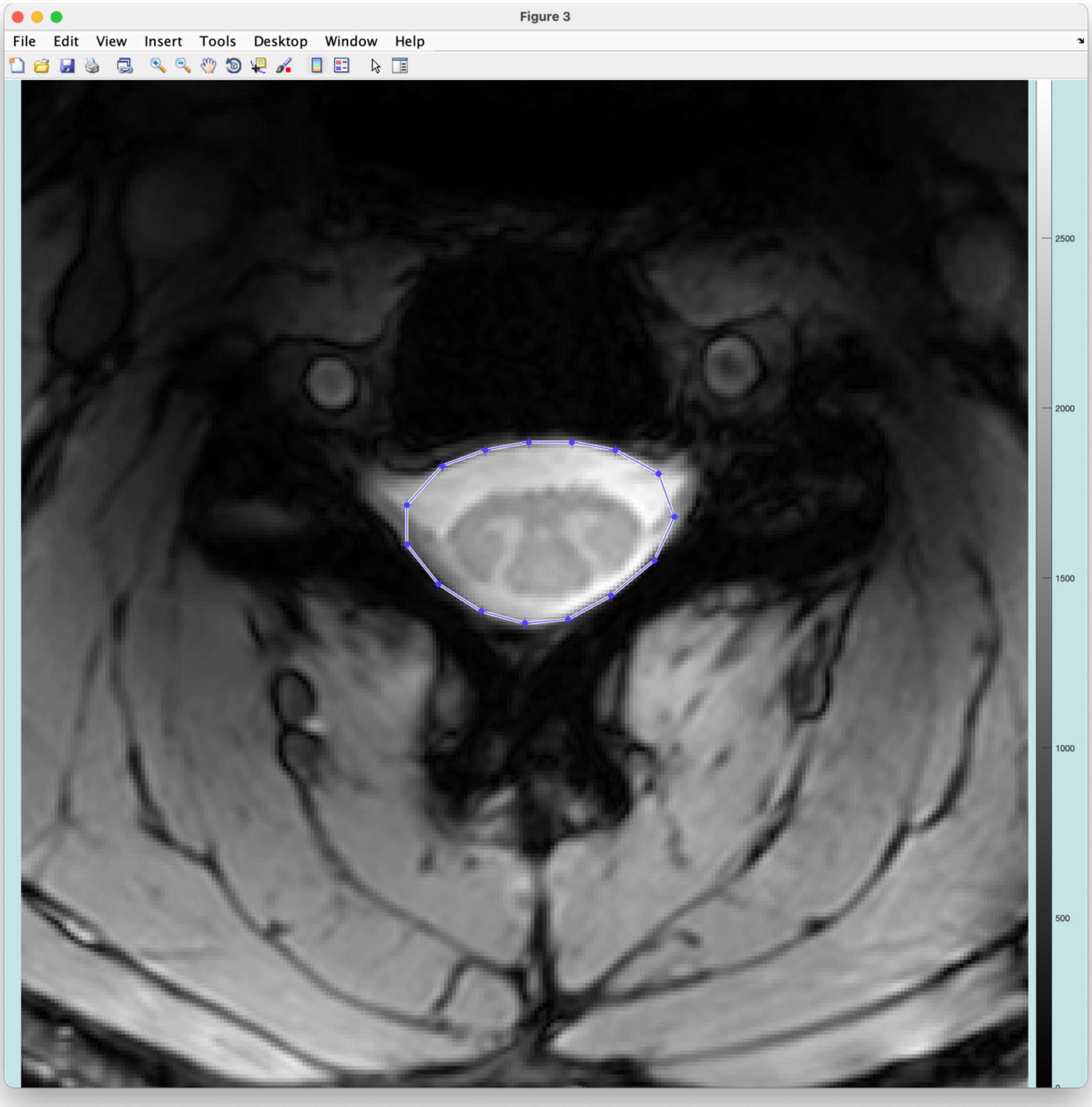


Drawing the CSF boundary (shown above) and WM boundary (shown below) are similar.

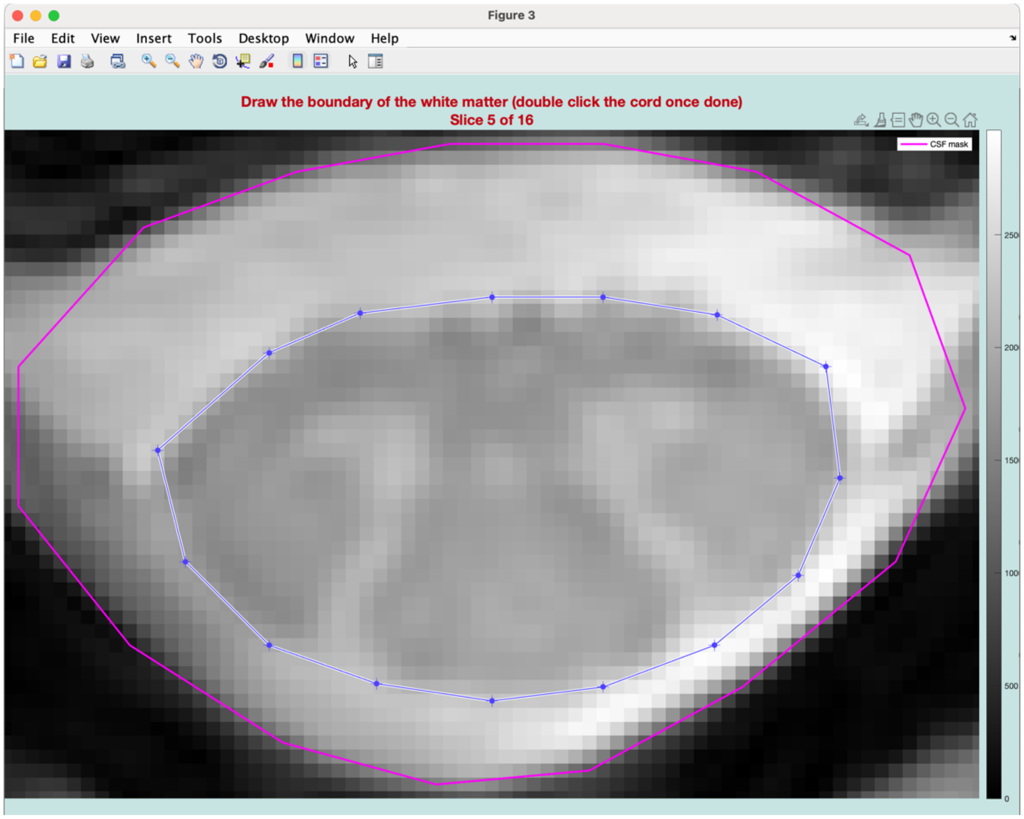


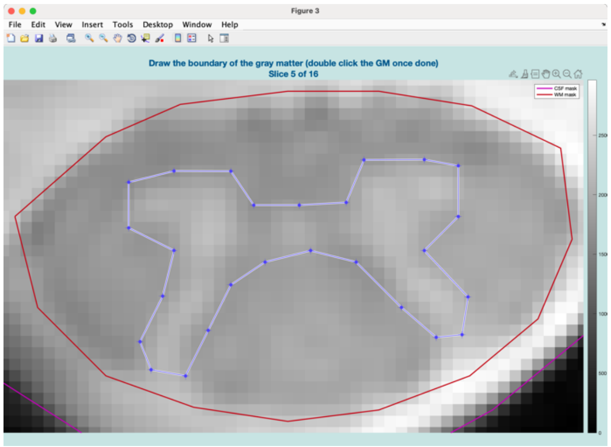


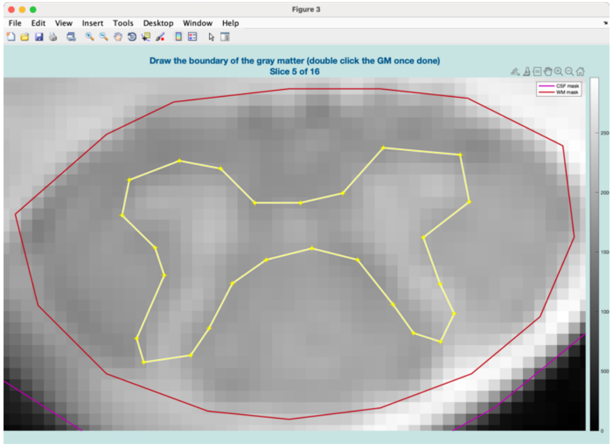


In the gray matter, if SCT *deepseg* was chosen, a blue boundary will appear (the figure on the left) showing the boundary estimated by *deepseg* that can be edited, or otherwise deleted and redrawn manually. If PAM50 co-registration was chosen, then Neptune will run both *deepseg* and SCT co-registration, and first display the blue *deepseg* boundary. If it is dissatisfactory, the user can right-click it and delete it, and then the PAM50 boundary appears in yellow (the figure on the right). If it is not satisfactory or the user is unsure, deleting it will show both the boundaries overlaid on each other to visualize the differences. The user can then delete one of the two and double-click on the other to confirm, or delete both and draw manually from scratch.

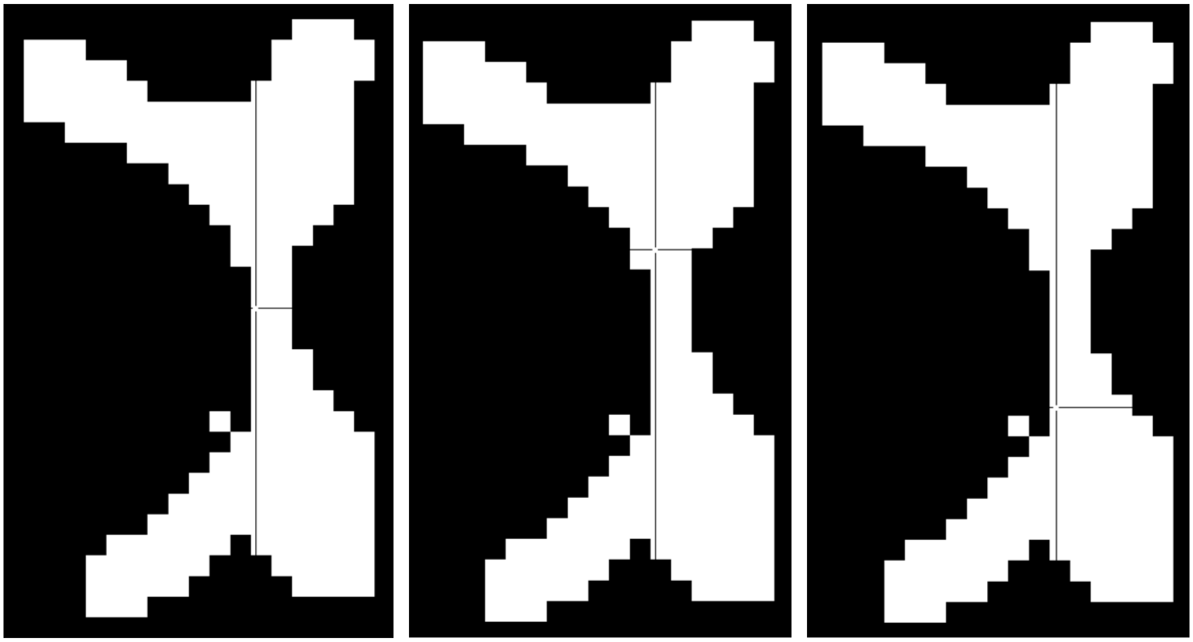

If step-16 (defining cord quadrants) is chosen, Neptune binarizes the GM mask, removes the boundary voxels shared with WM, and displays three interactive figures one after the other (left to right shown above). The GM is rotated right by 90 degrees to make this task easier (on the right half are ventral horns, and on the left are dorsal horns). In the first figure (left), the user must select the point that separates the cord into four quadrants. In the second figure (center), the user must select the point that isolates the left side of the cord, and in the third figure (right), the point that isolates the right side of the cord must be selected. After this, Neptune displays the identified quadrants in the gray and white matter:

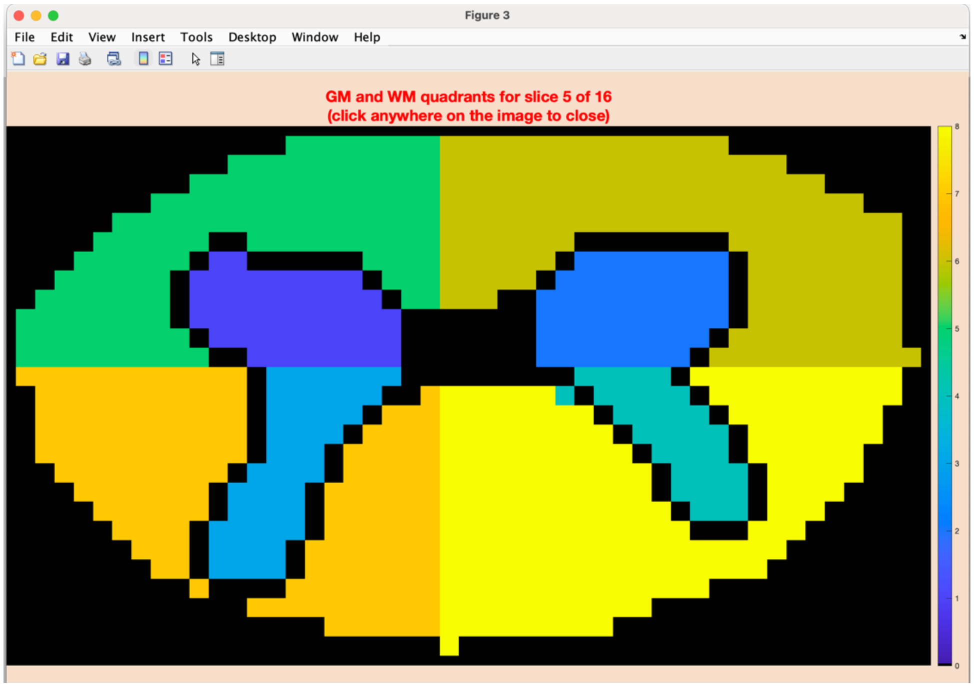

### 3. Post-processing in Neptune

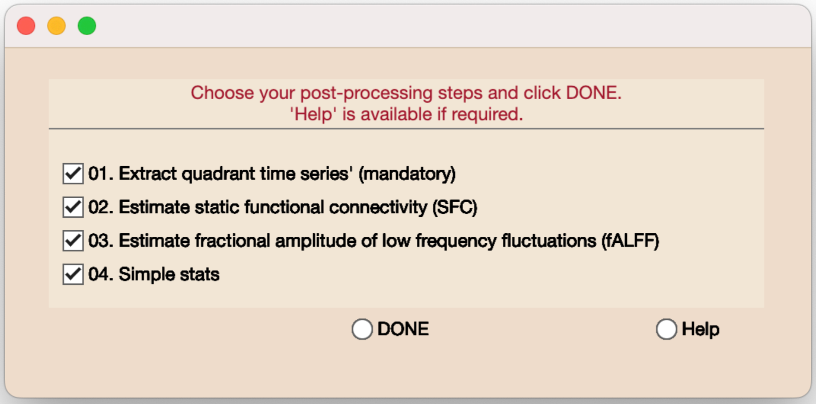


When post-processing begins, this main selection box will appear. Extracting quadrant time series is mandatory.

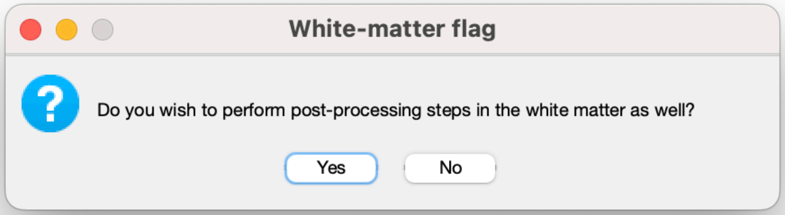


Although uncommon in the brain, WM data is sometimes processed in the cord. The larger veins can generate strong signals in WM, which could potentially be related to neural activity. Choosing 'yes' will perform post-processing in the WM as well.

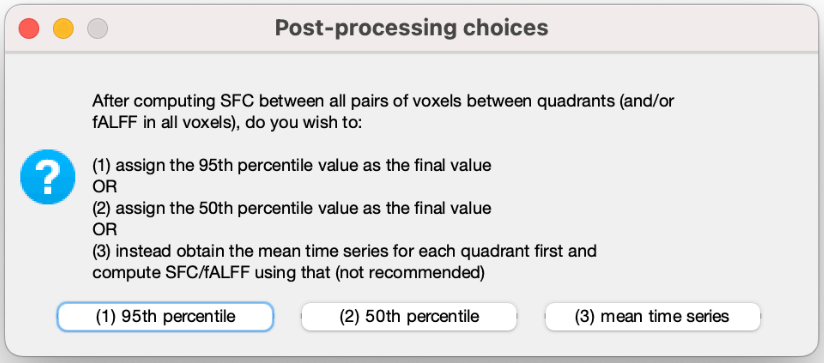


Neptune offers three ways to characterize functional connectivity (FC) between quadrants.

A simple way is computing mean time series across voxels from each quadrant and estimating FC between them (option 3 above). The first two options follow a different procedure commonly used in resting-state cord fMRI but not in the brain. Spinal cord fMRI is noisy and the anatomy is small; hence, there can be voxels affected more by noise, or voxels that are near the GM/WM boundary with questionable anatomical identity in the images. Either way, outliers are not uncommon. Averaging all the time series, including such ones, can lead to less reliable estimates. An alternative is to compute FC between all pairs of voxels between the quadrants and assign either the 95$^{th}$ percentile (option-1) or the 50$^{th}$ percentile (option-2) FC value as the final value. Users can choose based on their data quality and the research question under investigation.

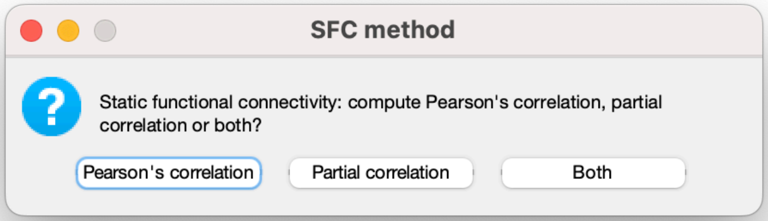


Pearson’s correlation is the default method to estimate FC. Partial correlation option is available, wherein a given correlation is controlled for FC with the rest of the quadrants.

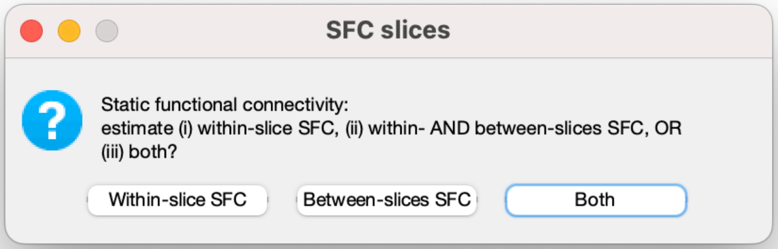


FC between different quadrants of the same slice (within-slice FC) has been the preeminent approach in the field. Between-slice FC has been found to be weak in the published studies so far. We present a way to compute and view between-slice FC. Users can choose to do both; the computational cost and execution time are minimal.

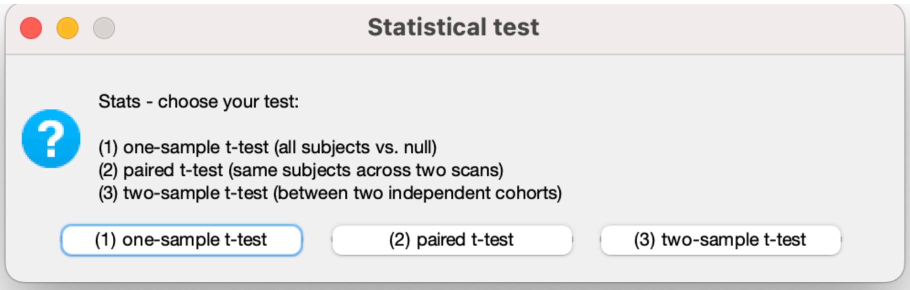


Neptune provides functionality for simple statistical analysis. We call it 'simple' because there is no option to use nuisance covariates, perform N-way or multivariate comparisons, do non-parametric statistics, or perform multiple comparisons correction. We chose to keep the statistics simple because there are elaborate statistical analysis packages both within and outside MATLAB, so Neptune's outputs (such as connectivity matrices) can be directly inputted into such software for further analyses. (From its inception, Neptune was never intended to be a statistical analysis toolbox.) Nevertheless, our simple stats tool can perform one-sample, two-sample and paired t-tests with a user-defined threshold. It can help gain preliminary insights before further elaborate statistical analyses.

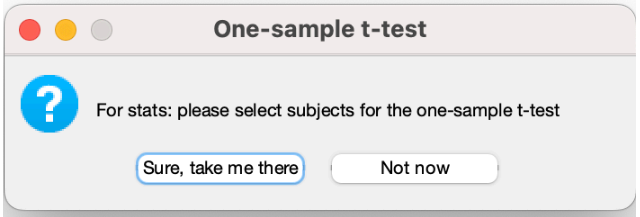


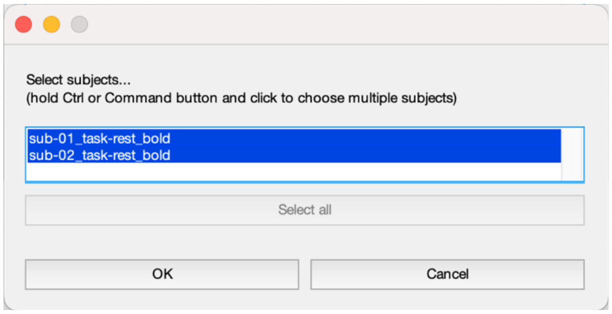


A one-sample t-test is appropriate in the presence of a single group compared against null. A dialog box allows users to choose the subjects. The p-value threshold can be defined as shown next:

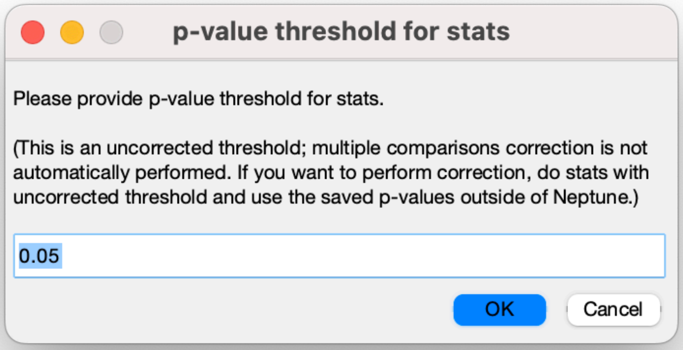

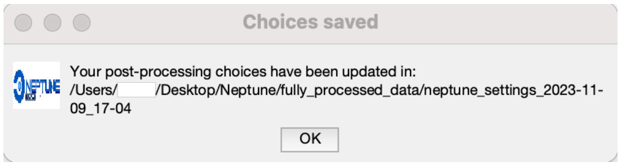


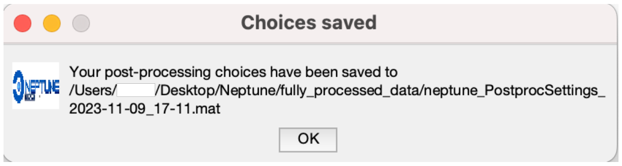


All pre- and post-processing choices are saved, after which Neptune begins execution.

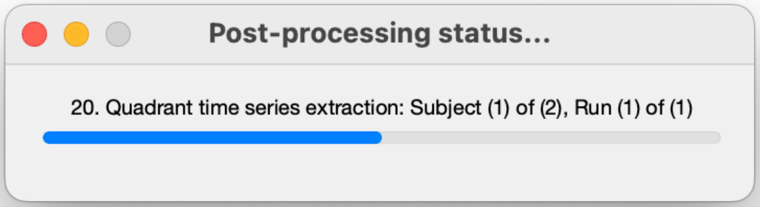


A status dialog will constantly update the users with the processing status.

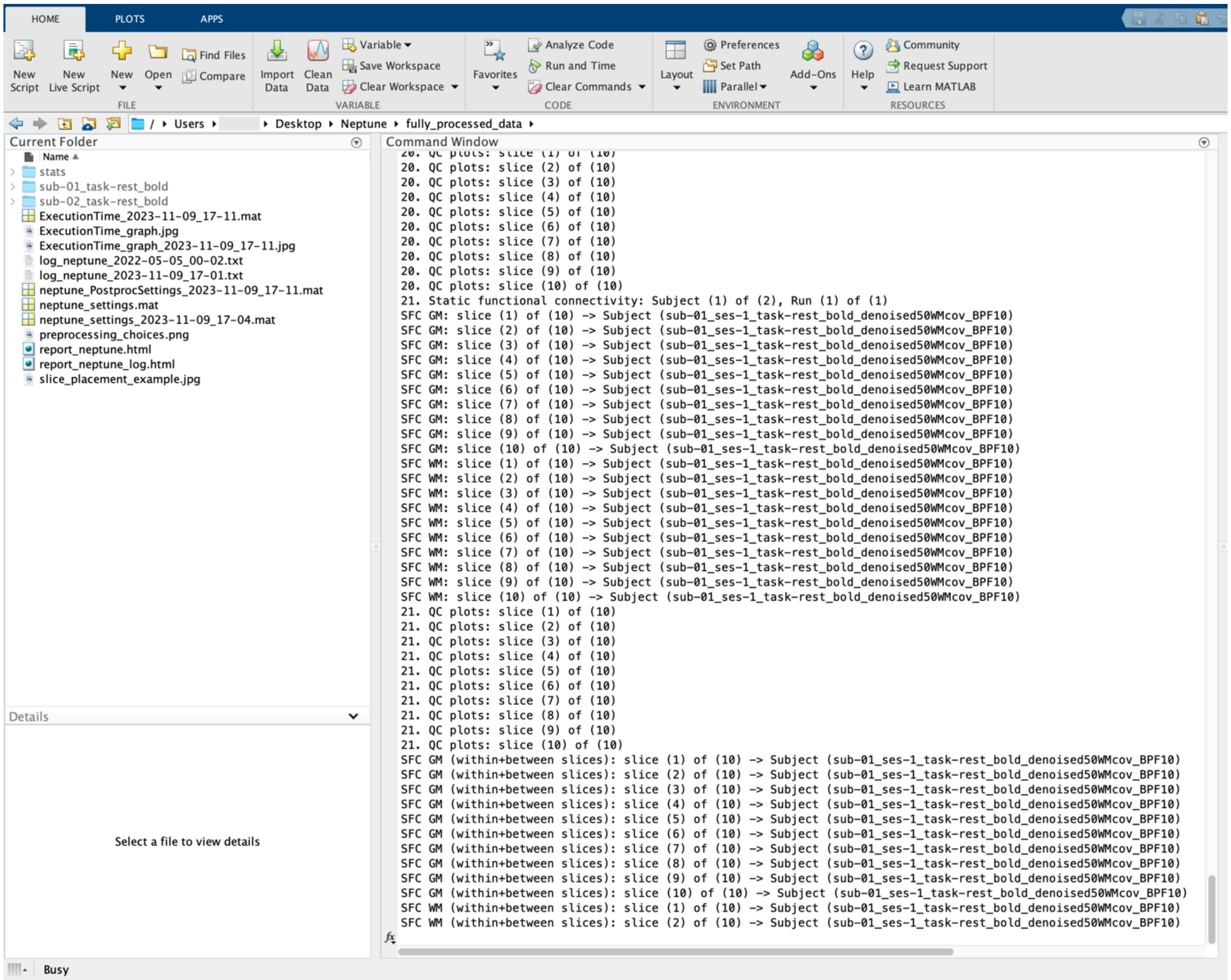


Thorough status messages are also displayed in the MATLAB command window.

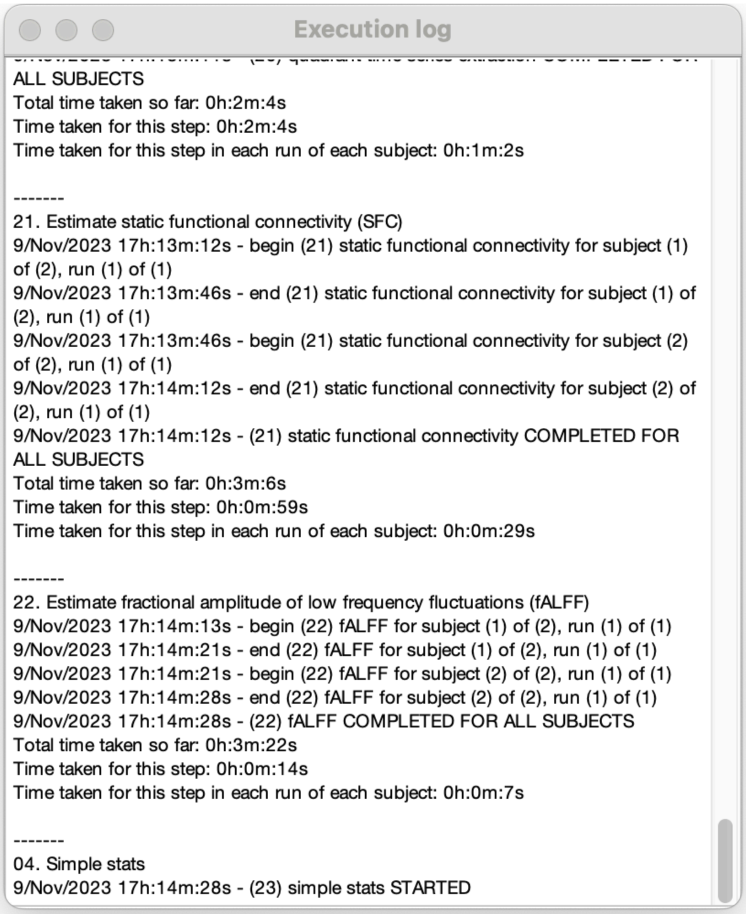


The execution log is constantly updated with the latest information. Users can scroll through it to find out exactly what choices were made, which steps were performed, beginning/ending times and total duration, starting from the beginning of execution until the current time. An effort has been made to log every possible information without omitting any details.

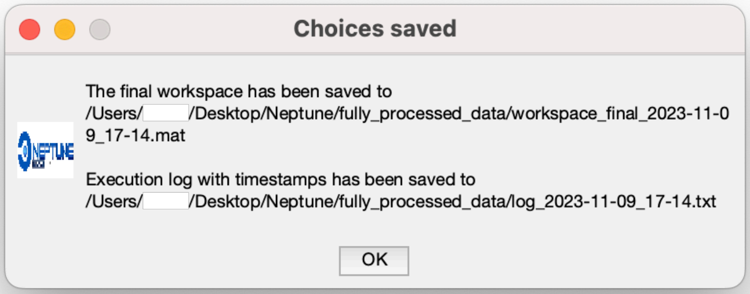


At the end of execution, the final workspace is saved. The entire execution log is also saved as a log file (*.txt*). The filename suffixes are timestamped so that they are never overwritten.

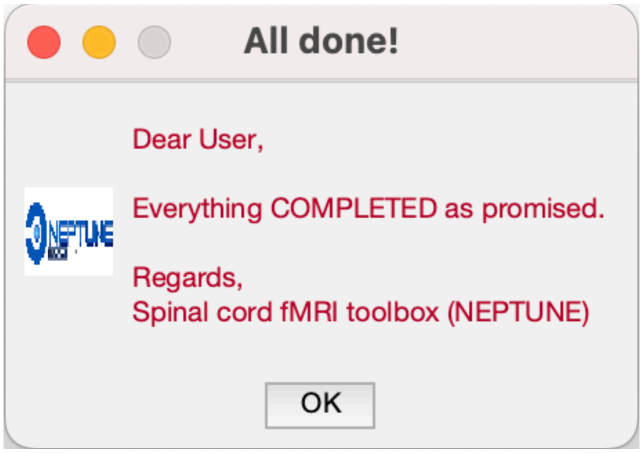


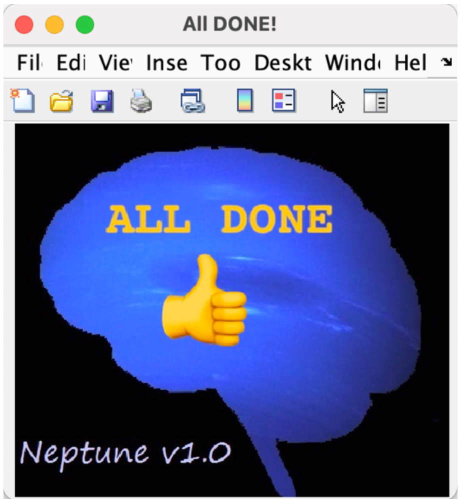


At the end of processing, Neptune politely informs the user that it is done.

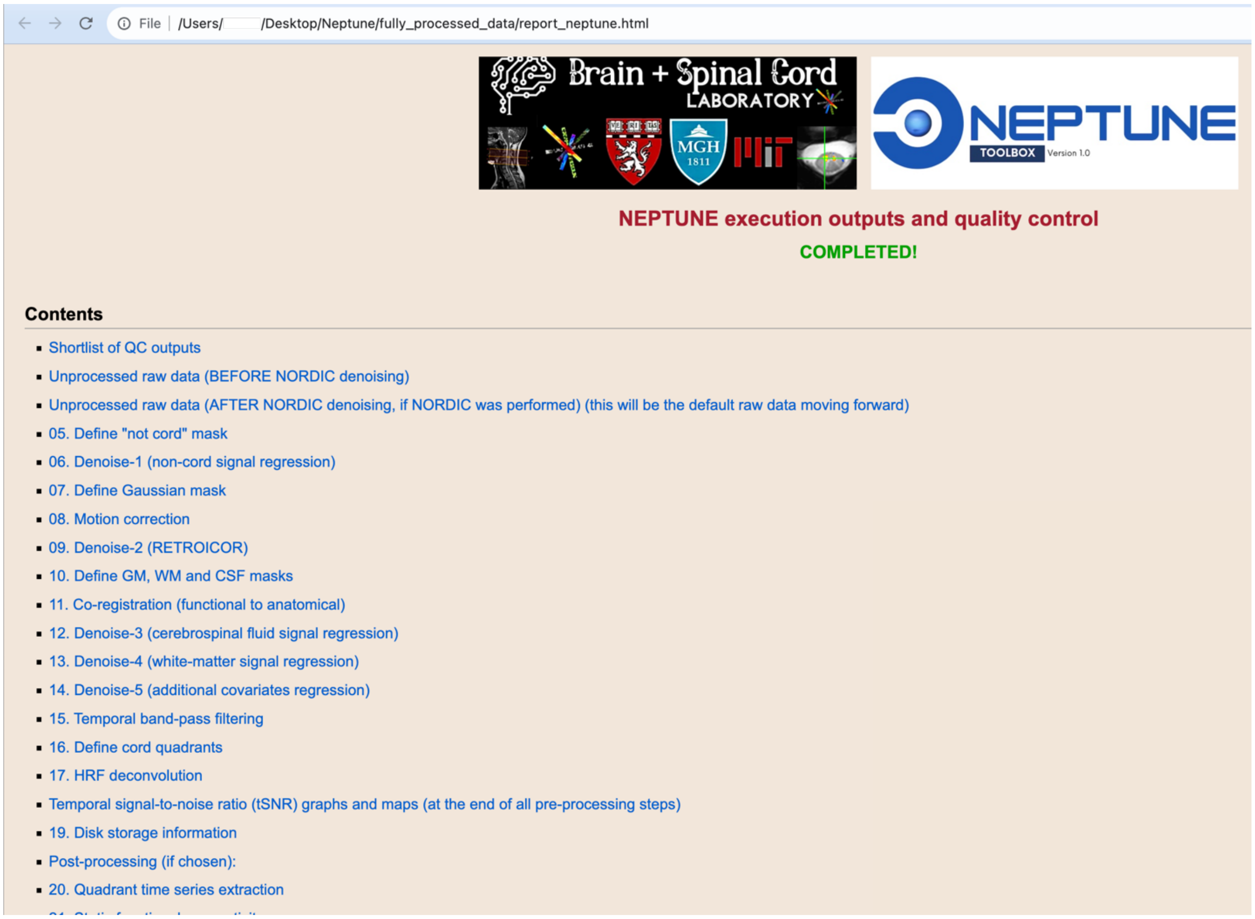


Neptune then generates a webpage with links to all QC outputs and opens them in the default browser. Users can assess their data quality and QC outputs conveniently using it.

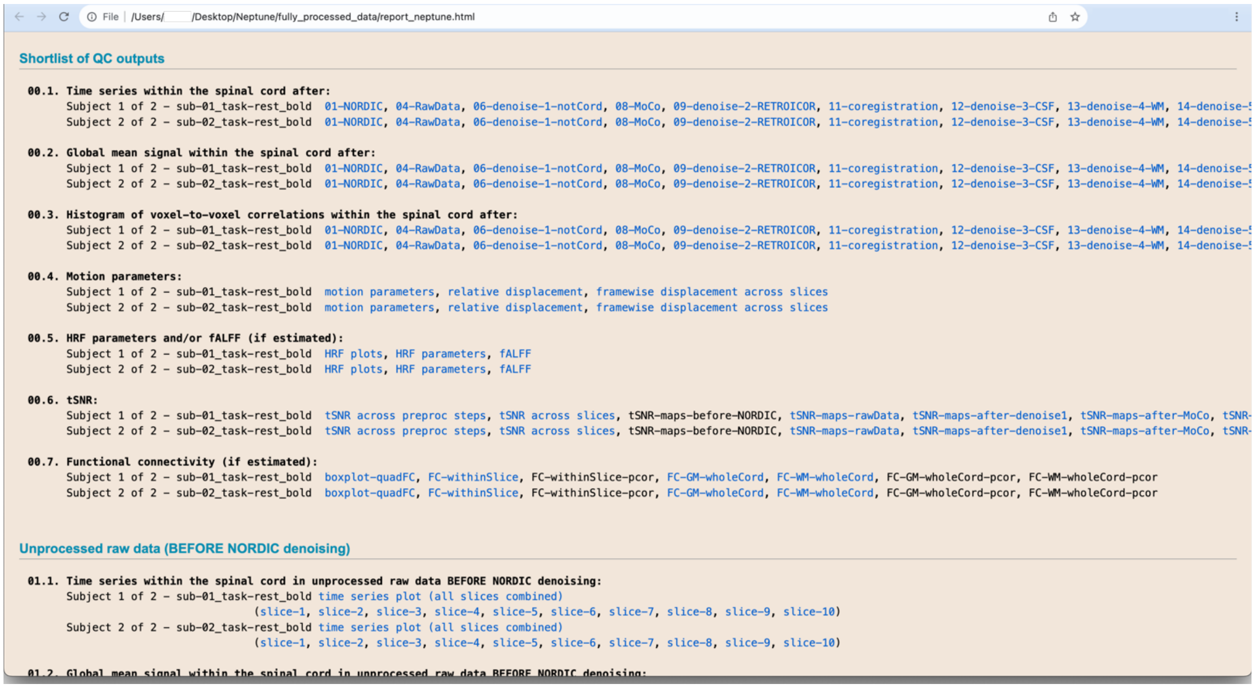

File | /Users/ /Desktop/Neptune/fully_processed_data/report_neptune.html

**01.2. Global mean signal within the spinal cord in unprocessed raw data BEFORE NORDIC denoising:**

Scrolling down shows the shortlisted QC outputs for quick assessment.

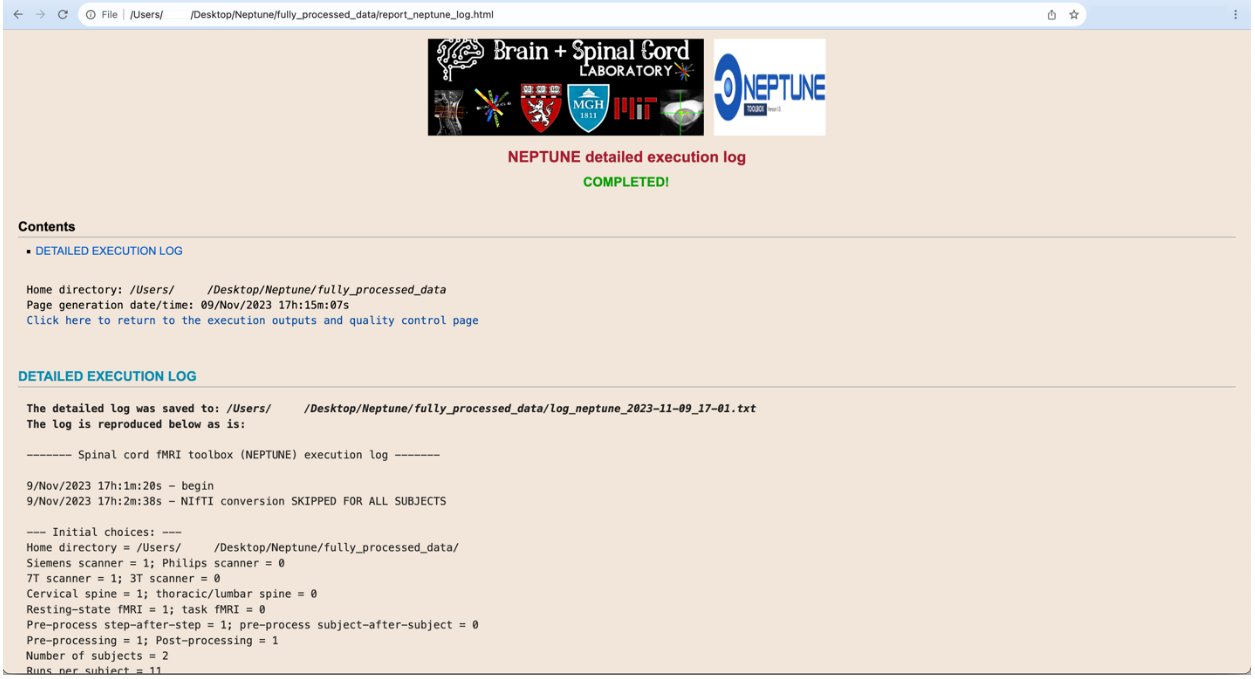

File | /Users/ /Desktop/Neptune/fully_processed_data/report_neptune_log.html



**NEPTUNE detailed execution log**

**COMPLETED!**

**Contents**

- DETAILED EXECUTION LOG

Home directory: /Users/ /Desktop/Neptune/fully_processed_data
Page generation date/time: 09/Nov/2023 17h:15m:07s
Click here to return to the execution outputs and quality control page

**DETAILED EXECUTION LOG**

**The detailed log was saved to: /Users/ /Desktop/Neptune/fully_processed_data/log_neptune_2023-11-09_17-01.txt**
**The log is reproduced below as is:**

```
------- Spinal cord fMRI toolbox (NEPTUNE) execution log -------

9/Nov/2023 17h:1m:20s - begin
9/Nov/2023 17h:2m:38s - NIfTI conversion SKIPPED FOR ALL SUBJECTS

--- Initial choices: ---
Home directory = /Users/     /Desktop/Neptune/fully_processed_data/
Siemens scanner = 1; Philips scanner = 0
7T scanner = 1; 3T scanner = 0
Cervical spine = 1; thoracic/lumbar spine = 0
Resting-state fMRI = 1; task fMRI = 0
Pre-process step-after-step = 1; pre-process subject-after-subject = 0
Pre-processing = 1; Post-processing = 1
Number of subjects = 2
Runs per subject = 11
```

Neptune also exports the execution log to the webpage for convenient browsing.

### 4. Understanding Neptune's outputs

We first describe the output files and filenames in section 4.1 and present QC outputs in 4.2.

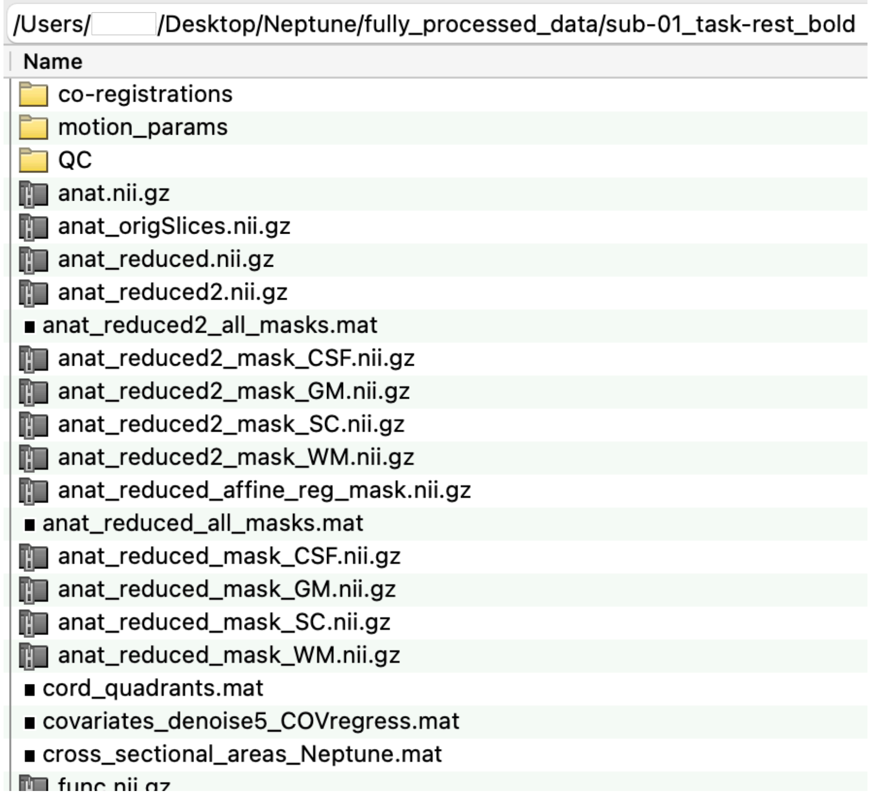


Each subject's fully processed directory typically contains three subdirectories and dozens of files. We describe the files first by using this subject's outputs as an example. For brevity, for the purpose of this section alone, anatomical files (*sub-01_ses-1_T2w**) were renamed as *anat**, and functional files (*sub-01_ses-1_task-rest_bold**) were renamed as *func**.

(*Screenshot of files continued from the previous figure*)

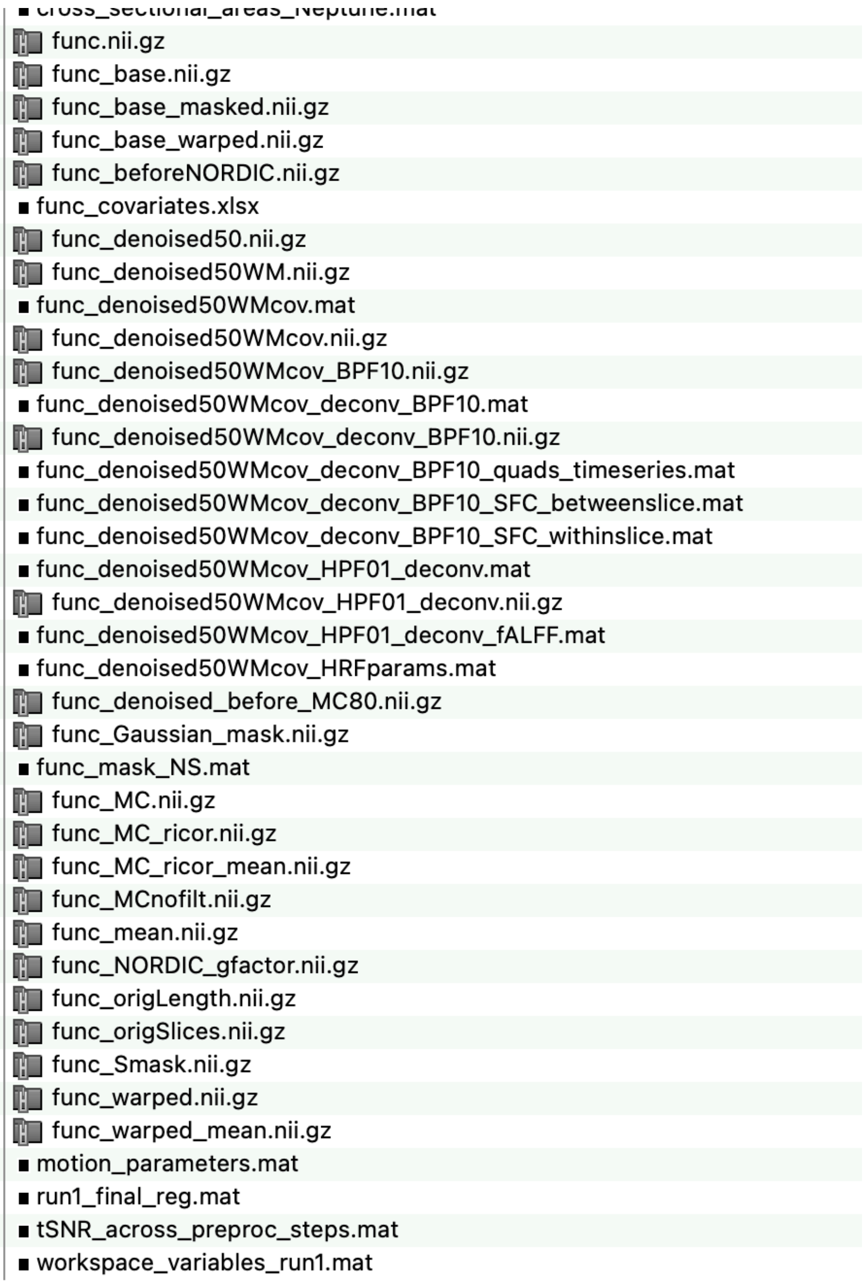

motion_parameters.mat

Slice-wise motion parameters (two translation parameters in the X-Y plane). These are matrices of size N×2 (N = no. of time points; 2 = two translation parameters). It also contains aggregate relative motion (framewise displacement, FD) values (size: #timepoints×#slices), as well as mean and maximum FD in each slice (size: #slices×1) and across all slices (one value). Absolute displacement values are also saved (size: #timepoints×#slices). Since motion correction is done slice-wise, we have only *x* and *y* translations, and in-plane rotation is poorly characterized and thus set to zero. All values are in millimeters.

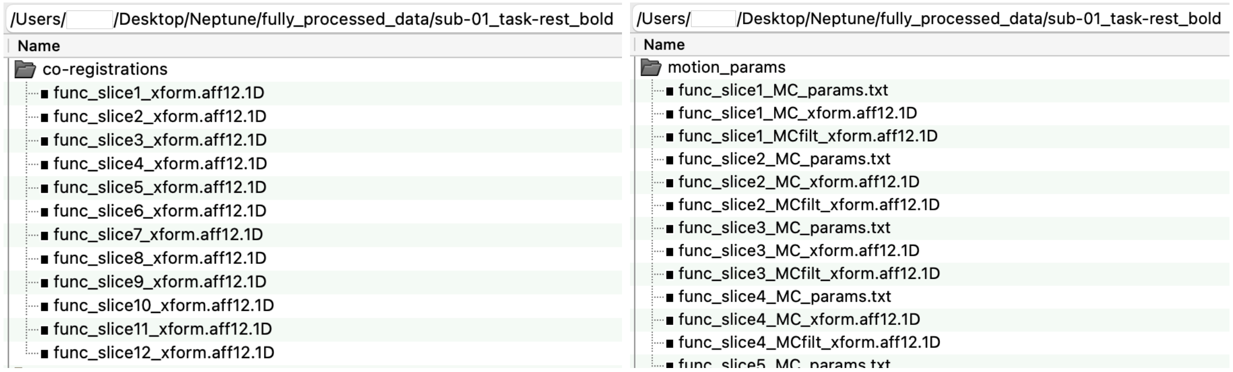

### 4.2. Neptune's QC outputs

Neptune saves QC outputs inside the *QC* subdirectory.

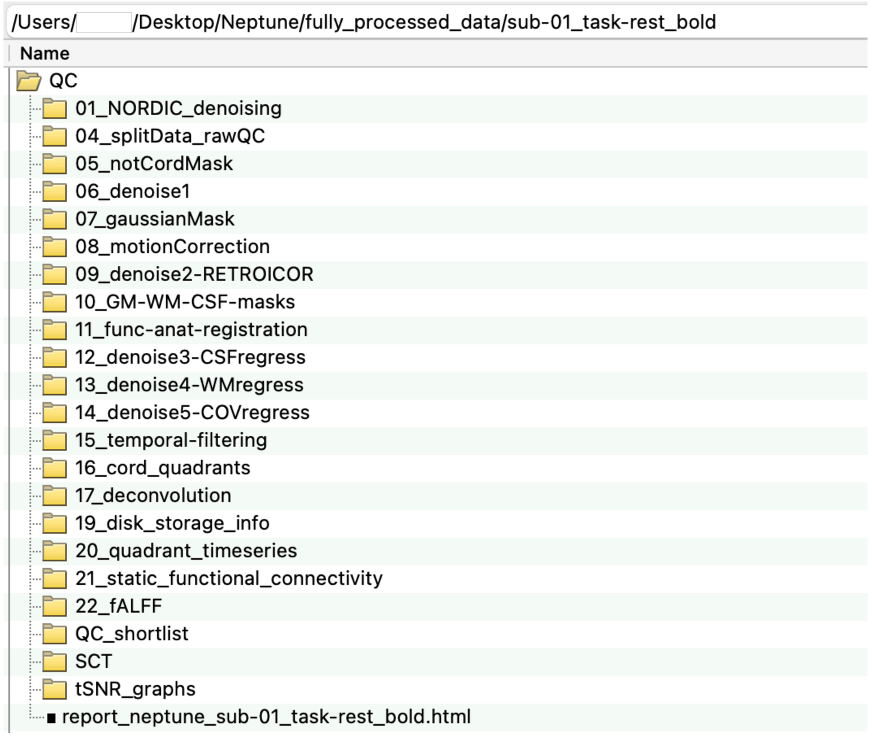


There are subdirectories within *QC* that correspond to the processing steps performed. Apart from the master webpage, Neptune also generates a webpage for each subject (*report_neptune_*.html*) where the given subject's QC outputs can be browsed. Alternatively, users can browse each subdirectory for the QC *.jpg* and *.mp4* files.

Each QC subdirectory contains automatically generated figures corresponding to each slice (with the suffix '*_slice##.jpg*'). Figures generated from data averaged across all slices are also provided ('*_allSlices.jpg*'). This is handy for two reasons. Firstly, they neatly summarize common patterns across all slices. It is often the case that some slices are bad (noisy, distorted), and there is noticeable variability in data quality and outcomes across slices exclusively driven by noise sources rather than neural information. Average information across all slices can highlight both consistent neural patterns prevalent across the FOV and significant effects of noise/distortion spread across all slices. This is meaningful because the data are processed slice-wise, and unlike the brain, there is inherent anatomical and functional similarity across axial slices (or spinal levels) in the spinal cord, making this operation sensible in the cord but not in the brain. Secondly, hundreds of slice-wise QC figures are generated in each subject, making it tedious to go through all of them in all the subjects, thus perhaps not justifying the cost-benefit ratio in terms of time and effort. Our general practice has been to go through all QC figures generated from data averaged across all slices, and delve into slice-wise figures only if we notice something unusual. Of course, exceptions exist, but this practice has been effective for us. Given this background, Neptune has a separate subdirectory (inside *QC*) called *QC_shortlist* that contains only the most essential QC outputs from data averaged across all slices. This should be the first stop for users to browse QC figures. We explain these shortlisted outputs next. In this section, corresponding figures are shown *below* their heading and description.

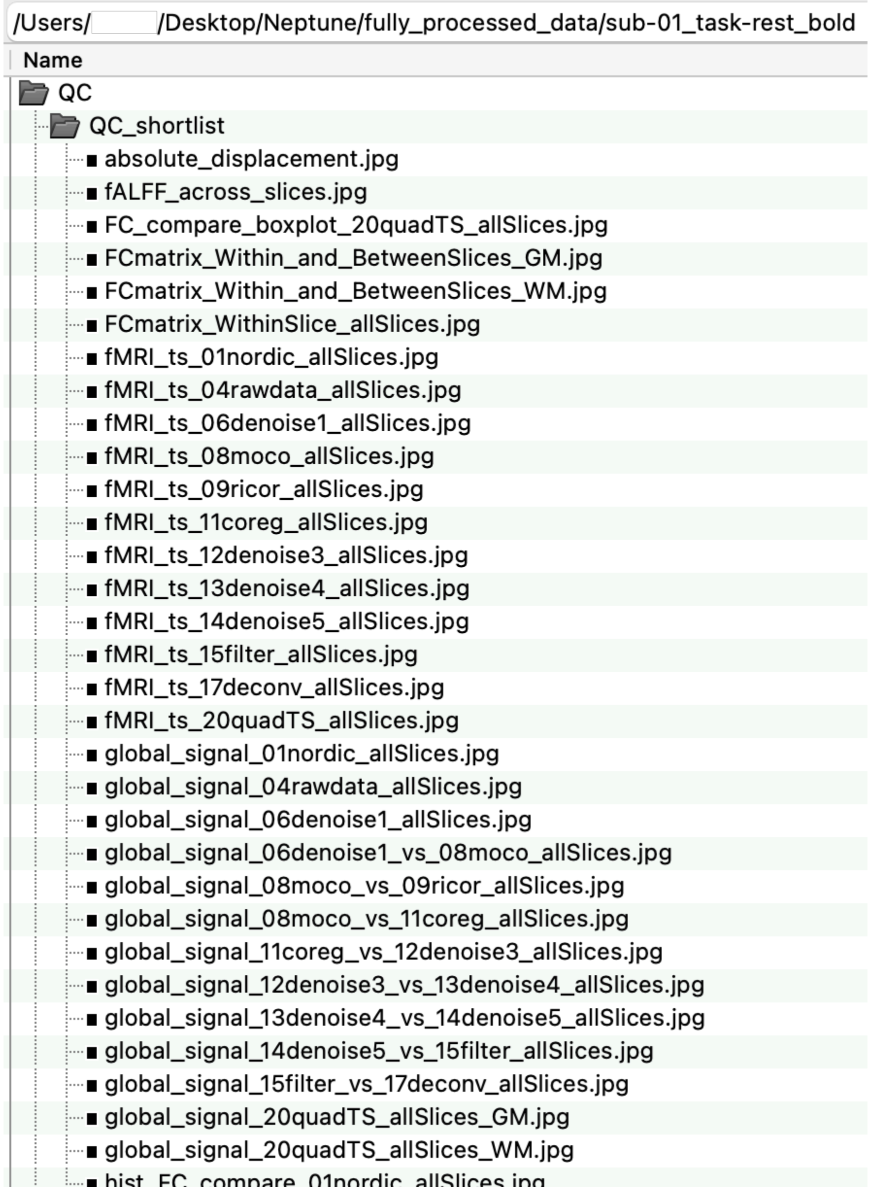

fMRI_ts_*_allSlices.jpg

These are carpet plots generated by pooling fMRI time series from all voxels within the mask (cord mask or anatomical tissues) and displaying the 2D matrix (#voxels×#time) as an image. The voxels are sorted in descending order of percentage signal change. These help identify systematic sources of noise that appear as vertical bands. Below, we show these plots for all processing steps, one after another, which also gives an idea of how much noise is removed in each step. The filenames and titles embedded in the figures are self-explanatory.

fMRI_ts_04rawdata_allSlices.jpg

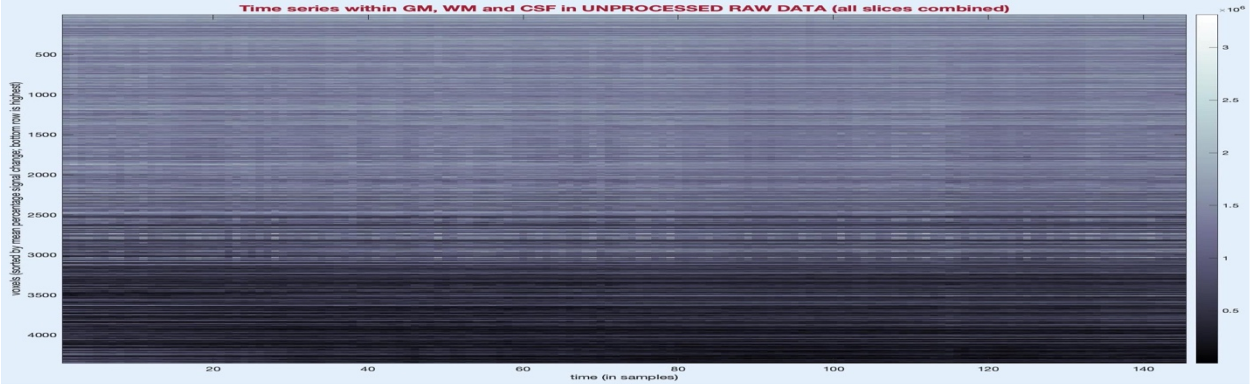


fMRI_ts_06denoise1_allSlices.jpg

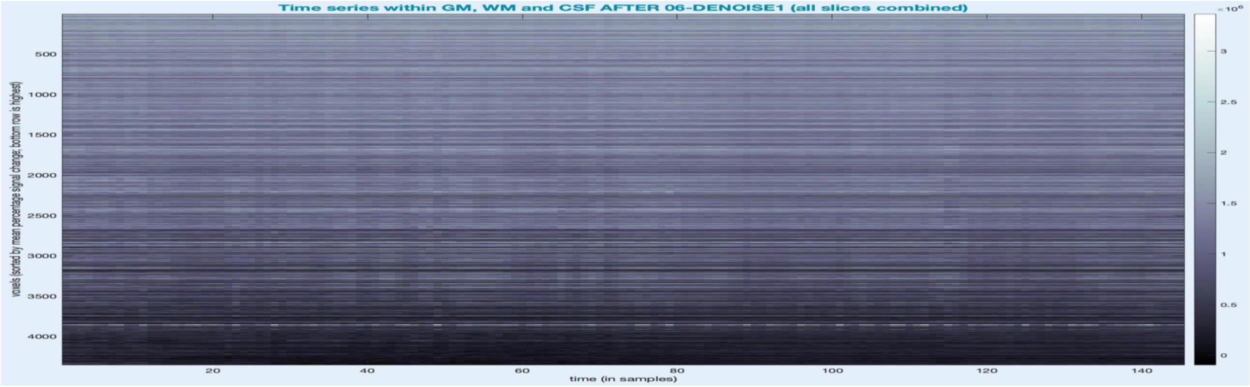


fMRI_ts_08moco_allSlices.jpg

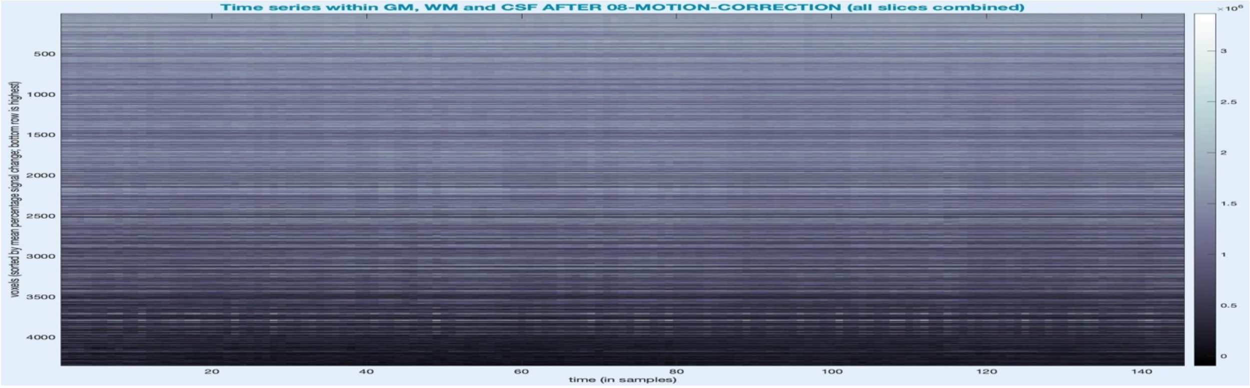

fMRI_ts_09ricor_allSlices.jpg

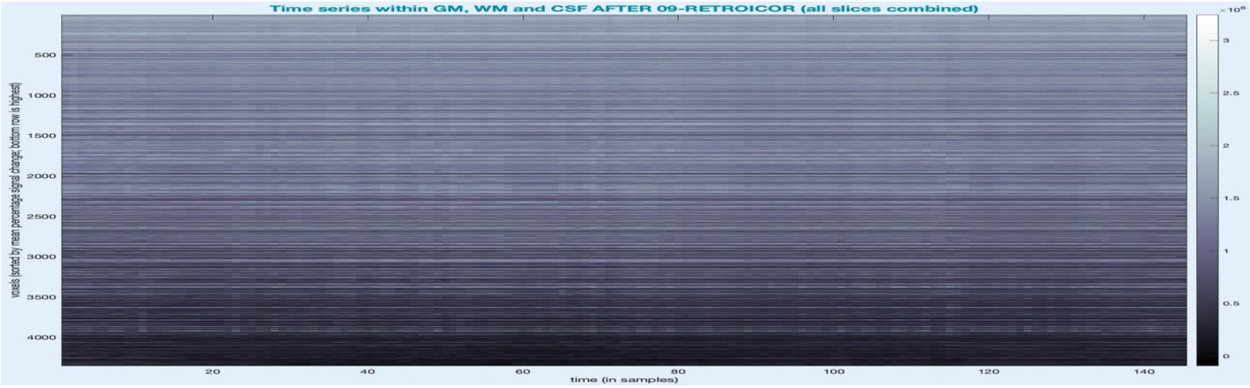


fMRI_ts_11coreg_allSlices.jpg

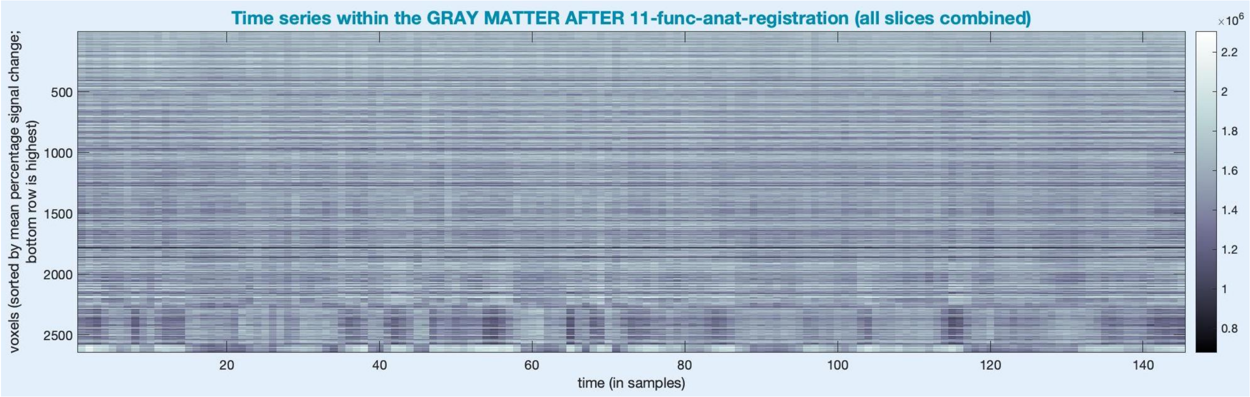


fMRI_ts_12denoise3_allSlices.jpg

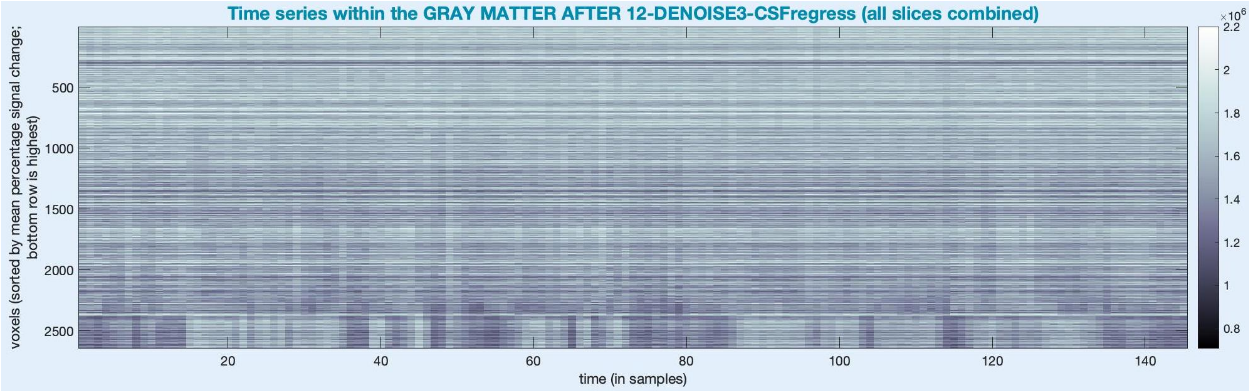

fMRI_ts_13denoise4_allSlices.jpg

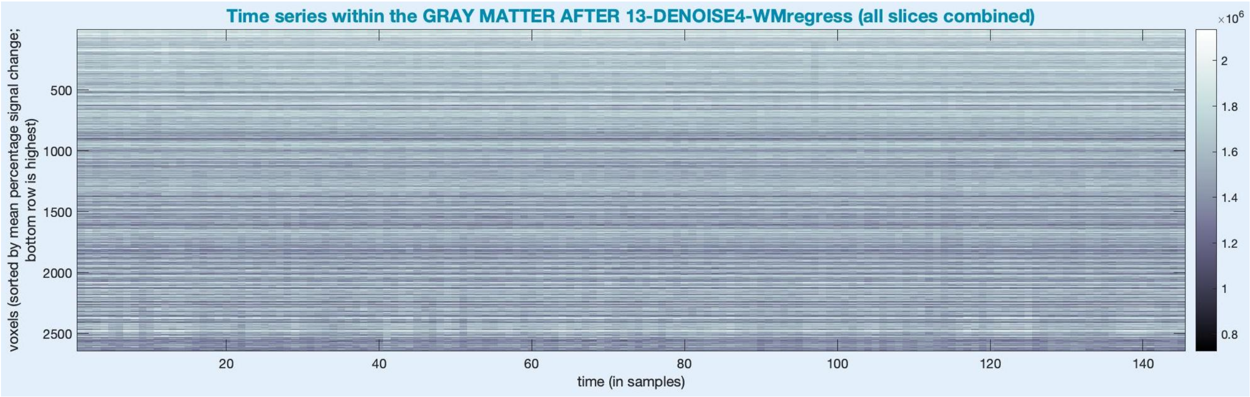


fMRI_ts_14denoise5_allSlices.jpg

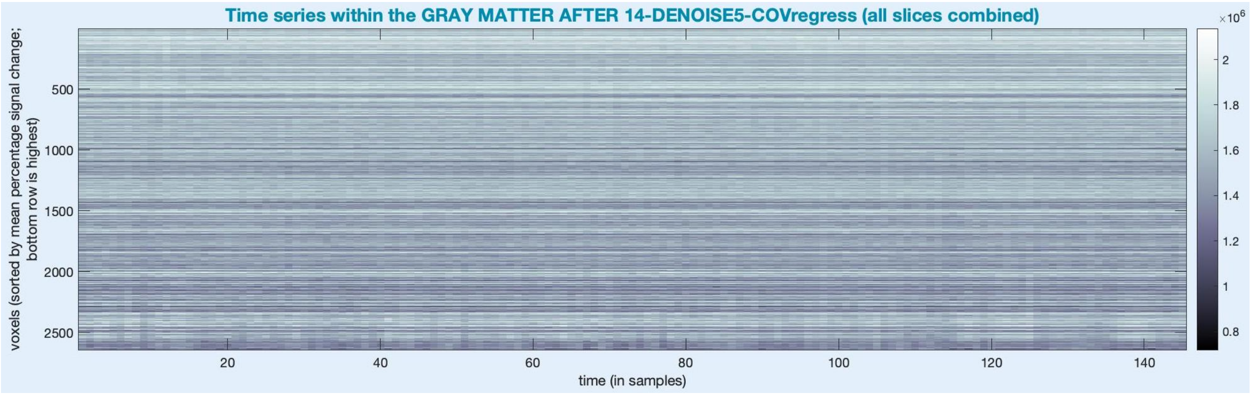


fMRI_ts_15filter_allSlices.jpg

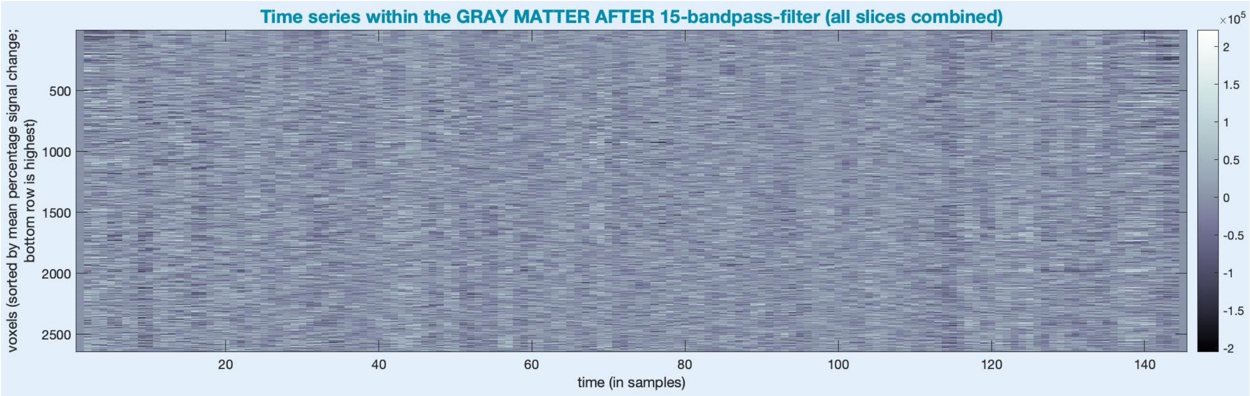

fMRI_ts_17deconv_allSlices.jpg

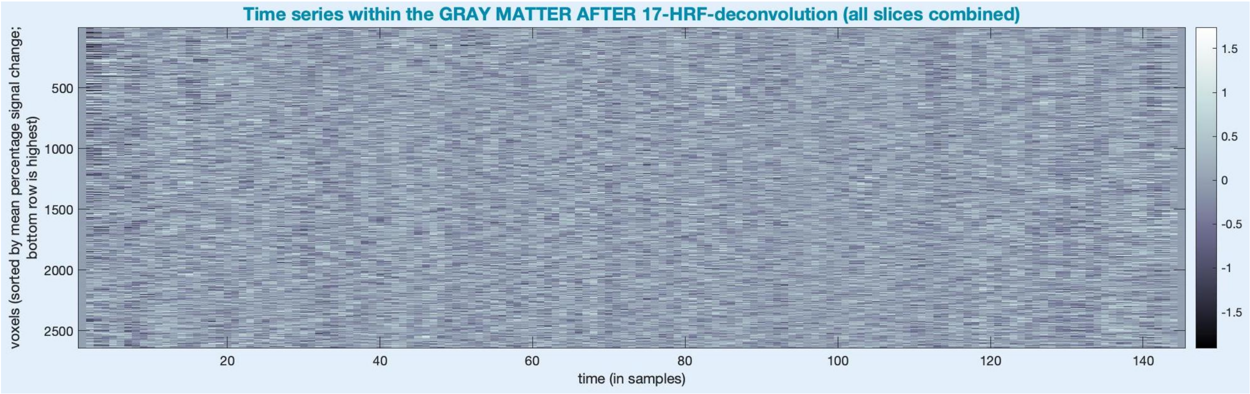


fMRI_ts_20quadTS_allSlices.jpg

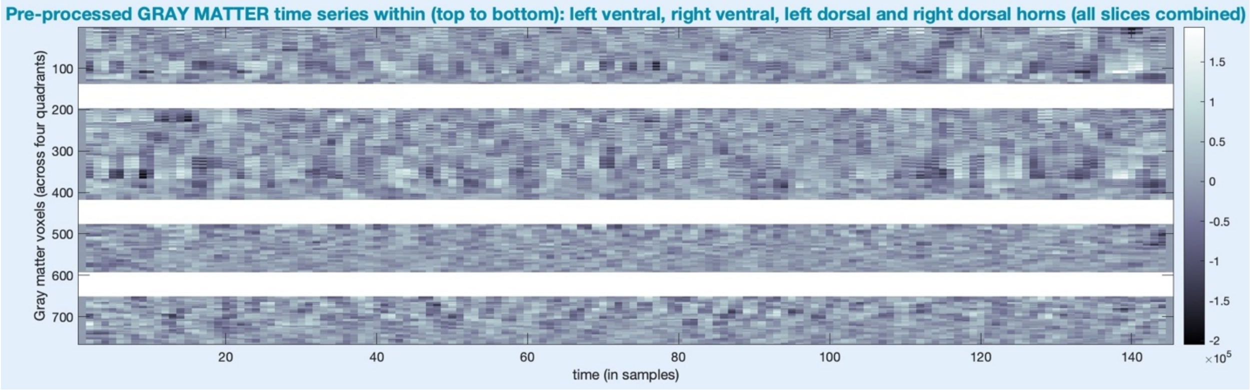


<u>global_signal_*_allSlices.jpg</u>

All voxel-level time series within a mask (e.g., cord or GM) are averaged and compared with those generated in the previous step. For example, below, the top figure shows unprocessed data in thick red and data after step-6 (not-cord regression) in a thin blue line. The bottom figure shows the same with the former in thin red and the latter in thick blue line, giving a clear visual picture.

global_signal_06denoise1_allSlices.jpg

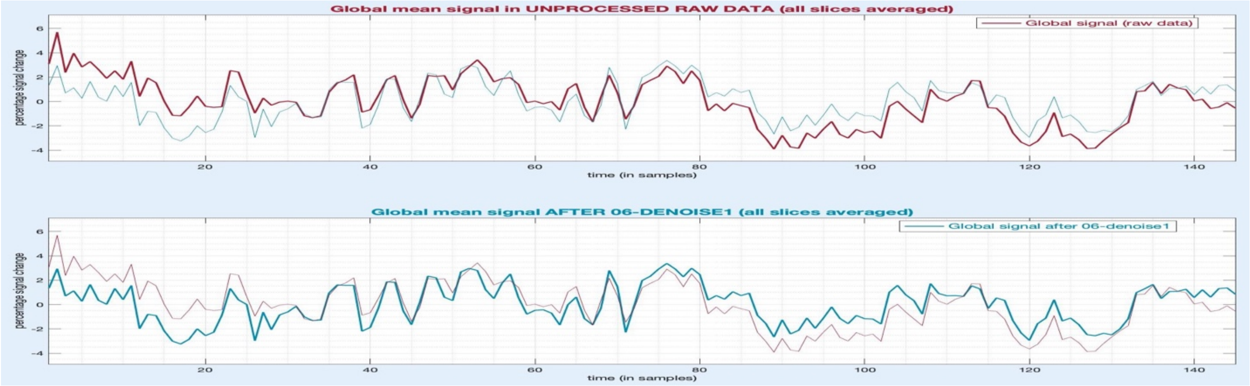

global_signal_06denoise1_vs_08moco_allSlices.jpg

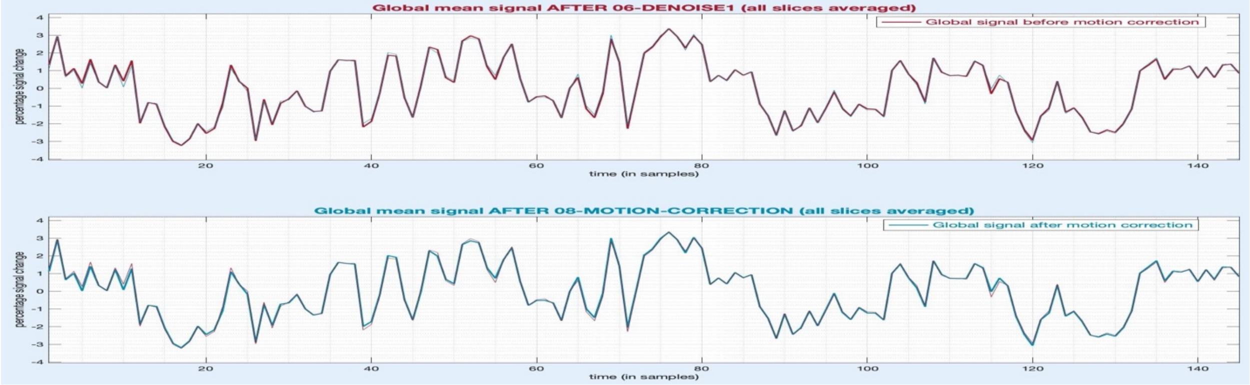


global_signal_08moco_vs_09ricor_allSlices.jpg

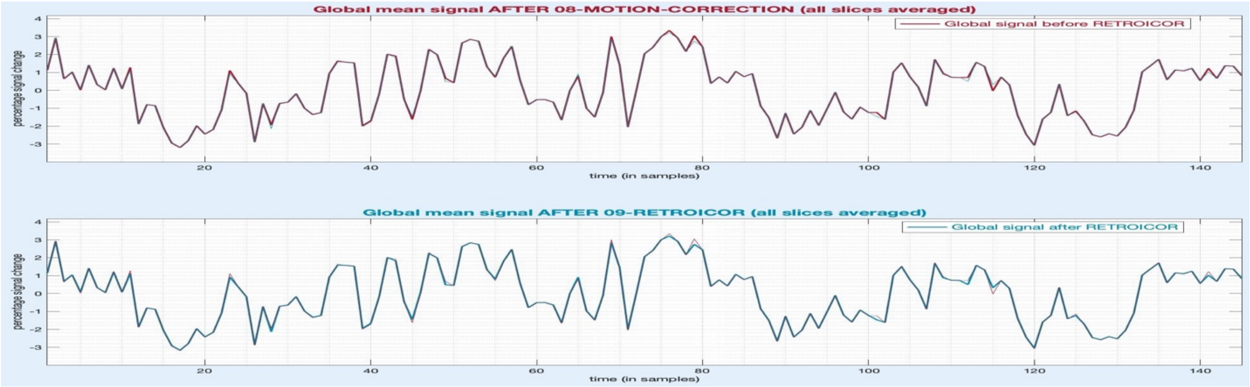


global_signal_08moco_vs_11coreg_allSlices.jpg

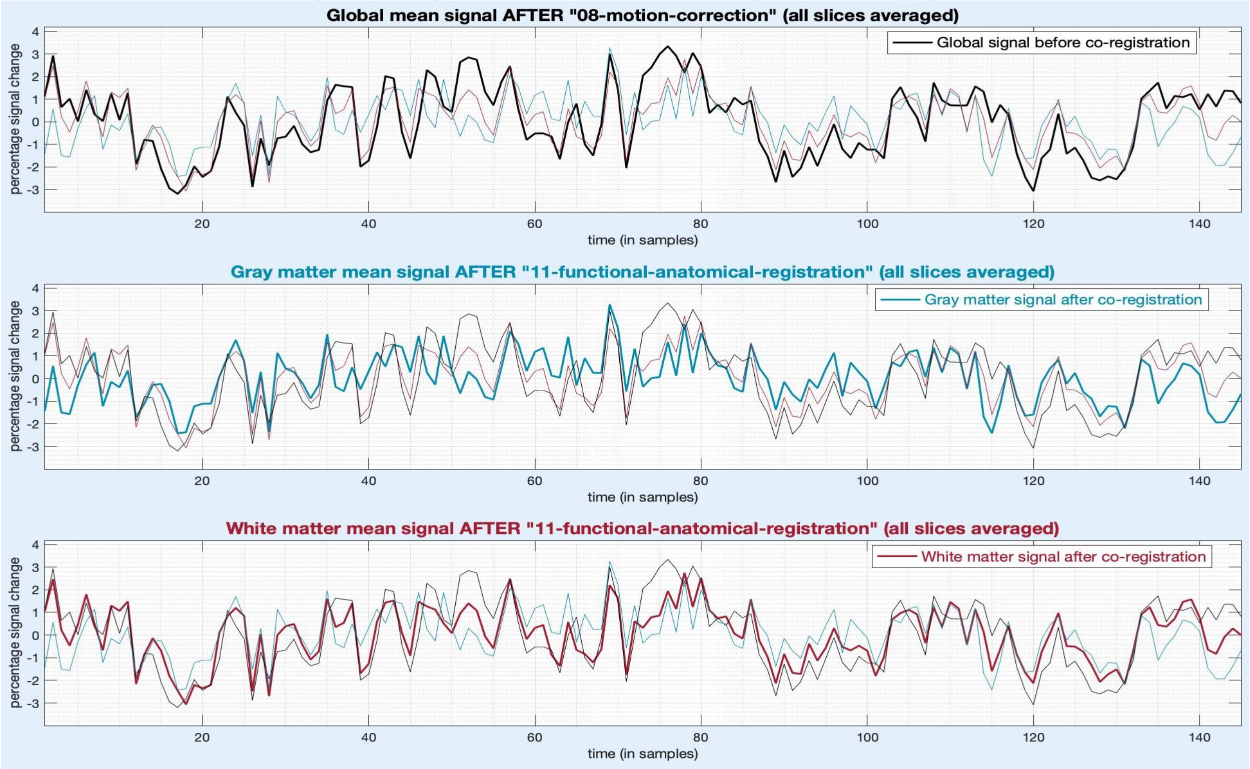

global_signal_11coreg_vs_12denoise3_allSlices.jpg

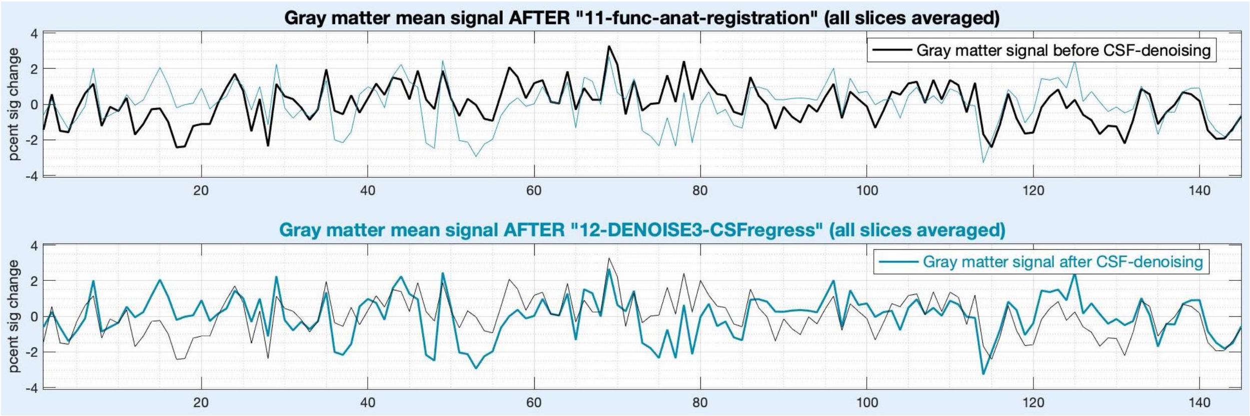


global_signal_12denoise3_vs_13denoise4_allSlices.jpg

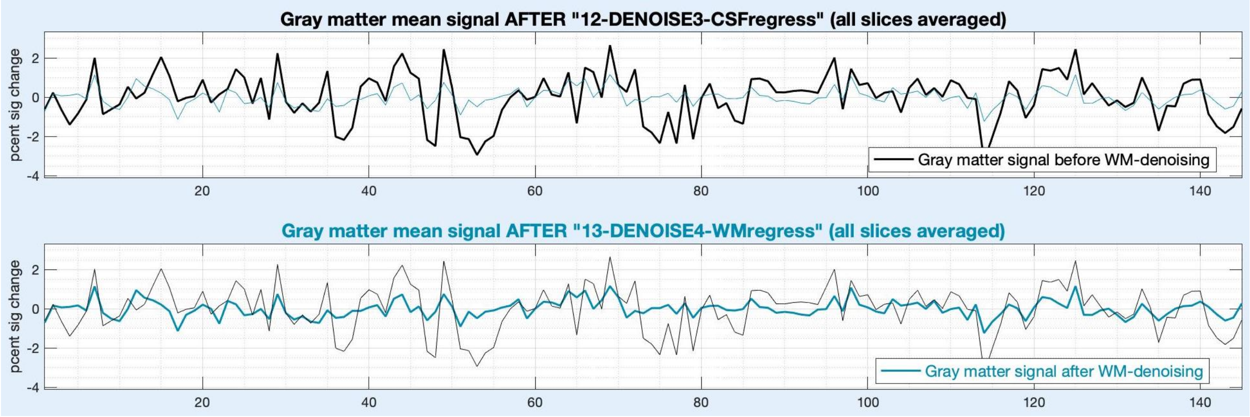


global_signal_13denoise4_vs_14denoise5_allSlices.jpg

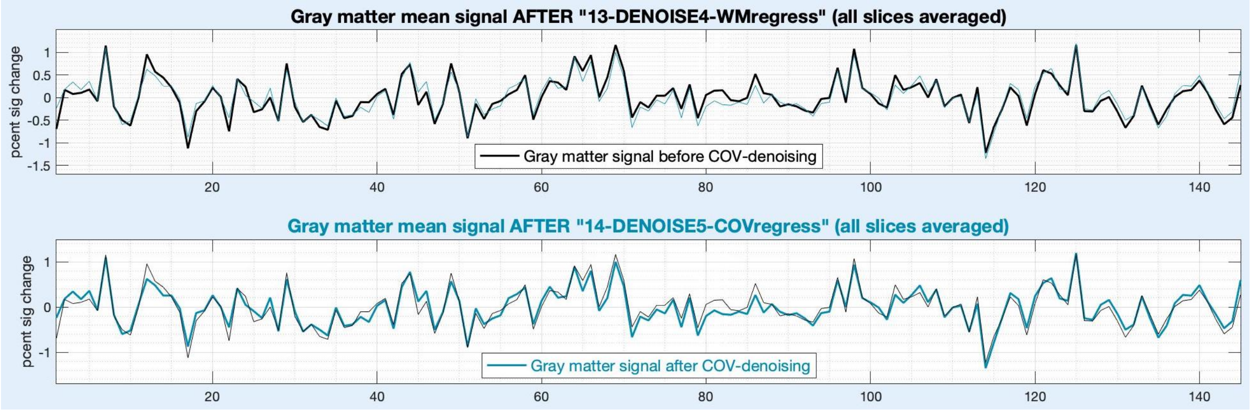

global_signal_14denoise5_vs_15filter_allSlices.jpg

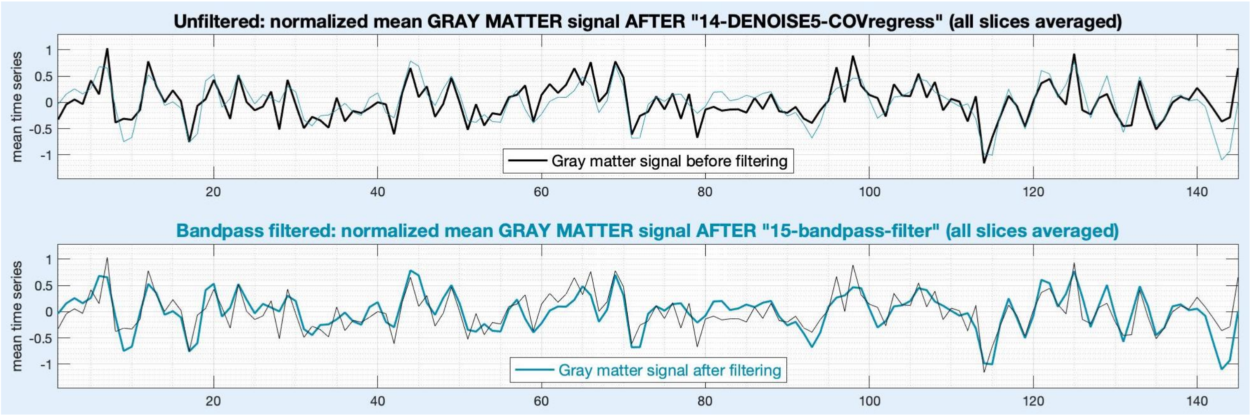


global_signal_15filter_vs_17deconv_allSlices.jpg

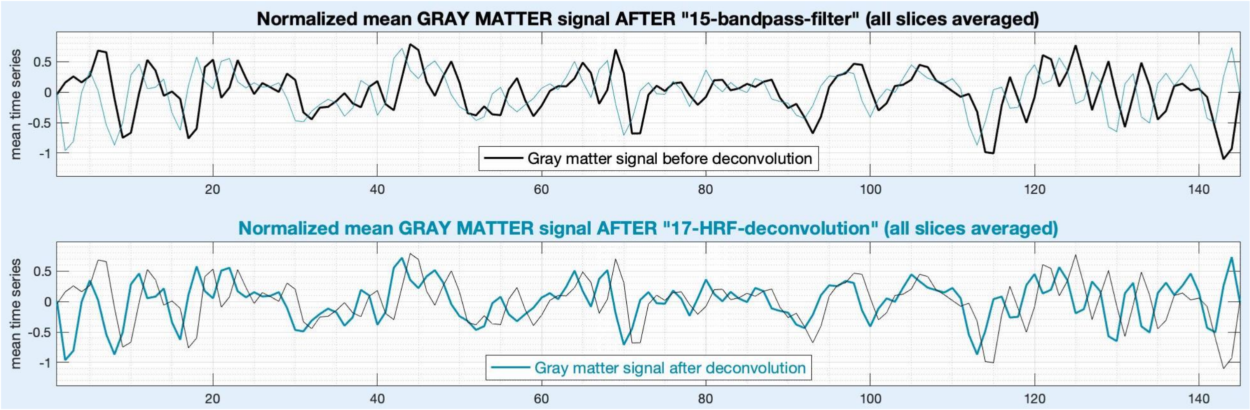


It must be noted that our sample data had a sampling time (volume acquisition time) of 3.34 seconds, which is insufficient to reliably estimate the HRF and conduct deconvolution (appropriate literature is cited in the main manuscript). We, however, performed deconvolution on this sample data merely to generate QC figures for the purpose of visualization in this document. Hence, it is not intended that any conclusions be drawn from figures related to the HRF or deconvolution in this document.

mean_timeseries_GM_20quadTS_allSlices.jpg

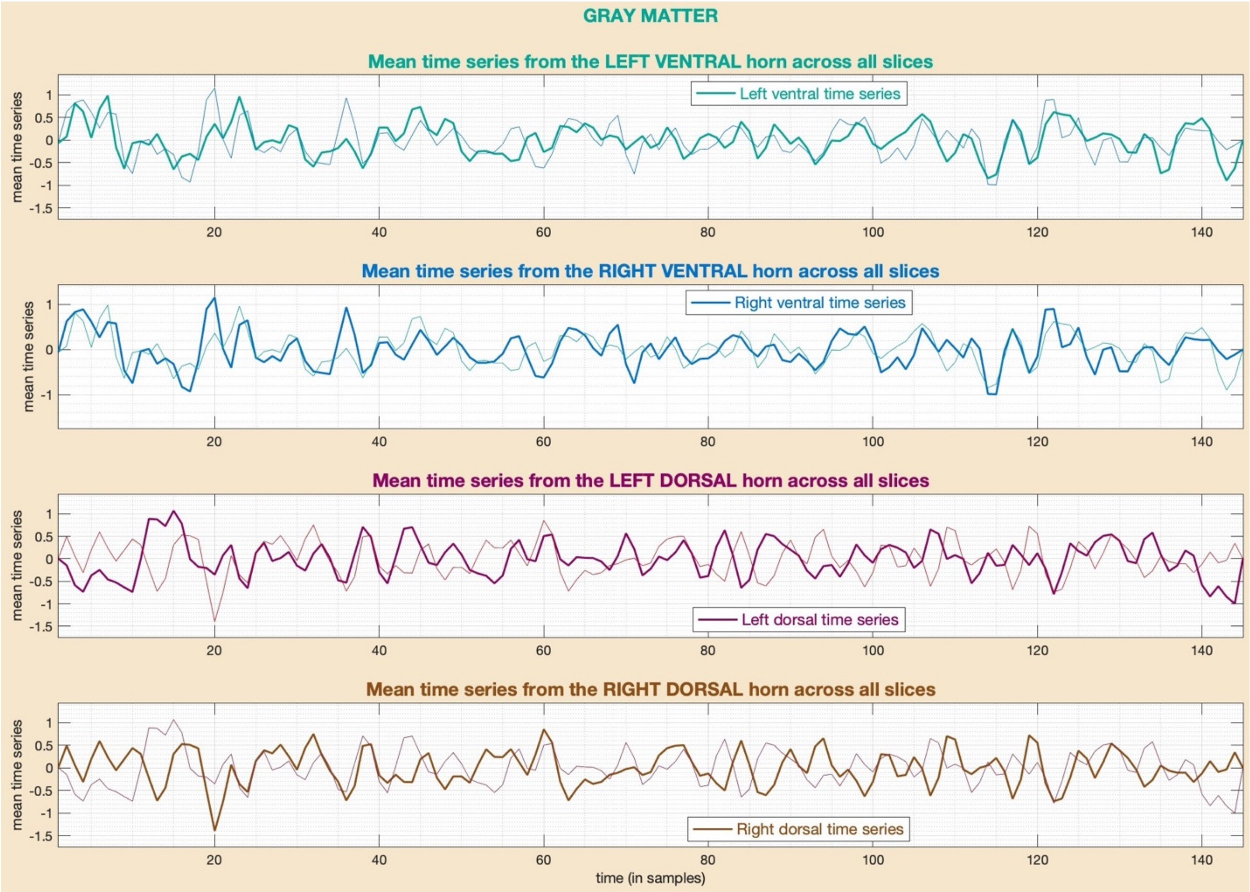


The above figure is slightly different from the others. It is the only mean time series plot generated during post-processing (as part of step-20). It shows the mean time series (averaged across all slices) within each of the four quadrants. Since we expect to find high connectivity between the left and right ventral horns (as well as the left and right dorsal horns), this figure is a visual aid to assess these trends in the data. In this figure, it is evident that the left/right ventral horn time series' have a decent overlap (the top two figures), and the case is the same with the dorsal horns (the bottom two figures).

(*Screenshot of files continued from the earlier figure*)

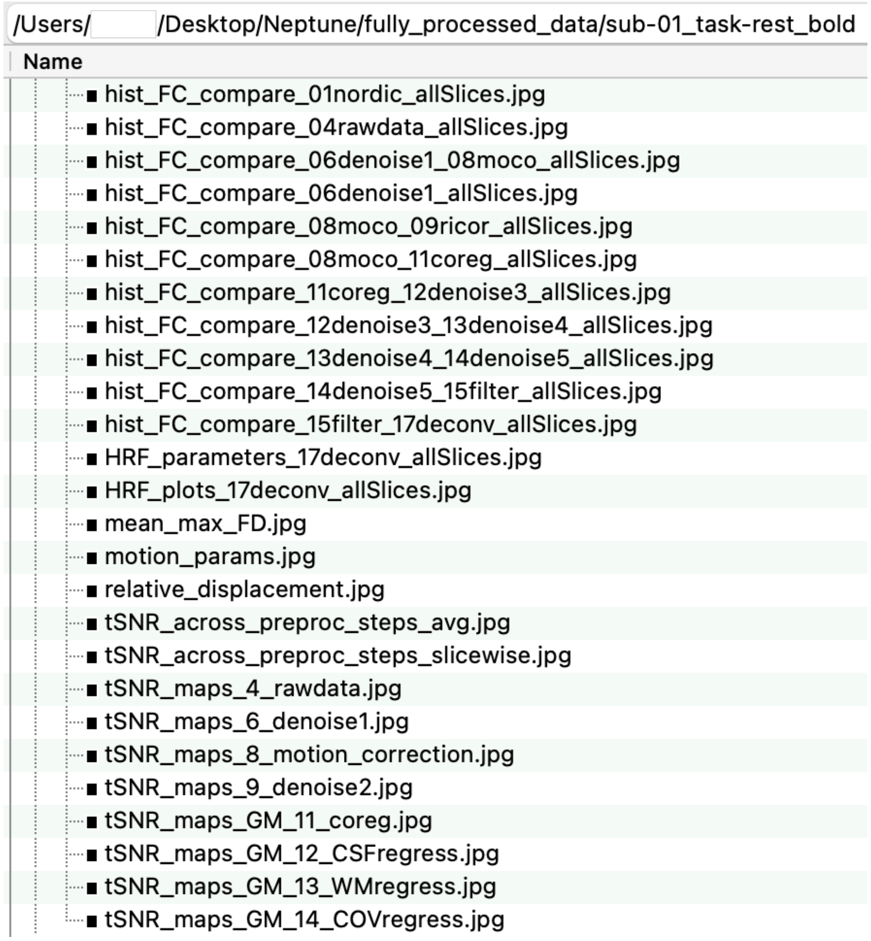


<u>hist_FC_compare_*_allSlices.jpg</u>

Neptune computes voxel-to-voxel correlations between all pairs of voxels within the mask, and generates a probability distribution of these values (i.e., the histogram divided by the number of correlations). It then plots a comparison of these across successive steps, one below the other.

hist_FC_compare_06denoise1_0allSlices.jpg

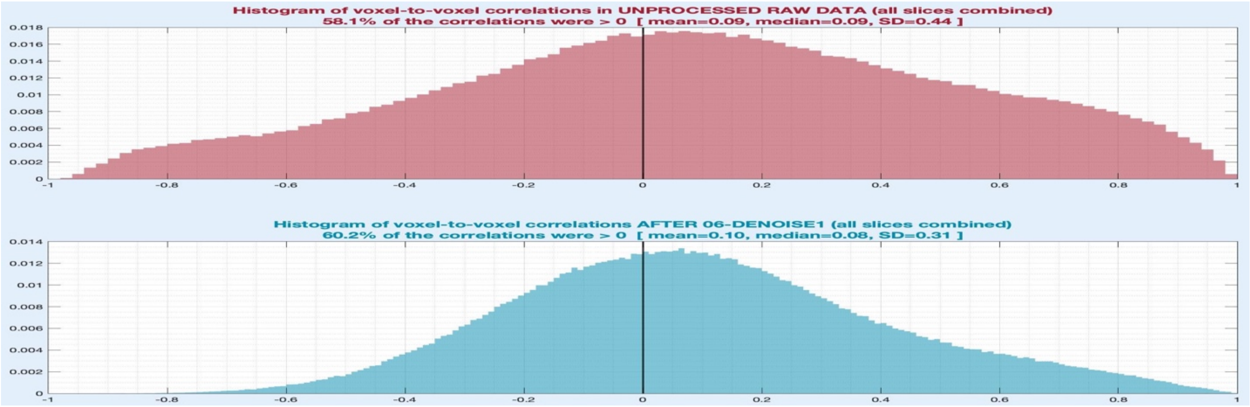

hist_FC_compare_06denoise1_08moco_allSlices.jpg

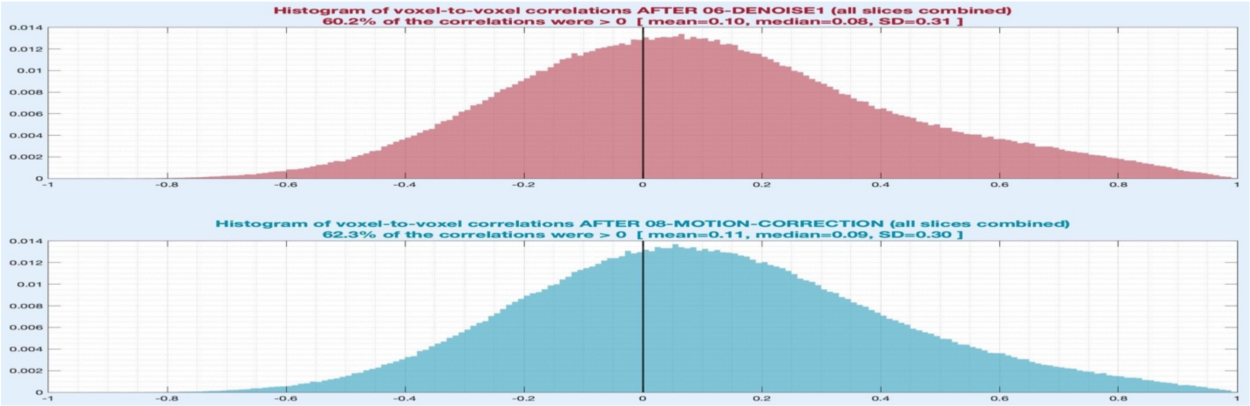


hist_FC_compare_08moco_09ricor_allSlices.jpg

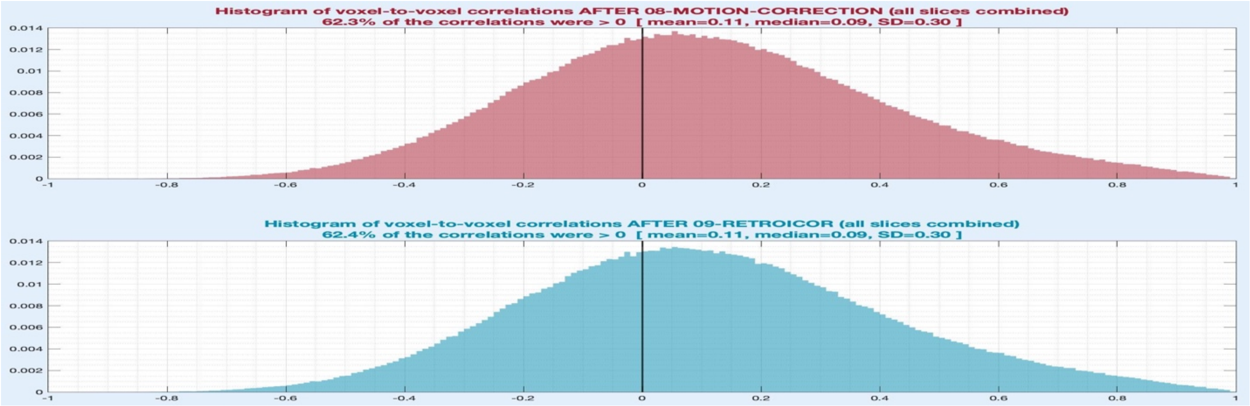


Wider histograms (i.e., those with larger standard deviations) indicate systematic, spatially distributed noise in the data. This is because the same noise source spread across different voxels induces highly correlated (or sometimes anticorrelated) voxels. This also tends to shift the histogram to the right (higher mean value). Thus, successive denoising steps are expected to lower the mean, median and standard deviation of the histogram. Any type of smoothing operation, however, will invariably widen the histogram, as can be observed after step-11 (co-registration) that performs spatial interpolation, and after step-15 (temporal filtering) that blurs the data in the time domain.

hist_FC_compare_08moco_11coreg_allSlices.jpg

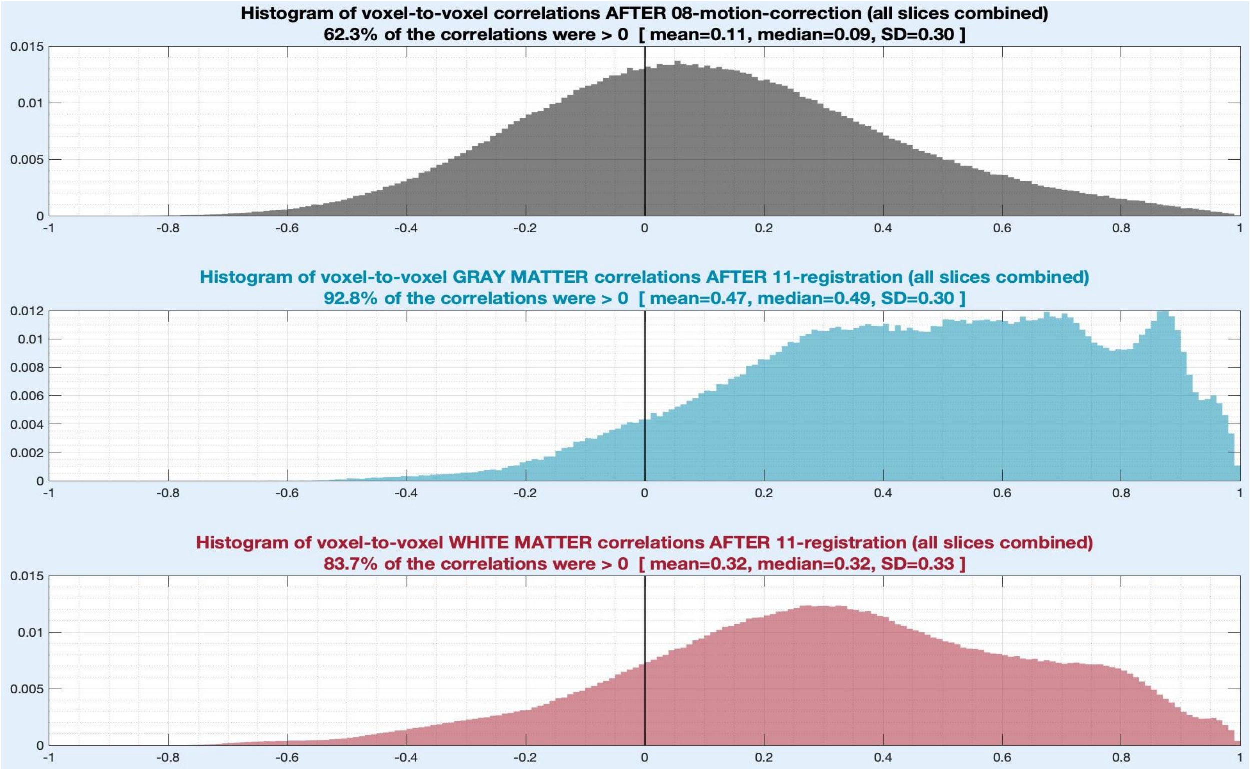


hist_FC_compare_11coreg_12denoise3_allSlices.jpg

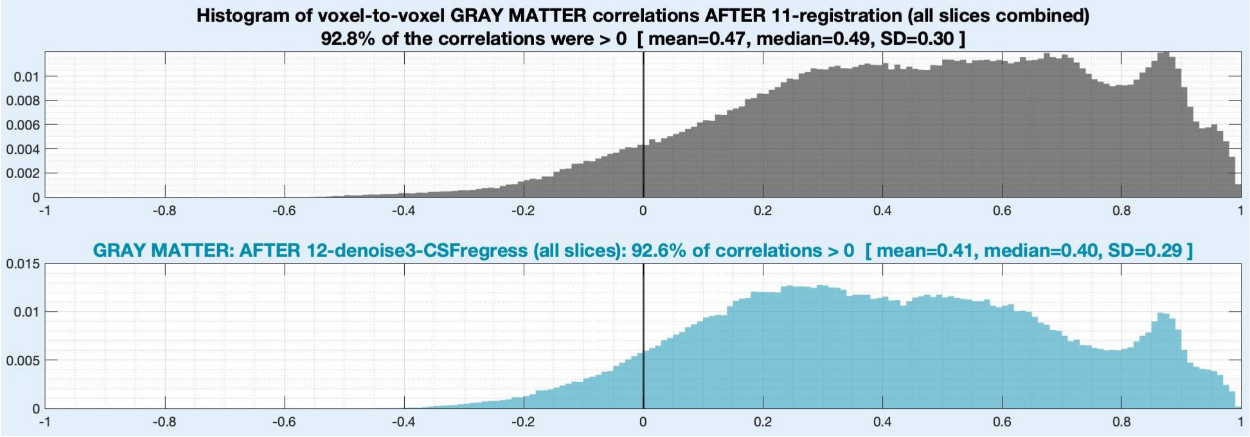

hist_FC_compare_12denoise3_13denoise4_allSlices.jpg

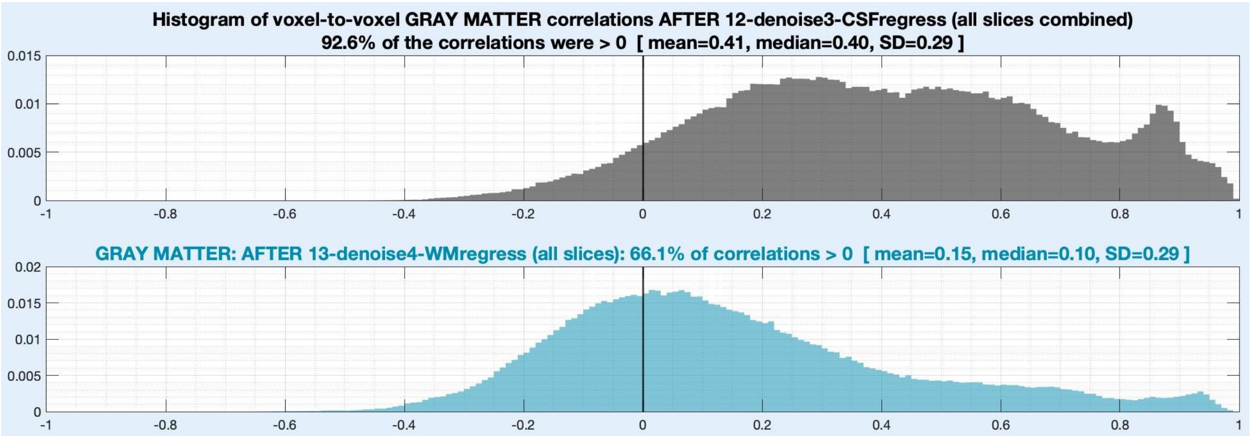


hist_FC_compare_13denoise4_14denoise5_allSlices.jpg

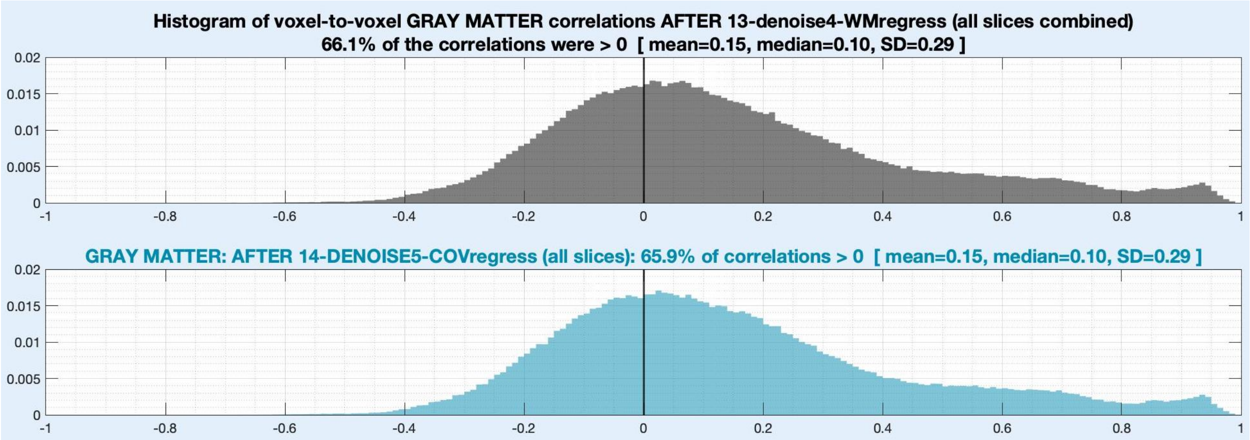


hist_FC_compare_14denoise5_15filter_allSlices.jpg

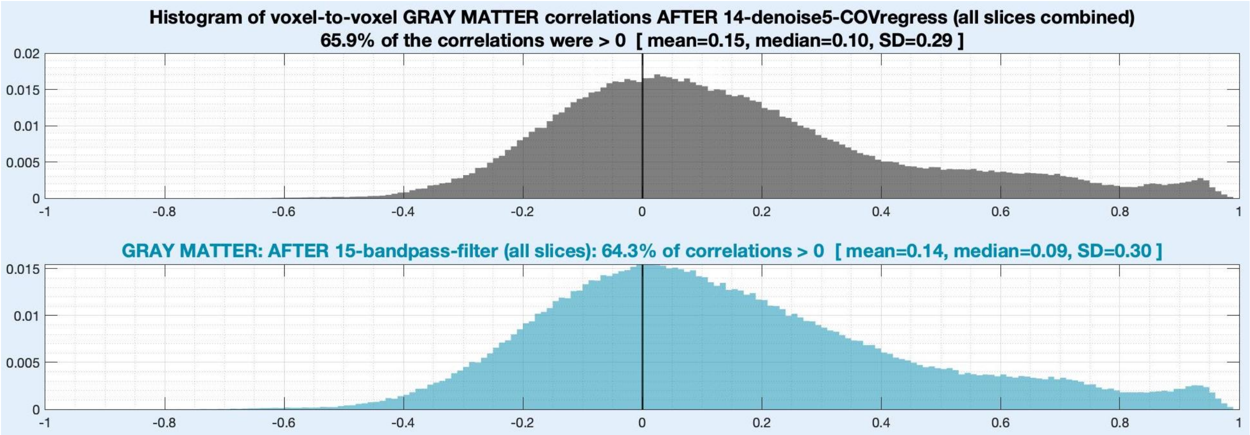

hist_FC_compare_15filter_17deconv_allSlices.jpg

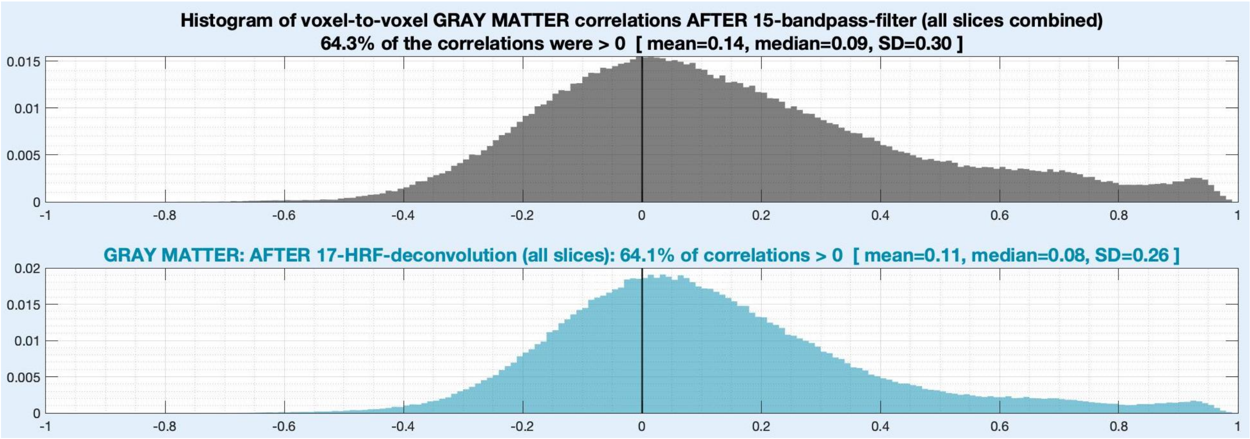


tSNR_across_preproc_steps_avg.jpg

Neptune computes tSNR from raw data as well as across denoising steps. The jumps in tSNR after a step indicate the amount of variance removed (or the noise minimized) from the time series by that step. This is because tSNR is the ratio of the signal mean to standard deviation, and a decrease in the latter results in a tSNR increase. Blurring operations (such as step-11 involving spatial interpolation) invariably cause a jump in tSNR values even though they do not involve explicit noise removal.

Neptune computes the mean tSNR value from all voxels and slices within a mask and displays them to the user to assess data quality and denoising performance. Since the cord mask is obtained in step-5 itself, Neptune computes tSNR in the whole cord as defined by the cord mask (GM+WM+CSF) all the way from steps 6 to 15. Since GM and WM tissue boundaries in the anatomical space are obtained only in step-10, tSNR within GM (or WM) is computed only from steps 11 to 15. The top half of the below figure shows tSNR across pre-processing steps in the cord in our sample data, and the bottom half shows the same within GM and WM. When step-13 (WM signal denoising) is chosen, GM tSNR tends to be higher because more variance is removed from GM but none from WM during this step.

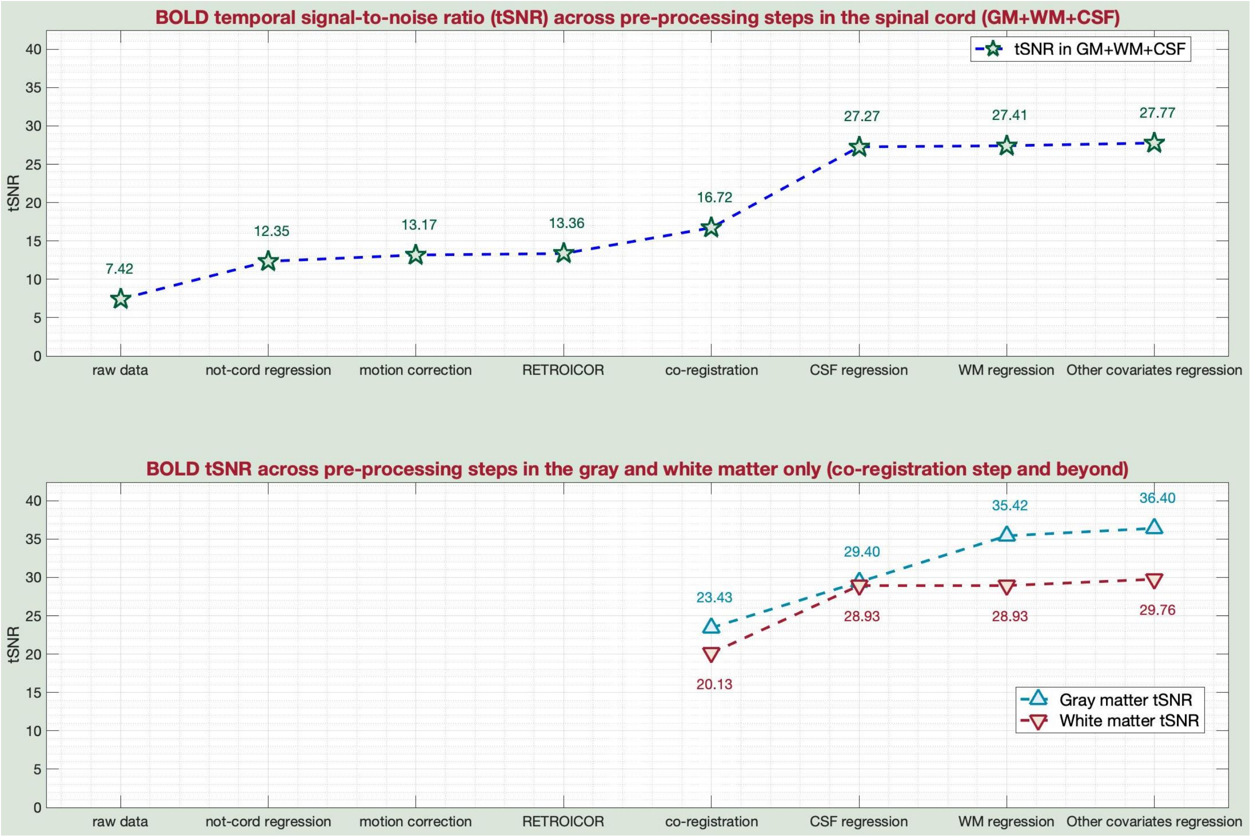


<u>tSNR_across_preproc_steps_slicewise.jpg</u>

Neptune also computes tSNR values within each slice in raw data and across all denoising steps, which are available in '*tSNR_across_preproc_steps.mat*'. For simplicity, it plots tSNR across slices only for raw data and at the end of all denoising steps, as shown in the figure below.

The tSNR value typically drops as we move from superior to inferior slices due to anatomy and distance from receive coils, which can be observed in this figure. In some acquisitions, there can be more dramatic decreases, either linearly or abruptly beyond a certain vertebral level. This knowledge can alter what we expect from our data's inferior slices. TSNR across slices can also show abrupt dips only in specific intermediate slices (and not elsewhere), which often happens due to signal dephasing at bone-disc interfaces. Taken together, keenly observing tSNR outputs is highly informative and helpful.

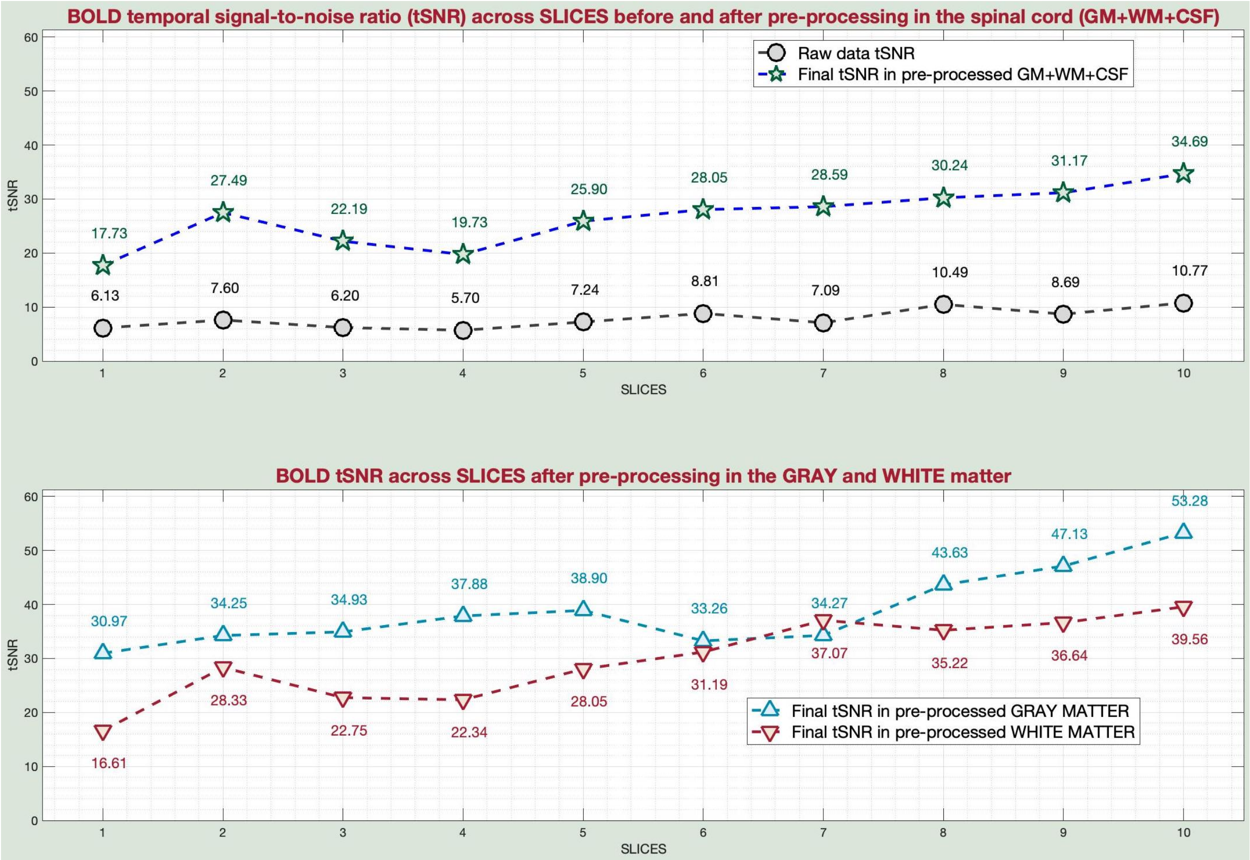


<u>tSNR_maps_*.jpg</u>

Neptune also generates slice-wise tSNR maps from raw data and after each denoising step. It shows us high and low tSNR zones in the data, both within parts of the same slice and between different slices. It informs which parts of the cord are getting significant jumps in tSNR across denoising steps. This is helpful because such information cannot be inferred from mean values or graphs.

tSNR_maps_4_rawdata.jpg

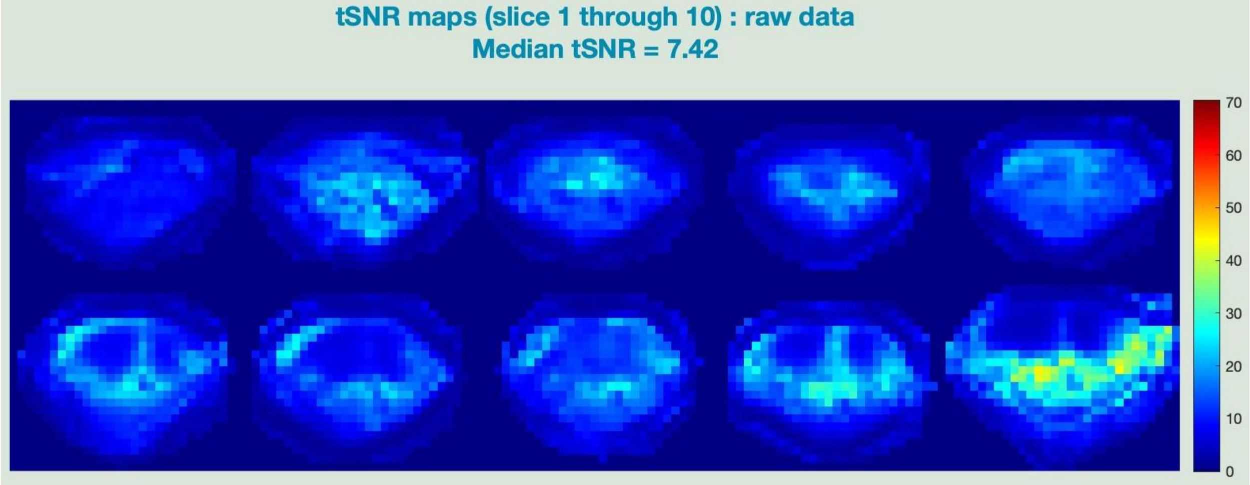

tSNR_maps_6_denoise1.jpg

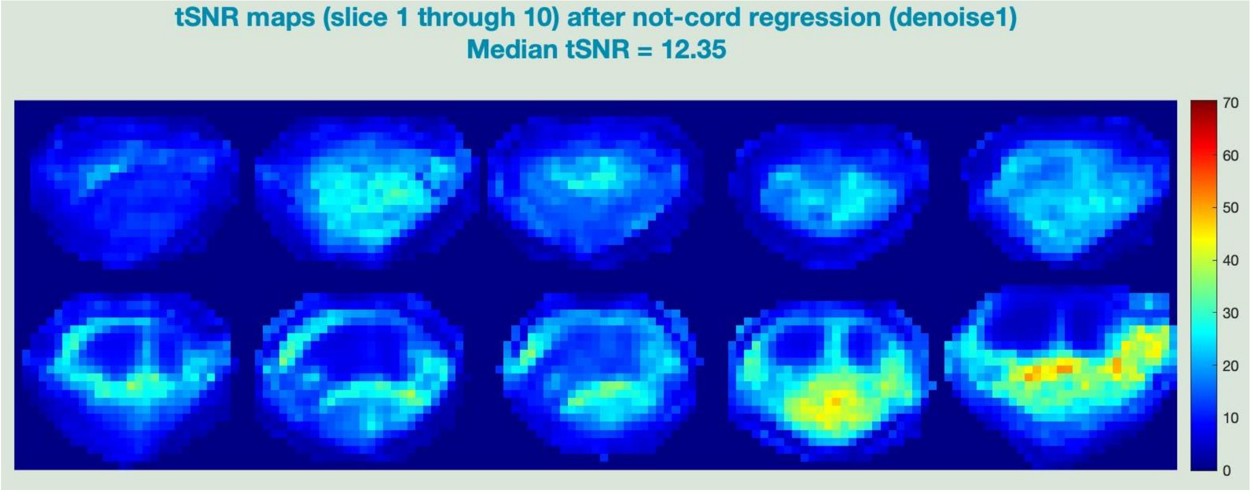


tSNR_maps_8_motion_correction.jpg

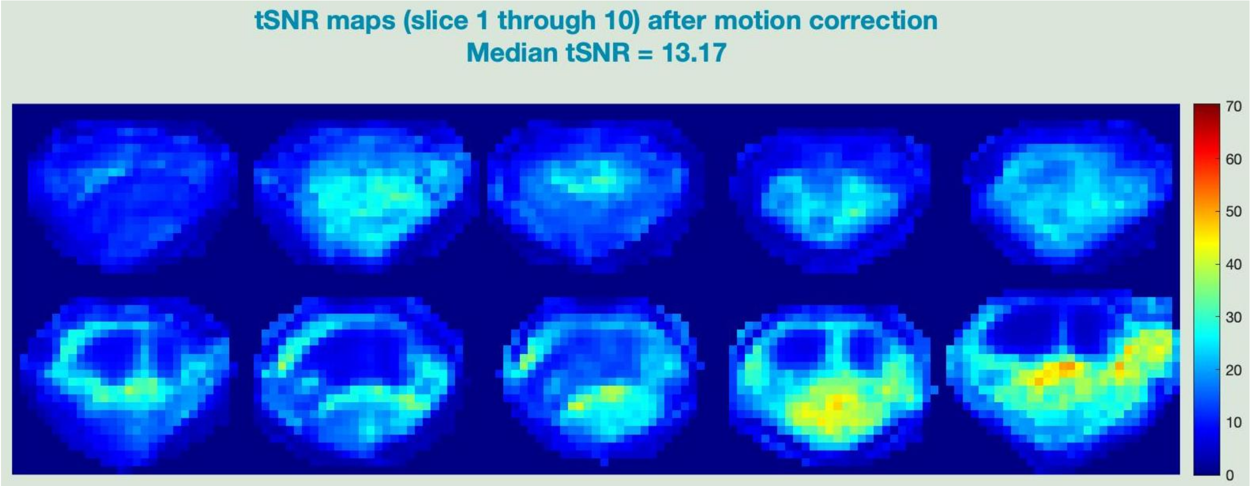


tSNR_maps_9_denoise2.jpg

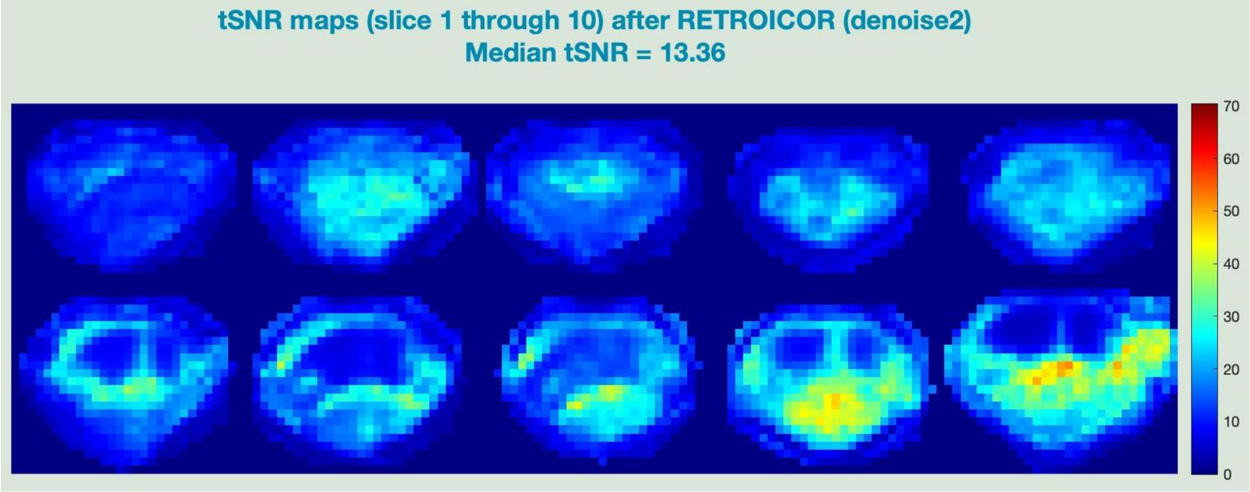

tSNR_maps_GM_11_coreg.jpg

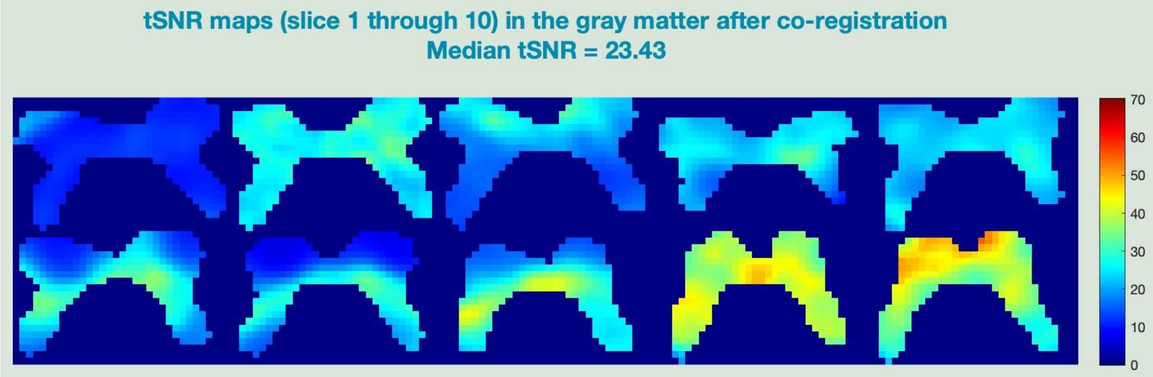


tSNR_maps_GM_12_CSFregress.jpg

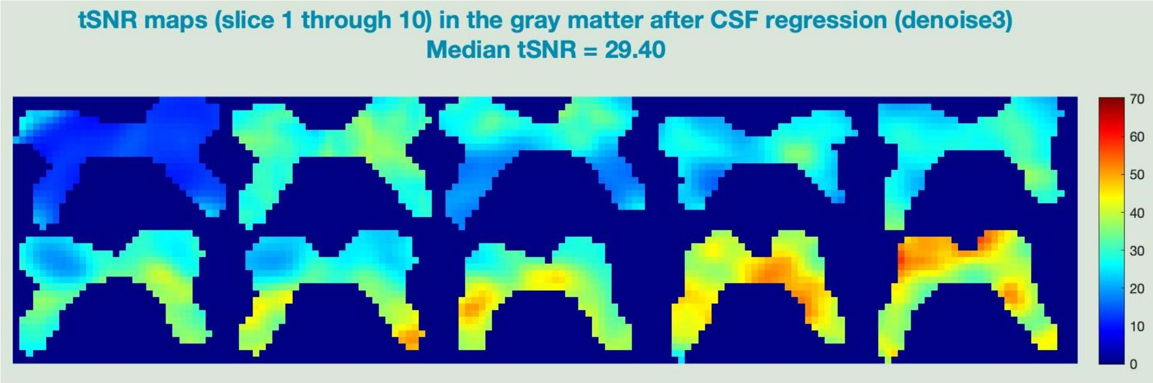


tSNR_maps_GM_13_WMregress.jpg

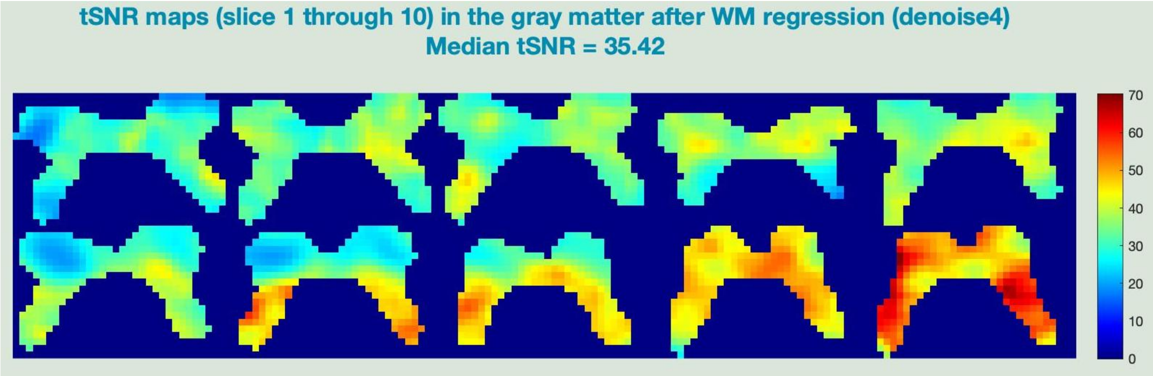


tSNR_maps_GM_14_COVregress.jpg

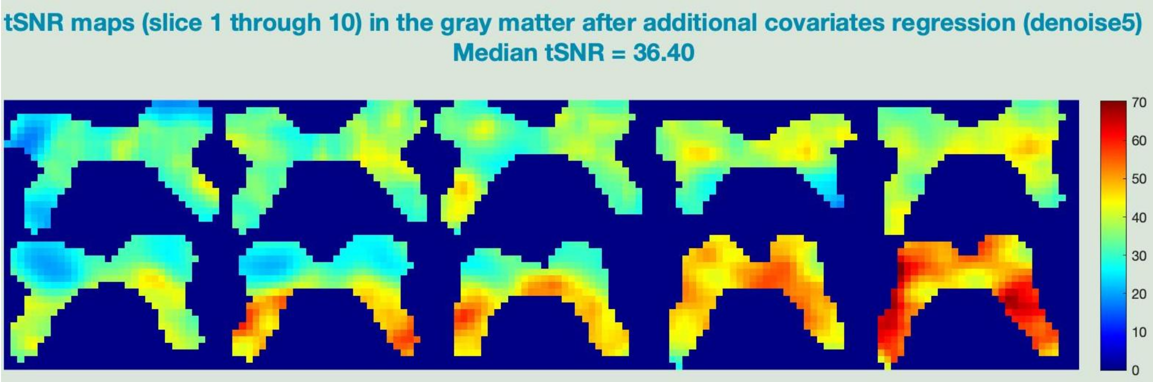

motion_params.jpg

Neptune displays raw *x* and *y* translational motion parameters for each slice to help clearly identify motion spikes and possibly their source. For example, motion spikes caused by swallowing result in larger spikes along the *y* direction (anterior-posterior), while those caused by subject movement tend to cause comparable changes in both the *x* and *y* directions. In this sample data, we do not see such motion spikes.

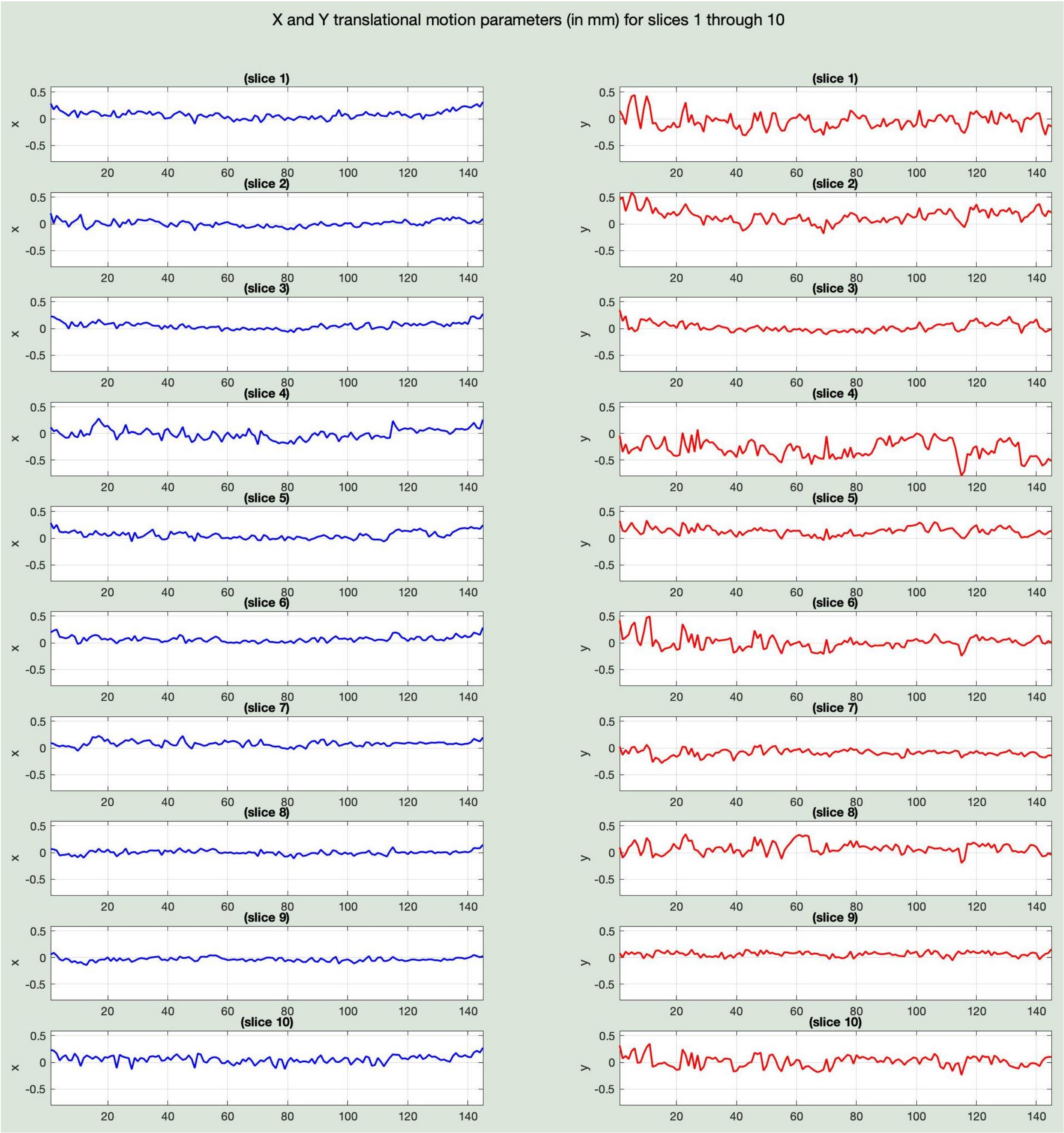

absolute_displacement.jpg and relative_displacement.jpg

Neptune also shows aggregate motion values for each slice – absolute displacement (left) and relative displacement (called framewise displacement, FD) (right). These help detect gradual drifts in subject position and abrupt motion events, respectively.

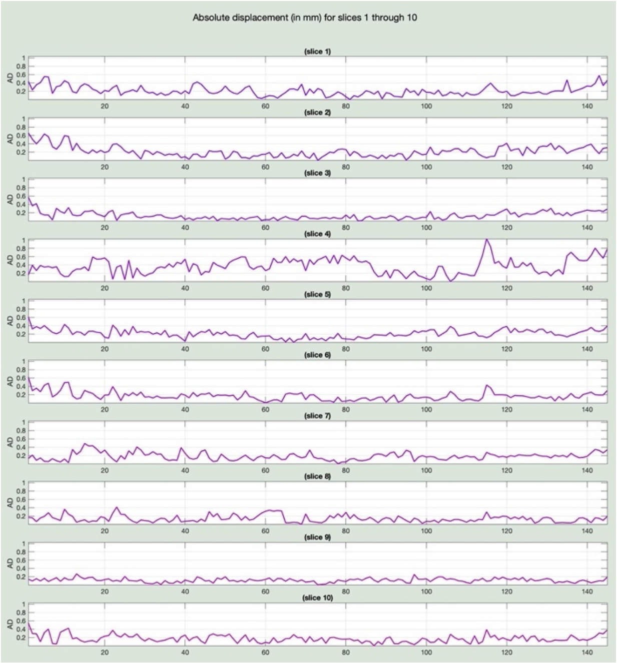


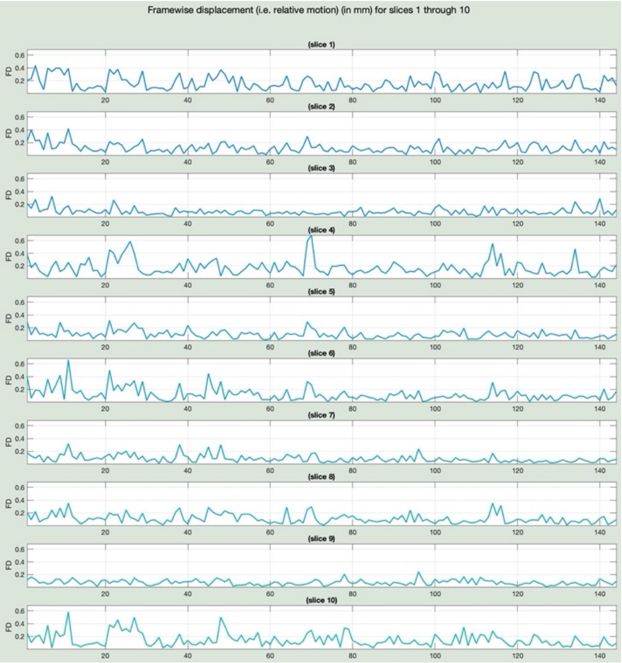


mean_max_FD.jpg

Neptune plots mean FD for each slice (top) as well as maximum FD (bottom). This helps compare the motion profile of different slices and identify problematic slices.

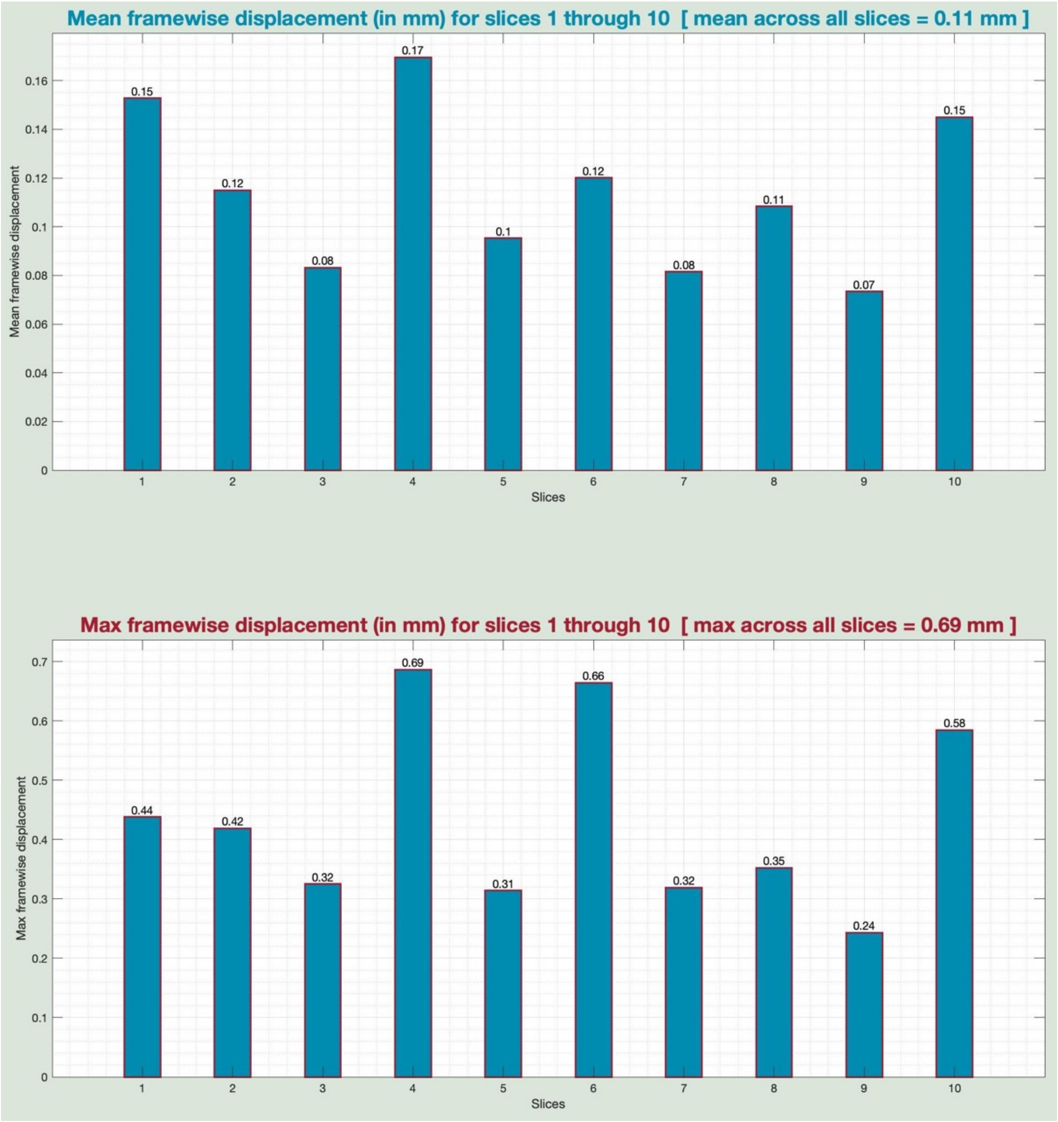


HRF_plots_17deconv_allSlices.jpg

As mentioned before, our HRF outcomes presented here are solely for illustrative purposes because we do not have the necessary temporal resolution. The below figure plots, from top to bottom, the HRF across slices (averaged across all quadrants) (in GM and WM), and the HRF across quadrants (averaged across all slices) (in GM and WM). With sufficient temporal resolution, it is possible to observe HRF differences across slices and quadrants.

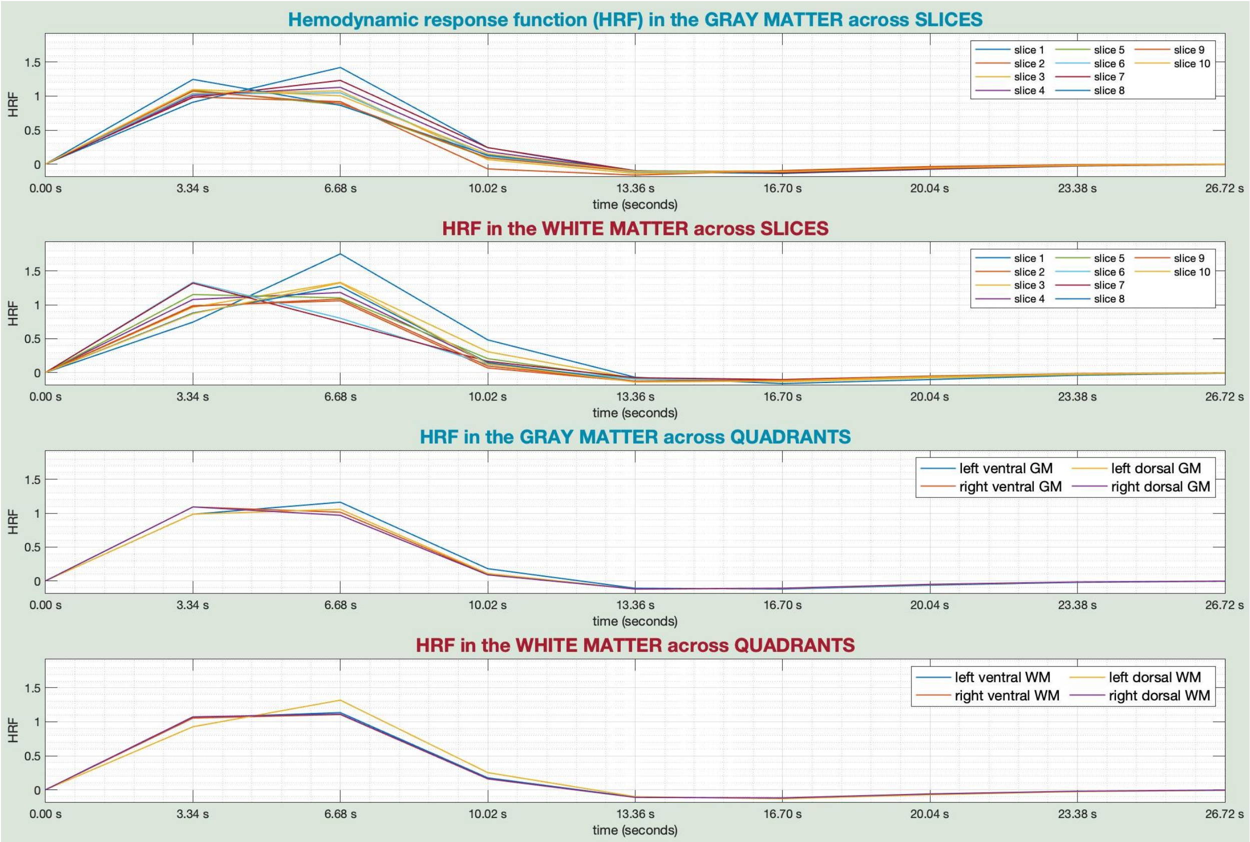


HRF_parameters_17deconv_allSlices.jpg

The below figure shows bar graphs of HRF shape parameters across slices and quadrants. The HRF shape (seen in the above figure) can be characterized by its amplitude (response height), latency (time-to-peak), and duration (full-width at half-max). These are computed from the estimated HRF curve during deconvolution. The left half of the below figure displays HRF parameters in the GM, and the right half displays in the WM. The values shown across slices are averaged across all voxels/quadrants within a given slice. The values shown across quadrants are averaged from voxels across all slices within a given quadrant. The scale of the *y*-axis is adjusted to visibly bring out the differences, which is something to note when viewing this figure.

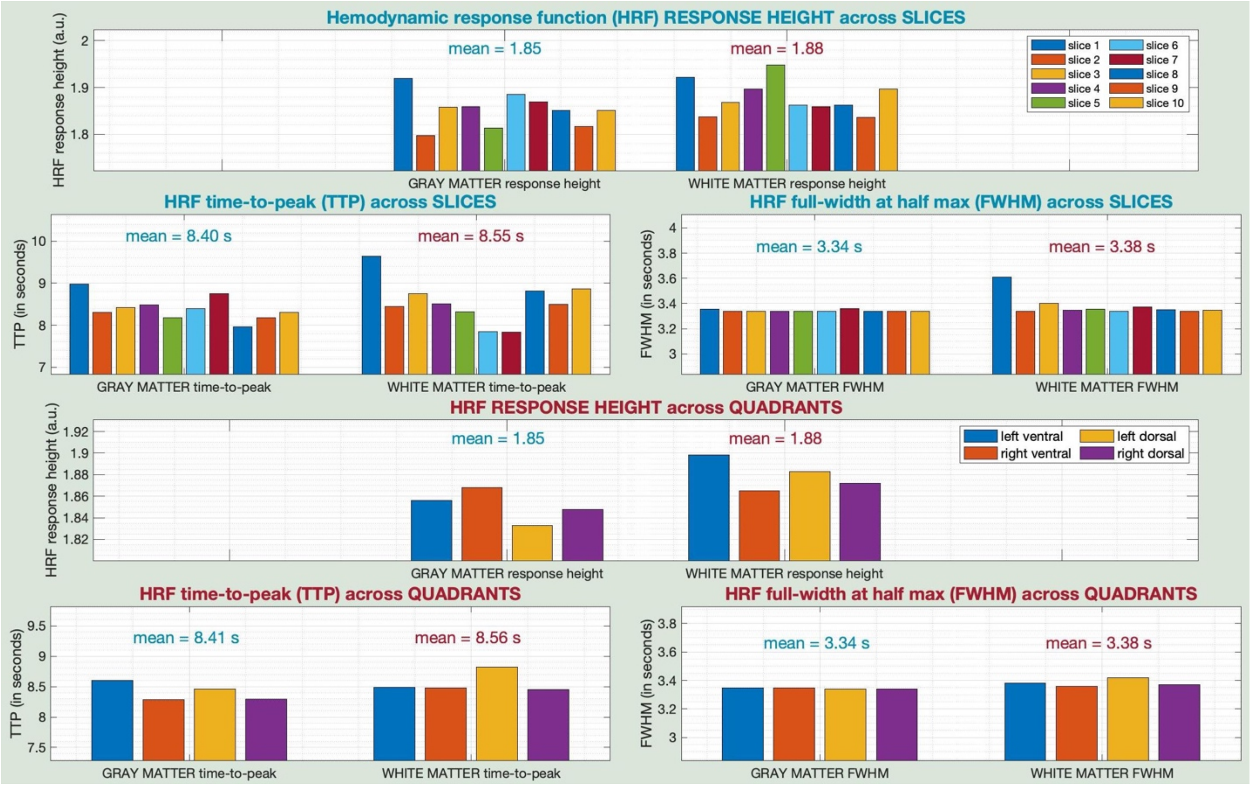


FC_compare_boxplot_20quadTS_allSlices.jpg

The most robust finding in resting-state spinal cord fMRI literature so far is the existence of high within-slice FC between left (L) and right (R) ventral (V) horns, as well as between L and R dorsal (D) horns (with the former being stronger than the latter), and the rest of the connections being weak (LV-LD, RV-RD, LV-RD, RV-LD). Therefore, reproducing these results in newly acquired data may be viewed as a litmus test for overall data quality. The below figure demonstrates this for our data by presenting boxplots of voxel-to-voxel correlations both within and between GM quadrants. The center mark is the median value, the edges of the boxes are 25$^{th}$ and 75$^{th}$ percentile values, the whiskers extend to the most extreme values not considered outliers, and the outliers are shown individually as red '+' marker symbols.

What we explained above is FC between quadrants, but it is also true that correlations between voxels of the same quadrant must have the highest value because of their proximity and functional homogeneity. In the figure below, the red boxes show within-quadrant FC, the green boxes represent LV-RV and LD-RD connectivity, and the blue boxes show the rest of the connections. We clearly observe the highest values among red, moderate values among green, and low values among the blue boxes.

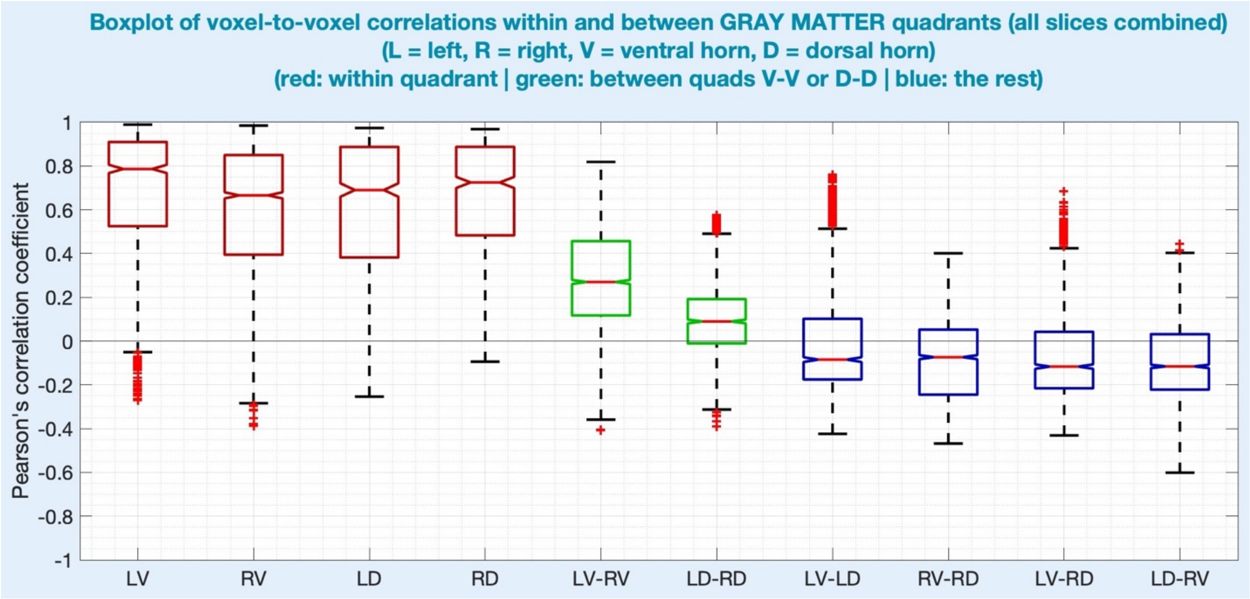


FCmatrix_WithinSlice_allSlices.jpg

This figure shows the all-important within-slice FC matrix, averaged across all slices in the given subject. It is symmetric because FC is non-directional. We observe the expected pattern of high LV-RV connectivity (R=0.41) and LD-RD connectivity (R=0.21), with the former being higher than the latter, and the rest of the connections being weak (range of R = [0.02–0.15]).

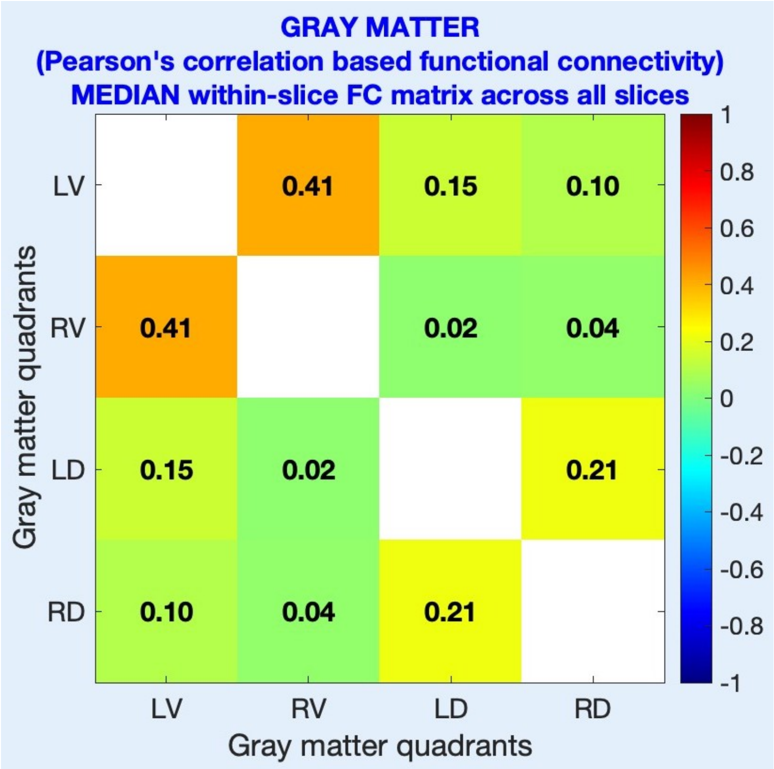

FCmatrix_Within_and_BetweenSlices_GM.jpg

This is the novel all-encompassing within- and between-slice FC matrix visualization, shown in a format similar to what we see in the brain connectome. The matrix goes from the first to the last slice linearly across both dimensions, and the slices are visually separated by blank white spaces. Within each unit, we can observe FC between corresponding quadrants (LV-LD-RV-RD, in that order). The diagonal units correspond to within-slice FC, and the off-diagonal units show corresponding between-slice FC. The matrix is symmetric as FC is non-directional. Understanding this visualization is helpful for both neuroscientific investigation (which we hope researchers will use) and identifying noisy/unreliable data (spurious correlations between random slices).

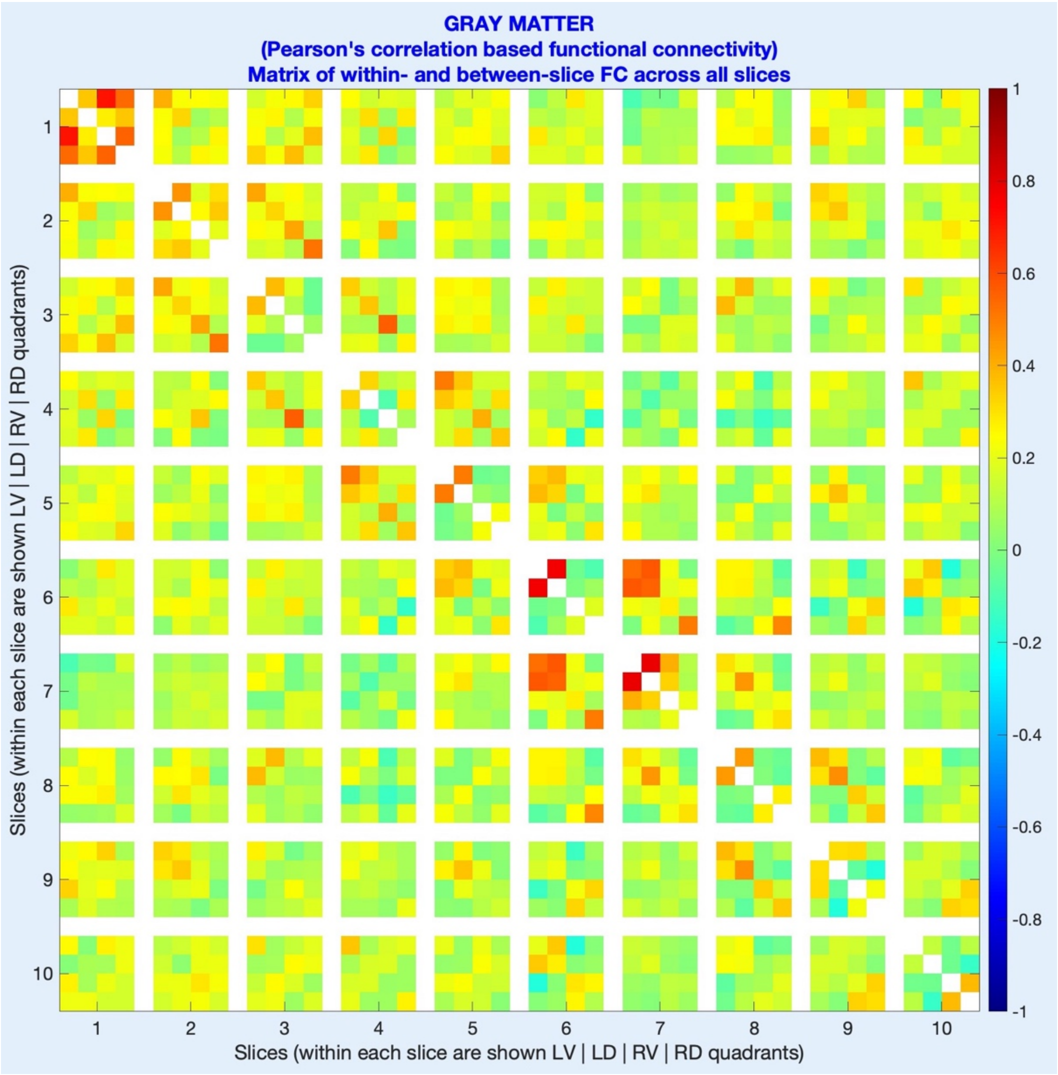

fALFF_across_slices.jpg

Neptune presents bar graphs of fALFF values across slices (in each quadrant separately) and across quadrants (averaged across all slices). FALFF is a neuroscientifically interesting metric and is also helpful as a test of data quality. Data with relatively low neural information has lower fALFF values, which can be used to compare different protocols. Severely distorted data or those slices having signal dropouts also present with dips in fALFF values. Since fALFF is the ratio of signal power in the fMRI band of interest (typically 0.01–0.1 Hz) to the signal power in the entire frequency band, any factor that decreases the fMRI signal of interest (loss of neural information) or increases the noise floor (higher distortions or noise) will show up as altered fALFF values, which users can benefit from. Since the data used here had low temporal resolution (=3.34s, corresponding to the highest frequency value being sampled at 0.15 Hz), fALFF is not informative.

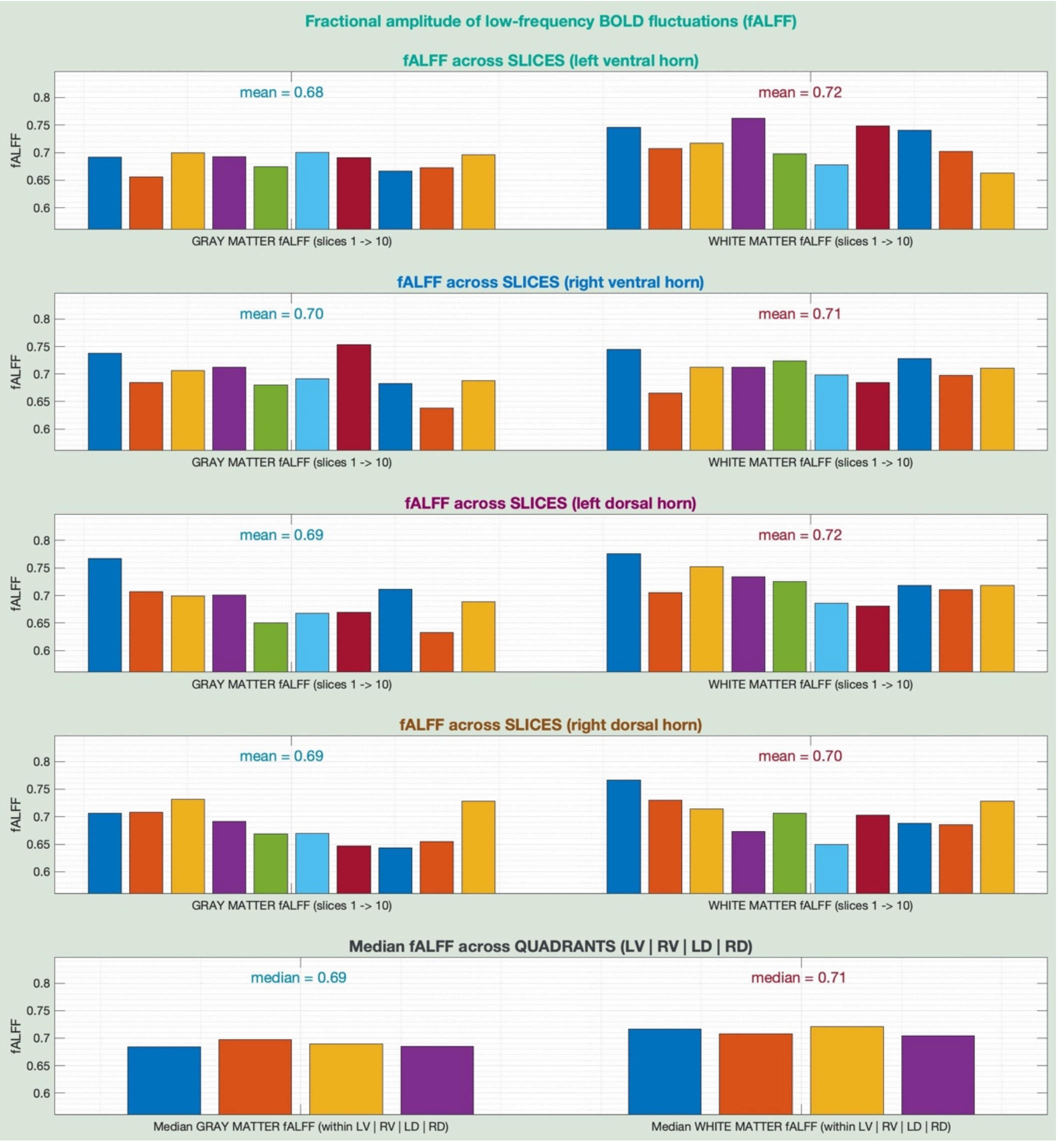

So far, we described QC outputs available in the *QC_shortlist* subdirectory. Next, we describe a few relevant QC outputs related to each step within other subdirectories.

<u>05_notCordMask / notCordMask_slice##.jpg</u>

It shows the boundary of the not-cord mask for each slice finalized by the user during step-5.

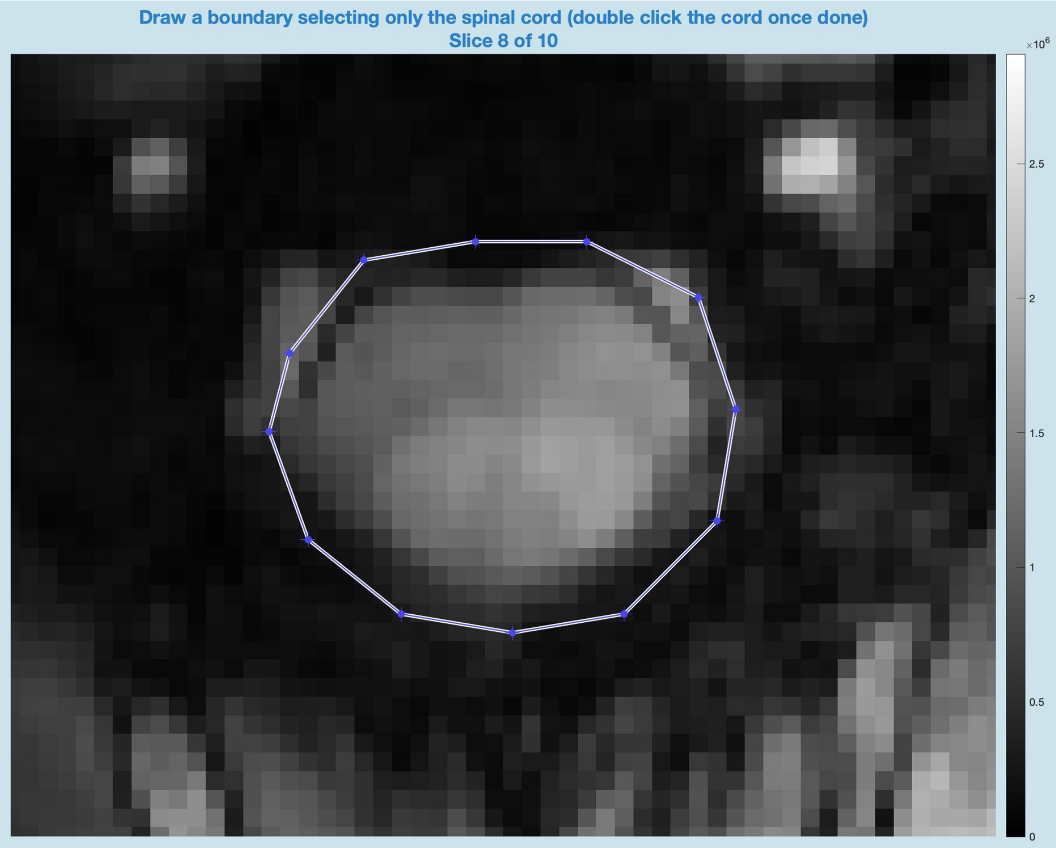


<u>10_GM-WM-CSF-masks / final_masks_slice##.jpg</u>

It shows the GM, WM and CSF boundaries for each slice finalized by the user during step-10. Two separate views are shown – one with full FOV and another zoomed into the cord. Boundary points drawn individually in GM, WM or CSF are also available (not shown here). These figures are handy for researchers to ensure that their lab members (perhaps newly hired) are finalizing these manual masks as intended.

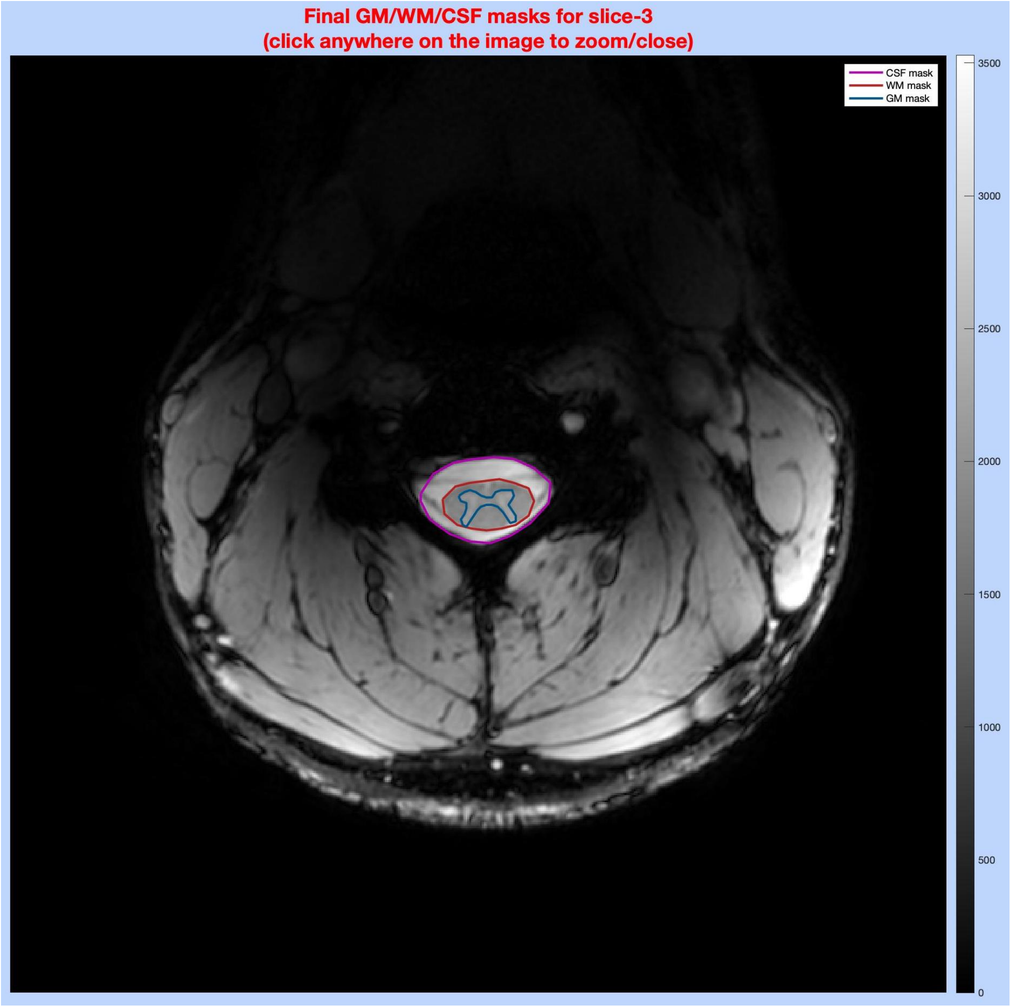
Final GM/WM/CSF masks for slice-3
(click anywhere on the image to zoom/close)
CSF mask
WM mask
GM mask
3500
3000
2500
2000
1500
1000
500
0

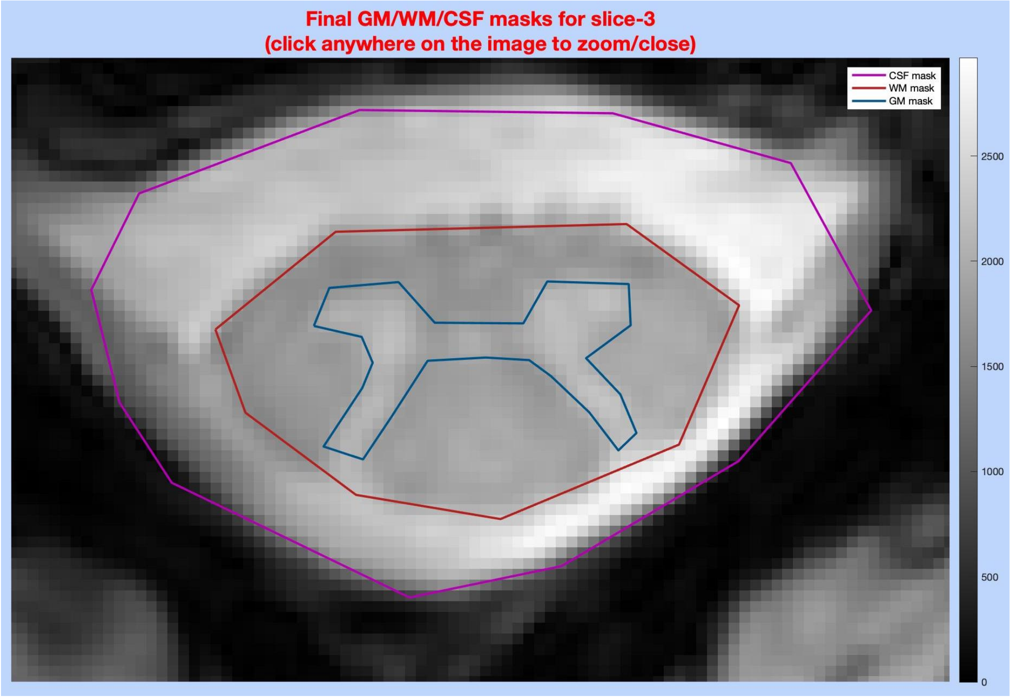
Final GM/WM/CSF masks for slice-3
(click anywhere on the image to zoom/close)
CSF mask
WM mask
GM mask
2500
2000
1500
1000
500
0

10_GM-WM-CSF-masks / cross_sectional_areas_across_slices.jpg

This figure shows cross-sectional areas (CSA) across slices – GM area, WM area, CSF area, and cord area (GM+WM). These are computed from the masks defined by the user. They are not validated measures, so, as previously mentioned, are not intended to be used in publications. This figure helps ensure the quality of anatomical images because average CSA across vertebral levels is published in the literature, which can be compared against the values shown here. For instance, the CSF area is expected to be highest in C1/C2 and gradually decrease as we go inferior. Sudden dips in CSA and zig-zag patterns are also causes of concern that indicate some form of distortion in anatomical images.

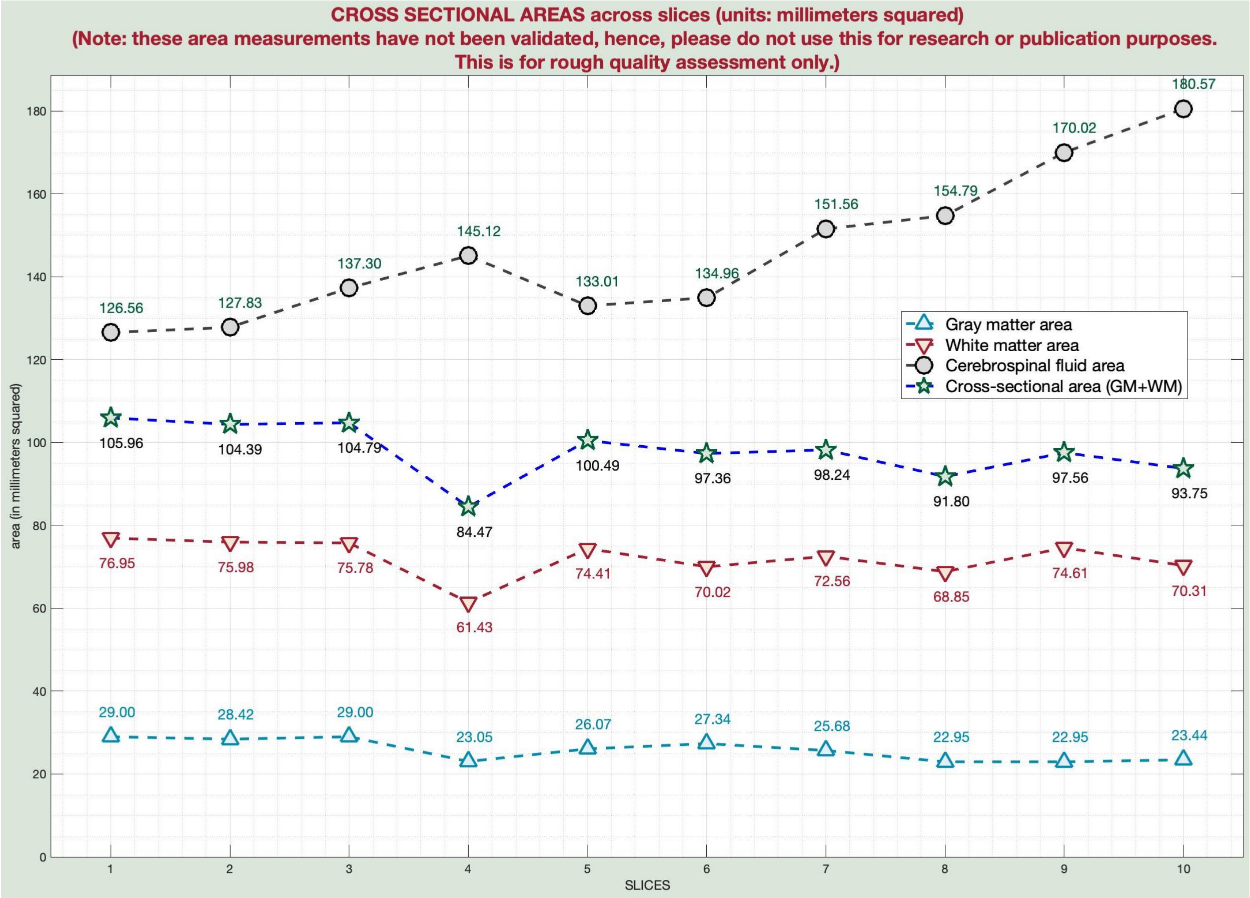


11_func-anat-registration / func_anat_reg_slice##.jpg

This figure shows the alignment of functional images with corresponding anatomical counterparts. The top two figures are zoomed out, while the bottom two are zoomed in and show tissue boundaries. It can be used to ascertain successful co-registration.

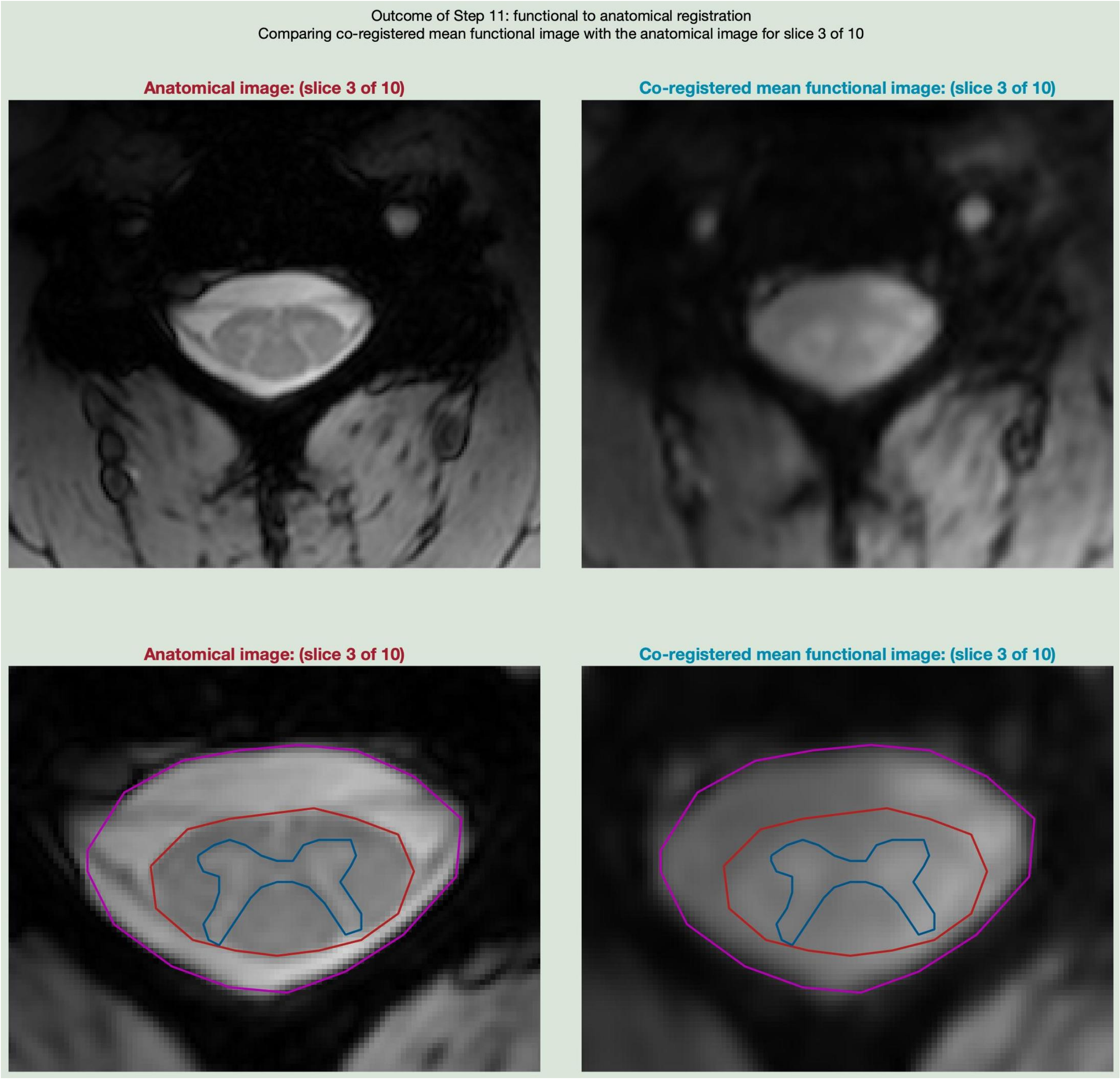


16_cord_quadrants / cord_quadrants_slice##.jpg

The figure below shows the four GM and WM quadrants in each slice obtained after step-16.

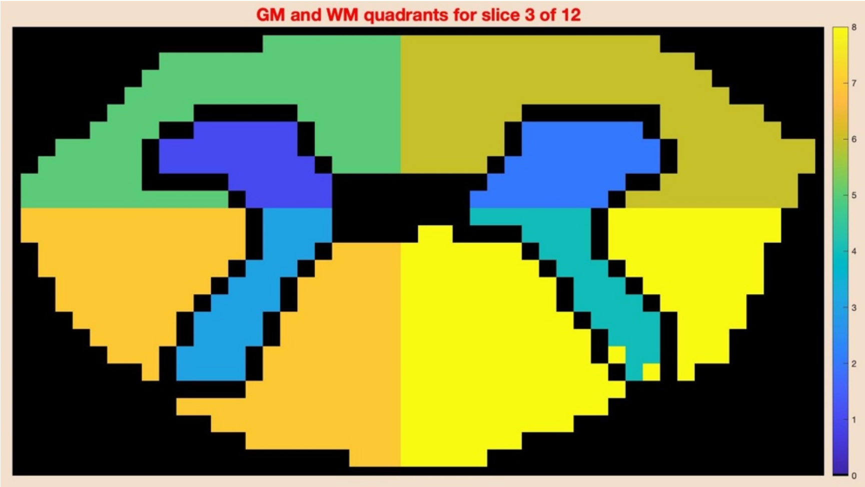


16_cord_quadrants / cross_sectional_areas_each_quad.jpg

This is a crowded figure, but with some attention, it can help detect segmentation errors. It shows the area of each quadrant across slices in both GM and WM. Sudden dips and zig-zag patterns are a matter of concern. In this example, we can observe a dip in the WM areas of slice 4 without any significant change in GM areas, indicating that the tissue boundary between WM and CSF was possibly not well defined (in this instance, it was because of signal dropout).

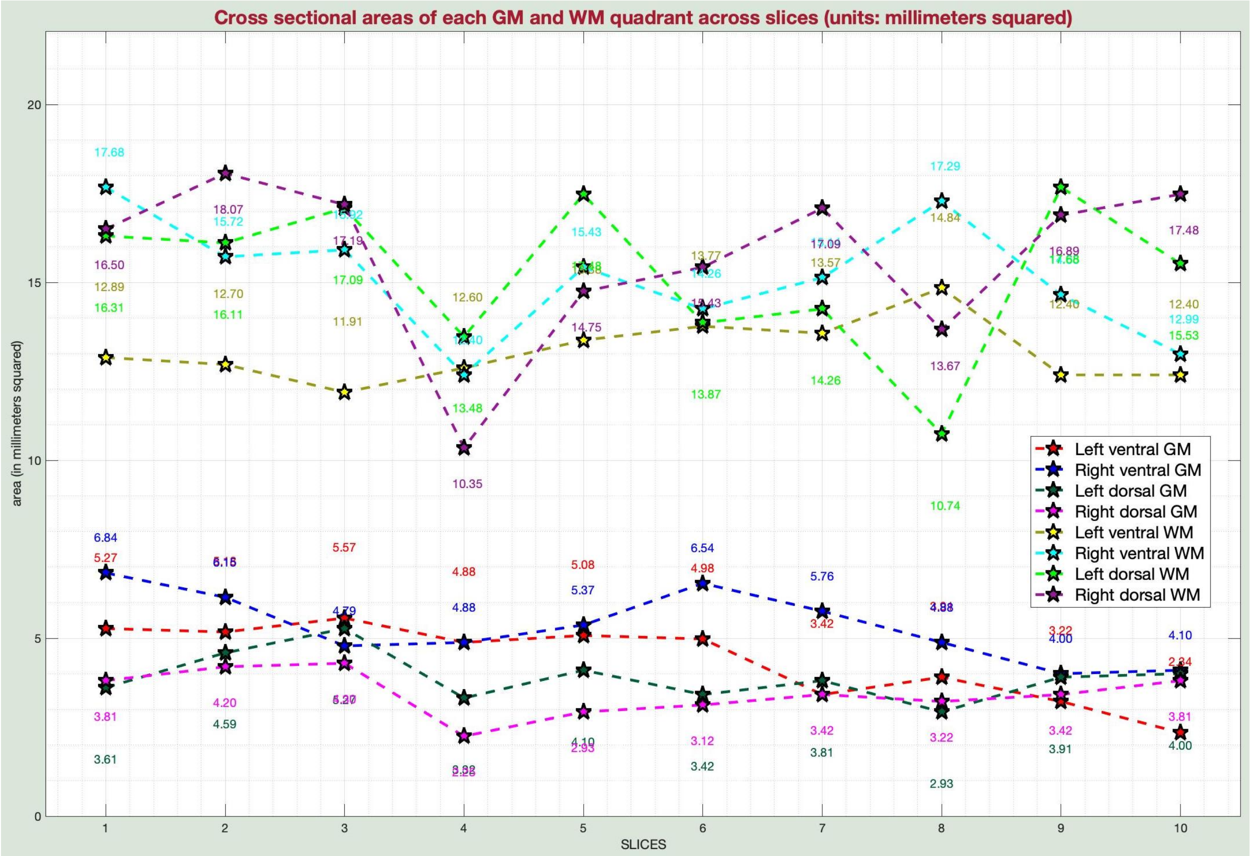

19_disk_storage_info / disk_storage_bargraph.jpg

This bar graph shows the benefits of Neptune's file management. Removing redundant slice-wise files significantly lowers the burden on data storage and movement (from the left to the central bars). Further compressing the *.nii* files to *.nii.gz* adds to the space savings (from the central bar to the one on the right).

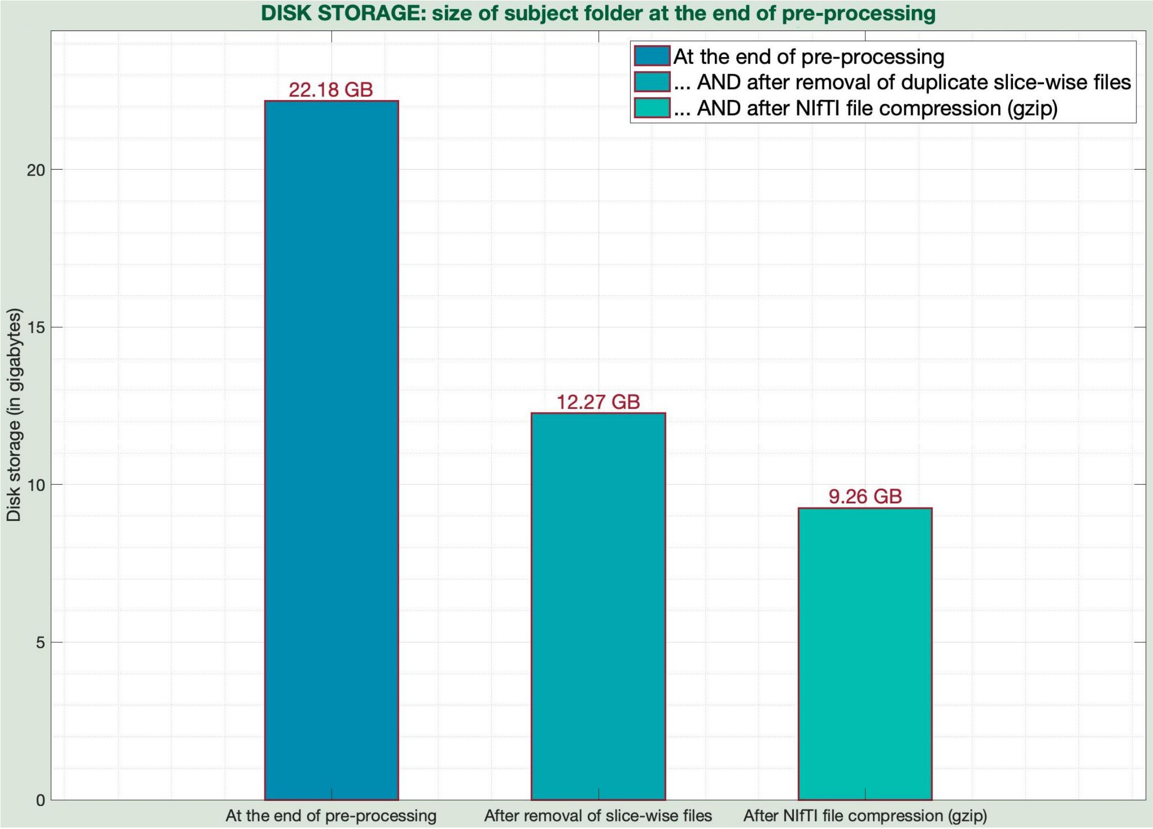


20_quadrant_timeseries / QuadMask_for_postproc_slice##.jpg

In step-20, Neptune takes the cord quadrant masks obtained from step-16 and erodes them further using a disc of radius 1.5 mm (usually 6 voxels). This further expels boundary voxels and avoids the possibility of having misplaced voxels in a given quadrant's mask. Neptune ensures that the erosion does not wipe out quadrants by not eroding if a given quadrant has fewer than 3 voxels. The below figure shows quadrant masks before and after erosion. The latter is used in further post-processing steps. Users can modify the disc radius in '*scfMRItb_20_quadMask.m*' (variable of interest: *wm_erode*), or altogether not erode by changing a parameter flag (*skip_erosion*). The code is well-commented to facilitate user modifications, if necessary.

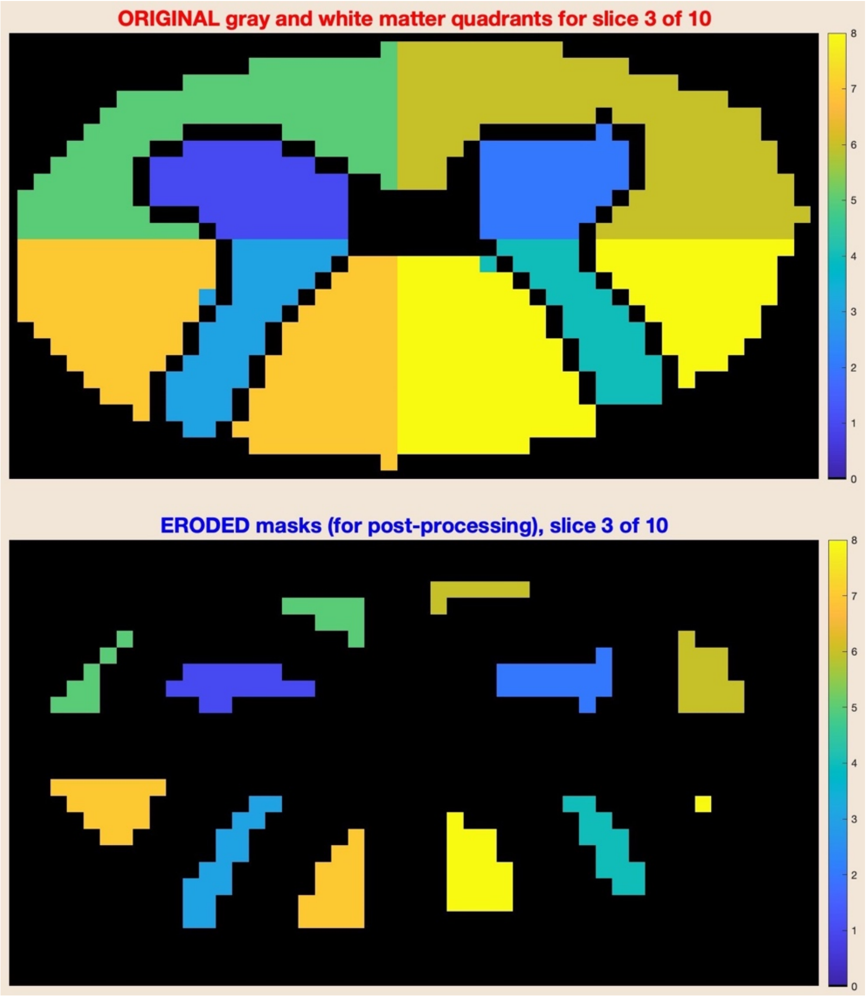


<home directory> / stats / figures / stats_SFC_WithinSlice_SlicesPooled.jpg
<home directory> / stats / figures / stats_SFC_Within_and_BetweenSlices_GM.jpg

The below figures are comparable to within-slice FC and within/between-slice FC figures described earlier. These figures show significant group-level T-statistics instead of FC values.

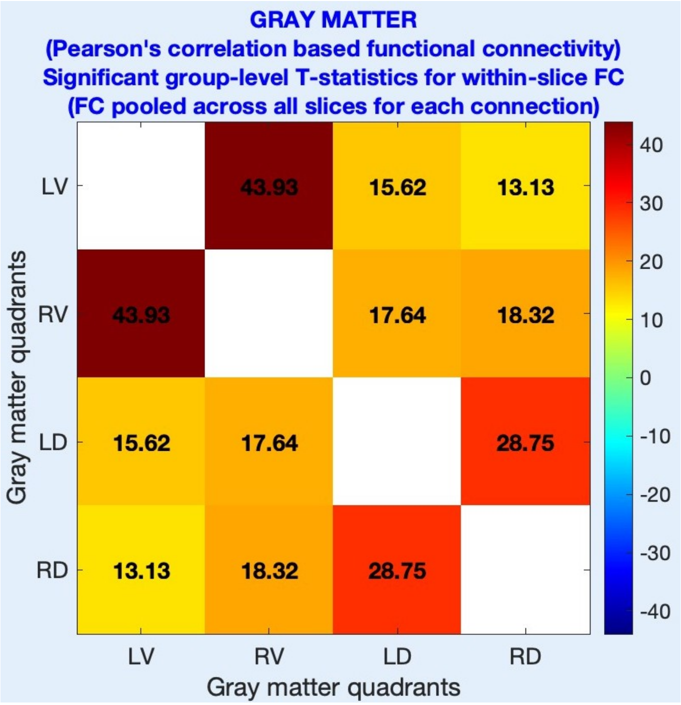
GRAY MATTER
(Pearson's correlation based functional connectivity)
Significant group-level T-statistics for within-slice FC
(FC pooled across all slices for each connection)
LV
RV
LD
RD
43.93
15.62
13.13
43.93
17.64
18.32
15.62
17.64
28.75
13.13
18.32
28.75
Gray matter quadrants
Gray matter quadrants
40
30
20
10
0
-10
-20
-30
-40

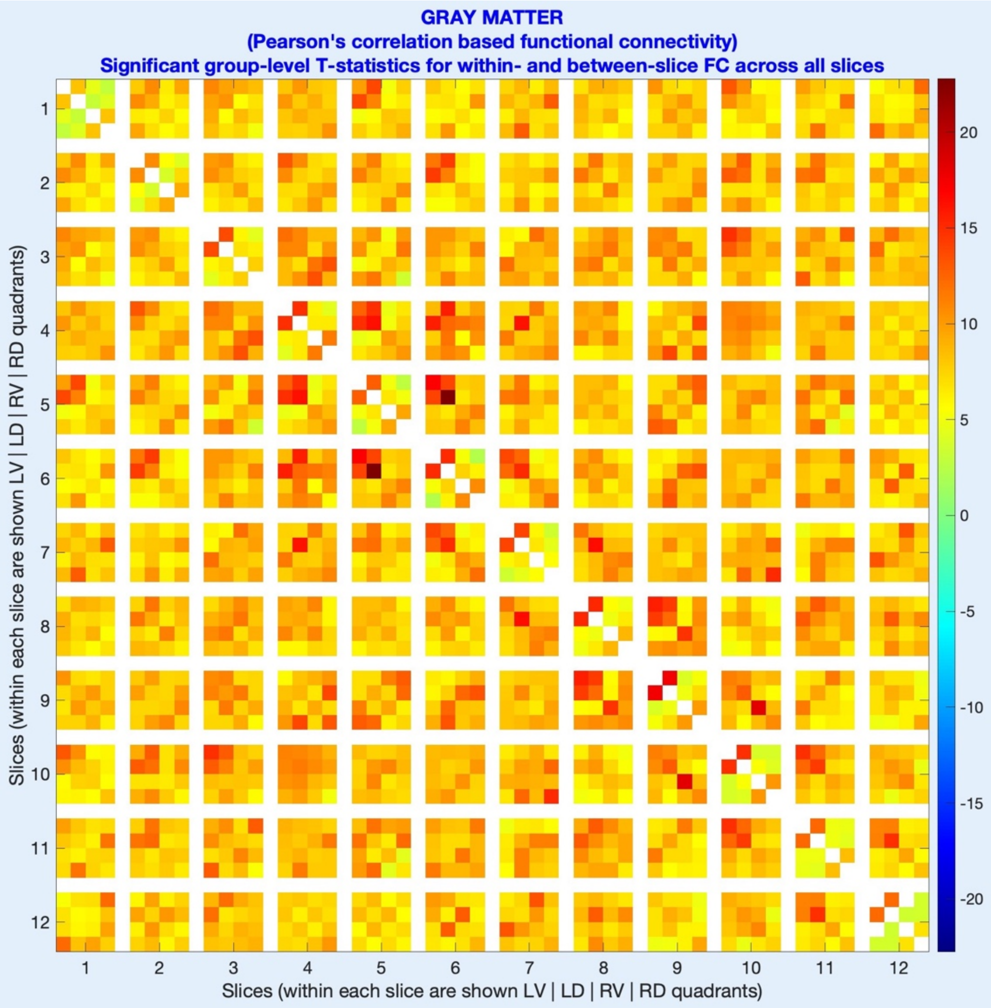
GRAY MATTER
(Pearson's correlation based functional connectivity)
Significant group-level T-statistics for within- and between-slice FC across all slices
Slices (within each slice are shown LV | LD | RV | RD quadrants)
Slices (within each slice are shown LV | LD | RV | RD quadrants)
1 2 3 4 5 6 7 8 9 10 11 12
20
15
10
5
0
-5
-10
-15
-20

<home directory> / stats / figures / stats_fALFF.jpg

Lastly, this figure shows significant group-level T-statistics for fALFF, and is comparable to the fALFF figure described before.

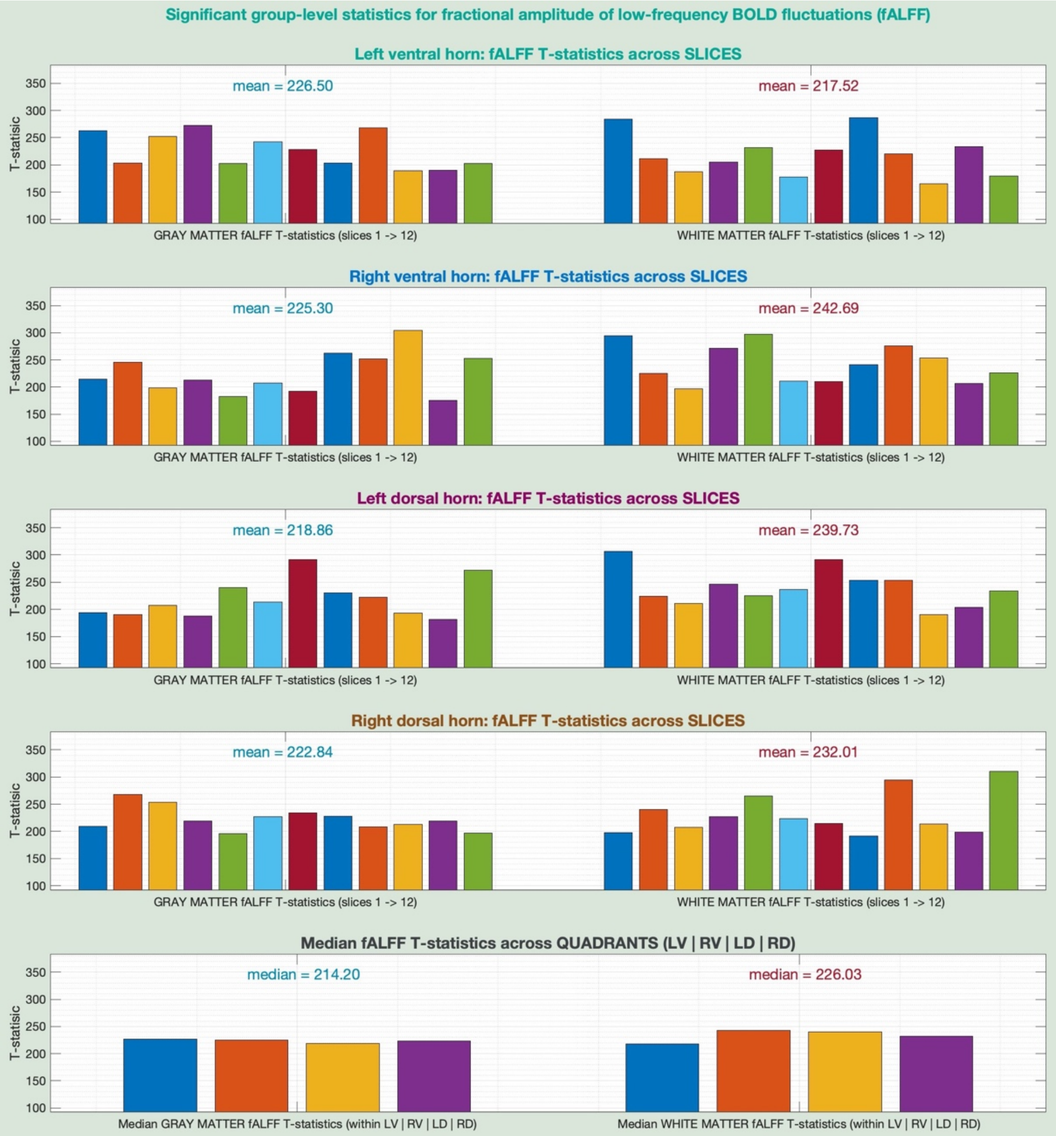


------- ------- -------
