## Supplementary material for "Neptune: a toolbox for spinal cord functional MRI data processing and quality assurance": Neptune software: scfMRItb_01_getNifti_philips.m

```
function scfMRItb_01_getNifti_philips(base_dir_sub, C_spine, scan7T, varargin)

if nargin<4
    wbar3 = waitbar(0,'01. PAR/REC to NIfTI conversion...','Name','Progress: PAR/REC to NIfTI conversion...','Units','normalized','Position',[3/5 0 1/5 1/17]);
else
    wbar3 = varargin{1};
end

scan3T = ~scan7T;
L_spine = ~C_spine;

current_dir = pwd;

cd(base_dir_sub)
sdir = dir('*.PAR');

for k=1:length(sdir)
    try waitbar((k/length(sdir)),wbar3,sprintf('01. PAR/REC to NIfTI conversion in this subject: file (%d) of (%d)',k,length(sdir))); catch, end
    if     scan3T && C_spine % 3T cervical
        vuPARRECtoNIfTI(sdir(k).name, base_dir_sub); % magnitude only
    elseif scan3T && L_spine % 3T lumbar
        vuPARRECtoNIfTI(sdir(k).name, base_dir_sub, 2, 3); % complex data
    else % 7T
        try % if data are complex
            vuPARRECtoNIfTI(sdir(k).name, base_dir_sub, 2, 3);
        catch % magnitude only
            vuPARRECtoNIfTI(sdir(k).name, base_dir_sub);
        end
    end
end
close(wbar3)
cd(current_dir)

end
```
