## Supplementary material for "Neptune: a toolbox for spinal cord functional MRI data processing and quality assurance": Neptune software: scfMRItb_01_getNifti_siemens.m

```
function scfMRItb_01_getNifti_siemens(base_dir_sub,varargin)

% The "dicom" folder must conform to one of the two following structures:
% (i) The "dicom" folder must have all the dicom files (.dcm or .ima or PAR/REC files) and no other folders
% Or, (ii) the "dicom" folder must have other folders inside it, and these folders must have the dicoms

if nargin<2
    wbar3 = waitbar(0,'01. Dicom to NIfTI conversion...','Name','Progress: Dicom to NIfTI conversion...','Units','normalized','Position',[3/5 0 1/5 1/17]);
else
    wbar3 = varargin{1};
end

directory = sprintf('%s/dicom',base_dir_sub);
cd(directory)

sdir1=dir; sdir1=sdir1(3:end);
for k=1:length(sdir1)
    sdirtype(k,1) = sdir1(k).isdir;
end; clear k

if sum(sdirtype)==0 % if there are only dicom files and no folders
    dicm2nii(directory, base_dir_sub, '.nii')
else % if there are folders containing dicom files
    flag=0;
    for k=1:length(sdir1)
        if sdirtype(k)==1
            flag=flag+1;
            try waitbar((flag/sum(sdirtype)),wbar3,sprintf('01. Dicom to NIfTI conversion in this subject: directory (%d) of (%d)',flag,sum(sdirtype))); catch, end
            cd(sdir1(k).name)
            dicm2nii(sprintf('%s/%s',directory,sdir1(k).name), base_dir_sub, 'nii')
            cd ..
        end
    end; clear k
    close(wbar3)
end

cd ..

end
```
