## Supplementary material for "Neptune: a toolbox for spinal cord functional MRI data processing and quality assurance": Neptune software: scfMRItb_01_init_NIfTIconvert.m

```
% Setting up dicom (or PAR/REC) to NIfTI conversion

if scanner~=3 % Siemens or Philips scanners only
    
if skip_NIfTIconvert_choice==0 % it will ask you if you want to perform NIfTI conversion, because the choice has not been made yet
    if scanner==1
        try waitbar(4/8,wbar1,'Dicom to NIfTI conversion...'); catch, end
        quest = sprintf('Convert raw data (dicom) to NIfTI?\n(not required if you already have the .nii files)');
    elseif scanner==2
        try waitbar(4/8,wbar1,'PAR/REC to NIfTI conversion...'); catch, end
        quest = sprintf('Convert raw data (PAR/REC) to NIfTI?\n(not required if you already have the .nii files)');
    end
    answer_NIfTIconvert = questdlg(quest,'NIfTI conversion','Yes, convert','No','No'); clear quest
end

switch answer_NIfTIconvert
    case 'Yes, convert'
        nifti_convert=1;
        if scanner==1
            clk=clock; L=L+1; logg{L,1}=sprintf('%d/%s/%d %dh:%dm:%ds - start (01) dicom to NIfTI conversion',clk(3),month{clk(2)},clk(1),clk(4),clk(5),round(clk(6))); fprintf(fileID,'%s\n',logg{L,1}); try set(S.tx,'string',logg); catch, end; clear clk
            quest = sprintf('NIfTI conversion can be automated for all subjects if you have a folder named "dicom" (that has the raw files) within each subject''s directory.\n\nDo you want to automate, or convert manually subject after subject?\nYou could also choose to open the dicm2nii GUI.\n');
            answer2 = questdlg(quest,'NIfTI conversion','Automatic','Manual','Open the GUI','Manual'); clear quest
            wbar2 = waitbar(0,'Dicom to NIfTI conversion...','Name','NIfTI conversion...','Units','normalized','Position',[4/5 0 1/5 1/17]);
        elseif scanner==2
            clk=clock; L=L+1; logg{L,1}=sprintf('%d/%s/%d %dh:%dm:%ds - start (01) PAR/REC to NIfTI conversion',clk(3),month{clk(2)},clk(1),clk(4),clk(5),round(clk(6))); fprintf(fileID,'%s\n',logg{L,1}); try set(S.tx,'string',logg); catch, end; clear clk
            quest = sprintf('NIfTI conversion can be automated for all subjects if you have your PAR/REC files in each subject''s directory.\n\nDo you want to automate, or convert manually subject after subject?\n');
            answer2 = questdlg(quest,'NIfTI conversion','Automatic','Manual','Manual'); clear quest
            wbar2 = waitbar(0,'PAR/REC to NIfTI conversion...','Name','NIfTI conversion...','Units','normalized','Position',[4/5 0 1/5 1/17]);
        end
        switch answer2
            case 'Automatic'
                if scanner==1 % Siemens
                    for k=1:length(sub_dir)
                        clk=clock; L=L+1; logg{L,1}=sprintf('%d/%s/%d %dh:%dm:%ds - begin (01) dicom to NIfTI conversion for subject (%d) of %d)',clk(3),month{clk(2)},clk(1),clk(4),clk(5),round(clk(6)),k,length(sub_dir)); fprintf(fileID,'%s\n',logg{L,1}); try set(S.tx,'string',logg); catch, end; clear clk
                        try waitbar((k/length(sub_dir)),wbar2,sprintf('Dicom to NIfTI conversion: Subject (%d) of (%d)\n',k,length(sub_dir))); catch, end
                        fprintf('01. Dicom to NIfTI conversion: Subject (%d) of (%d)\n',k,length(sub_dir))
                        time1 = tic;
                        scfMRItb_01_getNifti_siemens(sub_dir{k})
                        time_taken_01_dicom_to_nifti{k,1} = toc(time1);
                        clk=clock; L=L+1; logg{L,1}=sprintf('%d/%s/%d %dh:%dm:%ds - end (01) dicom to NIfTI conversion for subject (%d) of %d)',clk(3),month{clk(2)},clk(1),clk(4),clk(5),round(clk(6)),k,length(sub_dir)); fprintf(fileID,'%s\n',logg{L,1}); try set(S.tx,'string',logg); catch, end; clear clk time1
                        refresh_html_log
                    end; clear k2

                elseif scanner==2 % Philips
                    for k=1:length(sub_dir)
                        clk=clock; L=L+1; logg{L,1}=sprintf('%d/%s/%d %dh:%dm:%ds - begin (01) PAR/REC to NIfTI conversion for subject (%d) of %d)',clk(3),month{clk(2)},clk(1),clk(4),clk(5),round(clk(6)),k,length(sub_dir)); fprintf(fileID,'%s\n',logg{L,1}); try set(S.tx,'string',logg); catch, end; clear clk
                        try waitbar((k/length(sub_dir)),wbar2,sprintf('PAR/REC to NIfTI conversion: Subject (%d) of (%d)\n',k,length(sub_dir))); catch, end
                        fprintf('01. PAR/REC to NIfTI conversion: Subject (%d) of (%d)\n',k,length(sub_dir))
                        time1 = tic;
                        scfMRItb_01_getNifti_philips(sub_dir{k}, C_spine, scan7T)
                        time_taken_01_parrec_to_nifti{k,1} = toc(time1);
                        clk=clock; L=L+1; logg{L,1}=sprintf('%d/%s/%d %dh:%dm:%ds - end (01) PAR/REC to NIfTI conversion for subject (%d) of %d)',clk(3),month{clk(2)},clk(1),clk(4),clk(5),round(clk(6)),k,length(sub_dir)); fprintf(fileID,'%s\n',logg{L,1}); try set(S.tx,'string',logg); catch, end; clear clk time1
                        refresh_html_log
                    end; clear k2
                end
                
            case 'Manual'
                if scanner==1 % Siemens
                    fprintf('01. Dicom to NIfTI conversion\n')
                    answer3 = questdlg('Select dicom folder/files, OR enter wildcard search string','NIfTI conversion','Choose dicom folder/files','Wildcard search string','Choose dicom folder/files');
                    switch answer3
                        case 'Choose dicom folder/files'
                            for k=1:length(sub_dir)
                                fprintf('Choose dicom files/folder for NIfTI conversion: Subject (%d) of (%d)\n',k,length(sub_dir))
                                cd(sub_dir{k,1})
                                sdir3 = dir; sdir3name={sdir3.name};
                                prompt = sprintf('Subject %s ... Select dicom source file(s) or folder(s). It can be a zip or tgz file, a folder containing dicom files, or other convertible files...\n',sub_dir{k});
                                indx = listdlg('PromptString',{prompt,''},'SelectionMode','multiple','ListSize',[400 400],'ListString',sdir3);
                                for k3=1:length(indx)
                                    if sdir3(indx(k3)).isdir==1
                                        dicom_dir{k}{k3,1} = [sdir3name{indx(k3)} '/'];
                                    else
                                        dicom_dir{k}{k3,1} = sdir3name{indx(k3)};
                                    end
                                end; clear k3 indx prompt sdir3 sdir3name
                                cd ..
                            end; clear k2
                            
                        case 'Wildcard search string'
                            fprintf('Choose dicom files/folder for NIfTI conversion')
                            prompt = sprintf('Enter wild card string to choose dicom source file(s) or folder(s).\nDicoms can be zip or tgz files, folders containing dicom files, or other convertible files...\n');
                            answer4 = inputdlg(prompt,'NIfTI conversion',1,{'*.dcm'}); clear prompt
                            for k=1:length(sub_dir)
                                cd(sub_dir{k,1})
                                sdir3 = dir(sprintf('**/%s',answer4));
                                sdir3name={sdir3.name}; sdir3folder={sdir3.folder};
                                for k3=1:length(sdir3)
                                    dicom_dir{k}{k3,1} = sprintf('%s/%s',sdir3folder{k3},sdir3name{k3});
                                end
                                clear k3 sdir3 sdir3name sdir3folder
                                cd ..
                            end; clear k2 answer4
                    end; clear answer3
                    for k=1:length(sub_dir)
                        clk=clock; L=L+1; logg{L,1}=sprintf('%d/%s/%d %dh:%dm:%ds - begin (01) dicom to NIfTI conversion for subject (%d) of %d)',clk(3),month{clk(2)},clk(1),clk(4),clk(5),round(clk(6)),k,length(sub_dir)); fprintf(fileID,'%s\n',logg{L,1}); try set(S.tx,'string',logg); catch, end; clear clk
                        try waitbar((k/length(sub_dir)),wbar2,sprintf('Dicom to NIfTI conversion: Subject (%d) of (%d)\n',k,length(sub_dir))); catch, end
                        fprintf('01. Dicom to NIfTI conversion: Subject (%d) of (%d)\n',k,length(sub_dir))
                        time1 = tic;
                        dicm2nii(dicom_dir{k}, sub_dir{k,1}, '.nii')
                        time_taken_01_dicom_to_nifti{k,1} = toc(time1);
                        clk=clock; L=L+1; logg{L,1}=sprintf('%d/%s/%d %dh:%dm:%ds - end (01) dicom to NIfTI conversion for subject (%d) of %d)',clk(3),month{clk(2)},clk(1),clk(4),clk(5),round(clk(6)),k,length(sub_dir)); fprintf(fileID,'%s\n',logg{L,1}); try set(S.tx,'string',logg); catch, end; clear clk time1
                        refresh_html_log
                    end
                    
                elseif scanner==2 % Philips
                    fprintf('01. PAR/REC to NIfTI conversion\n')
                    answer3 = questdlg('Select PAR/REC files OR enter wildcard search string','NIfTI conversion','Choose PAR/REC files','Wildcard search string','Choose PAR/REC files');
                    switch answer3
                        case 'Choose PAR/REC files'
                            for k=1:length(sub_dir)
                                fprintf('Choose PAR/REC files for NIfTI conversion: Subject (%d) of (%d)\n',k,length(sub_dir))
                                cd(sub_dir{k,1})
                                sdir3 = dir('*.PAR'); sdir3={sdir3.name};
                                prompt = sprintf('Subject %s ... Select PAR file(s).',sub_dir{k});
                                indx = listdlg('PromptString',{prompt,''},'SelectionMode','multiple','ListSize',[400 400],'ListString',sdir3);
                                for k3=1:length(indx)
                                    parrec_dir{k}{k3,1} = sdir3{indx(k3)};
                                end; clear k3 indx prompt sdir3
                                cd ..
                            end; clear k2
                            
                        case 'Wildcard search string'
                            fprintf('Choose PAR/REC files for NIfTI conversion')
                            prompt = sprintf('Enter wild card string to choose PAR/REC files (use extension *.PAR).\n');
                            answer4 = inputdlg(prompt,'NIfTI conversion',1,{'*.PAR'}); clear prompt
                            for k=1:length(sub_dir)
                                cd(sub_dir{k,1})
                                sdir3 = dir(sprintf('**/%s',answer4{1}));
                                sdir3name={sdir3.name}; sdir3folder={sdir3.folder};
                                for k3=1:length(sdir3)
                                    if strcmp(sdir3folder{k3},sub_dir{k,1}(1:end-1))~=1
                                        movefile(sprintf('%s/%s',sdir3folder{k3},sdir3name{k3}),sprintf('%s%s',sub_dir{k,1},sdir3name{k3}));
                                    end
                                    parrec_dir{k}{k3,1} = sdir3name{k3};
                                end
                                clear k3 sdir3 sdir3name sdir3folder
                                cd ..
                            end; clear k2 answer4
                    end; clear answer3
                    for k=1:length(sub_dir)
                        clk=clock; L=L+1; logg{L,1}=sprintf('%d/%s/%d %dh:%dm:%ds - begin (01) PAR/REC to NIfTI conversion for subject (%d) of %d)',clk(3),month{clk(2)},clk(1),clk(4),clk(5),round(clk(6)),k,length(sub_dir)); fprintf(fileID,'%s\n',logg{L,1}); try set(S.tx,'string',logg); catch, end; clear clk
                        try waitbar((k/length(sub_dir)),wbar2,sprintf('PAR/REC to NIfTI conversion: Subject (%d) of (%d)\n',k,length(sub_dir))); catch, end
                        fprintf('01. PAR/REC to NIfTI conversion: Subject (%d) of (%d)\n',k,length(sub_dir))
                        cd(sub_dir{k})
                        time1 = tic;
                        for k3=1:length(parrec_dir{k})
                            if     scan3T && C_spine % 3T cervical
                                vuPARRECtoNIfTI(parrec_dir{k}{k3,1}, sub_dir{k,1}); % magnitude only
                            elseif scan3T && L_spine % 3T lumbar
                                vuPARRECtoNIfTI(parrec_dir{k}{k3,1}, sub_dir{k,1}, 2, 3); % complex data
                            else % 7T
                                try % if data are complex
                                    vuPARRECtoNIfTI(parrec_dir{k}{k3,1}, sub_dir{k,1}, 2, 3);
                                catch % magnitude only
                                    vuPARRECtoNIfTI(parrec_dir{k}{k3,1}, sub_dir{k,1});
                                end
                            end
                        end
                        time_taken_01_parrec_to_nifti{k,1} = toc(time1);
                        cd ..
                        clk=clock; L=L+1; logg{L,1}=sprintf('%d/%s/%d %dh:%dm:%ds - end (01) PAR/REC to NIfTI conversion for subject (%d) of %d)',clk(3),month{clk(2)},clk(1),clk(4),clk(5),round(clk(6)),k,length(sub_dir)); fprintf(fileID,'%s\n',logg{L,1}); try set(S.tx,'string',logg); catch, end; clear clk time1
                        refresh_html_log
                    end

                end
                
            case 'Open the GUI'
                dicm2nii
        end
        close(wbar2)
        clk=clock; L=L+1; logg{L,1}=sprintf('%d/%s/%d %dh:%dm:%ds - NIfTI conversion COMPLETED FOR ALL SUBJECTS',clk(3),month{clk(2)},clk(1),clk(4),clk(5),round(clk(6))); fprintf(fileID,'%s\n',logg{L,1}); try set(S.tx,'string',logg); catch, end; clear clk
        L=L+1; logg{L,1}=sprintf('Total time taken so far: %dh:%dm:%ds',round(toc(tc00)/3600),round(mod(toc(tc00),3600)/60),round(mod(toc(tc00),60))); fprintf(fileID,'%s\n',logg{L,1}); try set(S.tx,'string',logg); catch, end; clear clk

    case 'No'
        clk=clock; L=L+1; logg{L,1}=sprintf('%d/%s/%d %dh:%dm:%ds - NIfTI conversion SKIPPED FOR ALL SUBJECTS',clk(3),month{clk(2)},clk(1),clk(4),clk(5),round(clk(6))); fprintf(fileID,'%s\n',logg{L,1}); try set(S.tx,'string',logg); catch, end; clear clk
        nifti_convert=0;
end; clear answer2 answer3 sdir2 sdir3

end
```
