## Supplementary material for "Neptune: a toolbox for spinal cord functional MRI data processing and quality assurance": Neptune software: scfMRItb_01_NORDIC_denoising.m

```
function scfMRItb_01_NORDIC_denoising(base_dir_sub,fname,fname_anat,NORDIC_type,run, varargin)

do_NORDIC_on_anatomical = 0; % NORDIC did not work on anatomicals that we tested on, as of April 2022.

if nargin<6
    wbar3 = waitbar(0,'Perform NORDIC denoising...','Name','Progress: Perform NORDIC denoising...','Units','normalized','Position',[3/5 0 1/5 1/17]);
else
    wbar3 = varargin{1};
end

if nargin<7
    Rns = 0; % 0 = don't create runX subfolder; X = create runX subfolder
else
    Rns = varargin{2};
end

if ~(exist([base_dir_sub 'QC' '/01_NORDIC_denoising'],'dir'))
    mkdir([base_dir_sub 'QC' '/01_NORDIC_denoising'])
end
if Rns==0
%     QCpath3 = [base_dir_sub 'QC' '/01_NORDIC_denoising/'];
else
    if ~(exist([base_dir_sub 'QC' '/01_NORDIC_denoising' '/run' num2str(Rns)],'dir'))
        mkdir([base_dir_sub 'QC' '/01_NORDIC_denoising' '/run' num2str(Rns)])
    end
%     QCpath3 = [base_dir_sub 'QC' '/01_NORDIC_denoising' '/run' num2str(Rns) '/'];
end

%% Perform NORDIC

%% functional
scfMRItb_04_unzipFile(base_dir_sub,fname, '')

if NORDIC_type==1 % use magnitude-only data
    ARG.magnitude_only = 1; % use magnitude-only data
    ARG.DIROUT = base_dir_sub; % output directory
    ARG.save_gfactor_map = 1; % save the estimated g-factor map
    NIFTI_NORDIC([base_dir_sub fname '.nii'], [base_dir_sub fname '.nii'], [fname '_NORDIC'], ARG)
    clear ARG
elseif NORDIC_type==2 % use magnitude and phase data
    ARG.DIROUT = base_dir_sub; % output directory
    ARG.save_gfactor_map = 1; % save the estimated g-factor map
    scfMRItb_04_unzipFile(base_dir_sub,fname, '_ph')
    NIFTI_NORDIC([base_dir_sub fname '.nii'], [base_dir_sub fname '_ph' '.nii'], [fname '_NORDIC'], ARG)
    clear ARG
end

movefile([base_dir_sub fname '.nii'],[base_dir_sub fname '_beforeNORDIC.nii'])
if exist([base_dir_sub fname '.nii.gz'],'file')
    movefile([base_dir_sub fname '.nii.gz'],[base_dir_sub fname '_beforeNORDIC.nii.gz'])
else
    gzip([base_dir_sub fname '_beforeNORDIC.nii'])
end
movefile([base_dir_sub fname '_NORDIC.nii'],[base_dir_sub fname '.nii'])
if exist([base_dir_sub fname '_NORDIC.nii.gz'],'file')
    movefile([base_dir_sub fname '_NORDIC.nii.gz'],[base_dir_sub fname '.nii.gz'])
else
    gzip([base_dir_sub fname '.nii'])
end

%% anatomical (as of April 2022, NORDIC was not suitable for anatomical data, hence this section will not be executed or accounted for)
    scfMRItb_04_unzipFile(base_dir_sub,fname_anat, '')
    if run==1 && do_NORDIC_on_anatomical==1

    if NORDIC_type==1 % use magnitude-only data
        ARG.magnitude_only = 1; % use magnitude-only data
        ARG.DIROUT = base_dir_sub; % output directory
        ARG.save_gfactor_map = 1; % save the estimated g-factor map
        NIFTI_NORDIC([base_dir_sub fname_anat '.nii'], [base_dir_sub fname_anat '.nii'], [fname_anat '_NORDIC'], ARG)
        clear ARG
    elseif NORDIC_type==2 % use magnitude and phase data
        ARG.DIROUT = base_dir_sub; % output directory
        ARG.save_gfactor_map = 1; % save the estimated g-factor map
        scfMRItb_04_unzipFile(base_dir_sub,fname_anat, '_ph')
        NIFTI_NORDIC([base_dir_sub fname_anat '.nii'], [base_dir_sub fname_anat '_ph' '.nii'], [fname_anat '_NORDIC'], ARG)
        clear ARG
    end

    movefile([base_dir_sub fname_anat '.nii'],[base_dir_sub fname_anat '_beforeNORDIC.nii'])
    if exist([base_dir_sub fname_anat '.nii.gz'],'file')
        movefile([base_dir_sub fname_anat '.nii.gz'],[base_dir_sub fname_anat '_beforeNORDIC.nii.gz'])
    else
        gzip([base_dir_sub fname_anat '_beforeNORDIC.nii'])
    end
    movefile([base_dir_sub fname_anat '_NORDIC.nii'],[base_dir_sub fname_anat '.nii'])
    if exist([base_dir_sub fname_anat '_NORDIC.nii.gz'],'file')
        movefile([base_dir_sub fname_anat '_NORDIC.nii.gz'],[base_dir_sub fname_anat '.nii.gz'])
    else
        gzip([base_dir_sub fname_anat '.nii'])
    end

    end

%%
if nargin<5
    close(wbar3)
end

end
```
