## Supplementary material for "Neptune: a toolbox for spinal cord functional MRI data processing and quality assurance": Neptune software: scfMRItb_02_slicetimingcorrection.m

```
function scfMRItb_02_slicetimingcorrection(base_dir_sub,fname, sliceorder)

% http://andysbrainblog.blogspot.com/2012/11/slice-timing-correction-in-spm.html
% slice timing correction is not recommended for TR<=1s

% which software to use:
use_SPM = 1; % use SPM for slice timing correction, not AFNI (0 = use AFNI instead)

scfMRItb_04_unzipFile(base_dir_sub, fname, '_before_slicetiming')
if ~(exist([base_dir_sub fname '_before_slicetiming' '.nii'], 'file'))

refslice = 0; % The topmost slice is considered the first slice to be acquired
scfMRItb_04_unzipFile(base_dir_sub, fname, '')
filename = [base_dir_sub fname '.nii'];
scfMRItb_04_unzipFile(base_dir_sub, fname, '')
F = load_untouch_nii(filename);
TR = F.hdr.dime.pixdim(1,5); % volume acquisition time (repetition time)
num_slices = F.hdr.dime.dim(4); % number of slices
TA = TR - (TR/num_slices); % time at which the last slice was acquired
timing(1) = TA/(num_slices-1);
timing(2) = TR-TA;

% ----- SLICE TIMING CORRECTION ----------------------------------

if use_SPM==1
    try
    spm_slice_timing(filename, sliceorder, refslice, timing)
    
    catch % in case SPM is not installed, try AFNI
        if mod(length(sliceorder),2)~=0, sliceorder = sliceorder(end-1); end
        ww = sliceorder(round(length(sliceorder)/2)+1:end)-sliceorder(1:round(length(sliceorder)/2));
        if (((sum(ww(2:end)-ww(1:end-1)))==0) && (mean(ww)==1) && ((sum(sliceorder(2:end)-sliceorder(1:end-1)))>0)) % e.g., [1 3 5 7 9 2 4 6 8 10]
            unix(['3dTshift -tzero 0 -tpattern altplus -wsinc5 -prefix a ' filename])
        elseif (((sum(ww(2:end)-ww(1:end-1)))==0) && (mean(ww)==1) && ((sum(sliceorder(2:end)-sliceorder(1:end-1)))<0)) % e.g., [9 7 5 3 1 10 8 6 4 2]
            unix(['3dTshift -tzero 0 -tpattern altminus -wsinc5 -prefix a ' filename])
        elseif (((sum(ww(2:end)-ww(1:end-1)))==0) && (mean(ww)>0)) % e.g., [1 2 3 4 5 6 7 8 9 10]
            unix(['3dTshift -tzero 0 -tpattern seqplus -wsinc5 -prefix a ' filename])
        elseif (((sum(ww(2:end)-ww(1:end-1)))==0) && (mean(ww)<0)) % e.g., [10 9 8 7 6 5 4 3 2 1]
            unix(['3dTshift -tzero 0 -tpattern seqminus -wsinc5 -prefix a ' filename])
        else % read about the function here: https://afni.nimh.nih.gov/pub/dist/doc/program_help/3dTshift.html
            fprintf('Slice timing correction was not performed (in AFNI) due to confusion about slice order. Please check scfMRItb_02_slicetimingcorrection.m')
        end
    end
    
else
    if mod(length(sliceorder),2)~=0, sliceorder = sliceorder(end-1); end
    ww = sliceorder(round(length(sliceorder)/2)+1:end)-sliceorder(1:round(length(sliceorder)/2));
    if (((sum(ww(2:end)-ww(1:end-1)))==0) && (mean(ww)==1) && ((sum(sliceorder(2:end)-sliceorder(1:end-1)))>0)) % e.g., [1 3 5 7 9 2 4 6 8 10]
        unix(['3dTshift -tzero 0 -tpattern altplus -wsinc5 -prefix a ' filename])
    elseif (((sum(ww(2:end)-ww(1:end-1)))==0) && (mean(ww)==1) && ((sum(sliceorder(2:end)-sliceorder(1:end-1)))<0)) % e.g., [9 7 5 3 1 10 8 6 4 2]
        unix(['3dTshift -tzero 0 -tpattern altminus -wsinc5 -prefix a ' filename])
    elseif (((sum(ww(2:end)-ww(1:end-1)))==0) && (mean(ww)>0)) % e.g., [1 2 3 4 5 6 7 8 9 10]
        unix(['3dTshift -tzero 0 -tpattern seqplus -wsinc5 -prefix a ' filename])
    elseif (((sum(ww(2:end)-ww(1:end-1)))==0) && (mean(ww)<0)) % e.g., [10 9 8 7 6 5 4 3 2 1]
        unix(['3dTshift -tzero 0 -tpattern seqminus -wsinc5 -prefix a ' filename])
    else % read about the function here: https://afni.nimh.nih.gov/pub/dist/doc/program_help/3dTshift.html
        fprintf('Slice timing correction was not performed (in AFNI) due to confusion about slice order. Please check scfMRItb_02_slicetimingcorrection.m')
    end
end

movefile(filename,[base_dir_sub fname '_before_slicetiming' '.nii'],'f')
movefile([base_dir_sub 'a' fname '.nii'],filename,'f')

end

end
```
