## Supplementary material for "Neptune: a toolbox for spinal cord functional MRI data processing and quality assurance": Neptune software: scfMRItb_03_matchAnatFuncSlices.m

```
function [notunix, noAFNI] = scfMRItb_03_matchAnatFuncSlices(base_dir_sub, fname, fname_anat_orig)

% ----- REMOVE TOP & BOTTOM SLICES FROM ANATOMICAL IF TOO MANY ------------
% assumptions: if there is an unequal number of slices, then the anatomical
% will have more slices than the functionals. also, there will be an even
% number of extra slices (so an equal number is removed from top & bottom)

% add_AFNI_to_path
fname_anat = [fname_anat_orig '_reduced'];

scfMRItb_04_unzipFile(base_dir_sub, fname_anat_orig, '')
scfMRItb_04_unzipFile(base_dir_sub, fname, '')

A = load_untouch_nii([base_dir_sub fname_anat_orig '.nii']);
F = load_untouch_nii([base_dir_sub fname '.nii']);
reslice_anat_func = 2; % 1 = reslice the anatomical to the functional slice thickness. 2 = reslice the functional to the anatomical slice thickness


%% Reslice to match the slice thickness
if reslice_anat_func==1 % 1 = reslice the anatomical to the functional slice thickness
    % If slice thickness is different between anatomical and functional, then
    %  the anatomical will be resliced to the slice thickness of the functional.
    %  This is necessary for the toolbox to function as intended
    if A.hdr.dime.pixdim(4)~=F.hdr.dime.pixdim(4)
        scfMRItb_03_reslice_anat(base_dir_sub,fname_anat_orig,F.hdr.dime.pixdim(4))
        clear A
        A = load_untouch_nii([base_dir_sub fname_anat_orig '.nii']);
    end
    
elseif reslice_anat_func==2 % 2 = reslice the functional to the anatomical slice thickness
    % If slice thickness is different between anatomical and functional, then
    %  the functional will be resliced to the slice thickness of the anatomical.
    %  This is necessary for the toolbox to function as intended
    if A.hdr.dime.pixdim(4)~=F.hdr.dime.pixdim(4)
        scfMRItb_03_reslice_func(base_dir_sub,fname,A.hdr.dime.pixdim(4))
        clear F
        F = load_untouch_nii([base_dir_sub fname '.nii']);
    end
    
end

%%
num_slices_A = size(A.img,3);
num_slices_F = size(F.img,3);
clear A F;

if num_slices_A > num_slices_F
    if rem(num_slices_A - num_slices_F,2)==0 % if number of more slices in the anat is an even number
    z_shift = (num_slices_A - num_slices_F) / 2;
    z1 = z_shift;
    z2 = num_slices_A-z_shift-1;
    notunix = unix(['rm -f ' base_dir_sub fname_anat '.nii']);    
    noAFNI = unix(['3dZcutup -keep ' num2str(z1) ' ' num2str(z2) ' ' ...
        '-prefix ' base_dir_sub fname_anat '.nii' ' ' ...
        base_dir_sub fname_anat_orig '.nii']);
    clear z1 z2 z_shift;
    
    else % if number of more slices in the anat is an odd number
    z_shift = (num_slices_A - num_slices_F) / 2;
    z1 = z_shift+0.5;
    z2 = num_slices_A-(z_shift-0.5)-1;
    notunix = unix(['rm -f ' base_dir_sub fname_anat '.nii']);    
    noAFNI = unix(['3dZcutup -keep ' num2str(z1) ' ' num2str(z2) ' ' ...
        '-prefix ' base_dir_sub fname_anat '.nii' ' ' ...
        base_dir_sub fname_anat_orig '.nii']);
    clear z1 z2 z_shift;    
    end
    
elseif num_slices_A < num_slices_F
    if rem(num_slices_F - num_slices_A,2)==0 % if number of more slices in the anat is an even number
    z_shift = (num_slices_F - num_slices_A) / 2;
    z1 = z_shift;
    z2 = num_slices_F-z_shift-1;
    notunix = unix(['rm -f ' base_dir_sub fname '_reduced' '.nii']);    
    noAFNI = unix(['3dZcutup -keep ' num2str(z1) ' ' num2str(z2) ' ' ...
        '-prefix ' base_dir_sub fname '_reduced' '.nii' ' ' ...
        base_dir_sub fname '.nii']);
    clear z1 z2 z_shift;
    movefile([base_dir_sub fname '.nii'],[base_dir_sub fname '_beforeReduced_ORIGINAL' '.nii'],'f')
    unix(['rm -f ' base_dir_sub fname '.nii']);
    movefile([base_dir_sub fname '_reduced' '.nii'],[base_dir_sub fname '.nii'],'f')
    
    else % if number of more slices in the anat is an odd number
    z_shift = (num_slices_F - num_slices_A) / 2;
    z1 = z_shift+0.5;
    z2 = num_slices_F-(z_shift-0.5)-1;
    notunix = unix(['rm -f ' base_dir_sub fname '_reduced' '.nii']);
    noAFNI = unix(['3dZcutup -keep ' num2str(z1) ' ' num2str(z2) ' ' ...
        '-prefix ' base_dir_sub fname '_reduced' '.nii' ' ' ...
        base_dir_sub fname '.nii']);
    clear z1 z2 z_shift;    
    movefile([base_dir_sub fname '.nii'],[base_dir_sub fname '_beforeReduced_ORIGINAL' '.nii'],'f')
    unix(['rm -f ' base_dir_sub fname '.nii']);
    movefile([base_dir_sub fname '_reduced' '.nii'],[base_dir_sub fname '.nii'],'f')
    end    
    copyfile([base_dir_sub fname_anat_orig '.nii'], [base_dir_sub fname_anat '.nii'], 'f')
    
else
    copyfile([base_dir_sub fname_anat_orig '.nii'], [base_dir_sub fname_anat '.nii'], 'f')
    notunix = []; noAFNI = [];
end
clear num_slices_*;

end
```
