## Supplementary material for "Neptune: a toolbox for spinal cord functional MRI data processing and quality assurance": Neptune software: scfMRItb_03_reslice_anat.m

```
function scfMRItb_03_reslice_anat(base_dir_sub, fname_anat_orig, S_func)

% If slice thickness is different between anatomical and functional, then
%  the anatomical will be resliced to the slice thickness of the functional.
%  This is necessary for the toolbox to function as intended

scfMRItb_04_unzipFile(base_dir_sub, fname_anat_orig, '')
A = load_untouch_nii([base_dir_sub fname_anat_orig '.nii']);
dx = A.hdr.dime.pixdim(2);
dy = A.hdr.dime.pixdim(3);
dz = S_func;
fname_anat_resliced = [fname_anat_orig '_resliced'];

unix(['3dresample -dxyz ' num2str(dx) ' ' num2str(dy) ' ' num2str(dz) ' -prefix ' base_dir_sub fname_anat_resliced '.nii' ' -input ' base_dir_sub fname_anat_orig '.nii'])

movefile([base_dir_sub fname_anat_orig '.nii'],[base_dir_sub fname_anat_orig '_beforeReslice_ORIGINAL' '.nii'],'f')
unix(['rm -f ' base_dir_sub fname_anat_orig '.nii']);
movefile([base_dir_sub fname_anat_orig '_resliced' '.nii'],[base_dir_sub fname_anat_orig '.nii'],'f')

end
```
