## Supplementary material for "Neptune: a toolbox for spinal cord functional MRI data processing and quality assurance": Neptune software: scfMRItb_04_splitData.m

```
function [notunix, noAFNI] = scfMRItb_04_splitData(base_dir_sub,fname,fname_anat_orig,varargin)

fname_anat = [fname_anat_orig '_reduced'];

if nargin<4
    wbar3 = waitbar(0,'04. Split data...','Name','Progress(04): Split data...','Units','normalized','Position',[3/5 0 1/5 1/17]);
else
    wbar3 = varargin{1};
end

%% ----- SPLIT UP DATA -----------------------------------------------------
% ... and save each slice separately

scfMRItb_04_unzipFile(base_dir_sub, fname_anat, '')
A = load_untouch_nii([base_dir_sub fname_anat '.nii']); % anatomical
datatype_flag_A = 0;
siz3 = size(A.img,3);
if A.hdr.dime.datatype<16 || A.hdr.dime.bitpix<32 % if data type is integer (uint8, int16 or int32) then convert to single
    A.hdr.dime.datatype = 16; % 2 uint8; 4 int16; 8 int32; 16 single; 32 double (float32); 64 double (float64)
    A.hdr.dime.bitpix = 32; % 8 uint8; 16 int16; 32 int32; 32 single; 64 double (float32); 64 double (float64)
    A.img = single(A.img);
    save_untouch_nii(A,[base_dir_sub fname_anat '_tempp' '.nii']);
    datatype_flag_A = 1;
end
clear A

datatype_flag_F = 0;
scfMRItb_04_unzipFile(base_dir_sub, fname, '')
data = load_untouch_nii([base_dir_sub fname '.nii']);
if data.hdr.dime.datatype<16 || data.hdr.dime.bitpix<32 % if data type is integer (uint8, int16 or int32) then convert to single
    data.hdr.dime.datatype = 16; % 2 uint8; 4 int16; 8 int32; 16 single; 32 double (float32); 64 double (float64)
    data.hdr.dime.bitpix = 32; % 8 uint8; 16 int16; 32 int32; 32 single; 64 double (float32); 64 double (float64)
    data.img = single(data.img);
    save_untouch_nii(data,[base_dir_sub fname '_tempp' '.nii']);
    datatype_flag_F = 1;
end
clear data


for i3 = 1 : siz3
    try waitbar(0.5*(i3/siz3),wbar3,sprintf('04. Split data: slice (%d) of (%d)',i3,siz3)); catch, end
    fprintf('04. Split data: slice (%d) of (%d)\n',i3,siz3)
        
    % anatomical
    notunix = unix(['rm -f ' base_dir_sub fname_anat '_slice' num2str(i3) '.nii']);
    if datatype_flag_A==0
    noAFNI = unix(['3dZcutup -keep ' num2str(i3-1) ' ' num2str(i3-1) ' ' ...
        '-prefix ' base_dir_sub fname_anat '_slice' num2str(i3) '.nii' ' ' ...
        base_dir_sub fname_anat '.nii']);
    elseif datatype_flag_A==1
    noAFNI = unix(['3dZcutup -keep ' num2str(i3-1) ' ' num2str(i3-1) ' ' ...
        '-prefix ' base_dir_sub fname_anat '_slice' num2str(i3) '.nii' ' ' ...
        base_dir_sub fname_anat '_tempp' '.nii']);
    end
    
    % functional
    if datatype_flag_F==0
    unix(['3dZcutup -keep ' num2str(i3-1) ' ' num2str(i3-1) ' ' ...
        '-prefix ' base_dir_sub fname '_slice' num2str(i3) '.nii' ' ' ...
        base_dir_sub fname '.nii']);
    elseif datatype_flag_F==1
    unix(['3dZcutup -keep ' num2str(i3-1) ' ' num2str(i3-1) ' ' ...
        '-prefix ' base_dir_sub fname '_slice' num2str(i3) '.nii' ' ' ...
        base_dir_sub fname '_tempp' '.nii']);        
    end
end

if datatype_flag_A==1
    delete([base_dir_sub fname_anat '_tempp' '.nii'])
end
if datatype_flag_F==1
    delete([base_dir_sub fname '_tempp' '.nii'])
end


if nargin<4
    close(wbar3)
end

end
```
