## Supplementary material for "Neptune: a toolbox for spinal cord functional MRI data processing and quality assurance": Neptune software: scfMRItb_04_unzipFile.m

```
function scfMRItb_04_unzipFile(base_dir_sub, fname, suffix)

% if .nii file doesn't exist but .nii.gz exists then unzip it
if ~exist([base_dir_sub fname suffix '.nii'],'file') && exist([base_dir_sub fname suffix '.nii.gz'],'file')
    gunzip([base_dir_sub fname suffix '.nii.gz'])
end

end
```
