## Supplementary material for "Neptune: a toolbox for spinal cord functional MRI data processing and quality assurance": Neptune software: scfMRItb_05_notCordMask.m

```
function scfMRItb_05_notCordMask(base_dir_sub,fname,varargin)

scrsz = get(0,'ScreenSize');
pause on;
fname3 = [fname '_mean'];

if nargin<6
    exist_SCT = 1; % 1 = spinal cord toolbox exists, 0 = doesn't exist
else
    exist_SCT = varargin{4};
end
if nargin<4
    save_videos = 1; % 1 = save QC videos (generates sizeable .mp4 files and takes a few minutes to do it). This is turned on by default, but if you don't want to save videos then set it to 0
else
    save_videos = varargin{2};
end
if nargin<5
    Rns = 0; % 0 = don't create runX subfolder; X = create runX subfolder
else
    Rns = varargin{3};
end
if Rns==0, run=1; else, run=Rns; end

if ~(exist([base_dir_sub 'QC'],'dir'))
    mkdir([base_dir_sub 'QC'])
end
if ~(exist([base_dir_sub 'QC' '/05_notCordMask'],'dir'))
    mkdir([base_dir_sub 'QC' '/05_notCordMask'])
end
if Rns==0
    QCpath2 = [base_dir_sub 'QC' '/05_notCordMask/'];
else
    if ~(exist([base_dir_sub 'QC' '/05_notCordMask' '/run' num2str(Rns)],'dir'))
        mkdir([base_dir_sub 'QC' '/05_notCordMask' '/run' num2str(Rns)])
    end
    QCpath2 = [base_dir_sub 'QC' '/05_notCordMask' '/run' num2str(Rns) '/'];
end

if ~(exist([base_dir_sub 'QC' '/04_splitData_rawQC'],'dir'))
    mkdir([base_dir_sub 'QC' '/04_splitData_rawQC'])
end
if Rns==0
    QCpath = [base_dir_sub 'QC' '/04_splitData_rawQC/'];
else
    if ~(exist([base_dir_sub 'QC' '/04_splitData_rawQC' '/run' num2str(Rns)],'dir'))
        mkdir([base_dir_sub 'QC' '/04_splitData_rawQC' '/run' num2str(Rns)])
    end
    QCpath = [base_dir_sub 'QC' '/04_splitData_rawQC' '/run' num2str(Rns) '/'];
end

% if ~(exist([base_dir_sub 'QC' '/01_NORDIC_denoising'],'dir'))
%     mkdir([base_dir_sub 'QC' '/01_NORDIC_denoising'])
% end
if Rns==0
    QCpath3 = [base_dir_sub 'QC' '/01_NORDIC_denoising/'];
else
%     if ~(exist([base_dir_sub 'QC' '/01_NORDIC_denoising' '/run' num2str(Rns)],'dir'))
%         mkdir([base_dir_sub 'QC' '/01_NORDIC_denoising' '/run' num2str(Rns)])
%     end
    QCpath3 = [base_dir_sub 'QC' '/01_NORDIC_denoising' '/run' num2str(Rns) '/'];
end

if ~(exist([base_dir_sub 'QC' '/QC_shortlist'],'dir'))
    mkdir([base_dir_sub 'QC' '/QC_shortlist'])
end
if Rns==0
    QCpath4 = [base_dir_sub 'QC' '/QC_shortlist/'];
else
    if ~(exist([base_dir_sub 'QC' '/QC_shortlist' '/run' num2str(Rns)],'dir'))
        mkdir([base_dir_sub 'QC' '/QC_shortlist' '/run' num2str(Rns)])
    end
    QCpath4 = [base_dir_sub 'QC' '/QC_shortlist' '/run' num2str(Rns) '/'];
end

unix(['rm -f ' base_dir_sub fname3 '.nii']);
unix(['3dTstat -mean -prefix ' base_dir_sub fname3 '.nii ' base_dir_sub fname '.nii' ]);

%% Use SCT (if available) to get an estimate of CSF boundary

if exist_SCT==1 % this works on, at least, SCT v5.4 and later (I tested it with SCT v5.4 and v5.5 only)
    try
    fprintf('\nRunning SCT (if required) to estimate the boundary of the CSF...\n\n')
    if ~(exist([base_dir_sub 'QC' '/SCT'],'dir'))
        mkdir([base_dir_sub 'QC' '/SCT'])
    end
    if ~exist([base_dir_sub fname3 '_CSF_seg.nii'],'file')
        scfMRItb_04_unzipFile(base_dir_sub, fname3, '')
        unix(['sct_propseg -i ' base_dir_sub fname3 '.nii' ' -c t2s -ofolder ' base_dir_sub ' -CSF' ' -qc ' base_dir_sub 'QC/SCT'])
    end
    CSF_seg = load_untouch_nii([base_dir_sub fname3 '_CSF_seg.nii']); CSF_seg=CSF_seg.img;
    for i3 = 1 : size(CSF_seg,3)
        CSF_seg(:,:,i3) = imfill(CSF_seg(:,:,i3),'holes');
        CSF_seg(:,:,i3) = imdilate(CSF_seg(:,:,i3), strel('disk', 3));
        CSF_boundarymask = boundarymask(rot90(CSF_seg(:,:,i3)));
        [row,col] = find(CSF_boundarymask==1);
        if ~isempty(row)
        CSF_boundary{i3,1} = bwtraceboundary(CSF_boundarymask,[row(1),col(1)],'N');
        fctr = round(size(CSF_boundary{i3,1},1)/12);
        CSF_boundary{i3,1} = CSF_boundary{i3,1}(1:fctr:end,:);
        else
        CSF_boundary{i3,1} = [];
        end
        clear CSF_boundarymask row col fctr
    end
    clear CSF_seg
    fprintf('\nCompleted sct_propseg to estimate the boundary of the CSF.\n\n')
    catch
    end
end

%% ----- DEFINE NOT-CORD MASK ----------------------------------------------

if ~(exist([base_dir_sub fname '_mask_NS.mat'], 'file'))

if nargin<3
    wbar3 = waitbar(0,'05. Define "not cord" mask...','Name','Progress(05): "Not-cord" mask...','Units','normalized','Position',[3/5 0 1/5 1/17]);
else
    wbar3 = varargin{1};
end

scfMRItb_04_unzipFile(base_dir_sub, fname3, '')
A = load_untouch_nii([base_dir_sub fname3 '.nii']); % mean functional
mask_NS  = A;
siz3 = size(A.img,3);
new_size = round(size(A.img,1) / 2);

for i3 = 1 : siz3
    try waitbar((i3/siz3),wbar3,sprintf('05. Define "not-cord" mask: slice (%d) of (%d)',i3,siz3)); catch, end
    img = rot90(A.img(:,:,i3));
    temp = img(round((size(img,1)-new_size)/2)+1:end-round((size(img,1)-new_size)/2), ...
        round((size(img,2)-new_size)/2)+1:end-round((size(img,2)-new_size)/2));
    mmax = max(temp(:)); clear temp;
    fighndl = figure('Position',[round((scrsz(3)-scrsz(4))/2) 1 scrsz(4) scrsz(4)],'Color',[0.8,0.89,0.84],'InvertHardcopy','off'); try addToolbarExplorationButtons(fighndl); catch, end % [0.85,0.90,0.85] [361 1 scrsz(4) scrsz(4)]
    imshow(single(img), [0 mmax], 'InitialMagnification', 'fit'); colorbar;
    fighndl.Color = [0.8, 0.87, 0.84] + [0, 0.02*(1-exp(siz3/i3)/exp(siz3)), 0.1*exp(i3/siz3)/exp(1)]; % [0.85, 0.90, 0.85] + [0, 0, 0.1*i3/siz3];
    title({'Draw a boundary selecting only the spinal cord (double click the cord once done)';['Slice ' num2str(i3) ' of ' num2str(siz3)]},'Color',[i3/(5*siz3),0.5,i3/siz3],'FontSize',16);
    disp(['Slice ' num2str(i3) ' of ' num2str(siz3) ': Draw a boundary selecting only the spinal cord.']);
    if i3~=1
        xlim([lims_x1_(i3-1),lims_x2_(i3-1)]), ylim([lims_y1_(i3-1),lims_y2_(i3-1)]),
    end
    
    if exist('CSF_boundary','var') && ~isempty(CSF_boundary{i3,1})
        h = impoly(gca,[CSF_boundary{i3}(:,2),CSF_boundary{i3}(:,1)]);
    else
        h = impoly;
    end
    position = wait(h);
    bmask = poly2mask(position(:,1), position(:,2), size(A.img,2), size(A.img,1));
    bmask = ~bmask;
    mask_NS.img(:,:,i3) = rot90(bmask, -1);
    position_maskNS{i3} = position;
    saveas(fighndl,[QCpath2 'notCordMask_slice' sprintf('%.2d',i3) '.jpg'])
    close(fighndl)
    lims_x1(i3) = min(position(:,1)); lims_x2(i3) = max(position(:,1)); %#ok<*AGROW>
    lims_x1_(i3) = round(lims_x1(i3) - 0.5*(lims_x2(i3)-lims_x1(i3))); lims_x2_(i3) = round(lims_x2(i3) + 0.5*(lims_x2(i3)-lims_x1(i3)));
    lims_y1(i3) = min(position(:,2)); lims_y2(i3) = max(position(:,2));
    lims_y1_(i3) = round(lims_y1(i3) - 0.5*(lims_y2(i3)-lims_y1(i3))); lims_y2_(i3) = round(lims_y2(i3) + 0.5*(lims_y2(i3)-lims_y1(i3)));
end

eval(['save ' base_dir_sub fname '_mask_NS' '.mat mask_NS position_maskNS;']);

for i3 = 1 : siz3
    scfMRItb_04_resplitData(base_dir_sub, fname, '', siz3) % if slice-wise data doesn't exist but combined 4D data exists, then split the data to generate the needed slice-wise data
    scfMRItb_04_unzipFile(base_dir_sub, fname, ['_slice' num2str(i3)])
    func = load_untouch_nii([base_dir_sub fname '_slice' num2str(i3) '.nii']); func.img=[];
    unix(['rm -f ' base_dir_sub fname '_slice' num2str(i3) '_Smask.nii']);
    func.img = ~mask_NS.img(:,:,i3);
    func.hdr.dime.dim(1)=3; func.hdr.dime.dim(5)=1;
    func.hdr.dime.dim(2)=size(func.img,1); func.hdr.dime.dim(3)=size(func.img,2); func.hdr.dime.dim(4)=size(func.img,3);
    save_untouch_nii(func,  [base_dir_sub fname '_slice' num2str(i3) '_Smask.nii']);
    clear func
end

clear mask_NS position_maskNS new_size img temp mmax i3 func
end % if

%% Compute SNR
% Raw data SNR is computed as the ratio of signal variance in the cord
% (median value) to the signal variance outside the cord (median value)
% within a given slice, computed from raw NIfTI data.

load([base_dir_sub fname '_mask_NS' '.mat'])
mask_NS = mask_NS.img;
scfMRItb_04_unzipFile(base_dir_sub, fname, '')
F = load_untouch_nii([base_dir_sub fname '.nii']);
F = single(F.img);

if ~exist('siz3','var')
    scfMRItb_04_unzipFile(base_dir_sub, fname3, '')
    A = load_untouch_nii([base_dir_sub fname3 '.nii']); % mean functional
    siz3 = size(A.img,3);
end

for i3=1:siz3
    fprintf('05. Additional computations: slice (%d) of (%d)\n',i3,siz3)
    for w=1:size(F,4)
        temp = F(:,:,i3,w);
        ts_S(:,w)  = temp(find(mask_NS(:,:,i3)==0)); % spine
        ts_NS(:,w) = temp(find(mask_NS(:,:,i3)==1)); % not spine
    end
    if ~isempty(ts_S) && ~isempty(ts_NS)
    for w=1:size(ts_S,1)
        var1(w,1) = nanvar(ts_S(w,:)'); %#ok<*UDIM>
    end
    for w=1:size(ts_NS,1)
        var2(w,1) = nanvar(ts_NS(w,:)');
    end
    SNR(i3,1) = median(var1)/median(var2);
    else
    SNR(i3,1) = NaN;
    end
    clear ts_S ts_NS var1 var2 temp
end

SNR_in_dB = 10.*log(SNR);
info_SNR = sprintf('SNR = signal-to-noise ratio defined as the ratio of signal variance in the cord (median value) to the signal variance outside the cord (median value) within a given slice, computed from raw NIfTI data.\nThat is, SNR = median(var(timeseries_cord)) / median(var(timeseries_notcord));\nSNR_in_dB = 10.*log(SNR);\n');
save([base_dir_sub 'SNR_slicewise_run' num2str(run) '.mat'], 'SNR','SNR_in_dB','info_SNR');

% QC plot
for w=1:length(SNR)
    dispstring1{w} = sprintf('%.2f',SNR(w));
    dispstring2{w} = sprintf('%.2f dB',SNR_in_dB(w));
end; clear w
fighndl = figure('Position',[1 1 1280 800],'Color',[0.85,0.90,0.85],'InvertHardcopy','off','Visible','off'); % red [0.95,0.9,0.85], green [0.85,0.90,0.85], blue [0.89,0.94,0.99]
subplot2n(2,1,1), plot(SNR,'--o','color',[0.25,0.25,0.25],'linewidth',2.25, 'MarkerSize',14, 'MarkerEdgeColor','k', 'MarkerFaceColor',[0.85,0.85,0.85]),
    grid on,grid minor, ylabel('Median signal-to-noise ratio'), xlabel('SLICES'), xticks([1:length(SNR)]),
    xlim([0.5 length(SNR)+0.5]), ylim([0 1.2*max(SNR)])
    text([1:length(SNR)]-0.15, double(SNR'+0.1*max(SNR)), dispstring1,'Color',[0.05,0.05,0.05],'FontSize',12)
    legend({'Raw data SNR'},'FontSize',14,'TextColor','k','Location','best')
    title(sprintf('BOLD signal-to-noise ratio (SNR) across slices, defined as:\nthe ratio of BOLD signal variance in the cord (median value) to the BOLD signal variance outside the cord (median value)'),'Color',[0.4,0.1,0.1],'FontSize',16)
subplot2n(2,1,2), plot(SNR_in_dB,'--o','color',[0.25,0.25,0.25],'linewidth',2.25, 'MarkerSize',14, 'MarkerEdgeColor',[0.4,0.1,0.1], 'MarkerFaceColor',[0.85,0.75,0.75]),
    grid on,grid minor,
    ylabel('Median SNR in decibels'), xlabel('SLICES'), xticks([1:length(SNR_in_dB)]),
    xlim([0.5 length(SNR_in_dB)+0.5]), ylim([min([0,1.2*min(SNR_in_dB)]) 1.2*max(SNR_in_dB)])
    text([1:length(SNR_in_dB)]-0.25, double(SNR_in_dB'+0.1*max(SNR_in_dB)), dispstring2,'Color',[0.16,0.04,0.04],'FontSize',12)
    legend({'Raw data SNR in decibels'},'FontSize',14,'TextColor',[0.16,0.04,0.04],'Location','best')
    title(sprintf('BOLD SNR across slices in decibels:\nSNR in dB = 10 x log(SNR)'),'Color',[0.4,0.1,0.1],'FontSize',16)
saveas(fighndl,[QCpath 'SNR_graph_slicewise' '.jpg'])

close(fighndl)
clear mask_NS F info_SNR SNR SNR_in_dB dispstring1 dispstring2

%% time series plots, global signal and FC histogram
scrsz = get(0,'ScreenSize'); pause on;
cmap = colormap('bone'); cmap = (1-exp(-cmap))./(1-exp(-1));
if ~exist('siz3','var')
    A = load_untouch_nii([base_dir_sub fname3 '.nii']); % mean functional
    siz3 = size(A.img,3);
end

Smask_err=0;
try
    if exist([base_dir_sub fname '_Smask.nii'],'file')
    scfMRItb_04_resplitData(base_dir_sub, fname, '_Smask', siz3) % if slice-wise data doesn't exist but combined 4D data exists, then split the data to generate the needed slice-wise data
    else
	zzz 
    end
catch
    load([base_dir_sub fname '_mask_NS' '.mat'])
    Smask1_ = ~mask_NS.img; clear mask_NS
    Smask_err=1;
end

F2_=[]; mF2_=[]; C_=[];
for i3 = 1 : siz3
    try waitbar((i3/siz3),wbar3,sprintf('05. QC plots: slice (%d) of (%d)',i3,siz3)); catch, end
    fprintf('05. QC plots: slice (%d) of (%d)\n',i3,siz3)
    scfMRItb_04_resplitData(base_dir_sub, fname, '', siz3) % if slice-wise data doesn't exist but combined 4D data exists, then split the data to generate the needed slice-wise data
    F1 = load_untouch_nii([base_dir_sub fname '_slice' num2str(i3) '.nii']);
    TR = F1.hdr.dime.pixdim(5); F1 = single(squeeze(F1.img));
    
    if Smask_err==0
        scfMRItb_04_resplitData(base_dir_sub, fname, '_Smask', siz3) % if slice-wise data doesn't exist but combined 4D data exists, then split the data to generate the needed slice-wise data
        scfMRItb_04_unzipFile(base_dir_sub, fname, ['_slice' num2str(i3) '_Smask'])
        mask1 = load_untouch_nii([base_dir_sub fname '_slice' num2str(i3) '_Smask.nii']); mask1 = mask1.img;
    else
        mask1 = Smask1_(:,:,i3);
    end
    [m1,m2] = find(mask1==1); Mm1=median(m1); Mm2=median(m2);
    sdist = sqrt((m1-Mm1).^2 + (m2-Mm2).^2);
    clear im Mm1 Mm2 m1 m2
    for i5=1:size(F1,3)
        tmp = F1(:,:,i5);
        F2(:,i5) = tmp(find(mask1==1)); clear tmp
    end; clear i5 F1
    [~,svar] = sort(var(F2,0,2)); [~,sdist] = sort(sdist); sortorder{i3,1} = sdist; %#ok<*AGROW> % sort timeseries based on distance from the center of the cord (if you want to sort by signal variance then use svar)
    
    fighndl = figure('Position',[1 1 1152 720],'Color',[0.95,0.9,0.85],'InvertHardcopy','off','Visible','off'); % red [0.95,0.9,0.85], green [0.85,0.90,0.85], blue [0.89,0.94,0.99]
    imagesc(F2(sortorder{i3},:)),colormap(cmap),colorbar, xlabel('time (in samples)'),ylabel('voxels (sorted by distance from the center of the cord; top row is nearest)'),
        title(['Time series within GM, WM and CSF in UNPROCESSED RAW DATA, in slice ' num2str(i3) ' of ' num2str(siz3)],'Color',[0.65,0.11,0.19],'FontSize',14);
    saveas(fighndl,[QCpath 'fMRI_ts_rawdata_slice' sprintf('%.2d',i3) '.jpg'])
    close(fighndl)
    
    mF2=nanmean(F2,1); mF2 = 100*(mF2-mean(mF2))./mean(mF2);
    fighndl = figure('Position',[1 1 1152 720],'Color',[0.95,0.9,0.85],'InvertHardcopy','off','Visible','off'); % red [0.95,0.9,0.85], green [0.85,0.90,0.85], blue [0.89,0.94,0.99]
    plot(mF2,'color',[0.65,0.11,0.19],'linewidth',1.5), grid on,grid minor, xlim([1 length(mF2)]),ylim([1.25.*min(mF2) 1.25.*max(mF2)]), xlabel('time (in samples)'),ylabel('percentage signal change'),
      title(['Global mean signal in unprocessed raw data (slice ' num2str(i3) ' of ' num2str(siz3) ')'],'Color',[0.65,0.11,0.19],'FontSize',14),
    saveas(fighndl,[QCpath 'global_signal_rawdata_slice' sprintf('%.2d',i3) '.jpg'])
    close(fighndl)
    
    C = corrcoef(F2'); C=uppertriangle(C);
    fighndl = figure('Position',[1 1 1152 720],'Color',[0.95,0.9,0.85],'InvertHardcopy','off','Visible','off'); % red [0.95,0.9,0.85], green [0.85,0.90,0.85], blue [0.89,0.94,0.99]
    histogram(C,'EdgeColor','none','Normalization','probability'); grid on,grid minor, xlim([-1 1]),  % xlim([-max(abs(C)) max(abs(C))]),
        hold on, xline(0,'Color','k','linewidth',1.5); hold off
        title(sprintf('Histogram of voxel-to-voxel correlations in unprocessed raw data (slice %d of %d)\n%.1f%% of the correlations were > 0  [ mean=%.2f, median=%.2f, SD=%.2f ]',i3,siz3,100*length(find(C>0))/length(C),mean(C),median(C),std(C)),'Color',[0.65,0.11,0.19],'FontSize',14)
    saveas(fighndl,[QCpath 'hist_FC_rawdata_slice' sprintf('%.2d',i3) '.jpg'])
    close(fighndl)
    
    F2_=cat(1,F2_,F2); mF2_=cat(1,mF2_,mF2); C_=cat(1,C_,C);
    clear F2 TR mask1 svar sdist mF2 C
end


[~,svar] = sort(var(F2_,0,2)); [~,spsc] = sort(meanPSC(F2_));
sortorder2 = sortorder; sortorder=[]; f=0; %#ok<*ASGLU>
for im=1:length(sortorder2)
    temp = sortorder2{im}; temp = temp + f; f = f + length(temp);
    sortorder = cat(1,sortorder,temp); clear temp
end; clear im f
sortorder = spsc; % options: spsc for percentage signal change, svar for variance, (comment out the line if you want to use distance from the center of the cord separately for each slice)
for im=max(sortorder):-1:1, if isempty(find(sortorder==im)), sortorder(find(sortorder>im)) = sortorder(find(sortorder>im))-1; end, end; clear im %#ok<*EFIND>

fighndl = figure('Position',[1 1 1152 720],'Color',[0.89,0.94,0.99],'InvertHardcopy','off','Visible','off'); % red [0.95,0.9,0.85], green [0.85,0.90,0.85], blue [0.89,0.94,0.99]
imagesc(F2_(sortorder,:)),colormap(cmap),colorbar, xlabel('time (in samples)'),ylabel('voxels (sorted by mean percentage signal change; bottom row is highest)'),
  title(['Time series within GM, WM and CSF in UNPROCESSED RAW DATA (all slices combined)'],'Color',[0.65,0.11,0.19],'FontSize',14);
saveas(fighndl,[QCpath 'fMRI_ts_rawdata_allSlices' '.jpg'])
saveas(fighndl,[QCpath4 'fMRI_ts_04rawdata_allSlices' '.jpg'])
 close(fighndl)
mF2_=nanmean(mF2_,1);
fighndl = figure('Position',[1 1 1152 720],'Color',[0.89,0.94,0.99],'InvertHardcopy','off','Visible','off'); % red [0.95,0.9,0.85], green [0.85,0.90,0.85], blue [0.89,0.94,0.99]
plot(mF2_,'color',[0.65,0.11,0.19],'linewidth',1.5), grid on,grid minor, xlim([1 length(mF2_)]),ylim([1.25.*min(mF2_) 1.25.*max(mF2_)]), xlabel('time (in samples)'),ylabel('percentage signal change'),
  title(['Global mean signal in UNPROCESSED RAW DATA (all slices averaged)'],'Color',[0.65,0.11,0.19],'FontSize',14),
saveas(fighndl,[QCpath 'global_signal_rawdata_allSlices' '.jpg'])
saveas(fighndl,[QCpath4 'global_signal_04rawdata_allSlices' '.jpg'])
 close(fighndl)
fighndl = figure('Position',[1 1 1152 720],'Color',[0.89,0.94,0.99],'InvertHardcopy','off','Visible','off'); % red [0.95,0.9,0.85], green [0.85,0.90,0.85], blue [0.89,0.94,0.99]
histogram(C_,'EdgeColor','none','Normalization','probability'); grid on,grid minor, xlim([-1 1]),
  hold on, xline(0,'Color','k','linewidth',1.5); hold off
  title(sprintf('Histogram of voxel-to-voxel correlations in UNPROCESSED RAW DATA (all slices combined)\n%.1f%% of the correlations were > 0  [ mean=%.2f, median=%.2f, SD=%.2f ]',100*length(find(C_>0))/length(C_),mean(C_),median(C_),std(C_)),'Color',[0.65,0.11,0.19],'FontSize',14)
saveas(fighndl,[QCpath 'hist_FC_rawdata_allSlices' '.jpg'])
saveas(fighndl,[QCpath4 'hist_FC_compare_04rawdata_allSlices' '.jpg'])
 close(fighndl)
clear svar sortorder spsc F2_ mF2_ C_ cmap sortorder spsc

%% Make QC video (needs audio video toolbox in Matlab)
if ~isempty(which('VideoWriter')) && (save_videos==1) % check if Matlab's audio video toolbox exists

for i3 = 1 : siz3
    try waitbar((i3/siz3),wbar3,sprintf('05. Generate QC video: slice (%d) of (%d)',i3,siz3)); catch, end
    fprintf('05. Generate QC video: slice (%d) of (%d)\n',i3,siz3)
    F1 = load_untouch_nii([base_dir_sub fname '_slice' num2str(i3) '.nii']);
    TR = F1.hdr.dime.pixdim(5); F1 = rot90(single(squeeze(F1.img)));
    video = VideoWriter([QCpath 'fMRIvideo_rawdata_slice' sprintf('%.2d',i3) '.mp4'],'MPEG-4'); %#ok<*TNMLP> % create the video object
    video.FrameRate = round(30/TR); % 30 seconds of data in every second of the video
    video.Quality = 90; % very limited compression so that the images are shown as is (and thus avoid misunderstandings due to video quality)
    open(video); % open the file for writing

    mask1 = load_untouch_nii([base_dir_sub fname '_slice' num2str(i3) '_Smask.nii']);
    mask1 = rot90(mask1.img);
    [xmask,ymask] = find(mask1==1);
    minx=round(min(xmask)-0.1*(size(mask1,1))); maxx=round(max(xmask)+0.1*(size(mask1,1)));
    miny=round(min(ymask)-0.1*(size(mask1,2))); maxy=round(max(ymask)+0.1*(size(mask1,2)));
    if (maxx-minx)<(maxy-miny)
        factr = ((maxy-miny)-(maxx-minx))/2;
        if factr==round(factr)
            maxx=maxx+factr; minx=minx-factr;
        else
            maxx=maxx+factr+0.5; minx=minx-factr+0.5;
        end
    elseif (maxy-miny)<(maxx-minx)
        factr = ((maxx-minx)-(maxy-miny))/2;
        if factr==round(factr)
            maxy=maxy+factr; miny=miny-factr;
        else
            maxy=maxy+factr+0.5; miny=miny-factr+0.5;
        end
    end
    [sizeFx,sizeFy] = size(F1(:,:,1));
    F2 = F1(minx:maxx,miny:maxy,:);
    for i5=1:size(F1,3)
        F1(:,:,i5) = imresize(F2(:,:,i5),[sizeFx sizeFy]);
    end
    clear maxx maxy minx miny xmask ymask mask1 factr sizeFx sizeFy F2 i5
    for ii=1:size(F1,3) % loop across the number of images
        I = F1(:,:,ii); %read the next image
        I(find(isnan(I)))=0;
        I = (I - min(I(:))) ./ (max(I(:)) - min(I(:)));
        fighndl = figure('Position',[round((scrsz(3)-scrsz(4))/2) 1 round(0.75*scrsz(3)) round(0.75*scrsz(4))],'Color',[0.95,0.9,0.85],'InvertHardcopy','off','Visible','off'); % [0.85,0.90,0.85]
        imshow(I, [0 max(I(:))], 'InitialMagnification', 'fit'); title({['Unprocessed raw fMRI data (slice ' num2str(i3) ' of ' num2str(siz3) ')'];['Time point ' num2str(ii) ' of ' num2str(size(F1,3))]},'Color',[0.65,0.11,0.19],'FontSize',18);
        frame = getframe(gcf);
        writeVideo(video,frame); % write the image to file
        close(fighndl)
        clear I frame
    end
    close(video); %close the file
    clear F1 video ii TR
end; clear i3

end

%% Generate QC outputs if NORDIC was performed:

if exist([base_dir_sub 'QC' '/01_NORDIC_denoising'],'dir')
    
%% NORDIC: time series plots, global signal and FC histogram
scrsz = get(0,'ScreenSize'); pause on;
cmap = colormap('bone'); cmap = (1-exp(-cmap))./(1-exp(-1)); 
if ~exist('siz3','var')
    scfMRItb_04_unzipFile(base_dir_sub, fname3, '')
    A = load_untouch_nii([base_dir_sub fname3 '.nii']); % mean functional
    siz3 = size(A.img,3);
end

Smask_err=0;
try
    if exist([base_dir_sub fname '_Smask.nii'],'file')
    scfMRItb_04_resplitData(base_dir_sub, fname, '_Smask', siz3) % if slice-wise data doesn't exist but combined 4D data exists, then split the data to generate the needed slice-wise data
    else
	zzz 
    end
catch
    load([base_dir_sub fname '_mask_NS' '.mat'])
    Smask1_ = ~mask_NS.img; clear mask_NS
    Smask_err=1;
end

F2_=[]; mF2_=[]; C_=[]; F4_=[]; mF4_=[]; C2_=[];
if exist([base_dir_sub fname '_slice' num2str(1) '_beforeNORDIC' '.nii'],'file') || exist([base_dir_sub fname '_slice' num2str(1) '_beforeNORDIC' '.nii.gz'],'file') || exist([base_dir_sub fname '_beforeNORDIC' '.nii'],'file') || exist([base_dir_sub fname '_beforeNORDIC' '.nii.gz'],'file')
    F6_=[]; mF6_=[]; C3_=[];
end
for i3 = 1 : siz3
    try waitbar((i3/siz3),wbar3,sprintf('05. QC plots (NORDIC): slice (%d) of (%d)',i3,siz3)); catch, end
    fprintf('05. QC plots (NORDIC): slice (%d) of (%d)\n',i3,siz3)
    scfMRItb_04_resplitData(base_dir_sub, fname, '_beforeNORDIC', siz3) % if slice-wise data doesn't exist but combined 4D data exists, then split the data to generate the needed slice-wise data
    scfMRItb_04_unzipFile(base_dir_sub, fname, ['_slice' num2str(i3) '_beforeNORDIC'])
    F1 = load_untouch_nii([base_dir_sub fname '_slice' num2str(i3) '_beforeNORDIC.nii']);
    TR = F1.hdr.dime.pixdim(5); F1 = single(squeeze(F1.img));
    
    if Smask_err==0
        scfMRItb_04_resplitData(base_dir_sub, fname, '_Smask', siz3) % if slice-wise data doesn't exist but combined 4D data exists, then split the data to generate the needed slice-wise data
        scfMRItb_04_unzipFile(base_dir_sub, fname, ['_slice' num2str(i3) '_Smask'])
        mask1 = load_untouch_nii([base_dir_sub fname '_slice' num2str(i3) '_Smask.nii']); mask1 = mask1.img;
    else
        mask1 = Smask1_(:,:,i3);
    end
    [m1,m2] = find(mask1==1); Mm1=median(m1); Mm2=median(m2);
    sdist = sqrt((m1-Mm1).^2 + (m2-Mm2).^2);
    clear im Mm1 Mm2 m1 m2
    for i5=1:size(F1,3)
        tmp = F1(:,:,i5);
        F2(:,i5) = tmp(find(mask1==1)); clear tmp
    end; clear i5 F1
    [~,svar] = sort(var(F2,0,2)); [~,sdist] = sort(sdist); sortorder{i3,1} = sdist; %#ok<*AGROW> % sort timeseries based on distance from the center of the cord (if you want to sort by signal variance then use svar)
    
    fighndl = figure('Position',[1 1 1152 720],'Color',[0.95,0.9,0.85],'InvertHardcopy','off','Visible','off'); % red [0.95,0.9,0.85], green [0.85,0.90,0.85], blue [0.89,0.94,0.99]
    imagesc(F2(sortorder{i3},:)),colormap(cmap),colorbar, xlabel('time (in samples)'),ylabel('voxels (sorted by distance from the center of the cord; top row is nearest)'),
        title(['Time series within GM, WM and CSF before NORDIC denoising in slice ' num2str(i3) ' of ' num2str(siz3)],'Color',[0,0.55,0.69],'FontSize',14);
    saveas(fighndl,[QCpath3 'fMRI_ts_01nordic_slice' sprintf('%.2d',i3) '.jpg'])
    close(fighndl)
    
    scfMRItb_04_resplitData(base_dir_sub, fname, '', siz3) % if slice-wise data doesn't exist but combined 4D data exists, then split the data to generate the needed slice-wise data
    scfMRItb_04_unzipFile(base_dir_sub, fname, ['_slice' num2str(i3)])
    F3 = load_untouch_nii([base_dir_sub fname '_slice' num2str(i3) '.nii']); F3 = single(squeeze(F3.img));
    for i5=1:size(F3,3)
        tmp = F3(:,:,i5);
        F4(:,i5) = tmp(find(mask1==1)); clear tmp
    end; clear i5 F3
    mF2=mean(F2,1); mF2 = 100*(mF2-mean(mF2))./mean(mF2); mF4=mean(F4,1); mF4 = 100*(mF4-mean(mF4))./mean(mF4);
    fighndl = figure('Position',[1 1 1152 720],'Color',[0.95,0.9,0.85],'InvertHardcopy','off','Visible','off'); % red [0.95,0.9,0.85], green [0.85,0.90,0.85], blue [0.89,0.94,0.99]
    subplot2n(2,1,1), plot(mF2,'color',[0.65,0.11,0.19],'linewidth',1.5), hold on, plot(mF4,'color',[0,0.55,0.69],'linewidth',0.2), xlabel('time (in samples)'),ylabel('percentage signal change'),
      grid on,grid minor, xlim([1 length(mF2)]),ylim([1.25.*min([mF2,mF4]) 1.25.*max([mF2,mF4])]), title(['Global mean signal BEFORE NORDIC denoising (slice ' num2str(i3) ' of ' num2str(siz3) ')'],'Color',[0.65,0.11,0.19],'FontSize',14), legend({'Global signal before NORDIC denoising'},'FontSize',12,'TextColor',[0.65,0.11,0.19],'Location','best')
    subplot2n(2,1,2), plot(mF4,'color',[0,0.55,0.69],'linewidth',1.5), hold on, plot(mF2,'color',[0.65,0.11,0.19],'linewidth',0.2), xlabel('time (in samples)'),ylabel('percentage signal change'),
      grid on,grid minor, xlim([1 length(mF4)]),ylim([1.25.*min([mF2,mF4]) 1.25.*max([mF2,mF4])]), title(['Global mean signal AFTER NORDIC denoising (slice ' num2str(i3) ' of ' num2str(siz3) ')'],'Color',[0,0.55,0.69],'FontSize',14), legend({'Global signal after NORDIC denoising'},'FontSize',12,'TextColor',[0,0.55,0.69],'Location','best')
    saveas(fighndl,[QCpath3 'global_signal_01nordic_slice' sprintf('%.2d',i3) '.jpg'])
    close(fighndl)

    C  = corrcoef(F4'); C  = uppertriangle(C) ;
    C2 = corrcoef(F2'); C2 = uppertriangle(C2);
    fighndl = figure('Position',[1 1 1152 720],'Color',[0.95,0.9,0.85],'InvertHardcopy','off','Visible','off'); % red [0.95,0.9,0.85], green [0.85,0.90,0.85], blue [0.89,0.94,0.99]
    subplot2n(2,1,1), histogram(C2,'EdgeColor','none','FaceColor',[0.65,0.11,0.19],'FaceAlpha',0.5,'Normalization','probability'); grid on,grid minor, xlim([-1 1]), hold on, xline(0,'Color','k','linewidth',1.5); hold off  % xlim([-max(abs([C;C2])) max(abs([C;C2]))]),
      title(sprintf('Histogram of voxel-to-voxel correlations BEFORE NORDIC denoising (slice %d of %d)\n%.1f%% of the correlations were > 0  [ mean=%.2f, median=%.2f, SD=%.2f ]',i3,siz3,100*length(find(C2>0))/length(C2),mean(C2),median(C2),std(C2)),'Color',[0.65,0.11,0.19],'FontSize',14)
    subplot2n(2,1,2), histogram(C,'EdgeColor','none','FaceColor',[0,0.55,0.69],'FaceAlpha',0.5,'Normalization','probability'); grid on,grid minor, xlim([-1 1]), hold on, xline(0,'Color','k','linewidth',1.5); hold off
      title(sprintf('Histogram of voxel-to-voxel correlations AFTER NORDIC denoising (slice %d of %d)\n%.1f%% of the correlations were > 0  [ mean=%.2f, median=%.2f, SD=%.2f ]',i3,siz3,100*length(find(C>0))/length(C),mean(C),median(C),std(C)),'Color',[0,0.55,0.69],'FontSize',14)
    saveas(fighndl,[QCpath3 'hist_FC_compare_01nordic_slice' sprintf('%.2d',i3) '.jpg'])
    close(fighndl)
        
    F2_=cat(1,F2_,F2); mF2_=cat(1,mF2_,mF2); C_=cat(1,C_,C); F4_=cat(1,F4_,F4); mF4_=cat(1,mF4_,mF4); C2_=cat(1,C2_,C2);
    clear F2 F4 TR mask1 svar sdist mF2 mF4 C C2
end


[~,svar] = sort(var(F2_,0,2)); [~,spsc] = sort(meanPSC(F2_));
sortorder2 = sortorder; sortorder=[]; f=0; %#ok<*ASGLU>
for im=1:length(sortorder2)
    temp = sortorder2{im}; temp = temp + f; f = f + length(temp);
    sortorder = cat(1,sortorder,temp); clear temp
end; clear im f
sortorder = spsc; % options: spsc for percentage signal change, svar for variance, (comment out the line if you want to use distance from the center of the cord separately for each slice)
for im=max(sortorder):-1:1, if isempty(find(sortorder==im)), sortorder(find(sortorder>im)) = sortorder(find(sortorder>im))-1; end, end; clear im %#ok<*EFIND>

fighndl = figure('Position',[1 1 1152 720],'Color',[0.89,0.94,0.99],'InvertHardcopy','off','Visible','off'); % red [0.95,0.9,0.85], green [0.85,0.90,0.85], blue [0.89,0.94,0.99]
imagesc(F2_(sortorder,:)),colormap(cmap),colorbar, xlabel('time (in samples)'),ylabel('voxels (sorted by mean percentage signal change; bottom row is highest)'),
  title(['Time series within GM, WM and CSF BEFORE NORDIC denoising (all slices combined)'],'Color',[0,0.55,0.69],'FontSize',14);
saveas(fighndl,[QCpath3 'fMRI_ts_01nordic_allSlices' '.jpg'])
saveas(fighndl,[QCpath4 'fMRI_ts_01nordic_allSlices' '.jpg'])
 close(fighndl)
mF2_=mean(mF2_,1); mF4_=mean(mF4_,1);
fighndl = figure('Position',[1 1 1152 720],'Color',[0.89,0.94,0.99],'InvertHardcopy','off','Visible','off'); % red [0.95,0.9,0.85], green [0.85,0.90,0.85], blue [0.89,0.94,0.99]
subplot2n(2,1,1), plot(mF2_,'color',[0.65,0.11,0.19],'linewidth',1.5), hold on, plot(mF4_,'color',[0,0.55,0.69],'linewidth',0.2), xlabel('time (in samples)'),ylabel('percentage signal change'),
  grid on,grid minor, xlim([1 length(mF2_)]),ylim([1.25.*min([mF2_,mF4_]) 1.25.*max([mF2_,mF4_])]), title(['Global mean signal BEFORE NORDIC denoising (all slices averaged)'],'Color',[0.65,0.11,0.19],'FontSize',14), legend({'Global signal before NORDIC denoising'},'FontSize',12,'TextColor',[0.65,0.11,0.19],'Location','best')
subplot2n(2,1,2), plot(mF4_,'color',[0,0.55,0.69],'linewidth',1.5), hold on, plot(mF2_,'color',[0.65,0.11,0.19],'linewidth',0.2), xlabel('time (in samples)'),ylabel('percentage signal change'),
  grid on,grid minor, xlim([1 length(mF4_)]),ylim([1.25.*min([mF2_,mF4_]) 1.25.*max([mF2_,mF4_])]), title(['Global mean signal AFTER NORDIC denoising (all slices averaged)'],'Color',[0,0.55,0.69],'FontSize',14), legend({'Global signal after NORDIC denoising'},'FontSize',12,'TextColor',[0,0.55,0.69],'Location','best')
saveas(fighndl,[QCpath3 'global_signal_01nordic_allSlices' '.jpg'])
saveas(fighndl,[QCpath4 'global_signal_01nordic_allSlices' '.jpg'])
 close(fighndl)
fighndl = figure('Position',[1 1 1152 720],'Color',[0.89,0.94,0.99],'InvertHardcopy','off','Visible','off'); % red [0.95,0.9,0.85], green [0.85,0.90,0.85], blue [0.89,0.94,0.99]
subplot2n(2,1,1), histogram(C2_,'EdgeColor','none','FaceColor',[0.65,0.11,0.19],'FaceAlpha',0.5,'Normalization','probability'); grid on,grid minor, xlim([-1 1]), hold on, xline(0,'Color','k','linewidth',1.5); hold off
    title(sprintf('Histogram of voxel-to-voxel correlations BEFORE NORDIC denoising (all slices combined)\n%.1f%% of the correlations were > 0  [ mean=%.2f, median=%.2f, SD=%.2f ]',100*length(find(C2_>0))/length(C2_),mean(C2_),median(C2_),std(C2_)),'Color',[0.65,0.11,0.19],'FontSize',14)
subplot2n(2,1,2), histogram(C_,'EdgeColor','none','FaceColor',[0,0.55,0.69],'FaceAlpha',0.5,'Normalization','probability'); grid on,grid minor, xlim([-1 1]), hold on, xline(0,'Color','k','linewidth',1.5); hold off
    title(sprintf('Histogram of voxel-to-voxel correlations AFTER NORDIC denoising (all slices combined)\n%.1f%% of the correlations were > 0  [ mean=%.2f, median=%.2f, SD=%.2f ]',100*length(find(C_>0))/length(C_),mean(C_),median(C_),std(C_)),'Color',[0,0.55,0.69],'FontSize',14)
saveas(fighndl,[QCpath3 'hist_FC_compare_01nordic_allSlices' '.jpg'])
saveas(fighndl,[QCpath4 'hist_FC_compare_01nordic_allSlices' '.jpg'])
 close(fighndl)
clear svar spsc sortorder F2_ mF2_ C_ F4_ mF4_ C2_ F6_ mF6_ C3_ cmap

%% NORDIC: make QC video (needs audio video toolbox in Matlab)
if ~isempty(which('VideoWriter')) && (save_videos==1) % check if Matlab's audio video toolbox exists

subplot = @(m,n,p) subtightplot(m, n, p, [0.001 0.005], [0.01 0.001], [0.01 0.001]);
for i3 = 1 : siz3
    try waitbar((i3/siz3),wbar3,sprintf('05. Generate QC video (NORDIC): slice (%d) of (%d)',i3,siz3)); catch, end
    fprintf('05. Generate QC video: slice (%d) of (%d)\n',i3,siz3)
    scfMRItb_04_resplitData(base_dir_sub, fname, '_beforeNORDIC', siz3) % if slice-wise data doesn't exist but combined 4D data exists, then split the data to generate the needed slice-wise data
    F1 = load_untouch_nii([base_dir_sub fname '_slice' num2str(i3) '_beforeNORDIC.nii']);
    TR = F1.hdr.dime.pixdim(5); F1 = rot90(single(squeeze(F1.img)));
    video = VideoWriter([QCpath3 'fMRIvideo_01nordic_slice' sprintf('%.2d',i3) '.mp4'],'MPEG-4'); %#ok<*TNMLP> % create the video object
    video.FrameRate = round(30/TR); % 30 seconds of data in every second of the video
    video.Quality = 90; % very limited compression so that the images are shown as is (and thus avoid misunderstandings due to video quality)
    open(video); % open the file for writing
    
    if Smask_err==0
        scfMRItb_04_resplitData(base_dir_sub, fname, '_Smask', siz3) % if slice-wise data doesn't exist but combined 4D data exists, then split the data to generate the needed slice-wise data
        scfMRItb_04_unzipFile(base_dir_sub, fname, ['_slice' num2str(i3) '_Smask'])
        mask1 = load_untouch_nii([base_dir_sub fname '_slice' num2str(i3) '_Smask.nii']); mask1 = mask1.img;
    else
        mask1 = Smask1_(:,:,i3);
    end
    mask1 = rot90(mask1);
    [xmask,ymask] = find(mask1==1);
    minx=round(min(xmask)-0.1*(size(mask1,1))); maxx=round(max(xmask)+0.1*(size(mask1,1)));
    miny=round(min(ymask)-0.1*(size(mask1,2))); maxy=round(max(ymask)+0.1*(size(mask1,2)));
    if (maxx-minx)<(maxy-miny)
        factr = ((maxy-miny)-(maxx-minx))/2;
        if factr==round(factr)
            maxx=maxx+factr; minx=minx-factr;
        else
            maxx=maxx+factr+0.5; minx=minx-factr+0.5;
        end
    elseif (maxy-miny)<(maxx-minx)
        factr = ((maxx-minx)-(maxy-miny))/2;
        if factr==round(factr)
            maxy=maxy+factr; miny=miny-factr;
        else
            maxy=maxy+factr+0.5; miny=miny-factr+0.5;
        end
    end
    [sizeFx,sizeFy] = size(F1(:,:,1));
    F2 = F1(minx:maxx,miny:maxy,:);
    for i5=1:size(F1,3)
        F1(:,:,i5) = imresize(F2(:,:,i5),[sizeFx sizeFy]);
    end
    scfMRItb_04_resplitData(base_dir_sub, fname, '', siz3) % if slice-wise data doesn't exist but combined 4D data exists, then split the data to generate the needed slice-wise data
    F3 = load_untouch_nii([base_dir_sub fname '_slice' num2str(i3) '.nii']); F3 = rot90(single(squeeze(F3.img)));
    F4 = F3(minx:maxx,miny:maxy,:);
    for i5=1:size(F3,3)
        F3(:,:,i5) = imresize(F4(:,:,i5),[sizeFx sizeFy]);
    end
    clear maxx maxy minx miny xmask ymask mask1 factr sizeFx sizeFy F2 F4 i5
    
    for ii=1:size(F1,3) % loop across the number of images
        I = F1(:,:,ii); %read the next image
        I(find(isnan(I)))=0;
        I = (I - min(I(:))) ./ (max(I(:)) - min(I(:)));
        I2 = F3(:,:,ii); %read the next image
        I2(find(isnan(I2)))=0;
        I2 = (I2 - min(I2(:))) ./ (max(I2(:)) - min(I2(:)));
        fighndl = figure('Position',[round((scrsz(3)-scrsz(4))/2) 1 round(0.75*scrsz(3)) round(0.75*scrsz(4))],'Color',[0.95,0.9,0.85],'InvertHardcopy','off','Visible','off'); % [0.85,0.90,0.85]
        subplot(1,2,1), imshow(I, [0 max(I(:))], 'InitialMagnification', 'fit'); title({['fMRI data BEFORE NORDIC denoising (slice ' num2str(i3) ' of ' num2str(siz3) ')'];['Time point ' num2str(ii) ' of ' num2str(size(F1,3))]},'Color',[0.65,0.11,0.19],'FontSize',18);
        subplot(1,2,2), imshow(I2, [0 max(I2(:))], 'InitialMagnification', 'fit'); title({['fMRI data AFTER NORDIC denoising (slice ' num2str(i3) ' of ' num2str(siz3) ')'];['Time point ' num2str(ii) ' of ' num2str(size(F1,3))]},'Color',[0,0.55,0.69],'FontSize',18);
        frame = getframe(gcf);
        writeVideo(video,frame); % write the image to file
        close(fighndl)
        clear I I2 frame
    end
    close(video); %close the file
    clear F1 F3 video ii TR
end; clear i3

end

end

end
```
