## Supplementary material for "Neptune: a toolbox for spinal cord functional MRI data processing and quality assurance": Neptune software: scfMRItb_07_baseimage_if_no_06denoise1.m

```
function scfMRItb_07_baseimage_if_no_06denoise1(base_dir_sub,fname, varargin)

fname3 = [fname '_mean'];

if nargin<3
    wbar3 = waitbar(0,'08. get base image...','Name','Progress(08): Get base image...','Units','normalized','Position',[3/5 0 1/5 1/17]);
else
    wbar3 = varargin{1};
end
    
try
    scfMRItb_04_unzipFile(base_dir_sub, fname3, '')
    F = load_untouch_nii([base_dir_sub fname3 '.nii']); % functional
catch
    afni_error = unix(['3dTstat -mean -prefix ' base_dir_sub fname3 '.nii ' base_dir_sub fname '_resliced' '.nii' ]); % try, in case the functionals were resliced along the z-axis
    if afni_error~=0 % if functionals were not resliced along the z-axis
        unix(['3dTstat -mean -prefix ' base_dir_sub fname3 '.nii ' base_dir_sub fname '.nii' ]);
    end; clear afni_error
    scfMRItb_04_unzipFile(base_dir_sub, fname3, '')
    F = load_untouch_nii([base_dir_sub fname3 '.nii']); % functional
end
siz3 = size(F.img,3); clear F

%% ----- AUTOMATICALLY SELECT A BASE IMAGE FOR EACH SLICE ------------------

scfMRItb_04_resplitData(base_dir_sub, fname, '_base', siz3) % if slice-wise data doesn't exist but combined 4D data exists, then split the data to generate the needed slice-wise data

if ~(exist([base_dir_sub fname '_slice' num2str(siz3) '_base.nii'], 'file'))

eval(['load ' base_dir_sub fname '_mask_NS' '.mat;']);
mask_NS = ~mask_NS.img;
base_list = zeros([1 siz3]);

for i3 = 1 : siz3
    try waitbar((i3/siz3),wbar3,sprintf('08. get base image: slice (%d) of (%d)',i3,siz3)); catch, end
    scfMRItb_04_resplitData(base_dir_sub, fname, '', siz3) % if slice-wise data doesn't exist but combined 4D data exists, then split the data to generate the needed slice-wise data
    scfMRItb_04_unzipFile(base_dir_sub, fname, ['_slice' num2str(i3)])
    F = load_untouch_nii([base_dir_sub fname '_slice' num2str(i3) '.nii']);
    F = squeeze(F.img);
    % only consider the spinal cord
    F = F .* repmat(mask_NS(:,:,i3), [1 1 size(F,3)]);
    
    % median intensity for each time series
    median_img = zeros([size(F,1) size(F,2)]);
    for i1 = 1 : size(F,1)
        for i2 = 1 : size(F,2)
            if any(F(i1,i2,:),3)
                median_img(i1,i2) = median(squeeze(F(i1,i2,:)));
            end
        end
    end
    
    median_img_diff = zeros(size(F,3),1);
    for i4 = 1 : size(F,3)
        median_img_diff(i4) = sum(sum(abs(median_img - F(:,:,i4))));
    end
    optimal = find(min(median_img_diff) == median_img_diff);
    optimal = optimal(1,1); % in case there is more than one (unlikely, but...)

    base_list(1,i3) = optimal - 1; % first image = 0
end
clear mask_NS F median_img median_img_diff optimal;

base_list;
for i3 = 1 : siz3
    try waitbar((i3/siz3),wbar3,sprintf('08. get base image (last part): slice (%d) of (%d)',i3,siz3)); catch, end
    unix(['rm -f ' base_dir_sub fname '_slice' num2str(i3) '+orig.*']);
    str = ['3dcalc -a ' base_dir_sub fname '_slice' num2str(i3) '.nii ' ...
        '-expr ''a'' -prefix ' base_dir_sub fname '_slice' num2str(i3) '+orig'];
    unix(str);
    
    % switch from 'tlrc' back to 'orig' if necessary
    if exist([base_dir_sub fname '_slice' num2str(i3) '+tlrc.BRIK'], 'file')
        unix(['3drefit -view ''orig'' ' base_dir_sub fname '_slice' num2str(i3) '+tlrc']);
    end
    
    unix(['rm -f ' base_dir_sub fname '_slice' num2str(i3) '_base.nii']);
    str = ['3dTcat -prefix ' fname '_slice' num2str(i3) '_base.nii ' ...
        '-session ' base_dir_sub ' ''' base_dir_sub fname '_slice' num2str(i3) '+orig[' num2str(base_list(1,i3)) ']'''];
    unix(str);
    unix(['rm -f ' base_dir_sub fname '_slice' num2str(i3) '+orig.*']);
end

end

try
if nargin<3
    close(wbar3)
end
catch
end

end
```
