## Supplementary material for "Neptune: a toolbox for spinal cord functional MRI data processing and quality assurance": Neptune software: scfMRItb_07_gaussianMask.m

```
function scfMRItb_07_gaussianMask(base_dir_sub,fname,automatic, varargin)

scrsz = get(0,'ScreenSize');
pause on;
fname3 = [fname '_mean'];

if nargin<4
    wbar3 = waitbar(0,'07. Define Gaussian mask...','Name','Progress(07): Gaussian mask...','Units','normalized','Position',[4/5 0 1/5 1/17]);
else
    wbar3 = varargin{1};
end
if nargin<5
    run = 0; % 0 = don't create runX subfolder; X = create runX subfolder
else
    run = varargin{2};
end

if ~(exist([base_dir_sub 'QC'],'dir'))
    mkdir([base_dir_sub 'QC'])
end
if ~(exist([base_dir_sub 'QC' '/07_gaussianMask'],'dir'))
    mkdir([base_dir_sub 'QC' '/07_gaussianMask'])
end
if run==0
    QCpath = [base_dir_sub 'QC' '/07_gaussianMask/'];
else
    if ~(exist([base_dir_sub 'QC' '/07_gaussianMask' '/run' num2str(run)],'dir'))
        mkdir([base_dir_sub 'QC' '/07_gaussianMask' '/run' num2str(run)])
    end
    QCpath = [base_dir_sub 'QC' '/07_gaussianMask' '/run' num2str(run) '/'];
end


% ----- CREATE GAUSSIAN MASKS FOR FUNCTIONAL ------------------------------

%%
% define Gaussian mask manually (user input required), either if automatic setting is 0 or if the not-spine mask is unavailable (because doing automatic requires the not-spine mask)
if (automatic==0) || ~(exist([base_dir_sub fname '_mask_NS.mat'], 'file'))
    
scfMRItb_04_unzipFile(base_dir_sub, fname3, '')
F = load_untouch_nii([base_dir_sub fname3 '.nii']); % mean functional
siz3 = size(F.img,3);
Gmask = F; % Gaussian mask
new_size = round(size(F.img,1) / 2);

for i3 = 1 : siz3 % generate x and y limits for viewing the images
    try
        scfMRItb_04_resplitData(base_dir_sub, fname, '_Smask', siz3) % if slice-wise data doesn't exist but combined 4D data exists, then split the data to generate the needed slice-wise data
        scfMRItb_04_unzipFile(base_dir_sub, fname, ['_slice' num2str(i3) '_Smask'])
        Smask = load_untouch_nii([base_dir_sub fname '_slice' num2str(i3) '_Smask.nii']);
    catch
        Smask.img = ones(size(F,1),size(F,2));
    end
    Smask = Smask.img;
    [w2,w1] = find(Smask~=0);
    w1min(i3,1) = min(w1)/size(Smask,1); w1max(i3,1) = max(w1)/size(Smask,1);
    w2min(i3,1) = min(w2)/size(Smask,2); w2max(i3,1) = max(w2)/size(Smask,2);
    clear Smask w1 w2
end
clear F;

for i3 = 1 : siz3
    try waitbar((i3/siz3),wbar3,sprintf('07. Define Gaussian mask: slice (%d) of (%d)',i3,siz3)); catch, end
    scfMRItb_04_resplitData(base_dir_sub, fname, '_base', siz3) % if slice-wise data doesn't exist but combined 4D data exists, then split the data to generate the needed slice-wise data
    scfMRItb_04_unzipFile(base_dir_sub, fname, ['_slice' num2str(i3) '_base'])
    F = load_untouch_nii([base_dir_sub fname '_slice' num2str(i3) '_base.nii']);
    clf;
    img = F.img;
    temp = img(round((size(img,1)-new_size)/2)+1:end-round((size(img,1)-new_size)/2), ...
        round((size(img,2)-new_size)/2)+1:end-round((size(img,2)-new_size)/2));
    mmax = max(temp(:)); clear temp;
    fighndl = figure('Position',[round((scrsz(3)-scrsz(4))/2) 1 scrsz(4) scrsz(4)],'Color',[0.95,0.9,0.85],'InvertHardcopy','off'); try addToolbarExplorationButtons(fighndl); catch, end
    imshow(single(img), [0 mmax], 'InitialMagnification', 'fit'); colorbar;
    xlim([round(size(img,1)*0.9*w1min(i3)),round(size(img,1)*1.1*w1max(i3))]),ylim([round(size(img,2)*0.9*w2min(i3)),round(size(img,2)*1.1*w2max(i3))]),
    
    title(['Slice ' num2str(i3) ': Select center of spinal cord.']);
    disp(['Slice ' num2str(i3) ': Select center of spinal cord.']);
    [x_center, y_center] = ginput(1);
    title(['Slice ' num2str(i3) ': Select right extent of CSF.']);
    disp(['Slice ' num2str(i3) ': Select right extent of CSF.']);
    [x_right, y_right] = ginput(1);
    title(['Slice ' num2str(i3) ': Select top extent of CSF.']);
    disp(['Slice ' num2str(i3) ': Select top extent of CSF.']);
    [x_top, y_top] = ginput(1);
    
    x_center = round(x_center);
    y_center = round(y_center);
    x_right  = round(x_right);
    y_right  = round(y_right);
    x_top    = round(x_top);
    y_top    = round(y_top);
    
    x_fwhm = abs(x_right-x_center);
    y_fwhm = abs(y_center-y_top);
    x1 = repmat(((1:1:size(F.img,2))'-y_center),[1 size(F.img,2)]);
    y1 = repmat(((1:1:size(F.img,2))-x_center),[size(F.img,1) 1]);
    W = single(gauss2D_R(x1, y1, 2*y_fwhm, 2*x_fwhm, 1, 1));
    
    % suppress voxels less than 0.1
    min_weight = 0.1;
    W = W .* (W > min_weight);
    
    Gmask.img(:,:,i3) = W;
    clf;
    figmriF(F.img .* W); % briefly display masked image
        title(sprintf('Gaussian masked mean functional image - slice %d of %d',i3,siz3),'FontSize',14)
    saveas(fighndl,[QCpath 'MeanFunc_GaussianMasked_slice' sprintf('%.2d',i3) '.jpg'])
    pause(1);
    close(fighndl)
    clear x_center y_center x_right y_right x_top y_top x_fwhm y_fwhm x1 y1 W
end
clear new_size img temp mmax fighndl;

%%
elseif automatic==1 % determine Gaussian mask automatically using the not-spine mask

scfMRItb_04_unzipFile(base_dir_sub, fname3, '')
F = load_untouch_nii([base_dir_sub fname3 '.nii']); % mean functional
siz3 = size(F.img,3);
Gmask = F; % Gaussian mask
new_size = round(size(F.img,1) / 2);
clear F

load([base_dir_sub fname '_mask_NS.mat'])
mask_NS = mask_NS.img;

for i3 = 1 : siz3
    try waitbar((i3/siz3),wbar3,sprintf('07. Compute Gaussian mask: slice (%d) of (%d)',i3,siz3)); catch, end
    scfMRItb_04_resplitData(base_dir_sub, fname, '_base', siz3) % if slice-wise data doesn't exist but combined 4D data exists, then split the data to generate the needed slice-wise data
    scfMRItb_04_unzipFile(base_dir_sub, fname, ['_slice' num2str(i3) '_base'])
    F = load_untouch_nii([base_dir_sub fname '_slice' num2str(i3) '_base.nii']);
    img = mask_NS(:,:,i3);
    [i1,i2] = find(img==0);
    x_center = round(median(i2)); % in imagesc display, the vertical axis is the x-axis, hence i2 is used instead of i1
    y_center = round(median(i1)); % in imagesc display, the horizontal axis is the y-axis, hence i1 is used instead of i2
    x_right  = max(i2);
    x_left   = min(i2);
    y_top    = min(i1);
    y_bottom = max(i1);
    
    x_fwhm = round(abs(x_right-x_left)/2);
    y_fwhm = round(abs(y_top-y_bottom)/2);
    x1 = repmat(((1:1:size(F.img,1))'-y_center),[1 size(F.img,2)]);
    y1 = repmat(((1:1:size(F.img,2))-x_center),[size(F.img,1) 1]);
    W = single(gauss2D_R(x1, y1, 2*y_fwhm, 2*x_fwhm, 1, 1));
    
    % suppress voxels less than 0.1
    min_weight = 0.1;
    W = W .* (W > min_weight);
    
    Gmask.img(:,:,i3) = W;
    fighndl = figure('Position',[round((scrsz(3)-scrsz(4))/2) 1 scrsz(4) scrsz(4)],'InvertHardcopy','off','Visible','off'); try addToolbarExplorationButtons(fighndl); catch, end
    figmriF(single(F.img) .* W); % briefly display masked image
        title(sprintf('Gaussian masked mean functional image - slice %d of %d',i3,siz3),'FontSize',14)
    saveas(fighndl,[QCpath 'MeanFunc_GaussianMasked_slice' sprintf('%.2d',i3) '.jpg'])
    close(fighndl)
    % pause(0.75);
    clear x_center y_center x_right y_right x_top y_top x_left y_bottom x_fwhm y_fwhm x1 y1 W min_weight F img
end
clear new_size img temp mmax fighndl;

end

%%
F = load_untouch_nii([base_dir_sub fname3 '.nii']); % mean functional
unix(['rm -f ' base_dir_sub fname '_Gaussian_mask.nii']);
save_untouch_nii(Gmask, [base_dir_sub fname '_Gaussian_mask.nii']);
clear W Gmask x1 y1;

for i3 = 1 : size(F.img,3)
    try waitbar((i3/size(F.img,3)),wbar3,sprintf('07. Gaussian mask (final housekeeping): slice (%d) of (%d)',i3,size(F.img,3))); catch, end
    unix(['rm -f ' base_dir_sub fname '_slice' num2str(i3) '_Gaussian_mask.nii']);
    unix(['3dZcutup -keep ' num2str(i3-1) ' ' num2str(i3-1) ' ' ...
        '-prefix ' base_dir_sub fname '_slice' num2str(i3) '_Gaussian_mask.nii' ' ' ...
        base_dir_sub fname '_Gaussian_mask.nii']);
end

try
if nargin<4
    close(wbar3)
end
catch
end

end
```
