## Supplementary material for "Neptune: a toolbox for spinal cord functional MRI data processing and quality assurance": Neptune software: scfMRItb_08_motionCorrection.m

```
function scfMRItb_08_motionCorrection(base_dir_sub,fname, varargin)

if nargin<3
    motion_param_smoothing = 0; % do not smooth estimated motion parameters
else
    motion_param_smoothing = varargin{1};
end
if nargin<4
    wbar3 = waitbar(0,'08. Motion correction...','Name','Progress(08): Motion correction...','Units','normalized','Position',[3/5 0 1/5 1/17]);
else
    wbar3 = varargin{2};
end
if nargin<5
    save_videos = 1; % 1 = save QC videos (generates sizeable .mp4 files and takes a few minutes to do it). This is turned on by default, but if you don't want to save videos then set it to 0
else
    save_videos = varargin{3};
end
if nargin<6
    run = 0; % 0 = don't create runX subfolder; X = create runX subfolder
else
    run = varargin{4};
end

if ~(exist([base_dir_sub 'QC'],'dir'))
    mkdir([base_dir_sub 'QC'])
end
if ~(exist([base_dir_sub 'QC' '/08_motionCorrection'],'dir'))
    mkdir([base_dir_sub 'QC' '/08_motionCorrection'])
end
if run==0
    QCpath = [base_dir_sub 'QC' '/08_motionCorrection/'];
else
    if ~(exist([base_dir_sub 'QC' '/08_motionCorrection' '/run' num2str(run)],'dir'))
        mkdir([base_dir_sub 'QC' '/08_motionCorrection' '/run' num2str(run)])
    end
    QCpath = [base_dir_sub 'QC' '/08_motionCorrection' '/run' num2str(run) '/'];
end

if ~(exist([base_dir_sub 'QC' '/QC_shortlist'],'dir'))
    mkdir([base_dir_sub 'QC' '/QC_shortlist'])
end
if run==0
    QCpath2 = [base_dir_sub 'QC' '/QC_shortlist/'];
else
    if ~(exist([base_dir_sub 'QC' '/QC_shortlist' '/run' num2str(run)],'dir'))
        mkdir([base_dir_sub 'QC' '/QC_shortlist' '/run' num2str(run)])
    end
    QCpath2 = [base_dir_sub 'QC' '/QC_shortlist' '/run' num2str(run) '/'];
end

fname3 = [fname '_mean'];
scfMRItb_04_unzipFile(base_dir_sub, fname3, '')
A = load_untouch_nii([base_dir_sub fname3 '.nii']); % mean functional
siz3 = size(A.img,3);
cutoff1 = 0.80; cutoff1_str = '80';

%% ----- SLICE-BY-SLICE MOTION CORRECTION ----------------------------------

parfix = '-parfix 3 0 -parfix 4 0 -parfix 5 0 -parfix 6 0 -parfix 7 1 -parfix 8 1 -parfix 9 1 -parfix 10 0 -parfix 11 0 -parfix 12 0';
for i3 = 1 : siz3
    try waitbar((i3/siz3),wbar3,sprintf('08. Motion correction (initial housekeeping): slice (%d) of (%d)',i3,siz3)); catch, end
    unix(['rm -f ' base_dir_sub fname '_slice' num2str(i3) '_MCnofilt.nii']);    
    scfMRItb_04_resplitData(base_dir_sub, fname, '_Gaussian_mask', siz3) % if slice-wise data doesn't exist but combined 4D data exists, then split the data to generate the needed slice-wise data
    scfMRItb_04_resplitData(base_dir_sub, fname, '_base', siz3) % if slice-wise data doesn't exist but combined 4D data exists, then split the data to generate the needed slice-wise data
    scfMRItb_04_resplitData(base_dir_sub, fname, ['_denoised_before_MC' cutoff1_str], siz3) % if slice-wise data doesn't exist but combined 4D data exists, then split the data to generate the needed slice-wise data

    str = ['3dWarpDrive -affine_general ' parfix ' -quintic -final quintic ' ...
        '-base ' base_dir_sub fname '_slice' num2str(i3) '_base.nii ' ...
        '-prefix ' base_dir_sub fname '_slice' num2str(i3) '_MCnofilt.nii ' ...
        '-weight ' base_dir_sub fname '_slice' num2str(i3) '_Gaussian_mask.nii ' ...
        '-1Dfile ' base_dir_sub fname '_slice' num2str(i3) '_MC_params.txt ' ...
        '-1Dmatrix_save ' base_dir_sub fname '_slice' num2str(i3) '_MC_xform.aff12.1D ' ...
        base_dir_sub fname '_slice' num2str(i3) '_denoised_before_MC' cutoff1_str '.nii'];
    runerr = unix(str); clear str
    
    if runerr~=0 % if denoise1 was skipped or not performed, the above unix command will return an error upon which the following command will be executed using pre-denoise1 data
    scfMRItb_04_resplitData(base_dir_sub, fname, '', siz3) % if slice-wise data doesn't exist but combined 4D data exists, then split the data to generate the needed slice-wise data
    str = ['3dWarpDrive -affine_general ' parfix ' -quintic -final quintic ' ...
        '-base ' base_dir_sub fname '_slice' num2str(i3) '_base.nii ' ...
        '-prefix ' base_dir_sub fname '_slice' num2str(i3) '_MCnofilt.nii ' ...
        '-weight ' base_dir_sub fname '_slice' num2str(i3) '_Gaussian_mask.nii ' ...
        '-1Dfile ' base_dir_sub fname '_slice' num2str(i3) '_MC_params.txt ' ...
        '-1Dmatrix_save ' base_dir_sub fname '_slice' num2str(i3) '_MC_xform.aff12.1D ' ...
        base_dir_sub fname '_slice' num2str(i3) '.nii'];
    unix(str);
    end
    clear str runerr
end

for i3 = 1 : siz3
    try waitbar((i3/siz3),wbar3,sprintf('08. Motion correction: slice (%d) of (%d)',i3,siz3)); catch, end
    fprintf('Motion correction, slice (%d) of (%d)\n',i3,siz3)
    xform_file = [base_dir_sub fname '_slice' num2str(i3) '_MC_xform.aff12.1D'];
    
    % read in motion parameters
    xform_raw = read_1D_xform_file(xform_file);
    xform_orig = zeros(12,size(xform_raw{1},1));
    for i1 = 1 : 12
        xform_orig(i1,:) = xform_raw{i1};
    end
    clear xform_raw;
    
    % filter motion parameters (to suppress sporadic artifacts)
    for i1 = 1 : 2
        if i1 == 1
            xform = xform_orig(4,:); % L-R
        else
            xform = xform_orig(8,:); % A-P
        end
        xform_filt = xform;
        
        % N-point median filter (if chosen)
        if motion_param_smoothing~=0
            N_order = round((motion_param_smoothing-1)/2);
            for i2 = 1+N_order : size(xform,2)-N_order
                xform_filt(1,i2) = median(xform(1,i2-N_order:i2+N_order));
            end
            for i2 = 1 : N_order
                xform_filt(1,i2) = median(xform(1,1:i2+N_order));
            end
            for i2 = size(xform,2)-N_order+1 : size(xform,2)
                xform_filt(1,i2) = median(xform(1,i2-N_order:size(xform,2)));
            end
            clear N_order i2
        end
        
        if i1 == 1
            xform_orig(4,:) = xform_filt;
        else
            xform_orig(8,:) = xform_filt;
        end
        clear xform xform_filt;
    end
    
    % write out parameters to new file
    if motion_param_smoothing~=0
        fileID = fopen([base_dir_sub fname '_slice' num2str(i3) '_MCfilt_xform.aff12.1D'], 'w');
        fprintf(fileID,'%13.8f %13.8f %13.8f %13.8f %13.8f %13.8f %13.8f %13.8f %13.8f %13.8f %13.8f %13.8f\n', xform_orig);
        fclose(fileID);
    end

    % apply motion parameters
    unix(['rm -f ' base_dir_sub fname '_slice' num2str(i3) '_MC.nii']);

    if motion_param_smoothing~=0
	runerr = unix(['3dAllineate -verb -final wsinc5 ' ...
                '-input ' base_dir_sub fname '_slice' num2str(i3) '_denoised_before_MC' cutoff1_str '.nii ' ...
                '-prefix ' base_dir_sub fname '_slice' num2str(i3) '_MC.nii ' ...
                '-1Dmatrix_apply ' base_dir_sub fname '_slice' num2str(i3) '_MCfilt_xform.aff12.1D']);
    
    if runerr~=0 % if denoise1 was skipped or not performed, the above unix command will return an error upon which the following command will be executed using pre-denoise1 data
        unix(['3dAllineate -verb -final wsinc5 ' ...
            '-input ' base_dir_sub fname '_slice' num2str(i3) '.nii ' ...
            '-prefix ' base_dir_sub fname '_slice' num2str(i3) '_MC.nii ' ...
            '-1Dmatrix_apply ' base_dir_sub fname '_slice' num2str(i3) '_MCfilt_xform.aff12.1D']);
    end
    clear runerr

    else
    movefile([base_dir_sub fname '_slice' num2str(i3) '_MCnofilt.nii'],[base_dir_sub fname '_slice' num2str(i3) '_MC.nii'],'f'); % if motion parameter smoothing was not performed then there is no need to do realignment again
    end
end


%% Calculate framewise displacement and save motion correction QC photos

M_pool=[];
for i3=1:siz3
    M{i3,1} = importdata([base_dir_sub fname '_slice' num2str(i3) '_MC_params.txt']);
    M{i3} = M{i3}.data(:,1:2); M{i3}(:,3:6)=0; % M is a matrix of size Nx6 (N = no. of time points; 6 = 3 translation and 3 rotation parameters). We have only x and y translations taken into account. The rest of the degrees of freedom are zero padded.
    [FD(:,i3), mean_FD_perslice(i3,1), AD(:,i3)] = FD_metric2(M{i3},50);
    max_FD_perslice(i3,1) = max(FD(:,i3));
    M{i3}(:,3:end)=[]; 
    M_pool = cat(1,M_pool,M{i3}(:));
end; clear i3
mean_FD_total = mean(mean_FD_perslice);
max_FD_total = max(max_FD_perslice);
motion_parameters = M;

readme_framewise_displacement = sprintf('FD: Framewise Displacement; mean_FD: mean FD (one value per run); max_FD: max FD (one value per run); AD: Absolute Displacement\nmotion_parameters: a matrix of size Nx6 (N = no. of time points; 6 = 3 translation and 3 rotation parameters). We have only x and y translations taken into account. The rest of the degrees of freedom are zero padded.\nAll values in millimeters\nFor more info, type: help FD_metric2\n');
save([base_dir_sub 'motion_parameters.mat'],'motion_parameters','FD','AD','mean_FD_perslice','mean_FD_total','max_FD_perslice','max_FD_total','readme_framewise_displacement')

scrsz = get(0,'ScreenSize'); pause on;
fighndl = figure('Position',[round((scrsz(3)-scrsz(4))/2) 1 scrsz(4) scrsz(4)],'Color',[0.85,0.90,0.85],'InvertHardcopy','off','Visible','off'); % [0.85,0.90,0.85] [0.8, 0.89, 0.94]
for i3=1:siz3
    subplot2n(siz3,2,2*i3-1), plot(M{i3}(:,1),'b','linewidth',1.5), xlim([1 length(M{i3}(:,1))]),ylim([min(M_pool) max(M_pool)]), grid on, ylabel('x'),title(['(slice ' num2str(i3) ')'])
    subplot2n(siz3,2,2*i3),   plot(M{i3}(:,2),'r','linewidth',1.5), xlim([1 length(M{i3}(:,2))]),ylim([min(M_pool) max(M_pool)]), grid on, ylabel('y'),title(['(slice ' num2str(i3) ')'])
end; clear i3
try suptitle2(['X and Y translational motion parameters (in mm) for slices ' num2str(1) ' through ' num2str(siz3)]); fighndl.Visible='off'; catch, end
saveas(fighndl,[QCpath 'motion_params' '.jpg'])
saveas(fighndl,[QCpath2 'motion_params' '.jpg'])
close(fighndl)

fighndl = figure('Position',[round((scrsz(3)-scrsz(4))/2) 1 scrsz(4) scrsz(4)],'Color',[0.85,0.90,0.85],'InvertHardcopy','off','Visible','off'); % [0.85,0.90,0.85] [0.8, 0.89, 0.94]
for i3=1:siz3
    subplot2n(siz3,1,i3), plot(FD(:,i3),'Color',[0,0.5+0.25*i3/siz3,1-0.25*i3/siz3],'linewidth',1.5), xlim([1 length(FD(:,i3))]),ylim([min(FD(:)) max(FD(:))]), grid on, ylabel('FD'),title(['(slice ' num2str(i3) ')'])
end; clear i3
try suptitle2(['Framewise displacement (i.e. relative motion) (in mm) for slices ' num2str(1) ' through ' num2str(siz3)]); fighndl.Visible='off'; catch, end
saveas(fighndl,[QCpath 'relative_displacement' '.jpg'])
saveas(fighndl,[QCpath2 'relative_displacement' '.jpg'])
close(fighndl)

fighndl = figure('Position',[round((scrsz(3)-scrsz(4))/2) 1 scrsz(4) scrsz(4)],'Color',[0.85,0.90,0.85],'InvertHardcopy','off','Visible','off'); % [0.85,0.90,0.85] [0.8, 0.89, 0.94]
for i3=1:siz3
    subplot2n(siz3,1,i3), plot(AD(:,i3),'Color',[0.5+0.25*i3/siz3,0,1-0.25*i3/siz3],'linewidth',1.5), xlim([1 length(AD(:,i3))]),ylim([min(AD(:)) max(AD(:))]), grid on, ylabel('AD'),title(['(slice ' num2str(i3) ')'])
end; clear i3
try suptitle2(['Absolute displacement (in mm) for slices ' num2str(1) ' through ' num2str(siz3)]); fighndl.Visible='off'; catch, end
saveas(fighndl,[QCpath 'absolute_displacement' '.jpg'])
saveas(fighndl,[QCpath2 'absolute_displacement' '.jpg'])
close(fighndl)

fighndl = figure('Position',[round((scrsz(3)-scrsz(4))/2) 1 scrsz(4) scrsz(4)],'Color',[0.85,0.90,0.85],'InvertHardcopy','off','Visible','off'); % [0.85,0.90,0.85] [0.8, 0.89, 0.94]
subplot2n(2,1,1), b = bar(mean_FD_perslice,0.4,'FaceColor',[0,0.55,0.69],'EdgeColor',[0.65,0.11,0.19],'LineWidth',1.5);
  xtips1 = b(1).XEndPoints; ytips1 = b(1).YEndPoints; labels1 = string(round(b(1).YData,2)); text(xtips1,ytips1,labels1,'HorizontalAlignment','center','VerticalAlignment','bottom'); clear b xtips ytips labels1
  grid on, grid minor, ylim([0 max(mean_FD_perslice)+0.01]), ylabel('Mean framewise displacement'), xlabel('Slices'), title(['Mean framewise displacement (in mm) for slices ' num2str(1) ' through ' num2str(siz3) '  [ mean across all slices = ' sprintf('%.2f',mean_FD_total) ' mm ]'],'Color',[0,0.55,0.69],'FontSize',18);
subplot2n(2,1,2), b = bar(max_FD_perslice,0.4,'FaceColor',[0,0.55,0.69],'EdgeColor',[0.65,0.11,0.19],'LineWidth',1.5);
  xtips1 = b(1).XEndPoints; ytips1 = b(1).YEndPoints; labels1 = string(round(b(1).YData,2)); text(xtips1,ytips1,labels1,'HorizontalAlignment','center','VerticalAlignment','bottom'); clear b xtips ytips labels1
  grid on, grid minor, ylim([0 max(max_FD_perslice)+0.05]), ylabel('Max framewise displacement'), xlabel('Slices'), title(['Max framewise displacement (in mm) for slices ' num2str(1) ' through ' num2str(siz3) '  [ max across all slices = ' sprintf('%.2f',max_FD_total) ' mm ]'],'Color',[0.65,0.11,0.19],'FontSize',18);
saveas(fighndl,[QCpath 'mean_max_FD' '.jpg'])
saveas(fighndl,[QCpath2 'mean_max_FD' '.jpg'])
close(fighndl)


% Move motion parameters (and transformations) to a subdirectory within the subject's main directory
sdirm = cat(1,dir([base_dir_sub fname '*' 'MC_params.txt']),dir([base_dir_sub fname '*' '_MC_xform.aff12.1D']),dir([base_dir_sub fname '*' '_MCfilt_xform.aff12.1D']));
mkdir([base_dir_sub 'motion_params'])
for i5=1:length(sdirm)
    movefile([sdirm(i5).folder '/' sdirm(i5).name],[base_dir_sub 'motion_params' '/' sdirm(i5).name])
end; clear sdirm i5

%% time series plots, global signal and FC histogram
scrsz = get(0,'ScreenSize'); pause on;
cmap = colormap('bone'); cmap = (1-exp(-cmap))./(1-exp(-1)); 

Smask_err=0;
try
    scfMRItb_04_resplitData(base_dir_sub, fname, '_Smask', siz3) % if slice-wise data doesn't exist but combined 4D data exists, then split the data to generate the needed slice-wise data
catch
    load([base_dir_sub fname '_mask_NS' '.mat'])
    Smask1_ = ~mask_NS.img; clear mask_NS
    Smask_err=1;
end

F2_=[]; mF2_=[]; C_=[]; F4_=[]; mF4_=[]; C2_=[];
if exist([base_dir_sub fname '_slice' num2str(1) '_denoised_before_MC' cutoff1_str '.nii'],'file') || exist([base_dir_sub fname '_slice' num2str(1) '_denoised_before_MC' cutoff1_str '.nii.gz'],'file') || exist([base_dir_sub fname '_denoised_before_MC' cutoff1_str '.nii'],'file') || exist([base_dir_sub fname '_denoised_before_MC' cutoff1_str '.nii.gz'],'file')
    F6_=[]; mF6_=[]; C3_=[];
end
for i3 = 1 : siz3
    try waitbar((i3/siz3),wbar3,sprintf('08. QC plots: slice (%d) of (%d)',i3,siz3)); catch, end
    fprintf('08. QC plots: slice (%d) of (%d)\n',i3,siz3)
    scfMRItb_04_resplitData(base_dir_sub, fname, '_MC', siz3) % if slice-wise data doesn't exist but combined 4D data exists, then split the data to generate the needed slice-wise data
    scfMRItb_04_unzipFile(base_dir_sub, fname, ['_slice' num2str(i3) '_MC'])
    F1 = load_untouch_nii([base_dir_sub fname '_slice' num2str(i3) '_MC.nii']);
    TR = F1.hdr.dime.pixdim(5); F1 = single(squeeze(F1.img));
    
    if Smask_err==0
        scfMRItb_04_resplitData(base_dir_sub, fname, '_Smask', siz3) % if slice-wise data doesn't exist but combined 4D data exists, then split the data to generate the needed slice-wise data
        scfMRItb_04_unzipFile(base_dir_sub, fname, ['_slice' num2str(i3) '_Smask'])
        mask1 = load_untouch_nii([base_dir_sub fname '_slice' num2str(i3) '_Smask.nii']); mask1 = mask1.img;
    else
        mask1 = Smask1_(:,:,i3);
    end
    [m1,m2] = find(mask1==1); Mm1=median(m1); Mm2=median(m2);
    sdist = sqrt((m1-Mm1).^2 + (m2-Mm2).^2);
    clear im Mm1 Mm2 m1 m2
    for i5=1:size(F1,3)
        tmp = F1(:,:,i5);
        F2(:,i5) = tmp(find(mask1==1)); clear tmp
    end; clear i5 F1
    [~,svar] = sort(var(F2,0,2)); [~,sdist] = sort(sdist); sortorder{i3,1} = sdist; %#ok<*AGROW> % sort timeseries based on distance from the center of the cord (if you want to sort by signal variance then use svar)
    
    fighndl = figure('Position',[1 1 1152 720],'Color',[0.95,0.9,0.85],'InvertHardcopy','off','Visible','off'); % red [0.95,0.9,0.85], green [0.85,0.90,0.85], blue [0.89,0.94,0.99]
    imagesc(F2(sortorder{i3},:)),colormap(cmap),colorbar, xlabel('time (in samples)'),ylabel('voxels (sorted by distance from the center of the cord; top row is nearest)'),
        title(['Time series within GM, WM and CSF after "08-motion-correction" in slice ' num2str(i3) ' of ' num2str(siz3)],'Color',[0,0.55,0.69],'FontSize',14);
    saveas(fighndl,[QCpath 'fMRI_ts_08moco_slice' sprintf('%.2d',i3) '.jpg'])
    close(fighndl)
    
    scfMRItb_04_resplitData(base_dir_sub, fname, '', siz3) % if slice-wise data doesn't exist but combined 4D data exists, then split the data to generate the needed slice-wise data
    scfMRItb_04_unzipFile(base_dir_sub, fname, ['_slice' num2str(i3)])
    F3 = load_untouch_nii([base_dir_sub fname '_slice' num2str(i3) '.nii']); F3 = single(squeeze(F3.img));
    for i5=1:size(F3,3)
        tmp = F3(:,:,i5);
        F4(:,i5) = tmp(find(mask1==1)); clear tmp
    end; clear i5 F3
    mF2=mean(F2,1); mF2 = 100*(mF2-mean(mF2))./mean(mF2); mF4=mean(F4,1); mF4 = 100*(mF4-mean(mF4))./mean(mF4);
    fighndl = figure('Position',[1 1 1152 720],'Color',[0.95,0.9,0.85],'InvertHardcopy','off','Visible','off'); % red [0.95,0.9,0.85], green [0.85,0.90,0.85], blue [0.89,0.94,0.99]
    subplot2n(2,1,1), plot(mF4,'color',[0.65,0.11,0.19],'linewidth',1.5), hold on, plot(mF2,'color',[0,0.55,0.69],'linewidth',0.2), xlabel('time (in samples)'),ylabel('percentage signal change'),
      grid on,grid minor, xlim([1 length(mF4)]),ylim([1.25.*min([mF2,mF4]) 1.25.*max([mF2,mF4])]), title(['Global mean signal in UNPROCESSED RAW DATA (slice ' num2str(i3) ' of ' num2str(siz3) ')'],'Color',[0.65,0.11,0.19],'FontSize',14), legend({'Global signal (raw data)'},'FontSize',12,'TextColor',[0.65,0.11,0.19],'Location','best')
    subplot2n(2,1,2), plot(mF2,'color',[0,0.55,0.69],'linewidth',1.5), hold on, plot(mF4,'color',[0.65,0.11,0.19],'linewidth',0.2), xlabel('time (in samples)'),ylabel('percentage signal change'),
      grid on,grid minor, xlim([1 length(mF2)]),ylim([1.25.*min([mF2,mF4]) 1.25.*max([mF2,mF4])]), title(['Global mean signal AFTER 08-MOTION-CORRECTION (slice ' num2str(i3) ' of ' num2str(siz3) ')'],'Color',[0,0.55,0.69],'FontSize',14), legend({'Global signal after 08-motion-correction'},'FontSize',12,'TextColor',[0,0.55,0.69],'Location','best')
    saveas(fighndl,[QCpath 'global_signal_08moco_slice' sprintf('%.2d',i3) '.jpg'])
    close(fighndl)

    C  = corrcoef(F2'); C  = uppertriangle(C) ;
    C2 = corrcoef(F4'); C2 = uppertriangle(C2);
    fighndl = figure('Position',[1 1 1152 720],'Color',[0.95,0.9,0.85],'InvertHardcopy','off','Visible','off'); % red [0.95,0.9,0.85], green [0.85,0.90,0.85], blue [0.89,0.94,0.99]
    subplot2n(2,1,1), histogram(C2,'EdgeColor','none','FaceColor',[0.65,0.11,0.19],'FaceAlpha',0.5,'Normalization','probability'); grid on,grid minor, xlim([-1 1]), hold on, xline(0,'Color','k','linewidth',1.5); hold off  % xlim([-max(abs([C;C2])) max(abs([C;C2]))]),
      title(sprintf('Histogram of voxel-to-voxel correlations in UNPROCESSED RAW DATA (slice %d of %d)\n%.1f%% of the correlations were > 0  [ mean=%.2f, median=%.2f, SD=%.2f ]',i3,siz3,100*length(find(C2>0))/length(C2),mean(C2),median(C2),std(C2)),'Color',[0.65,0.11,0.19],'FontSize',14)
    subplot2n(2,1,2), histogram(C,'EdgeColor','none','FaceColor',[0,0.55,0.69],'FaceAlpha',0.5,'Normalization','probability'); grid on,grid minor, xlim([-1 1]), hold on, xline(0,'Color','k','linewidth',1.5); hold off
      title(sprintf('Histogram of voxel-to-voxel correlations AFTER 08-MOTION-CORRECTION (slice %d of %d)\n%.1f%% of the correlations were > 0  [ mean=%.2f, median=%.2f, SD=%.2f ]',i3,siz3,100*length(find(C>0))/length(C),mean(C),median(C),std(C)),'Color',[0,0.55,0.69],'FontSize',14)
    saveas(fighndl,[QCpath 'hist_FC_compare_08moco_slice' sprintf('%.2d',i3) '.jpg'])
    close(fighndl)
    
    if exist([base_dir_sub fname '_slice' num2str(1) '_denoised_before_MC' cutoff1_str '.nii'],'file') || exist([base_dir_sub fname '_slice' num2str(1) '_denoised_before_MC' cutoff1_str '.nii.gz'],'file') || exist([base_dir_sub fname '_denoised_before_MC' cutoff1_str '.nii'],'file') || exist([base_dir_sub fname '_denoised_before_MC' cutoff1_str '.nii.gz'],'file')
        scfMRItb_04_resplitData(base_dir_sub, fname, ['_denoised_before_MC' cutoff1_str], siz3) % if slice-wise data doesn't exist but combined 4D data exists, then split the data to generate the needed slice-wise data
        scfMRItb_04_unzipFile(base_dir_sub, fname, ['_slice' num2str(i3) '_denoised_before_MC' cutoff1_str])
        F5 = load_untouch_nii([base_dir_sub fname '_slice' num2str(i3) '_denoised_before_MC' cutoff1_str '.nii']); F5 = single(squeeze(F5.img));
        for i5=1:size(F5,3)
            tmp = F5(:,:,i5);
            F6(:,i5) = tmp(find(mask1==1)); clear tmp
        end; clear i5 F5
        mF6=mean(F6,1); mF6 = 100*(mF6-mean(mF6))./mean(mF6);
        fighndl = figure('Position',[1 1 1152 720],'Color',[0.95,0.9,0.85],'InvertHardcopy','off','Visible','off'); % red [0.95,0.9,0.85], green [0.85,0.90,0.85], blue [0.89,0.94,0.99]
        subplot2n(2,1,1), plot(mF6,'color',[0.65,0.11,0.19],'linewidth',1.5), hold on, plot(mF2,'color',[0,0.55,0.69],'linewidth',0.2), xlabel('time (in samples)'),ylabel('percentage signal change'),
          grid on,grid minor, xlim([1 length(mF6)]),ylim([min(1.25.*[mF2,mF6]) 1.25.*max([mF2,mF6])]), title(['Global mean signal after "06-denoise1" (slice ' num2str(i3) ' of ' num2str(siz3) ')'],'Color',[0.65,0.11,0.19],'FontSize',14), legend({'Global signal before motion correction'},'FontSize',12,'TextColor',[0.65,0.11,0.19],'Location','best')
        subplot2n(2,1,2), plot(mF2,'color',[0,0.55,0.69],'linewidth',1.5), hold on, plot(mF6,'color',[0.65,0.11,0.19],'linewidth',0.2), xlabel('time (in samples)'),ylabel('percentage signal change'),
          grid on,grid minor, xlim([1 length(mF2)]),ylim([min(1.25.*[mF2,mF6]) 1.25.*max([mF2,mF6])]), title(['Global mean signal after "08-motion-correction" (slice ' num2str(i3) ' of ' num2str(siz3) ')'],'Color',[0,0.55,0.69],'FontSize',14), legend({'Global signal after motion correction'},'FontSize',12,'TextColor',[0,0.55,0.69],'Location','best')
        saveas(fighndl,[QCpath 'global_signal_06denoise1_vs_08moco_slice' sprintf('%.2d',i3) '.jpg'])
        close(fighndl)
        
        C3 = corrcoef(F6'); C3=uppertriangle(C3);
        fighndl = figure('Position',[1 1 1152 720],'Color',[0.95,0.9,0.85],'InvertHardcopy','off','Visible','off'); % red [0.95,0.9,0.85], green [0.85,0.90,0.85], blue [0.89,0.94,0.99]
        subplot2n(2,1,1), histogram(C3,'EdgeColor','none','FaceColor',[0.65,0.11,0.19],'FaceAlpha',0.5,'Normalization','probability'); grid on,grid minor, xlim([-1 1]), hold on, xline(0,'Color','k','linewidth',1.5); hold off
          title(sprintf('Histogram of voxel-to-voxel correlations AFTER 06-DENOISE1 (slice %d of %d)\n%.1f%% of the correlations were > 0  [ mean=%.2f, median=%.2f, SD=%.2f ]',i3,siz3,100*length(find(C3>0))/length(C3),mean(C3),median(C3),std(C3)),'Color',[0.65,0.11,0.19],'FontSize',14)
        subplot2n(2,1,2), histogram(C,'EdgeColor','none','FaceColor',[0,0.55,0.69],'FaceAlpha',0.5,'Normalization','probability'); grid on,grid minor, xlim([-1 1]), hold on, xline(0,'Color','k','linewidth',1.5); hold off
          title(sprintf('Histogram of voxel-to-voxel correlations AFTER 08-MOTION-CORRECTION (slice %d of %d)\n%.1f%% of the correlations were > 0  [ mean=%.2f, median=%.2f, SD=%.2f ]',i3,siz3,100*length(find(C>0))/length(C),mean(C),median(C),std(C)),'Color',[0,0.55,0.69],'FontSize',14)
        saveas(fighndl,[QCpath 'hist_FC_compare_06denoise1_08moco_slice' sprintf('%.2d',i3) '.jpg'])
        close(fighndl)
        F6_=cat(1,F6_,F6); mF6_=cat(1,mF6_,mF6); C3_=cat(1,C3_,C3);
        clear F6 mF6 C3
    end
    
    F2_=cat(1,F2_,F2); mF2_=cat(1,mF2_,mF2); C_=cat(1,C_,C); F4_=cat(1,F4_,F4); mF4_=cat(1,mF4_,mF4); C2_=cat(1,C2_,C2);
    clear F2 F4 TR mask1 svar sdist mF2 mF4 C C2
end


[~,svar] = sort(var(F2_,0,2)); [~,spsc] = sort(meanPSC(F2_));
sortorder2 = sortorder; sortorder=[]; f=0; %#ok<*ASGLU>
for im=1:length(sortorder2)
    temp = sortorder2{im}; temp = temp + f; f = f + length(temp);
    sortorder = cat(1,sortorder,temp); clear temp
end; clear im f
sortorder = spsc; % options: spsc for percentage signal change, svar for variance, (comment out the line if you want to use distance from the center of the cord separately for each slice)
for im=max(sortorder):-1:1, if isempty(find(sortorder==im)), sortorder(find(sortorder>im)) = sortorder(find(sortorder>im))-1; end, end; clear im %#ok<*EFIND>

fighndl = figure('Position',[1 1 1152 720],'Color',[0.89,0.94,0.99],'InvertHardcopy','off','Visible','off'); % red [0.95,0.9,0.85], green [0.85,0.90,0.85], blue [0.89,0.94,0.99]
imagesc(F2_(sortorder,:)),colormap(cmap),colorbar, xlabel('time (in samples)'),ylabel('voxels (sorted by mean percentage signal change; bottom row is highest)'),
  title(['Time series within GM, WM and CSF AFTER 08-MOTION-CORRECTION (all slices combined)'],'Color',[0,0.55,0.69],'FontSize',14);
saveas(fighndl,[QCpath 'fMRI_ts_08moco_allSlices' '.jpg'])
saveas(fighndl,[QCpath2 'fMRI_ts_08moco_allSlices' '.jpg'])
 close(fighndl)
mF2_=mean(mF2_,1); mF4_=mean(mF4_,1);
fighndl = figure('Position',[1 1 1152 720],'Color',[0.89,0.94,0.99],'InvertHardcopy','off','Visible','off'); % red [0.95,0.9,0.85], green [0.85,0.90,0.85], blue [0.89,0.94,0.99]
subplot2n(2,1,1), plot(mF4_,'color',[0.65,0.11,0.19],'linewidth',1.5), hold on, plot(mF2_,'color',[0,0.55,0.69],'linewidth',0.2), xlabel('time (in samples)'),ylabel('percentage signal change'),
  grid on,grid minor, xlim([1 length(mF2_)]),ylim([1.25.*min([mF2_,mF4_]) 1.25.*max([mF2_,mF4_])]), title(['Global mean signal in UNPROCESSED RAW DATA (all slices averaged)'],'Color',[0.65,0.11,0.19],'FontSize',14), legend({'Global signal (raw data)'},'FontSize',12,'TextColor',[0.65,0.11,0.19],'Location','best')
subplot2n(2,1,2), plot(mF2_,'color',[0,0.55,0.69],'linewidth',1.5), hold on, plot(mF4_,'color',[0.65,0.11,0.19],'linewidth',0.2), xlabel('time (in samples)'),ylabel('percentage signal change'),
  grid on,grid minor, xlim([1 length(mF2_)]),ylim([1.25.*min([mF2_,mF4_]) 1.25.*max([mF2_,mF4_])]), title(['Global mean signal AFTER 08-MOTION-CORRECTION (all slices averaged)'],'Color',[0,0.55,0.69],'FontSize',14), legend({'Global signal after 08-motion-correction'},'FontSize',12,'TextColor',[0,0.55,0.69],'Location','best')
saveas(fighndl,[QCpath 'global_signal_08moco_allSlices' '.jpg'])
 close(fighndl)
fighndl = figure('Position',[1 1 1152 720],'Color',[0.89,0.94,0.99],'InvertHardcopy','off','Visible','off'); % red [0.95,0.9,0.85], green [0.85,0.90,0.85], blue [0.89,0.94,0.99]
subplot2n(2,1,1), histogram(C2_,'EdgeColor','none','FaceColor',[0.65,0.11,0.19],'FaceAlpha',0.5,'Normalization','probability'); grid on,grid minor, xlim([-1 1]), hold on, xline(0,'Color','k','linewidth',1.5); hold off
    title(sprintf('Histogram of voxel-to-voxel correlations in UNPROCESSED RAW DATA (all slices combined)\n%.1f%% of the correlations were > 0  [ mean=%.2f, median=%.2f, SD=%.2f ]',100*length(find(C2_>0))/length(C2_),mean(C2_),median(C2_),std(C2_)),'Color',[0.65,0.11,0.19],'FontSize',14)
subplot2n(2,1,2), histogram(C_,'EdgeColor','none','FaceColor',[0,0.55,0.69],'FaceAlpha',0.5,'Normalization','probability'); grid on,grid minor, xlim([-1 1]), hold on, xline(0,'Color','k','linewidth',1.5); hold off
    title(sprintf('Histogram of voxel-to-voxel correlations AFTER 08-MOTION-CORRECTION (all slices combined)\n%.1f%% of the correlations were > 0  [ mean=%.2f, median=%.2f, SD=%.2f ]',100*length(find(C_>0))/length(C_),mean(C_),median(C_),std(C_)),'Color',[0,0.55,0.69],'FontSize',14)
saveas(fighndl,[QCpath 'hist_FC_compare_08moco_allSlices' '.jpg'])
 close(fighndl)
if exist([base_dir_sub fname '_slice' num2str(1) '_denoised_before_MC' cutoff1_str '.nii'],'file') || exist([base_dir_sub fname '_slice' num2str(1) '_denoised_before_MC' cutoff1_str '.nii.gz'],'file') || exist([base_dir_sub fname '_denoised_before_MC' cutoff1_str '.nii'],'file') || exist([base_dir_sub fname '_denoised_before_MC' cutoff1_str '.nii.gz'],'file')
    mF6_=mean(mF6_,1);
    fighndl = figure('Position',[1 1 1152 720],'Color',[0.89,0.94,0.99],'InvertHardcopy','off','Visible','off'); % red [0.95,0.9,0.85], green [0.85,0.90,0.85], blue [0.89,0.94,0.99]
    subplot2n(2,1,1), plot(mF6_,'color',[0.65,0.11,0.19],'linewidth',1.5), hold on, plot(mF2_,'color',[0,0.55,0.69],'linewidth',0.2), xlabel('time (in samples)'),ylabel('percentage signal change'),
      grid on,grid minor, xlim([1 length(mF2_)]),ylim([1.25.*min([mF2_,mF6_]) 1.25.*max([mF2_,mF6_])]), title(['Global mean signal AFTER 06-DENOISE1 (all slices averaged)'],'Color',[0.65,0.11,0.19],'FontSize',14), legend({'Global signal before motion correction'},'FontSize',12,'TextColor',[0.65,0.11,0.19],'Location','best')
    subplot2n(2,1,2), plot(mF2_,'color',[0,0.55,0.69],'linewidth',1.5), hold on, plot(mF6_,'color',[0.65,0.11,0.19],'linewidth',0.2), xlabel('time (in samples)'),ylabel('percentage signal change'),
      grid on,grid minor, xlim([1 length(mF2_)]),ylim([1.25.*min([mF2_,mF6_]) 1.25.*max([mF2_,mF6_])]), title(['Global mean signal AFTER 08-MOTION-CORRECTION (all slices averaged)'],'Color',[0,0.55,0.69],'FontSize',14), legend({'Global signal after motion correction'},'FontSize',12,'TextColor',[0,0.55,0.69],'Location','best')
    saveas(fighndl,[QCpath 'global_signal_06denoise1_vs_08moco_allSlices' '.jpg'])
    saveas(fighndl,[QCpath2 'global_signal_06denoise1_vs_08moco_allSlices' '.jpg'])
     close(fighndl)
    fighndl = figure('Position',[1 1 1152 720],'Color',[0.89,0.94,0.99],'InvertHardcopy','off','Visible','off'); % red [0.95,0.9,0.85], green [0.85,0.90,0.85], blue [0.89,0.94,0.99]
    subplot2n(2,1,1), histogram(C3_,'EdgeColor','none','FaceColor',[0.65,0.11,0.19],'FaceAlpha',0.5,'Normalization','probability'); grid on,grid minor, xlim([-1 1]), hold on, xline(0,'Color','k','linewidth',1.5); hold off
      title(sprintf('Histogram of voxel-to-voxel correlations AFTER 06-DENOISE1 (all slices combined)\n%.1f%% of the correlations were > 0  [ mean=%.2f, median=%.2f, SD=%.2f ]',100*length(find(C3_>0))/length(C3_),mean(C3_),median(C3_),std(C3_)),'Color',[0.65,0.11,0.19],'FontSize',14)
    subplot2n(2,1,2), histogram(C_,'EdgeColor','none','FaceColor',[0,0.55,0.69],'FaceAlpha',0.5,'Normalization','probability'); grid on,grid minor, xlim([-1 1]), hold on, xline(0,'Color','k','linewidth',1.5); hold off
      title(sprintf('Histogram of voxel-to-voxel correlations AFTER 08-MOTION-CORRECTION (all slices combined)\n%.1f%% of the correlations were > 0  [ mean=%.2f, median=%.2f, SD=%.2f ]',100*length(find(C_>0))/length(C_),mean(C_),median(C_),std(C_)),'Color',[0,0.55,0.69],'FontSize',14)
    saveas(fighndl,[QCpath 'hist_FC_compare_06denoise1_08moco_allSlices' '.jpg'])
    saveas(fighndl,[QCpath2 'hist_FC_compare_06denoise1_08moco_allSlices' '.jpg'])
     close(fighndl)
end
clear svar spsc sortorder F2_ mF2_ C_ F4_ mF4_ C2_ F6_ mF6_ C3_ cmap

%% Make QC video (needs audio video toolbox in Matlab)
if ~isempty(which('VideoWriter')) && (save_videos==1) % check if Matlab's audio video toolbox exists

subplot = @(m,n,p) subtightplot(m, n, p, [0.001 0.005], [0.01 0.001], [0.01 0.001]);
for i3 = 1 : siz3
    try waitbar((i3/siz3),wbar3,sprintf('08. Generate QC video: slice (%d) of (%d)',i3,siz3)); catch, end
    fprintf('08. Generate QC video: slice (%d) of (%d)\n',i3,siz3)
    scfMRItb_04_resplitData(base_dir_sub, fname, '_MC', siz3) % if slice-wise data doesn't exist but combined 4D data exists, then split the data to generate the needed slice-wise data
    F1 = load_untouch_nii([base_dir_sub fname '_slice' num2str(i3) '_MC.nii']);
    TR = F1.hdr.dime.pixdim(5); F1 = rot90(single(squeeze(F1.img)));
    video = VideoWriter([QCpath 'fMRIvideo_08moco_slice' sprintf('%.2d',i3) '.mp4'],'MPEG-4'); %#ok<*TNMLP> % create the video object
    video.FrameRate = round(30/TR); % 30 seconds of data in every second of the video
    video.Quality = 90; % very limited compression so that the images are shown as is (and thus avoid misunderstandings due to video quality)
    open(video); % open the file for writing
    
    if Smask_err==0
        scfMRItb_04_resplitData(base_dir_sub, fname, '_Smask', siz3) % if slice-wise data doesn't exist but combined 4D data exists, then split the data to generate the needed slice-wise data
        scfMRItb_04_unzipFile(base_dir_sub, fname, ['_slice' num2str(i3) '_Smask'])
        mask1 = load_untouch_nii([base_dir_sub fname '_slice' num2str(i3) '_Smask.nii']); mask1 = mask1.img;
    else
        mask1 = Smask1_(:,:,i3);
    end
    mask1 = rot90(mask1);
    [xmask,ymask] = find(mask1==1);
    minx=round(min(xmask)-0.1*(size(mask1,1))); maxx=round(max(xmask)+0.1*(size(mask1,1)));
    miny=round(min(ymask)-0.1*(size(mask1,2))); maxy=round(max(ymask)+0.1*(size(mask1,2)));
    if (maxx-minx)<(maxy-miny)
        factr = ((maxy-miny)-(maxx-minx))/2;
        if factr==round(factr)
            maxx=maxx+factr; minx=minx-factr;
        else
            maxx=maxx+factr+0.5; minx=minx-factr+0.5;
        end
    elseif (maxy-miny)<(maxx-minx)
        factr = ((maxx-minx)-(maxy-miny))/2;
        if factr==round(factr)
            maxy=maxy+factr; miny=miny-factr;
        else
            maxy=maxy+factr+0.5; miny=miny-factr+0.5;
        end
    end
    [sizeFx,sizeFy] = size(F1(:,:,1));
    F2 = F1(minx:maxx,miny:maxy,:);
    for i5=1:size(F1,3)
        F1(:,:,i5) = imresize(F2(:,:,i5),[sizeFx sizeFy]);
    end
    scfMRItb_04_resplitData(base_dir_sub, fname, '', siz3) % if slice-wise data doesn't exist but combined 4D data exists, then split the data to generate the needed slice-wise data
    F3 = load_untouch_nii([base_dir_sub fname '_slice' num2str(i3) '.nii']); F3 = rot90(single(squeeze(F3.img)));
    F4 = F3(minx:maxx,miny:maxy,:);
    for i5=1:size(F3,3)
        F3(:,:,i5) = imresize(F4(:,:,i5),[sizeFx sizeFy]);
    end
    clear maxx maxy minx miny xmask ymask mask1 factr sizeFx sizeFy F2 F4 i5
    
    for ii=1:size(F1,3) % loop across the number of images
        I = F1(:,:,ii); %read the next image
        I(find(isnan(I)))=0;
        I = (I - min(I(:))) ./ (max(I(:)) - min(I(:)));
        I2 = F3(:,:,ii); %read the next image
        I2(find(isnan(I2)))=0;
        I2 = (I2 - min(I2(:))) ./ (max(I2(:)) - min(I2(:)));
        fighndl = figure('Position',[round((scrsz(3)-scrsz(4))/2) 1 round(0.75*scrsz(3)) round(0.75*scrsz(4))],'Color',[0.95,0.9,0.85],'InvertHardcopy','off','Visible','off'); % [0.85,0.90,0.85]
        subplot(1,2,1), imshow(I2, [0 max(I2(:))], 'InitialMagnification', 'fit'); title({['Unprocessed raw fMRI data (slice ' num2str(i3) ' of ' num2str(siz3) ')'];['Time point ' num2str(ii) ' of ' num2str(size(F1,3))]},'Color',[0.65,0.11,0.19],'FontSize',18);
        subplot(1,2,2), imshow(I, [0 max(I(:))], 'InitialMagnification', 'fit'); title({['fMRI data after "08. motion correction" (slice ' num2str(i3) ' of ' num2str(siz3) ')'];['Time point ' num2str(ii) ' of ' num2str(size(F1,3))]},'Color',[0,0.55,0.69],'FontSize',18);
        frame = getframe(gcf);
        writeVideo(video,frame); % write the image to file
        close(fighndl)
        clear I I2 frame
    end
    close(video); %close the file
    clear F1 F3 video ii TR
end; clear i3

end

try
if nargin<3
    close(wbar3)
end
catch
end

end
```
