## Supplementary material for "Neptune: a toolbox for spinal cord functional MRI data processing and quality assurance": Neptune software: scfMRItb_10_GMWMCSFmasks.m

```
function [time_taken] = scfMRItb_10_GMWMCSFmasks(base_dir_sub,fname,fname_anat_orig, varargin)

scrsz = get(0,'ScreenSize');
pause on;
fname3 = [fname '_mean'];
fname_anat = [fname_anat_orig '_reduced'];
time_taken = [];

if ~(exist([base_dir_sub 'QC'],'dir'))
    mkdir([base_dir_sub 'QC'])
end
if ~(exist([base_dir_sub 'QC' '/10_GM-WM-CSF-masks'],'dir'))
    mkdir([base_dir_sub 'QC' '/10_GM-WM-CSF-masks'])
end
QCpath = [base_dir_sub 'QC' '/10_GM-WM-CSF-masks' '/'];


if nargin<6
    mask_GM_flag = 1; % 1 = estimate GM boundary using SCT deepseg, 2 = estimate GM boundary by co-registration to PAM50 and transforming the standard GM mask in SCT.
else
    mask_GM_flag = varargin{3};
end
retry_GMseg = 0; % 1 = in case mask_GM_flag is 2 and if PAM50 registration fails then get the GM boundaries by deepseg instead; 0 = don't try deepseg if SCT registration fails (default)
GMseg_backupOption = 1; % 1 = even if PAM50 registration was done, perform deepseg to get a backup GMseg boundary for use if the user is not satisfied with the PAM50 boundary (default). 0 = don't do this.

if nargin<7
    choices_preproc = ones(19,1);
else
    choices_preproc = varargin{4};
end
if sum([choices_preproc(3);choices_preproc(5:9)])==0 % if you are running neptune only to do anatomical segmentation AND if neptune was not previously run on this data
    if ~exist([base_dir_sub fname_anat '.nii'], 'file') && exist([base_dir_sub fname_anat_orig '.nii'], 'file')
        copyfile([base_dir_sub fname_anat_orig '.nii'], [base_dir_sub fname_anat '.nii'], 'f')
    end
    if ~exist([base_dir_sub fname_anat '.nii.gz'], 'file') && exist([base_dir_sub fname_anat_orig '.nii.gz'], 'file')
        copyfile([base_dir_sub fname_anat_orig '.nii.gz'], [base_dir_sub fname_anat '.nii.gz'], 'f')
    end
end

if nargin<8
    C_spine = 1;
else
    C_spine = varargin{5};
end

if nargin<9
    SCT_vert_auto = 1; % 1 = assume that we will perform automatic labeling of vertebrae
else
    SCT_vert_auto = varargin{6};
end

if nargin<10
    SCT_manualreg_labels = '3,5'; % Vertebrae for manual labeling in SCT (if manual option was chosen), needed for registration
else
    SCT_manualreg_labels = varargin{7};
end

scfMRItb_04_unzipFile(base_dir_sub, fname_anat, '')
A = load_untouch_nii([base_dir_sub fname_anat '.nii']);
siz3 = size(A.img,3);

% By default, do NOT recreate these masks if they already exist (this is a time-consuming step)
if ~( (exist([base_dir_sub fname_anat '_mask_GM.nii'], 'file')) || (exist([base_dir_sub fname_anat '_mask_GM.nii.gz'], 'file')) )

if nargin<4
    wbar3 = waitbar(0,'10. Define GM, WM and CSF masks...','Name','Progress(10): Define GM, WM and CSF masks...','Units','normalized','Position',[4/5 0 1/5 1/17]);
else
    wbar3 = varargin{1};
end
if nargin<5
    exist_SCT = 1; % 1 = spinal cord toolbox exists, 0 = doesn't exist
else
    exist_SCT = varargin{2};
end

% ----- GENERATE X AND Y LIMITS FOR VIEWING THE IMAGES --------------------
for i3 = 1 : siz3
    try
        scfMRItb_04_resplitData(base_dir_sub, fname, '_Smask', siz3) % if slice-wise data doesn't exist but combined 4D data exists, then split the data to generate the needed slice-wise data
        scfMRItb_04_unzipFile(base_dir_sub, fname, ['_slice' num2str(i3) '_Smask'])
        Smask = load_untouch_nii([base_dir_sub fname '_slice' num2str(i3) '_Smask.nii']);
    catch
        Smask.img = zeros(size(A.img,1),size(A.img,2)); Smask.img(round(size(A.img,1)/4):round(3*size(A.img,1)/4),round(size(A.img,2)/4):round(3*size(A.img,2)/4))=1;
    end
    Smask = rot90(Smask.img);
    [w2,w1] = find(Smask~=0);
    w1min(i3,1) = min(w1)/size(Smask,1); w1max(i3,1) = max(w1)/size(Smask,1);
    w2min(i3,1) = min(w2)/size(Smask,2); w2max(i3,1) = max(w2)/size(Smask,2);
    clear Smask w1 w2
end

%% Use SCT (if available) to get an estimate of GM, WM and CSF boundaries

if exist_SCT==1 % this works on, at least, SCT v5.4 and later (I tested it with SCT v5.4 only)
    try
    fprintf('\n10. Running SCT (if required) to estimate the boundary of GM, WM and CSF...\n\n')
    if ~(exist([base_dir_sub 'QC' '/SCT'],'dir'))
        mkdir([base_dir_sub 'QC' '/SCT'])
    end
    
    if ~exist([base_dir_sub fname_anat '_seg.nii'],'file')
        unix(['sct_deepseg_sc -i ' base_dir_sub fname_anat '.nii' ' -c t2s -ofolder ' base_dir_sub ' -qc ' base_dir_sub 'QC/SCT'])
        WM_seg  = load_untouch_nii([base_dir_sub fname_anat '_seg.nii']); WM_seg=uint8(WM_seg.img);
        movefile([base_dir_sub fname_anat '_seg.nii'],[base_dir_sub fname_anat '_seg2.nii'],'f')
    else
        WM_seg  = load_untouch_nii([base_dir_sub fname_anat '_seg.nii']); WM_seg=uint8(WM_seg.img);
    end
    
    if ~exist([base_dir_sub fname_anat '_CSF_seg.nii'],'file')
        unix(['sct_propseg -i ' base_dir_sub fname_anat '.nii' ' -c t2s -ofolder ' base_dir_sub ' -CSF' ' -qc ' base_dir_sub 'QC/SCT'])
        CSF_seg = load_untouch_nii([base_dir_sub fname_anat '_CSF_seg.nii']); CSF_seg=uint8(CSF_seg.img);
        delete([base_dir_sub fname_anat '_seg.nii'])
        movefile([base_dir_sub fname_anat '_seg2.nii'],[base_dir_sub fname_anat '_seg.nii'])
    else
        CSF_seg = load_untouch_nii([base_dir_sub fname_anat '_CSF_seg.nii']); CSF_seg=uint8(CSF_seg.img);
    end
    
    
    if mask_GM_flag==1
    if ~exist([base_dir_sub fname_anat '_gmseg.nii'],'file')
        unix(['sct_deepseg_gm -i ' base_dir_sub fname_anat '.nii' ' -o ' base_dir_sub fname_anat '_gmseg.nii' ' -qc ' base_dir_sub 'QC/SCT'])
        unix(['sct_maths -i ' base_dir_sub fname_anat '_seg.nii' ' -sub ' base_dir_sub fname_anat '_gmseg.nii' ' -o ' base_dir_sub fname_anat '_wmseg.nii'])
    end
    GM_seg = load_untouch_nii([base_dir_sub fname_anat '_gmseg.nii']); GM_seg=uint8(GM_seg.img);
    
    
    elseif mask_GM_flag==2
	flag1 = 0;
	try %#ok<ALIGN>
    if ~( (exist([base_dir_sub 'SCT_template2anat_step10/' 'PAM50_gm.nii.gz'],'file')) || (exist([base_dir_sub 'SCT_template2anat_step10/' 'PAM50_gm.nii'],'file')) )
        scfMRItb_10_GMmask_PAM50(base_dir_sub,fname_anat, SCT_vert_auto,SCT_manualreg_labels,C_spine)
        gunzip([base_dir_sub 'SCT_template2anat_step10/' 'PAM50_gm.nii.gz'])
        GM_seg = load_untouch_nii([base_dir_sub 'SCT_template2anat_step10/' 'PAM50_gm.nii']); GM_seg=uint8(GM_seg.img);
    end
    catch
        if retry_GMseg==1 % 1 = in case mask_GM_flag is 2 and if PAM50 registration fails then get the GM boundaries by deepseg instead; 0 = don't try deepseg if SCT registration fails (default)
            if ~exist([base_dir_sub fname_anat '_gmseg.nii'],'file')
                unix(['sct_deepseg_gm -i ' base_dir_sub fname_anat '.nii' ' -o ' base_dir_sub fname_anat '_gmseg.nii' ' -qc ' base_dir_sub 'QC/SCT'])
                unix(['sct_maths -i ' base_dir_sub fname_anat '_seg.nii' ' -sub ' base_dir_sub fname_anat '_gmseg.nii' ' -o ' base_dir_sub fname_anat '_wmseg.nii'])
            end
            GM_seg = load_untouch_nii([base_dir_sub fname_anat '_gmseg.nii']); GM_seg=uint8(GM_seg.img);
            flag1 = 1;
        end
    end
    
    if (GMseg_backupOption==1) && (flag1==0)
    if ~exist([base_dir_sub fname_anat '_gmseg.nii'],'file')
        unix(['sct_deepseg_gm -i ' base_dir_sub fname_anat '.nii' ' -o ' base_dir_sub fname_anat '_gmseg.nii' ' -qc ' base_dir_sub 'QC/SCT'])
        unix(['sct_maths -i ' base_dir_sub fname_anat '_seg.nii' ' -sub ' base_dir_sub fname_anat '_gmseg.nii' ' -o ' base_dir_sub fname_anat '_wmseg.nii'])
    end
    GM_seg2 = load_untouch_nii([base_dir_sub fname_anat '_gmseg.nii']); GM_seg2=uint8(GM_seg2.img);
    end

    if ~exist('GM_seg','var') % happens if SCT PAM50 co-registration failed altogether
        GM_seg = GM_seg2;
    end
    end

    
    CSF_seg = CSF_seg + WM_seg; CSF_seg(find(CSF_seg~=0)) = 1;
    for i3 = 1 : size(CSF_seg,3)
        CSF_seg(:,:,i3) = imfill(CSF_seg(:,:,i3),'holes');
        CSF_boundarymask = boundarymask(rot90(CSF_seg(:,:,i3)));
         [row,col] = find(CSF_boundarymask==1);
         if ~isempty(row)
         CSF_boundary{i3,1} = bwtraceboundary(CSF_boundarymask,[row(1),col(1)],'N');
         fctr = round(size(CSF_boundary{i3,1},1)/15);
         CSF_boundary{i3,1} = CSF_boundary{i3,1}(1:fctr:end,:);
         else
         CSF_boundary{i3,1} = [];
         end
         clear CSF_boundarymask row col fctr
        WM_boundarymask = boundarymask(rot90(WM_seg(:,:,i3)));
         [row,col] = find(WM_boundarymask==1);
         if ~isempty(row)
         WM_boundary{i3,1} = bwtraceboundary(WM_boundarymask,[row(1),col(1)],'N');
         fctr = round(size(WM_boundary{i3,1},1)/15);
         WM_boundary{i3,1} = WM_boundary{i3,1}(1:fctr:end,:);
         else
         WM_boundary{i3,1} = [];
         end
         clear WM_boundarymask row col fctr
        GM_boundarymask = boundarymask(rot90(GM_seg(:,:,i3)));
         [row,col] = find(GM_boundarymask==1);
         if ~isempty(row)
         GM_boundary{i3,1} = bwtraceboundary(GM_boundarymask,[row(1),col(1)],'N');
         fctr = round(size(GM_boundary{i3,1},1)/30);
         GM_boundary{i3,1} = GM_boundary{i3,1}(1:fctr:end,:);
         else
         GM_boundary{i3,1} = [];
         end
         clear GM_boundarymask row col fctr
        if exist('GM_seg2','var')
        GM_boundarymask2 = boundarymask(rot90(GM_seg2(:,:,i3)));
         [row,col] = find(GM_boundarymask2==1);
         if ~isempty(row)
         GM_boundary2{i3,1} = bwtraceboundary(GM_boundarymask2,[row(1),col(1)],'N');
         fctr = round(size(GM_boundary2{i3,1},1)/30);
         GM_boundary2{i3,1} = GM_boundary2{i3,1}(1:fctr:end,:);
         else
         GM_boundary2{i3,1} = [];
         end
         clear GM_boundarymask2 row col fctr
        end
    end
    clear CSF_seg WM_seg GM_seg GM_seg2
    fprintf('\nCompleted sct_propseg, sct_deepseg_sc and sct_deepseg_gm to estimate the boundary of CSF, WM and GM.\n\n')
    catch
    end
end

%% ----- DEFINE GM / WM / CSF MASKS ----------------------------------------
if exist([base_dir_sub fname_anat '_all_masks.mat'], 'file')
    % load previous results (in case some masks need to be re-done)
    eval(['load ' base_dir_sub fname_anat '_all_masks.mat']);
else % this is the first time that this code has been executed
    mask_GM  = A;
    mask_WM  = A;
    mask_CSF = A;
end
new_size = round(size(A.img,1) / 2);
pause on;

%% CSF
position_CSF = cell(siz3,1);
for i3 = 1 : siz3
    try waitbar((0/3)+(i3/siz3)/3,wbar3,sprintf('10. Define CSF mask: slice (%d) of (%d)',i3,siz3)); catch, end
    img = rot90(A.img(:,:,i3));
    temp = img(round((size(img,1)-new_size)/2)+1:end-round((size(img,1)-new_size)/2), ...
        round((size(img,2)-new_size)/2)+1:end-round((size(img,2)-new_size)/2));
    mmax = max(temp(:)); clear temp;
    fighndl = figure('Position',[round((scrsz(3)-scrsz(4))/2) 1 scrsz(4) scrsz(4)],'InvertHardcopy','off'); fighndl.Color = [0.8, 0.87, 0.84]; try addToolbarExplorationButtons(fighndl); catch, end
    imshow(single(img), [0 mmax], 'InitialMagnification', 'fit'); colorbar;
    fighndl.Color = [0.8, 0.87, 0.84] + [0, 0.02*(1-exp(siz3/i3)/exp(siz3)), 0.1*exp(i3/siz3)/exp(1)]; % [0.85, 0.90, 0.85] + [0, 0, 0.1*i3/siz3];
    title({'Draw the boundary of CSF (double click the cord once done)';['Slice ' num2str(i3) ' of ' num2str(siz3)]},'Color',[0.7,0,0.7],'FontSize',16);
    disp(['Slice ' num2str(i3) ' of ' num2str(siz3) ': Draw the boundary of CSF.']);
    if i3==1
        xlim([round(size(img,1)*w1min(i3)),round(size(img,1)*w1max(i3))]),ylim([round(size(img,2)*w2min(i3)),round(size(img,2)*w2max(i3))]),
    else
        xlim([min([round(size(img,1)*w1min(i3)),lims_x1_(i3-1)]),max([round(size(img,1)*w1max(i3)),lims_x2_(i3-1)])]),
        ylim([min([round(size(img,2)*w2min(i3)),lims_y1_(i3-1)]),max([round(size(img,2)*w2max(i3)),lims_y2_(i3-1)])]),
    end
    time1 = tic;
    
    if exist('CSF_boundary','var') && ~isempty(CSF_boundary{i3})
        h = impoly(gca,[CSF_boundary{i3}(:,2),CSF_boundary{i3}(:,1)]);
    else
        h = impoly;
    end
    position = wait(h);
    if isempty(position)
        while isempty(position)
        h = impoly;
        position = wait(h);
        end
    end
    time_taken(i3,1) = toc(time1);
    
    bmask = poly2mask(position(:,1), position(:,2), size(A.img,2), size(A.img,1));
    mask_CSF.img(:,:,i3) = rot90(bmask, -1);
    saveas(fighndl,[base_dir_sub 'QC' '/10_GM-WM-CSF-masks' '/' 'CSFmask_slice' sprintf('%.2d',i3) '.jpg'])
    clear img mmax h bmask time1
    close(fighndl)
    lims_x1(i3) = min(position(:,1)); lims_x2(i3) = max(position(:,1)); %#ok<*AGROW>
    lims_x1_(i3) = round(lims_x1(i3) - 0.15*(lims_x2(i3)-lims_x1(i3))); lims_x2_(i3) = round(lims_x2(i3) + 0.15*(lims_x2(i3)-lims_x1(i3)));
    lims_y1(i3) = min(position(:,2)); lims_y2(i3) = max(position(:,2));
    lims_y1_(i3) = round(lims_y1(i3) - 0.15*(lims_y2(i3)-lims_y1(i3))); lims_y2_(i3) = round(lims_y2(i3) + 0.15*(lims_y2(i3)-lims_y1(i3)));
    position_CSF{i3,1} = position; clear position
    eval(['save ' base_dir_sub fname_anat '_all_masks_incomplete.mat ' 'mask_CSF position_CSF;']);
end

%% WM
position_WM = cell(siz3,1);
for i3 = 1 : siz3
    try waitbar((1/3)+(i3/siz3)/3,wbar3,sprintf('10. Define white matter mask: slice (%d) of (%d)',i3,siz3)); catch, end
    img = rot90(A.img(:,:,i3));
    temp = img(round((size(img,1)-new_size)/2)+1:end-round((size(img,1)-new_size)/2), ...
        round((size(img,2)-new_size)/2)+1:end-round((size(img,2)-new_size)/2));
    mmax = max(temp(:)); clear temp;
    fighndl = figure('Position',[round((scrsz(3)-scrsz(4))/2) 1 scrsz(4) scrsz(4)],'InvertHardcopy','off'); fighndl.Color = [0.8, 0.87, 0.84]; try addToolbarExplorationButtons(fighndl); catch, end
    imshow(single(img), [0 mmax], 'InitialMagnification', 'fit'); colorbar;
    fighndl.Color = [0.8, 0.87, 0.84] + [0, 0.02*(1-exp(siz3/i3)/exp(siz3)), 0.1*exp(i3/siz3)/exp(1)]; % [0.85, 0.90, 0.85] + [0, 0, 0.1*i3/siz3];
    title({'Draw the boundary of the white matter (double click the cord once done)';['Slice ' num2str(i3) ' of ' num2str(siz3)]},'Color',[0.70,0.13,0.13],'FontSize',16);
    disp(['Slice ' num2str(i3) ' of ' num2str(siz3) ': Draw the boundary of the white matter.']);
    hold on, plot([position_CSF{i3}(:,1);position_CSF{i3}(1,1)],[position_CSF{i3}(:,2);position_CSF{i3}(1,2)],'m','LineWidth',2), hold off
    xlim([round(lims_x1(i3))-1,round(lims_x2(i3))+1]), ylim([round(lims_y1(i3))-1,round(lims_y2(i3))+1]),
    legend('CSF mask')
    time1 = tic;

    if exist('WM_boundary','var') && ~isempty(WM_boundary{i3})
        h = impoly(gca,[WM_boundary{i3}(:,2),WM_boundary{i3}(:,1)]);
    else
        h = impoly;
    end
    position = wait(h);
    if isempty(position)
        while isempty(position)
        h = impoly;
        position = wait(h);
        end
    end
    time_taken(i3,2) = toc(time1);
    
    bmask = poly2mask(position(:,1), position(:,2), size(A.img,2), size(A.img,1));
    mask_WM.img(:,:,i3) = rot90(bmask, -1);
    saveas(fighndl,[base_dir_sub 'QC' '/10_GM-WM-CSF-masks' '/' 'WMmask_slice' sprintf('%.2d',i3) '.jpg'])
    clear img mmax h bmask
    close(fighndl)
    lims_x1(i3) = min(position(:,1)); lims_x2(i3) = max(position(:,1));
    lims_y1(i3) = min(position(:,2)); lims_y2(i3) = max(position(:,2));
    position_WM{i3,1} = position; clear position
    eval(['save ' base_dir_sub fname_anat '_all_masks_incomplete.mat ' 'mask_WM mask_CSF position_WM position_CSF;']);
end

%% GM
position_GM = cell(siz3,1);
for i3 = 1 : siz3
    try waitbar((2/3)+(i3/siz3)/3,wbar3,sprintf('10. Define gray matter mask: slice (%d) of (%d)',i3,siz3)); catch, end
    img = rot90(A.img(:,:,i3));
    temp = img(round((size(img,1)-new_size)/2)+1:end-round((size(img,1)-new_size)/2), ...
        round((size(img,2)-new_size)/2)+1:end-round((size(img,2)-new_size)/2));
    mmax = max(temp(:)); clear temp;
    fighndl = figure('Position',[round((scrsz(3)-scrsz(4))/2) 1 scrsz(4) scrsz(4)],'InvertHardcopy','off'); fighndl.Color = [0.8, 0.87, 0.84]; try addToolbarExplorationButtons(fighndl); catch, end
	imshow(single(img), [0 mmax], 'InitialMagnification', 'fit'); colorbar;
    fighndl.Color = [0.8, 0.87, 0.84] + [0, 0.02*(1-exp(siz3/i3)/exp(siz3)), 0.1*exp(i3/siz3)/exp(1)]; % [0.85, 0.90, 0.85] + [0, 0, 0.1*i3/siz3];
    title({'Draw the boundary of the gray matter (double click the GM once done)';['Slice ' num2str(i3) ' of ' num2str(siz3)]},'Color',[0,0.33,0.53],'FontSize',16);
    disp(['Slice ' num2str(i3) ' of ' num2str(siz3) ': Draw the boundary of the gray matter.']);
    hold on, plot([position_CSF{i3}(:,1);position_CSF{i3}(1,1)],[position_CSF{i3}(:,2);position_CSF{i3}(1,2)],'Color',[0.7,0,0.7],'LineWidth',2), plot([position_WM{i3}(:,1);position_WM{i3}(1,1)],[position_WM{i3}(:,2);position_WM{i3}(1,2)],'Color',[0.70,0.13,0.13],'LineWidth',2), hold off
    xlim([round(lims_x1(i3))-1,round(lims_x2(i3))+1]), ylim([round(lims_y1(i3))-1,round(lims_y2(i3))+1]),
    legend('CSF mask','WM mask')
    time1 = tic;
    
    
    if exist('GM_boundary','var') && ~isempty(GM_boundary{i3})
        h = impoly(gca,[GM_boundary{i3}(:,2),GM_boundary{i3}(:,1)]);
    else
        h = impoly;
    end
    position = wait(h);
    
    if isempty(position)
    if exist('GM_boundary2','var') && ~isempty(GM_boundary2{i3})
        h = impoly(gca,[GM_boundary2{i3}(:,2),GM_boundary2{i3}(:,1)]); setColor(h,"yellow")
    else
        h = impoly;
    end
    position = wait(h);
    end
    
    if isempty(position)
    if exist('GM_boundary','var') && ~isempty(GM_boundary{i3}) && exist('GM_boundary2','var') && ~isempty(GM_boundary2{i3})
        h = impoly(gca,[GM_boundary2{i3}(:,2),GM_boundary2{i3}(:,1)]); setColor(h,"yellow")
        h = impoly(gca,[GM_boundary{i3}(:,2),GM_boundary{i3}(:,1)]);
    else
        h = impoly;
    end
    position = wait(h);
    end
    
    if isempty(position)
        while isempty(position)
            close(fighndl)
            clear h position fighndl
            fighndl = figure('Position',[round((scrsz(3)-scrsz(4))/2) 1 scrsz(4) scrsz(4)],'InvertHardcopy','off'); fighndl.Color = [0.8, 0.87, 0.84]; try addToolbarExplorationButtons(fighndl); catch, end
	        imshow(single(img), [0 mmax], 'InitialMagnification', 'fit'); colorbar;
            fighndl.Color = [0.8, 0.87, 0.84] + [0, 0.02*(1-exp(siz3/i3)/exp(siz3)), 0.1*exp(i3/siz3)/exp(1)]; % [0.85, 0.90, 0.85] + [0, 0, 0.1*i3/siz3];
            title({'Draw the boundary of the gray matter (double click the GM once done)';['Slice ' num2str(i3) ' of ' num2str(siz3)]},'Color',[0,0.33,0.53],'FontSize',16);
            disp(['Slice ' num2str(i3) ' of ' num2str(siz3) ': Draw the boundary of the gray matter.']);
            hold on, plot([position_CSF{i3}(:,1);position_CSF{i3}(1,1)],[position_CSF{i3}(:,2);position_CSF{i3}(1,2)],'Color',[0.7,0,0.7],'LineWidth',2), plot([position_WM{i3}(:,1);position_WM{i3}(1,1)],[position_WM{i3}(:,2);position_WM{i3}(1,2)],'Color',[0.70,0.13,0.13],'LineWidth',2), hold off
            xlim([round(lims_x1(i3))-1,round(lims_x2(i3))+1]), ylim([round(lims_y1(i3))-1,round(lims_y2(i3))+1]),
            legend('CSF mask','WM mask')
        h = impoly;
        position = wait(h);
        end
    end
    time_taken(i3,3) = toc(time1);
    
    
    bmask = poly2mask(position(:,1), position(:,2), size(A.img,2), size(A.img,1));
    mask_GM.img(:,:,i3) = rot90(bmask, -1);
    saveas(fighndl,[base_dir_sub 'QC' '/10_GM-WM-CSF-masks' '/' 'GMmask_slice' sprintf('%.2d',i3) '.jpg'])
    close(fighndl)
    lims_x1(i3) = min(position(:,1)); lims_x2(i3) = max(position(:,1));
    lims_y1(i3) = min(position(:,2)); lims_y2(i3) = max(position(:,2));
    position_GM{i3,1} = position;
    eval(['save ' base_dir_sub fname_anat '_all_masks_incomplete.mat ' 'mask_GM mask_WM mask_CSF position_GM position_WM position_CSF;']);
    
    fighndl2 = figure('Position',[round((scrsz(3)-scrsz(4))/2) 1 scrsz(4) scrsz(4)],'InvertHardcopy','off');
	imshow(single(img), [0 mmax], 'InitialMagnification', 'fit'); colorbar;
    fighndl2.Color = [0.75, 0.84, 0.99];
    hold on, plot([position_CSF{i3}(:,1);position_CSF{i3}(1,1)],[position_CSF{i3}(:,2);position_CSF{i3}(1,2)],'Color',[0.7,0,0.7],'LineWidth',2), plot([position_WM{i3}(:,1);position_WM{i3}(1,1)],[position_WM{i3}(:,2);position_WM{i3}(1,2)],'Color',[0.70,0.13,0.13],'LineWidth',2), plot([position_GM{i3}(:,1);position_GM{i3}(1,1)],[position_GM{i3}(:,2);position_GM{i3}(1,2)],'Color',[0,0.33,0.53],'LineWidth',2), hold off
    legend('CSF mask','WM mask','GM mask'), title({sprintf('Final GM/WM/CSF masks for slice-%d',i3);'(click anywhere on the image to zoom/close)'},'Color','red','FontSize',18)
    saveas(fighndl2,[base_dir_sub 'QC' '/10_GM-WM-CSF-masks' '/' 'final_masks_slice' sprintf('%.2d',i3) '_fullFOV' '.jpg'])
    waitforbuttonpress
    xlim([lims_x1_(i3),lims_x2_(i3)]), ylim([lims_y1_(i3),lims_y2_(i3)]),
    saveas(fighndl2,[base_dir_sub 'QC' '/10_GM-WM-CSF-masks' '/' 'final_masks_slice' sprintf('%.2d',i3) '_zoomed' '.jpg'])
    waitforbuttonpress
    close(fighndl2)
    clear img mmax h bmask position fighndl2
end
clear fighndl lims_x1 lims_x2 lims_y1 lims_y2 lims_y2_ lims_y1_ lims_x2_ lims_x1_

%% save files
eval(['save ' base_dir_sub fname_anat '_all_masks.mat ' 'mask_GM mask_WM mask_CSF position_GM position_WM position_CSF;']);
eval(['delete ' base_dir_sub fname_anat '_all_masks_incomplete.mat;']);

mask_CSF.img = mask_CSF.img - mask_WM.img;
mask_WM.img = mask_WM.img - mask_GM.img;
% NOTE: check to ensure that there are no negative values in mask_CSF or mask_WM (which would indicate that the borders cross - a bad thing).
sum(sum(sum(mask_WM.img==-1)))
sum(sum(sum(mask_CSF.img==-1)))

for i1 = 1 : size(mask_GM.img,1)
    for i2 = 1 : size(mask_GM.img,2)
        for i3 = 1 : size(mask_GM.img,3)
            if mask_CSF.img(i1,i2,i3) == -1
                mask_CSF.img(i1,i2,i3) = 0;
            end
            if mask_WM.img(i1,i2,i3) == -1
                mask_WM.img(i1,i2,i3) = 0;
            end
        end
    end
end

% also create a mask of the spinal cord (GM + WM)
mask_SC = mask_GM;
mask_SC.img = or(mask_GM.img, mask_WM.img);

save_untouch_nii(mask_CSF, [base_dir_sub fname_anat '_mask_CSF.nii']);
save_untouch_nii(mask_WM,  [base_dir_sub fname_anat '_mask_WM.nii']);
save_untouch_nii(mask_GM,  [base_dir_sub fname_anat '_mask_GM.nii']);
save_untouch_nii(mask_SC,  [base_dir_sub fname_anat '_mask_SC.nii']);
clear A clear mask_* new_size img temp mmax;

end % if

%% ----- CREATE MASKS FOR ANATOMICAL & FUNCTIONALS -------------------------

if ~( exist([base_dir_sub fname_anat '_slice' num2str(siz3) '_affine_reg_mask.nii'], 'file') || exist([base_dir_sub fname_anat '_slice' num2str(siz3) '_affine_reg_mask.nii.gz'], 'file') || exist([base_dir_sub fname_anat '_affine_reg_mask.nii'], 'file') || exist([base_dir_sub fname_anat '_affine_reg_mask.nii.gz'], 'file') )  ||  ~( exist([base_dir_sub fname '_slice' num2str(siz3) '_base_masked.nii'], 'file') || exist([base_dir_sub fname '_slice' num2str(siz3) '_base_masked.nii.gz'], 'file') || exist([base_dir_sub fname '_base_masked.nii'], 'file') || exist([base_dir_sub fname '_base_masked.nii.gz'], 'file') )

scfMRItb_04_unzipFile(base_dir_sub, fname_anat, '_mask_GM')
scfMRItb_04_unzipFile(base_dir_sub, fname_anat, '_mask_WM')
scfMRItb_04_unzipFile(base_dir_sub, fname_anat, '_mask_CSF')
mask_GM  = load_untouch_nii([base_dir_sub fname_anat '_mask_GM.nii']);
mask_WM  = load_untouch_nii([base_dir_sub fname_anat '_mask_WM.nii']);
mask_CSF = load_untouch_nii([base_dir_sub fname_anat '_mask_CSF.nii']);

mask_all = or(or(mask_GM.img, mask_WM.img), mask_CSF.img) > 0;
mask_all_d = imdilate(mask_all, strel('disk', 20));

% consider only GM, WM, and a boundary condition (a ring around CSF)
affine_reg_mask = 5.00 .* single(mask_GM.img) + 5.00 .* single(mask_WM.img) + ...
    + 5.00 .* single(mask_CSF.img) + 5.00 .* xor(mask_all, mask_all_d);

try
scfMRItb_04_unzipFile(base_dir_sub, fname, '_Gaussian_mask')
F_Gmask = load_untouch_nii([base_dir_sub fname '_Gaussian_mask.nii']);
funct_mask = F_Gmask.img;
catch
end

% ----- SAVE MASKS FOR ANATOMICAL & FUNCTIONALS ---------------------------
% save affine_reg_mask and funct_mask

try    
for i3 = 1 : siz3
    try waitbar((i3/siz3),wbar3,sprintf('10. Save affine_reg_mask: slice (%d) of (%d)',i3,siz3)); catch, end
    unix(['rm -f ' base_dir_sub fname_anat '_slice' num2str(i3) '_affine_reg_mask.nii']);
    scfMRItb_04_resplitData(base_dir_sub, fname_anat, '', siz3) % if slice-wise data doesn't exist but combined 4D data exists, then split the data to generate the needed slice-wise data
    scfMRItb_04_unzipFile(base_dir_sub, fname_anat, ['_slice' num2str(i3)])
    AM = load_untouch_nii([base_dir_sub fname_anat '_slice' num2str(i3) '.nii']);
    AM.img = affine_reg_mask(:,:,i3);
    save_untouch_nii(AM, [base_dir_sub fname_anat '_slice' num2str(i3) '_affine_reg_mask.nii']);
    clear AM;
end
catch
end

try
scfMRItb_04_unzipFile(base_dir_sub, fname3, '')
F = load_untouch_nii([base_dir_sub fname3 '.nii']);
for i3 = 1 : size(F.img,3)
    try waitbar((i3/size(F.img,3)),wbar3,sprintf('10. Save masked functional: slice (%d) of (%d)',i3,size(F.img,3))); catch, end
    unix(['rm -f ' base_dir_sub fname '_slice' num2str(i3) '_base_masked.nii']);
    scfMRItb_04_resplitData(base_dir_sub, fname, '_base', siz3) % if slice-wise data doesn't exist but combined 4D data exists, then split the data to generate the needed slice-wise data
    scfMRItb_04_unzipFile(base_dir_sub, fname, ['_slice' num2str(i3) '_base'])
    FM = load_untouch_nii([base_dir_sub fname '_slice' num2str(i3) '_base.nii']);
    FM.img = single(FM.img) .* funct_mask(:,:,i3); % functional * mask
    save_untouch_nii(FM, [base_dir_sub fname '_slice' num2str(i3) '_base_masked.nii']);
    clear FM;
end
catch
end

end

%% ----- COMPUTE CROSS-SECTIONAL AREAS (CSA) FROM ANATOMICAL SEGMENTATIONS -------------------------

if ~(exist([base_dir_sub 'cross_sectional_areas_Neptune.mat'], 'file'))

mask_GM  = load_untouch_nii([base_dir_sub fname_anat '_mask_GM.nii']);
mask_WM  = load_untouch_nii([base_dir_sub fname_anat '_mask_WM.nii']);
mask_CSF = load_untouch_nii([base_dir_sub fname_anat '_mask_CSF.nii']);
pixdims(1) = mask_GM.hdr.dime.pixdim(2); pixdims(2) = mask_GM.hdr.dime.pixdim(3); % in mm

for i3=1:size(mask_GM.img,3)
    temp = length(find(mask_GM.img(:,:,i3)==1));
    GM_area(i3,1)  = temp * pixdims(1) * pixdims(2); clear temp
    temp = length(find(mask_WM.img(:,:,i3)==1));
    WM_area(i3,1)  = temp * pixdims(1) * pixdims(2); clear temp
    temp = length(find(mask_CSF.img(:,:,i3)==1));
    CSF_area(i3,1) = temp * pixdims(1) * pixdims(2); clear temp
    cross_sectional_area(i3,1) = GM_area(i3,1) + WM_area(i3,1);
end; clear i3

readme_units = sprintf('All areas are in millimeters squared\n(Note: these area measurements have not been validated, hence, please do not use this for research or publication purposes. This is for rough quality assessment only.)\n');
save([base_dir_sub 'cross_sectional_areas_Neptune.mat'],'GM_area','WM_area','CSF_area','cross_sectional_area','readme_units')

end

if ~exist([QCpath 'cross_sectional_areas_across_slices' '.jpg'],'file')
for w=1:size(mask_GM.img,3)
    dispstring1{w} = sprintf('%.2f',GM_area(w));
    dispstring2{w} = sprintf('%.2f',WM_area(w));
    dispstring3{w} = sprintf('%.2f',CSF_area(w));
    dispstring4{w} = sprintf('%.2f',cross_sectional_area(w));
end; clear w
fighndl = figure('Position',[1 1 1280 800],'Color',[0.85,0.90,0.85],'InvertHardcopy','off','Visible','off'); % red [0.95,0.9,0.85], green [0.85,0.90,0.85], blue [0.89,0.94,0.99]
plot(GM_area,'--^','color',[0,0.55,0.69],'linewidth',2.25, 'MarkerSize',12, 'MarkerEdgeColor',[0,0.55,0.69], 'MarkerFaceColor',[0.89,0.94,0.99]),
    hold on, plot(WM_area,'--v','color',[0.65,0.11,0.19],'linewidth',2.25, 'MarkerSize',12, 'MarkerEdgeColor',[0.65,0.11,0.19], 'MarkerFaceColor',[0.95,0.9,0.85]),
    plot(CSF_area,'--o','color',[0.25,0.25,0.25],'linewidth',2.25, 'MarkerSize',14, 'MarkerEdgeColor','k', 'MarkerFaceColor',[0.85,0.85,0.85]),
    plot(cross_sectional_area,'--p','color','b','linewidth',2.25, 'MarkerSize',18, 'MarkerEdgeColor',[0 0.37 0.22], 'MarkerFaceColor',[0.85,0.90,0.85]),
    grid on,grid minor, ylabel('area (in millimeters squared)'), xlabel('SLICES'), xticks([1:length(GM_area)]), 
    xlim([0.5 length(GM_area)+0.5]), ylim([0 max([GM_area;WM_area;CSF_area;cross_sectional_area])+8])
    text([1:length(GM_area)]-0.1,GM_area+6, dispstring1,'Color',[0,0.55,0.69],'FontSize',12)
    text([1:length(WM_area)]-0.1,WM_area-6, dispstring2,'Color',[0.65,0.11,0.19],'FontSize',12)
    text([1:length(CSF_area)]-0.1,CSF_area+6, dispstring3,'Color',[0 0.37 0.22],'FontSize',12)
    text([1:length(cross_sectional_area)]-0.1,cross_sectional_area-6, dispstring4,'Color',[0.05,0.05,0.05],'FontSize',12)
    legend({'Gray matter area';'White matter area';'Cerebrospinal fluid area';'Cross-sectional area (GM+WM)'},'FontSize',14,'TextColor','k','Location','best')
    title(sprintf('CROSS SECTIONAL AREAS across slices (units: millimeters squared)\n(Note: these area measurements have not been validated, hence, please do not use this for research or publication purposes. \nThis is for rough quality assessment only.)'),'Color',[0.65,0.11,0.19],'FontSize',16)
saveas(fighndl,[QCpath 'cross_sectional_areas_across_slices' '.jpg'])
close(fighndl)
end

try
if nargin<4
    close(wbar3)
end
catch
end

end
```
