## Supplementary material for "Neptune: a toolbox for spinal cord functional MRI data processing and quality assurance": Neptune software: scfMRItb_10_GMWMCSFmasks_run2.m

```
function scfMRItb_10_GMWMCSFmasks_run2(base_dir_sub,fname, varargin)

fname3 = [fname '_mean'];

if nargin<3
    wbar3 = waitbar(0,'10. Define GM, WM and CSF masks...','Name','Progress(10): Define GM, WM and CSF masks...','Units','normalized','Position',[3/5 0 1/5 1/17]);
else
    wbar3 = varargin{1};
end

% ----- CREATE MASKS FOR FUNCTIONALS --------------------------------------

scfMRItb_04_unzipFile(base_dir_sub, fname, '_Gaussian_mask')
F_Gmask = load_untouch_nii([base_dir_sub fname '_Gaussian_mask.nii']);
funct_mask = F_Gmask.img;

% ----- SAVE MASKS FOR FUNCTIONALS ----------------------------------------

scfMRItb_04_unzipFile(base_dir_sub, fname3, '')
F = load_untouch_nii([base_dir_sub fname3 '.nii']);
siz3 = size(F.img,3);
for i3 = 1 : siz3
    try waitbar((i3/size(F.img,3)),wbar3,sprintf('10. Save masked functional: slice (%d) of (%d)',i3,size(F.img,3))); catch, end
    unix(['rm -f ' base_dir_sub fname '_slice' num2str(i3) '_base_masked.nii']);
    scfMRItb_04_resplitData(base_dir_sub, fname, '_base', siz3) % if slice-wise data doesn't exist but combined 4D data exists, then split the data to generate the needed slice-wise data
    scfMRItb_04_unzipFile(base_dir_sub, fname3, ['_slice' num2str(i3) '_base'])
    FM = load_untouch_nii([base_dir_sub fname '_slice' num2str(i3) '_base.nii']);
    FM.img = FM.img .* funct_mask(:,:,i3); % functional * mask
    save_untouch_nii(FM, [base_dir_sub fname '_slice' num2str(i3) '_base_masked.nii']);
    clear FM
end

try
if nargin<3
    close(wbar3)
end
catch
end

end
```
