## Supplementary material for "Neptune: a toolbox for spinal cord functional MRI data processing and quality assurance": Neptune software: scfMRItb_11_FuncAnatRegistration.m

```
function scfMRItb_11_FuncAnatRegistration(base_dir_sub,fname,fname_anat_orig,run, varargin)

fname_anat = [fname_anat_orig '_reduced'];

scfMRItb_04_unzipFile(base_dir_sub, fname, '')
F = load_untouch_nii([base_dir_sub fname '.nii']);
siz3 = size(F.img,3);
siz4 = size(F.img,4);

if nargin<5
    wbar3 = waitbar(0,'11. Co-registration (functional to anatomical)...','Name','Progress(11): Co-registration (functional to anatomical)...','Units','normalized','Position',[3/5 0 1/5 1/17]);
else
    wbar3 = varargin{1};
end

if nargin<6
    save_videos = 1; % 1 = save QC videos (generates sizeable .mp4 files and takes a few minutes to do it). This is turned on by default, but if you don't want to save videos then set it to 0
else
    save_videos = varargin{2};
end

if nargin<7
    Rns = 0; % 0 = don't create runX subfolder; X = create runX subfolder
else
    Rns = varargin{3};
end

if ~(exist([base_dir_sub 'QC'],'dir'))
    mkdir([base_dir_sub 'QC'])
end
if ~(exist([base_dir_sub 'QC' '/11_func-anat-registration'],'dir'))
    mkdir([base_dir_sub 'QC' '/11_func-anat-registration'])
end
if Rns==0
    QCpath = [base_dir_sub 'QC' '/11_func-anat-registration/'];
else
    if ~(exist([base_dir_sub 'QC' '/11_func-anat-registration' '/run' num2str(Rns)],'dir'))
        mkdir([base_dir_sub 'QC' '/11_func-anat-registration' '/run' num2str(Rns)])
    end
    QCpath = [base_dir_sub 'QC' '/11_func-anat-registration' '/run' num2str(Rns) '/'];
end

if ~(exist([base_dir_sub 'QC' '/QC_shortlist'],'dir'))
    mkdir([base_dir_sub 'QC' '/QC_shortlist'])
end
if Rns==0
    QCpath2 = [base_dir_sub 'QC' '/QC_shortlist/'];
else
    if ~(exist([base_dir_sub 'QC' '/QC_shortlist' '/run' num2str(Rns)],'dir'))
        mkdir([base_dir_sub 'QC' '/QC_shortlist' '/run' num2str(Rns)])
    end
    QCpath2 = [base_dir_sub 'QC' '/QC_shortlist' '/run' num2str(Rns) '/'];
end

subplot = @(m,n,p) subtightplot(m, n, p, [0.001 0.005], [0.01 0.001], [0.01 0.001]);

%% ----- REGISTER AVERAGE FUNCTIONALS TO ANATOMICAL ------------------------

try waitbar(0.5,wbar3,'11. Co-registration'); catch, end
[final_reg, adj_final_trans,use_cmass] = spinal_cord_registration_9b(base_dir_sub, fname, fname_anat, siz3, siz4, wbar3);
save([base_dir_sub 'run' num2str(run) '_final_reg.mat'], 'final_reg', 'adj_final_trans'); % run number (1 or 2 or ...)

scrsz = get(0,'ScreenSize'); pause on;
eval(['load ' base_dir_sub fname_anat '_all_masks.mat ' 'position_GM position_WM position_CSF;']);
try
    scfMRItb_04_unzipFile(base_dir_sub, fname_anat, '_mask_SC')
    mask_SC2 = load_untouch_nii([base_dir_sub fname_anat '_mask_SC.nii']); mask_SC2 = mask_SC2.img;
catch
    scfMRItb_04_unzipFile(base_dir_sub, fname_anat, '_mask_GM')
    scfMRItb_04_unzipFile(base_dir_sub, fname_anat, '_mask_WM')
    mask_GM = load_untouch_nii([base_dir_sub fname_anat '_mask_GM.nii']);
    mask_WM = load_untouch_nii([base_dir_sub fname_anat '_mask_WM.nii']);
    mask_SC = mask_GM;
    mask_SC.img = or(mask_GM.img, mask_WM.img);
    save_untouch_nii(mask_SC, [base_dir_sub fname_anat '_mask_SC.nii']);
    mask_SC2 = mask_SC; clear mask_SC
end

for i3 = 1 : siz3
    try waitbar(2/4+(i3/siz3)/4,wbar3,sprintf('11. Co-registration (3/4) reg avg func to anat: slice (%d) of (%d)',i3,siz3)); catch, end
    if ~exist([base_dir_sub fname_anat '_slice' num2str(i3) '.nii'],'file')
        scfMRItb_04_resplitData(base_dir_sub, fname_anat, '', siz3);
    end
    if ~exist([base_dir_sub fname '_slice' num2str(i3) '_MC_ricor_mean.nii'],'file')
        suffix = '_MC_ricor_mean';
        scfMRItb_04_resplitData(base_dir_sub, fname, suffix, siz3); clear suffix
    end
    if ~exist([base_dir_sub fname '_slice' num2str(i3) '_warped_mean.nii'],'file')
        suffix = '_warped_mean';
        scfMRItb_04_resplitData(base_dir_sub, fname, suffix, siz3); clear suffix
    end
    unix(['touch ' base_dir_sub fname_anat '_slice' num2str(i3) '.nii']); % save time when viewing files
    unix(['rm -f ' base_dir_sub fname '_slice' num2str(i3) '_warped_mean.nii']);
    if use_cmass==0
    runerr = unix(['3dAllineate -verb -final wsinc5 ' ... 
                '-input ' base_dir_sub fname '_slice' num2str(i3) '_MC_ricor_mean.nii ' ...
                '-prefix ' base_dir_sub fname '_slice' num2str(i3) '_warped_mean.nii ' ...
                '-master ' base_dir_sub fname_anat '_slice' num2str(i3) '.nii ' ...
                '-1Dmatrix_apply ' base_dir_sub fname '_slice' num2str(i3) '_xform.aff12.1D']);
    else
    runerr = unix(['3dAllineate -verb -final wsinc5 ' ... 
                '-input ' base_dir_sub fname '_slice' num2str(i3) '_MC_ricor_mean.nii ' ...
                '-prefix ' base_dir_sub fname '_slice' num2str(i3) '_warped_mean.nii ' ...
                '-master ' base_dir_sub fname_anat '_slice' num2str(i3) '.nii ' ...
                '-cmass ' ...
                '-1Dmatrix_apply ' base_dir_sub fname '_slice' num2str(i3) '_xform.aff12.1D']);
    end
    
	if runerr~=0 %#ok<ALIGN>  if RETROICOR was skipped or not performed, the above unix command will return an error upon which the following command will be executed using pre-RETROICOR motion corrected data
        if use_cmass==0
        unix(['3dAllineate -verb -final wsinc5 ' ...
            '-input ' base_dir_sub fname '_slice' num2str(i3) '_MC_mean.nii ' ...
            '-prefix ' base_dir_sub fname '_slice' num2str(i3) '_warped_mean.nii ' ...
            '-master ' base_dir_sub fname_anat '_slice' num2str(i3) '.nii ' ...
            '-1Dmatrix_apply ' base_dir_sub fname '_slice' num2str(i3) '_xform.aff12.1D']);
        else
        unix(['3dAllineate -verb -final wsinc5 ' ...
            '-input ' base_dir_sub fname '_slice' num2str(i3) '_MC_mean.nii ' ...
            '-prefix ' base_dir_sub fname '_slice' num2str(i3) '_warped_mean.nii ' ...
            '-master ' base_dir_sub fname_anat '_slice' num2str(i3) '.nii ' ...
            '-cmass ' ...
            '-1Dmatrix_apply ' base_dir_sub fname '_slice' num2str(i3) '_xform.aff12.1D']);
        end
    end

    img_anat = load_untouch_nii([base_dir_sub fname_anat '_slice' num2str(i3) '.nii']); img_anat=img_anat.img;
    img_warped_mean = load_untouch_nii([base_dir_sub fname '_slice' num2str(i3) '_warped_mean.nii']); img_warped_mean=img_warped_mean.img;
    fighndl = figure('Position',[1 1 1120 1120],'Color',[0.85,0.90,0.85],'InvertHardcopy','off','Visible','off'); % [0.85,0.90,0.85] [0.8, 0.89, 0.94] [0.89, 0.94, 0.99] [361 1 scrsz(4) scrsz(4)]
    subtightplot(2,2,1,[0.035 0.035], [0.035 0.035], [0.035 0.035]), imshow(rot90(img_anat), [0 max(img_anat(:))]), title(['Anatomical image: (slice ' num2str(i3) ' of ' num2str(siz3) ')'],'Color',[0.65,0.11,0.19],'FontSize',16)
        try xlim([round(min(position_CSF{i3}(:,1))-0.125*(size(mask_SC2,1))),round(max(position_CSF{i3}(:,1))+0.125*(size(mask_SC2,1)))]), ylim([round(min(position_CSF{i3}(:,2))-0.125*(size(mask_SC2,2))),round(max(position_CSF{i3}(:,2))+0.125*(size(mask_SC2,2)))]), catch, end
    subtightplot(2,2,2,[0.035 0.035], [0.035 0.035], [0.035 0.035]), imshow(rot90(img_warped_mean), [0 max(img_warped_mean(:))]), title(['Co-registered mean functional image: (slice ' num2str(i3) ' of ' num2str(siz3) ')'],'Color',[0,0.55,0.69],'FontSize',16)
        try xlim([round(min(position_CSF{i3}(:,1))-0.125*(size(mask_SC2,1))),round(max(position_CSF{i3}(:,1))+0.125*(size(mask_SC2,1)))]), ylim([round(min(position_CSF{i3}(:,2))-0.125*(size(mask_SC2,2))),round(max(position_CSF{i3}(:,2))+0.125*(size(mask_SC2,2)))]), catch, end
    subtightplot(2,2,3,[0.035 0.035], [0.035 0.035], [0.035 0.035]), imshow(rot90(img_anat), [0 max(img_anat(:))]), 
        try xlim([round(size(img_anat,1)/3)+1,2*round(size(img_anat,1)/3),]), ylim([round(size(img_anat,2)/3)+1,2*round(size(img_anat,2)/3),]), title(['Anatomical image: (slice ' num2str(i3) ' of ' num2str(siz3) ')'],'Color',[0.65,0.11,0.19],'FontSize',16), catch, end
        hold on, try plot([position_CSF{i3}(:,1);position_CSF{i3}(1,1)],[position_CSF{i3}(:,2);position_CSF{i3}(1,2)],'Color',[0.7,0,0.7],'LineWidth',2), plot([position_WM{i3}(:,1);position_WM{i3}(1,1)],[position_WM{i3}(:,2);position_WM{i3}(1,2)],'Color',[0.70,0.13,0.13],'LineWidth',2), plot([position_GM{i3}(:,1);position_GM{i3}(1,1)],[position_GM{i3}(:,2);position_GM{i3}(1,2)],'Color',[0,0.33,0.53],'LineWidth',2), catch, end, hold off
        try xlim([round(min(position_CSF{i3}(:,1))-0.03*(size(mask_SC2,1))),round(max(position_CSF{i3}(:,1))+0.03*(size(mask_SC2,1)))]), ylim([round(min(position_CSF{i3}(:,2))-0.03*(size(mask_SC2,2))),round(max(position_CSF{i3}(:,2))+0.03*(size(mask_SC2,2)))]),  catch, end
    subtightplot(2,2,4,[0.035 0.035], [0.035 0.035], [0.035 0.035]), imshow(rot90(img_warped_mean), [0 max(img_warped_mean(:))]), 
        try xlim([round(size(img_warped_mean,1)/3)+1,2*round(size(img_warped_mean,1)/3)]), ylim([round(size(img_warped_mean,2)/3)+1,2*round(size(img_warped_mean,2)/3)]), title(['Co-registered mean functional image: (slice ' num2str(i3) ' of ' num2str(siz3) ')'],'Color',[0,0.55,0.69],'FontSize',16), catch, end
        hold on, try plot([position_CSF{i3}(:,1);position_CSF{i3}(1,1)],[position_CSF{i3}(:,2);position_CSF{i3}(1,2)],'Color',[0.7,0,0.7],'LineWidth',2), plot([position_WM{i3}(:,1);position_WM{i3}(1,1)],[position_WM{i3}(:,2);position_WM{i3}(1,2)],'Color',[0.70,0.13,0.13],'LineWidth',2), plot([position_GM{i3}(:,1);position_GM{i3}(1,1)],[position_GM{i3}(:,2);position_GM{i3}(1,2)],'Color',[0,0.33,0.53],'LineWidth',2), catch, end, hold off
        try xlim([round(min(position_CSF{i3}(:,1))-0.03*(size(mask_SC2,1))),round(max(position_CSF{i3}(:,1))+0.03*(size(mask_SC2,1)))]), ylim([round(min(position_CSF{i3}(:,2))-0.03*(size(mask_SC2,2))),round(max(position_CSF{i3}(:,2))+0.03*(size(mask_SC2,2)))]), catch, end
    try suptitle2(sprintf('Outcome of Step 11: functional to anatomical registration\nComparing co-registered mean functional image with the anatomical image for slice %s of %s',num2str(i3),num2str(siz3))); fighndl.Visible='off'; catch, end
    saveas(fighndl,[QCpath 'func_anat_reg_slice' sprintf('%.2d',i3) '.jpg'])
    close(fighndl)
    clear runerr img_anat img_warped_mean
end
clear maxx_SC maxy_SC minx_SC miny_SC

%% ----- SLICE-BY-SLICE AFFINE REGISTRATION --------------------------------

for i3 = 1 : siz3
    try waitbar(3/4+(i3/siz3)/4,wbar3,sprintf('11. Co-registration (4/4) slice-by-slice affine reg: slice (%d) of (%d)',i3,siz3)); catch, end
    unix(['rm -f ' base_dir_sub fname '_slice' num2str(i3) '_warped.nii']);
    if ~exist([base_dir_sub fname '_slice' num2str(i3) '_MC_ricor.nii'],'file')
        suffix = '_MC_ricor';
        scfMRItb_04_resplitData(base_dir_sub, fname, suffix, siz3); clear suffix
    end
    runerr = unix(['3dAllineate -verb -final wsinc5 ' ... 
                '-input ' base_dir_sub fname '_slice' num2str(i3) '_MC_ricor.nii ' ...
                '-prefix ' base_dir_sub fname '_slice' num2str(i3) '_warped.nii ' ...
                '-master ' base_dir_sub fname_anat '_slice' num2str(i3) '.nii ' ...
                '-1Dmatrix_apply ' base_dir_sub fname '_slice' num2str(i3) '_xform.aff12.1D']);

	if runerr~=0 %#ok<ALIGN>  if RETROICOR was skipped or not performed, the above unix command will return an error upon which the following command will be executed using pre-RETROICOR motion corrected data
        if ~exist([base_dir_sub fname '_slice' num2str(i3) '_MC.nii'],'file')
            suffix = '_MC';
            scfMRItb_04_resplitData(base_dir_sub, fname, suffix, siz3); clear suffix
        end
        unix(['3dAllineate -verb -final wsinc5 ' ...
            '-input ' base_dir_sub fname '_slice' num2str(i3) '_MC.nii ' ...
            '-prefix ' base_dir_sub fname '_slice' num2str(i3) '_warped.nii ' ...
            '-master ' base_dir_sub fname_anat '_slice' num2str(i3) '.nii ' ...
            '-1Dmatrix_apply ' base_dir_sub fname '_slice' num2str(i3) '_xform.aff12.1D']);
    end
    clear runerr
end


% Move affine registration transformations to a subdirectory within the subject's main directory
sdirm = dir([base_dir_sub fname '*' '_xform.aff12.1D']);
for i5=length(sdirm):-1:1
    if ~isempty(strfind(sdirm(i5).name,'MC'))
        sdirm(i5) = [];
    end
end
mkdir([base_dir_sub 'co-registrations'])
for i5=1:length(sdirm)
    movefile([sdirm(i5).folder '/' sdirm(i5).name],[base_dir_sub 'co-registrations' '/' sdirm(i5).name])
end; clear sdirm i5

%% time series plots, global signal and FC histogram
scrsz = get(0,'ScreenSize'); pause on;
cmap = colormap('bone'); cmap = (1-exp(-cmap))./(1-exp(-1)); 

scfMRItb_04_unzipFile(base_dir_sub, fname_anat, '_mask_GM')
scfMRItb_04_unzipFile(base_dir_sub, fname_anat, '_mask_WM')
mask_GM = load_untouch_nii([base_dir_sub fname_anat '_mask_GM.nii']); mask_GM = mask_GM.img;
mask_WM = load_untouch_nii([base_dir_sub fname_anat '_mask_WM.nii']); mask_WM = mask_WM.img;

Smask_err=0;
try
    scfMRItb_04_resplitData(base_dir_sub, fname, '_Smask', siz3) % if slice-wise data doesn't exist but combined 4D data exists, then split the data to generate the needed slice-wise data
catch
    load([base_dir_sub fname '_mask_NS' '.mat'])
    Smask1_ = ~mask_NS.img; clear mask_NS
    Smask_err=1;
end

F2g_=[]; mF2g_=[]; F2w_=[]; mF2w_=[]; Cg_=[]; Cw_=[]; F4_=[]; mF4_=[]; C2_=[];
if exist([base_dir_sub fname '_slice' num2str(i3) '_MC.nii'],'file') || exist([base_dir_sub fname '_slice' num2str(i3) '_MC.nii.gz'],'file') || exist([base_dir_sub fname '_MC.nii'],'file') || exist([base_dir_sub fname '_MC.nii.gz'],'file')
    F6_=[]; mF6_=[]; C3_=[];
end
for i3 = 1 : siz3
    try waitbar((i3/siz3),wbar3,sprintf('11. QC plots: slice (%d) of (%d)',i3,siz3)); catch, end
    fprintf('11. QC plots: slice (%d) of (%d)\n',i3,siz3)
    scfMRItb_04_resplitData(base_dir_sub, fname, '_warped', siz3) % if slice-wise data doesn't exist but combined 4D data exists, then split the data to generate the needed slice-wise data
    scfMRItb_04_unzipFile(base_dir_sub, fname, ['_slice' num2str(i3) '_warped'])
    F1 = load_untouch_nii([base_dir_sub fname '_slice' num2str(i3) '_warped.nii']);
    TR = F1.hdr.dime.pixdim(5); F1 = single(squeeze(F1.img));
    
    if Smask_err==0
        scfMRItb_04_resplitData(base_dir_sub, fname, '_Smask', siz3) % if slice-wise data doesn't exist but combined 4D data exists, then split the data to generate the needed slice-wise data
        scfMRItb_04_unzipFile(base_dir_sub, fname, ['_slice' num2str(i3) '_Smask'])
        Smask1 = load_untouch_nii([base_dir_sub fname '_slice' num2str(i3) '_Smask.nii']); Smask1 = Smask1.img;
    else
        Smask1 = Smask1_(:,:,i3);
    end
    
    for i5=1:size(F1,3)
        tmp = F1(:,:,i5);
        F2g(:,i5) = tmp(find(mask_GM(:,:,i3)==1));
        F2w(:,i5) = tmp(find(mask_WM(:,:,i3)==1));
        clear tmp
    end; clear i5 F1
    [~,svar1] = sort(var(F2g,0,2)); [~,sdist1] = sort(svar1); sortorder1{i3,1} = sdist1;
    [~,svar2] = sort(var(F2w,0,2)); [~,sdist2] = sort(svar2); sortorder2{i3,1} = sdist2;
    
    fighndl = figure('Position',[1 1 1152 720],'Color',[0.95,0.9,0.85],'InvertHardcopy','off','Visible','off'); % red [0.95,0.9,0.85], green [0.85,0.90,0.85], blue [0.89,0.94,0.99]
    subplot2n(2,1,1), imagesc(F2g(sortorder1{i3},:)),colormap(cmap),colorbar, xlabel('time (in samples)'),ylabel('Gray matter voxels (sorted by variance)'),
        title(['Time series within the GRAY MATTER after "11-functional-anatomical-registration" in slice ' num2str(i3) ' of ' num2str(siz3)],'Color',[0,0.55,0.69],'FontSize',14);
    subplot2n(2,1,2), imagesc(F2w(sortorder2{i3},:)),colormap(cmap),colorbar, xlabel('time (in samples)'),ylabel('White matter voxels (sorted by variance)'),
        title(['Time series within the WHITE MATTER after "11-functional-anatomical-registration" in slice ' num2str(i3) ' of ' num2str(siz3)],'Color',[0.65,0.11,0.19],'FontSize',14);
    saveas(fighndl,[QCpath 'fMRI_ts_11coreg_slice' sprintf('%.2d',i3) '.jpg'])
    close(fighndl)
    
    scfMRItb_04_resplitData(base_dir_sub, fname, '', siz3) % if slice-wise data doesn't exist but combined 4D data exists, then split the data to generate the needed slice-wise data
    scfMRItb_04_unzipFile(base_dir_sub, fname, ['_slice' num2str(i3)])
    F3 = load_untouch_nii([base_dir_sub fname '_slice' num2str(i3) '.nii']); F3 = single(squeeze(F3.img));
    for i5=1:size(F3,3)
        tmp = F3(:,:,i5);
        F4(:,i5) = tmp(find(Smask1==1)); clear tmp
    end; clear i5 F3
    mF2g=mean(F2g,1); mF2g = 100*(mF2g-mean(mF2g))./mean(mF2g); mF2w=mean(F2w,1); mF2w = 100*(mF2w-mean(mF2w))./mean(mF2w); mF4=mean(F4,1); mF4 = 100*(mF4-mean(mF4))./mean(mF4);
    fighndl = figure('Position',[1 1 1152 720],'Color',[0.95,0.9,0.85],'InvertHardcopy','off','Visible','off'); % red [0.95,0.9,0.85], green [0.85,0.90,0.85], blue [0.89,0.94,0.99]
    subplot2n(3,1,1), plot(mF4,'color','k','linewidth',1.5), hold on, plot(mF2g,'color',[0,0.55,0.69],'linewidth',0.2), plot(mF2w,'color',[0.65,0.11,0.19],'linewidth',0.2), xlabel('time (in samples)'),ylabel('percentage signal change'),
      grid on,grid minor, xlim([1 length(mF2g)]),ylim([min(1.25.*[mF2g,mF2w,mF4]) 1.25.*max([mF2g,mF2w,mF4])]), title(['Global mean signal in UNPROCESSED RAW DATA (slice ' num2str(i3) ' of ' num2str(siz3) ')'],'Color','k','FontSize',14), legend({'Global signal (raw data)'},'FontSize',12,'TextColor','k','Location','best')
    subplot2n(3,1,2), plot(mF2g,'color',[0,0.55,0.69],'linewidth',1.5), hold on, plot(mF4,'color','k','linewidth',0.2), xlabel('time (in samples)'),ylabel('percentage signal change'),
      grid on,grid minor, xlim([1 length(mF2g)]),ylim([min(1.25.*[mF2g,mF2w,mF4]) 1.25.*max([mF2g,mF2w,mF4])]), title(['Gray matter mean signal AFTER "11-functional-anatomical-registration" (slice ' num2str(i3) ' of ' num2str(siz3) ')'],'Color',[0,0.55,0.69],'FontSize',14), legend({'Gray matter signal after 11-registration'},'FontSize',12,'TextColor',[0,0.55,0.69],'Location','best')
    subplot2n(3,1,3), plot(mF2w,'color',[0.65,0.11,0.19],'linewidth',1.5), hold on, plot(mF4,'color','k','linewidth',0.2), xlabel('time (in samples)'),ylabel('percentage signal change'),
      grid on,grid minor, xlim([1 length(mF2g)]),ylim([min(1.25.*[mF2g,mF2w,mF4]) 1.25.*max([mF2g,mF2w,mF4])]), title(['White matter mean signal AFTER "11-functional-anatomical-registration" (slice ' num2str(i3) ' of ' num2str(siz3) ')'],'Color',[0.65,0.11,0.19],'FontSize',14), legend({'White matter signal after 11-registration'},'FontSize',12,'TextColor',[0.65,0.11,0.19],'Location','best')
    saveas(fighndl,[QCpath 'global_signal_11coreg_slice' sprintf('%.2d',i3) '.jpg'])
    close(fighndl)

    Cg = corrcoef(F2g'); Cg = uppertriangle(Cg);
    Cw = corrcoef(F2w'); Cw = uppertriangle(Cw);
    C2 = corrcoef(F4');  C2 = uppertriangle(C2);
    fighndl = figure('Position',[1 1 1152 720],'Color',[0.95,0.9,0.85],'InvertHardcopy','off','Visible','off'); % red [0.95,0.9,0.85], green [0.85,0.90,0.85], blue [0.89,0.94,0.99]
    subplot2n(3,1,1), histogram(C2,'EdgeColor','none','FaceColor','k','FaceAlpha',0.5,'Normalization','probability'); grid on,grid minor, xlim([-1 1]), hold on, xline(0,'Color','k','linewidth',1.5); hold off
      title(sprintf('Histogram of voxel-to-voxel correlations in UNPROCESSED RAW DATA (slice %d of %d)\n%.1f%% of the correlations were > 0  [ mean=%.2f, median=%.2f, SD=%.2f ]',i3,siz3,100*length(find(C2>0))/length(C2),mean(C2),median(C2),std(C2)),'Color','k','FontSize',14)
    subplot2n(3,1,2), histogram(Cg,'EdgeColor','none','FaceColor',[0,0.55,0.69],'FaceAlpha',0.5,'Normalization','probability'); grid on,grid minor, xlim([-1 1]), hold on, xline(0,'Color','k','linewidth',1.5); hold off
      title(sprintf('Histogram of voxel-to-voxel GRAY MATTER correlations AFTER 11-registration (slice %d of %d)\n%.1f%% of the correlations were > 0  [ mean=%.2f, median=%.2f, SD=%.2f ]',i3,siz3,100*length(find(Cg>0))/length(Cg),mean(Cg),median(Cg),std(Cg)),'Color',[0,0.55,0.69],'FontSize',14)
    subplot2n(3,1,3), histogram(Cw,'EdgeColor','none','FaceColor',[0.65,0.11,0.19],'FaceAlpha',0.5,'Normalization','probability'); grid on,grid minor, xlim([-1 1]), hold on, xline(0,'Color','k','linewidth',1.5); hold off
      title(sprintf('Histogram of voxel-to-voxel WHITE MATTER correlations AFTER 11-registration (slice %d of %d)\n%.1f%% of the correlations were > 0  [ mean=%.2f, median=%.2f, SD=%.2f ]',i3,siz3,100*length(find(Cw>0))/length(Cw),mean(Cw),median(Cw),std(Cw)),'Color',[0.65,0.11,0.19],'FontSize',14)
    saveas(fighndl,[QCpath 'hist_FC_compare_11coreg_slice' sprintf('%.2d',i3) '.jpg'])
    close(fighndl)
    
    if exist([base_dir_sub fname '_slice' num2str(i3) '_MC.nii'],'file') || exist([base_dir_sub fname '_slice' num2str(i3) '_MC.nii.gz'],'file') || exist([base_dir_sub fname '_MC.nii'],'file') || exist([base_dir_sub fname '_MC.nii.gz'],'file')
        scfMRItb_04_resplitData(base_dir_sub, fname, '_MC', siz3) % if slice-wise data doesn't exist but combined 4D data exists, then split the data to generate the needed slice-wise data
        scfMRItb_04_unzipFile(base_dir_sub, fname, ['_slice' num2str(i3) '_MC'])
        F5 = load_untouch_nii([base_dir_sub fname '_slice' num2str(i3) '_MC.nii']); F5 = single(squeeze(F5.img));
        for i5=1:size(F5,3)
            tmp = F5(:,:,i5);
            F6(:,i5) = tmp(find(Smask1==1)); clear tmp
        end; clear i5 F5
        mF6=mean(F6,1); mF6 = 100*(mF6-mean(mF6))./mean(mF6);
        fighndl = figure('Position',[1 1 1152 720],'Color',[0.95,0.9,0.85],'InvertHardcopy','off','Visible','off'); % red [0.95,0.9,0.85], green [0.85,0.90,0.85], blue [0.89,0.94,0.99]
        subplot2n(3,1,1), plot(mF6,'color','k','linewidth',1.5), hold on, plot(mF2g,'color',[0,0.55,0.69],'linewidth',0.2), plot(mF2w,'color',[0.65,0.11,0.19],'linewidth',0.2), xlabel('time (in samples)'),ylabel('percentage signal change'),
            grid on,grid minor, xlim([1 length(mF2g)]),ylim([1.25.*min([mF2g,mF2w,mF6]) 1.25.*max([mF2g,mF2w,mF6])]), title(['Global mean signal AFTER "08-motion-correction" (slice ' num2str(i3) ' of ' num2str(siz3) ')'],'Color','k','FontSize',14), legend({'Global signal before co-registration'},'FontSize',12,'TextColor','k','Location','best')
        subplot2n(3,1,2), plot(mF2g,'color',[0,0.55,0.69],'linewidth',1.5), hold on, plot(mF6,'color','k','linewidth',0.2), xlabel('time (in samples)'),ylabel('percentage signal change'),
            grid on,grid minor, xlim([1 length(mF2g)]),ylim([1.25.*min([mF2g,mF2w,mF6]) 1.25.*max([mF2g,mF2w,mF6])]), title(['Gray matter mean signal AFTER "11-functional-anatomical-registration" (slice ' num2str(i3) ' of ' num2str(siz3) ')'],'Color',[0,0.55,0.69],'FontSize',14), legend({'Gray matter signal after co-registration'},'FontSize',12,'TextColor',[0,0.55,0.69],'Location','best')
        subplot2n(3,1,3), plot(mF2w,'color',[0.65,0.11,0.19],'linewidth',1.5), hold on, plot(mF6,'color','k','linewidth',0.2), xlabel('time (in samples)'),ylabel('percentage signal change'),
            grid on,grid minor, xlim([1 length(mF2g)]),ylim([1.25.*min([mF2g,mF2w,mF6]) 1.25.*max([mF2g,mF2w,mF6])]), title(['White matter mean signal AFTER "11-functional-anatomical-registration" (slice ' num2str(i3) ' of ' num2str(siz3) ')'],'Color',[0.65,0.11,0.19],'FontSize',14), legend({'White matter signal after co-registration'},'FontSize',12,'TextColor',[0.65,0.11,0.19],'Location','best')
        saveas(fighndl,[QCpath 'global_signal_08moco_vs_11coreg_slice' sprintf('%.2d',i3) '.jpg'])
        close(fighndl)
        
        C3 = corrcoef(F6'); C3=uppertriangle(C3);
        fighndl = figure('Position',[1 1 1152 720],'Color',[0.95,0.9,0.85],'InvertHardcopy','off','Visible','off'); % red [0.95,0.9,0.85], green [0.85,0.90,0.85], blue [0.89,0.94,0.99]
        subplot2n(3,1,1), histogram(C3,'EdgeColor','none','FaceColor','k','FaceAlpha',0.5,'Normalization','probability'); grid on,grid minor, xlim([-1 1]), hold on, xline(0,'Color','k','linewidth',1.5); hold off
            title(sprintf('Histogram of voxel-to-voxel correlations AFTER 08-motion-correction (slice %d of %d)\n%.1f%% of the correlations were > 0  [ mean=%.2f, median=%.2f, SD=%.2f ]',i3,siz3,100*length(find(C3>0))/length(C3),mean(C3),median(C3),std(C3)),'Color','k','FontSize',14)
        subplot2n(3,1,2), histogram(Cg,'EdgeColor','none','FaceColor',[0,0.55,0.69],'FaceAlpha',0.5,'Normalization','probability'); grid on,grid minor, xlim([-1 1]), hold on, xline(0,'Color','k','linewidth',1.5); hold off
            title(sprintf('Histogram of voxel-to-voxel GRAY MATTER correlations AFTER 11-registration (slice %d of %d)\n%.1f%% of the correlations were > 0  [ mean=%.2f, median=%.2f, SD=%.2f ]',i3,siz3,100*length(find(Cg>0))/length(Cg),mean(Cg),median(Cg),std(Cg)),'Color',[0,0.55,0.69],'FontSize',14)
        subplot2n(3,1,3), histogram(Cw,'EdgeColor','none','FaceColor',[0.65,0.11,0.19],'FaceAlpha',0.5,'Normalization','probability'); grid on,grid minor, xlim([-1 1]), hold on, xline(0,'Color','k','linewidth',1.5); hold off
            title(sprintf('Histogram of voxel-to-voxel WHITE MATTER correlations AFTER 11-registration (slice %d of %d)\n%.1f%% of the correlations were > 0  [ mean=%.2f, median=%.2f, SD=%.2f ]',i3,siz3,100*length(find(Cw>0))/length(Cw),mean(Cw),median(Cw),std(Cw)),'Color',[0.65,0.11,0.19],'FontSize',14)
        saveas(fighndl,[QCpath 'hist_FC_compare_08moco_11coreg_slice' sprintf('%.2d',i3) '.jpg'])
        close(fighndl)
        F6_=cat(1,F6_,F6); mF6_=cat(1,mF6_,mF6); C3_=cat(1,C3_,C3);
        clear F6 mF6 C3
    end
    
    F2g_=cat(1,F2g_,F2g); mF2g_=cat(1,mF2g_,mF2g); F2w_=cat(1,F2w_,F2w); mF2w_=cat(1,mF2w_,mF2w); Cg_=cat(1,Cg_,Cg); Cw_=cat(1,Cw_,Cw); F4_=cat(1,F4_,F4); mF4_=cat(1,mF4_,mF4); C2_=cat(1,C2_,C2);
    clear F2g F2w F4 TR Smask1 svar1 svar2 mF2g mF2w mF4 Cg Cw C2
end


[~,svar1] = sort(var(F2g_,0,2)); [~,spsc1] = sort(meanPSC(F2g_));
[~,svar2] = sort(var(F2w_,0,2)); [~,spsc2] = sort(meanPSC(F2w_));
sortorder1b = sortorder1; sortorder1=[]; f1=0; %#ok<*ASGLU>
sortorder2b = sortorder2; sortorder2=[]; f2=0; %#ok<*ASGLU>
for im=1:length(sortorder1b)
    temp = sortorder1b{im}; temp = temp + f1; f1 = f1 + length(temp);
    sortorder1 = cat(1,sortorder1,temp); clear temp
    temp = sortorder2b{im}; temp = temp + f2; f2 = f2 + length(temp);
    sortorder2 = cat(1,sortorder2,temp); clear temp
end; clear im f1 f2
sortorder1 = spsc1; % options: spsc for percentage signal change, svar for variance, (comment out the line if you want to use distance from the center of the cord separately for each slice)
sortorder2 = spsc2; % options: spsc for percentage signal change, svar for variance, (comment out the line if you want to use distance from the center of the cord separately for each slice)
for im=max(sortorder1):-1:1, if isempty(find(sortorder1==im)), sortorder1(find(sortorder1>im)) = sortorder1(find(sortorder1>im))-1; end, end; clear im %#ok<*EFIND>
for im=max(sortorder2):-1:1, if isempty(find(sortorder2==im)), sortorder2(find(sortorder2>im)) = sortorder2(find(sortorder2>im))-1; end, end; clear im %#ok<*EFIND>

fighndl = figure('Position',[1 1 1152 720],'Color',[0.89,0.94,0.99],'InvertHardcopy','off','Visible','off'); % red [0.95,0.9,0.85], green [0.85,0.90,0.85], blue [0.89,0.94,0.99]
subplot2n(2,1,1), imagesc(F2g_(sortorder1,:)),colormap(cmap),colorbar, xlabel('time (in samples)'),ylabel(sprintf('voxels (sorted by mean percentage signal change;\nbottom row is highest)')),
  title(['Time series within the GRAY MATTER AFTER 11-func-anat-registration (all slices combined)'],'Color',[0,0.55,0.69],'FontSize',14);
subplot2n(2,1,2), imagesc(F2w_(sortorder2,:)),colormap(cmap),colorbar, xlabel('time (in samples)'),ylabel(sprintf('voxels (sorted by mean percentage signal change;\nbottom row is highest)')),
  title(['Time series within the WHITE MATTER AFTER 11-func-anat-registration (all slices combined)'],'Color',[0.65,0.11,0.19],'FontSize',14);
saveas(fighndl,[QCpath 'fMRI_ts_11coreg_allSlices' '.jpg'])
saveas(fighndl,[QCpath2 'fMRI_ts_11coreg_allSlices' '.jpg'])
 close(fighndl)
mF2g_=mean(mF2g_,1); mF2w_=mean(mF2w_,1); mF4_=mean(mF4_,1);
fighndl = figure('Position',[1 1 1152 720],'Color',[0.89,0.94,0.99],'InvertHardcopy','off','Visible','off'); % red [0.95,0.9,0.85], green [0.85,0.90,0.85], blue [0.89,0.94,0.99]
subplot2n(3,1,1), plot(mF4_,'color','k','linewidth',1.5), hold on, plot(mF2g_,'color',[0,0.55,0.69],'linewidth',0.2), plot(mF2w_,'color',[0.65,0.11,0.19],'linewidth',0.2), xlabel('time (in samples)'),ylabel('percentage signal change'),
    grid on,grid minor, xlim([1 length(mF2g_)]),ylim([1.25.*min([mF2g_,mF2w_,mF4_]) 1.25.*max([mF2g_,mF2w_,mF4_])]), title(['Global mean signal in UNPROCESSED RAW DATA (all slices averaged)'],'Color','k','FontSize',14), legend({'Global signal (raw data)'},'FontSize',12,'TextColor','k','Location','best')
subplot2n(3,1,2), plot(mF2g_,'color',[0,0.55,0.69],'linewidth',1.5), hold on, plot(mF4_,'color','k','linewidth',0.2), xlabel('time (in samples)'),ylabel('percentage signal change'),
    grid on,grid minor, xlim([1 length(mF2g_)]),ylim([1.25.*min([mF2g_,mF2w_,mF4_]) 1.25.*max([mF2g_,mF2w_,mF4_])]), title(['Gray matter mean signal AFTER "11-functional-anatomical-registration" (all slices averaged)'],'Color',[0,0.55,0.69],'FontSize',14), legend({'Gray matter signal after 11-registration'},'FontSize',12,'TextColor',[0,0.55,0.69],'Location','best')
subplot2n(3,1,3), plot(mF2w_,'color',[0.65,0.11,0.19],'linewidth',1.5), hold on, plot(mF4_,'color','k','linewidth',0.2), xlabel('time (in samples)'),ylabel('percentage signal change'),
    grid on,grid minor, xlim([1 length(mF2g_)]),ylim([1.25.*min([mF2g_,mF2w_,mF4_]) 1.25.*max([mF2g_,mF2w_,mF4_])]), title(['White matter mean signal AFTER "11-functional-anatomical-registration" (all slices averaged)'],'Color',[0.65,0.11,0.19],'FontSize',14), legend({'White matter signal after 11-registration'},'FontSize',12,'TextColor',[0.65,0.11,0.19],'Location','best')
saveas(fighndl,[QCpath 'global_signal_11coreg_allSlices' '.jpg'])
 close(fighndl)
fighndl = figure('Position',[1 1 1152 720],'Color',[0.89,0.94,0.99],'InvertHardcopy','off','Visible','off'); % red [0.95,0.9,0.85], green [0.85,0.90,0.85], blue [0.89,0.94,0.99]
subplot2n(3,1,1), histogram(C2_,'EdgeColor','none','FaceColor','k','FaceAlpha',0.5,'Normalization','probability'); grid on,grid minor, xlim([-1 1]), hold on, xline(0,'Color','k','linewidth',1.5); hold off
    title(sprintf('Histogram of voxel-to-voxel correlations in UNPROCESSED RAW DATA (all slices combined)\n%.1f%% of the correlations were > 0  [ mean=%.2f, median=%.2f, SD=%.2f ]',100*length(find(C2_>0))/length(C2_),mean(C2_),median(C2_),std(C2_)),'Color','k','FontSize',14)
subplot2n(3,1,2), histogram(Cg_,'EdgeColor','none','FaceColor',[0,0.55,0.69],'FaceAlpha',0.5,'Normalization','probability'); grid on,grid minor, xlim([-1 1]), hold on, xline(0,'Color','k','linewidth',1.5); hold off
    title(sprintf('Histogram of voxel-to-voxel GRAY MATTER correlations AFTER 11-registration (all slices combined)\n%.1f%% of the correlations were > 0  [ mean=%.2f, median=%.2f, SD=%.2f ]',100*length(find(Cg_>0))/length(Cg_),mean(Cg_),median(Cg_),std(Cg_)),'Color',[0,0.55,0.69],'FontSize',14)
subplot2n(3,1,3), histogram(Cw_,'EdgeColor','none','FaceColor',[0.65,0.11,0.19],'FaceAlpha',0.5,'Normalization','probability'); grid on,grid minor, xlim([-1 1]), hold on, xline(0,'Color','k','linewidth',1.5); hold off
    title(sprintf('Histogram of voxel-to-voxel WHITE MATTER correlations AFTER 11-registration (all slices combined)\n%.1f%% of the correlations were > 0  [ mean=%.2f, median=%.2f, SD=%.2f ]',100*length(find(Cw_>0))/length(Cw_),mean(Cw_),median(Cw_),std(Cw_)),'Color',[0.65,0.11,0.19],'FontSize',14)
saveas(fighndl,[QCpath 'hist_FC_compare_11coreg_allSlices' '.jpg'])
 close(fighndl)
if exist([base_dir_sub fname '_slice' num2str(i3) '_MC.nii'],'file') || exist([base_dir_sub fname '_slice' num2str(i3) '_MC.nii.gz'],'file') || exist([base_dir_sub fname '_MC.nii'],'file') || exist([base_dir_sub fname '_MC.nii.gz'],'file')
    mF6_=mean(mF6_,1);
    fighndl = figure('Position',[1 1 1152 720],'Color',[0.89,0.94,0.99],'InvertHardcopy','off','Visible','off'); % red [0.95,0.9,0.85], green [0.85,0.90,0.85], blue [0.89,0.94,0.99]
    subplot2n(3,1,1), plot(mF6_,'color','k','linewidth',1.5), hold on, plot(mF2g_,'color',[0,0.55,0.69],'linewidth',0.2), plot(mF2w_,'color',[0.65,0.11,0.19],'linewidth',0.2), xlabel('time (in samples)'),ylabel('percentage signal change'),
        grid on,grid minor, xlim([1 length(mF2g_)]),ylim([1.25.*min([mF2g_,mF2w_,mF6_]) 1.25.*max([mF2g_,mF2w_,mF6_])]), title(['Global mean signal AFTER "08-motion-correction" (all slices averaged)'],'Color','k','FontSize',14), legend({'Global signal before co-registration'},'FontSize',12,'TextColor','k','Location','best')
    subplot2n(3,1,2), plot(mF2g_,'color',[0,0.55,0.69],'linewidth',1.5), hold on, plot(mF6_,'color','k','linewidth',0.2), xlabel('time (in samples)'),ylabel('percentage signal change'),
        grid on,grid minor, xlim([1 length(mF2g_)]),ylim([1.25.*min([mF2g_,mF2w_,mF6_]) 1.25.*max([mF2g_,mF2w_,mF6_])]), title(['Gray matter mean signal AFTER "11-functional-anatomical-registration" (all slices averaged)'],'Color',[0,0.55,0.69],'FontSize',14), legend({'Gray matter signal after co-registration'},'FontSize',12,'TextColor',[0,0.55,0.69],'Location','best')
    subplot2n(3,1,3), plot(mF2w_,'color',[0.65,0.11,0.19],'linewidth',1.5), hold on, plot(mF6_,'color','k','linewidth',0.2), xlabel('time (in samples)'),ylabel('percentage signal change'),
        grid on,grid minor, xlim([1 length(mF2g_)]),ylim([1.25.*min([mF2g_,mF2w_,mF6_]) 1.25.*max([mF2g_,mF2w_,mF6_])]), title(['White matter mean signal AFTER "11-functional-anatomical-registration" (all slices averaged)'],'Color',[0.65,0.11,0.19],'FontSize',14), legend({'White matter signal after co-registration'},'FontSize',12,'TextColor',[0.65,0.11,0.19],'Location','best')
    saveas(fighndl,[QCpath 'global_signal_08moco_vs_11coreg_allSlices' '.jpg'])
    saveas(fighndl,[QCpath2 'global_signal_08moco_vs_11coreg_allSlices' '.jpg'])
     close(fighndl)
     fighndl = figure('Position',[1 1 1152 720],'Color',[0.89,0.94,0.99],'InvertHardcopy','off','Visible','off'); % red [0.95,0.9,0.85], green [0.85,0.90,0.85], blue [0.89,0.94,0.99]
    subplot2n(3,1,1), histogram(C3_,'EdgeColor','none','FaceColor','k','FaceAlpha',0.5,'Normalization','probability'); grid on,grid minor, xlim([-1 1]), hold on, xline(0,'Color','k','linewidth',1.5); hold off
       title(sprintf('Histogram of voxel-to-voxel correlations AFTER 08-motion-correction (all slices combined)\n%.1f%% of the correlations were > 0  [ mean=%.2f, median=%.2f, SD=%.2f ]',100*length(find(C3_>0))/length(C3_),mean(C3_),median(C3_),std(C3_)),'Color','k','FontSize',14)
    subplot2n(3,1,2), histogram(Cg_,'EdgeColor','none','FaceColor',[0,0.55,0.69],'FaceAlpha',0.5,'Normalization','probability'); grid on,grid minor, xlim([-1 1]), hold on, xline(0,'Color','k','linewidth',1.5); hold off
       title(sprintf('Histogram of voxel-to-voxel GRAY MATTER correlations AFTER 11-registration (all slices combined)\n%.1f%% of the correlations were > 0  [ mean=%.2f, median=%.2f, SD=%.2f ]',100*length(find(Cg_>0))/length(Cg_),mean(Cg_),median(Cg_),std(Cg_)),'Color',[0,0.55,0.69],'FontSize',14)
    subplot2n(3,1,3), histogram(Cw_,'EdgeColor','none','FaceColor',[0.65,0.11,0.19],'FaceAlpha',0.5,'Normalization','probability'); grid on,grid minor, xlim([-1 1]), hold on, xline(0,'Color','k','linewidth',1.5); hold off
       title(sprintf('Histogram of voxel-to-voxel WHITE MATTER correlations AFTER 11-registration (all slices combined)\n%.1f%% of the correlations were > 0  [ mean=%.2f, median=%.2f, SD=%.2f ]',100*length(find(Cw_>0))/length(Cw_),mean(Cw_),median(Cw_),std(Cw_)),'Color',[0.65,0.11,0.19],'FontSize',14)
    saveas(fighndl,[QCpath 'hist_FC_compare_08moco_11coreg_allSlices' '.jpg'])
    saveas(fighndl,[QCpath2 'hist_FC_compare_08moco_11coreg_allSlices' '.jpg'])
     close(fighndl)
end
clear svar1 spsc1 svar2 spsc2 sortorder1 sortorder2 F2g_ mF2g_ F2w_ mF2w_ Cg_ Cw_ F4_ mF4_ C2_ F6_ mF6_ C3_ cmap

%% Make QC video (needs audio video toolbox in Matlab)
if ~isempty(which('VideoWriter')) && (save_videos==1) % check if Matlab's audio video toolbox exists

subplot = @(m,n,p) subtightplot(m, n, p, [0.001 0.005], [0.01 0.001], [0.01 0.001]);
scfMRItb_04_unzipFile(base_dir_sub, fname_anat, '_mask_SC')
mask_SC2 = load_untouch_nii([base_dir_sub fname_anat '_mask_SC.nii']); mask_SC2 = mask_SC2.img;
for M=1:size(mask_SC2,3)
    mask_SC = rot90(mask_SC2(:,:,M));
    [xmask_SC,ymask_SC] = find(mask_SC==1);
    minx_SC(:,M) = round(min(xmask_SC)-0.1*(size(mask_SC,1))); maxx_SC(:,M)=round(max(xmask_SC)+0.1*(size(mask_SC,1)));
    miny_SC(:,M) = round(min(ymask_SC)-0.1*(size(mask_SC,2))); maxy_SC(:,M)=round(max(ymask_SC)+0.1*(size(mask_SC,2)));
    if (maxx_SC(:,M)-minx_SC(:,M))<(maxy_SC(:,M)-miny_SC(:,M))
        factr = ((maxy_SC(:,M)-miny_SC(:,M))-(maxx_SC(:,M)-minx_SC(:,M)))/2;
        if factr==round(factr)
            maxx_SC(:,M)=maxx_SC(:,M)+factr; minx_SC(:,M)=minx_SC(:,M)-factr;
        else
            maxx_SC(:,M)=maxx_SC(:,M)+factr+0.5; minx_SC(:,M)=minx_SC(:,M)-factr+0.5;
        end
    elseif (maxy_SC(:,M)-miny_SC(:,M))<(maxx_SC(:,M)-minx_SC(:,M))
        factr = ((maxx_SC(:,M)-minx_SC(:,M))-(maxy_SC(:,M)-miny_SC(:,M)))/2;
        if factr==round(factr)
            maxy_SC(:,M)=maxy_SC(:,M)+factr; miny_SC(:,M)=miny_SC(:,M)-factr;
        else
            maxy_SC(:,M)=maxy_SC(:,M)+factr+0.5; miny_SC(:,M)=miny_SC(:,M)-factr+0.5;
        end
    end
    clear mask_SC xmask_SC ymask_SC factr
end; clear M

for i3 = 1 : siz3
    try waitbar((i3/siz3),wbar3,sprintf('11. Generate QC video: slice (%d) of (%d)',i3,siz3)); catch, end
    fprintf('11. Generate QC video: slice (%d) of (%d)\n',i3,siz3)
    scfMRItb_04_resplitData(base_dir_sub, fname, '_warped', siz3) % if slice-wise data doesn't exist but combined 4D data exists, then split the data to generate the needed slice-wise data
    F1 = load_untouch_nii([base_dir_sub fname '_slice' num2str(i3) '_warped.nii']);
    TR = F1.hdr.dime.pixdim(5); F1 = rot90(single(squeeze(F1.img)));
    video = VideoWriter([QCpath 'fMRIvideo_11coreg_slice' sprintf('%.2d',i3) '.mp4'],'MPEG-4'); %#ok<*TNMLP> % create the video object
    video.FrameRate = round(30/TR); % 30 seconds of data in every second of the video
    video.Quality = 90; % very limited compression so that the images are shown as is (and thus avoid misunderstandings due to video quality)
    open(video); % open the file for writing
    
    [sizeFx_SC,sizeFy_SC] = size(F1(:,:,1));
    F2 = F1(minx_SC:maxx_SC,miny_SC:maxy_SC,:);
    for i5=1:size(F1,3)
        F1(:,:,i5) = imresize(F2(:,:,i5),[sizeFx_SC sizeFy_SC]);
    end
    
    if Smask_err==0
        scfMRItb_04_resplitData(base_dir_sub, fname, '_Smask', siz3) % if slice-wise data doesn't exist but combined 4D data exists, then split the data to generate the needed slice-wise data
        scfMRItb_04_unzipFile(base_dir_sub, fname, ['_slice' num2str(i3) '_Smask'])
        Smask1 = load_untouch_nii([base_dir_sub fname '_slice' num2str(i3) '_Smask.nii']); Smask1 = Smask1.img;
    else
        Smask1 = Smask1_(:,:,i3);
    end
    Smask1 = rot90(Smask1);
    [xmask,ymask] = find(Smask1==1);
    minx=round(min(xmask)-0.1*(size(Smask1,1))); maxx=round(max(xmask)+0.1*(size(Smask1,1)));
    miny=round(min(ymask)-0.1*(size(Smask1,2))); maxy=round(max(ymask)+0.1*(size(Smask1,2)));
    if (maxx-minx)<(maxy-miny)
        factr = ((maxy-miny)-(maxx-minx))/2;
        if factr==round(factr)
            maxx=maxx+factr; minx=minx-factr;
        else
            maxx=maxx+factr+0.5; minx=minx-factr+0.5;
        end
    elseif (maxy-miny)<(maxx-minx)
        factr = ((maxx-minx)-(maxy-miny))/2;
        if factr==round(factr)
            maxy=maxy+factr; miny=miny-factr;
        else
            maxy=maxy+factr+0.5; miny=miny-factr+0.5;
        end
    end
    scfMRItb_04_resplitData(base_dir_sub, fname, '', siz3) % if slice-wise data doesn't exist but combined 4D data exists, then split the data to generate the needed slice-wise data
    F3 = load_untouch_nii([base_dir_sub fname '_slice' num2str(i3) '.nii']); F3 = rot90(single(squeeze(F3.img)));
    [sizeFx,sizeFy] = size(F3(:,:,1));
    F4 = F3(minx:maxx,miny:maxy,:);
    for i5=1:size(F3,3)
        F3(:,:,i5) = imresize(F4(:,:,i5),[sizeFx sizeFy]);
    end
    clear maxx maxy minx miny xmask ymask Smask1 factr sizeFx sizeFy F2 F4 i5 sizeFx_SC sizeFy_SC
    
    for ii=1:size(F1,3) % loop across the number of images
        I = F1(:,:,ii); %read the next image
        I(find(isnan(I)))=0;
        I = (I - min(I(:))) ./ (max(I(:)) - min(I(:)));
        I2 = F3(:,:,ii); %read the next image
        I2(find(isnan(I2)))=0;
        I2 = (I2 - min(I2(:))) ./ (max(I2(:)) - min(I2(:)));
        fighndl = figure('Position',[round((scrsz(3)-scrsz(4))/2) 1 round(0.75*scrsz(3)) round(0.75*scrsz(4))],'Color',[0.95,0.9,0.85],'InvertHardcopy','off','Visible','off'); % [0.85,0.90,0.85]
        subplot(1,2,1), imshow(I2, [0 max(I2(:))], 'InitialMagnification', 'fit'); title({['Unprocessed raw fMRI data (slice ' num2str(i3) ' of ' num2str(siz3) ')'];['Time point ' num2str(ii) ' of ' num2str(size(F1,3))]},'Color','k','FontSize',18);
        subplot(1,2,2), imshow(I, [0 max(I(:))], 'InitialMagnification', 'fit'); title({['fMRI data after "11. func-anat-registration" (slice ' num2str(i3) ' of ' num2str(siz3) ')'];['Time point ' num2str(ii) ' of ' num2str(size(F1,3))]},'Color',[0,0.55,0.69],'FontSize',18);
        frame = getframe(gcf);
        writeVideo(video,frame); % write the image to file
        close(fighndl)
        clear I I2 frame
    end
    close(video); %close the file
    clear F1 F3 video ii TR
end; clear i3
clear maxx_SC maxy_SC minx_SC miny_SC F

end

try
if nargin<5
    close(wbar3)
end
catch
end

end
```
