## Supplementary material for "Neptune: a toolbox for spinal cord functional MRI data processing and quality assurance": Neptune software: scfMRItb_13_WMregress.m

```
function scfMRItb_13_WMregress(base_dir_sub,fname,fname_anat_orig, varargin)

fname_anat = [fname_anat_orig '_reduced'];
cutoff = 0.50; cutoff_str = '50';

if nargin<4
    wbar3 = waitbar(0,'13. Regress-out white-matter signal...','Name','Progress(13): Regress-out white-matter signal...','Units','normalized','Position',[3/5 0 1/5 1/17]);
else
    wbar3 = varargin{1};
end

if nargin<5
    save_videos = 1; % 1 = save QC videos (generates sizeable .mp4 files and takes a few minutes to do it). This is turned on by default, but if you don't want to save videos then set it to 0
else
    save_videos = varargin{2};
end
if nargin<6
    Rns = 0; % 0 = don't create runX subfolder; X = create runX subfolder
else
    Rns = varargin{3};
end

if ~(exist([base_dir_sub 'QC'],'dir'))
    mkdir([base_dir_sub 'QC'])
end
if ~(exist([base_dir_sub 'QC' '/13_denoise4-WMregress'],'dir'))
    mkdir([base_dir_sub 'QC' '/13_denoise4-WMregress'])
end
if Rns==0
    QCpath = [base_dir_sub 'QC' '/13_denoise4-WMregress/'];
else
    if ~(exist([base_dir_sub 'QC' '/13_denoise4-WMregress' '/run' num2str(Rns)],'dir'))
        mkdir([base_dir_sub 'QC' '/13_denoise4-WMregress' '/run' num2str(Rns)])
    end
    QCpath = [base_dir_sub 'QC' '/13_denoise4-WMregress' '/run' num2str(Rns) '/'];
end

if ~(exist([base_dir_sub 'QC' '/QC_shortlist'],'dir'))
    mkdir([base_dir_sub 'QC' '/QC_shortlist'])
end
if Rns==0
    QCpath2 = [base_dir_sub 'QC' '/QC_shortlist/'];
else
    if ~(exist([base_dir_sub 'QC' '/QC_shortlist' '/run' num2str(Rns)],'dir'))
        mkdir([base_dir_sub 'QC' '/QC_shortlist' '/run' num2str(Rns)])
    end
    QCpath2 = [base_dir_sub 'QC' '/QC_shortlist' '/run' num2str(Rns) '/'];
end

%% ----- REMOVE "GLOBAL" WM SIGNAL FROM EACH SLICE -------------------------

scfMRItb_04_unzipFile(base_dir_sub, fname_anat, '')
A = load_untouch_nii([base_dir_sub fname_anat '.nii']);
siz3 = size(A.img,3);

scfMRItb_04_unzipFile(base_dir_sub, fname_anat, '_mask_WM')
scfMRItb_04_unzipFile(base_dir_sub, fname_anat, '_mask_GM')
mask_WM = load_untouch_nii([base_dir_sub fname_anat '_mask_WM.nii']); mask_WM = mask_WM.img;
mask_GM = load_untouch_nii([base_dir_sub fname_anat '_mask_GM.nii']); mask_GM = mask_GM.img;
mask_any = single(mask_WM | mask_GM);

for i3 = 1 : siz3
    try waitbar((i3/siz3)/3,wbar3,sprintf('13. Regress-out WM signal: slice (%d) of (%d)',i3,siz3)); catch, end
    disp(sprintf('Regress-out white-matter signal: slice (%d) of (%d) -> subject %s',i3,siz3,fname)) %#ok<DSPS>
    flag=0;
    
    try % use denoised data ('denoise2') if that step was performed
        scfMRItb_04_resplitData(base_dir_sub, fname, ['_denoised' cutoff_str], siz3) % if slice-wise data doesn't exist but combined 4D data exists, then split the data to generate the needed slice-wise data
        scfMRItb_04_unzipFile(base_dir_sub, fname, ['_slice' num2str(i3) '_denoised' cutoff_str])
        F = load_untouch_nii([base_dir_sub fname '_slice' num2str(i3) '_denoised' cutoff_str '.nii']);
    catch % else use co-registered functional data
        scfMRItb_04_resplitData(base_dir_sub, fname, '_warped', siz3) % if slice-wise data doesn't exist but combined 4D data exists, then split the data to generate the needed slice-wise data
        scfMRItb_04_unzipFile(base_dir_sub, fname, ['_slice' num2str(i3) '_warped'])
        F = load_untouch_nii([base_dir_sub fname '_slice' num2str(i3) '_warped.nii']);
        flag=1;
    end
    F_denoised = F;
    F = single(squeeze(F.img));
    
    % only white matter mask
    mmask = single(mask_WM);
    B = F .* repmat(mmask(:,:,i3), [1 1 size(F,3)]);
    
    % use both white and gray matter masks?
%    mmask = mask_any;
%    B = F .* repmat(mmask(:,:,i3), [1 1 size(F,3)]);
    
clear mmask;
    Bv = zeros(sum(sum(any(B,3))), size(B,3), 'single');
    i4 = 0;
    for i1 = 1 : size(B,1)
        for i2 = 1 : size(B,2)
            if any(B(i1,i2,:),3)
                i4 = i4 + 1;
                Bv(i4,:) = reshape(B(i1,i2,:), [1 size(B,3)]);
            end
        end
    end
    clear B;
    Bv = Bv'; % N-by-P matrix:  N = rows = time points (observations)
              %   (N << P)      P = columns = voxels (variables)
              
    [~, B_score, B_latent] = pca(Bv);
    B_percent = B_latent ./ sum(B_latent);
    B_percum = B_percent;
    for i1 = 2 : size(B_percum,1)
        B_percum(i1,1) = B_percum(i1,1) + B_percum(i1-1,1);
    end
    B_percum(1,1)
    B_sscore = B_score(:,1);

    % actually do the regression now!
    for i1 = 1 : size(F,1)
        for i2 = 1 : size(F,2)
%             if mask_any(i1,i2,i3)
            if mask_GM(i1,i2,i3)
                temp = squeeze(F(i1,i2,:));              % original
                temp2 = B_sscore * lscov(B_sscore,temp); % noise regressor
                F(i1,i2,:) = reshape((temp - temp2), [1 1 size(F,3)]);
                clear temp temp2;
            end
        end
    end
    
    F_denoised.img = reshape(F, [size(F,1) size(F,2) 1 size(F,3)]);
    
    if flag==0 % if denoised data ('denoise2') was used
        unix(['rm -f ' base_dir_sub fname '_slice' num2str(i3) '_denoised' cutoff_str 'WM.nii']);
        save_untouch_nii(F_denoised, [base_dir_sub fname '_slice' num2str(i3) '_denoised' cutoff_str 'WM.nii']);
    elseif flag==1 % if co-registered pre-denoised data was used
        unix(['rm -f ' base_dir_sub fname '_slice' num2str(i3) '_WM.nii']);
        save_untouch_nii(F_denoised, [base_dir_sub fname '_slice' num2str(i3) '_WM.nii']);
    end
    clear F F_denoised B_*;
end
clear mask_WM mask_GM;

%% time series plots, global signal and FC histogram
scrsz = get(0,'ScreenSize'); pause on;
cmap = colormap('bone'); cmap = (1-exp(-cmap))./(1-exp(-1)); 

scfMRItb_04_unzipFile(base_dir_sub, fname_anat, '_mask_GM')
scfMRItb_04_unzipFile(base_dir_sub, fname_anat, '_mask_WM')
mask_GM = load_untouch_nii([base_dir_sub fname_anat '_mask_GM.nii']); mask_GM = mask_GM.img;
mask_WM = load_untouch_nii([base_dir_sub fname_anat '_mask_WM.nii']); mask_WM = mask_WM.img;

if exist([base_dir_sub fname '_slice' num2str(i3) '_denoised' cutoff_str '.nii'],'file') || exist([base_dir_sub fname '_slice' num2str(i3) '_denoised' cutoff_str '.nii.gz'],'file') || exist([base_dir_sub fname '_denoised' cutoff_str '.nii'],'file') || exist([base_dir_sub fname '_denoised' cutoff_str '.nii.gz'],'file')
    stp=2;
elseif exist([base_dir_sub fname '_slice' num2str(i3) '_warped.nii'],'file') || exist([base_dir_sub fname '_slice' num2str(i3) '_warped.nii.gz'],'file') || exist([base_dir_sub fname '_warped.nii'],'file') || exist([base_dir_sub fname '_warped.nii.gz'],'file')
    stp=1;
end

Smask_err=0;
try
    scfMRItb_04_resplitData(base_dir_sub, fname, '_Smask', siz3) % if slice-wise data doesn't exist but combined 4D data exists, then split the data to generate the needed slice-wise data
catch
    load([base_dir_sub fname '_mask_NS' '.mat'])
    Smask1_ = ~mask_NS.img; clear mask_NS
    Smask_err=1;
end

F2g_=[]; mF2g_=[]; F2w_=[]; mF2w_=[]; Cg_=[]; Cw_=[]; F4_=[]; mF4_=[]; C2_=[];
if stp~=0, F6g_=[]; mF6g_=[]; F6w_=[]; mF6w_=[]; C3g_=[]; C3w_=[]; end
for i3 = 1 : siz3
    try waitbar((i3/siz3),wbar3,sprintf('13. QC plots: slice (%d) of (%d)',i3,siz3)); catch, end
    fprintf('13. QC plots: slice (%d) of (%d)\n',i3,siz3)
    if stp==2
        scfMRItb_04_resplitData(base_dir_sub, fname, ['_denoised' cutoff_str 'WM'], siz3) % if slice-wise data doesn't exist but combined 4D data exists, then split the data to generate the needed slice-wise data
        scfMRItb_04_unzipFile(base_dir_sub, fname, ['_slice' num2str(i3) '_denoised' cutoff_str 'WM'])
        F1 = load_untouch_nii([base_dir_sub fname '_slice' num2str(i3) '_denoised' cutoff_str 'WM.nii']);
    elseif stp==1
        scfMRItb_04_resplitData(base_dir_sub, fname, '_WM', siz3) % if slice-wise data doesn't exist but combined 4D data exists, then split the data to generate the needed slice-wise data
        scfMRItb_04_unzipFile(base_dir_sub, fname, ['_slice' num2str(i3) '_WM'])
        F1 = load_untouch_nii([base_dir_sub fname '_slice' num2str(i3) '_WM.nii']);
    end
    TR = F1.hdr.dime.pixdim(5); F1 = single(squeeze(F1.img));
    
    if Smask_err==0
        scfMRItb_04_resplitData(base_dir_sub, fname, '_Smask', siz3) % if slice-wise data doesn't exist but combined 4D data exists, then split the data to generate the needed slice-wise data
        scfMRItb_04_unzipFile(base_dir_sub, fname, ['_slice' num2str(i3) '_Smask'])
        Smask1 = load_untouch_nii([base_dir_sub fname '_slice' num2str(i3) '_Smask.nii']); Smask1 = Smask1.img;
    else
        Smask1 = Smask1_(:,:,i3);
    end

    for i5=1:size(F1,3)
        tmp = F1(:,:,i5);
        F2g(:,i5) = tmp(find(mask_GM(:,:,i3)==1));
        F2w(:,i5) = tmp(find(mask_WM(:,:,i3)==1));
        clear tmp
    end; clear i5 F1
    [~,svar1] = sort(var(F2g,0,2)); [~,sdist1] = sort(svar1); sortorder1{i3,1} = sdist1;
    [~,svar2] = sort(var(F2w,0,2)); [~,sdist2] = sort(svar2); sortorder2{i3,1} = sdist2;
    
    fighndl = figure('Position',[1 1 1152 720],'Color',[0.95,0.9,0.85],'InvertHardcopy','off','Visible','off'); % red [0.95,0.9,0.85], green [0.85,0.90,0.85], blue [0.89,0.94,0.99]
    subplot2n(2,1,1), imagesc(F2g(sortorder1{i3},:)),colormap(cmap),colorbar, xlabel('time (in samples)'),ylabel('Gray matter voxels (sorted by variance)'),
        title(['Time series within the GRAY MATTER after "13-DENOISE4-WMregress" in slice ' num2str(i3) ' of ' num2str(siz3)],'Color',[0,0.55,0.69],'FontSize',14);
    subplot2n(2,1,2), imagesc(F2w(sortorder2{i3},:)),colormap(cmap),colorbar, xlabel('time (in samples)'),ylabel('White matter voxels (sorted by variance)'),
        title(['Time series within the WHITE MATTER after "13-DENOISE4-WMregress" in slice ' num2str(i3) ' of ' num2str(siz3)],'Color',[0.65,0.11,0.19],'FontSize',14);
    saveas(fighndl,[QCpath 'fMRI_ts_13denoise4_slice' sprintf('%.2d',i3) '.jpg'])
    close(fighndl)
    
    scfMRItb_04_resplitData(base_dir_sub, fname, '', siz3) % if slice-wise data doesn't exist but combined 4D data exists, then split the data to generate the needed slice-wise data
	scfMRItb_04_unzipFile(base_dir_sub, fname, ['_slice' num2str(i3)])
    F3 = load_untouch_nii([base_dir_sub fname '_slice' num2str(i3) '.nii']); F3 = single(squeeze(F3.img));
    for i5=1:size(F3,3)
        tmp = F3(:,:,i5);
        F4(:,i5) = tmp(find(Smask1==1)); clear tmp
    end; clear i5 F3
    mF2g=mean(F2g,1); mF2g = 100*(mF2g-mean(mF2g))./mean(mF2g); mF2w=mean(F2w,1); mF2w = 100*(mF2w-mean(mF2w))./mean(mF2w);
    mF4=mean(F4,1); mF4 = 100*(mF4-mean(mF4))./mean(mF4);
    fighndl = figure('Position',[1 1 1152 720],'Color',[0.95,0.9,0.85],'InvertHardcopy','off','Visible','off'); % red [0.95,0.9,0.85], green [0.85,0.90,0.85], blue [0.89,0.94,0.99]
    subplot2n(3,1,1), plot(mF4,'color','k','linewidth',1.5), hold on, plot(mF2g,'color',[0,0.55,0.69],'linewidth',0.2), plot(mF2w,'color',[0.65,0.11,0.19],'linewidth',0.2), xlabel('time (in samples)'),ylabel('percentage signal change'),
      grid on,grid minor, xlim([1 length(mF2g)]),ylim([min(1.25.*[mF2g,mF2w,mF4]) 1.25.*max([mF2g,mF2w,mF4])]), title(['Global mean signal in UNPROCESSED RAW DATA (slice ' num2str(i3) ' of ' num2str(siz3) ')'],'Color','k','FontSize',14), legend({'Global signal (raw data)'},'FontSize',12,'TextColor','k','Location','best')
    subplot2n(3,1,2), plot(mF2g,'color',[0,0.55,0.69],'linewidth',1.5), hold on, plot(mF2w,'color',[0.65,0.11,0.19],'linewidth',0.2), plot(mF4,'color','k','linewidth',0.2), xlabel('time (in samples)'),ylabel('percentage signal change'),
      grid on,grid minor, xlim([1 length(mF2g)]),ylim([min(1.25.*[mF2g,mF2w,mF4]) 1.25.*max([mF2g,mF2w,mF4])]), title(['Gray matter mean signal AFTER "13-DENOISE4-WMregress" (slice ' num2str(i3) ' of ' num2str(siz3) ')'],'Color',[0,0.55,0.69],'FontSize',14), legend({'Gray matter signal after 13-denoise4'},'FontSize',12,'TextColor',[0,0.55,0.69],'Location','best')
    subplot2n(3,1,3), plot(mF2w,'color',[0.65,0.11,0.19],'linewidth',1.5), hold on, plot(mF4,'color','k','linewidth',0.2), plot(mF2g,'color',[0,0.55,0.69],'linewidth',0.2), xlabel('time (in samples)'),ylabel('percentage signal change'),
      grid on,grid minor, xlim([1 length(mF2g)]),ylim([min(1.25.*[mF2g,mF2w,mF4]) 1.25.*max([mF2g,mF2w,mF4])]), title(['White matter mean signal AFTER "13-DENOISE4-WMregress" (slice ' num2str(i3) ' of ' num2str(siz3) ')'],'Color',[0.65,0.11,0.19],'FontSize',14), legend({'White matter signal after 13-denoise4'},'FontSize',12,'TextColor',[0.65,0.11,0.19],'Location','best')
    saveas(fighndl,[QCpath 'global_signal_13denoise4_slice' sprintf('%.2d',i3) '.jpg'])
    close(fighndl)

    Cg = corrcoef(F2g'); Cg = uppertriangle(Cg);
    Cw = corrcoef(F2w'); Cw = uppertriangle(Cw);
    C2 = corrcoef(F4');  C2 = uppertriangle(C2);
    fighndl = figure('Position',[1 1 1152 720],'Color',[0.95,0.9,0.85],'InvertHardcopy','off','Visible','off'); % red [0.95,0.9,0.85], green [0.85,0.90,0.85], blue [0.89,0.94,0.99]
    subplot2n(3,1,1), histogram(C2,'EdgeColor','none','FaceColor','k','FaceAlpha',0.5,'Normalization','probability'); grid on,grid minor, xlim([-1 1]), hold on, xline(0,'Color','k','linewidth',1.5); hold off
      title(sprintf('Histogram of voxel-to-voxel correlations in UNPROCESSED RAW DATA (slice %d of %d)\n%.1f%% of the correlations were > 0  [ mean=%.2f, median=%.2f, SD=%.2f ]',i3,siz3,100*length(find(C2>0))/length(C2),mean(C2),median(C2),std(C2)),'Color','k','FontSize',14)
    subplot2n(3,1,2), histogram(Cg,'EdgeColor','none','FaceColor',[0,0.55,0.69],'FaceAlpha',0.5,'Normalization','probability'); grid on,grid minor, xlim([-1 1]), hold on, xline(0,'Color','k','linewidth',1.5); hold off
      title(sprintf('Histogram of voxel-to-voxel GRAY MATTER correlations AFTER 13-DENOISE4-WMregress (slice %d of %d)\n%.1f%% of the correlations were > 0  [ mean=%.2f, median=%.2f, SD=%.2f ]',i3,siz3,100*length(find(Cg>0))/length(Cg),mean(Cg),median(Cg),std(Cg)),'Color',[0,0.55,0.69],'FontSize',14)
    subplot2n(3,1,3), histogram(Cw,'EdgeColor','none','FaceColor',[0.65,0.11,0.19],'FaceAlpha',0.5,'Normalization','probability'); grid on,grid minor, xlim([-1 1]), hold on, xline(0,'Color','k','linewidth',1.5); hold off
      title(sprintf('Histogram of voxel-to-voxel WHITE MATTER correlations AFTER 13-DENOISE4-WMregress (slice %d of %d)\n%.1f%% of the correlations were > 0  [ mean=%.2f, median=%.2f, SD=%.2f ]',i3,siz3,100*length(find(Cw>0))/length(Cw),mean(Cw),median(Cw),std(Cw)),'Color',[0.65,0.11,0.19],'FontSize',14)
    saveas(fighndl,[QCpath 'hist_FC_compare_13denoise4_slice' sprintf('%.2d',i3) '.jpg'])
    close(fighndl)
    
    if stp~=0
        if stp==1
            scfMRItb_04_resplitData(base_dir_sub, fname, '_warped', siz3) % if slice-wise data doesn't exist but combined 4D data exists, then split the data to generate the needed slice-wise data
            scfMRItb_04_unzipFile(base_dir_sub, fname, ['_slice' num2str(i3) '_warped'])
            F5 = load_untouch_nii([base_dir_sub fname '_slice' num2str(i3) '_warped.nii']); F5 = single(squeeze(F5.img));
        elseif stp==2
            scfMRItb_04_resplitData(base_dir_sub, fname, ['_denoised' cutoff_str], siz3) % if slice-wise data doesn't exist but combined 4D data exists, then split the data to generate the needed slice-wise data
            scfMRItb_04_unzipFile(base_dir_sub, fname, ['_slice' num2str(i3) '_denoised' cutoff_str ''])
            F5 = load_untouch_nii([base_dir_sub fname '_slice' num2str(i3) '_denoised' cutoff_str '.nii']); F5 = single(squeeze(F5.img));
        end
        for i5=1:size(F5,3)
            tmp = F5(:,:,i5);
            F6g(:,i5) = tmp(find(mask_GM(:,:,i3)==1));
            F6w(:,i5) = tmp(find(mask_WM(:,:,i3)==1));
            clear tmp
        end; clear i5 F5
        mF6g=mean(F6g,1); mF6g = 100*(mF6g-mean(mF6g))./mean(mF6g); mF6w=mean(F6w,1); mF6w = 100*(mF6w-mean(mF6w))./mean(mF6w);
        fighndl = figure('Position',[1 1 1152 720],'Color',[0.95,0.9,0.85],'InvertHardcopy','off','Visible','off'); % red [0.95,0.9,0.85], green [0.85,0.90,0.85], blue [0.89,0.94,0.99]
        if stp==1
            subplot2n(4,1,1), plot(mF6g,'color','k','linewidth',1.5), hold on, plot(mF2g,'color',[0,0.55,0.69],'linewidth',0.2), ylabel('pcent sig change'),
                grid on,grid minor, xlim([1 length(mF2g)]),ylim([min(1.25.*[mF2g,mF6g]) 1.25.*max([mF2g,mF6g])]), title(['Gray matter mean signal AFTER "11-func-anat-registration" (slice ' num2str(i3) ' of ' num2str(siz3) ')'],'Color','k','FontSize',14), legend({'Gray matter signal before WM-denoising'},'FontSize',12,'TextColor','k','Location','best')
            subplot2n(4,1,3), plot(mF6w,'color','k','linewidth',1.5), hold on, plot(mF2w,'color',[0.65,0.11,0.19],'linewidth',0.2), ylabel('pcent sig change'),
                grid on,grid minor, xlim([1 length(mF2g)]),ylim([min(1.25.*[mF2w,mF6w]) 1.25.*max([mF2w,mF6w])]), title(['White matter mean signal AFTER "11-func-anat-registration" (slice ' num2str(i3) ' of ' num2str(siz3) ')'],'Color','k','FontSize',14), legend({'White matter signal before WM-denoising'},'FontSize',12,'TextColor','k','Location','best')
        elseif stp==2
            subplot2n(4,1,1), plot(mF6g,'color','k','linewidth',1.5), hold on, plot(mF2g,'color',[0,0.55,0.69],'linewidth',0.2), ylabel('pcent sig change'),
                grid on,grid minor, xlim([1 length(mF2g)]),ylim([min(1.25.*[mF2g,mF6g]) 1.25.*max([mF2g,mF6g])]), title(['Gray matter mean signal AFTER "12-denoise3-CSFregress" (slice ' num2str(i3) ' of ' num2str(siz3) ')'],'Color','k','FontSize',14), legend({'Gray matter signal before WM-denoising'},'FontSize',12,'TextColor','k','Location','best')
            subplot2n(4,1,3), plot(mF6w,'color','k','linewidth',1.5), hold on, plot(mF2w,'color',[0.65,0.11,0.19],'linewidth',0.2), ylabel('pcent sig change'),
                grid on,grid minor, xlim([1 length(mF2g)]),ylim([min(1.25.*[mF2w,mF6w]) 1.25.*max([mF2w,mF6w])]), title(['White matter mean signal AFTER "12-denoise3-CSFregress" (slice ' num2str(i3) ' of ' num2str(siz3) ')'],'Color','k','FontSize',14), legend({'White matter signal before WM-denoising'},'FontSize',12,'TextColor','k','Location','best')
        end
        subplot2n(4,1,2), plot(mF2g,'color',[0,0.55,0.69],'linewidth',1.5), hold on, plot(mF6g,'color','k','linewidth',0.2), ylabel('pcent sig change'),
            grid on,grid minor, xlim([1 length(mF2g)]),ylim([min(1.25.*[mF2g,mF6g]) 1.25.*max([mF2g,mF6g])]), title(['Gray matter mean signal AFTER "13-denoise4-WMregress" (slice ' num2str(i3) ' of ' num2str(siz3) ')'],'Color',[0,0.55,0.69],'FontSize',14), legend({'Gray matter signal after WM-denoising'},'FontSize',12,'TextColor',[0,0.55,0.69],'Location','best')
        subplot2n(4,1,4), plot(mF2w,'color',[0.65,0.11,0.19],'linewidth',1.5), hold on, plot(mF6w,'color','k','linewidth',0.2), ylabel('pcent sig change'),
            grid on,grid minor, xlim([1 length(mF2g)]),ylim([min(1.25.*[mF2w,mF6w]) 1.25.*max([mF2w,mF6w])]), title(['White matter mean signal AFTER "13-denoise4-WMregress" (slice ' num2str(i3) ' of ' num2str(siz3) ')'],'Color',[0.65,0.11,0.19],'FontSize',14), legend({'White matter signal after WM-denoising'},'FontSize',12,'TextColor',[0.65,0.11,0.19],'Location','best')
        saveas(fighndl,[QCpath 'global_signal_12denoise3_vs_13denoise4_slice' sprintf('%.2d',i3) '.jpg'])
        close(fighndl)
        
        C3g = corrcoef(F6g'); C3g = uppertriangle(C3g); C3w = corrcoef(F6w'); C3w = uppertriangle(C3w);
        fighndl = figure('Position',[1 1 1152 720],'Color',[0.95,0.9,0.85],'InvertHardcopy','off','Visible','off'); % red [0.95,0.9,0.85], green [0.85,0.90,0.85], blue [0.89,0.94,0.99]
        if stp==1
            subplot2n(4,1,1), histogram(C3g,'EdgeColor','none','FaceColor','k','FaceAlpha',0.5,'Normalization','probability'); grid on,grid minor, xlim([-1 1]), hold on, xline(0,'Color','k','linewidth',1.5); hold off
                title(sprintf('Histogram of voxel-to-voxel GRAY MATTER correlations AFTER 11-func-anat-registration (slice %d of %d)\n%.1f%% of the correlations were > 0  [ mean=%.2f, median=%.2f, SD=%.2f ]',i3,siz3,100*length(find(C3g>0))/length(C3g),mean(C3g),median(C3g),std(C3g)),'Color','k','FontSize',14)
            subplot2n(4,1,3), histogram(C3w,'EdgeColor','none','FaceColor','k','FaceAlpha',0.5,'Normalization','probability'); grid on,grid minor, xlim([-1 1]), hold on, xline(0,'Color','k','linewidth',1.5); hold off
                title(sprintf('WHITE MATTER: AFTER 11-func-anat-registration (slice %d of %d): %.1f%% of the correlations were > 0  [ mean=%.2f, median=%.2f, SD=%.2f ]',i3,siz3,100*length(find(C3w>0))/length(C3w),mean(C3w),median(C3w),std(C3w)),'Color','k','FontSize',14)
        elseif stp==2
            subplot2n(4,1,1), histogram(C3g,'EdgeColor','none','FaceColor','k','FaceAlpha',0.5,'Normalization','probability'); grid on,grid minor, xlim([-1 1]), hold on, xline(0,'Color','k','linewidth',1.5); hold off
                title(sprintf('Histogram of voxel-to-voxel GRAY MATTER correlations AFTER 12-denoise3-CSFregress (slice %d of %d)\n%.1f%% of the correlations were > 0  [ mean=%.2f, median=%.2f, SD=%.2f ]',i3,siz3,100*length(find(C3g>0))/length(C3g),mean(C3g),median(C3g),std(C3g)),'Color','k','FontSize',14)
            subplot2n(4,1,3), histogram(C3w,'EdgeColor','none','FaceColor','k','FaceAlpha',0.5,'Normalization','probability'); grid on,grid minor, xlim([-1 1]), hold on, xline(0,'Color','k','linewidth',1.5); hold off
                title(sprintf('WHITE MATTER: AFTER 12-denoise3-CSFregress (slice %d of %d): %.1f%% of the correlations were > 0  [ mean=%.2f, median=%.2f, SD=%.2f ]',i3,siz3,100*length(find(C3w>0))/length(C3w),mean(C3w),median(C3w),std(C3w)),'Color','k','FontSize',14)
        end
        subplot2n(4,1,2), histogram(Cg,'EdgeColor','none','FaceColor',[0,0.55,0.69],'FaceAlpha',0.5,'Normalization','probability'); grid on,grid minor, xlim([-1 1]), hold on, xline(0,'Color','k','linewidth',1.5); hold off
            title(sprintf('GRAY MATTER: AFTER 13-denoise4-WMregress (slice %d of %d): %.1f%% of the correlations were > 0  [ mean=%.2f, median=%.2f, SD=%.2f ]',i3,siz3,100*length(find(Cg>0))/length(Cg),mean(Cg),median(Cg),std(Cg)),'Color',[0,0.55,0.69],'FontSize',14)
        subplot2n(4,1,4), histogram(Cw,'EdgeColor','none','FaceColor',[0.65,0.11,0.19],'FaceAlpha',0.5,'Normalization','probability'); grid on,grid minor, xlim([-1 1]), hold on, xline(0,'Color','k','linewidth',1.5); hold off
            title(sprintf('WHITE MATTER: AFTER 13-denoise4-WMregress (slice %d of %d): %.1f%% of the correlations were > 0  [ mean=%.2f, median=%.2f, SD=%.2f ]',i3,siz3,100*length(find(Cw>0))/length(Cw),mean(Cw),median(Cw),std(Cw)),'Color',[0.65,0.11,0.19],'FontSize',14)
        saveas(fighndl,[QCpath 'hist_FC_compare_12denoise3_13denoise4_slice' sprintf('%.2d',i3) '.jpg'])
        close(fighndl)
        F6g_=cat(1,F6g_,F6g); mF6g_=cat(1,mF6g_,mF6g); C3g_=cat(1,C3g_,C3g);
        F6w_=cat(1,F6w_,F6w); mF6w_=cat(1,mF6w_,mF6w); C3w_=cat(1,C3w_,C3w);
        clear F6g F6w mF6g mF6w C3g C3w
    end
    
    F2g_=cat(1,F2g_,F2g); mF2g_=cat(1,mF2g_,mF2g); F2w_=cat(1,F2w_,F2w); mF2w_=cat(1,mF2w_,mF2w); Cg_=cat(1,Cg_,Cg); Cw_=cat(1,Cw_,Cw); F4_=cat(1,F4_,F4); mF4_=cat(1,mF4_,mF4); C2_=cat(1,C2_,C2);
    clear F2g F2w F4 TR Smask1 svar1 svar2 mF2g mF2w mF4 Cg Cw C2
end


[~,svar1] = sort(var(F2g_,0,2)); [~,spsc1] = sort(meanPSC(F2g_));
[~,svar2] = sort(var(F2w_,0,2)); [~,spsc2] = sort(meanPSC(F2w_));
sortorder1b = sortorder1; sortorder1=[]; f1=0; %#ok<*ASGLU>
sortorder2b = sortorder2; sortorder2=[]; f2=0; %#ok<*ASGLU>
for im=1:length(sortorder1b)
    temp = sortorder1b{im}; temp = temp + f1; f1 = f1 + length(temp);
    sortorder1 = cat(1,sortorder1,temp); clear temp
    temp = sortorder2b{im}; temp = temp + f2; f2 = f2 + length(temp);
    sortorder2 = cat(1,sortorder2,temp); clear temp
end; clear im f1 f2
sortorder1 = spsc1; % options: spsc for percentage signal change, svar for variance, (comment out the line if you want to use distance from the center of the cord separately for each slice)
sortorder2 = spsc2; % options: spsc for percentage signal change, svar for variance, (comment out the line if you want to use distance from the center of the cord separately for each slice)
for im=max(sortorder1):-1:1, if isempty(find(sortorder1==im)), sortorder1(find(sortorder1>im)) = sortorder1(find(sortorder1>im))-1; end, end; clear im %#ok<*EFIND>
for im=max(sortorder2):-1:1, if isempty(find(sortorder2==im)), sortorder2(find(sortorder2>im)) = sortorder2(find(sortorder2>im))-1; end, end; clear im %#ok<*EFIND>

fighndl = figure('Position',[1 1 1152 720],'Color',[0.89,0.94,0.99],'InvertHardcopy','off','Visible','off'); % red [0.95,0.9,0.85], green [0.85,0.90,0.85], blue [0.89,0.94,0.99]
subplot2n(2,1,1), imagesc(F2g_(sortorder1,:)),colormap(cmap),colorbar, xlabel('time (in samples)'),ylabel(sprintf('voxels (sorted by mean percentage signal change;\nbottom row is highest)')),
  title(['Time series within the GRAY MATTER AFTER 13-DENOISE4-WMregress (all slices combined)'],'Color',[0,0.55,0.69],'FontSize',14);
subplot2n(2,1,2), imagesc(F2w_(sortorder2,:)),colormap(cmap),colorbar, xlabel('time (in samples)'),ylabel(sprintf('voxels (sorted by mean percentage signal change;\nbottom row is highest)')),
  title(['Time series within the WHITE MATTER AFTER 13-DENOISE4-WMregress (all slices combined)'],'Color',[0.65,0.11,0.19],'FontSize',14);
saveas(fighndl,[QCpath 'fMRI_ts_13denoise4_allSlices' '.jpg'])
saveas(fighndl,[QCpath2 'fMRI_ts_13denoise4_allSlices' '.jpg'])
 close(fighndl)
mF2g_=mean(mF2g_,1); mF2w_=mean(mF2w_,1); mF4_=mean(mF4_,1);
fighndl = figure('Position',[1 1 1152 720],'Color',[0.89,0.94,0.99],'InvertHardcopy','off','Visible','off'); % red [0.95,0.9,0.85], green [0.85,0.90,0.85], blue [0.89,0.94,0.99]
subplot2n(3,1,1), plot(mF4_,'color','k','linewidth',1.5), hold on, plot(mF2g_,'color',[0,0.55,0.69],'linewidth',0.2), plot(mF2w_,'color',[0.65,0.11,0.19],'linewidth',0.2), xlabel('time (in samples)'),ylabel('percentage signal change'),
    grid on,grid minor, xlim([1 length(mF2g_)]),ylim([1.25.*min([mF2g_,mF2w_,mF4_]) 1.25.*max([mF2g_,mF2w_,mF4_])]), title(['Global mean signal in UNPROCESSED RAW DATA (all slices averaged)'],'Color','k','FontSize',14), legend({'Global signal (raw data)'},'FontSize',12,'TextColor','k','Location','best')
subplot2n(3,1,2), plot(mF2g_,'color',[0,0.55,0.69],'linewidth',1.5), hold on, plot(mF2w_,'color',[0.65,0.11,0.19],'linewidth',0.2), plot(mF4_,'color','k','linewidth',0.2), xlabel('time (in samples)'),ylabel('percentage signal change'),
    grid on,grid minor, xlim([1 length(mF2g_)]),ylim([1.25.*min([mF2g_,mF2w_,mF4_]) 1.25.*max([mF2g_,mF2w_,mF4_])]), title(['Gray matter mean signal AFTER "13-DENOISE4-WMregress" (all slices averaged)'],'Color',[0,0.55,0.69],'FontSize',14), legend({'Gray matter signal after 13-denoise4'},'FontSize',12,'TextColor',[0,0.55,0.69],'Location','best')
subplot2n(3,1,3), plot(mF2w_,'color',[0.65,0.11,0.19],'linewidth',1.5), hold on, plot(mF4_,'color','k','linewidth',0.2), plot(mF2g_,'color',[0,0.55,0.69],'linewidth',0.2), xlabel('time (in samples)'),ylabel('percentage signal change'),
    grid on,grid minor, xlim([1 length(mF2g_)]),ylim([1.25.*min([mF2g_,mF2w_,mF4_]) 1.25.*max([mF2g_,mF2w_,mF4_])]), title(['White matter mean signal AFTER "13-DENOISE4-WMregress" (all slices averaged)'],'Color',[0.65,0.11,0.19],'FontSize',14), legend({'White matter signal after 13-denoise4'},'FontSize',12,'TextColor',[0.65,0.11,0.19],'Location','best')
saveas(fighndl,[QCpath 'global_signal_13denoise4_allSlices' '.jpg'])
 close(fighndl)
fighndl = figure('Position',[1 1 1152 720],'Color',[0.89,0.94,0.99],'InvertHardcopy','off','Visible','off'); % red [0.95,0.9,0.85], green [0.85,0.90,0.85], blue [0.89,0.94,0.99]
subplot2n(3,1,1), histogram(C2_,'EdgeColor','none','FaceColor','k','FaceAlpha',0.5,'Normalization','probability'); grid on,grid minor, xlim([-1 1]), hold on, xline(0,'Color','k','linewidth',1.5); hold off
    title(sprintf('Histogram of voxel-to-voxel correlations in UNPROCESSED RAW DATA (all slices combined)\n%.1f%% of the correlations were > 0  [ mean=%.2f, median=%.2f, SD=%.2f ]',100*length(find(C2_>0))/length(C2_),mean(C2_),median(C2_),std(C2_)),'Color','k','FontSize',14)
subplot2n(3,1,2), histogram(Cg_,'EdgeColor','none','FaceColor',[0,0.55,0.69],'FaceAlpha',0.5,'Normalization','probability'); grid on,grid minor, xlim([-1 1]), hold on, xline(0,'Color','k','linewidth',1.5); hold off
    title(sprintf('Histogram of voxel-to-voxel GRAY MATTER correlations AFTER 13-denoise4-WMregress (all slices combined)\n%.1f%% of the correlations were > 0  [ mean=%.2f, median=%.2f, SD=%.2f ]',100*length(find(Cg_>0))/length(Cg_),mean(Cg_),median(Cg_),std(Cg_)),'Color',[0,0.55,0.69],'FontSize',14)
subplot2n(3,1,3), histogram(Cw_,'EdgeColor','none','FaceColor',[0.65,0.11,0.19],'FaceAlpha',0.5,'Normalization','probability'); grid on,grid minor, xlim([-1 1]), hold on, xline(0,'Color','k','linewidth',1.5); hold off
    title(sprintf('Histogram of voxel-to-voxel WHITE MATTER correlations AFTER 13-denoise4-WMregress (all slices combined)\n%.1f%% of the correlations were > 0  [ mean=%.2f, median=%.2f, SD=%.2f ]',100*length(find(Cw_>0))/length(Cw_),mean(Cw_),median(Cw_),std(Cw_)),'Color',[0.65,0.11,0.19],'FontSize',14)
saveas(fighndl,[QCpath 'hist_FC_compare_13denoise4_allSlices' '.jpg'])
 close(fighndl)

if stp~=0
    mF6g_=mean(mF6g_,1); mF6w_=mean(mF6w_,1);
    fighndl = figure('Position',[1 1 1152 720],'Color',[0.89,0.94,0.99],'InvertHardcopy','off','Visible','off'); % red [0.95,0.9,0.85], green [0.85,0.90,0.85], blue [0.89,0.94,0.99]
    if stp==1
    	subplot2n(4,1,1), plot(mF6g_,'color','k','linewidth',1.5), hold on, plot(mF2g_,'color',[0,0.55,0.69],'linewidth',0.2), ylabel('pcent sig change'),
            grid on,grid minor, xlim([1 length(mF2g_)]),ylim([1.25.*min([mF2g_,mF6g_]) 1.25.*max([mF2g_,mF6g_])]), title(['Gray matter mean signal AFTER "11-func-anat-registration" (all slices averaged)'],'Color','k','FontSize',14), legend({'Gray matter signal before WM-denoising'},'FontSize',12,'TextColor','k','Location','best')
        subplot2n(4,1,3), plot(mF6w_,'color','k','linewidth',1.5), hold on, plot(mF2w_,'color',[0.65,0.11,0.19],'linewidth',0.2), ylabel('pcent sig change'),
            grid on,grid minor, xlim([1 length(mF2g_)]),ylim([1.25.*min([mF2w_,mF6w_]) 1.25.*max([mF2w_,mF6w_])]), title(['White matter mean signal AFTER "11-func-anat-registration" (all slices averaged)'],'Color','k','FontSize',14), legend({'White matter signal before WM-denoising'},'FontSize',12,'TextColor','k','Location','best')
    elseif stp==2
    	subplot2n(4,1,1), plot(mF6g_,'color','k','linewidth',1.5), hold on, plot(mF2g_,'color',[0,0.55,0.69],'linewidth',0.2), ylabel('pcent sig change'),
            grid on,grid minor, xlim([1 length(mF2g_)]),ylim([1.25.*min([mF2g_,mF6g_]) 1.25.*max([mF2g_,mF6g_])]), title(['Gray matter mean signal AFTER "12-DENOISE3-CSFregress" (all slices averaged)'],'Color','k','FontSize',14), legend({'Gray matter signal before WM-denoising'},'FontSize',12,'TextColor','k','Location','best')
        subplot2n(4,1,3), plot(mF6w_,'color','k','linewidth',1.5), hold on, plot(mF2w_,'color',[0.65,0.11,0.19],'linewidth',0.2), ylabel('pcent sig change'),
            grid on,grid minor, xlim([1 length(mF2g_)]),ylim([1.25.*min([mF2w_,mF6w_]) 1.25.*max([mF2w_,mF6w_])]), title(['White matter mean signal AFTER "12-DENOISE3-CSFregress" (all slices averaged)'],'Color','k','FontSize',14), legend({'White matter signal before WM-denoising'},'FontSize',12,'TextColor','k','Location','best')
    end
    subplot2n(4,1,2), plot(mF2g_,'color',[0,0.55,0.69],'linewidth',1.5), hold on, plot(mF6g_,'color','k','linewidth',0.2), ylabel('pcent sig change'),
        grid on,grid minor, xlim([1 length(mF2g_)]),ylim([1.25.*min([mF2g_,mF6g_]) 1.25.*max([mF2g_,mF6g_])]), title(['Gray matter mean signal AFTER "13-DENOISE4-WMregress" (all slices averaged)'],'Color',[0,0.55,0.69],'FontSize',14), legend({'Gray matter signal after WM-denoising'},'FontSize',12,'TextColor',[0,0.55,0.69],'Location','best')
    subplot2n(4,1,4), plot(mF2w_,'color',[0.65,0.11,0.19],'linewidth',1.5), hold on, plot(mF6w_,'color','k','linewidth',0.2), ylabel('pcent sig change'),
        grid on,grid minor, xlim([1 length(mF2g_)]),ylim([1.25.*min([mF2w_,mF6w_]) 1.25.*max([mF2w_,mF6w_])]), title(['White matter mean signal AFTER "13-DENOISE4-WMregress" (all slices averaged)'],'Color',[0.65,0.11,0.19],'FontSize',14), legend({'White matter signal after WM-denoising'},'FontSize',12,'TextColor',[0.65,0.11,0.19],'Location','best')
    saveas(fighndl,[QCpath 'global_signal_12denoise3_vs_13denoise4_allSlices' '.jpg'])
    saveas(fighndl,[QCpath2 'global_signal_12denoise3_vs_13denoise4_allSlices' '.jpg'])
     close(fighndl)
    
     fighndl = figure('Position',[1 1 1152 720],'Color',[0.89,0.94,0.99],'InvertHardcopy','off','Visible','off'); % red [0.95,0.9,0.85], green [0.85,0.90,0.85], blue [0.89,0.94,0.99]
    if stp==1
    	subplot2n(4,1,1), histogram(C3g_,'EdgeColor','none','FaceColor','k','FaceAlpha',0.5,'Normalization','probability'); grid on,grid minor, xlim([-1 1]), hold on, xline(0,'Color','k','linewidth',1.5); hold off
            title(sprintf('Histogram of voxel-to-voxel GRAY MATTER correlations AFTER 11-registration (all slices combined)\n%.1f%% of the correlations were > 0  [ mean=%.2f, median=%.2f, SD=%.2f ]',100*length(find(C3g_>0))/length(C3g_),mean(C3g_),median(C3g_),std(C3g_)),'Color','k','FontSize',14)
        subplot2n(4,1,3), histogram(C3w_,'EdgeColor','none','FaceColor','k','FaceAlpha',0.5,'Normalization','probability'); grid on,grid minor, xlim([-1 1]), hold on, xline(0,'Color','k','linewidth',1.5); hold off
            title(sprintf('WHITE MATTER: AFTER 11-registration (all slices): %.1f%% of the correlations were > 0  [ mean=%.2f, median=%.2f, SD=%.2f ]',100*length(find(C3w_>0))/length(C3w_),mean(C3w_),median(C3w_),std(C3w_)),'Color','k','FontSize',14)
    elseif stp==2
    	subplot2n(4,1,1), histogram(C3g_,'EdgeColor','none','FaceColor','k','FaceAlpha',0.5,'Normalization','probability'); grid on,grid minor, xlim([-1 1]), hold on, xline(0,'Color','k','linewidth',1.5); hold off
            title(sprintf('Histogram of voxel-to-voxel GRAY MATTER correlations AFTER 12-denoise3-CSFregress (all slices combined)\n%.1f%% of the correlations were > 0  [ mean=%.2f, median=%.2f, SD=%.2f ]',100*length(find(C3g_>0))/length(C3g_),mean(C3g_),median(C3g_),std(C3g_)),'Color','k','FontSize',14)
        subplot2n(4,1,3), histogram(C3w_,'EdgeColor','none','FaceColor','k','FaceAlpha',0.5,'Normalization','probability'); grid on,grid minor, xlim([-1 1]), hold on, xline(0,'Color','k','linewidth',1.5); hold off
            title(sprintf('WHITE MATTER: AFTER 12-denoise3-CSFregress (all slices): %.1f%% of the correlations were > 0  [ mean=%.2f, median=%.2f, SD=%.2f ]',100*length(find(C3w_>0))/length(C3w_),mean(C3w_),median(C3w_),std(C3w_)),'Color','k','FontSize',14)
    end
    subplot2n(4,1,2), histogram(Cg_,'EdgeColor','none','FaceColor',[0,0.55,0.69],'FaceAlpha',0.5,'Normalization','probability'); grid on,grid minor, xlim([-1 1]), hold on, xline(0,'Color','k','linewidth',1.5); hold off
       title(sprintf('GRAY MATTER: AFTER 13-denoise4-WMregress (all slices): %.1f%% of correlations > 0  [ mean=%.2f, median=%.2f, SD=%.2f ]',100*length(find(Cg_>0))/length(Cg_),mean(Cg_),median(Cg_),std(Cg_)),'Color',[0,0.55,0.69],'FontSize',14)
    subplot2n(4,1,4), histogram(Cw_,'EdgeColor','none','FaceColor',[0.65,0.11,0.19],'FaceAlpha',0.5,'Normalization','probability'); grid on,grid minor, xlim([-1 1]), hold on, xline(0,'Color','k','linewidth',1.5); hold off
       title(sprintf('WHITE MATTER: AFTER 13-denoise4-WMregress (all slices): %.1f%% of the correlations were > 0  [ mean=%.2f, median=%.2f, SD=%.2f ]',100*length(find(Cw_>0))/length(Cw_),mean(Cw_),median(Cw_),std(Cw_)),'Color',[0.65,0.11,0.19],'FontSize',14)
    saveas(fighndl,[QCpath 'hist_FC_compare_12denoise3_13denoise4_allSlices' '.jpg'])
    saveas(fighndl,[QCpath2 'hist_FC_compare_12denoise3_13denoise4_allSlices' '.jpg'])
     close(fighndl)
end
clear svar1 spsc1 svar2 spsc2 sortorder1 sortorder2 F2g_ mF2g_ F2w_ mF2w_ Cg_ Cw_ F4_ mF4_ C2_ F6g_ F6w_ mF6g_ mF6w_ C3g_ C3w_ cmap

%% Make QC video (needs audio video toolbox in Matlab)
if ~isempty(which('VideoWriter')) && ((save_videos==1)||(save_videos==2)) % check if Matlab's audio video toolbox exists

subplot = @(m,n,p) subtightplot(m, n, p, [0.001 0.005], [0.01 0.001], [0.01 0.001]);
scfMRItb_04_unzipFile(base_dir_sub, fname_anat, '_mask_SC')
mask_SC2 = load_untouch_nii([base_dir_sub fname_anat '_mask_SC.nii']); mask_SC2 = mask_SC2.img;
for M=1:size(mask_SC2,3)
    mask_SC = rot90(mask_SC2(:,:,M));
    [xmask_SC,ymask_SC] = find(mask_SC==1);
    minx_SC(:,M) = round(min(xmask_SC)-0.1*(size(mask_SC,1))); maxx_SC(:,M)=round(max(xmask_SC)+0.1*(size(mask_SC,1)));
    miny_SC(:,M) = round(min(ymask_SC)-0.1*(size(mask_SC,2))); maxy_SC(:,M)=round(max(ymask_SC)+0.1*(size(mask_SC,2)));
    if (maxx_SC(:,M)-minx_SC(:,M))<(maxy_SC(:,M)-miny_SC(:,M))
        factr = ((maxy_SC(:,M)-miny_SC(:,M))-(maxx_SC(:,M)-minx_SC(:,M)))/2;
        if factr==round(factr)
            maxx_SC(:,M)=maxx_SC(:,M)+factr; minx_SC(:,M)=minx_SC(:,M)-factr;
        else
            maxx_SC(:,M)=maxx_SC(:,M)+factr+0.5; minx_SC(:,M)=minx_SC(:,M)-factr+0.5;
        end
    elseif (maxy_SC(:,M)-miny_SC(:,M))<(maxx_SC(:,M)-minx_SC(:,M))
        factr = ((maxx_SC(:,M)-minx_SC(:,M))-(maxy_SC(:,M)-miny_SC(:,M)))/2;
        if factr==round(factr)
            maxy_SC(:,M)=maxy_SC(:,M)+factr; miny_SC(:,M)=miny_SC(:,M)-factr;
        else
            maxy_SC(:,M)=maxy_SC(:,M)+factr+0.5; miny_SC(:,M)=miny_SC(:,M)-factr+0.5;
        end
    end
    clear mask_SC xmask_SC ymask_SC factr
end; clear M

for i3 = 1 : siz3
    try waitbar((i3/siz3),wbar3,sprintf('13. Generate QC video: slice (%d) of (%d)',i3,siz3)); catch, end
    fprintf('13. Generate QC video: slice (%d) of (%d)\n',i3,siz3)
    if stp==2
        scfMRItb_04_resplitData(base_dir_sub, fname, ['_denoised' cutoff_str 'WM'], siz3) % if slice-wise data doesn't exist but combined 4D data exists, then split the data to generate the needed slice-wise data
        F1 = load_untouch_nii([base_dir_sub fname '_slice' num2str(i3) '_denoised' cutoff_str 'WM.nii']);
    elseif stp==1
        scfMRItb_04_resplitData(base_dir_sub, fname, '_WM', siz3) % if slice-wise data doesn't exist but combined 4D data exists, then split the data to generate the needed slice-wise data
        F1 = load_untouch_nii([base_dir_sub fname '_slice' num2str(i3) '_WM.nii']);
    end
    TR = F1.hdr.dime.pixdim(5); F1 = rot90(single(squeeze(F1.img)));
    video = VideoWriter([QCpath 'fMRIvideo_13denoise4_slice' sprintf('%.2d',i3) '.mp4'],'MPEG-4'); %#ok<*TNMLP> % create the video object
    video.FrameRate = round(30/TR); % 30 seconds of data in every second of the video
    video.Quality = 90; % very limited compression so that the images are shown as is (and thus avoid misunderstandings due to video quality)
    open(video); % open the file for writing
    
    [sizeFx_SC,sizeFy_SC] = size(F1(:,:,1));
    F2 = F1(minx_SC:maxx_SC,miny_SC:maxy_SC,:);
    for i5=1:size(F1,3)
        F1(:,:,i5) = imresize(F2(:,:,i5),[sizeFx_SC sizeFy_SC]);
    end
    
    if (save_videos==1)
    if stp==1
        scfMRItb_04_resplitData(base_dir_sub, fname, '_warped', siz3) % if slice-wise data doesn't exist but combined 4D data exists, then split the data to generate the needed slice-wise data
        F3 = load_untouch_nii([base_dir_sub fname '_slice' num2str(i3) '_warped.nii']); F3 = rot90(squeeze(F3.img));
    elseif stp==2
        scfMRItb_04_resplitData(base_dir_sub, fname, ['_denoised' cutoff_str], siz3) % if slice-wise data doesn't exist but combined 4D data exists, then split the data to generate the needed slice-wise data
        F3 = load_untouch_nii([base_dir_sub fname '_slice' num2str(i3) '_denoised' cutoff_str '.nii']); F3 = rot90(single(squeeze(F3.img)));
    end
    [sizeFx,sizeFy] = size(F3(:,:,1));
    F4 = F3(minx_SC:maxx_SC,miny_SC:maxy_SC,:);
    for i5=1:size(F3,3)
        F3(:,:,i5) = imresize(F4(:,:,i5),[sizeFx sizeFy]);
    end
    clear sizeFx sizeFy F2 F4 i5 sizeFx_SC sizeFy_SC
    
    elseif (save_videos==2)
    if Smask_err==0
        scfMRItb_04_resplitData(base_dir_sub, fname, '_Smask', siz3) % if slice-wise data doesn't exist but combined 4D data exists, then split the data to generate the needed slice-wise data
        scfMRItb_04_unzipFile(base_dir_sub, fname, ['_slice' num2str(i3) '_Smask'])
        Smask1 = load_untouch_nii([base_dir_sub fname '_slice' num2str(i3) '_Smask.nii']); Smask1 = Smask1.img;
    else
        Smask1 = Smask1_(:,:,i3);
    end
    Smask1 = rot90(Smask1);
    [xmask,ymask] = find(Smask1==1);
    minx=round(min(xmask)-0.1*(size(Smask1,1))); maxx=round(max(xmask)+0.1*(size(Smask1,1)));
    miny=round(min(ymask)-0.1*(size(Smask1,2))); maxy=round(max(ymask)+0.1*(size(Smask1,2)));
    if (maxx-minx)<(maxy-miny)
        factr = ((maxy-miny)-(maxx-minx))/2;
        if factr==round(factr)
            maxx=maxx+factr; minx=minx-factr;
        else
            maxx=maxx+factr+0.5; minx=minx-factr+0.5;
        end
    elseif (maxy-miny)<(maxx-minx)
        factr = ((maxx-minx)-(maxy-miny))/2;
        if factr==round(factr)
            maxy=maxy+factr; miny=miny-factr;
        else
            maxy=maxy+factr+0.5; miny=miny-factr+0.5;
        end
    end
    scfMRItb_04_resplitData(base_dir_sub, fname, '', siz3) % if slice-wise data doesn't exist but combined 4D data exists, then split the data to generate the needed slice-wise data
    F3 = load_untouch_nii([base_dir_sub fname '_slice' num2str(i3) '.nii']); F3 = rot90(single(squeeze(F3.img)));
    [sizeFx,sizeFy] = size(F3(:,:,1));
    F4 = F3(minx:maxx,miny:maxy,:);
    for i5=1:size(F3,3)
        F3(:,:,i5) = imresize(F4(:,:,i5),[sizeFx sizeFy]);
    end
    clear maxx maxy minx miny xmask ymask Smask1 factr sizeFx sizeFy F2 F4 i5 sizeFx_SC sizeFy_SC        
    end
    
    for ii=1:size(F1,3) % loop across the number of images
        I = F1(:,:,ii); %read the next image
        I(find(isnan(I)))=0;
        I = (I - min(I(:))) ./ (max(I(:)) - min(I(:)));
        I2 = F3(:,:,ii); %read the next image
        I2(find(isnan(I2)))=0;
        I2 = (I2 - min(I2(:))) ./ (max(I2(:)) - min(I2(:)));
        fighndl = figure('Position',[round((scrsz(3)-scrsz(4))/2) 1 round(0.75*scrsz(3)) round(0.75*scrsz(4))],'Color',[0.95,0.9,0.85],'InvertHardcopy','off','Visible','off'); % [0.85,0.90,0.85]
        if (save_videos==1)
        if stp==1
            subplot(1,2,1), imshow(I2, [0 max(I2(:))], 'InitialMagnification', 'fit'); title({['fMRI data after "11. func-anat-registration" (slice ' num2str(i3) ' of ' num2str(siz3) ')'];['Time point ' num2str(ii) ' of ' num2str(size(F1,3))]},'Color','k','FontSize',18);
        elseif stp==2
            subplot(1,2,1), imshow(I2, [0 max(I2(:))], 'InitialMagnification', 'fit'); title({['fMRI data after "12. denoise-3 (CSF regression)" (slice ' num2str(i3) ' of ' num2str(siz3) ')'];['Time point ' num2str(ii) ' of ' num2str(size(F1,3))]},'Color','k','FontSize',18);
        end
        elseif (save_videos==2)
            subplot(1,2,1), imshow(I2, [0 max(I2(:))], 'InitialMagnification', 'fit'); title({['Unprocessed raw fMRI data (slice ' num2str(i3) ' of ' num2str(siz3) ')'];['Time point ' num2str(ii) ' of ' num2str(size(F1,3))]},'Color','k','FontSize',18);
        end
        subplot(1,2,2), imshow(I, [0 max(I(:))], 'InitialMagnification', 'fit'); title({['fMRI data after "13. denoise-4 (WM regression)" (slice ' num2str(i3) ' of ' num2str(siz3) ')'];['Time point ' num2str(ii) ' of ' num2str(size(F1,3))]},'Color',[0,0.55,0.69],'FontSize',18);
        frame = getframe(gcf);
        writeVideo(video,frame); % write the image to file
        close(fighndl)
        clear I I2 frame
    end
    close(video); %close the file
    clear F1 F3 video ii TR
end; clear i3
clear maxx_SC maxy_SC minx_SC miny_SC F

end

try
if nargin<4
    close(wbar3)
end
catch
end

end
```
