## Supplementary material for "Neptune: a toolbox for spinal cord functional MRI data processing and quality assurance": Neptune software: scfMRItb_14_COVregress.m

```
function scfMRItb_14_COVregress(base_dir_sub,fname,fname_anat_orig,covregress_options,cov_files,slices, varargin)

fname_anat = [fname_anat_orig '_reduced'];
cutoff = 0.50; cutoff_str = '50';

if nargin<7
    wbar3 = waitbar(0,'14. Regress-out additional covariates...','Name','Progress(14): Regress-out additional covariate signals...','Units','normalized','Position',[3/5 0 1/5 1/17]);
else
    wbar3 = varargin{1};
end

if nargin<8
    save_videos = 1; % 1 = save QC videos (generates sizeable .mp4 files and takes a few minutes to do it). This is turned on by default, but if you don't want to save videos then set it to 0
else
    save_videos = varargin{2};
end
if nargin<9
    Rns = 0; % 0 = don't create runX subfolder; X = create runX subfolder
else
    Rns = varargin{3};
end

if ~(exist([base_dir_sub 'QC'],'dir'))
    mkdir([base_dir_sub 'QC'])
end
if ~(exist([base_dir_sub 'QC' '/14_denoise5-COVregress'],'dir'))
    mkdir([base_dir_sub 'QC' '/14_denoise5-COVregress'])
end
if Rns==0
    QCpath = [base_dir_sub 'QC' '/14_denoise5-COVregress/'];
else
    if ~(exist([base_dir_sub 'QC' '/14_denoise5-COVregress' '/run' num2str(Rns)],'dir'))
        mkdir([base_dir_sub 'QC' '/14_denoise5-COVregress' '/run' num2str(Rns)])
    end
    QCpath = [base_dir_sub 'QC' '/14_denoise5-COVregress' '/run' num2str(Rns) '/'];
end

if ~(exist([base_dir_sub 'QC' '/QC_shortlist'],'dir'))
    mkdir([base_dir_sub 'QC' '/QC_shortlist'])
end
if Rns==0
    QCpath2 = [base_dir_sub 'QC' '/QC_shortlist/'];
else
    if ~(exist([base_dir_sub 'QC' '/QC_shortlist' '/run' num2str(Rns)],'dir'))
        mkdir([base_dir_sub 'QC' '/QC_shortlist' '/run' num2str(Rns)])
    end
    QCpath2 = [base_dir_sub 'QC' '/QC_shortlist' '/run' num2str(Rns) '/'];
end
warning off

%% prepare covariates
covariates = []; % this will finally be of the following dimensions: #timepoints X #covariates X #slices

if covregress_options(1,1)==1
    % case 'Motion parameters (2 translation)'
    load([base_dir_sub 'motion_parameters.mat'],'motion_parameters')
    for i3=1:length(motion_parameters)
        m(:,:,i3) = motion_parameters{i3};
    end
    covariates = cat(2,covariates,m);
    clear motion_parameters m
elseif covregress_options(1,1)==2
    % case 'Motion parameters (2 translation + 2 derivatives)'
    load([base_dir_sub 'motion_parameters.mat'],'motion_parameters')
    for i3=1:length(motion_parameters)
        tmp = motion_parameters{i3};
        m(:,:,i3) = tmp;
        for j=2:size(tmp,1)
            delta_tmp((j-1),:) = tmp(j,:) - tmp((j-1),:); % temporal derivative in mm along x and y directions
        end
        delta_tmp = cat(1,median(delta_tmp,1).*ones(1,size(delta_tmp,2)),delta_tmp);
        m2(:,:,i3) = delta_tmp;
        clear j tmp delta_tmp
    end
    covariates = cat(2,covariates,m,m2);
    clear motion_parameters m m2
end

if (covregress_options(2,1)==1) || (covregress_options(2,1)==2)
    % case 'Scrubbing (volumes with motion > half voxel size)'  OR  case 'Scrubbing (user-defined motion threshold)'
    mmthresh = covregress_options(2,2);
    load([base_dir_sub 'motion_parameters.mat'],'FD')
    FD = cat(1,median(FD,1).*ones(1,size(FD,2)),FD);
    thFD = (FD >= mmthresh);
    covariates = cat(2,covariates,reshape(thFD,[size(thFD,1) 1 size(thFD,2)]));
    clear mmthresh FD thFD
end


if covregress_options(3,1)==1
    % case 'Regression with user-defined covariate(s)'
    if (strcmp(cov_files(end-3:end),'.txt')==1)||(strcmp(cov_files(end-3:end),'.csv')==1)||(strcmp(cov_files(end-3:end),'.xls')==1)||(strcmp(cov_files(end-4:end),'.xlsx')==1)||(strcmp(cov_files(end-3:end),'.dat')==1)
        if exist('readmatrix.m','file') % if MATLAB R2019a or later
            covs = readmatrix(cov_files);
        else
            if (strcmp(cov_files(end-3:end),'.txt')==1)
                covs = importdata(cov_files);
            elseif (strcmp(cov_files(end-3:end),'.csv')==1)||(strcmp(cov_files(end-3:end),'.xls')==1)||(strcmp(cov_files(end-4:end),'.xlsx')==1)
                try
                covs = readtable(cov_files,'ReadVariableNames',0); % need at least MATLAB R2013b for this function. The assumption is that nobody is using MATLAB versions older than this. But just in case somebody is, an option with xlsread is given next but I cannot guarantee that it will work as MATLAB phases out old functions.
                covs = covs{:,:};
                catch
                covs = xlsread(cov_files); % This will run if you have MATLAB older than R2013b and thus could not use readtable(). I cannot guarantee that it will work as MATLAB phases out old functions.
                end                    
            elseif strcmp(cov_files(end-3:end),'.dat')==1
                covs = load(cov_files);
            end
        end
        if (size(covs,2)~=slices) && (size(covs,3)==1) % if same covariate for all slices
            covs = repmat(covs,[1 1 slices]);
        elseif (size(covs,2)==slices) && (size(covs,3)==1) % if it is the same thing as a covariate but different for each slice such that the input was timeXslices, make it timeX1Xslices
            covs = reshape(covs,[size(covs,1) 1 size(covs,2)]);
        end
        if (size(covs,1)==1)&&(size(covs,2)~=1)&&(size(covs,3)==1)
            covs = permute(covs,[2 1 3]);
        end
        if size(covs,1)==1
            F = load_untouch_nii([base_dir_sub fname '.nii']);
            covs = repmat(cov,[F.hdr.dime.dim(5) 1 1]);
            clear F
        end
        
    elseif strcmp(cov_files(end-3:end),'.mat')==1
        covs = load(cov_files);
        covs = struct2cell(covs);
        covs = covs{1};
        if iscell(covs)
            for i3=1:length(covs)
                tmp(:,:,i3) = covs{i3};
            end
            if (size(tmp,1)==1)&&(size(tmp,2)~=1)&&(size(tmp,3)==1)
                tmp = permute(tmp,[2 1 3]);
            end
            clear covs
            if (size(tmp,2)~=slices) && (size(tmp,3)==1) % if same covariate for all slices
                covs = repmat(tmp,[1 1 slices]);
            elseif (size(tmp,2)==slices) && (size(tmp,3)==1) % if it is the same thing as a covariate but different for each slice such that the input was timeXslices, make it timeX1Xslices
                covs = reshape(tmp,[size(tmp,1) 1 size(tmp,2)]);
            else % separate covariate for each slice
                covs = tmp;
            end
            clear tmp
            if size(covs,1)==1
                F = load_untouch_nii([base_dir_sub fname '.nii']);
                covs = repmat(cov,[F.hdr.dime.dim(5) 1 1]);
                clear F
            end
        else
            if (size(covs,2)~=slices) && (size(covs,3)==1) % if same covariate for all slices
                covs = repmat(covs,[1 1 slices]);
            elseif (size(covs,2)==slices) && (size(covs,3)==1) % if it is the same thing as a covariate but different for each slice such that the input was timeXslices, make it timeX1Xslices
                covs = reshape(covs,[size(covs,1) 1 size(covs,2)]);
            end
            if (size(covs,1)==1)&&(size(covs,2)~=1)&&(size(covs,3)==1)
                covs = permute(covs,[2 1 3]);
            end
            if size(covs,1)==1
                F = load_untouch_nii([base_dir_sub fname '.nii']);
                covs = repmat(cov,[F.hdr.dime.dim(5) 1 1]);
                clear F
            end
        end
    end
    
    if size(covs,1)>size(covariates,1)
        covs = covs(size(covs,1)-size(covariates,1)+1:end,:,:);
    end
    covariates = cat(2,covariates,covs);
end

readme_covariates = sprintf('The variable ''covariates'' provides the final covariates used in step-14 (denoise-5, additional covariates regression). It''s dimensions are:\n#timepoints X #covariates X #slices\nIf slice-specific covariates were not specified, the same covariates were replicated across slices for execution\n');
save([base_dir_sub 'covariates_denoise5_COVregress.mat'],'covariates','readme_covariates')

%% ----- REGRESS OUT COVARIATE SIGNAL(S) FROM fMRI DATA (USING INPUT COVARIATES) ------------------

scfMRItb_04_unzipFile(base_dir_sub, fname_anat, '')
A = load_untouch_nii([base_dir_sub fname_anat '.nii']);
siz3 = size(A.img,3);

scfMRItb_04_unzipFile(base_dir_sub, fname_anat, '_mask_WM')
scfMRItb_04_unzipFile(base_dir_sub, fname_anat, '_mask_GM')
mask_WM = load_untouch_nii([base_dir_sub fname_anat '_mask_WM.nii']); mask_WM = mask_WM.img;
mask_GM = load_untouch_nii([base_dir_sub fname_anat '_mask_GM.nii']); mask_GM = mask_GM.img;
mask_any = single(mask_WM | mask_GM);

for i3 = 1 : siz3
    try waitbar((i3/siz3)/3,wbar3,sprintf('14. Regress-out covariates signal: slice (%d) of (%d)',i3,siz3)); catch, end
    disp(sprintf('Regress-out additional covariates signal: slice (%d) of (%d) -> subject %s',i3,siz3,fname)) %#ok<DSPS>
    
    try % use denoised data ('denoise2' and 'denoise3') if that step was performed
        scfMRItb_04_resplitData(base_dir_sub, fname, ['_denoised' cutoff_str 'WM'], siz3) % if slice-wise data doesn't exist but combined 4D data exists, then split the data to generate the needed slice-wise data
        scfMRItb_04_unzipFile(base_dir_sub, fname, ['_slice' num2str(i3) '_denoised' cutoff_str 'WM'])
        F = load_untouch_nii([base_dir_sub fname '_slice' num2str(i3) '_denoised' cutoff_str  'WM' '.nii']);
        flag=1;
    catch
        try % use denoised data ('denoise2' only) if that step was performed
            scfMRItb_04_resplitData(base_dir_sub, fname, ['_denoised' cutoff_str], siz3) % if slice-wise data doesn't exist but combined 4D data exists, then split the data to generate the needed slice-wise data
            scfMRItb_04_unzipFile(base_dir_sub, fname, ['_slice' num2str(i3) '_denoised' cutoff_str])
            F = load_untouch_nii([base_dir_sub fname '_slice' num2str(i3) '_denoised' cutoff_str '.nii']);
            flag=2;
        catch
            try % use denoised data ('denoise3' only) if that step was performed
                scfMRItb_04_resplitData(base_dir_sub, fname, '_WM', siz3) % if slice-wise data doesn't exist but combined 4D data exists, then split the data to generate the needed slice-wise data
                scfMRItb_04_unzipFile(base_dir_sub, fname, ['_slice' num2str(i3) 'WM'])
                F = load_untouch_nii([base_dir_sub fname '_slice' num2str(i3) 'WM' '.nii']);
                flag=3;
            catch % else use co-registered functional data
                scfMRItb_04_resplitData(base_dir_sub, fname, '_warped', siz3) % if slice-wise data doesn't exist but combined 4D data exists, then split the data to generate the needed slice-wise data
                scfMRItb_04_unzipFile(base_dir_sub, fname, ['_slice' num2str(i3) '_warped'])
                F = load_untouch_nii([base_dir_sub fname '_slice' num2str(i3) '_warped.nii']);
                flag=4;
            end
        end
    end
    F_denoised = F;
    F = single(squeeze(F.img));
    
    covariates_perslice = [covariates(:,:,i3) , ones(size(covariates,1),1)];

    % actually do the regression now!
    for i1 = 1 : size(F,1)
        for i2 = 1 : size(F,2)
            if mask_any(i1,i2,i3) % do in both GM and WM
                temp = squeeze(F(i1,i2,:));                     % original
%                 w = lscov(covariates_perslice,temp);            % Ordinary least squares
                w = regress(temp,covariates_perslice);          % Multiple linear regression using least squares
                F(i1,i2,:) = temp - covariates_perslice(:,1:end-1) * w(1:end-1);    % do denoising
                clear temp w
            end
        end
    end
    
    F_denoised.img = reshape(F, [size(F,1) size(F,2) 1 size(F,3)]);
    
    if flag==1 % if denoised data ('denoise2' and 'denoise3') was used
        unix(['rm -f ' base_dir_sub fname '_slice' num2str(i3) '_denoised' cutoff_str 'WMcov.nii']);
        save_untouch_nii(F_denoised, [base_dir_sub fname '_slice' num2str(i3) '_denoised' cutoff_str 'WMcov.nii']);
    elseif flag==2 % if denoised data ('denoise2' only) was used
        unix(['rm -f ' base_dir_sub fname '_slice' num2str(i3) '_denoised' cutoff_str 'cov.nii']);
        save_untouch_nii(F_denoised, [base_dir_sub fname '_slice' num2str(i3) '_denoised' cutoff_str 'cov.nii']);
    elseif flag==3 % if denoised data ('denoise3' only) was used
        unix(['rm -f ' base_dir_sub fname '_slice' num2str(i3) '_WMcov.nii']);
        save_untouch_nii(F_denoised, [base_dir_sub fname '_slice' num2str(i3) '_WMcov.nii']);
    elseif flag==4 % if co-registered pre-denoised data was used
        unix(['rm -f ' base_dir_sub fname '_slice' num2str(i3) '_cov.nii']);
        save_untouch_nii(F_denoised, [base_dir_sub fname '_slice' num2str(i3) '_cov.nii']);
    end
    clear F F_denoised covariates_perslice
end
warning on

%% time series plots, global signal and FC histogram
scrsz = get(0,'ScreenSize'); pause on;
cmap = colormap('bone'); cmap = (1-exp(-cmap))./(1-exp(-1)); 

scfMRItb_04_unzipFile(base_dir_sub, fname_anat, '_mask_GM')
scfMRItb_04_unzipFile(base_dir_sub, fname_anat, '_mask_WM')
mask_GM = load_untouch_nii([base_dir_sub fname_anat '_mask_GM.nii']); mask_GM = mask_GM.img;
mask_WM = load_untouch_nii([base_dir_sub fname_anat '_mask_WM.nii']); mask_WM = mask_WM.img;

if exist([base_dir_sub fname '_slice' num2str(i3) '_denoised' cutoff_str 'WM.nii'],'file') || exist([base_dir_sub fname '_slice' num2str(i3) '_denoised' cutoff_str 'WM.nii.gz'],'file') || exist([base_dir_sub fname '_denoised' cutoff_str 'WM.nii'],'file') || exist([base_dir_sub fname '_denoised' cutoff_str 'WM.nii.gz'],'file')
    stp=4;
elseif exist([base_dir_sub fname '_slice' num2str(i3) '_denoised' cutoff_str '.nii'],'file') || exist([base_dir_sub fname '_slice' num2str(i3) '_denoised' cutoff_str '.nii.gz'],'file') || exist([base_dir_sub fname '_denoised' cutoff_str '.nii'],'file') || exist([base_dir_sub fname '_denoised' cutoff_str '.nii.gz'],'file')
    stp=3;
elseif exist([base_dir_sub fname '_slice' num2str(i3) '_WM.nii'],'file') || exist([base_dir_sub fname '_slice' num2str(i3) '_WM.nii.gz'],'file') || exist([base_dir_sub fname '_WM.nii'],'file') || exist([base_dir_sub fname '_WM.nii.gz'],'file')
    stp=2;
elseif exist([base_dir_sub fname '_slice' num2str(i3) '_warped.nii'],'file') || exist([base_dir_sub fname '_slice' num2str(i3) '_warped.nii.gz'],'file') || exist([base_dir_sub fname '_warped.nii'],'file') || exist([base_dir_sub fname '_warped.nii.gz'],'file')
    stp=1;
end

Smask_err=0;
try
    scfMRItb_04_resplitData(base_dir_sub, fname, '_Smask', siz3) % if slice-wise data doesn't exist but combined 4D data exists, then split the data to generate the needed slice-wise data
catch
    load([base_dir_sub fname '_mask_NS' '.mat'])
    Smask1_ = ~mask_NS.img; clear mask_NS
    Smask_err=1;
end

F2g_=[]; mF2g_=[]; F2w_=[]; mF2w_=[]; Cg_=[]; Cw_=[]; F4_=[]; mF4_=[]; C2_=[];
if stp~=0, F6g_=[]; mF6g_=[]; F6w_=[]; mF6w_=[]; C3g_=[]; C3w_=[]; end
for i3 = 1 : siz3
    try waitbar((i3/siz3),wbar3,sprintf('14. QC plots: slice (%d) of (%d)',i3,siz3)); catch, end
    fprintf('14. QC plots: slice (%d) of (%d)\n',i3,siz3)
    if stp==4
        scfMRItb_04_resplitData(base_dir_sub, fname, ['_denoised' cutoff_str 'WMcov'], siz3) % if slice-wise data doesn't exist but combined 4D data exists, then split the data to generate the needed slice-wise data
        scfMRItb_04_unzipFile(base_dir_sub, fname, ['_slice' num2str(i3) '_denoised' cutoff_str 'WMcov'])
        F1 = load_untouch_nii([base_dir_sub fname '_slice' num2str(i3) '_denoised' cutoff_str 'WMcov.nii']);
    elseif stp==3
        scfMRItb_04_resplitData(base_dir_sub, fname, ['_denoised' cutoff_str 'cov'], siz3) % if slice-wise data doesn't exist but combined 4D data exists, then split the data to generate the needed slice-wise data
        scfMRItb_04_unzipFile(base_dir_sub, fname, ['_slice' num2str(i3) '_denoised' cutoff_str 'cov'])
        F1 = load_untouch_nii([base_dir_sub fname '_slice' num2str(i3) '_denoised' cutoff_str 'cov.nii']);
    elseif stp==2
        scfMRItb_04_resplitData(base_dir_sub, fname, '_WMcov', siz3) % if slice-wise data doesn't exist but combined 4D data exists, then split the data to generate the needed slice-wise data
        scfMRItb_04_unzipFile(base_dir_sub, fname, ['_slice' num2str(i3) '_WMcov'])
        F1 = load_untouch_nii([base_dir_sub fname '_slice' num2str(i3) '_WMcov.nii']);
    elseif stp==1
        scfMRItb_04_resplitData(base_dir_sub, fname, '_cov', siz3) % if slice-wise data doesn't exist but combined 4D data exists, then split the data to generate the needed slice-wise data
        scfMRItb_04_unzipFile(base_dir_sub, fname, ['_slice' num2str(i3) '_cov'])
        F1 = load_untouch_nii([base_dir_sub fname '_slice' num2str(i3) '_cov.nii']);
    end
    TR = F1.hdr.dime.pixdim(5); F1 = single(squeeze(F1.img));
    
    if Smask_err==0
        scfMRItb_04_resplitData(base_dir_sub, fname, '_Smask', siz3) % if slice-wise data doesn't exist but combined 4D data exists, then split the data to generate the needed slice-wise data
        scfMRItb_04_unzipFile(base_dir_sub, fname, ['_slice' num2str(i3) '_Smask'])
        Smask1 = load_untouch_nii([base_dir_sub fname '_slice' num2str(i3) '_Smask.nii']); Smask1 = Smask1.img;
    else
        Smask1 = Smask1_(:,:,i3);
    end

    for i5=1:size(F1,3)
        tmp = F1(:,:,i5);
        F2g(:,i5) = tmp(find(mask_GM(:,:,i3)==1));
        F2w(:,i5) = tmp(find(mask_WM(:,:,i3)==1));
        clear tmp
    end; clear i5 F1
    [~,svar1] = sort(var(F2g,0,2)); [~,sdist1] = sort(svar1); sortorder1{i3,1} = sdist1;
    [~,svar2] = sort(var(F2w,0,2)); [~,sdist2] = sort(svar2); sortorder2{i3,1} = sdist2;
    
    fighndl = figure('Position',[1 1 1152 720],'Color',[0.95,0.9,0.85],'InvertHardcopy','off','Visible','off'); % red [0.95,0.9,0.85], green [0.85,0.90,0.85], blue [0.89,0.94,0.99]
    subplot2n(2,1,1), imagesc(F2g(sortorder1{i3},:)),colormap(cmap),colorbar, xlabel('time (in samples)'),ylabel('Gray matter voxels (sorted by variance)'),
        title(['Time series within the GRAY MATTER after "14-DENOISE5-COVregress" in slice ' num2str(i3) ' of ' num2str(siz3)],'Color',[0,0.55,0.69],'FontSize',14);
    subplot2n(2,1,2), imagesc(F2w(sortorder2{i3},:)),colormap(cmap),colorbar, xlabel('time (in samples)'),ylabel('White matter voxels (sorted by variance)'),
        title(['Time series within the WHITE MATTER after "14-DENOISE5-COVregress" in slice ' num2str(i3) ' of ' num2str(siz3)],'Color',[0.65,0.11,0.19],'FontSize',14);
    saveas(fighndl,[QCpath 'fMRI_ts_14denoise5_slice' sprintf('%.2d',i3) '.jpg'])
    close(fighndl)
    
    scfMRItb_04_resplitData(base_dir_sub, fname, '', siz3) % if slice-wise data doesn't exist but combined 4D data exists, then split the data to generate the needed slice-wise data
	scfMRItb_04_unzipFile(base_dir_sub, fname, ['_slice' num2str(i3)])
    F3 = load_untouch_nii([base_dir_sub fname '_slice' num2str(i3) '.nii']); F3 = single(squeeze(F3.img));
    for i5=1:size(F3,3)
        tmp = F3(:,:,i5);
        F4(:,i5) = tmp(find(Smask1==1)); clear tmp
    end; clear i5 F3
    mF2g=mean(F2g,1); mF2g = 100*(mF2g-mean(mF2g))./mean(mF2g); mF2w=mean(F2w,1); mF2w = 100*(mF2w-mean(mF2w))./mean(mF2w);
    mF4=mean(F4,1); mF4 = 100*(mF4-mean(mF4))./mean(mF4);
    fighndl = figure('Position',[1 1 1152 720],'Color',[0.95,0.9,0.85],'InvertHardcopy','off','Visible','off'); % red [0.95,0.9,0.85], green [0.85,0.90,0.85], blue [0.89,0.94,0.99]
    subplot2n(3,1,1), plot(mF4,'color','k','linewidth',1.5), hold on, plot(mF2g,'color',[0,0.55,0.69],'linewidth',0.2), plot(mF2w,'color',[0.65,0.11,0.19],'linewidth',0.2), xlabel('time (in samples)'),ylabel('percentage signal change'),
      grid on,grid minor, xlim([1 length(mF2g)]),ylim([min(1.25.*[mF2g,mF2w,mF4]) 1.25.*max([mF2g,mF2w,mF4])]), title(['Global mean signal in UNPROCESSED RAW DATA (slice ' num2str(i3) ' of ' num2str(siz3) ')'],'Color','k','FontSize',14), legend({'Global signal (raw data)'},'FontSize',12,'TextColor','k','Location','best')
    subplot2n(3,1,2), plot(mF2g,'color',[0,0.55,0.69],'linewidth',1.5), hold on, plot(mF2w,'color',[0.65,0.11,0.19],'linewidth',0.2), plot(mF4,'color','k','linewidth',0.2), xlabel('time (in samples)'),ylabel('percentage signal change'),
      grid on,grid minor, xlim([1 length(mF2g)]),ylim([min(1.25.*[mF2g,mF2w,mF4]) 1.25.*max([mF2g,mF2w,mF4])]), title(['Gray matter mean signal AFTER "14-DENOISE5-COVregress" (slice ' num2str(i3) ' of ' num2str(siz3) ')'],'Color',[0,0.55,0.69],'FontSize',14), legend({'Gray matter signal after 14-denoise5'},'FontSize',12,'TextColor',[0,0.55,0.69],'Location','best')
    subplot2n(3,1,3), plot(mF2w,'color',[0.65,0.11,0.19],'linewidth',1.5), hold on, plot(mF4,'color','k','linewidth',0.2), plot(mF2g,'color',[0,0.55,0.69],'linewidth',0.2), xlabel('time (in samples)'),ylabel('percentage signal change'),
      grid on,grid minor, xlim([1 length(mF2g)]),ylim([min(1.25.*[mF2g,mF2w,mF4]) 1.25.*max([mF2g,mF2w,mF4])]), title(['White matter mean signal AFTER "14-DENOISE5-COVregress" (slice ' num2str(i3) ' of ' num2str(siz3) ')'],'Color',[0.65,0.11,0.19],'FontSize',14), legend({'White matter signal after 14-denoise5'},'FontSize',12,'TextColor',[0.65,0.11,0.19],'Location','best')
    saveas(fighndl,[QCpath 'global_signal_14denoise5_slice' sprintf('%.2d',i3) '.jpg'])
    close(fighndl)

    Cg = corrcoef(F2g'); Cg = uppertriangle(Cg);
    Cw = corrcoef(F2w'); Cw = uppertriangle(Cw);
    C2 = corrcoef(F4');  C2 = uppertriangle(C2);
    fighndl = figure('Position',[1 1 1152 720],'Color',[0.95,0.9,0.85],'InvertHardcopy','off','Visible','off'); % red [0.95,0.9,0.85], green [0.85,0.90,0.85], blue [0.89,0.94,0.99]
    subplot2n(3,1,1), histogram(C2,'EdgeColor','none','FaceColor','k','FaceAlpha',0.5,'Normalization','probability'); grid on,grid minor, xlim([-1 1]), hold on, xline(0,'Color','k','linewidth',1.5); hold off
      title(sprintf('Histogram of voxel-to-voxel correlations in UNPROCESSED RAW DATA (slice %d of %d)\n%.1f%% of the correlations were > 0  [ mean=%.2f, median=%.2f, SD=%.2f ]',i3,siz3,100*length(find(C2>0))/length(C2),mean(C2),median(C2),std(C2)),'Color','k','FontSize',14)
    subplot2n(3,1,2), histogram(Cg,'EdgeColor','none','FaceColor',[0,0.55,0.69],'FaceAlpha',0.5,'Normalization','probability'); grid on,grid minor, xlim([-1 1]), hold on, xline(0,'Color','k','linewidth',1.5); hold off
      title(sprintf('Histogram of voxel-to-voxel GRAY MATTER correlations AFTER 14-DENOISE5-COVregress (slice %d of %d)\n%.1f%% of the correlations were > 0  [ mean=%.2f, median=%.2f, SD=%.2f ]',i3,siz3,100*length(find(Cg>0))/length(Cg),mean(Cg),median(Cg),std(Cg)),'Color',[0,0.55,0.69],'FontSize',14)
    subplot2n(3,1,3), histogram(Cw,'EdgeColor','none','FaceColor',[0.65,0.11,0.19],'FaceAlpha',0.5,'Normalization','probability'); grid on,grid minor, xlim([-1 1]), hold on, xline(0,'Color','k','linewidth',1.5); hold off
      title(sprintf('Histogram of voxel-to-voxel WHITE MATTER correlations AFTER 14-DENOISE5-COVregress (slice %d of %d)\n%.1f%% of the correlations were > 0  [ mean=%.2f, median=%.2f, SD=%.2f ]',i3,siz3,100*length(find(Cw>0))/length(Cw),mean(Cw),median(Cw),std(Cw)),'Color',[0.65,0.11,0.19],'FontSize',14)
    saveas(fighndl,[QCpath 'hist_FC_compare_14denoise5_slice' sprintf('%.2d',i3) '.jpg'])
    close(fighndl)
    
    if stp~=0
        if stp==1
            scfMRItb_04_resplitData(base_dir_sub, fname, '_warped', siz3) % if slice-wise data doesn't exist but combined 4D data exists, then split the data to generate the needed slice-wise data
            scfMRItb_04_unzipFile(base_dir_sub, fname, ['_slice' num2str(i3) '_warped'])
            F5 = load_untouch_nii([base_dir_sub fname '_slice' num2str(i3) '_warped.nii']); F5 = single(squeeze(F5.img));
        elseif stp==2
            scfMRItb_04_resplitData(base_dir_sub, fname, '_WM', siz3) % if slice-wise data doesn't exist but combined 4D data exists, then split the data to generate the needed slice-wise data
            scfMRItb_04_unzipFile(base_dir_sub, fname, ['_slice' num2str(i3) '_WM'])
            F5 = load_untouch_nii([base_dir_sub fname '_slice' num2str(i3) '_WM.nii']); F5 = single(squeeze(F5.img));
        elseif stp==3
            scfMRItb_04_resplitData(base_dir_sub, fname, ['_denoised' cutoff_str], siz3) % if slice-wise data doesn't exist but combined 4D data exists, then split the data to generate the needed slice-wise data
            scfMRItb_04_unzipFile(base_dir_sub, fname, ['_slice' num2str(i3) '_denoised' cutoff_str ''])
            F5 = load_untouch_nii([base_dir_sub fname '_slice' num2str(i3) '_denoised' cutoff_str '.nii']); F5 = single(squeeze(F5.img));
        elseif stp==4
            scfMRItb_04_resplitData(base_dir_sub, fname, ['_denoised' cutoff_str 'WM'], siz3) % if slice-wise data doesn't exist but combined 4D data exists, then split the data to generate the needed slice-wise data
            scfMRItb_04_unzipFile(base_dir_sub, fname, ['_slice' num2str(i3) '_denoised' cutoff_str 'WM'])
            F5 = load_untouch_nii([base_dir_sub fname '_slice' num2str(i3) '_denoised' cutoff_str 'WM.nii']); F5 = single(squeeze(F5.img));
        end
        for i5=1:size(F5,3)
            tmp = F5(:,:,i5);
            F6g(:,i5) = tmp(find(mask_GM(:,:,i3)==1));
            F6w(:,i5) = tmp(find(mask_WM(:,:,i3)==1));
            clear tmp
        end; clear i5 F5
        mF6g=mean(F6g,1); mF6g = 100*(mF6g-mean(mF6g))./mean(mF6g); mF6w=mean(F6w,1); mF6w = 100*(mF6w-mean(mF6w))./mean(mF6w);
        fighndl = figure('Position',[1 1 1152 720],'Color',[0.95,0.9,0.85],'InvertHardcopy','off','Visible','off'); % red [0.95,0.9,0.85], green [0.85,0.90,0.85], blue [0.89,0.94,0.99]
        if stp==1
            subplot2n(4,1,1), plot(mF6g,'color','k','linewidth',1.5), hold on, plot(mF2g,'color',[0,0.55,0.69],'linewidth',0.2), ylabel('pcent sig change'),
                grid on,grid minor, xlim([1 length(mF2g)]),ylim([min(1.25.*[mF2g,mF6g]) 1.25.*max([mF2g,mF6g])]), title(['Gray matter mean signal AFTER "11-func-anat-registration" (slice ' num2str(i3) ' of ' num2str(siz3) ')'],'Color','k','FontSize',14), legend({'Gray matter signal before COV-denoising'},'FontSize',12,'TextColor','k','Location','best')
            subplot2n(4,1,3), plot(mF6w,'color','k','linewidth',1.5), hold on, plot(mF2w,'color',[0.65,0.11,0.19],'linewidth',0.2), ylabel('pcent sig change'),
                grid on,grid minor, xlim([1 length(mF2g)]),ylim([min(1.25.*[mF2w,mF6w]) 1.25.*max([mF2w,mF6w])]), title(['White matter mean signal AFTER "11-func-anat-registration" (slice ' num2str(i3) ' of ' num2str(siz3) ')'],'Color','k','FontSize',14), legend({'White matter signal before COV-denoising'},'FontSize',12,'TextColor','k','Location','best')
        elseif stp==2
            subplot2n(4,1,1), plot(mF6g,'color','k','linewidth',1.5), hold on, plot(mF2g,'color',[0,0.55,0.69],'linewidth',0.2), ylabel('pcent sig change'),
                grid on,grid minor, xlim([1 length(mF2g)]),ylim([min(1.25.*[mF2g,mF6g]) 1.25.*max([mF2g,mF6g])]), title(['Gray matter mean signal AFTER "13-denoise4-WMregress" (slice ' num2str(i3) ' of ' num2str(siz3) ')'],'Color','k','FontSize',14), legend({'Gray matter signal before COV-denoising'},'FontSize',12,'TextColor','k','Location','best')
            subplot2n(4,1,3), plot(mF6w,'color','k','linewidth',1.5), hold on, plot(mF2w,'color',[0.65,0.11,0.19],'linewidth',0.2), ylabel('pcent sig change'),
                grid on,grid minor, xlim([1 length(mF2g)]),ylim([min(1.25.*[mF2w,mF6w]) 1.25.*max([mF2w,mF6w])]), title(['White matter mean signal AFTER "13-denoise4-WMregress" (slice ' num2str(i3) ' of ' num2str(siz3) ')'],'Color','k','FontSize',14), legend({'White matter signal before COV-denoising'},'FontSize',12,'TextColor','k','Location','best')
        elseif stp==3
            subplot2n(4,1,1), plot(mF6g,'color','k','linewidth',1.5), hold on, plot(mF2g,'color',[0,0.55,0.69],'linewidth',0.2), ylabel('pcent sig change'),
                grid on,grid minor, xlim([1 length(mF2g)]),ylim([min(1.25.*[mF2g,mF6g]) 1.25.*max([mF2g,mF6g])]), title(['Gray matter mean signal AFTER "12-denoise3-CSFregress" (slice ' num2str(i3) ' of ' num2str(siz3) ')'],'Color','k','FontSize',14), legend({'Gray matter signal before COV-denoising'},'FontSize',12,'TextColor','k','Location','best')
            subplot2n(4,1,3), plot(mF6w,'color','k','linewidth',1.5), hold on, plot(mF2w,'color',[0.65,0.11,0.19],'linewidth',0.2), ylabel('pcent sig change'),
                grid on,grid minor, xlim([1 length(mF2g)]),ylim([min(1.25.*[mF2w,mF6w]) 1.25.*max([mF2w,mF6w])]), title(['White matter mean signal AFTER "12-denoise3-CSFregress" (slice ' num2str(i3) ' of ' num2str(siz3) ')'],'Color','k','FontSize',14), legend({'White matter signal before COV-denoising'},'FontSize',12,'TextColor','k','Location','best')
        elseif stp==4
            subplot2n(4,1,1), plot(mF6g,'color','k','linewidth',1.5), hold on, plot(mF2g,'color',[0,0.55,0.69],'linewidth',0.2), ylabel('pcent sig change'),
                grid on,grid minor, xlim([1 length(mF2g)]),ylim([min(1.25.*[mF2g,mF6g]) 1.25.*max([mF2g,mF6g])]), title(['Gray matter mean signal AFTER "13-denoise4-WMregress" (slice ' num2str(i3) ' of ' num2str(siz3) ')'],'Color','k','FontSize',14), legend({'Gray matter signal before COV-denoising'},'FontSize',12,'TextColor','k','Location','best')
            subplot2n(4,1,3), plot(mF6w,'color','k','linewidth',1.5), hold on, plot(mF2w,'color',[0.65,0.11,0.19],'linewidth',0.2), ylabel('pcent sig change'),
                grid on,grid minor, xlim([1 length(mF2g)]),ylim([min(1.25.*[mF2w,mF6w]) 1.25.*max([mF2w,mF6w])]), title(['White matter mean signal AFTER "13-denoise4-WMregress" (slice ' num2str(i3) ' of ' num2str(siz3) ')'],'Color','k','FontSize',14), legend({'White matter signal before COV-denoising'},'FontSize',12,'TextColor','k','Location','best')
        end
        subplot2n(4,1,2), plot(mF2g,'color',[0,0.55,0.69],'linewidth',1.5), hold on, plot(mF6g,'color','k','linewidth',0.2), ylabel('pcent sig change'),
            grid on,grid minor, xlim([1 length(mF2g)]),ylim([min(1.25.*[mF2g,mF6g]) 1.25.*max([mF2g,mF6g])]), title(['Gray matter mean signal AFTER "14-DENOISE5-COVregress" (slice ' num2str(i3) ' of ' num2str(siz3) ')'],'Color',[0,0.55,0.69],'FontSize',14), legend({'Gray matter signal after COV-denoising'},'FontSize',12,'TextColor',[0,0.55,0.69],'Location','best')
        subplot2n(4,1,4), plot(mF2w,'color',[0.65,0.11,0.19],'linewidth',1.5), hold on, plot(mF6w,'color','k','linewidth',0.2), ylabel('pcent sig change'),
            grid on,grid minor, xlim([1 length(mF2g)]),ylim([min(1.25.*[mF2w,mF6w]) 1.25.*max([mF2w,mF6w])]), title(['White matter mean signal AFTER "14-DENOISE5-COVregress" (slice ' num2str(i3) ' of ' num2str(siz3) ')'],'Color',[0.65,0.11,0.19],'FontSize',14), legend({'White matter signal after COV-denoising'},'FontSize',12,'TextColor',[0.65,0.11,0.19],'Location','best')
        saveas(fighndl,[QCpath 'global_signal_13denoise4_vs_14denoise5_slice' sprintf('%.2d',i3) '.jpg'])
        close(fighndl)
        
        C3g = corrcoef(F6g'); C3g = uppertriangle(C3g); C3w = corrcoef(F6w'); C3w = uppertriangle(C3w);
        fighndl = figure('Position',[1 1 1152 720],'Color',[0.95,0.9,0.85],'InvertHardcopy','off','Visible','off'); % red [0.95,0.9,0.85], green [0.85,0.90,0.85], blue [0.89,0.94,0.99]
        if stp==1
            subplot2n(4,1,1), histogram(C3g,'EdgeColor','none','FaceColor','k','FaceAlpha',0.5,'Normalization','probability'); grid on,grid minor, xlim([-1 1]), hold on, xline(0,'Color','k','linewidth',1.5); hold off
                title(sprintf('Histogram of voxel-to-voxel GRAY MATTER correlations AFTER 11-func-anat-registration (slice %d of %d)\n%.1f%% of the correlations were > 0  [ mean=%.2f, median=%.2f, SD=%.2f ]',i3,siz3,100*length(find(C3g>0))/length(C3g),mean(C3g),median(C3g),std(C3g)),'Color','k','FontSize',14)
            subplot2n(4,1,3), histogram(C3w,'EdgeColor','none','FaceColor','k','FaceAlpha',0.5,'Normalization','probability'); grid on,grid minor, xlim([-1 1]), hold on, xline(0,'Color','k','linewidth',1.5); hold off
                title(sprintf('WHITE MATTER: AFTER 11-func-anat-registration (slice %d of %d): %.1f%% of the correlations were > 0  [ mean=%.2f, median=%.2f, SD=%.2f ]',i3,siz3,100*length(find(C3w>0))/length(C3w),mean(C3w),median(C3w),std(C3w)),'Color','k','FontSize',14)
        elseif stp==2
            subplot2n(4,1,1), histogram(C3g,'EdgeColor','none','FaceColor','k','FaceAlpha',0.5,'Normalization','probability'); grid on,grid minor, xlim([-1 1]), hold on, xline(0,'Color','k','linewidth',1.5); hold off
                title(sprintf('Histogram of voxel-to-voxel GRAY MATTER correlations AFTER 13-denoise4-WMregress (slice %d of %d)\n%.1f%% of the correlations were > 0  [ mean=%.2f, median=%.2f, SD=%.2f ]',i3,siz3,100*length(find(C3g>0))/length(C3g),mean(C3g),median(C3g),std(C3g)),'Color','k','FontSize',14)
            subplot2n(4,1,3), histogram(C3w,'EdgeColor','none','FaceColor','k','FaceAlpha',0.5,'Normalization','probability'); grid on,grid minor, xlim([-1 1]), hold on, xline(0,'Color','k','linewidth',1.5); hold off
                title(sprintf('WHITE MATTER: AFTER 13-denoise4-WMregress (slice %d of %d): %.1f%% of the correlations were > 0  [ mean=%.2f, median=%.2f, SD=%.2f ]',i3,siz3,100*length(find(C3w>0))/length(C3w),mean(C3w),median(C3w),std(C3w)),'Color','k','FontSize',14)
        elseif stp==3
            subplot2n(4,1,1), histogram(C3g,'EdgeColor','none','FaceColor','k','FaceAlpha',0.5,'Normalization','probability'); grid on,grid minor, xlim([-1 1]), hold on, xline(0,'Color','k','linewidth',1.5); hold off
                title(sprintf('Histogram of voxel-to-voxel GRAY MATTER correlations AFTER 12-denoise3-CSFregress (slice %d of %d)\n%.1f%% of the correlations were > 0  [ mean=%.2f, median=%.2f, SD=%.2f ]',i3,siz3,100*length(find(C3g>0))/length(C3g),mean(C3g),median(C3g),std(C3g)),'Color','k','FontSize',14)
            subplot2n(4,1,3), histogram(C3w,'EdgeColor','none','FaceColor','k','FaceAlpha',0.5,'Normalization','probability'); grid on,grid minor, xlim([-1 1]), hold on, xline(0,'Color','k','linewidth',1.5); hold off
                title(sprintf('WHITE MATTER: AFTER 12-denoise3-CSFregress (slice %d of %d): %.1f%% of the correlations were > 0  [ mean=%.2f, median=%.2f, SD=%.2f ]',i3,siz3,100*length(find(C3w>0))/length(C3w),mean(C3w),median(C3w),std(C3w)),'Color','k','FontSize',14)
        elseif stp==4
            subplot2n(4,1,1), histogram(C3g,'EdgeColor','none','FaceColor','k','FaceAlpha',0.5,'Normalization','probability'); grid on,grid minor, xlim([-1 1]), hold on, xline(0,'Color','k','linewidth',1.5); hold off
                title(sprintf('Histogram of voxel-to-voxel GRAY MATTER correlations AFTER 13-denoise4-WMregress (slice %d of %d)\n%.1f%% of the correlations were > 0  [ mean=%.2f, median=%.2f, SD=%.2f ]',i3,siz3,100*length(find(C3g>0))/length(C3g),mean(C3g),median(C3g),std(C3g)),'Color','k','FontSize',14)
            subplot2n(4,1,3), histogram(C3w,'EdgeColor','none','FaceColor','k','FaceAlpha',0.5,'Normalization','probability'); grid on,grid minor, xlim([-1 1]), hold on, xline(0,'Color','k','linewidth',1.5); hold off
                title(sprintf('WHITE MATTER: AFTER 13-denoise4-WMregress (slice %d of %d): %.1f%% of the correlations were > 0  [ mean=%.2f, median=%.2f, SD=%.2f ]',i3,siz3,100*length(find(C3w>0))/length(C3w),mean(C3w),median(C3w),std(C3w)),'Color','k','FontSize',14)
        end
        subplot2n(4,1,2), histogram(Cg,'EdgeColor','none','FaceColor',[0,0.55,0.69],'FaceAlpha',0.5,'Normalization','probability'); grid on,grid minor, xlim([-1 1]), hold on, xline(0,'Color','k','linewidth',1.5); hold off
            title(sprintf('GRAY MATTER: AFTER 14-DENOISE5-COVregress (slice %d of %d): %.1f%% of the correlations were > 0  [ mean=%.2f, median=%.2f, SD=%.2f ]',i3,siz3,100*length(find(Cg>0))/length(Cg),mean(Cg),median(Cg),std(Cg)),'Color',[0,0.55,0.69],'FontSize',14)
        subplot2n(4,1,4), histogram(Cw,'EdgeColor','none','FaceColor',[0.65,0.11,0.19],'FaceAlpha',0.5,'Normalization','probability'); grid on,grid minor, xlim([-1 1]), hold on, xline(0,'Color','k','linewidth',1.5); hold off
            title(sprintf('WHITE MATTER: AFTER 14-DENOISE5-COVregress (slice %d of %d): %.1f%% of the correlations were > 0  [ mean=%.2f, median=%.2f, SD=%.2f ]',i3,siz3,100*length(find(Cw>0))/length(Cw),mean(Cw),median(Cw),std(Cw)),'Color',[0.65,0.11,0.19],'FontSize',14)
        saveas(fighndl,[QCpath 'hist_FC_compare_13denoise4_14denoise5_slice' sprintf('%.2d',i3) '.jpg'])
        close(fighndl)
        F6g_=cat(1,F6g_,F6g); mF6g_=cat(1,mF6g_,mF6g); C3g_=cat(1,C3g_,C3g);
        F6w_=cat(1,F6w_,F6w); mF6w_=cat(1,mF6w_,mF6w); C3w_=cat(1,C3w_,C3w);
        clear F6g F6w mF6g mF6w C3g C3w
    end
    
    F2g_=cat(1,F2g_,F2g); mF2g_=cat(1,mF2g_,mF2g); F2w_=cat(1,F2w_,F2w); mF2w_=cat(1,mF2w_,mF2w); Cg_=cat(1,Cg_,Cg); Cw_=cat(1,Cw_,Cw); F4_=cat(1,F4_,F4); mF4_=cat(1,mF4_,mF4); C2_=cat(1,C2_,C2);
    clear F2g F2w F4 TR Smask1 svar1 svar2 mF2g mF2w mF4 Cg Cw C2
end


[~,svar1] = sort(var(F2g_,0,2)); [~,spsc1] = sort(meanPSC(F2g_));
[~,svar2] = sort(var(F2w_,0,2)); [~,spsc2] = sort(meanPSC(F2w_));
sortorder1b = sortorder1; sortorder1=[]; f1=0; %#ok<*ASGLU>
sortorder2b = sortorder2; sortorder2=[]; f2=0; %#ok<*ASGLU>
for im=1:length(sortorder1b)
    temp = sortorder1b{im}; temp = temp + f1; f1 = f1 + length(temp);
    sortorder1 = cat(1,sortorder1,temp); clear temp
    temp = sortorder2b{im}; temp = temp + f2; f2 = f2 + length(temp);
    sortorder2 = cat(1,sortorder2,temp); clear temp
end; clear im f1 f2
sortorder1 = spsc1; % options: spsc for percentage signal change, svar for variance, (comment out the line if you want to use distance from the center of the cord separately for each slice)
sortorder2 = spsc2; % options: spsc for percentage signal change, svar for variance, (comment out the line if you want to use distance from the center of the cord separately for each slice)
for im=max(sortorder1):-1:1, if isempty(find(sortorder1==im)), sortorder1(find(sortorder1>im)) = sortorder1(find(sortorder1>im))-1; end, end; clear im %#ok<*EFIND>
for im=max(sortorder2):-1:1, if isempty(find(sortorder2==im)), sortorder2(find(sortorder2>im)) = sortorder2(find(sortorder2>im))-1; end, end; clear im %#ok<*EFIND>

fighndl = figure('Position',[1 1 1152 720],'Color',[0.89,0.94,0.99],'InvertHardcopy','off','Visible','off'); % red [0.95,0.9,0.85], green [0.85,0.90,0.85], blue [0.89,0.94,0.99]
subplot2n(2,1,1), imagesc(F2g_(sortorder1,:)),colormap(cmap),colorbar, xlabel('time (in samples)'),ylabel(sprintf('voxels (sorted by mean percentage signal change;\nbottom row is highest)')),
  title(['Time series within the GRAY MATTER AFTER 14-DENOISE5-COVregress (all slices combined)'],'Color',[0,0.55,0.69],'FontSize',14);
subplot2n(2,1,2), imagesc(F2w_(sortorder2,:)),colormap(cmap),colorbar, xlabel('time (in samples)'),ylabel(sprintf('voxels (sorted by mean percentage signal change;\nbottom row is highest)')),
  title(['Time series within the WHITE MATTER AFTER 14-DENOISE5-COVregress (all slices combined)'],'Color',[0.65,0.11,0.19],'FontSize',14);
saveas(fighndl,[QCpath 'fMRI_ts_14denoise5_allSlices' '.jpg'])
saveas(fighndl,[QCpath2 'fMRI_ts_14denoise5_allSlices' '.jpg'])
 close(fighndl)
mF2g_=mean(mF2g_,1); mF2w_=mean(mF2w_,1); mF4_=mean(mF4_,1);
fighndl = figure('Position',[1 1 1152 720],'Color',[0.89,0.94,0.99],'InvertHardcopy','off','Visible','off'); % red [0.95,0.9,0.85], green [0.85,0.90,0.85], blue [0.89,0.94,0.99]
subplot2n(3,1,1), plot(mF4_,'color','k','linewidth',1.5), hold on, plot(mF2g_,'color',[0,0.55,0.69],'linewidth',0.2), plot(mF2w_,'color',[0.65,0.11,0.19],'linewidth',0.2), xlabel('time (in samples)'),ylabel('percentage signal change'),
    grid on,grid minor, xlim([1 length(mF2g_)]),ylim([1.25.*min([mF2g_,mF2w_,mF4_]) 1.25.*max([mF2g_,mF2w_,mF4_])]), title(['Global mean signal in UNPROCESSED RAW DATA (all slices averaged)'],'Color','k','FontSize',14), legend({'Global signal (raw data)'},'FontSize',12,'TextColor','k','Location','best')
subplot2n(3,1,2), plot(mF2g_,'color',[0,0.55,0.69],'linewidth',1.5), hold on, plot(mF2w_,'color',[0.65,0.11,0.19],'linewidth',0.2), plot(mF4_,'color','k','linewidth',0.2), xlabel('time (in samples)'),ylabel('percentage signal change'),
    grid on,grid minor, xlim([1 length(mF2g_)]),ylim([1.25.*min([mF2g_,mF2w_,mF4_]) 1.25.*max([mF2g_,mF2w_,mF4_])]), title(['Gray matter mean signal AFTER "14-DENOISE5-COVregress" (all slices averaged)'],'Color',[0,0.55,0.69],'FontSize',14), legend({'Gray matter signal after 14-denoise5'},'FontSize',12,'TextColor',[0,0.55,0.69],'Location','best')
subplot2n(3,1,3), plot(mF2w_,'color',[0.65,0.11,0.19],'linewidth',1.5), hold on, plot(mF4_,'color','k','linewidth',0.2), plot(mF2g_,'color',[0,0.55,0.69],'linewidth',0.2), xlabel('time (in samples)'),ylabel('percentage signal change'),
    grid on,grid minor, xlim([1 length(mF2g_)]),ylim([1.25.*min([mF2g_,mF2w_,mF4_]) 1.25.*max([mF2g_,mF2w_,mF4_])]), title(['White matter mean signal AFTER "14-DENOISE5-COVregress" (all slices averaged)'],'Color',[0.65,0.11,0.19],'FontSize',14), legend({'White matter signal after 14-denoise5'},'FontSize',12,'TextColor',[0.65,0.11,0.19],'Location','best')
saveas(fighndl,[QCpath 'global_signal_14denoise5_allSlices' '.jpg'])
 close(fighndl)
fighndl = figure('Position',[1 1 1152 720],'Color',[0.89,0.94,0.99],'InvertHardcopy','off','Visible','off'); % red [0.95,0.9,0.85], green [0.85,0.90,0.85], blue [0.89,0.94,0.99]
subplot2n(3,1,1), histogram(C2_,'EdgeColor','none','FaceColor','k','FaceAlpha',0.5,'Normalization','probability'); grid on,grid minor, xlim([-1 1]), hold on, xline(0,'Color','k','linewidth',1.5); hold off
    title(sprintf('Histogram of voxel-to-voxel correlations in UNPROCESSED RAW DATA (all slices combined)\n%.1f%% of the correlations were > 0  [ mean=%.2f, median=%.2f, SD=%.2f ]',100*length(find(C2_>0))/length(C2_),mean(C2_),median(C2_),std(C2_)),'Color','k','FontSize',14)
subplot2n(3,1,2), histogram(Cg_,'EdgeColor','none','FaceColor',[0,0.55,0.69],'FaceAlpha',0.5,'Normalization','probability'); grid on,grid minor, xlim([-1 1]), hold on, xline(0,'Color','k','linewidth',1.5); hold off
    title(sprintf('Histogram of voxel-to-voxel GRAY MATTER correlations AFTER 14-DENOISE5-COVregress (all slices combined)\n%.1f%% of the correlations were > 0  [ mean=%.2f, median=%.2f, SD=%.2f ]',100*length(find(Cg_>0))/length(Cg_),mean(Cg_),median(Cg_),std(Cg_)),'Color',[0,0.55,0.69],'FontSize',14)
subplot2n(3,1,3), histogram(Cw_,'EdgeColor','none','FaceColor',[0.65,0.11,0.19],'FaceAlpha',0.5,'Normalization','probability'); grid on,grid minor, xlim([-1 1]), hold on, xline(0,'Color','k','linewidth',1.5); hold off
    title(sprintf('Histogram of voxel-to-voxel WHITE MATTER correlations AFTER 14-DENOISE5-COVregress (all slices combined)\n%.1f%% of the correlations were > 0  [ mean=%.2f, median=%.2f, SD=%.2f ]',100*length(find(Cw_>0))/length(Cw_),mean(Cw_),median(Cw_),std(Cw_)),'Color',[0.65,0.11,0.19],'FontSize',14)
saveas(fighndl,[QCpath 'hist_FC_compare_14denoise5_allSlices' '.jpg'])
 close(fighndl)

if stp~=0
    mF6g_=mean(mF6g_,1); mF6w_=mean(mF6w_,1);
    fighndl = figure('Position',[1 1 1152 720],'Color',[0.89,0.94,0.99],'InvertHardcopy','off','Visible','off'); % red [0.95,0.9,0.85], green [0.85,0.90,0.85], blue [0.89,0.94,0.99]
    if stp==1
    	subplot2n(4,1,1), plot(mF6g_,'color','k','linewidth',1.5), hold on, plot(mF2g_,'color',[0,0.55,0.69],'linewidth',0.2), ylabel('pcent sig change'),
            grid on,grid minor, xlim([1 length(mF2g_)]),ylim([1.25.*min([mF2g_,mF6g_]) 1.25.*max([mF2g_,mF6g_])]), title(['Gray matter mean signal AFTER "11-func-anat-registration" (all slices averaged)'],'Color','k','FontSize',14), legend({'Gray matter signal before COV-denoising'},'FontSize',12,'TextColor','k','Location','best')
        subplot2n(4,1,3), plot(mF6w_,'color','k','linewidth',1.5), hold on, plot(mF2w_,'color',[0.65,0.11,0.19],'linewidth',0.2), ylabel('pcent sig change'),
            grid on,grid minor, xlim([1 length(mF2g_)]),ylim([1.25.*min([mF2w_,mF6w_]) 1.25.*max([mF2w_,mF6w_])]), title(['White matter mean signal AFTER "11-func-anat-registration" (all slices averaged)'],'Color','k','FontSize',14), legend({'White matter signal before COV-denoising'},'FontSize',12,'TextColor','k','Location','best')
    elseif stp==2
    	subplot2n(4,1,1), plot(mF6g_,'color','k','linewidth',1.5), hold on, plot(mF2g_,'color',[0,0.55,0.69],'linewidth',0.2), ylabel('pcent sig change'),
            grid on,grid minor, xlim([1 length(mF2g_)]),ylim([1.25.*min([mF2g_,mF6g_]) 1.25.*max([mF2g_,mF6g_])]), title(['Gray matter mean signal AFTER "13-DENOISE4-WMregress" (all slices averaged)'],'Color','k','FontSize',14), legend({'Gray matter signal before COV-denoising'},'FontSize',12,'TextColor','k','Location','best')
        subplot2n(4,1,3), plot(mF6w_,'color','k','linewidth',1.5), hold on, plot(mF2w_,'color',[0.65,0.11,0.19],'linewidth',0.2), ylabel('pcent sig change'),
            grid on,grid minor, xlim([1 length(mF2g_)]),ylim([1.25.*min([mF2w_,mF6w_]) 1.25.*max([mF2w_,mF6w_])]), title(['White matter mean signal AFTER "13-DENOISE4-WMregress" (all slices averaged)'],'Color','k','FontSize',14), legend({'White matter signal before COV-denoising'},'FontSize',12,'TextColor','k','Location','best')
    elseif stp==3
    	subplot2n(4,1,1), plot(mF6g_,'color','k','linewidth',1.5), hold on, plot(mF2g_,'color',[0,0.55,0.69],'linewidth',0.2), ylabel('pcent sig change'),
            grid on,grid minor, xlim([1 length(mF2g_)]),ylim([1.25.*min([mF2g_,mF6g_]) 1.25.*max([mF2g_,mF6g_])]), title(['Gray matter mean signal AFTER "12-DENOISE3-CSFregress" (all slices averaged)'],'Color','k','FontSize',14), legend({'Gray matter signal before COV-denoising'},'FontSize',12,'TextColor','k','Location','best')
        subplot2n(4,1,3), plot(mF6w_,'color','k','linewidth',1.5), hold on, plot(mF2w_,'color',[0.65,0.11,0.19],'linewidth',0.2), ylabel('pcent sig change'),
            grid on,grid minor, xlim([1 length(mF2g_)]),ylim([1.25.*min([mF2w_,mF6w_]) 1.25.*max([mF2w_,mF6w_])]), title(['White matter mean signal AFTER "12-DENOISE3-CSFregress" (all slices averaged)'],'Color','k','FontSize',14), legend({'White matter signal before COV-denoising'},'FontSize',12,'TextColor','k','Location','best')
    elseif stp==4
    	subplot2n(4,1,1), plot(mF6g_,'color','k','linewidth',1.5), hold on, plot(mF2g_,'color',[0,0.55,0.69],'linewidth',0.2), ylabel('pcent sig change'),
            grid on,grid minor, xlim([1 length(mF2g_)]),ylim([1.25.*min([mF2g_,mF6g_]) 1.25.*max([mF2g_,mF6g_])]), title(['Gray matter mean signal AFTER "13-DENOISE4-WMregress" (all slices averaged)'],'Color','k','FontSize',14), legend({'Gray matter signal before COV-denoising'},'FontSize',12,'TextColor','k','Location','best')
        subplot2n(4,1,3), plot(mF6w_,'color','k','linewidth',1.5), hold on, plot(mF2w_,'color',[0.65,0.11,0.19],'linewidth',0.2), ylabel('pcent sig change'),
            grid on,grid minor, xlim([1 length(mF2g_)]),ylim([1.25.*min([mF2w_,mF6w_]) 1.25.*max([mF2w_,mF6w_])]), title(['White matter mean signal AFTER "13-DENOISE4-WMregress" (all slices averaged)'],'Color','k','FontSize',14), legend({'White matter signal before COV-denoising'},'FontSize',12,'TextColor','k','Location','best')
    end
    subplot2n(4,1,2), plot(mF2g_,'color',[0,0.55,0.69],'linewidth',1.5), hold on, plot(mF6g_,'color','k','linewidth',0.2), ylabel('pcent sig change'),
        grid on,grid minor, xlim([1 length(mF2g_)]),ylim([1.25.*min([mF2g_,mF6g_]) 1.25.*max([mF2g_,mF6g_])]), title(['Gray matter mean signal AFTER "14-DENOISE5-COVregress" (all slices averaged)'],'Color',[0,0.55,0.69],'FontSize',14), legend({'Gray matter signal after COV-denoising'},'FontSize',12,'TextColor',[0,0.55,0.69],'Location','best')
    subplot2n(4,1,4), plot(mF2w_,'color',[0.65,0.11,0.19],'linewidth',1.5), hold on, plot(mF6w_,'color','k','linewidth',0.2), ylabel('pcent sig change'),
        grid on,grid minor, xlim([1 length(mF2g_)]),ylim([1.25.*min([mF2w_,mF6w_]) 1.25.*max([mF2w_,mF6w_])]), title(['White matter mean signal AFTER "14-DENOISE5-COVregress" (all slices averaged)'],'Color',[0.65,0.11,0.19],'FontSize',14), legend({'White matter signal after COV-denoising'},'FontSize',12,'TextColor',[0.65,0.11,0.19],'Location','best')
    saveas(fighndl,[QCpath 'global_signal_13denoise4_vs_14denoise5_allSlices' '.jpg'])
    saveas(fighndl,[QCpath2 'global_signal_13denoise4_vs_14denoise5_allSlices' '.jpg'])
     close(fighndl)
    
    fighndl = figure('Position',[1 1 1152 720],'Color',[0.89,0.94,0.99],'InvertHardcopy','off','Visible','off'); % red [0.95,0.9,0.85], green [0.85,0.90,0.85], blue [0.89,0.94,0.99]
    if stp==1
    	subplot2n(4,1,1), histogram(C3g_,'EdgeColor','none','FaceColor','k','FaceAlpha',0.5,'Normalization','probability'); grid on,grid minor, xlim([-1 1]), hold on, xline(0,'Color','k','linewidth',1.5); hold off
            title(sprintf('Histogram of voxel-to-voxel GRAY MATTER correlations AFTER 11-registration (all slices combined)\n%.1f%% of the correlations were > 0  [ mean=%.2f, median=%.2f, SD=%.2f ]',100*length(find(C3g_>0))/length(C3g_),mean(C3g_),median(C3g_),std(C3g_)),'Color','k','FontSize',14)
        subplot2n(4,1,3), histogram(C3w_,'EdgeColor','none','FaceColor','k','FaceAlpha',0.5,'Normalization','probability'); grid on,grid minor, xlim([-1 1]), hold on, xline(0,'Color','k','linewidth',1.5); hold off
            title(sprintf('WHITE MATTER: AFTER 11-registration (all slices): %.1f%% of the correlations were > 0  [ mean=%.2f, median=%.2f, SD=%.2f ]',100*length(find(C3w_>0))/length(C3w_),mean(C3w_),median(C3w_),std(C3w_)),'Color','k','FontSize',14)
    elseif stp==2
    	subplot2n(4,1,1), histogram(C3g_,'EdgeColor','none','FaceColor','k','FaceAlpha',0.5,'Normalization','probability'); grid on,grid minor, xlim([-1 1]), hold on, xline(0,'Color','k','linewidth',1.5); hold off
            title(sprintf('Histogram of voxel-to-voxel GRAY MATTER correlations AFTER 13-denoise4-WMregress (all slices combined)\n%.1f%% of the correlations were > 0  [ mean=%.2f, median=%.2f, SD=%.2f ]',100*length(find(C3g_>0))/length(C3g_),mean(C3g_),median(C3g_),std(C3g_)),'Color','k','FontSize',14)
        subplot2n(4,1,3), histogram(C3w_,'EdgeColor','none','FaceColor','k','FaceAlpha',0.5,'Normalization','probability'); grid on,grid minor, xlim([-1 1]), hold on, xline(0,'Color','k','linewidth',1.5); hold off
            title(sprintf('WHITE MATTER: AFTER 13-denoise4-WMregress (all slices): %.1f%% of the correlations were > 0  [ mean=%.2f, median=%.2f, SD=%.2f ]',100*length(find(C3w_>0))/length(C3w_),mean(C3w_),median(C3w_),std(C3w_)),'Color','k','FontSize',14)
    elseif stp==3
    	subplot2n(4,1,1), histogram(C3g_,'EdgeColor','none','FaceColor','k','FaceAlpha',0.5,'Normalization','probability'); grid on,grid minor, xlim([-1 1]), hold on, xline(0,'Color','k','linewidth',1.5); hold off
            title(sprintf('Histogram of voxel-to-voxel GRAY MATTER correlations AFTER 12-denoise3-CSFregress (all slices combined)\n%.1f%% of the correlations were > 0  [ mean=%.2f, median=%.2f, SD=%.2f ]',100*length(find(C3g_>0))/length(C3g_),mean(C3g_),median(C3g_),std(C3g_)),'Color','k','FontSize',14)
        subplot2n(4,1,3), histogram(C3w_,'EdgeColor','none','FaceColor','k','FaceAlpha',0.5,'Normalization','probability'); grid on,grid minor, xlim([-1 1]), hold on, xline(0,'Color','k','linewidth',1.5); hold off
            title(sprintf('WHITE MATTER: AFTER 12-denoise3-CSFregress (all slices): %.1f%% of the correlations were > 0  [ mean=%.2f, median=%.2f, SD=%.2f ]',100*length(find(C3w_>0))/length(C3w_),mean(C3w_),median(C3w_),std(C3w_)),'Color','k','FontSize',14)
    elseif stp==4
    	subplot2n(4,1,1), histogram(C3g_,'EdgeColor','none','FaceColor','k','FaceAlpha',0.5,'Normalization','probability'); grid on,grid minor, xlim([-1 1]), hold on, xline(0,'Color','k','linewidth',1.5); hold off
            title(sprintf('Histogram of voxel-to-voxel GRAY MATTER correlations AFTER 13-denoise4-WMregress (all slices combined)\n%.1f%% of the correlations were > 0  [ mean=%.2f, median=%.2f, SD=%.2f ]',100*length(find(C3g_>0))/length(C3g_),mean(C3g_),median(C3g_),std(C3g_)),'Color','k','FontSize',14)
        subplot2n(4,1,3), histogram(C3w_,'EdgeColor','none','FaceColor','k','FaceAlpha',0.5,'Normalization','probability'); grid on,grid minor, xlim([-1 1]), hold on, xline(0,'Color','k','linewidth',1.5); hold off
            title(sprintf('WHITE MATTER: AFTER 13-denoise4-WMregress (all slices): %.1f%% of the correlations were > 0  [ mean=%.2f, median=%.2f, SD=%.2f ]',100*length(find(C3w_>0))/length(C3w_),mean(C3w_),median(C3w_),std(C3w_)),'Color','k','FontSize',14)
    end
    subplot2n(4,1,2), histogram(Cg_,'EdgeColor','none','FaceColor',[0,0.55,0.69],'FaceAlpha',0.5,'Normalization','probability'); grid on,grid minor, xlim([-1 1]), hold on, xline(0,'Color','k','linewidth',1.5); hold off
       title(sprintf('GRAY MATTER: AFTER 14-DENOISE5-COVregress (all slices): %.1f%% of correlations > 0  [ mean=%.2f, median=%.2f, SD=%.2f ]',100*length(find(Cg_>0))/length(Cg_),mean(Cg_),median(Cg_),std(Cg_)),'Color',[0,0.55,0.69],'FontSize',14)
    subplot2n(4,1,4), histogram(Cw_,'EdgeColor','none','FaceColor',[0.65,0.11,0.19],'FaceAlpha',0.5,'Normalization','probability'); grid on,grid minor, xlim([-1 1]), hold on, xline(0,'Color','k','linewidth',1.5); hold off
       title(sprintf('WHITE MATTER: AFTER 14-DENOISE5-COVregress (all slices): %.1f%% of the correlations were > 0  [ mean=%.2f, median=%.2f, SD=%.2f ]',100*length(find(Cw_>0))/length(Cw_),mean(Cw_),median(Cw_),std(Cw_)),'Color',[0.65,0.11,0.19],'FontSize',14)
    saveas(fighndl,[QCpath 'hist_FC_compare_13denoise4_14denoise5_allSlices' '.jpg'])
    saveas(fighndl,[QCpath2 'hist_FC_compare_13denoise4_14denoise5_allSlices' '.jpg'])
     close(fighndl)
end
clear svar1 spsc1 svar2 spsc2 sortorder1 sortorder2 F2g_ mF2g_ F2w_ mF2w_ Cg_ Cw_ F4_ mF4_ C2_ F6g_ F6w_ mF6g_ mF6w_ C3g_ C3w_ cmap

%% Make QC video (needs audio video toolbox in Matlab)
if ~isempty(which('VideoWriter')) && ((save_videos==1)||(save_videos==2)) % check if Matlab's audio video toolbox exists

subplot = @(m,n,p) subtightplot(m, n, p, [0.001 0.005], [0.01 0.001], [0.01 0.001]);
scfMRItb_04_unzipFile(base_dir_sub, fname_anat, '_mask_SC')
mask_SC2 = load_untouch_nii([base_dir_sub fname_anat '_mask_SC.nii']); mask_SC2 = mask_SC2.img;
for M=1:size(mask_SC2,3)
    mask_SC = rot90(mask_SC2(:,:,M));
    [xmask_SC,ymask_SC] = find(mask_SC==1);
    minx_SC(:,M) = round(min(xmask_SC)-0.1*(size(mask_SC,1))); maxx_SC(:,M)=round(max(xmask_SC)+0.1*(size(mask_SC,1)));
    miny_SC(:,M) = round(min(ymask_SC)-0.1*(size(mask_SC,2))); maxy_SC(:,M)=round(max(ymask_SC)+0.1*(size(mask_SC,2)));
    if (maxx_SC(:,M)-minx_SC(:,M))<(maxy_SC(:,M)-miny_SC(:,M))
        factr = ((maxy_SC(:,M)-miny_SC(:,M))-(maxx_SC(:,M)-minx_SC(:,M)))/2;
        if factr==round(factr)
            maxx_SC(:,M)=maxx_SC(:,M)+factr; minx_SC(:,M)=minx_SC(:,M)-factr;
        else
            maxx_SC(:,M)=maxx_SC(:,M)+factr+0.5; minx_SC(:,M)=minx_SC(:,M)-factr+0.5;
        end
    elseif (maxy_SC(:,M)-miny_SC(:,M))<(maxx_SC(:,M)-minx_SC(:,M))
        factr = ((maxx_SC(:,M)-minx_SC(:,M))-(maxy_SC(:,M)-miny_SC(:,M)))/2;
        if factr==round(factr)
            maxy_SC(:,M)=maxy_SC(:,M)+factr; miny_SC(:,M)=miny_SC(:,M)-factr;
        else
            maxy_SC(:,M)=maxy_SC(:,M)+factr+0.5; miny_SC(:,M)=miny_SC(:,M)-factr+0.5;
        end
    end
    clear mask_SC xmask_SC ymask_SC factr
end; clear M

for i3 = 1 : siz3
    try waitbar((i3/siz3),wbar3,sprintf('14. Generate QC video: slice (%d) of (%d)',i3,siz3)); catch, end
    fprintf('14. Generate QC video: slice (%d) of (%d)\n',i3,siz3)
    if stp==4
        scfMRItb_04_resplitData(base_dir_sub, fname, ['_denoised' cutoff_str 'WMcov'], siz3) % if slice-wise data doesn't exist but combined 4D data exists, then split the data to generate the needed slice-wise data
        F1 = load_untouch_nii([base_dir_sub fname '_slice' num2str(i3) '_denoised' cutoff_str 'WMcov.nii']);
    elseif stp==3
        scfMRItb_04_resplitData(base_dir_sub, fname, ['_denoised' cutoff_str 'cov'], siz3) % if slice-wise data doesn't exist but combined 4D data exists, then split the data to generate the needed slice-wise data
        F1 = load_untouch_nii([base_dir_sub fname '_slice' num2str(i3) '_denoised' cutoff_str 'cov.nii']);
    elseif stp==2
        scfMRItb_04_resplitData(base_dir_sub, fname, '_WMcov', siz3) % if slice-wise data doesn't exist but combined 4D data exists, then split the data to generate the needed slice-wise data
        F1 = load_untouch_nii([base_dir_sub fname '_slice' num2str(i3) '_WMcov.nii']);
    elseif stp==1
        scfMRItb_04_resplitData(base_dir_sub, fname, '_cov', siz3) % if slice-wise data doesn't exist but combined 4D data exists, then split the data to generate the needed slice-wise data
        F1 = load_untouch_nii([base_dir_sub fname '_slice' num2str(i3) '_cov.nii']);
    end
    TR = F1.hdr.dime.pixdim(5); F1 = rot90(single(squeeze(F1.img)));
    video = VideoWriter([QCpath 'fMRIvideo_14denoise5_slice' sprintf('%.2d',i3) '.mp4'],'MPEG-4'); %#ok<*TNMLP> % create the video object
    video.FrameRate = round(30/TR); % 30 seconds of data in every second of the video
    video.Quality = 90; % very limited compression so that the images are shown as is (and thus avoid misunderstandings due to video quality)
    open(video); % open the file for writing
    
    [sizeFx_SC,sizeFy_SC] = size(F1(:,:,1));
    F2 = F1(minx_SC:maxx_SC,miny_SC:maxy_SC,:);
    for i5=1:size(F1,3)
        F1(:,:,i5) = imresize(F2(:,:,i5),[sizeFx_SC sizeFy_SC]);
    end
    
    if (save_videos==1)
    if stp==1
        scfMRItb_04_resplitData(base_dir_sub, fname, '_warped', siz3) % if slice-wise data doesn't exist but combined 4D data exists, then split the data to generate the needed slice-wise data
        F3 = load_untouch_nii([base_dir_sub fname '_slice' num2str(i3) '_warped.nii']); F3 = rot90(squeeze(F3.img));
    elseif stp==2
        scfMRItb_04_resplitData(base_dir_sub, fname, '_WM', siz3) % if slice-wise data doesn't exist but combined 4D data exists, then split the data to generate the needed slice-wise data
        F3 = load_untouch_nii([base_dir_sub fname '_slice' num2str(i3) '_WM.nii']); F3 = rot90(single(squeeze(F3.img)));
    elseif stp==3
        scfMRItb_04_resplitData(base_dir_sub, fname, ['_denoised' cutoff_str], siz3) % if slice-wise data doesn't exist but combined 4D data exists, then split the data to generate the needed slice-wise data
        F3 = load_untouch_nii([base_dir_sub fname '_slice' num2str(i3) '_denoised' cutoff_str '.nii']); F3 = rot90(single(squeeze(F3.img)));
    elseif stp==4
        scfMRItb_04_resplitData(base_dir_sub, fname, ['_denoised' cutoff_str 'WM'], siz3) % if slice-wise data doesn't exist but combined 4D data exists, then split the data to generate the needed slice-wise data
        F3 = load_untouch_nii([base_dir_sub fname '_slice' num2str(i3) '_denoised' cutoff_str 'WM.nii']); F3 = rot90(single(squeeze(F3.img)));
    end
    [sizeFx,sizeFy] = size(F3(:,:,1));
    F4 = F3(minx_SC:maxx_SC,miny_SC:maxy_SC,:);
    for i5=1:size(F3,3)
        F3(:,:,i5) = imresize(F4(:,:,i5),[sizeFx sizeFy]);
    end
    clear sizeFx sizeFy F2 F4 i5 sizeFx_SC sizeFy_SC
    
    elseif (save_videos==2)
    if Smask_err==0
        scfMRItb_04_resplitData(base_dir_sub, fname, '_Smask', siz3) % if slice-wise data doesn't exist but combined 4D data exists, then split the data to generate the needed slice-wise data
        scfMRItb_04_unzipFile(base_dir_sub, fname, ['_slice' num2str(i3) '_Smask'])
        Smask1 = load_untouch_nii([base_dir_sub fname '_slice' num2str(i3) '_Smask.nii']); Smask1 = Smask1.img;
    else
        Smask1 = Smask1_(:,:,i3);
    end
    Smask1 = rot90(Smask1);
    [xmask,ymask] = find(Smask1==1);
    minx=round(min(xmask)-0.1*(size(Smask1,1))); maxx=round(max(xmask)+0.1*(size(Smask1,1)));
    miny=round(min(ymask)-0.1*(size(Smask1,2))); maxy=round(max(ymask)+0.1*(size(Smask1,2)));
    if (maxx-minx)<(maxy-miny)
        factr = ((maxy-miny)-(maxx-minx))/2;
        if factr==round(factr)
            maxx=maxx+factr; minx=minx-factr;
        else
            maxx=maxx+factr+0.5; minx=minx-factr+0.5;
        end
    elseif (maxy-miny)<(maxx-minx)
        factr = ((maxx-minx)-(maxy-miny))/2;
        if factr==round(factr)
            maxy=maxy+factr; miny=miny-factr;
        else
            maxy=maxy+factr+0.5; miny=miny-factr+0.5;
        end
    end
    scfMRItb_04_resplitData(base_dir_sub, fname, '', siz3) % if slice-wise data doesn't exist but combined 4D data exists, then split the data to generate the needed slice-wise data
    F3 = load_untouch_nii([base_dir_sub fname '_slice' num2str(i3) '.nii']); F3 = rot90(single(squeeze(F3.img)));
    [sizeFx,sizeFy] = size(F3(:,:,1));
    F4 = F3(minx:maxx,miny:maxy,:);
    for i5=1:size(F3,3)
        F3(:,:,i5) = imresize(F4(:,:,i5),[sizeFx sizeFy]);
    end
    clear maxx maxy minx miny xmask ymask Smask1 factr sizeFx sizeFy F2 F4 i5 sizeFx_SC sizeFy_SC        
    end
    
    for ii=1:size(F1,3) % loop across the number of images
        I = F1(:,:,ii); %read the next image
        I(find(isnan(I)))=0;
        I = (I - min(I(:))) ./ (max(I(:)) - min(I(:)));
        I2 = F3(:,:,ii); %read the next image
        I2(find(isnan(I2)))=0;
        I2 = (I2 - min(I2(:))) ./ (max(I2(:)) - min(I2(:)));
        fighndl = figure('Position',[round((scrsz(3)-scrsz(4))/2) 1 round(0.75*scrsz(3)) round(0.75*scrsz(4))],'Color',[0.95,0.9,0.85],'InvertHardcopy','off','Visible','off'); % [0.85,0.90,0.85]
        if (save_videos==1)
        if stp==1
            subplot(1,2,1), imshow(I2, [0 max(I2(:))], 'InitialMagnification', 'fit'); title({['fMRI data after "11. func-anat-registration" (slice ' num2str(i3) ' of ' num2str(siz3) ')'];['Time point ' num2str(ii) ' of ' num2str(size(F1,3))]},'Color','k','FontSize',18);
        elseif stp==2
            subplot(1,2,1), imshow(I2, [0 max(I2(:))], 'InitialMagnification', 'fit'); title({['fMRI data after "13. denoise-4 (WM regression)" (slice ' num2str(i3) ' of ' num2str(siz3) ')'];['Time point ' num2str(ii) ' of ' num2str(size(F1,3))]},'Color','k','FontSize',18);
        elseif stp==3
            subplot(1,2,1), imshow(I2, [0 max(I2(:))], 'InitialMagnification', 'fit'); title({['fMRI data after "12. denoise-3 (CSF regression)" (slice ' num2str(i3) ' of ' num2str(siz3) ')'];['Time point ' num2str(ii) ' of ' num2str(size(F1,3))]},'Color','k','FontSize',18);
        elseif stp==4
            subplot(1,2,1), imshow(I2, [0 max(I2(:))], 'InitialMagnification', 'fit'); title({['fMRI data after "13. denoise-4 (WM regression)" (slice ' num2str(i3) ' of ' num2str(siz3) ')'];['Time point ' num2str(ii) ' of ' num2str(size(F1,3))]},'Color','k','FontSize',18);
        end
        elseif (save_videos==2)
            subplot(1,2,1), imshow(I2, [0 max(I2(:))], 'InitialMagnification', 'fit'); title({['Unprocessed raw fMRI data (slice ' num2str(i3) ' of ' num2str(siz3) ')'];['Time point ' num2str(ii) ' of ' num2str(size(F1,3))]},'Color','k','FontSize',18);
        end
        subplot(1,2,2), imshow(I, [0 max(I(:))], 'InitialMagnification', 'fit'); title({['fMRI data after "14. denoise-5 (COV regression)" (slice ' num2str(i3) ' of ' num2str(siz3) ')'];['Time point ' num2str(ii) ' of ' num2str(size(F1,3))]},'Color',[0,0.55,0.69],'FontSize',18);
        frame = getframe(gcf);
        writeVideo(video,frame); % write the image to file
        close(fighndl)
        clear I I2 frame
    end
    close(video); %close the file
    clear F1 F3 video ii TR
end; clear i3
clear maxx_SC maxy_SC minx_SC miny_SC F

end

try
if nargin<7
    close(wbar3)
end
catch
end

end
```
