## Supplementary material for "Neptune: a toolbox for spinal cord functional MRI data processing and quality assurance": Neptune software: scfMRItb_15_filterData.m

```
function scfMRItb_15_filterData(base_dir_sub,fname,fname_anat_orig,bpf_hz,detrend_flag, varargin)

fname_anat = [fname_anat_orig '_reduced'];
cutoff = 0.50; cutoff_str = '50';

fullsize = 0; % If fullsize=1 then the pre-processed data will not be cropped. By default fullsize=0, so the images are cropped (suggested because the cord is often just a small portion of the field of view [to avoid warping along the phase encoding direction], so the images can be safely cropped without losing the cord. Rest assured that there are checks to ensure that the cord is not cropped off.)

if nargin<6
    wbar3 = waitbar(0,'15. Temporal filtering...','Name','Progress(15): Temporal filtering...','Units','normalized','Position',[3/5 0 1/5 1/17]);
else
    wbar3 = varargin{1};
end

if nargin<7
    save_videos = 1; % 1 = save QC videos (generates sizeable .mp4 files and takes a few minutes to do it). This is turned on by default, but if you don't want to save videos then set it to 0
else
    save_videos = varargin{2};
end
if nargin<8
    Rns = 0; % 0 = don't create runX subfolder; X = create runX subfolder
else
    Rns = varargin{3};
end

if ~(exist([base_dir_sub 'QC'],'dir'))
    mkdir([base_dir_sub 'QC'])
end
if ~(exist([base_dir_sub 'QC' '/15_temporal-filtering'],'dir'))
    mkdir([base_dir_sub 'QC' '/15_temporal-filtering'])
end
if Rns==0
    QCpath = [base_dir_sub 'QC' '/15_temporal-filtering/'];
else
    if ~(exist([base_dir_sub 'QC' '/15_temporal-filtering' '/run' num2str(Rns)],'dir'))
        mkdir([base_dir_sub 'QC' '/15_temporal-filtering' '/run' num2str(Rns)])
    end
    QCpath = [base_dir_sub 'QC' '/15_temporal-filtering' '/run' num2str(Rns) '/'];
end

if ~(exist([base_dir_sub 'QC' '/QC_shortlist'],'dir'))
    mkdir([base_dir_sub 'QC' '/QC_shortlist'])
end
if Rns==0
    QCpath2 = [base_dir_sub 'QC' '/QC_shortlist/'];
else
    if ~(exist([base_dir_sub 'QC' '/QC_shortlist' '/run' num2str(Rns)],'dir'))
        mkdir([base_dir_sub 'QC' '/QC_shortlist' '/run' num2str(Rns)])
    end
    QCpath2 = [base_dir_sub 'QC' '/QC_shortlist' '/run' num2str(Rns) '/'];
end

f_BPF_max_vec = bpf_hz;
f_BPF_max_vec = cat(1, f_BPF_max_vec, [ ]); % if you want filtering to be done for more than one cut-off value, then fill the [ ] with your desired value(s) in Hertz, separated by semicolons

for k=1:length(f_BPF_max_vec)
if f_BPF_max_vec(k)<0.1
    f_BPF_max_str_vec{k,1} = sprintf('0%.0f',f_BPF_max_vec(k)*100); %#ok<*AGROW>
else
    f_BPF_max_str_vec{k,1} = sprintf('%.0f',f_BPF_max_vec(k)*100);
end
end; clear k


if exist([base_dir_sub fname '_warped' '.nii'], 'file') || exist([base_dir_sub fname '_warped' '.nii.gz'], 'file') % if none of CSF, WM or COV regress steps were performed
    whichip=8;
end
if exist([base_dir_sub fname '_WM' '.nii'], 'file') || exist([base_dir_sub fname '_WM' '.nii.gz'], 'file') % if only WM regress step was performed
    whichip=7;
end
if exist([base_dir_sub fname '_denoised' cutoff_str '.nii'], 'file') || exist([base_dir_sub fname '_denoised' cutoff_str '.nii.gz'], 'file') % if only CSF regress step was performed
    whichip=6;
end
if exist([base_dir_sub fname '_cov' '.nii'], 'file') || exist([base_dir_sub fname '_cov' '.nii.gz'], 'file') % if only COV regress step was performed
    whichip=5;
end
if exist([base_dir_sub fname '_WMcov' '.nii'], 'file') || exist([base_dir_sub fname '_WMcov' '.nii.gz'], 'file') % if both WM and COV regress steps were performed
    whichip=4;
end
if exist([base_dir_sub fname '_denoised' cutoff_str 'WM' '.nii'], 'file') || exist([base_dir_sub fname '_denoised' cutoff_str 'WM' '.nii.gz'], 'file') % if both CSF and WM regress steps were performed
    whichip=3;
end
if exist([base_dir_sub fname '_denoised' cutoff_str 'cov' '.nii'], 'file') || exist([base_dir_sub fname '_denoised' cutoff_str 'cov' '.nii.gz'], 'file') % if both CSF and COV regress steps were performed
    whichip=2;
end
if exist([base_dir_sub fname '_denoised' cutoff_str 'WMcov' '.nii'], 'file') || exist([base_dir_sub fname '_denoised' cutoff_str 'WMcov' '.nii.gz'], 'file') % if CSF, WM and COV regress steps were performed
    whichip=1;
end

do_LPF = 0; % do not perform low-pass filtering because it is not useful, given that bandpass filtering is being done (1 = do LPF)
save_nifti = 1; % filtered data is saved as .mat by default, which is used in steps hereafter. This flag must be set to 1 if you also want to save band-pass filtered data as .nii, in addition to the .mat (this is set to 1 by default, so the NIfTIs of band-pass filtered data are saved. However, low-pass and high-pass filtered data are only saved as .mat) 

%% prep
if whichip==1
    load([base_dir_sub fname '_denoised' cutoff_str 'WMcov' '.mat']); %#ok<*LOAD>
elseif whichip==2
    load([base_dir_sub fname '_denoised' cutoff_str 'cov' '.mat']);
elseif whichip==3
    load([base_dir_sub fname '_denoised' cutoff_str 'WM' '.mat']);
elseif whichip==4
    load([base_dir_sub fname '_WMcov' '.mat']);
elseif whichip==5
    load([base_dir_sub fname '_cov' '.mat']);
elseif whichip==6
    load([base_dir_sub fname '_denoised' cutoff_str '.mat']);
elseif whichip==7
    load([base_dir_sub fname '_WM' '.mat']);
elseif whichip==8
    load([base_dir_sub fname '_warped' '.mat']);
end
F2 = F; clear F
[siz1, siz2, siz3, ~] = size(F2.img);

f_BPF_max = f_BPF_max_vec(1,1);
f_BPF_max_str = f_BPF_max_str_vec{1,1};
vat = F2.hdr.dime.pixdim(1,5); % volume acquisition time
f_sampling = 1 / vat; % Hz

f_BPF_max_str_vec(find(f_BPF_max_vec>=f_sampling/2))=[]; % cannot bandpass filter with upper cut-offs larger than half the data sampling rate
f_BPF_max_vec(find(f_BPF_max_vec>=f_sampling/2))=[]; % cannot bandpass filter with upper cut-offs larger than half the data sampling rate
if isempty(f_BPF_max_vec)
    f_BPF_max_vec = f_sampling/2; % for the sanity of future steps that depend on _BPF data, do BPF with the upper cutoff as the maximum frequency itself IF upper cut-off is larger than half the data sampling rate
    if f_BPF_max_vec<0.1
        f_BPF_max_str_vec = sprintf('0%.0f',f_BPF_max_vec*100); %#ok<*AGROW>
    else
        f_BPF_max_str_vec = sprintf('%.0f',f_BPF_max_vec*100);
    end
end

% define cord mask; filtering will be performed only on voxels within the mask
basestr = strfind(base_dir_sub,'/');
basestr = base_dir_sub(basestr(end-1)+1:end); % name of subject folder
try
    scfMRItb_04_unzipFile(base_dir_sub, fname_anat, '2_mask_GM')
    scfMRItb_04_unzipFile(base_dir_sub, fname_anat, '2_mask_WM')
    mask_GM_reduced  = load_untouch_nii([base_dir_sub fname_anat '2' '_mask_GM.nii']);
    mask_WM_reduced  = load_untouch_nii([base_dir_sub fname_anat '2' '_mask_WM.nii']);
catch
	scfMRItb_04_unzipFile(base_dir_sub, fname_anat, '_mask_GM')
	scfMRItb_04_unzipFile(base_dir_sub, fname_anat, '_mask_WM')
	scfMRItb_04_unzipFile(base_dir_sub, fname_anat, '_mask_CSF')
    mmask1 = load_untouch_nii([base_dir_sub fname_anat '_mask_GM.nii']); % gray matter
    mmask2 = load_untouch_nii([base_dir_sub fname_anat '_mask_WM.nii']); % white matter
    mmask3 = load_untouch_nii([base_dir_sub fname_anat '_mask_CSF.nii']); % white matter
    mmask_all = single(or(or(mmask1.img, mmask2.img), mmask3.img));    
    smmask = (mean(mmask_all,3) > 0);
    [s1,s2] = find(smmask~=0);
    s1median = median(s1); s2median = median(s2);
    s1min = min(s1); s2min = min(s2);
    s1max = max(s1); s2max = max(s2);
    smaxrange = max([s2max-s2min,s1max-s1min]);
    if round(size(mmask_all,1)/4) > smaxrange
        new_size = round(size(mmask_all,1)/4); % 128 - assume even for now
    elseif round(size(mmask_all,1)/3) > smaxrange
        new_size = round(size(mmask_all,1)/3); % 171
    elseif round(size(mmask_all,1)/2) > smaxrange
        new_size = round(size(mmask_all,1)/2); % 256
    else
        new_size = size(mmask_all,1); % 512
    end
    if fullsize==1
        new_size = size(mmask_all,1);
    end
    xmin = s1median - round(new_size/2) + 1; xmax = s1median + round(new_size/2);
    ymin = s2median - round(new_size/2) + 1; ymax = s2median + round(new_size/2);
    new_size = xmax - xmin + 1;
    if str2num(basestr(1:4)) <= 2017 %#ok<ST2NM> % if data were acquired prior to 2018 then don't change the bounding box of the reduced size images that were already processed before
        mask_GM_reduced.img = mmask1.img(round((size(A.img,1)-new_size)/2)+1:end-round((size(A.img,1)-new_size)/2), ...
            round((size(A.img,2)-new_size)/2)+1:end-round((size(A.img,2)-new_size)/2), :);
        mask_WM_reduced.img = mmask2.img(round((size(A.img,1)-new_size)/2)+1:end-round((size(A.img,1)-new_size)/2), ...
            round((size(A.img,2)-new_size)/2)+1:end-round((size(A.img,2)-new_size)/2), :);
    else % else if data were acquired after 2017 then use the new method to determine the bounding box by centering the new box around the centroid of the spinal cord mask
        mask_GM_reduced.img = mmask1.img(xmin:xmax, ymin:ymax, :);
        mask_WM_reduced.img = mmask2.img(xmin:xmax, ymin:ymax, :);
    end
end
% mask_filt = or(or(mask_GM_reduced.img, mask_WM_reduced.img), mask_CSF_reduced.img) > 0;
mask_filt = or(mask_GM_reduced.img, mask_WM_reduced.img) > 0;


%% ----- LOW PASS FILTER DATA ----------------------------------------------
% LPF not useful, hence not doing (do_LPF=0)
if do_LPF == 1
    F = F2;
    Wst = [0.01 f_BPF_max] / (f_sampling/2);
    stopband_attenuation = 60; % in dB
    passband_ripple = 3; % in dB
    minorder = ceil( acosh( sqrt( (10^(0.1*(stopband_attenuation+2*passband_ripple))-1)/(sqrt(10^(0.1*passband_ripple)-1)^2) )) ./ acosh( f_sampling/f_BPF_max )); % Refer to this link for the formula used here to determine the model order: https://electronics.stackexchange.com/questions/519144/designing-a-chebyshev-filter-am-i-getting-the-order-wrong
    cheby_order = min([minorder,round(size(F.img,4)/10)]); % The model order must be at least one-tenth of the total number of time points (this is a ballpark value, not determined analytically).
    [B_LPF, A_LPF] = cheby2(cheby_order, stopband_attenuation, Wst, 'low');
    clear minorder stopband_attenuation passband_ripple cheby_order Wst
    for i3 = 1 : siz3
        try waitbar((i3/siz3)/3,wbar3,sprintf('15. Temporal filtering (1/3: low-pass): slice (%d) of (%d)',i3,siz3)); catch, end
        for i1 = 1 : siz1
            for i2 = 1 : siz2
                if mask_filt(i1,i2,i3)==1
                if detrend_flag==1
                    F.img(i1,i2,i3,:) = single(reshape(filtfilt(B_LPF, A_LPF, detrend(double(squeeze(F2.img(i1,i2,i3,:))),1)'), [1 1 1 size(F2.img,4)]));
                elseif detrend_flag==2
                    F.img(i1,i2,i3,:) = single(reshape(filtfilt(B_LPF, A_LPF, detrend(double(squeeze(F2.img(i1,i2,i3,:))),2)'), [1 1 1 size(F2.img,4)]));
                elseif detrend_flag==0
                    F.img(i1,i2,i3,:) = single(reshape(filtfilt(B_LPF, A_LPF, double(squeeze(F2.img(i1,i2,i3,:)))'), [1 1 1 size(F2.img,4)]));
                end
                end
            end
        end
    end
    
    if whichip==1
        save([base_dir_sub fname '_denoised' cutoff_str 'WMcov' '_LPF' f_BPF_max_str '.mat'], 'F'); % save lowpass filtered data as .mat
    elseif whichip==2
        save([base_dir_sub fname '_denoised' cutoff_str 'cov' '_LPF' f_BPF_max_str '.mat'], 'F'); % save lowpass filtered data as .mat
    elseif whichip==3
        save([base_dir_sub fname '_denoised' cutoff_str 'WM' '_LPF' f_BPF_max_str '.mat'], 'F'); % save lowpass filtered data as .mat
    elseif whichip==4
        save([base_dir_sub fname '_WMcov' '_LPF' f_BPF_max_str '.mat'], 'F'); % save lowpass filtered data as .mat
    elseif whichip==5
        save([base_dir_sub fname '_cov' '_LPF' f_BPF_max_str '.mat'], 'F'); % save lowpass filtered data as .mat
    elseif whichip==6
        save([base_dir_sub fname '_denoised' cutoff_str '_LPF' f_BPF_max_str '.mat'], 'F'); % save lowpass filtered data as .mat
    elseif whichip==7
        save([base_dir_sub fname '_WM' '_LPF' f_BPF_max_str '.mat'], 'F'); % save lowpass filtered data as .mat
    elseif whichip==8
        save([base_dir_sub fname '_warped' '_LPF' f_BPF_max_str '.mat'], 'F'); % save lowpass filtered data as .mat
    end
    clear F;
    clear Wst B_LPF A_LPF
end

%% ----- BAND PASS FILTER DATA ---------------------------------------------
i4max = length(f_BPF_max_vec);
for i4 = 1:length(f_BPF_max_vec)

F = F2;
f_BPF_max = f_BPF_max_vec(1,i4);
f_BPF_max_str = f_BPF_max_str_vec{1,i4};

Wst = [0.01 f_BPF_max] / (f_sampling/2);
stopband_attenuation = 60; % in dB
passband_ripple = 3; % in dB
minorder = ceil( acosh( sqrt( (10^(0.1*(stopband_attenuation+2*passband_ripple))-1)/(sqrt(10^(0.1*passband_ripple)-1)^2) )) ./ acosh( f_sampling/f_BPF_max )); % Refer to this link for the formula used here to determine the model order: https://electronics.stackexchange.com/questions/519144/designing-a-chebyshev-filter-am-i-getting-the-order-wrong
cheby_order = min([minorder,round(size(F.img,4)/10)]); % The model order must be at least one-tenth of the total number of time points (this is a ballpark value, not determined analytically).
[B_BPF, A_BPF] = cheby2(cheby_order, stopband_attenuation, Wst);
clear minorder stopband_attenuation passband_ripple cheby_order Wst

filts = designfilt('bandpassfir', 'FilterOrder', round(30*f_sampling), 'CutoffFrequency1', 0.01, 'CutoffFrequency2', f_BPF_max, 'SampleRate', f_sampling);
coef = filts.Coefficients; coef=coef'; clear filts

for i3 = 1 : siz3
    try waitbar((1/3)+(i3/siz3)/3,wbar3,sprintf('15. Temporal filtering (%d/%d: band-pass): slice (%d) of (%d)',i4,i4max+1,i3,siz3)); catch, end
    for i1 = 1 : siz1
        for i2 = 1 : siz2
            if mask_filt(i1,i2,i3)==1
            try
            if detrend_flag==1
                F.img(i1,i2,i3,:) = filtfilt(coef, 1, detrend(double(squeeze(F2.img(i1,i2,i3,:))),1));
            elseif detrend_flag==2
                F.img(i1,i2,i3,:) = filtfilt(coef, 1, detrend(double(squeeze(F2.img(i1,i2,i3,:))),2));
            elseif detrend_flag==0
                F.img(i1,i2,i3,:) = filtfilt(coef, 1, double(squeeze(F2.img(i1,i2,i3,:))));
            end
            catch
            if detrend_flag==1
                F.img(i1,i2,i3,:) = filtfilt(B_BPF, A_BPF, detrend(double(squeeze(F2.img(i1,i2,i3,:))),1));
            elseif detrend_flag==2
                F.img(i1,i2,i3,:) = filtfilt(B_BPF, A_BPF, detrend(double(squeeze(F2.img(i1,i2,i3,:))),2));
            elseif detrend_flag==0
                F.img(i1,i2,i3,:) = filtfilt(B_BPF, A_BPF, double(squeeze(F2.img(i1,i2,i3,:))));
            end
            end
            end
        end
    end
end
F.img = single(F.img);

F.hdr.dime.dim(2) = size(F.img,1); F.hdr.dime.dim(3) = size(F.img,2);
try
    F.original;
    F.original.hdr.dime.dim(2) = size(F.img,1); F.original.hdr.dime.dim(3) = size(F.img,2);
catch
end

if whichip==1
    save([base_dir_sub fname '_denoised' cutoff_str 'WMcov' '_BPF' f_BPF_max_str '.mat'], 'F'); % save bandpass filtered data as .mat
elseif whichip==2
    save([base_dir_sub fname '_denoised' cutoff_str 'cov' '_BPF' f_BPF_max_str '.mat'], 'F'); % save bandpass filtered data as .mat
elseif whichip==3
    save([base_dir_sub fname '_denoised' cutoff_str 'WM' '_BPF' f_BPF_max_str '.mat'], 'F'); % save bandpass filtered data as .mat
elseif whichip==4
    save([base_dir_sub fname '_WMcov' '_BPF' f_BPF_max_str '.mat'], 'F'); % save bandpass filtered data as .mat
elseif whichip==5
    save([base_dir_sub fname '_cov' '_BPF' f_BPF_max_str '.mat'], 'F'); % save bandpass filtered data as .mat
elseif whichip==6
    save([base_dir_sub fname '_denoised' cutoff_str '_BPF' f_BPF_max_str '.mat'], 'F'); % save bandpass filtered data as .mat
elseif whichip==7
    save([base_dir_sub fname '_WM' '_BPF' f_BPF_max_str '.mat'], 'F'); % save bandpass filtered data as .mat
elseif whichip==8
    save([base_dir_sub fname '_warped' '_BPF' f_BPF_max_str '.mat'], 'F'); % save bandpass filtered data as .mat
end

if save_nifti==1
if whichip==1
    save_untouch_nii(F, [base_dir_sub fname '_denoised' cutoff_str 'WMcov' '_BPF' f_BPF_max_str '.nii']); % save bandpass filtered data as NIfTI
elseif whichip==2
    save_untouch_nii(F, [base_dir_sub fname '_denoised' cutoff_str 'cov' '_BPF' f_BPF_max_str '.nii']); % save bandpass filtered data as NIfTI
elseif whichip==3
    save_untouch_nii(F, [base_dir_sub fname '_denoised' cutoff_str 'WM' '_BPF' f_BPF_max_str '.nii']); % save bandpass filtered data as NIfTI
elseif whichip==4
    save_untouch_nii(F, [base_dir_sub fname '_WMcov' '_BPF' f_BPF_max_str '.nii']); % save bandpass filtered data as NIfTI
elseif whichip==5
    save_untouch_nii(F, [base_dir_sub fname '_cov' '_BPF' f_BPF_max_str '.nii']); % save bandpass filtered data as NIfTI
elseif whichip==6
    save_untouch_nii(F, [base_dir_sub fname '_denoised' cutoff_str '_BPF' f_BPF_max_str '.nii']); % save bandpass filtered data as NIfTI
elseif whichip==7
    save_untouch_nii(F, [base_dir_sub fname '_WM' '_BPF' f_BPF_max_str '.nii']); % save bandpass filtered data as NIfTI
elseif whichip==8
    save_untouch_nii(F, [base_dir_sub fname '_warped' '_BPF' f_BPF_max_str '.nii']); % save bandpass filtered data as NIfTI
end
end

clear F
clear Wst B_LPF A_LPF f_BPF_max coef
end % i4

%% ----- HIGH PASS FILTER DATA ---------------------------------------------

F = F2;

clear B_HPF A_HPF
Wst = 0.01 / (f_sampling/2);
stopband_attenuation = 60; % in dB
passband_ripple = 3; % in dB
minorder = ceil( acosh( sqrt( (10^(0.1*(stopband_attenuation+2*passband_ripple))-1)/(sqrt(10^(0.1*passband_ripple)-1)^2) )) ./ acosh( f_sampling/0.01 )); % Refer to this link for the formula used here to determine the model order: https://electronics.stackexchange.com/questions/519144/designing-a-chebyshev-filter-am-i-getting-the-order-wrong
cheby_order = min([minorder,round(size(F.img,4)/10)]); % The model order must be at least one-tenth of the total number of time points (this is a ballpark value, not determined analytically).
[B_HPF, A_HPF] = cheby2(cheby_order, stopband_attenuation, Wst, 'high');
clear minorder stopband_attenuation passband_ripple cheby_order Wst

filts = designfilt('highpassfir', 'FilterOrder', round(30*f_sampling), 'CutoffFrequency', 0.01, 'SampleRate', f_sampling);
coef = filts.Coefficients; coef=coef'; clear filts

for i3 = 1 : siz3
    try waitbar((2/3)+(i3/siz3)/3,wbar3,sprintf('15. Temporal filtering (%d/%d: high-pass): slice (%d) of (%d)',i4max+1,i4max+1,i3,siz3)); catch, end
    for i1 = 1 : siz1
        for i2 = 1 : siz2
            if mask_filt(i1,i2,i3)==1
            try
            if detrend_flag==1
                F.img(i1,i2,i3,:) = filtfilt(coef, 1, detrend(double(squeeze(F2.img(i1,i2,i3,:))),1));
            elseif detrend_flag==2
                F.img(i1,i2,i3,:) = filtfilt(coef, 1, detrend(double(squeeze(F2.img(i1,i2,i3,:))),2));
            elseif detrend_flag==0
                F.img(i1,i2,i3,:) = filtfilt(coef, 1, double(squeeze(F2.img(i1,i2,i3,:))));
            end
            catch
            if detrend_flag==1
                F.img(i1,i2,i3,:) = filtfilt(B_HPF, A_HPF, detrend(double(squeeze(F2.img(i1,i2,i3,:))),1));
            elseif detrend_flag==2
                F.img(i1,i2,i3,:) = filtfilt(B_HPF, A_HPF, detrend(double(squeeze(F2.img(i1,i2,i3,:))),2));
            elseif detrend_flag==0
                F.img(i1,i2,i3,:) = filtfilt(B_HPF, A_HPF, double(squeeze(F2.img(i1,i2,i3,:))));
            end
            end
            end
        end
    end
end
F.img = single(F.img);

F.hdr.dime.dim(2) = size(F.img,1); F.hdr.dime.dim(3) = size(F.img,2);
try
    F.original;
    F.original.hdr.dime.dim(2) = size(F.img,1); F.original.hdr.dime.dim(3) = size(F.img,2);
catch
end

if whichip==1
    save([base_dir_sub fname '_denoised' cutoff_str 'WMcov' '_HPF' '01' '.mat'], 'F'); % save highpass filtered data as .mat
elseif whichip==2
    save([base_dir_sub fname '_denoised' cutoff_str 'cov' '_HPF' '01' '.mat'], 'F'); % save highpass filtered data as .mat
elseif whichip==3
    save([base_dir_sub fname '_denoised' cutoff_str 'WM' '_HPF' '01' '.mat'], 'F'); % save highpass filtered data as .mat
elseif whichip==4
    save([base_dir_sub fname '_WMcov' '_HPF' '01' '.mat'], 'F'); % save highpass filtered data as .mat
elseif whichip==5
    save([base_dir_sub fname '_cov' '_HPF' '01' '.mat'], 'F'); % save highpass filtered data as .mat
elseif whichip==6
    save([base_dir_sub fname '_denoised' cutoff_str '_HPF' '01' '.mat'], 'F'); % save highpass filtered data as .mat
elseif whichip==7
    save([base_dir_sub fname '_WM' '_HPF' '01' '.mat'], 'F'); % save highpass filtered data as .mat
elseif whichip==8
    save([base_dir_sub fname '_warped' '_HPF' '01' '.mat'], 'F'); % save highpass filtered data as .mat
end

if 0 % saving high-pass filtered data as NIfTI is disabled by default. If you want the .nii in addition to .mat, set this to 1
if whichip==1 %#ok<UNRCH>
    save_untouch_nii(F, [base_dir_sub fname '_denoised' cutoff_str 'WMcov' '_HPF' '01' '.nii']); % save highpass filtered data as NIfTI
elseif whichip==2
    save_untouch_nii(F, [base_dir_sub fname '_denoised' cutoff_str 'cov' '_HPF' '01' '.nii']); % save highpass filtered data as NIfTI
elseif whichip==3
    save_untouch_nii(F, [base_dir_sub fname '_denoised' cutoff_str 'WM' '_HPF' '01' '.nii']); % save highpass filtered data as NIfTI
elseif whichip==4
    save_untouch_nii(F, [base_dir_sub fname '_WMcov' '_HPF' '01' '.nii']); % save highpass filtered data as NIfTI
elseif whichip==5
    save_untouch_nii(F, [base_dir_sub fname '_cov' '_HPF' '01' '.nii']); % save highpass filtered data as NIfTI
elseif whichip==6
    save_untouch_nii(F, [base_dir_sub fname '_denoised' cutoff_str '_HPF' '01' '.nii']); % save highpass filtered data as NIfTI
elseif whichip==7
    save_untouch_nii(F, [base_dir_sub fname '_WM' '_HPF' '01' '.nii']); % save highpass filtered data as NIfTI
elseif whichip==8
    save_untouch_nii(F, [base_dir_sub fname '_warped' '_HPF' '01' '.nii']); % save highpass filtered data as NIfTI
end
end

clear F coef;

TR = vat;

try
save([base_dir_sub 'workspace_variables_temp.mat'], 'base_dir_sub','fname*','cutoff*','f_BPF_max*','TR')
catch
save([base_dir_sub 'workspace_variables_temp.mat'])
end


%% time series plots, global signal and FC histogram
scrsz = get(0,'ScreenSize'); pause on;
cmap = colormap('bone'); cmap = (1-exp(-cmap))./(1-exp(-1)); 

scfMRItb_04_unzipFile(base_dir_sub, fname_anat, '_mask_GM')
scfMRItb_04_unzipFile(base_dir_sub, fname_anat, '_mask_WM')
mask_GM = load_untouch_nii([base_dir_sub fname_anat '_mask_GM.nii']); mask_GM = mask_GM.img;
mask_WM = load_untouch_nii([base_dir_sub fname_anat '_mask_WM.nii']); mask_WM = mask_WM.img;

scfMRItb_04_unzipFile(base_dir_sub, fname_anat, '2_mask_GM')
scfMRItb_04_unzipFile(base_dir_sub, fname_anat, '2_mask_WM')
mask_GM2 = load_untouch_nii([base_dir_sub fname_anat '2_mask_GM.nii']); mask_GM2 = mask_GM2.img;
mask_WM2 = load_untouch_nii([base_dir_sub fname_anat '2_mask_WM.nii']); mask_WM2 = mask_WM2.img;

if whichip==1
    load([base_dir_sub fname '_denoised' cutoff_str 'WMcov' '_BPF' f_BPF_max_str '.mat']); F1 = F; clear F %#ok<*LOAD>
    stp=8;
elseif whichip==2
    load([base_dir_sub fname '_denoised' cutoff_str 'cov' '_BPF' f_BPF_max_str '.mat']); F1 = F; clear F
    stp=7;
elseif whichip==3
    load([base_dir_sub fname '_denoised' cutoff_str 'WM' '_BPF' f_BPF_max_str '.mat']); F1 = F; clear F
    stp=6;
elseif whichip==4
    load([base_dir_sub fname '_WMcov' '_BPF' f_BPF_max_str '.mat']); F1 = F; clear F
    stp=5;
elseif whichip==5
    load([base_dir_sub fname '_cov' '_BPF' f_BPF_max_str '.mat']); F1 = F; clear F
    stp=4;
elseif whichip==6
    load([base_dir_sub fname '_denoised' cutoff_str '_BPF' f_BPF_max_str '.mat']); F1 = F; clear F
    stp=3;
elseif whichip==7
    load([base_dir_sub fname '_WM' '_BPF' f_BPF_max_str '.mat']); F1 = F; clear F
    stp=2;
elseif whichip==8
    load([base_dir_sub fname '_warped' '_BPF' f_BPF_max_str '.mat']); F1 = F; clear F
    stp=1;
end
TR = F1.hdr.dime.pixdim(5); F1 = single(squeeze(F1.img));


Smask_err=0;
try
    scfMRItb_04_resplitData(base_dir_sub, fname, '_Smask', siz3) % if slice-wise data doesn't exist but combined 4D data exists, then split the data to generate the needed slice-wise data
catch
    load([base_dir_sub fname '_mask_NS' '.mat'])
    Smask1_ = ~mask_NS.img; clear mask_NS
    Smask_err=1;
end

F2g_=[]; mF2g_=[]; F2w_=[]; mF2w_=[]; Cg_=[]; Cw_=[]; F4_=[]; mF4_=[]; C2_=[];
if stp~=0, F6g_=[]; mF6g_=[]; F6w_=[]; mF6w_=[]; C3g_=[]; C3w_=[]; end
for i3 = 1 : siz3
    try waitbar((i3/siz3),wbar3,sprintf('15. QC plots: slice (%d) of (%d)',i3,siz3)); catch, end
    fprintf('15. QC plots: slice (%d) of (%d)\n',i3,siz3)
    
    if Smask_err==0
        scfMRItb_04_resplitData(base_dir_sub, fname, '_Smask', siz3) % if slice-wise data doesn't exist but combined 4D data exists, then split the data to generate the needed slice-wise data
        scfMRItb_04_unzipFile(base_dir_sub, fname, ['_slice' num2str(i3) '_Smask'])
        Smask1 = load_untouch_nii([base_dir_sub fname '_slice' num2str(i3) '_Smask.nii']); Smask1 = Smask1.img;
    else
        Smask1 = Smask1_(:,:,i3);
    end

    for i5=1:size(F1,4)
        tmp = F1(:,:,i3,i5);
        F2g(:,i5) = tmp(find(mask_GM2(:,:,i3)==1));
        F2w(:,i5) = tmp(find(mask_WM2(:,:,i3)==1));
        clear tmp
    end; clear i5
    [~,svar1] = sort(var(F2g,0,2)); [~,sdist1] = sort(svar1); sortorder1{i3,1} = sdist1;
    [~,svar2] = sort(var(F2w,0,2)); [~,sdist2] = sort(svar2); sortorder2{i3,1} = sdist2;
    
    fighndl = figure('Position',[1 1 1152 720],'Color',[0.95,0.9,0.85],'InvertHardcopy','off','Visible','off'); % red [0.95,0.9,0.85], green [0.85,0.90,0.85], blue [0.89,0.94,0.99]
    subplot2n(2,1,1), imagesc(F2g(sortorder1{i3},:)),colormap(cmap),colorbar, xlabel('time (in samples)'),ylabel('Gray matter voxels (sorted by variance)'),
        title(['Time series within the GRAY MATTER after "15-bandpass-filter" in slice ' num2str(i3) ' of ' num2str(siz3)],'Color',[0,0.55,0.69],'FontSize',14);
    subplot2n(2,1,2), imagesc(F2w(sortorder2{i3},:)),colormap(cmap),colorbar, xlabel('time (in samples)'),ylabel('White matter voxels (sorted by variance)'),
        title(['Time series within the WHITE MATTER after "15-bandpass-filter" in slice ' num2str(i3) ' of ' num2str(siz3)],'Color',[0.65,0.11,0.19],'FontSize',14);
    saveas(fighndl,[QCpath 'fMRI_ts_15filter_slice' sprintf('%.2d',i3) '.jpg'])
    close(fighndl)
    
    scfMRItb_04_resplitData(base_dir_sub, fname, '', siz3) % if slice-wise data doesn't exist but combined 4D data exists, then split the data to generate the needed slice-wise data
	scfMRItb_04_unzipFile(base_dir_sub, fname, ['_slice' num2str(i3)])
    F3 = load_untouch_nii([base_dir_sub fname '_slice' num2str(i3) '.nii']); F3 = single(squeeze(F3.img));
    for i5=1:size(F3,3)
        tmp = F3(:,:,i5);
        F4(:,i5) = tmp(find(Smask1==1)); clear tmp
    end; clear i5 F3
    mF2g=mean(F2g,1); mF2w=mean(F2w,1);
    mF2g=zscore(mF2g); mF2w=zscore(mF2w);

    Cg = corrcoef(F2g'); Cg = uppertriangle(Cg);
    Cw = corrcoef(F2w'); Cw = uppertriangle(Cw);
    C2 = corrcoef(F4');  C2 = uppertriangle(C2);
    fighndl = figure('Position',[1 1 1152 720],'Color',[0.95,0.9,0.85],'InvertHardcopy','off','Visible','off'); % red [0.95,0.9,0.85], green [0.85,0.90,0.85], blue [0.89,0.94,0.99]
    subplot2n(3,1,1), histogram(C2,'EdgeColor','none','FaceColor','k','FaceAlpha',0.5,'Normalization','probability'); grid on,grid minor, xlim([-1 1]), hold on, xline(0,'Color','k','linewidth',1.5); hold off
      title(sprintf('Histogram of voxel-to-voxel correlations in UNPROCESSED RAW DATA (slice %d of %d)\n%.1f%% of the correlations were > 0  [ mean=%.2f, median=%.2f, SD=%.2f ]',i3,siz3,100*length(find(C2>0))/length(C2),mean(C2),median(C2),std(C2)),'Color','k','FontSize',14)
    subplot2n(3,1,2), histogram(Cg,'EdgeColor','none','FaceColor',[0,0.55,0.69],'FaceAlpha',0.5,'Normalization','probability'); grid on,grid minor, xlim([-1 1]), hold on, xline(0,'Color','k','linewidth',1.5); hold off
      title(sprintf('Histogram of voxel-to-voxel GRAY MATTER correlations AFTER 15-bandpass-filter [0.01-%.2fHz] (slice %d of %d)\n%.1f%% of the correlations were > 0  [ mean=%.2f, median=%.2f, SD=%.2f ]',f_BPF_max_vec(1),i3,siz3,100*length(find(Cg>0))/length(Cg),mean(Cg),median(Cg),std(Cg)),'Color',[0,0.55,0.69],'FontSize',14)
    subplot2n(3,1,3), histogram(Cw,'EdgeColor','none','FaceColor',[0.65,0.11,0.19],'FaceAlpha',0.5,'Normalization','probability'); grid on,grid minor, xlim([-1 1]), hold on, xline(0,'Color','k','linewidth',1.5); hold off
      title(sprintf('Histogram of voxel-to-voxel WHITE MATTER correlations AFTER 15-bandpass-filter [0.01-%.2fHz] (slice %d of %d)\n%.1f%% of the correlations were > 0  [ mean=%.2f, median=%.2f, SD=%.2f ]',f_BPF_max_vec(1),i3,siz3,100*length(find(Cw>0))/length(Cw),mean(Cw),median(Cw),std(Cw)),'Color',[0.65,0.11,0.19],'FontSize',14)
    saveas(fighndl,[QCpath 'hist_FC_compare_15filter_slice' sprintf('%.2d',i3) '.jpg'])
    close(fighndl)
    
    if stp~=0
        if stp==1
            scfMRItb_04_resplitData(base_dir_sub, fname, '_warped', siz3) % if slice-wise data doesn't exist but combined 4D data exists, then split the data to generate the needed slice-wise data
            scfMRItb_04_unzipFile(base_dir_sub, fname, ['_slice' num2str(i3) '_warped'])
            F5 = load_untouch_nii([base_dir_sub fname '_slice' num2str(i3) '_warped.nii']); F5 = single(squeeze(F5.img));
        elseif stp==2
            scfMRItb_04_resplitData(base_dir_sub, fname, '_WM', siz3) % if slice-wise data doesn't exist but combined 4D data exists, then split the data to generate the needed slice-wise data
            scfMRItb_04_unzipFile(base_dir_sub, fname, ['_slice' num2str(i3) '_WM'])
            F5 = load_untouch_nii([base_dir_sub fname '_slice' num2str(i3) '_WM.nii']); F5 = single(squeeze(F5.img));
        elseif stp==3
            scfMRItb_04_resplitData(base_dir_sub, fname, ['_denoised' cutoff_str], siz3) % if slice-wise data doesn't exist but combined 4D data exists, then split the data to generate the needed slice-wise data
            scfMRItb_04_unzipFile(base_dir_sub, fname, ['_slice' num2str(i3) '_denoised' cutoff_str ''])
            F5 = load_untouch_nii([base_dir_sub fname '_slice' num2str(i3) '_denoised' cutoff_str '.nii']); F5 = single(squeeze(F5.img));
        elseif stp==4
            scfMRItb_04_resplitData(base_dir_sub, fname, '_cov', siz3) % if slice-wise data doesn't exist but combined 4D data exists, then split the data to generate the needed slice-wise data
            scfMRItb_04_unzipFile(base_dir_sub, fname, ['_slice' num2str(i3) '_cov'])
            F5 = load_untouch_nii([base_dir_sub fname '_slice' num2str(i3) '_cov.nii']); F5 = single(squeeze(F5.img));
        elseif stp==5
            scfMRItb_04_resplitData(base_dir_sub, fname, '_WMcov', siz3) % if slice-wise data doesn't exist but combined 4D data exists, then split the data to generate the needed slice-wise data
            scfMRItb_04_unzipFile(base_dir_sub, fname, ['_slice' num2str(i3) '_WMcov'])
            F5 = load_untouch_nii([base_dir_sub fname '_slice' num2str(i3) '_WMcov.nii']); F5 = single(squeeze(F5.img));
        elseif stp==6
            scfMRItb_04_resplitData(base_dir_sub, fname, ['_denoised' cutoff_str 'WM'], siz3) % if slice-wise data doesn't exist but combined 4D data exists, then split the data to generate the needed slice-wise data
            scfMRItb_04_unzipFile(base_dir_sub, fname, ['_slice' num2str(i3) '_denoised' cutoff_str 'WM'])
            F5 = load_untouch_nii([base_dir_sub fname '_slice' num2str(i3) '_denoised' cutoff_str 'WM.nii']); F5 = single(squeeze(F5.img));
        elseif stp==7
            scfMRItb_04_resplitData(base_dir_sub, fname, ['_denoised' cutoff_str 'cov'], siz3) % if slice-wise data doesn't exist but combined 4D data exists, then split the data to generate the needed slice-wise data
            scfMRItb_04_unzipFile(base_dir_sub, fname, ['_slice' num2str(i3) '_denoised' cutoff_str 'cov'])
            F5 = load_untouch_nii([base_dir_sub fname '_slice' num2str(i3) '_denoised' cutoff_str 'cov.nii']); F5 = single(squeeze(F5.img));
        elseif stp==8
            scfMRItb_04_resplitData(base_dir_sub, fname, ['_denoised' cutoff_str 'WMcov'], siz3) % if slice-wise data doesn't exist but combined 4D data exists, then split the data to generate the needed slice-wise data
            scfMRItb_04_unzipFile(base_dir_sub, fname, ['_slice' num2str(i3) '_denoised' cutoff_str 'WMcov'])
            F5 = load_untouch_nii([base_dir_sub fname '_slice' num2str(i3) '_denoised' cutoff_str 'WMcov.nii']); F5 = single(squeeze(F5.img));
        end
        for i5=1:size(F5,3)
            tmp = F5(:,:,i5);
            F6g(:,i5) = tmp(find(mask_GM(:,:,i3)==1));
            F6w(:,i5) = tmp(find(mask_WM(:,:,i3)==1));
            clear tmp
        end; clear i5 F5
        mF6g=mean(F6g,1); mF6w=mean(F6w,1); 
        mF6g=zscore(mF6g); mF6w=zscore(mF6w);
        fighndl = figure('Position',[1 1 1152 720],'Color',[0.95,0.9,0.85],'InvertHardcopy','off','Visible','off'); % red [0.95,0.9,0.85], green [0.85,0.90,0.85], blue [0.89,0.94,0.99]
        if stp==1
            subplot2n(4,1,1), plot(mF6g,'color','k','linewidth',1.5), hold on, plot(mF2g,'color',[0,0.55,0.69],'linewidth',0.2), ylabel('mean time series'),
                grid on,grid minor, xlim([1 length(mF2g)]),ylim([min(1.25.*[mF2g,mF6g]) 1.25.*max([mF2g,mF6g])]), title(['Unfiltered: normalized mean GRAY MATTER signal AFTER "11-func-anat-registration" (slice ' num2str(i3) ' of ' num2str(siz3) ')'],'Color','k','FontSize',14), legend({'Gray matter signal before filtering'},'FontSize',12,'TextColor','k','Location','best')
            subplot2n(4,1,3), plot(mF6w,'color','k','linewidth',1.5), hold on, plot(mF2w,'color',[0.65,0.11,0.19],'linewidth',0.2), ylabel('mean time series'),
                grid on,grid minor, xlim([1 length(mF2g)]),ylim([min(1.25.*[mF2w,mF6w]) 1.25.*max([mF2w,mF6w])]), title(['Unfiltered: normalized mean WHITE MATTER signal AFTER "11-func-anat-registration" (slice ' num2str(i3) ' of ' num2str(siz3) ')'],'Color','k','FontSize',14), legend({'White matter signal before filtering'},'FontSize',12,'TextColor','k','Location','best')
        elseif stp==3
            subplot2n(4,1,1), plot(mF6g,'color','k','linewidth',1.5), hold on, plot(mF2g,'color',[0,0.55,0.69],'linewidth',0.2), ylabel('mean time series'),
                grid on,grid minor, xlim([1 length(mF2g)]),ylim([min(1.25.*[mF2g,mF6g]) 1.25.*max([mF2g,mF6g])]), title(['Unfiltered: normalized mean GRAY MATTER signal AFTER "12-denoise3-CSFregress" (slice ' num2str(i3) ' of ' num2str(siz3) ')'],'Color','k','FontSize',14), legend({'Gray matter signal before filtering'},'FontSize',12,'TextColor','k','Location','best')
            subplot2n(4,1,3), plot(mF6w,'color','k','linewidth',1.5), hold on, plot(mF2w,'color',[0.65,0.11,0.19],'linewidth',0.2), ylabel('mean time series'),
                grid on,grid minor, xlim([1 length(mF2g)]),ylim([min(1.25.*[mF2w,mF6w]) 1.25.*max([mF2w,mF6w])]), title(['Unfiltered: normalized mean WHITE MATTER signal AFTER "12-denoise3-CSFregress" (slice ' num2str(i3) ' of ' num2str(siz3) ')'],'Color','k','FontSize',14), legend({'White matter signal before filtering'},'FontSize',12,'TextColor','k','Location','best')
        elseif (stp==2)||(stp==6)
            subplot2n(4,1,1), plot(mF6g,'color','k','linewidth',1.5), hold on, plot(mF2g,'color',[0,0.55,0.69],'linewidth',0.2), ylabel('mean time series'),
                grid on,grid minor, xlim([1 length(mF2g)]),ylim([min(1.25.*[mF2g,mF6g]) 1.25.*max([mF2g,mF6g])]), title(['Unfiltered: normalized mean GRAY MATTER signal AFTER "13-denoise4-WMregress" (slice ' num2str(i3) ' of ' num2str(siz3) ')'],'Color','k','FontSize',14), legend({'Gray matter signal before filtering'},'FontSize',12,'TextColor','k','Location','best')
            subplot2n(4,1,3), plot(mF6w,'color','k','linewidth',1.5), hold on, plot(mF2w,'color',[0.65,0.11,0.19],'linewidth',0.2), ylabel('mean time series'),
                grid on,grid minor, xlim([1 length(mF2g)]),ylim([min(1.25.*[mF2w,mF6w]) 1.25.*max([mF2w,mF6w])]), title(['Unfiltered: normalized mean WHITE MATTER signal AFTER "13-denoise4-WMregress" (slice ' num2str(i3) ' of ' num2str(siz3) ')'],'Color','k','FontSize',14), legend({'White matter signal before filtering'},'FontSize',12,'TextColor','k','Location','best')
        elseif (stp==4)||(stp==5)||(stp==7)||(stp==8)
            subplot2n(4,1,1), plot(mF6g,'color','k','linewidth',1.5), hold on, plot(mF2g,'color',[0,0.55,0.69],'linewidth',0.2), ylabel('mean time series'),
                grid on,grid minor, xlim([1 length(mF2g)]),ylim([min(1.25.*[mF2g,mF6g]) 1.25.*max([mF2g,mF6g])]), title(['Unfiltered: normalized mean GRAY MATTER signal AFTER "14-denoise5-COVregress" (slice ' num2str(i3) ' of ' num2str(siz3) ')'],'Color','k','FontSize',14), legend({'Gray matter signal before filtering'},'FontSize',12,'TextColor','k','Location','best')
            subplot2n(4,1,3), plot(mF6w,'color','k','linewidth',1.5), hold on, plot(mF2w,'color',[0.65,0.11,0.19],'linewidth',0.2), ylabel('mean time series'),
                grid on,grid minor, xlim([1 length(mF2g)]),ylim([min(1.25.*[mF2w,mF6w]) 1.25.*max([mF2w,mF6w])]), title(['Unfiltered: normalized mean WHITE MATTER signal AFTER "14-denoise5-COVregress" (slice ' num2str(i3) ' of ' num2str(siz3) ')'],'Color','k','FontSize',14), legend({'White matter signal before filtering'},'FontSize',12,'TextColor','k','Location','best')
        end
        subplot2n(4,1,2), plot(mF2g,'color',[0,0.55,0.69],'linewidth',1.5), hold on, plot(mF6g,'color','k','linewidth',0.2), ylabel('mean time series'),
            grid on,grid minor, xlim([1 length(mF2g)]),ylim([min(1.25.*[mF2g,mF6g]) 1.25.*max([mF2g,mF6g])]), title(['Bandpass filtered: normalized mean GRAY MATTER signal AFTER "15-bandpass-filter" [0.01-' num2str(f_BPF_max_vec(1)) 'Hz] (slice ' num2str(i3) ' of ' num2str(siz3) ')'],'Color',[0,0.55,0.69],'FontSize',14), legend({'Gray matter signal after filtering'},'FontSize',12,'TextColor',[0,0.55,0.69],'Location','best')
        subplot2n(4,1,4), plot(mF2w,'color',[0.65,0.11,0.19],'linewidth',1.5), hold on, plot(mF6w,'color','k','linewidth',0.2), ylabel('mean time series'),
            grid on,grid minor, xlim([1 length(mF2g)]),ylim([min(1.25.*[mF2w,mF6w]) 1.25.*max([mF2w,mF6w])]), title(['Bandpass filtered: normalized mean WHITE MATTER signal AFTER "15-bandpass-filter" [0.01-' num2str(f_BPF_max_vec(1)) 'Hz] (slice ' num2str(i3) ' of ' num2str(siz3) ')'],'Color',[0.65,0.11,0.19],'FontSize',14), legend({'White matter signal after filtering'},'FontSize',12,'TextColor',[0.65,0.11,0.19],'Location','best')
        saveas(fighndl,[QCpath 'mean_timeseries_14denoise5_vs_15filter_slice' sprintf('%.2d',i3) '.jpg'])
        close(fighndl)
        
        C3g = corrcoef(F6g'); C3g = uppertriangle(C3g); C3w = corrcoef(F6w'); C3w = uppertriangle(C3w);
        fighndl = figure('Position',[1 1 1152 720],'Color',[0.95,0.9,0.85],'InvertHardcopy','off','Visible','off'); % red [0.95,0.9,0.85], green [0.85,0.90,0.85], blue [0.89,0.94,0.99]
        if stp==1
            subplot2n(4,1,1), histogram(C3g,'EdgeColor','none','FaceColor','k','FaceAlpha',0.5,'Normalization','probability'); grid on,grid minor, xlim([-1 1]), hold on, xline(0,'Color','k','linewidth',1.5); hold off
                title(sprintf('Histogram of voxel-to-voxel GRAY MATTER correlations AFTER 11-func-anat-registration (slice %d of %d)\n%.1f%% of the correlations were > 0  [ mean=%.2f, median=%.2f, SD=%.2f ]',i3,siz3,100*length(find(C3g>0))/length(C3g),mean(C3g),median(C3g),std(C3g)),'Color','k','FontSize',14)
            subplot2n(4,1,3), histogram(C3w,'EdgeColor','none','FaceColor','k','FaceAlpha',0.5,'Normalization','probability'); grid on,grid minor, xlim([-1 1]), hold on, xline(0,'Color','k','linewidth',1.5); hold off
                title(sprintf('WHITE MATTER: AFTER 11-func-anat-registration (slice %d of %d): %.1f%% of the correlations were > 0  [ mean=%.2f, median=%.2f, SD=%.2f ]',i3,siz3,100*length(find(C3w>0))/length(C3w),mean(C3w),median(C3w),std(C3w)),'Color','k','FontSize',14)
        elseif stp==3
            subplot2n(4,1,1), histogram(C3g,'EdgeColor','none','FaceColor','k','FaceAlpha',0.5,'Normalization','probability'); grid on,grid minor, xlim([-1 1]), hold on, xline(0,'Color','k','linewidth',1.5); hold off
                title(sprintf('Histogram of voxel-to-voxel GRAY MATTER correlations AFTER 12-denoise3-CSFregress (slice %d of %d)\n%.1f%% of the correlations were > 0  [ mean=%.2f, median=%.2f, SD=%.2f ]',i3,siz3,100*length(find(C3g>0))/length(C3g),mean(C3g),median(C3g),std(C3g)),'Color','k','FontSize',14)
            subplot2n(4,1,3), histogram(C3w,'EdgeColor','none','FaceColor','k','FaceAlpha',0.5,'Normalization','probability'); grid on,grid minor, xlim([-1 1]), hold on, xline(0,'Color','k','linewidth',1.5); hold off
                title(sprintf('WHITE MATTER: AFTER 12-denoise3-CSFregress (slice %d of %d): %.1f%% of the correlations were > 0  [ mean=%.2f, median=%.2f, SD=%.2f ]',i3,siz3,100*length(find(C3w>0))/length(C3w),mean(C3w),median(C3w),std(C3w)),'Color','k','FontSize',14)
        elseif (stp==2)||(stp==6)
            subplot2n(4,1,1), histogram(C3g,'EdgeColor','none','FaceColor','k','FaceAlpha',0.5,'Normalization','probability'); grid on,grid minor, xlim([-1 1]), hold on, xline(0,'Color','k','linewidth',1.5); hold off
                title(sprintf('Histogram of voxel-to-voxel GRAY MATTER correlations AFTER 13-denoise4-WMregress (slice %d of %d)\n%.1f%% of the correlations were > 0  [ mean=%.2f, median=%.2f, SD=%.2f ]',i3,siz3,100*length(find(C3g>0))/length(C3g),mean(C3g),median(C3g),std(C3g)),'Color','k','FontSize',14)
            subplot2n(4,1,3), histogram(C3w,'EdgeColor','none','FaceColor','k','FaceAlpha',0.5,'Normalization','probability'); grid on,grid minor, xlim([-1 1]), hold on, xline(0,'Color','k','linewidth',1.5); hold off
                title(sprintf('WHITE MATTER: AFTER 13-denoise4-WMregress (slice %d of %d): %.1f%% of the correlations were > 0  [ mean=%.2f, median=%.2f, SD=%.2f ]',i3,siz3,100*length(find(C3w>0))/length(C3w),mean(C3w),median(C3w),std(C3w)),'Color','k','FontSize',14)
        elseif (stp==4)||(stp==5)||(stp==7)||(stp==8)
            subplot2n(4,1,1), histogram(C3g,'EdgeColor','none','FaceColor','k','FaceAlpha',0.5,'Normalization','probability'); grid on,grid minor, xlim([-1 1]), hold on, xline(0,'Color','k','linewidth',1.5); hold off
                title(sprintf('Histogram of voxel-to-voxel GRAY MATTER correlations AFTER 14-denoise5-COVregress (slice %d of %d)\n%.1f%% of the correlations were > 0  [ mean=%.2f, median=%.2f, SD=%.2f ]',i3,siz3,100*length(find(C3g>0))/length(C3g),mean(C3g),median(C3g),std(C3g)),'Color','k','FontSize',14)
            subplot2n(4,1,3), histogram(C3w,'EdgeColor','none','FaceColor','k','FaceAlpha',0.5,'Normalization','probability'); grid on,grid minor, xlim([-1 1]), hold on, xline(0,'Color','k','linewidth',1.5); hold off
                title(sprintf('WHITE MATTER: AFTER 14-denoise5-COVregress (slice %d of %d): %.1f%% of the correlations were > 0  [ mean=%.2f, median=%.2f, SD=%.2f ]',i3,siz3,100*length(find(C3w>0))/length(C3w),mean(C3w),median(C3w),std(C3w)),'Color','k','FontSize',14)
        end
        subplot2n(4,1,2), histogram(Cg,'EdgeColor','none','FaceColor',[0,0.55,0.69],'FaceAlpha',0.5,'Normalization','probability'); grid on,grid minor, xlim([-1 1]), hold on, xline(0,'Color','k','linewidth',1.5); hold off
            title(sprintf('GRAY MATTER: AFTER 15-bandpass-filter (slice %d of %d): %.1f%% of the correlations were > 0  [ mean=%.2f, median=%.2f, SD=%.2f ]',i3,siz3,100*length(find(Cg>0))/length(Cg),mean(Cg),median(Cg),std(Cg)),'Color',[0,0.55,0.69],'FontSize',14)
        subplot2n(4,1,4), histogram(Cw,'EdgeColor','none','FaceColor',[0.65,0.11,0.19],'FaceAlpha',0.5,'Normalization','probability'); grid on,grid minor, xlim([-1 1]), hold on, xline(0,'Color','k','linewidth',1.5); hold off
            title(sprintf('WHITE MATTER: AFTER 15-bandpass-filter (slice %d of %d): %.1f%% of the correlations were > 0  [ mean=%.2f, median=%.2f, SD=%.2f ]',i3,siz3,100*length(find(Cw>0))/length(Cw),mean(Cw),median(Cw),std(Cw)),'Color',[0.65,0.11,0.19],'FontSize',14)
        saveas(fighndl,[QCpath 'hist_FC_compare_14denoise5_15filter_slice' sprintf('%.2d',i3) '.jpg'])
        close(fighndl)
        F6g_=cat(1,F6g_,F6g); mF6g_=cat(1,mF6g_,mF6g); C3g_=cat(1,C3g_,C3g);
        F6w_=cat(1,F6w_,F6w); mF6w_=cat(1,mF6w_,mF6w); C3w_=cat(1,C3w_,C3w);
        clear F6g F6w mF6g mF6w C3g C3w
    end
    
    F2g_=cat(1,F2g_,F2g); mF2g_=cat(1,mF2g_,mF2g); F2w_=cat(1,F2w_,F2w); mF2w_=cat(1,mF2w_,mF2w); Cg_=cat(1,Cg_,Cg); Cw_=cat(1,Cw_,Cw); F4_=cat(1,F4_,F4); C2_=cat(1,C2_,C2);
    clear F2g F2w F4 TR Smask1 svar1 svar2 mF2g mF2w mF4 Cg Cw C2
end


[~,svar1] = sort(var(F2g_,0,2)); [~,spsc1] = sort(meanPSC(F2g_));
[~,svar2] = sort(var(F2w_,0,2)); [~,spsc2] = sort(meanPSC(F2w_));
sortorder1b = sortorder1; sortorder1=[]; f1=0; %#ok<*ASGLU>
sortorder2b = sortorder2; sortorder2=[]; f2=0; %#ok<*ASGLU>
for im=1:length(sortorder1b)
    temp = sortorder1b{im}; temp = temp + f1; f1 = f1 + length(temp);
    sortorder1 = cat(1,sortorder1,temp); clear temp
    temp = sortorder2b{im}; temp = temp + f2; f2 = f2 + length(temp);
    sortorder2 = cat(1,sortorder2,temp); clear temp
end; clear im f1 f2
sortorder1 = spsc1; % options: spsc for percentage signal change, svar for variance, (comment out the line if you want to use distance from the center of the cord separately for each slice)
sortorder2 = spsc2; % options: spsc for percentage signal change, svar for variance, (comment out the line if you want to use distance from the center of the cord separately for each slice)
for im=max(sortorder1):-1:1, if isempty(find(sortorder1==im)), sortorder1(find(sortorder1>im)) = sortorder1(find(sortorder1>im))-1; end, end; clear im %#ok<*EFIND>
for im=max(sortorder2):-1:1, if isempty(find(sortorder2==im)), sortorder2(find(sortorder2>im)) = sortorder2(find(sortorder2>im))-1; end, end; clear im %#ok<*EFIND>

fighndl = figure('Position',[1 1 1152 720],'Color',[0.89,0.94,0.99],'InvertHardcopy','off','Visible','off'); % red [0.95,0.9,0.85], green [0.85,0.90,0.85], blue [0.89,0.94,0.99]
subplot2n(2,1,1), imagesc(F2g_(sortorder1,:)),colormap(cmap),colorbar, xlabel('time (in samples)'),ylabel(sprintf('voxels (sorted by mean percentage signal change;\nbottom row is highest)')),
  title(['Time series within the GRAY MATTER AFTER 15-bandpass-filter (all slices combined)'],'Color',[0,0.55,0.69],'FontSize',14);
subplot2n(2,1,2), imagesc(F2w_(sortorder2,:)),colormap(cmap),colorbar, xlabel('time (in samples)'),ylabel(sprintf('voxels (sorted by mean percentage signal change;\nbottom row is highest)')),
  title(['Time series within the WHITE MATTER AFTER 15-bandpass-filter (all slices combined)'],'Color',[0.65,0.11,0.19],'FontSize',14);
saveas(fighndl,[QCpath 'fMRI_ts_15filter_allSlices' '.jpg'])
saveas(fighndl,[QCpath2 'fMRI_ts_15filter_allSlices' '.jpg'])
 close(fighndl)
mF2g_=mean(mF2g_,1); mF2w_=mean(mF2w_,1); mF4_=mean(mF4_,1);
fighndl = figure('Position',[1 1 1152 720],'Color',[0.89,0.94,0.99],'InvertHardcopy','off','Visible','off'); % red [0.95,0.9,0.85], green [0.85,0.90,0.85], blue [0.89,0.94,0.99]
subplot2n(3,1,1), histogram(C2_,'EdgeColor','none','FaceColor','k','FaceAlpha',0.5,'Normalization','probability'); grid on,grid minor, xlim([-1 1]), hold on, xline(0,'Color','k','linewidth',1.5); hold off
    title(sprintf('Histogram of voxel-to-voxel correlations in UNPROCESSED RAW DATA (all slices combined)\n%.1f%% of the correlations were > 0  [ mean=%.2f, median=%.2f, SD=%.2f ]',100*length(find(C2_>0))/length(C2_),mean(C2_),median(C2_),std(C2_)),'Color','k','FontSize',14)
subplot2n(3,1,2), histogram(Cg_,'EdgeColor','none','FaceColor',[0,0.55,0.69],'FaceAlpha',0.5,'Normalization','probability'); grid on,grid minor, xlim([-1 1]), hold on, xline(0,'Color','k','linewidth',1.5); hold off
    title(sprintf('Histogram of voxel-to-voxel GRAY MATTER correlations AFTER 15-bandpass-filter [0.01-%.2fHz] (all slices combined)\n%.1f%% of the correlations were > 0  [ mean=%.2f, median=%.2f, SD=%.2f ]',f_BPF_max_vec(1),100*length(find(Cg_>0))/length(Cg_),mean(Cg_),median(Cg_),std(Cg_)),'Color',[0,0.55,0.69],'FontSize',14)
subplot2n(3,1,3), histogram(Cw_,'EdgeColor','none','FaceColor',[0.65,0.11,0.19],'FaceAlpha',0.5,'Normalization','probability'); grid on,grid minor, xlim([-1 1]), hold on, xline(0,'Color','k','linewidth',1.5); hold off
    title(sprintf('Histogram of voxel-to-voxel WHITE MATTER correlations AFTER 15-bandpass-filter [0.01-%.2fHz] (all slices combined)\n%.1f%% of the correlations were > 0  [ mean=%.2f, median=%.2f, SD=%.2f ]',f_BPF_max_vec(1),100*length(find(Cw_>0))/length(Cw_),mean(Cw_),median(Cw_),std(Cw_)),'Color',[0.65,0.11,0.19],'FontSize',14)
saveas(fighndl,[QCpath 'hist_FC_compare_15filter_allSlices' '.jpg'])
 close(fighndl)

if stp~=0
    mF6g_=mean(mF6g_,1); mF6w_=mean(mF6w_,1);
    fighndl = figure('Position',[1 1 1152 720],'Color',[0.89,0.94,0.99],'InvertHardcopy','off','Visible','off'); % red [0.95,0.9,0.85], green [0.85,0.90,0.85], blue [0.89,0.94,0.99]
    if stp==1
    	subplot2n(4,1,1), plot(mF6g_,'color','k','linewidth',1.5), hold on, plot(mF2g_,'color',[0,0.55,0.69],'linewidth',0.2), ylabel('mean time series'),
            grid on,grid minor, xlim([1 length(mF2g_)]),ylim([1.25.*min([mF2g_,mF6g_]) 1.25.*max([mF2g_,mF6g_])]), title(['Unfiltered: normalized mean GRAY MATTER signal AFTER "11-func-anat-registration" (all slices averaged)'],'Color','k','FontSize',14), legend({'Gray matter signal before filtering'},'FontSize',12,'TextColor','k','Location','best')
        subplot2n(4,1,3), plot(mF6w_,'color','k','linewidth',1.5), hold on, plot(mF2w_,'color',[0.65,0.11,0.19],'linewidth',0.2), ylabel('mean time series'),
            grid on,grid minor, xlim([1 length(mF2g_)]),ylim([1.25.*min([mF2w_,mF6w_]) 1.25.*max([mF2w_,mF6w_])]), title(['Unfiltered: normalized mean WHITE MATTER signal AFTER "11-func-anat-registration" (all slices averaged)'],'Color','k','FontSize',14), legend({'White matter signal before filtering'},'FontSize',12,'TextColor','k','Location','best')
    elseif stp==3
    	subplot2n(4,1,1), plot(mF6g_,'color','k','linewidth',1.5), hold on, plot(mF2g_,'color',[0,0.55,0.69],'linewidth',0.2), ylabel('mean time series'),
            grid on,grid minor, xlim([1 length(mF2g_)]),ylim([1.25.*min([mF2g_,mF6g_]) 1.25.*max([mF2g_,mF6g_])]), title(['Unfiltered: normalized mean GRAY MATTER signal AFTER "12-DENOISE3-CSFregress" (all slices averaged)'],'Color','k','FontSize',14), legend({'Gray matter signal before filtering'},'FontSize',12,'TextColor','k','Location','best')
        subplot2n(4,1,3), plot(mF6w_,'color','k','linewidth',1.5), hold on, plot(mF2w_,'color',[0.65,0.11,0.19],'linewidth',0.2), ylabel('mean time series'),
            grid on,grid minor, xlim([1 length(mF2g_)]),ylim([1.25.*min([mF2w_,mF6w_]) 1.25.*max([mF2w_,mF6w_])]), title(['Unfiltered: normalized mean WHITE MATTER signal AFTER "12-DENOISE3-CSFregress" (all slices averaged)'],'Color','k','FontSize',14), legend({'White matter signal before filtering'},'FontSize',12,'TextColor','k','Location','best')
    elseif (stp==2)||(stp==6)
    	subplot2n(4,1,1), plot(mF6g_,'color','k','linewidth',1.5), hold on, plot(mF2g_,'color',[0,0.55,0.69],'linewidth',0.2), ylabel('mean time series'),
            grid on,grid minor, xlim([1 length(mF2g_)]),ylim([1.25.*min([mF2g_,mF6g_]) 1.25.*max([mF2g_,mF6g_])]), title(['Unfiltered: normalized mean GRAY MATTER signal AFTER "13-DENOISE4-WMregress" (all slices averaged)'],'Color','k','FontSize',14), legend({'Gray matter signal before filtering'},'FontSize',12,'TextColor','k','Location','best')
        subplot2n(4,1,3), plot(mF6w_,'color','k','linewidth',1.5), hold on, plot(mF2w_,'color',[0.65,0.11,0.19],'linewidth',0.2), ylabel('mean time series'),
            grid on,grid minor, xlim([1 length(mF2g_)]),ylim([1.25.*min([mF2w_,mF6w_]) 1.25.*max([mF2w_,mF6w_])]), title(['Unfiltered: normalized mean WHITE MATTER signal AFTER "13-DENOISE4-WMregress" (all slices averaged)'],'Color','k','FontSize',14), legend({'White matter signal before filtering'},'FontSize',12,'TextColor','k','Location','best')
    elseif (stp==4)||(stp==5)||(stp==7)||(stp==8)
    	subplot2n(4,1,1), plot(mF6g_,'color','k','linewidth',1.5), hold on, plot(mF2g_,'color',[0,0.55,0.69],'linewidth',0.2), ylabel('mean time series'),
            grid on,grid minor, xlim([1 length(mF2g_)]),ylim([1.25.*min([mF2g_,mF6g_]) 1.25.*max([mF2g_,mF6g_])]), title(['Unfiltered: normalized mean GRAY MATTER signal AFTER "14-DENOISE5-COVregress" (all slices averaged)'],'Color','k','FontSize',14), legend({'Gray matter signal before filtering'},'FontSize',12,'TextColor','k','Location','best')
        subplot2n(4,1,3), plot(mF6w_,'color','k','linewidth',1.5), hold on, plot(mF2w_,'color',[0.65,0.11,0.19],'linewidth',0.2), ylabel('mean time series'),
            grid on,grid minor, xlim([1 length(mF2g_)]),ylim([1.25.*min([mF2w_,mF6w_]) 1.25.*max([mF2w_,mF6w_])]), title(['Unfiltered: normalized mean WHITE MATTER signal AFTER "14-DENOISE5-COVregress" (all slices averaged)'],'Color','k','FontSize',14), legend({'White matter signal before filtering'},'FontSize',12,'TextColor','k','Location','best')
    end
    subplot2n(4,1,2), plot(mF2g_,'color',[0,0.55,0.69],'linewidth',1.5), hold on, plot(mF6g_,'color','k','linewidth',0.2), ylabel('mean time series'),
        grid on,grid minor, xlim([1 length(mF2g_)]),ylim([1.25.*min([mF2g_,mF6g_]) 1.25.*max([mF2g_,mF6g_])]), title(['Bandpass filtered: normalized mean GRAY MATTER signal AFTER "15-bandpass-filter" (all slices averaged)'],'Color',[0,0.55,0.69],'FontSize',14), legend({'Gray matter signal after filtering'},'FontSize',12,'TextColor',[0,0.55,0.69],'Location','best')
    subplot2n(4,1,4), plot(mF2w_,'color',[0.65,0.11,0.19],'linewidth',1.5), hold on, plot(mF6w_,'color','k','linewidth',0.2), ylabel('mean time series'),
        grid on,grid minor, xlim([1 length(mF2g_)]),ylim([1.25.*min([mF2w_,mF6w_]) 1.25.*max([mF2w_,mF6w_])]), title(['Bandpass filtered: normalized mean WHITE MATTER signal AFTER "15-bandpass-filter" (all slices averaged)'],'Color',[0.65,0.11,0.19],'FontSize',14), legend({'White matter signal after filtering'},'FontSize',12,'TextColor',[0.65,0.11,0.19],'Location','best')
    saveas(fighndl,[QCpath 'mean_timeseries_14denoise5_vs_15filter_allSlices' '.jpg'])
    saveas(fighndl,[QCpath2 'global_signal_14denoise5_vs_15filter_allSlices' '.jpg'])
     close(fighndl)
    
    fighndl = figure('Position',[1 1 1152 720],'Color',[0.89,0.94,0.99],'InvertHardcopy','off','Visible','off'); % red [0.95,0.9,0.85], green [0.85,0.90,0.85], blue [0.89,0.94,0.99]
    if stp==1
    	subplot2n(4,1,1), histogram(C3g_,'EdgeColor','none','FaceColor','k','FaceAlpha',0.5,'Normalization','probability'); grid on,grid minor, xlim([-1 1]), hold on, xline(0,'Color','k','linewidth',1.5); hold off
            title(sprintf('Histogram of voxel-to-voxel GRAY MATTER correlations AFTER 11-registration (all slices combined)\n%.1f%% of the correlations were > 0  [ mean=%.2f, median=%.2f, SD=%.2f ]',100*length(find(C3g_>0))/length(C3g_),mean(C3g_),median(C3g_),std(C3g_)),'Color','k','FontSize',14)
        subplot2n(4,1,3), histogram(C3w_,'EdgeColor','none','FaceColor','k','FaceAlpha',0.5,'Normalization','probability'); grid on,grid minor, xlim([-1 1]), hold on, xline(0,'Color','k','linewidth',1.5); hold off
            title(sprintf('WHITE MATTER: AFTER 11-registration (all slices): %.1f%% of the correlations were > 0  [ mean=%.2f, median=%.2f, SD=%.2f ]',100*length(find(C3w_>0))/length(C3w_),mean(C3w_),median(C3w_),std(C3w_)),'Color','k','FontSize',14)
    elseif stp==3
    	subplot2n(4,1,1), histogram(C3g_,'EdgeColor','none','FaceColor','k','FaceAlpha',0.5,'Normalization','probability'); grid on,grid minor, xlim([-1 1]), hold on, xline(0,'Color','k','linewidth',1.5); hold off
            title(sprintf('Histogram of voxel-to-voxel GRAY MATTER correlations AFTER 12-denoise3-CSFregress (all slices combined)\n%.1f%% of the correlations were > 0  [ mean=%.2f, median=%.2f, SD=%.2f ]',100*length(find(C3g_>0))/length(C3g_),mean(C3g_),median(C3g_),std(C3g_)),'Color','k','FontSize',14)
        subplot2n(4,1,3), histogram(C3w_,'EdgeColor','none','FaceColor','k','FaceAlpha',0.5,'Normalization','probability'); grid on,grid minor, xlim([-1 1]), hold on, xline(0,'Color','k','linewidth',1.5); hold off
            title(sprintf('WHITE MATTER: AFTER 12-denoise3-CSFregress (all slices): %.1f%% of the correlations were > 0  [ mean=%.2f, median=%.2f, SD=%.2f ]',100*length(find(C3w_>0))/length(C3w_),mean(C3w_),median(C3w_),std(C3w_)),'Color','k','FontSize',14)
    elseif (stp==2)||(stp==6)
    	subplot2n(4,1,1), histogram(C3g_,'EdgeColor','none','FaceColor','k','FaceAlpha',0.5,'Normalization','probability'); grid on,grid minor, xlim([-1 1]), hold on, xline(0,'Color','k','linewidth',1.5); hold off
            title(sprintf('Histogram of voxel-to-voxel GRAY MATTER correlations AFTER 13-denoise4-WMregress (all slices combined)\n%.1f%% of the correlations were > 0  [ mean=%.2f, median=%.2f, SD=%.2f ]',100*length(find(C3g_>0))/length(C3g_),mean(C3g_),median(C3g_),std(C3g_)),'Color','k','FontSize',14)
        subplot2n(4,1,3), histogram(C3w_,'EdgeColor','none','FaceColor','k','FaceAlpha',0.5,'Normalization','probability'); grid on,grid minor, xlim([-1 1]), hold on, xline(0,'Color','k','linewidth',1.5); hold off
            title(sprintf('WHITE MATTER: AFTER 13-denoise4-WMregress (all slices): %.1f%% of the correlations were > 0  [ mean=%.2f, median=%.2f, SD=%.2f ]',100*length(find(C3w_>0))/length(C3w_),mean(C3w_),median(C3w_),std(C3w_)),'Color','k','FontSize',14)
    elseif (stp==4)||(stp==5)||(stp==7)||(stp==8)
    	subplot2n(4,1,1), histogram(C3g_,'EdgeColor','none','FaceColor','k','FaceAlpha',0.5,'Normalization','probability'); grid on,grid minor, xlim([-1 1]), hold on, xline(0,'Color','k','linewidth',1.5); hold off
            title(sprintf('Histogram of voxel-to-voxel GRAY MATTER correlations AFTER 14-denoise5-COVregress (all slices combined)\n%.1f%% of the correlations were > 0  [ mean=%.2f, median=%.2f, SD=%.2f ]',100*length(find(C3g_>0))/length(C3g_),mean(C3g_),median(C3g_),std(C3g_)),'Color','k','FontSize',14)
        subplot2n(4,1,3), histogram(C3w_,'EdgeColor','none','FaceColor','k','FaceAlpha',0.5,'Normalization','probability'); grid on,grid minor, xlim([-1 1]), hold on, xline(0,'Color','k','linewidth',1.5); hold off
            title(sprintf('WHITE MATTER: AFTER 14-denoise5-COVregress (all slices): %.1f%% of the correlations were > 0  [ mean=%.2f, median=%.2f, SD=%.2f ]',100*length(find(C3w_>0))/length(C3w_),mean(C3w_),median(C3w_),std(C3w_)),'Color','k','FontSize',14)
    end
    subplot2n(4,1,2), histogram(Cg_,'EdgeColor','none','FaceColor',[0,0.55,0.69],'FaceAlpha',0.5,'Normalization','probability'); grid on,grid minor, xlim([-1 1]), hold on, xline(0,'Color','k','linewidth',1.5); hold off
       title(sprintf('GRAY MATTER: AFTER 15-bandpass-filter (all slices): %.1f%% of correlations > 0  [ mean=%.2f, median=%.2f, SD=%.2f ]',100*length(find(Cg_>0))/length(Cg_),mean(Cg_),median(Cg_),std(Cg_)),'Color',[0,0.55,0.69],'FontSize',14)
    subplot2n(4,1,4), histogram(Cw_,'EdgeColor','none','FaceColor',[0.65,0.11,0.19],'FaceAlpha',0.5,'Normalization','probability'); grid on,grid minor, xlim([-1 1]), hold on, xline(0,'Color','k','linewidth',1.5); hold off
       title(sprintf('WHITE MATTER: AFTER 15-bandpass-filter (all slices): %.1f%% of the correlations were > 0  [ mean=%.2f, median=%.2f, SD=%.2f ]',100*length(find(Cw_>0))/length(Cw_),mean(Cw_),median(Cw_),std(Cw_)),'Color',[0.65,0.11,0.19],'FontSize',14)
    saveas(fighndl,[QCpath 'hist_FC_compare_14denoise5_15filter_allSlices' '.jpg'])
    saveas(fighndl,[QCpath2 'hist_FC_compare_14denoise5_15filter_allSlices' '.jpg'])
     close(fighndl)
end
clear svar1 spsc1 svar2 spsc2 sortorder1 sortorder2 F2g_ mF2g_ F2w_ mF2w_ Cg_ Cw_ F4_ mF4_ C2_ F6g_ F6w_ mF6g_ mF6w_ C3g_ C3w_ cmap

try
if nargin<6
    close(wbar3)
end
catch
end

end
```
