## Supplementary material for "Neptune: a toolbox for spinal cord functional MRI data processing and quality assurance": Neptune software: scfMRItb_16_cord_quadrants.m

```
function scfMRItb_16_cord_quadrants(base_dir_sub,fname,fname_anat_orig, varargin)

scrsz = get(0,'ScreenSize');
pause on;

fullsize = 0; % If fullsize=1 then the pre-processed data will not be cropped. By default fullsize=0, so the images are cropped (suggested because the cord is often just a small portion of the field of view [to avoid warping along the phase encoding direction], so the images can be safely cropped without losing the cord. Rest assured that there are checks to ensure that the cord is not cropped off.)

if nargin<6
    wbar3 = waitbar(0,'16. Define cord quadrants...','Name','Progress(16): Define cord quadrants...','Units','normalized','Position',[4/5 0 1/5 1/17]);
else
    wbar3 = varargin{3};
end

if ~(exist([base_dir_sub 'QC'],'dir'))
    mkdir([base_dir_sub 'QC'])
end
if ~(exist([base_dir_sub 'QC' '/16_cord_quadrants'],'dir'))
    mkdir([base_dir_sub 'QC' '/16_cord_quadrants'])
end
QCpath = [base_dir_sub 'QC' '/16_cord_quadrants' '/'];

if nargin<4
    scan7T = 1;
else
    scan7T = varargin{1};
end
if nargin<5
    C_spine = 1;
else
    C_spine = varargin{2};
end

scan3T = ~scan7T;
L_spine = ~C_spine;

fname_anat = [fname_anat_orig '_reduced'];


%% ----- PARAMETERS WITH WHICH NIFTI IMAGES WERE MADE SMALLER in step 14 (save 4D data) -------------------

scfMRItb_04_unzipFile(base_dir_sub, fname_anat, '')
A = load_untouch_nii([base_dir_sub fname_anat '.nii']);
siz3 = size(A.img,3);

scfMRItb_04_unzipFile(base_dir_sub, fname_anat, '_mask_GM')
scfMRItb_04_unzipFile(base_dir_sub, fname_anat, '_mask_WM')
scfMRItb_04_unzipFile(base_dir_sub, fname_anat, '_mask_CSF')
mmask1 = load_untouch_nii([base_dir_sub fname_anat '_mask_GM.nii']); % gray matter
mmask2 = load_untouch_nii([base_dir_sub fname_anat '_mask_WM.nii']); % white matter
mmask3 = load_untouch_nii([base_dir_sub fname_anat '_mask_CSF.nii']); % white matter
mmask_all = single(or(or(mmask1.img, mmask2.img), mmask3.img));

smmask = (mean(mmask_all,3) > 0); 
[s1,s2] = find(smmask~=0);
s1median = median(s1); s2median = median(s2);
s1min = min(s1); s2min = min(s2);
s1max = max(s1); s2max = max(s2);
smaxrange = max([s2max-s2min,s1max-s1min]);
if round(size(mmask_all,1)/4) > smaxrange
    new_size = round(size(mmask_all,1)/4); % 128 - assume even for now
elseif round(size(mmask_all,1)/3) > smaxrange
    new_size = round(size(mmask_all,1)/3); % 171
elseif round(size(mmask_all,1)/2) > smaxrange
    new_size = round(size(mmask_all,1)/2); % 256
else
    new_size = size(mmask_all,1); % 512
end
% new_size = round(size(mmask,1)/4); % 128 - assume even for now

if fullsize==1
    new_size = size(mmask_all,1);
end

xmin = s1median - round(new_size/2) + 1; xmax = s1median + round(new_size/2);
ymin = s2median - round(new_size/2) + 1; ymax = s2median + round(new_size/2);
new_size = xmax - xmin + 1;

mask_GM_reduced = mmask1; mask_WM_reduced = mmask2; mask_CSF_reduced = mmask3;
mask_GM_reduced.hdr.dime.dim(2)  = new_size; mask_GM_reduced.hdr.dime.dim(3)  = new_size;
mask_WM_reduced.hdr.dime.dim(2)  = new_size; mask_WM_reduced.hdr.dime.dim(3)  = new_size;
mask_CSF_reduced.hdr.dime.dim(2) = new_size; mask_CSF_reduced.hdr.dime.dim(3) = new_size;

basestr = strfind(base_dir_sub,'/');
basestr = base_dir_sub(basestr(end-1)+1:end); % name of subject folder
if str2num(basestr(1:4)) <= 2017 %#ok<ST2NM> % if data were acquired prior to 2018 then don't change the bounding box of the reduced size images that were already processed before
    mask_GM_reduced.img  = mmask1.img(round((size(A.img,1)-new_size)/2)+1:end-round((size(A.img,1)-new_size)/2), ...
        round((size(A.img,2)-new_size)/2)+1:end-round((size(A.img,2)-new_size)/2), :);
    mask_WM_reduced.img  = mmask2.img(round((size(A.img,1)-new_size)/2)+1:end-round((size(A.img,1)-new_size)/2), ...
        round((size(A.img,2)-new_size)/2)+1:end-round((size(A.img,2)-new_size)/2), :);
    mask_CSF_reduced.img = mmask3.img(round((size(A.img,1)-new_size)/2)+1:end-round((size(A.img,1)-new_size)/2), ...
        round((size(A.img,2)-new_size)/2)+1:end-round((size(A.img,2)-new_size)/2), :); %#ok<*NASGU>
else % else if data were acquired after 2017 then use the new method to determine the bounding box by centering the new box around the centroid of the spinal cord mask
    mask_GM_reduced.img  = mmask1.img(xmin:xmax, ymin:ymax, :);
    mask_WM_reduced.img  = mmask2.img(xmin:xmax, ymin:ymax, :);
    mask_CSF_reduced.img = mmask3.img(xmin:xmax, ymin:ymax, :);
end

FC_mask_quads = mask_GM_reduced; FC_mask_quads.img=[];
pixdims(1) = mask_GM_reduced.hdr.dime.pixdim(2); pixdims(2) = mask_GM_reduced.hdr.dime.pixdim(3); % in mm
anat_reduced2 = A; anat_reduced2.hdr.dime.dim(2)  = new_size; anat_reduced2.hdr.dime.dim(3)  = new_size;
anat_reduced2.img = A.img(xmin:xmax, ymin:ymax, :);


%% ----- FIND WHITE/GRAY MATTER BOUNDARIES ---------------------------------

% find gray matter boundary
anat_orig = mask_GM_reduced.img;
anat_edges_GM = zeros(size(anat_orig));
for i1 = 1 : siz3
    try waitbar((i1/siz3),wbar3,sprintf('16. Initial housekeeping (find gray matter boundary): slice (%d) of (%d)',i1,siz3)); catch, end
    disp(sprintf('16. Initial housekeeping (find gray matter boundary): slice (%d) of (%d)',i1,siz3))
    anat_mask = double(anat_orig(:,:,i1) > 0);
    anat_edges_GM(:,:,i1) = double(edge(anat_mask, 'zerocross')); clear anat_mask
end

% find white matter boundary
anat_edges_WM = zeros(size(anat_orig));
temp = (mask_WM_reduced.img + mask_GM_reduced.img) > 0;
for i1 = 1 : siz3
    try waitbar((i1/siz3),wbar3,sprintf('16. Initial housekeeping (find white matter boundary): slice (%d) of (%d)',i1,siz3)); catch, end
    disp(sprintf('16. Initial housekeeping (find white matter boundary): slice (%d) of (%d)',i1,siz3))
    ttemp = temp(:,:,i1);
    ttemp_b = bwboundaries(ttemp);
    ttemp_b = ttemp_b{1};
    ttemp2 = anat_edges_WM(:,:,i1);
    for z1 = 1 : size(ttemp_b,1)
        ttemp2(ttemp_b(z1,1),ttemp_b(z1,2)) = 1;
    end
    anat_edges_WM(:,:,i1) = ttemp2;
    clear ttemp*;
end
clear F temp z1;

FC_mask_GMedge = anat_edges_GM;
FC_mask_WMedge = anat_edges_WM;
clear anat_edges_GM anat_edges_WM anat_orig anat_mask

% make sure that GM and WM masks do not overlap (exclude edge voxels)
FC_mask_GM = double((single(mask_GM_reduced.img - mask_WM_reduced.img) - FC_mask_GMedge - FC_mask_WMedge) > 0);
FC_mask_WM = double((single(mask_WM_reduced.img - mask_GM_reduced.img) - FC_mask_WMedge - FC_mask_GMedge) > 0);

if sum(FC_mask_GM(:)) < sum(FC_mask_GMedge(:)) % if the voxel size is so large that it almost eroded the mask itself, then:
    clear FC_mask_GM
    FC_mask_GM = double((single(mask_GM_reduced.img - mask_WM_reduced.img)) > 0);
end
if sum(FC_mask_WM(:)) < sum(FC_mask_WMedge(:)) % if the voxel size is so large that it almost eroded the mask itself, then:
    clear FC_mask_WM
    FC_mask_WM = double((single(mask_WM_reduced.img - mask_GM_reduced.img)) > 0);
end
clear FC_mask_GMedge FC_mask_WMedge

%% ---------------------------- IDENTIFY CORD QUADRANTS ----------------------------

try
    eval(['load ' base_dir_sub fname_anat '_all_masks.mat ' 'mask_WM position_WM;']);
catch
    eval(['load ' base_dir_sub fname_anat '_all_masks.mat ' 'mask_WM;']);
end

clear A
A = zeros(size(FC_mask_GM));
siz3 = size(A,3);

for i3 = 1 : siz3
    try waitbar((i3/siz3),wbar3,sprintf('16. Define cord quadrants: slice (%d) of (%d)',i3,siz3)); catch, end
    disp(sprintf('16. Define cord quadrants: slice (%d) of (%d)',i3,siz3))
    GM = FC_mask_GM(:,:,i3);
    WM = FC_mask_WM(:,:,i3);

    fighndl = figure('Position',[round((scrsz(3)-scrsz(4))/2) 1 scrsz(4) scrsz(4)],'Color',[0.8,0.89,0.84],'InvertHardcopy','off'); try addToolbarExplorationButtons(fighndl); catch, end % [0.85,0.90,0.85]
    figmriF(GM);
    [w1,w2] = find(WM~=0);
    try xlim([min(w2),max(w2)]), ylim([min(w1),max(w1)]); catch, end
    fighndl.Color = [0.8, 0.87, 0.84] + [0, 0.02*(1-exp(siz3/i3)/exp(siz3)), 0.1*exp(i3/siz3)/exp(1)]; % [0.85, 0.90, 0.85] + [0, 0, 0.1*i3/siz3];
    
    title({'[ Define cord quadrants ]';['(Slice ' num2str(i3) ' of ' num2str(siz3) ') - Select the point that separates the cord into four quadrants.']},'Color',[0.13,0.13,0.83],'FontSize',16);
    disp(['Slice ' num2str(i3) ' of ' num2str(siz3) ': Select the point that separates the cord into four quadrants.']);
    [x_center, y_center] = ginput(1);

    title({'[ Define cord quadrants ]';['(Slice ' num2str(i3) ' of ' num2str(siz3) ') - Select the point that isolates left side of the spinal cord.']},'Color',[0.83,0.13,0.13],'FontSize',16);
    disp(['Slice ' num2str(i3) ' of ' num2str(siz3) ': Select the point that isolates left side of the spinal cord.']);
    [x_left, y_left] = ginput(1);
    
    title({'[ Define cord quadrants ]';['(Slice ' num2str(i3) ' of ' num2str(siz3) ') - Select the point that isolates right side of the spinal cord.']},'Color',[0.13,0.70,0.13],'FontSize',16);  % 'Color',[0.83,0.13,0.83]
    disp(['Slice ' num2str(i3) ' of ' num2str(siz3) ': Select the point that isolates right side of the spinal cord.']);
    [x_right, y_right] = ginput(1);
    
    close(fighndl)
    
    x_center = round(x_center);
    y_center = round(y_center);
    y_left   = round(y_left);
    y_right  = round(y_right);
    
    % divide spinal cord into eight segments:
    %     (1) = left ventral GM
    %     (2) = right ventral GM
    %     (3) = left dorsal GM
    %     (4) = right dorsal GM
    %     (5) = left ventral WM
    %     (6) = right ventral WM
    %     (7) = left dorsal WM
    %     (8) = right dorsal WM
    
    for i1 = 1 : size(A,1)
        for i2 = 1 : size(A,2)
            if     (i1 < y_left)  && (i2 >= x_center) && GM(i1,i2), A(i1,i2,i3) = 1;
            elseif (i1 >= y_right) && (i2 >= x_center) && GM(i1,i2), A(i1,i2,i3) = 2;
            elseif (i1 < y_left)  && (i2 < x_center)  && GM(i1,i2), A(i1,i2,i3) = 3;
            elseif (i1 >= y_right) && (i2 < x_center)  && GM(i1,i2), A(i1,i2,i3) = 4;
            elseif (i1 < y_center)  && (i2 >= x_center) && WM(i1,i2), A(i1,i2,i3) = 5;
            elseif (i1 >= y_center) && (i2 >= x_center) && WM(i1,i2), A(i1,i2,i3) = 6;
            elseif (i1 < y_center)  && (i2 < x_center)  && WM(i1,i2), A(i1,i2,i3) = 7;
            elseif (i1 >= y_center) && (i2 < x_center)  && WM(i1,i2), A(i1,i2,i3) = 8;
            end % if
        end
    end
    
    fighndl2 = figure('Position',[round((scrsz(3)-scrsz(4))/2) 1 scrsz(4) scrsz(4)],'InvertHardcopy','off');
    fighndl2.Color = [0.95, 0.875, 0.8];
    tmp = A(:,:,i3); tmp(find(tmp==0))=NaN;
    tmp=rot90(tmp); [w1,w2] = find(rot90(FC_mask_WM(:,:,i3))~=0);
	imshow(tmp, [0 max(tmp(:))], 'InitialMagnification', 'fit'); colormap('jet'),colorbar;
    colormap([0 0 0; parula(256)])
    xlim([min(w2)-1,max(w2)+1]), ylim([min(w1)-1,max(w1)+1]);
    title(sprintf('GM and WM quadrants for slice %d of %d',i3,siz3),'Color','red','FontSize',18)
    saveas(fighndl2,[QCpath 'cord_quadrants_slice' sprintf('%.2d',i3) '.jpg'])
    title({sprintf('GM and WM quadrants for slice %d of %d',i3,siz3);'(click anywhere on the image to close)'},'Color','red','FontSize',18)
    waitforbuttonpress
    close(fighndl2)
    clear x_left x_right x_center y_center y_left y_right GM WM w1 w2 WhiteMask tmp
end

%% save files
FC_mask_quads.img = A; clear A;
readme_cord_quadrants = sprintf('GM/WM quadrants are provided in "FC_mask_quads". GM/WM/CSF masks provided in "mask_*_reduced". The anatomical image is provided in "anat_reduced2".\nIn FC_mask_quads.img, the value in each voxel corresponds to the following:\n(1) = left ventral GM\n(2) = right ventral GM\n(3) = left dorsal GM\n(4) = right dorsal GM\n(5) = left ventral WM\n(6) = right ventral WM\n(7) = left dorsal WM\n(8) = right dorsal WM\n(0) = neither GM now WM\n\n');
readme_FC_mask_quads = readme_cord_quadrants;
save([base_dir_sub 'cord_quadrants.mat'],'FC_mask_quads','mask_GM_reduced','mask_WM_reduced','mask_CSF_reduced','anat_reduced2','new_size','readme_cord_quadrants')

for i3=1:siz3
    for i4=1:max(FC_mask_quads.img(:))
        temp = length(find(FC_mask_quads.img(:,:,i3)==i4));
        area_cord_quadrants(i3,i4) = temp * pixdims(1) * pixdims(2); clear temp
    end
end; clear i3
try
    save([base_dir_sub 'cross_sectional_areas_Neptune.mat'],'area_cord_quadrants','readme_cord_quadrants', '-append')
catch
    readme_units = sprintf('All areas are in millimeters squared\n(Note: these area measurements have not been validated, hence, please do not use this for research or publication purposes. This is for rough quality assessment only.)\n');
    save([base_dir_sub 'area_cord_quadrants_Neptune.mat'],'area_cord_quadrants','readme_cord_quadrants','readme_units')
end

for w=1:size(area_cord_quadrants,1)
    for w2=1:size(area_cord_quadrants,2)
        dispstring1{w,w2} = sprintf('%.2f',area_cord_quadrants(w,w2));
    end
end; clear w2
fighndl = figure('Position',[1 1 1280 800],'Color',[0.85,0.90,0.85],'InvertHardcopy','off','Visible','off'); % red [0.95,0.9,0.85], green [0.85,0.90,0.85], blue [0.89,0.94,0.99]
plot(area_cord_quadrants(:,1),'--p','color','r','linewidth',2.25, 'MarkerSize',14, 'MarkerEdgeColor','k', 'MarkerFaceColor','r'),
    hold on, plot(area_cord_quadrants(:,2),'--p','color','b','linewidth',2.25, 'MarkerSize',14, 'MarkerEdgeColor','k', 'MarkerFaceColor','b'),
    plot(area_cord_quadrants(:,3),'--p','color',[0 0.37 0.22],'linewidth',2.25, 'MarkerSize',14, 'MarkerEdgeColor','k', 'MarkerFaceColor',[0 0.37 0.22]),
    plot(area_cord_quadrants(:,4),'--p','color','m','linewidth',2.25, 'MarkerSize',14, 'MarkerEdgeColor','k', 'MarkerFaceColor','m'),
    plot(area_cord_quadrants(:,5),'--p','color',[0.6 0.6 0.1],'linewidth',2.25, 'MarkerSize',14, 'MarkerEdgeColor','k', 'MarkerFaceColor','y'),
    plot(area_cord_quadrants(:,6),'--p','color','c','linewidth',2.25, 'MarkerSize',14, 'MarkerEdgeColor','k', 'MarkerFaceColor','c'),
    plot(area_cord_quadrants(:,7),'--p','color','g','linewidth',2.25, 'MarkerSize',14, 'MarkerEdgeColor','k', 'MarkerFaceColor','g'),
    plot(area_cord_quadrants(:,8),'--p','color',[0.6 0.1 0.6],'linewidth',2.25, 'MarkerSize',14, 'MarkerEdgeColor','k', 'MarkerFaceColor',[0.6 0.1 0.6]),
    grid on,grid minor, ylabel('area (in millimeters squared)'), xlabel('SLICES'), xticks([1:size(area_cord_quadrants,1)]), 
    xlim([0.5 size(area_cord_quadrants,1)+0.5]), ylim([0 max(area_cord_quadrants(:))+4])
    text([1:size(area_cord_quadrants,1)]-0.1,area_cord_quadrants(:,1)+2, dispstring1(:,1),'Color','r','FontSize',10)
    text([1:size(area_cord_quadrants,1)]-0.1,area_cord_quadrants(:,2)+1, dispstring1(:,2),'Color','b','FontSize',10)
    text([1:size(area_cord_quadrants,1)]-0.1,max([0.5.*ones(size(area_cord_quadrants,1),1), area_cord_quadrants(:,3)-2]')', dispstring1(:,3),'Color',[0 0.37 0.22],'FontSize',10)
    text([1:size(area_cord_quadrants,1)]-0.1,max([0.5.*ones(size(area_cord_quadrants,1),1), area_cord_quadrants(:,4)-1]')', dispstring1(:,4),'Color','m','FontSize',10)
    text([1:size(area_cord_quadrants,1)]-0.1,area_cord_quadrants(:,5)+2, dispstring1(:,5),'Color',[0.6 0.6 0.1],'FontSize',10)
    text([1:size(area_cord_quadrants,1)]-0.1,area_cord_quadrants(:,6)+1, dispstring1(:,6),'Color','c','FontSize',10)
    text([1:size(area_cord_quadrants,1)]-0.1,max([0.5.*ones(size(area_cord_quadrants,1),1), area_cord_quadrants(:,7)-2]')', dispstring1(:,7),'Color','g','FontSize',10)
    text([1:size(area_cord_quadrants,1)]-0.1,max([0.5.*ones(size(area_cord_quadrants,1),1), area_cord_quadrants(:,8)-1]')', dispstring1(:,8),'Color',[0.6 0.1 0.6],'FontSize',10)
    legend({'Left ventral GM';'Right ventral GM';'Left dorsal GM';'Right dorsal GM';'Left ventral WM';'Right ventral WM';'Left dorsal WM';'Right dorsal WM'},'FontSize',14,'TextColor','k','Location','best')
    title('Cross sectional areas of each GM and WM quadrant across slices (units: millimeters squared)','Color',[0.65,0.11,0.19],'FontSize',16)
saveas(fighndl,[QCpath 'cross_sectional_areas_each_quad' '.jpg'])
close(fighndl)


try
    load([base_dir_sub 'workspace_variables_temp.mat'])
catch
    % save variables for next program...
    f_BPF_max_vec = 0.10;
    f_BPF_max_str_vec = {'10'};
    cutoff = 0.50; cutoff_str = '50';
    f_BPF_max = f_BPF_max_vec(1,1);
    f_BPF_max_str = f_BPF_max_str_vec{1,1};
    scfMRItb_04_unzipFile(base_dir_sub, fname, '')
    F = load_untouch_nii([base_dir_sub fname '.nii']); TR = F.hdr.dime.pixdim(1,5); clear F
end

try
    save([base_dir_sub 'workspace_variables_run1.mat'],'base_dir_sub','fname*','cutoff*','mask_*','f_BPF_max*','*FC_mask_*','*spine','TR','scan3T','new_size')
catch
    save([base_dir_sub 'workspace_variables_run1.mat'])
end

try
if nargin<6
    close(wbar3)
end
catch
end

end
```
