## Supplementary material for "Neptune: a toolbox for spinal cord functional MRI data processing and quality assurance": Neptune software: scfMRItb_17_deconvolution_rest.m

```
function scfMRItb_17_deconvolution_rest(base_dir_sub,fname,fname_anat_orig, varargin)

fname_anat = [fname_anat_orig '_reduced'];
try
    load([base_dir_sub sprintf('workspace_variables_run%d.mat',1)], 'cutoff_str')
catch
    cutoff_str = '50';
end

fullsize = 0; % If fullsize=1 then the pre-processed data will not be cropped. By default fullsize=0, so the images are cropped (suggested because the cord is often just a small portion of the field of view [to avoid warping along the phase encoding direction], so the images can be safely cropped without losing the cord. Rest assured that there are checks to ensure that the cord is not cropped off.)

if nargin<4
    run_on_HPF_data = 1; % by default, use high-pass filtered data to perform deconvolution. A value of 0 will perform deconvolution on band-pass filtered data
else
    run_on_HPF_data = varargin{1};
end

if nargin<5
    bpf_hz = 0.1;
else
    bpf_hz = varargin{2};
end
bpf_hz = bpf_hz(1);
if bpf_hz<0.1
    bpf_hz_str = sprintf('0%.0f',bpf_hz(1)*100);
else
    bpf_hz_str = sprintf('%.0f',bpf_hz(1)*100);
end

if nargin<6
    wbar3 = waitbar(0,'17. HRF deconvolution...','Name','Progress(17): HRF deconvolution...','Units','normalized','Position',[3/5 0 1/5 1/17]);
else
    wbar3 = varargin{3};
end

if nargin<7
    save_videos = 1; % 1 = save QC videos (generates sizeable .mp4 files and takes a few minutes to do it). This is turned on by default, but if you don't want to save videos then set it to 0
else
    save_videos = varargin{4};
end
if nargin<8
    Rns = 0; % 0 = don't create runX subfolder; X = create runX subfolder
else
    Rns = varargin{5};
end

if ~(exist([base_dir_sub 'QC'],'dir'))
    mkdir([base_dir_sub 'QC'])
end
if ~(exist([base_dir_sub 'QC' '/17_deconvolution'],'dir'))
    mkdir([base_dir_sub 'QC' '/17_deconvolution'])
end
if Rns==0
    QCpath = [base_dir_sub 'QC' '/17_deconvolution/'];
else
    if ~(exist([base_dir_sub 'QC' '/17_deconvolution' '/run' num2str(Rns)],'dir'))
        mkdir([base_dir_sub 'QC' '/17_deconvolution' '/run' num2str(Rns)])
    end
    QCpath = [base_dir_sub 'QC' '/17_deconvolution' '/run' num2str(Rns) '/'];
end

if ~(exist([base_dir_sub 'QC' '/QC_shortlist'],'dir'))
    mkdir([base_dir_sub 'QC' '/QC_shortlist'])
end
if Rns==0
    QCpath2 = [base_dir_sub 'QC' '/QC_shortlist/'];
else
    if ~(exist([base_dir_sub 'QC' '/QC_shortlist' '/run' num2str(Rns)],'dir'))
        mkdir([base_dir_sub 'QC' '/QC_shortlist' '/run' num2str(Rns)])
    end
    QCpath2 = [base_dir_sub 'QC' '/QC_shortlist' '/run' num2str(Rns) '/'];
end

%% Prepare cord mask to perform deconvolution only in voxels within the mask
basestr = strfind(base_dir_sub,'/');
basestr = base_dir_sub(basestr(end-1)+1:end); % name of subject folder
try
    scfMRItb_04_unzipFile(base_dir_sub, fname_anat, '2_mask_GM')
    scfMRItb_04_unzipFile(base_dir_sub, fname_anat, '2_mask_WM')
    mask_GM_reduced  = load_untouch_nii([base_dir_sub fname_anat '2' '_mask_GM.nii']);
    mask_WM_reduced  = load_untouch_nii([base_dir_sub fname_anat '2' '_mask_WM.nii']);
catch
    try
    load([base_dir_sub sprintf('workspace_variables_run%d.mat',1)], 'mask_GM_reduced','mask_WM_reduced')
    catch
	scfMRItb_04_unzipFile(base_dir_sub, fname_anat, '_mask_GM')
	scfMRItb_04_unzipFile(base_dir_sub, fname_anat, '_mask_WM')
	scfMRItb_04_unzipFile(base_dir_sub, fname_anat, '_mask_CSF')
    mmask1 = load_untouch_nii([base_dir_sub fname_anat '_mask_GM.nii']); % gray matter
    mmask2 = load_untouch_nii([base_dir_sub fname_anat '_mask_WM.nii']); % white matter
    mmask3 = load_untouch_nii([base_dir_sub fname_anat '_mask_CSF.nii']); % white matter
    mmask_all = single(or(or(mmask1.img, mmask2.img), mmask3.img));    
    smmask = (mean(mmask_all,3) > 0);
    [s1,s2] = find(smmask~=0);
    s1median = median(s1); s2median = median(s2);
    s1min = min(s1); s2min = min(s2);
    s1max = max(s1); s2max = max(s2);
    smaxrange = max([s2max-s2min,s1max-s1min]);
    if round(size(mmask_all,1)/4) > smaxrange
        new_size = round(size(mmask_all,1)/4); % 128 - assume even for now
    elseif round(size(mmask_all,1)/3) > smaxrange
        new_size = round(size(mmask_all,1)/3); % 171
    elseif round(size(mmask_all,1)/2) > smaxrange
        new_size = round(size(mmask_all,1)/2); % 256
    else
        new_size = size(mmask_all,1); % 512
    end
    if fullsize==1
        new_size = size(mmask_all,1);
    end
    xmin = s1median - round(new_size/2) + 1; xmax = s1median + round(new_size/2);
    ymin = s2median - round(new_size/2) + 1; ymax = s2median + round(new_size/2);
    new_size = xmax - xmin + 1;
    if str2num(basestr(1:4)) <= 2017 %#ok<ST2NM> % if data were acquired prior to 2018 then don't change the bounding box of the reduced size images that were already processed before
        mask_GM_reduced.img = mmask1.img(round((size(A.img,1)-new_size)/2)+1:end-round((size(A.img,1)-new_size)/2), ...
            round((size(A.img,2)-new_size)/2)+1:end-round((size(A.img,2)-new_size)/2), :);
        mask_WM_reduced.img = mmask2.img(round((size(A.img,1)-new_size)/2)+1:end-round((size(A.img,1)-new_size)/2), ...
            round((size(A.img,2)-new_size)/2)+1:end-round((size(A.img,2)-new_size)/2), :);
    else % else if data were acquired after 2017 then use the new method to determine the bounding box by centering the new box around the centroid of the spinal cord mask
        mask_GM_reduced.img = mmask1.img(xmin:xmax, ymin:ymax, :);
        mask_WM_reduced.img = mmask2.img(xmin:xmax, ymin:ymax, :);
    end
    end
end
mask_deconv = or(mask_GM_reduced.img, mask_WM_reduced.img) > 0;

%% LOAD DATA

try waitbar(0.1,wbar3,'17. Load 4D fMRI data'); catch, end

if exist([base_dir_sub fname '_warped' '.nii'], 'file') || exist([base_dir_sub fname '_warped' '.nii.gz'], 'file') % if none of CSF, WM or COV regress steps were performed
    whichip=8;
    suffix = '_warped';
end
if exist([base_dir_sub fname '_WM' '.nii'], 'file') || exist([base_dir_sub fname '_WM' '.nii.gz'], 'file') % if only WM regress step was performed
    whichip=7;
    suffix = '_WM';
end
if exist([base_dir_sub fname '_denoised' cutoff_str '.nii'], 'file') || exist([base_dir_sub fname '_denoised' cutoff_str '.nii.gz'], 'file') % if only CSF regress step was performed
    whichip=6;
    suffix = ['_denoised' cutoff_str];
end
if exist([base_dir_sub fname '_cov' '.nii'], 'file') || exist([base_dir_sub fname '_cov' '.nii.gz'], 'file') % if only COV regress step was performed
    whichip=5;
    suffix = '_cov';
end
if exist([base_dir_sub fname '_WMcov' '.nii'], 'file') || exist([base_dir_sub fname '_WMcov' '.nii.gz'], 'file') % if both WM and COV regress steps were performed
    whichip=4;
    suffix = '_WMcov';
end
if exist([base_dir_sub fname '_denoised' cutoff_str 'WM' '.nii'], 'file') || exist([base_dir_sub fname '_denoised' cutoff_str 'WM' '.nii.gz'], 'file') % if both CSF and WM regress steps were performed
    whichip=3;
    suffix = ['_denoised' cutoff_str 'WM'];
end
if exist([base_dir_sub fname '_denoised' cutoff_str 'cov' '.nii'], 'file') || exist([base_dir_sub fname '_denoised' cutoff_str 'cov' '.nii.gz'], 'file') % if both CSF and COV regress steps were performed
    whichip=2;
    suffix = ['_denoised' cutoff_str 'cov'];
end
if exist([base_dir_sub fname '_denoised' cutoff_str 'WMcov' '.nii'], 'file') || exist([base_dir_sub fname '_denoised' cutoff_str 'WMcov' '.nii.gz'], 'file') % if CSF, WM and COV regress steps were performed
    whichip=1;
    suffix = ['_denoised' cutoff_str 'WMcov'];
end


if run_on_HPF_data==1
    if whichip==1
        if exist([base_dir_sub fname '_denoised' cutoff_str 'WMcov' '_HPF' '01' '.mat'], 'file')
            load([base_dir_sub fname '_denoised' cutoff_str 'WMcov' '_HPF' '01' '.mat'], 'F');
        else
            load([base_dir_sub fname '_denoised' cutoff_str 'WMcov' '.mat'], 'F');
        end
    elseif whichip==2
        if exist([base_dir_sub fname '_denoised' cutoff_str 'cov' '_HPF' '01' '.mat'], 'file')
            load([base_dir_sub fname '_denoised' cutoff_str 'cov' '_HPF' '01' '.mat'], 'F');
        else
            load([base_dir_sub fname '_denoised' cutoff_str 'cov' '.mat'], 'F');
        end
    elseif whichip==3
        if exist([base_dir_sub fname '_denoised' cutoff_str 'WM' '_HPF' '01' '.mat'], 'file')
            load([base_dir_sub fname '_denoised' cutoff_str 'WM' '_HPF' '01' '.mat'], 'F');
        else
            load([base_dir_sub fname '_denoised' cutoff_str 'WM' '.mat'], 'F');
        end
    elseif whichip==4
        if exist([base_dir_sub fname '_WMcov' '_HPF' '01' '.mat'], 'file')
            load([base_dir_sub fname '_WMcov' '_HPF' '01' '.mat'], 'F');
        else
            load([base_dir_sub fname '_WMcov' '.mat'], 'F');
        end
    elseif whichip==5
        if exist([base_dir_sub fname '_cov' '_HPF' '01' '.mat'], 'file')
            load([base_dir_sub fname '_cov' '_HPF' '01' '.mat'], 'F');
        else
            load([base_dir_sub fname '_cov' '.mat'], 'F');
        end
    elseif whichip==6
        if exist([base_dir_sub fname '_denoised' cutoff_str '_HPF' '01' '.mat'], 'file')
            load([base_dir_sub fname '_denoised' cutoff_str '_HPF' '01' '.mat'], 'F');
        else
            load([base_dir_sub fname '_denoised' cutoff_str '.mat'], 'F');
        end
    elseif whichip==7
        if exist([base_dir_sub fname '_WM' '_HPF' '01' '.mat'], 'file')
            load([base_dir_sub fname '_WM' '_HPF' '01' '.mat'], 'F');
        else
            load([base_dir_sub fname '_WM' '.mat'], 'F');
        end
    elseif whichip==8
        if exist([base_dir_sub fname '_warped' '_HPF' '01' '.mat'], 'file')
            load([base_dir_sub fname '_warped' '_HPF' '01' '.mat'], 'F');
        else
            load([base_dir_sub fname '_warped' '.mat'], 'F');
        end
    end
    
elseif run_on_HPF_data==0
    try
        if whichip==1
            if exist([base_dir_sub fname '_denoised' cutoff_str 'WMcov' '_BPF' bpf_hz_str '.mat'], 'file')
                load([base_dir_sub fname '_denoised' cutoff_str 'WMcov' '_BPF' bpf_hz_str '.mat'], 'F');
            else
                load([base_dir_sub fname '_denoised' cutoff_str 'WMcov' '.mat'], 'F');
            end
        elseif whichip==2
            if exist([base_dir_sub fname '_denoised' cutoff_str 'cov' '_BPF' bpf_hz_str '.mat'], 'file')
                load([base_dir_sub fname '_denoised' cutoff_str 'cov' '_BPF' bpf_hz_str '.mat'], 'F');
            else
                load([base_dir_sub fname '_denoised' cutoff_str 'cov' '.mat'], 'F');
            end
        elseif whichip==3
            if exist([base_dir_sub fname '_denoised' cutoff_str 'WM' '_BPF' bpf_hz_str '.mat'], 'file')
                load([base_dir_sub fname '_denoised' cutoff_str 'WM' '_BPF' bpf_hz_str '.mat'], 'F');
            else
                load([base_dir_sub fname '_denoised' cutoff_str 'WM' '.mat'], 'F');
            end
        elseif whichip==4
            if exist([base_dir_sub fname '_WMcov' '_BPF' bpf_hz_str '.mat'], 'file')
                load([base_dir_sub fname '_WMcov' '_BPF' bpf_hz_str '.mat'], 'F');
            else
                load([base_dir_sub fname '_WMcov' '.mat'], 'F');
            end
        elseif whichip==5
            if exist([base_dir_sub fname '_cov' '_BPF' bpf_hz_str '.mat'], 'file')
                load([base_dir_sub fname '_cov' '_BPF' bpf_hz_str '.mat'], 'F');
            else
                load([base_dir_sub fname '_cov' '.mat'], 'F');
            end
        elseif whichip==6
            if exist([base_dir_sub fname '_denoised' cutoff_str '_BPF' bpf_hz_str '.mat'], 'file')
                load([base_dir_sub fname '_denoised' cutoff_str '_BPF' bpf_hz_str '.mat'], 'F');
            else
                load([base_dir_sub fname '_denoised' cutoff_str '.mat'], 'F');
            end
        elseif whichip==7
            if exist([base_dir_sub fname '_WM' '_BPF' bpf_hz_str '.mat'], 'file')
                load([base_dir_sub fname '_WM' '_BPF' bpf_hz_str '.mat'], 'F');
            else
                load([base_dir_sub fname '_WM' '.mat'], 'F');
            end
        elseif whichip==8
            if exist([base_dir_sub fname '_warped' '_BPF' bpf_hz_str '.mat'], 'file')
                load([base_dir_sub fname '_warped' '_BPF' bpf_hz_str '.mat'], 'F');
            else
                load([base_dir_sub fname '_warped' '.mat'], 'F');
            end
        end
        n1=1;
        
    catch
        if whichip==1
            if exist([base_dir_sub fname '_denoised' cutoff_str 'WMcov' '_BPF' '10' '.mat'], 'file')
                load([base_dir_sub fname '_denoised' cutoff_str 'WMcov' '_BPF' '10' '.mat'], 'F');
            else
                load([base_dir_sub fname '_denoised' cutoff_str 'WMcov' '.mat'], 'F');
            end
        elseif whichip==2
            if exist([base_dir_sub fname '_denoised' cutoff_str 'cov' '_BPF' '10' '.mat'], 'file')
                load([base_dir_sub fname '_denoised' cutoff_str 'cov' '_BPF' '10' '.mat'], 'F');
            else
                load([base_dir_sub fname '_denoised' cutoff_str 'cov' '.mat'], 'F');
            end
        elseif whichip==3
            if exist([base_dir_sub fname '_denoised' cutoff_str 'WM' '_BPF' '10' '.mat'], 'file')
                load([base_dir_sub fname '_denoised' cutoff_str 'WM' '_BPF' '10' '.mat'], 'F');
            else
                load([base_dir_sub fname '_denoised' cutoff_str 'WM' '.mat'], 'F');
            end
        elseif whichip==4
            if exist([base_dir_sub fname '_WMcov' '_BPF' '10' '.mat'], 'file')
                load([base_dir_sub fname '_WMcov' '_BPF' '10' '.mat'], 'F');
            else
                load([base_dir_sub fname '_WMcov' '.mat'], 'F');
            end
        elseif whichip==5
            if exist([base_dir_sub fname '_cov' '_BPF' '10' '.mat'], 'file')
                load([base_dir_sub fname '_cov' '_BPF' '10' '.mat'], 'F');
            else
                load([base_dir_sub fname '_cov' '.mat'], 'F');
            end
        elseif whichip==6
            if exist([base_dir_sub fname '_denoised' cutoff_str '_BPF' '10' '.mat'], 'file')
                load([base_dir_sub fname '_denoised' cutoff_str '_BPF' '10' '.mat'], 'F');
            else
                load([base_dir_sub fname '_denoised' cutoff_str '.mat'], 'F');
            end
        elseif whichip==7
            if exist([base_dir_sub fname '_WM' '_BPF' '10' '.mat'], 'file')
                load([base_dir_sub fname '_WM' '_BPF' '10' '.mat'], 'F');
            else
                load([base_dir_sub fname '_WM' '.mat'], 'F');
            end
        elseif whichip==8
            if exist([base_dir_sub fname '_warped' '_BPF' '10' '.mat'], 'file')
                load([base_dir_sub fname '_warped' '_BPF' '10' '.mat'], 'F');
            else
                load([base_dir_sub fname '_warped' '.mat'], 'F');
            end
        end
        n1=2;
    end
end

[siz1, siz2, siz3, ~] = size(F.img);

x = F;
x.hdr.dime.dim(2) = size(x.img,1); x.hdr.dime.dim(3) = size(x.img,2);
try
    x.original;
    x.original.hdr.dime.dim(2) = size(x.img,1); x.original.hdr.dime.dim(3) = size(x.img,2);
catch
end
x.img=[];

TR = F.hdr.dime.pixdim(1,5); % volume acquisition time (repetition time)

% generating Chebyshev bandpass filter coefficients (0.01-0.1 Hz)
f_sampling = 1 / TR; % Hz

Wst = [0.01 bpf_hz] / (f_sampling/2); % bpf_hz is the upper cutoff frequency
stopband_attenuation = 60; % in dB
passband_ripple = 3; % in dB
minorder = ceil( acosh( sqrt( (10^(0.1*(stopband_attenuation+2*passband_ripple))-1)/(sqrt(10^(0.1*passband_ripple)-1)^2) )) ./ acosh( f_sampling/bpf_hz )); % Refer to this link for the formula used here to determine the model order: https://electronics.stackexchange.com/questions/519144/designing-a-chebyshev-filter-am-i-getting-the-order-wrong
cheby_order = min([minorder,round(size(F.img,4)/10)]); % The model order must be at least one-tenth of the total number of time points (this is a ballpark value, not determined analytically).
[B_BPF, A_BPF] = cheby2(cheby_order, stopband_attenuation, Wst);
clear minorder stopband_attenuation passband_ripple cheby_order Wst

% The following two lines are to generate FIR filter coefficients
L = round(30*f_sampling); filts = designfilt('bandpassfir','FilterOrder',L,'CutoffFrequency1',0.01,'CutoffFrequency2',bpf_hz,'SampleRate',1/TR);
coef = filts.Coefficients; coef=coef'; clear L filts % coefficients for bandpass filtering

%% ----- HRF DECONVOLUTION ----------------------------------
% https://pubmed.ncbi.nlm.nih.gov/23422254/
% https://pubmed.ncbi.nlm.nih.gov/29656446/

F = F.img;
if run_on_HPF_data==1, F2 = zeros(size(F,1),size(F,2),size(F,3),size(F,4)); F3=F2; end
F4 = zeros(size(F,1),size(F,2),size(F,3),size(F,4));
HRF = zeros(siz1,siz2,siz3,round(30/TR)); RH = zeros(siz1,siz2,siz3); TTP = RH; FWHM = RH;

for i3 = 1 : siz3
    try waitbar((i3/siz3),wbar3,sprintf('17. HRF deconvolution: slice (%d) of (%d) - %s',i3,siz3,fname)); catch, end
    fprintf('Step 17: HRF deconvolution - slice (%d) of (%d) - %s\n',i3,siz3,fname)
    for i1 = 1:siz1
        for i2 = 1:siz2
            if mask_deconv(i1,i2,i3)==1
                Fdata = double(squeeze(F(i1,i2,i3,:)));
                [Fdeconv,HRF(i1,i2,i3,:),~,PARA] = exec_deconv(Fdata,TR);
                Fdeconv = double(Fdeconv);
                RH(i1,i2,i3) = PARA(1);
                TTP(i1,i2,i3) = PARA(2);
                FWHM(i1,i2,i3) = PARA(3);

                if run_on_HPF_data==1
                    if ~ ( exist([base_dir_sub fname suffix '_BPF' bpf_hz_str '.nii'],'file') || exist([base_dir_sub fname suffix '_BPF' bpf_hz_str '.nii.gz'],'file') )
                        try
                        F2(i1,i2,i3,:) = filtfilt(coef, 1, Fdata); % non-deconvolved filtered data
                        catch
                        F2(i1,i2,i3,:) = filtfilt(B_BPF, A_BPF, Fdata); % non-deconvolved filtered data
                        end
                    end
                    F3(i1,i2,i3,:) = Fdeconv; % deconvolved unfiltered data
                end
                
                try
                F4(i1,i2,i3,:) = filtfilt(coef, 1, Fdeconv); % deconvolved filtered data
                catch
                F4(i1,i2,i3,:) = filtfilt(B_BPF, A_BPF, Fdeconv); % deconvolved filtered data
                end
                clear Fdata Fdeconv PARA
            end
        end
    end
end

fprintf('Step 17: HRF deconvolution: saving deconvolved data\n')

%% SAVE DATA

% ----- SAVE NON-DECONVOLVED FILTERED DATA if needed ----------------------------------

try waitbar(1,wbar3,'17. HRF deconvolution: saving deconvolved data'); catch, end

if run_on_HPF_data==1
if ~ ( exist([base_dir_sub fname suffix '_BPF' bpf_hz_str '.nii'],'file') || exist([base_dir_sub fname suffix '_BPF' bpf_hz_str '.nii.gz'],'file') )

x.img = F2;
if whichip==1
    save_untouch_nii(x, [base_dir_sub fname '_denoised' cutoff_str 'WMcov' '_BPF' bpf_hz_str '.nii']) % save non-deconvolved filtered data as NIfTI
    clear F; F=x; x.img=[];
    save([base_dir_sub fname '_denoised' cutoff_str 'WMcov' '_BPF' bpf_hz_str '.mat'], 'F'); % save non-deconvolved filtered data as .mat
elseif whichip==2
    save_untouch_nii(x, [base_dir_sub fname '_denoised' cutoff_str 'cov' '_BPF' bpf_hz_str '.nii']) % save non-deconvolved filtered data as NIfTI
    clear F; F=x; x.img=[];
    save([base_dir_sub fname '_denoised' cutoff_str 'cov' '_BPF' bpf_hz_str '.mat'], 'F'); % save non-deconvolved filtered data as .mat
elseif whichip==3
    save_untouch_nii(x, [base_dir_sub fname '_denoised' cutoff_str 'WM' '_BPF' bpf_hz_str '.nii']) % save non-deconvolved filtered data as NIfTI
    clear F; F=x; x.img=[];
    save([base_dir_sub fname '_denoised' cutoff_str 'WM' '_BPF' bpf_hz_str '.mat'], 'F'); % save non-deconvolved filtered data as .mat
elseif whichip==4
    save_untouch_nii(x, [base_dir_sub fname '_WMcov' '_BPF' bpf_hz_str '.nii']) % save non-deconvolved filtered data as NIfTI
    clear F; F=x; x.img=[];
    save([base_dir_sub fname '_WMcov' '_BPF' bpf_hz_str '.mat'], 'F'); % save non-deconvolved filtered data as .mat
elseif whichip==5
    save_untouch_nii(x, [base_dir_sub fname '_cov' '_BPF' bpf_hz_str '.nii']) % save non-deconvolved filtered data as NIfTI
    clear F; F=x; x.img=[];
    save([base_dir_sub fname '_cov' '_BPF' bpf_hz_str '.mat'], 'F'); % save non-deconvolved filtered data as .mat
elseif whichip==6
    save_untouch_nii(x, [base_dir_sub fname '_denoised' cutoff_str '_BPF' bpf_hz_str '.nii']) % save non-deconvolved filtered data as NIfTI
    clear F; F=x; x.img=[];
    save([base_dir_sub fname '_denoised' cutoff_str '_BPF' bpf_hz_str '.mat'], 'F'); % save non-deconvolved filtered data as .mat
elseif whichip==7
    save_untouch_nii(x, [base_dir_sub fname '_WM' '_BPF' bpf_hz_str '.nii']) % save non-deconvolved filtered data as NIfTI
    clear F; F=x; x.img=[];
    save([base_dir_sub fname '_WM' '_BPF' bpf_hz_str '.mat'], 'F'); % save non-deconvolved filtered data as .mat
elseif whichip==8
    save_untouch_nii(x, [base_dir_sub fname '_warped' '_BPF' bpf_hz_str '.nii']) % save non-deconvolved filtered data as NIfTI
    clear F; F=x; x.img=[];
    save([base_dir_sub fname '_warped' '_BPF' bpf_hz_str '.mat'], 'F'); % save non-deconvolved filtered data as .mat
end

end
end

% ----- SAVE DECONVOLVED UNFILTERED DATA ----------------------------------

if run_on_HPF_data==1
    
x.img = F3;
if whichip==1
    save_untouch_nii(x, [base_dir_sub fname '_denoised' cutoff_str 'WMcov' '_HPF' '01' '_deconv' '.nii']) % save deconvolved unfiltered data as NIfTI
    clear F; F=x; x.img=[];
    save([base_dir_sub fname '_denoised' cutoff_str 'WMcov' '_HPF' '01' '_deconv' '.mat'], 'F'); % save deconvolved unfiltered data as .mat
elseif whichip==2
    save_untouch_nii(x, [base_dir_sub fname '_denoised' cutoff_str 'cov' '_HPF' '01' '_deconv' '.nii']) % save deconvolved unfiltered data as NIfTI
    clear F; F=x; x.img=[];
    save([base_dir_sub fname '_denoised' cutoff_str 'cov' '_HPF' '01' '_deconv' '.mat'], 'F'); % save deconvolved unfiltered data as .mat
elseif whichip==3
    save_untouch_nii(x, [base_dir_sub fname '_denoised' cutoff_str 'WM' '_HPF' '01' '_deconv' '.nii']) % save deconvolved unfiltered data as NIfTI
    clear F; F=x; x.img=[];
    save([base_dir_sub fname '_denoised' cutoff_str 'WM' '_HPF' '01' '_deconv' '.mat'], 'F'); % save deconvolved unfiltered data as .mat
elseif whichip==4
    save_untouch_nii(x, [base_dir_sub fname '_WMcov' '_HPF' '01' '_deconv' '.nii']) % save deconvolved unfiltered data as NIfTI
    clear F; F=x; x.img=[];
    save([base_dir_sub fname '_WMcov' '_HPF' '01' '_deconv' '.mat'], 'F'); % save deconvolved unfiltered data as .mat
elseif whichip==5
    save_untouch_nii(x, [base_dir_sub fname '_cov' '_HPF' '01' '_deconv' '.nii']) % save deconvolved unfiltered data as NIfTI
    clear F; F=x; x.img=[];
    save([base_dir_sub fname '_cov' '_HPF' '01' '_deconv' '.mat'], 'F'); % save deconvolved unfiltered data as .mat
elseif whichip==6
    save_untouch_nii(x, [base_dir_sub fname '_denoised' cutoff_str '_HPF' '01' '_deconv' '.nii']) % save deconvolved unfiltered data as NIfTI
    clear F; F=x; x.img=[];
    save([base_dir_sub fname '_denoised' cutoff_str '_HPF' '01' '_deconv' '.mat'], 'F'); % save deconvolved unfiltered data as .mat
elseif whichip==7
    save_untouch_nii(x, [base_dir_sub fname '_WM' '_HPF' '01' '_deconv' '.nii']) % save deconvolved unfiltered data as NIfTI
    clear F; F=x; x.img=[];
    save([base_dir_sub fname '_WM' '_HPF' '01' '_deconv' '.mat'], 'F'); % save deconvolved unfiltered data as .mat
elseif whichip==8
    save_untouch_nii(x, [base_dir_sub fname '_warped' '_HPF' '01' '_deconv' '.nii']) % save deconvolved unfiltered data as NIfTI
    clear F; F=x; x.img=[];
    save([base_dir_sub fname '_warped' '_HPF' '01' '_deconv' '.mat'], 'F'); % save deconvolved unfiltered data as .mat
end

end

% ----- SAVE DECONVOLVED FILTERED DATA -----

x.img = F4;
if run_on_HPF_data==1
    if whichip==1
        save_untouch_nii(x, [base_dir_sub fname '_denoised' cutoff_str 'WMcov' '_deconv' '_BPF' bpf_hz_str '.nii']) % save deconvolved filtered data as NIfTI
        clear F; F=x; x.img=[];
        save([base_dir_sub fname '_denoised' cutoff_str 'WMcov' '_deconv' '_BPF' bpf_hz_str '.mat'], 'F'); % save deconvolved filtered data as .mat
    elseif whichip==2
        save_untouch_nii(x, [base_dir_sub fname '_denoised' cutoff_str 'cov' '_deconv' '_BPF' bpf_hz_str '.nii']) % save deconvolved filtered data as NIfTI
        clear F; F=x; x.img=[];
        save([base_dir_sub fname '_denoised' cutoff_str 'cov' '_deconv' '_BPF' bpf_hz_str '.mat'], 'F'); % save deconvolved filtered data as .mat
    elseif whichip==3
        save_untouch_nii(x, [base_dir_sub fname '_denoised' cutoff_str 'WM' '_deconv' '_BPF' bpf_hz_str '.nii']) % save deconvolved filtered data as NIfTI
        clear F; F=x; x.img=[];
        save([base_dir_sub fname '_denoised' cutoff_str 'WM' '_deconv' '_BPF' bpf_hz_str '.mat'], 'F'); % save deconvolved filtered data as .mat
    elseif whichip==4
        save_untouch_nii(x, [base_dir_sub fname '_WMcov' '_deconv' '_BPF' bpf_hz_str '.nii']) % save deconvolved filtered data as NIfTI
        clear F; F=x; x.img=[];
        save([base_dir_sub fname '_WMcov' '_deconv' '_BPF' bpf_hz_str '.mat'], 'F'); % save deconvolved filtered data as .mat
    elseif whichip==5
        save_untouch_nii(x, [base_dir_sub fname '_cov' '_deconv' '_BPF' bpf_hz_str '.nii']) % save deconvolved filtered data as NIfTI
        clear F; F=x; x.img=[];
        save([base_dir_sub fname '_cov' '_deconv' '_BPF' bpf_hz_str '.mat'], 'F'); % save deconvolved filtered data as .mat
    elseif whichip==6
        save_untouch_nii(x, [base_dir_sub fname '_denoised' cutoff_str '_deconv' '_BPF' bpf_hz_str '.nii']) % save deconvolved filtered data as NIfTI
        clear F; F=x; x.img=[];
        save([base_dir_sub fname '_denoised' cutoff_str '_deconv' '_BPF' bpf_hz_str '.mat'], 'F'); % save deconvolved filtered data as .mat
    elseif whichip==7
        save_untouch_nii(x, [base_dir_sub fname '_WM' '_deconv' '_BPF' bpf_hz_str '.nii']) % save deconvolved filtered data as NIfTI
        clear F; F=x; x.img=[];
        save([base_dir_sub fname '_WM' '_deconv' '_BPF' bpf_hz_str '.mat'], 'F'); % save deconvolved filtered data as .mat
    elseif whichip==8
        save_untouch_nii(x, [base_dir_sub fname '_warped' '_deconv' '_BPF' bpf_hz_str '.nii']) % save deconvolved filtered data as NIfTI
        clear F; F=x; x.img=[];
        save([base_dir_sub fname '_warped' '_deconv' '_BPF' bpf_hz_str '.mat'], 'F'); % save deconvolved filtered data as .mat
    end
    
elseif run_on_HPF_data==0
    if n1==1
        if whichip==1
            save_untouch_nii(x, [base_dir_sub fname '_denoised' cutoff_str 'WMcov' '_BPF' bpf_hz_str '_deconv' '.nii']) % save deconvolved filtered data as NIfTI
            clear F; F=x; x.img=[];
            save([base_dir_sub fname '_denoised' cutoff_str 'WMcov' '_BPF' bpf_hz_str '_deconv' '.mat'], 'F'); % save deconvolved filtered data as .mat
        elseif whichip==2
            save_untouch_nii(x, [base_dir_sub fname '_denoised' cutoff_str 'cov' '_BPF' bpf_hz_str '_deconv' '.nii']) % save deconvolved filtered data as NIfTI
            clear F; F=x; x.img=[];
            save([base_dir_sub fname '_denoised' cutoff_str 'cov' '_BPF' bpf_hz_str '_deconv' '.mat'], 'F'); % save deconvolved filtered data as .mat
        elseif whichip==3
            save_untouch_nii(x, [base_dir_sub fname '_denoised' cutoff_str 'WM' '_BPF' bpf_hz_str '_deconv' '.nii']) % save deconvolved filtered data as NIfTI
            clear F; F=x; x.img=[];
            save([base_dir_sub fname '_denoised' cutoff_str 'WM' '_BPF' bpf_hz_str '_deconv' '.mat'], 'F'); % save deconvolved filtered data as .mat
        elseif whichip==4
            save_untouch_nii(x, [base_dir_sub fname '_WMcov' '_BPF' bpf_hz_str '_deconv' '.nii']) % save deconvolved filtered data as NIfTI
            clear F; F=x; x.img=[];
            save([base_dir_sub fname '_WMcov' '_BPF' bpf_hz_str '_deconv' '.mat'], 'F'); % save deconvolved filtered data as .mat
        elseif whichip==5
            save_untouch_nii(x, [base_dir_sub fname '_cov' '_BPF' bpf_hz_str '_deconv' '.nii']) % save deconvolved filtered data as NIfTI
            clear F; F=x; x.img=[];
            save([base_dir_sub fname '_cov' '_BPF' bpf_hz_str '_deconv' '.mat'], 'F'); % save deconvolved filtered data as .mat
        elseif whichip==6
            save_untouch_nii(x, [base_dir_sub fname '_denoised' cutoff_str '_BPF' bpf_hz_str '_deconv' '.nii']) % save deconvolved filtered data as NIfTI
            clear F; F=x; x.img=[];
            save([base_dir_sub fname '_denoised' cutoff_str '_BPF' bpf_hz_str '_deconv' '.mat'], 'F'); % save deconvolved filtered data as .mat
        elseif whichip==7
            save_untouch_nii(x, [base_dir_sub fname '_WM' '_BPF' bpf_hz_str '_deconv' '.nii']) % save deconvolved filtered data as NIfTI
            clear F; F=x; x.img=[];
            save([base_dir_sub fname '_WM' '_BPF' bpf_hz_str '_deconv' '.mat'], 'F'); % save deconvolved filtered data as .mat
        elseif whichip==8
            save_untouch_nii(x, [base_dir_sub fname '_warped' '_BPF' bpf_hz_str '_deconv' '.nii']) % save deconvolved filtered data as NIfTI
            clear F; F=x; x.img=[];
            save([base_dir_sub fname '_warped' '_BPF' bpf_hz_str '_deconv' '.mat'], 'F'); % save deconvolved filtered data as .mat
        end
    
    elseif n1==2
        if whichip==1
            save_untouch_nii(x, [base_dir_sub fname '_denoised' cutoff_str 'WMcov' '_BPF' '10' '_deconv' '.nii']) % save deconvolved filtered data as NIfTI
            clear F; F=x; x.img=[];
            save([base_dir_sub fname '_denoised' cutoff_str 'WMcov' '_BPF' '10' '_deconv' '.mat'], 'F'); % save deconvolved filtered data as .mat
        elseif whichip==2
            save_untouch_nii(x, [base_dir_sub fname '_denoised' cutoff_str 'cov' '_BPF' '10' '_deconv' '.nii']) % save deconvolved filtered data as NIfTI
            clear F; F=x; x.img=[];
            save([base_dir_sub fname '_denoised' cutoff_str 'cov' '_BPF' '10' '_deconv' '.mat'], 'F'); % save deconvolved filtered data as .mat
        elseif whichip==3
            save_untouch_nii(x, [base_dir_sub fname '_denoised' cutoff_str 'WM' '_BPF' '10' '_deconv' '.nii']) % save deconvolved filtered data as NIfTI
            clear F; F=x; x.img=[];
            save([base_dir_sub fname '_denoised' cutoff_str 'WM' '_BPF' '10' '_deconv' '.mat'], 'F'); % save deconvolved filtered data as .mat
        elseif whichip==4
            save_untouch_nii(x, [base_dir_sub fname '_WMcov' '_BPF' '10' '_deconv' '.nii']) % save deconvolved filtered data as NIfTI
            clear F; F=x; x.img=[];
            save([base_dir_sub fname '_WMcov' '_BPF' '10' '_deconv' '.mat'], 'F'); % save deconvolved filtered data as .mat
        elseif whichip==5
            save_untouch_nii(x, [base_dir_sub fname '_cov' '_BPF' '10' '_deconv' '.nii']) % save deconvolved filtered data as NIfTI
            clear F; F=x; x.img=[];
            save([base_dir_sub fname '_cov' '_BPF' '10' '_deconv' '.mat'], 'F'); % save deconvolved filtered data as .mat
        elseif whichip==6
            save_untouch_nii(x, [base_dir_sub fname '_denoised' cutoff_str '_BPF' '10' '_deconv' '.nii']) % save deconvolved filtered data as NIfTI
            clear F; F=x; x.img=[];
            save([base_dir_sub fname '_denoised' cutoff_str '_BPF' '10' '_deconv' '.mat'], 'F'); % save deconvolved filtered data as .mat
        elseif whichip==7
            save_untouch_nii(x, [base_dir_sub fname '_WM' '_BPF' '10' '_deconv' '.nii']) % save deconvolved filtered data as NIfTI
            clear F; F=x; x.img=[];
            save([base_dir_sub fname '_WM' '_BPF' '10' '_deconv' '.mat'], 'F'); % save deconvolved filtered data as .mat
        elseif whichip==8
            save_untouch_nii(x, [base_dir_sub fname '_warped' '_BPF' '10' '_deconv' '.nii']) % save deconvolved filtered data as NIfTI
            clear F; F=x; x.img=[];
            save([base_dir_sub fname '_warped' '_BPF' '10' '_deconv' '.mat'], 'F'); % save deconvolved filtered data as .mat
        end
        
    end
end

% ----- SAVE HRF PARAMETERS ----------------------------------

if run_on_HPF_data==1
    if whichip==1
        save([base_dir_sub fname '_denoised' cutoff_str 'WMcov' '_HRFparams' '.mat'], 'RH','TTP','FWHM','HRF'); % save HRF parameters
    elseif whichip==2
        save([base_dir_sub fname '_denoised' cutoff_str 'cov' '_HRFparams' '.mat'], 'RH','TTP','FWHM','HRF'); % save HRF parameters
    elseif whichip==3
        save([base_dir_sub fname '_denoised' cutoff_str 'WM' '_HRFparams' '.mat'], 'RH','TTP','FWHM','HRF'); % save HRF parameters
    elseif whichip==4
        save([base_dir_sub fname '_WMcov' '_HRFparams' '.mat'], 'RH','TTP','FWHM','HRF'); % save HRF parameters
    elseif whichip==5
        save([base_dir_sub fname '_cov' '_HRFparams' '.mat'], 'RH','TTP','FWHM','HRF'); % save HRF parameters
    elseif whichip==6
        save([base_dir_sub fname '_denoised' cutoff_str '_HRFparams' '.mat'], 'RH','TTP','FWHM','HRF'); % save HRF parameters
    elseif whichip==7
        save([base_dir_sub fname '_WM' '_HRFparams' '.mat'], 'RH','TTP','FWHM','HRF'); % save HRF parameters
    elseif whichip==8
        save([base_dir_sub fname '_warped' '_HRFparams' '.mat'], 'RH','TTP','FWHM','HRF'); % save HRF parameters
    end
    
elseif run_on_HPF_data==0
    if whichip==1
        save([base_dir_sub fname '_denoised' cutoff_str 'WMcov' '_BPF' '_HRFparams' '.mat'], 'RH','TTP','FWHM','HRF'); % save HRF parameters
    elseif whichip==2
        save([base_dir_sub fname '_denoised' cutoff_str 'cov' '_BPF' '_HRFparams' '.mat'], 'RH','TTP','FWHM','HRF'); % save HRF parameters
    elseif whichip==3
        save([base_dir_sub fname '_denoised' cutoff_str 'WM' '_BPF' '_HRFparams' '.mat'], 'RH','TTP','FWHM','HRF'); % save HRF parameters
    elseif whichip==4
        save([base_dir_sub fname '_WMcov' '_BPF' '_HRFparams' '.mat'], 'RH','TTP','FWHM','HRF'); % save HRF parameters
    elseif whichip==5
        save([base_dir_sub fname '_cov' '_BPF' '_HRFparams' '.mat'], 'RH','TTP','FWHM','HRF'); % save HRF parameters
    elseif whichip==6
        save([base_dir_sub fname '_denoised' cutoff_str '_BPF' '_HRFparams' '.mat'], 'RH','TTP','FWHM','HRF'); % save HRF parameters
    elseif whichip==7
        save([base_dir_sub fname '_WM' '_BPF' '_HRFparams' '.mat'], 'RH','TTP','FWHM','HRF'); % save HRF parameters
    elseif whichip==8
        save([base_dir_sub fname '_warped' '_BPF' '_HRFparams' '.mat'], 'RH','TTP','FWHM','HRF'); % save HRF parameters
    end
end

clear F1 F2 F3 F4

%% time series plots, global signal and FC histogram
scrsz = get(0,'ScreenSize'); pause on;
cmap = colormap('bone'); cmap = (1-exp(-cmap))./(1-exp(-1)); 

scfMRItb_04_unzipFile(base_dir_sub, fname_anat, '_mask_GM')
scfMRItb_04_unzipFile(base_dir_sub, fname_anat, '_mask_WM')
mask_GM = load_untouch_nii([base_dir_sub fname_anat '_mask_GM.nii']); mask_GM = mask_GM.img;
mask_WM = load_untouch_nii([base_dir_sub fname_anat '_mask_WM.nii']); mask_WM = mask_WM.img;

scfMRItb_04_unzipFile(base_dir_sub, fname_anat, '2_mask_GM')
scfMRItb_04_unzipFile(base_dir_sub, fname_anat, '2_mask_WM')
mask_GM2 = load_untouch_nii([base_dir_sub fname_anat '2_mask_GM.nii']); mask_GM2 = mask_GM2.img;
mask_WM2 = load_untouch_nii([base_dir_sub fname_anat '2_mask_WM.nii']); mask_WM2 = mask_WM2.img;

if Rns==0, run=1; else, run=Rns; end
try
load([base_dir_sub 'cord_quadrants.mat'],'FC_mask_quads');
catch
eval(sprintf('load([base_dir_sub ''workspace_variables_run%d.mat''],''FC_mask_quads'');',run))
end
if size(FC_mask_quads)==1
    FC_mask_quads = FC_mask_quads.img;
end
    % (1) = left ventral GM; (2) = right ventral GM; (3) = left dorsal GM; (4) = right dorsal GM
    % (5) = left ventral WM; (6) = right ventral WM; (7) = left dorsal WM; (8) = right dorsal WM
FC_mask_GM = FC_mask_quads; FC_mask_GM(find(FC_mask_GM>=5))=0; FC_mask_GM(find(FC_mask_GM>=1))=1;
FC_mask_WM = FC_mask_quads; FC_mask_WM(find(FC_mask_WM<=4))=0; FC_mask_WM(find(FC_mask_WM>=5))=1;

if run_on_HPF_data==1
    if whichip==1
        load([base_dir_sub fname '_denoised' cutoff_str 'WMcov' '_deconv' '_BPF' bpf_hz_str '.mat']); F1 = F; clear F %#ok<*LOAD>
        load([base_dir_sub fname '_denoised' cutoff_str 'WMcov' '_HRFparams' '.mat'], 'RH','TTP','FWHM','HRF'); % load HRF parameters
    elseif whichip==2
        load([base_dir_sub fname '_denoised' cutoff_str 'cov' '_deconv' '_BPF' bpf_hz_str '.mat']); F1 = F; clear F
        load([base_dir_sub fname '_denoised' cutoff_str 'cov' '_HRFparams' '.mat'], 'RH','TTP','FWHM','HRF'); % load HRF parameters
    elseif whichip==3
        load([base_dir_sub fname '_denoised' cutoff_str 'WM' '_deconv' '_BPF' bpf_hz_str '.mat']); F1 = F; clear F
        load([base_dir_sub fname '_denoised' cutoff_str 'WM' '_HRFparams' '.mat'], 'RH','TTP','FWHM','HRF'); % load HRF parameters
    elseif whichip==4
        load([base_dir_sub fname '_WMcov' '_deconv' '_BPF' bpf_hz_str '.mat']); F1 = F; clear F
        load([base_dir_sub fname '_WMcov' '_HRFparams' '.mat'], 'RH','TTP','FWHM','HRF'); % load HRF parameters
    elseif whichip==5
        load([base_dir_sub fname '_cov' '_deconv' '_BPF' bpf_hz_str '.mat']); F1 = F; clear F
        load([base_dir_sub fname '_cov' '_HRFparams' '.mat'], 'RH','TTP','FWHM','HRF'); % load HRF parameters
    elseif whichip==6
        load([base_dir_sub fname '_denoised' cutoff_str '_deconv' '_BPF' bpf_hz_str '.mat']); F1 = F; clear F
        load([base_dir_sub fname '_denoised' cutoff_str '_HRFparams' '.mat'], 'RH','TTP','FWHM','HRF'); % load HRF parameters
    elseif whichip==7
        load([base_dir_sub fname '_WM' '_deconv' '_BPF' bpf_hz_str '.mat']); F1 = F; clear F
        load([base_dir_sub fname '_WM' '_HRFparams' '.mat'], 'RH','TTP','FWHM','HRF'); % load HRF parameters
    elseif whichip==8
        load([base_dir_sub fname '_warped' '_deconv' '_BPF' bpf_hz_str '.mat']); F1 = F; clear F
        load([base_dir_sub fname '_warped' '_HRFparams' '.mat'], 'RH','TTP','FWHM','HRF'); % load HRF parameters
    end
elseif run_on_HPF_data==0
    if whichip==1
        load([base_dir_sub fname '_denoised' cutoff_str 'WMcov' '_BPF' bpf_hz_str '_deconv' '.mat']); F1 = F; clear F
        load([base_dir_sub fname '_denoised' cutoff_str 'WMcov' '_BPF' '_HRFparams' '.mat'], 'RH','TTP','FWHM','HRF'); % load HRF parameters
    elseif whichip==2
        load([base_dir_sub fname '_denoised' cutoff_str 'cov' '_BPF' bpf_hz_str '_deconv' '.mat']); F1 = F; clear F
        load([base_dir_sub fname '_denoised' cutoff_str 'cov' '_BPF' '_HRFparams' '.mat'], 'RH','TTP','FWHM','HRF'); % load HRF parameters
    elseif whichip==3
        load([base_dir_sub fname '_denoised' cutoff_str 'WM' '_BPF' bpf_hz_str '_deconv' '.mat']); F1 = F; clear F
        load([base_dir_sub fname '_denoised' cutoff_str 'WM' '_BPF' '_HRFparams' '.mat'], 'RH','TTP','FWHM','HRF'); % load HRF parameters
    elseif whichip==4
        load([base_dir_sub fname '_WMcov' '_BPF' bpf_hz_str '_deconv' '.mat']); F1 = F; clear F
        load([base_dir_sub fname '_WMcov' '_BPF' '_HRFparams' '.mat'], 'RH','TTP','FWHM','HRF'); % load HRF parameters
    elseif whichip==5
        load([base_dir_sub fname '_cov' '_BPF' bpf_hz_str '_deconv' '.mat']); F1 = F; clear F
        load([base_dir_sub fname '_cov' '_BPF' '_HRFparams' '.mat'], 'RH','TTP','FWHM','HRF'); % load HRF parameters
    elseif whichip==6
        load([base_dir_sub fname '_denoised' cutoff_str '_BPF' bpf_hz_str '_deconv' '.mat']); F1 = F; clear F
        load([base_dir_sub fname '_denoised' cutoff_str '_BPF' '_HRFparams' '.mat'], 'RH','TTP','FWHM','HRF'); % load HRF parameters
    elseif whichip==7
        load([base_dir_sub fname '_WM' '_BPF' bpf_hz_str '_deconv' '.mat']); F1 = F; clear F
        load([base_dir_sub fname '_WM' '_BPF' '_HRFparams' '.mat'], 'RH','TTP','FWHM','HRF'); % load HRF parameters
    elseif whichip==8
        load([base_dir_sub fname '_warped' '_BPF' bpf_hz_str '_deconv' '.mat']); F1 = F; clear F
        load([base_dir_sub fname '_warped' '_BPF' '_HRFparams' '.mat'], 'RH','TTP','FWHM','HRF'); % load HRF parameters
    end
end

if exist([base_dir_sub fname '_denoised' cutoff_str 'WMcov' '_BPF' bpf_hz_str '.nii'],'file') || exist([base_dir_sub fname '_denoised' cutoff_str 'WMcov' '_BPF' bpf_hz_str '.nii.gz'],'file')
    load([base_dir_sub fname '_denoised' cutoff_str 'WMcov' '_BPF' bpf_hz_str '.mat']); F5all=single(F.img); clear F
    stp=8;
elseif exist([base_dir_sub fname '_denoised' cutoff_str 'cov' '_BPF' bpf_hz_str '.nii'],'file') || exist([base_dir_sub fname '_denoised' cutoff_str 'cov' '_BPF' bpf_hz_str '.nii.gz'],'file')
    load([base_dir_sub fname '_denoised' cutoff_str 'cov' '_BPF' bpf_hz_str '.mat']); F5all=single(F.img); clear F
    stp=7;
elseif exist([base_dir_sub fname '_denoised' cutoff_str 'WM' '_BPF' bpf_hz_str '.nii'],'file') || exist([base_dir_sub fname '_denoised' cutoff_str 'WM' '_BPF' bpf_hz_str '.nii.gz'],'file')
    load([base_dir_sub fname '_denoised' cutoff_str 'WM' '_BPF' bpf_hz_str '.mat']); F5all=single(F.img); clear F
    stp=6;
elseif exist([base_dir_sub fname '_WMcov' '_BPF' bpf_hz_str '.nii'],'file') || exist([base_dir_sub fname '_WMcov' '_BPF' bpf_hz_str '.nii.gz'],'file')
    load([base_dir_sub fname '_WMcov' '_BPF' bpf_hz_str '.mat']); F5all=single(F.img); clear F
    stp=5;
elseif exist([base_dir_sub fname '_cov' '_BPF' bpf_hz_str '.nii'],'file') || exist([base_dir_sub fname '_cov' '_BPF' bpf_hz_str '.nii.gz'],'file')
    load([base_dir_sub fname '_cov' '_BPF' bpf_hz_str '.mat']); F5all=single(F.img); clear F
    stp=4;
elseif exist([base_dir_sub fname '_denoised' cutoff_str '_BPF' bpf_hz_str '.nii'],'file') || exist([base_dir_sub fname '_denoised' cutoff_str '_BPF' bpf_hz_str '.nii.gz'],'file')
    load([base_dir_sub fname '_denoised' cutoff_str '_BPF' bpf_hz_str '.mat']); F5all=single(F.img); clear F
    stp=3;
elseif exist([base_dir_sub fname '_WM' '_BPF' bpf_hz_str '.nii'],'file') || exist([base_dir_sub fname '_WM' '_BPF' bpf_hz_str '.nii.gz'],'file')
    load([base_dir_sub fname '_WM' '_BPF' bpf_hz_str '.mat']); F5all=single(F.img); clear F
    stp=2;
elseif exist([base_dir_sub fname '_warped' '_BPF' bpf_hz_str '.nii'],'file') || exist([base_dir_sub fname '_warped' '_BPF' bpf_hz_str '.nii.gz'],'file')
    load([base_dir_sub fname '_warped' '_BPF' bpf_hz_str '.mat']); F5all=single(F.img); clear F
    stp=1;
end
TR = F1.hdr.dime.pixdim(5); F1 = single(squeeze(F1.img));


for w1=1:size(HRF,3)
    for w2=1:size(HRF,4)
        tmp = HRF(:,:,w1,w2);
        HRF1_GM(w1,w2) = mean(tmp(find(FC_mask_GM(:,:,w1)==1)));
        HRF1_WM(w1,w2) = mean(tmp(find(FC_mask_WM(:,:,w1)==1)));
        clear tmp
    end
end; clear w1 w2
for w2=1:size(HRF,4)
    tmp = HRF(:,:,:,w2);
    HRF2_GM(1,w2) = mean(tmp(find(FC_mask_quads==1))); HRF2_GM(2,w2) = mean(tmp(find(FC_mask_quads==2)));
    HRF2_GM(3,w2) = mean(tmp(find(FC_mask_quads==3))); HRF2_GM(4,w2) = mean(tmp(find(FC_mask_quads==4)));
    HRF2_WM(1,w2) = mean(tmp(find(FC_mask_quads==5))); HRF2_WM(2,w2) = mean(tmp(find(FC_mask_quads==6)));
    HRF2_WM(3,w2) = mean(tmp(find(FC_mask_quads==7))); HRF2_WM(4,w2) = mean(tmp(find(FC_mask_quads==8)));
    clear tmp
end; clear w2
for w=1:size(HRF1_GM,1)
    legstr{w} = sprintf('slice %d',w);
    xlabl{w}  = sprintf('%.2f s',(w-1)*TR);
end; clear w
    % (1) = left ventral GM; (2) = right ventral GM; (3) = left dorsal GM; (4) = right dorsal GM
    % (5) = left ventral WM; (6) = right ventral WM; (7) = left dorsal WM; (8) = right dorsal WM
fighndl = figure('Position',[1 1 1280 800],'Color',[0.85,0.90,0.85],'InvertHardcopy','off','Visible','off'); % red [0.95,0.9,0.85], green [0.85,0.90,0.85], blue [0.89,0.94,0.99]
subplot2n(4,1,1), plot(HRF1_GM','linewidth',1), xlabel('time (seconds)'), ylabel('HRF'), xticks([1:length(HRF1_GM')]),xticklabels(xlabl), legend(legstr,'NumColumns',3)
    grid on,grid minor, ylim([1.1.*min([HRF1_GM(:);HRF1_WM(:);HRF2_GM(:);HRF2_WM(:)]), 1.1.*max([HRF1_GM(:);HRF1_WM(:);HRF2_GM(:);HRF2_WM(:)])]), title(['Hemodynamic response function (HRF) in the GRAY MATTER across SLICES'],'Color',[0,0.55,0.69],'FontSize',16),
subplot2n(4,1,2), plot(HRF1_WM','linewidth',1), xlabel('time (seconds)'), ylabel('HRF'), xticks([1:length(HRF1_WM')]),xticklabels(xlabl), legend(legstr,'NumColumns',3)
    grid on,grid minor, ylim([1.1.*min([HRF1_GM(:);HRF1_WM(:);HRF2_GM(:);HRF2_WM(:)]), 1.1.*max([HRF1_GM(:);HRF1_WM(:);HRF2_GM(:);HRF2_WM(:)])]), title(['HRF in the WHITE MATTER across SLICES'],'Color',[0.65,0.11,0.19],'FontSize',16),
subplot2n(4,1,3), plot(HRF2_GM','linewidth',1), xlabel('time (seconds)'), ylabel('HRF'), xticks([1:length(HRF2_GM')]),xticklabels(xlabl), legend({'left ventral GM','right ventral GM','left dorsal GM','right dorsal GM'},'FontSize',12,'NumColumns',2)
    grid on,grid minor, ylim([1.1.*min([HRF1_GM(:);HRF1_WM(:);HRF2_GM(:);HRF2_WM(:)]), 1.1.*max([HRF1_GM(:);HRF1_WM(:);HRF2_GM(:);HRF2_WM(:)])]), title(['HRF in the GRAY MATTER across QUADRANTS'],'Color',[0,0.55,0.69],'FontSize',16),
subplot2n(4,1,4), plot(HRF2_WM','linewidth',1), xlabel('time (seconds)'), ylabel('HRF'), xticks([1:length(HRF2_WM')]),xticklabels(xlabl), legend({'left ventral WM','right ventral WM','left dorsal WM','right dorsal WM'},'FontSize',12,'NumColumns',2)
    grid on,grid minor, ylim([1.1.*min([HRF1_GM(:);HRF1_WM(:);HRF2_GM(:);HRF2_WM(:)]), 1.1.*max([HRF1_GM(:);HRF1_WM(:);HRF2_GM(:);HRF2_WM(:)])]), title(['HRF in the WHITE MATTER across QUADRANTS'],'Color',[0.65,0.11,0.19],'FontSize',16),
saveas(fighndl,[QCpath 'HRF_plots_17deconv_allSlices' '.jpg'])
saveas(fighndl,[QCpath2 'HRF_plots_17deconv_allSlices' '.jpg'])
clear HRF1_GM HRF1_WM HRF2_GM HRF2_WM
close(fighndl)

for w1=1:size(RH,3)
    tmp1 = RH(:,:,w1); tmp2 = TTP(:,:,w1); tmp3 = FWHM(:,:,w1);
    RH1_GM(w1,1)   = mean(tmp1(find(FC_mask_GM(:,:,w1)==1))); RH1_WM(w1,1)   = mean(tmp1(find(FC_mask_WM(:,:,w1)==1)));
    TTP1_GM(w1,1)  = TR.*mean(tmp2(find(FC_mask_GM(:,:,w1)==1))); TTP1_WM(w1,1)  = TR.*mean(tmp2(find(FC_mask_WM(:,:,w1)==1)));
    FWHM1_GM(w1,1) = TR.*mean(tmp3(find(FC_mask_GM(:,:,w1)==1))); FWHM1_WM(w1,1) = TR.*mean(tmp3(find(FC_mask_WM(:,:,w1)==1)));
    clear tmp1 tmp2 tmp3
end; clear w1
RH2_GM(1,1) = mean(RH(find(FC_mask_quads==1))); RH2_GM(2,1) = mean(RH(find(FC_mask_quads==2))); RH2_GM(3,1) = mean(RH(find(FC_mask_quads==3))); RH2_GM(4,1) = mean(RH(find(FC_mask_quads==4)));
RH2_WM(1,1) = mean(RH(find(FC_mask_quads==5))); RH2_WM(2,1) = mean(RH(find(FC_mask_quads==6))); RH2_WM(3,1) = mean(RH(find(FC_mask_quads==7))); RH2_WM(4,1) = mean(RH(find(FC_mask_quads==8)));
TTP2_GM(1,1) = TR.*mean(TTP(find(FC_mask_quads==1))); TTP2_GM(2,1) = TR.*mean(TTP(find(FC_mask_quads==2))); TTP2_GM(3,1) = TR.*mean(TTP(find(FC_mask_quads==3))); TTP2_GM(4,1) = TR.*mean(TTP(find(FC_mask_quads==4)));
TTP2_WM(1,1) = TR.*mean(TTP(find(FC_mask_quads==5))); TTP2_WM(2,1) = TR.*mean(TTP(find(FC_mask_quads==6))); TTP2_WM(3,1) = TR.*mean(TTP(find(FC_mask_quads==7))); TTP2_WM(4,1) = TR.*mean(TTP(find(FC_mask_quads==8)));
FWHM2_GM(1,1) = TR.*mean(FWHM(find(FC_mask_quads==1))); FWHM2_GM(2,1) = TR.*mean(FWHM(find(FC_mask_quads==2))); FWHM2_GM(3,1) = TR.*mean(FWHM(find(FC_mask_quads==3))); FWHM2_GM(4,1) = TR.*mean(FWHM(find(FC_mask_quads==4)));
FWHM2_WM(1,1) = TR.*mean(FWHM(find(FC_mask_quads==5))); FWHM2_WM(2,1) = TR.*mean(FWHM(find(FC_mask_quads==6))); FWHM2_WM(3,1) = TR.*mean(FWHM(find(FC_mask_quads==7))); FWHM2_WM(4,1) = TR.*mean(FWHM(find(FC_mask_quads==8)));
    % (1) = left ventral GM; (2) = right ventral GM; (3) = left dorsal GM; (4) = right dorsal GM
    % (5) = left ventral WM; (6) = right ventral WM; (7) = left dorsal WM; (8) = right dorsal WM
fighndl = figure('Position',[1 1 1280 800],'Color',[0.85,0.90,0.85],'InvertHardcopy','off','Visible','off'); % red [0.95,0.9,0.85], green [0.85,0.90,0.85], blue [0.89,0.94,0.99]
subplot2n(4,2,[1,2]), bar([zeros(size(RH1_GM)),RH1_GM,RH1_WM,zeros(size(RH1_WM))]'), ylabel('HRF response height (a.u.)'), xticklabels({'','GRAY MATTER response height','WHITE MATTER response height',''}),
    a = get(gca,'xticklabels'); set(gca,'xticklabels',a,'fontsize',12); clear a
    grid on,grid minor, ylim([min([RH1_GM(:);RH1_WM(:)])-(max([RH1_GM(:);RH1_WM(:)])-min([RH1_GM(:);RH1_WM(:)]))/2, max([RH1_GM(:);RH1_WM(:)])+(max([RH1_GM(:);RH1_WM(:)])-min([RH1_GM(:);RH1_WM(:)]))/2]),
    title(['Hemodynamic response function (HRF) RESPONSE HEIGHT across SLICES'],'Color',[0,0.55,0.69],'FontSize',16), legend(legstr,'Location','northeast','NumColumns',2,'FontSize',10)
    text(1.8,max([RH1_GM(:);RH1_WM(:)])+(max([RH1_GM(:);RH1_WM(:)])-min([RH1_GM(:);RH1_WM(:)]))/4, sprintf('mean = %.2f',mean(RH1_GM(:))),'Color',[0,0.55,0.69],'FontSize',15)
    text(2.8,max([RH1_GM(:);RH1_WM(:)])+(max([RH1_GM(:);RH1_WM(:)])-min([RH1_GM(:);RH1_WM(:)]))/4, sprintf('mean = %.2f',mean(RH1_WM(:))),'Color',[0.65,0.11,0.19],'FontSize',15)
subtightplot(4,2,3,0.05,[0.08,0.0525],[0.1,0.065]), bar([TTP1_GM,TTP1_WM]'), ylabel('TTP (in seconds)'), xticklabels({'GRAY MATTER time-to-peak','WHITE MATTER time-to-peak'}),
    a = get(gca,'xticklabels'); set(gca,'xticklabels',a,'fontsize',12); clear a
    grid on,grid minor, ylim([min([TTP1_GM(:);TTP1_WM(:)])-1, max([TTP1_GM(:);TTP1_WM(:)])+1]), title(['HRF time-to-peak (TTP) across SLICES'],'Color',[0,0.55,0.69],'FontSize',16),
    text(0.8,max([TTP1_GM(:);TTP1_WM(:)])+0.5, sprintf('mean = %.2f s',mean(TTP1_GM(:))),'Color',[0,0.55,0.69],'FontSize',15)
    text(1.8,max([TTP1_GM(:);TTP1_WM(:)])+0.5, sprintf('mean = %.2f s',mean(TTP1_WM(:))),'Color',[0.65,0.11,0.19],'FontSize',15)
subtightplot(4,2,4,0.05,[0.08,0.0525],[0.1,0.065]), bar([FWHM1_GM,FWHM1_WM]'), ylabel('FWHM (in seconds)'), xticklabels({'GRAY MATTER FWHM','WHITE MATTER FWHM'}),
    a = get(gca,'xticklabels'); set(gca,'xticklabels',a,'fontsize',12); clear a
    grid on,grid minor, ylim([min([FWHM1_GM(:);FWHM1_WM(:)])-0.5, max([FWHM1_GM(:);FWHM1_WM(:)])+0.5]), title(['HRF full-width at half max (FWHM) across SLICES'],'Color',[0,0.55,0.69],'FontSize',16), 
    text(0.8,max([FWHM1_GM(:);FWHM1_WM(:)])+0.25, sprintf('mean = %.2f s',mean(FWHM1_GM(:))),'Color',[0,0.55,0.69],'FontSize',15)
    text(1.8,max([FWHM1_GM(:);FWHM1_WM(:)])+0.25, sprintf('mean = %.2f s',mean(FWHM1_WM(:))),'Color',[0.65,0.11,0.19],'FontSize',15)
subplot2n(4,2,[5,6]), bar([zeros(size(RH2_GM)),RH2_GM,RH2_WM,zeros(size(RH2_WM))]'), ylabel('HRF response height (a.u.)'), xticklabels({'','GRAY MATTER response height','WHITE MATTER response height',''}),
    a = get(gca,'xticklabels'); set(gca,'xticklabels',a,'fontsize',12); clear a
    grid on,grid minor, ylim([min([RH2_GM(:);RH2_WM(:)])-(max([RH2_GM(:);RH2_WM(:)])-min([RH2_GM(:);RH2_WM(:)]))/2, max([RH2_GM(:);RH2_WM(:)])+(max([RH2_GM(:);RH2_WM(:)])-min([RH2_GM(:);RH2_WM(:)]))/2]),
    title(['HRF RESPONSE HEIGHT across QUADRANTS'],'Color',[0.65,0.11,0.19],'FontSize',16), legend({'left ventral','right ventral','left dorsal','right dorsal'},'Location','northeast','NumColumns',2,'FontSize',12)
    text(1.8,max([RH2_GM(:);RH2_WM(:)])+(max([RH2_GM(:);RH2_WM(:)])-min([RH2_GM(:);RH2_WM(:)]))/4, sprintf('mean = %.2f',mean(RH2_GM(:))),'Color',[0,0.55,0.69],'FontSize',15)
    text(2.8,max([RH2_GM(:);RH2_WM(:)])+(max([RH2_GM(:);RH2_WM(:)])-min([RH2_GM(:);RH2_WM(:)]))/4, sprintf('mean = %.2f',mean(RH2_WM(:))),'Color',[0.65,0.11,0.19],'FontSize',15)
subtightplot(4,2,7,0.05,[0.105,0.055],[0.1,0.065]), bar([TTP2_GM,TTP2_WM]'), ylabel('TTP (in seconds)'), xticklabels({'GRAY MATTER time-to-peak','WHITE MATTER time-to-peak'}),
    a = get(gca,'xticklabels'); set(gca,'xticklabels',a,'fontsize',12); clear a
    grid on,grid minor, ylim([min([TTP2_GM(:);TTP2_WM(:)])-1, max([TTP2_GM(:);TTP2_WM(:)])+1]), title(['HRF time-to-peak (TTP) across QUADRANTS'],'Color',[0.65,0.11,0.19],'FontSize',16),
    text(0.8,max([TTP2_GM(:);TTP2_WM(:)])+0.5, sprintf('mean = %.2f s',mean(TTP2_GM(:))),'Color',[0,0.55,0.69],'FontSize',15)
    text(1.8,max([TTP2_GM(:);TTP2_WM(:)])+0.5, sprintf('mean = %.2f s',mean(TTP2_WM(:))),'Color',[0.65,0.11,0.19],'FontSize',15)
subtightplot(4,2,8,0.05,[0.105,0.055],[0.1,0.065]), bar([FWHM2_GM,FWHM2_WM]'), ylabel('FWHM (in seconds)'), xticklabels({'GRAY MATTER FWHM','WHITE MATTER FWHM'}),
    a = get(gca,'xticklabels'); set(gca,'xticklabels',a,'fontsize',12); clear a
    grid on,grid minor, ylim([min([FWHM2_GM(:);FWHM2_WM(:)])-0.5, max([FWHM2_GM(:);FWHM2_WM(:)])+0.5]), title(['HRF full-width at half max (FWHM) across QUADRANTS'],'Color',[0.65,0.11,0.19],'FontSize',16), 
    text(0.8,max([FWHM2_GM(:);FWHM2_WM(:)])+0.25, sprintf('mean = %.2f s',mean(FWHM2_GM(:))),'Color',[0,0.55,0.69],'FontSize',15)
    text(1.8,max([FWHM2_GM(:);FWHM2_WM(:)])+0.25, sprintf('mean = %.2f s',mean(FWHM2_WM(:))),'Color',[0.65,0.11,0.19],'FontSize',15)
saveas(fighndl,[QCpath 'HRF_parameters_17deconv_allSlices' '.jpg'])
saveas(fighndl,[QCpath2 'HRF_parameters_17deconv_allSlices' '.jpg'])
clear HRF1_GM HRF1_WM HRF2_GM HRF2_WM RH1_GM RH1_WM RH2_GM RH2_WM TTP1_GM TTP1_WM TTP2_WM TTP2_GM FWHM2_WM FWHM2_GM FWHM1_WM FWHM1_GM
close(fighndl)


Smask_err=0;
try
    scfMRItb_04_resplitData(base_dir_sub, fname, '_Smask', siz3) % if slice-wise data doesn't exist but combined 4D data exists, then split the data to generate the needed slice-wise data
catch
    load([base_dir_sub fname '_mask_NS' '.mat'])
    Smask1_ = ~mask_NS.img; clear mask_NS
    Smask_err=1;
end

F2g_=[]; mF2g_=[]; F2w_=[]; mF2w_=[]; Cg_=[]; Cw_=[]; F4_=[]; mF4_=[]; C2_=[];
if stp~=0, F6g_=[]; mF6g_=[]; F6w_=[]; mF6w_=[]; C3g_=[]; C3w_=[]; end
for i3 = 1 : siz3
    try waitbar((i3/siz3),wbar3,sprintf('17. QC plots: slice (%d) of (%d)',i3,siz3)); catch, end
    fprintf('17. QC plots: slice (%d) of (%d)\n',i3,siz3)
    
    if Smask_err==0
        scfMRItb_04_resplitData(base_dir_sub, fname, '_Smask', siz3) % if slice-wise data doesn't exist but combined 4D data exists, then split the data to generate the needed slice-wise data
        scfMRItb_04_unzipFile(base_dir_sub, fname, ['_slice' num2str(i3) '_Smask'])
        Smask1 = load_untouch_nii([base_dir_sub fname '_slice' num2str(i3) '_Smask.nii']); Smask1 = Smask1.img;
    else
        Smask1 = Smask1_(:,:,i3);
    end

    for i5=1:size(F1,4)
        tmp = F1(:,:,i3,i5);
        F2g(:,i5) = tmp(find(mask_GM2(:,:,i3)==1));
        F2w(:,i5) = tmp(find(mask_WM2(:,:,i3)==1));
        clear tmp
    end; clear i5
    [~,svar1] = sort(var(F2g,0,2)); [~,sdist1] = sort(svar1); sortorder1{i3,1} = sdist1;
    [~,svar2] = sort(var(F2w,0,2)); [~,sdist2] = sort(svar2); sortorder2{i3,1} = sdist2;
    
    fighndl = figure('Position',[1 1 1152 720],'Color',[0.95,0.9,0.85],'InvertHardcopy','off','Visible','off'); % red [0.95,0.9,0.85], green [0.85,0.90,0.85], blue [0.89,0.94,0.99]
    subplot2n(2,1,1), imagesc(F2g(sortorder1{i3},:)),colormap(cmap),colorbar, xlabel('time (in samples)'),ylabel('Gray matter voxels (sorted by variance)'),
        title(['Time series within the GRAY MATTER after "17-HRF-deconvolution" in slice ' num2str(i3) ' of ' num2str(siz3)],'Color',[0,0.55,0.69],'FontSize',14);
    subplot2n(2,1,2), imagesc(F2w(sortorder2{i3},:)),colormap(cmap),colorbar, xlabel('time (in samples)'),ylabel('White matter voxels (sorted by variance)'),
        title(['Time series within the WHITE MATTER after "17-HRF-deconvolution" in slice ' num2str(i3) ' of ' num2str(siz3)],'Color',[0.65,0.11,0.19],'FontSize',14);
    saveas(fighndl,[QCpath 'fMRI_ts_17deconv_slice' sprintf('%.2d',i3) '.jpg'])
    close(fighndl)
    
    scfMRItb_04_resplitData(base_dir_sub, fname, '', siz3) % if slice-wise data doesn't exist but combined 4D data exists, then split the data to generate the needed slice-wise data
	scfMRItb_04_unzipFile(base_dir_sub, fname, ['_slice' num2str(i3)])
    F3 = load_untouch_nii([base_dir_sub fname '_slice' num2str(i3) '.nii']); F3 = single(squeeze(F3.img));
    for i5=1:size(F3,3)
        tmp = F3(:,:,i5);
        F4(:,i5) = tmp(find(Smask1==1)); clear tmp
    end; clear i5 F3
    mF2g=mean(F2g,1); mF2w=mean(F2w,1);
    mF2g=zscore(mF2g); mF2w=zscore(mF2w);

    Cg = corrcoef(F2g'); Cg = uppertriangle(Cg);
    Cw = corrcoef(F2w'); Cw = uppertriangle(Cw);
    C2 = corrcoef(F4');  C2 = uppertriangle(C2);
    fighndl = figure('Position',[1 1 1152 720],'Color',[0.95,0.9,0.85],'InvertHardcopy','off','Visible','off'); % red [0.95,0.9,0.85], green [0.85,0.90,0.85], blue [0.89,0.94,0.99]
    subplot2n(3,1,1), histogram(C2,'EdgeColor','none','FaceColor','k','FaceAlpha',0.5,'Normalization','probability'); grid on,grid minor, xlim([-1 1]), hold on, xline(0,'Color','k','linewidth',1.5); hold off
      title(sprintf('Histogram of voxel-to-voxel correlations in UNPROCESSED RAW DATA (slice %d of %d)\n%.1f%% of the correlations were > 0  [ mean=%.2f, median=%.2f, SD=%.2f ]',i3,siz3,100*length(find(C2>0))/length(C2),mean(C2),median(C2),std(C2)),'Color','k','FontSize',14)
    subplot2n(3,1,2), histogram(Cg,'EdgeColor','none','FaceColor',[0,0.55,0.69],'FaceAlpha',0.5,'Normalization','probability'); grid on,grid minor, xlim([-1 1]), hold on, xline(0,'Color','k','linewidth',1.5); hold off
      title(sprintf('Histogram of voxel-to-voxel GRAY MATTER correlations AFTER 17-HRF-deconvolution (slice %d of %d)\n%.1f%% of the correlations were > 0  [ mean=%.2f, median=%.2f, SD=%.2f ]',i3,siz3,100*length(find(Cg>0))/length(Cg),mean(Cg),median(Cg),std(Cg)),'Color',[0,0.55,0.69],'FontSize',14)
    subplot2n(3,1,3), histogram(Cw,'EdgeColor','none','FaceColor',[0.65,0.11,0.19],'FaceAlpha',0.5,'Normalization','probability'); grid on,grid minor, xlim([-1 1]), hold on, xline(0,'Color','k','linewidth',1.5); hold off
      title(sprintf('Histogram of voxel-to-voxel WHITE MATTER correlations AFTER 17-HRF-deconvolution (slice %d of %d)\n%.1f%% of the correlations were > 0  [ mean=%.2f, median=%.2f, SD=%.2f ]',i3,siz3,100*length(find(Cw>0))/length(Cw),mean(Cw),median(Cw),std(Cw)),'Color',[0.65,0.11,0.19],'FontSize',14)
    saveas(fighndl,[QCpath 'hist_FC_compare_17deconv_slice' sprintf('%.2d',i3) '.jpg'])
    close(fighndl)
    
    if stp~=0
        msk=1;
        if exist('F5all','var')
            F5 = single(squeeze(F5all(:,:,i3,:)));
            msk=2;
        else
            if stp==1
                scfMRItb_04_resplitData(base_dir_sub, fname, '_warped', siz3) % if slice-wise data doesn't exist but combined 4D data exists, then split the data to generate the needed slice-wise data
                scfMRItb_04_unzipFile(base_dir_sub, fname, ['_slice' num2str(i3) '_warped'])
                F5 = load_untouch_nii([base_dir_sub fname '_slice' num2str(i3) '_warped.nii']); F5 = single(squeeze(F5.img));
            elseif stp==2
                scfMRItb_04_resplitData(base_dir_sub, fname, '_WM', siz3) % if slice-wise data doesn't exist but combined 4D data exists, then split the data to generate the needed slice-wise data
                scfMRItb_04_unzipFile(base_dir_sub, fname, ['_slice' num2str(i3) '_WM'])
                F5 = load_untouch_nii([base_dir_sub fname '_slice' num2str(i3) '_WM.nii']); F5 = single(squeeze(F5.img));
            elseif stp==3
                scfMRItb_04_resplitData(base_dir_sub, fname, ['_denoised' cutoff_str], siz3) % if slice-wise data doesn't exist but combined 4D data exists, then split the data to generate the needed slice-wise data
                scfMRItb_04_unzipFile(base_dir_sub, fname, ['_slice' num2str(i3) '_denoised' cutoff_str ''])
                F5 = load_untouch_nii([base_dir_sub fname '_slice' num2str(i3) '_denoised' cutoff_str '.nii']); F5 = single(squeeze(F5.img));
            elseif stp==4
                scfMRItb_04_resplitData(base_dir_sub, fname, '_cov', siz3) % if slice-wise data doesn't exist but combined 4D data exists, then split the data to generate the needed slice-wise data
                scfMRItb_04_unzipFile(base_dir_sub, fname, ['_slice' num2str(i3) '_cov'])
                F5 = load_untouch_nii([base_dir_sub fname '_slice' num2str(i3) '_cov.nii']); F5 = single(squeeze(F5.img));
            elseif stp==5
                scfMRItb_04_resplitData(base_dir_sub, fname, '_WMcov', siz3) % if slice-wise data doesn't exist but combined 4D data exists, then split the data to generate the needed slice-wise data
                scfMRItb_04_unzipFile(base_dir_sub, fname, ['_slice' num2str(i3) '_WMcov'])
                F5 = load_untouch_nii([base_dir_sub fname '_slice' num2str(i3) '_WMcov.nii']); F5 = single(squeeze(F5.img));
            elseif stp==6
                scfMRItb_04_resplitData(base_dir_sub, fname, ['_denoised' cutoff_str 'WM'], siz3) % if slice-wise data doesn't exist but combined 4D data exists, then split the data to generate the needed slice-wise data
                scfMRItb_04_unzipFile(base_dir_sub, fname, ['_slice' num2str(i3) '_denoised' cutoff_str 'WM'])
                F5 = load_untouch_nii([base_dir_sub fname '_slice' num2str(i3) '_denoised' cutoff_str 'WM.nii']); F5 = single(squeeze(F5.img));
            elseif stp==7
                scfMRItb_04_resplitData(base_dir_sub, fname, ['_denoised' cutoff_str 'cov'], siz3) % if slice-wise data doesn't exist but combined 4D data exists, then split the data to generate the needed slice-wise data
                scfMRItb_04_unzipFile(base_dir_sub, fname, ['_slice' num2str(i3) '_denoised' cutoff_str 'cov'])
                F5 = load_untouch_nii([base_dir_sub fname '_slice' num2str(i3) '_denoised' cutoff_str 'cov.nii']); F5 = single(squeeze(F5.img));
            elseif stp==8
                scfMRItb_04_resplitData(base_dir_sub, fname, ['_denoised' cutoff_str 'WMcov'], siz3) % if slice-wise data doesn't exist but combined 4D data exists, then split the data to generate the needed slice-wise data
                scfMRItb_04_unzipFile(base_dir_sub, fname, ['_slice' num2str(i3) '_denoised' cutoff_str 'WMcov'])
                F5 = load_untouch_nii([base_dir_sub fname '_slice' num2str(i3) '_denoised' cutoff_str 'WMcov.nii']); F5 = single(squeeze(F5.img));
            end
        end
        
        for i5=1:size(F5,3)
            tmp = F5(:,:,i5);
            if msk==1
                F6g(:,i5) = tmp(find(mask_GM(:,:,i3)==1));
                F6w(:,i5) = tmp(find(mask_WM(:,:,i3)==1));
            else
                F6g(:,i5) = tmp(find(mask_GM2(:,:,i3)==1));
                F6w(:,i5) = tmp(find(mask_WM2(:,:,i3)==1));
            end
            clear tmp
        end; clear i5 F5
        mF6g=mean(F6g,1); mF6w=mean(F6w,1); 
        mF6g=zscore(mF6g); mF6w=zscore(mF6w);
        fighndl = figure('Position',[1 1 1152 720],'Color',[0.95,0.9,0.85],'InvertHardcopy','off','Visible','off'); % red [0.95,0.9,0.85], green [0.85,0.90,0.85], blue [0.89,0.94,0.99]
        if (stp==1)&&(msk==1)
            subplot2n(4,1,1), plot(mF6g,'color','k','linewidth',1.5), hold on, plot(mF2g,'color',[0,0.55,0.69],'linewidth',0.2), ylabel('mean time series'),
                grid on,grid minor, xlim([1 length(mF2g)]),ylim([min(1.25.*[mF2g,mF6g]) 1.25.*max([mF2g,mF6g])]), title(['Unfiltered: normalized mean GRAY MATTER signal AFTER "11-func-anat-registration" (slice ' num2str(i3) ' of ' num2str(siz3) ')'],'Color','k','FontSize',14), legend({'Gray matter signal before deconvolution'},'FontSize',12,'TextColor','k','Location','best')
            subplot2n(4,1,3), plot(mF6w,'color','k','linewidth',1.5), hold on, plot(mF2w,'color',[0.65,0.11,0.19],'linewidth',0.2), ylabel('mean time series'),
                grid on,grid minor, xlim([1 length(mF2g)]),ylim([min(1.25.*[mF2w,mF6w]) 1.25.*max([mF2w,mF6w])]), title(['Unfiltered: normalized mean WHITE MATTER signal AFTER "11-func-anat-registration" (slice ' num2str(i3) ' of ' num2str(siz3) ')'],'Color','k','FontSize',14), legend({'White matter signal before deconvolution'},'FontSize',12,'TextColor','k','Location','best')
        elseif (stp==3)&&(msk==1)
            subplot2n(4,1,1), plot(mF6g,'color','k','linewidth',1.5), hold on, plot(mF2g,'color',[0,0.55,0.69],'linewidth',0.2), ylabel('mean time series'),
                grid on,grid minor, xlim([1 length(mF2g)]),ylim([min(1.25.*[mF2g,mF6g]) 1.25.*max([mF2g,mF6g])]), title(['Unfiltered: normalized mean GRAY MATTER signal AFTER "12-denoise3-CSFregress" (slice ' num2str(i3) ' of ' num2str(siz3) ')'],'Color','k','FontSize',14), legend({'Gray matter signal before deconvolution'},'FontSize',12,'TextColor','k','Location','best')
            subplot2n(4,1,3), plot(mF6w,'color','k','linewidth',1.5), hold on, plot(mF2w,'color',[0.65,0.11,0.19],'linewidth',0.2), ylabel('mean time series'),
                grid on,grid minor, xlim([1 length(mF2g)]),ylim([min(1.25.*[mF2w,mF6w]) 1.25.*max([mF2w,mF6w])]), title(['Unfiltered: normalized mean WHITE MATTER signal AFTER "12-denoise3-CSFregress" (slice ' num2str(i3) ' of ' num2str(siz3) ')'],'Color','k','FontSize',14), legend({'White matter signal before deconvolution'},'FontSize',12,'TextColor','k','Location','best')
        elseif ((stp==2)||(stp==6))&&(msk==1)
            subplot2n(4,1,1), plot(mF6g,'color','k','linewidth',1.5), hold on, plot(mF2g,'color',[0,0.55,0.69],'linewidth',0.2), ylabel('mean time series'),
                grid on,grid minor, xlim([1 length(mF2g)]),ylim([min(1.25.*[mF2g,mF6g]) 1.25.*max([mF2g,mF6g])]), title(['Unfiltered: normalized mean GRAY MATTER signal AFTER "13-denoise4-WMregress" (slice ' num2str(i3) ' of ' num2str(siz3) ')'],'Color','k','FontSize',14), legend({'Gray matter signal before deconvolution'},'FontSize',12,'TextColor','k','Location','best')
            subplot2n(4,1,3), plot(mF6w,'color','k','linewidth',1.5), hold on, plot(mF2w,'color',[0.65,0.11,0.19],'linewidth',0.2), ylabel('mean time series'),
                grid on,grid minor, xlim([1 length(mF2g)]),ylim([min(1.25.*[mF2w,mF6w]) 1.25.*max([mF2w,mF6w])]), title(['Unfiltered: normalized mean WHITE MATTER signal AFTER "13-denoise4-WMregress" (slice ' num2str(i3) ' of ' num2str(siz3) ')'],'Color','k','FontSize',14), legend({'White matter signal before deconvolution'},'FontSize',12,'TextColor','k','Location','best')
        elseif ((stp==4)||(stp==5)||(stp==7)||(stp==8))&&(msk==1)
            subplot2n(4,1,1), plot(mF6g,'color','k','linewidth',1.5), hold on, plot(mF2g,'color',[0,0.55,0.69],'linewidth',0.2), ylabel('mean time series'),
                grid on,grid minor, xlim([1 length(mF2g)]),ylim([min(1.25.*[mF2g,mF6g]) 1.25.*max([mF2g,mF6g])]), title(['Bandpass filtered: normalized mean GRAY MATTER signal AFTER "14-denoise5-COVregress" (slice ' num2str(i3) ' of ' num2str(siz3) ')'],'Color','k','FontSize',14), legend({'Gray matter signal before deconvolution'},'FontSize',12,'TextColor','k','Location','best')
            subplot2n(4,1,3), plot(mF6w,'color','k','linewidth',1.5), hold on, plot(mF2w,'color',[0.65,0.11,0.19],'linewidth',0.2), ylabel('mean time series'),
                grid on,grid minor, xlim([1 length(mF2g)]),ylim([min(1.25.*[mF2w,mF6w]) 1.25.*max([mF2w,mF6w])]), title(['Bandpass filtered: normalized mean WHITE MATTER signal AFTER "14-denoise5-COVregress" (slice ' num2str(i3) ' of ' num2str(siz3) ')'],'Color','k','FontSize',14), legend({'White matter signal before deconvolution'},'FontSize',12,'TextColor','k','Location','best')
        elseif (msk==2)
            subplot2n(4,1,1), plot(mF6g,'color','k','linewidth',1.5), hold on, plot(mF2g,'color',[0,0.55,0.69],'linewidth',0.2), ylabel('mean time series'),
                grid on,grid minor, xlim([1 length(mF2g)]),ylim([min(1.25.*[mF2g,mF6g]) 1.25.*max([mF2g,mF6g])]), title(['Bandpass filtered: normalized mean GRAY MATTER signal AFTER "15-bandpass-filter" (slice ' num2str(i3) ' of ' num2str(siz3) ')'],'Color','k','FontSize',14), legend({'Gray matter signal before deconvolution'},'FontSize',12,'TextColor','k','Location','best')
            subplot2n(4,1,3), plot(mF6w,'color','k','linewidth',1.5), hold on, plot(mF2w,'color',[0.65,0.11,0.19],'linewidth',0.2), ylabel('mean time series'),
                grid on,grid minor, xlim([1 length(mF2g)]),ylim([min(1.25.*[mF2w,mF6w]) 1.25.*max([mF2w,mF6w])]), title(['Bandpass filtered: normalized mean WHITE MATTER signal AFTER "15-bandpass-filter" (slice ' num2str(i3) ' of ' num2str(siz3) ')'],'Color','k','FontSize',14), legend({'White matter signal before deconvolution'},'FontSize',12,'TextColor','k','Location','best')
        end
        subplot2n(4,1,2), plot(mF2g,'color',[0,0.55,0.69],'linewidth',1.5), hold on, plot(mF6g,'color','k','linewidth',0.2), ylabel('mean time series'),
            grid on,grid minor, xlim([1 length(mF2g)]),ylim([min(1.25.*[mF2g,mF6g]) 1.25.*max([mF2g,mF6g])]), title(['Deconvolved bandpass filtered: normalized mean GRAY MATTER signal AFTER "17-HRF-deconvolution" [0.01-' num2str(bpf_hz_str) 'Hz] (slice ' num2str(i3) ' of ' num2str(siz3) ')'],'Color',[0,0.55,0.69],'FontSize',14), legend({'Gray matter signal after deconvolution'},'FontSize',12,'TextColor',[0,0.55,0.69],'Location','best')
        subplot2n(4,1,4), plot(mF2w,'color',[0.65,0.11,0.19],'linewidth',1.5), hold on, plot(mF6w,'color','k','linewidth',0.2), xlabel('time (in samples)'), ylabel('mean time series'),
            grid on,grid minor, xlim([1 length(mF2g)]),ylim([min(1.25.*[mF2w,mF6w]) 1.25.*max([mF2w,mF6w])]), title(['Deconvolved bandpass filtered: normalized mean WHITE MATTER signal AFTER "17-HRF-deconvolution" [0.01-' num2str(bpf_hz_str) 'Hz] (slice ' num2str(i3) ' of ' num2str(siz3) ')'],'Color',[0.65,0.11,0.19],'FontSize',14), legend({'White matter signal after deconvolution'},'FontSize',12,'TextColor',[0.65,0.11,0.19],'Location','best')
        saveas(fighndl,[QCpath 'mean_timeseries_15filter_vs_17deconv_slice' sprintf('%.2d',i3) '.jpg'])
        close(fighndl)
        
        C3g = corrcoef(F6g'); C3g = uppertriangle(C3g); C3w = corrcoef(F6w'); C3w = uppertriangle(C3w);
        fighndl = figure('Position',[1 1 1152 720],'Color',[0.95,0.9,0.85],'InvertHardcopy','off','Visible','off'); % red [0.95,0.9,0.85], green [0.85,0.90,0.85], blue [0.89,0.94,0.99]
        if (stp==1)&&(msk==1)
            subplot2n(4,1,1), histogram(C3g,'EdgeColor','none','FaceColor','k','FaceAlpha',0.5,'Normalization','probability'); grid on,grid minor, xlim([-1 1]), hold on, xline(0,'Color','k','linewidth',1.5); hold off
                title(sprintf('Histogram of voxel-to-voxel GRAY MATTER correlations AFTER 11-func-anat-registration (slice %d of %d)\n%.1f%% of the correlations were > 0  [ mean=%.2f, median=%.2f, SD=%.2f ]',i3,siz3,100*length(find(C3g>0))/length(C3g),mean(C3g),median(C3g),std(C3g)),'Color','k','FontSize',14)
            subplot2n(4,1,3), histogram(C3w,'EdgeColor','none','FaceColor','k','FaceAlpha',0.5,'Normalization','probability'); grid on,grid minor, xlim([-1 1]), hold on, xline(0,'Color','k','linewidth',1.5); hold off
                title(sprintf('WHITE MATTER: AFTER 11-func-anat-registration (slice %d of %d): %.1f%% of the correlations were > 0  [ mean=%.2f, median=%.2f, SD=%.2f ]',i3,siz3,100*length(find(C3w>0))/length(C3w),mean(C3w),median(C3w),std(C3w)),'Color','k','FontSize',14)
        elseif (stp==3)&&(msk==1)
            subplot2n(4,1,1), histogram(C3g,'EdgeColor','none','FaceColor','k','FaceAlpha',0.5,'Normalization','probability'); grid on,grid minor, xlim([-1 1]), hold on, xline(0,'Color','k','linewidth',1.5); hold off
                title(sprintf('Histogram of voxel-to-voxel GRAY MATTER correlations AFTER 12-denoise3-CSFregress (slice %d of %d)\n%.1f%% of the correlations were > 0  [ mean=%.2f, median=%.2f, SD=%.2f ]',i3,siz3,100*length(find(C3g>0))/length(C3g),mean(C3g),median(C3g),std(C3g)),'Color','k','FontSize',14)
            subplot2n(4,1,3), histogram(C3w,'EdgeColor','none','FaceColor','k','FaceAlpha',0.5,'Normalization','probability'); grid on,grid minor, xlim([-1 1]), hold on, xline(0,'Color','k','linewidth',1.5); hold off
                title(sprintf('WHITE MATTER: AFTER 12-denoise3-CSFregress (slice %d of %d): %.1f%% of the correlations were > 0  [ mean=%.2f, median=%.2f, SD=%.2f ]',i3,siz3,100*length(find(C3w>0))/length(C3w),mean(C3w),median(C3w),std(C3w)),'Color','k','FontSize',14)
        elseif ((stp==2)||(stp==6))&&(msk==1)
            subplot2n(4,1,1), histogram(C3g,'EdgeColor','none','FaceColor','k','FaceAlpha',0.5,'Normalization','probability'); grid on,grid minor, xlim([-1 1]), hold on, xline(0,'Color','k','linewidth',1.5); hold off
                title(sprintf('Histogram of voxel-to-voxel GRAY MATTER correlations AFTER 13-denoise4-WMregress (slice %d of %d)\n%.1f%% of the correlations were > 0  [ mean=%.2f, median=%.2f, SD=%.2f ]',i3,siz3,100*length(find(C3g>0))/length(C3g),mean(C3g),median(C3g),std(C3g)),'Color','k','FontSize',14)
            subplot2n(4,1,3), histogram(C3w,'EdgeColor','none','FaceColor','k','FaceAlpha',0.5,'Normalization','probability'); grid on,grid minor, xlim([-1 1]), hold on, xline(0,'Color','k','linewidth',1.5); hold off
                title(sprintf('WHITE MATTER: AFTER 13-denoise4-WMregress (slice %d of %d): %.1f%% of the correlations were > 0  [ mean=%.2f, median=%.2f, SD=%.2f ]',i3,siz3,100*length(find(C3w>0))/length(C3w),mean(C3w),median(C3w),std(C3w)),'Color','k','FontSize',14)
        elseif ((stp==4)||(stp==5)||(stp==7)||(stp==8))&&(msk==1)
            subplot2n(4,1,1), histogram(C3g,'EdgeColor','none','FaceColor','k','FaceAlpha',0.5,'Normalization','probability'); grid on,grid minor, xlim([-1 1]), hold on, xline(0,'Color','k','linewidth',1.5); hold off
                title(sprintf('Histogram of voxel-to-voxel GRAY MATTER correlations AFTER 14-denoise5-COVregress (slice %d of %d)\n%.1f%% of the correlations were > 0  [ mean=%.2f, median=%.2f, SD=%.2f ]',i3,siz3,100*length(find(C3g>0))/length(C3g),mean(C3g),median(C3g),std(C3g)),'Color','k','FontSize',14)
            subplot2n(4,1,3), histogram(C3w,'EdgeColor','none','FaceColor','k','FaceAlpha',0.5,'Normalization','probability'); grid on,grid minor, xlim([-1 1]), hold on, xline(0,'Color','k','linewidth',1.5); hold off
                title(sprintf('WHITE MATTER: AFTER 14-denoise5-COVregress (slice %d of %d): %.1f%% of the correlations were > 0  [ mean=%.2f, median=%.2f, SD=%.2f ]',i3,siz3,100*length(find(C3w>0))/length(C3w),mean(C3w),median(C3w),std(C3w)),'Color','k','FontSize',14)
        elseif (msk==2)
            subplot2n(4,1,1), histogram(C3g,'EdgeColor','none','FaceColor','k','FaceAlpha',0.5,'Normalization','probability'); grid on,grid minor, xlim([-1 1]), hold on, xline(0,'Color','k','linewidth',1.5); hold off
                title(sprintf('Histogram of voxel-to-voxel GRAY MATTER correlations AFTER 15-bandpass-filter (slice %d of %d)\n%.1f%% of the correlations were > 0  [ mean=%.2f, median=%.2f, SD=%.2f ]',i3,siz3,100*length(find(C3g>0))/length(C3g),mean(C3g),median(C3g),std(C3g)),'Color','k','FontSize',14)
            subplot2n(4,1,3), histogram(C3w,'EdgeColor','none','FaceColor','k','FaceAlpha',0.5,'Normalization','probability'); grid on,grid minor, xlim([-1 1]), hold on, xline(0,'Color','k','linewidth',1.5); hold off
                title(sprintf('WHITE MATTER: AFTER 15-bandpass-filter (slice %d of %d): %.1f%% of the correlations were > 0  [ mean=%.2f, median=%.2f, SD=%.2f ]',i3,siz3,100*length(find(C3w>0))/length(C3w),mean(C3w),median(C3w),std(C3w)),'Color','k','FontSize',14)
        end
        subplot2n(4,1,2), histogram(Cg,'EdgeColor','none','FaceColor',[0,0.55,0.69],'FaceAlpha',0.5,'Normalization','probability'); grid on,grid minor, xlim([-1 1]), hold on, xline(0,'Color','k','linewidth',1.5); hold off
            title(sprintf('GRAY MATTER: AFTER 17-HRF-deconvolution (slice %d of %d): %.1f%% of the correlations were > 0  [ mean=%.2f, median=%.2f, SD=%.2f ]',i3,siz3,100*length(find(Cg>0))/length(Cg),mean(Cg),median(Cg),std(Cg)),'Color',[0,0.55,0.69],'FontSize',14)
        subplot2n(4,1,4), histogram(Cw,'EdgeColor','none','FaceColor',[0.65,0.11,0.19],'FaceAlpha',0.5,'Normalization','probability'); grid on,grid minor, xlim([-1 1]), hold on, xline(0,'Color','k','linewidth',1.5); hold off
            title(sprintf('WHITE MATTER: AFTER 17-HRF-deconvolution (slice %d of %d): %.1f%% of the correlations were > 0  [ mean=%.2f, median=%.2f, SD=%.2f ]',i3,siz3,100*length(find(Cw>0))/length(Cw),mean(Cw),median(Cw),std(Cw)),'Color',[0.65,0.11,0.19],'FontSize',14)
        saveas(fighndl,[QCpath 'hist_FC_compare_15filter_17deconv_slice' sprintf('%.2d',i3) '.jpg'])
        close(fighndl)
        F6g_=cat(1,F6g_,F6g); mF6g_=cat(1,mF6g_,mF6g); C3g_=cat(1,C3g_,C3g);
        F6w_=cat(1,F6w_,F6w); mF6w_=cat(1,mF6w_,mF6w); C3w_=cat(1,C3w_,C3w);
        clear F6g F6w mF6g mF6w C3g C3w
    end
    
    F2g_=cat(1,F2g_,F2g); mF2g_=cat(1,mF2g_,mF2g); F2w_=cat(1,F2w_,F2w); mF2w_=cat(1,mF2w_,mF2w); Cg_=cat(1,Cg_,Cg); Cw_=cat(1,Cw_,Cw); F4_=cat(1,F4_,F4); C2_=cat(1,C2_,C2);
    clear F2g F2w F4 TR Smask1 svar1 svar2 mF2g mF2w mF4 Cg Cw C2
end


[~,svar1] = sort(var(F2g_,0,2)); [~,spsc1] = sort(meanPSC(F2g_));
[~,svar2] = sort(var(F2w_,0,2)); [~,spsc2] = sort(meanPSC(F2w_));
sortorder1b = sortorder1; sortorder1=[]; f1=0; %#ok<*ASGLU>
sortorder2b = sortorder2; sortorder2=[]; f2=0; %#ok<*ASGLU>
for im=1:length(sortorder1b)
    temp = sortorder1b{im}; temp = temp + f1; f1 = f1 + length(temp);
    sortorder1 = cat(1,sortorder1,temp); clear temp
    temp = sortorder2b{im}; temp = temp + f2; f2 = f2 + length(temp);
    sortorder2 = cat(1,sortorder2,temp); clear temp
end; clear im f1 f2
sortorder1 = spsc1; % options: spsc for percentage signal change, svar for variance, (comment out the line if you want to use distance from the center of the cord separately for each slice)
sortorder2 = spsc2; % options: spsc for percentage signal change, svar for variance, (comment out the line if you want to use distance from the center of the cord separately for each slice)
for im=max(sortorder1):-1:1, if isempty(find(sortorder1==im)), sortorder1(find(sortorder1>im)) = sortorder1(find(sortorder1>im))-1; end, end; clear im %#ok<*EFIND>
for im=max(sortorder2):-1:1, if isempty(find(sortorder2==im)), sortorder2(find(sortorder2>im)) = sortorder2(find(sortorder2>im))-1; end, end; clear im %#ok<*EFIND>

fighndl = figure('Position',[1 1 1152 720],'Color',[0.89,0.94,0.99],'InvertHardcopy','off','Visible','off'); % red [0.95,0.9,0.85], green [0.85,0.90,0.85], blue [0.89,0.94,0.99]
subplot2n(2,1,1), imagesc(F2g_(sortorder1,:)),colormap(cmap),colorbar, xlabel('time (in samples)'),ylabel(sprintf('voxels (sorted by mean percentage signal change;\nbottom row is highest)')),
  title(['Time series within the GRAY MATTER AFTER 17-HRF-deconvolution (all slices combined)'],'Color',[0,0.55,0.69],'FontSize',14);
subplot2n(2,1,2), imagesc(F2w_(sortorder2,:)),colormap(cmap),colorbar, xlabel('time (in samples)'),ylabel(sprintf('voxels (sorted by mean percentage signal change;\nbottom row is highest)')),
  title(['Time series within the WHITE MATTER AFTER 17-HRF-deconvolution (all slices combined)'],'Color',[0.65,0.11,0.19],'FontSize',14);
saveas(fighndl,[QCpath 'fMRI_ts_17deconv_allSlices' '.jpg'])
saveas(fighndl,[QCpath2 'fMRI_ts_17deconv_allSlices' '.jpg'])
 close(fighndl)
mF2g_=mean(mF2g_,1); mF2w_=mean(mF2w_,1); mF4_=mean(mF4_,1);
fighndl = figure('Position',[1 1 1152 720],'Color',[0.89,0.94,0.99],'InvertHardcopy','off','Visible','off'); % red [0.95,0.9,0.85], green [0.85,0.90,0.85], blue [0.89,0.94,0.99]
subplot2n(3,1,1), histogram(C2_,'EdgeColor','none','FaceColor','k','FaceAlpha',0.5,'Normalization','probability'); grid on,grid minor, xlim([-1 1]), hold on, xline(0,'Color','k','linewidth',1.5); hold off
    title(sprintf('Histogram of voxel-to-voxel correlations in UNPROCESSED RAW DATA (all slices combined)\n%.1f%% of the correlations were > 0  [ mean=%.2f, median=%.2f, SD=%.2f ]',100*length(find(C2_>0))/length(C2_),mean(C2_),median(C2_),std(C2_)),'Color','k','FontSize',14)
subplot2n(3,1,2), histogram(Cg_,'EdgeColor','none','FaceColor',[0,0.55,0.69],'FaceAlpha',0.5,'Normalization','probability'); grid on,grid minor, xlim([-1 1]), hold on, xline(0,'Color','k','linewidth',1.5); hold off
    title(sprintf('Histogram of voxel-to-voxel GRAY MATTER correlations AFTER 17-HRF-deconvolution (all slices combined)\n%.1f%% of the correlations were > 0  [ mean=%.2f, median=%.2f, SD=%.2f ]',100*length(find(Cg_>0))/length(Cg_),mean(Cg_),median(Cg_),std(Cg_)),'Color',[0,0.55,0.69],'FontSize',14)
subplot2n(3,1,3), histogram(Cw_,'EdgeColor','none','FaceColor',[0.65,0.11,0.19],'FaceAlpha',0.5,'Normalization','probability'); grid on,grid minor, xlim([-1 1]), hold on, xline(0,'Color','k','linewidth',1.5); hold off
    title(sprintf('Histogram of voxel-to-voxel WHITE MATTER correlations AFTER 17-HRF-deconvolution (all slices combined)\n%.1f%% of the correlations were > 0  [ mean=%.2f, median=%.2f, SD=%.2f ]',100*length(find(Cw_>0))/length(Cw_),mean(Cw_),median(Cw_),std(Cw_)),'Color',[0.65,0.11,0.19],'FontSize',14)
saveas(fighndl,[QCpath 'hist_FC_compare_17deconv_allSlices' '.jpg'])
 close(fighndl)

if stp~=0
    mF6g_=mean(mF6g_,1); mF6w_=mean(mF6w_,1);
    fighndl = figure('Position',[1 1 1152 720],'Color',[0.89,0.94,0.99],'InvertHardcopy','off','Visible','off'); % red [0.95,0.9,0.85], green [0.85,0.90,0.85], blue [0.89,0.94,0.99]
    if (stp==1)&&(msk==1)
    	subplot2n(4,1,1), plot(mF6g_,'color','k','linewidth',1.5), hold on, plot(mF2g_,'color',[0,0.55,0.69],'linewidth',0.2), ylabel('mean time series'),
            grid on,grid minor, xlim([1 length(mF2g_)]),ylim([1.25.*min([mF2g_,mF6g_]) 1.25.*max([mF2g_,mF6g_])]), title(['Normalized mean GRAY MATTER signal AFTER "11-func-anat-registration" (all slices averaged)'],'Color','k','FontSize',14), legend({'Gray matter signal before deconvolution'},'FontSize',12,'TextColor','k','Location','best')
        subplot2n(4,1,3), plot(mF6w_,'color','k','linewidth',1.5), hold on, plot(mF2w_,'color',[0.65,0.11,0.19],'linewidth',0.2), ylabel('mean time series'),
            grid on,grid minor, xlim([1 length(mF2g_)]),ylim([1.25.*min([mF2w_,mF6w_]) 1.25.*max([mF2w_,mF6w_])]), title(['Normalized mean WHITE MATTER signal AFTER "11-func-anat-registration" (all slices averaged)'],'Color','k','FontSize',14), legend({'White matter signal before deconvolution'},'FontSize',12,'TextColor','k','Location','best')
    elseif (stp==3)&&(msk==1)
    	subplot2n(4,1,1), plot(mF6g_,'color','k','linewidth',1.5), hold on, plot(mF2g_,'color',[0,0.55,0.69],'linewidth',0.2), ylabel('mean time series'),
            grid on,grid minor, xlim([1 length(mF2g_)]),ylim([1.25.*min([mF2g_,mF6g_]) 1.25.*max([mF2g_,mF6g_])]), title(['Normalized mean GRAY MATTER signal AFTER "12-DENOISE3-CSFregress" (all slices averaged)'],'Color','k','FontSize',14), legend({'Gray matter signal before deconvolution'},'FontSize',12,'TextColor','k','Location','best')
        subplot2n(4,1,3), plot(mF6w_,'color','k','linewidth',1.5), hold on, plot(mF2w_,'color',[0.65,0.11,0.19],'linewidth',0.2), ylabel('mean time series'),
            grid on,grid minor, xlim([1 length(mF2g_)]),ylim([1.25.*min([mF2w_,mF6w_]) 1.25.*max([mF2w_,mF6w_])]), title(['Normalized mean WHITE MATTER signal AFTER "12-DENOISE3-CSFregress" (all slices averaged)'],'Color','k','FontSize',14), legend({'White matter signal before deconvolution'},'FontSize',12,'TextColor','k','Location','best')
    elseif ((stp==2)||(stp==6))&&(msk==1)
    	subplot2n(4,1,1), plot(mF6g_,'color','k','linewidth',1.5), hold on, plot(mF2g_,'color',[0,0.55,0.69],'linewidth',0.2), ylabel('mean time series'),
            grid on,grid minor, xlim([1 length(mF2g_)]),ylim([1.25.*min([mF2g_,mF6g_]) 1.25.*max([mF2g_,mF6g_])]), title(['Normalized mean GRAY MATTER signal AFTER "13-DENOISE4-WMregress" (all slices averaged)'],'Color','k','FontSize',14), legend({'Gray matter signal before deconvolution'},'FontSize',12,'TextColor','k','Location','best')
        subplot2n(4,1,3), plot(mF6w_,'color','k','linewidth',1.5), hold on, plot(mF2w_,'color',[0.65,0.11,0.19],'linewidth',0.2), ylabel('mean time series'),
            grid on,grid minor, xlim([1 length(mF2g_)]),ylim([1.25.*min([mF2w_,mF6w_]) 1.25.*max([mF2w_,mF6w_])]), title(['Normalized mean WHITE MATTER signal AFTER "13-DENOISE4-WMregress" (all slices averaged)'],'Color','k','FontSize',14), legend({'White matter signal before deconvolution'},'FontSize',12,'TextColor','k','Location','best')
    elseif ((stp==4)||(stp==5)||(stp==7)||(stp==8))&&(msk==1)
    	subplot2n(4,1,1), plot(mF6g_,'color','k','linewidth',1.5), hold on, plot(mF2g_,'color',[0,0.55,0.69],'linewidth',0.2), ylabel('mean time series'),
            grid on,grid minor, xlim([1 length(mF2g_)]),ylim([1.25.*min([mF2g_,mF6g_]) 1.25.*max([mF2g_,mF6g_])]), title(['Normalized mean GRAY MATTER signal AFTER "14-DENOISE5-COVregress" (all slices averaged)'],'Color','k','FontSize',14), legend({'Gray matter signal before deconvolution'},'FontSize',12,'TextColor','k','Location','best')
        subplot2n(4,1,3), plot(mF6w_,'color','k','linewidth',1.5), hold on, plot(mF2w_,'color',[0.65,0.11,0.19],'linewidth',0.2), ylabel('mean time series'),
            grid on,grid minor, xlim([1 length(mF2g_)]),ylim([1.25.*min([mF2w_,mF6w_]) 1.25.*max([mF2w_,mF6w_])]), title(['Normalized mean WHITE MATTER signal AFTER "14-DENOISE5-COVregress" (all slices averaged)'],'Color','k','FontSize',14), legend({'White matter signal before deconvolution'},'FontSize',12,'TextColor','k','Location','best')
    elseif (msk==2)
    	subplot2n(4,1,1), plot(mF6g_,'color','k','linewidth',1.5), hold on, plot(mF2g_,'color',[0,0.55,0.69],'linewidth',0.2), ylabel('mean time series'),
            grid on,grid minor, xlim([1 length(mF2g_)]),ylim([1.25.*min([mF2g_,mF6g_]) 1.25.*max([mF2g_,mF6g_])]), title(['Normalized mean GRAY MATTER signal AFTER "15-bandpass-filter" (all slices averaged)'],'Color','k','FontSize',14), legend({'Gray matter signal before deconvolution'},'FontSize',12,'TextColor','k','Location','best')
        subplot2n(4,1,3), plot(mF6w_,'color','k','linewidth',1.5), hold on, plot(mF2w_,'color',[0.65,0.11,0.19],'linewidth',0.2), ylabel('mean time series'),
            grid on,grid minor, xlim([1 length(mF2g_)]),ylim([1.25.*min([mF2w_,mF6w_]) 1.25.*max([mF2w_,mF6w_])]), title(['Normalized mean WHITE MATTER signal AFTER "15-bandpass-filter" (all slices averaged)'],'Color','k','FontSize',14), legend({'White matter signal before deconvolution'},'FontSize',12,'TextColor','k','Location','best')
    end
    subplot2n(4,1,2), plot(mF2g_,'color',[0,0.55,0.69],'linewidth',1.5), hold on, plot(mF6g_,'color','k','linewidth',0.2), ylabel('mean time series'),
        grid on,grid minor, xlim([1 length(mF2g_)]),ylim([1.25.*min([mF2g_,mF6g_]) 1.25.*max([mF2g_,mF6g_])]), title(['Normalized mean GRAY MATTER signal AFTER "17-HRF-deconvolution" (all slices averaged)'],'Color',[0,0.55,0.69],'FontSize',14), legend({'Gray matter signal after deconvolution'},'FontSize',12,'TextColor',[0,0.55,0.69],'Location','best')
    subplot2n(4,1,4), plot(mF2w_,'color',[0.65,0.11,0.19],'linewidth',1.5), hold on, plot(mF6w_,'color','k','linewidth',0.2), ylabel('mean time series'),
        grid on,grid minor, xlim([1 length(mF2g_)]),ylim([1.25.*min([mF2w_,mF6w_]) 1.25.*max([mF2w_,mF6w_])]), title(['Normalized mean WHITE MATTER signal AFTER "17-HRF-deconvolution" (all slices averaged)'],'Color',[0.65,0.11,0.19],'FontSize',14), legend({'White matter signal after deconvolution'},'FontSize',12,'TextColor',[0.65,0.11,0.19],'Location','best')
    saveas(fighndl,[QCpath 'mean_timeseries_15filter_vs_17deconv_allSlices' '.jpg'])
    saveas(fighndl,[QCpath2 'global_signal_15filter_vs_17deconv_allSlices' '.jpg'])
     close(fighndl)
    
    fighndl = figure('Position',[1 1 1152 720],'Color',[0.89,0.94,0.99],'InvertHardcopy','off','Visible','off'); % red [0.95,0.9,0.85], green [0.85,0.90,0.85], blue [0.89,0.94,0.99]
    if (stp==1)&&(msk==1)
    	subplot2n(4,1,1), histogram(C3g_,'EdgeColor','none','FaceColor','k','FaceAlpha',0.5,'Normalization','probability'); grid on,grid minor, xlim([-1 1]), hold on, xline(0,'Color','k','linewidth',1.5); hold off
            title(sprintf('Histogram of voxel-to-voxel GRAY MATTER correlations AFTER 11-registration (all slices combined)\n%.1f%% of the correlations were > 0  [ mean=%.2f, median=%.2f, SD=%.2f ]',100*length(find(C3g_>0))/length(C3g_),mean(C3g_),median(C3g_),std(C3g_)),'Color','k','FontSize',14)
        subplot2n(4,1,3), histogram(C3w_,'EdgeColor','none','FaceColor','k','FaceAlpha',0.5,'Normalization','probability'); grid on,grid minor, xlim([-1 1]), hold on, xline(0,'Color','k','linewidth',1.5); hold off
            title(sprintf('WHITE MATTER: AFTER 11-registration (all slices): %.1f%% of the correlations were > 0  [ mean=%.2f, median=%.2f, SD=%.2f ]',100*length(find(C3w_>0))/length(C3w_),mean(C3w_),median(C3w_),std(C3w_)),'Color','k','FontSize',14)
    elseif (stp==3)&&(msk==1)
    	subplot2n(4,1,1), histogram(C3g_,'EdgeColor','none','FaceColor','k','FaceAlpha',0.5,'Normalization','probability'); grid on,grid minor, xlim([-1 1]), hold on, xline(0,'Color','k','linewidth',1.5); hold off
            title(sprintf('Histogram of voxel-to-voxel GRAY MATTER correlations AFTER 12-denoise3-CSFregress (all slices combined)\n%.1f%% of the correlations were > 0  [ mean=%.2f, median=%.2f, SD=%.2f ]',100*length(find(C3g_>0))/length(C3g_),mean(C3g_),median(C3g_),std(C3g_)),'Color','k','FontSize',14)
        subplot2n(4,1,3), histogram(C3w_,'EdgeColor','none','FaceColor','k','FaceAlpha',0.5,'Normalization','probability'); grid on,grid minor, xlim([-1 1]), hold on, xline(0,'Color','k','linewidth',1.5); hold off
            title(sprintf('WHITE MATTER: AFTER 12-denoise3-CSFregress (all slices): %.1f%% of the correlations were > 0  [ mean=%.2f, median=%.2f, SD=%.2f ]',100*length(find(C3w_>0))/length(C3w_),mean(C3w_),median(C3w_),std(C3w_)),'Color','k','FontSize',14)
    elseif ((stp==2)||(stp==6))&&(msk==1)
    	subplot2n(4,1,1), histogram(C3g_,'EdgeColor','none','FaceColor','k','FaceAlpha',0.5,'Normalization','probability'); grid on,grid minor, xlim([-1 1]), hold on, xline(0,'Color','k','linewidth',1.5); hold off
            title(sprintf('Histogram of voxel-to-voxel GRAY MATTER correlations AFTER 13-denoise4-WMregress (all slices combined)\n%.1f%% of the correlations were > 0  [ mean=%.2f, median=%.2f, SD=%.2f ]',100*length(find(C3g_>0))/length(C3g_),mean(C3g_),median(C3g_),std(C3g_)),'Color','k','FontSize',14)
        subplot2n(4,1,3), histogram(C3w_,'EdgeColor','none','FaceColor','k','FaceAlpha',0.5,'Normalization','probability'); grid on,grid minor, xlim([-1 1]), hold on, xline(0,'Color','k','linewidth',1.5); hold off
            title(sprintf('WHITE MATTER: AFTER 13-denoise4-WMregress (all slices): %.1f%% of the correlations were > 0  [ mean=%.2f, median=%.2f, SD=%.2f ]',100*length(find(C3w_>0))/length(C3w_),mean(C3w_),median(C3w_),std(C3w_)),'Color','k','FontSize',14)
    elseif ((stp==4)||(stp==5)||(stp==7)||(stp==8))&&(msk==1)
    	subplot2n(4,1,1), histogram(C3g_,'EdgeColor','none','FaceColor','k','FaceAlpha',0.5,'Normalization','probability'); grid on,grid minor, xlim([-1 1]), hold on, xline(0,'Color','k','linewidth',1.5); hold off
            title(sprintf('Histogram of voxel-to-voxel GRAY MATTER correlations AFTER 14-denoise5-COVregress (all slices combined)\n%.1f%% of the correlations were > 0  [ mean=%.2f, median=%.2f, SD=%.2f ]',100*length(find(C3g_>0))/length(C3g_),mean(C3g_),median(C3g_),std(C3g_)),'Color','k','FontSize',14)
        subplot2n(4,1,3), histogram(C3w_,'EdgeColor','none','FaceColor','k','FaceAlpha',0.5,'Normalization','probability'); grid on,grid minor, xlim([-1 1]), hold on, xline(0,'Color','k','linewidth',1.5); hold off
            title(sprintf('WHITE MATTER: AFTER 14-denoise5-COVregress (all slices): %.1f%% of the correlations were > 0  [ mean=%.2f, median=%.2f, SD=%.2f ]',100*length(find(C3w_>0))/length(C3w_),mean(C3w_),median(C3w_),std(C3w_)),'Color','k','FontSize',14)
    elseif (msk==2)
    	subplot2n(4,1,1), histogram(C3g_,'EdgeColor','none','FaceColor','k','FaceAlpha',0.5,'Normalization','probability'); grid on,grid minor, xlim([-1 1]), hold on, xline(0,'Color','k','linewidth',1.5); hold off
            title(sprintf('Histogram of voxel-to-voxel GRAY MATTER correlations AFTER 15-bandpass-filter (all slices combined)\n%.1f%% of the correlations were > 0  [ mean=%.2f, median=%.2f, SD=%.2f ]',100*length(find(C3g_>0))/length(C3g_),mean(C3g_),median(C3g_),std(C3g_)),'Color','k','FontSize',14)
        subplot2n(4,1,3), histogram(C3w_,'EdgeColor','none','FaceColor','k','FaceAlpha',0.5,'Normalization','probability'); grid on,grid minor, xlim([-1 1]), hold on, xline(0,'Color','k','linewidth',1.5); hold off
            title(sprintf('WHITE MATTER: AFTER 15-bandpass-filter (all slices): %.1f%% of the correlations were > 0  [ mean=%.2f, median=%.2f, SD=%.2f ]',100*length(find(C3w_>0))/length(C3w_),mean(C3w_),median(C3w_),std(C3w_)),'Color','k','FontSize',14)
    end
    subplot2n(4,1,2), histogram(Cg_,'EdgeColor','none','FaceColor',[0,0.55,0.69],'FaceAlpha',0.5,'Normalization','probability'); grid on,grid minor, xlim([-1 1]), hold on, xline(0,'Color','k','linewidth',1.5); hold off
       title(sprintf('GRAY MATTER: AFTER 17-HRF-deconvolution (all slices): %.1f%% of correlations > 0  [ mean=%.2f, median=%.2f, SD=%.2f ]',100*length(find(Cg_>0))/length(Cg_),mean(Cg_),median(Cg_),std(Cg_)),'Color',[0,0.55,0.69],'FontSize',14)
    subplot2n(4,1,4), histogram(Cw_,'EdgeColor','none','FaceColor',[0.65,0.11,0.19],'FaceAlpha',0.5,'Normalization','probability'); grid on,grid minor, xlim([-1 1]), hold on, xline(0,'Color','k','linewidth',1.5); hold off
       title(sprintf('WHITE MATTER: AFTER 17-HRF-deconvolution (all slices): %.1f%% of the correlations were > 0  [ mean=%.2f, median=%.2f, SD=%.2f ]',100*length(find(Cw_>0))/length(Cw_),mean(Cw_),median(Cw_),std(Cw_)),'Color',[0.65,0.11,0.19],'FontSize',14)
    saveas(fighndl,[QCpath 'hist_FC_compare_15filter_17deconv_allSlices' '.jpg'])
    saveas(fighndl,[QCpath2 'hist_FC_compare_15filter_17deconv_allSlices' '.jpg'])
     close(fighndl)
end
clear svar1 spsc1 svar2 spsc2 sortorder1 sortorder2 F2g_ mF2g_ F2w_ mF2w_ Cg_ Cw_ F4_ mF4_ C2_ F6g_ F6w_ mF6g_ mF6w_ C3g_ C3w_ cmap

try
if nargin<6
    close(wbar3)
end
catch
end

end
```
