## Supplementary material for "Neptune: a toolbox for spinal cord functional MRI data processing and quality assurance": Neptune software: scfMRItb_18_PAM50registration.m

```
function scfMRItb_18_PAM50registration(base_dir_sub,fname_anat_orig, varargin)

% this works on, at least, SCT v5.4 and later (I tested it with SCT v5.4 only)
fname_anat = [fname_anat_orig '_reduced' '2']; % reduced2 is the anatomical with clipped sides to focus mainly on the spinal cord
scfMRItb_04_unzipFile(base_dir_sub, fname_anat, '')

if nargin<3
    wbar3 = waitbar(0,'11. Co-registration to standard space (PAM50)...','Name','Progress(11): Co-registration to standard space (PAM50)...','Units','normalized','Position',[3/5 0 1/5 1/17]);
else
    wbar3 = varargin{1};
end

if nargin<4
    C_spine = 1; % 1 = assume that we are dealing with the cervical spinal cord
else
    C_spine = varargin{2};
end

if nargin<5
    SCT_vert_auto = 1; % 1 = assume that we will perform automatic labeling of vertebrae
else
    SCT_vert_auto = varargin{3};
end

if nargin<6
    SCT_manualreg_labels = '3,5'; % Vertebrae for manual labeling in SCT (if manual option was chosen), needed for registration
else
    SCT_manualreg_labels = varargin{4};
end

try waitbar(0.5,wbar3,'18. Co-registration PAM50'); catch, end

%% Use SCT to get an estimate of GM, WM and CSF boundaries

fprintf('\n18. Running SCT to estimate the spinal cord boundary...\n\n')
if ~(exist([base_dir_sub 'QC' '/SCT'],'dir'))
    mkdir([base_dir_sub 'QC' '/SCT'])
end
if ~exist([base_dir_sub fname_anat '_seg.nii'],'file')
    unix(['sct_deepseg_sc -i ' base_dir_sub fname_anat '.nii' ' -c t2s -ofolder ' base_dir_sub ' -qc ' base_dir_sub 'QC/SCT'])
end
fprintf('\n18. Completed running SCT to estimate the spinal cord boundary.\n\n')


% Vertebral labeling
if SCT_vert_auto==1 % automatic labeling
try
% Check QC report once done at: ./QC/SCT/index.html
% Note: Here, two files are output: _seg_labeled, which represents the 
% labeled segmentation (i.e., the value corresponds to the vertebral level), 
% and _seg_labeled_discs, which only has a single point for each inter-vertebral disc level.
% The convention is: Value 3 ?> C2-C3 disc, Value 4 ?> C3-C4 disc, etc.
unix(['sct_label_vertebrae -i ' base_dir_sub fname_anat '.nii' ' -s ' base_dir_sub fname_anat '_seg.nii' ' -c t2 -ofolder ' base_dir_sub ' -qc ' base_dir_sub 'QC/SCT'])

% If automatic labeling did not work, then perform manual identification of C2-C3 disc 
% and then redo vertebral labeling
if ~( exist([base_dir_sub fname_anat '_seg_labeled.nii'],'file') || exist([base_dir_sub fname_anat '_seg_labeled.nii.gz'],'file') )
    fprintf('\n18. Automatic labeling of vertebrae in SCT did not work; hence, manual identification of C2-C3 disc has been initiated.\nA viewer has popped out and you just have to click at the posterior tip of C2-C3 inter-vertebral disc.\n\n')
    unix(['sct_label_utils -i ' base_dir_sub fname_anat '.nii' ' -create-viewer 3 -o ' base_dir_sub fname_anat '_label_c2c3.nii.gz' ' -msg "Click at the posterior tip of C2/C3 inter-vertebral disc"'])
    unix(['sct_label_vertebrae -i ' base_dir_sub fname_anat '.nii' ' -s ' base_dir_sub fname_anat '_seg.nii' ' -c t2 -initlabel ' base_dir_sub fname_anat '_label_c2c3.nii.gz' ' -ofolder ' base_dir_sub ' -qc ' base_dir_sub 'QC/SCT'])
end
seg_labeled_discs = load_untouch_nii([base_dir_sub fname_anat '_seg_labeled_discs.nii']); seg_labeled_discs=seg_labeled_discs.img;

% Create labels at C3 (if c-spine) or T2 (if t-spine) mid-vertebral levels. These labels are needed for template registration.
if C_spine==1
vertbody2 = 3+length(find(seg_labeled_discs(:)~=0))-1; clear seg_labeled_discs
unix(['sct_label_utils -i ' base_dir_sub fname_anat '_seg_labeled.nii' ' -vert-body 3,' num2str(vertbody2) ' -o ' base_dir_sub fname_anat '_labels_vert.nii.gz' ' -qc ' base_dir_sub 'QC/SCT'])
else
vertbody2 = 9+length(find(seg_labeled_discs(:)~=0))-1; clear seg_labeled_discs
unix(['sct_label_utils -i ' base_dir_sub fname_anat '_seg_labeled.nii' ' -vert-body 9,' num2str(vertbody2) ' -o ' base_dir_sub fname_anat '_labels_vert.nii.gz' ' -qc ' base_dir_sub 'QC/SCT'])
end
% Generate a QC report to visualize the two selected labels on the anatomical image
unix(['sct_qc -i ' base_dir_sub fname_anat '.nii' ' -s ' base_dir_sub fname_anat '_labels_vert.nii.gz' ' -p sct_label_utils ' ' -qc ' base_dir_sub 'QC/SCT'])


catch
% If the automatic labeling procedure fails then completely bypass sct_label_vertebrae and do the labeling manually.
% In that case, SCT provides a viewer to do so conveniently.
unix(['sct_label_utils -i ' base_dir_sub fname_anat '.nii' ' -create-viewer ' SCT_manualreg_labels ' -o ' base_dir_sub fname_anat '_labels_disc.nii.gz' ' -msg "Place labels at the posterior tip of each inter-vertebral disc. E.g. Label 3: C2/C3, Label 4: C3/C4, etc."' ' -qc ' base_dir_sub 'QC/SCT'])
% WARNING: If you provide more than 2 labels, there will be a non-linear transformation along z, which implies that everything above the top label and below the bottom label
% will be lost in the transformation. Therefore, if you are interested in regions outside of the specified labels, only use one or two labels. No more.
end


% Register t2->template.
unix(['sct_register_to_template -i ' base_dir_sub fname_anat '.nii' ' -s ' base_dir_sub fname_anat '_seg.nii' ' -ldisc ' base_dir_sub fname_anat '_labels_disc.nii.gz'  ' -c t2s' ' -ofolder ' base_dir_sub ' -qc ' base_dir_sub 'QC/SCT'])


elseif SCT_vert_auto==0 %manual labeling
% You might want to completely bypass sct_label_vertebrae and do the labeling manually.
% In that case, SCT provides a viewer to do so conveniently.
unix(['sct_label_utils -i ' base_dir_sub fname_anat '.nii' ' -create-viewer ' SCT_manualreg_labels ' -o ' base_dir_sub fname_anat '_labels_disc.nii.gz' ' -msg "Place labels at the posterior tip of each inter-vertebral disc. E.g. Label 3: C2/C3, Label 4: C3/C4, etc."' ' -qc ' base_dir_sub 'QC/SCT'])
% WARNING: If you provide more than 2 labels, there will be a non-linear transformation along z, which implies that everything above the top label and below the bottom label
% will be lost in the transformation. Therefore, if you are interested in regions outside of the specified labels, only use one or two labels. No more.

% Register t2->template.
unix(['sct_register_to_template -i ' base_dir_sub fname_anat '.nii' ' -s ' base_dir_sub fname_anat '_seg.nii' ' -ldisc ' base_dir_sub fname_anat '_labels_disc.nii.gz'  ' -c t2s' ' -ofolder ' base_dir_sub ' -qc ' base_dir_sub 'QC/SCT'])

end


% Warp template objects (T2, cord segmentation, vertebral levels, etc.).
unix(['sct_warp_template -d ' base_dir_sub fname_anat '.nii' ' -w ' base_dir_sub 'warp_template2anat.nii.gz' ' -a 1' ' -s 1' ' -ofolder ' base_dir_sub ' -qc ' base_dir_sub 'QC/SCT'])

% Compute cross-sectional area (CSA) of spinal cord (separately per level and per slice)
unix(['sct_process_segmentation -i ' base_dir_sub fname_anat '_seg.nii' ' -vert 1:24 -perlevel 1' ' -vertfile ' base_dir_sub 'template/PAM50_levels.nii.gz' ' -o ' base_dir_sub 'CSA_perlevel_SCT.csv' ' -qc ' base_dir_sub 'QC/SCT'])
unix(['sct_process_segmentation -i ' base_dir_sub fname_anat '_seg.nii' ' -perslice 1' ' -vertfile ' base_dir_sub 'template/PAM50_levels.nii.gz' ' -o ' base_dir_sub 'CSA_perslice_SCT.csv' ' -qc ' base_dir_sub 'QC/SCT'])

try
if exist([base_dir_sub 'SCT_PAM50registration_step18'],'dir')
    tmp=clock; clk_settings = sprintf('%.4d-%.2d-%.2d_%.2d-%.2d',tmp(1),tmp(2),tmp(3),tmp(4),tmp(5)); clear tmp
    movefile([base_dir_sub 'SCT_PAM50registration_step18'],[base_dir_sub sprintf('SCT_PAM50registration_step18_old_%s',clk_settings)],'f')
    clear clk_settings
end
movefile([base_dir_sub 'template'],[base_dir_sub 'SCT_PAM50registration_step18'],'f')
movefile([base_dir_sub 'spinal_levels'],[base_dir_sub 'SCT_PAM50registration_step18/spinal_levels'],'f')
movefile([base_dir_sub 'atlas'],[base_dir_sub 'SCT_PAM50registration_step18/atlas'],'f')

gzip([base_dir_sub 'anat2template.nii']); delete([base_dir_sub 'anat2template.nii']);
movefile([base_dir_sub 'anat2template.nii.gz'],[base_dir_sub 'SCT_PAM50registration_step18/' 'anat2template.nii.gz'],'f')

gzip([base_dir_sub 'template2anat.nii']); delete([base_dir_sub 'template2anat.nii']);
movefile([base_dir_sub 'template2anat.nii.gz'],[base_dir_sub 'SCT_PAM50registration_step18/' 'template2anat.nii.gz'],'f')

movefile([base_dir_sub 'warp_anat2template.nii.gz'],[base_dir_sub 'SCT_PAM50registration_step18/' 'warp_anat2template.nii.gz'],'f')
movefile([base_dir_sub 'warp_curve2straight.nii.gz'],[base_dir_sub 'SCT_PAM50registration_step18/' 'warp_curve2straight.nii.gz'],'f')
movefile([base_dir_sub 'warp_straight2curve.nii.gz'],[base_dir_sub 'SCT_PAM50registration_step18/' 'warp_straight2curve.nii.gz'],'f')
movefile([base_dir_sub 'warp_template2anat.nii.gz'],[base_dir_sub 'SCT_PAM50registration_step18/' 'warp_template2anat.nii.gz'],'f')
movefile([base_dir_sub 'straight_ref.nii.gz'],[base_dir_sub 'SCT_PAM50registration_step18/' 'straight_ref.nii.gz'],'f')
movefile([base_dir_sub 'straightening.cache'],[base_dir_sub 'SCT_PAM50registration_step18/' 'straightening.cache'],'f')
catch
end

try
if nargin<3
    close(wbar3)
end
catch
end

end
```
