## Supplementary material for "Neptune: a toolbox for spinal cord functional MRI data processing and quality assurance": Neptune software: scfMRItb_19_gunzip_niigz.m

```
function skip = scfMRItb_19_gunzip_niigz(subdir, varargin)

if nargin<2
    wbar3 = waitbar(0,'00. unzip .nii.gz files...','Name','Progress(19): unzip .nii.gz files...','Units','normalized','Position',[3/5 0 1/5 1/17]);
else
    wbar3 = varargin{1};
end

%% ----- GUNZIP .nii.gz FILES ----------------------------------

sdir = dir([subdir '*.nii.gz']);

skip=0;
for k=1:length(sdir)
    if ~(exist([subdir sdir(k).name(1:end-3)], 'file'))
        skip=skip+1;
        try waitbar((k/length(sdir)),wbar3,sprintf('00. unzip .nii.gz files: file (%d) of (%d)',k,length(sdir))); catch, end
        fprintf('00. unzip .nii.gz files: file (%d) of (%d)\n',k,length(sdir))
        gunzip([sdir(k).folder '/' sdir(k).name])
    end
end

try
if nargin<2
    close(wbar3)
end
catch
end

end
```
