## Supplementary material for "Neptune: a toolbox for spinal cord functional MRI data processing and quality assurance": Neptune software: scfMRItb_19_gzip_nii.m

```
function skip = scfMRItb_19_gzip_nii(base_dir_sub, varargin)

if nargin<2
    wbar3 = waitbar(0,'19. gzip NIfTI files...','Name','Progress(19): gzip NIfTI files...','Units','normalized','Position',[3/5 0 1/5 1/17]);
else
    wbar3 = varargin{1};
end

%% ----- GZIP NIFTI FILES ----------------------------------

sdir = dir([base_dir_sub '*.nii']);

skip=0;
for i5=1:length(sdir)
    if ~(exist([base_dir_sub sdir(i5).name(1:end-4) '.nii.gz'], 'file'))
        skip=skip+1;
        try waitbar((i5/length(sdir)),wbar3,sprintf('19. gzip NIfTI files: file (%d) of (%d)',i5,length(sdir))); catch, end
        fprintf('\t19. gzip NIfTI files: file (%d) of (%d)\n',i5,length(sdir))
        gzip([sdir(i5).folder '/' sdir(i5).name])
    end
end

try
if nargin<2
    close(wbar3)
end
catch
end

end
```
