## Supplementary material for "Neptune: a toolbox for spinal cord functional MRI data processing and quality assurance": Neptune software: scfMRItb_19_remove_slicewise_files.m

```
function scfMRItb_19_remove_slicewise_files(base_dir_sub, fname4, fname_anat_orig, varargin)

% All slice-wise images have already been combined for all slices and saved 
% as 4D NIfTI images in step-14 (or as 3D images in case of anatomical images). 
% Hence, this code will delete those redundant slice-wise files.
%
% If you run any part of Neptune in the future on this subject again, it
% will automatically regenerate the slice-wise files as required.

if nargin<4
    wbar3 = waitbar(0,'19. Remove slice-wise files...','Name','Progress(19): Remove slice-wise files...','Units','normalized','Position',[3/5 0 1/5 1/17]);
else
    wbar3 = varargin{1};
end

fname_anat = [fname_anat_orig '_reduced'];
try
    A = load_untouch_nii([base_dir_sub fname_anat '.nii']);
catch
    gunzip([base_dir_sub fname_anat '.nii.gz'])
    A = load_untouch_nii([base_dir_sub fname_anat '.nii']);
    if exist([base_dir_sub fname_anat '.nii.gz'],'file') && exist([base_dir_sub fname_anat '.nii'],'file')
        delete([base_dir_sub fname_anat '.nii'])
    end
end
slices = size(A.img,3);
warning off

%% delete slice-wise images because they have already been combined for all slices and saved as 4D NIfTI images (or as 3D NIfTI images in case of anatomical files)

% anatomical *_reduced.nii
sdir1 = []; sdir2 = []; sdir1z = []; sdir2z = [];
for i3=1:slices
    sdir1  = cat(1,sdir1,dir([base_dir_sub fname_anat '_slice' num2str(i3) '.nii']));
    sdir1z = cat(1,sdir1z,dir([base_dir_sub fname_anat '_slice' num2str(i3) '.nii.gz']));
end
sdir2  = cat(1,sdir2,dir([base_dir_sub fname_anat '.nii']));
sdir2z = cat(1,sdir2z,dir([base_dir_sub fname_anat '.nii.gz']));
if (~(isempty(sdir1)&&isempty(sdir1z))) && (~(isempty(sdir2)&&isempty(sdir2z))) && ( (length(sdir1)==slices*length(sdir2))||(length(sdir1)==slices*length(sdir2z))||(length(sdir1z)==slices*length(sdir2))||(length(sdir1z)==slices*length(sdir2z)) )  % if the number of slice-wise .nii files is equal to the number of 4D .nii files of same suffix times the number of slices, then it is safe to delete the slice-wise images (as they have been safely combined and saved as 4D images). Slice-wise images can be regenerated from the 4D images later if needed.
    for i5=1:length(sdir1)
        delete([sdir1(i5).folder '/' sdir1(i5).name])
    end
    for i5=1:length(sdir1z)
        delete([sdir1z(i5).folder '/' sdir1z(i5).name])
    end
end
clear sdir1 sdir2 sdir1z sdir2z i3 i5

% _beforeNORDIC
sdir1 = []; sdir2 = []; sdir1z = []; sdir2z = [];
for k2=1:length(fname4)
    for i3=1:slices
        sdir1  = cat(1,sdir1,dir([base_dir_sub fname4{k2} '_slice' num2str(i3) '_beforeNORDIC.nii']));
        sdir1z = cat(1,sdir1z,dir([base_dir_sub fname4{k2} '_slice' num2str(i3) '_beforeNORDIC.nii.gz']));
    end
    sdir2  = cat(1,sdir2,dir([base_dir_sub fname4{k2} '_beforeNORDIC.nii']));
    sdir2z = cat(1,sdir2z,dir([base_dir_sub fname4{k2} '_beforeNORDIC.nii.gz']));
end
if (~(isempty(sdir1)&&isempty(sdir1z))) && (~(isempty(sdir2)&&isempty(sdir2z))) && ( (length(sdir1)==slices*length(sdir2))||(length(sdir1)==slices*length(sdir2z))||(length(sdir1z)==slices*length(sdir2))||(length(sdir1z)==slices*length(sdir2z)) )  % if the number of slice-wise .nii files is equal to the number of 4D .nii files of same suffix times the number of slices, then it is safe to delete the slice-wise images (as they have been safely combined and saved as 4D images). Slice-wise images can be regenerated from the 4D images later if needed.
    for i5=1:length(sdir1)
        delete([sdir1(i5).folder '/' sdir1(i5).name])
    end
    for i5=1:length(sdir1z)
        delete([sdir1z(i5).folder '/' sdir1z(i5).name])
    end
end
clear sdir1 sdir2 sdir1z sdir2z k2 i3 i5

% _slice*.nii
sdir1 = []; sdir2 = []; sdir1z = []; sdir2z = [];
for k2=1:length(fname4)
    for i3=1:slices
        sdir1  = cat(1,sdir1,dir([base_dir_sub fname4{k2} '_slice' num2str(i3) '.nii']));
        sdir1z = cat(1,sdir1z,dir([base_dir_sub fname4{k2} '_slice' num2str(i3) '.nii.gz']));
    end
    sdir2  = cat(1,sdir2,dir([base_dir_sub fname4{k2} '.nii']));
    sdir2z = cat(1,sdir2z,dir([base_dir_sub fname4{k2} '.nii.gz']));
end
if ~(isempty(sdir1)&&isempty(sdir1z)) && (~(isempty(sdir2)&&isempty(sdir2z))) 
    for i5=1:length(sdir1)
        delete([sdir1(i5).folder '/' sdir1(i5).name])
    end
    for i5=1:length(sdir1z)
        delete([sdir1z(i5).folder '/' sdir1z(i5).name])
    end
end
clear sdir1 sdir2 sdir1z sdir2z k2 i3 i5

% _Smask
sdir1 = []; sdir2 = []; sdir1z = []; sdir2z = [];
for k2=1:length(fname4)
    for i3=1:slices
        sdir1  = cat(1,sdir1,dir([base_dir_sub fname4{k2} '_slice' num2str(i3) '_Smask.nii']));
        sdir1z = cat(1,sdir1z,dir([base_dir_sub fname4{k2} '_slice' num2str(i3) '_Smask.nii.gz']));
    end
    sdir2  = cat(1,sdir2,dir([base_dir_sub fname4{k2} '_Smask.nii']));
    sdir2z = cat(1,sdir2z,dir([base_dir_sub fname4{k2} '_Smask.nii.gz']));
end
if (~(isempty(sdir1)&&isempty(sdir1z))) && (~(isempty(sdir2)&&isempty(sdir2z))) && ( (length(sdir1)==slices*length(sdir2))||(length(sdir1)==slices*length(sdir2z))||(length(sdir1z)==slices*length(sdir2))||(length(sdir1z)==slices*length(sdir2z)) )  % if the number of slice-wise .nii files is equal to the number of 4D .nii files of same suffix times the number of slices, then it is safe to delete the slice-wise images (as they have been safely combined and saved as 4D images). Slice-wise images can be regenerated from the 4D images later if needed.
    for i5=1:length(sdir1)
        delete([sdir1(i5).folder '/' sdir1(i5).name])
    end
    for i5=1:length(sdir1z)
        delete([sdir1z(i5).folder '/' sdir1z(i5).name])
    end
end
clear sdir1 sdir2 sdir1z sdir2z k2 i3 i5

% _base
sdir1 = []; sdir2 = []; sdir1z = []; sdir2z = [];
for k2=1:length(fname4)
    for i3=1:slices
        sdir1  = cat(1,sdir1,dir([base_dir_sub fname4{k2} '_slice' num2str(i3) '_base.nii']));
        sdir1z = cat(1,sdir1z,dir([base_dir_sub fname4{k2} '_slice' num2str(i3) '_base.nii.gz']));
    end
    sdir2  = cat(1,sdir2,dir([base_dir_sub fname4{k2} '_base.nii']));
    sdir2z = cat(1,sdir2z,dir([base_dir_sub fname4{k2} '_base.nii.gz']));
end
if (~(isempty(sdir1)&&isempty(sdir1z))) && (~(isempty(sdir2)&&isempty(sdir2z))) && ( (length(sdir1)==slices*length(sdir2))||(length(sdir1)==slices*length(sdir2z))||(length(sdir1z)==slices*length(sdir2))||(length(sdir1z)==slices*length(sdir2z)) )  % if the number of slice-wise .nii files is equal to the number of 4D .nii files of same suffix times the number of slices, then it is safe to delete the slice-wise images (as they have been safely combined and saved as 4D images). Slice-wise images can be regenerated from the 4D images later if needed.
    for i5=1:length(sdir1)
        delete([sdir1(i5).folder '/' sdir1(i5).name])
    end
    for i5=1:length(sdir1z)
        delete([sdir1z(i5).folder '/' sdir1z(i5).name])
    end
end
clear sdir1 sdir2 sdir1z sdir2z k2 i3 i5

% _base_masked
sdir1 = []; sdir2 = []; sdir1z = []; sdir2z = [];
for k2=1:length(fname4)
    for i3=1:slices
        sdir1  = cat(1,sdir1,dir([base_dir_sub fname4{k2} '_slice' num2str(i3) '_base_masked.nii']));
        sdir1z = cat(1,sdir1z,dir([base_dir_sub fname4{k2} '_slice' num2str(i3) '_base_masked.nii.gz']));
    end
    sdir2  = cat(1,sdir2,dir([base_dir_sub fname4{k2} '_base_masked.nii']));
    sdir2z = cat(1,sdir2z,dir([base_dir_sub fname4{k2} '_base_masked.nii.gz']));
end
if (~(isempty(sdir1)&&isempty(sdir1z))) && (~(isempty(sdir2)&&isempty(sdir2z))) && ( (length(sdir1)==slices*length(sdir2))||(length(sdir1)==slices*length(sdir2z))||(length(sdir1z)==slices*length(sdir2))||(length(sdir1z)==slices*length(sdir2z)) )  % if the number of slice-wise .nii files is equal to the number of 4D .nii files of same suffix times the number of slices, then it is safe to delete the slice-wise images (as they have been safely combined and saved as 4D images). Slice-wise images can be regenerated from the 4D images later if needed.
    for i5=1:length(sdir1)
        delete([sdir1(i5).folder '/' sdir1(i5).name])
    end
    for i5=1:length(sdir1z)
        delete([sdir1z(i5).folder '/' sdir1z(i5).name])
    end
end
clear sdir1 sdir2 sdir1z sdir2z k2 i3 i5

% _denoised_before_MC80
sdir1 = []; sdir2 = []; sdir1z = []; sdir2z = [];
for k2=1:length(fname4)
    for i3=1:slices
        sdir1  = cat(1,sdir1,dir([base_dir_sub fname4{k2} '_slice' num2str(i3) '_denoised_before_MC80.nii']));
        sdir1z = cat(1,sdir1z,dir([base_dir_sub fname4{k2} '_slice' num2str(i3) '_denoised_before_MC80.nii.gz']));
    end
    sdir2  = cat(1,sdir2,dir([base_dir_sub fname4{k2} '_denoised_before_MC80.nii']));
    sdir2z = cat(1,sdir2z,dir([base_dir_sub fname4{k2} '_denoised_before_MC80.nii.gz']));
end
if (~(isempty(sdir1)&&isempty(sdir1z))) && (~(isempty(sdir2)&&isempty(sdir2z))) && ( (length(sdir1)==slices*length(sdir2))||(length(sdir1)==slices*length(sdir2z))||(length(sdir1z)==slices*length(sdir2))||(length(sdir1z)==slices*length(sdir2z)) )  % if the number of slice-wise .nii files is equal to the number of 4D .nii files of same suffix times the number of slices, then it is safe to delete the slice-wise images (as they have been safely combined and saved as 4D images). Slice-wise images can be regenerated from the 4D images later if needed.
    for i5=1:length(sdir1)
        delete([sdir1(i5).folder '/' sdir1(i5).name])
    end
    for i5=1:length(sdir1z)
        delete([sdir1z(i5).folder '/' sdir1z(i5).name])
    end
end
clear sdir1 sdir2 sdir1z sdir2z k2 i3 i5

% _Gaussian_mask
sdir1 = []; sdir2 = []; sdir1z = []; sdir2z = [];
for k2=1:length(fname4)
    for i3=1:slices
        sdir1  = cat(1,sdir1,dir([base_dir_sub fname4{k2} '_slice' num2str(i3) '_Gaussian_mask.nii']));
        sdir1z = cat(1,sdir1z,dir([base_dir_sub fname4{k2} '_slice' num2str(i3) '_Gaussian_mask.nii.gz']));
    end
    sdir2  = cat(1,sdir2,dir([base_dir_sub fname4{k2} '_Gaussian_mask.nii']));
    sdir2z = cat(1,sdir2z,dir([base_dir_sub fname4{k2} '_Gaussian_mask.nii.gz']));
end
if (~(isempty(sdir1)&&isempty(sdir1z))) && (~(isempty(sdir2)&&isempty(sdir2z))) && ( (length(sdir1)==slices*length(sdir2))||(length(sdir1)==slices*length(sdir2z))||(length(sdir1z)==slices*length(sdir2))||(length(sdir1z)==slices*length(sdir2z)) )  % if the number of slice-wise .nii files is equal to the number of 4D .nii files of same suffix times the number of slices, then it is safe to delete the slice-wise images (as they have been safely combined and saved as 4D images). Slice-wise images can be regenerated from the 4D images later if needed.
    for i5=1:length(sdir1)
        delete([sdir1(i5).folder '/' sdir1(i5).name])
    end
    for i5=1:length(sdir1z)
        delete([sdir1z(i5).folder '/' sdir1z(i5).name])
    end
end
clear sdir1 sdir2 sdir1z sdir2z k2 i3 i5

% _MC
sdir1 = []; sdir2 = []; sdir1z = []; sdir2z = [];
for k2=1:length(fname4)
    for i3=1:slices
        sdir1  = cat(1,sdir1,dir([base_dir_sub fname4{k2} '_slice' num2str(i3) '_MC.nii']));
        sdir1z = cat(1,sdir1z,dir([base_dir_sub fname4{k2} '_slice' num2str(i3) '_MC.nii.gz']));
    end
    sdir2  = cat(1,sdir2,dir([base_dir_sub fname4{k2} '_MC.nii']));
    sdir2z = cat(1,sdir2z,dir([base_dir_sub fname4{k2} '_MC.nii.gz']));
end
if (~(isempty(sdir1)&&isempty(sdir1z))) && (~(isempty(sdir2)&&isempty(sdir2z))) && ( (length(sdir1)==slices*length(sdir2))||(length(sdir1)==slices*length(sdir2z))||(length(sdir1z)==slices*length(sdir2))||(length(sdir1z)==slices*length(sdir2z)) )  % if the number of slice-wise .nii files is equal to the number of 4D .nii files of same suffix times the number of slices, then it is safe to delete the slice-wise images (as they have been safely combined and saved as 4D images). Slice-wise images can be regenerated from the 4D images later if needed.
    for i5=1:length(sdir1)
        delete([sdir1(i5).folder '/' sdir1(i5).name])
    end
    for i5=1:length(sdir1z)
        delete([sdir1z(i5).folder '/' sdir1z(i5).name])
    end
end
clear sdir1 sdir2 sdir1z sdir2z k2 i3 i5

% _MC_mean
sdir1 = []; sdir2 = []; sdir1z = []; sdir2z = [];
for k2=1:length(fname4)
    for i3=1:slices
        sdir1  = cat(1,sdir1,dir([base_dir_sub fname4{k2} '_slice' num2str(i3) '_MC_mean.nii']));
        sdir1z = cat(1,sdir1z,dir([base_dir_sub fname4{k2} '_slice' num2str(i3) '_MC_mean.nii.gz']));
    end
    sdir2  = cat(1,sdir2,dir([base_dir_sub fname4{k2} '_MC_mean.nii']));
    sdir2z = cat(1,sdir2z,dir([base_dir_sub fname4{k2} '_MC_mean.nii.gz']));
end
if (~(isempty(sdir1)&&isempty(sdir1z))) && (~(isempty(sdir2)&&isempty(sdir2z))) && ( (length(sdir1)==slices*length(sdir2))||(length(sdir1)==slices*length(sdir2z))||(length(sdir1z)==slices*length(sdir2))||(length(sdir1z)==slices*length(sdir2z)) )  % if the number of slice-wise .nii files is equal to the number of 4D .nii files of same suffix times the number of slices, then it is safe to delete the slice-wise images (as they have been safely combined and saved as 4D images). Slice-wise images can be regenerated from the 4D images later if needed.
    for i5=1:length(sdir1)
        delete([sdir1(i5).folder '/' sdir1(i5).name])
    end
    for i5=1:length(sdir1z)
        delete([sdir1z(i5).folder '/' sdir1z(i5).name])
    end
end
clear sdir1 sdir2 sdir1z sdir2z k2 i3 i5

% _MCnofilt
sdir1 = []; sdir2 = []; sdir1z = []; sdir2z = [];
for k2=1:length(fname4)
    for i3=1:slices
        sdir1  = cat(1,sdir1,dir([base_dir_sub fname4{k2} '_slice' num2str(i3) '_MCnofilt.nii']));
        sdir1z = cat(1,sdir1z,dir([base_dir_sub fname4{k2} '_slice' num2str(i3) '_MCnofilt.nii.gz']));
    end
    sdir2  = cat(1,sdir2,dir([base_dir_sub fname4{k2} '_MCnofilt.nii']));
    sdir2z = cat(1,sdir2z,dir([base_dir_sub fname4{k2} '_MCnofilt.nii.gz']));
end
if (~(isempty(sdir1)&&isempty(sdir1z))) && (~(isempty(sdir2)&&isempty(sdir2z))) && ( (length(sdir1)==slices*length(sdir2))||(length(sdir1)==slices*length(sdir2z))||(length(sdir1z)==slices*length(sdir2))||(length(sdir1z)==slices*length(sdir2z)) )  % if the number of slice-wise .nii files is equal to the number of 4D .nii files of same suffix times the number of slices, then it is safe to delete the slice-wise images (as they have been safely combined and saved as 4D images). Slice-wise images can be regenerated from the 4D images later if needed.
    for i5=1:length(sdir1)
        delete([sdir1(i5).folder '/' sdir1(i5).name])
    end
    for i5=1:length(sdir1z)
        delete([sdir1z(i5).folder '/' sdir1z(i5).name])
    end
end
clear sdir1 sdir2 sdir1z sdir2z k2 i3 i5

% _MC_ricor
sdir1 = []; sdir2 = []; sdir1z = []; sdir2z = [];
for k2=1:length(fname4)
    for i3=1:slices
        sdir1  = cat(1,sdir1,dir([base_dir_sub fname4{k2} '_slice' num2str(i3) '_MC_ricor.nii']));
        sdir1z = cat(1,sdir1z,dir([base_dir_sub fname4{k2} '_slice' num2str(i3) '_MC_ricor.nii.gz']));
    end
    sdir2  = cat(1,sdir2,dir([base_dir_sub fname4{k2} '_MC_ricor.nii']));
    sdir2z = cat(1,sdir2z,dir([base_dir_sub fname4{k2} '_MC_ricor.nii.gz']));
end
if (~(isempty(sdir1)&&isempty(sdir1z))) && (~(isempty(sdir2)&&isempty(sdir2z))) && ( (length(sdir1)==slices*length(sdir2))||(length(sdir1)==slices*length(sdir2z))||(length(sdir1z)==slices*length(sdir2))||(length(sdir1z)==slices*length(sdir2z)) )  % if the number of slice-wise .nii files is equal to the number of 4D .nii files of same suffix times the number of slices, then it is safe to delete the slice-wise images (as they have been safely combined and saved as 4D images). Slice-wise images can be regenerated from the 4D images later if needed.
    for i5=1:length(sdir1)
        delete([sdir1(i5).folder '/' sdir1(i5).name])
    end
    for i5=1:length(sdir1z)
        delete([sdir1z(i5).folder '/' sdir1z(i5).name])
    end
end
clear sdir1 sdir2 sdir1z sdir2z k2 i3 i5

% _MC_ricor_mean
sdir1 = []; sdir2 = []; sdir1z = []; sdir2z = [];
for k2=1:length(fname4)
    for i3=1:slices
        sdir1  = cat(1,sdir1,dir([base_dir_sub fname4{k2} '_slice' num2str(i3) '_MC_ricor_mean.nii']));
        sdir1z = cat(1,sdir1z,dir([base_dir_sub fname4{k2} '_slice' num2str(i3) '_MC_ricor_mean.nii.gz']));
    end
    sdir2  = cat(1,sdir2,dir([base_dir_sub fname4{k2} '_MC_ricor_mean.nii']));
    sdir2z = cat(1,sdir2z,dir([base_dir_sub fname4{k2} '_MC_ricor_mean.nii.gz']));
end
if (~(isempty(sdir1)&&isempty(sdir1z))) && (~(isempty(sdir2)&&isempty(sdir2z))) && ( (length(sdir1)==slices*length(sdir2))||(length(sdir1)==slices*length(sdir2z))||(length(sdir1z)==slices*length(sdir2))||(length(sdir1z)==slices*length(sdir2z)) )  % if the number of slice-wise .nii files is equal to the number of 4D .nii files of same suffix times the number of slices, then it is safe to delete the slice-wise images (as they have been safely combined and saved as 4D images). Slice-wise images can be regenerated from the 4D images later if needed.
    for i5=1:length(sdir1)
        delete([sdir1(i5).folder '/' sdir1(i5).name])
    end
    for i5=1:length(sdir1z)
        delete([sdir1z(i5).folder '/' sdir1z(i5).name])
    end
end
clear sdir1 sdir2 sdir1z sdir2z k2 i3 i5

% _warped
sdir1 = []; sdir2 = []; sdir1z = []; sdir2z = [];
for k2=1:length(fname4)
    for i3=1:slices
        sdir1  = cat(1,sdir1,dir([base_dir_sub fname4{k2} '_slice' num2str(i3) '_warped.nii']));
        sdir1z = cat(1,sdir1z,dir([base_dir_sub fname4{k2} '_slice' num2str(i3) '_warped.nii.gz']));
    end
    sdir2  = cat(1,sdir2,dir([base_dir_sub fname4{k2} '_warped.nii']));
    sdir2z = cat(1,sdir2z,dir([base_dir_sub fname4{k2} '_warped.nii.gz']));
end
if (~(isempty(sdir1)&&isempty(sdir1z))) && (~(isempty(sdir2)&&isempty(sdir2z))) && ( (length(sdir1)==slices*length(sdir2))||(length(sdir1)==slices*length(sdir2z))||(length(sdir1z)==slices*length(sdir2))||(length(sdir1z)==slices*length(sdir2z)) )  % if the number of slice-wise .nii files is equal to the number of 4D .nii files of same suffix times the number of slices, then it is safe to delete the slice-wise images (as they have been safely combined and saved as 4D images). Slice-wise images can be regenerated from the 4D images later if needed.
    for i5=1:length(sdir1)
        delete([sdir1(i5).folder '/' sdir1(i5).name])
    end
    for i5=1:length(sdir1z)
        delete([sdir1z(i5).folder '/' sdir1z(i5).name])
    end
end
clear sdir1 sdir2 sdir1z sdir2z k2 i3 i5

% _base_warped
sdir1 = []; sdir2 = []; sdir1z = []; sdir2z = [];
for k2=1:length(fname4)
    for i3=1:slices
        sdir1  = cat(1,sdir1,dir([base_dir_sub fname4{k2} '_slice' num2str(i3) '_base_warped.nii']));
        sdir1z = cat(1,sdir1z,dir([base_dir_sub fname4{k2} '_slice' num2str(i3) '_base_warped.nii.gz']));
    end
    sdir2  = cat(1,sdir2,dir([base_dir_sub fname4{k2} '_base_warped.nii']));
    sdir2z = cat(1,sdir2z,dir([base_dir_sub fname4{k2} '_base_warped.nii.gz']));
end
if (~(isempty(sdir1)&&isempty(sdir1z))) && (~(isempty(sdir2)&&isempty(sdir2z))) && ( (length(sdir1)==slices*length(sdir2))||(length(sdir1)==slices*length(sdir2z))||(length(sdir1z)==slices*length(sdir2))||(length(sdir1z)==slices*length(sdir2z)) )  % if the number of slice-wise .nii files is equal to the number of 4D .nii files of same suffix times the number of slices, then it is safe to delete the slice-wise images (as they have been safely combined and saved as 4D images). Slice-wise images can be regenerated from the 4D images later if needed.
    for i5=1:length(sdir1)
        delete([sdir1(i5).folder '/' sdir1(i5).name])
    end
    for i5=1:length(sdir1z)
        delete([sdir1z(i5).folder '/' sdir1z(i5).name])
    end
end
clear sdir1 sdir2 sdir1z sdir2z k2 i3 i5

% _warped_mean
sdir1 = []; sdir2 = []; sdir1z = []; sdir2z = [];
for k2=1:length(fname4)
    for i3=1:slices
        sdir1  = cat(1,sdir1,dir([base_dir_sub fname4{k2} '_slice' num2str(i3) '_warped_mean.nii']));
        sdir1z = cat(1,sdir1z,dir([base_dir_sub fname4{k2} '_slice' num2str(i3) '_warped_mean.nii.gz']));
    end
    sdir2  = cat(1,sdir2,dir([base_dir_sub fname4{k2} '_warped_mean.nii']));
    sdir2z = cat(1,sdir2z,dir([base_dir_sub fname4{k2} '_warped_mean.nii.gz']));
end
if (~(isempty(sdir1)&&isempty(sdir1z))) && (~(isempty(sdir2)&&isempty(sdir2z))) && ( (length(sdir1)==slices*length(sdir2))||(length(sdir1)==slices*length(sdir2z))||(length(sdir1z)==slices*length(sdir2))||(length(sdir1z)==slices*length(sdir2z)) )  % if the number of slice-wise .nii files is equal to the number of 4D .nii files of same suffix times the number of slices, then it is safe to delete the slice-wise images (as they have been safely combined and saved as 4D images). Slice-wise images can be regenerated from the 4D images later if needed.
    for i5=1:length(sdir1)
        delete([sdir1(i5).folder '/' sdir1(i5).name])
    end
    for i5=1:length(sdir1z)
        delete([sdir1z(i5).folder '/' sdir1z(i5).name])
    end
end
clear sdir1 sdir2 sdir1z sdir2z k2 i3 i5

% anatomical *_affine_reg_mask
sdir1 = []; sdir2 = []; sdir1z = []; sdir2z = [];
for i3=1:slices
    sdir1  = cat(1,sdir1,dir([base_dir_sub fname_anat '_slice' num2str(i3) '_affine_reg_mask.nii']));
    sdir1z = cat(1,sdir1z,dir([base_dir_sub fname_anat '_slice' num2str(i3) '_affine_reg_mask.nii.gz']));
end
sdir2  = cat(1,sdir2,dir([base_dir_sub fname_anat '_affine_reg_mask.nii']));
sdir2z = cat(1,sdir2z,dir([base_dir_sub fname_anat '_affine_reg_mask.nii.gz']));
if (~(isempty(sdir1)&&isempty(sdir1z))) && (~(isempty(sdir2)&&isempty(sdir2z))) && ( (length(sdir1)==slices*length(sdir2))||(length(sdir1)==slices*length(sdir2z))||(length(sdir1z)==slices*length(sdir2))||(length(sdir1z)==slices*length(sdir2z)) )  % if the number of slice-wise .nii files is equal to the number of 4D .nii files of same suffix times the number of slices, then it is safe to delete the slice-wise images (as they have been safely combined and saved as 4D images). Slice-wise images can be regenerated from the 4D images later if needed.
    for i5=1:length(sdir1)
        delete([sdir1(i5).folder '/' sdir1(i5).name])
    end
    for i5=1:length(sdir1z)
        delete([sdir1z(i5).folder '/' sdir1z(i5).name])
    end
end
clear sdir1 sdir2 sdir1z sdir2z i3 i5

% anatomical *_reduced_0.nii
sdir1 = []; sdir2 = []; sdir1z = []; sdir2z = [];
for i3=1:slices
    sdir1  = cat(1,sdir1,dir([base_dir_sub fname_anat '_slice' num2str(i3) '_' num2str(0) '.nii']));
    sdir1z = cat(1,sdir1z,dir([base_dir_sub fname_anat '_slice' num2str(i3) '_' num2str(0) '.nii.gz']));
end
sdir2  = cat(1,sdir2,dir([base_dir_sub fname_anat '_' num2str(0) '.nii']));
sdir2z = cat(1,sdir2z,dir([base_dir_sub fname_anat '_' num2str(0) '.nii.gz']));
if (~(isempty(sdir1)&&isempty(sdir1z))) && (~(isempty(sdir2)&&isempty(sdir2z))) && ( (length(sdir1)==slices*length(sdir2))||(length(sdir1)==slices*length(sdir2z))||(length(sdir1z)==slices*length(sdir2))||(length(sdir1z)==slices*length(sdir2z)) )  % if the number of slice-wise .nii files is equal to the number of 4D .nii files of same suffix times the number of slices, then it is safe to delete the slice-wise images (as they have been safely combined and saved as 4D images). Slice-wise images can be regenerated from the 4D images later if needed.
    for i5=1:length(sdir1)
        delete([sdir1(i5).folder '/' sdir1(i5).name])
    end
    for i5=1:length(sdir1z)
        delete([sdir1z(i5).folder '/' sdir1z(i5).name])
    end
end
clear sdir1 sdir2 sdir1z sdir2z i3 i5

% _denoised50
try load([base_dir_sub 'workspace_variables_run1.mat'], 'cutoff_str'), catch, cutoff_str='50'; end
sdir1 = []; sdir2 = []; sdir1z = []; sdir2z = [];
for k2=1:length(fname4)
    for i3=1:slices
        sdir1  = cat(1,sdir1,dir([base_dir_sub fname4{k2} '_slice' num2str(i3) '_denoised' cutoff_str '.nii']));
        sdir1z = cat(1,sdir1z,dir([base_dir_sub fname4{k2} '_slice' num2str(i3) '_denoised' cutoff_str '.nii.gz']));
    end
    sdir2  = cat(1,sdir2,dir([base_dir_sub fname4{k2} '_denoised' cutoff_str '.nii']));
    sdir2z = cat(1,sdir2z,dir([base_dir_sub fname4{k2} '_denoised' cutoff_str '.nii.gz']));
end
if (~(isempty(sdir1)&&isempty(sdir1z))) && (~(isempty(sdir2)&&isempty(sdir2z))) && ( (length(sdir1)==slices*length(sdir2))||(length(sdir1)==slices*length(sdir2z))||(length(sdir1z)==slices*length(sdir2))||(length(sdir1z)==slices*length(sdir2z)) )  % if the number of slice-wise .nii files is equal to the number of 4D .nii files of same suffix times the number of slices, then it is safe to delete the slice-wise images (as they have been safely combined and saved as 4D images). Slice-wise images can be regenerated from the 4D images later if needed.
    for i5=1:length(sdir1)
        delete([sdir1(i5).folder '/' sdir1(i5).name])
    end
    for i5=1:length(sdir1z)
        delete([sdir1z(i5).folder '/' sdir1z(i5).name])
    end
end
clear sdir1 sdir2 sdir1z sdir2z k2 i3 i5

% _WM
sdir1 = []; sdir2 = []; sdir1z = []; sdir2z = [];
for k2=1:length(fname4)
    for i3=1:slices
        sdir1  = cat(1,sdir1,dir([base_dir_sub fname4{k2} '_slice' num2str(i3) '_WM.nii']));
        sdir1z = cat(1,sdir1z,dir([base_dir_sub fname4{k2} '_slice' num2str(i3) '_WM.nii.gz']));
    end
    sdir2  = cat(1,sdir2,dir([base_dir_sub fname4{k2} '_WM.nii']));
    sdir2z = cat(1,sdir2z,dir([base_dir_sub fname4{k2} '_WM.nii.gz']));
end
if (~(isempty(sdir1)&&isempty(sdir1z))) && (~(isempty(sdir2)&&isempty(sdir2z))) && ( (length(sdir1)==slices*length(sdir2))||(length(sdir1)==slices*length(sdir2z))||(length(sdir1z)==slices*length(sdir2))||(length(sdir1z)==slices*length(sdir2z)) )  % if the number of slice-wise .nii files is equal to the number of 4D .nii files of same suffix times the number of slices, then it is safe to delete the slice-wise images (as they have been safely combined and saved as 4D images). Slice-wise images can be regenerated from the 4D images later if needed.
    for i5=1:length(sdir1)
        delete([sdir1(i5).folder '/' sdir1(i5).name])
    end
    for i5=1:length(sdir1z)
        delete([sdir1z(i5).folder '/' sdir1z(i5).name])
    end
end
clear sdir1 sdir2 sdir1z sdir2z k2 i3 i5

% _cov
sdir1 = []; sdir2 = []; sdir1z = []; sdir2z = [];
for k2=1:length(fname4)
    for i3=1:slices
        sdir1  = cat(1,sdir1,dir([base_dir_sub fname4{k2} '_slice' num2str(i3) '_cov.nii']));
        sdir1z = cat(1,sdir1z,dir([base_dir_sub fname4{k2} '_slice' num2str(i3) '_cov.nii.gz']));
    end
    sdir2  = cat(1,sdir2,dir([base_dir_sub fname4{k2} '_cov.nii']));
    sdir2z = cat(1,sdir2z,dir([base_dir_sub fname4{k2} '_cov.nii.gz']));
end
if (~(isempty(sdir1)&&isempty(sdir1z))) && (~(isempty(sdir2)&&isempty(sdir2z))) && ( (length(sdir1)==slices*length(sdir2))||(length(sdir1)==slices*length(sdir2z))||(length(sdir1z)==slices*length(sdir2))||(length(sdir1z)==slices*length(sdir2z)) )  % if the number of slice-wise .nii files is equal to the number of 4D .nii files of same suffix times the number of slices, then it is safe to delete the slice-wise images (as they have been safely combined and saved as 4D images). Slice-wise images can be regenerated from the 4D images later if needed.
    for i5=1:length(sdir1)
        delete([sdir1(i5).folder '/' sdir1(i5).name])
    end
    for i5=1:length(sdir1z)
        delete([sdir1z(i5).folder '/' sdir1z(i5).name])
    end
end
clear sdir1 sdir2 sdir1z sdir2z k2 i3 i5

% _WMcov
sdir1 = []; sdir2 = []; sdir1z = []; sdir2z = [];
for k2=1:length(fname4)
    for i3=1:slices
        sdir1  = cat(1,sdir1,dir([base_dir_sub fname4{k2} '_slice' num2str(i3) '_WMcov.nii']));
        sdir1z = cat(1,sdir1z,dir([base_dir_sub fname4{k2} '_slice' num2str(i3) '_WMcov.nii.gz']));
    end
    sdir2  = cat(1,sdir2,dir([base_dir_sub fname4{k2} '_WMcov.nii']));
    sdir2z = cat(1,sdir2z,dir([base_dir_sub fname4{k2} '_WMcov.nii.gz']));
end
if (~(isempty(sdir1)&&isempty(sdir1z))) && (~(isempty(sdir2)&&isempty(sdir2z))) && ( (length(sdir1)==slices*length(sdir2))||(length(sdir1)==slices*length(sdir2z))||(length(sdir1z)==slices*length(sdir2))||(length(sdir1z)==slices*length(sdir2z)) )  % if the number of slice-wise .nii files is equal to the number of 4D .nii files of same suffix times the number of slices, then it is safe to delete the slice-wise images (as they have been safely combined and saved as 4D images). Slice-wise images can be regenerated from the 4D images later if needed.
    for i5=1:length(sdir1)
        delete([sdir1(i5).folder '/' sdir1(i5).name])
    end
    for i5=1:length(sdir1z)
        delete([sdir1z(i5).folder '/' sdir1z(i5).name])
    end
end
clear sdir1 sdir2 sdir1z sdir2z k2 i3 i5

% _denoised50WM
try load([base_dir_sub 'workspace_variables_run1.mat'], 'cutoff_str'), catch, cutoff_str='50'; end
sdir1 = []; sdir2 = []; sdir1z = []; sdir2z = [];
for k2=1:length(fname4)
    for i3=1:slices
        sdir1  = cat(1,sdir1,dir([base_dir_sub fname4{k2} '_slice' num2str(i3) '_denoised' cutoff_str 'WM.nii']));
        sdir1z = cat(1,sdir1z,dir([base_dir_sub fname4{k2} '_slice' num2str(i3) '_denoised' cutoff_str 'WM.nii.gz']));
    end
    sdir2  = cat(1,sdir2,dir([base_dir_sub fname4{k2} '_denoised' cutoff_str 'WM.nii']));
    sdir2z = cat(1,sdir2z,dir([base_dir_sub fname4{k2} '_denoised' cutoff_str 'WM.nii.gz']));
end
if (~(isempty(sdir1)&&isempty(sdir1z))) && (~(isempty(sdir2)&&isempty(sdir2z))) && ( (length(sdir1)==slices*length(sdir2))||(length(sdir1)==slices*length(sdir2z))||(length(sdir1z)==slices*length(sdir2))||(length(sdir1z)==slices*length(sdir2z)) )  % if the number of slice-wise .nii files is equal to the number of 4D .nii files of same suffix times the number of slices, then it is safe to delete the slice-wise images (as they have been safely combined and saved as 4D images). Slice-wise images can be regenerated from the 4D images later if needed.
    for i5=1:length(sdir1)
        delete([sdir1(i5).folder '/' sdir1(i5).name])
    end
    for i5=1:length(sdir1z)
        delete([sdir1z(i5).folder '/' sdir1z(i5).name])
    end
end
clear sdir1 sdir2 sdir1z sdir2z k2 i3 i5

% _denoised50cov
try load([base_dir_sub 'workspace_variables_run1.mat'], 'cutoff_str'), catch, cutoff_str='50'; end
sdir1 = []; sdir2 = []; sdir1z = []; sdir2z = [];
for k2=1:length(fname4)
    for i3=1:slices
        sdir1  = cat(1,sdir1,dir([base_dir_sub fname4{k2} '_slice' num2str(i3) '_denoised' cutoff_str 'cov.nii']));
        sdir1z = cat(1,sdir1z,dir([base_dir_sub fname4{k2} '_slice' num2str(i3) '_denoised' cutoff_str 'cov.nii.gz']));
    end
    sdir2  = cat(1,sdir2,dir([base_dir_sub fname4{k2} '_denoised' cutoff_str 'cov.nii']));
    sdir2z = cat(1,sdir2z,dir([base_dir_sub fname4{k2} '_denoised' cutoff_str 'cov.nii.gz']));
end
if (~(isempty(sdir1)&&isempty(sdir1z))) && (~(isempty(sdir2)&&isempty(sdir2z))) && ( (length(sdir1)==slices*length(sdir2))||(length(sdir1)==slices*length(sdir2z))||(length(sdir1z)==slices*length(sdir2))||(length(sdir1z)==slices*length(sdir2z)) )  % if the number of slice-wise .nii files is equal to the number of 4D .nii files of same suffix times the number of slices, then it is safe to delete the slice-wise images (as they have been safely combined and saved as 4D images). Slice-wise images can be regenerated from the 4D images later if needed.
    for i5=1:length(sdir1)
        delete([sdir1(i5).folder '/' sdir1(i5).name])
    end
    for i5=1:length(sdir1z)
        delete([sdir1z(i5).folder '/' sdir1z(i5).name])
    end
end
clear sdir1 sdir2 sdir1z sdir2z k2 i3 i5

% _denoised50WMcov
try load([base_dir_sub 'workspace_variables_run1.mat'], 'cutoff_str'), catch, cutoff_str='50'; end
sdir1 = []; sdir2 = []; sdir1z = []; sdir2z = [];
for k2=1:length(fname4)
    for i3=1:slices
        sdir1  = cat(1,sdir1,dir([base_dir_sub fname4{k2} '_slice' num2str(i3) '_denoised' cutoff_str 'WMcov.nii']));
        sdir1z = cat(1,sdir1z,dir([base_dir_sub fname4{k2} '_slice' num2str(i3) '_denoised' cutoff_str 'WMcov.nii.gz']));
    end
    sdir2  = cat(1,sdir2,dir([base_dir_sub fname4{k2} '_denoised' cutoff_str 'WMcov.nii']));
    sdir2z = cat(1,sdir2z,dir([base_dir_sub fname4{k2} '_denoised' cutoff_str 'WMcov.nii.gz']));
end
if (~(isempty(sdir1)&&isempty(sdir1z))) && (~(isempty(sdir2)&&isempty(sdir2z))) && ( (length(sdir1)==slices*length(sdir2))||(length(sdir1)==slices*length(sdir2z))||(length(sdir1z)==slices*length(sdir2))||(length(sdir1z)==slices*length(sdir2z)) )  % if the number of slice-wise .nii files is equal to the number of 4D .nii files of same suffix times the number of slices, then it is safe to delete the slice-wise images (as they have been safely combined and saved as 4D images). Slice-wise images can be regenerated from the 4D images later if needed.
    for i5=1:length(sdir1)
        delete([sdir1(i5).folder '/' sdir1(i5).name])
    end
    for i5=1:length(sdir1z)
        delete([sdir1z(i5).folder '/' sdir1z(i5).name])
    end
end
clear sdir1 sdir2 sdir1z sdir2z k2 i3 i5

try
if nargin<4
    close(wbar3)
end
catch
end
warning on

end
```
