## Supplementary material for "Neptune: a toolbox for spinal cord functional MRI data processing and quality assurance": Neptune software: scfMRItb_20_quadMask.m

```
function scfMRItb_20_quadMask(base_dir_sub,fname4,WM_flag,Rns)

scrsz = get(0,'ScreenSize');
pause on;

if ~(exist([base_dir_sub 'QC' '/20_quadrant_timeseries'],'dir'))
    mkdir([base_dir_sub 'QC' '/20_quadrant_timeseries'])
end
if Rns==0
    QCpath = [base_dir_sub 'QC' '/20_quadrant_timeseries/'];
else
    if ~(exist([base_dir_sub 'QC' '/20_quadrant_timeseries' '/run' num2str(Rns)],'dir'))
        mkdir([base_dir_sub 'QC' '/20_quadrant_timeseries' '/run' num2str(Rns)])
    end
    QCpath = [base_dir_sub 'QC' '/20_quadrant_timeseries' '/run' num2str(Rns) '/'];
end

if ~(exist([base_dir_sub 'QC' '/QC_shortlist'],'dir'))
    mkdir([base_dir_sub 'QC' '/QC_shortlist'])
end
if Rns==0
    QCpath2 = [base_dir_sub 'QC' '/QC_shortlist/'];
else
    if ~(exist([base_dir_sub 'QC' '/QC_shortlist' '/run' num2str(Rns)],'dir'))
        mkdir([base_dir_sub 'QC' '/QC_shortlist' '/run' num2str(Rns)])
    end
    QCpath2 = [base_dir_sub 'QC' '/QC_shortlist' '/run' num2str(Rns) '/'];
end

if Rns==0, run=1; else, run=Rns; end


% If this is set to 1, then QuadMask and time series are obtained for the
% white matter by default (future steps however depend on the original WM_flag
% setting)
do_WM_by_default = 1;

if do_WM_by_default==1
    WM_flag = 1;
end

%% ----- OBTAIN MASKS FOR EACH OF THE 4 QUADRANTS IN EACH SLICE ------------

load([base_dir_sub fname4 '.mat']) %#ok<*LOAD> % variable F
siz3 = size(F.img,3); siz4 = size(F.img,4);

load([base_dir_sub 'workspace_variables_run1' '.mat'],'FC_mask_quads','FC_mask_WM');
try FC_mask_quads=FC_mask_quads.img; catch, end

skip_erosion = 0; % erode gray matter mask (1 = do not erode; -1 = dilate)
wm_erode = 6; % larger the number, more eroded the WM masks. 1 is the smallest value that can be assigned (least erosion).

if max([F.hdr.dime.pixdim(2),F.hdr.dime.pixdim(3)])<=1 % less than 1mm in-plane voxel size
    dil1 = 2;
else
    dil1 = 1;
end

%% gray matter

for i3 = siz3:-1:1 
    F_oneslice = squeeze(F.img(:,:,i3,:));
    for i9 = 1 : 4 % for each quadrant (GM)
        mmask_orig = (FC_mask_quads(:,:,i3) == i9);
        if     i9 == 1 % left ventral
            mmask_ipsi = (FC_mask_quads(:,:,i3) == 3) | (FC_mask_quads(:,:,i3) == 2);
        elseif i9 == 2 % right ventral
            mmask_ipsi = (FC_mask_quads(:,:,i3) == 4) | (FC_mask_quads(:,:,i3) == 1);
        elseif i9 == 3 % left dorsal
            mmask_ipsi = (FC_mask_quads(:,:,i3) == 1) | (FC_mask_quads(:,:,i3) == 4);
        elseif i9 == 4 % right dorsal
            mmask_ipsi = (FC_mask_quads(:,:,i3) == 2) | (FC_mask_quads(:,:,i3) == 3);
        end % ipsilateral mask
        mmask_dilated = imdilate(mmask_orig, strel('disk', dil1));
        mmask_ipsi_dilated = imdilate(mmask_ipsi, strel('disk', dil1));
        mmask_overlap = (mmask_dilated + mmask_ipsi_dilated) > 1;
        mmask_overlap_dilated = imdilate(mmask_overlap, strel('disk', 1));
        mmask_valid = ~mmask_overlap_dilated;
        clear mmask_dilated mmask_ipsi mmask_ipsi_dilated mmask_overlap mmask_overlap_dilated;
        
        if skip_erosion == 0 % erode mask
            quadmask = imerode(mmask_orig, strel('disk', 1));
            if sum(sum( and(quadmask, mmask_valid) )) <= 3
                quadmask = mmask_orig; % mask is already tiny, so don't erode
            end
        elseif skip_erosion == 1 % don't erode mask
            quadmask = mmask_orig;
        elseif skip_erosion == -1 % a secret option --> dilate mask!
            quadmask = imdilate(mmask_orig, strel('disk', 1));
        end % ~skip_erosion
        
        % exclude region near central gray matter
        if ~( sum(sum( and(quadmask, mmask_valid) )) <= 3 ) % don't do this if mask is already tiny
            quadmask = and(quadmask, mmask_valid);
        end
        
        num_mask = sum(quadmask(:));
        [r, c] = find(quadmask);
        index = [r, c];

        ttemp = F_oneslice .* repmat(quadmask, [1 1 siz4]);
        temp = zeros(num_mask, siz4);
        for i1 = 1 : num_mask
            temp(i1,:) = reshape(ttemp(r(i1),c(i1),:), [1 siz4]);
        end
        clear r c;
        
        % save results
        timeseries_GM{i3,i9} = temp;
        coordinates_GM{i3,i9} = index;
        mask_quads_eroded_GM(:,:,i9,i3) = quadmask;
        
        clear temp mmask_valid mmask_orig mmask num_mask index quadmask
    end % i9
    clear F_oneslice
end % i3

%% white matter
if WM_flag == 1 % perform steps in the white matter (0 = don't)
for i3 = siz3:-1:1
    F_oneslice = squeeze(F.img(:,:,i3,:));
    for i9 = 5 : 8 % for each quadrant (WM)
        mmask_orig = FC_mask_quads(:,:,i3) == i9;
        
        if ~( sum(sum(mmask_orig)) <= 3 ) % don't do this if mask is already tiny
        imask = wm_erode;
        quadmask = imerode(mmask_orig, strel('disk', imask));
        while sum(quadmask(:)) <= 1
            if imask>=2
                imask = imask - 1;
                quadmask = imerode(mmask_orig, strel('disk', imask));
            else
                break;
            end
        end
        if (sum(quadmask(:))<=3), quadmask=mmask_orig; end
        
        % --- keeping the next two lines will create slightly larger WM quads than otherwise ---
        if imask>=2
            imask = imask - 1;
            quadmask = imerode(mmask_orig, strel('disk', imask));
        end
        if (sum(quadmask(:))<=3), quadmask=mmask_orig; end
        % --- ---
        else
        quadmask=mmask_orig;
        end
        
        num_mask = sum(quadmask(:));
        ttemp = F_oneslice .* repmat(quadmask, [1 1 siz4]);
        [r, c] = find(quadmask);
        index = [r, c];

        temp = zeros(num_mask, siz4);
        for i1 = 1 : num_mask
            temp(i1,:) = reshape(ttemp(r(i1),c(i1),:), [1 siz4]);
        end
        clear r c;
        
        % save results
        timeseries_WM{i3,i9-4} = temp;
        coordinates_WM{i3,i9-4} = index;
        mask_quads_eroded_WM(:,:,i9-4,i3) = quadmask;
        
        clear temp mmask_valid mmask_orig mmask num_mask index imask quadmask
    end % i9
    clear F_oneslice
end % i3
end % WM_flag

%% save
info_quads = sprintf('\nColumns/quadrants:\n1 = left ventral,\n2 = right ventral\n3 = left dorsal\n4 = right dorsal\n\ntimeseries = {slices x quadrants} cells with VxT matrix in each,\nwhere V = voxels in each quadrant, and T = time points\n\ncoordinates = matrix coordinates of the timeseries voxels in the pre-processed 3D+time data\n\nmask_quads_eroded = mask of each of the quadrants in the original 3D space\nwhere voxels of corresponding quadrants carry values 1 (LV), 2 (RV), 3 (LD) and 4 (RD), and a value of 0 everywhere else.\n\n');

if WM_flag == 1
save([base_dir_sub fname4 '_quads_timeseries' '.mat'], 'timeseries_GM', 'timeseries_WM', 'coordinates_GM', 'coordinates_WM', 'mask_quads_eroded_GM', 'mask_quads_eroded_WM', 'info_quads')
else
save([base_dir_sub fname4 '_quads_timeseries' '.mat'], 'timeseries_GM', 'coordinates_GM', 'mask_quads_eroded_GM', 'info_quads')
end

%% QC plots of masks
if run==1
quadmask = zeros(size(FC_mask_quads));
for i9=1:size(mask_quads_eroded_GM,3)
    quadmask = quadmask + i9.*single(squeeze(mask_quads_eroded_GM(:,:,i9,:)));
end
if WM_flag == 1
for i9=1:size(mask_quads_eroded_WM,3)
    quadmask = quadmask + (i9+4).*single(squeeze(mask_quads_eroded_WM(:,:,i9,:)));
end
end

for i3=1:siz3
    fprintf('20. QC masks: slice (%d) of (%d)\n',i3,siz3)
    tmp1 = FC_mask_quads(:,:,i3); tmp1(find(tmp1==0))=NaN;
    tmp1 = rot90(tmp1); [w1,w2] = find(rot90(FC_mask_WM(:,:,i3))~=0);
    tmp2 = quadmask(:,:,i3); tmp2(find(tmp2==0))=NaN;
    tmp2 = rot90(tmp2);
    fighndl = figure('Position',[round((scrsz(3)-scrsz(4))/2) 1 scrsz(4) scrsz(4)],'Color',[0.95,0.9,0.85],'InvertHardcopy','off','Visible','off'); % red [0.95,0.9,0.85], green [0.85,0.90,0.85], blue [0.89,0.94,0.99]
    subplot2n(2,1,1), imshow(tmp1, [0 max(tmp1(:))], 'InitialMagnification', 'fit'); colormap('jet'),colorbar;
        colormap([0 0 0; parula(256)])
        xlim([min(w2)-1,max(w2)+1]), ylim([min(w1)-1,max(w1)+1]);
        title(sprintf('ORIGINAL gray and white matter quadrants for slice %d of %d',i3,siz3),'Color','red','FontSize',18)    
    subplot2n(2,1,2), imshow(tmp2, [0 max(tmp1(:))], 'InitialMagnification', 'fit'); colormap('jet'),colorbar;
        colormap([0 0 0; parula(256)])
        xlim([min(w2)-1,max(w2)+1]), ylim([min(w1)-1,max(w1)+1]);
        title(sprintf('ERODED masks (for post-processing), slice %d of %d',i3,siz3),'Color','blue','FontSize',18)
    saveas(fighndl,[QCpath 'QuadMask_for_postproc_slice' sprintf('%.2d',i3) '.jpg'])
    close(fighndl)
    clear fighndl tmp1 tmp2 w1 w2
end
end

%% time series plots, global signal and FC histogram
    % (1) = left ventral GM; (2) = right ventral GM; (3) = left dorsal GM; (4) = right dorsal GM
    % (5) = left ventral WM; (6) = right ventral WM; (7) = left dorsal WM; (8) = right dorsal WM

load([base_dir_sub fname4 '.mat']) % variable F
F1 = single(F.img); clear F
cmap = colormap('bone'); cmap = (1-exp(-cmap))./(1-exp(-1)); 

F2g1_=[]; F2g2_=[]; F2g3_=[]; F2g4_=[]; mF2g1_=[]; mF2g2_=[]; mF2g3_=[]; mF2g4_=[]; 
Cg11_=[]; Cg22_=[]; Cg33_=[]; Cg44_=[]; Cg12_=[]; Cg13_=[]; Cg14_=[]; Cg23_=[]; Cg24_=[]; Cg34_=[];
if WM_flag==1
F2w1_=[]; F2w2_=[]; F2w3_=[]; F2w4_=[]; mF2w1_=[]; mF2w2_=[]; mF2w3_=[]; mF2w4_=[]; 
Cw11_=[]; Cw22_=[]; Cw33_=[]; Cw44_=[]; Cw12_=[]; Cw13_=[]; Cw14_=[]; Cw23_=[]; Cw24_=[]; Cw34_=[];
end

for i3 = 1 : siz3
    fprintf('20. QC plots: slice (%d) of (%d)\n',i3,siz3)
    
    for i5=1:size(F1,4)
        tmp = F1(:,:,i3,i5);
        F2g1(:,i5) = tmp(find(mask_quads_eroded_GM(:,:,1,i3)==1));
        F2g2(:,i5) = tmp(find(mask_quads_eroded_GM(:,:,2,i3)==1));
        F2g3(:,i5) = tmp(find(mask_quads_eroded_GM(:,:,3,i3)==1));
        F2g4(:,i5) = tmp(find(mask_quads_eroded_GM(:,:,4,i3)==1));
        if WM_flag==1
        F2w1(:,i5) = tmp(find(mask_quads_eroded_WM(:,:,1,i3)==1));
        F2w2(:,i5) = tmp(find(mask_quads_eroded_WM(:,:,2,i3)==1));
        F2w3(:,i5) = tmp(find(mask_quads_eroded_WM(:,:,3,i3)==1));
        F2w4(:,i5) = tmp(find(mask_quads_eroded_WM(:,:,4,i3)==1));
        end
        clear tmp
    end; clear i5
    emptysize = round(size(cat(1,F2g1,F2g2,F2g3,F2g4),1)/10); emptyweight = max(max(cat(1,F2g1,F2g2,F2g3,F2g4)));
    emptymat = emptyweight .* ones(emptysize,size(F2g1,2));
    F2g = cat(1,F2g1,emptymat,F2g2,emptymat,F2g3,emptymat,F2g4); clear emptysize emptyweight emptymat
    if WM_flag==1
    emptysize = round(size(cat(1,F2w1,F2w2,F2w3,F2w4),1)/10); emptyweight = max(max(cat(1,F2w1,F2w2,F2w3,F2w4)));
    emptymat = emptyweight .* ones(emptysize,size(F2w1,2));
    F2w = cat(1,F2w1,emptymat,F2w2,emptymat,F2w3,emptymat,F2w4); clear emptysize emptyweight emptymat
    end
    
    fighndl = figure('Position',[1 1 1152 720],'Color',[0.95,0.9,0.85],'InvertHardcopy','off','Visible','off'); % red [0.95,0.9,0.85], green [0.85,0.90,0.85], blue [0.89,0.94,0.99]
    subplot2n(2,1,1), imagesc(F2g),colormap(cmap),colorbar, xlabel('time (in samples)'),ylabel('Gray matter voxels (across four quadrants)'),
        title(['Pre-processed GRAY MATTER time series within (top to bottom): left ventral, right ventral, left dorsal and right dorsal horns (slice ' num2str(i3) ' of ' num2str(siz3) ')'],'Color',[0,0.55,0.69],'FontSize',14);
	if WM_flag==1 %#ok<*ALIGN>
    subplot2n(2,1,2), imagesc(F2w),colormap(cmap),colorbar, xlabel('time (in samples)'),ylabel('White matter voxels (across four quadrants)'),
        title(['Pre-processed WHITE MATTER time series within (top to bottom): left ventral, right ventral, left dorsal and right dorsal horns (slice ' num2str(i3) ' of ' num2str(siz3) ')'],'Color',[0.65,0.11,0.19],'FontSize',14);
    end
    saveas(fighndl,[QCpath 'fMRI_ts_20quadTS_slice' sprintf('%.2d',i3) '.jpg'])
    close(fighndl)
    clear F2g F2w
    
    mF2g1 = zscore(mean(F2g1,1)); mF2g2 = zscore(mean(F2g2,1)); mF2g3 = zscore(mean(F2g3,1)); mF2g4 = zscore(mean(F2g4,1));
	if WM_flag==1
    mF2w1 = zscore(mean(F2w1,1)); mF2w2 = zscore(mean(F2w2,1)); mF2w3 = zscore(mean(F2w3,1)); mF2w4 = zscore(mean(F2w4,1));
    end
    
    Cg11 = uppertriangle(corrcoef(F2g1')); Cg22 = uppertriangle(corrcoef(F2g2')); Cg33 = uppertriangle(corrcoef(F2g3')); Cg44 = uppertriangle(corrcoef(F2g4'));
    Cg12=[]; Cg13=[]; Cg14=[]; Cg23=[]; Cg24=[]; Cg34=[];
    f=0;
    for v1=1:size(F2g1,1), for v2=1:size(F2g2,1)
            f=f+1;
            Cg12(f,1) = corr2(F2g1(v1,:)',F2g2(v2,:)');
    end, end; clear v1 v2 f
    f=0;
    for v1=1:size(F2g1,1), for v2=1:size(F2g3,1)
            f=f+1;
            Cg13(f,1) = corr2(F2g1(v1,:)',F2g3(v2,:)');
    end, end; clear v1 v2 f
    f=0;
    for v1=1:size(F2g1,1), for v2=1:size(F2g4,1)
            f=f+1;
            Cg14(f,1) = corr2(F2g1(v1,:)',F2g4(v2,:)');
    end, end; clear v1 v2 f
    f=0;
    for v1=1:size(F2g2,1), for v2=1:size(F2g3,1)
            f=f+1;
            Cg23(f,1) = corr2(F2g2(v1,:)',F2g3(v2,:)');
    end, end; clear v1 v2 f
    f=0;
    for v1=1:size(F2g2,1), for v2=1:size(F2g4,1)
            f=f+1;
            Cg24(f,1) = corr2(F2g2(v1,:)',F2g4(v2,:)');
    end, end; clear v1 v2 f
    f=0;
    for v1=1:size(F2g3,1), for v2=1:size(F2g4,1)
            f=f+1;
            Cg34(f,1) = corr2(F2g3(v1,:)',F2g4(v2,:)');
    end, end; clear v1 v2 f
    Cg = cat(1, Cg11, Cg22, Cg33, Cg44, Cg12, Cg34, Cg13, Cg24, Cg14, Cg23);
    grpCg = cat(1, ones(length(Cg11),1), 2.*ones(length(Cg22),1), 3.*ones(length(Cg33),1), 4.*ones(length(Cg44),1), 5.*ones(length(Cg12),1), 6.*ones(length(Cg34),1), 7.*ones(length(Cg13),1), 8.*ones(length(Cg24),1), 9.*ones(length(Cg14),1), 10.*ones(length(Cg23),1));
    
	if WM_flag==1
    Cw11 = uppertriangle(corrcoef(F2w1')); Cw22 = uppertriangle(corrcoef(F2w2')); Cw33 = uppertriangle(corrcoef(F2w3')); Cw44 = uppertriangle(corrcoef(F2w4'));
    Cw12=[]; Cw13=[]; Cw14=[]; Cw23=[]; Cw24=[]; Cw34=[];
    f=0;
    for v1=1:size(F2w1,1), for v2=1:size(F2w2,1)
            f=f+1;
            Cw12(f,1) = corr2(F2w1(v1,:)',F2w2(v2,:)');
    end, end; clear v1 v2 f
    f=0;
    for v1=1:size(F2w1,1), for v2=1:size(F2w3,1)
            f=f+1;
            Cw13(f,1) = corr2(F2w1(v1,:)',F2w3(v2,:)');
    end, end; clear v1 v2 f
    f=0;
    for v1=1:size(F2w1,1), for v2=1:size(F2w4,1)
            f=f+1;
            Cw14(f,1) = corr2(F2w1(v1,:)',F2w4(v2,:)');
    end, end; clear v1 v2 f
    f=0;
    for v1=1:size(F2w2,1), for v2=1:size(F2w3,1)
            f=f+1;
            Cw23(f,1) = corr2(F2w2(v1,:)',F2w3(v2,:)');
    end, end; clear v1 v2 f
    f=0;
    for v1=1:size(F2w2,1), for v2=1:size(F2w4,1)
            f=f+1;
            Cw24(f,1) = corr2(F2w2(v1,:)',F2w4(v2,:)');
    end, end; clear v1 v2 f
    f=0;
    for v1=1:size(F2w3,1), for v2=1:size(F2w4,1)
            f=f+1;
            Cw34(f,1) = corr2(F2w3(v1,:)',F2w4(v2,:)');
    end, end; clear v1 v2 f
    Cw = cat(1, Cw11, Cw22, Cw33, Cw44, Cw12, Cw34, Cw13, Cw24, Cw14, Cw23);
    grpCw = cat(1, ones(length(Cw11),1), 2.*ones(length(Cw22),1), 3.*ones(length(Cw33),1), 4.*ones(length(Cw44),1), 5.*ones(length(Cw12),1), 6.*ones(length(Cw34),1), 7.*ones(length(Cw13),1), 8.*ones(length(Cw24),1), 9.*ones(length(Cw14),1), 10.*ones(length(Cw23),1));
    end

    fighndl = figure('Position',[1 1 1120 1120],'Color',[0.95,0.9,0.85],'InvertHardcopy','off','Visible','off'); % red [0.95,0.9,0.85], green [0.85,0.90,0.85], blue [0.89,0.94,0.99]
    labelsC = {'LV','RV','LD','RD','LV-RV','LD-RD','LV-LD','RV-RD','LV-RD','LD-RV'};
    try
    subplot2n(2,1,1), b = boxplot(Cg,grpCg,'Notch','on','Labels',labelsC,'ColorGroup',[1,1,1,1,2,2,3,3,3,3]);
        set(gca,'FontSize',15); set(b,'LineWidth', 2); clear b
        grid on, grid minor, ylim([-1 1]), ylabel('Pearson''s correlation coefficient'), hold on, yline(0,'Color','k'); hold off
        title(sprintf('Boxplot of voxel-to-voxel correlations within and between GRAY MATTER quadrants (slice %d of %d)\n(L = left, R = right, V = ventral horn, D = dorsal horn)\n(red: within quadrant | green: between quads V-V or D-D | blue: the rest)\n',i3,siz3),'Color',[0,0.55,0.69],'FontSize',17)
    catch
    end
	if WM_flag==1
    try
    subplot2n(2,1,2), b = boxplot(Cw,grpCw,'Notch','on','Labels',labelsC,'ColorGroup',[1,1,1,1,2,2,3,3,3,3]);
        set(gca,'FontSize',15); set(b,'LineWidth', 2); clear b
        grid on, grid minor, ylim([-1 1]), ylabel('Pearson''s correlation coefficient'), hold on, yline(0,'Color','k'); hold off
        title(sprintf('Boxplot of voxel-to-voxel correlations within and between WHITE MATTER quadrants (slice %d of %d)\n(L = left, R = right, V = ventral horn, D = dorsal horn)\n(red: within quadrant | green: between quads V-V or D-D | blue: the rest)\n',i3,siz3),'Color',[0.65,0.11,0.19],'FontSize',17)
    catch
    end
    end
    saveas(fighndl,[QCpath 'boxplot_FC_compare_20quadTS_slice' sprintf('%.2d',i3) '.jpg'])
    close(fighndl)
    
    fighndl = figure('Position',[1 1 1152 720],'Color',[0.96,0.9,0.84],'InvertHardcopy','off','Visible','off'); % red [0.95,0.9,0.85], green [0.85,0.90,0.85], blue [0.89,0.94,0.99]
    subplot2n(4,1,1), plot(mF2g1,'color',[0,0.65,0.59],'linewidth',1.5), hold on, plot(mF2g2,'color',[0,0.45,0.79],'linewidth',0.15), ylabel('mean time series'),
        try ylim([min(min(1.25.*[mF2g1,mF2g2,mF2g3,mF2g4])) 1.25.*max(max([mF2g1,mF2g2,mF2g3,mF2g4]))]), catch end
        grid on,grid minor, xlim([1 length(mF2g1)]), title(sprintf('GRAY MATTER\n\nMean time series from the LEFT VENTRAL horn (slice %d of %d)',i3,siz3),'Color',[0,0.65,0.59],'FontSize',14), legend({'Left ventral time series'},'FontSize',12,'TextColor',[0,0.65,0.59],'Location','best')
    subplot2n(4,1,2), plot(mF2g2,'color',[0,0.45,0.79],'linewidth',1.5), hold on, plot(mF2g1,'color',[0,0.65,0.59],'linewidth',0.15), ylabel('mean time series'),
        try ylim([min(min(1.25.*[mF2g1,mF2g2,mF2g3,mF2g4])) 1.25.*max(max([mF2g1,mF2g2,mF2g3,mF2g4]))]), catch end
        grid on,grid minor, xlim([1 length(mF2g1)]), title(['Mean time series from the RIGHT VENTRAL horn (slice ' num2str(i3) ' of ' num2str(siz3) ')'],'Color',[0,0.45,0.79],'FontSize',14), legend({'Right ventral time series'},'FontSize',12,'TextColor',[0,0.45,0.79],'Location','best')
    subplot2n(4,1,3), plot(mF2g3,'color',[0.55,0.01,0.39],'linewidth',1.5), hold on, plot(mF2g4,'color',[0.75,0.06,0.09],'linewidth',0.15), ylabel('mean time series'),
        try ylim([min(min(1.25.*[mF2g1,mF2g2,mF2g3,mF2g4])) 1.25.*max(max([mF2g1,mF2g2,mF2g3,mF2g4]))]), catch end
        grid on,grid minor, xlim([1 length(mF2g1)]), title(['Mean time series from the LEFT DORSAL horn (slice ' num2str(i3) ' of ' num2str(siz3) ')'],'Color',[0.55,0.01,0.39],'FontSize',14), legend({'Left dorsal time series'},'FontSize',12,'TextColor',[0.55,0.01,0.39],'Location','best')
    subplot2n(4,1,4), plot(mF2g4,'color',[0.55,0.31,0.09],'linewidth',1.5), hold on, plot(mF2g3,'color',[0.55,0.16,0.29],'linewidth',0.15), xlabel('time (in samples)'), ylabel('mean time series'),
        try ylim([min(min(1.25.*[mF2g1,mF2g2,mF2g3,mF2g4])) 1.25.*max(max([mF2g1,mF2g2,mF2g3,mF2g4]))]), catch end
        grid on,grid minor, xlim([1 length(mF2g1)]), title(['Mean time series from the RIGHT DORSAL horn (slice ' num2str(i3) ' of ' num2str(siz3) ')'],'Color',[0.55,0.31,0.09],'FontSize',14), legend({'Right dorsal time series'},'FontSize',12,'TextColor',[0.55,0.31,0.09],'Location','best')
    saveas(fighndl,[QCpath 'mean_timeseries_GM_20quadTS_slice' sprintf('%.2d',i3) '.jpg'])
    close(fighndl)
	
    if WM_flag==1
    fighndl = figure('Position',[1 1 1152 720],'Color',[0.92,0.9,0.88],'InvertHardcopy','off','Visible','off'); % red [0.95,0.9,0.85], green [0.85,0.90,0.85], blue [0.89,0.94,0.99]
    subplot2n(4,1,1), plot(mF2w1,'color',[0,0.65,0.59],'linewidth',1.5), hold on, plot(mF2w2,'color',[0,0.45,0.79],'linewidth',0.15), ylabel('mean time series'),
        try ylim([min(min(1.25.*[mF2w1,mF2w2,mF2w3,mF2w4])) 1.25.*max(max([mF2w1,mF2w2,mF2w3,mF2w4]))]), catch, end
        grid on,grid minor, xlim([1 length(mF2w1)]), title(sprintf('WHITE MATTER\n\nMean time series from the LEFT VENTRAL horn (slice %d of %d)',i3,siz3),'Color',[0,0.65,0.59],'FontSize',14), legend({'Left ventral time series'},'FontSize',12,'TextColor',[0,0.65,0.59],'Location','best')
    subplot2n(4,1,2), plot(mF2w2,'color',[0,0.45,0.79],'linewidth',1.5), hold on, plot(mF2w1,'color',[0,0.65,0.59],'linewidth',0.15), ylabel('mean time series'),
        try ylim([min(min(1.25.*[mF2w1,mF2w2,mF2w3,mF2w4])) 1.25.*max(max([mF2w1,mF2w2,mF2w3,mF2w4]))]), catch, end
        grid on,grid minor, xlim([1 length(mF2w1)]), title(['Mean time series from the RIGHT VENTRAL horn (slice ' num2str(i3) ' of ' num2str(siz3) ')'],'Color',[0,0.45,0.79],'FontSize',14), legend({'Right ventral time series'},'FontSize',12,'TextColor',[0,0.45,0.79],'Location','best')
    subplot2n(4,1,3), plot(mF2w3,'color',[0.55,0.01,0.39],'linewidth',1.5), hold on, plot(mF2w4,'color',[0.75,0.06,0.09],'linewidth',0.15), ylabel('mean time series'),
        try ylim([min(min(1.25.*[mF2w1,mF2w2,mF2w3,mF2w4])) 1.25.*max(max([mF2w1,mF2w2,mF2w3,mF2w4]))]), catch, end
        grid on,grid minor, xlim([1 length(mF2w1)]), title(['Mean time series from the LEFT DORSAL horn (slice ' num2str(i3) ' of ' num2str(siz3) ')'],'Color',[0.55,0.01,0.39],'FontSize',14), legend({'Left dorsal time series'},'FontSize',12,'TextColor',[0.55,0.01,0.39],'Location','best')
    subplot2n(4,1,4), plot(mF2w4,'color',[0.55,0.31,0.09],'linewidth',1.5), hold on, plot(mF2w3,'color',[0.55,0.16,0.29],'linewidth',0.15), xlabel('time (in samples)'), ylabel('mean time series'),
        try ylim([min(min(1.25.*[mF2w1,mF2w2,mF2w3,mF2w4])) 1.25.*max(max([mF2w1,mF2w2,mF2w3,mF2w4]))]), catch, end
        grid on,grid minor, xlim([1 length(mF2w1)]), title(['Mean time series from the RIGHT DORSAL horn (slice ' num2str(i3) ' of ' num2str(siz3) ')'],'Color',[0.55,0.31,0.09],'FontSize',14), legend({'Right dorsal time series'},'FontSize',12,'TextColor',[0.55,0.31,0.09],'Location','best')
    saveas(fighndl,[QCpath 'mean_timeseries_WM_20quadTS_slice' sprintf('%.2d',i3) '.jpg'])
    close(fighndl)
    end
    
    F2g1_=cat(1,F2g1_,F2g1); F2g2_=cat(1,F2g2_,F2g2); F2g3_=cat(1,F2g3_,F2g3); F2g4_=cat(1,F2g4_,F2g4);
    mF2g1_=cat(1,mF2g1_,mF2g1); mF2g2_=cat(1,mF2g2_,mF2g2); mF2g3_=cat(1,mF2g3_,mF2g3); mF2g4_=cat(1,mF2g4_,mF2g4);    
    Cg11_=cat(1,Cg11_,Cg11); Cg22_=cat(1,Cg22_,Cg22); Cg33_=cat(1,Cg33_,Cg33); Cg44_=cat(1,Cg44_,Cg44); Cg12_=cat(1,Cg12_,Cg12); Cg13_=cat(1,Cg13_,Cg13); Cg14_=cat(1,Cg14_,Cg14); Cg23_=cat(1,Cg23_,Cg23); Cg24_=cat(1,Cg24_,Cg24); Cg34_=cat(1,Cg34_,Cg34);
    if WM_flag==1
    F2w1_=cat(1,F2w1_,F2w1); F2w2_=cat(1,F2w2_,F2w2); F2w3_=cat(1,F2w3_,F2w3); F2w4_=cat(1,F2w4_,F2w4);
    mF2w1_=cat(1,mF2w1_,mF2w1); mF2w2_=cat(1,mF2w2_,mF2w2); mF2w3_=cat(1,mF2w3_,mF2w3); mF2w4_=cat(1,mF2w4_,mF2w4);    
    Cw11_=cat(1,Cw11_,Cw11); Cw22_=cat(1,Cw22_,Cw22); Cw33_=cat(1,Cw33_,Cw33); Cw44_=cat(1,Cw44_,Cw44); Cw12_=cat(1,Cw12_,Cw12); Cw13_=cat(1,Cw13_,Cw13); Cw14_=cat(1,Cw14_,Cw14); Cw23_=cat(1,Cw23_,Cw23); Cw24_=cat(1,Cw24_,Cw24); Cw34_=cat(1,Cw34_,Cw34);
    end
    clear F2w4 F2g F2g1 F2g2 F2g3 F2g4 F2w F2w1 F2w2 F2w3 Cg44 Cg11 Cg12 Cg13 Cg14 Cg22 Cg23 Cg24 Cg33 Cg34 Cg Cw44 Cw Cw11 Cw12 Cw13 Cw14 Cw22 Cw23 Cw24 Cw33 Cw34 mF2w4 mF2g1 mF2g2 mF2g3 mF2g4 mF2w1 mF2w2 mF2w3
end


emptysize = round(size(cat(1,F2g1_,F2g2_,F2g3_,F2g4_),1)/10); emptyweight = max(max(cat(1,F2g1_,F2g2_,F2g3_,F2g4_)));
emptymat = emptyweight .* ones(emptysize,size(F2g1_,2));
F2g_ = cat(1,F2g1_,emptymat,F2g2_,emptymat,F2g3_,emptymat,F2g4_); clear emptysize emptyweight emptymat
if WM_flag==1
    emptysize = round(size(cat(1,F2w1_,F2w2_,F2w3_,F2w4_),1)/10); emptyweight = max(max(cat(1,F2w1_,F2w2_,F2w3_,F2w4_)));
    emptymat = emptyweight .* ones(emptysize,size(F2w1_,2));
    F2w_ = cat(1,F2w1_,emptymat,F2w2_,emptymat,F2w3_,emptymat,F2w4_); clear emptysize emptyweight emptymat
end

fighndl = figure('Position',[1 1 1152 720],'Color',[0.89,0.94,0.99],'InvertHardcopy','off','Visible','off'); % red [0.95,0.9,0.85], green [0.85,0.90,0.85], blue [0.89,0.94,0.99]
subplot2n(2,1,1), imagesc(F2g_),colormap(cmap),colorbar, xlabel('time (in samples)'),ylabel('Gray matter voxels (across four quadrants)'),
    title(['Pre-processed GRAY MATTER time series within (top to bottom): left ventral, right ventral, left dorsal and right dorsal horns (all slices combined)'],'Color',[0,0.55,0.69],'FontSize',14);
if WM_flag==1
subplot2n(2,1,2), imagesc(F2w_),colormap(cmap),colorbar, xlabel('time (in samples)'),ylabel('White matter voxels (across four quadrants)'),
    title(['Pre-processed WHITE MATTER time series within (top to bottom): left ventral, right ventral, left dorsal and right dorsal horns (all slices combined)'],'Color',[0.65,0.11,0.19],'FontSize',14);
end
saveas(fighndl,[QCpath 'fMRI_ts_20quadTS_allSlices' '.jpg'])
saveas(fighndl,[QCpath2 'fMRI_ts_20quadTS_allSlices' '.jpg'])
close(fighndl)
clear F2g_ F2w_

Cg_ = cat(1, Cg11_, Cg22_, Cg33_, Cg44_, Cg12_, Cg34_, Cg13_, Cg24_, Cg14_, Cg23_);
grpCg_ = cat(1, ones(length(Cg11_),1), 2.*ones(length(Cg22_),1), 3.*ones(length(Cg33_),1), 4.*ones(length(Cg44_),1), 5.*ones(length(Cg12_),1), 6.*ones(length(Cg34_),1), 7.*ones(length(Cg13_),1), 8.*ones(length(Cg24_),1), 9.*ones(length(Cg14_),1), 10.*ones(length(Cg23_),1));
Cw_ = cat(1, Cw11_, Cw22_, Cw33_, Cw44_, Cw12_, Cw34_, Cw13_, Cw24_, Cw14_, Cw23_);
grpCw_ = cat(1, ones(length(Cw11_),1), 2.*ones(length(Cw22_),1), 3.*ones(length(Cw33_),1), 4.*ones(length(Cw44_),1), 5.*ones(length(Cw12_),1), 6.*ones(length(Cw34_),1), 7.*ones(length(Cw13_),1), 8.*ones(length(Cw24_),1), 9.*ones(length(Cw14_),1), 10.*ones(length(Cw23_),1));

fighndl = figure('Position',[1 1 1120 1120],'Color',[0.89,0.94,0.99],'InvertHardcopy','off','Visible','off'); % red [0.95,0.9,0.85], green [0.85,0.90,0.85], blue [0.89,0.94,0.99]
labelsC = {'LV','RV','LD','RD','LV-RV','LD-RD','LV-LD','RV-RD','LV-RD','LD-RV'};
subplot2n(2,1,1), b = boxplot(Cg_,grpCg_,'Notch','on','Labels',labelsC,'ColorGroup',[1,1,1,1,2,2,3,3,3,3]);
    set(gca,'FontSize',15); set(b,'LineWidth', 2); clear b
    grid on, grid minor, ylim([-1 1]), ylabel('Pearson''s correlation coefficient'), hold on, yline(0,'Color','k'); hold off
    title(sprintf('Boxplot of voxel-to-voxel correlations within and between GRAY MATTER quadrants (all slices combined)\n(L = left, R = right, V = ventral horn, D = dorsal horn)\n(red: within quadrant | green: between quads V-V or D-D | blue: the rest)\n'),'Color',[0,0.55,0.69],'FontSize',17)
if WM_flag==1
subplot2n(2,1,2), b = boxplot(Cw_,grpCw_,'Notch','on','Labels',labelsC,'ColorGroup',[1,1,1,1,2,2,3,3,3,3]);
    set(gca,'FontSize',15); set(b,'LineWidth', 2); clear b
    grid on, grid minor, ylim([-1 1]), ylabel('Pearson''s correlation coefficient'), hold on, yline(0,'Color','k'); hold off
    title(sprintf('Boxplot of voxel-to-voxel correlations within and between WHITE MATTER quadrants (all slices combined)\n(L = left, R = right, V = ventral horn, D = dorsal horn)\n(red: within quadrant | green: between quads V-V or D-D | blue: the rest)\n'),'Color',[0.65,0.11,0.19],'FontSize',17)
end
saveas(fighndl,[QCpath 'boxplot_FC_compare_20quadTS_allSlices' '.jpg'])
saveas(fighndl,[QCpath2 'FC_compare_boxplot_20quadTS_allSlices' '.jpg'])
close(fighndl)

mF2g1_=mean(mF2g1_,1); mF2g2_=mean(mF2g2_,1); mF2g3_=mean(mF2g3_,1); mF2g4_=mean(mF2g4_,1);
mF2w1_=mean(mF2w1_,1); mF2w2_=mean(mF2w2_,1); mF2w3_=mean(mF2w3_,1); mF2w4_=mean(mF2w4_,1);
fighndl = figure('Position',[1 1 1152 720],'Color',[0.96,0.9,0.80],'InvertHardcopy','off','Visible','off'); % red [0.95,0.9,0.85], green [0.85,0.90,0.85], blue [0.89,0.94,0.99]
subplot2n(4,1,1), plot(mF2g1_,'color',[0,0.65,0.59],'linewidth',1.5), hold on, plot(mF2g2_,'color',[0,0.45,0.79],'linewidth',0.15), ylabel('mean time series'),
    try ylim([min(min(1.25.*[mF2g1_,mF2g2_,mF2g3_,mF2g4_])) 1.25.*max(max([mF2g1_,mF2g2_,mF2g3_,mF2g4_]))]), catch, end
    grid on,grid minor, xlim([1 length(mF2g1_)]), title(sprintf('GRAY MATTER\n\nMean time series from the LEFT VENTRAL horn across all slices'),'Color',[0,0.65,0.59],'FontSize',14), legend({'Left ventral time series'},'FontSize',12,'TextColor',[0,0.65,0.59],'Location','best')
subplot2n(4,1,2), plot(mF2g2_,'color',[0,0.45,0.79],'linewidth',1.5), hold on, plot(mF2g1_,'color',[0,0.65,0.59],'linewidth',0.15), ylabel('mean time series'),
    try ylim([min(min(1.25.*[mF2g1_,mF2g2_,mF2g3_,mF2g4_])) 1.25.*max(max([mF2g1_,mF2g2_,mF2g3_,mF2g4_]))]), catch, end
    grid on,grid minor, xlim([1 length(mF2g1_)]), title(['Mean time series from the RIGHT VENTRAL horn across all slices'],'Color',[0,0.45,0.79],'FontSize',14), legend({'Right ventral time series'},'FontSize',12,'TextColor',[0,0.45,0.79],'Location','best')
subplot2n(4,1,3), plot(mF2g3_,'color',[0.55,0.01,0.39],'linewidth',1.5), hold on, plot(mF2g4_,'color',[0.75,0.06,0.09],'linewidth',0.15), ylabel('mean time series'),
    try ylim([min(min(1.25.*[mF2g1_,mF2g2_,mF2g3_,mF2g4_])) 1.25.*max(max([mF2g1_,mF2g2_,mF2g3_,mF2g4_]))]), catch, end
    grid on,grid minor, xlim([1 length(mF2g1_)]), title(['Mean time series from the LEFT DORSAL horn across all slices'],'Color',[0.55,0.01,0.39],'FontSize',14), legend({'Left dorsal time series'},'FontSize',12,'TextColor',[0.55,0.01,0.39],'Location','best')
subplot2n(4,1,4), plot(mF2g4_,'color',[0.55,0.31,0.09],'linewidth',1.5), hold on, plot(mF2g3_,'color',[0.55,0.16,0.29],'linewidth',0.15), xlabel('time (in samples)'), ylabel('mean time series'),
    try ylim([min(min(1.25.*[mF2g1_,mF2g2_,mF2g3_,mF2g4_])) 1.25.*max(max([mF2g1_,mF2g2_,mF2g3_,mF2g4_]))]), catch, end
    grid on,grid minor, xlim([1 length(mF2g1_)]), title(['Mean time series from the RIGHT DORSAL horn across all slices'],'Color',[0.55,0.31,0.09],'FontSize',14), legend({'Right dorsal time series'},'FontSize',12,'TextColor',[0.55,0.31,0.09],'Location','best')
saveas(fighndl,[QCpath 'mean_timeseries_GM_20quadTS_allSlices' '.jpg'])
saveas(fighndl,[QCpath2 'global_signal_20quadTS_allSlices_GM' '.jpg'])
close(fighndl)

if WM_flag==1
fighndl = figure('Position',[1 1 1152 720],'Color',[0.91,0.91,0.87],'InvertHardcopy','off','Visible','off'); % red [0.95,0.9,0.85], green [0.85,0.90,0.85], blue [0.89,0.94,0.99]
subplot2n(4,1,1), plot(mF2w1_,'color',[0,0.65,0.59],'linewidth',1.5), hold on, plot(mF2w2_,'color',[0,0.45,0.79],'linewidth',0.15), ylabel('mean time series'),
    try ylim([min(min(1.25.*[mF2w1_,mF2w2_,mF2w3_,mF2w4_])) 1.25.*max(max([mF2w1_,mF2w2_,mF2w3_,mF2w4_]))]), catch, end
    grid on,grid minor, xlim([1 length(mF2w1_)]), title(sprintf('WHITE MATTER\n\nMean time series from the LEFT VENTRAL horn across all slices'),'Color',[0,0.65,0.59],'FontSize',14), legend({'Left ventral time series'},'FontSize',12,'TextColor',[0,0.65,0.59],'Location','best')
subplot2n(4,1,2), plot(mF2w2_,'color',[0,0.45,0.79],'linewidth',1.5), hold on, plot(mF2w1_,'color',[0,0.65,0.59],'linewidth',0.15), ylabel('mean time series'),
    try ylim([min(min(1.25.*[mF2w1_,mF2w2_,mF2w3_,mF2w4_])) 1.25.*max(max([mF2w1_,mF2w2_,mF2w3_,mF2w4_]))]), catch, end
    grid on,grid minor, xlim([1 length(mF2w1_)]), title(['Mean time series from the RIGHT VENTRAL horn across all slices'],'Color',[0,0.45,0.79],'FontSize',14), legend({'Right ventral time series'},'FontSize',12,'TextColor',[0,0.45,0.79],'Location','best')
subplot2n(4,1,3), plot(mF2w3_,'color',[0.55,0.01,0.39],'linewidth',1.5), hold on, plot(mF2w4_,'color',[0.75,0.06,0.09],'linewidth',0.15), ylabel('mean time series'),
    try ylim([min(min(1.25.*[mF2w1_,mF2w2_,mF2w3_,mF2w4_])) 1.25.*max(max([mF2w1_,mF2w2_,mF2w3_,mF2w4_]))]), catch, end
    grid on,grid minor, xlim([1 length(mF2w1_)]), title(['Mean time series from the LEFT DORSAL horn across all slices'],'Color',[0.55,0.01,0.39],'FontSize',14), legend({'Left dorsal time series'},'FontSize',12,'TextColor',[0.55,0.01,0.39],'Location','best')
subplot2n(4,1,4), plot(mF2w4_,'color',[0.55,0.31,0.09],'linewidth',1.5), hold on, plot(mF2w3_,'color',[0.55,0.16,0.29],'linewidth',0.15), xlabel('time (in samples)'), ylabel('mean time series'),
    try ylim([min(min(1.25.*[mF2w1_,mF2w2_,mF2w3_,mF2w4_])) 1.25.*max(max([mF2w1_,mF2w2_,mF2w3_,mF2w4_]))]), catch, end
    grid on,grid minor, xlim([1 length(mF2w1_)]), title(['Mean time series from the RIGHT DORSAL horn across all slices'],'Color',[0.55,0.31,0.09],'FontSize',14), legend({'Right dorsal time series'},'FontSize',12,'TextColor',[0.55,0.31,0.09],'Location','best')
saveas(fighndl,[QCpath 'mean_timeseries_WM_20quadTS_allSlices' '.jpg'])
saveas(fighndl,[QCpath2 'global_signal_20quadTS_allSlices_WM' '.jpg'])
close(fighndl)
end

clear F2g4_ F2g1_ F2g2_ F2g3_ Cg44_ Cg11_ Cg12_ Cg13_ Cg14_ Cg22_ Cg23_ Cg24_ Cg33_ Cg34_ mF2g4_ mF2g1_ mF2g2_ mF2g3_ F2w4_ F2w1_ F2w2_ F2w3_ Cw44_ Cw11_ Cw12_ Cw13_ Cw14_ Cw22_ Cw23_ Cw24_ Cw33_ Cw34_ mF2w4_ mF2w1_ mF2w2_ mF2w3_ cmap

end
```
