## Supplementary material for "Neptune: a toolbox for spinal cord functional MRI data processing and quality assurance": Neptune software: scfMRItb_21_SFC_betweenslice.m

```
function scfMRItb_21_SFC_betweenslice(base_dir_sub,fname4,naming, SFC_type,WM_flag,assign_value,Rns, dont_do_pcor_betweenslice)

if nargin < 6
    assign_value = 1; % computing FC between all pairs of voxels in the quadrants and assign the 95th percentile value as the final FC value. If this flag is set to 1, then mean time series is computed for each quadrant instead and FC computed between them
    if nargin < 5
        WM_flag = 1; % compute SFC in the white matter (0 = don't)
        if nargin < 4
            SFC_type = 1; % compute only Pearson's correlation as a measure of FC (2 = compute partial correlation only; 3 = compute both separately)
        end
    end
end

scrsz = get(0,'ScreenSize');

if ~(exist([base_dir_sub 'QC' '/21_static_functional_connectivity'],'dir'))
    mkdir([base_dir_sub 'QC' '/21_static_functional_connectivity'])
end
if Rns==0
    QCpath = [base_dir_sub 'QC' '/21_static_functional_connectivity/'];
else
    if ~(exist([base_dir_sub 'QC' '/21_static_functional_connectivity' '/run' num2str(Rns)],'dir'))
        mkdir([base_dir_sub 'QC' '/21_static_functional_connectivity' '/run' num2str(Rns)])
    end
    QCpath = [base_dir_sub 'QC' '/21_static_functional_connectivity' '/run' num2str(Rns) '/'];
end

if ~(exist([base_dir_sub 'QC' '/QC_shortlist'],'dir'))
    mkdir([base_dir_sub 'QC' '/QC_shortlist'])
end
if Rns==0
    QCpath2 = [base_dir_sub 'QC' '/QC_shortlist/'];
else
    if ~(exist([base_dir_sub 'QC' '/QC_shortlist' '/run' num2str(Rns)],'dir'))
        mkdir([base_dir_sub 'QC' '/QC_shortlist' '/run' num2str(Rns)])
    end
    QCpath2 = [base_dir_sub 'QC' '/QC_shortlist' '/run' num2str(Rns) '/'];
end

%% ----- COMPUTE FULL AND/OR PARTIAL CORRELATION-BASED STATIC FUNCTIONAL CONNECTIVITY (SFC) -----

load([base_dir_sub fname4 '.mat'], 'timeseries_GM')
[L1,L2] = size(timeseries_GM);
if WM_flag==1
    load([base_dir_sub fname4 '.mat'], 'timeseries_WM')
end

if SFC_type == 1 % do only Pearson's correlation
    pearson = 1; partial = 0;
elseif SFC_type == 2 % do only partial correlation
%     pearson = 0; partial = 1;
    pearson = 0;
    if dont_do_pcor_betweenslice==1
        partial = 0;
    else
        partial = 1;
    end
elseif SFC_type == 3 % do both Pearson's correlation and partial correlation separately
%     pearson = 1; partial = 1;
    pearson = 1;
    if dont_do_pcor_betweenslice==1
        partial = 0;
    else
        partial = 1;
    end
end

pcor_covariates_type = 1; % 1 = partial correlation using covariates only from the slice(s) involved; 2 = partial correlation using covariates from all the slices

dof_correction = 1; % correct for degrees of freedom during z-value calculation (0 = do not correct)

if assign_value==1
    JLTM_cutoff = 0.95; % perform SFC between all pairs of voxels in the two quadrants and assign the 95th percentile value as the final SFC
elseif assign_value==2
    JLTM_cutoff = 0.5; % perform SFC between all pairs of voxels in the two quadrants and assign the 50th percentile value as the final SFC    
end

%% Pearson's correlation based SFC
if pearson ==1
    
SFC_GM_Zval = zeros(L2,L2,L1,L1); SFC_GM_Rval = zeros(L2,L2,L1,L1); SFC_GM_timeseries = cell(L2,L2,L1,L1);
SFC_GM_Zval_mat = zeros(L1*L2,L1*L2); SFC_GM_Rval_mat=SFC_GM_Zval_mat;
for i3 = 1:L1
    fprintf('SFC GM (within+between slices): slice (%d) of (%d) -> Subject (%s)\n',i3,L1,naming)
    for k1 = 1:L2
        temp1 = timeseries_GM{i3,k1};
        if assign_value == 3 % if this flag is 1 then use mean time series from each quadrant instead of computing SFC between all voxels and choosing the 95th percentile value
            temp1 = mean(temp1,1);
        end
        for i4 = 1:L1
%             fprintf('SFC GM: slice-1 (%d) of (%d), slice-2 (%d) of (%d), quadrant (%d) of (%d) -> Subject (%s)\n',i3,L1,i4,L1,k1,L2,naming)
            for k2 = 1:L2
                if ((i3==i4) && (k1<k2)) || ((i3<i4) && (k1<=k2))
                    temp2 = timeseries_GM{i4,k2};
                    if assign_value == 3 % if this flag is 1 then use mean time series from each quadrant instead of computing SFC between all voxels and choosing the 95th percentile value
                        temp2 = mean(temp2,1);
                    end
                    try %#ok<*ALIGN>
                    [z, ts1, ts2, r] = xcorr_zscore_vec(temp1, temp2, dof_correction, size(temp1,2), JLTM_cutoff);
                    SFC_GM_Zval(k1,k2,i3,i4) = z; SFC_GM_Zval(k2,k1,i3,i4) = z; SFC_GM_Zval(k1,k2,i4,i3) = z; SFC_GM_Zval(k2,k1,i4,i3) = z;
                    SFC_GM_Rval(k1,k2,i3,i4) = r; SFC_GM_Rval(k2,k1,i3,i4) = r; SFC_GM_Rval(k1,k2,i4,i3) = r; SFC_GM_Rval(k2,k1,i4,i3) = r;
                    SFC_GM_timeseries{k1,k2,i3,i4}(:,1) = ts1; SFC_GM_timeseries{k1,k2,i3,i4}(:,2) = ts2;
                    SFC_GM_Zval_mat((i3-1)*L2+k1,(i4-1)*L2+k2) = z; SFC_GM_Zval_mat((i4-1)*L2+k2,(i3-1)*L2+k1) = z; SFC_GM_Zval_mat((i3-1)*L2+k2,(i4-1)*L2+k1) = z; SFC_GM_Zval_mat((i4-1)*L2+k1,(i3-1)*L2+k2) = z;
                    SFC_GM_Rval_mat((i3-1)*L2+k1,(i4-1)*L2+k2) = r; SFC_GM_Rval_mat((i4-1)*L2+k2,(i3-1)*L2+k1) = r; SFC_GM_Rval_mat((i3-1)*L2+k2,(i4-1)*L2+k1) = r; SFC_GM_Rval_mat((i4-1)*L2+k1,(i3-1)*L2+k2) = r;
                    catch
                    SFC_GM_Zval(k1,k2,i3,i4) = NaN; SFC_GM_Zval(k2,k1,i3,i4) = NaN; SFC_GM_Zval(k1,k2,i4,i3) = NaN; SFC_GM_Zval(k2,k1,i4,i3) = NaN;
                    SFC_GM_Rval(k1,k2,i3,i4) = NaN; SFC_GM_Rval(k2,k1,i3,i4) = NaN; SFC_GM_Rval(k1,k2,i4,i3) = NaN; SFC_GM_Rval(k2,k1,i4,i3) = NaN;
                    SFC_GM_timeseries{k1,k2,i3,i4}(:,1) = NaN; SFC_GM_timeseries{k1,k2,i3,i4}(:,2) = NaN;
                    SFC_GM_Zval_mat((i3-1)*L2+k1,(i4-1)*L2+k2) = NaN; SFC_GM_Zval_mat((i4-1)*L2+k2,(i3-1)*L2+k1) = NaN; SFC_GM_Zval_mat((i3-1)*L2+k2,(i4-1)*L2+k1) = NaN; SFC_GM_Zval_mat((i4-1)*L2+k1,(i3-1)*L2+k2) = NaN;
                    SFC_GM_Rval_mat((i3-1)*L2+k1,(i4-1)*L2+k2) = NaN; SFC_GM_Rval_mat((i4-1)*L2+k2,(i3-1)*L2+k1) = NaN; SFC_GM_Rval_mat((i3-1)*L2+k2,(i4-1)*L2+k1) = NaN; SFC_GM_Rval_mat((i4-1)*L2+k1,(i3-1)*L2+k2) = NaN;
                    end
                    clear temp2 z r ts1 ts2
                end
            end
        end
        SFC_GM_Rval(k1,k1,i3,i3) = 1;
        clear temp1
    end
end

if WM_flag == 1 % compute SFC in the white matter (0 = don't)
SFC_WM_Zval = zeros(L2,L2,L1); SFC_WM_Rval = zeros(L2,L2,L1); SFC_WM_timeseries = cell(L2,L2,L1);
SFC_WM_Zval_mat = zeros(L1*L2,L1*L2); SFC_WM_Rval_mat=SFC_GM_Zval_mat;
for i3 = 1:L1
    fprintf('SFC WM (within+between slices): slice (%d) of (%d) -> Subject (%s)\n',i3,L1,naming)
    for k1 = 1:L2
        temp1 = timeseries_WM{i3,k1};
        if assign_value == 3 % if this flag is 1 then use mean time series from each quadrant instead of computing SFC between all voxels and choosing the 95th percentile value
            temp1 = mean(temp1,1);
        end
        for i4 = 1:L1
%             fprintf('SFC WM: slice-1 (%d) of (%d), slice-2 (%d) of (%d), quadrant (%d) of (%d) -> Subject (%s)\n',i3,L1,i4,L1,k1,L2,naming)
            for k2 = 1:L2
                if ((i3==i4) && (k1<k2)) || ((i3<i4) && (k1<=k2))
                    temp2 = timeseries_WM{i4,k2};
                    if assign_value == 3 % if this flag is 1 then use mean time series from each quadrant instead of computing SFC between all voxels and choosing the 95th percentile value
                        temp2 = mean(temp2,1);
                    end
                    try
                    [z, ts1, ts2, r] = xcorr_zscore_vec(temp1, temp2, dof_correction, size(temp1,2), JLTM_cutoff);
                    SFC_WM_Zval(k1,k2,i3,i4) = z; SFC_WM_Zval(k2,k1,i3,i4) = z; SFC_WM_Zval(k1,k2,i4,i3) = z; SFC_WM_Zval(k2,k1,i4,i3) = z;
                    SFC_WM_Rval(k1,k2,i3,i4) = r; SFC_WM_Rval(k2,k1,i3,i4) = r; SFC_WM_Rval(k1,k2,i4,i3) = r; SFC_WM_Rval(k2,k1,i4,i3) = r;
                    SFC_WM_timeseries{k1,k2,i3,i4}(:,1) = ts1; SFC_WM_timeseries{k1,k2,i3,i4}(:,2) = ts2;
                    SFC_WM_Zval_mat((i3-1)*L2+k1,(i4-1)*L2+k2) = z; SFC_WM_Zval_mat((i4-1)*L2+k2,(i3-1)*L2+k1) = z; SFC_WM_Zval_mat((i3-1)*L2+k2,(i4-1)*L2+k1) = z; SFC_WM_Zval_mat((i4-1)*L2+k1,(i3-1)*L2+k2) = z;
                    SFC_WM_Rval_mat((i3-1)*L2+k1,(i4-1)*L2+k2) = r; SFC_WM_Rval_mat((i4-1)*L2+k2,(i3-1)*L2+k1) = r; SFC_WM_Rval_mat((i3-1)*L2+k2,(i4-1)*L2+k1) = r; SFC_WM_Rval_mat((i4-1)*L2+k1,(i3-1)*L2+k2) = r;
                    catch
                    SFC_WM_Zval(k1,k2,i3,i4) = NaN; SFC_WM_Zval(k2,k1,i3,i4) = NaN; SFC_WM_Zval(k1,k2,i4,i3) = NaN; SFC_WM_Zval(k2,k1,i4,i3) = NaN;
                    SFC_WM_Rval(k1,k2,i3,i4) = NaN; SFC_WM_Rval(k2,k1,i3,i4) = NaN; SFC_WM_Rval(k1,k2,i4,i3) = NaN; SFC_WM_Rval(k2,k1,i4,i3) = NaN;
                    SFC_WM_timeseries{k1,k2,i3,i4}(:,1) = NaN; SFC_WM_timeseries{k1,k2,i3,i4}(:,2) = NaN;
                    SFC_WM_Zval_mat((i3-1)*L2+k1,(i4-1)*L2+k2) = NaN; SFC_WM_Zval_mat((i4-1)*L2+k2,(i3-1)*L2+k1) = NaN; SFC_WM_Zval_mat((i3-1)*L2+k2,(i4-1)*L2+k1) = NaN; SFC_WM_Zval_mat((i4-1)*L2+k1,(i3-1)*L2+k2) = NaN;
                    SFC_WM_Rval_mat((i3-1)*L2+k1,(i4-1)*L2+k2) = NaN; SFC_WM_Rval_mat((i4-1)*L2+k2,(i3-1)*L2+k1) = NaN; SFC_WM_Rval_mat((i3-1)*L2+k2,(i4-1)*L2+k1) = NaN; SFC_WM_Rval_mat((i4-1)*L2+k1,(i3-1)*L2+k2) = NaN;
                    end
                    clear temp2 z r ts1 ts2
                end
            end
        end
        SFC_WM_Rval(k1,k1,i3,i3) = 1;
        clear temp1
    end
end
end

% save
info_SFC = sprintf('\nZval = Z-value obtained after Fischers R-to-Z transformation of Pearsons correlation values; R-val = correlation (R) value\nDimensions: 4 quadrants x 4 quadrants x slices x slices\nIn each slice pair, the 4x4 matrix represents the SFC value between corresponding quadrants:\nQuadrants: 1 = left ventral, 2 = right ventral, 3 = left dorsal, 4 = right dorsal\nExample: the element (1,3,6,9) represents the SFC between (1)left ventral quadrant in slice-6 and (3)left dorsal quadrant in slice-9\nNote that only upper triangle data is available for some variables as FC is non-directional and symmetric\n\n');
if WM_flag == 1
save([base_dir_sub naming '_SFC_betweenslice' '.mat'], 'SFC_GM_Zval', 'SFC_GM_Rval', 'SFC_GM_timeseries', 'SFC_WM_Zval', 'SFC_WM_Rval', 'SFC_WM_timeseries', 'SFC_GM_Zval_mat', 'SFC_GM_Rval_mat', 'SFC_WM_Zval_mat', 'SFC_WM_Rval_mat', 'info_SFC')
else
save([base_dir_sub naming '_SFC_betweenslice' '.mat'], 'SFC_GM_Zval', 'SFC_GM_Rval', 'SFC_GM_timeseries', 'SFC_GM_Zval_mat', 'SFC_GM_Rval_mat', 'info_SFC')
end

end % Pearson's correlation

%% Partial correlation based SFC
if partial == 1
warning off

if pcor_covariates_type == 2
    covariates_ts_GM = zeros(L1,L2,size(timeseries_GM{1,1},2));
    for i3 = 1:L1
        for k1 = 1:L2
            covariates_ts_GM(i3,k1,:) = mean(timeseries_GM{i3,k1},1);
        end
    end
end

SFC_pcor_GM_Zval = zeros(L2,L2,L1); SFC_pcor_GM_Rval = zeros(L2,L2,L1); SFC_pcor_GM_timeseries = cell(L2,L2,L1);
SFC_pcor_GM_Zval_mat = zeros(L1*L2,L1*L2); SFC_pcor_GM_Rval_mat=SFC_pcor_GM_Zval_mat;
for i3 = 1:L1
    for k1 = 1:L2
        fprintf('SFC GM (within+between slices partial corr): slice-1 (%d) of (%d), quadrant (%d) of (%d) -> Subject (%s)\n',i3,L1,k1,L2,naming)
        temp1 = timeseries_GM{i3,k1};
        if assign_value == 3 % if this flag is 1 then use mean time series from each quadrant instead of computing SFC between all voxels and choosing the 95th percentile value
            temp1 = mean(temp1,1);
        end
        for i4 = 1:L1
%             fprintf('SFC GM (partial corr): slice-1 (%d) of (%d), slice-2 (%d) of (%d), quadrant (%d) of (%d) -> Subject (%s)\n',i3,L1,i4,L1,k1,L2,naming)
            for k2 = 1:L2
                if ((i3==i4) && (k1<k2)) || ((i3<i4) && (k1<=k2))
                    temp2 = timeseries_GM{i4,k2};
                    if assign_value == 3 % if this flag is 1 then use mean time series from each quadrant instead of computing SFC between all voxels and choosing the 95th percentile value
                        temp2 = mean(temp2,1);
                    end
                    
                    if pcor_covariates_type == 1 % partial correlation using covariates only from the slice(s) involved
                        if i3==i4
                            ind = find(([1;2;3;4]~=k1)&([1;2;3;4]~=k2));
                            temp3 = mean(timeseries_GM{i3,ind(1)},1);
                            temp4 = mean(timeseries_GM{i3,ind(2)},1);
                            junk = [temp3; temp4];
                            clear ind temp3 temp4
                        else
                            ind1 = find([1;2;3;4]~=k1); ind2 = find([1;2;3;4]~=k2);
                            temp31 = mean(timeseries_GM{i3,ind1(1)},1); temp32 = mean(timeseries_GM{i3,ind1(2)},1); temp33 = mean(timeseries_GM{i3,ind1(3)},1);
                            temp41 = mean(timeseries_GM{i4,ind2(1)},1); temp42 = mean(timeseries_GM{i4,ind2(2)},1); temp43 = mean(timeseries_GM{i4,ind2(3)},1);
                            junk = [temp31; temp32; temp33; temp41; temp42; temp43];
                            clear ind1 ind2 temp31 temp32 temp33 temp41 temp42 temp43
                        end
                    elseif pcor_covariates_type == 2 % partial correlation using covariates from all the slices
                        if i3==i4
                            ind = find(([1;2;3;4]~=k1)&([1;2;3;4]~=k2));
                            temp3 = mean(timeseries_GM{i3,ind(1)},1);
                            temp4 = mean(timeseries_GM{i3,ind(2)},1);
                            junk = [temp3; temp4];
                            temp5 = covariates_ts_GM; temp5(i3,:,:) = [];
                            temp5 = reshape(temp5,size(temp5,1)*size(temp5,2),size(temp5,3));
                            junk = cat(1,junk,temp5);
                            clear ind temp3 temp4 temp5
                        else
                            ind1 = find([1;2;3;4]~=k1); ind2 = find([1;2;3;4]~=k2);
                            temp31 = mean(timeseries_GM{i3,ind1(1)},1); temp32 = mean(timeseries_GM{i3,ind1(2)},1); temp33 = mean(timeseries_GM{i3,ind1(3)},1);
                            temp41 = mean(timeseries_GM{i4,ind2(1)},1); temp42 = mean(timeseries_GM{i4,ind2(2)},1); temp43 = mean(timeseries_GM{i4,ind2(3)},1);
                            junk = [temp31; temp32; temp33; temp41; temp42; temp43];
                            temp5 = covariates_ts_GM; temp5([i3,i4],:,:) = [];
                            temp5 = reshape(temp5,size(temp5,1)*size(temp5,2),size(temp5,3));
                            junk = cat(1,junk,temp5);
                            clear ind1 ind2 temp31 temp32 temp33 temp41 temp42 temp43 temp5                        
                        end
                    end
                    
                    try
                    [z, ts1, ts2, r] = pcorr_zscore_vec(temp1, temp2, dof_correction, junk, JLTM_cutoff);
                    SFC_pcor_GM_Zval(k1,k2,i3,i4) = z; SFC_pcor_GM_Zval(k2,k1,i3,i4) = z; SFC_pcor_GM_Zval(k1,k2,i4,i3) = z; SFC_pcor_GM_Zval(k2,k1,i4,i3) = z;
                    SFC_pcor_GM_Rval(k1,k2,i3,i4) = r; SFC_pcor_GM_Rval(k2,k1,i3,i4) = r; SFC_pcor_GM_Rval(k1,k2,i4,i3) = r; SFC_pcor_GM_Rval(k2,k1,i4,i3) = r;
                    SFC_pcor_GM_timeseries{k1,k2,i3,i4}(:,1) = ts1; SFC_pcor_GM_timeseries{k1,k2,i3,i4}(:,2) = ts2;
                    SFC_pcor_GM_Zval_mat((i3-1)*L2+k1,(i4-1)*L2+k2) = z; SFC_pcor_GM_Zval_mat((i4-1)*L2+k2,(i3-1)*L2+k1) = z; SFC_pcor_GM_Zval_mat((i3-1)*L2+k2,(i4-1)*L2+k1) = z; SFC_pcor_GM_Zval_mat((i4-1)*L2+k1,(i3-1)*L2+k2) = z;
                    SFC_pcor_GM_Rval_mat((i3-1)*L2+k1,(i4-1)*L2+k2) = r; SFC_pcor_GM_Rval_mat((i4-1)*L2+k2,(i3-1)*L2+k1) = r; SFC_pcor_GM_Rval_mat((i3-1)*L2+k2,(i4-1)*L2+k1) = r; SFC_pcor_GM_Rval_mat((i4-1)*L2+k1,(i3-1)*L2+k2) = r;
                    catch
                    SFC_pcor_GM_Zval(k1,k2,i3,i4) = NaN; SFC_pcor_GM_Zval(k2,k1,i3,i4) = NaN; SFC_pcor_GM_Zval(k1,k2,i4,i3) = NaN; SFC_pcor_GM_Zval(k2,k1,i4,i3) = NaN;
                    SFC_pcor_GM_Rval(k1,k2,i3,i4) = NaN; SFC_pcor_GM_Rval(k2,k1,i3,i4) = NaN; SFC_pcor_GM_Rval(k1,k2,i4,i3) = NaN; SFC_pcor_GM_Rval(k2,k1,i4,i3) = NaN;
                    SFC_pcor_GM_timeseries{k1,k2,i3,i4}(:,1) = NaN; SFC_pcor_GM_timeseries{k1,k2,i3,i4}(:,2) = NaN;
                    SFC_pcor_GM_Zval_mat((i3-1)*L2+k1,(i4-1)*L2+k2) = NaN; SFC_pcor_GM_Zval_mat((i4-1)*L2+k2,(i3-1)*L2+k1) = NaN; SFC_pcor_GM_Zval_mat((i3-1)*L2+k2,(i4-1)*L2+k1) = NaN; SFC_pcor_GM_Zval_mat((i4-1)*L2+k1,(i3-1)*L2+k2) = NaN;
                    SFC_pcor_GM_Rval_mat((i3-1)*L2+k1,(i4-1)*L2+k2) = NaN; SFC_pcor_GM_Rval_mat((i4-1)*L2+k2,(i3-1)*L2+k1) = NaN; SFC_pcor_GM_Rval_mat((i3-1)*L2+k2,(i4-1)*L2+k1) = NaN; SFC_pcor_GM_Rval_mat((i4-1)*L2+k1,(i3-1)*L2+k2) = NaN;
                    end
                    clear temp2 temp3 temp4 ind junk z r ts1 ts2
                end
            end
        end
        SFC_pcor_GM_Rval(k1,k1,i3,i3) = 1;
        clear temp1
    end
end

if WM_flag == 1 % compute SFC in the white matter (0 = don't)
if pcor_covariates_type == 2
    covariates_ts_WM = zeros(L1,L2,size(timeseries_WM{1,1},2));
    for i3 = 1:L1
        for k1 = 1:L2
            covariates_ts_WM(i3,k1,:) = mean(timeseries_WM{i3,k1},1);
        end
    end
end
SFC_pcor_WM_Zval = zeros(L2,L2,L1); SFC_pcor_WM_Rval = zeros(L2,L2,L1); SFC_pcor_WM_timeseries = cell(L2,L2,L1);
SFC_pcor_WM_Zval_mat = zeros(L1*L2,L1*L2); SFC_pcor_WM_Rval_mat=SFC_pcor_WM_Zval_mat;
for i3 = 1:L1
    for k1 = 1:L2
        fprintf('SFC WM (within+between slices partial corr): slice-1 (%d) of (%d), quadrant (%d) of (%d) -> Subject (%s)\n',i3,L1,k1,L2,naming)
        temp1 = timeseries_WM{i3,k1};
        if assign_value == 3 % if this flag is 1 then use mean time series from each quadrant instead of computing SFC between all voxels and choosing the 95th percentile value
            temp1 = mean(temp1,1);
        end
        for i4 = 1:L1
%             fprintf('SFC WM (partial corr): slice-1 (%d) of (%d), slice-2 (%d) of (%d), quadrant (%d) of (%d) -> Subject (%s)\n',i3,L1,i4,L1,k1,L2,naming)
            for k2 = 1:L2
                if ((i3==i4) && (k1<k2)) || ((i3<i4) && (k1<=k2))
                    temp2 = timeseries_WM{i4,k2};
                    if assign_value == 3 % if this flag is 1 then use mean time series from each quadrant instead of computing SFC between all voxels and choosing the 95th percentile value
                        temp2 = mean(temp2,1);
                    end

                    if pcor_covariates_type == 1 % partial correlation using covariates only from the slice(s) involved
                        if i3==i4
                            ind = find(([1;2;3;4]~=k1)&([1;2;3;4]~=k2));
                            temp3 = mean(timeseries_WM{i3,ind(1)},1);
                            temp4 = mean(timeseries_WM{i3,ind(2)},1);
                            junk = [temp3; temp4];
                            clear ind temp3 temp4
                        else
                            ind1 = find([1;2;3;4]~=k1); ind2 = find([1;2;3;4]~=k2);
                            temp31 = mean(timeseries_WM{i3,ind1(1)},1); temp32 = mean(timeseries_WM{i3,ind1(2)},1); temp33 = mean(timeseries_WM{i3,ind1(3)},1);
                            temp41 = mean(timeseries_WM{i4,ind2(1)},1); temp42 = mean(timeseries_WM{i4,ind2(2)},1); temp43 = mean(timeseries_WM{i4,ind2(3)},1);
                            junk = [temp31; temp32; temp33; temp41; temp42; temp43];
                            clear ind1 ind2 temp31 temp32 temp33 temp41 temp42 temp43
                        end
                    elseif pcor_covariates_type == 2 % partial correlation using covariates from all the slices
                        if i3==i4
                            ind = find(([1;2;3;4]~=k1)&([1;2;3;4]~=k2));
                            temp3 = mean(timeseries_WM{i3,ind(1)},1);
                            temp4 = mean(timeseries_WM{i3,ind(2)},1);
                            junk = [temp3; temp4];
                            temp5 = covariates_ts_WM; temp5(i3,:,:) = [];
                            temp5 = reshape(temp5,size(temp5,1)*size(temp5,2),size(temp5,3));
                            junk = cat(1,junk,temp5);
                            clear ind temp3 temp4 temp5
                        else
                            ind1 = find([1;2;3;4]~=k1); ind2 = find([1;2;3;4]~=k2);
                            temp31 = mean(timeseries_WM{i3,ind1(1)},1); temp32 = mean(timeseries_WM{i3,ind1(2)},1); temp33 = mean(timeseries_WM{i3,ind1(3)},1);
                            temp41 = mean(timeseries_WM{i4,ind2(1)},1); temp42 = mean(timeseries_WM{i4,ind2(2)},1); temp43 = mean(timeseries_WM{i4,ind2(3)},1);
                            junk = [temp31; temp32; temp33; temp41; temp42; temp43];
                            temp5 = covariates_ts_WM; temp5([i3,i4],:,:) = [];
                            temp5 = reshape(temp5,size(temp5,1)*size(temp5,2),size(temp5,3));
                            junk = cat(1,junk,temp5);
                            clear ind1 ind2 temp31 temp32 temp33 temp41 temp42 temp43 temp5                        
                        end
                    end
                    
                    try
                    [z, ts1, ts2, r] = pcorr_zscore_vec(temp1, temp2, dof_correction, junk, JLTM_cutoff);
                    SFC_pcor_WM_Zval(k1,k2,i3,i4) = z; SFC_pcor_WM_Zval(k2,k1,i3,i4) = z; SFC_pcor_WM_Zval(k1,k2,i4,i3) = z; SFC_pcor_WM_Zval(k2,k1,i4,i3) = z;
                    SFC_pcor_WM_Rval(k1,k2,i3,i4) = r; SFC_pcor_WM_Rval(k2,k1,i3,i4) = r; SFC_pcor_WM_Rval(k1,k2,i4,i3) = r; SFC_pcor_WM_Rval(k2,k1,i4,i3) = r;
                    SFC_pcor_WM_timeseries{k1,k2,i3,i4}(:,1) = ts1; SFC_pcor_WM_timeseries{k1,k2,i3,i4}(:,2) = ts2;
                    SFC_pcor_WM_Zval_mat((i3-1)*L2+k1,(i4-1)*L2+k2) = z; SFC_pcor_WM_Zval_mat((i4-1)*L2+k2,(i3-1)*L2+k1) = z; SFC_pcor_WM_Zval_mat((i3-1)*L2+k2,(i4-1)*L2+k1) = z; SFC_pcor_WM_Zval_mat((i4-1)*L2+k1,(i3-1)*L2+k2) = z;
                    SFC_pcor_WM_Rval_mat((i3-1)*L2+k1,(i4-1)*L2+k2) = r; SFC_pcor_WM_Rval_mat((i4-1)*L2+k2,(i3-1)*L2+k1) = r; SFC_pcor_WM_Rval_mat((i3-1)*L2+k2,(i4-1)*L2+k1) = r; SFC_pcor_WM_Rval_mat((i4-1)*L2+k1,(i3-1)*L2+k2) = r;
                    catch
                    SFC_pcor_WM_Zval(k1,k2,i3,i4) = NaN; SFC_pcor_WM_Zval(k2,k1,i3,i4) = NaN; SFC_pcor_WM_Zval(k1,k2,i4,i3) = NaN; SFC_pcor_WM_Zval(k2,k1,i4,i3) = NaN;
                    SFC_pcor_WM_Rval(k1,k2,i3,i4) = NaN; SFC_pcor_WM_Rval(k2,k1,i3,i4) = NaN; SFC_pcor_WM_Rval(k1,k2,i4,i3) = NaN; SFC_pcor_WM_Rval(k2,k1,i4,i3) = NaN;
                    SFC_pcor_WM_timeseries{k1,k2,i3,i4}(:,1) = NaN; SFC_pcor_WM_timeseries{k1,k2,i3,i4}(:,2) = NaN;
                    SFC_pcor_WM_Zval_mat((i3-1)*L2+k1,(i4-1)*L2+k2) = NaN; SFC_pcor_WM_Zval_mat((i4-1)*L2+k2,(i3-1)*L2+k1) = NaN; SFC_pcor_WM_Zval_mat((i3-1)*L2+k2,(i4-1)*L2+k1) = NaN; SFC_pcor_WM_Zval_mat((i4-1)*L2+k1,(i3-1)*L2+k2) = NaN;
                    SFC_pcor_WM_Rval_mat((i3-1)*L2+k1,(i4-1)*L2+k2) = NaN; SFC_pcor_WM_Rval_mat((i4-1)*L2+k2,(i3-1)*L2+k1) = NaN; SFC_pcor_WM_Rval_mat((i3-1)*L2+k2,(i4-1)*L2+k1) = NaN; SFC_pcor_WM_Rval_mat((i4-1)*L2+k1,(i3-1)*L2+k2) = NaN;
                    end
                    clear temp2 junk z r ts1 ts2
                end
            end
        end
        SFC_pcor_WM_Rval(k1,k1,i3,i3) = 1;
        clear temp1
    end
end
end

% save
info_SFC_pcor = sprintf('\nZval = Z-value obtained after Fischers R-to-Z transformation of partial correlation values; R-val = partial correlation (R) value\nDimensions: 4 quadrants x 4 quadrants x slices\nIn each slice pair, the 4x4 matrix represents the SFC value between corresponding quadrants:\nQuadrants: 1 = left ventral, 2 = right ventral, 3 = left dorsal, 4 = right dorsal\nExample: the element (1,3,6,9) represents the SFC between (1)left ventral quadrant in slice-6 and (3)left dorsal quadrant in slice-9\nNote that only upper triangle data is available for some variables as FC is non-directional and symmetric\n\n');
if WM_flag == 1
save([base_dir_sub naming '_SFC_betweenslice_pcor' '.mat'], 'SFC_pcor_GM_Zval', 'SFC_pcor_GM_Rval', 'SFC_pcor_GM_timeseries', 'SFC_pcor_WM_Zval', 'SFC_pcor_WM_Rval', 'SFC_pcor_WM_timeseries', 'SFC_pcor_GM_Zval_mat', 'SFC_pcor_GM_Rval_mat', 'SFC_pcor_WM_Zval_mat', 'SFC_pcor_WM_Rval_mat', 'info_SFC_pcor')
else
save([base_dir_sub naming '_SFC_betweenslice_pcor' '.mat'], 'SFC_pcor_GM_Zval', 'SFC_pcor_GM_Rval', 'SFC_pcor_GM_timeseries', 'SFC_pcor_GM_Zval_mat', 'SFC_pcor_GM_Rval_mat', 'info_SFC_pcor')
end

warning on
end % partial correlation

%% QC plots of within+between-slice FC
siz3 = L1;
fprintf('21. QC plots: within+between slices FC -> subject (%s)\n',naming)

if pearson==1
Cg = SFC_GM_Rval_mat;
Cg2 = []; Cg3 = [];
for v=1:siz3
    Cg2 = cat(1,Cg2,Cg(4*(v-1)+1:4*v,:),zeros(1,size(Cg,2)));
end
for v=1:siz3
    Cg3 = cat(2,Cg3,Cg2(:,4*(v-1)+1:4*v),zeros(size(Cg2,1),1));
end
clear Cg Cg2
Cg = Cg3; clear Cg3
Cg = Cg(1:end-1,1:end-1);

WhiteMask = ones(size(Cg)); WhiteMask(eye(size(Cg,1))==1) = 0;
BlackMask = ones(size(Cg)); BlackMask(find(and((Cg==0),(WhiteMask~=0)))) = 0;

fighndl = figure('Position',[1 1 1080 1080],'Color',[0.89,0.94,0.99],'InvertHardcopy','off','Visible','off'); % red [0.95,0.9,0.85], green [0.85,0.90,0.85], blue [0.89,0.94,0.99]
imagesc(Cg,'AlphaData',WhiteMask.*BlackMask,[-1 1]), axis square
	colormap('jet'),colorbar;
    for v=1:siz3, labelsC{v,1} = sprintf('%d',v); end
    xticks(2.5:5:size(Cg,1)), xticklabels(labelsC), a = get(gca,'xticklabels'); set(gca,'xticklabels',a,'fontsize',16); clear a
    yticks(2.5:5:size(Cg,1)), yticklabels(labelsC), a = get(gca,'yticklabels'); set(gca,'yticklabels',a,'fontsize',16); clear a
    xlabel('Slices (within each slice are shown LV | LD | RV | RD quadrants)'),ylabel('Slices (within each slice are shown LV | LD | RV | RD quadrants)')
    title(sprintf('GRAY MATTER\n(Pearson''s correlation based functional connectivity)\nMatrix of within- and between-slice FC across all slices'),'Color','blue','FontSize',18)
saveas(fighndl,[QCpath 'FCmatrix_Within_and_BetweenSlices_GM' '.jpg'])
saveas(fighndl,[QCpath2 'FCmatrix_Within_and_BetweenSlices_GM' '.jpg'])
close(fighndl)
clear fighndl Cg WhiteMask BlackMask

if WM_flag == 1
Cw = SFC_WM_Rval_mat;
Cw2 = []; Cw3 = [];
for v=1:siz3
    Cw2 = cat(1,Cw2,Cw(4*(v-1)+1:4*v,:),zeros(1,size(Cw,2)));
end
for v=1:siz3
    Cw3 = cat(2,Cw3,Cw2(:,4*(v-1)+1:4*v),zeros(size(Cw2,1),1));
end
clear Cw Cw2
Cw = Cw3; clear Cw3
Cw = Cw(1:end-1,1:end-1);

WhiteMask = ones(size(Cw)); WhiteMask(eye(size(Cw,1))==1) = 0;
BlackMask = ones(size(Cw)); BlackMask(find(and((Cw==0),(WhiteMask~=0)))) = 0;

fighndl = figure('Position',[1 1 1080 1080],'Color',[0.89,0.94,0.99],'InvertHardcopy','off','Visible','off'); % red [0.95,0.9,0.85], green [0.85,0.90,0.85], blue [0.89,0.94,0.99]
imagesc(Cw,'AlphaData',WhiteMask.*BlackMask,[-1 1]), axis square
	colormap('jet'),colorbar;
    for v=1:siz3, labelsC{v,1} = sprintf('%d',v); end
    xticks(2.5:5:size(Cw,1)), xticklabels(labelsC), a = get(gca,'xticklabels'); set(gca,'xticklabels',a,'fontsize',16); clear a
    yticks(2.5:5:size(Cw,1)), yticklabels(labelsC), a = get(gca,'yticklabels'); set(gca,'yticklabels',a,'fontsize',16); clear a
    xlabel('Slices (within each slice are shown LV | LD | RV | RD quadrants)'),ylabel('Slices (within each slice are shown LV | LD | RV | RD quadrants)')
    title(sprintf('WHITE MATTER\n(Pearson''s correlation based functional connectivity)\nMatrix of within- and between-slice FC across all slices'),'Color','red','FontSize',18)
saveas(fighndl,[QCpath 'FCmatrix_Within_and_BetweenSlices_WM' '.jpg'])
saveas(fighndl,[QCpath2 'FCmatrix_Within_and_BetweenSlices_WM' '.jpg'])
close(fighndl)
clear fighndl Cw WhiteMask BlackMask
end
end


if partial==1
Cg = SFC_pcor_GM_Rval_mat;
Cg2 = []; Cg3 = [];
for v=1:siz3
    Cg2 = cat(1,Cg2,Cg(4*(v-1)+1:4*v,:),zeros(1,size(Cg,2)));
end
for v=1:siz3
    Cg3 = cat(2,Cg3,Cg2(:,4*(v-1)+1:4*v),zeros(size(Cg2,1),1));
end
clear Cg Cg2
Cg = Cg3; clear Cg3
Cg = Cg(1:end-1,1:end-1);

WhiteMask = ones(size(Cg)); WhiteMask(eye(size(Cg,1))==1) = 0;
BlackMask = ones(size(Cg)); BlackMask(find(and((Cg==0),(WhiteMask~=0)))) = 0;

fighndl = figure('Position',[1 1 1080 1080],'Color',[0.89,0.94,0.99],'InvertHardcopy','off','Visible','off'); % red [0.95,0.9,0.85], green [0.85,0.90,0.85], blue [0.89,0.94,0.99]
imagesc(Cg,'AlphaData',WhiteMask.*BlackMask,[-1 1]), axis square
	colormap('jet'),colorbar;
    for v=1:siz3, labelsC{v,1} = sprintf('%d',v); end
    xticks(2.5:5:size(Cg,1)), xticklabels(labelsC), a = get(gca,'xticklabels'); set(gca,'xticklabels',a,'fontsize',16); clear a
    yticks(2.5:5:size(Cg,1)), yticklabels(labelsC), a = get(gca,'yticklabels'); set(gca,'yticklabels',a,'fontsize',16); clear a
    xlabel('Slices (within each slice are shown LV | LD | RV | RD quadrants)'),ylabel('Slices (within each slice are shown LV | LD | RV | RD quadrants)')
    title(sprintf('GRAY MATTER\n(Partial correlation based functional connectivity)\nMatrix of within- and between-slice FC across all slices'),'Color','blue','FontSize',18)
saveas(fighndl,[QCpath 'FCmatrix_partialcorr_Within_and_BetweenSlices_GM' '.jpg'])
saveas(fighndl,[QCpath2 'FCmatrix_partialcorr_Within_and_BetweenSlices_GM' '.jpg'])
close(fighndl)
clear fighndl Cg WhiteMask BlackMask

if WM_flag == 1
Cw = SFC_pcor_WM_Rval_mat;
Cw2 = []; Cw3 = [];
for v=1:siz3
    Cw2 = cat(1,Cw2,Cw(4*(v-1)+1:4*v,:),zeros(1,size(Cw,2)));
end
for v=1:siz3
    Cw3 = cat(2,Cw3,Cw2(:,4*(v-1)+1:4*v),zeros(size(Cw2,1),1));
end
clear Cw Cw2
Cw = Cw3; clear Cw3
Cw = Cw(1:end-1,1:end-1);

WhiteMask = ones(size(Cw)); WhiteMask(eye(size(Cw,1))==1) = 0;
BlackMask = ones(size(Cw)); BlackMask(find(and((Cw==0),(WhiteMask~=0)))) = 0;

fighndl = figure('Position',[1 1 1080 1080],'Color',[0.89,0.94,0.99],'InvertHardcopy','off','Visible','off'); % red [0.95,0.9,0.85], green [0.85,0.90,0.85], blue [0.89,0.94,0.99]
imagesc(Cw,'AlphaData',WhiteMask.*BlackMask,[-1 1]), axis square
	colormap('jet'),colorbar;
    for v=1:siz3, labelsC{v,1} = sprintf('%d',v); end
    xticks(2.5:5:size(Cw,1)), xticklabels(labelsC), a = get(gca,'xticklabels'); set(gca,'xticklabels',a,'fontsize',16); clear a
    yticks(2.5:5:size(Cw,1)), yticklabels(labelsC), a = get(gca,'yticklabels'); set(gca,'yticklabels',a,'fontsize',16); clear a
    xlabel('Slices (within each slice are shown LV | LD | RV | RD quadrants)'),ylabel('Slices (within each slice are shown LV | LD | RV | RD quadrants)')
    title(sprintf('WHITE MATTER\n(Partial correlation based functional connectivity)\nMatrix of within- and between-slice FC across all slices'),'Color','red','FontSize',18)
saveas(fighndl,[QCpath 'FCmatrix_partialcorr_Within_and_BetweenSlices_WM' '.jpg'])
saveas(fighndl,[QCpath2 'FCmatrix_partialcorr_Within_and_BetweenSlices_WM' '.jpg'])
close(fighndl)
clear fighndl Cw WhiteMask BlackMask
end
end


end
```
