## Supplementary material for "Neptune: a toolbox for spinal cord functional MRI data processing and quality assurance": Neptune software: scfMRItb_22_fALFF.m

```
function scfMRItb_22_fALFF(base_dir_sub,fname4, WM_flag,assign_value,Rns)

if nargin < 4
    assign_value = 1; % computing FC between all pairs of voxels in the quadrants and assign the 95th percentile value as the final FC value. If this flag is set to 1, then mean time series is computed for each quadrant instead and FC computed between them
    if nargin < 3
        WM_flag = 1; % compute SFC in the white matter (0 = don't)
    end
end

scrsz = get(0,'ScreenSize');
if Rns==0, run=1; else, run=Rns; end

if ~(exist([base_dir_sub 'QC' '/22_fALFF'],'dir'))
    mkdir([base_dir_sub 'QC' '/22_fALFF'])
end
if Rns==0
    QCpath = [base_dir_sub 'QC' '/22_fALFF/'];
else
    if ~(exist([base_dir_sub 'QC' '/22_fALFF' '/run' num2str(Rns)],'dir'))
        mkdir([base_dir_sub 'QC' '/22_fALFF' '/run' num2str(Rns)])
    end
    QCpath = [base_dir_sub 'QC' '/22_fALFF' '/run' num2str(Rns) '/'];
end

if ~(exist([base_dir_sub 'QC' '/QC_shortlist'],'dir'))
    mkdir([base_dir_sub 'QC' '/QC_shortlist'])
end
if Rns==0
    QCpath2 = [base_dir_sub 'QC' '/QC_shortlist/'];
else
    if ~(exist([base_dir_sub 'QC' '/QC_shortlist' '/run' num2str(Rns)],'dir'))
        mkdir([base_dir_sub 'QC' '/QC_shortlist' '/run' num2str(Rns)])
    end
    QCpath2 = [base_dir_sub 'QC' '/QC_shortlist' '/run' num2str(Rns) '/'];
end

%% ----- COMPUTE FRACTIONAL AMPLITUDE OF LOW FREQUENCY FLUCTUATIONS (fALFF) -----

load([base_dir_sub fname4 '.mat']) %#ok<*LOAD> % variable F
TR = F.hdr.dime.pixdim(5);
siz3 = size(F.img,3); siz4 = size(F.img,4);

if assign_value==1
    JLTM_cutoff = 0.95; % perform SFC between all pairs of voxels in the two quadrants and assign the 95th percentile value as the final SFC
elseif assign_value==2
    JLTM_cutoff = 0.5; % perform SFC between all pairs of voxels in the two quadrants and assign the 50th percentile value as the final SFC    
end

load([base_dir_sub 'workspace_variables_run' num2str(run) '.mat'],'FC_mask_quads');
try FC_mask_quads=FC_mask_quads.img; catch, end

skip_erosion = 0; % erode gray matter mask (1 = do not erode; -1 = dilate)
wm_erode = 6; % larger the number, more eroded the WM masks. 1 is the smallest value that can be assigned (least erosion).

if max([F.hdr.dime.pixdim(2),F.hdr.dime.pixdim(3)])<=1 % less than 1mm in-plane voxel size
    dil1 = 2;
else
    dil1 = 1;
end

%% extract time series from high-pass filtered data exclusively for fALFF

%%% gray matter

for i3 = siz3:-1:1 
    F_oneslice = squeeze(F.img(:,:,i3,:));
    for i9 = 1 : 4 % for each quadrant (GM)
        mmask_orig = (FC_mask_quads(:,:,i3) == i9);
        if     i9 == 1 % left ventral
            mmask_ipsi = (FC_mask_quads(:,:,i3) == 3) | (FC_mask_quads(:,:,i3) == 2);
        elseif i9 == 2 % right ventral
            mmask_ipsi = (FC_mask_quads(:,:,i3) == 4) | (FC_mask_quads(:,:,i3) == 1);
        elseif i9 == 3 % left dorsal
            mmask_ipsi = (FC_mask_quads(:,:,i3) == 1) | (FC_mask_quads(:,:,i3) == 4);
        elseif i9 == 4 % right dorsal
            mmask_ipsi = (FC_mask_quads(:,:,i3) == 2) | (FC_mask_quads(:,:,i3) == 3);
        end % ipsilateral mask
        mmask_dilated = imdilate(mmask_orig, strel('disk', dil1));
        mmask_ipsi_dilated = imdilate(mmask_ipsi, strel('disk', dil1));
        mmask_overlap = (mmask_dilated + mmask_ipsi_dilated) > 1;
        mmask_overlap_dilated = imdilate(mmask_overlap, strel('disk', 1));
        mmask_valid = ~mmask_overlap_dilated;
        clear mmask_dilated mmask_ipsi mmask_ipsi_dilated mmask_overlap mmask_overlap_dilated;
        
        if skip_erosion == 0 % erode mask
            quadmask = imerode(mmask_orig, strel('disk', 1));
            if sum(sum( and(quadmask, mmask_valid) )) <= 3
                quadmask = mmask_orig; % mask is already tiny, so don't erode
            end
        elseif skip_erosion == 1 % don't erode mask
            quadmask = mmask_orig;
        elseif skip_erosion == -1 % a secret option --> dilate mask!
            quadmask = imdilate(mmask_orig, strel('disk', 1));
        end % ~skip_erosion
        % exclude region near central gray matter
        if ~( sum(sum( and(quadmask, mmask_valid) )) <= 3 ) % don't do this if mask is already tiny
            quadmask = and(quadmask, mmask_valid);
        end
        
        num_mask = sum(quadmask(:));
        [r, c] = find(quadmask);
        index = [r, c];
        ttemp = F_oneslice .* repmat(quadmask, [1 1 siz4]);
        temp = zeros(num_mask, siz4);
        for i1 = 1 : num_mask
            temp(i1,:) = reshape(ttemp(r(i1),c(i1),:), [1 siz4]);
        end
        % results
        timeseries_GM{i3,i9} = temp;
        clear temp mmask_valid mmask_orig mmask num_mask index r c
    end % i9
    clear F_oneslice
end % i3

%%% white matter
if WM_flag == 1 % perform steps in the white matter (0 = don't)
for i3 = siz3:-1:1
    F_oneslice = squeeze(F.img(:,:,i3,:));
    for i9 = 5 : 8 % for each quadrant (WM)
        mmask_orig = FC_mask_quads(:,:,i3) == i9;
        imask = wm_erode;
        quadmask = imerode(mmask_orig, strel('disk', imask));
        while sum(quadmask(:)) <= 1
            imask = imask - 1;
            quadmask = imerode(mmask_orig, strel('disk', imask));
        end
        % --- keeping the next two lines will create slightly larger WM quads than otherwise ---
        if imask>=2
            imask = imask - 1;
            quadmask = imerode(mmask_orig, strel('disk', imask));
        end
        % --- ---
        
        num_mask = sum(quadmask(:));
        ttemp = F_oneslice .* repmat(quadmask, [1 1 siz4]);
        [r, c] = find(quadmask);
        index = [r, c];
        temp = zeros(num_mask, siz4);
        for i1 = 1 : num_mask
            temp(i1,:) = reshape(ttemp(r(i1),c(i1),:), [1 siz4]);
        end
        % save results
        timeseries_WM{i3,i9-4} = temp;
        clear temp mmask_valid mmask_orig mmask num_mask index imask r c
    end % i9
    clear F_oneslice
end % i3
end % WM_flag

[L1,L2] = size(timeseries_GM);

%% fALFF
fALFF_GM = zeros(L1,L2);
for i3 = 1:L1
    fprintf('fALFF GM: slice (%d) of (%d) -> Subject (%s)\n',i3,L1,fname4)
    for k1 = 1:L2
        temp1 = timeseries_GM{i3,k1};
        if assign_value == 3 % if this flag is 1 then use mean time series from each quadrant instead of computing fALFF between all voxels and choosing the 95th percentile value
            temp1 = mean(temp1,1);
        end
        try %#ok<*ALIGN>
        temp2 = falff2(temp1', TR, 0.1, 0.01);
        temp2 = sort(temp2);
        fALFF_GM(i3,k1) = temp2(ceil(length(temp2)*JLTM_cutoff));
        catch, end
        clear temp1 temp2
    end
end

if WM_flag == 1 % compute fALFF in the white matter (0 = don't)
fALFF_WM = zeros(L1,L2);
for i3 = 1:L1
    fprintf('fALFF WM: slice (%d) of (%d) -> Subject (%s)\n',i3,L1,fname4)
    for k1 = 1:L2
        temp1 = timeseries_WM{i3,k1};
        if assign_value == 3 % if this flag is 1 then use mean time series from each quadrant instead of computing fALFF between all voxels and choosing the 95th percentile value
            temp1 = mean(temp1,1);
        end
        try
        temp2 = falff2(temp1', TR, 0.1, 0.01);
        temp2 = sort(temp2);
        fALFF_WM(i3,k1) = temp2(ceil(length(temp2)*JLTM_cutoff));
        catch, end
        clear temp1 temp2
    end
end
end

% save
info_fALFF = sprintf('\nDimensions: slices x 4 quadrants\nIn each slice, the four values correspond to the fALFF value of corresponding quadrants:\nQuadrants: 1 = left ventral, 2 = right ventral, 3 = left dorsal, 4 = right dorsal\n\n');
if WM_flag == 1
save([base_dir_sub fname4 '_fALFF' '.mat'], 'fALFF_GM', 'fALFF_WM', 'info_fALFF')
else
save([base_dir_sub fname4 '_fALFF' '.mat'], 'fALFF_GM', 'info_fALFF')
end

%% QC plots of fALFF
fprintf('22. QC plot: fALFF -> subject (%s)\n',fname4)

if WM_flag == 0
    fALFF_WM = zeros(size(fALFF_GM));
    fALFF_WM(find(fALFF_WM==0)) = NaN;
end

fighndl = figure('Position',[1 1 1280 1280],'Color',[0.85,0.90,0.85],'InvertHardcopy','off','Visible','off'); % red [0.95,0.9,0.85], green [0.85,0.90,0.85], blue [0.89,0.94,0.99]
subplot2n(5,1,1), bar([fALFF_GM(:,1),fALFF_WM(:,1)]'), ylabel('fALFF'), xticklabels({sprintf('GRAY MATTER fALFF (slices 1 -> %d)',siz3),sprintf('WHITE MATTER fALFF (slices 1 -> %d)',siz3)}),
    a = get(gca,'xticklabels'); set(gca,'xticklabels',a,'fontsize',12); clear a
    grid on,grid minor, 
    try ylim([min([fALFF_GM(:);fALFF_WM(:)])-(max([fALFF_GM(:);fALFF_WM(:)])-min([fALFF_GM(:);fALFF_WM(:)]))/2, max([fALFF_GM(:);fALFF_WM(:)])+(max([fALFF_GM(:);fALFF_WM(:)])-min([fALFF_GM(:);fALFF_WM(:)]))/2]), catch, end
    title(sprintf('Fractional amplitude of low-frequency BOLD fluctuations (fALFF)\n\nfALFF across SLICES (left ventral horn)'),'Color',[0,0.65,0.59],'FontSize',16), 
    text(0.9,max([fALFF_GM(:);fALFF_WM(:)])+(max([fALFF_GM(:);fALFF_WM(:)])-min([fALFF_GM(:);fALFF_WM(:)]))/4, sprintf('mean = %.2f',mean(fALFF_GM(:,1))),'Color',[0,0.55,0.69],'FontSize',15)
    text(1.9,max([fALFF_GM(:);fALFF_WM(:)])+(max([fALFF_GM(:);fALFF_WM(:)])-min([fALFF_GM(:);fALFF_WM(:)]))/4, sprintf('mean = %.2f',mean(fALFF_WM(:,1))),'Color',[0.65,0.11,0.19],'FontSize',15)
subplot2n(5,1,2), bar([fALFF_GM(:,2),fALFF_WM(:,2)]'), ylabel('fALFF'), xticklabels({sprintf('GRAY MATTER fALFF (slices 1 -> %d)',siz3),sprintf('WHITE MATTER fALFF (slices 1 -> %d)',siz3)}),
    a = get(gca,'xticklabels'); set(gca,'xticklabels',a,'fontsize',12); clear a
    grid on,grid minor, 
    try ylim([min([fALFF_GM(:);fALFF_WM(:)])-(max([fALFF_GM(:);fALFF_WM(:)])-min([fALFF_GM(:);fALFF_WM(:)]))/2, max([fALFF_GM(:);fALFF_WM(:)])+(max([fALFF_GM(:);fALFF_WM(:)])-min([fALFF_GM(:);fALFF_WM(:)]))/2]), catch, end
    title(['fALFF across SLICES (right ventral horn)'],'Color',[0,0.45,0.79],'FontSize',16), 
    text(0.9,max([fALFF_GM(:);fALFF_WM(:)])+(max([fALFF_GM(:);fALFF_WM(:)])-min([fALFF_GM(:);fALFF_WM(:)]))/4, sprintf('mean = %.2f',mean(fALFF_GM(:,2))),'Color',[0,0.55,0.69],'FontSize',15)
    text(1.9,max([fALFF_GM(:);fALFF_WM(:)])+(max([fALFF_GM(:);fALFF_WM(:)])-min([fALFF_GM(:);fALFF_WM(:)]))/4, sprintf('mean = %.2f',mean(fALFF_WM(:,2))),'Color',[0.65,0.11,0.19],'FontSize',15)
subplot2n(5,1,3), bar([fALFF_GM(:,3),fALFF_WM(:,3)]'), ylabel('fALFF'), xticklabels({sprintf('GRAY MATTER fALFF (slices 1 -> %d)',siz3),sprintf('WHITE MATTER fALFF (slices 1 -> %d)',siz3)}),
    a = get(gca,'xticklabels'); set(gca,'xticklabels',a,'fontsize',12); clear a
    grid on,grid minor, 
    try ylim([min([fALFF_GM(:);fALFF_WM(:)])-(max([fALFF_GM(:);fALFF_WM(:)])-min([fALFF_GM(:);fALFF_WM(:)]))/2, max([fALFF_GM(:);fALFF_WM(:)])+(max([fALFF_GM(:);fALFF_WM(:)])-min([fALFF_GM(:);fALFF_WM(:)]))/2]), catch, end
    title(['fALFF across SLICES (left dorsal horn)'],'Color',[0.55,0.01,0.39],'FontSize',16), 
    text(0.9,max([fALFF_GM(:);fALFF_WM(:)])+(max([fALFF_GM(:);fALFF_WM(:)])-min([fALFF_GM(:);fALFF_WM(:)]))/4, sprintf('mean = %.2f',mean(fALFF_GM(:,3))),'Color',[0,0.55,0.69],'FontSize',15)
    text(1.9,max([fALFF_GM(:);fALFF_WM(:)])+(max([fALFF_GM(:);fALFF_WM(:)])-min([fALFF_GM(:);fALFF_WM(:)]))/4, sprintf('mean = %.2f',mean(fALFF_WM(:,3))),'Color',[0.65,0.11,0.19],'FontSize',15)
subplot2n(5,1,4), bar([fALFF_GM(:,4),fALFF_WM(:,4)]'), ylabel('fALFF'), xticklabels({sprintf('GRAY MATTER fALFF (slices 1 -> %d)',siz3),sprintf('WHITE MATTER fALFF (slices 1 -> %d)',siz3)}),
    a = get(gca,'xticklabels'); set(gca,'xticklabels',a,'fontsize',12); clear a
    grid on,grid minor, 
    try ylim([min([fALFF_GM(:);fALFF_WM(:)])-(max([fALFF_GM(:);fALFF_WM(:)])-min([fALFF_GM(:);fALFF_WM(:)]))/2, max([fALFF_GM(:);fALFF_WM(:)])+(max([fALFF_GM(:);fALFF_WM(:)])-min([fALFF_GM(:);fALFF_WM(:)]))/2]), catch, end
    title(['fALFF across SLICES (right dorsal horn)'],'Color',[0.55,0.31,0.09],'FontSize',16), 
    text(0.9,max([fALFF_GM(:);fALFF_WM(:)])+(max([fALFF_GM(:);fALFF_WM(:)])-min([fALFF_GM(:);fALFF_WM(:)]))/4, sprintf('mean = %.2f',mean(fALFF_GM(:,4))),'Color',[0,0.55,0.69],'FontSize',15)
    text(1.9,max([fALFF_GM(:);fALFF_WM(:)])+(max([fALFF_GM(:);fALFF_WM(:)])-min([fALFF_GM(:);fALFF_WM(:)]))/4, sprintf('mean = %.2f',mean(fALFF_WM(:,4))),'Color',[0.65,0.11,0.19],'FontSize',15)
subplot2n(5,1,5), bar([mean(fALFF_GM,1)',mean(fALFF_WM,1)']'), ylabel('fALFF'), xticklabels({sprintf('Median GRAY MATTER fALFF (within LV | RV | LD | RD)'),sprintf('Median WHITE MATTER fALFF (within LV | RV | LD | RD)')}),
    a = get(gca,'xticklabels'); set(gca,'xticklabels',a,'fontsize',12); clear a
    grid on,grid minor, 
    try ylim([min([fALFF_GM(:);fALFF_WM(:)])-(max([fALFF_GM(:);fALFF_WM(:)])-min([fALFF_GM(:);fALFF_WM(:)]))/2, max([fALFF_GM(:);fALFF_WM(:)])+(max([fALFF_GM(:);fALFF_WM(:)])-min([fALFF_GM(:);fALFF_WM(:)]))/2]), catch, end
    title(['Median fALFF across QUADRANTS (LV | RV | LD | RD)'],'Color',[0.24,0.25,0.26],'FontSize',16), 
    text(0.9,max([fALFF_GM(:);fALFF_WM(:)])+(max([fALFF_GM(:);fALFF_WM(:)])-min([fALFF_GM(:);fALFF_WM(:)]))/4, sprintf('median = %.2f',median(fALFF_GM(:))),'Color',[0,0.55,0.69],'FontSize',15)
    text(1.9,max([fALFF_GM(:);fALFF_WM(:)])+(max([fALFF_GM(:);fALFF_WM(:)])-min([fALFF_GM(:);fALFF_WM(:)]))/4, sprintf('median = %.2f',median(fALFF_WM(:))),'Color',[0.65,0.11,0.19],'FontSize',15)
saveas(fighndl,[QCpath 'fALFF_across_slices' '.jpg'])
saveas(fighndl,[QCpath2 'fALFF_across_slices' '.jpg'])
close(fighndl)
    
end
```
