## Supplementary material for "Neptune: a toolbox for spinal cord functional MRI data processing and quality assurance": Neptune software: scfMRItb_23_stats1.m

```
function scfMRItb_23_stats1(sub_dir,func_files,postproc_filesuffix,choices_postproc, indx_group1,indx_group2,ttest_type,alpha_stats, WM_flag,slices,SFC_type,SFC_withinslice_flag,SFC_betweenslice_flag, use_deconv_data_for_postproc,dont_do_pcor_betweenslice)

scrsz = get(0,'ScreenSize');
siz3 = slices;

home_dir = sub_dir{1}(1:max(strfind(sub_dir{1}(1:end-1),'/')));
if isempty(home_dir)
    home_dir = sub_dir{1}(1:max(strfind(sub_dir{1}(1:end-1),'\')));
end

tmp=clock; clk_stats = sprintf('%.4d-%.2d-%.2d_%.2d-%.2d',tmp(1),tmp(2),tmp(3),tmp(4),tmp(5)); clear tmp

if ~(exist([home_dir 'stats/figures/'],'dir'))
    mkdir([home_dir 'stats/figures/'])
end
QCpath = [home_dir 'stats/figures/'];

if SFC_type == 1 % do only Pearson's correlation
    pearson = 1; partial = 0;
elseif SFC_type == 2 % do only partial correlation
    pearson = 0; partial = 1;
elseif SFC_type == 3 % do both Pearson's correlation and partial correlation separately
    pearson = 1; partial = 1;
end

for k=1:length(sub_dir)
    for k2=1:size(func_files,2)
        if (exist([sub_dir{k} func_files{k,k2} postproc_filesuffix{1} '.mat'], 'file'))
            ip_file{k,k2} = [func_files{k,k2} postproc_filesuffix{1}];
        elseif (exist([sub_dir{k} func_files{k,k2} postproc_filesuffix{2} '.mat'], 'file'))
            ip_file{k,k2} = [func_files{k,k2} postproc_filesuffix{2}];
        elseif (exist([sub_dir{k} func_files{k,k2} postproc_filesuffix{3} '.mat'], 'file'))
            ip_file{k,k2} = [func_files{k,k2} postproc_filesuffix{3}];
        elseif (exist([sub_dir{k} func_files{k,k2} postproc_filesuffix{4} '.mat'], 'file'))
            ip_file{k,k2} = [func_files{k,k2} postproc_filesuffix{4}];
        elseif (exist([sub_dir{k} func_files{k,k2} postproc_filesuffix{5} '.mat'], 'file'))
            ip_file{k,k2} = [func_files{k,k2} postproc_filesuffix{5}];
        elseif (exist([sub_dir{k} func_files{k,k2} postproc_filesuffix{6} '.mat'], 'file'))
            ip_file{k,k2} = [func_files{k,k2} postproc_filesuffix{6}];
        end
    end
end

for k=1:length(sub_dir)
    for k2=1:size(func_files,2)
        if use_deconv_data_for_postproc == 1
            if (exist([sub_dir{k} func_files{k,k2} postproc_filesuffix{3} '.mat'], 'file'))
                ip_file2{k,k2} = [func_files{k,k2} postproc_filesuffix{3}];
            elseif (exist([sub_dir{k} func_files{k,k2} postproc_filesuffix{5} '.mat'], 'file'))
                ip_file2{k,k2} = [func_files{k,k2} postproc_filesuffix{5}];
            elseif (exist([sub_dir{k} func_files{k,k2} postproc_filesuffix{6} '.mat'], 'file'))
                ip_file2{k,k2} = [func_files{k,k2} postproc_filesuffix{6}];
            end
        elseif use_deconv_data_for_postproc == 0
            if (exist([sub_dir{k} func_files{k,k2} postproc_filesuffix{2} '.mat'], 'file'))
                ip_file2{k,k2} = [func_files{k,k2} postproc_filesuffix{2}];
            elseif (exist([sub_dir{k} func_files{k,k2} postproc_filesuffix{5} '.mat'], 'file'))
                ip_file2{k,k2} = [func_files{k,k2} postproc_filesuffix{5}];
            elseif (exist([sub_dir{k} func_files{k,k2} postproc_filesuffix{6} '.mat'], 'file'))
                ip_file2{k,k2} = [func_files{k,k2} postproc_filesuffix{6}];
            end
        end
    end
end

alpha = alpha_stats; % 0.05; % statistical threshold (uncorrected 0.05)

%% SFC stats Pearson's correlation
if ((choices_postproc(2)==1)||exist([sub_dir{1} ip_file{1,1} '_SFC_withinslice' '.mat'],'file')||exist([sub_dir{1} ip_file{1,1} '_SFC_betweenslice' '.mat'],'file')) && (pearson==1)

if SFC_withinslice_flag==1
    
if ttest_type==1 % one-sample t-test
f=0;
for k=1:length(sub_dir)
    if ismember(k,indx_group1)
    f=f+1;
    for k2=1:size(func_files,2)
        if WM_flag == 1
            load([sub_dir{k} ip_file{k,k2} '_SFC_withinslice' '.mat'], 'SFC_GM_Zval', 'SFC_WM_Zval')
            tempg(:,:,:,k2) = SFC_GM_Zval; clear SFC_GM_Zval
            tempw(:,:,:,k2) = SFC_WM_Zval; clear SFC_WM_Zval
        else
            load([sub_dir{k} ip_file{k,k2} '_SFC_withinslice' '.mat'], 'SFC_GM_Zval')
            tempg(:,:,:,k2) = SFC_GM_Zval; clear SFC_GM_Zval
        end
    end
    SFC_GM_all(:,:,:,f) = mean(tempg,4);
    if WM_flag == 1
    SFC_WM_all(:,:,:,f) = mean(tempw,4);
    end
    clear tempg tempw
    end
end; clear f k k2
for w1=1:size(SFC_GM_all,1)
    for w2=1:size(SFC_GM_all,2)
        for w3=1:size(SFC_GM_all,3)
            [h_SFC_GM_all_WS(w1,w2,w3),p_SFC_GM_all_WS(w1,w2,w3),~,stats] = ttest(squeeze(SFC_GM_all(w1,w2,w3,:)),0,'Alpha',alpha);
            tstat_SFC_GM_all_WS(w1,w2,w3) = stats.tstat; clear stats
            if WM_flag == 1
            [h_SFC_WM_all_WS(w1,w2,w3),p_SFC_WM_all_WS(w1,w2,w3),~,stats] = ttest(squeeze(SFC_WM_all(w1,w2,w3,:)),0,'Alpha',alpha);
            tstat_SFC_WM_all_WS(w1,w2,w3) = stats.tstat; clear stats
            end
        end
        [h_SFC_GM_all_WS_SlicesPooled(w1,w2),p_SFC_GM_all_WS_SlicesPooled(w1,w2),~,stats] = ttest(reshape(squeeze(SFC_GM_all(w1,w2,:,:)),[],1),0,'Alpha',alpha);
        tstat_SFC_GM_all_WS_SlicesPooled(w1,w2) = stats.tstat; clear stats
        if WM_flag == 1
        [h_SFC_WM_all_WS_SlicesPooled(w1,w2),p_SFC_WM_all_WS_SlicesPooled(w1,w2),~,stats] = ttest(reshape(squeeze(SFC_WM_all(w1,w2,:,:)),[],1),0,'Alpha',alpha);
        tstat_SFC_WM_all_WS_SlicesPooled(w1,w2) = stats.tstat; clear stats
        end
    end
end

elseif ttest_type==2 % paired t-test
    if sum(abs(indx_group1-indx_group2))==0
        f=0;
        for k=1:length(sub_dir)
            if ismember(k,indx_group1)
                f=f+1;
                if WM_flag == 1
                    load([sub_dir{k} ip_file{k,1} '_SFC_withinslice' '.mat'], 'SFC_GM_Zval', 'SFC_WM_Zval')
                    SFC_GM_all_group1(:,:,:,f) = SFC_GM_Zval; clear SFC_GM_Zval
                    SFC_WM_all_group1(:,:,:,f) = SFC_WM_Zval; clear SFC_WM_Zval
                    load([sub_dir{k} ip_file{k,2} '_SFC_withinslice' '.mat'], 'SFC_GM_Zval', 'SFC_WM_Zval')
                    SFC_GM_all_group2(:,:,:,f) = SFC_GM_Zval; clear SFC_GM_Zval
                    SFC_WM_all_group2(:,:,:,f) = SFC_WM_Zval; clear SFC_WM_Zval
                else
                    load([sub_dir{k} ip_file{k,1} '_SFC_withinslice' '.mat'], 'SFC_GM_Zval')
                    SFC_GM_all_group1(:,:,:,f) = SFC_GM_Zval; clear SFC_GM_Zval
                    load([sub_dir{k} ip_file{k,2} '_SFC_withinslice' '.mat'], 'SFC_GM_Zval')
                    SFC_GM_all_group2(:,:,:,f) = SFC_GM_Zval; clear SFC_GM_Zval
                end
            end
        end; clear f k k2
        for w1=1:size(SFC_GM_all_group1,1)
            for w2=1:size(SFC_GM_all_group1,2)
                for w3=1:size(SFC_GM_all_group1,3)
                    [h_SFC_GM_all_WS(w1,w2,w3),p_SFC_GM_all_WS(w1,w2,w3),~,stats] = ttest(squeeze(SFC_GM_all_group1(w1,w2,w3,:)),squeeze(SFC_GM_all_group2(w1,w2,w3,:)),'Alpha',alpha);
                    tstat_SFC_GM_all_WS(w1,w2,w3) = stats.tstat; clear stats
                    if WM_flag == 1
                    [h_SFC_WM_all_WS(w1,w2,w3),p_SFC_WM_all_WS(w1,w2,w3),~,stats] = ttest(squeeze(SFC_WM_all_group1(w1,w2,w3,:)),squeeze(SFC_WM_all_group2(w1,w2,w3,:)),'Alpha',alpha);
                    tstat_SFC_WM_all_WS(w1,w2,w3) = stats.tstat; clear stats
                    end
                end
                [h_SFC_GM_all_WS_SlicesPooled(w1,w2),p_SFC_GM_all_WS_SlicesPooled(w1,w2),~,stats] = ttest(reshape(squeeze(SFC_GM_all_group1(w1,w2,:,:)),squeeze(SFC_GM_all_group2(w1,w2,:,:)),[],1),'Alpha',alpha);
                tstat_SFC_GM_all_WS_SlicesPooled(w1,w2) = stats.tstat; clear stats
                if WM_flag == 1
                [h_SFC_WM_all_WS_SlicesPooled(w1,w2),p_SFC_WM_all_WS_SlicesPooled(w1,w2),~,stats] = ttest(reshape(squeeze(SFC_WM_all_group1(w1,w2,:,:)),squeeze(SFC_WM_all_group2(w1,w2,:,:)),[],1),'Alpha',alpha);
                tstat_SFC_WM_all_WS_SlicesPooled(w1,w2) = stats.tstat; clear stats
                end
            end
        end
        
    else
        f=0;
        for k=1:length(sub_dir)
            if ismember(k,indx_group1)
                f=f+1;
                for k2=1:size(func_files,2)
                    if WM_flag == 1
                        load([sub_dir{k} ip_file{k,k2} '_SFC_withinslice' '.mat'], 'SFC_GM_Zval', 'SFC_WM_Zval')
                        tempg(:,:,:,k2) = SFC_GM_Zval; clear SFC_GM_Zval
                        tempw(:,:,:,k2) = SFC_WM_Zval; clear SFC_WM_Zval
                    else
                        load([sub_dir{k} ip_file{k,k2} '_SFC_withinslice' '.mat'], 'SFC_GM_Zval')
                        tempg(:,:,:,k2) = SFC_GM_Zval; clear SFC_GM_Zval
                    end
                end
                SFC_GM_all_group1(:,:,:,f) = mean(tempg,4);
                if WM_flag == 1
                SFC_WM_all_group1(:,:,:,f) = mean(tempw,4);
                end
                clear tempg tempw
            end
        end; clear f k k2
        f=0;
        for k=1:length(sub_dir)
            if ismember(k,indx_group2)
                f=f+1;
                for k2=1:size(func_files,2)
                    if WM_flag == 1
                        load([sub_dir{k} ip_file{k,k2} '_SFC_withinslice' '.mat'], 'SFC_GM_Zval', 'SFC_WM_Zval')
                        tempg(:,:,:,k2) = SFC_GM_Zval; clear SFC_GM_Zval
                        tempw(:,:,:,k2) = SFC_WM_Zval; clear SFC_WM_Zval
                    else
                        load([sub_dir{k} ip_file{k,k2} '_SFC_withinslice' '.mat'], 'SFC_GM_Zval')
                        tempg(:,:,:,k2) = SFC_GM_Zval; clear SFC_GM_Zval
                    end
                end
                SFC_GM_all_group2(:,:,:,f) = mean(tempg,4);
                if WM_flag == 1
                SFC_WM_all_group2(:,:,:,f) = mean(tempw,4);
                end
                clear tempg tempw
            end
        end; clear f k k2
        for w1=1:size(SFC_GM_all_group1,1)
            for w2=1:size(SFC_GM_all_group1,2)
                for w3=1:size(SFC_GM_all_group1,3)
                    [h_SFC_GM_all_WS(w1,w2,w3),p_SFC_GM_all_WS(w1,w2,w3),~,stats] = ttest(squeeze(SFC_GM_all_group1(w1,w2,w3,:)),squeeze(SFC_GM_all_group2(w1,w2,w3,:)),'Alpha',alpha);
                    tstat_SFC_GM_all_WS(w1,w2,w3) = stats.tstat; clear stats
                    if WM_flag == 1
                    [h_SFC_WM_all_WS(w1,w2,w3),p_SFC_WM_all_WS(w1,w2,w3),~,stats] = ttest(squeeze(SFC_WM_all_group1(w1,w2,w3,:)),squeeze(SFC_WM_all_group2(w1,w2,w3,:)),'Alpha',alpha);
                    tstat_SFC_WM_all_WS(w1,w2,w3) = stats.tstat; clear stats
                    end
                end
                [h_SFC_GM_all_WS_SlicesPooled(w1,w2),p_SFC_GM_all_WS_SlicesPooled(w1,w2),~,stats] = ttest(reshape(squeeze(SFC_GM_all_group1(w1,w2,:,:)),squeeze(SFC_GM_all_group2(w1,w2,:,:)),[],1),'Alpha',alpha);
                tstat_SFC_GM_all_WS_SlicesPooled(w1,w2) = stats.tstat; clear stats
                if WM_flag == 1
                [h_SFC_WM_all_WS_SlicesPooled(w1,w2),p_SFC_WM_all_WS_SlicesPooled(w1,w2),~,stats] = ttest(reshape(squeeze(SFC_WM_all_group1(w1,w2,:,:)),squeeze(SFC_WM_all_group2(w1,w2,:,:)),[],1),'Alpha',alpha);
                tstat_SFC_WM_all_WS_SlicesPooled(w1,w2) = stats.tstat; clear stats
                end
            end
        end
    end
    
elseif ttest_type==3 % two-sample t-test
    f=0;
    for k=1:length(sub_dir)
        if ismember(k,indx_group1)
            f=f+1;
            for k2=1:size(func_files,2)
                if WM_flag == 1
                    load([sub_dir{k} ip_file{k,k2} '_SFC_withinslice' '.mat'], 'SFC_GM_Zval', 'SFC_WM_Zval')
                    tempg(:,:,:,k2) = SFC_GM_Zval; clear SFC_GM_Zval
                    tempw(:,:,:,k2) = SFC_WM_Zval; clear SFC_WM_Zval
                else
                    load([sub_dir{k} ip_file{k,k2} '_SFC_withinslice' '.mat'], 'SFC_GM_Zval')
                    tempg(:,:,:,k2) = SFC_GM_Zval; clear SFC_GM_Zval
                end
            end
            SFC_GM_all_group1(:,:,:,f) = mean(tempg,4);
            if WM_flag == 1
            SFC_WM_all_group1(:,:,:,f) = mean(tempw,4);
            end
            clear tempg tempw
        end
    end; clear f k k2
    f=0;
    for k=1:length(sub_dir)
        if ismember(k,indx_group2)
            f=f+1;
            for k2=1:size(func_files,2)
                if WM_flag == 1
                    load([sub_dir{k} ip_file{k,k2} '_SFC_withinslice' '.mat'], 'SFC_GM_Zval', 'SFC_WM_Zval')
                    tempg(:,:,:,k2) = SFC_GM_Zval; clear SFC_GM_Zval
                    tempw(:,:,:,k2) = SFC_WM_Zval; clear SFC_WM_Zval
                else
                    load([sub_dir{k} ip_file{k,k2} '_SFC_withinslice' '.mat'], 'SFC_GM_Zval')
                    tempg(:,:,:,k2) = SFC_GM_Zval; clear SFC_GM_Zval
                end
            end
            SFC_GM_all_group2(:,:,:,f) = mean(tempg,4);
            if WM_flag == 1
            SFC_WM_all_group2(:,:,:,f) = mean(tempw,4);
            end
            clear tempg tempw
        end
    end; clear f k k2
    for w1=1:size(SFC_GM_all_group1,1)
        for w2=1:size(SFC_GM_all_group1,2)
            for w3=1:size(SFC_GM_all_group1,3)
                [h_SFC_GM_all_WS(w1,w2,w3),p_SFC_GM_all_WS(w1,w2,w3),~,stats] = ttest2(squeeze(SFC_GM_all_group1(w1,w2,w3,:)),squeeze(SFC_GM_all_group2(w1,w2,w3,:)),'Alpha',alpha);
                tstat_SFC_GM_all_WS(w1,w2,w3) = stats.tstat; clear stats
                if WM_flag == 1
                [h_SFC_WM_all_WS(w1,w2,w3),p_SFC_WM_all_WS(w1,w2,w3),~,stats] = ttest2(squeeze(SFC_WM_all_group1(w1,w2,w3,:)),squeeze(SFC_WM_all_group2(w1,w2,w3,:)),'Alpha',alpha);
                tstat_SFC_WM_all_WS(w1,w2,w3) = stats.tstat; clear stats
                end
            end
            [h_SFC_GM_all_WS_SlicesPooled(w1,w2),p_SFC_GM_all_WS_SlicesPooled(w1,w2),~,stats] = ttest2(reshape(squeeze(SFC_GM_all_group1(w1,w2,:,:)),squeeze(SFC_GM_all_group2(w1,w2,:,:)),[],1),'Alpha',alpha);
            tstat_SFC_GM_all_WS_SlicesPooled(w1,w2) = stats.tstat; clear stats
            if WM_flag == 1
            [h_SFC_WM_all_WS_SlicesPooled(w1,w2),p_SFC_WM_all_WS_SlicesPooled(w1,w2),~,stats] = ttest2(reshape(squeeze(SFC_WM_all_group1(w1,w2,:,:)),squeeze(SFC_WM_all_group2(w1,w2,:,:)),[],1),'Alpha',alpha);
            tstat_SFC_WM_all_WS_SlicesPooled(w1,w2) = stats.tstat; clear stats
            end
        end
    end
end

if exist('SFC_GM_all','var')
if WM_flag == 1
save([home_dir 'stats/' 'stats' '_SFC_withinslice_' clk_stats '.mat'],'SFC_GM_all','SFC_WM_all','h_SFC_GM_all_WS','p_SFC_GM_all_WS','tstat_SFC_GM_all_WS', 'h_SFC_WM_all_WS','p_SFC_WM_all_WS','tstat_SFC_WM_all_WS','h_SFC_GM_all_WS_SlicesPooled','p_SFC_GM_all_WS_SlicesPooled','tstat_SFC_GM_all_WS_SlicesPooled','h_SFC_WM_all_WS_SlicesPooled','p_SFC_WM_all_WS_SlicesPooled','tstat_SFC_WM_all_WS_SlicesPooled', 'ip_file','indx_group1','indx_group2','ttest_type')
else
save([home_dir 'stats/' 'stats' '_SFC_withinslice_' clk_stats '.mat'],'SFC_GM_all','h_SFC_GM_all_WS','p_SFC_GM_all_WS','tstat_SFC_GM_all_WS','h_SFC_GM_all_WS_SlicesPooled','p_SFC_GM_all_WS_SlicesPooled','tstat_SFC_GM_all_WS_SlicesPooled', 'ip_file','indx_group1','indx_group2','ttest_type')
end
end
if exist('SFC_GM_all_group1','var')
if WM_flag == 1
save([home_dir 'stats/' 'stats' '_SFC_withinslice_' clk_stats '.mat'],'SFC_GM_all_group1','SFC_GM_all_group2','SFC_WM_all_group1','SFC_WM_all_group2','h_SFC_GM_all_WS','p_SFC_GM_all_WS','tstat_SFC_GM_all_WS', 'h_SFC_WM_all_WS','p_SFC_WM_all_WS','tstat_SFC_WM_all_WS','h_SFC_GM_all_WS_SlicesPooled','p_SFC_GM_all_WS_SlicesPooled','tstat_SFC_GM_all_WS_SlicesPooled','h_SFC_WM_all_WS_SlicesPooled','p_SFC_WM_all_WS_SlicesPooled','tstat_SFC_WM_all_WS_SlicesPooled', 'ip_file','indx_group1','indx_group2','ttest_type')
else
save([home_dir 'stats/' 'stats' '_SFC_withinslice_' clk_stats '.mat'],'SFC_GM_all_group1','SFC_GM_all_group2','h_SFC_GM_all_WS','p_SFC_GM_all_WS','tstat_SFC_GM_all_WS','h_SFC_GM_all_WS_SlicesPooled','p_SFC_GM_all_WS_SlicesPooled','tstat_SFC_GM_all_WS_SlicesPooled', 'ip_file','indx_group1','indx_group2','ttest_type')
end
end
clear SFC_GM_all SFC_WM_all SFC_GM_all_group1 SFC_GM_all_group2 SFC_WM_all_group1 SFC_WM_all_group2
end


if SFC_betweenslice_flag==1
    
if ttest_type==1 % one-sample t-test
f=0;
for k=1:length(sub_dir)
    if ismember(k,indx_group1)
    f=f+1;
    for k2=1:size(func_files,2)
        if WM_flag == 1
            load([sub_dir{k} ip_file{k,k2} '_SFC_betweenslice' '.mat'], 'SFC_GM_Zval_mat', 'SFC_WM_Zval_mat')
            tempg(:,:,k2) = SFC_GM_Zval_mat; clear SFC_GM_Zval_mat
            tempw(:,:,k2) = SFC_WM_Zval_mat; clear SFC_WM_Zval_mat
        else
            load([sub_dir{k} ip_file{k,k2} '_SFC_betweenslice' '.mat'], 'SFC_GM_Zval_mat')
            tempg(:,:,k2) = SFC_GM_Zval_mat; clear SFC_GM_Zval_mat
        end
    end
    SFC_GM_all(:,:,f) = mean(tempg,3);
    if WM_flag == 1
    SFC_WM_all(:,:,f) = mean(tempw,3);
    end
    clear tempg tempw
    end
end; clear f k k2
for w1=1:size(SFC_GM_all,1)
    for w2=1:size(SFC_GM_all,2)
        [h_SFC_GM_all_BS(w1,w2),p_SFC_GM_all_BS(w1,w2),~,stats] = ttest(squeeze(SFC_GM_all(w1,w2,:)),0,'Alpha',alpha);
        tstat_SFC_GM_all_BS(w1,w2) = stats.tstat; clear stats
        if WM_flag == 1
        [h_SFC_WM_all_BS(w1,w2),p_SFC_WM_all_BS(w1,w2),~,stats] = ttest(squeeze(SFC_WM_all(w1,w2,:)),0,'Alpha',alpha);
        tstat_SFC_WM_all_BS(w1,w2) = stats.tstat; clear stats
        end
    end
end

elseif ttest_type==2 % paired t-test
    if sum(abs(indx_group1-indx_group2))==0
        f=0;
        for k=1:length(sub_dir)
            if ismember(k,indx_group1)
                f=f+1;
                if WM_flag == 1
                    load([sub_dir{k} ip_file{k,1} '_SFC_betweenslice' '.mat'], 'SFC_GM_Zval_mat', 'SFC_WM_Zval_mat')
                    SFC_GM_all_group1(:,:,f) = SFC_GM_Zval_mat; clear SFC_GM_Zval_mat
                    SFC_WM_all_group1(:,:,f) = SFC_WM_Zval_mat; clear SFC_WM_Zval_mat
                    load([sub_dir{k} ip_file{k,2} '_SFC_betweenslice' '.mat'], 'SFC_GM_Zval_mat', 'SFC_WM_Zval_mat')
                    SFC_GM_all_group2(:,:,f) = SFC_GM_Zval_mat; clear SFC_GM_Zval_mat
                    SFC_WM_all_group2(:,:,f) = SFC_WM_Zval_mat; clear SFC_WM_Zval_mat
                else
                    load([sub_dir{k} ip_file{k,1} '_SFC_betweenslice' '.mat'], 'SFC_GM_Zval_mat')
                    SFC_GM_all_group1(:,:,f) = SFC_GM_Zval_mat; clear SFC_GM_Zval_mat
                    load([sub_dir{k} ip_file{k,2} '_SFC_betweenslice' '.mat'], 'SFC_GM_Zval_mat')
                    SFC_GM_all_group2(:,:,f) = SFC_GM_Zval_mat; clear SFC_GM_Zval_mat
                end
            end
        end; clear f k k2
        for w1=1:size(SFC_GM_all_group1,1)
            for w2=1:size(SFC_GM_all_group1,2)
                [h_SFC_GM_all_BS(w1,w2),p_SFC_GM_all_BS(w1,w2),~,stats] = ttest(squeeze(SFC_GM_all_group1(w1,w2,:)),squeeze(SFC_GM_all_group2(w1,w2,:)),'Alpha',alpha);
                tstat_SFC_GM_all_BS(w1,w2) = stats.tstat; clear stats
                if WM_flag == 1
                [h_SFC_WM_all_BS(w1,w2),p_SFC_WM_all_BS(w1,w2),~,stats] = ttest(squeeze(SFC_WM_all_group1(w1,w2,:)),squeeze(SFC_WM_all_group2(w1,w2,:)),'Alpha',alpha);
                tstat_SFC_WM_all_BS(w1,w2) = stats.tstat; clear stats
                end
            end
        end
        
    else
        f=0;
        for k=1:length(sub_dir)
            if ismember(k,indx_group1)
                f=f+1;
                for k2=1:size(func_files,2)
                    if WM_flag == 1
                        load([sub_dir{k} ip_file{k,k2} '_SFC_betweenslice' '.mat'], 'SFC_GM_Zval_mat', 'SFC_WM_Zval_mat')
                        tempg(:,:,k2) = SFC_GM_Zval_mat; clear SFC_GM_Zval_mat
                        tempw(:,:,k2) = SFC_WM_Zval_mat; clear SFC_WM_Zval_mat
                    else
                        load([sub_dir{k} ip_file{k,k2} '_SFC_betweenslice' '.mat'], 'SFC_GM_Zval_mat')
                        tempg(:,:,k2) = SFC_GM_Zval_mat; clear SFC_GM_Zval_mat
                    end
                end
                SFC_GM_all_group1(:,:,f) = mean(tempg,3);
                if WM_flag == 1
                SFC_WM_all_group1(:,:,f) = mean(tempw,3);
                end
                clear tempg tempw
            end
        end; clear f k k2
        f=0;
        for k=1:length(sub_dir)
            if ismember(k,indx_group2)
                f=f+1;
                for k2=1:size(func_files,2)
                    if WM_flag == 1
                        load([sub_dir{k} ip_file{k,k2} '_SFC_betweenslice' '.mat'], 'SFC_GM_Zval_mat', 'SFC_WM_Zval_mat')
                        tempg(:,:,k2) = SFC_GM_Zval_mat; clear SFC_GM_Zval_mat
                        tempw(:,:,k2) = SFC_WM_Zval_mat; clear SFC_WM_Zval_mat
                    else
                        load([sub_dir{k} ip_file{k,k2} '_SFC_betweenslice' '.mat'], 'SFC_GM_Zval_mat')
                        tempg(:,:,k2) = SFC_GM_Zval_mat; clear SFC_GM_Zval_mat
                    end
                end
                SFC_GM_all_group2(:,:,f) = mean(tempg,3);
                if WM_flag == 1
                SFC_WM_all_group2(:,:,f) = mean(tempw,3);
                end
                clear tempg tempw
            end
        end; clear f k k2
        for w1=1:size(SFC_GM_all_group1,1)
            for w2=1:size(SFC_GM_all_group1,2)
                [h_SFC_GM_all_BS(w1,w2),p_SFC_GM_all_BS(w1,w2),~,stats] = ttest(squeeze(SFC_GM_all_group1(w1,w2,:)),squeeze(SFC_GM_all_group2(w1,w2,:)),'Alpha',alpha);
                tstat_SFC_GM_all_BS(w1,w2) = stats.tstat; clear stats
                if WM_flag == 1
                [h_SFC_WM_all_BS(w1,w2),p_SFC_WM_all_BS(w1,w2),~,stats] = ttest(squeeze(SFC_WM_all_group1(w1,w2,:)),squeeze(SFC_WM_all_group2(w1,w2,:)),'Alpha',alpha);
                tstat_SFC_WM_all_BS(w1,w2) = stats.tstat; clear stats
                end
            end
        end
    end
    
elseif ttest_type==3 % two-sample t-test
    f=0;
    for k=1:length(sub_dir)
        if ismember(k,indx_group1)
            f=f+1;
            for k2=1:size(func_files,2)
                if WM_flag == 1
                    load([sub_dir{k} ip_file{k,k2} '_SFC_betweenslice' '.mat'], 'SFC_GM_Zval_mat', 'SFC_WM_Zval_mat')
                    tempg(:,:,k2) = SFC_GM_Zval_mat; clear SFC_GM_Zval_mat
                    tempw(:,:,k2) = SFC_WM_Zval_mat; clear SFC_WM_Zval_mat
                else
                    load([sub_dir{k} ip_file{k,k2} '_SFC_betweenslice' '.mat'], 'SFC_GM_Zval_mat')
                    tempg(:,:,k2) = SFC_GM_Zval_mat; clear SFC_GM_Zval_mat
                end
            end
            SFC_GM_all_group1(:,:,f) = mean(tempg,3);
            if WM_flag == 1
            SFC_WM_all_group1(:,:,f) = mean(tempw,3);
            end
            clear tempg tempw
        end
    end; clear f k k2
    f=0;
    for k=1:length(sub_dir)
        if ismember(k,indx_group2)
            f=f+1;
            for k2=1:size(func_files,2)
                if WM_flag == 1
                    load([sub_dir{k} ip_file{k,k2} '_SFC_betweenslice' '.mat'], 'SFC_GM_Zval_mat', 'SFC_WM_Zval_mat')
                    tempg(:,:,k2) = SFC_GM_Zval_mat; clear SFC_GM_Zval_mat
                    tempw(:,:,k2) = SFC_WM_Zval_mat; clear SFC_WM_Zval_mat
                else
                    load([sub_dir{k} ip_file{k,k2} '_SFC_betweenslice' '.mat'], 'SFC_GM_Zval_mat')
                    tempg(:,:,k2) = SFC_GM_Zval_mat; clear SFC_GM_Zval_mat
                end
            end
            SFC_GM_all_group2(:,:,f) = mean(tempg,3);
            if WM_flag == 1
            SFC_WM_all_group2(:,:,f) = mean(tempw,3);
            end
            clear tempg tempw
        end
    end; clear f k k2
    for w1=1:size(SFC_GM_all_group1,1)
        for w2=1:size(SFC_GM_all_group1,2)
            [h_SFC_GM_all_BS(w1,w2),p_SFC_GM_all_BS(w1,w2),~,stats] = ttest2(squeeze(SFC_GM_all_group1(w1,w2,:)),squeeze(SFC_GM_all_group2(w1,w2,:)),'Alpha',alpha);
            tstat_SFC_GM_all_BS(w1,w2) = stats.tstat; clear stats
            if WM_flag == 1
            [h_SFC_WM_all_BS(w1,w2),p_SFC_WM_all_BS(w1,w2),~,stats] = ttest2(squeeze(SFC_WM_all_group1(w1,w2,:)),squeeze(SFC_WM_all_group2(w1,w2,:)),'Alpha',alpha);
            tstat_SFC_WM_all_BS(w1,w2) = stats.tstat; clear stats
            end
        end
    end
end

if exist('SFC_GM_all','var')
if WM_flag == 1
save([home_dir 'stats/' 'stats' '_SFC_betweenslice_' clk_stats '.mat'],'SFC_GM_all','SFC_WM_all','h_SFC_GM_all_BS','p_SFC_GM_all_BS','tstat_SFC_GM_all_BS', 'h_SFC_WM_all_BS','p_SFC_WM_all_BS','tstat_SFC_WM_all_BS','ip_file','indx_group1','indx_group2','ttest_type')
else
save([home_dir 'stats/' 'stats' '_SFC_betweenslice_' clk_stats '.mat'],'SFC_GM_all','h_SFC_GM_all_BS','p_SFC_GM_all_BS','tstat_SFC_GM_all_BS', 'ip_file','indx_group1','indx_group2','ttest_type')
end
end
if exist('SFC_GM_all_group1','var')
if WM_flag == 1
save([home_dir 'stats/' 'stats' '_SFC_betweenslice_' clk_stats '.mat'],'SFC_GM_all_group1','SFC_GM_all_group2','SFC_WM_all_group1','SFC_WM_all_group2','h_SFC_GM_all_BS','p_SFC_GM_all_BS','tstat_SFC_GM_all_BS', 'h_SFC_WM_all_BS','p_SFC_WM_all_BS','tstat_SFC_WM_all_BS','ip_file','indx_group1','indx_group2','ttest_type')
else
save([home_dir 'stats/' 'stats' '_SFC_betweenslice_' clk_stats '.mat'],'SFC_GM_all_group1','SFC_GM_all_group2','h_SFC_GM_all_BS','p_SFC_GM_all_BS','tstat_SFC_GM_all_BS', 'ip_file','indx_group1','indx_group2','ttest_type')
end
end
clear SFC_GM_all SFC_WM_all SFC_GM_all_group1 SFC_GM_all_group2 SFC_WM_all_group1 SFC_WM_all_group2
end

end

%% SFC stats Partial correlation
if ((choices_postproc(2)==1)||exist([sub_dir{1} ip_file{1,1} '_SFC_withinslice_pcor' '.mat'],'file')||exist([sub_dir{1} ip_file{1,1} '_SFC_betweenslice_pcor' '.mat'],'file')) && (partial==1)
if SFC_withinslice_flag==1
    
if ttest_type==1 % one-sample t-test
f=0;
for k=1:length(sub_dir)
    if ismember(k,indx_group1)
    f=f+1;
    for k2=1:size(func_files,2)
        if WM_flag == 1
            load([sub_dir{k} ip_file{k,k2} '_SFC_withinslice_pcor' '.mat'], 'SFC_pcor_GM_Zval', 'SFC_pcor_WM_Zval')
            tempg(:,:,:,k2) = SFC_pcor_GM_Zval; clear SFC_pcor_GM_Zval
            tempw(:,:,:,k2) = SFC_pcor_WM_Zval; clear SFC_pcor_WM_Zval
        else
            load([sub_dir{k} ip_file{k,k2} '_SFC_withinslice_pcor' '.mat'], 'SFC_pcor_GM_Zval')
            tempg(:,:,:,k2) = SFC_pcor_GM_Zval; clear SFC_pcor_GM_Zval
        end
    end
    SFC_pcor_GM_all(:,:,:,f) = mean(tempg,4);
    if WM_flag == 1
    SFC_pcor_WM_all(:,:,:,f) = mean(tempw,4);
    end
    clear tempg tempw
    end
end; clear f k k2
for w1=1:size(SFC_pcor_GM_all,1)
    for w2=1:size(SFC_pcor_GM_all,2)
        for w3=1:size(SFC_pcor_GM_all,3)
            [h_SFC_pcor_GM_all_WS(w1,w2,w3),p_SFC_pcor_GM_all_WS(w1,w2,w3),~,stats] = ttest(squeeze(SFC_pcor_GM_all(w1,w2,w3,:)),0,'Alpha',alpha);
            tstat_SFC_pcor_GM_all_WS(w1,w2,w3) = stats.tstat; clear stats
            if WM_flag == 1
            [h_SFC_pcor_WM_all_WS(w1,w2,w3),p_SFC_pcor_WM_all_WS(w1,w2,w3),~,stats] = ttest(squeeze(SFC_pcor_WM_all(w1,w2,w3,:)),0,'Alpha',alpha);
            tstat_SFC_pcor_WM_all_WS(w1,w2,w3) = stats.tstat; clear stats
            end
        end
        [h_SFC_pcor_GM_all_WS_SlicesPooled(w1,w2),p_SFC_pcor_GM_all_WS_SlicesPooled(w1,w2),~,stats] = ttest(reshape(squeeze(SFC_pcor_GM_all(w1,w2,:,:)),[],1),0,'Alpha',alpha);
        tstat_SFC_pcor_GM_all_WS_SlicesPooled(w1,w2) = stats.tstat; clear stats
        if WM_flag == 1
        [h_SFC_pcor_WM_all_WS_SlicesPooled(w1,w2),p_SFC_pcor_WM_all_WS_SlicesPooled(w1,w2),~,stats] = ttest(reshape(squeeze(SFC_pcor_WM_all(w1,w2,:,:)),[],1),0,'Alpha',alpha);
        tstat_SFC_pcor_WM_all_WS_SlicesPooled(w1,w2) = stats.tstat; clear stats
        end
    end
end

elseif ttest_type==2 % paired t-test
    if sum(abs(indx_group1-indx_group2))==0
        f=0;
        for k=1:length(sub_dir)
            if ismember(k,indx_group1)
                f=f+1;
                if WM_flag == 1
                    load([sub_dir{k} ip_file{k,1} '_SFC_withinslice_pcor' '.mat'], 'SFC_pcor_GM_Zval', 'SFC_pcor_WM_Zval')
                    SFC_pcor_GM_all_group1(:,:,:,f) = SFC_pcor_GM_Zval; clear SFC_pcor_GM_Zval
                    SFC_pcor_WM_all_group1(:,:,:,f) = SFC_pcor_WM_Zval; clear SFC_pcor_WM_Zval
                    load([sub_dir{k} ip_file{k,2} '_SFC_withinslice_pcor' '.mat'], 'SFC_pcor_GM_Zval', 'SFC_pcor_WM_Zval')
                    SFC_pcor_GM_all_group2(:,:,:,f) = SFC_pcor_GM_Zval; clear SFC_pcor_GM_Zval
                    SFC_pcor_WM_all_group2(:,:,:,f) = SFC_pcor_WM_Zval; clear SFC_pcor_WM_Zval
                else
                    load([sub_dir{k} ip_file{k,1} '_SFC_withinslice_pcor' '.mat'], 'SFC_pcor_GM_Zval')
                    SFC_pcor_GM_all_group1(:,:,:,f) = SFC_pcor_GM_Zval; clear SFC_pcor_GM_Zval
                    load([sub_dir{k} ip_file{k,2} '_SFC_withinslice_pcor' '.mat'], 'SFC_pcor_GM_Zval')
                    SFC_pcor_GM_all_group2(:,:,:,f) = SFC_pcor_GM_Zval; clear SFC_pcor_GM_Zval
                end
            end
        end; clear f k k2
        for w1=1:size(SFC_pcor_GM_all_group1,1)
            for w2=1:size(SFC_pcor_GM_all_group1,2)
                for w3=1:size(SFC_pcor_GM_all_group1,3)
                    [h_SFC_pcor_GM_all_WS(w1,w2,w3),p_SFC_pcor_GM_all_WS(w1,w2,w3),~,stats] = ttest(squeeze(SFC_pcor_GM_all_group1(w1,w2,w3,:)),squeeze(SFC_pcor_GM_all_group2(w1,w2,w3,:)),'Alpha',alpha);
                    tstat_SFC_pcor_GM_all_WS(w1,w2,w3) = stats.tstat; clear stats
                    if WM_flag == 1
                    [h_SFC_pcor_WM_all_WS(w1,w2,w3),p_SFC_pcor_WM_all_WS(w1,w2,w3),~,stats] = ttest(squeeze(SFC_pcor_WM_all_group1(w1,w2,w3,:)),squeeze(SFC_pcor_WM_all_group2(w1,w2,w3,:)),'Alpha',alpha);
                    tstat_SFC_pcor_WM_all_WS(w1,w2,w3) = stats.tstat; clear stats
                    end
                end
                [h_SFC_pcor_GM_all_WS_SlicesPooled(w1,w2),p_SFC_pcor_GM_all_WS_SlicesPooled(w1,w2),~,stats] = ttest(reshape(squeeze(SFC_pcor_GM_all_group1(w1,w2,:,:)),squeeze(SFC_pcor_GM_all_group2(w1,w2,:,:)),[],1),'Alpha',alpha);
                tstat_SFC_pcor_GM_all_WS_SlicesPooled(w1,w2) = stats.tstat; clear stats
                if WM_flag == 1
                [h_SFC_pcor_WM_all_WS_SlicesPooled(w1,w2),p_SFC_pcor_WM_all_WS_SlicesPooled(w1,w2),~,stats] = ttest(reshape(squeeze(SFC_pcor_WM_all_group1(w1,w2,:,:)),squeeze(SFC_pcor_WM_all_group2(w1,w2,:,:)),[],1),'Alpha',alpha);
                tstat_SFC_pcor_WM_all_WS_SlicesPooled(w1,w2) = stats.tstat; clear stats
                end
            end
        end
        
    else
        f=0;
        for k=1:length(sub_dir)
            if ismember(k,indx_group1)
                f=f+1;
                for k2=1:size(func_files,2)
                    if WM_flag == 1
                        load([sub_dir{k} ip_file{k,k2} '_SFC_withinslice_pcor' '.mat'], 'SFC_pcor_GM_Zval', 'SFC_pcor_WM_Zval')
                        tempg(:,:,:,k2) = SFC_pcor_GM_Zval; clear SFC_pcor_GM_Zval
                        tempw(:,:,:,k2) = SFC_pcor_WM_Zval; clear SFC_pcor_WM_Zval
                    else
                        load([sub_dir{k} ip_file{k,k2} '_SFC_withinslice_pcor' '.mat'], 'SFC_pcor_GM_Zval')
                        tempg(:,:,:,k2) = SFC_pcor_GM_Zval; clear SFC_pcor_GM_Zval
                    end
                end
                SFC_pcor_GM_all_group1(:,:,:,f) = mean(tempg,4);
                if WM_flag == 1
                SFC_pcor_WM_all_group1(:,:,:,f) = mean(tempw,4);
                end
                clear tempg tempw
            end
        end; clear f k k2
        f=0;
        for k=1:length(sub_dir)
            if ismember(k,indx_group2)
                f=f+1;
                for k2=1:size(func_files,2)
                    if WM_flag == 1
                        load([sub_dir{k} ip_file{k,k2} '_SFC_withinslice_pcor' '.mat'], 'SFC_pcor_GM_Zval', 'SFC_pcor_WM_Zval')
                        tempg(:,:,:,k2) = SFC_pcor_GM_Zval; clear SFC_pcor_GM_Zval
                        tempw(:,:,:,k2) = SFC_pcor_WM_Zval; clear SFC_pcor_WM_Zval
                    else
                        load([sub_dir{k} ip_file{k,k2} '_SFC_withinslice_pcor' '.mat'], 'SFC_pcor_GM_Zval')
                        tempg(:,:,:,k2) = SFC_pcor_GM_Zval; clear SFC_pcor_GM_Zval
                    end
                end
                SFC_pcor_GM_all_group2(:,:,:,f) = mean(tempg,4);
                if WM_flag == 1
                SFC_pcor_WM_all_group2(:,:,:,f) = mean(tempw,4);
                end
                clear tempg tempw
            end
        end; clear f k k2
        for w1=1:size(SFC_pcor_GM_all_group1,1)
            for w2=1:size(SFC_pcor_GM_all_group1,2)
                for w3=1:size(SFC_pcor_GM_all_group1,3)
                    [h_SFC_pcor_GM_all_WS(w1,w2,w3),p_SFC_pcor_GM_all_WS(w1,w2,w3),~,stats] = ttest(squeeze(SFC_pcor_GM_all_group1(w1,w2,w3,:)),squeeze(SFC_pcor_GM_all_group2(w1,w2,w3,:)),'Alpha',alpha);
                    tstat_SFC_pcor_GM_all_WS(w1,w2,w3) = stats.tstat; clear stats
                    if WM_flag == 1
                    [h_SFC_pcor_WM_all_WS(w1,w2,w3),p_SFC_pcor_WM_all_WS(w1,w2,w3),~,stats] = ttest(squeeze(SFC_pcor_WM_all_group1(w1,w2,w3,:)),squeeze(SFC_pcor_WM_all_group2(w1,w2,w3,:)),'Alpha',alpha);
                    tstat_SFC_pcor_WM_all_WS(w1,w2,w3) = stats.tstat; clear stats
                    end
                end
                [h_SFC_pcor_GM_all_WS_SlicesPooled(w1,w2),p_SFC_pcor_GM_all_WS_SlicesPooled(w1,w2),~,stats] = ttest(reshape(squeeze(SFC_pcor_GM_all_group1(w1,w2,:,:)),squeeze(SFC_pcor_GM_all_group2(w1,w2,:,:)),[],1),'Alpha',alpha);
                tstat_SFC_pcor_GM_all_WS_SlicesPooled(w1,w2) = stats.tstat; clear stats
                if WM_flag == 1
                [h_SFC_pcor_WM_all_WS_SlicesPooled(w1,w2),p_SFC_pcor_WM_all_WS_SlicesPooled(w1,w2),~,stats] = ttest(reshape(squeeze(SFC_pcor_WM_all_group1(w1,w2,:,:)),squeeze(SFC_pcor_WM_all_group2(w1,w2,:,:)),[],1),'Alpha',alpha);
                tstat_SFC_pcor_WM_all_WS_SlicesPooled(w1,w2) = stats.tstat; clear stats
                end
            end
        end
    end
    
elseif ttest_type==3 % two-sample t-test
    f=0;
    for k=1:length(sub_dir)
        if ismember(k,indx_group1)
            f=f+1;
            for k2=1:size(func_files,2)
                if WM_flag == 1
                    load([sub_dir{k} ip_file{k,k2} '_SFC_withinslice_pcor' '.mat'], 'SFC_pcor_GM_Zval', 'SFC_pcor_WM_Zval')
                    tempg(:,:,:,k2) = SFC_pcor_GM_Zval; clear SFC_pcor_GM_Zval
                    tempw(:,:,:,k2) = SFC_pcor_WM_Zval; clear SFC_pcor_WM_Zval
                else
                    load([sub_dir{k} ip_file{k,k2} '_SFC_withinslice_pcor' '.mat'], 'SFC_pcor_GM_Zval')
                    tempg(:,:,:,k2) = SFC_pcor_GM_Zval; clear SFC_pcor_GM_Zval
                end
            end
            SFC_pcor_GM_all_group1(:,:,:,f) = mean(tempg,4);
            if WM_flag == 1
            SFC_pcor_WM_all_group1(:,:,:,f) = mean(tempw,4);
            end
            clear tempg tempw
        end
    end; clear f k k2
    f=0;
    for k=1:length(sub_dir)
        if ismember(k,indx_group2)
            f=f+1;
            for k2=1:size(func_files,2)
                if WM_flag == 1
                    load([sub_dir{k} ip_file{k,k2} '_SFC_withinslice_pcor' '.mat'], 'SFC_pcor_GM_Zval', 'SFC_pcor_WM_Zval')
                    tempg(:,:,:,k2) = SFC_pcor_GM_Zval; clear SFC_pcor_GM_Zval
                    tempw(:,:,:,k2) = SFC_pcor_WM_Zval; clear SFC_pcor_WM_Zval
                else
                    load([sub_dir{k} ip_file{k,k2} '_SFC_withinslice_pcor' '.mat'], 'SFC_pcor_GM_Zval')
                    tempg(:,:,:,k2) = SFC_pcor_GM_Zval; clear SFC_pcor_GM_Zval
                end
            end
            SFC_pcor_GM_all_group2(:,:,:,f) = mean(tempg,4);
            if WM_flag == 1
            SFC_pcor_WM_all_group2(:,:,:,f) = mean(tempw,4);
            end
            clear tempg tempw
        end
    end; clear f k k2
    for w1=1:size(SFC_pcor_GM_all_group1,1)
        for w2=1:size(SFC_pcor_GM_all_group1,2)
            for w3=1:size(SFC_pcor_GM_all_group1,3)
                [h_SFC_pcor_GM_all_WS(w1,w2,w3),p_SFC_pcor_GM_all_WS(w1,w2,w3),~,stats] = ttest2(squeeze(SFC_pcor_GM_all_group1(w1,w2,w3,:)),squeeze(SFC_pcor_GM_all_group2(w1,w2,w3,:)),'Alpha',alpha);
                tstat_SFC_pcor_GM_all_WS(w1,w2,w3) = stats.tstat; clear stats
                if WM_flag == 1
                [h_SFC_pcor_WM_all_WS(w1,w2,w3),p_SFC_pcor_WM_all_WS(w1,w2,w3),~,stats] = ttest2(squeeze(SFC_pcor_WM_all_group1(w1,w2,w3,:)),squeeze(SFC_pcor_WM_all_group2(w1,w2,w3,:)),'Alpha',alpha);
                tstat_SFC_pcor_WM_all_WS(w1,w2,w3) = stats.tstat; clear stats
                end
            end
            [h_SFC_pcor_GM_all_WS_SlicesPooled(w1,w2),p_SFC_pcor_GM_all_WS_SlicesPooled(w1,w2),~,stats] = ttest2(reshape(squeeze(SFC_pcor_GM_all_group1(w1,w2,:,:)),squeeze(SFC_pcor_GM_all_group2(w1,w2,:,:)),[],1),'Alpha',alpha);
            tstat_SFC_pcor_GM_all_WS_SlicesPooled(w1,w2) = stats.tstat; clear stats
            if WM_flag == 1
            [h_SFC_pcor_WM_all_WS_SlicesPooled(w1,w2),p_SFC_pcor_WM_all_WS_SlicesPooled(w1,w2),~,stats] = ttest2(reshape(squeeze(SFC_pcor_WM_all_group1(w1,w2,:,:)),squeeze(SFC_pcor_WM_all_group2(w1,w2,:,:)),[],1),'Alpha',alpha);
            tstat_SFC_pcor_WM_all_WS_SlicesPooled(w1,w2) = stats.tstat; clear stats
            end
        end
    end
end

if exist('SFC_pcor_GM_all','var')
if WM_flag == 1
save([home_dir 'stats/' 'stats' '_SFC_withinslice_pcor_' clk_stats '.mat'],'SFC_pcor_GM_all','SFC_pcor_WM_all','h_SFC_pcor_GM_all_WS','p_SFC_pcor_GM_all_WS','tstat_SFC_pcor_GM_all_WS', 'h_SFC_pcor_WM_all_WS','p_SFC_pcor_WM_all_WS','tstat_SFC_pcor_WM_all_WS','h_SFC_pcor_GM_all_WS_SlicesPooled','p_SFC_pcor_GM_all_WS_SlicesPooled','tstat_SFC_pcor_GM_all_WS_SlicesPooled','h_SFC_pcor_WM_all_WS_SlicesPooled','p_SFC_pcor_WM_all_WS_SlicesPooled','tstat_SFC_pcor_WM_all_WS_SlicesPooled', 'ip_file','indx_group1','indx_group2','ttest_type')
else
save([home_dir 'stats/' 'stats' '_SFC_withinslice_pcor_' clk_stats '.mat'],'SFC_pcor_GM_all','h_SFC_pcor_GM_all_WS','p_SFC_pcor_GM_all_WS','tstat_SFC_pcor_GM_all_WS','h_SFC_pcor_GM_all_WS_SlicesPooled','p_SFC_pcor_GM_all_WS_SlicesPooled','tstat_SFC_pcor_GM_all_WS_SlicesPooled', 'ip_file','indx_group1','indx_group2','ttest_type')
end
end
if exist('SFC_pcor_GM_all_group1','var')
if WM_flag == 1
save([home_dir 'stats/' 'stats' '_SFC_withinslice_pcor_' clk_stats '.mat'],'SFC_pcor_GM_all_group1','SFC_pcor_GM_all_group2','SFC_pcor_WM_all_group1','SFC_pcor_WM_all_group2','h_SFC_pcor_GM_all_WS','p_SFC_pcor_GM_all_WS','tstat_SFC_pcor_GM_all_WS', 'h_SFC_pcor_WM_all_WS','p_SFC_pcor_WM_all_WS','tstat_SFC_pcor_WM_all_WS','h_SFC_pcor_GM_all_WS_SlicesPooled','p_SFC_pcor_GM_all_WS_SlicesPooled','tstat_SFC_pcor_GM_all_WS_SlicesPooled','h_SFC_pcor_WM_all_WS_SlicesPooled','p_SFC_pcor_WM_all_WS_SlicesPooled','tstat_SFC_pcor_WM_all_WS_SlicesPooled', 'ip_file','indx_group1','indx_group2','ttest_type')
else
save([home_dir 'stats/' 'stats' '_SFC_withinslice_pcor_' clk_stats '.mat'],'SFC_pcor_GM_all_group1','SFC_pcor_GM_all_group2','h_SFC_pcor_GM_all_WS','p_SFC_pcor_GM_all_WS','tstat_SFC_pcor_GM_all_WS','h_SFC_pcor_GM_all_WS_SlicesPooled','p_SFC_pcor_GM_all_WS_SlicesPooled','tstat_SFC_pcor_GM_all_WS_SlicesPooled', 'ip_file','indx_group1','indx_group2','ttest_type')
end
end
clear SFC_pcor_GM_all SFC_pcor_WM_all SFC_pcor_GM_all_group1 SFC_pcor_GM_all_group2 SFC_pcor_WM_all_group1 SFC_pcor_WM_all_group2
end


if SFC_betweenslice_flag==1
    
if ttest_type==1 % one-sample t-test
f=0;
for k=1:length(sub_dir)
    if ismember(k,indx_group1)
    f=f+1;
    for k2=1:size(func_files,2)
        if WM_flag == 1
            load([sub_dir{k} ip_file{k,k2} '_SFC_betweenslice_pcor' '.mat'], 'SFC_pcor_GM_Zval_mat', 'SFC_pcor_WM_Zval_mat')
            tempg(:,:,k2) = SFC_pcor_GM_Zval_mat; clear SFC_pcor_GM_Zval_mat
            tempw(:,:,k2) = SFC_pcor_WM_Zval_mat; clear SFC_pcor_WM_Zval_mat
        else
            load([sub_dir{k} ip_file{k,k2} '_SFC_betweenslice_pcor' '.mat'], 'SFC_pcor_GM_Zval_mat')
            tempg(:,:,k2) = SFC_pcor_GM_Zval_mat; clear SFC_pcor_GM_Zval_mat
        end
    end
    SFC_pcor_GM_all(:,:,f) = mean(tempg,3);
    if WM_flag == 1
    SFC_pcor_WM_all(:,:,f) = mean(tempw,3);
    end
    clear tempg tempw
    end
end; clear f k k2
for w1=1:size(SFC_pcor_GM_all,1)
    for w2=1:size(SFC_pcor_GM_all,2)
        [h_SFC_pcor_GM_all_BS(w1,w2),p_SFC_pcor_GM_all_BS(w1,w2),~,stats] = ttest(squeeze(SFC_pcor_GM_all(w1,w2,:)),0,'Alpha',alpha);
        tstat_SFC_pcor_GM_all_BS(w1,w2) = stats.tstat; clear stats
        if WM_flag == 1
        [h_SFC_pcor_WM_all_BS(w1,w2),p_SFC_pcor_WM_all_BS(w1,w2),~,stats] = ttest(squeeze(SFC_pcor_WM_all(w1,w2,:)),0,'Alpha',alpha);
        tstat_SFC_pcor_WM_all_BS(w1,w2) = stats.tstat; clear stats
        end
    end
end

elseif ttest_type==2 % paired t-test
    if sum(abs(indx_group1-indx_group2))==0
        f=0;
        for k=1:length(sub_dir)
            if ismember(k,indx_group1)
                f=f+1;
                if WM_flag == 1
                    load([sub_dir{k} ip_file{k,1} '_SFC_betweenslice_pcor' '.mat'], 'SFC_pcor_GM_Zval_mat', 'SFC_pcor_WM_Zval_mat')
                    SFC_pcor_GM_all_group1(:,:,f) = SFC_pcor_GM_Zval_mat; clear SFC_pcor_GM_Zval_mat
                    SFC_pcor_WM_all_group1(:,:,f) = SFC_pcor_WM_Zval_mat; clear SFC_pcor_WM_Zval_mat
                    load([sub_dir{k} ip_file{k,2} '_SFC_betweenslice_pcor' '.mat'], 'SFC_pcor_GM_Zval_mat', 'SFC_pcor_WM_Zval_mat')
                    SFC_pcor_GM_all_group2(:,:,f) = SFC_pcor_GM_Zval_mat; clear SFC_pcor_GM_Zval_mat
                    SFC_pcor_WM_all_group2(:,:,f) = SFC_pcor_WM_Zval_mat; clear SFC_pcor_WM_Zval_mat
                else
                    load([sub_dir{k} ip_file{k,1} '_SFC_betweenslice_pcor' '.mat'], 'SFC_pcor_GM_Zval_mat')
                    SFC_pcor_GM_all_group1(:,:,f) = SFC_pcor_GM_Zval_mat; clear SFC_pcor_GM_Zval_mat
                    load([sub_dir{k} ip_file{k,2} '_SFC_betweenslice_pcor' '.mat'], 'SFC_pcor_GM_Zval_mat')
                    SFC_pcor_GM_all_group2(:,:,f) = SFC_pcor_GM_Zval_mat; clear SFC_pcor_GM_Zval_mat
                end
            end
        end; clear f k k2
        for w1=1:size(SFC_pcor_GM_all_group1,1)
            for w2=1:size(SFC_pcor_GM_all_group1,2)
                [h_SFC_pcor_GM_all_BS(w1,w2),p_SFC_pcor_GM_all_BS(w1,w2),~,stats] = ttest(squeeze(SFC_pcor_GM_all_group1(w1,w2,:)),squeeze(SFC_pcor_GM_all_group2(w1,w2,:)),'Alpha',alpha);
                tstat_SFC_pcor_GM_all_BS(w1,w2) = stats.tstat; clear stats
                if WM_flag == 1
                [h_SFC_pcor_WM_all_BS(w1,w2),p_SFC_pcor_WM_all_BS(w1,w2),~,stats] = ttest(squeeze(SFC_pcor_WM_all_group1(w1,w2,:)),squeeze(SFC_pcor_WM_all_group2(w1,w2,:)),'Alpha',alpha);
                tstat_SFC_pcor_WM_all_BS(w1,w2) = stats.tstat; clear stats
                end
            end
        end
        
    else
        f=0;
        for k=1:length(sub_dir)
            if ismember(k,indx_group1)
                f=f+1;
                for k2=1:size(func_files,2)
                    if WM_flag == 1
                        load([sub_dir{k} ip_file{k,k2} '_SFC_betweenslice_pcor' '.mat'], 'SFC_pcor_GM_Zval_mat', 'SFC_pcor_WM_Zval_mat')
                        tempg(:,:,k2) = SFC_pcor_GM_Zval_mat; clear SFC_pcor_GM_Zval_mat
                        tempw(:,:,k2) = SFC_pcor_WM_Zval_mat; clear SFC_pcor_WM_Zval_mat
                    else
                        load([sub_dir{k} ip_file{k,k2} '_SFC_betweenslice_pcor' '.mat'], 'SFC_pcor_GM_Zval_mat')
                        tempg(:,:,k2) = SFC_pcor_GM_Zval_mat; clear SFC_pcor_GM_Zval_mat
                    end
                end
                SFC_pcor_GM_all_group1(:,:,f) = mean(tempg,3);
                if WM_flag == 1
                SFC_pcor_WM_all_group1(:,:,f) = mean(tempw,3);
                end
                clear tempg tempw
            end
        end; clear f k k2
        f=0;
        for k=1:length(sub_dir)
            if ismember(k,indx_group2)
                f=f+1;
                for k2=1:size(func_files,2)
                    if WM_flag == 1
                        load([sub_dir{k} ip_file{k,k2} '_SFC_betweenslice_pcor' '.mat'], 'SFC_pcor_GM_Zval_mat', 'SFC_pcor_WM_Zval_mat')
                        tempg(:,:,k2) = SFC_pcor_GM_Zval_mat; clear SFC_pcor_GM_Zval_mat
                        tempw(:,:,k2) = SFC_pcor_WM_Zval_mat; clear SFC_pcor_WM_Zval_mat
                    else
                        load([sub_dir{k} ip_file{k,k2} '_SFC_betweenslice_pcor' '.mat'], 'SFC_pcor_GM_Zval_mat')
                        tempg(:,:,k2) = SFC_pcor_GM_Zval_mat; clear SFC_pcor_GM_Zval_mat
                    end
                end
                SFC_pcor_GM_all_group2(:,:,f) = mean(tempg,3);
                if WM_flag == 1
                SFC_pcor_WM_all_group2(:,:,f) = mean(tempw,3);
                end
                clear tempg tempw
            end
        end; clear f k k2
        for w1=1:size(SFC_pcor_GM_all_group1,1)
            for w2=1:size(SFC_pcor_GM_all_group1,2)
                [h_SFC_pcor_GM_all_BS(w1,w2),p_SFC_pcor_GM_all_BS(w1,w2),~,stats] = ttest(squeeze(SFC_pcor_GM_all_group1(w1,w2,:)),squeeze(SFC_pcor_GM_all_group2(w1,w2,:)),'Alpha',alpha);
                tstat_SFC_pcor_GM_all_BS(w1,w2) = stats.tstat; clear stats
                if WM_flag == 1
                [h_SFC_pcor_WM_all_BS(w1,w2),p_SFC_pcor_WM_all_BS(w1,w2),~,stats] = ttest(squeeze(SFC_pcor_WM_all_group1(w1,w2,:)),squeeze(SFC_pcor_WM_all_group2(w1,w2,:)),'Alpha',alpha);
                tstat_SFC_pcor_WM_all_BS(w1,w2) = stats.tstat; clear stats
                end
            end
        end
    end
    
elseif ttest_type==3 % two-sample t-test
    f=0;
    for k=1:length(sub_dir)
        if ismember(k,indx_group1)
            f=f+1;
            for k2=1:size(func_files,2)
                if WM_flag == 1
                    load([sub_dir{k} ip_file{k,k2} '_SFC_betweenslice_pcor' '.mat'], 'SFC_pcor_GM_Zval_mat', 'SFC_pcor_WM_Zval_mat')
                    tempg(:,:,k2) = SFC_pcor_GM_Zval_mat; clear SFC_pcor_GM_Zval_mat
                    tempw(:,:,k2) = SFC_pcor_WM_Zval_mat; clear SFC_pcor_WM_Zval_mat
                else
                    load([sub_dir{k} ip_file{k,k2} '_SFC_betweenslice_pcor' '.mat'], 'SFC_pcor_GM_Zval_mat')
                    tempg(:,:,k2) = SFC_pcor_GM_Zval_mat; clear SFC_pcor_GM_Zval_mat
                end
            end
            SFC_pcor_GM_all_group1(:,:,f) = mean(tempg,3);
            if WM_flag == 1
            SFC_pcor_WM_all_group1(:,:,f) = mean(tempw,3);
            end
            clear tempg tempw
        end
    end; clear f k k2
    f=0;
    for k=1:length(sub_dir)
        if ismember(k,indx_group2)
            f=f+1;
            for k2=1:size(func_files,2)
                if WM_flag == 1
                    load([sub_dir{k} ip_file{k,k2} '_SFC_betweenslice_pcor' '.mat'], 'SFC_pcor_GM_Zval_mat', 'SFC_pcor_WM_Zval_mat')
                    tempg(:,:,k2) = SFC_pcor_GM_Zval_mat; clear SFC_pcor_GM_Zval_mat
                    tempw(:,:,k2) = SFC_pcor_WM_Zval_mat; clear SFC_pcor_WM_Zval_mat
                else
                    load([sub_dir{k} ip_file{k,k2} '_SFC_betweenslice_pcor' '.mat'], 'SFC_pcor_GM_Zval_mat')
                    tempg(:,:,k2) = SFC_pcor_GM_Zval_mat; clear SFC_pcor_GM_Zval_mat
                end
            end
            SFC_pcor_GM_all_group2(:,:,f) = mean(tempg,3);
            if WM_flag == 1
            SFC_pcor_WM_all_group2(:,:,f) = mean(tempw,3);
            end
            clear tempg tempw
        end
    end; clear f k k2
    for w1=1:size(SFC_pcor_GM_all_group1,1)
        for w2=1:size(SFC_pcor_GM_all_group1,2)
            [h_SFC_pcor_GM_all_BS(w1,w2),p_SFC_pcor_GM_all_BS(w1,w2),~,stats] = ttest2(squeeze(SFC_pcor_GM_all_group1(w1,w2,:)),squeeze(SFC_pcor_GM_all_group2(w1,w2,:)),'Alpha',alpha);
            tstat_SFC_pcor_GM_all_BS(w1,w2) = stats.tstat; clear stats
            if WM_flag == 1
            [h_SFC_pcor_WM_all_BS(w1,w2),p_SFC_pcor_WM_all_BS(w1,w2),~,stats] = ttest2(squeeze(SFC_pcor_WM_all_group1(w1,w2,:)),squeeze(SFC_pcor_WM_all_group2(w1,w2,:)),'Alpha',alpha);
            tstat_SFC_pcor_WM_all_BS(w1,w2) = stats.tstat; clear stats
            end
        end
    end
end

if exist('SFC_pcor_GM_all','var')
if WM_flag == 1
save([home_dir 'stats/' 'stats' '_SFC_betweenslice_pcor_' clk_stats '.mat'],'SFC_pcor_GM_all','SFC_pcor_WM_all','h_SFC_pcor_GM_all_BS','p_SFC_pcor_GM_all_BS','tstat_SFC_pcor_GM_all_BS', 'h_SFC_pcor_WM_all_BS','p_SFC_pcor_WM_all_BS','tstat_SFC_pcor_WM_all_BS','ip_file','indx_group1','indx_group2','ttest_type')
else
save([home_dir 'stats/' 'stats' '_SFC_betweenslice_pcor_' clk_stats '.mat'],'SFC_pcor_GM_all','h_SFC_pcor_GM_all_BS','p_SFC_pcor_GM_all_BS','tstat_SFC_pcor_GM_all_BS', 'ip_file','indx_group1','indx_group2','ttest_type')
end
end
if exist('SFC_pcor_GM_all_group1','var')
if WM_flag == 1
save([home_dir 'stats/' 'stats' '_SFC_betweenslice_pcor_' clk_stats '.mat'],'SFC_pcor_GM_all_group1','SFC_pcor_GM_all_group2','SFC_pcor_WM_all_group1','SFC_pcor_WM_all_group2','h_SFC_pcor_GM_all_BS','p_SFC_pcor_GM_all_BS','tstat_SFC_pcor_GM_all_BS', 'h_SFC_pcor_WM_all_BS','p_SFC_pcor_WM_all_BS','tstat_SFC_pcor_WM_all_BS','ip_file','indx_group1','indx_group2','ttest_type')
else
save([home_dir 'stats/' 'stats' '_SFC_betweenslice_pcor_' clk_stats '.mat'],'SFC_pcor_GM_all_group1','SFC_pcor_GM_all_group2','h_SFC_pcor_GM_all_BS','p_SFC_pcor_GM_all_BS','tstat_SFC_pcor_GM_all_BS', 'ip_file','indx_group1','indx_group2','ttest_type')
end
end
clear SFC_pcor_GM_all SFC_pcor_WM_all SFC_pcor_GM_all_group1 SFC_pcor_GM_all_group2 SFC_pcor_WM_all_group1 SFC_pcor_WM_all_group2
end

end

%% fALFF stats
if ((choices_postproc(3)==1)||exist([sub_dir{1} ip_file2{1,1} '_fALFF' '.mat'],'file'))
    
if ttest_type==1 % one-sample t-test
f=0;
for k=1:length(sub_dir)
    if ismember(k,indx_group1)
    f=f+1;
    for k2=1:size(func_files,2)
        if WM_flag == 1
            load([sub_dir{k} ip_file2{k,k2} '_fALFF' '.mat'], 'fALFF_GM', 'fALFF_WM')
            tempg(:,:,k2) = fALFF_GM; clear fALFF_GM
            tempw(:,:,k2) = fALFF_WM; clear fALFF_WM
        else
            load([sub_dir{k} ip_file2{k,k2} '_fALFF' '.mat'], 'fALFF_GM')
            tempg(:,:,k2) = fALFF_GM; clear fALFF_GM
        end
    end
    fALFF_GM_all(:,:,f) = mean(tempg,3);
    if WM_flag == 1
    fALFF_WM_all(:,:,f) = mean(tempw,3);
    end
    clear tempg tempw
    end
end; clear f k k2
for w1=1:size(fALFF_GM_all,1)
    for w2=1:size(fALFF_GM_all,2)
        [h_fALFF_GM_all(w1,w2),p_fALFF_GM_all(w1,w2),~,stats] = ttest(squeeze(fALFF_GM_all(w1,w2,:)),0,'Alpha',alpha);
        tstat_fALFF_GM_all(w1,w2) = stats.tstat; clear stats
        if WM_flag == 1
        [h_fALFF_WM_all(w1,w2),p_fALFF_WM_all(w1,w2),~,stats] = ttest(squeeze(fALFF_WM_all(w1,w2,:)),0,'Alpha',alpha);
        tstat_fALFF_WM_all(w1,w2) = stats.tstat; clear stats
        end
    end
end

elseif ttest_type==2 % paired t-test
    if sum(abs(indx_group1-indx_group2))==0
        f=0;
        for k=1:length(sub_dir)
            if ismember(k,indx_group1)
                f=f+1;
                if WM_flag == 1
                    load([sub_dir{k} ip_file2{k,1} '_fALFF' '.mat'], 'fALFF_GM', 'fALFF_WM')
                    fALFF_GM_all_group1(:,:,f) = fALFF_GM; clear fALFF_GM
                    fALFF_WM_all_group1(:,:,f) = fALFF_WM; clear fALFF_WM
                    load([sub_dir{k} ip_file2{k,2} '_fALFF' '.mat'], 'fALFF_GM', 'fALFF_WM')
                    fALFF_GM_all_group2(:,:,f) = fALFF_GM; clear fALFF_GM
                    fALFF_WM_all_group2(:,:,f) = fALFF_WM; clear fALFF_WM
                else
                    load([sub_dir{k} ip_file2{k,1} '_fALFF' '.mat'], 'fALFF_GM')
                    fALFF_GM_all_group1(:,:,f) = fALFF_GM; clear fALFF_GM
                    load([sub_dir{k} ip_file2{k,2} '_fALFF' '.mat'], 'fALFF_GM')
                    fALFF_GM_all_group2(:,:,f) = fALFF_GM; clear fALFF_GM
                end
            end
        end; clear f k k2
        for w1=1:size(fALFF_GM_all_group1,1)
            for w2=1:size(fALFF_GM_all_group1,2)
                [h_fALFF_GM_all(w1,w2),p_fALFF_GM_all(w1,w2),~,stats] = ttest(squeeze(fALFF_GM_all_group1(w1,w2,:)),squeeze(fALFF_GM_all_group2(w1,w2,:)),'Alpha',alpha);
                tstat_fALFF_GM_all(w1,w2) = stats.tstat; clear stats
                if WM_flag == 1
                [h_fALFF_WM_all(w1,w2),p_fALFF_WM_all(w1,w2),~,stats] = ttest(squeeze(fALFF_WM_all_group1(w1,w2,:)),squeeze(fALFF_WM_all_group2(w1,w2,:)),'Alpha',alpha);
                tstat_fALFF_WM_all(w1,w2) = stats.tstat; clear stats
                end
            end
        end
        
    else
        f=0;
        for k=1:length(sub_dir)
            if ismember(k,indx_group1)
                f=f+1;
                for k2=1:size(func_files,2)
                    if WM_flag == 1
                        load([sub_dir{k} ip_file2{k,k2} '_fALFF' '.mat'], 'fALFF_GM', 'fALFF_WM')
                        tempg(:,:,k2) = fALFF_GM; clear fALFF_GM
                        tempw(:,:,k2) = fALFF_WM; clear fALFF_WM
                    else
                        load([sub_dir{k} ip_file2{k,k2} '_fALFF' '.mat'], 'fALFF_GM')
                        tempg(:,:,k2) = fALFF_GM; clear fALFF_GM
                    end
                end
                fALFF_GM_all_group1(:,:,f) = mean(tempg,3);
                if WM_flag == 1
                fALFF_WM_all_group1(:,:,f) = mean(tempw,3);
                end
                clear tempg tempw
            end
        end; clear f k k2
        f=0;
        for k=1:length(sub_dir)
            if ismember(k,indx_group2)
                f=f+1;
                for k2=1:size(func_files,2)
                    if WM_flag == 1
                        load([sub_dir{k} ip_file2{k,k2} '_fALFF' '.mat'], 'fALFF_GM', 'fALFF_WM')
                        tempg(:,:,k2) = fALFF_GM; clear fALFF_GM
                        tempw(:,:,k2) = fALFF_WM; clear fALFF_WM
                    else
                        load([sub_dir{k} ip_file2{k,k2} '_fALFF' '.mat'], 'fALFF_GM')
                        tempg(:,:,k2) = fALFF_GM; clear fALFF_GM
                    end
                end
                fALFF_GM_all_group2(:,:,f) = mean(tempg,3);
                if WM_flag == 1
                fALFF_WM_all_group2(:,:,f) = mean(tempw,3);
                end
                clear tempg tempw
            end
        end; clear f k k2
        for w1=1:size(fALFF_GM_all_group1,1)
            for w2=1:size(fALFF_GM_all_group1,2)
                [h_fALFF_GM_all(w1,w2),p_fALFF_GM_all(w1,w2),~,stats] = ttest(squeeze(fALFF_GM_all_group1(w1,w2,:)),squeeze(fALFF_GM_all_group2(w1,w2,:)),'Alpha',alpha);
                tstat_fALFF_GM_all(w1,w2) = stats.tstat; clear stats
                if WM_flag == 1
                [h_fALFF_WM_all(w1,w2),p_fALFF_WM_all(w1,w2),~,stats] = ttest(squeeze(fALFF_WM_all_group1(w1,w2,:)),squeeze(fALFF_WM_all_group2(w1,w2,:)),'Alpha',alpha);
                tstat_fALFF_WM_all(w1,w2) = stats.tstat; clear stats
                end
            end
        end
    end
    
elseif ttest_type==3 % two-sample t-test
    f=0;
    for k=1:length(sub_dir)
        if ismember(k,indx_group1)
            f=f+1;
            for k2=1:size(func_files,2)
                if WM_flag == 1
                    load([sub_dir{k} ip_file2{k,k2} '_fALFF' '.mat'], 'fALFF_GM', 'fALFF_WM')
                    tempg(:,:,k2) = fALFF_GM; clear fALFF_GM
                    tempw(:,:,k2) = fALFF_WM; clear fALFF_WM
                else
                    load([sub_dir{k} ip_file2{k,k2} '_fALFF' '.mat'], 'fALFF_GM')
                    tempg(:,:,k2) = fALFF_GM; clear fALFF_GM
                end
            end
            fALFF_GM_all_group1(:,:,f) = mean(tempg,3);
            if WM_flag == 1
            fALFF_WM_all_group1(:,:,f) = mean(tempw,3);
            end
            clear tempg tempw
        end
    end; clear f k k2
    f=0;
    for k=1:length(sub_dir)
        if ismember(k,indx_group2)
            f=f+1;
            for k2=1:size(func_files,2)
                if WM_flag == 1
                    load([sub_dir{k} ip_file2{k,k2} '_fALFF' '.mat'], 'fALFF_GM', 'fALFF_WM')
                    tempg(:,:,k2) = fALFF_GM; clear fALFF_GM
                    tempw(:,:,k2) = fALFF_WM; clear fALFF_WM
                else
                    load([sub_dir{k} ip_file2{k,k2} '_fALFF' '.mat'], 'fALFF_GM')
                    tempg(:,:,k2) = fALFF_GM; clear fALFF_GM
                end
            end
            fALFF_GM_all_group2(:,:,f) = mean(tempg,3);
            if WM_flag == 1
            fALFF_WM_all_group2(:,:,f) = mean(tempw,3);
            end
            clear tempg tempw
        end
    end; clear f k k2
    for w1=1:size(fALFF_GM_all_group1,1)
        for w2=1:size(fALFF_GM_all_group1,2)
            [h_fALFF_GM_all(w1,w2),p_fALFF_GM_all(w1,w2),~,stats] = ttest2(squeeze(fALFF_GM_all_group1(w1,w2,:)),squeeze(fALFF_GM_all_group2(w1,w2,:)),'Alpha',alpha);
            tstat_fALFF_GM_all(w1,w2) = stats.tstat; clear stats
            if WM_flag == 1
            [h_fALFF_WM_all(w1,w2),p_fALFF_WM_all(w1,w2),~,stats] = ttest2(squeeze(fALFF_WM_all_group1(w1,w2,:)),squeeze(fALFF_WM_all_group2(w1,w2,:)),'Alpha',alpha);
            tstat_fALFF_WM_all(w1,w2) = stats.tstat; clear stats
            end
        end
    end
end

if exist('fALFF_GM_all','var')
if WM_flag == 1
save([home_dir 'stats/' 'stats' '_fALFF_' clk_stats '.mat'],'fALFF_GM_all','fALFF_WM_all','h_fALFF_GM_all','p_fALFF_GM_all','tstat_fALFF_GM_all', 'h_fALFF_WM_all','p_fALFF_WM_all','tstat_fALFF_WM_all','ip_file2','indx_group1','indx_group2','ttest_type')
else
save([home_dir 'stats/' 'stats' '_fALFF_' clk_stats '.mat'],'fALFF_GM_all','h_fALFF_GM_all','p_fALFF_GM_all','tstat_fALFF_GM_all', 'ip_file2','indx_group1','indx_group2','ttest_type')
end
end
if exist('fALFF_GM_all_group1','var')
if WM_flag == 1
save([home_dir 'stats/' 'stats' '_fALFF_' clk_stats '.mat'],'fALFF_GM_all_group1','fALFF_GM_all_group2','fALFF_WM_all_group1','fALFF_WM_all_group2','h_fALFF_GM_all','p_fALFF_GM_all','tstat_fALFF_GM_all', 'h_fALFF_WM_all','p_fALFF_WM_all','tstat_fALFF_WM_all','ip_file2','indx_group1','indx_group2','ttest_type')
else
save([home_dir 'stats/' 'stats' '_fALFF_' clk_stats '.mat'],'fALFF_GM_all_group1','fALFF_GM_all_group2','h_fALFF_GM_all','p_fALFF_GM_all','tstat_fALFF_GM_all', 'ip_file2','indx_group1','indx_group2','ttest_type')
end
end
clear fALFF_GM_all fALFF_WM_all fALFF_GM_all_group1 fALFF_GM_all_group2 fALFF_WM_all_group1 fALFF_WM_all_group2

end

%% QC plots of stats
fprintf('23. QC plots: simple stats\n')

%% QC plots of stats: within-slice SFC
if ((choices_postproc(2)==1)||exist([sub_dir{1} ip_file{1,1} '_SFC_withinslice' '.mat'],'file')||exist([sub_dir{1} ip_file{1,1} '_SFC_withinslice_pcor' '.mat'],'file')) && (SFC_withinslice_flag==1)

WhiteMask = ones(4,4); WhiteMask(eye(4)==1) = 0;
for i3=1:siz3
    fprintf('23. QC plots: simple stats - SFC_WithinSlice - slice (%d) of (%d)\n',i3,siz3)
    
    if pearson==1
    Cg = real(tstat_SFC_GM_all_WS(:,:,i3));
    if WM_flag == 0
    fighndl = figure('Position',[1 1 500 500],'Color',[0.95,0.9,0.85],'InvertHardcopy','off','Visible','off'); % red [0.95,0.9,0.85], green [0.85,0.90,0.85], blue [0.89,0.94,0.99]
    imagesc(Cg,'AlphaData',WhiteMask.*h_SFC_GM_all_WS(:,:,i3),[-max(abs(tstat_SFC_GM_all_WS(:)))-10^-30 max(abs(tstat_SFC_GM_all_WS(:)))+10^-30]), axis square
        colormap('jet'),colorbar;
        for v1=1:size(Cg,1), for v2=1:size(Cg,2), if v1~=v2 %#ok<*ALIGN>
        if h_SFC_GM_all_WS(v1,v2,i3)==1
        text(v1,v2,sprintf('%.2f',(Cg(v1,v2))),'HorizontalAlignment','center','VerticalAlignment','middle','fontsize',13','FontWeight','bold','color','k');
        else
        text(v1,v2,sprintf('%.2f',(Cg(v1,v2))),'HorizontalAlignment','center','VerticalAlignment','middle','fontsize',11','color',[0.1,0.1,0.1]);
        end
        end, end, end
        xticks([1:4]), xticklabels({'LV','RV','LD','RD'}), a = get(gca,'xticklabels'); set(gca,'xticklabels',a,'fontsize',13); clear a
        yticks([1:4]), yticklabels({'LV','RV','LD','RD'}), a = get(gca,'yticklabels'); set(gca,'yticklabels',a,'fontsize',13); clear a
        xlabel('Gray matter quadrants'),ylabel('Gray matter quadrants')
        title(sprintf('GRAY MATTER\n(Pearson''s correlation based functional connectivity)\nSignificant group-level T-statistics for within-slice FC (slice %d of %d)',i3,siz3),'Color','blue','FontSize',13)
    else
    Cw = real(tstat_SFC_WM_all_WS(:,:,i3));
    fighndl = figure('Position',[1 1 1000 500],'Color',[0.95,0.9,0.85],'InvertHardcopy','off','Visible','off'); % red [0.95,0.9,0.85], green [0.85,0.90,0.85], blue [0.89,0.94,0.99]
    subplot2n(1,2,1), imagesc(Cg,'AlphaData',WhiteMask.*h_SFC_GM_all_WS(:,:,i3),[-max(abs(tstat_SFC_GM_all_WS(:)))-10^-30 max(abs(tstat_SFC_GM_all_WS(:)))+10^-30]), axis square
        colormap('jet'),colorbar;
        for v1=1:size(Cg,1), for v2=1:size(Cg,2), if v1~=v2 %#ok<*ALIGN>
        if h_SFC_GM_all_WS(v1,v2,i3)==1
        text(v1,v2,sprintf('%.2f',(Cg(v1,v2))),'HorizontalAlignment','center','VerticalAlignment','middle','fontsize',13','FontWeight','bold','color','k');
        else
        text(v1,v2,sprintf('%.2f',(Cg(v1,v2))),'HorizontalAlignment','center','VerticalAlignment','middle','fontsize',11','color',[0.1,0.1,0.1]);
        end
        end, end, end
        xticks([1:4]), xticklabels({'LV','RV','LD','RD'}), a = get(gca,'xticklabels'); set(gca,'xticklabels',a,'fontsize',13); clear a
        yticks([1:4]), yticklabels({'LV','RV','LD','RD'}), a = get(gca,'yticklabels'); set(gca,'yticklabels',a,'fontsize',13); clear a
        xlabel('Gray matter quadrants'),ylabel('Gray matter quadrants')
        title(sprintf('GRAY MATTER\n(Pearson''s correlation based functional connectivity)\nSignificant group-level T-statistics for within-slice FC (slice %d of %d)',i3,siz3),'Color','blue','FontSize',13)
    subplot2n(1,2,2), imagesc(Cw,'AlphaData',WhiteMask.*h_SFC_WM_all_WS(:,:,i3),[-max(abs(tstat_SFC_WM_all_WS(:)))-10^-30 max(abs(tstat_SFC_WM_all_WS(:)))+10^-30]), axis square
        colormap('jet'),colorbar;
        for v1=1:size(Cw,1), for v2=1:size(Cw,2), if v1~=v2 %#ok<*ALIGN>
        if h_SFC_WM_all_WS(v1,v2,i3)==1
        text(v1,v2,sprintf('%.2f',(Cw(v1,v2))),'HorizontalAlignment','center','VerticalAlignment','middle','fontsize',13','FontWeight','bold','color','k');
        else
        text(v1,v2,sprintf('%.2f',(Cw(v1,v2))),'HorizontalAlignment','center','VerticalAlignment','middle','fontsize',11','color',[0.1,0.1,0.1]);
        end
        end, end, end
        xticks([1:4]), xticklabels({'LV','RV','LD','RD'}), a = get(gca,'xticklabels'); set(gca,'xticklabels',a,'fontsize',13); clear a
        yticks([1:4]), yticklabels({'LV','RV','LD','RD'}), a = get(gca,'yticklabels'); set(gca,'yticklabels',a,'fontsize',13); clear a
        xlabel('White matter quadrants'),ylabel('White matter quadrants')
        title(sprintf('WHITE MATTER\n(Pearson''s correlation based functional connectivity)\nSignificant group-level T-statistics for within-slice FC (slice %d of %d)',i3,siz3),'Color','red','FontSize',13)
    end
    saveas(fighndl,[QCpath 'stats_SFC_WithinSlice_slice' sprintf('%.2d',i3) '.jpg'])
    close(fighndl)
    clear fighndl Cg Cw v1 v2
    end
    
    if partial==1
    Cg = real(tstat_SFC_pcor_GM_all_WS(:,:,i3));
    if WM_flag == 0
    fighndl = figure('Position',[1 1 500 500],'Color',[0.95,0.9,0.85],'InvertHardcopy','off','Visible','off'); % red [0.95,0.9,0.85], green [0.85,0.90,0.85], blue [0.89,0.94,0.99]
    imagesc(Cg,'AlphaData',WhiteMask.*h_SFC_pcor_GM_all_WS(:,:,i3),[-max(abs(tstat_SFC_pcor_GM_all_WS(:)))-10^-30 max(abs(tstat_SFC_pcor_GM_all_WS(:)))+10^-30]), axis square
        colormap('jet'),colorbar;
        for v1=1:size(Cg,1), for v2=1:size(Cg,2), if v1~=v2 %#ok<*ALIGN>
        if h_SFC_pcor_GM_all_WS(v1,v2,i3)==1
        text(v1,v2,sprintf('%.2f',(Cg(v1,v2))),'HorizontalAlignment','center','VerticalAlignment','middle','fontsize',13','FontWeight','bold','color','k');
        else
        text(v1,v2,sprintf('%.2f',(Cg(v1,v2))),'HorizontalAlignment','center','VerticalAlignment','middle','fontsize',11','color',[0.1,0.1,0.1]);
        end
        end, end, end
        xticks([1:4]), xticklabels({'LV','RV','LD','RD'}), a = get(gca,'xticklabels'); set(gca,'xticklabels',a,'fontsize',13); clear a
        yticks([1:4]), yticklabels({'LV','RV','LD','RD'}), a = get(gca,'yticklabels'); set(gca,'yticklabels',a,'fontsize',13); clear a
        xlabel('Gray matter quadrants'),ylabel('Gray matter quadrants')
        title(sprintf('GRAY MATTER\n(Partial correlation based functional connectivity)\nSignificant group-level T-statistics for within-slice FC (slice %d of %d)',i3,siz3),'Color','blue','FontSize',13)
    else
    Cw = real(tstat_SFC_pcor_WM_all_WS(:,:,i3));
    fighndl = figure('Position',[1 1 1000 500],'Color',[0.95,0.9,0.85],'InvertHardcopy','off','Visible','off'); % red [0.95,0.9,0.85], green [0.85,0.90,0.85], blue [0.89,0.94,0.99]
    subplot2n(1,2,1), imagesc(Cg,'AlphaData',WhiteMask.*h_SFC_pcor_GM_all_WS(:,:,i3),[-max(abs(tstat_SFC_pcor_GM_all_WS(:)))-10^-30 max(abs(tstat_SFC_pcor_GM_all_WS(:)))+10^-30]), axis square
        colormap('jet'),colorbar;
        for v1=1:size(Cg,1), for v2=1:size(Cg,2), if v1~=v2 %#ok<*ALIGN>
        if h_SFC_pcor_GM_all_WS(v1,v2,i3)==1
        text(v1,v2,sprintf('%.2f',(Cg(v1,v2))),'HorizontalAlignment','center','VerticalAlignment','middle','fontsize',13','FontWeight','bold','color','k');
        else
        text(v1,v2,sprintf('%.2f',(Cg(v1,v2))),'HorizontalAlignment','center','VerticalAlignment','middle','fontsize',11','color',[0.1,0.1,0.1]);
        end
        end, end, end
        xticks([1:4]), xticklabels({'LV','RV','LD','RD'}), a = get(gca,'xticklabels'); set(gca,'xticklabels',a,'fontsize',13); clear a
        yticks([1:4]), yticklabels({'LV','RV','LD','RD'}), a = get(gca,'yticklabels'); set(gca,'yticklabels',a,'fontsize',13); clear a
        xlabel('Gray matter quadrants'),ylabel('Gray matter quadrants')
        title(sprintf('GRAY MATTER\n(Partial correlation based functional connectivity)\nSignificant group-level T-statistics for within-slice FC (slice %d of %d)',i3,siz3),'Color','blue','FontSize',13)
    subplot2n(1,2,2), imagesc(Cw,'AlphaData',WhiteMask.*h_SFC_pcor_WM_all_WS(:,:,i3),[-max(abs(tstat_SFC_pcor_WM_all_WS(:)))-10^-30 max(abs(tstat_SFC_pcor_WM_all_WS(:)))+10^-30]), axis square
        colormap('jet'),colorbar;
        for v1=1:size(Cw,1), for v2=1:size(Cw,2), if v1~=v2 %#ok<*ALIGN>
        if h_SFC_pcor_WM_all_WS(v1,v2,i3)==1
        text(v1,v2,sprintf('%.2f',(Cw(v1,v2))),'HorizontalAlignment','center','VerticalAlignment','middle','fontsize',13','FontWeight','bold','color','k');
        else
        text(v1,v2,sprintf('%.2f',(Cw(v1,v2))),'HorizontalAlignment','center','VerticalAlignment','middle','fontsize',11','color',[0.1,0.1,0.1]);
        end
        end, end, end
        xticks([1:4]), xticklabels({'LV','RV','LD','RD'}), a = get(gca,'xticklabels'); set(gca,'xticklabels',a,'fontsize',13); clear a
        yticks([1:4]), yticklabels({'LV','RV','LD','RD'}), a = get(gca,'yticklabels'); set(gca,'yticklabels',a,'fontsize',13); clear a
        xlabel('White matter quadrants'),ylabel('White matter quadrants')
        title(sprintf('WHITE MATTER\n(Partial correlation based functional connectivity)\nSignificant group-level T-statistics for within-slice FC (slice %d of %d)',i3,siz3),'Color','red','FontSize',13)
    end
    saveas(fighndl,[QCpath 'stats_SFC_pcor_WithinSlice_slice' sprintf('%.2d',i3) '.jpg'])
    close(fighndl)
    clear fighndl Cg Cw v1 v2
    end
end

    if pearson==1
    Cg = real(tstat_SFC_GM_all_WS_SlicesPooled);
    if WM_flag == 0
    fighndl = figure('Position',[1 1 500 500],'Color',[0.89,0.94,0.99],'InvertHardcopy','off','Visible','off'); % red [0.95,0.9,0.85], green [0.85,0.90,0.85], blue [0.89,0.94,0.99]
    imagesc(Cg,'AlphaData',WhiteMask.*h_SFC_GM_all_WS_SlicesPooled,[-max(abs(Cg(:)))-10^-30 max(abs(Cg(:)))+10^-30]), axis square
        colormap('jet'),colorbar;
        for v1=1:size(Cg,1), for v2=1:size(Cg,2), if v1~=v2 %#ok<*ALIGN>
        if h_SFC_GM_all_WS_SlicesPooled(v1,v2)==1
        text(v1,v2,sprintf('%.2f',(Cg(v1,v2))),'HorizontalAlignment','center','VerticalAlignment','middle','fontsize',13','FontWeight','bold','color','k');
        else
        text(v1,v2,sprintf('%.2f',(Cg(v1,v2))),'HorizontalAlignment','center','VerticalAlignment','middle','fontsize',11','color',[0.1,0.1,0.1]);
        end
        end, end, end
        xticks([1:4]), xticklabels({'LV','RV','LD','RD'}), a = get(gca,'xticklabels'); set(gca,'xticklabels',a,'fontsize',13); clear a
        yticks([1:4]), yticklabels({'LV','RV','LD','RD'}), a = get(gca,'yticklabels'); set(gca,'yticklabels',a,'fontsize',13); clear a
        xlabel('Gray matter quadrants'),ylabel('Gray matter quadrants')
        title(sprintf('GRAY MATTER\n(Pearson''s correlation based functional connectivity)\nSignificant group-level T-statistics for within-slice FC\n(FC pooled across all slices for each connection)'),'Color','blue','FontSize',13)
    else
    Cw = real(tstat_SFC_WM_all_WS_SlicesPooled);
    fighndl = figure('Position',[1 1 1000 500],'Color',[0.89,0.94,0.99],'InvertHardcopy','off','Visible','off'); % red [0.95,0.9,0.85], green [0.85,0.90,0.85], blue [0.89,0.94,0.99]
    subplot2n(1,2,1), imagesc(Cg,'AlphaData',WhiteMask.*h_SFC_GM_all_WS_SlicesPooled,[-max(abs(Cg(:)))-10^-30 max(abs(Cg(:)))+10^-30]), axis square
        colormap('jet'),colorbar;
        for v1=1:size(Cg,1), for v2=1:size(Cg,2), if v1~=v2 %#ok<*ALIGN>
        if h_SFC_GM_all_WS_SlicesPooled(v1,v2)==1
        text(v1,v2,sprintf('%.2f',(Cg(v1,v2))),'HorizontalAlignment','center','VerticalAlignment','middle','fontsize',13','FontWeight','bold','color','k');
        else
        text(v1,v2,sprintf('%.2f',(Cg(v1,v2))),'HorizontalAlignment','center','VerticalAlignment','middle','fontsize',11','color',[0.1,0.1,0.1]);
        end
        end, end, end
        xticks([1:4]), xticklabels({'LV','RV','LD','RD'}), a = get(gca,'xticklabels'); set(gca,'xticklabels',a,'fontsize',13); clear a
        yticks([1:4]), yticklabels({'LV','RV','LD','RD'}), a = get(gca,'yticklabels'); set(gca,'yticklabels',a,'fontsize',13); clear a
        xlabel('Gray matter quadrants'),ylabel('Gray matter quadrants')
        title(sprintf('GRAY MATTER\n(Pearson''s correlation based functional connectivity)\nSignificant group-level T-statistics for within-slice FC\n(FC pooled across all slices for each connection)'),'Color','blue','FontSize',13)
    subplot2n(1,2,2), imagesc(Cw,'AlphaData',WhiteMask.*h_SFC_WM_all_WS_SlicesPooled,[-max(abs(Cw(:)))-10^-30 max(abs(Cw(:)))+10^-30]), axis square
        colormap('jet'),colorbar;
        for v1=1:size(Cw,1), for v2=1:size(Cw,2), if v1~=v2 %#ok<*ALIGN>
        if h_SFC_WM_all_WS_SlicesPooled(v1,v2)==1
        text(v1,v2,sprintf('%.2f',(Cw(v1,v2))),'HorizontalAlignment','center','VerticalAlignment','middle','fontsize',13','FontWeight','bold','color','k');
        else
        text(v1,v2,sprintf('%.2f',(Cw(v1,v2))),'HorizontalAlignment','center','VerticalAlignment','middle','fontsize',11','color',[0.1,0.1,0.1]);
        end
        end, end, end
        xticks([1:4]), xticklabels({'LV','RV','LD','RD'}), a = get(gca,'xticklabels'); set(gca,'xticklabels',a,'fontsize',13); clear a
        yticks([1:4]), yticklabels({'LV','RV','LD','RD'}), a = get(gca,'yticklabels'); set(gca,'yticklabels',a,'fontsize',13); clear a
        xlabel('White matter quadrants'),ylabel('White matter quadrants')
        title(sprintf('WHITE MATTER\n(Pearson''s correlation based functional connectivity)\nSignificant group-level T-statistics for within-slice FC\n(FC pooled across all slices for each connection)'),'Color','red','FontSize',13)
    end
    
    saveas(fighndl,[QCpath 'stats_SFC_WithinSlice_SlicesPooled' '.jpg'])
    close(fighndl)
    clear fighndl Cg Cw v1 v2
    end

    if partial==1
    Cg = real(tstat_SFC_pcor_GM_all_WS_SlicesPooled);
    if WM_flag == 0
    fighndl = figure('Position',[1 1 500 500],'Color',[0.89,0.94,0.99],'InvertHardcopy','off','Visible','off'); % red [0.95,0.9,0.85], green [0.85,0.90,0.85], blue [0.89,0.94,0.99]
    imagesc(Cg,'AlphaData',WhiteMask.*h_SFC_pcor_GM_all_WS_SlicesPooled,[-max(abs(Cg(:)))-10^-30 max(abs(Cg(:)))+10^-30]), axis square
        colormap('jet'),colorbar;
        for v1=1:size(Cg,1), for v2=1:size(Cg,2), if v1~=v2 %#ok<*ALIGN>
        text(v1,v2,sprintf('%.2f',(Cg(v1,v2))),'HorizontalAlignment','center','VerticalAlignment','middle','fontsize',13','FontWeight','bold','color','k');
        end, end, end
        xticks([1:4]), xticklabels({'LV','RV','LD','RD'}), a = get(gca,'xticklabels'); set(gca,'xticklabels',a,'fontsize',13); clear a
        yticks([1:4]), yticklabels({'LV','RV','LD','RD'}), a = get(gca,'yticklabels'); set(gca,'yticklabels',a,'fontsize',13); clear a
        xlabel('Gray matter quadrants'),ylabel('Gray matter quadrants')
        title(sprintf('GRAY MATTER\n(Partial correlation based functional connectivity)\nSignificant group-level T-statistics for within-slice FC\n(FC pooled across all slices for each connection)'),'Color','blue','FontSize',13)
    else
    Cw = real(tstat_SFC_pcor_WM_all_WS_SlicesPooled);
    fighndl = figure('Position',[1 1 1000 500],'Color',[0.89,0.94,0.99],'InvertHardcopy','off','Visible','off'); % red [0.95,0.9,0.85], green [0.85,0.90,0.85], blue [0.89,0.94,0.99]
    subplot2n(1,2,1), imagesc(Cg,'AlphaData',WhiteMask.*h_SFC_pcor_GM_all_WS_SlicesPooled,[-max(abs(Cg(:)))-10^-30 max(abs(Cg(:)))+10^-30]), axis square
        colormap('jet'),colorbar;
        for v1=1:size(Cg,1), for v2=1:size(Cg,2), if v1~=v2 %#ok<*ALIGN>
        text(v1,v2,sprintf('%.2f',(Cg(v1,v2))),'HorizontalAlignment','center','VerticalAlignment','middle','fontsize',13','FontWeight','bold','color','k');
        end, end, end
        xticks([1:4]), xticklabels({'LV','RV','LD','RD'}), a = get(gca,'xticklabels'); set(gca,'xticklabels',a,'fontsize',13); clear a
        yticks([1:4]), yticklabels({'LV','RV','LD','RD'}), a = get(gca,'yticklabels'); set(gca,'yticklabels',a,'fontsize',13); clear a
        xlabel('Gray matter quadrants'),ylabel('Gray matter quadrants')
        title(sprintf('GRAY MATTER\n(Partial correlation based functional connectivity)\nSignificant group-level T-statistics for within-slice FC\n(FC pooled across all slices for each connection)'),'Color','blue','FontSize',13)
    subplot2n(1,2,2), imagesc(Cw,'AlphaData',WhiteMask.*h_SFC_pcor_WM_all_WS_SlicesPooled,[-max(abs(Cw(:)))-10^-30 max(abs(Cw(:)))+10^-30]), axis square
        colormap('jet'),colorbar;
        for v1=1:size(Cw,1), for v2=1:size(Cw,2), if v1~=v2 %#ok<*ALIGN>
        text(v1,v2,sprintf('%.2f',(Cw(v1,v2))),'HorizontalAlignment','center','VerticalAlignment','middle','fontsize',13','FontWeight','bold','color','k');
        end, end, end
        xticks([1:4]), xticklabels({'LV','RV','LD','RD'}), a = get(gca,'xticklabels'); set(gca,'xticklabels',a,'fontsize',13); clear a
        yticks([1:4]), yticklabels({'LV','RV','LD','RD'}), a = get(gca,'yticklabels'); set(gca,'yticklabels',a,'fontsize',13); clear a
        xlabel('White matter quadrants'),ylabel('White matter quadrants')
        title(sprintf('WHITE MATTER\n(Partial correlation based functional connectivity)\nSignificant group-level T-statistics for within-slice FC\n(FC pooled across all slices for each connection)'),'Color','red','FontSize',13)
    end
    saveas(fighndl,[QCpath 'stats_SFC_pcor_WithinSlice_SlicesPooled' '.jpg'])
    close(fighndl)
    clear fighndl Cg Cw v1 v2
    end

end

%% QC plots of stats: within+between-slice FC
if ((choices_postproc(2)==1)||exist([sub_dir{1} ip_file{1,1} '_SFC_betweenslice' '.mat'],'file')||exist([sub_dir{1} ip_file{1,1} '_SFC_betweenslice_pcor' '.mat'],'file')) && (SFC_betweenslice_flag==1)
fprintf('23. QC plots: simple stats - SFC_BetweenSlices\n')

if pearson==1
Cg = real(tstat_SFC_GM_all_BS.*h_SFC_GM_all_BS);
Cg2 = []; Cg3 = [];
for v=1:siz3
    Cg2 = cat(1,Cg2,Cg(4*(v-1)+1:4*v,:),zeros(1,size(Cg,2)));
end
for v=1:siz3
    Cg3 = cat(2,Cg3,Cg2(:,4*(v-1)+1:4*v),zeros(size(Cg2,1),1));
end
clear Cg Cg2
Cg = Cg3; clear Cg3
Cg = Cg(1:end-1,1:end-1);

WhiteMask = ones(size(Cg)); WhiteMask(eye(size(Cg,1))==1) = 0;
BlackMask = ones(size(Cg)); BlackMask(find(and((Cg==0),(WhiteMask~=0)))) = 0;

fighndl = figure('Position',[1 1 1080 1080],'Color',[0.89,0.94,0.99],'InvertHardcopy','off','Visible','off'); % red [0.95,0.9,0.85], green [0.85,0.90,0.85], blue [0.89,0.94,0.99]
imagesc(Cg,'AlphaData',WhiteMask.*BlackMask,[-max(abs(Cg(:)))-10^-30 max(abs(Cg(:)))+10^-30]), axis square
	colormap('jet'),colorbar;
    for v=1:siz3, labelsC{v,1} = sprintf('%d',v); end
    xticks(2.5:5:size(Cg,1)), xticklabels(labelsC), a = get(gca,'xticklabels'); set(gca,'xticklabels',a,'fontsize',16); clear a
    yticks(2.5:5:size(Cg,1)), yticklabels(labelsC), a = get(gca,'yticklabels'); set(gca,'yticklabels',a,'fontsize',16); clear a
    xlabel('Slices (within each slice are shown LV | LD | RV | RD quadrants)'),ylabel('Slices (within each slice are shown LV | LD | RV | RD quadrants)')
    title(sprintf('GRAY MATTER\n(Pearson''s correlation based functional connectivity)\nSignificant group-level T-statistics for within- and between-slice FC across all slices'),'Color','blue','FontSize',18)
saveas(fighndl,[QCpath 'stats_SFC_Within_and_BetweenSlices_GM' '.jpg'])
close(fighndl)
clear fighndl Cg WhiteMask BlackMask

if WM_flag == 1
Cw = real(tstat_SFC_WM_all_BS.*h_SFC_WM_all_BS);
Cw2 = []; Cw3 = [];
for v=1:siz3
    Cw2 = cat(1,Cw2,Cw(4*(v-1)+1:4*v,:),zeros(1,size(Cw,2)));
end
for v=1:siz3
    Cw3 = cat(2,Cw3,Cw2(:,4*(v-1)+1:4*v),zeros(size(Cw2,1),1));
end
clear Cw Cw2
Cw = Cw3; clear Cw3
Cw = Cw(1:end-1,1:end-1);

WhiteMask = ones(size(Cw)); WhiteMask(eye(size(Cw,1))==1) = 0;
BlackMask = ones(size(Cw)); BlackMask(find(and((Cw==0),(WhiteMask~=0)))) = 0;

fighndl = figure('Position',[1 1 1080 1080],'Color',[0.89,0.94,0.99],'InvertHardcopy','off','Visible','off'); % red [0.95,0.9,0.85], green [0.85,0.90,0.85], blue [0.89,0.94,0.99]
imagesc(Cw,'AlphaData',WhiteMask.*BlackMask,[-max(abs(Cw(:)))-10^-30 max(abs(Cw(:)))+10^-30]), axis square
	colormap('jet'),colorbar;
    for v=1:siz3, labelsC{v,1} = sprintf('%d',v); end
    xticks(2.5:5:size(Cw,1)), xticklabels(labelsC), a = get(gca,'xticklabels'); set(gca,'xticklabels',a,'fontsize',16); clear a
    yticks(2.5:5:size(Cw,1)), yticklabels(labelsC), a = get(gca,'yticklabels'); set(gca,'yticklabels',a,'fontsize',16); clear a
    xlabel('Slices (within each slice are shown LV | LD | RV | RD quadrants)'),ylabel('Slices (within each slice are shown LV | LD | RV | RD quadrants)')
    title(sprintf('WHITE MATTER\n(Pearson''s correlation based functional connectivity)\nSignificant group-level T-statistics for within- and between-slice FC across all slices'),'Color','red','FontSize',18)
saveas(fighndl,[QCpath 'stats_SFC_Within_and_BetweenSlices_WM' '.jpg'])
close(fighndl)
clear fighndl Cw WhiteMask BlackMask
end
end


if (partial==1) && ~(dont_do_pcor_betweenslice==1)
Cg = real(tstat_SFC_pcor_GM_all_BS.*h_SFC_pcor_GM_all_BS);
Cg2 = []; Cg3 = [];
for v=1:siz3
    Cg2 = cat(1,Cg2,Cg(4*(v-1)+1:4*v,:),zeros(1,size(Cg,2)));
end
for v=1:siz3
    Cg3 = cat(2,Cg3,Cg2(:,4*(v-1)+1:4*v),zeros(size(Cg2,1),1));
end
clear Cg Cg2
Cg = Cg3; clear Cg3
Cg = Cg(1:end-1,1:end-1);

WhiteMask = ones(size(Cg)); WhiteMask(eye(size(Cg,1))==1) = 0;
BlackMask = ones(size(Cg)); BlackMask(find(and((Cg==0),(WhiteMask~=0)))) = 0;

fighndl = figure('Position',[1 1 1080 1080],'Color',[0.89,0.94,0.99],'InvertHardcopy','off','Visible','off'); % red [0.95,0.9,0.85], green [0.85,0.90,0.85], blue [0.89,0.94,0.99]
imagesc(Cg,'AlphaData',WhiteMask.*BlackMask,[-max(abs(Cg(:)))-10^-30 max(abs(Cg(:)))+10^-30]), axis square
	colormap('jet'),colorbar;
    for v=1:siz3, labelsC{v,1} = sprintf('%d',v); end
    xticks(2.5:5:size(Cg,1)), xticklabels(labelsC), a = get(gca,'xticklabels'); set(gca,'xticklabels',a,'fontsize',16); clear a
    yticks(2.5:5:size(Cg,1)), yticklabels(labelsC), a = get(gca,'yticklabels'); set(gca,'yticklabels',a,'fontsize',16); clear a
    xlabel('Slices (within each slice are shown LV | LD | RV | RD quadrants)'),ylabel('Slices (within each slice are shown LV | LD | RV | RD quadrants)')
    title(sprintf('GRAY MATTER\n(Partial correlation based functional connectivity)\nSignificant group-level T-statistics for within- and between-slice FC across all slices'),'Color','blue','FontSize',18)
saveas(fighndl,[QCpath 'stats_SFC_pcor_Within_and_BetweenSlices_GM' '.jpg'])
close(fighndl)
clear fighndl Cg WhiteMask BlackMask

if WM_flag == 1
Cw = real(tstat_SFC_pcor_WM_all_BS.*h_SFC_pcor_WM_all_BS);
Cw2 = []; Cw3 = [];
for v=1:siz3
    Cw2 = cat(1,Cw2,Cw(4*(v-1)+1:4*v,:),zeros(1,size(Cw,2)));
end
for v=1:siz3
    Cw3 = cat(2,Cw3,Cw2(:,4*(v-1)+1:4*v),zeros(size(Cw2,1),1));
end
clear Cw Cw2
Cw = Cw3; clear Cw3
Cw = Cw(1:end-1,1:end-1);

WhiteMask = ones(size(Cw)); WhiteMask(eye(size(Cw,1))==1) = 0;
BlackMask = ones(size(Cw)); BlackMask(find(and((Cw==0),(WhiteMask~=0)))) = 0;

fighndl = figure('Position',[1 1 1080 1080],'Color',[0.89,0.94,0.99],'InvertHardcopy','off','Visible','off'); % red [0.95,0.9,0.85], green [0.85,0.90,0.85], blue [0.89,0.94,0.99]
imagesc(Cw,'AlphaData',WhiteMask.*BlackMask,[-max(abs(Cw(:)))-10^-30 max(abs(Cw(:)))+10^-30]), axis square
	colormap('jet'),colorbar;
    for v=1:siz3, labelsC{v,1} = sprintf('%d',v); end
    xticks(2.5:5:size(Cw,1)), xticklabels(labelsC), a = get(gca,'xticklabels'); set(gca,'xticklabels',a,'fontsize',16); clear a
    yticks(2.5:5:size(Cw,1)), yticklabels(labelsC), a = get(gca,'yticklabels'); set(gca,'yticklabels',a,'fontsize',16); clear a
    xlabel('Slices (within each slice are shown LV | LD | RV | RD quadrants)'),ylabel('Slices (within each slice are shown LV | LD | RV | RD quadrants)')
    title(sprintf('WHITE MATTER\n(Partial correlation based functional connectivity)\nSignificant group-level T-statistics for within- and between-slice FC across all slices'),'Color','red','FontSize',18)
saveas(fighndl,[QCpath 'stats_SFC_pcor_Within_and_BetweenSlices_WM' '.jpg'])
close(fighndl)
clear fighndl Cw WhiteMask BlackMask
end
end


% Generate unmasked T-statistic images next:

if pearson==1
Cg = real(tstat_SFC_GM_all_BS);
Cg2 = []; Cg3 = [];
for v=1:siz3
    Cg2 = cat(1,Cg2,Cg(4*(v-1)+1:4*v,:),zeros(1,size(Cg,2)));
end
for v=1:siz3
    Cg3 = cat(2,Cg3,Cg2(:,4*(v-1)+1:4*v),zeros(size(Cg2,1),1));
end
clear Cg Cg2
Cg = Cg3; clear Cg3
Cg = Cg(1:end-1,1:end-1);

WhiteMask = ones(size(Cg)); WhiteMask(eye(size(Cg,1))==1) = 0;
BlackMask = ones(size(Cg)); BlackMask(find(and((Cg==0),(WhiteMask~=0)))) = 0;

fighndl = figure('Position',[1 1 1080 1080],'Color',[0.89,0.94,0.99],'InvertHardcopy','off','Visible','off'); % red [0.95,0.9,0.85], green [0.85,0.90,0.85], blue [0.89,0.94,0.99]
imagesc(Cg,'AlphaData',WhiteMask.*BlackMask,[-max(abs(Cg(:)))-10^-30 max(abs(Cg(:)))+10^-30]), axis square
	colormap('jet'),colorbar;
    for v=1:siz3, labelsC{v,1} = sprintf('%d',v); end
    xticks(2.5:5:size(Cg,1)), xticklabels(labelsC), a = get(gca,'xticklabels'); set(gca,'xticklabels',a,'fontsize',16); clear a
    yticks(2.5:5:size(Cg,1)), yticklabels(labelsC), a = get(gca,'yticklabels'); set(gca,'yticklabels',a,'fontsize',16); clear a
    xlabel('Slices (within each slice are shown LV | LD | RV | RD quadrants)'),ylabel('Slices (within each slice are shown LV | LD | RV | RD quadrants)')
    title(sprintf('GRAY MATTER\n(Pearson''s correlation based functional connectivity)\nUnthresholded group-level T-statistics for within- and between-slice FC across all slices'),'Color','blue','FontSize',18)
saveas(fighndl,[QCpath 'stats_SFC_Within_and_BetweenSlices_GM_unthresholded' '.jpg'])
close(fighndl)
clear fighndl Cg WhiteMask BlackMask

if WM_flag == 1
Cw = real(tstat_SFC_WM_all_BS);
Cw2 = []; Cw3 = [];
for v=1:siz3
    Cw2 = cat(1,Cw2,Cw(4*(v-1)+1:4*v,:),zeros(1,size(Cw,2)));
end
for v=1:siz3
    Cw3 = cat(2,Cw3,Cw2(:,4*(v-1)+1:4*v),zeros(size(Cw2,1),1));
end
clear Cw Cw2
Cw = Cw3; clear Cw3
Cw = Cw(1:end-1,1:end-1);

WhiteMask = ones(size(Cw)); WhiteMask(eye(size(Cw,1))==1) = 0;
BlackMask = ones(size(Cw)); BlackMask(find(and((Cw==0),(WhiteMask~=0)))) = 0;

fighndl = figure('Position',[1 1 1080 1080],'Color',[0.89,0.94,0.99],'InvertHardcopy','off','Visible','off'); % red [0.95,0.9,0.85], green [0.85,0.90,0.85], blue [0.89,0.94,0.99]
imagesc(Cw,'AlphaData',WhiteMask.*BlackMask,[-max(abs(Cw(:)))-10^-30 max(abs(Cw(:)))+10^-30]), axis square
	colormap('jet'),colorbar;
    for v=1:siz3, labelsC{v,1} = sprintf('%d',v); end
    xticks(2.5:5:size(Cw,1)), xticklabels(labelsC), a = get(gca,'xticklabels'); set(gca,'xticklabels',a,'fontsize',16); clear a
    yticks(2.5:5:size(Cw,1)), yticklabels(labelsC), a = get(gca,'yticklabels'); set(gca,'yticklabels',a,'fontsize',16); clear a
    xlabel('Slices (within each slice are shown LV | LD | RV | RD quadrants)'),ylabel('Slices (within each slice are shown LV | LD | RV | RD quadrants)')
    title(sprintf('WHITE MATTER\n(Pearson''s correlation based functional connectivity)\nUnthresholded group-level T-statistics for within- and between-slice FC across all slices'),'Color','red','FontSize',18)
saveas(fighndl,[QCpath 'stats_SFC_Within_and_BetweenSlices_WM_unthresholded' '.jpg'])
close(fighndl)
clear fighndl Cw WhiteMask BlackMask
end
end


if (partial==1) && ~(dont_do_pcor_betweenslice==1)
Cg = real(tstat_SFC_pcor_GM_all_BS);
Cg2 = []; Cg3 = [];
for v=1:siz3
    Cg2 = cat(1,Cg2,Cg(4*(v-1)+1:4*v,:),zeros(1,size(Cg,2)));
end
for v=1:siz3
    Cg3 = cat(2,Cg3,Cg2(:,4*(v-1)+1:4*v),zeros(size(Cg2,1),1));
end
clear Cg Cg2
Cg = Cg3; clear Cg3
Cg = Cg(1:end-1,1:end-1);

WhiteMask = ones(size(Cg)); WhiteMask(eye(size(Cg,1))==1) = 0;
BlackMask = ones(size(Cg)); BlackMask(find(and((Cg==0),(WhiteMask~=0)))) = 0;

fighndl = figure('Position',[1 1 1080 1080],'Color',[0.89,0.94,0.99],'InvertHardcopy','off','Visible','off'); % red [0.95,0.9,0.85], green [0.85,0.90,0.85], blue [0.89,0.94,0.99]
imagesc(Cg,'AlphaData',WhiteMask.*BlackMask,[-max(abs(Cg(:)))-10^-30 max(abs(Cg(:)))+10^-30]), axis square
	colormap('jet'),colorbar;
    for v=1:siz3, labelsC{v,1} = sprintf('%d',v); end
    xticks(2.5:5:size(Cg,1)), xticklabels(labelsC), a = get(gca,'xticklabels'); set(gca,'xticklabels',a,'fontsize',16); clear a
    yticks(2.5:5:size(Cg,1)), yticklabels(labelsC), a = get(gca,'yticklabels'); set(gca,'yticklabels',a,'fontsize',16); clear a
    xlabel('Slices (within each slice are shown LV | LD | RV | RD quadrants)'),ylabel('Slices (within each slice are shown LV | LD | RV | RD quadrants)')
    title(sprintf('GRAY MATTER\n(Partial correlation based functional connectivity)\nUnthresholded group-level T-statistics for within- and between-slice FC across all slices'),'Color','blue','FontSize',18)
saveas(fighndl,[QCpath 'stats_SFC_pcor_Within_and_BetweenSlices_GM_unthresholded' '.jpg'])
close(fighndl)
clear fighndl Cg WhiteMask BlackMask

if WM_flag == 1
Cw = real(tstat_SFC_pcor_WM_all_BS);
Cw2 = []; Cw3 = [];
for v=1:siz3
    Cw2 = cat(1,Cw2,Cw(4*(v-1)+1:4*v,:),zeros(1,size(Cw,2)));
end
for v=1:siz3
    Cw3 = cat(2,Cw3,Cw2(:,4*(v-1)+1:4*v),zeros(size(Cw2,1),1));
end
clear Cw Cw2
Cw = Cw3; clear Cw3
Cw = Cw(1:end-1,1:end-1);

WhiteMask = ones(size(Cw)); WhiteMask(eye(size(Cw,1))==1) = 0;
BlackMask = ones(size(Cw)); BlackMask(find(and((Cw==0),(WhiteMask~=0)))) = 0;

fighndl = figure('Position',[1 1 1080 1080],'Color',[0.89,0.94,0.99],'InvertHardcopy','off','Visible','off'); % red [0.95,0.9,0.85], green [0.85,0.90,0.85], blue [0.89,0.94,0.99]
imagesc(Cw,'AlphaData',WhiteMask.*BlackMask,[-max(abs(Cw(:)))-10^-30 max(abs(Cw(:)))+10^-30]), axis square
	colormap('jet'),colorbar;
    for v=1:siz3, labelsC{v,1} = sprintf('%d',v); end
    xticks(2.5:5:size(Cw,1)), xticklabels(labelsC), a = get(gca,'xticklabels'); set(gca,'xticklabels',a,'fontsize',16); clear a
    yticks(2.5:5:size(Cw,1)), yticklabels(labelsC), a = get(gca,'yticklabels'); set(gca,'yticklabels',a,'fontsize',16); clear a
    xlabel('Slices (within each slice are shown LV | LD | RV | RD quadrants)'),ylabel('Slices (within each slice are shown LV | LD | RV | RD quadrants)')
    title(sprintf('WHITE MATTER\n(Partial correlation based functional connectivity)\nUnthresholded group-level T-statistics for within- and between-slice FC across all slices'),'Color','red','FontSize',18)
saveas(fighndl,[QCpath 'stats_SFC_pcor_Within_and_BetweenSlices_WM_unthresholded' '.jpg'])
close(fighndl)
clear fighndl Cw WhiteMask BlackMask
end
end

end

%% QC plots of stats: fALFF
if ((choices_postproc(3)==1)||exist([sub_dir{1} ip_file2{1,1} '_fALFF' '.mat'],'file'))
fprintf('23. QC plots: simple stats - fALFF\n\n')

tstat_fALFF_GM_all = tstat_fALFF_GM_all.*h_fALFF_GM_all; tstat_fALFF_GM_all(find(tstat_fALFF_GM_all==0))=NaN;
if WM_flag == 0
    tstat_fALFF_WM_all = zeros(size(tstat_fALFF_GM_all));
    tstat_fALFF_WM_all(find(tstat_fALFF_WM_all==0)) = NaN;
else
tstat_fALFF_WM_all = tstat_fALFF_WM_all.*h_fALFF_WM_all; tstat_fALFF_WM_all(find(tstat_fALFF_WM_all==0))=NaN;
end

fighndl = figure('Position',[1 1 1280 1280],'Color',[0.85,0.90,0.85],'InvertHardcopy','off','Visible','off'); % red [0.95,0.9,0.85], green [0.85,0.90,0.85], blue [0.89,0.94,0.99]
subplot2n(5,1,1), bar([tstat_fALFF_GM_all(:,1),tstat_fALFF_WM_all(:,1)]'), ylabel('T-statisic'), xticklabels({sprintf('GRAY MATTER fALFF T-statistics (slices 1 -> %d)',siz3),sprintf('WHITE MATTER fALFF T-statistics (slices 1 -> %d)',siz3)}),
    a = get(gca,'xticklabels'); set(gca,'xticklabels',a,'fontsize',12); clear a
    grid on,grid minor, 
    try ylim([nanmin([tstat_fALFF_GM_all(:);tstat_fALFF_WM_all(:)])-(nanmax([tstat_fALFF_GM_all(:);tstat_fALFF_WM_all(:)])-nanmin([tstat_fALFF_GM_all(:);tstat_fALFF_WM_all(:)]))/2, nanmax([tstat_fALFF_GM_all(:);tstat_fALFF_WM_all(:)])+(nanmax([tstat_fALFF_GM_all(:);tstat_fALFF_WM_all(:)])-nanmin([tstat_fALFF_GM_all(:);tstat_fALFF_WM_all(:)]))/2]), catch, end
    title(sprintf('Significant group-level statistics for fractional amplitude of low-frequency BOLD fluctuations (fALFF)\n\nLeft ventral horn: fALFF T-statistics across SLICES'),'Color',[0,0.65,0.59],'FontSize',16), 
    text(0.9,nanmax([tstat_fALFF_GM_all(:);tstat_fALFF_WM_all(:)])+(nanmax([tstat_fALFF_GM_all(:);tstat_fALFF_WM_all(:)])-nanmin([tstat_fALFF_GM_all(:);tstat_fALFF_WM_all(:)]))/4, sprintf('mean = %.2f',nanmean(tstat_fALFF_GM_all(:,1))),'Color',[0,0.55,0.69],'FontSize',15)
    text(1.9,nanmax([tstat_fALFF_GM_all(:);tstat_fALFF_WM_all(:)])+(nanmax([tstat_fALFF_GM_all(:);tstat_fALFF_WM_all(:)])-nanmin([tstat_fALFF_GM_all(:);tstat_fALFF_WM_all(:)]))/4, sprintf('mean = %.2f',nanmean(tstat_fALFF_WM_all(:,1))),'Color',[0.65,0.11,0.19],'FontSize',15)
subplot2n(5,1,2), bar([tstat_fALFF_GM_all(:,2),tstat_fALFF_WM_all(:,2)]'), ylabel('T-statisic'), xticklabels({sprintf('GRAY MATTER fALFF T-statistics (slices 1 -> %d)',siz3),sprintf('WHITE MATTER fALFF T-statistics (slices 1 -> %d)',siz3)}),
    a = get(gca,'xticklabels'); set(gca,'xticklabels',a,'fontsize',12); clear a
    grid on,grid minor, 
    try ylim([nanmin([tstat_fALFF_GM_all(:);tstat_fALFF_WM_all(:)])-(nanmax([tstat_fALFF_GM_all(:);tstat_fALFF_WM_all(:)])-nanmin([tstat_fALFF_GM_all(:);tstat_fALFF_WM_all(:)]))/2, nanmax([tstat_fALFF_GM_all(:);tstat_fALFF_WM_all(:)])+(nanmax([tstat_fALFF_GM_all(:);tstat_fALFF_WM_all(:)])-nanmin([tstat_fALFF_GM_all(:);tstat_fALFF_WM_all(:)]))/2]), catch, end
    title(['Right ventral horn: fALFF T-statistics across SLICES'],'Color',[0,0.45,0.79],'FontSize',16), 
    text(0.9,nanmax([tstat_fALFF_GM_all(:);tstat_fALFF_WM_all(:)])+(nanmax([tstat_fALFF_GM_all(:);tstat_fALFF_WM_all(:)])-nanmin([tstat_fALFF_GM_all(:);tstat_fALFF_WM_all(:)]))/4, sprintf('mean = %.2f',nanmean(tstat_fALFF_GM_all(:,2))),'Color',[0,0.55,0.69],'FontSize',15)
    text(1.9,nanmax([tstat_fALFF_GM_all(:);tstat_fALFF_WM_all(:)])+(nanmax([tstat_fALFF_GM_all(:);tstat_fALFF_WM_all(:)])-nanmin([tstat_fALFF_GM_all(:);tstat_fALFF_WM_all(:)]))/4, sprintf('mean = %.2f',nanmean(tstat_fALFF_WM_all(:,2))),'Color',[0.65,0.11,0.19],'FontSize',15)
subplot2n(5,1,3), bar([tstat_fALFF_GM_all(:,3),tstat_fALFF_WM_all(:,3)]'), ylabel('T-statisic'), xticklabels({sprintf('GRAY MATTER fALFF T-statistics (slices 1 -> %d)',siz3),sprintf('WHITE MATTER fALFF T-statistics (slices 1 -> %d)',siz3)}),
    a = get(gca,'xticklabels'); set(gca,'xticklabels',a,'fontsize',12); clear a
    grid on,grid minor, 
    try ylim([nanmin([tstat_fALFF_GM_all(:);tstat_fALFF_WM_all(:)])-(nanmax([tstat_fALFF_GM_all(:);tstat_fALFF_WM_all(:)])-nanmin([tstat_fALFF_GM_all(:);tstat_fALFF_WM_all(:)]))/2, nanmax([tstat_fALFF_GM_all(:);tstat_fALFF_WM_all(:)])+(nanmax([tstat_fALFF_GM_all(:);tstat_fALFF_WM_all(:)])-nanmin([tstat_fALFF_GM_all(:);tstat_fALFF_WM_all(:)]))/2]), catch, end
    title(['Left dorsal horn: fALFF T-statistics across SLICES'],'Color',[0.55,0.01,0.39],'FontSize',16), 
    text(0.9,nanmax([tstat_fALFF_GM_all(:);tstat_fALFF_WM_all(:)])+(nanmax([tstat_fALFF_GM_all(:);tstat_fALFF_WM_all(:)])-nanmin([tstat_fALFF_GM_all(:);tstat_fALFF_WM_all(:)]))/4, sprintf('mean = %.2f',nanmean(tstat_fALFF_GM_all(:,3))),'Color',[0,0.55,0.69],'FontSize',15)
    text(1.9,nanmax([tstat_fALFF_GM_all(:);tstat_fALFF_WM_all(:)])+(nanmax([tstat_fALFF_GM_all(:);tstat_fALFF_WM_all(:)])-nanmin([tstat_fALFF_GM_all(:);tstat_fALFF_WM_all(:)]))/4, sprintf('mean = %.2f',nanmean(tstat_fALFF_WM_all(:,3))),'Color',[0.65,0.11,0.19],'FontSize',15)
subplot2n(5,1,4), bar([tstat_fALFF_GM_all(:,4),tstat_fALFF_WM_all(:,4)]'), ylabel('T-statisic'), xticklabels({sprintf('GRAY MATTER fALFF T-statistics (slices 1 -> %d)',siz3),sprintf('WHITE MATTER fALFF T-statistics (slices 1 -> %d)',siz3)}),
    a = get(gca,'xticklabels'); set(gca,'xticklabels',a,'fontsize',12); clear a
    grid on,grid minor, 
    try ylim([nanmin([tstat_fALFF_GM_all(:);tstat_fALFF_WM_all(:)])-(nanmax([tstat_fALFF_GM_all(:);tstat_fALFF_WM_all(:)])-nanmin([tstat_fALFF_GM_all(:);tstat_fALFF_WM_all(:)]))/2, nanmax([tstat_fALFF_GM_all(:);tstat_fALFF_WM_all(:)])+(nanmax([tstat_fALFF_GM_all(:);tstat_fALFF_WM_all(:)])-nanmin([tstat_fALFF_GM_all(:);tstat_fALFF_WM_all(:)]))/2]), catch, end
    title(['Right dorsal horn: fALFF T-statistics across SLICES'],'Color',[0.55,0.31,0.09],'FontSize',16), 
    text(0.9,nanmax([tstat_fALFF_GM_all(:);tstat_fALFF_WM_all(:)])+(nanmax([tstat_fALFF_GM_all(:);tstat_fALFF_WM_all(:)])-nanmin([tstat_fALFF_GM_all(:);tstat_fALFF_WM_all(:)]))/4, sprintf('mean = %.2f',nanmean(tstat_fALFF_GM_all(:,4))),'Color',[0,0.55,0.69],'FontSize',15)
    text(1.9,nanmax([tstat_fALFF_GM_all(:);tstat_fALFF_WM_all(:)])+(nanmax([tstat_fALFF_GM_all(:);tstat_fALFF_WM_all(:)])-nanmin([tstat_fALFF_GM_all(:);tstat_fALFF_WM_all(:)]))/4, sprintf('mean = %.2f',nanmean(tstat_fALFF_WM_all(:,4))),'Color',[0.65,0.11,0.19],'FontSize',15)
subplot2n(5,1,5), bar([nanmean(tstat_fALFF_GM_all,1)',nanmean(tstat_fALFF_WM_all,1)']'), ylabel('T-statisic'), xticklabels({sprintf('Median GRAY MATTER fALFF T-statistics (within LV | RV | LD | RD)'),sprintf('Median WHITE MATTER fALFF T-statistics (within LV | RV | LD | RD)')}),
    a = get(gca,'xticklabels'); set(gca,'xticklabels',a,'fontsize',12); clear a
    grid on,grid minor, 
    try ylim([nanmin([tstat_fALFF_GM_all(:);tstat_fALFF_WM_all(:)])-(nanmax([tstat_fALFF_GM_all(:);tstat_fALFF_WM_all(:)])-nanmin([tstat_fALFF_GM_all(:);tstat_fALFF_WM_all(:)]))/2, nanmax([tstat_fALFF_GM_all(:);tstat_fALFF_WM_all(:)])+(nanmax([tstat_fALFF_GM_all(:);tstat_fALFF_WM_all(:)])-nanmin([tstat_fALFF_GM_all(:);tstat_fALFF_WM_all(:)]))/2]), catch, end
    title(['Median fALFF T-statistics across QUADRANTS (LV | RV | LD | RD)'],'Color',[0.24,0.25,0.26],'FontSize',16), 
    text(0.9,nanmax([tstat_fALFF_GM_all(:);tstat_fALFF_WM_all(:)])+(nanmax([tstat_fALFF_GM_all(:);tstat_fALFF_WM_all(:)])-nanmin([tstat_fALFF_GM_all(:);tstat_fALFF_WM_all(:)]))/4, sprintf('median = %.2f',nanmedian(tstat_fALFF_GM_all(:))),'Color',[0,0.55,0.69],'FontSize',15)
    text(1.9,nanmax([tstat_fALFF_GM_all(:);tstat_fALFF_WM_all(:)])+(nanmax([tstat_fALFF_GM_all(:);tstat_fALFF_WM_all(:)])-nanmin([tstat_fALFF_GM_all(:);tstat_fALFF_WM_all(:)]))/4, sprintf('median = %.2f',nanmedian(tstat_fALFF_WM_all(:))),'Color',[0.65,0.11,0.19],'FontSize',15)
saveas(fighndl,[QCpath 'stats_fALFF' '.jpg'])
close(fighndl)

end

    
end
```
