## Supplementary material for "Neptune: a toolbox for spinal cord functional MRI data processing and quality assurance": Neptune software: generate_QC_plots_only.m

```
function generate_QC_plots_only(sub_dir,func_files,anat_files,RUNS,slices,choices_preproc, cutoff_str,bpf_hz,save_videos_, varargin)

% use this function to generate (or regenerate) QC plots only, 
% if you have already done pre-processing and have the corresponding outputs.
% (This is for pre-processing QC outputs only)

siz3 = slices;
runs = RUNS;

home_dir = sub_dir{1}(1:max(strfind(sub_dir{1}(1:end-1),'/')));
if isempty(home_dir)
    home_dir = sub_dir{1}(1:max(strfind(sub_dir{1}(1:end-1),'\')));
end

if nargin>=10
    run_on_HPF_data = varargin{1};
else
    run_on_HPF_data = 1; % assume
end

if nargin>=11
    choices_postproc = varargin{2};
else
    choices_postproc = 0; % assume
end
if exist('choices_postproc','var')
    postproc=1;
else 
    postproc=0;
end

if nargin>=12
    time_taken_preproc = varargin{3};
end

if bpf_hz(1)<0.1
    bpf_hz_str = sprintf('0%.0f',bpf_hz(1)*100);
else
    bpf_hz_str = sprintf('%.0f',bpf_hz(1)*100);
end

scrsz = get(0,'ScreenSize'); pause on;
save_videos_steps = zeros(8,1);

%%
for k=1:length(sub_dir)
    for k2=1:RUNS(k)
%%  
disp(sprintf('subject (%d) of (%d), run (%d) of (%d)',k,length(sub_dir),k2,RUNS(k)))
base_dir_sub = sub_dir{k};
fname = func_files{k,k2};
fname3 = [fname '_mean'];
fname_anat_orig = anat_files{k};
fname_anat = [fname_anat_orig '_reduced'];

if RUNS(k)==1, Rns=0; else, Rns=k2; end

cmap = colormap('bone'); cmap = (1-exp(-cmap))./(1-exp(-1)); 

if ~(exist([base_dir_sub 'QC'],'dir'))
    mkdir([base_dir_sub 'QC'])
end

Smask_err=0;
try
    scfMRItb_04_resplitData(base_dir_sub, fname, '_Smask', siz3) % if slice-wise data doesn't exist but combined 4D data exists, then split the data to generate the needed slice-wise data
catch
    load([base_dir_sub fname '_mask_NS' '.mat'])
    Smask1_ = ~mask_NS.img; clear mask_NS
    Smask_err=1;
end

try
    scfMRItb_04_unzipFile(base_dir_sub, fname_anat, '_mask_GM')
    scfMRItb_04_unzipFile(base_dir_sub, fname_anat, '_mask_WM')
    mask_GM = load_untouch_nii([base_dir_sub fname_anat '_mask_GM.nii']); mask_GM = mask_GM.img;
    mask_WM = load_untouch_nii([base_dir_sub fname_anat '_mask_WM.nii']); mask_WM = mask_WM.img;
catch
end

%% scfMRItb_05_notCordMask
if choices_preproc(5)==1
if ~(exist([base_dir_sub 'QC' '/05_notCordMask'],'dir'))
    mkdir([base_dir_sub 'QC' '/05_notCordMask'])
end
if Rns==0
    QCpath2 = [base_dir_sub 'QC' '/05_notCordMask/'];
else
    if ~(exist([base_dir_sub 'QC' '/05_notCordMask' '/run' num2str(Rns)],'dir'))
        mkdir([base_dir_sub 'QC' '/05_notCordMask' '/run' num2str(Rns)])
    end
    QCpath2 = [base_dir_sub 'QC' '/05_notCordMask' '/run' num2str(Rns) '/'];
end

if ~(exist([base_dir_sub 'QC' '/04_splitData_rawQC'],'dir'))
    mkdir([base_dir_sub 'QC' '/04_splitData_rawQC'])
end
if Rns==0
    QCpath = [base_dir_sub 'QC' '/04_splitData_rawQC/'];
else
    if ~(exist([base_dir_sub 'QC' '/04_splitData_rawQC' '/run' num2str(Rns)],'dir'))
        mkdir([base_dir_sub 'QC' '/04_splitData_rawQC' '/run' num2str(Rns)])
    end
    QCpath = [base_dir_sub 'QC' '/04_splitData_rawQC' '/run' num2str(Rns) '/'];
end

if (save_videos_==0)||(save_videos_==1)
    save_videos = save_videos_;
elseif (save_videos_==2)&&(sum([choices_preproc(6);choices_preproc(8:9);choices_preproc(11:14)])==0)
    save_videos = 1;
else
    save_videos = 0;
end
if save_videos==1
    save_videos_steps(1)=1;
end


try
    load([base_dir_sub fname '_mask_NS.mat'],'position_maskNS')
    scfMRItb_04_unzipFile(base_dir_sub, fname3, '')
    A = load_untouch_nii([base_dir_sub fname3 '.nii']); % mean functional
    new_size = round(size(A.img,1) / 2);
    for i3 = 1 : siz3
        try waitbar((i3/siz3),wbar3,sprintf('05. Define "not-cord" mask: slice (%d) of (%d)',i3,siz3)); catch, end
        img = rot90(A.img(:,:,i3));
        temp = img(round((size(img,1)-new_size)/2)+1:end-round((size(img,1)-new_size)/2), ...
            round((size(img,2)-new_size)/2)+1:end-round((size(img,2)-new_size)/2));
        mmax = max(temp(:)); clear temp;
        fighndl = figure('Position',[round((scrsz(3)-scrsz(4))/2) 1 scrsz(4) scrsz(4)],'Color',[0.8,0.89,0.84],'InvertHardcopy','off','Visible','off'); % [0.85,0.90,0.85] [361 1 scrsz(4) scrsz(4)]
        imshow(single(img), [0 mmax], 'InitialMagnification', 'fit'); colorbar;
        fighndl.Color = [0.8, 0.87, 0.84] + [0, 0.02*(1-exp(siz3/i3)/exp(siz3)), 0.1*exp(i3/siz3)/exp(1)]; % [0.85, 0.90, 0.85] + [0, 0, 0.1*i3/siz3];
        title({'Draw a boundary selecting only the spinal cord (double click the cord once done)';['Slice ' num2str(i3) ' of ' num2str(siz3)]},'Color',[i3/(5*siz3),0.5,i3/siz3],'FontSize',16);
        hold on, plot([position_maskNS{i3}(:,1);position_maskNS{i3}(1,1)],[position_maskNS{i3}(:,2);position_maskNS{i3}(1,2)],'m','LineWidth',2), hold off
        lims_x1(i3) = min(position_maskNS{i3}(:,1)); lims_x2(i3) = max(position_maskNS{i3}(:,1)); %#ok<*AGROW>
        lims_x1_(i3) = round(lims_x1(i3) - 0.5*(lims_x2(i3)-lims_x1(i3))); lims_x2_(i3) = round(lims_x2(i3) + 0.5*(lims_x2(i3)-lims_x1(i3)));
        lims_y1(i3) = min(position_maskNS{i3}(:,2)); lims_y2(i3) = max(position_maskNS{i3}(:,2));
        lims_y1_(i3) = round(lims_y1(i3) - 0.5*(lims_y2(i3)-lims_y1(i3))); lims_y2_(i3) = round(lims_y2(i3) + 0.5*(lims_y2(i3)-lims_y1(i3)));
        xlim([min([round(size(img,1)*w1min(i3)),lims_x1_(i3)]),max([round(size(img,1)*w1max(i3)),lims_x2_(i3)])]),
        ylim([min([round(size(img,2)*w2min(i3)),lims_y1_(i3)]),max([round(size(img,2)*w2max(i3)),lims_y2_(i3)])]),
        saveas(fighndl,[QCpath2 'notCordMask_slice' sprintf('%.2d',i3) '.jpg'])
        close(fighndl)
        clear img mmax        
    end
    clear A position_maskNS fighndl lims_x1 lims_x2 lims_y1 lims_y2 lims_y2_ lims_y1_ lims_x2_ lims_x1_
catch
end


%%% SNR graph
% Raw data SNR is computed as the ratio of signal variance in the cord (median value) to the signal variance outside the cord (median value) within a given slice, computed from raw NIfTI data.
try
    load([base_dir_sub 'SNR_slicewise_run' num2str(k2) '.mat'], 'SNR','SNR_in_dB')
catch
load([base_dir_sub fname '_mask_NS' '.mat'])
mask_NS = mask_NS.img;
scfMRItb_04_unzipFile(base_dir_sub, fname, '')
F = load_untouch_nii([base_dir_sub fname '.nii']);
F = F.img;
for i3=1:siz3
    fprintf('05. SNR computation: slice (%d) of (%d)\n',i3,siz3)
    for w=1:size(F,4)
        temp = F(:,:,i3,w);
        ts_S(:,w)  = temp(find(mask_NS(:,:,i3)==0)); % spine
        ts_NS(:,w) = temp(find(mask_NS(:,:,i3)==1)); % not spine
        clear temp
    end
    for w=1:size(ts_S,1)
        var1(w,1) = nanvar(ts_S(w,:)'); %#ok<*UDIM>
    end
    for w=1:size(ts_NS,1)
        var2(w,1) = nanvar(ts_NS(w,:)');
    end
    SNR(i3,1) = median(var1)/median(var2);
    clear ts_S ts_NS var1 var2
end
SNR_in_dB = 10.*log(SNR);
info_SNR = sprintf('SNR = signal-to-noise ratio defined as the ratio of signal variance in the cord (median value) to the signal variance outside the cord (median value) within a given slice, computed from raw NIfTI data.\nThat is, SNR = median(var(timeseries_cord)) / median(var(timeseries_notcord));\nSNR_in_dB = 10.*log(SNR);\n');
save([base_dir_sub 'SNR_slicewise_run' num2str(run) '.mat'], 'SNR','SNR_in_dB','info_SNR');
end

% SNR QC plot
for w=1:length(SNR)
    dispstring1{w} = sprintf('%.2f',SNR(w));
    dispstring2{w} = sprintf('%.2f dB',SNR_in_dB(w));
end; clear w
fighndl = figure('Position',[1 1 1280 800],'Color',[0.85,0.90,0.85],'InvertHardcopy','off','Visible','off'); % red [0.95,0.9,0.85], green [0.85,0.90,0.85], blue [0.89,0.94,0.99]
subplot2n(2,1,1), plot(SNR,'--o','color',[0.25,0.25,0.25],'linewidth',2.25, 'MarkerSize',14, 'MarkerEdgeColor','k', 'MarkerFaceColor',[0.85,0.85,0.85]),
    grid on,grid minor, ylabel('Median signal-to-noise ratio'), xlabel('SLICES'), xticks([1:length(SNR)]),
    xlim([0.5 length(SNR)+0.5]), ylim([0 1.2*max(SNR)])
    text([1:length(SNR)]-0.15, double(SNR'+0.1*max(SNR)), dispstring1,'Color',[0.05,0.05,0.05],'FontSize',12)
    legend({'Raw data SNR'},'FontSize',14,'TextColor','k','Location','best')
    title(sprintf('BOLD signal-to-noise ratio (SNR) across slices, defined as:\nthe ratio of BOLD signal variance in the cord (median value) to the BOLD signal variance outside the cord (median value)'),'Color',[0.4,0.1,0.1],'FontSize',16)
subplot2n(2,1,2), plot(SNR_in_dB,'--o','color',[0.25,0.25,0.25],'linewidth',2.25, 'MarkerSize',14, 'MarkerEdgeColor',[0.4,0.1,0.1], 'MarkerFaceColor',[0.85,0.75,0.75]),
    grid on,grid minor,
    ylabel('Median SNR in decibels'), xlabel('SLICES'), xticks([1:length(SNR_in_dB)]),
    xlim([0.5 length(SNR_in_dB)+0.5]), ylim([min([0,1.2*min(SNR_in_dB)]) 1.2*max(SNR_in_dB)])
    text([1:length(SNR_in_dB)]-0.25, double(SNR_in_dB'+0.1*max(SNR_in_dB)), dispstring2,'Color',[0.16,0.04,0.04],'FontSize',12)
    legend({'Raw data SNR in decibels'},'FontSize',14,'TextColor',[0.16,0.04,0.04],'Location','best')
    title(sprintf('BOLD SNR across slices in decibels:\nSNR in dB = 10 x log(SNR)'),'Color',[0.4,0.1,0.1],'FontSize',16)
saveas(fighndl,[QCpath 'SNR_dB_slicewise' '.jpg'])
close(fighndl)
clear mask_NS info_SNR SNR SNR_in_dB F fighndl dispstring1 dispstring2


F2_=[]; mF2_=[]; C_=[];
for i3 = 1 : siz3
    try waitbar((i3/siz3),wbar3,sprintf('05. QC plots: slice (%d) of (%d)',i3,siz3)); catch, end
    fprintf('05. QC plots: slice (%d) of (%d), subject (%d) of (%d), run (%d) of (%d)\n',i3,siz3,k,size(func_files,1),k2,size(func_files,2))
    scfMRItb_04_resplitData(base_dir_sub, fname, '', siz3) % if slice-wise data doesn't exist but combined 4D data exists, then split the data to generate the needed slice-wise data
    F1 = load_untouch_nii([base_dir_sub fname '_slice' num2str(i3) '.nii']);
    TR = F1.hdr.dime.pixdim(5); F1 = single(squeeze(F1.img));
    
    if Smask_err==0
        scfMRItb_04_resplitData(base_dir_sub, fname, '_Smask', siz3) % if slice-wise data doesn't exist but combined 4D data exists, then split the data to generate the needed slice-wise data
        scfMRItb_04_unzipFile(base_dir_sub, fname, ['_slice' num2str(i3) '_Smask'])
        mask1 = load_untouch_nii([base_dir_sub fname '_slice' num2str(i3) '_Smask.nii']); mask1 = mask1.img;
    else
        mask1 = Smask1_(:,:,i3);
    end
    [m1,m2] = find(mask1==1); Mm1=median(m1); Mm2=median(m2);
    sdist = sqrt((m1-Mm1).^2 + (m2-Mm2).^2);
    clear im Mm1 Mm2 m1 m2
    for i5=1:size(F1,3)
        tmp = F1(:,:,i5);
        F2(:,i5) = tmp(find(mask1==1)); clear tmp
    end; clear i5 F1
    [~,svar] = sort(var(F2,0,2)); [~,sdist] = sort(sdist); sortorder{i3,1} = sdist; %#ok<*AGROW> % sort timeseries based on distance from the center of the cord (if you want to sort by signal variance then use svar)
    
    fighndl = figure('Position',[1 1 1152 720],'Color',[0.95,0.9,0.85],'InvertHardcopy','off','Visible','off'); % red [0.95,0.9,0.85], green [0.85,0.90,0.85], blue [0.89,0.94,0.99]
    imagesc(F2(sortorder{i3},:)),colormap(cmap),colorbar, xlabel('time (in samples)'),ylabel('voxels (sorted by distance from the center of the cord; top row is nearest)'),
        title(['Time series within GM, WM and CSF in UNPROCESSED RAW DATA, in slice ' num2str(i3) ' of ' num2str(siz3)],'Color',[0.65,0.11,0.19],'FontSize',14);
    saveas(fighndl,[QCpath 'fMRI_ts_rawdata_slice' sprintf('%.2d',i3) '.jpg'])
    close(fighndl)
    
    mF2=mean(F2,1); mF2 = 100*(mF2-mean(mF2))./mean(mF2);
    fighndl = figure('Position',[1 1 1152 720],'Color',[0.95,0.9,0.85],'InvertHardcopy','off','Visible','off'); % red [0.95,0.9,0.85], green [0.85,0.90,0.85], blue [0.89,0.94,0.99]
    plot(mF2,'color',[0.65,0.11,0.19],'linewidth',1.5), grid on,grid minor, xlim([1 length(mF2)]),ylim([1.25.*min(mF2) 1.25.*max(mF2)]), xlabel('time (in samples)'),ylabel('percentage signal change'),
      title(['Global mean signal in unprocessed raw data (slice ' num2str(i3) ' of ' num2str(siz3) ')'],'Color',[0.65,0.11,0.19],'FontSize',14),
    saveas(fighndl,[QCpath 'global_signal_rawdata_slice' sprintf('%.2d',i3) '.jpg'])
    close(fighndl)
    
    C = corrcoef(F2'); C=uppertriangle(C);
    fighndl = figure('Position',[1 1 1152 720],'Color',[0.95,0.9,0.85],'InvertHardcopy','off','Visible','off'); % red [0.95,0.9,0.85], green [0.85,0.90,0.85], blue [0.89,0.94,0.99]
    histogram(C,'EdgeColor','none','Normalization','probability'); grid on,grid minor, xlim([-1 1]),  % xlim([-max(abs(C)) max(abs(C))]),
        hold on, xline(0,'Color','k','linewidth',1.5); hold off
        title(sprintf('Histogram of voxel-to-voxel correlations in unprocessed raw data (slice %d of %d)\n%.1f%% of the correlations were > 0  [ mean=%.2f, median=%.2f, SD=%.2f ]',i3,siz3,100*length(find(C>0))/length(C),mean(C),median(C),std(C)),'Color',[0.65,0.11,0.19],'FontSize',14)
    saveas(fighndl,[QCpath 'hist_FC_rawdata_slice' sprintf('%.2d',i3) '.jpg'])
    close(fighndl)
    
    F2_=cat(1,F2_,F2); mF2_=cat(1,mF2_,mF2); C_=cat(1,C_,C);
    clear F2 TR mask1 svar sdist mF2 C
end

[~,svar] = sort(var(F2_,0,2)); [~,spsc] = sort(meanPSC(F2_));
sortorder2 = sortorder; sortorder=[]; f=0; %#ok<*ASGLU>
for im=1:length(sortorder2)
    temp = sortorder2{im}; temp = temp + f; f = f + length(temp);
    sortorder = cat(1,sortorder,temp); clear temp
end; clear im f
sortorder = spsc; % options: spsc for percentage signal change, svar for variance, (comment out the line if you want to use distance from the center of the cord separately for each slice)
for im=max(sortorder):-1:1, if isempty(find(sortorder==im)), sortorder(find(sortorder>im)) = sortorder(find(sortorder>im))-1; end, end; clear im %#ok<*EFIND>

fighndl = figure('Position',[1 1 1152 720],'Color',[0.89,0.94,0.99],'InvertHardcopy','off','Visible','off'); % red [0.95,0.9,0.85], green [0.85,0.90,0.85], blue [0.89,0.94,0.99]
imagesc(F2_(sortorder,:)),colormap(cmap),colorbar, xlabel('time (in samples)'),ylabel('voxels (sorted by mean percentage signal change; bottom row is highest)'),
  title(['Time series within GM, WM and CSF in UNPROCESSED RAW DATA (all slices combined)'],'Color',[0.65,0.11,0.19],'FontSize',14);
saveas(fighndl,[QCpath 'fMRI_ts_rawdata_allSlices' '.jpg'])
 close(fighndl)
mF2_=mean(mF2_,1);
fighndl = figure('Position',[1 1 1152 720],'Color',[0.89,0.94,0.99],'InvertHardcopy','off','Visible','off'); % red [0.95,0.9,0.85], green [0.85,0.90,0.85], blue [0.89,0.94,0.99]
plot(mF2_,'color',[0.65,0.11,0.19],'linewidth',1.5), grid on,grid minor, xlim([1 length(mF2_)]),ylim([1.25.*min(mF2_) 1.25.*max(mF2_)]), xlabel('time (in samples)'),ylabel('percentage signal change'),
  title(['Global mean signal in UNPROCESSED RAW DATA (all slices averaged)'],'Color',[0.65,0.11,0.19],'FontSize',14),
saveas(fighndl,[QCpath 'global_signal_rawdata_allSlices' '.jpg'])
 close(fighndl)
fighndl = figure('Position',[1 1 1152 720],'Color',[0.89,0.94,0.99],'InvertHardcopy','off','Visible','off'); % red [0.95,0.9,0.85], green [0.85,0.90,0.85], blue [0.89,0.94,0.99]
histogram(C_,'EdgeColor','none','Normalization','probability'); grid on,grid minor, xlim([-1 1]),
  hold on, xline(0,'Color','k','linewidth',1.5); hold off
  title(sprintf('Histogram of voxel-to-voxel correlations in UNPROCESSED RAW DATA (all slices combined)\n%.1f%% of the correlations were > 0  [ mean=%.2f, median=%.2f, SD=%.2f ]',100*length(find(C_>0))/length(C_),mean(C_),median(C_),std(C_)),'Color',[0.65,0.11,0.19],'FontSize',14)
saveas(fighndl,[QCpath 'hist_FC_rawdata_allSlices' '.jpg'])
 close(fighndl)
clear svar sortorder spsc F2_ mF2_ C_ sortorder2 spsc

%%% Make QC video (needs audio video toolbox in Matlab)
if ~isempty(which('VideoWriter')) && (save_videos==1) % check if Matlab's audio video toolbox exists

for i3 = 1 : siz3
    try waitbar((i3/siz3),wbar3,sprintf('05. Generate QC video: slice (%d) of (%d)',i3,siz3)); catch, end
    fprintf('05. Generate QC video: slice (%d) of (%d), subject (%d) of (%d), run (%d) of (%d)\n',i3,siz3,k,size(func_files,1),k2,size(func_files,2))
    F1 = load_untouch_nii([base_dir_sub fname '_slice' num2str(i3) '.nii']);
    TR = F1.hdr.dime.pixdim(5); F1 = rot90(single(squeeze(F1.img)));
    video = VideoWriter([QCpath 'fMRIvideo_rawdata_slice' sprintf('%.2d',i3) '.mp4'],'MPEG-4'); %#ok<*TNMLP> % create the video object
    video.FrameRate = round(30/TR); % 30 seconds of data in every second of the video
    video.Quality = 90; % very limited compression so that the images are shown as is (and thus avoid misunderstandings due to video quality)
    open(video); % open the file for writing

    mask1 = load_untouch_nii([base_dir_sub fname '_slice' num2str(i3) '_Smask.nii']);
    mask1 = rot90(mask1.img);
    [xmask,ymask] = find(mask1==1);
    minx=round(min(xmask)-0.1*(size(mask1,1))); maxx=round(max(xmask)+0.1*(size(mask1,1)));
    miny=round(min(ymask)-0.1*(size(mask1,2))); maxy=round(max(ymask)+0.1*(size(mask1,2)));
    if (maxx-minx)<(maxy-miny)
        factr = ((maxy-miny)-(maxx-minx))/2;
        if factr==round(factr)
            maxx=maxx+factr; minx=minx-factr;
        else
            maxx=maxx+factr+0.5; minx=minx-factr+0.5;
        end
    elseif (maxy-miny)<(maxx-minx)
        factr = ((maxx-minx)-(maxy-miny))/2;
        if factr==round(factr)
            maxy=maxy+factr; miny=miny-factr;
        else
            maxy=maxy+factr+0.5; miny=miny-factr+0.5;
        end
    end
    [sizeFx,sizeFy] = size(F1(:,:,1));
    F2 = F1(minx:maxx,miny:maxy,:);
    for i5=1:size(F1,3)
        F1(:,:,i5) = imresize(F2(:,:,i5),[sizeFx sizeFy]);
    end
    clear maxx maxy minx miny xmask ymask mask1 factr sizeFx sizeFy F2 i5
    for ii=1:size(F1,3) % loop across the number of images
        I = F1(:,:,ii); %read the next image
        I(find(isnan(I)))=0;
        I = (I - min(I(:))) ./ (max(I(:)) - min(I(:)));
        fighndl = figure('Position',[round((scrsz(3)-scrsz(4))/2) 1 round(0.75*scrsz(3)) round(0.75*scrsz(4))],'Color',[0.95,0.9,0.85],'InvertHardcopy','off','Visible','off'); % [0.85,0.90,0.85]
        imshow(I, [0 max(I(:))], 'InitialMagnification', 'fit'); title({['Unprocessed raw fMRI data (slice ' num2str(i3) ' of ' num2str(siz3) ')'];['Time point ' num2str(ii) ' of ' num2str(size(F1,3))]},'Color',[0.65,0.11,0.19],'FontSize',18);
        frame = getframe(gcf);
        writeVideo(video,frame); % write the image to file
        close(fighndl)
        clear I frame
    end
    close(video); %close the file
    clear F1 video ii TR
end; clear i3

end
end

%% scfMRItb_06_denoise1
if choices_preproc(6)==1
clear QCpath QCpath2 save_videos fighndl
if ~(exist([base_dir_sub 'QC' '/06_denoise1'],'dir'))
    mkdir([base_dir_sub 'QC' '/06_denoise1'])
end
if Rns==0
    QCpath = [base_dir_sub 'QC' '/06_denoise1/'];
else
    if ~(exist([base_dir_sub 'QC' '/06_denoise1' '/run' num2str(Rns)],'dir'))
        mkdir([base_dir_sub 'QC' '/06_denoise1' '/run' num2str(Rns)])
    end
    QCpath = [base_dir_sub 'QC' '/06_denoise1' '/run' num2str(Rns) '/'];
end
cutoff1_str = '80';

if (save_videos_==0)||(save_videos_==1)
    save_videos = save_videos_;
elseif (save_videos_==2)&&(sum([choices_preproc(8:9);choices_preproc(11:14)])==0)
    save_videos = 1;
else
    save_videos = 0;
end
if save_videos==1
    save_videos_steps(2)=1;
end

F2_=[]; mF2_=[]; C_=[]; F4_=[]; mF4_=[]; C2_=[];
for i3 = 1 : siz3
    try waitbar((i3/siz3),wbar3,sprintf('06. QC plots: slice (%d) of (%d)',i3,siz3)); catch, end
    fprintf('06. QC plots: slice (%d) of (%d), subject (%d) of (%d), run (%d) of (%d)\n',i3,siz3,k,size(func_files,1),k2,size(func_files,2))
    scfMRItb_04_resplitData(base_dir_sub, fname, ['_denoised_before_MC' cutoff1_str], siz3) % if slice-wise data doesn't exist but combined 4D data exists, then split the data to generate the needed slice-wise data
    scfMRItb_04_unzipFile(base_dir_sub, fname, ['_slice' num2str(i3) '_denoised_before_MC' cutoff1_str])
    F1 = load_untouch_nii([base_dir_sub fname '_slice' num2str(i3) '_denoised_before_MC' cutoff1_str '.nii']);
    TR = F1.hdr.dime.pixdim(5); F1 = single(squeeze(F1.img));
    
    if Smask_err==0
        scfMRItb_04_resplitData(base_dir_sub, fname, '_Smask', siz3) % if slice-wise data doesn't exist but combined 4D data exists, then split the data to generate the needed slice-wise data
        scfMRItb_04_unzipFile(base_dir_sub, fname, ['_slice' num2str(i3) '_Smask'])
        mask1 = load_untouch_nii([base_dir_sub fname '_slice' num2str(i3) '_Smask.nii']); mask1 = mask1.img;
    else
        mask1 = Smask1_(:,:,i3);
    end
    [m1,m2] = find(mask1==1); Mm1=median(m1); Mm2=median(m2);
    sdist = sqrt((m1-Mm1).^2 + (m2-Mm2).^2);
    clear im Mm1 Mm2 m1 m2
    for i5=1:size(F1,3)
        tmp = F1(:,:,i5);
        F2(:,i5) = tmp(find(mask1==1)); clear tmp
    end; clear i5 F1
    [~,svar] = sort(var(F2,0,2)); [~,sdist] = sort(sdist); sortorder{i3,1} = sdist; %#ok<*AGROW> % sort timeseries based on distance from the center of the cord (if you want to sort by signal variance then use svar)
    
    fighndl = figure('Position',[1 1 1152 720],'Color',[0.95,0.9,0.85],'InvertHardcopy','off','Visible','off'); % red [0.95,0.9,0.85], green [0.85,0.90,0.85], blue [0.89,0.94,0.99]
    imagesc(F2(sortorder{i3},:)),colormap(cmap),colorbar, xlabel('time (in samples)'),ylabel('voxels (sorted by distance from the center of the cord; top row is nearest)'),
        title(['Time series within GM, WM and CSF after "06-denoise1" in slice ' num2str(i3) ' of ' num2str(siz3)],'Color',[0,0.55,0.69],'FontSize',14);
    saveas(fighndl,[QCpath 'fMRI_ts_06denoise1_slice' sprintf('%.2d',i3) '.jpg'])
    close(fighndl)
    
    scfMRItb_04_resplitData(base_dir_sub, fname, '', siz3) % if slice-wise data doesn't exist but combined 4D data exists, then split the data to generate the needed slice-wise data
    scfMRItb_04_unzipFile(base_dir_sub, fname, ['_slice' num2str(i3)])
    F3 = load_untouch_nii([base_dir_sub fname '_slice' num2str(i3) '.nii']); F3 = single(squeeze(F3.img));
    for i5=1:size(F3,3)
        tmp = F3(:,:,i5);
        F4(:,i5) = tmp(find(mask1==1)); clear tmp
    end; clear i5 F3
    mF2=mean(F2,1); mF2 = 100*(mF2-mean(mF2))./mean(mF2); mF4=mean(F4,1); mF4 = 100*(mF4-mean(mF4))./mean(mF4);
    fighndl = figure('Position',[1 1 1152 720],'Color',[0.95,0.9,0.85],'InvertHardcopy','off','Visible','off'); % red [0.95,0.9,0.85], green [0.85,0.90,0.85], blue [0.89,0.94,0.99]
    subplot2n(2,1,1), plot(mF4,'color',[0.65,0.11,0.19],'linewidth',1.5), hold on, plot(mF2,'color',[0,0.55,0.69],'linewidth',0.2), xlabel('time (in samples)'),ylabel('percentage signal change'),
      grid on,grid minor, xlim([1 length(mF4)]),ylim([1.25.*min([mF2,mF4]) 1.25.*max([mF2,mF4])]), title(['Global mean signal in unprocessed raw data (slice ' num2str(i3) ' of ' num2str(siz3) ')'],'Color',[0.65,0.11,0.19],'FontSize',14), legend({'Global signal (raw data)'},'FontSize',12,'TextColor',[0.65,0.11,0.19],'Location','best')
    subplot2n(2,1,2), plot(mF2,'color',[0,0.55,0.69],'linewidth',1.5), hold on, plot(mF4,'color',[0.65,0.11,0.19],'linewidth',0.2), xlabel('time (in samples)'),ylabel('percentage signal change'),
      grid on,grid minor, xlim([1 length(mF4)]),ylim([1.25.*min([mF2,mF4]) 1.25.*max([mF2,mF4])]), title(['Global mean signal after "06-denoise1" (slice ' num2str(i3) ' of ' num2str(siz3) ')'],'Color',[0,0.55,0.69],'FontSize',14), legend({'Global signal after 06-denoise1'},'FontSize',12,'TextColor',[0,0.55,0.69],'Location','best')
    saveas(fighndl,[QCpath 'global_signal_06denoise1_slice' sprintf('%.2d',i3) '.jpg'])
    close(fighndl)

    C  = corrcoef(F2'); C  = uppertriangle(C) ;
    C2 = corrcoef(F4'); C2 = uppertriangle(C2);
    fighndl = figure('Position',[1 1 1152 720],'Color',[0.95,0.9,0.85],'InvertHardcopy','off','Visible','off'); % red [0.95,0.9,0.85], green [0.85,0.90,0.85], blue [0.89,0.94,0.99]
    subplot2n(2,1,1), histogram(C2,'EdgeColor','none','FaceColor',[0.65,0.11,0.19],'FaceAlpha',0.5,'Normalization','probability'); grid on,grid minor, xlim([-1 1]), hold on, xline(0,'Color','k','linewidth',1.5); hold off
      title(sprintf('Histogram of voxel-to-voxel correlations in UNPROCESSED RAW DATA (slice %d of %d)\n%.1f%% of the correlations were > 0  [ mean=%.2f, median=%.2f, SD=%.2f ]',i3,siz3,100*length(find(C2>0))/length(C2),mean(C2),median(C2),std(C2)),'Color',[0.65,0.11,0.19],'FontSize',14)
    subplot2n(2,1,2), histogram(C,'EdgeColor','none','FaceColor',[0,0.55,0.69],'FaceAlpha',0.5,'Normalization','probability'); grid on,grid minor, xlim([-1 1]), hold on, xline(0,'Color','k','linewidth',1.5); hold off
      title(sprintf('Histogram of voxel-to-voxel correlations AFTER 06-DENOISE1 (slice %d of %d)\n%.1f%% of the correlations were > 0  [ mean=%.2f, median=%.2f, SD=%.2f ]',i3,siz3,100*length(find(C>0))/length(C),mean(C),median(C),std(C)),'Color',[0,0.55,0.69],'FontSize',14)
    saveas(fighndl,[QCpath 'hist_FC_compare_06denoise1_slice' sprintf('%.2d',i3) '.jpg'])
    close(fighndl)
    
    F2_=cat(1,F2_,F2); mF2_=cat(1,mF2_,mF2); C_=cat(1,C_,C); F4_=cat(1,F4_,F4); mF4_=cat(1,mF4_,mF4); C2_=cat(1,C2_,C2);
    clear F2 F4 TR mask1 svar sdist mF2 mF4 C C2
end

[~,svar] = sort(var(F2_,0,2)); [~,spsc] = sort(meanPSC(F2_));
sortorder2 = sortorder; sortorder=[]; f=0; %#ok<*ASGLU>
for im=1:length(sortorder2)
    temp = sortorder2{im}; temp = temp + f; f = f + length(temp);
    sortorder = cat(1,sortorder,temp); clear temp
end; clear im f
sortorder = spsc; % options: spsc for percentage signal change, svar for variance, (comment out the line if you want to use distance from the center of the cord separately for each slice)
for im=max(sortorder):-1:1, if isempty(find(sortorder==im)), sortorder(find(sortorder>im)) = sortorder(find(sortorder>im))-1; end, end; clear im %#ok<*EFIND>

fighndl = figure('Position',[1 1 1152 720],'Color',[0.89,0.94,0.99],'InvertHardcopy','off','Visible','off'); % red [0.95,0.9,0.85], green [0.85,0.90,0.85], blue [0.89,0.94,0.99]
imagesc(F2_(sortorder,:)),colormap(cmap),colorbar, xlabel('time (in samples)'),ylabel('voxels (sorted by mean percentage signal change; bottom row is highest)'),
  title(['Time series within GM, WM and CSF AFTER 06-DENOISE1 (all slices combined)'],'Color',[0,0.55,0.69],'FontSize',14);
saveas(fighndl,[QCpath 'fMRI_ts_06denoise1_allSlices' '.jpg'])
 close(fighndl)
mF2_=mean(mF2_,1); mF4_=mean(mF4_,1);
fighndl = figure('Position',[1 1 1152 720],'Color',[0.89,0.94,0.99],'InvertHardcopy','off','Visible','off'); % red [0.95,0.9,0.85], green [0.85,0.90,0.85], blue [0.89,0.94,0.99]
subplot2n(2,1,1), plot(mF4_,'color',[0.65,0.11,0.19],'linewidth',1.5), hold on, plot(mF2_,'color',[0,0.55,0.69],'linewidth',0.2), xlabel('time (in samples)'),ylabel('percentage signal change'),
  grid on,grid minor, xlim([1 length(mF4_)]),ylim([1.25.*min([mF2_,mF4_]) 1.25.*max([mF2_,mF4_])]), title(['Global mean signal in UNPROCESSED RAW DATA (all slices averaged)'],'Color',[0.65,0.11,0.19],'FontSize',14), legend({'Global signal (raw data)'},'FontSize',12,'TextColor',[0.65,0.11,0.19],'Location','best')
subplot2n(2,1,2), plot(mF2_,'color',[0,0.55,0.69],'linewidth',1.5), hold on, plot(mF4_,'color',[0.65,0.11,0.19],'linewidth',0.2), xlabel('time (in samples)'),ylabel('percentage signal change'),
  grid on,grid minor, xlim([1 length(mF2_)]),ylim([1.25.*min([mF2_,mF4_]) 1.25.*max([mF2_,mF4_])]), title(['Global mean signal AFTER 06-DENOISE1 (all slices averaged)'],'Color',[0,0.55,0.69],'FontSize',14), legend({'Global signal after 06-denoise1'},'FontSize',12,'TextColor',[0,0.55,0.69],'Location','best')
saveas(fighndl,[QCpath 'global_signal_06denoise1_allSlices' '.jpg'])
 close(fighndl)
fighndl = figure('Position',[1 1 1152 720],'Color',[0.89,0.94,0.99],'InvertHardcopy','off','Visible','off'); % red [0.95,0.9,0.85], green [0.85,0.90,0.85], blue [0.89,0.94,0.99]
subplot2n(2,1,1), histogram(C2_,'EdgeColor','none','FaceColor',[0.65,0.11,0.19],'FaceAlpha',0.5,'Normalization','probability'); grid on,grid minor, xlim([-1 1]), hold on, xline(0,'Color','k','linewidth',1.5); hold off
 title(sprintf('Histogram of voxel-to-voxel correlations in UNPROCESSED RAW DATA (all slices combined)\n%.1f%% of the correlations were > 0  [ mean=%.2f, median=%.2f, SD=%.2f ]',100*length(find(C2_>0))/length(C2_),mean(C2_),median(C2_),std(C2_)),'Color',[0.65,0.11,0.19],'FontSize',14)
subplot2n(2,1,2), histogram(C_,'EdgeColor','none','FaceColor',[0,0.55,0.69],'FaceAlpha',0.5,'Normalization','probability'); grid on,grid minor, xlim([-1 1]), hold on, xline(0,'Color','k','linewidth',1.5); hold off
 title(sprintf('Histogram of voxel-to-voxel correlations AFTER 06-DENOISE1 (all slices combined)\n%.1f%% of the correlations were > 0  [ mean=%.2f, median=%.2f, SD=%.2f ]',100*length(find(C_>0))/length(C_),mean(C_),median(C_),std(C_)),'Color',[0,0.55,0.69],'FontSize',14)
saveas(fighndl,[QCpath 'hist_FC_compare_06denoise1_allSlices' '.jpg'])
 close(fighndl)
clear svar sortorder spsc F2_ mF2_ C_ F4_ mF4_ C2_

%%% Make QC video (needs audio video toolbox in Matlab)
if ~isempty(which('VideoWriter')) && (save_videos==1) % check if Matlab's audio video toolbox exists

subplot = @(m,n,p) subtightplot(m, n, p, [0.001 0.005], [0.01 0.001], [0.01 0.001]);
for i3 = 1 : siz3
    try waitbar((i3/siz3),wbar3,sprintf('06. Generate QC video: slice (%d) of (%d)',i3,siz3)); catch, end
    fprintf('06. Generate QC video: slice (%d) of (%d), subject (%d) of (%d), run (%d) of (%d)\n',i3,siz3,k,size(func_files,1),k2,size(func_files,2))
    scfMRItb_04_resplitData(base_dir_sub, fname, ['_denoised_before_MC' cutoff1_str], siz3) % if slice-wise data doesn't exist but combined 4D data exists, then split the data to generate the needed slice-wise data
    scfMRItb_04_unzipFile(base_dir_sub, fname, ['_slice' num2str(i3) '_denoised_before_MC' cutoff1_str])
    F1 = load_untouch_nii([base_dir_sub fname '_slice' num2str(i3) '_denoised_before_MC' cutoff1_str '.nii']);
    TR = F1.hdr.dime.pixdim(5); F1 = rot90(single(squeeze(F1.img)));
    video = VideoWriter([QCpath 'fMRIvideo_06denoise1_slice' sprintf('%.2d',i3) '.mp4'],'MPEG-4'); %#ok<*TNMLP> % create the video object
    video.FrameRate = round(30/TR); % 30 seconds of data in every second of the video
    video.Quality = 90; % very limited compression so that the images are shown as is (and thus avoid misunderstandings due to video quality)
    open(video); % open the file for writing
    
    if Smask_err==0
        scfMRItb_04_resplitData(base_dir_sub, fname, '_Smask', siz3) % if slice-wise data doesn't exist but combined 4D data exists, then split the data to generate the needed slice-wise data
        scfMRItb_04_unzipFile(base_dir_sub, fname, ['_slice' num2str(i3) '_Smask'])
        mask1 = load_untouch_nii([base_dir_sub fname '_slice' num2str(i3) '_Smask.nii']); mask1 = mask1.img;
    else
        mask1 = Smask1_(:,:,i3);
    end
    mask1 = rot90(mask1);
    [xmask,ymask] = find(mask1==1);
    minx=round(min(xmask)-0.1*(size(mask1,1))); maxx=round(max(xmask)+0.1*(size(mask1,1)));
    miny=round(min(ymask)-0.1*(size(mask1,2))); maxy=round(max(ymask)+0.1*(size(mask1,2)));
    if (maxx-minx)<(maxy-miny)
        factr = ((maxy-miny)-(maxx-minx))/2;
        if factr==round(factr)
            maxx=maxx+factr; minx=minx-factr;
        else
            maxx=maxx+factr+0.5; minx=minx-factr+0.5;
        end
    elseif (maxy-miny)<(maxx-minx)
        factr = ((maxx-minx)-(maxy-miny))/2;
        if factr==round(factr)
            maxy=maxy+factr; miny=miny-factr;
        else
            maxy=maxy+factr+0.5; miny=miny-factr+0.5;
        end
    end
    [sizeFx,sizeFy] = size(F1(:,:,1));
    F2 = F1(minx:maxx,miny:maxy,:);
    for i5=1:size(F1,3)
        F1(:,:,i5) = imresize(F2(:,:,i5),[sizeFx sizeFy]);
    end
    scfMRItb_04_resplitData(base_dir_sub, fname, '', siz3) % if slice-wise data doesn't exist but combined 4D data exists, then split the data to generate the needed slice-wise data
    F3 = load_untouch_nii([base_dir_sub fname '_slice' num2str(i3) '.nii']); F3 = rot90(single(squeeze(F3.img)));
    F4 = F3(minx:maxx,miny:maxy,:);
    for i5=1:size(F3,3)
        F3(:,:,i5) = imresize(F4(:,:,i5),[sizeFx sizeFy]);
    end
    clear maxx maxy minx miny xmask ymask mask1 factr sizeFx sizeFy F2 F4 i5
    
    for ii=1:size(F1,3) % loop across the number of images
        I = F1(:,:,ii); %read the next image
        I(find(isnan(I)))=0;
        I = (I - min(I(:))) ./ (max(I(:)) - min(I(:)));
        I2 = F3(:,:,ii); %read the next image
        I2(find(isnan(I2)))=0;
        I2 = (I2 - min(I2(:))) ./ (max(I2(:)) - min(I2(:)));
        fighndl = figure('Position',[round((scrsz(3)-scrsz(4))/2) 1 round(0.75*scrsz(3)) round(0.75*scrsz(4))],'Color',[0.95,0.9,0.85],'InvertHardcopy','off','Visible','off'); % [0.85,0.90,0.85]
        subplot(1,2,1), imshow(I2, [0 max(I2(:))], 'InitialMagnification', 'fit'); title({['Unprocessed raw fMRI data (slice ' num2str(i3) ' of ' num2str(siz3) ')'];['Time point ' num2str(ii) ' of ' num2str(size(F1,3))]},'Color',[0.65,0.11,0.19],'FontSize',18);
        subplot(1,2,2), imshow(I, [0 max(I(:))], 'InitialMagnification', 'fit'); title({['fMRI data after "06. denoise1" (slice ' num2str(i3) ' of ' num2str(siz3) ')'];['Time point ' num2str(ii) ' of ' num2str(size(F1,3))]},'Color',[0,0.55,0.69],'FontSize',18);
        frame = getframe(gcf);
        writeVideo(video,frame); % write the image to file
        close(fighndl)
        clear I I2 frame
    end
    close(video); %close the file
    clear F1 F3 video ii TR
end
clear i3 cutoff1_str

end
end

%% scfMRItb_07_gaussianMask
if choices_preproc(7)==1
clear QCpath QCpath2 save_videos fighndl
if ~(exist([base_dir_sub 'QC' '/07_gaussianMask'],'dir'))
    mkdir([base_dir_sub 'QC' '/07_gaussianMask'])
end
if Rns==0
    QCpath = [base_dir_sub 'QC' '/07_gaussianMask/'];
else
    if ~(exist([base_dir_sub 'QC' '/07_gaussianMask' '/run' num2str(Rns)],'dir'))
        mkdir([base_dir_sub 'QC' '/07_gaussianMask' '/run' num2str(Rns)])
    end
    QCpath = [base_dir_sub 'QC' '/07_gaussianMask' '/run' num2str(Rns) '/'];
end

try
    Gmask = load_untouch_nii([base_dir_sub fname '_Gaussian_mask.nii']);
    for i3 = 1 : siz3
        try waitbar((i3/siz3),wbar3,sprintf('07. Compute Gaussian mask: slice (%d) of (%d)',i3,siz3)); catch, end
        scfMRItb_04_resplitData(base_dir_sub, fname, '_base', siz3) % if slice-wise data doesn't exist but combined 4D data exists, then split the data to generate the needed slice-wise data
        scfMRItb_04_unzipFile(base_dir_sub, fname, ['_slice' num2str(i3) '_base'])
        F = load_untouch_nii([base_dir_sub fname '_slice' num2str(i3) '_base.nii']);
        fighndl = figure('Position',[round((scrsz(3)-scrsz(4))/2) 1 scrsz(4) scrsz(4)],'InvertHardcopy','off','Visible','off');
        figmriF(single(F.img) .* Gmask(:,:,i3).img); % briefly display masked image
        title(sprintf('Gaussian masked mean functional image - slice %d of %d',i3,siz3),'FontSize',14)
        saveas(fighndl,[QCpath 'MeanFunc_GaussianMasked_slice' sprintf('%.2d',i3) '.jpg'])
        close(fighndl)
        clear F
    end
    
catch
    scfMRItb_04_unzipFile(base_dir_sub, fname3, '')
    F = load_untouch_nii([base_dir_sub fname3 '.nii']); % mean functional
    Gmask = F; % Gaussian mask
    new_size = round(size(F.img,1) / 2);
    clear F
    load([base_dir_sub fname '_mask_NS.mat'])
    mask_NS = mask_NS.img;
    for i3 = 1 : siz3
        scfMRItb_04_resplitData(base_dir_sub, fname, '_base', siz3) % if slice-wise data doesn't exist but combined 4D data exists, then split the data to generate the needed slice-wise data
        scfMRItb_04_unzipFile(base_dir_sub, fname, ['_slice' num2str(i3) '_base'])
        F = load_untouch_nii([base_dir_sub fname '_slice' num2str(i3) '_base.nii']);
        img = mask_NS(:,:,i3);
        [i1,i2] = find(img==0);
        x_center = round(median(i2)); % in imagesc display, the vertical axis is the x-axis, hence i2 is used instead of i1
        y_center = round(median(i1)); % in imagesc display, the horizontal axis is the y-axis, hence i1 is used instead of i2
        x_right  = max(i2);        x_left   = min(i2);
        y_top    = min(i1);        y_bottom = max(i1);
        x_fwhm = round(abs(x_right-x_left)/2); y_fwhm = round(abs(y_top-y_bottom)/2);
        x1 = repmat(((1:1:size(F.img,1))'-y_center),[1 size(F.img,2)]);
        y1 = repmat(((1:1:size(F.img,2))-x_center),[size(F.img,1) 1]);
        W = single(gauss2D_R(x1, y1, 2*y_fwhm, 2*x_fwhm, 1, 1));
        % suppress voxels less than 0.1
        min_weight = 0.1;
        W = W .* (W > min_weight);

        Gmask.img(:,:,i3) = W;
        fighndl = figure('Position',[round((scrsz(3)-scrsz(4))/2) 1 scrsz(4) scrsz(4)],'InvertHardcopy','off','Visible','off'); try addToolbarExplorationButtons(fighndl); catch, end
        figmriF(single(F.img) .* W); % briefly display masked image
        title(sprintf('Gaussian masked mean functional image - slice %d of %d',i3,siz3),'FontSize',14)
        saveas(fighndl,[QCpath 'MeanFunc_GaussianMasked_slice' sprintf('%.2d',i3) '.jpg'])
        close(fighndl)
        clear x_center y_center x_right y_right x_top y_top x_left y_bottom x_fwhm y_fwhm x1 y1 W min_weight F img
    end
    clear new_size img temp mmax fighndl;
    
    unix(['rm -f ' base_dir_sub fname '_Gaussian_mask.nii']);
    save_untouch_nii(Gmask, [base_dir_sub fname '_Gaussian_mask.nii']);
    clear W Gmask x1 y1;
    for i3 = 1 : siz3
        unix(['rm -f ' base_dir_sub fname '_slice' num2str(i3) '_Gaussian_mask.nii']);
        unix(['3dZcutup -keep ' num2str(i3-1) ' ' num2str(i3-1) ' ' ...
            '-prefix ' base_dir_sub fname '_slice' num2str(i3) '_Gaussian_mask.nii' ' ' ...
            base_dir_sub fname '_Gaussian_mask.nii']);
    end
end

clear fighndl Gmask

end

%% scfMRItb_08_motionCorrection
if choices_preproc(8)==1
clear QCpath save_videos fighndl
if ~(exist([base_dir_sub 'QC' '/08_motionCorrection'],'dir'))
    mkdir([base_dir_sub 'QC' '/08_motionCorrection'])
end
if Rns==0
    QCpath = [base_dir_sub 'QC' '/08_motionCorrection/'];
else
    if ~(exist([base_dir_sub 'QC' '/08_motionCorrection' '/run' num2str(Rns)],'dir'))
        mkdir([base_dir_sub 'QC' '/08_motionCorrection' '/run' num2str(Rns)])
    end
    QCpath = [base_dir_sub 'QC' '/08_motionCorrection' '/run' num2str(Rns) '/'];
end

if (save_videos_==0)||(save_videos_==1)
    save_videos = save_videos_;
elseif (save_videos_==2)&&(sum([choices_preproc(9);choices_preproc(11:14)])==0)
    save_videos = 1;
else
    save_videos = 0;
end
if save_videos==1
    save_videos_steps(3)=1;
end


try
    load([base_dir_sub 'motion_parameters.mat'],'motion_parameters','FD','AD','mean_FD_perslice','mean_FD_total','max_FD_perslice','max_FD_total')
catch
    for i3=1:siz3
        motion_parameters{i3,1} = importdata([base_dir_sub fname '_slice' num2str(i3) '_MC_params.txt']);
        motion_parameters{i3} = motion_parameters{i3}.data(:,1:2); motion_parameters{i3}(:,3:6)=0; % motion_parameters is a matrix of size Nx6 (N = no. of time points; 6 = 3 translation and 3 rotation parameters). We have only x and y translations taken into account. The rest of the degrees of freedom are zero padded.
        [FD(:,i3), mean_FD_perslice(i3,1), AD(:,i3)] = FD_metric2(motion_parameters{i3},50);
        max_FD_perslice(i3,1) = max(FD(:,i3));
        motion_parameters{i3}(:,3:end)=[];
    end; clear i3
    mean_FD_total = mean(mean_FD_perslice);
    max_FD_total = max(max_FD_perslice);
    readme_framewise_displacement = sprintf('FD: Framewise Displacement; mean_FD: mean FD (one value per run); max_FD: max FD (one value per run); AD: Absolute Displacement\nmotion_parameters: a matrix of size Nx6 (N = no. of time points; 6 = 3 translation and 3 rotation parameters). We have only x and y translations taken into account. The rest of the degrees of freedom are zero padded.\nAll values in millimeters\nFor more info, type: help FD_metric2\n');
    save([base_dir_sub 'motion_parameters.mat'],'motion_parameters','FD','AD','mean_FD_perslice','mean_FD_total','max_FD_perslice','max_FD_total','readme_framewise_displacement')
    % Move motion parameters (and transformations) to a subdirectory within the subject's main directory
    sdirm = cat(1,dir([base_dir_sub fname '*' 'MC_params.txt']),dir([base_dir_sub fname '*' '_MC_xform.aff12.1D']),dir([base_dir_sub fname '*' '_MCfilt_xform.aff12.1D']));
    mkdir([base_dir_sub 'motion_params'])
    for i5=1:length(sdirm)
        movefile([sdirm(i5).folder '/' sdirm(i5).name],[base_dir_sub 'motion_params' '/' sdirm(i5).name])
    end; clear sdirm i5
end
M_pool=[];
for i3=1:siz3
    M_pool = cat(1,M_pool,motion_parameters{i3}(:));
end; clear i3
M = motion_parameters;
fighndl = figure('Position',[round((scrsz(3)-scrsz(4))/2) 1 scrsz(4) scrsz(4)],'Color',[0.85,0.90,0.85],'InvertHardcopy','off','Visible','off'); % [0.85,0.90,0.85] [0.8, 0.89, 0.94]
for i3=1:siz3
    subplot2n(siz3,2,2*i3-1), plot(M{i3}(:,1),'b','linewidth',1.5), xlim([1 length(M{i3}(:,1))]),ylim([min(M_pool) max(M_pool)]), grid on, ylabel('x'),title(['(slice ' num2str(i3) ')'])
    subplot2n(siz3,2,2*i3),   plot(M{i3}(:,2),'r','linewidth',1.5), xlim([1 length(M{i3}(:,2))]),ylim([min(M_pool) max(M_pool)]), grid on, ylabel('y'),title(['(slice ' num2str(i3) ')'])
end; clear i3
try suptitle2(['X and Y translational motion parameters (in mm) for slices ' num2str(1) ' through ' num2str(siz3)]); fighndl.Visible='off'; catch, end
saveas(fighndl,[QCpath 'motion_params' '.jpg'])
clf
for i3=1:siz3
    subplot2n(siz3,1,i3), plot(FD(:,i3),'Color',[0,0.5+0.25*i3/siz3,1-0.25*i3/siz3],'linewidth',1.5), xlim([1 length(FD(:,i3))]),ylim([min(FD(:)) max(FD(:))]), grid on, ylabel('FD'),title(['(slice ' num2str(i3) ')'])
end; clear i3
try suptitle2(['Framewise displacement (i.e. relative motion) (in mm) for slices ' num2str(1) ' through ' num2str(siz3)]); fighndl.Visible='off'; catch, end
saveas(fighndl,[QCpath 'relative_displacement' '.jpg'])
clf
for i3=1:siz3
    subplot2n(siz3,1,i3), plot(AD(:,i3),'Color',[0.5+0.25*i3/siz3,0,1-0.25*i3/siz3],'linewidth',1.5), xlim([1 length(AD(:,i3))]),ylim([min(AD(:)) max(AD(:))]), grid on, ylabel('AD'),title(['(slice ' num2str(i3) ')'])
end; clear i3
try suptitle2(['Absolute displacement (in mm) for slices ' num2str(1) ' through ' num2str(siz3)]); fighndl.Visible='off'; catch, end
saveas(fighndl,[QCpath 'absolute_displacement' '.jpg'])
close(fighndl)
fighndl = figure('Position',[round((scrsz(3)-scrsz(4))/2) 1 scrsz(4) scrsz(4)],'Color',[0.85,0.90,0.85],'InvertHardcopy','off','Visible','off'); % [0.85,0.90,0.85] [0.8, 0.89, 0.94]
subplot2n(2,1,1), b = bar(mean_FD_perslice,0.4,'FaceColor',[0,0.55,0.69],'EdgeColor',[0.65,0.11,0.19],'LineWidth',1.5);
  xtips1 = b(1).XEndPoints; ytips1 = b(1).YEndPoints; labels1 = string(round(b(1).YData,2)); text(xtips1,ytips1,labels1,'HorizontalAlignment','center','VerticalAlignment','bottom'); clear b xtips ytips labels1
  grid on, grid minor, ylim([0 max(mean_FD_perslice)+0.01]), ylabel('Mean framewise displacement'), xlabel('Slices'), title(['Mean framewise displacement (in mm) for slices ' num2str(1) ' through ' num2str(siz3) '  [ mean across all slices = ' sprintf('%.2f',mean_FD_total) ' mm ]'],'Color',[0,0.55,0.69],'FontSize',18);
subplot2n(2,1,2), b = bar(max_FD_perslice,0.4,'FaceColor',[0,0.55,0.69],'EdgeColor',[0.65,0.11,0.19],'LineWidth',1.5);
  xtips1 = b(1).XEndPoints; ytips1 = b(1).YEndPoints; labels1 = string(round(b(1).YData,2)); text(xtips1,ytips1,labels1,'HorizontalAlignment','center','VerticalAlignment','bottom'); clear b xtips ytips labels1
  grid on, grid minor, ylim([0 max(max_FD_perslice)+0.05]), ylabel('Max framewise displacement'), xlabel('Slices'), title(['Max framewise displacement (in mm) for slices ' num2str(1) ' through ' num2str(siz3) '  [ max across all slices = ' sprintf('%.2f',max_FD_total) ' mm ]'],'Color',[0.65,0.11,0.19],'FontSize',18);
saveas(fighndl,[QCpath 'mean_max_FD' '.jpg'])
close(fighndl)
clear M motion_parameters FD AD mean_FD_perslice mean_FD_total max_FD_perslice max_FD_total


F2_=[]; mF2_=[]; C_=[]; F4_=[]; mF4_=[]; C2_=[];
if exist([base_dir_sub fname '_slice' num2str(1) '_denoised_before_MC' cutoff1_str '.nii'],'file') || exist([base_dir_sub fname '_slice' num2str(1) '_denoised_before_MC' cutoff1_str '.nii.gz'],'file') || exist([base_dir_sub fname '_denoised_before_MC' cutoff1_str '.nii'],'file') || exist([base_dir_sub fname '_denoised_before_MC' cutoff1_str '.nii.gz'],'file')
    F6_=[]; mF6_=[]; C3_=[];
end
for i3 = 1 : siz3
    try waitbar((i3/siz3),wbar3,sprintf('08. QC plots: slice (%d) of (%d)',i3,siz3)); catch, end
    fprintf('08. QC plots: slice (%d) of (%d), subject (%d) of (%d), run (%d) of (%d)\n',i3,siz3,k,size(func_files,1),k2,size(func_files,2))
    scfMRItb_04_resplitData(base_dir_sub, fname, '_MC', siz3) % if slice-wise data doesn't exist but combined 4D data exists, then split the data to generate the needed slice-wise data
    scfMRItb_04_unzipFile(base_dir_sub, fname, ['_slice' num2str(i3) '_MC'])
    F1 = load_untouch_nii([base_dir_sub fname '_slice' num2str(i3) '_MC.nii']);
    TR = F1.hdr.dime.pixdim(5); F1 = single(squeeze(F1.img));
    
    if Smask_err==0
        scfMRItb_04_resplitData(base_dir_sub, fname, '_Smask', siz3) % if slice-wise data doesn't exist but combined 4D data exists, then split the data to generate the needed slice-wise data
        scfMRItb_04_unzipFile(base_dir_sub, fname, ['_slice' num2str(i3) '_Smask'])
        mask1 = load_untouch_nii([base_dir_sub fname '_slice' num2str(i3) '_Smask.nii']); mask1 = mask1.img;
    else
        mask1 = Smask1_(:,:,i3);
    end
    [m1,m2] = find(mask1==1); Mm1=median(m1); Mm2=median(m2);
    sdist = sqrt((m1-Mm1).^2 + (m2-Mm2).^2);
    clear im Mm1 Mm2 m1 m2
    for i5=1:size(F1,3)
        tmp = F1(:,:,i5);
        F2(:,i5) = tmp(find(mask1==1)); clear tmp
    end; clear i5 F1
    [~,svar] = sort(var(F2,0,2)); [~,sdist] = sort(sdist); sortorder{i3,1} = sdist; %#ok<*AGROW> % sort timeseries based on distance from the center of the cord (if you want to sort by signal variance then use svar)
    
    fighndl = figure('Position',[1 1 1152 720],'Color',[0.95,0.9,0.85],'InvertHardcopy','off','Visible','off'); % red [0.95,0.9,0.85], green [0.85,0.90,0.85], blue [0.89,0.94,0.99]
    imagesc(F2(sortorder{i3},:)),colormap(cmap),colorbar, xlabel('time (in samples)'),ylabel('voxels (sorted by distance from the center of the cord; top row is nearest)'),
        title(['Time series within GM, WM and CSF after "08-motion-correction" in slice ' num2str(i3) ' of ' num2str(siz3)],'Color',[0,0.55,0.69],'FontSize',14);
    saveas(fighndl,[QCpath 'fMRI_ts_08moco_slice' sprintf('%.2d',i3) '.jpg'])
    close(fighndl)
    
    scfMRItb_04_resplitData(base_dir_sub, fname, '', siz3) % if slice-wise data doesn't exist but combined 4D data exists, then split the data to generate the needed slice-wise data
    scfMRItb_04_unzipFile(base_dir_sub, fname, ['_slice' num2str(i3)])
    F3 = load_untouch_nii([base_dir_sub fname '_slice' num2str(i3) '.nii']); F3 = single(squeeze(F3.img));
    for i5=1:size(F3,3)
        tmp = F3(:,:,i5);
        F4(:,i5) = tmp(find(mask1==1)); clear tmp
    end; clear i5 F3
    mF2=mean(F2,1); mF2 = 100*(mF2-mean(mF2))./mean(mF2); mF4=mean(F4,1); mF4 = 100*(mF4-mean(mF4))./mean(mF4);
    fighndl = figure('Position',[1 1 1152 720],'Color',[0.95,0.9,0.85],'InvertHardcopy','off','Visible','off'); % red [0.95,0.9,0.85], green [0.85,0.90,0.85], blue [0.89,0.94,0.99]
    subplot2n(2,1,1), plot(mF4,'color',[0.65,0.11,0.19],'linewidth',1.5), hold on, plot(mF2,'color',[0,0.55,0.69],'linewidth',0.2), xlabel('time (in samples)'),ylabel('percentage signal change'),
      grid on,grid minor, xlim([1 length(mF4)]),ylim([1.25.*min([mF2,mF4]) 1.25.*max([mF2,mF4])]), title(['Global mean signal in UNPROCESSED RAW DATA (slice ' num2str(i3) ' of ' num2str(siz3) ')'],'Color',[0.65,0.11,0.19],'FontSize',14), legend({'Global signal (raw data)'},'FontSize',12,'TextColor',[0.65,0.11,0.19],'Location','best')
    subplot2n(2,1,2), plot(mF2,'color',[0,0.55,0.69],'linewidth',1.5), hold on, plot(mF4,'color',[0.65,0.11,0.19],'linewidth',0.2), xlabel('time (in samples)'),ylabel('percentage signal change'),
      grid on,grid minor, xlim([1 length(mF2)]),ylim([1.25.*min([mF2,mF4]) 1.25.*max([mF2,mF4])]), title(['Global mean signal AFTER 08-MOTION-CORRECTION (slice ' num2str(i3) ' of ' num2str(siz3) ')'],'Color',[0,0.55,0.69],'FontSize',14), legend({'Global signal after 08-motion-correction'},'FontSize',12,'TextColor',[0,0.55,0.69],'Location','best')
    saveas(fighndl,[QCpath 'global_signal_08moco_slice' sprintf('%.2d',i3) '.jpg'])
    close(fighndl)

    C  = corrcoef(F2'); C  = uppertriangle(C) ;
    C2 = corrcoef(F4'); C2 = uppertriangle(C2);
    fighndl = figure('Position',[1 1 1152 720],'Color',[0.95,0.9,0.85],'InvertHardcopy','off','Visible','off'); % red [0.95,0.9,0.85], green [0.85,0.90,0.85], blue [0.89,0.94,0.99]
    subplot2n(2,1,1), histogram(C2,'EdgeColor','none','FaceColor',[0.65,0.11,0.19],'FaceAlpha',0.5,'Normalization','probability'); grid on,grid minor, xlim([-1 1]), hold on, xline(0,'Color','k','linewidth',1.5); hold off  % xlim([-max(abs([C;C2])) max(abs([C;C2]))]),
      title(sprintf('Histogram of voxel-to-voxel correlations in UNPROCESSED RAW DATA (slice %d of %d)\n%.1f%% of the correlations were > 0  [ mean=%.2f, median=%.2f, SD=%.2f ]',i3,siz3,100*length(find(C2>0))/length(C2),mean(C2),median(C2),std(C2)),'Color',[0.65,0.11,0.19],'FontSize',14)
    subplot2n(2,1,2), histogram(C,'EdgeColor','none','FaceColor',[0,0.55,0.69],'FaceAlpha',0.5,'Normalization','probability'); grid on,grid minor, xlim([-1 1]), hold on, xline(0,'Color','k','linewidth',1.5); hold off
      title(sprintf('Histogram of voxel-to-voxel correlations AFTER 08-MOTION-CORRECTION (slice %d of %d)\n%.1f%% of the correlations were > 0  [ mean=%.2f, median=%.2f, SD=%.2f ]',i3,siz3,100*length(find(C>0))/length(C),mean(C),median(C),std(C)),'Color',[0,0.55,0.69],'FontSize',14)
    saveas(fighndl,[QCpath 'hist_FC_compare_08moco_slice' sprintf('%.2d',i3) '.jpg'])
    close(fighndl)
    
    if exist([base_dir_sub fname '_slice' num2str(1) '_denoised_before_MC' cutoff1_str '.nii'],'file') || exist([base_dir_sub fname '_slice' num2str(1) '_denoised_before_MC' cutoff1_str '.nii.gz'],'file') || exist([base_dir_sub fname '_denoised_before_MC' cutoff1_str '.nii'],'file') || exist([base_dir_sub fname '_denoised_before_MC' cutoff1_str '.nii.gz'],'file')
        scfMRItb_04_resplitData(base_dir_sub, fname, ['_denoised_before_MC' cutoff1_str], siz3) % if slice-wise data doesn't exist but combined 4D data exists, then split the data to generate the needed slice-wise data
        scfMRItb_04_unzipFile(base_dir_sub, fname, ['_slice' num2str(i3) '_denoised_before_MC' cutoff1_str])
        F5 = load_untouch_nii([base_dir_sub fname '_slice' num2str(i3) '_denoised_before_MC' cutoff1_str '.nii']); F5 = single(squeeze(F5.img));
        for i5=1:size(F5,3)
            tmp = F5(:,:,i5);
            F6(:,i5) = tmp(find(mask1==1)); clear tmp
        end; clear i5 F5
        mF6=mean(F6,1); mF6 = 100*(mF6-mean(mF6))./mean(mF6);
        fighndl = figure('Position',[1 1 1152 720],'Color',[0.95,0.9,0.85],'InvertHardcopy','off','Visible','off'); % red [0.95,0.9,0.85], green [0.85,0.90,0.85], blue [0.89,0.94,0.99]
        subplot2n(2,1,1), plot(mF6,'color',[0.65,0.11,0.19],'linewidth',1.5), hold on, plot(mF2,'color',[0,0.55,0.69],'linewidth',0.2), xlabel('time (in samples)'),ylabel('percentage signal change'),
          grid on,grid minor, xlim([1 length(mF6)]),ylim([min(1.25.*[mF2,mF6]) 1.25.*max([mF2,mF6])]), title(['Global mean signal after "06-denoise1" (slice ' num2str(i3) ' of ' num2str(siz3) ')'],'Color',[0.65,0.11,0.19],'FontSize',14), legend({'Global signal before motion correction'},'FontSize',12,'TextColor',[0.65,0.11,0.19],'Location','best')
        subplot2n(2,1,2), plot(mF2,'color',[0,0.55,0.69],'linewidth',1.5), hold on, plot(mF6,'color',[0.65,0.11,0.19],'linewidth',0.2), xlabel('time (in samples)'),ylabel('percentage signal change'),
          grid on,grid minor, xlim([1 length(mF2)]),ylim([min(1.25.*[mF2,mF6]) 1.25.*max([mF2,mF6])]), title(['Global mean signal after "08-motion-correction" (slice ' num2str(i3) ' of ' num2str(siz3) ')'],'Color',[0,0.55,0.69],'FontSize',14), legend({'Global signal after motion correction'},'FontSize',12,'TextColor',[0,0.55,0.69],'Location','best')
        saveas(fighndl,[QCpath 'global_signal_06denoise1_vs_08moco_slice' sprintf('%.2d',i3) '.jpg'])
        close(fighndl)
        
        C3 = corrcoef(F6'); C3=uppertriangle(C3);
        fighndl = figure('Position',[1 1 1152 720],'Color',[0.95,0.9,0.85],'InvertHardcopy','off','Visible','off'); % red [0.95,0.9,0.85], green [0.85,0.90,0.85], blue [0.89,0.94,0.99]
        subplot2n(2,1,1), histogram(C3,'EdgeColor','none','FaceColor',[0.65,0.11,0.19],'FaceAlpha',0.5,'Normalization','probability'); grid on,grid minor, xlim([-1 1]), hold on, xline(0,'Color','k','linewidth',1.5); hold off
          title(sprintf('Histogram of voxel-to-voxel correlations AFTER 06-DENOISE1 (slice %d of %d)\n%.1f%% of the correlations were > 0  [ mean=%.2f, median=%.2f, SD=%.2f ]',i3,siz3,100*length(find(C3>0))/length(C3),mean(C3),median(C3),std(C3)),'Color',[0.65,0.11,0.19],'FontSize',14)
        subplot2n(2,1,2), histogram(C,'EdgeColor','none','FaceColor',[0,0.55,0.69],'FaceAlpha',0.5,'Normalization','probability'); grid on,grid minor, xlim([-1 1]), hold on, xline(0,'Color','k','linewidth',1.5); hold off
          title(sprintf('Histogram of voxel-to-voxel correlations AFTER 08-MOTION-CORRECTION (slice %d of %d)\n%.1f%% of the correlations were > 0  [ mean=%.2f, median=%.2f, SD=%.2f ]',i3,siz3,100*length(find(C>0))/length(C),mean(C),median(C),std(C)),'Color',[0,0.55,0.69],'FontSize',14)
        saveas(fighndl,[QCpath 'hist_FC_compare_06denoise1_08moco_slice' sprintf('%.2d',i3) '.jpg'])
        close(fighndl)
        F6_=cat(1,F6_,F6); mF6_=cat(1,mF6_,mF6); C3_=cat(1,C3_,C3);
        clear F6 mF6 C3
    end
    
    F2_=cat(1,F2_,F2); mF2_=cat(1,mF2_,mF2); C_=cat(1,C_,C); F4_=cat(1,F4_,F4); mF4_=cat(1,mF4_,mF4); C2_=cat(1,C2_,C2);
    clear F2 F4 TR mask1 svar sdist mF2 mF4 C C2
end

[~,svar] = sort(var(F2_,0,2)); [~,spsc] = sort(meanPSC(F2_));
sortorder2 = sortorder; sortorder=[]; f=0; %#ok<*ASGLU>
for im=1:length(sortorder2)
    temp = sortorder2{im}; temp = temp + f; f = f + length(temp);
    sortorder = cat(1,sortorder,temp); clear temp
end; clear im f
sortorder = spsc; % options: spsc for percentage signal change, svar for variance, (comment out the line if you want to use distance from the center of the cord separately for each slice)
for im=max(sortorder):-1:1, if isempty(find(sortorder==im)), sortorder(find(sortorder>im)) = sortorder(find(sortorder>im))-1; end, end; clear im %#ok<*EFIND>

fighndl = figure('Position',[1 1 1152 720],'Color',[0.89,0.94,0.99],'InvertHardcopy','off','Visible','off'); % red [0.95,0.9,0.85], green [0.85,0.90,0.85], blue [0.89,0.94,0.99]
imagesc(F2_(sortorder,:)),colormap(cmap),colorbar, xlabel('time (in samples)'),ylabel('voxels (sorted by mean percentage signal change; bottom row is highest)'),
  title(['Time series within GM, WM and CSF AFTER 08-MOTION-CORRECTION (all slices combined)'],'Color',[0,0.55,0.69],'FontSize',14);
saveas(fighndl,[QCpath 'fMRI_ts_08moco_allSlices' '.jpg'])
 close(fighndl)
mF2_=mean(mF2_,1); mF4_=mean(mF4_,1);
fighndl = figure('Position',[1 1 1152 720],'Color',[0.89,0.94,0.99],'InvertHardcopy','off','Visible','off'); % red [0.95,0.9,0.85], green [0.85,0.90,0.85], blue [0.89,0.94,0.99]
subplot2n(2,1,1), plot(mF4_,'color',[0.65,0.11,0.19],'linewidth',1.5), hold on, plot(mF2_,'color',[0,0.55,0.69],'linewidth',0.2), xlabel('time (in samples)'),ylabel('percentage signal change'),
  grid on,grid minor, xlim([1 length(mF2_)]),ylim([1.25.*min([mF2_,mF4_]) 1.25.*max([mF2_,mF4_])]), title(['Global mean signal in UNPROCESSED RAW DATA (all slices averaged)'],'Color',[0.65,0.11,0.19],'FontSize',14), legend({'Global signal (raw data)'},'FontSize',12,'TextColor',[0.65,0.11,0.19],'Location','best')
subplot2n(2,1,2), plot(mF2_,'color',[0,0.55,0.69],'linewidth',1.5), hold on, plot(mF4_,'color',[0.65,0.11,0.19],'linewidth',0.2), xlabel('time (in samples)'),ylabel('percentage signal change'),
  grid on,grid minor, xlim([1 length(mF2_)]),ylim([1.25.*min([mF2_,mF4_]) 1.25.*max([mF2_,mF4_])]), title(['Global mean signal AFTER 08-MOTION-CORRECTION (all slices averaged)'],'Color',[0,0.55,0.69],'FontSize',14), legend({'Global signal after 08-motion-correction'},'FontSize',12,'TextColor',[0,0.55,0.69],'Location','best')
saveas(fighndl,[QCpath 'global_signal_08moco_allSlices' '.jpg'])
 close(fighndl)
fighndl = figure('Position',[1 1 1152 720],'Color',[0.89,0.94,0.99],'InvertHardcopy','off','Visible','off'); % red [0.95,0.9,0.85], green [0.85,0.90,0.85], blue [0.89,0.94,0.99]
subplot2n(2,1,1), histogram(C2_,'EdgeColor','none','FaceColor',[0.65,0.11,0.19],'FaceAlpha',0.5,'Normalization','probability'); grid on,grid minor, xlim([-1 1]), hold on, xline(0,'Color','k','linewidth',1.5); hold off
    title(sprintf('Histogram of voxel-to-voxel correlations in UNPROCESSED RAW DATA (all slices combined)\n%.1f%% of the correlations were > 0  [ mean=%.2f, median=%.2f, SD=%.2f ]',100*length(find(C2_>0))/length(C2_),mean(C2_),median(C2_),std(C2_)),'Color',[0.65,0.11,0.19],'FontSize',14)
subplot2n(2,1,2), histogram(C_,'EdgeColor','none','FaceColor',[0,0.55,0.69],'FaceAlpha',0.5,'Normalization','probability'); grid on,grid minor, xlim([-1 1]), hold on, xline(0,'Color','k','linewidth',1.5); hold off
    title(sprintf('Histogram of voxel-to-voxel correlations AFTER 08-MOTION-CORRECTION (all slices combined)\n%.1f%% of the correlations were > 0  [ mean=%.2f, median=%.2f, SD=%.2f ]',100*length(find(C_>0))/length(C_),mean(C_),median(C_),std(C_)),'Color',[0,0.55,0.69],'FontSize',14)
saveas(fighndl,[QCpath 'hist_FC_compare_08moco_allSlices' '.jpg'])
 close(fighndl)
if exist([base_dir_sub fname '_slice' num2str(1) '_denoised_before_MC' cutoff1_str '.nii'],'file') || exist([base_dir_sub fname '_slice' num2str(1) '_denoised_before_MC' cutoff1_str '.nii.gz'],'file') || exist([base_dir_sub fname '_denoised_before_MC' cutoff1_str '.nii'],'file') || exist([base_dir_sub fname '_denoised_before_MC' cutoff1_str '.nii.gz'],'file')
    mF6_=mean(mF6_,1);
    fighndl = figure('Position',[1 1 1152 720],'Color',[0.89,0.94,0.99],'InvertHardcopy','off','Visible','off'); % red [0.95,0.9,0.85], green [0.85,0.90,0.85], blue [0.89,0.94,0.99]
    subplot2n(2,1,1), plot(mF6_,'color',[0.65,0.11,0.19],'linewidth',1.5), hold on, plot(mF2_,'color',[0,0.55,0.69],'linewidth',0.2), xlabel('time (in samples)'),ylabel('percentage signal change'),
      grid on,grid minor, xlim([1 length(mF2_)]),ylim([1.25.*min([mF2_,mF6_]) 1.25.*max([mF2_,mF6_])]), title(['Global mean signal AFTER 06-DENOISE1 (all slices averaged)'],'Color',[0.65,0.11,0.19],'FontSize',14), legend({'Global signal before motion correction'},'FontSize',12,'TextColor',[0.65,0.11,0.19],'Location','best')
    subplot2n(2,1,2), plot(mF2_,'color',[0,0.55,0.69],'linewidth',1.5), hold on, plot(mF6_,'color',[0.65,0.11,0.19],'linewidth',0.2), xlabel('time (in samples)'),ylabel('percentage signal change'),
      grid on,grid minor, xlim([1 length(mF2_)]),ylim([1.25.*min([mF2_,mF6_]) 1.25.*max([mF2_,mF6_])]), title(['Global mean signal AFTER 08-MOTION-CORRECTION (all slices averaged)'],'Color',[0,0.55,0.69],'FontSize',14), legend({'Global signal after motion correction'},'FontSize',12,'TextColor',[0,0.55,0.69],'Location','best')
    saveas(fighndl,[QCpath 'global_signal_06denoise1_vs_08moco_allSlices' '.jpg'])
     close(fighndl)
    fighndl = figure('Position',[1 1 1152 720],'Color',[0.89,0.94,0.99],'InvertHardcopy','off','Visible','off'); % red [0.95,0.9,0.85], green [0.85,0.90,0.85], blue [0.89,0.94,0.99]
    subplot2n(2,1,1), histogram(C3_,'EdgeColor','none','FaceColor',[0.65,0.11,0.19],'FaceAlpha',0.5,'Normalization','probability'); grid on,grid minor, xlim([-1 1]), hold on, xline(0,'Color','k','linewidth',1.5); hold off
      title(sprintf('Histogram of voxel-to-voxel correlations AFTER 06-DENOISE1 (all slices combined)\n%.1f%% of the correlations were > 0  [ mean=%.2f, median=%.2f, SD=%.2f ]',100*length(find(C3_>0))/length(C3_),mean(C3_),median(C3_),std(C3_)),'Color',[0.65,0.11,0.19],'FontSize',14)
    subplot2n(2,1,2), histogram(C_,'EdgeColor','none','FaceColor',[0,0.55,0.69],'FaceAlpha',0.5,'Normalization','probability'); grid on,grid minor, xlim([-1 1]), hold on, xline(0,'Color','k','linewidth',1.5); hold off
      title(sprintf('Histogram of voxel-to-voxel correlations AFTER 08-MOTION-CORRECTION (all slices combined)\n%.1f%% of the correlations were > 0  [ mean=%.2f, median=%.2f, SD=%.2f ]',100*length(find(C_>0))/length(C_),mean(C_),median(C_),std(C_)),'Color',[0,0.55,0.69],'FontSize',14)
    saveas(fighndl,[QCpath 'hist_FC_compare_06denoise1_08moco_allSlices' '.jpg'])
     close(fighndl)
end
clear svar spsc sortorder F2_ mF2_ C_ F4_ mF4_ C2_ F6_ mF6_ C3_

%%% Make QC video (needs audio video toolbox in Matlab)
if ~isempty(which('VideoWriter')) && (save_videos==1) % check if Matlab's audio video toolbox exists

subplot = @(m,n,p) subtightplot(m, n, p, [0.001 0.005], [0.01 0.001], [0.01 0.001]);
for i3 = 1 : siz3
    try waitbar((i3/siz3),wbar3,sprintf('08. Generate QC video: slice (%d) of (%d)',i3,siz3)); catch, end
    fprintf('08. Generate QC video: slice (%d) of (%d), subject (%d) of (%d), run (%d) of (%d)\n',i3,siz3,k,size(func_files,1),k2,size(func_files,2))
    scfMRItb_04_resplitData(base_dir_sub, fname, '_MC', siz3) % if slice-wise data doesn't exist but combined 4D data exists, then split the data to generate the needed slice-wise data
    F1 = load_untouch_nii([base_dir_sub fname '_slice' num2str(i3) '_MC.nii']);
    TR = F1.hdr.dime.pixdim(5); F1 = rot90(single(squeeze(F1.img)));
    video = VideoWriter([QCpath 'fMRIvideo_08moco_slice' sprintf('%.2d',i3) '.mp4'],'MPEG-4'); %#ok<*TNMLP> % create the video object
    video.FrameRate = round(30/TR); % 30 seconds of data in every second of the video
    video.Quality = 90; % very limited compression so that the images are shown as is (and thus avoid misunderstandings due to video quality)
    open(video); % open the file for writing
    
    if Smask_err==0
        scfMRItb_04_resplitData(base_dir_sub, fname, '_Smask', siz3) % if slice-wise data doesn't exist but combined 4D data exists, then split the data to generate the needed slice-wise data
        scfMRItb_04_unzipFile(base_dir_sub, fname, ['_slice' num2str(i3) '_Smask'])
        mask1 = load_untouch_nii([base_dir_sub fname '_slice' num2str(i3) '_Smask.nii']); mask1 = mask1.img;
    else
        mask1 = Smask1_(:,:,i3);
    end
    mask1 = rot90(mask1);
    [xmask,ymask] = find(mask1==1);
    minx=round(min(xmask)-0.1*(size(mask1,1))); maxx=round(max(xmask)+0.1*(size(mask1,1)));
    miny=round(min(ymask)-0.1*(size(mask1,2))); maxy=round(max(ymask)+0.1*(size(mask1,2)));
    if (maxx-minx)<(maxy-miny)
        factr = ((maxy-miny)-(maxx-minx))/2;
        if factr==round(factr)
            maxx=maxx+factr; minx=minx-factr;
        else
            maxx=maxx+factr+0.5; minx=minx-factr+0.5;
        end
    elseif (maxy-miny)<(maxx-minx)
        factr = ((maxx-minx)-(maxy-miny))/2;
        if factr==round(factr)
            maxy=maxy+factr; miny=miny-factr;
        else
            maxy=maxy+factr+0.5; miny=miny-factr+0.5;
        end
    end
    [sizeFx,sizeFy] = size(F1(:,:,1));
    F2 = F1(minx:maxx,miny:maxy,:);
    for i5=1:size(F1,3)
        F1(:,:,i5) = imresize(F2(:,:,i5),[sizeFx sizeFy]);
    end
    scfMRItb_04_resplitData(base_dir_sub, fname, '', siz3) % if slice-wise data doesn't exist but combined 4D data exists, then split the data to generate the needed slice-wise data
    F3 = load_untouch_nii([base_dir_sub fname '_slice' num2str(i3) '.nii']); F3 = rot90(single(squeeze(F3.img)));
    F4 = F3(minx:maxx,miny:maxy,:);
    for i5=1:size(F3,3)
        F3(:,:,i5) = imresize(F4(:,:,i5),[sizeFx sizeFy]);
    end
    clear maxx maxy minx miny xmask ymask mask1 factr sizeFx sizeFy F2 F4 i5
    
    for ii=1:size(F1,3) % loop across the number of images
        I = F1(:,:,ii); %read the next image
        I(find(isnan(I)))=0;
        I = (I - min(I(:))) ./ (max(I(:)) - min(I(:)));
        I2 = F3(:,:,ii); %read the next image
        I2(find(isnan(I2)))=0;
        I2 = (I2 - min(I2(:))) ./ (max(I2(:)) - min(I2(:)));
        fighndl = figure('Position',[round((scrsz(3)-scrsz(4))/2) 1 round(0.75*scrsz(3)) round(0.75*scrsz(4))],'Color',[0.95,0.9,0.85],'InvertHardcopy','off','Visible','off'); % [0.85,0.90,0.85]
        subplot(1,2,1), imshow(I2, [0 max(I2(:))], 'InitialMagnification', 'fit'); title({['Unprocessed raw fMRI data (slice ' num2str(i3) ' of ' num2str(siz3) ')'];['Time point ' num2str(ii) ' of ' num2str(size(F1,3))]},'Color',[0.65,0.11,0.19],'FontSize',18);
        subplot(1,2,2), imshow(I, [0 max(I(:))], 'InitialMagnification', 'fit'); title({['fMRI data after "08. motion correction" (slice ' num2str(i3) ' of ' num2str(siz3) ')'];['Time point ' num2str(ii) ' of ' num2str(size(F1,3))]},'Color',[0,0.55,0.69],'FontSize',18);
        frame = getframe(gcf);
        writeVideo(video,frame); % write the image to file
        close(fighndl)
        clear I I2 frame
    end
    close(video); %close the file
    clear F1 F3 video ii TR
end; clear i3

end
end

%% scfMRItb_09_retroicor
if choices_preproc(9)==1
clear QCpath save_videos fighndl
if ~(exist([base_dir_sub 'QC' '/09_denoise2-RETROICOR'],'dir'))
    mkdir([base_dir_sub 'QC' '/09_denoise2-RETROICOR'])
end
if Rns==0
    QCpath = [base_dir_sub 'QC' '/09_denoise2-RETROICOR/'];
else
    if ~(exist([base_dir_sub 'QC' '/09_denoise2-RETROICOR' '/run' num2str(Rns)],'dir'))
        mkdir([base_dir_sub 'QC' '/09_denoise2-RETROICOR' '/run' num2str(Rns)])
    end
    QCpath = [base_dir_sub 'QC' '/09_denoise2-RETROICOR' '/run' num2str(Rns) '/'];
end

if (save_videos_==0)||(save_videos_==1)
    save_videos = save_videos_;
elseif (save_videos_==2)&&(sum([choices_preproc(11:14)])==0)
    save_videos = 1;
else
    save_videos = 0;
end
if save_videos==1
    save_videos_steps(4)=1;
end


F2_=[]; mF2_=[]; C_=[]; F4_=[]; mF4_=[]; C2_=[];
if exist([base_dir_sub fname '_slice' num2str(siz3) '_MC.nii'],'file') || exist([base_dir_sub fname '_slice' num2str(i3) '_MC.nii.gz'],'file') || exist([base_dir_sub fname '_MC.nii'],'file') || exist([base_dir_sub fname '_MC.nii.gz'],'file')
    F6_=[]; mF6_=[]; C3_=[];
end
for i3 = 1 : siz3
    try waitbar((i3/siz3),wbar3,sprintf('09. QC plots: slice (%d) of (%d)',i3,siz3)); catch, end
    fprintf('09. QC plots: slice (%d) of (%d), subject (%d) of (%d), run (%d) of (%d)\n',i3,siz3,k,size(func_files,1),k2,size(func_files,2))
    scfMRItb_04_resplitData(base_dir_sub, fname, '_MC_ricor', siz3) % if slice-wise data doesn't exist but combined 4D data exists, then split the data to generate the needed slice-wise data
    scfMRItb_04_unzipFile(base_dir_sub, fname, ['_slice' num2str(i3) '_MC_ricor'])
    F1 = load_untouch_nii([base_dir_sub fname '_slice' num2str(i3) '_MC_ricor.nii']);
    TR = F1.hdr.dime.pixdim(5); F1 = single(squeeze(F1.img));
    
    if Smask_err==0
        scfMRItb_04_resplitData(base_dir_sub, fname, '_Smask', siz3) % if slice-wise data doesn't exist but combined 4D data exists, then split the data to generate the needed slice-wise data
        scfMRItb_04_unzipFile(base_dir_sub, fname, ['_slice' num2str(i3) '_Smask'])
        mask1 = load_untouch_nii([base_dir_sub fname '_slice' num2str(i3) '_Smask.nii']); mask1 = mask1.img;
    else
        mask1 = Smask1_(:,:,i3);
    end
    [m1,m2] = find(mask1==1); Mm1=median(m1); Mm2=median(m2);
    sdist = sqrt((m1-Mm1).^2 + (m2-Mm2).^2);
    clear im Mm1 Mm2 m1 m2
    for i5=1:size(F1,3)
        tmp = F1(:,:,i5);
        F2(:,i5) = tmp(find(mask1==1)); clear tmp
    end; clear i5 F1
    [~,svar] = sort(var(F2,0,2)); [~,sdist] = sort(sdist); sortorder{i3,1} = sdist; %#ok<*AGROW> % sort timeseries based on distance from the center of the cord (if you want to sort by signal variance then use svar)
    
    fighndl = figure('Position',[1 1 1152 720],'Color',[0.95,0.9,0.85],'InvertHardcopy','off','Visible','off'); % red [0.95,0.9,0.85], green [0.85,0.90,0.85], blue [0.89,0.94,0.99]
    imagesc(F2(sortorder{i3},:)),colormap(cmap),colorbar, xlabel('time (in samples)'),ylabel('voxels (sorted by distance from the center of the cord; top row is nearest)'),
        title(['Time series within GM, WM and CSF after "09-RETROICOR" in slice ' num2str(i3) ' of ' num2str(siz3)],'Color',[0,0.55,0.69],'FontSize',14);
    saveas(fighndl,[QCpath 'fMRI_ts_09ricor_slice' sprintf('%.2d',i3) '.jpg'])
    close(fighndl)
    
    scfMRItb_04_resplitData(base_dir_sub, fname, '', siz3) % if slice-wise data doesn't exist but combined 4D data exists, then split the data to generate the needed slice-wise data
    scfMRItb_04_unzipFile(base_dir_sub, fname, ['_slice' num2str(i3)])
    F3 = load_untouch_nii([base_dir_sub fname '_slice' num2str(i3) '.nii']); F3 = single(squeeze(F3.img));
    for i5=1:size(F3,3)
        tmp = F3(:,:,i5);
        F4(:,i5) = tmp(find(mask1==1)); clear tmp
    end; clear i5 F3
    mF2=mean(F2,1); mF2 = 100*(mF2-mean(mF2))./mean(mF2); mF4=mean(F4,1); mF4 = 100*(mF4-mean(mF4))./mean(mF4);
    fighndl = figure('Position',[1 1 1152 720],'Color',[0.95,0.9,0.85],'InvertHardcopy','off','Visible','off'); % red [0.95,0.9,0.85], green [0.85,0.90,0.85], blue [0.89,0.94,0.99]
    subplot2n(2,1,1), plot(mF4,'color',[0.65,0.11,0.19],'linewidth',1.5), hold on, plot(mF2,'color',[0,0.55,0.69],'linewidth',0.2), xlabel('time (in samples)'),ylabel('percentage signal change'),
      grid on,grid minor, xlim([1 length(mF2)]),ylim([1.25.*min([mF2,mF4]) 1.25.*max([mF2,mF4])]), title(['Global mean signal in UNPROCESSED RAW DATA (slice ' num2str(i3) ' of ' num2str(siz3) ')'],'Color',[0.65,0.11,0.19],'FontSize',14), legend({'Global signal (raw data)'},'FontSize',12,'TextColor',[0.65,0.11,0.19],'Location','best')
    subplot2n(2,1,2), plot(mF2,'color',[0,0.55,0.69],'linewidth',1.5), hold on, plot(mF4,'color',[0.65,0.11,0.19],'linewidth',0.2), xlabel('time (in samples)'),ylabel('percentage signal change'),
      grid on,grid minor, xlim([1 length(mF2)]),ylim([1.25.*min([mF2,mF4]) 1.25.*max([mF2,mF4])]), title(['Global mean signal AFTER "09-RETROICOR" (slice ' num2str(i3) ' of ' num2str(siz3) ')'],'Color',[0,0.55,0.69],'FontSize',14), legend({'Global signal after 09-RETROICOR'},'FontSize',12,'TextColor',[0,0.55,0.69],'Location','best')
    saveas(fighndl,[QCpath 'global_signal_09ricor_slice' sprintf('%.2d',i3) '.jpg'])
    close(fighndl)

    C  = corrcoef(F2'); C  = uppertriangle(C) ;
    C2 = corrcoef(F4'); C2 = uppertriangle(C2);
    fighndl = figure('Position',[1 1 1152 720],'Color',[0.95,0.9,0.85],'InvertHardcopy','off','Visible','off'); % red [0.95,0.9,0.85], green [0.85,0.90,0.85], blue [0.89,0.94,0.99]
    subplot2n(2,1,1), histogram(C2,'EdgeColor','none','FaceColor',[0.65,0.11,0.19],'FaceAlpha',0.5,'Normalization','probability'); grid on,grid minor, xlim([-1 1]), hold on, xline(0,'Color','k','linewidth',1.5); hold off
      title(sprintf('Histogram of voxel-to-voxel correlations in UNPROCESSED RAW DATA (slice %d of %d)\n%.1f%% of the correlations were > 0  [ mean=%.2f, median=%.2f, SD=%.2f ]',i3,siz3,100*length(find(C2>0))/length(C2),mean(C2),median(C2),std(C2)),'Color',[0.65,0.11,0.19],'FontSize',14)
    subplot2n(2,1,2), histogram(C,'EdgeColor','none','FaceColor',[0,0.55,0.69],'FaceAlpha',0.5,'Normalization','probability'); grid on,grid minor, xlim([-1 1]), hold on, xline(0,'Color','k','linewidth',1.5); hold off
      title(sprintf('Histogram of voxel-to-voxel correlations AFTER 09-RETROICOR (slice %d of %d)\n%.1f%% of the correlations were > 0  [ mean=%.2f, median=%.2f, SD=%.2f ]',i3,siz3,100*length(find(C>0))/length(C),mean(C),median(C),std(C)),'Color',[0,0.55,0.69],'FontSize',14)
    saveas(fighndl,[QCpath 'hist_FC_compare_09ricor_slice' sprintf('%.2d',i3) '.jpg'])
    close(fighndl)
    
    if exist([base_dir_sub fname '_slice' num2str(i3) '_MC.nii'],'file') || exist([base_dir_sub fname '_slice' num2str(i3) '_MC.nii.gz'],'file') || exist([base_dir_sub fname '_MC.nii'],'file') || exist([base_dir_sub fname '_MC.nii.gz'],'file')
        scfMRItb_04_resplitData(base_dir_sub, fname, '_MC', siz3) % if slice-wise data doesn't exist but combined 4D data exists, then split the data to generate the needed slice-wise data
        scfMRItb_04_unzipFile(base_dir_sub, fname, ['_slice' num2str(i3) '_MC'])
        F5 = load_untouch_nii([base_dir_sub fname '_slice' num2str(i3) '_MC.nii']); F5 = single(squeeze(F5.img));
        for i5=1:size(F5,3)
            tmp = F5(:,:,i5);
            F6(:,i5) = tmp(find(mask1==1)); clear tmp
        end; clear i5 F5
        mF6=mean(F6,1); mF6 = 100*(mF6-mean(mF6))./mean(mF6);
        fighndl = figure('Position',[1 1 1152 720],'Color',[0.95,0.9,0.85],'InvertHardcopy','off','Visible','off'); % red [0.95,0.9,0.85], green [0.85,0.90,0.85], blue [0.89,0.94,0.99]
        subplot2n(2,1,1), plot(mF6,'color',[0.65,0.11,0.19],'linewidth',1.5), hold on, plot(mF2,'color',[0,0.55,0.69],'linewidth',0.2), xlabel('time (in samples)'),ylabel('percentage signal change'),
          grid on,grid minor, xlim([1 length(mF2)]),ylim([1.25.*min([mF2,mF6]) 1.25.*max([mF2,mF6])]), title(['Global mean signal AFTER "08-motion-correction" (slice ' num2str(i3) ' of ' num2str(siz3) ')'],'Color',[0.65,0.11,0.19],'FontSize',14), legend({'Global signal before RETROICOR'},'FontSize',12,'TextColor',[0.65,0.11,0.19],'Location','best')
        subplot2n(2,1,2), plot(mF2,'color',[0,0.55,0.69],'linewidth',1.5), hold on, plot(mF6,'color',[0.65,0.11,0.19],'linewidth',0.2), xlabel('time (in samples)'),ylabel('percentage signal change'),
          grid on,grid minor, xlim([1 length(mF2)]),ylim([1.25.*min([mF2,mF6]) 1.25.*max([mF2,mF6])]), title(['Global mean signal AFTER "09-RETROICOR" (slice ' num2str(i3) ' of ' num2str(siz3) ')'],'Color',[0,0.55,0.69],'FontSize',14), legend({'Global signal after RETROICOR'},'FontSize',12,'TextColor',[0,0.55,0.69],'Location','best')
        saveas(fighndl,[QCpath 'global_signal_08moco_vs_09ricor_slice' sprintf('%.2d',i3) '.jpg'])
        close(fighndl)
        
        C3 = corrcoef(F6'); C3=uppertriangle(C3);
        fighndl = figure('Position',[1 1 1152 720],'Color',[0.95,0.9,0.85],'InvertHardcopy','off','Visible','off'); % red [0.95,0.9,0.85], green [0.85,0.90,0.85], blue [0.89,0.94,0.99]
        subplot2n(2,1,1), histogram(C3,'EdgeColor','none','FaceColor',[0.65,0.11,0.19],'FaceAlpha',0.5,'Normalization','probability'); grid on,grid minor, xlim([-1 1]), hold on, xline(0,'Color','k','linewidth',1.5); hold off
          title(sprintf('Histogram of voxel-to-voxel correlations AFTER 08-motion-correction (slice %d of %d)\n%.1f%% of the correlations were > 0  [ mean=%.2f, median=%.2f, SD=%.2f ]',i3,siz3,100*length(find(C3>0))/length(C3),mean(C3),median(C3),std(C3)),'Color',[0.65,0.11,0.19],'FontSize',14)
        subplot2n(2,1,2), histogram(C,'EdgeColor','none','FaceColor',[0,0.55,0.69],'FaceAlpha',0.5,'Normalization','probability'); grid on,grid minor, xlim([-1 1]), hold on, xline(0,'Color','k','linewidth',1.5); hold off
          title(sprintf('Histogram of voxel-to-voxel correlations AFTER 09-RETROICOR (slice %d of %d)\n%.1f%% of the correlations were > 0  [ mean=%.2f, median=%.2f, SD=%.2f ]',i3,siz3,100*length(find(C>0))/length(C),mean(C),median(C),std(C)),'Color',[0,0.55,0.69],'FontSize',14)
        saveas(fighndl,[QCpath 'hist_FC_compare_08moco_09ricor_slice' sprintf('%.2d',i3) '.jpg'])
        close(fighndl)
        F6_=cat(1,F6_,F6); mF6_=cat(1,mF6_,mF6); C3_=cat(1,C3_,C3);
        clear F6 mF6 C3
    end
    
    F2_=cat(1,F2_,F2); mF2_=cat(1,mF2_,mF2); C_=cat(1,C_,C); F4_=cat(1,F4_,F4); mF4_=cat(1,mF4_,mF4); C2_=cat(1,C2_,C2);
    clear F2 F4 TR mask1 svar sdist mF2 mF4 C C2
end


[~,svar] = sort(var(F2_,0,2)); [~,spsc] = sort(meanPSC(F2_));
sortorder2 = sortorder; sortorder=[]; f=0; %#ok<*ASGLU>
for im=1:length(sortorder2)
    temp = sortorder2{im}; temp = temp + f; f = f + length(temp);
    sortorder = cat(1,sortorder,temp); clear temp
end; clear im f
sortorder = spsc; % options: spsc for percentage signal change, svar for variance, (comment out the line if you want to use distance from the center of the cord separately for each slice)
for im=max(sortorder):-1:1, if isempty(find(sortorder==im)), sortorder(find(sortorder>im)) = sortorder(find(sortorder>im))-1; end, end; clear im %#ok<*EFIND>

fighndl = figure('Position',[1 1 1152 720],'Color',[0.89,0.94,0.99],'InvertHardcopy','off','Visible','off'); % red [0.95,0.9,0.85], green [0.85,0.90,0.85], blue [0.89,0.94,0.99]
imagesc(F2_(sortorder,:)),colormap(cmap),colorbar, xlabel('time (in samples)'),ylabel('voxels (sorted by mean percentage signal change; bottom row is highest)'),
  title(['Time series within GM, WM and CSF AFTER 09-RETROICOR (all slices combined)'],'Color',[0,0.55,0.69],'FontSize',14);
saveas(fighndl,[QCpath 'fMRI_ts_09ricor_allSlices' '.jpg'])
 close(fighndl)
mF2_=mean(mF2_,1); mF4_=mean(mF4_,1);
fighndl = figure('Position',[1 1 1152 720],'Color',[0.89,0.94,0.99],'InvertHardcopy','off','Visible','off'); % red [0.95,0.9,0.85], green [0.85,0.90,0.85], blue [0.89,0.94,0.99]
subplot2n(2,1,1), plot(mF4_,'color',[0.65,0.11,0.19],'linewidth',1.5), hold on, plot(mF2_,'color',[0,0.55,0.69],'linewidth',0.2), xlabel('time (in samples)'),ylabel('percentage signal change'),
  grid on,grid minor, xlim([1 length(mF2_)]),ylim([1.25.*min([mF2_,mF4_]) 1.25.*max([mF2_,mF4_])]), title(['Global mean signal in UNPROCESSED RAW DATA (all slices averaged)'],'Color',[0.65,0.11,0.19],'FontSize',14), legend({'Global signal (raw data)'},'FontSize',12,'TextColor',[0.65,0.11,0.19],'Location','best')
subplot2n(2,1,2), plot(mF2_,'color',[0,0.55,0.69],'linewidth',1.5), hold on, plot(mF4_,'color',[0.65,0.11,0.19],'linewidth',0.2), xlabel('time (in samples)'),ylabel('percentage signal change'),
  grid on,grid minor, xlim([1 length(mF2_)]),ylim([1.25.*min([mF2_,mF4_]) 1.25.*max([mF2_,mF4_])]), title(['Global mean signal AFTER 09-RETROICOR (all slices averaged)'],'Color',[0,0.55,0.69],'FontSize',14), legend({'Global signal after 09-RETROICOR'},'FontSize',12,'TextColor',[0,0.55,0.69],'Location','best')
saveas(fighndl,[QCpath 'global_signal_09ricor_allSlices' '.jpg'])
 close(fighndl)
fighndl = figure('Position',[1 1 1152 720],'Color',[0.89,0.94,0.99],'InvertHardcopy','off','Visible','off'); % red [0.95,0.9,0.85], green [0.85,0.90,0.85], blue [0.89,0.94,0.99]
subplot2n(2,1,1), histogram(C2_,'EdgeColor','none','FaceColor',[0.65,0.11,0.19],'FaceAlpha',0.5,'Normalization','probability'); grid on,grid minor, xlim([-1 1]), hold on, xline(0,'Color','k','linewidth',1.5); hold off
    title(sprintf('Histogram of voxel-to-voxel correlations in UNPROCESSED RAW DATA (all slices combined)\n%.1f%% of the correlations were > 0  [ mean=%.2f, median=%.2f, SD=%.2f ]',100*length(find(C2_>0))/length(C2_),mean(C2_),median(C2_),std(C2_)),'Color',[0.65,0.11,0.19],'FontSize',14)
subplot2n(2,1,2), histogram(C_,'EdgeColor','none','FaceColor',[0,0.55,0.69],'FaceAlpha',0.5,'Normalization','probability'); grid on,grid minor, xlim([-1 1]), hold on, xline(0,'Color','k','linewidth',1.5); hold off
    title(sprintf('Histogram of voxel-to-voxel correlations AFTER 09-RETROICOR (all slices combined)\n%.1f%% of the correlations were > 0  [ mean=%.2f, median=%.2f, SD=%.2f ]',100*length(find(C_>0))/length(C_),mean(C_),median(C_),std(C_)),'Color',[0,0.55,0.69],'FontSize',14)
saveas(fighndl,[QCpath 'hist_FC_compare_09ricor_allSlices' '.jpg'])
 close(fighndl)
if exist([base_dir_sub fname '_slice' num2str(i3) '_MC.nii'],'file') || exist([base_dir_sub fname '_slice' num2str(i3) '_MC.nii.gz'],'file') || exist([base_dir_sub fname '_MC.nii'],'file') || exist([base_dir_sub fname '_MC.nii.gz'],'file')
    mF6_=mean(mF6_,1);
    fighndl = figure('Position',[1 1 1152 720],'Color',[0.89,0.94,0.99],'InvertHardcopy','off','Visible','off'); % red [0.95,0.9,0.85], green [0.85,0.90,0.85], blue [0.89,0.94,0.99]
    subplot2n(2,1,1), plot(mF6_,'color',[0.65,0.11,0.19],'linewidth',1.5), hold on, plot(mF2_,'color',[0,0.55,0.69],'linewidth',0.2), xlabel('time (in samples)'),ylabel('percentage signal change'),
      grid on,grid minor, xlim([1 length(mF2_)]),ylim([1.25.*min([mF2_,mF6_]) 1.25.*max([mF2_,mF6_])]), title(['Global mean signal AFTER 08-MOTION-CORRECTION (all slices averaged)'],'Color',[0.65,0.11,0.19],'FontSize',14), legend({'Global signal before RETROICOR'},'FontSize',12,'TextColor',[0.65,0.11,0.19],'Location','best')
    subplot2n(2,1,2), plot(mF2_,'color',[0,0.55,0.69],'linewidth',1.5), hold on, plot(mF6_,'color',[0.65,0.11,0.19],'linewidth',0.2), xlabel('time (in samples)'),ylabel('percentage signal change'),
      grid on,grid minor, xlim([1 length(mF2_)]),ylim([1.25.*min([mF2_,mF6_]) 1.25.*max([mF2_,mF6_])]), title(['Global mean signal AFTER 09-RETROICOR (all slices averaged)'],'Color',[0,0.55,0.69],'FontSize',14), legend({'Global signal after RETROICOR'},'FontSize',12,'TextColor',[0,0.55,0.69],'Location','best')
    saveas(fighndl,[QCpath 'global_signal_08moco_vs_09ricor_allSlices' '.jpg'])
     close(fighndl)
     fighndl = figure('Position',[1 1 1152 720],'Color',[0.89,0.94,0.99],'InvertHardcopy','off','Visible','off'); % red [0.95,0.9,0.85], green [0.85,0.90,0.85], blue [0.89,0.94,0.99]
    subplot2n(2,1,1), histogram(C3_,'EdgeColor','none','FaceColor',[0.65,0.11,0.19],'FaceAlpha',0.5,'Normalization','probability'); grid on,grid minor, xlim([-1 1]), hold on, xline(0,'Color','k','linewidth',1.5); hold off
      title(sprintf('Histogram of voxel-to-voxel correlations AFTER 08-MOTION-CORRECTION (all slices combined)\n%.1f%% of the correlations were > 0  [ mean=%.2f, median=%.2f, SD=%.2f ]',100*length(find(C3_>0))/length(C3_),mean(C3_),median(C3_),std(C3_)),'Color',[0.65,0.11,0.19],'FontSize',14)
    subplot2n(2,1,2), histogram(C_,'EdgeColor','none','FaceColor',[0,0.55,0.69],'FaceAlpha',0.5,'Normalization','probability'); grid on,grid minor, xlim([-1 1]), hold on, xline(0,'Color','k','linewidth',1.5); hold off
      title(sprintf('Histogram of voxel-to-voxel correlations AFTER 09-RETROICOR (all slices combined)\n%.1f%% of the correlations were > 0  [ mean=%.2f, median=%.2f, SD=%.2f ]',100*length(find(C_>0))/length(C_),mean(C_),median(C_),std(C_)),'Color',[0,0.55,0.69],'FontSize',14)
    saveas(fighndl,[QCpath 'hist_FC_compare_08moco_09ricor_allSlices' '.jpg'])
     close(fighndl)
end
clear svar spsc sortorder F2_ mF2_ C_ F4_ mF4_ C2_ F6_ mF6_ C3_


%%% Make QC video (needs audio video toolbox in Matlab)
if ~isempty(which('VideoWriter')) && (save_videos==1) % check if Matlab's audio video toolbox exists

subplot = @(m,n,p) subtightplot(m, n, p, [0.001 0.005], [0.01 0.001], [0.01 0.001]);
for i3 = 1 : siz3
    try waitbar((i3/siz3),wbar3,sprintf('09. Generate QC video: slice (%d) of (%d)',i3,siz3)); catch, end
    fprintf('09. Generate QC video: slice (%d) of (%d), subject (%d) of (%d), run (%d) of (%d)\n',i3,siz3,k,size(func_files,1),k2,size(func_files,2))
    scfMRItb_04_resplitData(base_dir_sub, fname, '_MC_ricor', siz3) % if slice-wise data doesn't exist but combined 4D data exists, then split the data to generate the needed slice-wise data
    F1 = load_untouch_nii([base_dir_sub fname '_slice' num2str(i3) '_MC_ricor.nii']);
    TR = F1.hdr.dime.pixdim(5); F1 = rot90(single(squeeze(F1.img)));
    video = VideoWriter([QCpath 'fMRIvideo_09ricor_slice' sprintf('%.2d',i3) '.mp4'],'MPEG-4'); %#ok<*TNMLP> % create the video object
    video.FrameRate = round(30/TR); % 30 seconds of data in every second of the video
    video.Quality = 90; % very limited compression so that the images are shown as is (and thus avoid misunderstandings due to video quality)
    open(video); % open the file for writing
    
    if Smask_err==0
        scfMRItb_04_resplitData(base_dir_sub, fname, '_Smask', siz3) % if slice-wise data doesn't exist but combined 4D data exists, then split the data to generate the needed slice-wise data
        scfMRItb_04_unzipFile(base_dir_sub, fname, ['_slice' num2str(i3) '_Smask'])
        mask1 = load_untouch_nii([base_dir_sub fname '_slice' num2str(i3) '_Smask.nii']); mask1 = mask1.img;
    else
        mask1 = Smask1_(:,:,i3);
    end
    mask1 = rot90(mask1);
    [xmask,ymask] = find(mask1==1);
    minx=round(min(xmask)-0.1*(size(mask1,1))); maxx=round(max(xmask)+0.1*(size(mask1,1)));
    miny=round(min(ymask)-0.1*(size(mask1,2))); maxy=round(max(ymask)+0.1*(size(mask1,2)));
    if (maxx-minx)<(maxy-miny)
        factr = ((maxy-miny)-(maxx-minx))/2;
        if factr==round(factr)
            maxx=maxx+factr; minx=minx-factr;
        else
            maxx=maxx+factr+0.5; minx=minx-factr+0.5;
        end
    elseif (maxy-miny)<(maxx-minx)
        factr = ((maxx-minx)-(maxy-miny))/2;
        if factr==round(factr)
            maxy=maxy+factr; miny=miny-factr;
        else
            maxy=maxy+factr+0.5; miny=miny-factr+0.5;
        end
    end
    [sizeFx,sizeFy] = size(F1(:,:,1));
    F2 = F1(minx:maxx,miny:maxy,:);
    for i5=1:size(F1,3)
        F1(:,:,i5) = imresize(F2(:,:,i5),[sizeFx sizeFy]);
    end
    scfMRItb_04_resplitData(base_dir_sub, fname, '', siz3) % if slice-wise data doesn't exist but combined 4D data exists, then split the data to generate the needed slice-wise data
    F3 = load_untouch_nii([base_dir_sub fname '_slice' num2str(i3) '.nii']); F3 = rot90(single(squeeze(F3.img)));
    F4 = F3(minx:maxx,miny:maxy,:);
    for i5=1:size(F3,3)
        F3(:,:,i5) = imresize(F4(:,:,i5),[sizeFx sizeFy]);
    end
    clear maxx maxy minx miny xmask ymask mask1 factr sizeFx sizeFy F2 F4 i5
    
    for ii=1:size(F1,3) % loop across the number of images
        I = F1(:,:,ii); %read the next image
        I(find(isnan(I)))=0;
        I = (I - min(I(:))) ./ (max(I(:)) - min(I(:)));
        I2 = F3(:,:,ii); %read the next image
        I2(find(isnan(I2)))=0;
        I2 = (I2 - min(I2(:))) ./ (max(I2(:)) - min(I2(:)));
        fighndl = figure('Position',[round((scrsz(3)-scrsz(4))/2) 1 round(0.75*scrsz(3)) round(0.75*scrsz(4))],'Color',[0.95,0.9,0.85],'InvertHardcopy','off','Visible','off'); % [0.85,0.90,0.85]
        subplot(1,2,1), imshow(I2, [0 max(I2(:))], 'InitialMagnification', 'fit'); title({['Unprocessed raw fMRI data (slice ' num2str(i3) ' of ' num2str(siz3) ')'];['Time point ' num2str(ii) ' of ' num2str(size(F1,3))]},'Color',[0.65,0.11,0.19],'FontSize',18);
        subplot(1,2,2), imshow(I, [0 max(I(:))], 'InitialMagnification', 'fit'); title({['fMRI data after "09. RETROICOR" (slice ' num2str(i3) ' of ' num2str(siz3) ')'];['Time point ' num2str(ii) ' of ' num2str(size(F1,3))]},'Color',[0,0.55,0.69],'FontSize',18);
        frame = getframe(gcf);
        writeVideo(video,frame); % write the image to file
        close(fighndl)
        clear I I2 frame
    end
    close(video); %close the file
    clear F1 F3 video ii TR
end; clear i3

end
end

%% scfMRItb_10_GMWMCSFmasks
if (choices_preproc(10)==1)&&(k2==1)
clear QCpath save_videos fighndl
if ~(exist([base_dir_sub 'QC' '/10_GM-WM-CSF-masks'],'dir'))
    mkdir([base_dir_sub 'QC' '/10_GM-WM-CSF-masks'])
end
QCpath = [base_dir_sub 'QC' '/10_GM-WM-CSF-masks' '/'];

try
    load([base_dir_sub fname_anat '_all_masks.mat'],'position_GM','position_WM','position_CSF')
    scfMRItb_04_unzipFile(base_dir_sub, fname_anat, '')
    A = load_untouch_nii([base_dir_sub fname_anat '.nii']);
    new_size = round(size(A.img,1) / 2);
    for i3 = 1 : siz3
        try
            scfMRItb_04_resplitData(base_dir_sub, fname, '_Smask', siz3) % if slice-wise data doesn't exist but combined 4D data exists, then split the data to generate the needed slice-wise data
            scfMRItb_04_unzipFile(base_dir_sub, fname, ['_slice' num2str(i3) '_Smask'])
            Smask = load_untouch_nii([base_dir_sub fname '_slice' num2str(i3) '_Smask.nii']);
        catch
            Smask.img = zeros(size(A.img,1),size(A.img,2)); Smask.img(round(size(A.img,1)/4):round(3*size(A.img,1)/4),round(size(A.img,2)/4):round(3*size(A.img,2)/4))=1;
        end
        Smask = rot90(Smask.img);
        [w2,w1] = find(Smask~=0);
        w1min(i3,1) = min(w1)/size(Smask,1); w1max(i3,1) = max(w1)/size(Smask,1);
        w2min(i3,1) = min(w2)/size(Smask,2); w2max(i3,1) = max(w2)/size(Smask,2);
        clear Smask w1 w2
        
        img = rot90(A.img(:,:,i3));
        temp = img(round((size(img,1)-new_size)/2)+1:end-round((size(img,1)-new_size)/2), ...
            round((size(img,2)-new_size)/2)+1:end-round((size(img,2)-new_size)/2));
        mmax = max(temp(:)); clear temp;
        
        % mask_CSF
        fighndl = figure('Position',[round((scrsz(3)-scrsz(4))/2) 1 scrsz(4) scrsz(4)],'InvertHardcopy','off','Visible','off'); fighndl.Color = [0.8, 0.87, 0.84];
        imshow(single(img), [0 mmax], 'InitialMagnification', 'fit'); colorbar;
        fighndl.Color = [0.8, 0.87, 0.84] + [0, 0.02*(1-exp(siz3/i3)/exp(siz3)), 0.1*exp(i3/siz3)/exp(1)]; % [0.85, 0.90, 0.85] + [0, 0, 0.1*i3/siz3];
        title({'Draw the boundary of CSF (double click the cord once done)';['Slice ' num2str(i3) ' of ' num2str(siz3)]},'Color',[0.7,0,0.7],'FontSize',16);
        hold on, plot([position_CSF{i3}(:,1);position_CSF{i3}(1,1)],[position_CSF{i3}(:,2);position_CSF{i3}(1,2)],'m','LineWidth',2), hold off
        legend('CSF mask')
        lims_x1(i3) = min(position_CSF{i3}(:,1)); lims_x2(i3) = max(position_CSF{i3}(:,1)); %#ok<*AGROW>
        lims_x1_(i3) = round(lims_x1(i3) - 0.15*(lims_x2(i3)-lims_x1(i3))); lims_x2_(i3) = round(lims_x2(i3) + 0.15*(lims_x2(i3)-lims_x1(i3)));
        lims_y1(i3) = min(position_CSF{i3}(:,2)); lims_y2(i3) = max(position_CSF{i3}(:,2));
        lims_y1_(i3) = round(lims_y1(i3) - 0.15*(lims_y2(i3)-lims_y1(i3))); lims_y2_(i3) = round(lims_y2(i3) + 0.15*(lims_y2(i3)-lims_y1(i3)));
        xlim([min([round(size(img,1)*w1min(i3)),lims_x1_(i3)]),max([round(size(img,1)*w1max(i3)),lims_x2_(i3)])]),
        ylim([min([round(size(img,2)*w2min(i3)),lims_y1_(i3)]),max([round(size(img,2)*w2max(i3)),lims_y2_(i3)])]),
        saveas(fighndl,[QCpath 'CSFmask_slice' sprintf('%.2d',i3) '.jpg'])
        close(fighndl)
        
        % mask_WM
        fighndl = figure('Position',[round((scrsz(3)-scrsz(4))/2) 1 scrsz(4) scrsz(4)],'InvertHardcopy','off','Visible','off'); fighndl.Color = [0.8, 0.87, 0.84];
        imshow(single(img), [0 mmax], 'InitialMagnification', 'fit'); colorbar;
        fighndl.Color = [0.8, 0.87, 0.84] + [0, 0.02*(1-exp(siz3/i3)/exp(siz3)), 0.1*exp(i3/siz3)/exp(1)]; % [0.85, 0.90, 0.85] + [0, 0, 0.1*i3/siz3];
        title({'Draw the boundary of the white matter (double click the cord once done)';['Slice ' num2str(i3) ' of ' num2str(siz3)]},'Color',[0.70,0.13,0.13],'FontSize',16);
        hold on, plot([position_CSF{i3}(:,1);position_CSF{i3}(1,1)],[position_CSF{i3}(:,2);position_CSF{i3}(1,2)],'Color',[0.7,0,0.7],'LineWidth',2),
            plot([position_WM{i3}(:,1);position_WM{i3}(1,1)],[position_WM{i3}(:,2);position_WM{i3}(1,2)],'Color',[0.70,0.13,0.13],'LineWidth',2), hold off
        xlim([round(lims_x1(i3))-1,round(lims_x2(i3))+1]), ylim([round(lims_y1(i3))-1,round(lims_y2(i3))+1]),
        legend('CSF mask','WM mask')
        saveas(fighndl,[QCpath 'WMmask_slice' sprintf('%.2d',i3) '.jpg'])
        lims_x1(i3) = min(position_WM{i3}(:,1)); lims_x2(i3) = max(position_WM{i3}(:,1));
        lims_y1(i3) = min(position_WM{i3}(:,2)); lims_y2(i3) = max(position_WM{i3}(:,2));
        close(fighndl)
        
        % mask_GM
        fighndl = figure('Position',[round((scrsz(3)-scrsz(4))/2) 1 scrsz(4) scrsz(4)],'InvertHardcopy','off','Visible','off'); fighndl.Color = [0.8, 0.87, 0.84];
        imshow(single(img), [0 mmax], 'InitialMagnification', 'fit'); colorbar;
        fighndl.Color = [0.8, 0.87, 0.84] + [0, 0.02*(1-exp(siz3/i3)/exp(siz3)), 0.1*exp(i3/siz3)/exp(1)]; % [0.85, 0.90, 0.85] + [0, 0, 0.1*i3/siz3];
        title({'Draw the boundary of the gray matter (double click the GM once done)';['Slice ' num2str(i3) ' of ' num2str(siz3)]},'Color',[0,0.33,0.53],'FontSize',16);
        hold on, plot([position_CSF{i3}(:,1);position_CSF{i3}(1,1)],[position_CSF{i3}(:,2);position_CSF{i3}(1,2)],'Color',[0.7,0,0.7],'LineWidth',2),
            plot([position_WM{i3}(:,1);position_WM{i3}(1,1)],[position_WM{i3}(:,2);position_WM{i3}(1,2)],'Color',[0.70,0.13,0.13],'LineWidth',2),
            plot([position_GM{i3}(:,1);position_GM{i3}(1,1)],[position_GM{i3}(:,2);position_GM{i3}(1,2)],'Color',[0,0.33,0.53],'LineWidth',2), hold off
        xlim([round(lims_x1(i3))-1,round(lims_x2(i3))+1]), ylim([round(lims_y1(i3))-1,round(lims_y2(i3))+1]),
        legend('CSF mask','WM mask','GM mask')
        saveas(fighndl,[QCpath 'GMmask_slice' sprintf('%.2d',i3) '.jpg'])
        close(fighndl)
        
        fighndl2 = figure('Position',[round((scrsz(3)-scrsz(4))/2) 1 scrsz(4) scrsz(4)],'InvertHardcopy','off','Visible','off');
        imshow(single(img), [0 mmax], 'InitialMagnification', 'fit'); colorbar;
        fighndl2.Color = [0.75, 0.84, 0.99];
        hold on, plot([position_CSF{i3}(:,1);position_CSF{i3}(1,1)],[position_CSF{i3}(:,2);position_CSF{i3}(1,2)],'Color',[0.7,0,0.7],'LineWidth',2), 
            plot([position_WM{i3}(:,1);position_WM{i3}(1,1)],[position_WM{i3}(:,2);position_WM{i3}(1,2)],'Color',[0.70,0.13,0.13],'LineWidth',2), 
            plot([position_GM{i3}(:,1);position_GM{i3}(1,1)],[position_GM{i3}(:,2);position_GM{i3}(1,2)],'Color',[0,0.33,0.53],'LineWidth',2), hold off
        legend('CSF mask','WM mask','GM mask'), title({sprintf('Final GM/WM/CSF masks for slice-%d',i3);'(click anywhere on the image to zoom/close)'},'Color','red','FontSize',18)
        saveas(fighndl2,[QCpath 'final_masks_slice' sprintf('%.2d',i3) '_fullFOV' '.jpg'])
        xlim([lims_x1_(i3),lims_x2_(i3)]), ylim([lims_y1_(i3),lims_y2_(i3)]),
        saveas(fighndl2,[QCpath 'final_masks_slice' sprintf('%.2d',i3) '_zoomed' '.jpg'])
        close(fighndl2)
        clear img mmax position
    end
    clear A position_GM position_WM position_CSF fighndl fighndl2 lims_x1 lims_x2 lims_y1 lims_y2 lims_y2_ lims_y1_ lims_x2_ lims_x1_
catch
end

% cross sectional areas fig
try
load([base_dir_sub 'cross_sectional_areas.mat'],'GM_area','WM_area','CSF_area','cross_sectional_area')
for w=1:length(GM_area)
    dispstring1{w} = sprintf('%.2f',GM_area(w));
    dispstring2{w} = sprintf('%.2f',WM_area(w));
    dispstring3{w} = sprintf('%.2f',CSF_area(w));
    dispstring4{w} = sprintf('%.2f',cross_sectional_area(w));
end; clear w
fighndl = figure('Position',[1 1 1280 800],'Color',[0.85,0.90,0.85],'InvertHardcopy','off','Visible','off'); % red [0.95,0.9,0.85], green [0.85,0.90,0.85], blue [0.89,0.94,0.99]
plot(GM_area,'--^','color',[0,0.55,0.69],'linewidth',2.25, 'MarkerSize',12, 'MarkerEdgeColor',[0,0.55,0.69], 'MarkerFaceColor',[0.89,0.94,0.99]),
    hold on, plot(WM_area,'--v','color',[0.65,0.11,0.19],'linewidth',2.25, 'MarkerSize',12, 'MarkerEdgeColor',[0.65,0.11,0.19], 'MarkerFaceColor',[0.95,0.9,0.85]),
    plot(CSF_area,'--o','color',[0.25,0.25,0.25],'linewidth',2.25, 'MarkerSize',14, 'MarkerEdgeColor','k', 'MarkerFaceColor',[0.85,0.85,0.85]),
    plot(cross_sectional_area,'--p','color','b','linewidth',2.25, 'MarkerSize',18, 'MarkerEdgeColor',[0 0.37 0.22], 'MarkerFaceColor',[0.85,0.90,0.85]),
    grid on,grid minor, ylabel('area (in millimeters squared)'), xlabel('SLICES'), xticks([1:length(GM_area)]), 
    xlim([0.5 length(GM_area)+0.5]), ylim([0 max([GM_area;WM_area;CSF_area;cross_sectional_area])+8])
    text([1:length(GM_area)]-0.1,GM_area+6, dispstring1,'Color',[0,0.55,0.69],'FontSize',12)
    text([1:length(WM_area)]-0.1,WM_area-6, dispstring2,'Color',[0.65,0.11,0.19],'FontSize',12)
    text([1:length(CSF_area)]-0.1,CSF_area+6, dispstring3,'Color',[0 0.37 0.22],'FontSize',12)
    text([1:length(cross_sectional_area)]-0.1,cross_sectional_area-6, dispstring4,'Color',[0.05,0.05,0.05],'FontSize',12)
    legend({'Gray matter area';'White matter area';'Cerebrospinal fluid area';'Cross-sectional area (GM+WM)'},'FontSize',14,'TextColor','k','Location','best')
    title(sprintf('CROSS SECTIONAL AREAS across slices (units: millimeters squared)\n(Note: these area measurements have not been validated, hence, please do not use this for research or publication purposes. \nThis is for rough quality assessment only.)'),'Color',[0.65,0.11,0.19],'FontSize',16)
saveas(fighndl,[QCpath 'cross_sectional_areas_across_slices' '.jpg'])
clear GM_area WM_area CSF_area cross_sectional_area dispstring1 dispstring2 dispstring3 dispstring4
close(fighndl)
catch
end
end

%% scfMRItb_11_FuncAnatRegistration
if choices_preproc(11)==1
clear QCpath save_videos fighndl
if ~(exist([base_dir_sub 'QC' '/11_func-anat-registration'],'dir'))
    mkdir([base_dir_sub 'QC' '/11_func-anat-registration'])
end
if Rns==0
    QCpath = [base_dir_sub 'QC' '/11_func-anat-registration/'];
else
    if ~(exist([base_dir_sub 'QC' '/11_func-anat-registration' '/run' num2str(Rns)],'dir'))
        mkdir([base_dir_sub 'QC' '/11_func-anat-registration' '/run' num2str(Rns)])
    end
    QCpath = [base_dir_sub 'QC' '/11_func-anat-registration' '/run' num2str(Rns) '/'];
end

if (save_videos_==0)||(save_videos_==1)
    save_videos = save_videos_;
elseif (save_videos_==2)&&(sum([choices_preproc(12:14)])==0)
    save_videos = 1;
else
    save_videos = 0;
end
if save_videos==1
    save_videos_steps(5)=1;
end


try
    try eval(['load ' base_dir_sub fname_anat '_all_masks.mat ' 'position_GM position_WM position_CSF;']); catch, end
    try
        scfMRItb_04_unzipFile(base_dir_sub, fname_anat, '_mask_SC')
        mask_SC2 = load_untouch_nii([base_dir_sub fname_anat '_mask_SC.nii']); mask_SC2 = mask_SC2.img;
    catch
        mask_SC = load_untouch_nii([base_dir_sub fname_anat '_mask_GM.nii']);
        mask_SC.img = or(mask_GM, mask_WM);
        save_untouch_nii(mask_SC, [base_dir_sub fname_anat '_mask_SC.nii']);
        mask_SC2 = mask_SC.img; clear mask_SC
    end
    
    for i3 = 1 : siz3
        if ~exist([base_dir_sub fname_anat '_slice' num2str(i3) '.nii'],'file')
            scfMRItb_04_resplitData(base_dir_sub, fname_anat, '', siz3);
        end
        if ~exist([base_dir_sub fname '_slice' num2str(i3) '_warped_mean.nii'],'file')
            scfMRItb_04_resplitData(base_dir_sub, fname, '_warped_mean', siz3);
        end
        img_anat = load_untouch_nii([base_dir_sub fname_anat '_slice' num2str(i3) '.nii']); img_anat=img_anat.img;
        img_warped_mean = load_untouch_nii([base_dir_sub fname '_slice' num2str(i3) '_warped_mean.nii']); img_warped_mean=img_warped_mean.img;
        fighndl = figure('Position',[1 1 1120 1120],'Color',[0.85,0.90,0.85],'InvertHardcopy','off','Visible','off'); % [0.85,0.90,0.85] [0.8, 0.89, 0.94] [0.89, 0.94, 0.99] [361 1 scrsz(4) scrsz(4)]
        subtightplot(2,2,1,[0.035 0.035], [0.035 0.035], [0.035 0.035]), imshow(rot90(img_anat), [0 max(img_anat(:))]), title(['Anatomical image: (slice ' num2str(i3) ' of ' num2str(siz3) ')'],'Color',[0.65,0.11,0.19],'FontSize',16)
            try xlim([round(min(position_CSF{i3}(:,1))-0.125*(size(mask_SC2,1))),round(max(position_CSF{i3}(:,1))+0.125*(size(mask_SC2,1)))]), ylim([round(min(position_CSF{i3}(:,2))-0.125*(size(mask_SC2,2))),round(max(position_CSF{i3}(:,2))+0.125*(size(mask_SC2,2)))]), catch, end
        subtightplot(2,2,2,[0.035 0.035], [0.035 0.035], [0.035 0.035]), imshow(rot90(img_warped_mean), [0 max(img_warped_mean(:))]), title(['Co-registered mean functional image: (slice ' num2str(i3) ' of ' num2str(siz3) ')'],'Color',[0,0.55,0.69],'FontSize',16)
            try xlim([round(min(position_CSF{i3}(:,1))-0.125*(size(mask_SC2,1))),round(max(position_CSF{i3}(:,1))+0.125*(size(mask_SC2,1)))]), ylim([round(min(position_CSF{i3}(:,2))-0.125*(size(mask_SC2,2))),round(max(position_CSF{i3}(:,2))+0.125*(size(mask_SC2,2)))]), catch, end
        subtightplot(2,2,3,[0.035 0.035], [0.035 0.035], [0.035 0.035]), imshow(rot90(img_anat), [0 max(img_anat(:))]), 
            xlim([round(size(img_anat,1)/3)+1,2*round(size(img_anat,1)/3),]), ylim([round(size(img_anat,2)/3)+1,2*round(size(img_anat,2)/3),]), title(['Anatomical image: (slice ' num2str(i3) ' of ' num2str(siz3) ')'],'Color',[0.65,0.11,0.19],'FontSize',16)
            try hold on, plot([position_CSF{i3}(:,1);position_CSF{i3}(1,1)],[position_CSF{i3}(:,2);position_CSF{i3}(1,2)],'Color',[0.7,0,0.7],'LineWidth',2), plot([position_WM{i3}(:,1);position_WM{i3}(1,1)],[position_WM{i3}(:,2);position_WM{i3}(1,2)],'Color',[0.70,0.13,0.13],'LineWidth',2), plot([position_GM{i3}(:,1);position_GM{i3}(1,1)],[position_GM{i3}(:,2);position_GM{i3}(1,2)],'Color',[0,0.33,0.53],'LineWidth',2), hold off, catch, end
            try xlim([round(min(position_CSF{i3}(:,1))-0.03*(size(mask_SC2,1))),round(max(position_CSF{i3}(:,1))+0.03*(size(mask_SC2,1)))]), ylim([round(min(position_CSF{i3}(:,2))-0.03*(size(mask_SC2,2))),round(max(position_CSF{i3}(:,2))+0.03*(size(mask_SC2,2)))]), catch, end
        subtightplot(2,2,4,[0.035 0.035], [0.035 0.035], [0.035 0.035]), imshow(rot90(img_warped_mean), [0 max(img_warped_mean(:))]), 
            xlim([round(size(img_warped_mean,1)/3)+1,2*round(size(img_warped_mean,1)/3)]), ylim([round(size(img_warped_mean,2)/3)+1,2*round(size(img_warped_mean,2)/3)]), title(['Co-registered mean functional image: (slice ' num2str(i3) ' of ' num2str(siz3) ')'],'Color',[0,0.55,0.69],'FontSize',16)
            try hold on, plot([position_CSF{i3}(:,1);position_CSF{i3}(1,1)],[position_CSF{i3}(:,2);position_CSF{i3}(1,2)],'Color',[0.7,0,0.7],'LineWidth',2), plot([position_WM{i3}(:,1);position_WM{i3}(1,1)],[position_WM{i3}(:,2);position_WM{i3}(1,2)],'Color',[0.70,0.13,0.13],'LineWidth',2), plot([position_GM{i3}(:,1);position_GM{i3}(1,1)],[position_GM{i3}(:,2);position_GM{i3}(1,2)],'Color',[0,0.33,0.53],'LineWidth',2), hold off, catch, end
            try xlim([round(min(position_CSF{i3}(:,1))-0.03*(size(mask_SC2,1))),round(max(position_CSF{i3}(:,1))+0.03*(size(mask_SC2,1)))]), ylim([round(min(position_CSF{i3}(:,2))-0.03*(size(mask_SC2,2))),round(max(position_CSF{i3}(:,2))+0.03*(size(mask_SC2,2)))]), catch, end
        try suptitle2(sprintf('Outcome of Step 11: functional to anatomical registration\nComparing co-registered mean functional image with the anatomical image for slice %s of %s',num2str(i3),num2str(siz3))); fighndl.Visible='off'; catch, end
        saveas(fighndl,[QCpath 'func_anat_reg_slice' sprintf('%.2d',i3) '.jpg'])
        close(fighndl)
        clear img_anat img_warped_mean
    end
    clear mask_SC2 fighndl
catch
end


F2g_=[]; mF2g_=[]; F2w_=[]; mF2w_=[]; Cg_=[]; Cw_=[]; F4_=[]; mF4_=[]; C2_=[];
if exist([base_dir_sub fname '_slice' num2str(i3) '_MC.nii'],'file') || exist([base_dir_sub fname '_slice' num2str(i3) '_MC.nii.gz'],'file') || exist([base_dir_sub fname '_MC.nii'],'file') || exist([base_dir_sub fname '_MC.nii.gz'],'file')
    F6_=[]; mF6_=[]; C3_=[];
end
for i3 = 1 : siz3
    try waitbar((i3/siz3),wbar3,sprintf('11. QC plots: slice (%d) of (%d)',i3,siz3)); catch, end
    fprintf('11. QC plots: slice (%d) of (%d), subject (%d) of (%d), run (%d) of (%d)\n',i3,siz3,k,size(func_files,1),k2,size(func_files,2))
    scfMRItb_04_resplitData(base_dir_sub, fname, '_warped', siz3) % if slice-wise data doesn't exist but combined 4D data exists, then split the data to generate the needed slice-wise data
    scfMRItb_04_unzipFile(base_dir_sub, fname, ['_slice' num2str(i3) '_warped'])
    F1 = load_untouch_nii([base_dir_sub fname '_slice' num2str(i3) '_warped.nii']);
    TR = F1.hdr.dime.pixdim(5); F1 = single(squeeze(F1.img));
    
    if Smask_err==0
        scfMRItb_04_resplitData(base_dir_sub, fname, '_Smask', siz3) % if slice-wise data doesn't exist but combined 4D data exists, then split the data to generate the needed slice-wise data
        scfMRItb_04_unzipFile(base_dir_sub, fname, ['_slice' num2str(i3) '_Smask'])
        Smask1 = load_untouch_nii([base_dir_sub fname '_slice' num2str(i3) '_Smask.nii']); Smask1 = Smask1.img;
    else
        Smask1 = Smask1_(:,:,i3);
    end
    
    for i5=1:size(F1,3)
        tmp = F1(:,:,i5);
        F2g(:,i5) = tmp(find(mask_GM(:,:,i3)==1));
        F2w(:,i5) = tmp(find(mask_WM(:,:,i3)==1));
        clear tmp
    end; clear i5 F1
    [~,svar1] = sort(var(F2g,0,2)); [~,sdist1] = sort(svar1); sortorder1{i3,1} = sdist1;
    [~,svar2] = sort(var(F2w,0,2)); [~,sdist2] = sort(svar2); sortorder2{i3,1} = sdist2;
    
    fighndl = figure('Position',[1 1 1152 720],'Color',[0.95,0.9,0.85],'InvertHardcopy','off','Visible','off'); % red [0.95,0.9,0.85], green [0.85,0.90,0.85], blue [0.89,0.94,0.99]
    subplot2n(2,1,1), imagesc(F2g(sortorder1{i3},:)),colormap(cmap),colorbar, xlabel('time (in samples)'),ylabel('Gray matter voxels (sorted by variance)'),
        title(['Time series within the GRAY MATTER after "11-functional-anatomical-registration" in slice ' num2str(i3) ' of ' num2str(siz3)],'Color',[0,0.55,0.69],'FontSize',14);
    subplot2n(2,1,2), imagesc(F2w(sortorder2{i3},:)),colormap(cmap),colorbar, xlabel('time (in samples)'),ylabel('White matter voxels (sorted by variance)'),
        title(['Time series within the WHITE MATTER after "11-functional-anatomical-registration" in slice ' num2str(i3) ' of ' num2str(siz3)],'Color',[0.65,0.11,0.19],'FontSize',14);
    saveas(fighndl,[QCpath 'fMRI_ts_11coreg_slice' sprintf('%.2d',i3) '.jpg'])
    close(fighndl)
    
    scfMRItb_04_resplitData(base_dir_sub, fname, '', siz3) % if slice-wise data doesn't exist but combined 4D data exists, then split the data to generate the needed slice-wise data
    scfMRItb_04_unzipFile(base_dir_sub, fname, ['_slice' num2str(i3)])
    F3 = load_untouch_nii([base_dir_sub fname '_slice' num2str(i3) '.nii']); F3 = single(squeeze(F3.img));
    for i5=1:size(F3,3)
        tmp = F3(:,:,i5);
        F4(:,i5) = tmp(find(Smask1==1)); clear tmp
    end; clear i5 F3
    mF2g=mean(F2g,1); mF2g = 100*(mF2g-mean(mF2g))./mean(mF2g); mF2w=mean(F2w,1); mF2w = 100*(mF2w-mean(mF2w))./mean(mF2w); mF4=mean(F4,1); mF4 = 100*(mF4-mean(mF4))./mean(mF4);
    fighndl = figure('Position',[1 1 1152 720],'Color',[0.95,0.9,0.85],'InvertHardcopy','off','Visible','off'); % red [0.95,0.9,0.85], green [0.85,0.90,0.85], blue [0.89,0.94,0.99]
    subplot2n(3,1,1), plot(mF4,'color','k','linewidth',1.5), hold on, plot(mF2g,'color',[0,0.55,0.69],'linewidth',0.2), plot(mF2w,'color',[0.65,0.11,0.19],'linewidth',0.2), xlabel('time (in samples)'),ylabel('percentage signal change'),
      grid on,grid minor, xlim([1 length(mF2g)]),ylim([min(1.25.*[mF2g,mF2w,mF4]) 1.25.*max([mF2g,mF2w,mF4])]), title(['Global mean signal in UNPROCESSED RAW DATA (slice ' num2str(i3) ' of ' num2str(siz3) ')'],'Color','k','FontSize',14), legend({'Global signal (raw data)'},'FontSize',12,'TextColor','k','Location','best')
    subplot2n(3,1,2), plot(mF2g,'color',[0,0.55,0.69],'linewidth',1.5), hold on, plot(mF2w,'color',[0.65,0.11,0.19],'linewidth',0.2), plot(mF4,'color','k','linewidth',0.2), xlabel('time (in samples)'),ylabel('percentage signal change'),
      grid on,grid minor, xlim([1 length(mF2g)]),ylim([min(1.25.*[mF2g,mF2w,mF4]) 1.25.*max([mF2g,mF2w,mF4])]), title(['Gray matter mean signal AFTER "11-functional-anatomical-registration" (slice ' num2str(i3) ' of ' num2str(siz3) ')'],'Color',[0,0.55,0.69],'FontSize',14), legend({'Gray matter signal after 11-registration'},'FontSize',12,'TextColor',[0,0.55,0.69],'Location','best')
    subplot2n(3,1,3), plot(mF2w,'color',[0.65,0.11,0.19],'linewidth',1.5), hold on, plot(mF4,'color','k','linewidth',0.2), plot(mF2g,'color',[0,0.55,0.69],'linewidth',0.2), xlabel('time (in samples)'),ylabel('percentage signal change'),
      grid on,grid minor, xlim([1 length(mF2g)]),ylim([min(1.25.*[mF2g,mF2w,mF4]) 1.25.*max([mF2g,mF2w,mF4])]), title(['White matter mean signal AFTER "11-functional-anatomical-registration" (slice ' num2str(i3) ' of ' num2str(siz3) ')'],'Color',[0.65,0.11,0.19],'FontSize',14), legend({'White matter signal after 11-registration'},'FontSize',12,'TextColor',[0.65,0.11,0.19],'Location','best')
    saveas(fighndl,[QCpath 'global_signal_11coreg_slice' sprintf('%.2d',i3) '.jpg'])
    close(fighndl)

    Cg = corrcoef(F2g'); Cg = uppertriangle(Cg);
    Cw = corrcoef(F2w'); Cw = uppertriangle(Cw);
    C2 = corrcoef(F4');  C2 = uppertriangle(C2);
    fighndl = figure('Position',[1 1 1152 720],'Color',[0.95,0.9,0.85],'InvertHardcopy','off','Visible','off'); % red [0.95,0.9,0.85], green [0.85,0.90,0.85], blue [0.89,0.94,0.99]
    subplot2n(3,1,1), histogram(C2,'EdgeColor','none','FaceColor','k','FaceAlpha',0.5,'Normalization','probability'); grid on,grid minor, xlim([-1 1]), hold on, xline(0,'Color','k','linewidth',1.5); hold off
      title(sprintf('Histogram of voxel-to-voxel correlations in UNPROCESSED RAW DATA (slice %d of %d)\n%.1f%% of the correlations were > 0  [ mean=%.2f, median=%.2f, SD=%.2f ]',i3,siz3,100*length(find(C2>0))/length(C2),mean(C2),median(C2),std(C2)),'Color','k','FontSize',14)
    subplot2n(3,1,2), histogram(Cg,'EdgeColor','none','FaceColor',[0,0.55,0.69],'FaceAlpha',0.5,'Normalization','probability'); grid on,grid minor, xlim([-1 1]), hold on, xline(0,'Color','k','linewidth',1.5); hold off
      title(sprintf('Histogram of voxel-to-voxel GRAY MATTER correlations AFTER 11-registration (slice %d of %d)\n%.1f%% of the correlations were > 0  [ mean=%.2f, median=%.2f, SD=%.2f ]',i3,siz3,100*length(find(Cg>0))/length(Cg),mean(Cg),median(Cg),std(Cg)),'Color',[0,0.55,0.69],'FontSize',14)
    subplot2n(3,1,3), histogram(Cw,'EdgeColor','none','FaceColor',[0.65,0.11,0.19],'FaceAlpha',0.5,'Normalization','probability'); grid on,grid minor, xlim([-1 1]), hold on, xline(0,'Color','k','linewidth',1.5); hold off
      title(sprintf('Histogram of voxel-to-voxel WHITE MATTER correlations AFTER 11-registration (slice %d of %d)\n%.1f%% of the correlations were > 0  [ mean=%.2f, median=%.2f, SD=%.2f ]',i3,siz3,100*length(find(Cw>0))/length(Cw),mean(Cw),median(Cw),std(Cw)),'Color',[0.65,0.11,0.19],'FontSize',14)
    saveas(fighndl,[QCpath 'hist_FC_compare_11coreg_slice' sprintf('%.2d',i3) '.jpg'])
    close(fighndl)
    
    if exist([base_dir_sub fname '_slice' num2str(i3) '_MC.nii'],'file') || exist([base_dir_sub fname '_slice' num2str(i3) '_MC.nii.gz'],'file') || exist([base_dir_sub fname '_MC.nii'],'file') || exist([base_dir_sub fname '_MC.nii.gz'],'file')
        scfMRItb_04_resplitData(base_dir_sub, fname, '_MC', siz3) % if slice-wise data doesn't exist but combined 4D data exists, then split the data to generate the needed slice-wise data
        scfMRItb_04_unzipFile(base_dir_sub, fname, ['_slice' num2str(i3) '_MC'])
        F5 = load_untouch_nii([base_dir_sub fname '_slice' num2str(i3) '_MC.nii']); F5 = single(squeeze(F5.img));
        for i5=1:size(F5,3)
            tmp = F5(:,:,i5);
            F6(:,i5) = tmp(find(Smask1==1)); clear tmp
        end; clear i5 F5
        mF6=mean(F6,1); mF6 = 100*(mF6-mean(mF6))./mean(mF6);
        fighndl = figure('Position',[1 1 1152 720],'Color',[0.95,0.9,0.85],'InvertHardcopy','off','Visible','off'); % red [0.95,0.9,0.85], green [0.85,0.90,0.85], blue [0.89,0.94,0.99]
        subplot2n(3,1,1), plot(mF6,'color','k','linewidth',1.5), hold on, plot(mF2g,'color',[0,0.55,0.69],'linewidth',0.2), plot(mF2w,'color',[0.65,0.11,0.19],'linewidth',0.2), xlabel('time (in samples)'),ylabel('percentage signal change'),
            grid on,grid minor, xlim([1 length(mF2g)]),ylim([1.25.*min([mF2g,mF2w,mF6]) 1.25.*max([mF2g,mF2w,mF6])]), title(['Global mean signal AFTER "08-motion-correction" (slice ' num2str(i3) ' of ' num2str(siz3) ')'],'Color','k','FontSize',14), legend({'Global signal before co-registration'},'FontSize',12,'TextColor','k','Location','best')
        subplot2n(3,1,2), plot(mF2g,'color',[0,0.55,0.69],'linewidth',1.5), hold on, plot(mF2w,'color',[0.65,0.11,0.19],'linewidth',0.2), plot(mF6,'color','k','linewidth',0.2), xlabel('time (in samples)'),ylabel('percentage signal change'),
            grid on,grid minor, xlim([1 length(mF2g)]),ylim([1.25.*min([mF2g,mF2w,mF6]) 1.25.*max([mF2g,mF2w,mF6])]), title(['Gray matter mean signal AFTER "11-functional-anatomical-registration" (slice ' num2str(i3) ' of ' num2str(siz3) ')'],'Color',[0,0.55,0.69],'FontSize',14), legend({'Gray matter signal after co-registration'},'FontSize',12,'TextColor',[0,0.55,0.69],'Location','best')
        subplot2n(3,1,3), plot(mF2w,'color',[0.65,0.11,0.19],'linewidth',1.5), hold on, plot(mF6,'color','k','linewidth',0.2), plot(mF2g,'color',[0,0.55,0.69],'linewidth',0.2), xlabel('time (in samples)'),ylabel('percentage signal change'),
            grid on,grid minor, xlim([1 length(mF2g)]),ylim([1.25.*min([mF2g,mF2w,mF6]) 1.25.*max([mF2g,mF2w,mF6])]), title(['White matter mean signal AFTER "11-functional-anatomical-registration" (slice ' num2str(i3) ' of ' num2str(siz3) ')'],'Color',[0.65,0.11,0.19],'FontSize',14), legend({'White matter signal after co-registration'},'FontSize',12,'TextColor',[0.65,0.11,0.19],'Location','best')
        saveas(fighndl,[QCpath 'global_signal_08moco_vs_11coreg_slice' sprintf('%.2d',i3) '.jpg'])
        close(fighndl)
        
        C3 = corrcoef(F6'); C3=uppertriangle(C3);
        fighndl = figure('Position',[1 1 1152 720],'Color',[0.95,0.9,0.85],'InvertHardcopy','off','Visible','off'); % red [0.95,0.9,0.85], green [0.85,0.90,0.85], blue [0.89,0.94,0.99]
        subplot2n(3,1,1), histogram(C3,'EdgeColor','none','FaceColor','k','FaceAlpha',0.5,'Normalization','probability'); grid on,grid minor, xlim([-1 1]), hold on, xline(0,'Color','k','linewidth',1.5); hold off
            title(sprintf('Histogram of voxel-to-voxel correlations AFTER 08-motion-correction (slice %d of %d)\n%.1f%% of the correlations were > 0  [ mean=%.2f, median=%.2f, SD=%.2f ]',i3,siz3,100*length(find(C3>0))/length(C3),mean(C3),median(C3),std(C3)),'Color','k','FontSize',14)
        subplot2n(3,1,2), histogram(Cg,'EdgeColor','none','FaceColor',[0,0.55,0.69],'FaceAlpha',0.5,'Normalization','probability'); grid on,grid minor, xlim([-1 1]), hold on, xline(0,'Color','k','linewidth',1.5); hold off
            title(sprintf('Histogram of voxel-to-voxel GRAY MATTER correlations AFTER 11-registration (slice %d of %d)\n%.1f%% of the correlations were > 0  [ mean=%.2f, median=%.2f, SD=%.2f ]',i3,siz3,100*length(find(Cg>0))/length(Cg),mean(Cg),median(Cg),std(Cg)),'Color',[0,0.55,0.69],'FontSize',14)
        subplot2n(3,1,3), histogram(Cw,'EdgeColor','none','FaceColor',[0.65,0.11,0.19],'FaceAlpha',0.5,'Normalization','probability'); grid on,grid minor, xlim([-1 1]), hold on, xline(0,'Color','k','linewidth',1.5); hold off
            title(sprintf('Histogram of voxel-to-voxel WHITE MATTER correlations AFTER 11-registration (slice %d of %d)\n%.1f%% of the correlations were > 0  [ mean=%.2f, median=%.2f, SD=%.2f ]',i3,siz3,100*length(find(Cw>0))/length(Cw),mean(Cw),median(Cw),std(Cw)),'Color',[0.65,0.11,0.19],'FontSize',14)
        saveas(fighndl,[QCpath 'hist_FC_compare_08moco_11coreg_slice' sprintf('%.2d',i3) '.jpg'])
        close(fighndl)
        F6_=cat(1,F6_,F6); mF6_=cat(1,mF6_,mF6); C3_=cat(1,C3_,C3);
        clear F6 mF6 C3
    end
    
    F2g_=cat(1,F2g_,F2g); mF2g_=cat(1,mF2g_,mF2g); F2w_=cat(1,F2w_,F2w); mF2w_=cat(1,mF2w_,mF2w); Cg_=cat(1,Cg_,Cg); Cw_=cat(1,Cw_,Cw); F4_=cat(1,F4_,F4); mF4_=cat(1,mF4_,mF4); C2_=cat(1,C2_,C2);
    clear F2g F2w F4 TR Smask1 svar1 svar2 mF2g mF2w mF4 Cg Cw C2
end

[~,svar1] = sort(var(F2g_,0,2)); [~,spsc1] = sort(meanPSC(F2g_));
[~,svar2] = sort(var(F2w_,0,2)); [~,spsc2] = sort(meanPSC(F2w_));
sortorder1b = sortorder1; sortorder1=[]; f1=0; %#ok<*ASGLU>
sortorder2b = sortorder2; sortorder2=[]; f2=0; %#ok<*ASGLU>
for im=1:length(sortorder1b)
    temp = sortorder1b{im}; temp = temp + f1; f1 = f1 + length(temp);
    sortorder1 = cat(1,sortorder1,temp); clear temp
    temp = sortorder2b{im}; temp = temp + f2; f2 = f2 + length(temp);
    sortorder2 = cat(1,sortorder2,temp); clear temp
end; clear im f1 f2
sortorder1 = spsc1; % options: spsc for percentage signal change, svar for variance, (comment out the line if you want to use distance from the center of the cord separately for each slice)
sortorder2 = spsc2; % options: spsc for percentage signal change, svar for variance, (comment out the line if you want to use distance from the center of the cord separately for each slice)
for im=max(sortorder1):-1:1, if isempty(find(sortorder1==im)), sortorder1(find(sortorder1>im)) = sortorder1(find(sortorder1>im))-1; end, end; clear im %#ok<*EFIND>
for im=max(sortorder2):-1:1, if isempty(find(sortorder2==im)), sortorder2(find(sortorder2>im)) = sortorder2(find(sortorder2>im))-1; end, end; clear im %#ok<*EFIND>

fighndl = figure('Position',[1 1 1152 720],'Color',[0.89,0.94,0.99],'InvertHardcopy','off','Visible','off'); % red [0.95,0.9,0.85], green [0.85,0.90,0.85], blue [0.89,0.94,0.99]
subplot2n(2,1,1), imagesc(F2g_(sortorder1,:)),colormap(cmap),colorbar, xlabel('time (in samples)'),ylabel(sprintf('voxels (sorted by mean percentage signal change;\nbottom row is highest)')),
  title(['Time series within the GRAY MATTER AFTER 11-func-anat-registration (all slices combined)'],'Color',[0,0.55,0.69],'FontSize',14);
subplot2n(2,1,2), imagesc(F2w_(sortorder2,:)),colormap(cmap),colorbar, xlabel('time (in samples)'),ylabel(sprintf('voxels (sorted by mean percentage signal change;\nbottom row is highest)')),
  title(['Time series within the WHITE MATTER AFTER 11-func-anat-registration (all slices combined)'],'Color',[0.65,0.11,0.19],'FontSize',14);
saveas(fighndl,[QCpath 'fMRI_ts_11coreg_allSlices' '.jpg'])
 close(fighndl)
mF2g_=mean(mF2g_,1); mF2w_=mean(mF2w_,1); mF4_=mean(mF4_,1);
fighndl = figure('Position',[1 1 1152 720],'Color',[0.89,0.94,0.99],'InvertHardcopy','off','Visible','off'); % red [0.95,0.9,0.85], green [0.85,0.90,0.85], blue [0.89,0.94,0.99]
subplot2n(3,1,1), plot(mF4_,'color','k','linewidth',1.5), hold on, plot(mF2g_,'color',[0,0.55,0.69],'linewidth',0.2), plot(mF2w_,'color',[0.65,0.11,0.19],'linewidth',0.2), xlabel('time (in samples)'),ylabel('percentage signal change'),
    grid on,grid minor, xlim([1 length(mF2g_)]),ylim([1.25.*min([mF2g_,mF2w_,mF4_]) 1.25.*max([mF2g_,mF2w_,mF4_])]), title(['Global mean signal in UNPROCESSED RAW DATA (all slices averaged)'],'Color','k','FontSize',14), legend({'Global signal (raw data)'},'FontSize',12,'TextColor','k','Location','best')
subplot2n(3,1,2), plot(mF2g_,'color',[0,0.55,0.69],'linewidth',1.5), hold on, plot(mF2w_,'color',[0.65,0.11,0.19],'linewidth',0.2), plot(mF4_,'color','k','linewidth',0.2), xlabel('time (in samples)'),ylabel('percentage signal change'),
    grid on,grid minor, xlim([1 length(mF2g_)]),ylim([1.25.*min([mF2g_,mF2w_,mF4_]) 1.25.*max([mF2g_,mF2w_,mF4_])]), title(['Gray matter mean signal AFTER "11-functional-anatomical-registration" (all slices averaged)'],'Color',[0,0.55,0.69],'FontSize',14), legend({'Gray matter signal after 11-registration'},'FontSize',12,'TextColor',[0,0.55,0.69],'Location','best')
subplot2n(3,1,3), plot(mF2w_,'color',[0.65,0.11,0.19],'linewidth',1.5), hold on, plot(mF4_,'color','k','linewidth',0.2), plot(mF2g_,'color',[0,0.55,0.69],'linewidth',0.2), xlabel('time (in samples)'),ylabel('percentage signal change'),
    grid on,grid minor, xlim([1 length(mF2g_)]),ylim([1.25.*min([mF2g_,mF2w_,mF4_]) 1.25.*max([mF2g_,mF2w_,mF4_])]), title(['White matter mean signal AFTER "11-functional-anatomical-registration" (all slices averaged)'],'Color',[0.65,0.11,0.19],'FontSize',14), legend({'White matter signal after 11-registration'},'FontSize',12,'TextColor',[0.65,0.11,0.19],'Location','best')
saveas(fighndl,[QCpath 'global_signal_11coreg_allSlices' '.jpg'])
 close(fighndl)
fighndl = figure('Position',[1 1 1152 720],'Color',[0.89,0.94,0.99],'InvertHardcopy','off','Visible','off'); % red [0.95,0.9,0.85], green [0.85,0.90,0.85], blue [0.89,0.94,0.99]
subplot2n(3,1,1), histogram(C2_,'EdgeColor','none','FaceColor','k','FaceAlpha',0.5,'Normalization','probability'); grid on,grid minor, xlim([-1 1]), hold on, xline(0,'Color','k','linewidth',1.5); hold off
    title(sprintf('Histogram of voxel-to-voxel correlations in UNPROCESSED RAW DATA (all slices combined)\n%.1f%% of the correlations were > 0  [ mean=%.2f, median=%.2f, SD=%.2f ]',100*length(find(C2_>0))/length(C2_),mean(C2_),median(C2_),std(C2_)),'Color','k','FontSize',14)
subplot2n(3,1,2), histogram(Cg_,'EdgeColor','none','FaceColor',[0,0.55,0.69],'FaceAlpha',0.5,'Normalization','probability'); grid on,grid minor, xlim([-1 1]), hold on, xline(0,'Color','k','linewidth',1.5); hold off
    title(sprintf('Histogram of voxel-to-voxel GRAY MATTER correlations AFTER 11-registration (all slices combined)\n%.1f%% of the correlations were > 0  [ mean=%.2f, median=%.2f, SD=%.2f ]',100*length(find(Cg_>0))/length(Cg_),mean(Cg_),median(Cg_),std(Cg_)),'Color',[0,0.55,0.69],'FontSize',14)
subplot2n(3,1,3), histogram(Cw_,'EdgeColor','none','FaceColor',[0.65,0.11,0.19],'FaceAlpha',0.5,'Normalization','probability'); grid on,grid minor, xlim([-1 1]), hold on, xline(0,'Color','k','linewidth',1.5); hold off
    title(sprintf('Histogram of voxel-to-voxel WHITE MATTER correlations AFTER 11-registration (all slices combined)\n%.1f%% of the correlations were > 0  [ mean=%.2f, median=%.2f, SD=%.2f ]',100*length(find(Cw_>0))/length(Cw_),mean(Cw_),median(Cw_),std(Cw_)),'Color',[0.65,0.11,0.19],'FontSize',14)
saveas(fighndl,[QCpath 'hist_FC_compare_11coreg_allSlices' '.jpg'])
 close(fighndl)
if exist([base_dir_sub fname '_slice' num2str(i3) '_MC.nii'],'file') || exist([base_dir_sub fname '_slice' num2str(i3) '_MC.nii.gz'],'file') || exist([base_dir_sub fname '_MC.nii'],'file') || exist([base_dir_sub fname '_MC.nii.gz'],'file')
    mF6_=mean(mF6_,1);
    fighndl = figure('Position',[1 1 1152 720],'Color',[0.89,0.94,0.99],'InvertHardcopy','off','Visible','off'); % red [0.95,0.9,0.85], green [0.85,0.90,0.85], blue [0.89,0.94,0.99]
    subplot2n(3,1,1), plot(mF6_,'color','k','linewidth',1.5), hold on, plot(mF2g_,'color',[0,0.55,0.69],'linewidth',0.2), plot(mF2w_,'color',[0.65,0.11,0.19],'linewidth',0.2), xlabel('time (in samples)'),ylabel('percentage signal change'),
        grid on,grid minor, xlim([1 length(mF2g_)]),ylim([1.25.*min([mF2g_,mF2w_,mF6_]) 1.25.*max([mF2g_,mF2w_,mF6_])]), title(['Global mean signal AFTER "08-motion-correction" (all slices averaged)'],'Color','k','FontSize',14), legend({'Global signal before co-registration'},'FontSize',12,'TextColor','k','Location','best')
    subplot2n(3,1,2), plot(mF2g_,'color',[0,0.55,0.69],'linewidth',1.5), hold on, plot(mF2w_,'color',[0.65,0.11,0.19],'linewidth',0.2), plot(mF6_,'color','k','linewidth',0.2), xlabel('time (in samples)'),ylabel('percentage signal change'),
        grid on,grid minor, xlim([1 length(mF2g_)]),ylim([1.25.*min([mF2g_,mF2w_,mF6_]) 1.25.*max([mF2g_,mF2w_,mF6_])]), title(['Gray matter mean signal AFTER "11-functional-anatomical-registration" (all slices averaged)'],'Color',[0,0.55,0.69],'FontSize',14), legend({'Gray matter signal after co-registration'},'FontSize',12,'TextColor',[0,0.55,0.69],'Location','best')
    subplot2n(3,1,3), plot(mF2w_,'color',[0.65,0.11,0.19],'linewidth',1.5), hold on, plot(mF6_,'color','k','linewidth',0.2), plot(mF2g_,'color',[0,0.55,0.69],'linewidth',0.2), xlabel('time (in samples)'),ylabel('percentage signal change'),
        grid on,grid minor, xlim([1 length(mF2g_)]),ylim([1.25.*min([mF2g_,mF2w_,mF6_]) 1.25.*max([mF2g_,mF2w_,mF6_])]), title(['White matter mean signal AFTER "11-functional-anatomical-registration" (all slices averaged)'],'Color',[0.65,0.11,0.19],'FontSize',14), legend({'White matter signal after co-registration'},'FontSize',12,'TextColor',[0.65,0.11,0.19],'Location','best')
    saveas(fighndl,[QCpath 'global_signal_08moco_vs_11coreg_allSlices' '.jpg'])
     close(fighndl)
     fighndl = figure('Position',[1 1 1152 720],'Color',[0.89,0.94,0.99],'InvertHardcopy','off','Visible','off'); % red [0.95,0.9,0.85], green [0.85,0.90,0.85], blue [0.89,0.94,0.99]
    subplot2n(3,1,1), histogram(C3_,'EdgeColor','none','FaceColor','k','FaceAlpha',0.5,'Normalization','probability'); grid on,grid minor, xlim([-1 1]), hold on, xline(0,'Color','k','linewidth',1.5); hold off
       title(sprintf('Histogram of voxel-to-voxel correlations AFTER 08-motion-correction (all slices combined)\n%.1f%% of the correlations were > 0  [ mean=%.2f, median=%.2f, SD=%.2f ]',100*length(find(C3_>0))/length(C3_),mean(C3_),median(C3_),std(C3_)),'Color','k','FontSize',14)
    subplot2n(3,1,2), histogram(Cg_,'EdgeColor','none','FaceColor',[0,0.55,0.69],'FaceAlpha',0.5,'Normalization','probability'); grid on,grid minor, xlim([-1 1]), hold on, xline(0,'Color','k','linewidth',1.5); hold off
       title(sprintf('Histogram of voxel-to-voxel GRAY MATTER correlations AFTER 11-registration (all slices combined)\n%.1f%% of the correlations were > 0  [ mean=%.2f, median=%.2f, SD=%.2f ]',100*length(find(Cg_>0))/length(Cg_),mean(Cg_),median(Cg_),std(Cg_)),'Color',[0,0.55,0.69],'FontSize',14)
    subplot2n(3,1,3), histogram(Cw_,'EdgeColor','none','FaceColor',[0.65,0.11,0.19],'FaceAlpha',0.5,'Normalization','probability'); grid on,grid minor, xlim([-1 1]), hold on, xline(0,'Color','k','linewidth',1.5); hold off
       title(sprintf('Histogram of voxel-to-voxel WHITE MATTER correlations AFTER 11-registration (all slices combined)\n%.1f%% of the correlations were > 0  [ mean=%.2f, median=%.2f, SD=%.2f ]',100*length(find(Cw_>0))/length(Cw_),mean(Cw_),median(Cw_),std(Cw_)),'Color',[0.65,0.11,0.19],'FontSize',14)
    saveas(fighndl,[QCpath 'hist_FC_compare_08moco_11coreg_allSlices' '.jpg'])
     close(fighndl)
end
clear svar1 spsc1 svar2 spsc2 sortorder1 sortorder2 F2g_ mF2g_ F2w_ mF2w_ Cg_ Cw_ F4_ mF4_ C2_ F6_ mF6_ C3_

%%% Make QC video (needs audio video toolbox in Matlab)
if ~isempty(which('VideoWriter')) && (save_videos==1) % check if Matlab's audio video toolbox exists

subplot = @(m,n,p) subtightplot(m, n, p, [0.001 0.005], [0.01 0.001], [0.01 0.001]);
scfMRItb_04_unzipFile(base_dir_sub, fname_anat, '_mask_SC')
try
    scfMRItb_04_unzipFile(base_dir_sub, fname_anat, '_mask_SC')
    mask_SC2 = load_untouch_nii([base_dir_sub fname_anat '_mask_SC.nii']); mask_SC2 = mask_SC2.img;
catch
    mask_SC = load_untouch_nii([base_dir_sub fname_anat '_mask_GM.nii']);
    mask_SC.img = or(mask_GM, mask_WM);
    save_untouch_nii(mask_SC, [base_dir_sub fname_anat '_mask_SC.nii']);
    mask_SC2 = mask_SC.img; clear mask_SC
end
for M=1:size(mask_SC2,3)
    mask_SC = rot90(mask_SC2(:,:,M));
    [xmask_SC,ymask_SC] = find(mask_SC==1);
    minx_SC(:,M) = round(min(xmask_SC)-0.1*(size(mask_SC,1))); maxx_SC(:,M)=round(max(xmask_SC)+0.1*(size(mask_SC,1)));
    miny_SC(:,M) = round(min(ymask_SC)-0.1*(size(mask_SC,2))); maxy_SC(:,M)=round(max(ymask_SC)+0.1*(size(mask_SC,2)));
    if (maxx_SC(:,M)-minx_SC(:,M))<(maxy_SC(:,M)-miny_SC(:,M))
        factr = ((maxy_SC(:,M)-miny_SC(:,M))-(maxx_SC(:,M)-minx_SC(:,M)))/2;
        if factr==round(factr)
            maxx_SC(:,M)=maxx_SC(:,M)+factr; minx_SC(:,M)=minx_SC(:,M)-factr;
        else
            maxx_SC(:,M)=maxx_SC(:,M)+factr+0.5; minx_SC(:,M)=minx_SC(:,M)-factr+0.5;
        end
    elseif (maxy_SC(:,M)-miny_SC(:,M))<(maxx_SC(:,M)-minx_SC(:,M))
        factr = ((maxx_SC(:,M)-minx_SC(:,M))-(maxy_SC(:,M)-miny_SC(:,M)))/2;
        if factr==round(factr)
            maxy_SC(:,M)=maxy_SC(:,M)+factr; miny_SC(:,M)=miny_SC(:,M)-factr;
        else
            maxy_SC(:,M)=maxy_SC(:,M)+factr+0.5; miny_SC(:,M)=miny_SC(:,M)-factr+0.5;
        end
    end
    clear mask_SC xmask_SC ymask_SC factr
end; clear M

for i3 = 1 : siz3
    try waitbar((i3/siz3),wbar3,sprintf('11. Generate QC video: slice (%d) of (%d)',i3,siz3)); catch, end
    fprintf('11. Generate QC video: slice (%d) of (%d), subject (%d) of (%d), run (%d) of (%d)\n',i3,siz3,k,size(func_files,1),k2,size(func_files,2))
    scfMRItb_04_resplitData(base_dir_sub, fname, '_warped', siz3) % if slice-wise data doesn't exist but combined 4D data exists, then split the data to generate the needed slice-wise data
    F1 = load_untouch_nii([base_dir_sub fname '_slice' num2str(i3) '_warped.nii']);
    TR = F1.hdr.dime.pixdim(5); F1 = rot90(single(squeeze(F1.img)));
    video = VideoWriter([QCpath 'fMRIvideo_11coreg_slice' sprintf('%.2d',i3) '.mp4'],'MPEG-4'); %#ok<*TNMLP> % create the video object
    video.FrameRate = round(30/TR); % 30 seconds of data in every second of the video
    video.Quality = 90; % very limited compression so that the images are shown as is (and thus avoid misunderstandings due to video quality)
    open(video); % open the file for writing
    
    [sizeFx_SC,sizeFy_SC] = size(F1(:,:,1));
    F2 = F1(minx_SC:maxx_SC,miny_SC:maxy_SC,:);
    for i5=1:size(F1,3)
        F1(:,:,i5) = imresize(F2(:,:,i5),[sizeFx_SC sizeFy_SC]);
    end
    
    if Smask_err==0
        scfMRItb_04_resplitData(base_dir_sub, fname, '_Smask', siz3) % if slice-wise data doesn't exist but combined 4D data exists, then split the data to generate the needed slice-wise data
        scfMRItb_04_unzipFile(base_dir_sub, fname, ['_slice' num2str(i3) '_Smask'])
        Smask1 = load_untouch_nii([base_dir_sub fname '_slice' num2str(i3) '_Smask.nii']); Smask1 = Smask1.img;
    else
        Smask1 = Smask1_(:,:,i3);
    end
    Smask1 = rot90(Smask1);
    [xmask,ymask] = find(Smask1==1);
    minx=round(min(xmask)-0.1*(size(Smask1,1))); maxx=round(max(xmask)+0.1*(size(Smask1,1)));
    miny=round(min(ymask)-0.1*(size(Smask1,2))); maxy=round(max(ymask)+0.1*(size(Smask1,2)));
    if (maxx-minx)<(maxy-miny)
        factr = ((maxy-miny)-(maxx-minx))/2;
        if factr==round(factr)
            maxx=maxx+factr; minx=minx-factr;
        else
            maxx=maxx+factr+0.5; minx=minx-factr+0.5;
        end
    elseif (maxy-miny)<(maxx-minx)
        factr = ((maxx-minx)-(maxy-miny))/2;
        if factr==round(factr)
            maxy=maxy+factr; miny=miny-factr;
        else
            maxy=maxy+factr+0.5; miny=miny-factr+0.5;
        end
    end
    scfMRItb_04_resplitData(base_dir_sub, fname, '', siz3) % if slice-wise data doesn't exist but combined 4D data exists, then split the data to generate the needed slice-wise data
    F3 = load_untouch_nii([base_dir_sub fname '_slice' num2str(i3) '.nii']); F3 = rot90(single(squeeze(F3.img)));
    [sizeFx,sizeFy] = size(F3(:,:,1));
    F4 = F3(minx:maxx,miny:maxy,:);
    for i5=1:size(F3,3)
        F3(:,:,i5) = imresize(F4(:,:,i5),[sizeFx sizeFy]);
    end
    clear maxx maxy minx miny xmask ymask Smask1 factr sizeFx sizeFy F2 F4 i5 sizeFx_SC sizeFy_SC
    
    for ii=1:size(F1,3) % loop across the number of images
        I = F1(:,:,ii); %read the next image
        I(find(isnan(I)))=0;
        I = (I - min(I(:))) ./ (max(I(:)) - min(I(:)));
        I2 = F3(:,:,ii); %read the next image
        I2(find(isnan(I2)))=0;
        I2 = (I2 - min(I2(:))) ./ (max(I2(:)) - min(I2(:)));
        fighndl = figure('Position',[round((scrsz(3)-scrsz(4))/2) 1 round(0.75*scrsz(3)) round(0.75*scrsz(4))],'Color',[0.95,0.9,0.85],'InvertHardcopy','off','Visible','off'); % [0.85,0.90,0.85]
        subplot(1,2,1), imshow(I2, [0 max(I2(:))], 'InitialMagnification', 'fit'); title({['Unprocessed raw fMRI data (slice ' num2str(i3) ' of ' num2str(siz3) ')'];['Time point ' num2str(ii) ' of ' num2str(size(F1,3))]},'Color','k','FontSize',18);
        subplot(1,2,2), imshow(I, [0 max(I(:))], 'InitialMagnification', 'fit'); title({['fMRI data after "11. func-anat-registration" (slice ' num2str(i3) ' of ' num2str(siz3) ')'];['Time point ' num2str(ii) ' of ' num2str(size(F1,3))]},'Color',[0,0.55,0.69],'FontSize',18);
        frame = getframe(gcf);
        writeVideo(video,frame); % write the image to file
        close(fighndl)
        clear I I2 frame
    end
    close(video); %close the file
    clear F1 F3 video ii TR
end; clear i3
clear maxx_SC maxy_SC minx_SC miny_SC F
end

end

%% scfMRItb_12_CSFregress
if choices_preproc(12)==1
clear QCpath save_videos fighndl
if ~(exist([base_dir_sub 'QC' '/12_denoise3-CSFregress'],'dir'))
    mkdir([base_dir_sub 'QC' '/12_denoise3-CSFregress'])
end
if Rns==0
    QCpath = [base_dir_sub 'QC' '/12_denoise3-CSFregress/'];
else
    if ~(exist([base_dir_sub 'QC' '/12_denoise3-CSFregress' '/run' num2str(Rns)],'dir'))
        mkdir([base_dir_sub 'QC' '/12_denoise3-CSFregress' '/run' num2str(Rns)])
    end
    QCpath = [base_dir_sub 'QC' '/12_denoise3-CSFregress' '/run' num2str(Rns) '/'];
end

if (save_videos_==0)||(save_videos_==1)
    save_videos = save_videos_;
elseif (save_videos_==2)&&(sum([choices_preproc(13:14)])==0)
    save_videos = 2;
else
    save_videos = 0;
end
if save_videos==1
    save_videos_steps(6)=1;
end


F2g_=[]; mF2g_=[]; F2w_=[]; mF2w_=[]; Cg_=[]; Cw_=[]; F4_=[]; mF4_=[]; C2_=[];
if exist([base_dir_sub fname '_slice' num2str(i3) '_warped.nii'],'file') || exist([base_dir_sub fname '_slice' num2str(i3) '_warped.nii.gz'],'file') || exist([base_dir_sub fname '_warped.nii'],'file') || exist([base_dir_sub fname '_warped.nii.gz'],'file')
    F6g_=[]; mF6g_=[]; F6w_=[]; mF6w_=[]; C3g_=[]; C3w_=[];
end
for i3 = 1 : siz3
    try waitbar((i3/siz3),wbar3,sprintf('12. QC plots: slice (%d) of (%d)',i3,siz3)); catch, end
    fprintf('12. QC plots: slice (%d) of (%d), subject (%d) of (%d), run (%d) of (%d)\n',i3,siz3,k,size(func_files,1),k2,size(func_files,2))
    scfMRItb_04_resplitData(base_dir_sub, fname, ['_denoised' cutoff_str], siz3) % if slice-wise data doesn't exist but combined 4D data exists, then split the data to generate the needed slice-wise data
    scfMRItb_04_unzipFile(base_dir_sub, fname, ['_slice' num2str(i3) '_denoised' cutoff_str])
    F1 = load_untouch_nii([base_dir_sub fname '_slice' num2str(i3) '_denoised' cutoff_str '.nii']);
    TR = F1.hdr.dime.pixdim(5); F1 = single(squeeze(F1.img));
    
    if Smask_err==0
        scfMRItb_04_resplitData(base_dir_sub, fname, '_Smask', siz3) % if slice-wise data doesn't exist but combined 4D data exists, then split the data to generate the needed slice-wise data
        scfMRItb_04_unzipFile(base_dir_sub, fname, ['_slice' num2str(i3) '_Smask'])
        Smask1 = load_untouch_nii([base_dir_sub fname '_slice' num2str(i3) '_Smask.nii']); Smask1 = Smask1.img;
    else
        Smask1 = Smask1_(:,:,i3);
    end

    for i5=1:size(F1,3)
        tmp = F1(:,:,i5);
        F2g(:,i5) = tmp(find(mask_GM(:,:,i3)==1));
        F2w(:,i5) = tmp(find(mask_WM(:,:,i3)==1));
        clear tmp
    end; clear i5 F1
    [~,svar1] = sort(var(F2g,0,2)); [~,sdist1] = sort(svar1); sortorder1{i3,1} = sdist1;
    [~,svar2] = sort(var(F2w,0,2)); [~,sdist2] = sort(svar2); sortorder2{i3,1} = sdist2;
    
    fighndl = figure('Position',[1 1 1152 720],'Color',[0.95,0.9,0.85],'InvertHardcopy','off','Visible','off'); % red [0.95,0.9,0.85], green [0.85,0.90,0.85], blue [0.89,0.94,0.99]
    subplot2n(2,1,1), imagesc(F2g(sortorder1{i3},:)),colormap(cmap),colorbar, xlabel('time (in samples)'),ylabel('Gray matter voxels (sorted by variance)'),
        title(['Time series within the GRAY MATTER after "12-DENOISE3-CSFregress" in slice ' num2str(i3) ' of ' num2str(siz3)],'Color',[0,0.55,0.69],'FontSize',14);
    subplot2n(2,1,2), imagesc(F2w(sortorder2{i3},:)),colormap(cmap),colorbar, xlabel('time (in samples)'),ylabel('White matter voxels (sorted by variance)'),
        title(['Time series within the WHITE MATTER after "12-DENOISE3-CSFregress" in slice ' num2str(i3) ' of ' num2str(siz3)],'Color',[0.65,0.11,0.19],'FontSize',14);
    saveas(fighndl,[QCpath 'fMRI_ts_12denoise3_slice' sprintf('%.2d',i3) '.jpg'])
    close(fighndl)
    
    scfMRItb_04_resplitData(base_dir_sub, fname, '', siz3) % if slice-wise data doesn't exist but combined 4D data exists, then split the data to generate the needed slice-wise data
	scfMRItb_04_unzipFile(base_dir_sub, fname, ['_slice' num2str(i3)])
    F3 = load_untouch_nii([base_dir_sub fname '_slice' num2str(i3) '.nii']); F3 = single(squeeze(F3.img));
    for i5=1:size(F3,3)
        tmp = F3(:,:,i5);
        F4(:,i5) = tmp(find(Smask1==1)); clear tmp
    end; clear i5 F3
    mF2g=mean(F2g,1); mF2g = 100*(mF2g-mean(mF2g))./mean(mF2g); mF2w=mean(F2w,1); mF2w = 100*(mF2w-mean(mF2w))./mean(mF2w);
    mF4=mean(F4,1); mF4 = 100*(mF4-mean(mF4))./mean(mF4);
    fighndl = figure('Position',[1 1 1152 720],'Color',[0.95,0.9,0.85],'InvertHardcopy','off','Visible','off'); % red [0.95,0.9,0.85], green [0.85,0.90,0.85], blue [0.89,0.94,0.99]
    subplot2n(3,1,1), plot(mF4,'color','k','linewidth',1.5), hold on, plot(mF2g,'color',[0,0.55,0.69],'linewidth',0.2), plot(mF2w,'color',[0.65,0.11,0.19],'linewidth',0.2), xlabel('time (in samples)'),ylabel('percentage signal change'),
      grid on,grid minor, xlim([1 length(mF2g)]),ylim([min(1.25.*[mF2g,mF2w,mF4]) 1.25.*max([mF2g,mF2w,mF4])]), title(['Global mean signal in UNPROCESSED RAW DATA (slice ' num2str(i3) ' of ' num2str(siz3) ')'],'Color','k','FontSize',14), legend({'Global signal (raw data)'},'FontSize',12,'TextColor','k','Location','best')
    subplot2n(3,1,2), plot(mF2g,'color',[0,0.55,0.69],'linewidth',1.5), hold on, plot(mF2w,'color',[0.65,0.11,0.19],'linewidth',0.2), plot(mF4,'color','k','linewidth',0.2), xlabel('time (in samples)'),ylabel('percentage signal change'),
      grid on,grid minor, xlim([1 length(mF2g)]),ylim([min(1.25.*[mF2g,mF2w,mF4]) 1.25.*max([mF2g,mF2w,mF4])]), title(['Gray matter mean signal AFTER "12-DENOISE3-CSFregress" (slice ' num2str(i3) ' of ' num2str(siz3) ')'],'Color',[0,0.55,0.69],'FontSize',14), legend({'Gray matter signal after 12-denoise3'},'FontSize',12,'TextColor',[0,0.55,0.69],'Location','best')
    subplot2n(3,1,3), plot(mF2w,'color',[0.65,0.11,0.19],'linewidth',1.5), hold on, plot(mF4,'color','k','linewidth',0.2), plot(mF2g,'color',[0,0.55,0.69],'linewidth',0.2), xlabel('time (in samples)'),ylabel('percentage signal change'),
      grid on,grid minor, xlim([1 length(mF2g)]),ylim([min(1.25.*[mF2g,mF2w,mF4]) 1.25.*max([mF2g,mF2w,mF4])]), title(['White matter mean signal AFTER "12-DENOISE3-CSFregress" (slice ' num2str(i3) ' of ' num2str(siz3) ')'],'Color',[0.65,0.11,0.19],'FontSize',14), legend({'White matter signal after 12-denoise3'},'FontSize',12,'TextColor',[0.65,0.11,0.19],'Location','best')
    saveas(fighndl,[QCpath 'global_signal_12denoise3_slice' sprintf('%.2d',i3) '.jpg'])
    close(fighndl)

    Cg = corrcoef(F2g'); Cg = uppertriangle(Cg);
    Cw = corrcoef(F2w'); Cw = uppertriangle(Cw);
    C2 = corrcoef(F4');  C2 = uppertriangle(C2);
    fighndl = figure('Position',[1 1 1152 720],'Color',[0.95,0.9,0.85],'InvertHardcopy','off','Visible','off'); % red [0.95,0.9,0.85], green [0.85,0.90,0.85], blue [0.89,0.94,0.99]
    subplot2n(3,1,1), histogram(C2,'EdgeColor','none','FaceColor','k','FaceAlpha',0.5,'Normalization','probability'); grid on,grid minor, xlim([-1 1]), hold on, xline(0,'Color','k','linewidth',1.5); hold off
      title(sprintf('Histogram of voxel-to-voxel correlations in UNPROCESSED RAW DATA (slice %d of %d)\n%.1f%% of the correlations were > 0  [ mean=%.2f, median=%.2f, SD=%.2f ]',i3,siz3,100*length(find(C2>0))/length(C2),mean(C2),median(C2),std(C2)),'Color','k','FontSize',14)
    subplot2n(3,1,2), histogram(Cg,'EdgeColor','none','FaceColor',[0,0.55,0.69],'FaceAlpha',0.5,'Normalization','probability'); grid on,grid minor, xlim([-1 1]), hold on, xline(0,'Color','k','linewidth',1.5); hold off
      title(sprintf('Histogram of voxel-to-voxel GRAY MATTER correlations AFTER 12-DENOISE3-CSFregress (slice %d of %d)\n%.1f%% of the correlations were > 0  [ mean=%.2f, median=%.2f, SD=%.2f ]',i3,siz3,100*length(find(Cg>0))/length(Cg),mean(Cg),median(Cg),std(Cg)),'Color',[0,0.55,0.69],'FontSize',14)
    subplot2n(3,1,3), histogram(Cw,'EdgeColor','none','FaceColor',[0.65,0.11,0.19],'FaceAlpha',0.5,'Normalization','probability'); grid on,grid minor, xlim([-1 1]), hold on, xline(0,'Color','k','linewidth',1.5); hold off
      title(sprintf('Histogram of voxel-to-voxel WHITE MATTER correlations AFTER 12-DENOISE3-CSFregress (slice %d of %d)\n%.1f%% of the correlations were > 0  [ mean=%.2f, median=%.2f, SD=%.2f ]',i3,siz3,100*length(find(Cw>0))/length(Cw),mean(Cw),median(Cw),std(Cw)),'Color',[0.65,0.11,0.19],'FontSize',14)
    saveas(fighndl,[QCpath 'hist_FC_compare_12denoise3_slice' sprintf('%.2d',i3) '.jpg'])
    close(fighndl)
    
    if exist([base_dir_sub fname '_slice' num2str(i3) '_warped.nii'],'file') || exist([base_dir_sub fname '_slice' num2str(i3) '_warped.nii.gz'],'file') || exist([base_dir_sub fname '_warped.nii'],'file') || exist([base_dir_sub fname '_warped.nii.gz'],'file')
        scfMRItb_04_resplitData(base_dir_sub, fname, '_warped', siz3) % if slice-wise data doesn't exist but combined 4D data exists, then split the data to generate the needed slice-wise data
        scfMRItb_04_unzipFile(base_dir_sub, fname, ['_slice' num2str(i3) '_warped'])
        F5 = load_untouch_nii([base_dir_sub fname '_slice' num2str(i3) '_warped.nii']); F5 = single(squeeze(F5.img));
        for i5=1:size(F5,3)
            tmp = F5(:,:,i5);
            F6g(:,i5) = tmp(find(mask_GM(:,:,i3)==1));
            F6w(:,i5) = tmp(find(mask_WM(:,:,i3)==1));
            clear tmp
        end; clear i5 F5
        mF6g=mean(F6g,1); mF6g = 100*(mF6g-mean(mF6g))./mean(mF6g); mF6w=mean(F6w,1); mF6w = 100*(mF6w-mean(mF6w))./mean(mF6w);
        fighndl = figure('Position',[1 1 1152 720],'Color',[0.95,0.9,0.85],'InvertHardcopy','off','Visible','off'); % red [0.95,0.9,0.85], green [0.85,0.90,0.85], blue [0.89,0.94,0.99]
        subplot2n(4,1,1), plot(mF6g,'color','k','linewidth',1.5), hold on, plot(mF2g,'color',[0,0.55,0.69],'linewidth',0.2), ylabel('pcent sig change'),
            grid on,grid minor, xlim([1 length(mF2g)]),ylim([min(1.25.*[mF2g,mF6g]) 1.25.*max([mF2g,mF6g])]), title(['Gray matter mean signal AFTER "11-func-anat-registration" (slice ' num2str(i3) ' of ' num2str(siz3) ')'],'Color','k','FontSize',14), legend({'Gray matter signal before CSF-denoising'},'FontSize',12,'TextColor','k','Location','best')
        subplot2n(4,1,2), plot(mF2g,'color',[0,0.55,0.69],'linewidth',1.5), hold on, plot(mF6g,'color','k','linewidth',0.2), ylabel('pcent sig change'),
            grid on,grid minor, xlim([1 length(mF2g)]),ylim([min(1.25.*[mF2g,mF6g]) 1.25.*max([mF2g,mF6g])]), title(['Gray matter mean signal AFTER "12-denoise3-CSFregress" (slice ' num2str(i3) ' of ' num2str(siz3) ')'],'Color',[0,0.55,0.69],'FontSize',14), legend({'Gray matter signal after CSF-denoising'},'FontSize',12,'TextColor',[0,0.55,0.69],'Location','best')
        subplot2n(4,1,3), plot(mF6w,'color','k','linewidth',1.5), hold on, plot(mF2w,'color',[0.65,0.11,0.19],'linewidth',0.2), ylabel('pcent sig change'),
            grid on,grid minor, xlim([1 length(mF2g)]),ylim([min(1.25.*[mF2w,mF6w]) 1.25.*max([mF2w,mF6w])]), title(['White matter mean signal AFTER "11-func-anat-registration" (slice ' num2str(i3) ' of ' num2str(siz3) ')'],'Color','k','FontSize',14), legend({'White matter signal before CSF-denoising'},'FontSize',12,'TextColor','k','Location','best')
        subplot2n(4,1,4), plot(mF2w,'color',[0.65,0.11,0.19],'linewidth',1.5), hold on, plot(mF6w,'color','k','linewidth',0.2), ylabel('pcent sig change'),
            grid on,grid minor, xlim([1 length(mF2g)]),ylim([min(1.25.*[mF2w,mF6w]) 1.25.*max([mF2w,mF6w])]), title(['White matter mean signal AFTER "12-denoise3-CSFregress" (slice ' num2str(i3) ' of ' num2str(siz3) ')'],'Color',[0.65,0.11,0.19],'FontSize',14), legend({'White matter signal after CSF-denoising'},'FontSize',12,'TextColor',[0.65,0.11,0.19],'Location','best')
        saveas(fighndl,[QCpath 'global_signal_11coreg_vs_12denoise3_slice' sprintf('%.2d',i3) '.jpg'])
        close(fighndl)
        
        C3g = corrcoef(F6g'); C3g = uppertriangle(C3g); C3w = corrcoef(F6w'); C3w = uppertriangle(C3w);
        fighndl = figure('Position',[1 1 1152 720],'Color',[0.95,0.9,0.85],'InvertHardcopy','off','Visible','off'); % red [0.95,0.9,0.85], green [0.85,0.90,0.85], blue [0.89,0.94,0.99]
        subplot2n(4,1,1), histogram(C3g,'EdgeColor','none','FaceColor','k','FaceAlpha',0.5,'Normalization','probability'); grid on,grid minor, xlim([-1 1]), hold on, xline(0,'Color','k','linewidth',1.5); hold off
            title(sprintf('Histogram of voxel-to-voxel GRAY MATTER correlations AFTER 11-func-anat-registration (slice %d of %d)\n%.1f%% of the correlations were > 0  [ mean=%.2f, median=%.2f, SD=%.2f ]',i3,siz3,100*length(find(C3g>0))/length(C3g),mean(C3g),median(C3g),std(C3g)),'Color','k','FontSize',14)
        subplot2n(4,1,2), histogram(Cg,'EdgeColor','none','FaceColor',[0,0.55,0.69],'FaceAlpha',0.5,'Normalization','probability'); grid on,grid minor, xlim([-1 1]), hold on, xline(0,'Color','k','linewidth',1.5); hold off
            title(sprintf('GRAY MATTER: AFTER 12-denoise3-CSFregress (slice %d of %d): %.1f%% of the correlations were > 0  [ mean=%.2f, median=%.2f, SD=%.2f ]',i3,siz3,100*length(find(Cg>0))/length(Cg),mean(Cg),median(Cg),std(Cg)),'Color',[0,0.55,0.69],'FontSize',14)
        subplot2n(4,1,3), histogram(C3w,'EdgeColor','none','FaceColor','k','FaceAlpha',0.5,'Normalization','probability'); grid on,grid minor, xlim([-1 1]), hold on, xline(0,'Color','k','linewidth',1.5); hold off
            title(sprintf('WHITE MATTER: AFTER 11-func-anat-registration (slice %d of %d): %.1f%% of the correlations were > 0  [ mean=%.2f, median=%.2f, SD=%.2f ]',i3,siz3,100*length(find(C3w>0))/length(C3w),mean(C3w),median(C3w),std(C3w)),'Color','k','FontSize',14)
        subplot2n(4,1,4), histogram(Cw,'EdgeColor','none','FaceColor',[0.65,0.11,0.19],'FaceAlpha',0.5,'Normalization','probability'); grid on,grid minor, xlim([-1 1]), hold on, xline(0,'Color','k','linewidth',1.5); hold off
            title(sprintf('WHITE MATTER: AFTER 12-denoise3-CSFregress (slice %d of %d): %.1f%% of the correlations were > 0  [ mean=%.2f, median=%.2f, SD=%.2f ]',i3,siz3,100*length(find(Cw>0))/length(Cw),mean(Cw),median(Cw),std(Cw)),'Color',[0.65,0.11,0.19],'FontSize',14)
        saveas(fighndl,[QCpath 'hist_FC_compare_11coreg_12denoise3_slice' sprintf('%.2d',i3) '.jpg'])
        close(fighndl)
        F6g_=cat(1,F6g_,F6g); mF6g_=cat(1,mF6g_,mF6g); C3g_=cat(1,C3g_,C3g);
        F6w_=cat(1,F6w_,F6w); mF6w_=cat(1,mF6w_,mF6w); C3w_=cat(1,C3w_,C3w);
        clear F6g F6w mF6g mF6w C3g C3w
    end
    
    F2g_=cat(1,F2g_,F2g); mF2g_=cat(1,mF2g_,mF2g); F2w_=cat(1,F2w_,F2w); mF2w_=cat(1,mF2w_,mF2w); Cg_=cat(1,Cg_,Cg); Cw_=cat(1,Cw_,Cw); F4_=cat(1,F4_,F4); mF4_=cat(1,mF4_,mF4); C2_=cat(1,C2_,C2);
    clear F2g F2w F4 TR Smask1 svar1 svar2 mF2g mF2w mF4 Cg Cw C2
end

[~,svar1] = sort(var(F2g_,0,2)); [~,spsc1] = sort(meanPSC(F2g_));
[~,svar2] = sort(var(F2w_,0,2)); [~,spsc2] = sort(meanPSC(F2w_));
sortorder1b = sortorder1; sortorder1=[]; f1=0; %#ok<*ASGLU>
sortorder2b = sortorder2; sortorder2=[]; f2=0; %#ok<*ASGLU>
for im=1:length(sortorder1b)
    temp = sortorder1b{im}; temp = temp + f1; f1 = f1 + length(temp);
    sortorder1 = cat(1,sortorder1,temp); clear temp
    temp = sortorder2b{im}; temp = temp + f2; f2 = f2 + length(temp);
    sortorder2 = cat(1,sortorder2,temp); clear temp
end; clear im f1 f2
sortorder1 = spsc1; % options: spsc for percentage signal change, svar for variance, (comment out the line if you want to use distance from the center of the cord separately for each slice)
sortorder2 = spsc2; % options: spsc for percentage signal change, svar for variance, (comment out the line if you want to use distance from the center of the cord separately for each slice)
for im=max(sortorder1):-1:1, if isempty(find(sortorder1==im)), sortorder1(find(sortorder1>im)) = sortorder1(find(sortorder1>im))-1; end, end; clear im %#ok<*EFIND>
for im=max(sortorder2):-1:1, if isempty(find(sortorder2==im)), sortorder2(find(sortorder2>im)) = sortorder2(find(sortorder2>im))-1; end, end; clear im %#ok<*EFIND>

fighndl = figure('Position',[1 1 1152 720],'Color',[0.89,0.94,0.99],'InvertHardcopy','off','Visible','off'); % red [0.95,0.9,0.85], green [0.85,0.90,0.85], blue [0.89,0.94,0.99]
subplot2n(2,1,1), imagesc(F2g_(sortorder1,:)),colormap(cmap),colorbar, xlabel('time (in samples)'),ylabel(sprintf('voxels (sorted by mean percentage signal change;\nbottom row is highest)')),
  title(['Time series within the GRAY MATTER AFTER 12-DENOISE3-CSFregress (all slices combined)'],'Color',[0,0.55,0.69],'FontSize',14);
subplot2n(2,1,2), imagesc(F2w_(sortorder2,:)),colormap(cmap),colorbar, xlabel('time (in samples)'),ylabel(sprintf('voxels (sorted by mean percentage signal change;\nbottom row is highest)')),
  title(['Time series within the WHITE MATTER AFTER 12-DENOISE3-CSFregress (all slices combined)'],'Color',[0.65,0.11,0.19],'FontSize',14);
saveas(fighndl,[QCpath 'fMRI_ts_12denoise3_allSlices' '.jpg'])
 close(fighndl)
mF2g_=mean(mF2g_,1); mF2w_=mean(mF2w_,1); mF4_=mean(mF4_,1);
fighndl = figure('Position',[1 1 1152 720],'Color',[0.89,0.94,0.99],'InvertHardcopy','off','Visible','off'); % red [0.95,0.9,0.85], green [0.85,0.90,0.85], blue [0.89,0.94,0.99]
subplot2n(3,1,1), plot(mF4_,'color','k','linewidth',1.5), hold on, plot(mF2g_,'color',[0,0.55,0.69],'linewidth',0.2), plot(mF2w_,'color',[0.65,0.11,0.19],'linewidth',0.2), xlabel('time (in samples)'),ylabel('percentage signal change'),
    grid on,grid minor, xlim([1 length(mF2g_)]),ylim([1.25.*min([mF2g_,mF2w_,mF4_]) 1.25.*max([mF2g_,mF2w_,mF4_])]), title(['Global mean signal in UNPROCESSED RAW DATA (all slices averaged)'],'Color','k','FontSize',14), legend({'Global signal (raw data)'},'FontSize',12,'TextColor','k','Location','best')
subplot2n(3,1,2), plot(mF2g_,'color',[0,0.55,0.69],'linewidth',1.5), hold on, plot(mF2w_,'color',[0.65,0.11,0.19],'linewidth',0.2), plot(mF4_,'color','k','linewidth',0.2), xlabel('time (in samples)'),ylabel('percentage signal change'),
    grid on,grid minor, xlim([1 length(mF2g_)]),ylim([1.25.*min([mF2g_,mF2w_,mF4_]) 1.25.*max([mF2g_,mF2w_,mF4_])]), title(['Gray matter mean signal AFTER "12-DENOISE3-CSFregress" (all slices averaged)'],'Color',[0,0.55,0.69],'FontSize',14), legend({'Gray matter signal after 12-denoise3'},'FontSize',12,'TextColor',[0,0.55,0.69],'Location','best')
subplot2n(3,1,3), plot(mF2w_,'color',[0.65,0.11,0.19],'linewidth',1.5), hold on, plot(mF4_,'color','k','linewidth',0.2), plot(mF2g_,'color',[0,0.55,0.69],'linewidth',0.2), xlabel('time (in samples)'),ylabel('percentage signal change'),
    grid on,grid minor, xlim([1 length(mF2g_)]),ylim([1.25.*min([mF2g_,mF2w_,mF4_]) 1.25.*max([mF2g_,mF2w_,mF4_])]), title(['White matter mean signal AFTER "12-DENOISE3-CSFregress" (all slices averaged)'],'Color',[0.65,0.11,0.19],'FontSize',14), legend({'White matter signal after 12-denoise3'},'FontSize',12,'TextColor',[0.65,0.11,0.19],'Location','best')
saveas(fighndl,[QCpath 'global_signal_12denoise3_allSlices' '.jpg'])
 close(fighndl)
fighndl = figure('Position',[1 1 1152 720],'Color',[0.89,0.94,0.99],'InvertHardcopy','off','Visible','off'); % red [0.95,0.9,0.85], green [0.85,0.90,0.85], blue [0.89,0.94,0.99]
subplot2n(3,1,1), histogram(C2_,'EdgeColor','none','FaceColor','k','FaceAlpha',0.5,'Normalization','probability'); grid on,grid minor, xlim([-1 1]), hold on, xline(0,'Color','k','linewidth',1.5); hold off
    title(sprintf('Histogram of voxel-to-voxel correlations in UNPROCESSED RAW DATA (all slices combined)\n%.1f%% of the correlations were > 0  [ mean=%.2f, median=%.2f, SD=%.2f ]',100*length(find(C2_>0))/length(C2_),mean(C2_),median(C2_),std(C2_)),'Color','k','FontSize',14)
subplot2n(3,1,2), histogram(Cg_,'EdgeColor','none','FaceColor',[0,0.55,0.69],'FaceAlpha',0.5,'Normalization','probability'); grid on,grid minor, xlim([-1 1]), hold on, xline(0,'Color','k','linewidth',1.5); hold off
    title(sprintf('Histogram of voxel-to-voxel GRAY MATTER correlations AFTER 12-denoise3-CSFregress (all slices combined)\n%.1f%% of the correlations were > 0  [ mean=%.2f, median=%.2f, SD=%.2f ]',100*length(find(Cg_>0))/length(Cg_),mean(Cg_),median(Cg_),std(Cg_)),'Color',[0,0.55,0.69],'FontSize',14)
subplot2n(3,1,3), histogram(Cw_,'EdgeColor','none','FaceColor',[0.65,0.11,0.19],'FaceAlpha',0.5,'Normalization','probability'); grid on,grid minor, xlim([-1 1]), hold on, xline(0,'Color','k','linewidth',1.5); hold off
    title(sprintf('Histogram of voxel-to-voxel WHITE MATTER correlations AFTER 12-denoise3-CSFregress (all slices combined)\n%.1f%% of the correlations were > 0  [ mean=%.2f, median=%.2f, SD=%.2f ]',100*length(find(Cw_>0))/length(Cw_),mean(Cw_),median(Cw_),std(Cw_)),'Color',[0.65,0.11,0.19],'FontSize',14)
saveas(fighndl,[QCpath 'hist_FC_compare_12denoise3_allSlices' '.jpg'])
 close(fighndl)
if exist([base_dir_sub fname '_slice' num2str(i3) '_warped.nii'],'file') || exist([base_dir_sub fname '_slice' num2str(i3) '_warped.nii.gz'],'file') || exist([base_dir_sub fname '_warped.nii'],'file') || exist([base_dir_sub fname '_warped.nii.gz'],'file')
    mF6g_=mean(mF6g_,1); mF6w_=mean(mF6w_,1);
    fighndl = figure('Position',[1 1 1152 720],'Color',[0.89,0.94,0.99],'InvertHardcopy','off','Visible','off'); % red [0.95,0.9,0.85], green [0.85,0.90,0.85], blue [0.89,0.94,0.99]
	subplot2n(4,1,1), plot(mF6g_,'color','k','linewidth',1.5), hold on, plot(mF2g_,'color',[0,0.55,0.69],'linewidth',0.2), ylabel('pcent sig change'),
        grid on,grid minor, xlim([1 length(mF2g_)]),ylim([1.25.*min([mF2g_,mF6g_]) 1.25.*max([mF2g_,mF6g_])]), title(['Gray matter mean signal AFTER "11-func-anat-registration" (all slices averaged)'],'Color','k','FontSize',14), legend({'Gray matter signal before CSF-denoising'},'FontSize',12,'TextColor','k','Location','best')
    subplot2n(4,1,2), plot(mF2g_,'color',[0,0.55,0.69],'linewidth',1.5), hold on, plot(mF6g_,'color','k','linewidth',0.2), ylabel('pcent sig change'),
        grid on,grid minor, xlim([1 length(mF2g_)]),ylim([1.25.*min([mF2g_,mF6g_]) 1.25.*max([mF2g_,mF6g_])]), title(['Gray matter mean signal AFTER "12-DENOISE3-CSFregress" (all slices averaged)'],'Color',[0,0.55,0.69],'FontSize',14), legend({'Gray matter signal after CSF-denoising'},'FontSize',12,'TextColor',[0,0.55,0.69],'Location','best')
	subplot2n(4,1,3), plot(mF6w_,'color','k','linewidth',1.5), hold on, plot(mF2w_,'color',[0.65,0.11,0.19],'linewidth',0.2), ylabel('pcent sig change'),
        grid on,grid minor, xlim([1 length(mF2g_)]),ylim([1.25.*min([mF2w_,mF6w_]) 1.25.*max([mF2w_,mF6w_])]), title(['White matter mean signal AFTER "11-func-anat-registration" (all slices averaged)'],'Color','k','FontSize',14), legend({'White matter signal before CSF-denoising'},'FontSize',12,'TextColor','k','Location','best')
    subplot2n(4,1,4), plot(mF2w_,'color',[0.65,0.11,0.19],'linewidth',1.5), hold on, plot(mF6w_,'color','k','linewidth',0.2), ylabel('pcent sig change'),
        grid on,grid minor, xlim([1 length(mF2g_)]),ylim([1.25.*min([mF2w_,mF6w_]) 1.25.*max([mF2w_,mF6w_])]), title(['White matter mean signal AFTER "12-DENOISE3-CSFregress" (all slices averaged)'],'Color',[0.65,0.11,0.19],'FontSize',14), legend({'White matter signal after CSF-denoising'},'FontSize',12,'TextColor',[0.65,0.11,0.19],'Location','best')
    saveas(fighndl,[QCpath 'global_signal_11coreg_vs_12denoise3_allSlices' '.jpg'])
     close(fighndl)
     fighndl = figure('Position',[1 1 1152 720],'Color',[0.89,0.94,0.99],'InvertHardcopy','off','Visible','off'); % red [0.95,0.9,0.85], green [0.85,0.90,0.85], blue [0.89,0.94,0.99]
	subplot2n(4,1,1), histogram(C3g_,'EdgeColor','none','FaceColor','k','FaceAlpha',0.5,'Normalization','probability'); grid on,grid minor, xlim([-1 1]), hold on, xline(0,'Color','k','linewidth',1.5); hold off
       title(sprintf('Histogram of voxel-to-voxel GRAY MATTER correlations AFTER 11-registration (all slices combined)\n%.1f%% of the correlations were > 0  [ mean=%.2f, median=%.2f, SD=%.2f ]',100*length(find(C3g_>0))/length(C3g_),mean(C3g_),median(C3g_),std(C3g_)),'Color','k','FontSize',14)
    subplot2n(4,1,2), histogram(Cg_,'EdgeColor','none','FaceColor',[0,0.55,0.69],'FaceAlpha',0.5,'Normalization','probability'); grid on,grid minor, xlim([-1 1]), hold on, xline(0,'Color','k','linewidth',1.5); hold off
       title(sprintf('GRAY MATTER: AFTER 12-denoise3-CSFregress (all slices): %.1f%% of correlations > 0  [ mean=%.2f, median=%.2f, SD=%.2f ]',100*length(find(Cg_>0))/length(Cg_),mean(Cg_),median(Cg_),std(Cg_)),'Color',[0,0.55,0.69],'FontSize',14)
	subplot2n(4,1,3), histogram(C3w_,'EdgeColor','none','FaceColor','k','FaceAlpha',0.5,'Normalization','probability'); grid on,grid minor, xlim([-1 1]), hold on, xline(0,'Color','k','linewidth',1.5); hold off
       title(sprintf('WHITE MATTER: AFTER 11-registration (all slices): %.1f%% of the correlations were > 0  [ mean=%.2f, median=%.2f, SD=%.2f ]',100*length(find(C3w_>0))/length(C3w_),mean(C3w_),median(C3w_),std(C3w_)),'Color','k','FontSize',14)
    subplot2n(4,1,4), histogram(Cw_,'EdgeColor','none','FaceColor',[0.65,0.11,0.19],'FaceAlpha',0.5,'Normalization','probability'); grid on,grid minor, xlim([-1 1]), hold on, xline(0,'Color','k','linewidth',1.5); hold off
       title(sprintf('WHITE MATTER: AFTER 12-denoise3-CSFregress (all slices): %.1f%% of the correlations were > 0  [ mean=%.2f, median=%.2f, SD=%.2f ]',100*length(find(Cw_>0))/length(Cw_),mean(Cw_),median(Cw_),std(Cw_)),'Color',[0.65,0.11,0.19],'FontSize',14)
    saveas(fighndl,[QCpath 'hist_FC_compare_11coreg_12denoise3_allSlices' '.jpg'])
     close(fighndl)
end
clear svar1 spsc1 svar2 spsc2 sortorder1 sortorder2 F2g_ mF2g_ F2w_ mF2w_ Cg_ Cw_ F4_ mF4_ C2_ F6g_ F6w_ mF6g_ mF6w_ C3g_ C3w_

%%% Make QC video (needs audio video toolbox in Matlab)
if ~isempty(which('VideoWriter')) && ((save_videos==1)||(save_videos==2)) % check if Matlab's audio video toolbox exists

subplot = @(m,n,p) subtightplot(m, n, p, [0.001 0.005], [0.01 0.001], [0.01 0.001]);
scfMRItb_04_unzipFile(base_dir_sub, fname_anat, '_mask_SC')
try
    scfMRItb_04_unzipFile(base_dir_sub, fname_anat, '_mask_SC')
    mask_SC2 = load_untouch_nii([base_dir_sub fname_anat '_mask_SC.nii']); mask_SC2 = mask_SC2.img;
catch
    mask_SC = load_untouch_nii([base_dir_sub fname_anat '_mask_GM.nii']);
    mask_SC.img = or(mask_GM, mask_WM);
    save_untouch_nii(mask_SC, [base_dir_sub fname_anat '_mask_SC.nii']);
    mask_SC2 = mask_SC.img; clear mask_SC
end
for M=1:size(mask_SC2,3)
    mask_SC = rot90(mask_SC2(:,:,M));
    [xmask_SC,ymask_SC] = find(mask_SC==1);
    minx_SC(:,M) = round(min(xmask_SC)-0.1*(size(mask_SC,1))); maxx_SC(:,M)=round(max(xmask_SC)+0.1*(size(mask_SC,1)));
    miny_SC(:,M) = round(min(ymask_SC)-0.1*(size(mask_SC,2))); maxy_SC(:,M)=round(max(ymask_SC)+0.1*(size(mask_SC,2)));
    if (maxx_SC(:,M)-minx_SC(:,M))<(maxy_SC(:,M)-miny_SC(:,M))
        factr = ((maxy_SC(:,M)-miny_SC(:,M))-(maxx_SC(:,M)-minx_SC(:,M)))/2;
        if factr==round(factr)
            maxx_SC(:,M)=maxx_SC(:,M)+factr; minx_SC(:,M)=minx_SC(:,M)-factr;
        else
            maxx_SC(:,M)=maxx_SC(:,M)+factr+0.5; minx_SC(:,M)=minx_SC(:,M)-factr+0.5;
        end
    elseif (maxy_SC(:,M)-miny_SC(:,M))<(maxx_SC(:,M)-minx_SC(:,M))
        factr = ((maxx_SC(:,M)-minx_SC(:,M))-(maxy_SC(:,M)-miny_SC(:,M)))/2;
        if factr==round(factr)
            maxy_SC(:,M)=maxy_SC(:,M)+factr; miny_SC(:,M)=miny_SC(:,M)-factr;
        else
            maxy_SC(:,M)=maxy_SC(:,M)+factr+0.5; miny_SC(:,M)=miny_SC(:,M)-factr+0.5;
        end
    end
    clear mask_SC xmask_SC ymask_SC factr
end; clear M

for i3 = 1 : siz3
    try waitbar((i3/siz3),wbar3,sprintf('12. Generate QC video: slice (%d) of (%d)',i3,siz3)); catch, end
    fprintf('12. Generate QC video: slice (%d) of (%d), subject (%d) of (%d), run (%d) of (%d)\n',i3,siz3,k,size(func_files,1),k2,size(func_files,2))
    scfMRItb_04_resplitData(base_dir_sub, fname, ['_denoised' cutoff_str], siz3) % if slice-wise data doesn't exist but combined 4D data exists, then split the data to generate the needed slice-wise data
    F1 = load_untouch_nii([base_dir_sub fname '_slice' num2str(i3) '_denoised' cutoff_str '.nii']);
    TR = F1.hdr.dime.pixdim(5); F1 = rot90(single(squeeze(F1.img)));
    video = VideoWriter([QCpath 'fMRIvideo_12denoise3_slice' sprintf('%.2d',i3) '.mp4'],'MPEG-4'); %#ok<*TNMLP> % create the video object
    video.FrameRate = round(30/TR); % 30 seconds of data in every second of the video
    video.Quality = 90; % very limited compression so that the images are shown as is (and thus avoid misunderstandings due to video quality)
    open(video); % open the file for writing
    
    [sizeFx_SC,sizeFy_SC] = size(F1(:,:,1));
    F2 = F1(minx_SC:maxx_SC,miny_SC:maxy_SC,:);
    for i5=1:size(F1,3)
        F1(:,:,i5) = imresize(F2(:,:,i5),[sizeFx_SC sizeFy_SC]);
    end
    
    if (save_videos==1)
    scfMRItb_04_resplitData(base_dir_sub, fname, '_warped', siz3) % if slice-wise data doesn't exist but combined 4D data exists, then split the data to generate the needed slice-wise data
    F3 = load_untouch_nii([base_dir_sub fname '_slice' num2str(i3) '_warped.nii']); F3 = rot90(single(squeeze(F3.img)));
    [sizeFx,sizeFy] = size(F3(:,:,1));
    F4 = F3(minx_SC:maxx_SC,miny_SC:maxy_SC,:);
    for i5=1:size(F3,3)
        F3(:,:,i5) = imresize(F4(:,:,i5),[sizeFx sizeFy]);
    end
    clear sizeFx sizeFy F2 F4 i5 sizeFx_SC sizeFy_SC
    
    elseif (save_videos==2)
    if Smask_err==0
        scfMRItb_04_resplitData(base_dir_sub, fname, '_Smask', siz3) % if slice-wise data doesn't exist but combined 4D data exists, then split the data to generate the needed slice-wise data
        scfMRItb_04_unzipFile(base_dir_sub, fname, ['_slice' num2str(i3) '_Smask'])
        Smask1 = load_untouch_nii([base_dir_sub fname '_slice' num2str(i3) '_Smask.nii']); Smask1 = Smask1.img;
    else
        Smask1 = Smask1_(:,:,i3);
    end
    Smask1 = rot90(Smask1);
    [xmask,ymask] = find(Smask1==1);
    minx=round(min(xmask)-0.1*(size(Smask1,1))); maxx=round(max(xmask)+0.1*(size(Smask1,1)));
    miny=round(min(ymask)-0.1*(size(Smask1,2))); maxy=round(max(ymask)+0.1*(size(Smask1,2)));
    if (maxx-minx)<(maxy-miny)
        factr = ((maxy-miny)-(maxx-minx))/2;
        if factr==round(factr)
            maxx=maxx+factr; minx=minx-factr;
        else
            maxx=maxx+factr+0.5; minx=minx-factr+0.5;
        end
    elseif (maxy-miny)<(maxx-minx)
        factr = ((maxx-minx)-(maxy-miny))/2;
        if factr==round(factr)
            maxy=maxy+factr; miny=miny-factr;
        else
            maxy=maxy+factr+0.5; miny=miny-factr+0.5;
        end
    end
    scfMRItb_04_resplitData(base_dir_sub, fname, '', siz3) % if slice-wise data doesn't exist but combined 4D data exists, then split the data to generate the needed slice-wise data
    F3 = load_untouch_nii([base_dir_sub fname '_slice' num2str(i3) '.nii']); F3 = rot90(single(squeeze(F3.img)));
    [sizeFx,sizeFy] = size(F3(:,:,1));
    F4 = F3(minx:maxx,miny:maxy,:);
    for i5=1:size(F3,3)
        F3(:,:,i5) = imresize(F4(:,:,i5),[sizeFx sizeFy]);
    end
    clear maxx maxy minx miny xmask ymask Smask1 factr sizeFx sizeFy F2 F4 i5 sizeFx_SC sizeFy_SC        
    end
    
    for ii=1:size(F1,3) % loop across the number of images
        I = F1(:,:,ii); %read the next image
        I(find(isnan(I)))=0;
        I = (I - min(I(:))) ./ (max(I(:)) - min(I(:)));
        I2 = F3(:,:,ii); %read the next image
        I2(find(isnan(I2)))=0;
        I2 = (I2 - min(I2(:))) ./ (max(I2(:)) - min(I2(:)));
        fighndl = figure('Position',[round((scrsz(3)-scrsz(4))/2) 1 round(0.75*scrsz(3)) round(0.75*scrsz(4))],'Color',[0.95,0.9,0.85],'InvertHardcopy','off','Visible','off'); % [0.85,0.90,0.85]
        if (save_videos==1)
            subplot(1,2,1), imshow(I2, [0 max(I2(:))], 'InitialMagnification', 'fit'); title({['fMRI data after "11. func-anat-registration" (slice ' num2str(i3) ' of ' num2str(siz3) ')'];['Time point ' num2str(ii) ' of ' num2str(size(F1,3))]},'Color','k','FontSize',18);
        elseif (save_videos==2)
            subplot(1,2,1), imshow(I2, [0 max(I2(:))], 'InitialMagnification', 'fit'); title({['Unprocessed raw fMRI data (slice ' num2str(i3) ' of ' num2str(siz3) ')'];['Time point ' num2str(ii) ' of ' num2str(size(F1,3))]},'Color','k','FontSize',18);
        end
        subplot(1,2,2), imshow(I, [0 max(I(:))], 'InitialMagnification', 'fit'); title({['fMRI data after "12. denoise-3 (CSF regression)" (slice ' num2str(i3) ' of ' num2str(siz3) ')'];['Time point ' num2str(ii) ' of ' num2str(size(F1,3))]},'Color',[0,0.55,0.69],'FontSize',18);
        frame = getframe(gcf);
        writeVideo(video,frame); % write the image to file
        close(fighndl)
        clear I I2 frame
    end
    close(video); %close the file
    clear F1 F3 video ii TR
end; clear i3
clear maxx_SC maxy_SC minx_SC miny_SC F
end

end

%% scfMRItb_13_WMregress
if choices_preproc(13)==1
clear QCpath save_videos fighndl
if ~(exist([base_dir_sub 'QC' '/13_denoise4-WMregress'],'dir'))
    mkdir([base_dir_sub 'QC' '/13_denoise4-WMregress'])
end
if Rns==0
    QCpath = [base_dir_sub 'QC' '/13_denoise4-WMregress/'];
else
    if ~(exist([base_dir_sub 'QC' '/13_denoise4-WMregress' '/run' num2str(Rns)],'dir'))
        mkdir([base_dir_sub 'QC' '/13_denoise4-WMregress' '/run' num2str(Rns)])
    end
    QCpath = [base_dir_sub 'QC' '/13_denoise4-WMregress' '/run' num2str(Rns) '/'];
end

if (save_videos_==0)||(save_videos_==1)
    save_videos = save_videos_;
elseif (save_videos_==2)&&(sum([choices_preproc(14)])==0)
    save_videos = 2;
else
    save_videos = 0;
end
if save_videos==1
    save_videos_steps(7)=1;
end

if exist([base_dir_sub fname '_slice' num2str(i3) '_denoised' cutoff_str '.nii'],'file') || exist([base_dir_sub fname '_slice' num2str(i3) '_denoised' cutoff_str '.nii.gz'],'file') || exist([base_dir_sub fname '_denoised' cutoff_str '.nii'],'file') || exist([base_dir_sub fname '_denoised' cutoff_str '.nii.gz'],'file')
    stp=2;
elseif exist([base_dir_sub fname '_slice' num2str(i3) '_warped.nii'],'file') || exist([base_dir_sub fname '_slice' num2str(i3) '_warped.nii.gz'],'file') || exist([base_dir_sub fname '_warped.nii'],'file') || exist([base_dir_sub fname '_warped.nii.gz'],'file')
    stp=1;
end


F2g_=[]; mF2g_=[]; F2w_=[]; mF2w_=[]; Cg_=[]; Cw_=[]; F4_=[]; mF4_=[]; C2_=[];
if stp~=0, F6g_=[]; mF6g_=[]; F6w_=[]; mF6w_=[]; C3g_=[]; C3w_=[]; end
for i3 = 1 : siz3
    try waitbar((i3/siz3),wbar3,sprintf('13. QC plots: slice (%d) of (%d)',i3,siz3)); catch, end
    fprintf('13. QC plots: slice (%d) of (%d), subject (%d) of (%d), run (%d) of (%d)\n',i3,siz3,k,size(func_files,1),k2,size(func_files,2))
    if stp==2
        scfMRItb_04_resplitData(base_dir_sub, fname, ['_denoised' cutoff_str 'WM'], siz3) % if slice-wise data doesn't exist but combined 4D data exists, then split the data to generate the needed slice-wise data
        scfMRItb_04_unzipFile(base_dir_sub, fname, ['_slice' num2str(i3) '_denoised' cutoff_str 'WM'])
        F1 = load_untouch_nii([base_dir_sub fname '_slice' num2str(i3) '_denoised' cutoff_str 'WM.nii']);
    elseif stp==1
        scfMRItb_04_resplitData(base_dir_sub, fname, '_WM', siz3) % if slice-wise data doesn't exist but combined 4D data exists, then split the data to generate the needed slice-wise data
        scfMRItb_04_unzipFile(base_dir_sub, fname, ['_slice' num2str(i3) '_WM'])
        F1 = load_untouch_nii([base_dir_sub fname '_slice' num2str(i3) '_WM.nii']);
    end
    TR = F1.hdr.dime.pixdim(5); F1 = single(squeeze(F1.img));
    
    if Smask_err==0
        scfMRItb_04_resplitData(base_dir_sub, fname, '_Smask', siz3) % if slice-wise data doesn't exist but combined 4D data exists, then split the data to generate the needed slice-wise data
        scfMRItb_04_unzipFile(base_dir_sub, fname, ['_slice' num2str(i3) '_Smask'])
        Smask1 = load_untouch_nii([base_dir_sub fname '_slice' num2str(i3) '_Smask.nii']); Smask1 = Smask1.img;
    else
        Smask1 = Smask1_(:,:,i3);
    end

    for i5=1:size(F1,3)
        tmp = F1(:,:,i5);
        F2g(:,i5) = tmp(find(mask_GM(:,:,i3)==1));
        F2w(:,i5) = tmp(find(mask_WM(:,:,i3)==1));
        clear tmp
    end; clear i5 F1
    [~,svar1] = sort(var(F2g,0,2)); [~,sdist1] = sort(svar1); sortorder1{i3,1} = sdist1;
    [~,svar2] = sort(var(F2w,0,2)); [~,sdist2] = sort(svar2); sortorder2{i3,1} = sdist2;
    
    fighndl = figure('Position',[1 1 1152 720],'Color',[0.95,0.9,0.85],'InvertHardcopy','off','Visible','off'); % red [0.95,0.9,0.85], green [0.85,0.90,0.85], blue [0.89,0.94,0.99]
    subplot2n(2,1,1), imagesc(F2g(sortorder1{i3},:)),colormap(cmap),colorbar, xlabel('time (in samples)'),ylabel('Gray matter voxels (sorted by variance)'),
        title(['Time series within the GRAY MATTER after "13-DENOISE4-WMregress" in slice ' num2str(i3) ' of ' num2str(siz3)],'Color',[0,0.55,0.69],'FontSize',14);
    subplot2n(2,1,2), imagesc(F2w(sortorder2{i3},:)),colormap(cmap),colorbar, xlabel('time (in samples)'),ylabel('White matter voxels (sorted by variance)'),
        title(['Time series within the WHITE MATTER after "13-DENOISE4-WMregress" in slice ' num2str(i3) ' of ' num2str(siz3)],'Color',[0.65,0.11,0.19],'FontSize',14);
    saveas(fighndl,[QCpath 'fMRI_ts_13denoise4_slice' sprintf('%.2d',i3) '.jpg'])
    close(fighndl)
    
    scfMRItb_04_resplitData(base_dir_sub, fname, '', siz3) % if slice-wise data doesn't exist but combined 4D data exists, then split the data to generate the needed slice-wise data
	scfMRItb_04_unzipFile(base_dir_sub, fname, ['_slice' num2str(i3)])
    F3 = load_untouch_nii([base_dir_sub fname '_slice' num2str(i3) '.nii']); F3 = single(squeeze(F3.img));
    for i5=1:size(F3,3)
        tmp = F3(:,:,i5);
        F4(:,i5) = tmp(find(Smask1==1)); clear tmp
    end; clear i5 F3
    mF2g=mean(F2g,1); mF2g = 100*(mF2g-mean(mF2g))./mean(mF2g); mF2w=mean(F2w,1); mF2w = 100*(mF2w-mean(mF2w))./mean(mF2w);
    mF4=mean(F4,1); mF4 = 100*(mF4-mean(mF4))./mean(mF4);
    fighndl = figure('Position',[1 1 1152 720],'Color',[0.95,0.9,0.85],'InvertHardcopy','off','Visible','off'); % red [0.95,0.9,0.85], green [0.85,0.90,0.85], blue [0.89,0.94,0.99]
    subplot2n(3,1,1), plot(mF4,'color','k','linewidth',1.5), hold on, plot(mF2g,'color',[0,0.55,0.69],'linewidth',0.2), plot(mF2w,'color',[0.65,0.11,0.19],'linewidth',0.2), xlabel('time (in samples)'),ylabel('percentage signal change'),
      grid on,grid minor, xlim([1 length(mF2g)]),ylim([min(1.25.*[mF2g,mF2w,mF4]) 1.25.*max([mF2g,mF2w,mF4])]), title(['Global mean signal in UNPROCESSED RAW DATA (slice ' num2str(i3) ' of ' num2str(siz3) ')'],'Color','k','FontSize',14), legend({'Global signal (raw data)'},'FontSize',12,'TextColor','k','Location','best')
    subplot2n(3,1,2), plot(mF2g,'color',[0,0.55,0.69],'linewidth',1.5), hold on, plot(mF2w,'color',[0.65,0.11,0.19],'linewidth',0.2), plot(mF4,'color','k','linewidth',0.2), xlabel('time (in samples)'),ylabel('percentage signal change'),
      grid on,grid minor, xlim([1 length(mF2g)]),ylim([min(1.25.*[mF2g,mF2w,mF4]) 1.25.*max([mF2g,mF2w,mF4])]), title(['Gray matter mean signal AFTER "13-DENOISE4-WMregress" (slice ' num2str(i3) ' of ' num2str(siz3) ')'],'Color',[0,0.55,0.69],'FontSize',14), legend({'Gray matter signal after 13-denoise4'},'FontSize',12,'TextColor',[0,0.55,0.69],'Location','best')
    subplot2n(3,1,3), plot(mF2w,'color',[0.65,0.11,0.19],'linewidth',1.5), hold on, plot(mF4,'color','k','linewidth',0.2), plot(mF2g,'color',[0,0.55,0.69],'linewidth',0.2), xlabel('time (in samples)'),ylabel('percentage signal change'),
      grid on,grid minor, xlim([1 length(mF2g)]),ylim([min(1.25.*[mF2g,mF2w,mF4]) 1.25.*max([mF2g,mF2w,mF4])]), title(['White matter mean signal AFTER "13-DENOISE4-WMregress" (slice ' num2str(i3) ' of ' num2str(siz3) ')'],'Color',[0.65,0.11,0.19],'FontSize',14), legend({'White matter signal after 13-denoise4'},'FontSize',12,'TextColor',[0.65,0.11,0.19],'Location','best')
    saveas(fighndl,[QCpath 'global_signal_13denoise4_slice' sprintf('%.2d',i3) '.jpg'])
    close(fighndl)

    Cg = corrcoef(F2g'); Cg = uppertriangle(Cg);
    Cw = corrcoef(F2w'); Cw = uppertriangle(Cw);
    C2 = corrcoef(F4');  C2 = uppertriangle(C2);
    fighndl = figure('Position',[1 1 1152 720],'Color',[0.95,0.9,0.85],'InvertHardcopy','off','Visible','off'); % red [0.95,0.9,0.85], green [0.85,0.90,0.85], blue [0.89,0.94,0.99]
    subplot2n(3,1,1), histogram(C2,'EdgeColor','none','FaceColor','k','FaceAlpha',0.5,'Normalization','probability'); grid on,grid minor, xlim([-1 1]), hold on, xline(0,'Color','k','linewidth',1.5); hold off
      title(sprintf('Histogram of voxel-to-voxel correlations in UNPROCESSED RAW DATA (slice %d of %d)\n%.1f%% of the correlations were > 0  [ mean=%.2f, median=%.2f, SD=%.2f ]',i3,siz3,100*length(find(C2>0))/length(C2),mean(C2),median(C2),std(C2)),'Color','k','FontSize',14)
    subplot2n(3,1,2), histogram(Cg,'EdgeColor','none','FaceColor',[0,0.55,0.69],'FaceAlpha',0.5,'Normalization','probability'); grid on,grid minor, xlim([-1 1]), hold on, xline(0,'Color','k','linewidth',1.5); hold off
      title(sprintf('Histogram of voxel-to-voxel GRAY MATTER correlations AFTER 13-DENOISE4-WMregress (slice %d of %d)\n%.1f%% of the correlations were > 0  [ mean=%.2f, median=%.2f, SD=%.2f ]',i3,siz3,100*length(find(Cg>0))/length(Cg),mean(Cg),median(Cg),std(Cg)),'Color',[0,0.55,0.69],'FontSize',14)
    subplot2n(3,1,3), histogram(Cw,'EdgeColor','none','FaceColor',[0.65,0.11,0.19],'FaceAlpha',0.5,'Normalization','probability'); grid on,grid minor, xlim([-1 1]), hold on, xline(0,'Color','k','linewidth',1.5); hold off
      title(sprintf('Histogram of voxel-to-voxel WHITE MATTER correlations AFTER 13-DENOISE4-WMregress (slice %d of %d)\n%.1f%% of the correlations were > 0  [ mean=%.2f, median=%.2f, SD=%.2f ]',i3,siz3,100*length(find(Cw>0))/length(Cw),mean(Cw),median(Cw),std(Cw)),'Color',[0.65,0.11,0.19],'FontSize',14)
    saveas(fighndl,[QCpath 'hist_FC_compare_13denoise4_slice' sprintf('%.2d',i3) '.jpg'])
    close(fighndl)
    
    if stp~=0
        if stp==1
            scfMRItb_04_resplitData(base_dir_sub, fname, '_warped', siz3) % if slice-wise data doesn't exist but combined 4D data exists, then split the data to generate the needed slice-wise data
            scfMRItb_04_unzipFile(base_dir_sub, fname, ['_slice' num2str(i3) '_warped'])
            F5 = load_untouch_nii([base_dir_sub fname '_slice' num2str(i3) '_warped.nii']); F5 = single(squeeze(F5.img));
        elseif stp==2
            scfMRItb_04_resplitData(base_dir_sub, fname, ['_denoised' cutoff_str], siz3) % if slice-wise data doesn't exist but combined 4D data exists, then split the data to generate the needed slice-wise data
            scfMRItb_04_unzipFile(base_dir_sub, fname, ['_slice' num2str(i3) '_denoised' cutoff_str ''])
            F5 = load_untouch_nii([base_dir_sub fname '_slice' num2str(i3) '_denoised' cutoff_str '.nii']); F5 = single(squeeze(F5.img));
        end
        for i5=1:size(F5,3)
            tmp = F5(:,:,i5);
            F6g(:,i5) = tmp(find(mask_GM(:,:,i3)==1));
            F6w(:,i5) = tmp(find(mask_WM(:,:,i3)==1));
            clear tmp
        end; clear i5 F5
        mF6g=mean(F6g,1); mF6g = 100*(mF6g-mean(mF6g))./mean(mF6g); mF6w=mean(F6w,1); mF6w = 100*(mF6w-mean(mF6w))./mean(mF6w);
        fighndl = figure('Position',[1 1 1152 720],'Color',[0.95,0.9,0.85],'InvertHardcopy','off','Visible','off'); % red [0.95,0.9,0.85], green [0.85,0.90,0.85], blue [0.89,0.94,0.99]
        if stp==1
            subplot2n(4,1,1), plot(mF6g,'color','k','linewidth',1.5), hold on, plot(mF2g,'color',[0,0.55,0.69],'linewidth',0.2), ylabel('pcent sig change'),
                grid on,grid minor, xlim([1 length(mF2g)]),ylim([min(1.25.*[mF2g,mF6g]) 1.25.*max([mF2g,mF6g])]), title(['Gray matter mean signal AFTER "11-func-anat-registration" (slice ' num2str(i3) ' of ' num2str(siz3) ')'],'Color','k','FontSize',14), legend({'Gray matter signal before WM-denoising'},'FontSize',12,'TextColor','k','Location','best')
            subplot2n(4,1,3), plot(mF6w,'color','k','linewidth',1.5), hold on, plot(mF2w,'color',[0.65,0.11,0.19],'linewidth',0.2), ylabel('pcent sig change'),
                grid on,grid minor, xlim([1 length(mF2g)]),ylim([min(1.25.*[mF2w,mF6w]) 1.25.*max([mF2w,mF6w])]), title(['White matter mean signal AFTER "11-func-anat-registration" (slice ' num2str(i3) ' of ' num2str(siz3) ')'],'Color','k','FontSize',14), legend({'White matter signal before WM-denoising'},'FontSize',12,'TextColor','k','Location','best')
        elseif stp==2
            subplot2n(4,1,1), plot(mF6g,'color','k','linewidth',1.5), hold on, plot(mF2g,'color',[0,0.55,0.69],'linewidth',0.2), ylabel('pcent sig change'),
                grid on,grid minor, xlim([1 length(mF2g)]),ylim([min(1.25.*[mF2g,mF6g]) 1.25.*max([mF2g,mF6g])]), title(['Gray matter mean signal AFTER "12-denoise3-CSFregress" (slice ' num2str(i3) ' of ' num2str(siz3) ')'],'Color','k','FontSize',14), legend({'Gray matter signal before WM-denoising'},'FontSize',12,'TextColor','k','Location','best')
            subplot2n(4,1,3), plot(mF6w,'color','k','linewidth',1.5), hold on, plot(mF2w,'color',[0.65,0.11,0.19],'linewidth',0.2), ylabel('pcent sig change'),
                grid on,grid minor, xlim([1 length(mF2g)]),ylim([min(1.25.*[mF2w,mF6w]) 1.25.*max([mF2w,mF6w])]), title(['White matter mean signal AFTER "12-denoise3-CSFregress" (slice ' num2str(i3) ' of ' num2str(siz3) ')'],'Color','k','FontSize',14), legend({'White matter signal before WM-denoising'},'FontSize',12,'TextColor','k','Location','best')
        end
        subplot2n(4,1,2), plot(mF2g,'color',[0,0.55,0.69],'linewidth',1.5), hold on, plot(mF6g,'color','k','linewidth',0.2), ylabel('pcent sig change'),
            grid on,grid minor, xlim([1 length(mF2g)]),ylim([min(1.25.*[mF2g,mF6g]) 1.25.*max([mF2g,mF6g])]), title(['Gray matter mean signal AFTER "13-denoise4-WMregress" (slice ' num2str(i3) ' of ' num2str(siz3) ')'],'Color',[0,0.55,0.69],'FontSize',14), legend({'Gray matter signal after WM-denoising'},'FontSize',12,'TextColor',[0,0.55,0.69],'Location','best')
        subplot2n(4,1,4), plot(mF2w,'color',[0.65,0.11,0.19],'linewidth',1.5), hold on, plot(mF6w,'color','k','linewidth',0.2), ylabel('pcent sig change'),
            grid on,grid minor, xlim([1 length(mF2g)]),ylim([min(1.25.*[mF2w,mF6w]) 1.25.*max([mF2w,mF6w])]), title(['White matter mean signal AFTER "13-denoise4-WMregress" (slice ' num2str(i3) ' of ' num2str(siz3) ')'],'Color',[0.65,0.11,0.19],'FontSize',14), legend({'White matter signal after WM-denoising'},'FontSize',12,'TextColor',[0.65,0.11,0.19],'Location','best')
        saveas(fighndl,[QCpath 'global_signal_12denoise3_vs_13denoise4_slice' sprintf('%.2d',i3) '.jpg'])
        close(fighndl)
        
        C3g = corrcoef(F6g'); C3g = uppertriangle(C3g); C3w = corrcoef(F6w'); C3w = uppertriangle(C3w);
        fighndl = figure('Position',[1 1 1152 720],'Color',[0.95,0.9,0.85],'InvertHardcopy','off','Visible','off'); % red [0.95,0.9,0.85], green [0.85,0.90,0.85], blue [0.89,0.94,0.99]
        if stp==1
            subplot2n(4,1,1), histogram(C3g,'EdgeColor','none','FaceColor','k','FaceAlpha',0.5,'Normalization','probability'); grid on,grid minor, xlim([-1 1]), hold on, xline(0,'Color','k','linewidth',1.5); hold off
                title(sprintf('Histogram of voxel-to-voxel GRAY MATTER correlations AFTER 11-func-anat-registration (slice %d of %d)\n%.1f%% of the correlations were > 0  [ mean=%.2f, median=%.2f, SD=%.2f ]',i3,siz3,100*length(find(C3g>0))/length(C3g),mean(C3g),median(C3g),std(C3g)),'Color','k','FontSize',14)
            subplot2n(4,1,3), histogram(C3w,'EdgeColor','none','FaceColor','k','FaceAlpha',0.5,'Normalization','probability'); grid on,grid minor, xlim([-1 1]), hold on, xline(0,'Color','k','linewidth',1.5); hold off
                title(sprintf('WHITE MATTER: AFTER 11-func-anat-registration (slice %d of %d): %.1f%% of the correlations were > 0  [ mean=%.2f, median=%.2f, SD=%.2f ]',i3,siz3,100*length(find(C3w>0))/length(C3w),mean(C3w),median(C3w),std(C3w)),'Color','k','FontSize',14)
        elseif stp==2
            subplot2n(4,1,1), histogram(C3g,'EdgeColor','none','FaceColor','k','FaceAlpha',0.5,'Normalization','probability'); grid on,grid minor, xlim([-1 1]), hold on, xline(0,'Color','k','linewidth',1.5); hold off
                title(sprintf('Histogram of voxel-to-voxel GRAY MATTER correlations AFTER 12-denoise3-CSFregress (slice %d of %d)\n%.1f%% of the correlations were > 0  [ mean=%.2f, median=%.2f, SD=%.2f ]',i3,siz3,100*length(find(C3g>0))/length(C3g),mean(C3g),median(C3g),std(C3g)),'Color','k','FontSize',14)
            subplot2n(4,1,3), histogram(C3w,'EdgeColor','none','FaceColor','k','FaceAlpha',0.5,'Normalization','probability'); grid on,grid minor, xlim([-1 1]), hold on, xline(0,'Color','k','linewidth',1.5); hold off
                title(sprintf('WHITE MATTER: AFTER 12-denoise3-CSFregress (slice %d of %d): %.1f%% of the correlations were > 0  [ mean=%.2f, median=%.2f, SD=%.2f ]',i3,siz3,100*length(find(C3w>0))/length(C3w),mean(C3w),median(C3w),std(C3w)),'Color','k','FontSize',14)
        end
        subplot2n(4,1,2), histogram(Cg,'EdgeColor','none','FaceColor',[0,0.55,0.69],'FaceAlpha',0.5,'Normalization','probability'); grid on,grid minor, xlim([-1 1]), hold on, xline(0,'Color','k','linewidth',1.5); hold off
            title(sprintf('GRAY MATTER: AFTER 13-denoise4-WMregress (slice %d of %d): %.1f%% of the correlations were > 0  [ mean=%.2f, median=%.2f, SD=%.2f ]',i3,siz3,100*length(find(Cg>0))/length(Cg),mean(Cg),median(Cg),std(Cg)),'Color',[0,0.55,0.69],'FontSize',14)
        subplot2n(4,1,4), histogram(Cw,'EdgeColor','none','FaceColor',[0.65,0.11,0.19],'FaceAlpha',0.5,'Normalization','probability'); grid on,grid minor, xlim([-1 1]), hold on, xline(0,'Color','k','linewidth',1.5); hold off
            title(sprintf('WHITE MATTER: AFTER 13-denoise4-WMregress (slice %d of %d): %.1f%% of the correlations were > 0  [ mean=%.2f, median=%.2f, SD=%.2f ]',i3,siz3,100*length(find(Cw>0))/length(Cw),mean(Cw),median(Cw),std(Cw)),'Color',[0.65,0.11,0.19],'FontSize',14)
        saveas(fighndl,[QCpath 'hist_FC_compare_12denoise3_13denoise4_slice' sprintf('%.2d',i3) '.jpg'])
        close(fighndl)
        F6g_=cat(1,F6g_,F6g); mF6g_=cat(1,mF6g_,mF6g); C3g_=cat(1,C3g_,C3g);
        F6w_=cat(1,F6w_,F6w); mF6w_=cat(1,mF6w_,mF6w); C3w_=cat(1,C3w_,C3w);
        clear F6g F6w mF6g mF6w C3g C3w
    end
    
    F2g_=cat(1,F2g_,F2g); mF2g_=cat(1,mF2g_,mF2g); F2w_=cat(1,F2w_,F2w); mF2w_=cat(1,mF2w_,mF2w); Cg_=cat(1,Cg_,Cg); Cw_=cat(1,Cw_,Cw); F4_=cat(1,F4_,F4); mF4_=cat(1,mF4_,mF4); C2_=cat(1,C2_,C2);
    clear F2g F2w F4 TR Smask1 svar1 svar2 mF2g mF2w mF4 Cg Cw C2
end

[~,svar1] = sort(var(F2g_,0,2)); [~,spsc1] = sort(meanPSC(F2g_));
[~,svar2] = sort(var(F2w_,0,2)); [~,spsc2] = sort(meanPSC(F2w_));
sortorder1b = sortorder1; sortorder1=[]; f1=0; %#ok<*ASGLU>
sortorder2b = sortorder2; sortorder2=[]; f2=0; %#ok<*ASGLU>
for im=1:length(sortorder1b)
    temp = sortorder1b{im}; temp = temp + f1; f1 = f1 + length(temp);
    sortorder1 = cat(1,sortorder1,temp); clear temp
    temp = sortorder2b{im}; temp = temp + f2; f2 = f2 + length(temp);
    sortorder2 = cat(1,sortorder2,temp); clear temp
end; clear im f1 f2
sortorder1 = spsc1; % options: spsc for percentage signal change, svar for variance, (comment out the line if you want to use distance from the center of the cord separately for each slice)
sortorder2 = spsc2; % options: spsc for percentage signal change, svar for variance, (comment out the line if you want to use distance from the center of the cord separately for each slice)
for im=max(sortorder1):-1:1, if isempty(find(sortorder1==im)), sortorder1(find(sortorder1>im)) = sortorder1(find(sortorder1>im))-1; end, end; clear im %#ok<*EFIND>
for im=max(sortorder2):-1:1, if isempty(find(sortorder2==im)), sortorder2(find(sortorder2>im)) = sortorder2(find(sortorder2>im))-1; end, end; clear im %#ok<*EFIND>

fighndl = figure('Position',[1 1 1152 720],'Color',[0.89,0.94,0.99],'InvertHardcopy','off','Visible','off'); % red [0.95,0.9,0.85], green [0.85,0.90,0.85], blue [0.89,0.94,0.99]
subplot2n(2,1,1), imagesc(F2g_(sortorder1,:)),colormap(cmap),colorbar, xlabel('time (in samples)'),ylabel(sprintf('voxels (sorted by mean percentage signal change;\nbottom row is highest)')),
  title(['Time series within the GRAY MATTER AFTER 13-DENOISE4-WMregress (all slices combined)'],'Color',[0,0.55,0.69],'FontSize',14);
subplot2n(2,1,2), imagesc(F2w_(sortorder2,:)),colormap(cmap),colorbar, xlabel('time (in samples)'),ylabel(sprintf('voxels (sorted by mean percentage signal change;\nbottom row is highest)')),
  title(['Time series within the WHITE MATTER AFTER 13-DENOISE4-WMregress (all slices combined)'],'Color',[0.65,0.11,0.19],'FontSize',14);
saveas(fighndl,[QCpath 'fMRI_ts_13denoise4_allSlices' '.jpg'])
 close(fighndl)
mF2g_=mean(mF2g_,1); mF2w_=mean(mF2w_,1); mF4_=mean(mF4_,1);
fighndl = figure('Position',[1 1 1152 720],'Color',[0.89,0.94,0.99],'InvertHardcopy','off','Visible','off'); % red [0.95,0.9,0.85], green [0.85,0.90,0.85], blue [0.89,0.94,0.99]
subplot2n(3,1,1), plot(mF4_,'color','k','linewidth',1.5), hold on, plot(mF2g_,'color',[0,0.55,0.69],'linewidth',0.2), plot(mF2w_,'color',[0.65,0.11,0.19],'linewidth',0.2), xlabel('time (in samples)'),ylabel('percentage signal change'),
    grid on,grid minor, xlim([1 length(mF2g_)]),ylim([1.25.*min([mF2g_,mF2w_,mF4_]) 1.25.*max([mF2g_,mF2w_,mF4_])]), title(['Global mean signal in UNPROCESSED RAW DATA (all slices averaged)'],'Color','k','FontSize',14), legend({'Global signal (raw data)'},'FontSize',12,'TextColor','k','Location','best')
subplot2n(3,1,2), plot(mF2g_,'color',[0,0.55,0.69],'linewidth',1.5), hold on, plot(mF2w_,'color',[0.65,0.11,0.19],'linewidth',0.2), plot(mF4_,'color','k','linewidth',0.2), xlabel('time (in samples)'),ylabel('percentage signal change'),
    grid on,grid minor, xlim([1 length(mF2g_)]),ylim([1.25.*min([mF2g_,mF2w_,mF4_]) 1.25.*max([mF2g_,mF2w_,mF4_])]), title(['Gray matter mean signal AFTER "13-DENOISE4-WMregress" (all slices averaged)'],'Color',[0,0.55,0.69],'FontSize',14), legend({'Gray matter signal after 13-denoise4'},'FontSize',12,'TextColor',[0,0.55,0.69],'Location','best')
subplot2n(3,1,3), plot(mF2w_,'color',[0.65,0.11,0.19],'linewidth',1.5), hold on, plot(mF4_,'color','k','linewidth',0.2), plot(mF2g_,'color',[0,0.55,0.69],'linewidth',0.2), xlabel('time (in samples)'),ylabel('percentage signal change'),
    grid on,grid minor, xlim([1 length(mF2g_)]),ylim([1.25.*min([mF2g_,mF2w_,mF4_]) 1.25.*max([mF2g_,mF2w_,mF4_])]), title(['White matter mean signal AFTER "13-DENOISE4-WMregress" (all slices averaged)'],'Color',[0.65,0.11,0.19],'FontSize',14), legend({'White matter signal after 13-denoise4'},'FontSize',12,'TextColor',[0.65,0.11,0.19],'Location','best')
saveas(fighndl,[QCpath 'global_signal_13denoise4_allSlices' '.jpg'])
 close(fighndl)
fighndl = figure('Position',[1 1 1152 720],'Color',[0.89,0.94,0.99],'InvertHardcopy','off','Visible','off'); % red [0.95,0.9,0.85], green [0.85,0.90,0.85], blue [0.89,0.94,0.99]
subplot2n(3,1,1), histogram(C2_,'EdgeColor','none','FaceColor','k','FaceAlpha',0.5,'Normalization','probability'); grid on,grid minor, xlim([-1 1]), hold on, xline(0,'Color','k','linewidth',1.5); hold off
    title(sprintf('Histogram of voxel-to-voxel correlations in UNPROCESSED RAW DATA (all slices combined)\n%.1f%% of the correlations were > 0  [ mean=%.2f, median=%.2f, SD=%.2f ]',100*length(find(C2_>0))/length(C2_),mean(C2_),median(C2_),std(C2_)),'Color','k','FontSize',14)
subplot2n(3,1,2), histogram(Cg_,'EdgeColor','none','FaceColor',[0,0.55,0.69],'FaceAlpha',0.5,'Normalization','probability'); grid on,grid minor, xlim([-1 1]), hold on, xline(0,'Color','k','linewidth',1.5); hold off
    title(sprintf('Histogram of voxel-to-voxel GRAY MATTER correlations AFTER 13-denoise4-WMregress (all slices combined)\n%.1f%% of the correlations were > 0  [ mean=%.2f, median=%.2f, SD=%.2f ]',100*length(find(Cg_>0))/length(Cg_),mean(Cg_),median(Cg_),std(Cg_)),'Color',[0,0.55,0.69],'FontSize',14)
subplot2n(3,1,3), histogram(Cw_,'EdgeColor','none','FaceColor',[0.65,0.11,0.19],'FaceAlpha',0.5,'Normalization','probability'); grid on,grid minor, xlim([-1 1]), hold on, xline(0,'Color','k','linewidth',1.5); hold off
    title(sprintf('Histogram of voxel-to-voxel WHITE MATTER correlations AFTER 13-denoise4-WMregress (all slices combined)\n%.1f%% of the correlations were > 0  [ mean=%.2f, median=%.2f, SD=%.2f ]',100*length(find(Cw_>0))/length(Cw_),mean(Cw_),median(Cw_),std(Cw_)),'Color',[0.65,0.11,0.19],'FontSize',14)
saveas(fighndl,[QCpath 'hist_FC_compare_13denoise4_allSlices' '.jpg'])
 close(fighndl)

if stp~=0
    mF6g_=mean(mF6g_,1); mF6w_=mean(mF6w_,1);
    fighndl = figure('Position',[1 1 1152 720],'Color',[0.89,0.94,0.99],'InvertHardcopy','off','Visible','off'); % red [0.95,0.9,0.85], green [0.85,0.90,0.85], blue [0.89,0.94,0.99]
    if stp==1
    	subplot2n(4,1,1), plot(mF6g_,'color','k','linewidth',1.5), hold on, plot(mF2g_,'color',[0,0.55,0.69],'linewidth',0.2), ylabel('pcent sig change'),
            grid on,grid minor, xlim([1 length(mF2g_)]),ylim([1.25.*min([mF2g_,mF6g_]) 1.25.*max([mF2g_,mF6g_])]), title(['Gray matter mean signal AFTER "11-func-anat-registration" (all slices averaged)'],'Color','k','FontSize',14), legend({'Gray matter signal before WM-denoising'},'FontSize',12,'TextColor','k','Location','best')
        subplot2n(4,1,3), plot(mF6w_,'color','k','linewidth',1.5), hold on, plot(mF2w_,'color',[0.65,0.11,0.19],'linewidth',0.2), ylabel('pcent sig change'),
            grid on,grid minor, xlim([1 length(mF2g_)]),ylim([1.25.*min([mF2w_,mF6w_]) 1.25.*max([mF2w_,mF6w_])]), title(['White matter mean signal AFTER "11-func-anat-registration" (all slices averaged)'],'Color','k','FontSize',14), legend({'White matter signal before WM-denoising'},'FontSize',12,'TextColor','k','Location','best')
    elseif stp==2
    	subplot2n(4,1,1), plot(mF6g_,'color','k','linewidth',1.5), hold on, plot(mF2g_,'color',[0,0.55,0.69],'linewidth',0.2), ylabel('pcent sig change'),
            grid on,grid minor, xlim([1 length(mF2g_)]),ylim([1.25.*min([mF2g_,mF6g_]) 1.25.*max([mF2g_,mF6g_])]), title(['Gray matter mean signal AFTER "12-DENOISE3-CSFregress" (all slices averaged)'],'Color','k','FontSize',14), legend({'Gray matter signal before WM-denoising'},'FontSize',12,'TextColor','k','Location','best')
        subplot2n(4,1,3), plot(mF6w_,'color','k','linewidth',1.5), hold on, plot(mF2w_,'color',[0.65,0.11,0.19],'linewidth',0.2), ylabel('pcent sig change'),
            grid on,grid minor, xlim([1 length(mF2g_)]),ylim([1.25.*min([mF2w_,mF6w_]) 1.25.*max([mF2w_,mF6w_])]), title(['White matter mean signal AFTER "12-DENOISE3-CSFregress" (all slices averaged)'],'Color','k','FontSize',14), legend({'White matter signal before WM-denoising'},'FontSize',12,'TextColor','k','Location','best')
    end
    subplot2n(4,1,2), plot(mF2g_,'color',[0,0.55,0.69],'linewidth',1.5), hold on, plot(mF6g_,'color','k','linewidth',0.2), ylabel('pcent sig change'),
        grid on,grid minor, xlim([1 length(mF2g_)]),ylim([1.25.*min([mF2g_,mF6g_]) 1.25.*max([mF2g_,mF6g_])]), title(['Gray matter mean signal AFTER "13-DENOISE4-WMregress" (all slices averaged)'],'Color',[0,0.55,0.69],'FontSize',14), legend({'Gray matter signal after WM-denoising'},'FontSize',12,'TextColor',[0,0.55,0.69],'Location','best')
    subplot2n(4,1,4), plot(mF2w_,'color',[0.65,0.11,0.19],'linewidth',1.5), hold on, plot(mF6w_,'color','k','linewidth',0.2), ylabel('pcent sig change'),
        grid on,grid minor, xlim([1 length(mF2g_)]),ylim([1.25.*min([mF2w_,mF6w_]) 1.25.*max([mF2w_,mF6w_])]), title(['White matter mean signal AFTER "13-DENOISE4-WMregress" (all slices averaged)'],'Color',[0.65,0.11,0.19],'FontSize',14), legend({'White matter signal after WM-denoising'},'FontSize',12,'TextColor',[0.65,0.11,0.19],'Location','best')
    saveas(fighndl,[QCpath 'global_signal_12denoise3_vs_13denoise4_allSlices' '.jpg'])
     close(fighndl)
    
     fighndl = figure('Position',[1 1 1152 720],'Color',[0.89,0.94,0.99],'InvertHardcopy','off','Visible','off'); % red [0.95,0.9,0.85], green [0.85,0.90,0.85], blue [0.89,0.94,0.99]
    if stp==1
    	subplot2n(4,1,1), histogram(C3g_,'EdgeColor','none','FaceColor','k','FaceAlpha',0.5,'Normalization','probability'); grid on,grid minor, xlim([-1 1]), hold on, xline(0,'Color','k','linewidth',1.5); hold off
            title(sprintf('Histogram of voxel-to-voxel GRAY MATTER correlations AFTER 11-registration (all slices combined)\n%.1f%% of the correlations were > 0  [ mean=%.2f, median=%.2f, SD=%.2f ]',100*length(find(C3g_>0))/length(C3g_),mean(C3g_),median(C3g_),std(C3g_)),'Color','k','FontSize',14)
        subplot2n(4,1,3), histogram(C3w_,'EdgeColor','none','FaceColor','k','FaceAlpha',0.5,'Normalization','probability'); grid on,grid minor, xlim([-1 1]), hold on, xline(0,'Color','k','linewidth',1.5); hold off
            title(sprintf('WHITE MATTER: AFTER 11-registration (all slices): %.1f%% of the correlations were > 0  [ mean=%.2f, median=%.2f, SD=%.2f ]',100*length(find(C3w_>0))/length(C3w_),mean(C3w_),median(C3w_),std(C3w_)),'Color','k','FontSize',14)
    elseif stp==2
    	subplot2n(4,1,1), histogram(C3g_,'EdgeColor','none','FaceColor','k','FaceAlpha',0.5,'Normalization','probability'); grid on,grid minor, xlim([-1 1]), hold on, xline(0,'Color','k','linewidth',1.5); hold off
            title(sprintf('Histogram of voxel-to-voxel GRAY MATTER correlations AFTER 12-denoise3-CSFregress (all slices combined)\n%.1f%% of the correlations were > 0  [ mean=%.2f, median=%.2f, SD=%.2f ]',100*length(find(C3g_>0))/length(C3g_),mean(C3g_),median(C3g_),std(C3g_)),'Color','k','FontSize',14)
        subplot2n(4,1,3), histogram(C3w_,'EdgeColor','none','FaceColor','k','FaceAlpha',0.5,'Normalization','probability'); grid on,grid minor, xlim([-1 1]), hold on, xline(0,'Color','k','linewidth',1.5); hold off
            title(sprintf('WHITE MATTER: AFTER 12-denoise3-CSFregress (all slices): %.1f%% of the correlations were > 0  [ mean=%.2f, median=%.2f, SD=%.2f ]',100*length(find(C3w_>0))/length(C3w_),mean(C3w_),median(C3w_),std(C3w_)),'Color','k','FontSize',14)
    end
    subplot2n(4,1,2), histogram(Cg_,'EdgeColor','none','FaceColor',[0,0.55,0.69],'FaceAlpha',0.5,'Normalization','probability'); grid on,grid minor, xlim([-1 1]), hold on, xline(0,'Color','k','linewidth',1.5); hold off
       title(sprintf('GRAY MATTER: AFTER 13-denoise4-WMregress (all slices): %.1f%% of correlations > 0  [ mean=%.2f, median=%.2f, SD=%.2f ]',100*length(find(Cg_>0))/length(Cg_),mean(Cg_),median(Cg_),std(Cg_)),'Color',[0,0.55,0.69],'FontSize',14)
    subplot2n(4,1,4), histogram(Cw_,'EdgeColor','none','FaceColor',[0.65,0.11,0.19],'FaceAlpha',0.5,'Normalization','probability'); grid on,grid minor, xlim([-1 1]), hold on, xline(0,'Color','k','linewidth',1.5); hold off
       title(sprintf('WHITE MATTER: AFTER 13-denoise4-WMregress (all slices): %.1f%% of the correlations were > 0  [ mean=%.2f, median=%.2f, SD=%.2f ]',100*length(find(Cw_>0))/length(Cw_),mean(Cw_),median(Cw_),std(Cw_)),'Color',[0.65,0.11,0.19],'FontSize',14)
    saveas(fighndl,[QCpath 'hist_FC_compare_12denoise3_13denoise4_allSlices' '.jpg'])
     close(fighndl)
end
clear svar1 spsc1 svar2 spsc2 sortorder1 sortorder2 F2g_ mF2g_ F2w_ mF2w_ Cg_ Cw_ F4_ mF4_ C2_ F6g_ F6w_ mF6g_ mF6w_ C3g_ C3w_

%%% Make QC video (needs audio video toolbox in Matlab)
if ~isempty(which('VideoWriter')) && ((save_videos==1)||(save_videos==2)) % check if Matlab's audio video toolbox exists

subplot = @(m,n,p) subtightplot(m, n, p, [0.001 0.005], [0.01 0.001], [0.01 0.001]);
scfMRItb_04_unzipFile(base_dir_sub, fname_anat, '_mask_SC')
try
    scfMRItb_04_unzipFile(base_dir_sub, fname_anat, '_mask_SC')
    mask_SC2 = load_untouch_nii([base_dir_sub fname_anat '_mask_SC.nii']); mask_SC2 = mask_SC2.img;
catch
    mask_SC = load_untouch_nii([base_dir_sub fname_anat '_mask_GM.nii']);
    mask_SC.img = or(mask_GM, mask_WM);
    save_untouch_nii(mask_SC, [base_dir_sub fname_anat '_mask_SC.nii']);
    mask_SC2 = mask_SC.img; clear mask_SC
end
for M=1:size(mask_SC2,3)
    mask_SC = rot90(mask_SC2(:,:,M));
    [xmask_SC,ymask_SC] = find(mask_SC==1);
    minx_SC(:,M) = round(min(xmask_SC)-0.1*(size(mask_SC,1))); maxx_SC(:,M)=round(max(xmask_SC)+0.1*(size(mask_SC,1)));
    miny_SC(:,M) = round(min(ymask_SC)-0.1*(size(mask_SC,2))); maxy_SC(:,M)=round(max(ymask_SC)+0.1*(size(mask_SC,2)));
    if (maxx_SC(:,M)-minx_SC(:,M))<(maxy_SC(:,M)-miny_SC(:,M))
        factr = ((maxy_SC(:,M)-miny_SC(:,M))-(maxx_SC(:,M)-minx_SC(:,M)))/2;
        if factr==round(factr)
            maxx_SC(:,M)=maxx_SC(:,M)+factr; minx_SC(:,M)=minx_SC(:,M)-factr;
        else
            maxx_SC(:,M)=maxx_SC(:,M)+factr+0.5; minx_SC(:,M)=minx_SC(:,M)-factr+0.5;
        end
    elseif (maxy_SC(:,M)-miny_SC(:,M))<(maxx_SC(:,M)-minx_SC(:,M))
        factr = ((maxx_SC(:,M)-minx_SC(:,M))-(maxy_SC(:,M)-miny_SC(:,M)))/2;
        if factr==round(factr)
            maxy_SC(:,M)=maxy_SC(:,M)+factr; miny_SC(:,M)=miny_SC(:,M)-factr;
        else
            maxy_SC(:,M)=maxy_SC(:,M)+factr+0.5; miny_SC(:,M)=miny_SC(:,M)-factr+0.5;
        end
    end
    clear mask_SC xmask_SC ymask_SC factr
end; clear M

for i3 = 1 : siz3
    try waitbar((i3/siz3),wbar3,sprintf('13. Generate QC video: slice (%d) of (%d)',i3,siz3)); catch, end
    fprintf('13. Generate QC video: slice (%d) of (%d), subject (%d) of (%d), run (%d) of (%d)\n',i3,siz3,k,size(func_files,1),k2,size(func_files,2))
    if stp==2
        scfMRItb_04_resplitData(base_dir_sub, fname, ['_denoised' cutoff_str 'WM'], siz3) % if slice-wise data doesn't exist but combined 4D data exists, then split the data to generate the needed slice-wise data
        F1 = load_untouch_nii([base_dir_sub fname '_slice' num2str(i3) '_denoised' cutoff_str 'WM.nii']);
    elseif stp==1
        scfMRItb_04_resplitData(base_dir_sub, fname, '_WM', siz3) % if slice-wise data doesn't exist but combined 4D data exists, then split the data to generate the needed slice-wise data
        F1 = load_untouch_nii([base_dir_sub fname '_slice' num2str(i3) '_WM.nii']);
    end
    TR = F1.hdr.dime.pixdim(5); F1 = rot90(single(squeeze(F1.img)));
    video = VideoWriter([QCpath 'fMRIvideo_13denoise4_slice' sprintf('%.2d',i3) '.mp4'],'MPEG-4'); %#ok<*TNMLP> % create the video object
    video.FrameRate = round(30/TR); % 30 seconds of data in every second of the video
    video.Quality = 90; % very limited compression so that the images are shown as is (and thus avoid misunderstandings due to video quality)
    open(video); % open the file for writing
    
    [sizeFx_SC,sizeFy_SC] = size(F1(:,:,1));
    F2 = F1(minx_SC:maxx_SC,miny_SC:maxy_SC,:);
    for i5=1:size(F1,3)
        F1(:,:,i5) = imresize(F2(:,:,i5),[sizeFx_SC sizeFy_SC]);
    end
    
    if (save_videos==1)
    if stp==1
        scfMRItb_04_resplitData(base_dir_sub, fname, '_warped', siz3) % if slice-wise data doesn't exist but combined 4D data exists, then split the data to generate the needed slice-wise data
        F3 = load_untouch_nii([base_dir_sub fname '_slice' num2str(i3) '_warped.nii']); F3 = rot90(squeeze(F3.img));
    elseif stp==2
        scfMRItb_04_resplitData(base_dir_sub, fname, ['_denoised' cutoff_str], siz3) % if slice-wise data doesn't exist but combined 4D data exists, then split the data to generate the needed slice-wise data
        F3 = load_untouch_nii([base_dir_sub fname '_slice' num2str(i3) '_denoised' cutoff_str '.nii']); F3 = rot90(single(squeeze(F3.img)));
    end
    [sizeFx,sizeFy] = size(F3(:,:,1));
    F4 = F3(minx_SC:maxx_SC,miny_SC:maxy_SC,:);
    for i5=1:size(F3,3)
        F3(:,:,i5) = imresize(F4(:,:,i5),[sizeFx sizeFy]);
    end
    clear sizeFx sizeFy F2 F4 i5 sizeFx_SC sizeFy_SC
    
    elseif (save_videos==2)
    if Smask_err==0
        scfMRItb_04_resplitData(base_dir_sub, fname, '_Smask', siz3) % if slice-wise data doesn't exist but combined 4D data exists, then split the data to generate the needed slice-wise data
        scfMRItb_04_unzipFile(base_dir_sub, fname, ['_slice' num2str(i3) '_Smask'])
        Smask1 = load_untouch_nii([base_dir_sub fname '_slice' num2str(i3) '_Smask.nii']); Smask1 = Smask1.img;
    else
        Smask1 = Smask1_(:,:,i3);
    end
    Smask1 = rot90(Smask1);
    [xmask,ymask] = find(Smask1==1);
    minx=round(min(xmask)-0.1*(size(Smask1,1))); maxx=round(max(xmask)+0.1*(size(Smask1,1)));
    miny=round(min(ymask)-0.1*(size(Smask1,2))); maxy=round(max(ymask)+0.1*(size(Smask1,2)));
    if (maxx-minx)<(maxy-miny)
        factr = ((maxy-miny)-(maxx-minx))/2;
        if factr==round(factr)
            maxx=maxx+factr; minx=minx-factr;
        else
            maxx=maxx+factr+0.5; minx=minx-factr+0.5;
        end
    elseif (maxy-miny)<(maxx-minx)
        factr = ((maxx-minx)-(maxy-miny))/2;
        if factr==round(factr)
            maxy=maxy+factr; miny=miny-factr;
        else
            maxy=maxy+factr+0.5; miny=miny-factr+0.5;
        end
    end
    scfMRItb_04_resplitData(base_dir_sub, fname, '', siz3) % if slice-wise data doesn't exist but combined 4D data exists, then split the data to generate the needed slice-wise data
    F3 = load_untouch_nii([base_dir_sub fname '_slice' num2str(i3) '.nii']); F3 = rot90(single(squeeze(F3.img)));
    [sizeFx,sizeFy] = size(F3(:,:,1));
    F4 = F3(minx:maxx,miny:maxy,:);
    for i5=1:size(F3,3)
        F3(:,:,i5) = imresize(F4(:,:,i5),[sizeFx sizeFy]);
    end
    clear maxx maxy minx miny xmask ymask Smask1 factr sizeFx sizeFy F2 F4 i5 sizeFx_SC sizeFy_SC
    end
    
    for ii=1:size(F1,3) % loop across the number of images
        I = F1(:,:,ii); %read the next image
        I(find(isnan(I)))=0;
        I = (I - min(I(:))) ./ (max(I(:)) - min(I(:)));
        I2 = F3(:,:,ii); %read the next image
        I2(find(isnan(I2)))=0;
        I2 = (I2 - min(I2(:))) ./ (max(I2(:)) - min(I2(:)));
        fighndl = figure('Position',[round((scrsz(3)-scrsz(4))/2) 1 round(0.75*scrsz(3)) round(0.75*scrsz(4))],'Color',[0.95,0.9,0.85],'InvertHardcopy','off','Visible','off'); % [0.85,0.90,0.85]
        if (save_videos==1)
        if stp==1
            subplot(1,2,1), imshow(I2, [0 max(I2(:))], 'InitialMagnification', 'fit'); title({['fMRI data after "11. func-anat-registration" (slice ' num2str(i3) ' of ' num2str(siz3) ')'];['Time point ' num2str(ii) ' of ' num2str(size(F1,3))]},'Color','k','FontSize',18);
        elseif stp==2
            subplot(1,2,1), imshow(I2, [0 max(I2(:))], 'InitialMagnification', 'fit'); title({['fMRI data after "12. denoise-3 (CSF regression)" (slice ' num2str(i3) ' of ' num2str(siz3) ')'];['Time point ' num2str(ii) ' of ' num2str(size(F1,3))]},'Color','k','FontSize',18);
        end
        elseif (save_videos==2)
            subplot(1,2,1), imshow(I2, [0 max(I2(:))], 'InitialMagnification', 'fit'); title({['Unprocessed raw fMRI data (slice ' num2str(i3) ' of ' num2str(siz3) ')'];['Time point ' num2str(ii) ' of ' num2str(size(F1,3))]},'Color','k','FontSize',18);
        end
        subplot(1,2,2), imshow(I, [0 max(I(:))], 'InitialMagnification', 'fit'); title({['fMRI data after "13. denoise-4 (WM regression)" (slice ' num2str(i3) ' of ' num2str(siz3) ')'];['Time point ' num2str(ii) ' of ' num2str(size(F1,3))]},'Color',[0,0.55,0.69],'FontSize',18);
        frame = getframe(gcf);
        writeVideo(video,frame); % write the image to file
        close(fighndl)
        clear I I2 frame
    end
    close(video); %close the file
    clear F1 F3 video ii TR
end; clear i3
clear maxx_SC maxy_SC minx_SC miny_SC F
end

end

%% scfMRItb_14_COVregress
if choices_preproc(14)==1
clear QCpath save_videos fighndl stp
if ~(exist([base_dir_sub 'QC' '/14_denoise5-COVregress'],'dir'))
    mkdir([base_dir_sub 'QC' '/14_denoise5-COVregress'])
end
if Rns==0
    QCpath = [base_dir_sub 'QC' '/14_denoise5-COVregress/'];
else
    if ~(exist([base_dir_sub 'QC' '/14_denoise5-COVregress' '/run' num2str(Rns)],'dir'))
        mkdir([base_dir_sub 'QC' '/14_denoise5-COVregress' '/run' num2str(Rns)])
    end
    QCpath = [base_dir_sub 'QC' '/14_denoise5-COVregress' '/run' num2str(Rns) '/'];
end
save_videos_steps(8)=1;

if exist([base_dir_sub fname '_slice' num2str(i3) '_denoised' cutoff_str 'WM.nii'],'file') || exist([base_dir_sub fname '_slice' num2str(i3) '_denoised' cutoff_str 'WM.nii.gz'],'file') || exist([base_dir_sub fname '_denoised' cutoff_str 'WM.nii'],'file') || exist([base_dir_sub fname '_denoised' cutoff_str 'WM.nii.gz'],'file')
    stp=4;
elseif exist([base_dir_sub fname '_slice' num2str(i3) '_denoised' cutoff_str '.nii'],'file') || exist([base_dir_sub fname '_slice' num2str(i3) '_denoised' cutoff_str '.nii.gz'],'file') || exist([base_dir_sub fname '_denoised' cutoff_str '.nii'],'file') || exist([base_dir_sub fname '_denoised' cutoff_str '.nii.gz'],'file')
    stp=3;
elseif exist([base_dir_sub fname '_slice' num2str(i3) '_WM.nii'],'file') || exist([base_dir_sub fname '_slice' num2str(i3) '_WM.nii.gz'],'file') || exist([base_dir_sub fname '_WM.nii'],'file') || exist([base_dir_sub fname '_WM.nii.gz'],'file')
    stp=2;
elseif exist([base_dir_sub fname '_slice' num2str(i3) '_warped.nii'],'file') || exist([base_dir_sub fname '_slice' num2str(i3) '_warped.nii.gz'],'file') || exist([base_dir_sub fname '_warped.nii'],'file') || exist([base_dir_sub fname '_warped.nii.gz'],'file')
    stp=1;
end


F2g_=[]; mF2g_=[]; F2w_=[]; mF2w_=[]; Cg_=[]; Cw_=[]; F4_=[]; mF4_=[]; C2_=[];
if stp~=0, F6g_=[]; mF6g_=[]; F6w_=[]; mF6w_=[]; C3g_=[]; C3w_=[]; end
for i3 = 1 : siz3
    try waitbar((i3/siz3),wbar3,sprintf('14. QC plots: slice (%d) of (%d)',i3,siz3)); catch, end
    fprintf('14. QC plots: slice (%d) of (%d), subject (%d) of (%d), run (%d) of (%d)\n',i3,siz3,k,size(func_files,1),k2,size(func_files,2))
    if stp==4
        scfMRItb_04_resplitData(base_dir_sub, fname, ['_denoised' cutoff_str 'WMcov'], siz3) % if slice-wise data doesn't exist but combined 4D data exists, then split the data to generate the needed slice-wise data
        scfMRItb_04_unzipFile(base_dir_sub, fname, ['_slice' num2str(i3) '_denoised' cutoff_str 'WMcov'])
        F1 = load_untouch_nii([base_dir_sub fname '_slice' num2str(i3) '_denoised' cutoff_str 'WMcov.nii']);
    elseif stp==3
        scfMRItb_04_resplitData(base_dir_sub, fname, ['_denoised' cutoff_str 'cov'], siz3) % if slice-wise data doesn't exist but combined 4D data exists, then split the data to generate the needed slice-wise data
        scfMRItb_04_unzipFile(base_dir_sub, fname, ['_slice' num2str(i3) '_denoised' cutoff_str 'cov'])
        F1 = load_untouch_nii([base_dir_sub fname '_slice' num2str(i3) '_denoised' cutoff_str 'cov.nii']);
    elseif stp==2
        scfMRItb_04_resplitData(base_dir_sub, fname, '_WMcov', siz3) % if slice-wise data doesn't exist but combined 4D data exists, then split the data to generate the needed slice-wise data
        scfMRItb_04_unzipFile(base_dir_sub, fname, ['_slice' num2str(i3) '_WMcov'])
        F1 = load_untouch_nii([base_dir_sub fname '_slice' num2str(i3) '_WMcov.nii']);
    elseif stp==1
        scfMRItb_04_resplitData(base_dir_sub, fname, '_cov', siz3) % if slice-wise data doesn't exist but combined 4D data exists, then split the data to generate the needed slice-wise data
        scfMRItb_04_unzipFile(base_dir_sub, fname, ['_slice' num2str(i3) '_cov'])
        F1 = load_untouch_nii([base_dir_sub fname '_slice' num2str(i3) '_cov.nii']);
    end
    TR = F1.hdr.dime.pixdim(5); F1 = single(squeeze(F1.img));
    
    if Smask_err==0
        scfMRItb_04_resplitData(base_dir_sub, fname, '_Smask', siz3) % if slice-wise data doesn't exist but combined 4D data exists, then split the data to generate the needed slice-wise data
        scfMRItb_04_unzipFile(base_dir_sub, fname, ['_slice' num2str(i3) '_Smask'])
        Smask1 = load_untouch_nii([base_dir_sub fname '_slice' num2str(i3) '_Smask.nii']); Smask1 = Smask1.img;
    else
        Smask1 = Smask1_(:,:,i3);
    end

    for i5=1:size(F1,3)
        tmp = F1(:,:,i5);
        F2g(:,i5) = tmp(find(mask_GM(:,:,i3)==1));
        F2w(:,i5) = tmp(find(mask_WM(:,:,i3)==1));
        clear tmp
    end; clear i5 F1
    [~,svar1] = sort(var(F2g,0,2)); [~,sdist1] = sort(svar1); sortorder1{i3,1} = sdist1;
    [~,svar2] = sort(var(F2w,0,2)); [~,sdist2] = sort(svar2); sortorder2{i3,1} = sdist2;
    
    fighndl = figure('Position',[1 1 1152 720],'Color',[0.95,0.9,0.85],'InvertHardcopy','off','Visible','off'); % red [0.95,0.9,0.85], green [0.85,0.90,0.85], blue [0.89,0.94,0.99]
    subplot2n(2,1,1), imagesc(F2g(sortorder1{i3},:)),colormap(cmap),colorbar, xlabel('time (in samples)'),ylabel('Gray matter voxels (sorted by variance)'),
        title(['Time series within the GRAY MATTER after "14-DENOISE5-COVregress" in slice ' num2str(i3) ' of ' num2str(siz3)],'Color',[0,0.55,0.69],'FontSize',14);
    subplot2n(2,1,2), imagesc(F2w(sortorder2{i3},:)),colormap(cmap),colorbar, xlabel('time (in samples)'),ylabel('White matter voxels (sorted by variance)'),
        title(['Time series within the WHITE MATTER after "14-DENOISE5-COVregress" in slice ' num2str(i3) ' of ' num2str(siz3)],'Color',[0.65,0.11,0.19],'FontSize',14);
    saveas(fighndl,[QCpath 'fMRI_ts_14denoise5_slice' sprintf('%.2d',i3) '.jpg'])
    close(fighndl)
    
    scfMRItb_04_resplitData(base_dir_sub, fname, '', siz3) % if slice-wise data doesn't exist but combined 4D data exists, then split the data to generate the needed slice-wise data
	scfMRItb_04_unzipFile(base_dir_sub, fname, ['_slice' num2str(i3)])
    F3 = load_untouch_nii([base_dir_sub fname '_slice' num2str(i3) '.nii']); F3 = single(squeeze(F3.img));
    for i5=1:size(F3,3)
        tmp = F3(:,:,i5);
        F4(:,i5) = tmp(find(Smask1==1)); clear tmp
    end; clear i5 F3
    mF2g=mean(F2g,1); mF2g = 100*(mF2g-mean(mF2g))./mean(mF2g); mF2w=mean(F2w,1); mF2w = 100*(mF2w-mean(mF2w))./mean(mF2w);
    mF4=mean(F4,1); mF4 = 100*(mF4-mean(mF4))./mean(mF4);
    fighndl = figure('Position',[1 1 1152 720],'Color',[0.95,0.9,0.85],'InvertHardcopy','off','Visible','off'); % red [0.95,0.9,0.85], green [0.85,0.90,0.85], blue [0.89,0.94,0.99]
    subplot2n(3,1,1), plot(mF4,'color','k','linewidth',1.5), hold on, plot(mF2g,'color',[0,0.55,0.69],'linewidth',0.2), plot(mF2w,'color',[0.65,0.11,0.19],'linewidth',0.2), xlabel('time (in samples)'),ylabel('percentage signal change'),
      grid on,grid minor, xlim([1 length(mF2g)]),ylim([min(1.25.*[mF2g,mF2w,mF4]) 1.25.*max([mF2g,mF2w,mF4])]), title(['Global mean signal in UNPROCESSED RAW DATA (slice ' num2str(i3) ' of ' num2str(siz3) ')'],'Color','k','FontSize',14), legend({'Global signal (raw data)'},'FontSize',12,'TextColor','k','Location','best')
    subplot2n(3,1,2), plot(mF2g,'color',[0,0.55,0.69],'linewidth',1.5), hold on, plot(mF2w,'color',[0.65,0.11,0.19],'linewidth',0.2), plot(mF4,'color','k','linewidth',0.2), xlabel('time (in samples)'),ylabel('percentage signal change'),
      grid on,grid minor, xlim([1 length(mF2g)]),ylim([min(1.25.*[mF2g,mF2w,mF4]) 1.25.*max([mF2g,mF2w,mF4])]), title(['Gray matter mean signal AFTER "14-DENOISE5-COVregress" (slice ' num2str(i3) ' of ' num2str(siz3) ')'],'Color',[0,0.55,0.69],'FontSize',14), legend({'Gray matter signal after 14-denoise5'},'FontSize',12,'TextColor',[0,0.55,0.69],'Location','best')
    subplot2n(3,1,3), plot(mF2w,'color',[0.65,0.11,0.19],'linewidth',1.5), hold on, plot(mF4,'color','k','linewidth',0.2), plot(mF2g,'color',[0,0.55,0.69],'linewidth',0.2), xlabel('time (in samples)'),ylabel('percentage signal change'),
      grid on,grid minor, xlim([1 length(mF2g)]),ylim([min(1.25.*[mF2g,mF2w,mF4]) 1.25.*max([mF2g,mF2w,mF4])]), title(['White matter mean signal AFTER "14-DENOISE5-COVregress" (slice ' num2str(i3) ' of ' num2str(siz3) ')'],'Color',[0.65,0.11,0.19],'FontSize',14), legend({'White matter signal after 14-denoise5'},'FontSize',12,'TextColor',[0.65,0.11,0.19],'Location','best')
    saveas(fighndl,[QCpath 'global_signal_14denoise5_slice' sprintf('%.2d',i3) '.jpg'])
    close(fighndl)

    Cg = corrcoef(F2g'); Cg = uppertriangle(Cg);
    Cw = corrcoef(F2w'); Cw = uppertriangle(Cw);
    C2 = corrcoef(F4');  C2 = uppertriangle(C2);
    fighndl = figure('Position',[1 1 1152 720],'Color',[0.95,0.9,0.85],'InvertHardcopy','off','Visible','off'); % red [0.95,0.9,0.85], green [0.85,0.90,0.85], blue [0.89,0.94,0.99]
    subplot2n(3,1,1), histogram(C2,'EdgeColor','none','FaceColor','k','FaceAlpha',0.5,'Normalization','probability'); grid on,grid minor, xlim([-1 1]), hold on, xline(0,'Color','k','linewidth',1.5); hold off
      title(sprintf('Histogram of voxel-to-voxel correlations in UNPROCESSED RAW DATA (slice %d of %d)\n%.1f%% of the correlations were > 0  [ mean=%.2f, median=%.2f, SD=%.2f ]',i3,siz3,100*length(find(C2>0))/length(C2),mean(C2),median(C2),std(C2)),'Color','k','FontSize',14)
    subplot2n(3,1,2), histogram(Cg,'EdgeColor','none','FaceColor',[0,0.55,0.69],'FaceAlpha',0.5,'Normalization','probability'); grid on,grid minor, xlim([-1 1]), hold on, xline(0,'Color','k','linewidth',1.5); hold off
      title(sprintf('Histogram of voxel-to-voxel GRAY MATTER correlations AFTER 14-DENOISE5-COVregress (slice %d of %d)\n%.1f%% of the correlations were > 0  [ mean=%.2f, median=%.2f, SD=%.2f ]',i3,siz3,100*length(find(Cg>0))/length(Cg),mean(Cg),median(Cg),std(Cg)),'Color',[0,0.55,0.69],'FontSize',14)
    subplot2n(3,1,3), histogram(Cw,'EdgeColor','none','FaceColor',[0.65,0.11,0.19],'FaceAlpha',0.5,'Normalization','probability'); grid on,grid minor, xlim([-1 1]), hold on, xline(0,'Color','k','linewidth',1.5); hold off
      title(sprintf('Histogram of voxel-to-voxel WHITE MATTER correlations AFTER 14-DENOISE5-COVregress (slice %d of %d)\n%.1f%% of the correlations were > 0  [ mean=%.2f, median=%.2f, SD=%.2f ]',i3,siz3,100*length(find(Cw>0))/length(Cw),mean(Cw),median(Cw),std(Cw)),'Color',[0.65,0.11,0.19],'FontSize',14)
    saveas(fighndl,[QCpath 'hist_FC_compare_14denoise5_slice' sprintf('%.2d',i3) '.jpg'])
    close(fighndl)
    
    if stp~=0
        if stp==1
            scfMRItb_04_resplitData(base_dir_sub, fname, '_warped', siz3) % if slice-wise data doesn't exist but combined 4D data exists, then split the data to generate the needed slice-wise data
            scfMRItb_04_unzipFile(base_dir_sub, fname, ['_slice' num2str(i3) '_warped'])
            F5 = load_untouch_nii([base_dir_sub fname '_slice' num2str(i3) '_warped.nii']); F5 = single(squeeze(F5.img));
        elseif stp==2
            scfMRItb_04_resplitData(base_dir_sub, fname, '_WM', siz3) % if slice-wise data doesn't exist but combined 4D data exists, then split the data to generate the needed slice-wise data
            scfMRItb_04_unzipFile(base_dir_sub, fname, ['_slice' num2str(i3) '_WM'])
            F5 = load_untouch_nii([base_dir_sub fname '_slice' num2str(i3) '_WM.nii']); F5 = single(squeeze(F5.img));
        elseif stp==3
            scfMRItb_04_resplitData(base_dir_sub, fname, ['_denoised' cutoff_str], siz3) % if slice-wise data doesn't exist but combined 4D data exists, then split the data to generate the needed slice-wise data
            scfMRItb_04_unzipFile(base_dir_sub, fname, ['_slice' num2str(i3) '_denoised' cutoff_str ''])
            F5 = load_untouch_nii([base_dir_sub fname '_slice' num2str(i3) '_denoised' cutoff_str '.nii']); F5 = single(squeeze(F5.img));
        elseif stp==4
            scfMRItb_04_resplitData(base_dir_sub, fname, ['_denoised' cutoff_str 'WM'], siz3) % if slice-wise data doesn't exist but combined 4D data exists, then split the data to generate the needed slice-wise data
            scfMRItb_04_unzipFile(base_dir_sub, fname, ['_slice' num2str(i3) '_denoised' cutoff_str 'WM'])
            F5 = load_untouch_nii([base_dir_sub fname '_slice' num2str(i3) '_denoised' cutoff_str 'WM.nii']); F5 = single(squeeze(F5.img));
        end
        for i5=1:size(F5,3)
            tmp = F5(:,:,i5);
            F6g(:,i5) = tmp(find(mask_GM(:,:,i3)==1));
            F6w(:,i5) = tmp(find(mask_WM(:,:,i3)==1));
            clear tmp
        end; clear i5 F5
        mF6g=mean(F6g,1); mF6g = 100*(mF6g-mean(mF6g))./mean(mF6g); mF6w=mean(F6w,1); mF6w = 100*(mF6w-mean(mF6w))./mean(mF6w);
        fighndl = figure('Position',[1 1 1152 720],'Color',[0.95,0.9,0.85],'InvertHardcopy','off','Visible','off'); % red [0.95,0.9,0.85], green [0.85,0.90,0.85], blue [0.89,0.94,0.99]
        if stp==1
            subplot2n(4,1,1), plot(mF6g,'color','k','linewidth',1.5), hold on, plot(mF2g,'color',[0,0.55,0.69],'linewidth',0.2), ylabel('pcent sig change'),
                grid on,grid minor, xlim([1 length(mF2g)]),ylim([min(1.25.*[mF2g,mF6g]) 1.25.*max([mF2g,mF6g])]), title(['Gray matter mean signal AFTER "11-func-anat-registration" (slice ' num2str(i3) ' of ' num2str(siz3) ')'],'Color','k','FontSize',14), legend({'Gray matter signal before COV-denoising'},'FontSize',12,'TextColor','k','Location','best')
            subplot2n(4,1,3), plot(mF6w,'color','k','linewidth',1.5), hold on, plot(mF2w,'color',[0.65,0.11,0.19],'linewidth',0.2), ylabel('pcent sig change'),
                grid on,grid minor, xlim([1 length(mF2g)]),ylim([min(1.25.*[mF2w,mF6w]) 1.25.*max([mF2w,mF6w])]), title(['White matter mean signal AFTER "11-func-anat-registration" (slice ' num2str(i3) ' of ' num2str(siz3) ')'],'Color','k','FontSize',14), legend({'White matter signal before COV-denoising'},'FontSize',12,'TextColor','k','Location','best')
        elseif stp==2
            subplot2n(4,1,1), plot(mF6g,'color','k','linewidth',1.5), hold on, plot(mF2g,'color',[0,0.55,0.69],'linewidth',0.2), ylabel('pcent sig change'),
                grid on,grid minor, xlim([1 length(mF2g)]),ylim([min(1.25.*[mF2g,mF6g]) 1.25.*max([mF2g,mF6g])]), title(['Gray matter mean signal AFTER "13-denoise4-WMregress" (slice ' num2str(i3) ' of ' num2str(siz3) ')'],'Color','k','FontSize',14), legend({'Gray matter signal before COV-denoising'},'FontSize',12,'TextColor','k','Location','best')
            subplot2n(4,1,3), plot(mF6w,'color','k','linewidth',1.5), hold on, plot(mF2w,'color',[0.65,0.11,0.19],'linewidth',0.2), ylabel('pcent sig change'),
                grid on,grid minor, xlim([1 length(mF2g)]),ylim([min(1.25.*[mF2w,mF6w]) 1.25.*max([mF2w,mF6w])]), title(['White matter mean signal AFTER "13-denoise4-WMregress" (slice ' num2str(i3) ' of ' num2str(siz3) ')'],'Color','k','FontSize',14), legend({'White matter signal before COV-denoising'},'FontSize',12,'TextColor','k','Location','best')
        elseif stp==3
            subplot2n(4,1,1), plot(mF6g,'color','k','linewidth',1.5), hold on, plot(mF2g,'color',[0,0.55,0.69],'linewidth',0.2), ylabel('pcent sig change'),
                grid on,grid minor, xlim([1 length(mF2g)]),ylim([min(1.25.*[mF2g,mF6g]) 1.25.*max([mF2g,mF6g])]), title(['Gray matter mean signal AFTER "12-denoise3-CSFregress" (slice ' num2str(i3) ' of ' num2str(siz3) ')'],'Color','k','FontSize',14), legend({'Gray matter signal before COV-denoising'},'FontSize',12,'TextColor','k','Location','best')
            subplot2n(4,1,3), plot(mF6w,'color','k','linewidth',1.5), hold on, plot(mF2w,'color',[0.65,0.11,0.19],'linewidth',0.2), ylabel('pcent sig change'),
                grid on,grid minor, xlim([1 length(mF2g)]),ylim([min(1.25.*[mF2w,mF6w]) 1.25.*max([mF2w,mF6w])]), title(['White matter mean signal AFTER "12-denoise3-CSFregress" (slice ' num2str(i3) ' of ' num2str(siz3) ')'],'Color','k','FontSize',14), legend({'White matter signal before COV-denoising'},'FontSize',12,'TextColor','k','Location','best')
        elseif stp==4
            subplot2n(4,1,1), plot(mF6g,'color','k','linewidth',1.5), hold on, plot(mF2g,'color',[0,0.55,0.69],'linewidth',0.2), ylabel('pcent sig change'),
                grid on,grid minor, xlim([1 length(mF2g)]),ylim([min(1.25.*[mF2g,mF6g]) 1.25.*max([mF2g,mF6g])]), title(['Gray matter mean signal AFTER "13-denoise4-WMregress" (slice ' num2str(i3) ' of ' num2str(siz3) ')'],'Color','k','FontSize',14), legend({'Gray matter signal before COV-denoising'},'FontSize',12,'TextColor','k','Location','best')
            subplot2n(4,1,3), plot(mF6w,'color','k','linewidth',1.5), hold on, plot(mF2w,'color',[0.65,0.11,0.19],'linewidth',0.2), ylabel('pcent sig change'),
                grid on,grid minor, xlim([1 length(mF2g)]),ylim([min(1.25.*[mF2w,mF6w]) 1.25.*max([mF2w,mF6w])]), title(['White matter mean signal AFTER "13-denoise4-WMregress" (slice ' num2str(i3) ' of ' num2str(siz3) ')'],'Color','k','FontSize',14), legend({'White matter signal before COV-denoising'},'FontSize',12,'TextColor','k','Location','best')
        end
        subplot2n(4,1,2), plot(mF2g,'color',[0,0.55,0.69],'linewidth',1.5), hold on, plot(mF6g,'color','k','linewidth',0.2), ylabel('pcent sig change'),
            grid on,grid minor, xlim([1 length(mF2g)]),ylim([min(1.25.*[mF2g,mF6g]) 1.25.*max([mF2g,mF6g])]), title(['Gray matter mean signal AFTER "14-DENOISE5-COVregress" (slice ' num2str(i3) ' of ' num2str(siz3) ')'],'Color',[0,0.55,0.69],'FontSize',14), legend({'Gray matter signal after COV-denoising'},'FontSize',12,'TextColor',[0,0.55,0.69],'Location','best')
        subplot2n(4,1,4), plot(mF2w,'color',[0.65,0.11,0.19],'linewidth',1.5), hold on, plot(mF6w,'color','k','linewidth',0.2), ylabel('pcent sig change'),
            grid on,grid minor, xlim([1 length(mF2g)]),ylim([min(1.25.*[mF2w,mF6w]) 1.25.*max([mF2w,mF6w])]), title(['White matter mean signal AFTER "14-DENOISE5-COVregress" (slice ' num2str(i3) ' of ' num2str(siz3) ')'],'Color',[0.65,0.11,0.19],'FontSize',14), legend({'White matter signal after COV-denoising'},'FontSize',12,'TextColor',[0.65,0.11,0.19],'Location','best')
        saveas(fighndl,[QCpath 'global_signal_13denoise4_vs_14denoise5_slice' sprintf('%.2d',i3) '.jpg'])
        close(fighndl)
        
        C3g = corrcoef(F6g'); C3g = uppertriangle(C3g); C3w = corrcoef(F6w'); C3w = uppertriangle(C3w);
        fighndl = figure('Position',[1 1 1152 720],'Color',[0.95,0.9,0.85],'InvertHardcopy','off','Visible','off'); % red [0.95,0.9,0.85], green [0.85,0.90,0.85], blue [0.89,0.94,0.99]
        if stp==1
            subplot2n(4,1,1), histogram(C3g,'EdgeColor','none','FaceColor','k','FaceAlpha',0.5,'Normalization','probability'); grid on,grid minor, xlim([-1 1]), hold on, xline(0,'Color','k','linewidth',1.5); hold off
                title(sprintf('Histogram of voxel-to-voxel GRAY MATTER correlations AFTER 11-func-anat-registration (slice %d of %d)\n%.1f%% of the correlations were > 0  [ mean=%.2f, median=%.2f, SD=%.2f ]',i3,siz3,100*length(find(C3g>0))/length(C3g),mean(C3g),median(C3g),std(C3g)),'Color','k','FontSize',14)
            subplot2n(4,1,3), histogram(C3w,'EdgeColor','none','FaceColor','k','FaceAlpha',0.5,'Normalization','probability'); grid on,grid minor, xlim([-1 1]), hold on, xline(0,'Color','k','linewidth',1.5); hold off
                title(sprintf('WHITE MATTER: AFTER 11-func-anat-registration (slice %d of %d): %.1f%% of the correlations were > 0  [ mean=%.2f, median=%.2f, SD=%.2f ]',i3,siz3,100*length(find(C3w>0))/length(C3w),mean(C3w),median(C3w),std(C3w)),'Color','k','FontSize',14)
        elseif stp==2
            subplot2n(4,1,1), histogram(C3g,'EdgeColor','none','FaceColor','k','FaceAlpha',0.5,'Normalization','probability'); grid on,grid minor, xlim([-1 1]), hold on, xline(0,'Color','k','linewidth',1.5); hold off
                title(sprintf('Histogram of voxel-to-voxel GRAY MATTER correlations AFTER 13-denoise4-WMregress (slice %d of %d)\n%.1f%% of the correlations were > 0  [ mean=%.2f, median=%.2f, SD=%.2f ]',i3,siz3,100*length(find(C3g>0))/length(C3g),mean(C3g),median(C3g),std(C3g)),'Color','k','FontSize',14)
            subplot2n(4,1,3), histogram(C3w,'EdgeColor','none','FaceColor','k','FaceAlpha',0.5,'Normalization','probability'); grid on,grid minor, xlim([-1 1]), hold on, xline(0,'Color','k','linewidth',1.5); hold off
                title(sprintf('WHITE MATTER: AFTER 13-denoise4-WMregress (slice %d of %d): %.1f%% of the correlations were > 0  [ mean=%.2f, median=%.2f, SD=%.2f ]',i3,siz3,100*length(find(C3w>0))/length(C3w),mean(C3w),median(C3w),std(C3w)),'Color','k','FontSize',14)
        elseif stp==3
            subplot2n(4,1,1), histogram(C3g,'EdgeColor','none','FaceColor','k','FaceAlpha',0.5,'Normalization','probability'); grid on,grid minor, xlim([-1 1]), hold on, xline(0,'Color','k','linewidth',1.5); hold off
                title(sprintf('Histogram of voxel-to-voxel GRAY MATTER correlations AFTER 12-denoise3-CSFregress (slice %d of %d)\n%.1f%% of the correlations were > 0  [ mean=%.2f, median=%.2f, SD=%.2f ]',i3,siz3,100*length(find(C3g>0))/length(C3g),mean(C3g),median(C3g),std(C3g)),'Color','k','FontSize',14)
            subplot2n(4,1,3), histogram(C3w,'EdgeColor','none','FaceColor','k','FaceAlpha',0.5,'Normalization','probability'); grid on,grid minor, xlim([-1 1]), hold on, xline(0,'Color','k','linewidth',1.5); hold off
                title(sprintf('WHITE MATTER: AFTER 12-denoise3-CSFregress (slice %d of %d): %.1f%% of the correlations were > 0  [ mean=%.2f, median=%.2f, SD=%.2f ]',i3,siz3,100*length(find(C3w>0))/length(C3w),mean(C3w),median(C3w),std(C3w)),'Color','k','FontSize',14)
        elseif stp==4
            subplot2n(4,1,1), histogram(C3g,'EdgeColor','none','FaceColor','k','FaceAlpha',0.5,'Normalization','probability'); grid on,grid minor, xlim([-1 1]), hold on, xline(0,'Color','k','linewidth',1.5); hold off
                title(sprintf('Histogram of voxel-to-voxel GRAY MATTER correlations AFTER 13-denoise4-WMregress (slice %d of %d)\n%.1f%% of the correlations were > 0  [ mean=%.2f, median=%.2f, SD=%.2f ]',i3,siz3,100*length(find(C3g>0))/length(C3g),mean(C3g),median(C3g),std(C3g)),'Color','k','FontSize',14)
            subplot2n(4,1,3), histogram(C3w,'EdgeColor','none','FaceColor','k','FaceAlpha',0.5,'Normalization','probability'); grid on,grid minor, xlim([-1 1]), hold on, xline(0,'Color','k','linewidth',1.5); hold off
                title(sprintf('WHITE MATTER: AFTER 13-denoise4-WMregress (slice %d of %d): %.1f%% of the correlations were > 0  [ mean=%.2f, median=%.2f, SD=%.2f ]',i3,siz3,100*length(find(C3w>0))/length(C3w),mean(C3w),median(C3w),std(C3w)),'Color','k','FontSize',14)
        end
        subplot2n(4,1,2), histogram(Cg,'EdgeColor','none','FaceColor',[0,0.55,0.69],'FaceAlpha',0.5,'Normalization','probability'); grid on,grid minor, xlim([-1 1]), hold on, xline(0,'Color','k','linewidth',1.5); hold off
            title(sprintf('GRAY MATTER: AFTER 14-DENOISE5-COVregress (slice %d of %d): %.1f%% of the correlations were > 0  [ mean=%.2f, median=%.2f, SD=%.2f ]',i3,siz3,100*length(find(Cg>0))/length(Cg),mean(Cg),median(Cg),std(Cg)),'Color',[0,0.55,0.69],'FontSize',14)
        subplot2n(4,1,4), histogram(Cw,'EdgeColor','none','FaceColor',[0.65,0.11,0.19],'FaceAlpha',0.5,'Normalization','probability'); grid on,grid minor, xlim([-1 1]), hold on, xline(0,'Color','k','linewidth',1.5); hold off
            title(sprintf('WHITE MATTER: AFTER 14-DENOISE5-COVregress (slice %d of %d): %.1f%% of the correlations were > 0  [ mean=%.2f, median=%.2f, SD=%.2f ]',i3,siz3,100*length(find(Cw>0))/length(Cw),mean(Cw),median(Cw),std(Cw)),'Color',[0.65,0.11,0.19],'FontSize',14)
        saveas(fighndl,[QCpath 'hist_FC_compare_13denoise4_14denoise5_slice' sprintf('%.2d',i3) '.jpg'])
        close(fighndl)
        F6g_=cat(1,F6g_,F6g); mF6g_=cat(1,mF6g_,mF6g); C3g_=cat(1,C3g_,C3g);
        F6w_=cat(1,F6w_,F6w); mF6w_=cat(1,mF6w_,mF6w); C3w_=cat(1,C3w_,C3w);
        clear F6g F6w mF6g mF6w C3g C3w
    end
    
    F2g_=cat(1,F2g_,F2g); mF2g_=cat(1,mF2g_,mF2g); F2w_=cat(1,F2w_,F2w); mF2w_=cat(1,mF2w_,mF2w); Cg_=cat(1,Cg_,Cg); Cw_=cat(1,Cw_,Cw); F4_=cat(1,F4_,F4); mF4_=cat(1,mF4_,mF4); C2_=cat(1,C2_,C2);
    clear F2g F2w F4 TR Smask1 svar1 svar2 mF2g mF2w mF4 Cg Cw C2
end

[~,svar1] = sort(var(F2g_,0,2)); [~,spsc1] = sort(meanPSC(F2g_));
[~,svar2] = sort(var(F2w_,0,2)); [~,spsc2] = sort(meanPSC(F2w_));
sortorder1b = sortorder1; sortorder1=[]; f1=0; %#ok<*ASGLU>
sortorder2b = sortorder2; sortorder2=[]; f2=0; %#ok<*ASGLU>
for im=1:length(sortorder1b)
    temp = sortorder1b{im}; temp = temp + f1; f1 = f1 + length(temp);
    sortorder1 = cat(1,sortorder1,temp); clear temp
    temp = sortorder2b{im}; temp = temp + f2; f2 = f2 + length(temp);
    sortorder2 = cat(1,sortorder2,temp); clear temp
end; clear im f1 f2
sortorder1 = spsc1; % options: spsc for percentage signal change, svar for variance, (comment out the line if you want to use distance from the center of the cord separately for each slice)
sortorder2 = spsc2; % options: spsc for percentage signal change, svar for variance, (comment out the line if you want to use distance from the center of the cord separately for each slice)
for im=max(sortorder1):-1:1, if isempty(find(sortorder1==im)), sortorder1(find(sortorder1>im)) = sortorder1(find(sortorder1>im))-1; end, end; clear im %#ok<*EFIND>
for im=max(sortorder2):-1:1, if isempty(find(sortorder2==im)), sortorder2(find(sortorder2>im)) = sortorder2(find(sortorder2>im))-1; end, end; clear im %#ok<*EFIND>

fighndl = figure('Position',[1 1 1152 720],'Color',[0.89,0.94,0.99],'InvertHardcopy','off','Visible','off'); % red [0.95,0.9,0.85], green [0.85,0.90,0.85], blue [0.89,0.94,0.99]
subplot2n(2,1,1), imagesc(F2g_(sortorder1,:)),colormap(cmap),colorbar, xlabel('time (in samples)'),ylabel(sprintf('voxels (sorted by mean percentage signal change;\nbottom row is highest)')),
  title(['Time series within the GRAY MATTER AFTER 14-DENOISE5-COVregress (all slices combined)'],'Color',[0,0.55,0.69],'FontSize',14);
subplot2n(2,1,2), imagesc(F2w_(sortorder2,:)),colormap(cmap),colorbar, xlabel('time (in samples)'),ylabel(sprintf('voxels (sorted by mean percentage signal change;\nbottom row is highest)')),
  title(['Time series within the WHITE MATTER AFTER 14-DENOISE5-COVregress (all slices combined)'],'Color',[0.65,0.11,0.19],'FontSize',14);
saveas(fighndl,[QCpath 'fMRI_ts_14denoise5_allSlices' '.jpg'])
 close(fighndl)
mF2g_=mean(mF2g_,1); mF2w_=mean(mF2w_,1); mF4_=mean(mF4_,1);
fighndl = figure('Position',[1 1 1152 720],'Color',[0.89,0.94,0.99],'InvertHardcopy','off','Visible','off'); % red [0.95,0.9,0.85], green [0.85,0.90,0.85], blue [0.89,0.94,0.99]
subplot2n(3,1,1), plot(mF4_,'color','k','linewidth',1.5), hold on, plot(mF2g_,'color',[0,0.55,0.69],'linewidth',0.2), plot(mF2w_,'color',[0.65,0.11,0.19],'linewidth',0.2), xlabel('time (in samples)'),ylabel('percentage signal change'),
    grid on,grid minor, xlim([1 length(mF2g_)]),ylim([1.25.*min([mF2g_,mF2w_,mF4_]) 1.25.*max([mF2g_,mF2w_,mF4_])]), title(['Global mean signal in UNPROCESSED RAW DATA (all slices averaged)'],'Color','k','FontSize',14), legend({'Global signal (raw data)'},'FontSize',12,'TextColor','k','Location','best')
subplot2n(3,1,2), plot(mF2g_,'color',[0,0.55,0.69],'linewidth',1.5), hold on, plot(mF2w_,'color',[0.65,0.11,0.19],'linewidth',0.2), plot(mF4_,'color','k','linewidth',0.2), xlabel('time (in samples)'),ylabel('percentage signal change'),
    grid on,grid minor, xlim([1 length(mF2g_)]),ylim([1.25.*min([mF2g_,mF2w_,mF4_]) 1.25.*max([mF2g_,mF2w_,mF4_])]), title(['Gray matter mean signal AFTER "14-DENOISE5-COVregress" (all slices averaged)'],'Color',[0,0.55,0.69],'FontSize',14), legend({'Gray matter signal after 14-denoise5'},'FontSize',12,'TextColor',[0,0.55,0.69],'Location','best')
subplot2n(3,1,3), plot(mF2w_,'color',[0.65,0.11,0.19],'linewidth',1.5), hold on, plot(mF4_,'color','k','linewidth',0.2), plot(mF2g_,'color',[0,0.55,0.69],'linewidth',0.2), xlabel('time (in samples)'),ylabel('percentage signal change'),
    grid on,grid minor, xlim([1 length(mF2g_)]),ylim([1.25.*min([mF2g_,mF2w_,mF4_]) 1.25.*max([mF2g_,mF2w_,mF4_])]), title(['White matter mean signal AFTER "14-DENOISE5-COVregress" (all slices averaged)'],'Color',[0.65,0.11,0.19],'FontSize',14), legend({'White matter signal after 14-denoise5'},'FontSize',12,'TextColor',[0.65,0.11,0.19],'Location','best')
saveas(fighndl,[QCpath 'global_signal_14denoise5_allSlices' '.jpg'])
 close(fighndl)
fighndl = figure('Position',[1 1 1152 720],'Color',[0.89,0.94,0.99],'InvertHardcopy','off','Visible','off'); % red [0.95,0.9,0.85], green [0.85,0.90,0.85], blue [0.89,0.94,0.99]
subplot2n(3,1,1), histogram(C2_,'EdgeColor','none','FaceColor','k','FaceAlpha',0.5,'Normalization','probability'); grid on,grid minor, xlim([-1 1]), hold on, xline(0,'Color','k','linewidth',1.5); hold off
    title(sprintf('Histogram of voxel-to-voxel correlations in UNPROCESSED RAW DATA (all slices combined)\n%.1f%% of the correlations were > 0  [ mean=%.2f, median=%.2f, SD=%.2f ]',100*length(find(C2_>0))/length(C2_),mean(C2_),median(C2_),std(C2_)),'Color','k','FontSize',14)
subplot2n(3,1,2), histogram(Cg_,'EdgeColor','none','FaceColor',[0,0.55,0.69],'FaceAlpha',0.5,'Normalization','probability'); grid on,grid minor, xlim([-1 1]), hold on, xline(0,'Color','k','linewidth',1.5); hold off
    title(sprintf('Histogram of voxel-to-voxel GRAY MATTER correlations AFTER 14-DENOISE5-COVregress (all slices combined)\n%.1f%% of the correlations were > 0  [ mean=%.2f, median=%.2f, SD=%.2f ]',100*length(find(Cg_>0))/length(Cg_),mean(Cg_),median(Cg_),std(Cg_)),'Color',[0,0.55,0.69],'FontSize',14)
subplot2n(3,1,3), histogram(Cw_,'EdgeColor','none','FaceColor',[0.65,0.11,0.19],'FaceAlpha',0.5,'Normalization','probability'); grid on,grid minor, xlim([-1 1]), hold on, xline(0,'Color','k','linewidth',1.5); hold off
    title(sprintf('Histogram of voxel-to-voxel WHITE MATTER correlations AFTER 14-DENOISE5-COVregress (all slices combined)\n%.1f%% of the correlations were > 0  [ mean=%.2f, median=%.2f, SD=%.2f ]',100*length(find(Cw_>0))/length(Cw_),mean(Cw_),median(Cw_),std(Cw_)),'Color',[0.65,0.11,0.19],'FontSize',14)
saveas(fighndl,[QCpath 'hist_FC_compare_14denoise5_allSlices' '.jpg'])
 close(fighndl)

if stp~=0
    mF6g_=mean(mF6g_,1); mF6w_=mean(mF6w_,1);
    fighndl = figure('Position',[1 1 1152 720],'Color',[0.89,0.94,0.99],'InvertHardcopy','off','Visible','off'); % red [0.95,0.9,0.85], green [0.85,0.90,0.85], blue [0.89,0.94,0.99]
    if stp==1
    	subplot2n(4,1,1), plot(mF6g_,'color','k','linewidth',1.5), hold on, plot(mF2g_,'color',[0,0.55,0.69],'linewidth',0.2), ylabel('pcent sig change'),
            grid on,grid minor, xlim([1 length(mF2g_)]),ylim([1.25.*min([mF2g_,mF6g_]) 1.25.*max([mF2g_,mF6g_])]), title(['Gray matter mean signal AFTER "11-func-anat-registration" (all slices averaged)'],'Color','k','FontSize',14), legend({'Gray matter signal before COV-denoising'},'FontSize',12,'TextColor','k','Location','best')
        subplot2n(4,1,3), plot(mF6w_,'color','k','linewidth',1.5), hold on, plot(mF2w_,'color',[0.65,0.11,0.19],'linewidth',0.2), ylabel('pcent sig change'),
            grid on,grid minor, xlim([1 length(mF2g_)]),ylim([1.25.*min([mF2w_,mF6w_]) 1.25.*max([mF2w_,mF6w_])]), title(['White matter mean signal AFTER "11-func-anat-registration" (all slices averaged)'],'Color','k','FontSize',14), legend({'White matter signal before COV-denoising'},'FontSize',12,'TextColor','k','Location','best')
    elseif stp==2
    	subplot2n(4,1,1), plot(mF6g_,'color','k','linewidth',1.5), hold on, plot(mF2g_,'color',[0,0.55,0.69],'linewidth',0.2), ylabel('pcent sig change'),
            grid on,grid minor, xlim([1 length(mF2g_)]),ylim([1.25.*min([mF2g_,mF6g_]) 1.25.*max([mF2g_,mF6g_])]), title(['Gray matter mean signal AFTER "13-DENOISE4-WMregress" (all slices averaged)'],'Color','k','FontSize',14), legend({'Gray matter signal before COV-denoising'},'FontSize',12,'TextColor','k','Location','best')
        subplot2n(4,1,3), plot(mF6w_,'color','k','linewidth',1.5), hold on, plot(mF2w_,'color',[0.65,0.11,0.19],'linewidth',0.2), ylabel('pcent sig change'),
            grid on,grid minor, xlim([1 length(mF2g_)]),ylim([1.25.*min([mF2w_,mF6w_]) 1.25.*max([mF2w_,mF6w_])]), title(['White matter mean signal AFTER "13-DENOISE4-WMregress" (all slices averaged)'],'Color','k','FontSize',14), legend({'White matter signal before COV-denoising'},'FontSize',12,'TextColor','k','Location','best')
    elseif stp==3
    	subplot2n(4,1,1), plot(mF6g_,'color','k','linewidth',1.5), hold on, plot(mF2g_,'color',[0,0.55,0.69],'linewidth',0.2), ylabel('pcent sig change'),
            grid on,grid minor, xlim([1 length(mF2g_)]),ylim([1.25.*min([mF2g_,mF6g_]) 1.25.*max([mF2g_,mF6g_])]), title(['Gray matter mean signal AFTER "12-DENOISE3-CSFregress" (all slices averaged)'],'Color','k','FontSize',14), legend({'Gray matter signal before COV-denoising'},'FontSize',12,'TextColor','k','Location','best')
        subplot2n(4,1,3), plot(mF6w_,'color','k','linewidth',1.5), hold on, plot(mF2w_,'color',[0.65,0.11,0.19],'linewidth',0.2), ylabel('pcent sig change'),
            grid on,grid minor, xlim([1 length(mF2g_)]),ylim([1.25.*min([mF2w_,mF6w_]) 1.25.*max([mF2w_,mF6w_])]), title(['White matter mean signal AFTER "12-DENOISE3-CSFregress" (all slices averaged)'],'Color','k','FontSize',14), legend({'White matter signal before COV-denoising'},'FontSize',12,'TextColor','k','Location','best')
    elseif stp==4
    	subplot2n(4,1,1), plot(mF6g_,'color','k','linewidth',1.5), hold on, plot(mF2g_,'color',[0,0.55,0.69],'linewidth',0.2), ylabel('pcent sig change'),
            grid on,grid minor, xlim([1 length(mF2g_)]),ylim([1.25.*min([mF2g_,mF6g_]) 1.25.*max([mF2g_,mF6g_])]), title(['Gray matter mean signal AFTER "13-DENOISE4-WMregress" (all slices averaged)'],'Color','k','FontSize',14), legend({'Gray matter signal before COV-denoising'},'FontSize',12,'TextColor','k','Location','best')
        subplot2n(4,1,3), plot(mF6w_,'color','k','linewidth',1.5), hold on, plot(mF2w_,'color',[0.65,0.11,0.19],'linewidth',0.2), ylabel('pcent sig change'),
            grid on,grid minor, xlim([1 length(mF2g_)]),ylim([1.25.*min([mF2w_,mF6w_]) 1.25.*max([mF2w_,mF6w_])]), title(['White matter mean signal AFTER "13-DENOISE4-WMregress" (all slices averaged)'],'Color','k','FontSize',14), legend({'White matter signal before COV-denoising'},'FontSize',12,'TextColor','k','Location','best')
    end
    subplot2n(4,1,2), plot(mF2g_,'color',[0,0.55,0.69],'linewidth',1.5), hold on, plot(mF6g_,'color','k','linewidth',0.2), ylabel('pcent sig change'),
        grid on,grid minor, xlim([1 length(mF2g_)]),ylim([1.25.*min([mF2g_,mF6g_]) 1.25.*max([mF2g_,mF6g_])]), title(['Gray matter mean signal AFTER "14-DENOISE5-COVregress" (all slices averaged)'],'Color',[0,0.55,0.69],'FontSize',14), legend({'Gray matter signal after COV-denoising'},'FontSize',12,'TextColor',[0,0.55,0.69],'Location','best')
    subplot2n(4,1,4), plot(mF2w_,'color',[0.65,0.11,0.19],'linewidth',1.5), hold on, plot(mF6w_,'color','k','linewidth',0.2), ylabel('pcent sig change'),
        grid on,grid minor, xlim([1 length(mF2g_)]),ylim([1.25.*min([mF2w_,mF6w_]) 1.25.*max([mF2w_,mF6w_])]), title(['White matter mean signal AFTER "14-DENOISE5-COVregress" (all slices averaged)'],'Color',[0.65,0.11,0.19],'FontSize',14), legend({'White matter signal after COV-denoising'},'FontSize',12,'TextColor',[0.65,0.11,0.19],'Location','best')
    saveas(fighndl,[QCpath 'global_signal_13denoise4_vs_14denoise5_allSlices' '.jpg'])
     close(fighndl)
    
    fighndl = figure('Position',[1 1 1152 720],'Color',[0.89,0.94,0.99],'InvertHardcopy','off','Visible','off'); % red [0.95,0.9,0.85], green [0.85,0.90,0.85], blue [0.89,0.94,0.99]
    if stp==1
    	subplot2n(4,1,1), histogram(C3g_,'EdgeColor','none','FaceColor','k','FaceAlpha',0.5,'Normalization','probability'); grid on,grid minor, xlim([-1 1]), hold on, xline(0,'Color','k','linewidth',1.5); hold off
            title(sprintf('Histogram of voxel-to-voxel GRAY MATTER correlations AFTER 11-registration (all slices combined)\n%.1f%% of the correlations were > 0  [ mean=%.2f, median=%.2f, SD=%.2f ]',100*length(find(C3g_>0))/length(C3g_),mean(C3g_),median(C3g_),std(C3g_)),'Color','k','FontSize',14)
        subplot2n(4,1,3), histogram(C3w_,'EdgeColor','none','FaceColor','k','FaceAlpha',0.5,'Normalization','probability'); grid on,grid minor, xlim([-1 1]), hold on, xline(0,'Color','k','linewidth',1.5); hold off
            title(sprintf('WHITE MATTER: AFTER 11-registration (all slices): %.1f%% of the correlations were > 0  [ mean=%.2f, median=%.2f, SD=%.2f ]',100*length(find(C3w_>0))/length(C3w_),mean(C3w_),median(C3w_),std(C3w_)),'Color','k','FontSize',14)
    elseif stp==2
    	subplot2n(4,1,1), histogram(C3g_,'EdgeColor','none','FaceColor','k','FaceAlpha',0.5,'Normalization','probability'); grid on,grid minor, xlim([-1 1]), hold on, xline(0,'Color','k','linewidth',1.5); hold off
            title(sprintf('Histogram of voxel-to-voxel GRAY MATTER correlations AFTER 13-denoise4-WMregress (all slices combined)\n%.1f%% of the correlations were > 0  [ mean=%.2f, median=%.2f, SD=%.2f ]',100*length(find(C3g_>0))/length(C3g_),mean(C3g_),median(C3g_),std(C3g_)),'Color','k','FontSize',14)
        subplot2n(4,1,3), histogram(C3w_,'EdgeColor','none','FaceColor','k','FaceAlpha',0.5,'Normalization','probability'); grid on,grid minor, xlim([-1 1]), hold on, xline(0,'Color','k','linewidth',1.5); hold off
            title(sprintf('WHITE MATTER: AFTER 13-denoise4-WMregress (all slices): %.1f%% of the correlations were > 0  [ mean=%.2f, median=%.2f, SD=%.2f ]',100*length(find(C3w_>0))/length(C3w_),mean(C3w_),median(C3w_),std(C3w_)),'Color','k','FontSize',14)
    elseif stp==3
    	subplot2n(4,1,1), histogram(C3g_,'EdgeColor','none','FaceColor','k','FaceAlpha',0.5,'Normalization','probability'); grid on,grid minor, xlim([-1 1]), hold on, xline(0,'Color','k','linewidth',1.5); hold off
            title(sprintf('Histogram of voxel-to-voxel GRAY MATTER correlations AFTER 12-denoise3-CSFregress (all slices combined)\n%.1f%% of the correlations were > 0  [ mean=%.2f, median=%.2f, SD=%.2f ]',100*length(find(C3g_>0))/length(C3g_),mean(C3g_),median(C3g_),std(C3g_)),'Color','k','FontSize',14)
        subplot2n(4,1,3), histogram(C3w_,'EdgeColor','none','FaceColor','k','FaceAlpha',0.5,'Normalization','probability'); grid on,grid minor, xlim([-1 1]), hold on, xline(0,'Color','k','linewidth',1.5); hold off
            title(sprintf('WHITE MATTER: AFTER 12-denoise3-CSFregress (all slices): %.1f%% of the correlations were > 0  [ mean=%.2f, median=%.2f, SD=%.2f ]',100*length(find(C3w_>0))/length(C3w_),mean(C3w_),median(C3w_),std(C3w_)),'Color','k','FontSize',14)
    elseif stp==4
    	subplot2n(4,1,1), histogram(C3g_,'EdgeColor','none','FaceColor','k','FaceAlpha',0.5,'Normalization','probability'); grid on,grid minor, xlim([-1 1]), hold on, xline(0,'Color','k','linewidth',1.5); hold off
            title(sprintf('Histogram of voxel-to-voxel GRAY MATTER correlations AFTER 13-denoise4-WMregress (all slices combined)\n%.1f%% of the correlations were > 0  [ mean=%.2f, median=%.2f, SD=%.2f ]',100*length(find(C3g_>0))/length(C3g_),mean(C3g_),median(C3g_),std(C3g_)),'Color','k','FontSize',14)
        subplot2n(4,1,3), histogram(C3w_,'EdgeColor','none','FaceColor','k','FaceAlpha',0.5,'Normalization','probability'); grid on,grid minor, xlim([-1 1]), hold on, xline(0,'Color','k','linewidth',1.5); hold off
            title(sprintf('WHITE MATTER: AFTER 13-denoise4-WMregress (all slices): %.1f%% of the correlations were > 0  [ mean=%.2f, median=%.2f, SD=%.2f ]',100*length(find(C3w_>0))/length(C3w_),mean(C3w_),median(C3w_),std(C3w_)),'Color','k','FontSize',14)
    end
    subplot2n(4,1,2), histogram(Cg_,'EdgeColor','none','FaceColor',[0,0.55,0.69],'FaceAlpha',0.5,'Normalization','probability'); grid on,grid minor, xlim([-1 1]), hold on, xline(0,'Color','k','linewidth',1.5); hold off
       title(sprintf('GRAY MATTER: AFTER 14-DENOISE5-COVregress (all slices): %.1f%% of correlations > 0  [ mean=%.2f, median=%.2f, SD=%.2f ]',100*length(find(Cg_>0))/length(Cg_),mean(Cg_),median(Cg_),std(Cg_)),'Color',[0,0.55,0.69],'FontSize',14)
    subplot2n(4,1,4), histogram(Cw_,'EdgeColor','none','FaceColor',[0.65,0.11,0.19],'FaceAlpha',0.5,'Normalization','probability'); grid on,grid minor, xlim([-1 1]), hold on, xline(0,'Color','k','linewidth',1.5); hold off
       title(sprintf('WHITE MATTER: AFTER 14-DENOISE5-COVregress (all slices): %.1f%% of the correlations were > 0  [ mean=%.2f, median=%.2f, SD=%.2f ]',100*length(find(Cw_>0))/length(Cw_),mean(Cw_),median(Cw_),std(Cw_)),'Color',[0.65,0.11,0.19],'FontSize',14)
    saveas(fighndl,[QCpath 'hist_FC_compare_13denoise4_14denoise5_allSlices' '.jpg'])
     close(fighndl)
end
clear svar1 spsc1 svar2 spsc2 sortorder1 sortorder2 F2g_ mF2g_ F2w_ mF2w_ Cg_ Cw_ F4_ mF4_ C2_ F6g_ F6w_ mF6g_ mF6w_ C3g_ C3w_

%%% Make QC video (needs audio video toolbox in Matlab)
if ~isempty(which('VideoWriter')) && ((save_videos==1)||(save_videos==2)) % check if Matlab's audio video toolbox exists

subplot = @(m,n,p) subtightplot(m, n, p, [0.001 0.005], [0.01 0.001], [0.01 0.001]);
scfMRItb_04_unzipFile(base_dir_sub, fname_anat, '_mask_SC')
try
    scfMRItb_04_unzipFile(base_dir_sub, fname_anat, '_mask_SC')
    mask_SC2 = load_untouch_nii([base_dir_sub fname_anat '_mask_SC.nii']); mask_SC2 = mask_SC2.img;
catch
    mask_SC = load_untouch_nii([base_dir_sub fname_anat '_mask_GM.nii']);
    mask_SC.img = or(mask_GM, mask_WM);
    save_untouch_nii(mask_SC, [base_dir_sub fname_anat '_mask_SC.nii']);
    mask_SC2 = mask_SC.img; clear mask_SC
end
for M=1:size(mask_SC2,3)
    mask_SC = rot90(mask_SC2(:,:,M));
    [xmask_SC,ymask_SC] = find(mask_SC==1);
    minx_SC(:,M) = round(min(xmask_SC)-0.1*(size(mask_SC,1))); maxx_SC(:,M)=round(max(xmask_SC)+0.1*(size(mask_SC,1)));
    miny_SC(:,M) = round(min(ymask_SC)-0.1*(size(mask_SC,2))); maxy_SC(:,M)=round(max(ymask_SC)+0.1*(size(mask_SC,2)));
    if (maxx_SC(:,M)-minx_SC(:,M))<(maxy_SC(:,M)-miny_SC(:,M))
        factr = ((maxy_SC(:,M)-miny_SC(:,M))-(maxx_SC(:,M)-minx_SC(:,M)))/2;
        if factr==round(factr)
            maxx_SC(:,M)=maxx_SC(:,M)+factr; minx_SC(:,M)=minx_SC(:,M)-factr;
        else
            maxx_SC(:,M)=maxx_SC(:,M)+factr+0.5; minx_SC(:,M)=minx_SC(:,M)-factr+0.5;
        end
    elseif (maxy_SC(:,M)-miny_SC(:,M))<(maxx_SC(:,M)-minx_SC(:,M))
        factr = ((maxx_SC(:,M)-minx_SC(:,M))-(maxy_SC(:,M)-miny_SC(:,M)))/2;
        if factr==round(factr)
            maxy_SC(:,M)=maxy_SC(:,M)+factr; miny_SC(:,M)=miny_SC(:,M)-factr;
        else
            maxy_SC(:,M)=maxy_SC(:,M)+factr+0.5; miny_SC(:,M)=miny_SC(:,M)-factr+0.5;
        end
    end
    clear mask_SC xmask_SC ymask_SC factr
end; clear M

for i3 = 1 : siz3
    try waitbar((i3/siz3),wbar3,sprintf('14. Generate QC video: slice (%d) of (%d)',i3,siz3)); catch, end
    fprintf('14. Generate QC video: slice (%d) of (%d), subject (%d) of (%d), run (%d) of (%d)\n',i3,siz3,k,size(func_files,1),k2,size(func_files,2))
    if stp==4
        scfMRItb_04_resplitData(base_dir_sub, fname, ['_denoised' cutoff_str 'WMcov'], siz3) % if slice-wise data doesn't exist but combined 4D data exists, then split the data to generate the needed slice-wise data
        F1 = load_untouch_nii([base_dir_sub fname '_slice' num2str(i3) '_denoised' cutoff_str 'WMcov.nii']);
    elseif stp==3
        scfMRItb_04_resplitData(base_dir_sub, fname, ['_denoised' cutoff_str 'cov'], siz3) % if slice-wise data doesn't exist but combined 4D data exists, then split the data to generate the needed slice-wise data
        F1 = load_untouch_nii([base_dir_sub fname '_slice' num2str(i3) '_denoised' cutoff_str 'cov.nii']);
    elseif stp==2
        scfMRItb_04_resplitData(base_dir_sub, fname, '_WMcov', siz3) % if slice-wise data doesn't exist but combined 4D data exists, then split the data to generate the needed slice-wise data
        F1 = load_untouch_nii([base_dir_sub fname '_slice' num2str(i3) '_WMcov.nii']);
    elseif stp==1
        scfMRItb_04_resplitData(base_dir_sub, fname, '_cov', siz3) % if slice-wise data doesn't exist but combined 4D data exists, then split the data to generate the needed slice-wise data
        F1 = load_untouch_nii([base_dir_sub fname '_slice' num2str(i3) '_cov.nii']);
    end
    TR = F1.hdr.dime.pixdim(5); F1 = rot90(single(squeeze(F1.img)));
    video = VideoWriter([QCpath 'fMRIvideo_14denoise5_slice' sprintf('%.2d',i3) '.mp4'],'MPEG-4'); %#ok<*TNMLP> % create the video object
    video.FrameRate = round(30/TR); % 30 seconds of data in every second of the video
    video.Quality = 90; % very limited compression so that the images are shown as is (and thus avoid misunderstandings due to video quality)
    open(video); % open the file for writing
    
    [sizeFx_SC,sizeFy_SC] = size(F1(:,:,1));
    F2 = F1(minx_SC:maxx_SC,miny_SC:maxy_SC,:);
    for i5=1:size(F1,3)
        F1(:,:,i5) = imresize(F2(:,:,i5),[sizeFx_SC sizeFy_SC]);
    end
    
    if (save_videos==1)
    if stp==1
        scfMRItb_04_resplitData(base_dir_sub, fname, '_warped', siz3) % if slice-wise data doesn't exist but combined 4D data exists, then split the data to generate the needed slice-wise data
        F3 = load_untouch_nii([base_dir_sub fname '_slice' num2str(i3) '_warped.nii']); F3 = rot90(squeeze(F3.img));
    elseif stp==2
        scfMRItb_04_resplitData(base_dir_sub, fname, '_WM', siz3) % if slice-wise data doesn't exist but combined 4D data exists, then split the data to generate the needed slice-wise data
        F3 = load_untouch_nii([base_dir_sub fname '_slice' num2str(i3) '_WM.nii']); F3 = rot90(single(squeeze(F3.img)));
    elseif stp==3
        scfMRItb_04_resplitData(base_dir_sub, fname, ['_denoised' cutoff_str], siz3) % if slice-wise data doesn't exist but combined 4D data exists, then split the data to generate the needed slice-wise data
        F3 = load_untouch_nii([base_dir_sub fname '_slice' num2str(i3) '_denoised' cutoff_str '.nii']); F3 = rot90(single(squeeze(F3.img)));
    elseif stp==4
        scfMRItb_04_resplitData(base_dir_sub, fname, ['_denoised' cutoff_str 'WM'], siz3) % if slice-wise data doesn't exist but combined 4D data exists, then split the data to generate the needed slice-wise data
        F3 = load_untouch_nii([base_dir_sub fname '_slice' num2str(i3) '_denoised' cutoff_str 'WM.nii']); F3 = rot90(single(squeeze(F3.img)));
    end
    [sizeFx,sizeFy] = size(F3(:,:,1));
    F4 = F3(minx_SC:maxx_SC,miny_SC:maxy_SC,:);
    for i5=1:size(F3,3)
        F3(:,:,i5) = imresize(F4(:,:,i5),[sizeFx sizeFy]);
    end
    clear sizeFx sizeFy F2 F4 i5 sizeFx_SC sizeFy_SC
    
    elseif (save_videos==2)
    if Smask_err==0
        scfMRItb_04_resplitData(base_dir_sub, fname, '_Smask', siz3) % if slice-wise data doesn't exist but combined 4D data exists, then split the data to generate the needed slice-wise data
        scfMRItb_04_unzipFile(base_dir_sub, fname, ['_slice' num2str(i3) '_Smask'])
        Smask1 = load_untouch_nii([base_dir_sub fname '_slice' num2str(i3) '_Smask.nii']); Smask1 = Smask1.img;
    else
        Smask1 = Smask1_(:,:,i3);
    end
    Smask1 = rot90(Smask1);
    [xmask,ymask] = find(Smask1==1);
    minx=round(min(xmask)-0.1*(size(Smask1,1))); maxx=round(max(xmask)+0.1*(size(Smask1,1)));
    miny=round(min(ymask)-0.1*(size(Smask1,2))); maxy=round(max(ymask)+0.1*(size(Smask1,2)));
    if (maxx-minx)<(maxy-miny)
        factr = ((maxy-miny)-(maxx-minx))/2;
        if factr==round(factr)
            maxx=maxx+factr; minx=minx-factr;
        else
            maxx=maxx+factr+0.5; minx=minx-factr+0.5;
        end
    elseif (maxy-miny)<(maxx-minx)
        factr = ((maxx-minx)-(maxy-miny))/2;
        if factr==round(factr)
            maxy=maxy+factr; miny=miny-factr;
        else
            maxy=maxy+factr+0.5; miny=miny-factr+0.5;
        end
    end
    scfMRItb_04_resplitData(base_dir_sub, fname, '', siz3) % if slice-wise data doesn't exist but combined 4D data exists, then split the data to generate the needed slice-wise data
    F3 = load_untouch_nii([base_dir_sub fname '_slice' num2str(i3) '.nii']); F3 = rot90(single(squeeze(F3.img)));
    [sizeFx,sizeFy] = size(F3(:,:,1));
    F4 = F3(minx:maxx,miny:maxy,:);
    for i5=1:size(F3,3)
        F3(:,:,i5) = imresize(F4(:,:,i5),[sizeFx sizeFy]);
    end
    clear maxx maxy minx miny xmask ymask Smask1 factr sizeFx sizeFy F2 F4 i5 sizeFx_SC sizeFy_SC        
    end
    
    for ii=1:size(F1,3) % loop across the number of images
        I = F1(:,:,ii); %read the next image
        I(find(isnan(I)))=0;
        I = (I - min(I(:))) ./ (max(I(:)) - min(I(:)));
        I2 = F3(:,:,ii); %read the next image
        I2(find(isnan(I2)))=0;
        I2 = (I2 - min(I2(:))) ./ (max(I2(:)) - min(I2(:)));
        fighndl = figure('Position',[round((scrsz(3)-scrsz(4))/2) 1 round(0.75*scrsz(3)) round(0.75*scrsz(4))],'Color',[0.95,0.9,0.85],'InvertHardcopy','off','Visible','off'); % [0.85,0.90,0.85]
        if (save_videos==1)
        if stp==1
            subplot(1,2,1), imshow(I2, [0 max(I2(:))], 'InitialMagnification', 'fit'); title({['fMRI data after "11. func-anat-registration" (slice ' num2str(i3) ' of ' num2str(siz3) ')'];['Time point ' num2str(ii) ' of ' num2str(size(F1,3))]},'Color','k','FontSize',18);
        elseif stp==2
            subplot(1,2,1), imshow(I2, [0 max(I2(:))], 'InitialMagnification', 'fit'); title({['fMRI data after "13. denoise-4 (WM regression)" (slice ' num2str(i3) ' of ' num2str(siz3) ')'];['Time point ' num2str(ii) ' of ' num2str(size(F1,3))]},'Color','k','FontSize',18);
        elseif stp==3
            subplot(1,2,1), imshow(I2, [0 max(I2(:))], 'InitialMagnification', 'fit'); title({['fMRI data after "12. denoise-3 (CSF regression)" (slice ' num2str(i3) ' of ' num2str(siz3) ')'];['Time point ' num2str(ii) ' of ' num2str(size(F1,3))]},'Color','k','FontSize',18);
        elseif stp==4
            subplot(1,2,1), imshow(I2, [0 max(I2(:))], 'InitialMagnification', 'fit'); title({['fMRI data after "13. denoise-4 (WM regression)" (slice ' num2str(i3) ' of ' num2str(siz3) ')'];['Time point ' num2str(ii) ' of ' num2str(size(F1,3))]},'Color','k','FontSize',18);
        end
        elseif (save_videos==2)
            subplot(1,2,1), imshow(I2, [0 max(I2(:))], 'InitialMagnification', 'fit'); title({['Unprocessed raw fMRI data (slice ' num2str(i3) ' of ' num2str(siz3) ')'];['Time point ' num2str(ii) ' of ' num2str(size(F1,3))]},'Color','k','FontSize',18);
        end
        subplot(1,2,2), imshow(I, [0 max(I(:))], 'InitialMagnification', 'fit'); title({['fMRI data after "14. denoise-5 (COV regression)" (slice ' num2str(i3) ' of ' num2str(siz3) ')'];['Time point ' num2str(ii) ' of ' num2str(size(F1,3))]},'Color',[0,0.55,0.69],'FontSize',18);
        frame = getframe(gcf);
        writeVideo(video,frame); % write the image to file
        close(fighndl)
        clear I I2 frame
    end
    close(video); %close the file
    clear F1 F3 video ii TR
end; clear i3
clear maxx_SC maxy_SC minx_SC miny_SC F
end

end

%% scfMRItb_15_filterData
if choices_preproc(15)==1
    try
clear QCpath save_videos fighndl stp
if ~(exist([base_dir_sub 'QC' '/15_temporal-filtering'],'dir'))
    mkdir([base_dir_sub 'QC' '/15_temporal-filtering'])
end
if Rns==0
    QCpath = [base_dir_sub 'QC' '/15_temporal-filtering/'];
else
    if ~(exist([base_dir_sub 'QC' '/15_temporal-filtering' '/run' num2str(Rns)],'dir'))
        mkdir([base_dir_sub 'QC' '/15_temporal-filtering' '/run' num2str(Rns)])
    end
    QCpath = [base_dir_sub 'QC' '/15_temporal-filtering' '/run' num2str(Rns) '/'];
end

f_BPF_max_vec = bpf_hz(1); f_BPF_max = bpf_hz(1);
if f_BPF_max_vec<0.1
    f_BPF_max_str_vec = sprintf('0%.0f',f_BPF_max_vec*100); f_BPF_max_str=f_BPF_max_str_vec; %#ok<*AGROW>
else
    f_BPF_max_str_vec = sprintf('%.0f',f_BPF_max_vec*100); f_BPF_max_str=f_BPF_max_str_vec; 
end

try
    scfMRItb_04_unzipFile(base_dir_sub, fname_anat, '2_mask_GM')
    scfMRItb_04_unzipFile(base_dir_sub, fname_anat, '2_mask_WM')
    mask_GM2 = load_untouch_nii([base_dir_sub fname_anat '2_mask_GM.nii']); mask_GM2 = mask_GM2.img;
    mask_WM2 = load_untouch_nii([base_dir_sub fname_anat '2_mask_WM.nii']); mask_WM2 = mask_WM2.img;
catch
    scfMRItb_04_unzipFile(base_dir_sub, fname_anat, '')
    A = load_untouch_nii([base_dir_sub fname_anat '.nii']);
    scfMRItb_04_unzipFile(base_dir_sub, fname_anat, '_mask_GM')
    scfMRItb_04_unzipFile(base_dir_sub, fname_anat, '_mask_WM')
    scfMRItb_04_unzipFile(base_dir_sub, fname_anat, '_mask_CSF')
    mmask1 = load_untouch_nii([base_dir_sub fname_anat '_mask_GM.nii']); % gray matter
    mmask2 = load_untouch_nii([base_dir_sub fname_anat '_mask_WM.nii']); % white matter
    mmask3 = load_untouch_nii([base_dir_sub fname_anat '_mask_CSF.nii']); % white matter
    mmask_all = single(or(or(mmask1.img, mmask2.img), mmask3.img));
    smmask = (mean(mmask_all,3) > 0);
    [s1,s2] = find(smmask~=0);
    s1median = median(s1); s2median = median(s2);
    s1min = min(s1); s2min = min(s2);
    s1max = max(s1); s2max = max(s2);
    smaxrange = max([s2max-s2min,s1max-s1min]);
    if round(size(mmask_all,1)/4) > smaxrange
        new_size = round(size(mmask_all,1)/4); % 128 - assume even for now
    elseif round(size(mmask_all,1)/3) > smaxrange
        new_size = round(size(mmask_all,1)/3); % 171
    elseif round(size(mmask_all,1)/2) > smaxrange
        new_size = round(size(mmask_all,1)/2); % 256
    else
        new_size = size(mmask_all,1); % 512
    end
    try if fullsize==1, new_size = size(mmask_all,1); end, catch, end    
    xmin = s1median - round(new_size/2) + 1; xmax = s1median + round(new_size/2);
    ymin = s2median - round(new_size/2) + 1; ymax = s2median + round(new_size/2);
    mask_GM_reduced = mmask1; mask_WM_reduced = mmask2; mask_CSF_reduced = mmask3;
    mask_GM_reduced.hdr.dime.dim(2)  = new_size; mask_GM_reduced.hdr.dime.dim(3)  = new_size;
    mask_WM_reduced.hdr.dime.dim(2)  = new_size; mask_WM_reduced.hdr.dime.dim(3)  = new_size;
    mask_CSF_reduced.hdr.dime.dim(2) = new_size; mask_CSF_reduced.hdr.dime.dim(3) = new_size;
    
    basestr = strfind(base_dir_sub,'/');
    basestr = base_dir_sub(basestr(end-1)+1:end); % name of subject folder
    if str2num(basestr(1:4)) <= 2017 %#ok<ST2NM> % if data were acquired prior to 2018 then don't change the bounding box of the reduced size images that were already processed before
        mask_GM_reduced.img  = mmask1.img(round((size(A.img,1)-new_size)/2)+1:end-round((size(A.img,1)-new_size)/2), ...
            round((size(A.img,1)-new_size)/2)+1:end-round((size(A.img,1)-new_size)/2), :);
        mask_WM_reduced.img  = mmask2.img(round((size(A.img,1)-new_size)/2)+1:end-round((size(A.img,1)-new_size)/2), ...
            round((size(A.img,1)-new_size)/2)+1:end-round((size(A.img,1)-new_size)/2), :);
        mask_CSF_reduced.img = mmask3.img(round((size(A.img,1)-new_size)/2)+1:end-round((size(A.img,1)-new_size)/2), ...
            round((size(A.img,1)-new_size)/2)+1:end-round((size(A.img,1)-new_size)/2), :); %#ok<*NASGU>
    else % else if data were acquired after 2017 then use the new method to determine the bounding box by centering the new box around the centroid of the spinal cord mask
        mask_GM_reduced.img  = mmask1.img(xmin:xmax, ymin:ymax, :);
        mask_WM_reduced.img  = mmask2.img(xmin:xmax, ymin:ymax, :);
        mask_CSF_reduced.img = mmask3.img(xmin:xmax, ymin:ymax, :);
    end    
    mask_SC_reduced = mask_GM_reduced;
    mask_SC_reduced.img = or(mask_GM_reduced.img, mask_WM_reduced.img);
    if ~exist([base_dir_sub fname_anat '2' '_mask_GM.nii'],'file')
        save_untouch_nii(mask_GM_reduced,  [base_dir_sub fname_anat '2' '_mask_GM.nii']); % gray matter
    end
    if ~exist([base_dir_sub fname_anat '2' '_mask_WM.nii'],'file')
        save_untouch_nii(mask_WM_reduced,  [base_dir_sub fname_anat '2' '_mask_WM.nii']); % white matter
    end
    if ~exist([base_dir_sub fname_anat '2' '_mask_CSF.nii'],'file')
        save_untouch_nii(mask_CSF_reduced, [base_dir_sub fname_anat '2' '_mask_CSF.nii']); % white matter
    end
    if ~exist([base_dir_sub fname_anat '2' '_mask_SC.nii'],'file')
        save_untouch_nii(mask_SC_reduced,  [base_dir_sub fname_anat '2' '_mask_SC.nii']); % gray matter
    end
    if ~exist([base_dir_sub fname_anat '2' '_all_masks.mat'],'file')
        save([base_dir_sub fname_anat '2' '_all_masks.mat'],'mask_GM_reduced','mask_WM_reduced','mask_CSF_reduced','mask_SC_reduced')
    end
    if ~exist([base_dir_sub fname_anat '2' '.nii'],'file')
        anat_reduced2 = A; anat_reduced2.hdr.dime.dim(2) = new_size; anat_reduced2.hdr.dime.dim(3) = new_size;
        anat_reduced2.img = A.img(xmin:xmax, ymin:ymax, :);
        save_untouch_nii(anat_reduced2, [base_dir_sub fname_anat '2' '.nii'])
    end
    mask_GM2 = mask_GM_reduced; mask_WM2 = mask_WM_reduced; mask_CSF2 = mask_CSF_reduced;
    clear mmask1 mmask2 mmask3 mmask_all smmask s1 s2 s1median s2median s1min s1max s2min s2max smaxrange new_size xmin xmax ymin ymax mask_GM_reduced mask_WM_reduced mask_CSF_reduced basestr mask_SC_reduced anat_reduced2 A
end

if exist([base_dir_sub fname '_warped' '.nii'], 'file') || exist([base_dir_sub fname '_warped' '.nii.gz'], 'file') % if none of CSF, WM or COV regress steps were performed
    whichip=8;
end
if exist([base_dir_sub fname '_WM' '.nii'], 'file') || exist([base_dir_sub fname '_WM' '.nii.gz'], 'file') % if only WM regress step was performed
    whichip=7;
end
if exist([base_dir_sub fname '_denoised' cutoff_str '.nii'], 'file') || exist([base_dir_sub fname '_denoised' cutoff_str '.nii.gz'], 'file') % if only CSF regress step was performed
    whichip=6;
end
if exist([base_dir_sub fname '_cov' '.nii'], 'file') || exist([base_dir_sub fname '_cov' '.nii.gz'], 'file') % if only COV regress step was performed
    whichip=5;
end
if exist([base_dir_sub fname '_WMcov' '.nii'], 'file') || exist([base_dir_sub fname '_WMcov' '.nii.gz'], 'file') % if both WM and COV regress steps were performed
    whichip=4;
end
if exist([base_dir_sub fname '_denoised' cutoff_str 'WM' '.nii'], 'file') || exist([base_dir_sub fname '_denoised' cutoff_str 'WM' '.nii.gz'], 'file') % if both CSF and WM regress steps were performed
    whichip=3;
end
if exist([base_dir_sub fname '_denoised' cutoff_str 'cov' '.nii'], 'file') || exist([base_dir_sub fname '_denoised' cutoff_str 'cov' '.nii.gz'], 'file') % if both CSF and COV regress steps were performed
    whichip=2;
end
if exist([base_dir_sub fname '_denoised' cutoff_str 'WMcov' '.nii'], 'file') || exist([base_dir_sub fname '_denoised' cutoff_str 'WMcov' '.nii.gz'], 'file') % if CSF, WM and COV regress steps were performed
    whichip=1;
end

if whichip==1
    load([base_dir_sub fname '_denoised' cutoff_str 'WMcov' '_BPF' f_BPF_max_str '.mat']); F1 = F; clear F %#ok<*LOAD>
    stp=8;
elseif whichip==2
    load([base_dir_sub fname '_denoised' cutoff_str 'cov' '_BPF' f_BPF_max_str '.mat']); F1 = F; clear F
    stp=7;
elseif whichip==3
    load([base_dir_sub fname '_denoised' cutoff_str 'WM' '_BPF' f_BPF_max_str '.mat']); F1 = F; clear F
    stp=6;
elseif whichip==4
    load([base_dir_sub fname '_WMcov' '_BPF' f_BPF_max_str '.mat']); F1 = F; clear F
    stp=5;
elseif whichip==5
    load([base_dir_sub fname '_cov' '_BPF' f_BPF_max_str '.mat']); F1 = F; clear F
    stp=4;
elseif whichip==6
    load([base_dir_sub fname '_denoised' cutoff_str '_BPF' f_BPF_max_str '.mat']); F1 = F; clear F
    stp=3;
elseif whichip==7
    load([base_dir_sub fname '_WM' '_BPF' f_BPF_max_str '.mat']); F1 = F; clear F
    stp=2;
elseif whichip==8
    load([base_dir_sub fname '_warped' '_BPF' f_BPF_max_str '.mat']); F1 = F; clear F
    stp=1;
end
TR = F1.hdr.dime.pixdim(5); F1 = single(squeeze(F1.img));


F2g_=[]; mF2g_=[]; F2w_=[]; mF2w_=[]; Cg_=[]; Cw_=[]; F4_=[]; mF4_=[]; C2_=[];
if stp~=0, F6g_=[]; mF6g_=[]; F6w_=[]; mF6w_=[]; C3g_=[]; C3w_=[]; end
for i3 = 1 : siz3
    try waitbar((i3/siz3),wbar3,sprintf('15. QC plots: slice (%d) of (%d)',i3,siz3)); catch, end
    fprintf('15. QC plots: slice (%d) of (%d), subject (%d) of (%d), run (%d) of (%d)\n',i3,siz3,k,size(func_files,1),k2,size(func_files,2))
    
    if Smask_err==0
        scfMRItb_04_resplitData(base_dir_sub, fname, '_Smask', siz3) % if slice-wise data doesn't exist but combined 4D data exists, then split the data to generate the needed slice-wise data
        scfMRItb_04_unzipFile(base_dir_sub, fname, ['_slice' num2str(i3) '_Smask'])
        Smask1 = load_untouch_nii([base_dir_sub fname '_slice' num2str(i3) '_Smask.nii']); Smask1 = Smask1.img;
    else
        Smask1 = Smask1_(:,:,i3);
    end

    for i5=1:size(F1,4)
        tmp = F1(:,:,i3,i5);
        F2g(:,i5) = tmp(find(mask_GM2(:,:,i3)==1));
        F2w(:,i5) = tmp(find(mask_WM2(:,:,i3)==1));
        clear tmp
    end; clear i5
    [~,svar1] = sort(var(F2g,0,2)); [~,sdist1] = sort(svar1); sortorder1{i3,1} = sdist1;
    [~,svar2] = sort(var(F2w,0,2)); [~,sdist2] = sort(svar2); sortorder2{i3,1} = sdist2;
    
    fighndl = figure('Position',[1 1 1152 720],'Color',[0.95,0.9,0.85],'InvertHardcopy','off','Visible','off'); % red [0.95,0.9,0.85], green [0.85,0.90,0.85], blue [0.89,0.94,0.99]
    subplot2n(2,1,1), imagesc(F2g(sortorder1{i3},:)),colormap(cmap),colorbar, xlabel('time (in samples)'),ylabel('Gray matter voxels (sorted by variance)'),
        title(['Time series within the GRAY MATTER after "15-bandpass-filter" in slice ' num2str(i3) ' of ' num2str(siz3)],'Color',[0,0.55,0.69],'FontSize',14);
    subplot2n(2,1,2), imagesc(F2w(sortorder2{i3},:)),colormap(cmap),colorbar, xlabel('time (in samples)'),ylabel('White matter voxels (sorted by variance)'),
        title(['Time series within the WHITE MATTER after "15-bandpass-filter" in slice ' num2str(i3) ' of ' num2str(siz3)],'Color',[0.65,0.11,0.19],'FontSize',14);
    saveas(fighndl,[QCpath 'fMRI_ts_15filter_slice' sprintf('%.2d',i3) '.jpg'])
    close(fighndl)
    
    scfMRItb_04_resplitData(base_dir_sub, fname, '', siz3) % if slice-wise data doesn't exist but combined 4D data exists, then split the data to generate the needed slice-wise data
	scfMRItb_04_unzipFile(base_dir_sub, fname, ['_slice' num2str(i3)])
    F3 = load_untouch_nii([base_dir_sub fname '_slice' num2str(i3) '.nii']); F3 = single(squeeze(F3.img));
    for i5=1:size(F3,3)
        tmp = F3(:,:,i5);
        F4(:,i5) = tmp(find(Smask1==1)); clear tmp
    end; clear i5 F3
    mF2g=mean(F2g,1); mF2w=mean(F2w,1);
    mF2g=zscore(mF2g); mF2w=zscore(mF2w);

    Cg = corrcoef(F2g'); Cg = uppertriangle(Cg);
    Cw = corrcoef(F2w'); Cw = uppertriangle(Cw);
    C2 = corrcoef(F4');  C2 = uppertriangle(C2);
    fighndl = figure('Position',[1 1 1152 720],'Color',[0.95,0.9,0.85],'InvertHardcopy','off','Visible','off'); % red [0.95,0.9,0.85], green [0.85,0.90,0.85], blue [0.89,0.94,0.99]
    subplot2n(3,1,1), histogram(C2,'EdgeColor','none','FaceColor','k','FaceAlpha',0.5,'Normalization','probability'); grid on,grid minor, xlim([-1 1]), hold on, xline(0,'Color','k','linewidth',1.5); hold off
      title(sprintf('Histogram of voxel-to-voxel correlations in UNPROCESSED RAW DATA (slice %d of %d)\n%.1f%% of the correlations were > 0  [ mean=%.2f, median=%.2f, SD=%.2f ]',i3,siz3,100*length(find(C2>0))/length(C2),mean(C2),median(C2),std(C2)),'Color','k','FontSize',14)
    subplot2n(3,1,2), histogram(Cg,'EdgeColor','none','FaceColor',[0,0.55,0.69],'FaceAlpha',0.5,'Normalization','probability'); grid on,grid minor, xlim([-1 1]), hold on, xline(0,'Color','k','linewidth',1.5); hold off
      title(sprintf('Histogram of voxel-to-voxel GRAY MATTER correlations AFTER 15-bandpass-filter [0.01-%.2fHz] (slice %d of %d)\n%.1f%% of the correlations were > 0  [ mean=%.2f, median=%.2f, SD=%.2f ]',f_BPF_max_vec(1),i3,siz3,100*length(find(Cg>0))/length(Cg),mean(Cg),median(Cg),std(Cg)),'Color',[0,0.55,0.69],'FontSize',14)
    subplot2n(3,1,3), histogram(Cw,'EdgeColor','none','FaceColor',[0.65,0.11,0.19],'FaceAlpha',0.5,'Normalization','probability'); grid on,grid minor, xlim([-1 1]), hold on, xline(0,'Color','k','linewidth',1.5); hold off
      title(sprintf('Histogram of voxel-to-voxel WHITE MATTER correlations AFTER 15-bandpass-filter [0.01-%.2fHz] (slice %d of %d)\n%.1f%% of the correlations were > 0  [ mean=%.2f, median=%.2f, SD=%.2f ]',f_BPF_max_vec(1),i3,siz3,100*length(find(Cw>0))/length(Cw),mean(Cw),median(Cw),std(Cw)),'Color',[0.65,0.11,0.19],'FontSize',14)
    saveas(fighndl,[QCpath 'hist_FC_compare_15filter_slice' sprintf('%.2d',i3) '.jpg'])
    close(fighndl)
    
    if stp~=0
        if stp==1
            scfMRItb_04_resplitData(base_dir_sub, fname, '_warped', siz3) % if slice-wise data doesn't exist but combined 4D data exists, then split the data to generate the needed slice-wise data
            scfMRItb_04_unzipFile(base_dir_sub, fname, ['_slice' num2str(i3) '_warped'])
            F5 = load_untouch_nii([base_dir_sub fname '_slice' num2str(i3) '_warped.nii']); F5 = single(squeeze(F5.img));
        elseif stp==2
            scfMRItb_04_resplitData(base_dir_sub, fname, '_WM', siz3) % if slice-wise data doesn't exist but combined 4D data exists, then split the data to generate the needed slice-wise data
            scfMRItb_04_unzipFile(base_dir_sub, fname, ['_slice' num2str(i3) '_WM'])
            F5 = load_untouch_nii([base_dir_sub fname '_slice' num2str(i3) '_WM.nii']); F5 = single(squeeze(F5.img));
        elseif stp==3
            scfMRItb_04_resplitData(base_dir_sub, fname, ['_denoised' cutoff_str], siz3) % if slice-wise data doesn't exist but combined 4D data exists, then split the data to generate the needed slice-wise data
            scfMRItb_04_unzipFile(base_dir_sub, fname, ['_slice' num2str(i3) '_denoised' cutoff_str ''])
            F5 = load_untouch_nii([base_dir_sub fname '_slice' num2str(i3) '_denoised' cutoff_str '.nii']); F5 = single(squeeze(F5.img));
        elseif stp==4
            scfMRItb_04_resplitData(base_dir_sub, fname, '_cov', siz3) % if slice-wise data doesn't exist but combined 4D data exists, then split the data to generate the needed slice-wise data
            scfMRItb_04_unzipFile(base_dir_sub, fname, ['_slice' num2str(i3) '_cov'])
            F5 = load_untouch_nii([base_dir_sub fname '_slice' num2str(i3) '_cov.nii']); F5 = single(squeeze(F5.img));
        elseif stp==5
            scfMRItb_04_resplitData(base_dir_sub, fname, '_WMcov', siz3) % if slice-wise data doesn't exist but combined 4D data exists, then split the data to generate the needed slice-wise data
            scfMRItb_04_unzipFile(base_dir_sub, fname, ['_slice' num2str(i3) '_WMcov'])
            F5 = load_untouch_nii([base_dir_sub fname '_slice' num2str(i3) '_WMcov.nii']); F5 = single(squeeze(F5.img));
        elseif stp==6
            scfMRItb_04_resplitData(base_dir_sub, fname, ['_denoised' cutoff_str 'WM'], siz3) % if slice-wise data doesn't exist but combined 4D data exists, then split the data to generate the needed slice-wise data
            scfMRItb_04_unzipFile(base_dir_sub, fname, ['_slice' num2str(i3) '_denoised' cutoff_str 'WM'])
            F5 = load_untouch_nii([base_dir_sub fname '_slice' num2str(i3) '_denoised' cutoff_str 'WM.nii']); F5 = single(squeeze(F5.img));
        elseif stp==7
            scfMRItb_04_resplitData(base_dir_sub, fname, ['_denoised' cutoff_str 'cov'], siz3) % if slice-wise data doesn't exist but combined 4D data exists, then split the data to generate the needed slice-wise data
            scfMRItb_04_unzipFile(base_dir_sub, fname, ['_slice' num2str(i3) '_denoised' cutoff_str 'cov'])
            F5 = load_untouch_nii([base_dir_sub fname '_slice' num2str(i3) '_denoised' cutoff_str 'cov.nii']); F5 = single(squeeze(F5.img));
        elseif stp==8
            scfMRItb_04_resplitData(base_dir_sub, fname, ['_denoised' cutoff_str 'WMcov'], siz3) % if slice-wise data doesn't exist but combined 4D data exists, then split the data to generate the needed slice-wise data
            scfMRItb_04_unzipFile(base_dir_sub, fname, ['_slice' num2str(i3) '_denoised' cutoff_str 'WMcov'])
            F5 = load_untouch_nii([base_dir_sub fname '_slice' num2str(i3) '_denoised' cutoff_str 'WMcov.nii']); F5 = single(squeeze(F5.img));
        end
        for i5=1:size(F5,3)
            tmp = F5(:,:,i5);
            F6g(:,i5) = tmp(find(mask_GM(:,:,i3)==1));
            F6w(:,i5) = tmp(find(mask_WM(:,:,i3)==1));
            clear tmp
        end; clear i5 F5
        mF6g=mean(F6g,1); mF6w=mean(F6w,1); 
        mF6g=zscore(mF6g); mF6w=zscore(mF6w);
        fighndl = figure('Position',[1 1 1152 720],'Color',[0.95,0.9,0.85],'InvertHardcopy','off','Visible','off'); % red [0.95,0.9,0.85], green [0.85,0.90,0.85], blue [0.89,0.94,0.99]
        if stp==1
            subplot2n(4,1,1), plot(mF6g,'color','k','linewidth',1.5), hold on, plot(mF2g,'color',[0,0.55,0.69],'linewidth',0.2), ylabel('mean time series'),
                grid on,grid minor, xlim([1 length(mF2g)]),ylim([min(1.25.*[mF2g,mF6g]) 1.25.*max([mF2g,mF6g])]), title(['Unfiltered: normalized mean GRAY MATTER signal AFTER "11-func-anat-registration" (slice ' num2str(i3) ' of ' num2str(siz3) ')'],'Color','k','FontSize',14), legend({'Gray matter signal before filtering'},'FontSize',12,'TextColor','k','Location','best')
            subplot2n(4,1,3), plot(mF6w,'color','k','linewidth',1.5), hold on, plot(mF2w,'color',[0.65,0.11,0.19],'linewidth',0.2), ylabel('mean time series'),
                grid on,grid minor, xlim([1 length(mF2g)]),ylim([min(1.25.*[mF2w,mF6w]) 1.25.*max([mF2w,mF6w])]), title(['Unfiltered: normalized mean WHITE MATTER signal AFTER "11-func-anat-registration" (slice ' num2str(i3) ' of ' num2str(siz3) ')'],'Color','k','FontSize',14), legend({'White matter signal before filtering'},'FontSize',12,'TextColor','k','Location','best')
        elseif stp==3
            subplot2n(4,1,1), plot(mF6g,'color','k','linewidth',1.5), hold on, plot(mF2g,'color',[0,0.55,0.69],'linewidth',0.2), ylabel('mean time series'),
                grid on,grid minor, xlim([1 length(mF2g)]),ylim([min(1.25.*[mF2g,mF6g]) 1.25.*max([mF2g,mF6g])]), title(['Unfiltered: normalized mean GRAY MATTER signal AFTER "12-denoise3-CSFregress" (slice ' num2str(i3) ' of ' num2str(siz3) ')'],'Color','k','FontSize',14), legend({'Gray matter signal before filtering'},'FontSize',12,'TextColor','k','Location','best')
            subplot2n(4,1,3), plot(mF6w,'color','k','linewidth',1.5), hold on, plot(mF2w,'color',[0.65,0.11,0.19],'linewidth',0.2), ylabel('mean time series'),
                grid on,grid minor, xlim([1 length(mF2g)]),ylim([min(1.25.*[mF2w,mF6w]) 1.25.*max([mF2w,mF6w])]), title(['Unfiltered: normalized mean WHITE MATTER signal AFTER "12-denoise3-CSFregress" (slice ' num2str(i3) ' of ' num2str(siz3) ')'],'Color','k','FontSize',14), legend({'White matter signal before filtering'},'FontSize',12,'TextColor','k','Location','best')
        elseif (stp==2)||(stp==6)
            subplot2n(4,1,1), plot(mF6g,'color','k','linewidth',1.5), hold on, plot(mF2g,'color',[0,0.55,0.69],'linewidth',0.2), ylabel('mean time series'),
                grid on,grid minor, xlim([1 length(mF2g)]),ylim([min(1.25.*[mF2g,mF6g]) 1.25.*max([mF2g,mF6g])]), title(['Unfiltered: normalized mean GRAY MATTER signal AFTER "13-denoise4-WMregress" (slice ' num2str(i3) ' of ' num2str(siz3) ')'],'Color','k','FontSize',14), legend({'Gray matter signal before filtering'},'FontSize',12,'TextColor','k','Location','best')
            subplot2n(4,1,3), plot(mF6w,'color','k','linewidth',1.5), hold on, plot(mF2w,'color',[0.65,0.11,0.19],'linewidth',0.2), ylabel('mean time series'),
                grid on,grid minor, xlim([1 length(mF2g)]),ylim([min(1.25.*[mF2w,mF6w]) 1.25.*max([mF2w,mF6w])]), title(['Unfiltered: normalized mean WHITE MATTER signal AFTER "13-denoise4-WMregress" (slice ' num2str(i3) ' of ' num2str(siz3) ')'],'Color','k','FontSize',14), legend({'White matter signal before filtering'},'FontSize',12,'TextColor','k','Location','best')
        elseif (stp==4)||(stp==5)||(stp==7)||(stp==8)
            subplot2n(4,1,1), plot(mF6g,'color','k','linewidth',1.5), hold on, plot(mF2g,'color',[0,0.55,0.69],'linewidth',0.2), ylabel('mean time series'),
                grid on,grid minor, xlim([1 length(mF2g)]),ylim([min(1.25.*[mF2g,mF6g]) 1.25.*max([mF2g,mF6g])]), title(['Unfiltered: normalized mean GRAY MATTER signal AFTER "14-denoise5-COVregress" (slice ' num2str(i3) ' of ' num2str(siz3) ')'],'Color','k','FontSize',14), legend({'Gray matter signal before filtering'},'FontSize',12,'TextColor','k','Location','best')
            subplot2n(4,1,3), plot(mF6w,'color','k','linewidth',1.5), hold on, plot(mF2w,'color',[0.65,0.11,0.19],'linewidth',0.2), ylabel('mean time series'),
                grid on,grid minor, xlim([1 length(mF2g)]),ylim([min(1.25.*[mF2w,mF6w]) 1.25.*max([mF2w,mF6w])]), title(['Unfiltered: normalized mean WHITE MATTER signal AFTER "14-denoise5-COVregress" (slice ' num2str(i3) ' of ' num2str(siz3) ')'],'Color','k','FontSize',14), legend({'White matter signal before filtering'},'FontSize',12,'TextColor','k','Location','best')
        end
        subplot2n(4,1,2), plot(mF2g,'color',[0,0.55,0.69],'linewidth',1.5), hold on, plot(mF6g,'color','k','linewidth',0.2), ylabel('mean time series'),
            grid on,grid minor, xlim([1 length(mF2g)]),ylim([min(1.25.*[mF2g,mF6g]) 1.25.*max([mF2g,mF6g])]), title(['Bandpass filtered: normalized mean GRAY MATTER signal AFTER "15-bandpass-filter" [0.01-' num2str(f_BPF_max_vec(1)) 'Hz] (slice ' num2str(i3) ' of ' num2str(siz3) ')'],'Color',[0,0.55,0.69],'FontSize',14), legend({'Gray matter signal after filtering'},'FontSize',12,'TextColor',[0,0.55,0.69],'Location','best')
        subplot2n(4,1,4), plot(mF2w,'color',[0.65,0.11,0.19],'linewidth',1.5), hold on, plot(mF6w,'color','k','linewidth',0.2), ylabel('mean time series'),
            grid on,grid minor, xlim([1 length(mF2g)]),ylim([min(1.25.*[mF2w,mF6w]) 1.25.*max([mF2w,mF6w])]), title(['Bandpass filtered: normalized mean WHITE MATTER signal AFTER "15-bandpass-filter" [0.01-' num2str(f_BPF_max_vec(1)) 'Hz] (slice ' num2str(i3) ' of ' num2str(siz3) ')'],'Color',[0.65,0.11,0.19],'FontSize',14), legend({'White matter signal after filtering'},'FontSize',12,'TextColor',[0.65,0.11,0.19],'Location','best')
        saveas(fighndl,[QCpath 'mean_timeseries_14denoise5_vs_15filter_slice' sprintf('%.2d',i3) '.jpg'])
        close(fighndl)
        
        C3g = corrcoef(F6g'); C3g = uppertriangle(C3g); C3w = corrcoef(F6w'); C3w = uppertriangle(C3w);
        fighndl = figure('Position',[1 1 1152 720],'Color',[0.95,0.9,0.85],'InvertHardcopy','off','Visible','off'); % red [0.95,0.9,0.85], green [0.85,0.90,0.85], blue [0.89,0.94,0.99]
        if stp==1
            subplot2n(4,1,1), histogram(C3g,'EdgeColor','none','FaceColor','k','FaceAlpha',0.5,'Normalization','probability'); grid on,grid minor, xlim([-1 1]), hold on, xline(0,'Color','k','linewidth',1.5); hold off
                title(sprintf('Histogram of voxel-to-voxel GRAY MATTER correlations AFTER 11-func-anat-registration (slice %d of %d)\n%.1f%% of the correlations were > 0  [ mean=%.2f, median=%.2f, SD=%.2f ]',i3,siz3,100*length(find(C3g>0))/length(C3g),mean(C3g),median(C3g),std(C3g)),'Color','k','FontSize',14)
            subplot2n(4,1,3), histogram(C3w,'EdgeColor','none','FaceColor','k','FaceAlpha',0.5,'Normalization','probability'); grid on,grid minor, xlim([-1 1]), hold on, xline(0,'Color','k','linewidth',1.5); hold off
                title(sprintf('WHITE MATTER: AFTER 11-func-anat-registration (slice %d of %d): %.1f%% of the correlations were > 0  [ mean=%.2f, median=%.2f, SD=%.2f ]',i3,siz3,100*length(find(C3w>0))/length(C3w),mean(C3w),median(C3w),std(C3w)),'Color','k','FontSize',14)
        elseif stp==3
            subplot2n(4,1,1), histogram(C3g,'EdgeColor','none','FaceColor','k','FaceAlpha',0.5,'Normalization','probability'); grid on,grid minor, xlim([-1 1]), hold on, xline(0,'Color','k','linewidth',1.5); hold off
                title(sprintf('Histogram of voxel-to-voxel GRAY MATTER correlations AFTER 12-denoise3-CSFregress (slice %d of %d)\n%.1f%% of the correlations were > 0  [ mean=%.2f, median=%.2f, SD=%.2f ]',i3,siz3,100*length(find(C3g>0))/length(C3g),mean(C3g),median(C3g),std(C3g)),'Color','k','FontSize',14)
            subplot2n(4,1,3), histogram(C3w,'EdgeColor','none','FaceColor','k','FaceAlpha',0.5,'Normalization','probability'); grid on,grid minor, xlim([-1 1]), hold on, xline(0,'Color','k','linewidth',1.5); hold off
                title(sprintf('WHITE MATTER: AFTER 12-denoise3-CSFregress (slice %d of %d): %.1f%% of the correlations were > 0  [ mean=%.2f, median=%.2f, SD=%.2f ]',i3,siz3,100*length(find(C3w>0))/length(C3w),mean(C3w),median(C3w),std(C3w)),'Color','k','FontSize',14)
        elseif (stp==2)||(stp==6)
            subplot2n(4,1,1), histogram(C3g,'EdgeColor','none','FaceColor','k','FaceAlpha',0.5,'Normalization','probability'); grid on,grid minor, xlim([-1 1]), hold on, xline(0,'Color','k','linewidth',1.5); hold off
                title(sprintf('Histogram of voxel-to-voxel GRAY MATTER correlations AFTER 13-denoise4-WMregress (slice %d of %d)\n%.1f%% of the correlations were > 0  [ mean=%.2f, median=%.2f, SD=%.2f ]',i3,siz3,100*length(find(C3g>0))/length(C3g),mean(C3g),median(C3g),std(C3g)),'Color','k','FontSize',14)
            subplot2n(4,1,3), histogram(C3w,'EdgeColor','none','FaceColor','k','FaceAlpha',0.5,'Normalization','probability'); grid on,grid minor, xlim([-1 1]), hold on, xline(0,'Color','k','linewidth',1.5); hold off
                title(sprintf('WHITE MATTER: AFTER 13-denoise4-WMregress (slice %d of %d): %.1f%% of the correlations were > 0  [ mean=%.2f, median=%.2f, SD=%.2f ]',i3,siz3,100*length(find(C3w>0))/length(C3w),mean(C3w),median(C3w),std(C3w)),'Color','k','FontSize',14)
        elseif (stp==4)||(stp==5)||(stp==7)||(stp==8)
            subplot2n(4,1,1), histogram(C3g,'EdgeColor','none','FaceColor','k','FaceAlpha',0.5,'Normalization','probability'); grid on,grid minor, xlim([-1 1]), hold on, xline(0,'Color','k','linewidth',1.5); hold off
                title(sprintf('Histogram of voxel-to-voxel GRAY MATTER correlations AFTER 14-denoise5-COVregress (slice %d of %d)\n%.1f%% of the correlations were > 0  [ mean=%.2f, median=%.2f, SD=%.2f ]',i3,siz3,100*length(find(C3g>0))/length(C3g),mean(C3g),median(C3g),std(C3g)),'Color','k','FontSize',14)
            subplot2n(4,1,3), histogram(C3w,'EdgeColor','none','FaceColor','k','FaceAlpha',0.5,'Normalization','probability'); grid on,grid minor, xlim([-1 1]), hold on, xline(0,'Color','k','linewidth',1.5); hold off
                title(sprintf('WHITE MATTER: AFTER 14-denoise5-COVregress (slice %d of %d): %.1f%% of the correlations were > 0  [ mean=%.2f, median=%.2f, SD=%.2f ]',i3,siz3,100*length(find(C3w>0))/length(C3w),mean(C3w),median(C3w),std(C3w)),'Color','k','FontSize',14)
        end
        subplot2n(4,1,2), histogram(Cg,'EdgeColor','none','FaceColor',[0,0.55,0.69],'FaceAlpha',0.5,'Normalization','probability'); grid on,grid minor, xlim([-1 1]), hold on, xline(0,'Color','k','linewidth',1.5); hold off
            title(sprintf('GRAY MATTER: AFTER 15-bandpass-filter (slice %d of %d): %.1f%% of the correlations were > 0  [ mean=%.2f, median=%.2f, SD=%.2f ]',i3,siz3,100*length(find(Cg>0))/length(Cg),mean(Cg),median(Cg),std(Cg)),'Color',[0,0.55,0.69],'FontSize',14)
        subplot2n(4,1,4), histogram(Cw,'EdgeColor','none','FaceColor',[0.65,0.11,0.19],'FaceAlpha',0.5,'Normalization','probability'); grid on,grid minor, xlim([-1 1]), hold on, xline(0,'Color','k','linewidth',1.5); hold off
            title(sprintf('WHITE MATTER: AFTER 15-bandpass-filter (slice %d of %d): %.1f%% of the correlations were > 0  [ mean=%.2f, median=%.2f, SD=%.2f ]',i3,siz3,100*length(find(Cw>0))/length(Cw),mean(Cw),median(Cw),std(Cw)),'Color',[0.65,0.11,0.19],'FontSize',14)
        saveas(fighndl,[QCpath 'hist_FC_compare_14denoise5_15filter_slice' sprintf('%.2d',i3) '.jpg'])
        close(fighndl)
        F6g_=cat(1,F6g_,F6g); mF6g_=cat(1,mF6g_,mF6g); C3g_=cat(1,C3g_,C3g);
        F6w_=cat(1,F6w_,F6w); mF6w_=cat(1,mF6w_,mF6w); C3w_=cat(1,C3w_,C3w);
        clear F6g F6w mF6g mF6w C3g C3w
    end
    
    F2g_=cat(1,F2g_,F2g); mF2g_=cat(1,mF2g_,mF2g); F2w_=cat(1,F2w_,F2w); mF2w_=cat(1,mF2w_,mF2w); Cg_=cat(1,Cg_,Cg); Cw_=cat(1,Cw_,Cw); F4_=cat(1,F4_,F4); C2_=cat(1,C2_,C2);
    clear F2g F2w F4 TR Smask1 svar1 svar2 mF2g mF2w mF4 Cg Cw C2
end

[~,svar1] = sort(var(F2g_,0,2)); [~,spsc1] = sort(meanPSC(F2g_));
[~,svar2] = sort(var(F2w_,0,2)); [~,spsc2] = sort(meanPSC(F2w_));
sortorder1b = sortorder1; sortorder1=[]; f1=0; %#ok<*ASGLU>
sortorder2b = sortorder2; sortorder2=[]; f2=0; %#ok<*ASGLU>
for im=1:length(sortorder1b)
    temp = sortorder1b{im}; temp = temp + f1; f1 = f1 + length(temp);
    sortorder1 = cat(1,sortorder1,temp); clear temp
    temp = sortorder2b{im}; temp = temp + f2; f2 = f2 + length(temp);
    sortorder2 = cat(1,sortorder2,temp); clear temp
end; clear im f1 f2
sortorder1 = spsc1; % options: spsc for percentage signal change, svar for variance, (comment out the line if you want to use distance from the center of the cord separately for each slice)
sortorder2 = spsc2; % options: spsc for percentage signal change, svar for variance, (comment out the line if you want to use distance from the center of the cord separately for each slice)
for im=max(sortorder1):-1:1, if isempty(find(sortorder1==im)), sortorder1(find(sortorder1>im)) = sortorder1(find(sortorder1>im))-1; end, end; clear im %#ok<*EFIND>
for im=max(sortorder2):-1:1, if isempty(find(sortorder2==im)), sortorder2(find(sortorder2>im)) = sortorder2(find(sortorder2>im))-1; end, end; clear im %#ok<*EFIND>

fighndl = figure('Position',[1 1 1152 720],'Color',[0.89,0.94,0.99],'InvertHardcopy','off','Visible','off'); % red [0.95,0.9,0.85], green [0.85,0.90,0.85], blue [0.89,0.94,0.99]
subplot2n(2,1,1), imagesc(F2g_(sortorder1,:)),colormap(cmap),colorbar, xlabel('time (in samples)'),ylabel(sprintf('voxels (sorted by mean percentage signal change;\nbottom row is highest)')),
  title(['Time series within the GRAY MATTER AFTER 15-bandpass-filter (all slices combined)'],'Color',[0,0.55,0.69],'FontSize',14);
subplot2n(2,1,2), imagesc(F2w_(sortorder2,:)),colormap(cmap),colorbar, xlabel('time (in samples)'),ylabel(sprintf('voxels (sorted by mean percentage signal change;\nbottom row is highest)')),
  title(['Time series within the WHITE MATTER AFTER 15-bandpass-filter (all slices combined)'],'Color',[0.65,0.11,0.19],'FontSize',14);
saveas(fighndl,[QCpath 'fMRI_ts_15filter_allSlices' '.jpg'])
 close(fighndl)
mF2g_=mean(mF2g_,1); mF2w_=mean(mF2w_,1); mF4_=mean(mF4_,1);
fighndl = figure('Position',[1 1 1152 720],'Color',[0.89,0.94,0.99],'InvertHardcopy','off','Visible','off'); % red [0.95,0.9,0.85], green [0.85,0.90,0.85], blue [0.89,0.94,0.99]
subplot2n(3,1,1), histogram(C2_,'EdgeColor','none','FaceColor','k','FaceAlpha',0.5,'Normalization','probability'); grid on,grid minor, xlim([-1 1]), hold on, xline(0,'Color','k','linewidth',1.5); hold off
    title(sprintf('Histogram of voxel-to-voxel correlations in UNPROCESSED RAW DATA (all slices combined)\n%.1f%% of the correlations were > 0  [ mean=%.2f, median=%.2f, SD=%.2f ]',100*length(find(C2_>0))/length(C2_),mean(C2_),median(C2_),std(C2_)),'Color','k','FontSize',14)
subplot2n(3,1,2), histogram(Cg_,'EdgeColor','none','FaceColor',[0,0.55,0.69],'FaceAlpha',0.5,'Normalization','probability'); grid on,grid minor, xlim([-1 1]), hold on, xline(0,'Color','k','linewidth',1.5); hold off
    title(sprintf('Histogram of voxel-to-voxel GRAY MATTER correlations AFTER 15-bandpass-filter [0.01-%.2fHz] (all slices combined)\n%.1f%% of the correlations were > 0  [ mean=%.2f, median=%.2f, SD=%.2f ]',f_BPF_max_vec(1),100*length(find(Cg_>0))/length(Cg_),mean(Cg_),median(Cg_),std(Cg_)),'Color',[0,0.55,0.69],'FontSize',14)
subplot2n(3,1,3), histogram(Cw_,'EdgeColor','none','FaceColor',[0.65,0.11,0.19],'FaceAlpha',0.5,'Normalization','probability'); grid on,grid minor, xlim([-1 1]), hold on, xline(0,'Color','k','linewidth',1.5); hold off
    title(sprintf('Histogram of voxel-to-voxel WHITE MATTER correlations AFTER 15-bandpass-filter [0.01-%.2fHz] (all slices combined)\n%.1f%% of the correlations were > 0  [ mean=%.2f, median=%.2f, SD=%.2f ]',f_BPF_max_vec(1),100*length(find(Cw_>0))/length(Cw_),mean(Cw_),median(Cw_),std(Cw_)),'Color',[0.65,0.11,0.19],'FontSize',14)
saveas(fighndl,[QCpath 'hist_FC_compare_15filter_allSlices' '.jpg'])
 close(fighndl)

if stp~=0
    mF6g_=mean(mF6g_,1); mF6w_=mean(mF6w_,1);
    fighndl = figure('Position',[1 1 1152 720],'Color',[0.89,0.94,0.99],'InvertHardcopy','off','Visible','off'); % red [0.95,0.9,0.85], green [0.85,0.90,0.85], blue [0.89,0.94,0.99]
    if stp==1
    	subplot2n(4,1,1), plot(mF6g_,'color','k','linewidth',1.5), hold on, plot(mF2g_,'color',[0,0.55,0.69],'linewidth',0.2), ylabel('mean time series'),
            grid on,grid minor, xlim([1 length(mF2g_)]),ylim([1.25.*min([mF2g_,mF6g_]) 1.25.*max([mF2g_,mF6g_])]), title(['Unfiltered: normalized mean GRAY MATTER signal AFTER "11-func-anat-registration" (all slices averaged)'],'Color','k','FontSize',14), legend({'Gray matter signal before filtering'},'FontSize',12,'TextColor','k','Location','best')
        subplot2n(4,1,3), plot(mF6w_,'color','k','linewidth',1.5), hold on, plot(mF2w_,'color',[0.65,0.11,0.19],'linewidth',0.2), ylabel('mean time series'),
            grid on,grid minor, xlim([1 length(mF2g_)]),ylim([1.25.*min([mF2w_,mF6w_]) 1.25.*max([mF2w_,mF6w_])]), title(['Unfiltered: normalized mean WHITE MATTER signal AFTER "11-func-anat-registration" (all slices averaged)'],'Color','k','FontSize',14), legend({'White matter signal before filtering'},'FontSize',12,'TextColor','k','Location','best')
    elseif stp==3
    	subplot2n(4,1,1), plot(mF6g_,'color','k','linewidth',1.5), hold on, plot(mF2g_,'color',[0,0.55,0.69],'linewidth',0.2), ylabel('mean time series'),
            grid on,grid minor, xlim([1 length(mF2g_)]),ylim([1.25.*min([mF2g_,mF6g_]) 1.25.*max([mF2g_,mF6g_])]), title(['Unfiltered: normalized mean GRAY MATTER signal AFTER "12-DENOISE3-CSFregress" (all slices averaged)'],'Color','k','FontSize',14), legend({'Gray matter signal before filtering'},'FontSize',12,'TextColor','k','Location','best')
        subplot2n(4,1,3), plot(mF6w_,'color','k','linewidth',1.5), hold on, plot(mF2w_,'color',[0.65,0.11,0.19],'linewidth',0.2), ylabel('mean time series'),
            grid on,grid minor, xlim([1 length(mF2g_)]),ylim([1.25.*min([mF2w_,mF6w_]) 1.25.*max([mF2w_,mF6w_])]), title(['Unfiltered: normalized mean WHITE MATTER signal AFTER "12-DENOISE3-CSFregress" (all slices averaged)'],'Color','k','FontSize',14), legend({'White matter signal before filtering'},'FontSize',12,'TextColor','k','Location','best')
    elseif (stp==2)||(stp==6)
    	subplot2n(4,1,1), plot(mF6g_,'color','k','linewidth',1.5), hold on, plot(mF2g_,'color',[0,0.55,0.69],'linewidth',0.2), ylabel('mean time series'),
            grid on,grid minor, xlim([1 length(mF2g_)]),ylim([1.25.*min([mF2g_,mF6g_]) 1.25.*max([mF2g_,mF6g_])]), title(['Unfiltered: normalized mean GRAY MATTER signal AFTER "13-DENOISE4-WMregress" (all slices averaged)'],'Color','k','FontSize',14), legend({'Gray matter signal before filtering'},'FontSize',12,'TextColor','k','Location','best')
        subplot2n(4,1,3), plot(mF6w_,'color','k','linewidth',1.5), hold on, plot(mF2w_,'color',[0.65,0.11,0.19],'linewidth',0.2), ylabel('mean time series'),
            grid on,grid minor, xlim([1 length(mF2g_)]),ylim([1.25.*min([mF2w_,mF6w_]) 1.25.*max([mF2w_,mF6w_])]), title(['Unfiltered: normalized mean WHITE MATTER signal AFTER "13-DENOISE4-WMregress" (all slices averaged)'],'Color','k','FontSize',14), legend({'White matter signal before filtering'},'FontSize',12,'TextColor','k','Location','best')
    elseif (stp==4)||(stp==5)||(stp==7)||(stp==8)
    	subplot2n(4,1,1), plot(mF6g_,'color','k','linewidth',1.5), hold on, plot(mF2g_,'color',[0,0.55,0.69],'linewidth',0.2), ylabel('mean time series'),
            grid on,grid minor, xlim([1 length(mF2g_)]),ylim([1.25.*min([mF2g_,mF6g_]) 1.25.*max([mF2g_,mF6g_])]), title(['Unfiltered: normalized mean GRAY MATTER signal AFTER "14-DENOISE5-COVregress" (all slices averaged)'],'Color','k','FontSize',14), legend({'Gray matter signal before filtering'},'FontSize',12,'TextColor','k','Location','best')
        subplot2n(4,1,3), plot(mF6w_,'color','k','linewidth',1.5), hold on, plot(mF2w_,'color',[0.65,0.11,0.19],'linewidth',0.2), ylabel('mean time series'),
            grid on,grid minor, xlim([1 length(mF2g_)]),ylim([1.25.*min([mF2w_,mF6w_]) 1.25.*max([mF2w_,mF6w_])]), title(['Unfiltered: normalized mean WHITE MATTER signal AFTER "14-DENOISE5-COVregress" (all slices averaged)'],'Color','k','FontSize',14), legend({'White matter signal before filtering'},'FontSize',12,'TextColor','k','Location','best')
    end
    subplot2n(4,1,2), plot(mF2g_,'color',[0,0.55,0.69],'linewidth',1.5), hold on, plot(mF6g_,'color','k','linewidth',0.2), ylabel('mean time series'),
        grid on,grid minor, xlim([1 length(mF2g_)]),ylim([1.25.*min([mF2g_,mF6g_]) 1.25.*max([mF2g_,mF6g_])]), title(['Bandpass filtered: normalized mean GRAY MATTER signal AFTER "15-bandpass-filter" (all slices averaged)'],'Color',[0,0.55,0.69],'FontSize',14), legend({'Gray matter signal after filtering'},'FontSize',12,'TextColor',[0,0.55,0.69],'Location','best')
    subplot2n(4,1,4), plot(mF2w_,'color',[0.65,0.11,0.19],'linewidth',1.5), hold on, plot(mF6w_,'color','k','linewidth',0.2), ylabel('mean time series'),
        grid on,grid minor, xlim([1 length(mF2g_)]),ylim([1.25.*min([mF2w_,mF6w_]) 1.25.*max([mF2w_,mF6w_])]), title(['Bandpass filtered: normalized mean WHITE MATTER signal AFTER "15-bandpass-filter" (all slices averaged)'],'Color',[0.65,0.11,0.19],'FontSize',14), legend({'White matter signal after filtering'},'FontSize',12,'TextColor',[0.65,0.11,0.19],'Location','best')
    saveas(fighndl,[QCpath 'mean_timeseries_14denoise5_vs_15filter_allSlices' '.jpg'])
     close(fighndl)
    
    fighndl = figure('Position',[1 1 1152 720],'Color',[0.89,0.94,0.99],'InvertHardcopy','off','Visible','off'); % red [0.95,0.9,0.85], green [0.85,0.90,0.85], blue [0.89,0.94,0.99]
    if stp==1
    	subplot2n(4,1,1), histogram(C3g_,'EdgeColor','none','FaceColor','k','FaceAlpha',0.5,'Normalization','probability'); grid on,grid minor, xlim([-1 1]), hold on, xline(0,'Color','k','linewidth',1.5); hold off
            title(sprintf('Histogram of voxel-to-voxel GRAY MATTER correlations AFTER 11-registration (all slices combined)\n%.1f%% of the correlations were > 0  [ mean=%.2f, median=%.2f, SD=%.2f ]',100*length(find(C3g_>0))/length(C3g_),mean(C3g_),median(C3g_),std(C3g_)),'Color','k','FontSize',14)
        subplot2n(4,1,3), histogram(C3w_,'EdgeColor','none','FaceColor','k','FaceAlpha',0.5,'Normalization','probability'); grid on,grid minor, xlim([-1 1]), hold on, xline(0,'Color','k','linewidth',1.5); hold off
            title(sprintf('WHITE MATTER: AFTER 11-registration (all slices): %.1f%% of the correlations were > 0  [ mean=%.2f, median=%.2f, SD=%.2f ]',100*length(find(C3w_>0))/length(C3w_),mean(C3w_),median(C3w_),std(C3w_)),'Color','k','FontSize',14)
    elseif stp==3
    	subplot2n(4,1,1), histogram(C3g_,'EdgeColor','none','FaceColor','k','FaceAlpha',0.5,'Normalization','probability'); grid on,grid minor, xlim([-1 1]), hold on, xline(0,'Color','k','linewidth',1.5); hold off
            title(sprintf('Histogram of voxel-to-voxel GRAY MATTER correlations AFTER 12-denoise3-CSFregress (all slices combined)\n%.1f%% of the correlations were > 0  [ mean=%.2f, median=%.2f, SD=%.2f ]',100*length(find(C3g_>0))/length(C3g_),mean(C3g_),median(C3g_),std(C3g_)),'Color','k','FontSize',14)
        subplot2n(4,1,3), histogram(C3w_,'EdgeColor','none','FaceColor','k','FaceAlpha',0.5,'Normalization','probability'); grid on,grid minor, xlim([-1 1]), hold on, xline(0,'Color','k','linewidth',1.5); hold off
            title(sprintf('WHITE MATTER: AFTER 12-denoise3-CSFregress (all slices): %.1f%% of the correlations were > 0  [ mean=%.2f, median=%.2f, SD=%.2f ]',100*length(find(C3w_>0))/length(C3w_),mean(C3w_),median(C3w_),std(C3w_)),'Color','k','FontSize',14)
    elseif (stp==2)||(stp==6)
    	subplot2n(4,1,1), histogram(C3g_,'EdgeColor','none','FaceColor','k','FaceAlpha',0.5,'Normalization','probability'); grid on,grid minor, xlim([-1 1]), hold on, xline(0,'Color','k','linewidth',1.5); hold off
            title(sprintf('Histogram of voxel-to-voxel GRAY MATTER correlations AFTER 13-denoise4-WMregress (all slices combined)\n%.1f%% of the correlations were > 0  [ mean=%.2f, median=%.2f, SD=%.2f ]',100*length(find(C3g_>0))/length(C3g_),mean(C3g_),median(C3g_),std(C3g_)),'Color','k','FontSize',14)
        subplot2n(4,1,3), histogram(C3w_,'EdgeColor','none','FaceColor','k','FaceAlpha',0.5,'Normalization','probability'); grid on,grid minor, xlim([-1 1]), hold on, xline(0,'Color','k','linewidth',1.5); hold off
            title(sprintf('WHITE MATTER: AFTER 13-denoise4-WMregress (all slices): %.1f%% of the correlations were > 0  [ mean=%.2f, median=%.2f, SD=%.2f ]',100*length(find(C3w_>0))/length(C3w_),mean(C3w_),median(C3w_),std(C3w_)),'Color','k','FontSize',14)
    elseif (stp==4)||(stp==5)||(stp==7)||(stp==8)
    	subplot2n(4,1,1), histogram(C3g_,'EdgeColor','none','FaceColor','k','FaceAlpha',0.5,'Normalization','probability'); grid on,grid minor, xlim([-1 1]), hold on, xline(0,'Color','k','linewidth',1.5); hold off
            title(sprintf('Histogram of voxel-to-voxel GRAY MATTER correlations AFTER 14-denoise5-COVregress (all slices combined)\n%.1f%% of the correlations were > 0  [ mean=%.2f, median=%.2f, SD=%.2f ]',100*length(find(C3g_>0))/length(C3g_),mean(C3g_),median(C3g_),std(C3g_)),'Color','k','FontSize',14)
        subplot2n(4,1,3), histogram(C3w_,'EdgeColor','none','FaceColor','k','FaceAlpha',0.5,'Normalization','probability'); grid on,grid minor, xlim([-1 1]), hold on, xline(0,'Color','k','linewidth',1.5); hold off
            title(sprintf('WHITE MATTER: AFTER 14-denoise5-COVregress (all slices): %.1f%% of the correlations were > 0  [ mean=%.2f, median=%.2f, SD=%.2f ]',100*length(find(C3w_>0))/length(C3w_),mean(C3w_),median(C3w_),std(C3w_)),'Color','k','FontSize',14)
    end
    subplot2n(4,1,2), histogram(Cg_,'EdgeColor','none','FaceColor',[0,0.55,0.69],'FaceAlpha',0.5,'Normalization','probability'); grid on,grid minor, xlim([-1 1]), hold on, xline(0,'Color','k','linewidth',1.5); hold off
       title(sprintf('GRAY MATTER: AFTER 15-bandpass-filter (all slices): %.1f%% of correlations > 0  [ mean=%.2f, median=%.2f, SD=%.2f ]',100*length(find(Cg_>0))/length(Cg_),mean(Cg_),median(Cg_),std(Cg_)),'Color',[0,0.55,0.69],'FontSize',14)
    subplot2n(4,1,4), histogram(Cw_,'EdgeColor','none','FaceColor',[0.65,0.11,0.19],'FaceAlpha',0.5,'Normalization','probability'); grid on,grid minor, xlim([-1 1]), hold on, xline(0,'Color','k','linewidth',1.5); hold off
       title(sprintf('WHITE MATTER: AFTER 15-bandpass-filter (all slices): %.1f%% of the correlations were > 0  [ mean=%.2f, median=%.2f, SD=%.2f ]',100*length(find(Cw_>0))/length(Cw_),mean(Cw_),median(Cw_),std(Cw_)),'Color',[0.65,0.11,0.19],'FontSize',14)
    saveas(fighndl,[QCpath 'hist_FC_compare_14denoise5_15filter_allSlices' '.jpg'])
     close(fighndl)
end
clear svar1 spsc1 svar2 spsc2 sortorder1 sortorder2 F2g_ mF2g_ F2w_ mF2w_ Cg_ Cw_ F4_ mF4_ C2_ F6g_ F6w_ mF6g_ mF6w_ C3g_ C3w_

    catch
    end
end

%% scfMRItb_16_cord_quadrants
if (choices_preproc(16)==1)&&(k2==1)
clear QCpath save_videos fighndl stp
if ~(exist([base_dir_sub 'QC' '/16_cord_quadrants'],'dir'))
    mkdir([base_dir_sub 'QC' '/16_cord_quadrants'])
end
QCpath = [base_dir_sub 'QC' '/16_cord_quadrants' '/'];

try
    load([base_dir_sub 'cord_quadrants.mat'],'FC_mask_quads')
    FC_mask_quads = FC_mask_quads.img;
catch
    load([base_dir_sub sprintf('workspace_variables_run%d.mat',k2)],'FC_mask_quads')
end
if length(FC_mask_quads)==1
    FC_mask_quads = FC_mask_quads.img;
end    

for i3 = 1 : siz3
    try waitbar((i3/siz3),wbar3,sprintf('16. Define cord quadrants: slice (%d) of (%d)',i3,siz3)); catch, end
    disp(sprintf('16. Define cord quadrants: slice (%d) of (%d), subject (%d) of (%d)',i3,siz3,k,size(func_files,1)))
    fighndl2 = figure('Position',[round((scrsz(3)-scrsz(4))/2) 1 scrsz(4) scrsz(4)],'InvertHardcopy','off','Visible','off');
    fighndl2.Color = [0.95, 0.875, 0.8];
    tmp = FC_mask_quads(:,:,i3); tmp(find(tmp==0))=NaN;
    tmp = rot90(tmp);
    [w1,w2] = find(rot90(FC_mask_quads(:,:,i3))~=0);
	imshow(tmp, [0 max(tmp(:))], 'InitialMagnification', 'fit'); colormap('jet'),colorbar;
    colormap([0 0 0; parula(256)])
    xlim([min(w2)-1,max(w2)+1]), ylim([min(w1)-1,max(w1)+1]);
    title(sprintf('GM and WM quadrants for slice %d of %d',i3,siz3),'Color','red','FontSize',18)
    saveas(fighndl2,[QCpath 'cord_quadrants_slice' sprintf('%.2d',i3) '.jpg'])
    close(fighndl2)
    clear w1 w2 tmp fighndl2
end
clear FC_mask_quads

try
    load([base_dir_sub 'cross_sectional_areas.mat'],'area_cord_quadrants')
    for w=1:size(area_cord_quadrants,1)
        for w2=1:size(area_cord_quadrants,2)
            dispstring1{w,w2} = sprintf('%.2f',area_cord_quadrants(w,w2));
        end
    end; clear w2
    fighndl = figure('Position',[1 1 1280 800],'Color',[0.85,0.90,0.85],'InvertHardcopy','off','Visible','off'); % red [0.95,0.9,0.85], green [0.85,0.90,0.85], blue [0.89,0.94,0.99]
    plot(area_cord_quadrants(:,1),'--p','color','r','linewidth',2.25, 'MarkerSize',14, 'MarkerEdgeColor','k', 'MarkerFaceColor','r'),
        hold on, plot(area_cord_quadrants(:,2),'--p','color','b','linewidth',2.25, 'MarkerSize',14, 'MarkerEdgeColor','k', 'MarkerFaceColor','b'),
        plot(area_cord_quadrants(:,3),'--p','color',[0 0.37 0.22],'linewidth',2.25, 'MarkerSize',14, 'MarkerEdgeColor','k', 'MarkerFaceColor',[0 0.37 0.22]),
        plot(area_cord_quadrants(:,4),'--p','color','m','linewidth',2.25, 'MarkerSize',14, 'MarkerEdgeColor','k', 'MarkerFaceColor','m'),
        plot(area_cord_quadrants(:,5),'--p','color',[0.6 0.6 0.1],'linewidth',2.25, 'MarkerSize',14, 'MarkerEdgeColor','k', 'MarkerFaceColor','y'),
        plot(area_cord_quadrants(:,6),'--p','color','c','linewidth',2.25, 'MarkerSize',14, 'MarkerEdgeColor','k', 'MarkerFaceColor','c'),
        plot(area_cord_quadrants(:,7),'--p','color','g','linewidth',2.25, 'MarkerSize',14, 'MarkerEdgeColor','k', 'MarkerFaceColor','g'),
        plot(area_cord_quadrants(:,8),'--p','color',[0.6 0.1 0.6],'linewidth',2.25, 'MarkerSize',14, 'MarkerEdgeColor','k', 'MarkerFaceColor',[0.6 0.1 0.6]),
        grid on,grid minor, ylabel('area (in millimeters squared)'), xlabel('SLICES'), xticks([1:size(area_cord_quadrants,1)]),
        xlim([0.5 size(area_cord_quadrants,1)+0.5]), ylim([0 max(area_cord_quadrants(:))+4])
        text([1:size(area_cord_quadrants,1)]-0.1,area_cord_quadrants(:,1)+2, dispstring1(:,1),'Color','r','FontSize',10)
        text([1:size(area_cord_quadrants,1)]-0.1,area_cord_quadrants(:,2)+1, dispstring1(:,2),'Color','b','FontSize',10)
        text([1:size(area_cord_quadrants,1)]-0.1,max([0.5.*ones(size(area_cord_quadrants,1),1), area_cord_quadrants(:,3)-2]')', dispstring1(:,3),'Color',[0 0.37 0.22],'FontSize',10)
        text([1:size(area_cord_quadrants,1)]-0.1,max([0.5.*ones(size(area_cord_quadrants,1),1), area_cord_quadrants(:,4)-1]')', dispstring1(:,4),'Color','m','FontSize',10)
        text([1:size(area_cord_quadrants,1)]-0.1,area_cord_quadrants(:,5)+2, dispstring1(:,5),'Color',[0.6 0.6 0.1],'FontSize',10)
        text([1:size(area_cord_quadrants,1)]-0.1,area_cord_quadrants(:,6)+1, dispstring1(:,6),'Color','c','FontSize',10)
        text([1:size(area_cord_quadrants,1)]-0.1,max([0.5.*ones(size(area_cord_quadrants,1),1), area_cord_quadrants(:,7)-2]')', dispstring1(:,7),'Color','g','FontSize',10)
        text([1:size(area_cord_quadrants,1)]-0.1,max([0.5.*ones(size(area_cord_quadrants,1),1), area_cord_quadrants(:,8)-1]')', dispstring1(:,8),'Color',[0.6 0.1 0.6],'FontSize',10)
        legend({'Left ventral GM';'Right ventral GM';'Left dorsal GM';'Right dorsal GM';'Left ventral WM';'Right ventral WM';'Left dorsal WM';'Right dorsal WM'},'FontSize',14,'TextColor','k','Location','best')
        title('Cross sectional areas of each GM and WM quadrant across slices (units: millimeters squared)','Color',[0.65,0.11,0.19],'FontSize',16)
    saveas(fighndl,[QCpath 'cross_sectional_areas_each_quad' '.jpg'])
    close(fighndl)
    clear dispstring1 w
catch
end

end

%% scfMRItb_17_deconvolution_rest
if choices_preproc(17)==1
    try
clear QCpath save_videos fighndl stp
if ~(exist([base_dir_sub 'QC' '/17_deconvolution'],'dir'))
    mkdir([base_dir_sub 'QC' '/17_deconvolution'])
end
if Rns==0
    QCpath = [base_dir_sub 'QC' '/17_deconvolution/'];
else
    if ~(exist([base_dir_sub 'QC' '/17_deconvolution' '/run' num2str(Rns)],'dir'))
        mkdir([base_dir_sub 'QC' '/17_deconvolution' '/run' num2str(Rns)])
    end
    QCpath = [base_dir_sub 'QC' '/17_deconvolution' '/run' num2str(Rns) '/'];
end

f_BPF_max_vec = bpf_hz(1);
if f_BPF_max_vec<0.1
    f_BPF_max_str_vec = sprintf('0%.0f',f_BPF_max_vec*100); %#ok<*AGROW>
else
    f_BPF_max_str_vec = sprintf('%.0f',f_BPF_max_vec*100);    
end

try
    scfMRItb_04_unzipFile(base_dir_sub, fname_anat, '2_mask_GM')
    scfMRItb_04_unzipFile(base_dir_sub, fname_anat, '2_mask_WM')
    mask_GM2 = load_untouch_nii([base_dir_sub fname_anat '2_mask_GM.nii']); mask_GM2 = mask_GM2.img;
    mask_WM2 = load_untouch_nii([base_dir_sub fname_anat '2_mask_WM.nii']); mask_WM2 = mask_WM2.img;
catch
    scfMRItb_04_unzipFile(base_dir_sub, fname_anat, '')
    A = load_untouch_nii([base_dir_sub fname_anat '.nii']);
    scfMRItb_04_unzipFile(base_dir_sub, fname_anat, '_mask_GM')
    scfMRItb_04_unzipFile(base_dir_sub, fname_anat, '_mask_WM')
    scfMRItb_04_unzipFile(base_dir_sub, fname_anat, '_mask_CSF')
    mmask1 = load_untouch_nii([base_dir_sub fname_anat '_mask_GM.nii']); % gray matter
    mmask2 = load_untouch_nii([base_dir_sub fname_anat '_mask_WM.nii']); % white matter
    mmask3 = load_untouch_nii([base_dir_sub fname_anat '_mask_CSF.nii']); % white matter
    mmask_all = single(or(or(mmask1.img, mmask2.img), mmask3.img));
    smmask = (mean(mmask_all,3) > 0);
    [s1,s2] = find(smmask~=0);
    s1median = median(s1); s2median = median(s2);
    s1min = min(s1); s2min = min(s2);
    s1max = max(s1); s2max = max(s2);
    smaxrange = max([s2max-s2min,s1max-s1min]);
    if round(size(mmask_all,1)/4) > smaxrange
        new_size = round(size(mmask_all,1)/4); % 128 - assume even for now
    elseif round(size(mmask_all,1)/3) > smaxrange
        new_size = round(size(mmask_all,1)/3); % 171
    elseif round(size(mmask_all,1)/2) > smaxrange
        new_size = round(size(mmask_all,1)/2); % 256
    else
        new_size = size(mmask_all,1); % 512
    end
    try if fullsize==1, new_size = size(mmask_all,1); end, catch, end    
    xmin = s1median - round(new_size/2) + 1; xmax = s1median + round(new_size/2);
    ymin = s2median - round(new_size/2) + 1; ymax = s2median + round(new_size/2);
    mask_GM_reduced = mmask1; mask_WM_reduced = mmask2; mask_CSF_reduced = mmask3;
    mask_GM_reduced.hdr.dime.dim(2)  = new_size; mask_GM_reduced.hdr.dime.dim(3)  = new_size;
    mask_WM_reduced.hdr.dime.dim(2)  = new_size; mask_WM_reduced.hdr.dime.dim(3)  = new_size;
    mask_CSF_reduced.hdr.dime.dim(2) = new_size; mask_CSF_reduced.hdr.dime.dim(3) = new_size;
    
    basestr = strfind(base_dir_sub,'/');
    basestr = base_dir_sub(basestr(end-1)+1:end); % name of subject folder
    if str2num(basestr(1:4)) <= 2017 %#ok<ST2NM> % if data were acquired prior to 2018 then don't change the bounding box of the reduced size images that were already processed before
        mask_GM_reduced.img  = mmask1.img(round((size(A.img,1)-new_size)/2)+1:end-round((size(A.img,1)-new_size)/2), ...
            round((size(A.img,1)-new_size)/2)+1:end-round((size(A.img,1)-new_size)/2), :);
        mask_WM_reduced.img  = mmask2.img(round((size(A.img,1)-new_size)/2)+1:end-round((size(A.img,1)-new_size)/2), ...
            round((size(A.img,1)-new_size)/2)+1:end-round((size(A.img,1)-new_size)/2), :);
        mask_CSF_reduced.img = mmask3.img(round((size(A.img,1)-new_size)/2)+1:end-round((size(A.img,1)-new_size)/2), ...
            round((size(A.img,1)-new_size)/2)+1:end-round((size(A.img,1)-new_size)/2), :); %#ok<*NASGU>
    else % else if data were acquired after 2017 then use the new method to determine the bounding box by centering the new box around the centroid of the spinal cord mask
        mask_GM_reduced.img  = mmask1.img(xmin:xmax, ymin:ymax, :);
        mask_WM_reduced.img  = mmask2.img(xmin:xmax, ymin:ymax, :);
        mask_CSF_reduced.img = mmask3.img(xmin:xmax, ymin:ymax, :);
    end    
    mask_SC_reduced = mask_GM_reduced;
    mask_SC_reduced.img = or(mask_GM_reduced.img, mask_WM_reduced.img);
    if ~exist([base_dir_sub fname_anat '2' '_mask_GM.nii'],'file')
        save_untouch_nii(mask_GM_reduced,  [base_dir_sub fname_anat '2' '_mask_GM.nii']); % gray matter
    end
    if ~exist([base_dir_sub fname_anat '2' '_mask_WM.nii'],'file')
        save_untouch_nii(mask_WM_reduced,  [base_dir_sub fname_anat '2' '_mask_WM.nii']); % white matter
    end
    if ~exist([base_dir_sub fname_anat '2' '_mask_CSF.nii'],'file')
        save_untouch_nii(mask_CSF_reduced, [base_dir_sub fname_anat '2' '_mask_CSF.nii']); % white matter
    end
    if ~exist([base_dir_sub fname_anat '2' '_mask_SC.nii'],'file')
        save_untouch_nii(mask_SC_reduced,  [base_dir_sub fname_anat '2' '_mask_SC.nii']); % gray matter
    end
    if ~exist([base_dir_sub fname_anat '2' '_all_masks.mat'],'file')
        save([base_dir_sub fname_anat '2' '_all_masks.mat'],'mask_GM_reduced','mask_WM_reduced','mask_CSF_reduced','mask_SC_reduced')
    end
    if ~exist([base_dir_sub fname_anat '2' '.nii'],'file')
        anat_reduced2 = A; anat_reduced2.hdr.dime.dim(2) = new_size; anat_reduced2.hdr.dime.dim(3) = new_size;
        anat_reduced2.img = A.img(xmin:xmax, ymin:ymax, :);
        save_untouch_nii(anat_reduced2, [base_dir_sub fname_anat '2' '.nii'])
    end
    mask_GM2 = mask_GM_reduced; mask_WM2 = mask_WM_reduced; mask_CSF2 = mask_CSF_reduced;
    clear mmask1 mmask2 mmask3 mmask_all smmask s1 s2 s1median s2median s1min s1max s2min s2max smaxrange new_size xmin xmax ymin ymax mask_GM_reduced mask_WM_reduced mask_CSF_reduced basestr mask_SC_reduced anat_reduced2 A
end

eval(sprintf('load([base_dir_sub ''workspace_variables_run%d.mat''],''FC_mask_quads'');',RUNS(k)))
if size(FC_mask_quads)==1
    FC_mask_quads = FC_mask_quads.img;
end
    % (1) = left ventral GM; (2) = right ventral GM; (3) = left dorsal GM; (4) = right dorsal GM
    % (5) = left ventral WM; (6) = right ventral WM; (7) = left dorsal WM; (8) = right dorsal WM
FC_mask_GM = FC_mask_quads; FC_mask_GM(find(FC_mask_GM>=5))=0; FC_mask_GM(find(FC_mask_GM>=1))=1;
FC_mask_WM = FC_mask_quads; FC_mask_WM(find(FC_mask_WM<=4))=0; FC_mask_WM(find(FC_mask_WM>=5))=1;

if exist([base_dir_sub fname '_warped' '.nii'], 'file') || exist([base_dir_sub fname '_warped' '.nii.gz'], 'file') % if none of CSF, WM or COV regress steps were performed
    whichip=8;
end
if exist([base_dir_sub fname '_WM' '.nii'], 'file') || exist([base_dir_sub fname '_WM' '.nii.gz'], 'file') % if only WM regress step was performed
    whichip=7;
end
if exist([base_dir_sub fname '_denoised' cutoff_str '.nii'], 'file') || exist([base_dir_sub fname '_denoised' cutoff_str '.nii.gz'], 'file') % if only CSF regress step was performed
    whichip=6;
end
if exist([base_dir_sub fname '_cov' '.nii'], 'file') || exist([base_dir_sub fname '_cov' '.nii.gz'], 'file') % if only COV regress step was performed
    whichip=5;
end
if exist([base_dir_sub fname '_WMcov' '.nii'], 'file') || exist([base_dir_sub fname '_WMcov' '.nii.gz'], 'file') % if both WM and COV regress steps were performed
    whichip=4;
end
if exist([base_dir_sub fname '_denoised' cutoff_str 'WM' '.nii'], 'file') || exist([base_dir_sub fname '_denoised' cutoff_str 'WM' '.nii.gz'], 'file') % if both CSF and WM regress steps were performed
    whichip=3;
end
if exist([base_dir_sub fname '_denoised' cutoff_str 'cov' '.nii'], 'file') || exist([base_dir_sub fname '_denoised' cutoff_str 'cov' '.nii.gz'], 'file') % if both CSF and COV regress steps were performed
    whichip=2;
end
if exist([base_dir_sub fname '_denoised' cutoff_str 'WMcov' '.nii'], 'file') || exist([base_dir_sub fname '_denoised' cutoff_str 'WMcov' '.nii.gz'], 'file') % if CSF, WM and COV regress steps were performed
    whichip=1;
end

if run_on_HPF_data==1
    if whichip==1
        load([base_dir_sub fname '_denoised' cutoff_str 'WMcov' '_deconv' '_BPF' bpf_hz_str '.mat']); F1 = F; clear F
        load([base_dir_sub fname '_denoised' cutoff_str 'WMcov' '_HRFparams' '.mat'], 'RH','TTP','FWHM','HRF'); % load HRF parameters
    elseif whichip==2
        load([base_dir_sub fname '_denoised' cutoff_str 'cov' '_deconv' '_BPF' bpf_hz_str '.mat']); F1 = F; clear F
        load([base_dir_sub fname '_denoised' cutoff_str 'cov' '_HRFparams' '.mat'], 'RH','TTP','FWHM','HRF'); % load HRF parameters
    elseif whichip==3
        load([base_dir_sub fname '_denoised' cutoff_str 'WM' '_deconv' '_BPF' bpf_hz_str '.mat']); F1 = F; clear F
        load([base_dir_sub fname '_denoised' cutoff_str 'WM' '_HRFparams' '.mat'], 'RH','TTP','FWHM','HRF'); % load HRF parameters
    elseif whichip==4
        load([base_dir_sub fname '_WMcov' '_deconv' '_BPF' bpf_hz_str '.mat']); F1 = F; clear F
        load([base_dir_sub fname '_WMcov' '_HRFparams' '.mat'], 'RH','TTP','FWHM','HRF'); % load HRF parameters
    elseif whichip==5
        load([base_dir_sub fname '_cov' '_deconv' '_BPF' bpf_hz_str '.mat']); F1 = F; clear F
        load([base_dir_sub fname '_cov' '_HRFparams' '.mat'], 'RH','TTP','FWHM','HRF'); % load HRF parameters
    elseif whichip==6
        load([base_dir_sub fname '_denoised' cutoff_str '_deconv' '_BPF' bpf_hz_str '.mat']); F1 = F; clear F
        load([base_dir_sub fname '_denoised' cutoff_str '_HRFparams' '.mat'], 'RH','TTP','FWHM','HRF'); % load HRF parameters
    elseif whichip==7
        load([base_dir_sub fname '_WM' '_deconv' '_BPF' bpf_hz_str '.mat']); F1 = F; clear F
        load([base_dir_sub fname '_WM' '_HRFparams' '.mat'], 'RH','TTP','FWHM','HRF'); % load HRF parameters
    elseif whichip==8
        load([base_dir_sub fname '_warped' '_deconv' '_BPF' bpf_hz_str '.mat']); F1 = F; clear F
        load([base_dir_sub fname '_warped' '_HRFparams' '.mat'], 'RH','TTP','FWHM','HRF'); % load HRF parameters
    end
elseif run_on_HPF_data==0
    if whichip==1
        load([base_dir_sub fname '_denoised' cutoff_str 'WMcov' '_BPF' bpf_hz_str '_deconv' '.mat']); F1 = F; clear F
        load([base_dir_sub fname '_denoised' cutoff_str 'WMcov' '_BPF' '_HRFparams' '.mat'], 'RH','TTP','FWHM','HRF'); % load HRF parameters
    elseif whichip==2
        load([base_dir_sub fname '_denoised' cutoff_str 'cov' '_BPF' bpf_hz_str '_deconv' '.mat']); F1 = F; clear F
        load([base_dir_sub fname '_denoised' cutoff_str 'cov' '_BPF' '_HRFparams' '.mat'], 'RH','TTP','FWHM','HRF'); % load HRF parameters
    elseif whichip==3
        load([base_dir_sub fname '_denoised' cutoff_str 'WM' '_BPF' bpf_hz_str '_deconv' '.mat']); F1 = F; clear F
        load([base_dir_sub fname '_denoised' cutoff_str 'WM' '_BPF' '_HRFparams' '.mat'], 'RH','TTP','FWHM','HRF'); % load HRF parameters
    elseif whichip==4
        load([base_dir_sub fname '_WMcov' '_BPF' bpf_hz_str '_deconv' '.mat']); F1 = F; clear F
        load([base_dir_sub fname '_WMcov' '_BPF' '_HRFparams' '.mat'], 'RH','TTP','FWHM','HRF'); % load HRF parameters
    elseif whichip==5
        load([base_dir_sub fname '_cov' '_BPF' bpf_hz_str '_deconv' '.mat']); F1 = F; clear F
        load([base_dir_sub fname '_cov' '_BPF' '_HRFparams' '.mat'], 'RH','TTP','FWHM','HRF'); % load HRF parameters
    elseif whichip==6
        load([base_dir_sub fname '_denoised' cutoff_str '_BPF' bpf_hz_str '_deconv' '.mat']); F1 = F; clear F
        load([base_dir_sub fname '_denoised' cutoff_str '_BPF' '_HRFparams' '.mat'], 'RH','TTP','FWHM','HRF'); % load HRF parameters
    elseif whichip==7
        load([base_dir_sub fname '_WM' '_BPF' bpf_hz_str '_deconv' '.mat']); F1 = F; clear F
        load([base_dir_sub fname '_WM' '_BPF' '_HRFparams' '.mat'], 'RH','TTP','FWHM','HRF'); % load HRF parameters
    elseif whichip==8
        load([base_dir_sub fname '_warped' '_BPF' bpf_hz_str '_deconv' '.mat']); F1 = F; clear F
        load([base_dir_sub fname '_warped' '_BPF' '_HRFparams' '.mat'], 'RH','TTP','FWHM','HRF'); % load HRF parameters
    end
end

if exist([base_dir_sub fname '_denoised' cutoff_str 'WMcov' '_BPF' bpf_hz_str '.nii'],'file') || exist([base_dir_sub fname '_denoised' cutoff_str 'WMcov' '_BPF' bpf_hz_str '.nii.gz'],'file')
    load([base_dir_sub fname '_denoised' cutoff_str 'WMcov' '_BPF' bpf_hz_str '.mat']); F5all=single(F.img); clear F
    stp=8;
elseif exist([base_dir_sub fname '_denoised' cutoff_str 'cov' '_BPF' bpf_hz_str '.nii'],'file') || exist([base_dir_sub fname '_denoised' cutoff_str 'cov' '_BPF' bpf_hz_str '.nii.gz'],'file')
    load([base_dir_sub fname '_denoised' cutoff_str 'cov' '_BPF' bpf_hz_str '.mat']); F5all=single(F.img); clear F
    stp=7;
elseif exist([base_dir_sub fname '_denoised' cutoff_str 'WM' '_BPF' bpf_hz_str '.nii'],'file') || exist([base_dir_sub fname '_denoised' cutoff_str 'WM' '_BPF' bpf_hz_str '.nii.gz'],'file')
    load([base_dir_sub fname '_denoised' cutoff_str 'WM' '_BPF' bpf_hz_str '.mat']); F5all=single(F.img); clear F
    stp=6;
elseif exist([base_dir_sub fname '_WMcov' '_BPF' bpf_hz_str '.nii'],'file') || exist([base_dir_sub fname '_WMcov' '_BPF' bpf_hz_str '.nii.gz'],'file')
    load([base_dir_sub fname '_WMcov' '_BPF' bpf_hz_str '.mat']); F5all=single(F.img); clear F
    stp=5;
elseif exist([base_dir_sub fname '_cov' '_BPF' bpf_hz_str '.nii'],'file') || exist([base_dir_sub fname '_cov' '_BPF' bpf_hz_str '.nii.gz'],'file')
    load([base_dir_sub fname '_cov' '_BPF' bpf_hz_str '.mat']); F5all=single(F.img); clear F
    stp=4;
elseif exist([base_dir_sub fname '_denoised' cutoff_str '_BPF' bpf_hz_str '.nii'],'file') || exist([base_dir_sub fname '_denoised' cutoff_str '_BPF' bpf_hz_str '.nii.gz'],'file')
    load([base_dir_sub fname '_denoised' cutoff_str '_BPF' bpf_hz_str '.mat']); F5all=single(F.img); clear F
    stp=3;
elseif exist([base_dir_sub fname '_WM' '_BPF' bpf_hz_str '.nii'],'file') || exist([base_dir_sub fname '_WM' '_BPF' bpf_hz_str '.nii.gz'],'file')
    load([base_dir_sub fname '_WM' '_BPF' bpf_hz_str '.mat']); F5all=single(F.img); clear F
    stp=2;
elseif exist([base_dir_sub fname '_warped' '_BPF' bpf_hz_str '.nii'],'file') || exist([base_dir_sub fname '_warped' '_BPF' bpf_hz_str '.nii.gz'],'file')
    load([base_dir_sub fname '_warped' '_BPF' bpf_hz_str '.mat']); F5all=single(F.img); clear F
    stp=1;
end
TR = F1.hdr.dime.pixdim(5); F1 = single(squeeze(F1.img));


for w1=1:size(HRF,3)
    for w2=1:size(HRF,4)
        tmp = HRF(:,:,w1,w2);
        HRF1_GM(w1,w2) = mean(tmp(find(FC_mask_GM(:,:,w1)==1)));
        HRF1_WM(w1,w2) = mean(tmp(find(FC_mask_WM(:,:,w1)==1)));
        clear tmp
    end
end; clear w1 w2
for w2=1:size(HRF,4)
    tmp = HRF(:,:,:,w2);
    HRF2_GM(1,w2) = mean(tmp(find(FC_mask_quads==1))); HRF2_GM(2,w2) = mean(tmp(find(FC_mask_quads==2)));
    HRF2_GM(3,w2) = mean(tmp(find(FC_mask_quads==3))); HRF2_GM(4,w2) = mean(tmp(find(FC_mask_quads==4)));
    HRF2_WM(1,w2) = mean(tmp(find(FC_mask_quads==5))); HRF2_WM(2,w2) = mean(tmp(find(FC_mask_quads==6)));
    HRF2_WM(3,w2) = mean(tmp(find(FC_mask_quads==7))); HRF2_WM(4,w2) = mean(tmp(find(FC_mask_quads==8)));
    clear tmp
end; clear w2
for w=1:size(HRF1_GM,1)
    legstr{w} = sprintf('slice %d',w);
    xlabl{w}  = sprintf('%.2f s',(w-1)*TR);
end; clear w
    % (1) = left ventral GM; (2) = right ventral GM; (3) = left dorsal GM; (4) = right dorsal GM
    % (5) = left ventral WM; (6) = right ventral WM; (7) = left dorsal WM; (8) = right dorsal WM
fighndl = figure('Position',[1 1 1280 800],'Color',[0.85,0.90,0.85],'InvertHardcopy','off','Visible','off'); % red [0.95,0.9,0.85], green [0.85,0.90,0.85], blue [0.89,0.94,0.99]
subplot2n(4,1,1), plot(HRF1_GM','linewidth',1), xlabel('time (seconds)'), ylabel('HRF'), xticks([1:length(HRF1_GM')]),xticklabels(xlabl), legend(legstr,'NumColumns',3)
    grid on,grid minor, ylim([1.1.*min([HRF1_GM(:);HRF1_WM(:);HRF2_GM(:);HRF2_WM(:)]), 1.1.*max([HRF1_GM(:);HRF1_WM(:);HRF2_GM(:);HRF2_WM(:)])]), title(['Hemodynamic response function (HRF) in the GRAY MATTER across SLICES'],'Color',[0,0.55,0.69],'FontSize',16),
subplot2n(4,1,2), plot(HRF1_WM','linewidth',1), xlabel('time (seconds)'), ylabel('HRF'), xticks([1:length(HRF1_WM')]),xticklabels(xlabl), legend(legstr,'NumColumns',3)
    grid on,grid minor, ylim([1.1.*min([HRF1_GM(:);HRF1_WM(:);HRF2_GM(:);HRF2_WM(:)]), 1.1.*max([HRF1_GM(:);HRF1_WM(:);HRF2_GM(:);HRF2_WM(:)])]), title(['HRF in the WHITE MATTER across SLICES'],'Color',[0.65,0.11,0.19],'FontSize',16),
subplot2n(4,1,3), plot(HRF2_GM','linewidth',1), xlabel('time (seconds)'), ylabel('HRF'), xticks([1:length(HRF2_GM')]),xticklabels(xlabl), legend({'left ventral GM','right ventral GM','left dorsal GM','right dorsal GM'},'FontSize',12,'NumColumns',2)
    grid on,grid minor, ylim([1.1.*min([HRF1_GM(:);HRF1_WM(:);HRF2_GM(:);HRF2_WM(:)]), 1.1.*max([HRF1_GM(:);HRF1_WM(:);HRF2_GM(:);HRF2_WM(:)])]), title(['HRF in the GRAY MATTER across QUADRANTS'],'Color',[0,0.55,0.69],'FontSize',16),
subplot2n(4,1,4), plot(HRF2_WM','linewidth',1), xlabel('time (seconds)'), ylabel('HRF'), xticks([1:length(HRF2_WM')]),xticklabels(xlabl), legend({'left ventral WM','right ventral WM','left dorsal WM','right dorsal WM'},'FontSize',12,'NumColumns',2)
    grid on,grid minor, ylim([1.1.*min([HRF1_GM(:);HRF1_WM(:);HRF2_GM(:);HRF2_WM(:)]), 1.1.*max([HRF1_GM(:);HRF1_WM(:);HRF2_GM(:);HRF2_WM(:)])]), title(['HRF in the WHITE MATTER across QUADRANTS'],'Color',[0.65,0.11,0.19],'FontSize',16),
saveas(fighndl,[QCpath 'HRF_plots_17deconv_allSlices' '.jpg'])
clear HRF1_GM HRF1_WM HRF2_GM HRF2_WM
close(fighndl)

for w1=1:size(RH,3)
    tmp1 = RH(:,:,w1); tmp2 = TTP(:,:,w1); tmp3 = FWHM(:,:,w1);
    RH1_GM(w1,1)   = mean(tmp1(find(FC_mask_GM(:,:,w1)==1))); RH1_WM(w1,1)   = mean(tmp1(find(FC_mask_WM(:,:,w1)==1)));
    TTP1_GM(w1,1)  = TR.*mean(tmp2(find(FC_mask_GM(:,:,w1)==1))); TTP1_WM(w1,1)  = TR.*mean(tmp2(find(FC_mask_WM(:,:,w1)==1)));
    FWHM1_GM(w1,1) = TR.*mean(tmp3(find(FC_mask_GM(:,:,w1)==1))); FWHM1_WM(w1,1) = TR.*mean(tmp3(find(FC_mask_WM(:,:,w1)==1)));
    clear tmp1 tmp2 tmp3
end; clear w1
RH2_GM(1,1) = mean(RH(find(FC_mask_quads==1))); RH2_GM(2,1) = mean(RH(find(FC_mask_quads==2))); RH2_GM(3,1) = mean(RH(find(FC_mask_quads==3))); RH2_GM(4,1) = mean(RH(find(FC_mask_quads==4)));
RH2_WM(1,1) = mean(RH(find(FC_mask_quads==5))); RH2_WM(2,1) = mean(RH(find(FC_mask_quads==6))); RH2_WM(3,1) = mean(RH(find(FC_mask_quads==7))); RH2_WM(4,1) = mean(RH(find(FC_mask_quads==8)));
TTP2_GM(1,1) = TR.*mean(TTP(find(FC_mask_quads==1))); TTP2_GM(2,1) = TR.*mean(TTP(find(FC_mask_quads==2))); TTP2_GM(3,1) = TR.*mean(TTP(find(FC_mask_quads==3))); TTP2_GM(4,1) = TR.*mean(TTP(find(FC_mask_quads==4)));
TTP2_WM(1,1) = TR.*mean(TTP(find(FC_mask_quads==5))); TTP2_WM(2,1) = TR.*mean(TTP(find(FC_mask_quads==6))); TTP2_WM(3,1) = TR.*mean(TTP(find(FC_mask_quads==7))); TTP2_WM(4,1) = TR.*mean(TTP(find(FC_mask_quads==8)));
FWHM2_GM(1,1) = TR.*mean(FWHM(find(FC_mask_quads==1))); FWHM2_GM(2,1) = TR.*mean(FWHM(find(FC_mask_quads==2))); FWHM2_GM(3,1) = TR.*mean(FWHM(find(FC_mask_quads==3))); FWHM2_GM(4,1) = TR.*mean(FWHM(find(FC_mask_quads==4)));
FWHM2_WM(1,1) = TR.*mean(FWHM(find(FC_mask_quads==5))); FWHM2_WM(2,1) = TR.*mean(FWHM(find(FC_mask_quads==6))); FWHM2_WM(3,1) = TR.*mean(FWHM(find(FC_mask_quads==7))); FWHM2_WM(4,1) = TR.*mean(FWHM(find(FC_mask_quads==8)));
    % (1) = left ventral GM; (2) = right ventral GM; (3) = left dorsal GM; (4) = right dorsal GM
    % (5) = left ventral WM; (6) = right ventral WM; (7) = left dorsal WM; (8) = right dorsal WM
fighndl = figure('Position',[1 1 1280 800],'Color',[0.85,0.90,0.85],'InvertHardcopy','off','Visible','off'); % red [0.95,0.9,0.85], green [0.85,0.90,0.85], blue [0.89,0.94,0.99]
subplot2n(4,2,[1,2]), bar([zeros(size(RH1_GM)),RH1_GM,RH1_WM,zeros(size(RH1_WM))]'), ylabel('HRF response height (a.u.)'), xticklabels({'','GRAY MATTER response height','WHITE MATTER response height',''}),
    a = get(gca,'xticklabels'); set(gca,'xticklabels',a,'fontsize',12); clear a
    grid on,grid minor, ylim([min([RH1_GM(:);RH1_WM(:)])-(max([RH1_GM(:);RH1_WM(:)])-min([RH1_GM(:);RH1_WM(:)]))/2, max([RH1_GM(:);RH1_WM(:)])+(max([RH1_GM(:);RH1_WM(:)])-min([RH1_GM(:);RH1_WM(:)]))/2]),
    title(['Hemodynamic response function (HRF) RESPONSE HEIGHT across SLICES'],'Color',[0,0.55,0.69],'FontSize',16), legend(legstr,'Location','northeast','NumColumns',2,'FontSize',10)
    text(1.8,max([RH1_GM(:);RH1_WM(:)])+(max([RH1_GM(:);RH1_WM(:)])-min([RH1_GM(:);RH1_WM(:)]))/4, sprintf('mean = %.2f',mean(RH1_GM(:))),'Color',[0,0.55,0.69],'FontSize',15)
    text(2.8,max([RH1_GM(:);RH1_WM(:)])+(max([RH1_GM(:);RH1_WM(:)])-min([RH1_GM(:);RH1_WM(:)]))/4, sprintf('mean = %.2f',mean(RH1_WM(:))),'Color',[0.65,0.11,0.19],'FontSize',15)
subtightplot(4,2,3,0.05,[0.08,0.0525],[0.1,0.065]), bar([TTP1_GM,TTP1_WM]'), ylabel('TTP (in seconds)'), xticklabels({'GRAY MATTER time-to-peak','WHITE MATTER time-to-peak'}),
    a = get(gca,'xticklabels'); set(gca,'xticklabels',a,'fontsize',12); clear a
    grid on,grid minor, ylim([min([TTP1_GM(:);TTP1_WM(:)])-1, max([TTP1_GM(:);TTP1_WM(:)])+1]), title(['HRF time-to-peak (TTP) across SLICES'],'Color',[0,0.55,0.69],'FontSize',16),
    text(0.8,max([TTP1_GM(:);TTP1_WM(:)])+0.5, sprintf('mean = %.2f s',mean(TTP1_GM(:))),'Color',[0,0.55,0.69],'FontSize',15)
    text(1.8,max([TTP1_GM(:);TTP1_WM(:)])+0.5, sprintf('mean = %.2f s',mean(TTP1_WM(:))),'Color',[0.65,0.11,0.19],'FontSize',15)
subtightplot(4,2,4,0.05,[0.08,0.0525],[0.1,0.065]), bar([FWHM1_GM,FWHM1_WM]'), ylabel('FWHM (in seconds)'), xticklabels({'GRAY MATTER FWHM','WHITE MATTER FWHM'}),
    a = get(gca,'xticklabels'); set(gca,'xticklabels',a,'fontsize',12); clear a
    grid on,grid minor, ylim([min([FWHM1_GM(:);FWHM1_WM(:)])-0.5, max([FWHM1_GM(:);FWHM1_WM(:)])+0.5]), title(['HRF full-width at half max (FWHM) across SLICES'],'Color',[0,0.55,0.69],'FontSize',16), 
    text(0.8,max([FWHM1_GM(:);FWHM1_WM(:)])+0.25, sprintf('mean = %.2f s',mean(FWHM1_GM(:))),'Color',[0,0.55,0.69],'FontSize',15)
    text(1.8,max([FWHM1_GM(:);FWHM1_WM(:)])+0.25, sprintf('mean = %.2f s',mean(FWHM1_WM(:))),'Color',[0.65,0.11,0.19],'FontSize',15)
subplot2n(4,2,[5,6]), bar([zeros(size(RH2_GM)),RH2_GM,RH2_WM,zeros(size(RH2_WM))]'), ylabel('HRF response height (a.u.)'), xticklabels({'','GRAY MATTER response height','WHITE MATTER response height',''}),
    a = get(gca,'xticklabels'); set(gca,'xticklabels',a,'fontsize',12); clear a
    grid on,grid minor, ylim([min([RH2_GM(:);RH2_WM(:)])-(max([RH2_GM(:);RH2_WM(:)])-min([RH2_GM(:);RH2_WM(:)]))/2, max([RH2_GM(:);RH2_WM(:)])+(max([RH2_GM(:);RH2_WM(:)])-min([RH2_GM(:);RH2_WM(:)]))/2]),
    title(['HRF RESPONSE HEIGHT across QUADRANTS'],'Color',[0.65,0.11,0.19],'FontSize',16), legend({'left ventral','right ventral','left dorsal','right dorsal'},'Location','northeast','NumColumns',2,'FontSize',12)
    text(1.8,max([RH2_GM(:);RH2_WM(:)])+(max([RH2_GM(:);RH2_WM(:)])-min([RH2_GM(:);RH2_WM(:)]))/4, sprintf('mean = %.2f',mean(RH2_GM(:))),'Color',[0,0.55,0.69],'FontSize',15)
    text(2.8,max([RH2_GM(:);RH2_WM(:)])+(max([RH2_GM(:);RH2_WM(:)])-min([RH2_GM(:);RH2_WM(:)]))/4, sprintf('mean = %.2f',mean(RH2_WM(:))),'Color',[0.65,0.11,0.19],'FontSize',15)
subtightplot(4,2,7,0.05,[0.105,0.055],[0.1,0.065]), bar([TTP2_GM,TTP2_WM]'), ylabel('TTP (in seconds)'), xticklabels({'GRAY MATTER time-to-peak','WHITE MATTER time-to-peak'}),
    a = get(gca,'xticklabels'); set(gca,'xticklabels',a,'fontsize',12); clear a
    grid on,grid minor, ylim([min([TTP2_GM(:);TTP2_WM(:)])-1, max([TTP2_GM(:);TTP2_WM(:)])+1]), title(['HRF time-to-peak (TTP) across QUADRANTS'],'Color',[0.65,0.11,0.19],'FontSize',16),
    text(0.8,max([TTP2_GM(:);TTP2_WM(:)])+0.5, sprintf('mean = %.2f s',mean(TTP2_GM(:))),'Color',[0,0.55,0.69],'FontSize',15)
    text(1.8,max([TTP2_GM(:);TTP2_WM(:)])+0.5, sprintf('mean = %.2f s',mean(TTP2_WM(:))),'Color',[0.65,0.11,0.19],'FontSize',15)
subtightplot(4,2,8,0.05,[0.105,0.055],[0.1,0.065]), bar([FWHM2_GM,FWHM2_WM]'), ylabel('FWHM (in seconds)'), xticklabels({'GRAY MATTER FWHM','WHITE MATTER FWHM'}),
    a = get(gca,'xticklabels'); set(gca,'xticklabels',a,'fontsize',12); clear a
    grid on,grid minor, ylim([min([FWHM2_GM(:);FWHM2_WM(:)])-0.5, max([FWHM2_GM(:);FWHM2_WM(:)])+0.5]), title(['HRF full-width at half max (FWHM) across QUADRANTS'],'Color',[0.65,0.11,0.19],'FontSize',16), 
    text(0.8,max([FWHM2_GM(:);FWHM2_WM(:)])+0.25, sprintf('mean = %.2f s',mean(FWHM2_GM(:))),'Color',[0,0.55,0.69],'FontSize',15)
    text(1.8,max([FWHM2_GM(:);FWHM2_WM(:)])+0.25, sprintf('mean = %.2f s',mean(FWHM2_WM(:))),'Color',[0.65,0.11,0.19],'FontSize',15)
saveas(fighndl,[QCpath 'HRF_parameters_17deconv_allSlices' '.jpg'])
clear HRF1_GM HRF1_WM HRF2_GM HRF2_WM RH1_GM RH1_WM RH2_GM RH2_WM TTP1_GM TTP1_WM TTP2_WM TTP2_GM FWHM2_WM FWHM2_GM FWHM1_WM FWHM1_GM
close(fighndl)


F2g_=[]; mF2g_=[]; F2w_=[]; mF2w_=[]; Cg_=[]; Cw_=[]; F4_=[]; mF4_=[]; C2_=[];
if stp~=0, F6g_=[]; mF6g_=[]; F6w_=[]; mF6w_=[]; C3g_=[]; C3w_=[]; end
for i3 = 1 : siz3
    try waitbar((i3/siz3),wbar3,sprintf('17. QC plots: slice (%d) of (%d)',i3,siz3)); catch, end
    fprintf('17. QC plots: slice (%d) of (%d), subject (%d) of (%d), run (%d) of (%d)\n',i3,siz3,k,size(func_files,1),k2,size(func_files,2))
    
    if Smask_err==0
        scfMRItb_04_resplitData(base_dir_sub, fname, '_Smask', siz3) % if slice-wise data doesn't exist but combined 4D data exists, then split the data to generate the needed slice-wise data
        scfMRItb_04_unzipFile(base_dir_sub, fname, ['_slice' num2str(i3) '_Smask'])
        Smask1 = load_untouch_nii([base_dir_sub fname '_slice' num2str(i3) '_Smask.nii']); Smask1 = Smask1.img;
    else
        Smask1 = Smask1_(:,:,i3);
    end

    for i5=1:size(F1,4)
        tmp = F1(:,:,i3,i5);
        F2g(:,i5) = tmp(find(mask_GM2(:,:,i3)==1));
        F2w(:,i5) = tmp(find(mask_WM2(:,:,i3)==1));
        clear tmp
    end; clear i5
    [~,svar1] = sort(var(F2g,0,2)); [~,sdist1] = sort(svar1); sortorder1{i3,1} = sdist1;
    [~,svar2] = sort(var(F2w,0,2)); [~,sdist2] = sort(svar2); sortorder2{i3,1} = sdist2;
    
    fighndl = figure('Position',[1 1 1152 720],'Color',[0.95,0.9,0.85],'InvertHardcopy','off','Visible','off'); % red [0.95,0.9,0.85], green [0.85,0.90,0.85], blue [0.89,0.94,0.99]
    subplot2n(2,1,1), imagesc(F2g(sortorder1{i3},:)),colormap(cmap),colorbar, xlabel('time (in samples)'),ylabel('Gray matter voxels (sorted by variance)'),
        title(['Time series within the GRAY MATTER after "17-HRF-deconvolution" in slice ' num2str(i3) ' of ' num2str(siz3)],'Color',[0,0.55,0.69],'FontSize',14);
    subplot2n(2,1,2), imagesc(F2w(sortorder2{i3},:)),colormap(cmap),colorbar, xlabel('time (in samples)'),ylabel('White matter voxels (sorted by variance)'),
        title(['Time series within the WHITE MATTER after "17-HRF-deconvolution" in slice ' num2str(i3) ' of ' num2str(siz3)],'Color',[0.65,0.11,0.19],'FontSize',14);
    saveas(fighndl,[QCpath 'fMRI_ts_17deconv_slice' sprintf('%.2d',i3) '.jpg'])
    close(fighndl)
    
    scfMRItb_04_resplitData(base_dir_sub, fname, '', siz3) % if slice-wise data doesn't exist but combined 4D data exists, then split the data to generate the needed slice-wise data
	scfMRItb_04_unzipFile(base_dir_sub, fname, ['_slice' num2str(i3)])
    F3 = load_untouch_nii([base_dir_sub fname '_slice' num2str(i3) '.nii']); F3 = single(squeeze(F3.img));
    for i5=1:size(F3,3)
        tmp = F3(:,:,i5);
        F4(:,i5) = tmp(find(Smask1==1)); clear tmp
    end; clear i5 F3
    mF2g=mean(F2g,1); mF2w=mean(F2w,1);
    mF2g=zscore(mF2g); mF2w=zscore(mF2w);

    Cg = corrcoef(F2g'); Cg = uppertriangle(Cg);
    Cw = corrcoef(F2w'); Cw = uppertriangle(Cw);
    C2 = corrcoef(F4');  C2 = uppertriangle(C2);
    fighndl = figure('Position',[1 1 1152 720],'Color',[0.95,0.9,0.85],'InvertHardcopy','off','Visible','off'); % red [0.95,0.9,0.85], green [0.85,0.90,0.85], blue [0.89,0.94,0.99]
    subplot2n(3,1,1), histogram(C2,'EdgeColor','none','FaceColor','k','FaceAlpha',0.5,'Normalization','probability'); grid on,grid minor, xlim([-1 1]), hold on, xline(0,'Color','k','linewidth',1.5); hold off
      title(sprintf('Histogram of voxel-to-voxel correlations in UNPROCESSED RAW DATA (slice %d of %d)\n%.1f%% of the correlations were > 0  [ mean=%.2f, median=%.2f, SD=%.2f ]',i3,siz3,100*length(find(C2>0))/length(C2),mean(C2),median(C2),std(C2)),'Color','k','FontSize',14)
    subplot2n(3,1,2), histogram(Cg,'EdgeColor','none','FaceColor',[0,0.55,0.69],'FaceAlpha',0.5,'Normalization','probability'); grid on,grid minor, xlim([-1 1]), hold on, xline(0,'Color','k','linewidth',1.5); hold off
      title(sprintf('Histogram of voxel-to-voxel GRAY MATTER correlations AFTER 17-HRF-deconvolution (slice %d of %d)\n%.1f%% of the correlations were > 0  [ mean=%.2f, median=%.2f, SD=%.2f ]',i3,siz3,100*length(find(Cg>0))/length(Cg),mean(Cg),median(Cg),std(Cg)),'Color',[0,0.55,0.69],'FontSize',14)
    subplot2n(3,1,3), histogram(Cw,'EdgeColor','none','FaceColor',[0.65,0.11,0.19],'FaceAlpha',0.5,'Normalization','probability'); grid on,grid minor, xlim([-1 1]), hold on, xline(0,'Color','k','linewidth',1.5); hold off
      title(sprintf('Histogram of voxel-to-voxel WHITE MATTER correlations AFTER 17-HRF-deconvolution (slice %d of %d)\n%.1f%% of the correlations were > 0  [ mean=%.2f, median=%.2f, SD=%.2f ]',i3,siz3,100*length(find(Cw>0))/length(Cw),mean(Cw),median(Cw),std(Cw)),'Color',[0.65,0.11,0.19],'FontSize',14)
    saveas(fighndl,[QCpath 'hist_FC_compare_17deconv_slice' sprintf('%.2d',i3) '.jpg'])
    close(fighndl)
    
    if stp~=0
        msk=1;
        if exist('F5all','var')
            F5 = single(squeeze(F5all(:,:,i3,:)));
            msk=2;
        else
            if stp==1
                scfMRItb_04_resplitData(base_dir_sub, fname, '_warped', siz3) % if slice-wise data doesn't exist but combined 4D data exists, then split the data to generate the needed slice-wise data
                scfMRItb_04_unzipFile(base_dir_sub, fname, ['_slice' num2str(i3) '_warped'])
                F5 = load_untouch_nii([base_dir_sub fname '_slice' num2str(i3) '_warped.nii']); F5 = single(squeeze(F5.img));
            elseif stp==2
                scfMRItb_04_resplitData(base_dir_sub, fname, '_WM', siz3) % if slice-wise data doesn't exist but combined 4D data exists, then split the data to generate the needed slice-wise data
                scfMRItb_04_unzipFile(base_dir_sub, fname, ['_slice' num2str(i3) '_WM'])
                F5 = load_untouch_nii([base_dir_sub fname '_slice' num2str(i3) '_WM.nii']); F5 = single(squeeze(F5.img));
            elseif stp==3
                scfMRItb_04_resplitData(base_dir_sub, fname, ['_denoised' cutoff_str], siz3) % if slice-wise data doesn't exist but combined 4D data exists, then split the data to generate the needed slice-wise data
                scfMRItb_04_unzipFile(base_dir_sub, fname, ['_slice' num2str(i3) '_denoised' cutoff_str ''])
                F5 = load_untouch_nii([base_dir_sub fname '_slice' num2str(i3) '_denoised' cutoff_str '.nii']); F5 = single(squeeze(F5.img));
            elseif stp==4
                scfMRItb_04_resplitData(base_dir_sub, fname, '_cov', siz3) % if slice-wise data doesn't exist but combined 4D data exists, then split the data to generate the needed slice-wise data
                scfMRItb_04_unzipFile(base_dir_sub, fname, ['_slice' num2str(i3) '_cov'])
                F5 = load_untouch_nii([base_dir_sub fname '_slice' num2str(i3) '_cov.nii']); F5 = single(squeeze(F5.img));
            elseif stp==5
                scfMRItb_04_resplitData(base_dir_sub, fname, '_WMcov', siz3) % if slice-wise data doesn't exist but combined 4D data exists, then split the data to generate the needed slice-wise data
                scfMRItb_04_unzipFile(base_dir_sub, fname, ['_slice' num2str(i3) '_WMcov'])
                F5 = load_untouch_nii([base_dir_sub fname '_slice' num2str(i3) '_WMcov.nii']); F5 = single(squeeze(F5.img));
            elseif stp==6
                scfMRItb_04_resplitData(base_dir_sub, fname, ['_denoised' cutoff_str 'WM'], siz3) % if slice-wise data doesn't exist but combined 4D data exists, then split the data to generate the needed slice-wise data
                scfMRItb_04_unzipFile(base_dir_sub, fname, ['_slice' num2str(i3) '_denoised' cutoff_str 'WM'])
                F5 = load_untouch_nii([base_dir_sub fname '_slice' num2str(i3) '_denoised' cutoff_str 'WM.nii']); F5 = single(squeeze(F5.img));
            elseif stp==7
                scfMRItb_04_resplitData(base_dir_sub, fname, ['_denoised' cutoff_str 'cov'], siz3) % if slice-wise data doesn't exist but combined 4D data exists, then split the data to generate the needed slice-wise data
                scfMRItb_04_unzipFile(base_dir_sub, fname, ['_slice' num2str(i3) '_denoised' cutoff_str 'cov'])
                F5 = load_untouch_nii([base_dir_sub fname '_slice' num2str(i3) '_denoised' cutoff_str 'cov.nii']); F5 = single(squeeze(F5.img));
            elseif stp==8
                scfMRItb_04_resplitData(base_dir_sub, fname, ['_denoised' cutoff_str 'WMcov'], siz3) % if slice-wise data doesn't exist but combined 4D data exists, then split the data to generate the needed slice-wise data
                scfMRItb_04_unzipFile(base_dir_sub, fname, ['_slice' num2str(i3) '_denoised' cutoff_str 'WMcov'])
                F5 = load_untouch_nii([base_dir_sub fname '_slice' num2str(i3) '_denoised' cutoff_str 'WMcov.nii']); F5 = single(squeeze(F5.img));
            end
        end
        
        for i5=1:size(F5,3)
            tmp = F5(:,:,i5);
            if msk==1
                F6g(:,i5) = tmp(find(mask_GM(:,:,i3)==1));
                F6w(:,i5) = tmp(find(mask_WM(:,:,i3)==1));
            else
                F6g(:,i5) = tmp(find(mask_GM2(:,:,i3)==1));
                F6w(:,i5) = tmp(find(mask_WM2(:,:,i3)==1));
            end
            clear tmp
        end; clear i5 F5
        mF6g=mean(F6g,1); mF6w=mean(F6w,1); 
        mF6g=zscore(mF6g); mF6w=zscore(mF6w);
        fighndl = figure('Position',[1 1 1152 720],'Color',[0.95,0.9,0.85],'InvertHardcopy','off','Visible','off'); % red [0.95,0.9,0.85], green [0.85,0.90,0.85], blue [0.89,0.94,0.99]
        if (stp==1)&&(msk==1)
            subplot2n(4,1,1), plot(mF6g,'color','k','linewidth',1.5), hold on, plot(mF2g,'color',[0,0.55,0.69],'linewidth',0.2), ylabel('mean time series'),
                grid on,grid minor, xlim([1 length(mF2g)]),ylim([min(1.25.*[mF2g,mF6g]) 1.25.*max([mF2g,mF6g])]), title(['Unfiltered: normalized mean GRAY MATTER signal AFTER "11-func-anat-registration" (slice ' num2str(i3) ' of ' num2str(siz3) ')'],'Color','k','FontSize',14), legend({'Gray matter signal before deconvolution'},'FontSize',12,'TextColor','k','Location','best')
            subplot2n(4,1,3), plot(mF6w,'color','k','linewidth',1.5), hold on, plot(mF2w,'color',[0.65,0.11,0.19],'linewidth',0.2), ylabel('mean time series'),
                grid on,grid minor, xlim([1 length(mF2g)]),ylim([min(1.25.*[mF2w,mF6w]) 1.25.*max([mF2w,mF6w])]), title(['Unfiltered: normalized mean WHITE MATTER signal AFTER "11-func-anat-registration" (slice ' num2str(i3) ' of ' num2str(siz3) ')'],'Color','k','FontSize',14), legend({'White matter signal before deconvolution'},'FontSize',12,'TextColor','k','Location','best')
        elseif (stp==3)&&(msk==1)
            subplot2n(4,1,1), plot(mF6g,'color','k','linewidth',1.5), hold on, plot(mF2g,'color',[0,0.55,0.69],'linewidth',0.2), ylabel('mean time series'),
                grid on,grid minor, xlim([1 length(mF2g)]),ylim([min(1.25.*[mF2g,mF6g]) 1.25.*max([mF2g,mF6g])]), title(['Unfiltered: normalized mean GRAY MATTER signal AFTER "12-denoise3-CSFregress" (slice ' num2str(i3) ' of ' num2str(siz3) ')'],'Color','k','FontSize',14), legend({'Gray matter signal before deconvolution'},'FontSize',12,'TextColor','k','Location','best')
            subplot2n(4,1,3), plot(mF6w,'color','k','linewidth',1.5), hold on, plot(mF2w,'color',[0.65,0.11,0.19],'linewidth',0.2), ylabel('mean time series'),
                grid on,grid minor, xlim([1 length(mF2g)]),ylim([min(1.25.*[mF2w,mF6w]) 1.25.*max([mF2w,mF6w])]), title(['Unfiltered: normalized mean WHITE MATTER signal AFTER "12-denoise3-CSFregress" (slice ' num2str(i3) ' of ' num2str(siz3) ')'],'Color','k','FontSize',14), legend({'White matter signal before deconvolution'},'FontSize',12,'TextColor','k','Location','best')
        elseif ((stp==2)||(stp==6))&&(msk==1)
            subplot2n(4,1,1), plot(mF6g,'color','k','linewidth',1.5), hold on, plot(mF2g,'color',[0,0.55,0.69],'linewidth',0.2), ylabel('mean time series'),
                grid on,grid minor, xlim([1 length(mF2g)]),ylim([min(1.25.*[mF2g,mF6g]) 1.25.*max([mF2g,mF6g])]), title(['Unfiltered: normalized mean GRAY MATTER signal AFTER "13-denoise4-WMregress" (slice ' num2str(i3) ' of ' num2str(siz3) ')'],'Color','k','FontSize',14), legend({'Gray matter signal before deconvolution'},'FontSize',12,'TextColor','k','Location','best')
            subplot2n(4,1,3), plot(mF6w,'color','k','linewidth',1.5), hold on, plot(mF2w,'color',[0.65,0.11,0.19],'linewidth',0.2), ylabel('mean time series'),
                grid on,grid minor, xlim([1 length(mF2g)]),ylim([min(1.25.*[mF2w,mF6w]) 1.25.*max([mF2w,mF6w])]), title(['Unfiltered: normalized mean WHITE MATTER signal AFTER "13-denoise4-WMregress" (slice ' num2str(i3) ' of ' num2str(siz3) ')'],'Color','k','FontSize',14), legend({'White matter signal before deconvolution'},'FontSize',12,'TextColor','k','Location','best')
        elseif ((stp==4)||(stp==5)||(stp==7)||(stp==8))&&(msk==1)
            subplot2n(4,1,1), plot(mF6g,'color','k','linewidth',1.5), hold on, plot(mF2g,'color',[0,0.55,0.69],'linewidth',0.2), ylabel('mean time series'),
                grid on,grid minor, xlim([1 length(mF2g)]),ylim([min(1.25.*[mF2g,mF6g]) 1.25.*max([mF2g,mF6g])]), title(['Bandpass filtered: normalized mean GRAY MATTER signal AFTER "14-denoise5-COVregress" (slice ' num2str(i3) ' of ' num2str(siz3) ')'],'Color','k','FontSize',14), legend({'Gray matter signal before deconvolution'},'FontSize',12,'TextColor','k','Location','best')
            subplot2n(4,1,3), plot(mF6w,'color','k','linewidth',1.5), hold on, plot(mF2w,'color',[0.65,0.11,0.19],'linewidth',0.2), ylabel('mean time series'),
                grid on,grid minor, xlim([1 length(mF2g)]),ylim([min(1.25.*[mF2w,mF6w]) 1.25.*max([mF2w,mF6w])]), title(['Bandpass filtered: normalized mean WHITE MATTER signal AFTER "14-denoise5-COVregress" (slice ' num2str(i3) ' of ' num2str(siz3) ')'],'Color','k','FontSize',14), legend({'White matter signal before deconvolution'},'FontSize',12,'TextColor','k','Location','best')
        elseif (msk==2)
            subplot2n(4,1,1), plot(mF6g,'color','k','linewidth',1.5), hold on, plot(mF2g,'color',[0,0.55,0.69],'linewidth',0.2), ylabel('mean time series'),
                grid on,grid minor, xlim([1 length(mF2g)]),ylim([min(1.25.*[mF2g,mF6g]) 1.25.*max([mF2g,mF6g])]), title(['Bandpass filtered: normalized mean GRAY MATTER signal AFTER "15-bandpass-filter" (slice ' num2str(i3) ' of ' num2str(siz3) ')'],'Color','k','FontSize',14), legend({'Gray matter signal before deconvolution'},'FontSize',12,'TextColor','k','Location','best')
            subplot2n(4,1,3), plot(mF6w,'color','k','linewidth',1.5), hold on, plot(mF2w,'color',[0.65,0.11,0.19],'linewidth',0.2), ylabel('mean time series'),
                grid on,grid minor, xlim([1 length(mF2g)]),ylim([min(1.25.*[mF2w,mF6w]) 1.25.*max([mF2w,mF6w])]), title(['Bandpass filtered: normalized mean WHITE MATTER signal AFTER "15-bandpass-filter" (slice ' num2str(i3) ' of ' num2str(siz3) ')'],'Color','k','FontSize',14), legend({'White matter signal before deconvolution'},'FontSize',12,'TextColor','k','Location','best')
        end
        subplot2n(4,1,2), plot(mF2g,'color',[0,0.55,0.69],'linewidth',1.5), hold on, plot(mF6g,'color','k','linewidth',0.2), ylabel('mean time series'),
            grid on,grid minor, xlim([1 length(mF2g)]),ylim([min(1.25.*[mF2g,mF6g]) 1.25.*max([mF2g,mF6g])]), title(['Deconvolved bandpass filtered: normalized mean GRAY MATTER signal AFTER "17-HRF-deconvolution" [0.01-' num2str(bpf_hz_str) 'Hz] (slice ' num2str(i3) ' of ' num2str(siz3) ')'],'Color',[0,0.55,0.69],'FontSize',14), legend({'Gray matter signal after deconvolution'},'FontSize',12,'TextColor',[0,0.55,0.69],'Location','best')
        subplot2n(4,1,4), plot(mF2w,'color',[0.65,0.11,0.19],'linewidth',1.5), hold on, plot(mF6w,'color','k','linewidth',0.2), xlabel('time (in samples)'), ylabel('mean time series'),
            grid on,grid minor, xlim([1 length(mF2g)]),ylim([min(1.25.*[mF2w,mF6w]) 1.25.*max([mF2w,mF6w])]), title(['Deconvolved bandpass filtered: normalized mean WHITE MATTER signal AFTER "17-HRF-deconvolution" [0.01-' num2str(bpf_hz_str) 'Hz] (slice ' num2str(i3) ' of ' num2str(siz3) ')'],'Color',[0.65,0.11,0.19],'FontSize',14), legend({'White matter signal after deconvolution'},'FontSize',12,'TextColor',[0.65,0.11,0.19],'Location','best')
        saveas(fighndl,[QCpath 'mean_timeseries_15filter_vs_17deconv_slice' sprintf('%.2d',i3) '.jpg'])
        close(fighndl)
        
        C3g = corrcoef(F6g'); C3g = uppertriangle(C3g); C3w = corrcoef(F6w'); C3w = uppertriangle(C3w);
        fighndl = figure('Position',[1 1 1152 720],'Color',[0.95,0.9,0.85],'InvertHardcopy','off','Visible','off'); % red [0.95,0.9,0.85], green [0.85,0.90,0.85], blue [0.89,0.94,0.99]
        if (stp==1)&&(msk==1)
            subplot2n(4,1,1), histogram(C3g,'EdgeColor','none','FaceColor','k','FaceAlpha',0.5,'Normalization','probability'); grid on,grid minor, xlim([-1 1]), hold on, xline(0,'Color','k','linewidth',1.5); hold off
                title(sprintf('Histogram of voxel-to-voxel GRAY MATTER correlations AFTER 11-func-anat-registration (slice %d of %d)\n%.1f%% of the correlations were > 0  [ mean=%.2f, median=%.2f, SD=%.2f ]',i3,siz3,100*length(find(C3g>0))/length(C3g),mean(C3g),median(C3g),std(C3g)),'Color','k','FontSize',14)
            subplot2n(4,1,3), histogram(C3w,'EdgeColor','none','FaceColor','k','FaceAlpha',0.5,'Normalization','probability'); grid on,grid minor, xlim([-1 1]), hold on, xline(0,'Color','k','linewidth',1.5); hold off
                title(sprintf('WHITE MATTER: AFTER 11-func-anat-registration (slice %d of %d): %.1f%% of the correlations were > 0  [ mean=%.2f, median=%.2f, SD=%.2f ]',i3,siz3,100*length(find(C3w>0))/length(C3w),mean(C3w),median(C3w),std(C3w)),'Color','k','FontSize',14)
        elseif (stp==3)&&(msk==1)
            subplot2n(4,1,1), histogram(C3g,'EdgeColor','none','FaceColor','k','FaceAlpha',0.5,'Normalization','probability'); grid on,grid minor, xlim([-1 1]), hold on, xline(0,'Color','k','linewidth',1.5); hold off
                title(sprintf('Histogram of voxel-to-voxel GRAY MATTER correlations AFTER 12-denoise3-CSFregress (slice %d of %d)\n%.1f%% of the correlations were > 0  [ mean=%.2f, median=%.2f, SD=%.2f ]',i3,siz3,100*length(find(C3g>0))/length(C3g),mean(C3g),median(C3g),std(C3g)),'Color','k','FontSize',14)
            subplot2n(4,1,3), histogram(C3w,'EdgeColor','none','FaceColor','k','FaceAlpha',0.5,'Normalization','probability'); grid on,grid minor, xlim([-1 1]), hold on, xline(0,'Color','k','linewidth',1.5); hold off
                title(sprintf('WHITE MATTER: AFTER 12-denoise3-CSFregress (slice %d of %d): %.1f%% of the correlations were > 0  [ mean=%.2f, median=%.2f, SD=%.2f ]',i3,siz3,100*length(find(C3w>0))/length(C3w),mean(C3w),median(C3w),std(C3w)),'Color','k','FontSize',14)
        elseif ((stp==2)||(stp==6))&&(msk==1)
            subplot2n(4,1,1), histogram(C3g,'EdgeColor','none','FaceColor','k','FaceAlpha',0.5,'Normalization','probability'); grid on,grid minor, xlim([-1 1]), hold on, xline(0,'Color','k','linewidth',1.5); hold off
                title(sprintf('Histogram of voxel-to-voxel GRAY MATTER correlations AFTER 13-denoise4-WMregress (slice %d of %d)\n%.1f%% of the correlations were > 0  [ mean=%.2f, median=%.2f, SD=%.2f ]',i3,siz3,100*length(find(C3g>0))/length(C3g),mean(C3g),median(C3g),std(C3g)),'Color','k','FontSize',14)
            subplot2n(4,1,3), histogram(C3w,'EdgeColor','none','FaceColor','k','FaceAlpha',0.5,'Normalization','probability'); grid on,grid minor, xlim([-1 1]), hold on, xline(0,'Color','k','linewidth',1.5); hold off
                title(sprintf('WHITE MATTER: AFTER 13-denoise4-WMregress (slice %d of %d): %.1f%% of the correlations were > 0  [ mean=%.2f, median=%.2f, SD=%.2f ]',i3,siz3,100*length(find(C3w>0))/length(C3w),mean(C3w),median(C3w),std(C3w)),'Color','k','FontSize',14)
        elseif ((stp==4)||(stp==5)||(stp==7)||(stp==8))&&(msk==1)
            subplot2n(4,1,1), histogram(C3g,'EdgeColor','none','FaceColor','k','FaceAlpha',0.5,'Normalization','probability'); grid on,grid minor, xlim([-1 1]), hold on, xline(0,'Color','k','linewidth',1.5); hold off
                title(sprintf('Histogram of voxel-to-voxel GRAY MATTER correlations AFTER 14-denoise5-COVregress (slice %d of %d)\n%.1f%% of the correlations were > 0  [ mean=%.2f, median=%.2f, SD=%.2f ]',i3,siz3,100*length(find(C3g>0))/length(C3g),mean(C3g),median(C3g),std(C3g)),'Color','k','FontSize',14)
            subplot2n(4,1,3), histogram(C3w,'EdgeColor','none','FaceColor','k','FaceAlpha',0.5,'Normalization','probability'); grid on,grid minor, xlim([-1 1]), hold on, xline(0,'Color','k','linewidth',1.5); hold off
                title(sprintf('WHITE MATTER: AFTER 14-denoise5-COVregress (slice %d of %d): %.1f%% of the correlations were > 0  [ mean=%.2f, median=%.2f, SD=%.2f ]',i3,siz3,100*length(find(C3w>0))/length(C3w),mean(C3w),median(C3w),std(C3w)),'Color','k','FontSize',14)
        elseif (msk==2)
            subplot2n(4,1,1), histogram(C3g,'EdgeColor','none','FaceColor','k','FaceAlpha',0.5,'Normalization','probability'); grid on,grid minor, xlim([-1 1]), hold on, xline(0,'Color','k','linewidth',1.5); hold off
                title(sprintf('Histogram of voxel-to-voxel GRAY MATTER correlations AFTER 15-bandpass-filter (slice %d of %d)\n%.1f%% of the correlations were > 0  [ mean=%.2f, median=%.2f, SD=%.2f ]',i3,siz3,100*length(find(C3g>0))/length(C3g),mean(C3g),median(C3g),std(C3g)),'Color','k','FontSize',14)
            subplot2n(4,1,3), histogram(C3w,'EdgeColor','none','FaceColor','k','FaceAlpha',0.5,'Normalization','probability'); grid on,grid minor, xlim([-1 1]), hold on, xline(0,'Color','k','linewidth',1.5); hold off
                title(sprintf('WHITE MATTER: AFTER 15-bandpass-filter (slice %d of %d): %.1f%% of the correlations were > 0  [ mean=%.2f, median=%.2f, SD=%.2f ]',i3,siz3,100*length(find(C3w>0))/length(C3w),mean(C3w),median(C3w),std(C3w)),'Color','k','FontSize',14)
        end
        subplot2n(4,1,2), histogram(Cg,'EdgeColor','none','FaceColor',[0,0.55,0.69],'FaceAlpha',0.5,'Normalization','probability'); grid on,grid minor, xlim([-1 1]), hold on, xline(0,'Color','k','linewidth',1.5); hold off
            title(sprintf('GRAY MATTER: AFTER 17-HRF-deconvolution (slice %d of %d): %.1f%% of the correlations were > 0  [ mean=%.2f, median=%.2f, SD=%.2f ]',i3,siz3,100*length(find(Cg>0))/length(Cg),mean(Cg),median(Cg),std(Cg)),'Color',[0,0.55,0.69],'FontSize',14)
        subplot2n(4,1,4), histogram(Cw,'EdgeColor','none','FaceColor',[0.65,0.11,0.19],'FaceAlpha',0.5,'Normalization','probability'); grid on,grid minor, xlim([-1 1]), hold on, xline(0,'Color','k','linewidth',1.5); hold off
            title(sprintf('WHITE MATTER: AFTER 17-HRF-deconvolution (slice %d of %d): %.1f%% of the correlations were > 0  [ mean=%.2f, median=%.2f, SD=%.2f ]',i3,siz3,100*length(find(Cw>0))/length(Cw),mean(Cw),median(Cw),std(Cw)),'Color',[0.65,0.11,0.19],'FontSize',14)
        saveas(fighndl,[QCpath 'hist_FC_compare_15filter_17deconv_slice' sprintf('%.2d',i3) '.jpg'])
        close(fighndl)
        F6g_=cat(1,F6g_,F6g); mF6g_=cat(1,mF6g_,mF6g); C3g_=cat(1,C3g_,C3g);
        F6w_=cat(1,F6w_,F6w); mF6w_=cat(1,mF6w_,mF6w); C3w_=cat(1,C3w_,C3w);
        clear F6g F6w mF6g mF6w C3g C3w
    end
    
    F2g_=cat(1,F2g_,F2g); mF2g_=cat(1,mF2g_,mF2g); F2w_=cat(1,F2w_,F2w); mF2w_=cat(1,mF2w_,mF2w); Cg_=cat(1,Cg_,Cg); Cw_=cat(1,Cw_,Cw); F4_=cat(1,F4_,F4); C2_=cat(1,C2_,C2);
    clear F2g F2w F4 TR Smask1 svar1 svar2 mF2g mF2w mF4 Cg Cw C2
end


[~,svar1] = sort(var(F2g_,0,2)); [~,spsc1] = sort(meanPSC(F2g_));
[~,svar2] = sort(var(F2w_,0,2)); [~,spsc2] = sort(meanPSC(F2w_));
sortorder1b = sortorder1; sortorder1=[]; f1=0; %#ok<*ASGLU>
sortorder2b = sortorder2; sortorder2=[]; f2=0; %#ok<*ASGLU>
for im=1:length(sortorder1b)
    temp = sortorder1b{im}; temp = temp + f1; f1 = f1 + length(temp);
    sortorder1 = cat(1,sortorder1,temp); clear temp
    temp = sortorder2b{im}; temp = temp + f2; f2 = f2 + length(temp);
    sortorder2 = cat(1,sortorder2,temp); clear temp
end; clear im f1 f2
sortorder1 = spsc1; % options: spsc for percentage signal change, svar for variance, (comment out the line if you want to use distance from the center of the cord separately for each slice)
sortorder2 = spsc2; % options: spsc for percentage signal change, svar for variance, (comment out the line if you want to use distance from the center of the cord separately for each slice)
for im=max(sortorder1):-1:1, if isempty(find(sortorder1==im)), sortorder1(find(sortorder1>im)) = sortorder1(find(sortorder1>im))-1; end, end; clear im %#ok<*EFIND>
for im=max(sortorder2):-1:1, if isempty(find(sortorder2==im)), sortorder2(find(sortorder2>im)) = sortorder2(find(sortorder2>im))-1; end, end; clear im %#ok<*EFIND>

fighndl = figure('Position',[1 1 1152 720],'Color',[0.89,0.94,0.99],'InvertHardcopy','off','Visible','off'); % red [0.95,0.9,0.85], green [0.85,0.90,0.85], blue [0.89,0.94,0.99]
subplot2n(2,1,1), imagesc(F2g_(sortorder1,:)),colormap(cmap),colorbar, xlabel('time (in samples)'),ylabel(sprintf('voxels (sorted by mean percentage signal change;\nbottom row is highest)')),
  title(['Time series within the GRAY MATTER AFTER 17-HRF-deconvolution (all slices combined)'],'Color',[0,0.55,0.69],'FontSize',14);
subplot2n(2,1,2), imagesc(F2w_(sortorder2,:)),colormap(cmap),colorbar, xlabel('time (in samples)'),ylabel(sprintf('voxels (sorted by mean percentage signal change;\nbottom row is highest)')),
  title(['Time series within the WHITE MATTER AFTER 17-HRF-deconvolution (all slices combined)'],'Color',[0.65,0.11,0.19],'FontSize',14);
saveas(fighndl,[QCpath 'fMRI_ts_17deconv_allSlices' '.jpg'])
 close(fighndl)
mF2g_=mean(mF2g_,1); mF2w_=mean(mF2w_,1); mF4_=mean(mF4_,1);
fighndl = figure('Position',[1 1 1152 720],'Color',[0.89,0.94,0.99],'InvertHardcopy','off','Visible','off'); % red [0.95,0.9,0.85], green [0.85,0.90,0.85], blue [0.89,0.94,0.99]
subplot2n(3,1,1), histogram(C2_,'EdgeColor','none','FaceColor','k','FaceAlpha',0.5,'Normalization','probability'); grid on,grid minor, xlim([-1 1]), hold on, xline(0,'Color','k','linewidth',1.5); hold off
    title(sprintf('Histogram of voxel-to-voxel correlations in UNPROCESSED RAW DATA (all slices combined)\n%.1f%% of the correlations were > 0  [ mean=%.2f, median=%.2f, SD=%.2f ]',100*length(find(C2_>0))/length(C2_),mean(C2_),median(C2_),std(C2_)),'Color','k','FontSize',14)
subplot2n(3,1,2), histogram(Cg_,'EdgeColor','none','FaceColor',[0,0.55,0.69],'FaceAlpha',0.5,'Normalization','probability'); grid on,grid minor, xlim([-1 1]), hold on, xline(0,'Color','k','linewidth',1.5); hold off
    title(sprintf('Histogram of voxel-to-voxel GRAY MATTER correlations AFTER 17-HRF-deconvolution (all slices combined)\n%.1f%% of the correlations were > 0  [ mean=%.2f, median=%.2f, SD=%.2f ]',100*length(find(Cg_>0))/length(Cg_),mean(Cg_),median(Cg_),std(Cg_)),'Color',[0,0.55,0.69],'FontSize',14)
subplot2n(3,1,3), histogram(Cw_,'EdgeColor','none','FaceColor',[0.65,0.11,0.19],'FaceAlpha',0.5,'Normalization','probability'); grid on,grid minor, xlim([-1 1]), hold on, xline(0,'Color','k','linewidth',1.5); hold off
    title(sprintf('Histogram of voxel-to-voxel WHITE MATTER correlations AFTER 17-HRF-deconvolution (all slices combined)\n%.1f%% of the correlations were > 0  [ mean=%.2f, median=%.2f, SD=%.2f ]',100*length(find(Cw_>0))/length(Cw_),mean(Cw_),median(Cw_),std(Cw_)),'Color',[0.65,0.11,0.19],'FontSize',14)
saveas(fighndl,[QCpath 'hist_FC_compare_17deconv_allSlices' '.jpg'])
 close(fighndl)

if stp~=0
    mF6g_=mean(mF6g_,1); mF6w_=mean(mF6w_,1);
    fighndl = figure('Position',[1 1 1152 720],'Color',[0.89,0.94,0.99],'InvertHardcopy','off','Visible','off'); % red [0.95,0.9,0.85], green [0.85,0.90,0.85], blue [0.89,0.94,0.99]
    if (stp==1)&&(msk==1)
    	subplot2n(4,1,1), plot(mF6g_,'color','k','linewidth',1.5), hold on, plot(mF2g_,'color',[0,0.55,0.69],'linewidth',0.2), ylabel('mean time series'),
            grid on,grid minor, xlim([1 length(mF2g_)]),ylim([1.25.*min([mF2g_,mF6g_]) 1.25.*max([mF2g_,mF6g_])]), title(['Normalized mean GRAY MATTER signal AFTER "11-func-anat-registration" (all slices averaged)'],'Color','k','FontSize',14), legend({'Gray matter signal before deconvolution'},'FontSize',12,'TextColor','k','Location','best')
        subplot2n(4,1,3), plot(mF6w_,'color','k','linewidth',1.5), hold on, plot(mF2w_,'color',[0.65,0.11,0.19],'linewidth',0.2), ylabel('mean time series'),
            grid on,grid minor, xlim([1 length(mF2g_)]),ylim([1.25.*min([mF2w_,mF6w_]) 1.25.*max([mF2w_,mF6w_])]), title(['Normalized mean WHITE MATTER signal AFTER "11-func-anat-registration" (all slices averaged)'],'Color','k','FontSize',14), legend({'White matter signal before deconvolution'},'FontSize',12,'TextColor','k','Location','best')
    elseif (stp==3)&&(msk==1)
    	subplot2n(4,1,1), plot(mF6g_,'color','k','linewidth',1.5), hold on, plot(mF2g_,'color',[0,0.55,0.69],'linewidth',0.2), ylabel('mean time series'),
            grid on,grid minor, xlim([1 length(mF2g_)]),ylim([1.25.*min([mF2g_,mF6g_]) 1.25.*max([mF2g_,mF6g_])]), title(['Normalized mean GRAY MATTER signal AFTER "12-DENOISE3-CSFregress" (all slices averaged)'],'Color','k','FontSize',14), legend({'Gray matter signal before deconvolution'},'FontSize',12,'TextColor','k','Location','best')
        subplot2n(4,1,3), plot(mF6w_,'color','k','linewidth',1.5), hold on, plot(mF2w_,'color',[0.65,0.11,0.19],'linewidth',0.2), ylabel('mean time series'),
            grid on,grid minor, xlim([1 length(mF2g_)]),ylim([1.25.*min([mF2w_,mF6w_]) 1.25.*max([mF2w_,mF6w_])]), title(['Normalized mean WHITE MATTER signal AFTER "12-DENOISE3-CSFregress" (all slices averaged)'],'Color','k','FontSize',14), legend({'White matter signal before deconvolution'},'FontSize',12,'TextColor','k','Location','best')
    elseif ((stp==2)||(stp==6))&&(msk==1)
    	subplot2n(4,1,1), plot(mF6g_,'color','k','linewidth',1.5), hold on, plot(mF2g_,'color',[0,0.55,0.69],'linewidth',0.2), ylabel('mean time series'),
            grid on,grid minor, xlim([1 length(mF2g_)]),ylim([1.25.*min([mF2g_,mF6g_]) 1.25.*max([mF2g_,mF6g_])]), title(['Normalized mean GRAY MATTER signal AFTER "13-DENOISE4-WMregress" (all slices averaged)'],'Color','k','FontSize',14), legend({'Gray matter signal before deconvolution'},'FontSize',12,'TextColor','k','Location','best')
        subplot2n(4,1,3), plot(mF6w_,'color','k','linewidth',1.5), hold on, plot(mF2w_,'color',[0.65,0.11,0.19],'linewidth',0.2), ylabel('mean time series'),
            grid on,grid minor, xlim([1 length(mF2g_)]),ylim([1.25.*min([mF2w_,mF6w_]) 1.25.*max([mF2w_,mF6w_])]), title(['Normalized mean WHITE MATTER signal AFTER "13-DENOISE4-WMregress" (all slices averaged)'],'Color','k','FontSize',14), legend({'White matter signal before deconvolution'},'FontSize',12,'TextColor','k','Location','best')
    elseif ((stp==4)||(stp==5)||(stp==7)||(stp==8))&&(msk==1)
    	subplot2n(4,1,1), plot(mF6g_,'color','k','linewidth',1.5), hold on, plot(mF2g_,'color',[0,0.55,0.69],'linewidth',0.2), ylabel('mean time series'),
            grid on,grid minor, xlim([1 length(mF2g_)]),ylim([1.25.*min([mF2g_,mF6g_]) 1.25.*max([mF2g_,mF6g_])]), title(['Normalized mean GRAY MATTER signal AFTER "14-DENOISE5-COVregress" (all slices averaged)'],'Color','k','FontSize',14), legend({'Gray matter signal before deconvolution'},'FontSize',12,'TextColor','k','Location','best')
        subplot2n(4,1,3), plot(mF6w_,'color','k','linewidth',1.5), hold on, plot(mF2w_,'color',[0.65,0.11,0.19],'linewidth',0.2), ylabel('mean time series'),
            grid on,grid minor, xlim([1 length(mF2g_)]),ylim([1.25.*min([mF2w_,mF6w_]) 1.25.*max([mF2w_,mF6w_])]), title(['Normalized mean WHITE MATTER signal AFTER "14-DENOISE5-COVregress" (all slices averaged)'],'Color','k','FontSize',14), legend({'White matter signal before deconvolution'},'FontSize',12,'TextColor','k','Location','best')
    elseif (msk==2)
    	subplot2n(4,1,1), plot(mF6g_,'color','k','linewidth',1.5), hold on, plot(mF2g_,'color',[0,0.55,0.69],'linewidth',0.2), ylabel('mean time series'),
            grid on,grid minor, xlim([1 length(mF2g_)]),ylim([1.25.*min([mF2g_,mF6g_]) 1.25.*max([mF2g_,mF6g_])]), title(['Normalized mean GRAY MATTER signal AFTER "15-bandpass-filter" (all slices averaged)'],'Color','k','FontSize',14), legend({'Gray matter signal before deconvolution'},'FontSize',12,'TextColor','k','Location','best')
        subplot2n(4,1,3), plot(mF6w_,'color','k','linewidth',1.5), hold on, plot(mF2w_,'color',[0.65,0.11,0.19],'linewidth',0.2), ylabel('mean time series'),
            grid on,grid minor, xlim([1 length(mF2g_)]),ylim([1.25.*min([mF2w_,mF6w_]) 1.25.*max([mF2w_,mF6w_])]), title(['Normalized mean WHITE MATTER signal AFTER "15-bandpass-filter" (all slices averaged)'],'Color','k','FontSize',14), legend({'White matter signal before deconvolution'},'FontSize',12,'TextColor','k','Location','best')
    end
    subplot2n(4,1,2), plot(mF2g_,'color',[0,0.55,0.69],'linewidth',1.5), hold on, plot(mF6g_,'color','k','linewidth',0.2), ylabel('mean time series'),
        grid on,grid minor, xlim([1 length(mF2g_)]),ylim([1.25.*min([mF2g_,mF6g_]) 1.25.*max([mF2g_,mF6g_])]), title(['Normalized mean GRAY MATTER signal AFTER "17-HRF-deconvolution" (all slices averaged)'],'Color',[0,0.55,0.69],'FontSize',14), legend({'Gray matter signal after deconvolution'},'FontSize',12,'TextColor',[0,0.55,0.69],'Location','best')
    subplot2n(4,1,4), plot(mF2w_,'color',[0.65,0.11,0.19],'linewidth',1.5), hold on, plot(mF6w_,'color','k','linewidth',0.2), ylabel('mean time series'),
        grid on,grid minor, xlim([1 length(mF2g_)]),ylim([1.25.*min([mF2w_,mF6w_]) 1.25.*max([mF2w_,mF6w_])]), title(['Normalized mean WHITE MATTER signal AFTER "17-HRF-deconvolution" (all slices averaged)'],'Color',[0.65,0.11,0.19],'FontSize',14), legend({'White matter signal after deconvolution'},'FontSize',12,'TextColor',[0.65,0.11,0.19],'Location','best')
    saveas(fighndl,[QCpath 'mean_timeseries_15filter_vs_17deconv_allSlices' '.jpg'])
     close(fighndl)
    
    fighndl = figure('Position',[1 1 1152 720],'Color',[0.89,0.94,0.99],'InvertHardcopy','off','Visible','off'); % red [0.95,0.9,0.85], green [0.85,0.90,0.85], blue [0.89,0.94,0.99]
    if (stp==1)&&(msk==1)
    	subplot2n(4,1,1), histogram(C3g_,'EdgeColor','none','FaceColor','k','FaceAlpha',0.5,'Normalization','probability'); grid on,grid minor, xlim([-1 1]), hold on, xline(0,'Color','k','linewidth',1.5); hold off
            title(sprintf('Histogram of voxel-to-voxel GRAY MATTER correlations AFTER 11-registration (all slices combined)\n%.1f%% of the correlations were > 0  [ mean=%.2f, median=%.2f, SD=%.2f ]',100*length(find(C3g_>0))/length(C3g_),mean(C3g_),median(C3g_),std(C3g_)),'Color','k','FontSize',14)
        subplot2n(4,1,3), histogram(C3w_,'EdgeColor','none','FaceColor','k','FaceAlpha',0.5,'Normalization','probability'); grid on,grid minor, xlim([-1 1]), hold on, xline(0,'Color','k','linewidth',1.5); hold off
            title(sprintf('WHITE MATTER: AFTER 11-registration (all slices): %.1f%% of the correlations were > 0  [ mean=%.2f, median=%.2f, SD=%.2f ]',100*length(find(C3w_>0))/length(C3w_),mean(C3w_),median(C3w_),std(C3w_)),'Color','k','FontSize',14)
    elseif (stp==3)&&(msk==1)
    	subplot2n(4,1,1), histogram(C3g_,'EdgeColor','none','FaceColor','k','FaceAlpha',0.5,'Normalization','probability'); grid on,grid minor, xlim([-1 1]), hold on, xline(0,'Color','k','linewidth',1.5); hold off
            title(sprintf('Histogram of voxel-to-voxel GRAY MATTER correlations AFTER 12-denoise3-CSFregress (all slices combined)\n%.1f%% of the correlations were > 0  [ mean=%.2f, median=%.2f, SD=%.2f ]',100*length(find(C3g_>0))/length(C3g_),mean(C3g_),median(C3g_),std(C3g_)),'Color','k','FontSize',14)
        subplot2n(4,1,3), histogram(C3w_,'EdgeColor','none','FaceColor','k','FaceAlpha',0.5,'Normalization','probability'); grid on,grid minor, xlim([-1 1]), hold on, xline(0,'Color','k','linewidth',1.5); hold off
            title(sprintf('WHITE MATTER: AFTER 12-denoise3-CSFregress (all slices): %.1f%% of the correlations were > 0  [ mean=%.2f, median=%.2f, SD=%.2f ]',100*length(find(C3w_>0))/length(C3w_),mean(C3w_),median(C3w_),std(C3w_)),'Color','k','FontSize',14)
    elseif ((stp==2)||(stp==6))&&(msk==1)
    	subplot2n(4,1,1), histogram(C3g_,'EdgeColor','none','FaceColor','k','FaceAlpha',0.5,'Normalization','probability'); grid on,grid minor, xlim([-1 1]), hold on, xline(0,'Color','k','linewidth',1.5); hold off
            title(sprintf('Histogram of voxel-to-voxel GRAY MATTER correlations AFTER 13-denoise4-WMregress (all slices combined)\n%.1f%% of the correlations were > 0  [ mean=%.2f, median=%.2f, SD=%.2f ]',100*length(find(C3g_>0))/length(C3g_),mean(C3g_),median(C3g_),std(C3g_)),'Color','k','FontSize',14)
        subplot2n(4,1,3), histogram(C3w_,'EdgeColor','none','FaceColor','k','FaceAlpha',0.5,'Normalization','probability'); grid on,grid minor, xlim([-1 1]), hold on, xline(0,'Color','k','linewidth',1.5); hold off
            title(sprintf('WHITE MATTER: AFTER 13-denoise4-WMregress (all slices): %.1f%% of the correlations were > 0  [ mean=%.2f, median=%.2f, SD=%.2f ]',100*length(find(C3w_>0))/length(C3w_),mean(C3w_),median(C3w_),std(C3w_)),'Color','k','FontSize',14)
    elseif ((stp==4)||(stp==5)||(stp==7)||(stp==8))&&(msk==1)
    	subplot2n(4,1,1), histogram(C3g_,'EdgeColor','none','FaceColor','k','FaceAlpha',0.5,'Normalization','probability'); grid on,grid minor, xlim([-1 1]), hold on, xline(0,'Color','k','linewidth',1.5); hold off
            title(sprintf('Histogram of voxel-to-voxel GRAY MATTER correlations AFTER 14-denoise5-COVregress (all slices combined)\n%.1f%% of the correlations were > 0  [ mean=%.2f, median=%.2f, SD=%.2f ]',100*length(find(C3g_>0))/length(C3g_),mean(C3g_),median(C3g_),std(C3g_)),'Color','k','FontSize',14)
        subplot2n(4,1,3), histogram(C3w_,'EdgeColor','none','FaceColor','k','FaceAlpha',0.5,'Normalization','probability'); grid on,grid minor, xlim([-1 1]), hold on, xline(0,'Color','k','linewidth',1.5); hold off
            title(sprintf('WHITE MATTER: AFTER 14-denoise5-COVregress (all slices): %.1f%% of the correlations were > 0  [ mean=%.2f, median=%.2f, SD=%.2f ]',100*length(find(C3w_>0))/length(C3w_),mean(C3w_),median(C3w_),std(C3w_)),'Color','k','FontSize',14)
    elseif (msk==2)
    	subplot2n(4,1,1), histogram(C3g_,'EdgeColor','none','FaceColor','k','FaceAlpha',0.5,'Normalization','probability'); grid on,grid minor, xlim([-1 1]), hold on, xline(0,'Color','k','linewidth',1.5); hold off
            title(sprintf('Histogram of voxel-to-voxel GRAY MATTER correlations AFTER 15-bandpass-filter (all slices combined)\n%.1f%% of the correlations were > 0  [ mean=%.2f, median=%.2f, SD=%.2f ]',100*length(find(C3g_>0))/length(C3g_),mean(C3g_),median(C3g_),std(C3g_)),'Color','k','FontSize',14)
        subplot2n(4,1,3), histogram(C3w_,'EdgeColor','none','FaceColor','k','FaceAlpha',0.5,'Normalization','probability'); grid on,grid minor, xlim([-1 1]), hold on, xline(0,'Color','k','linewidth',1.5); hold off
            title(sprintf('WHITE MATTER: AFTER 15-bandpass-filter (all slices): %.1f%% of the correlations were > 0  [ mean=%.2f, median=%.2f, SD=%.2f ]',100*length(find(C3w_>0))/length(C3w_),mean(C3w_),median(C3w_),std(C3w_)),'Color','k','FontSize',14)
    end
    subplot2n(4,1,2), histogram(Cg_,'EdgeColor','none','FaceColor',[0,0.55,0.69],'FaceAlpha',0.5,'Normalization','probability'); grid on,grid minor, xlim([-1 1]), hold on, xline(0,'Color','k','linewidth',1.5); hold off
       title(sprintf('GRAY MATTER: AFTER 17-HRF-deconvolution (all slices): %.1f%% of correlations > 0  [ mean=%.2f, median=%.2f, SD=%.2f ]',100*length(find(Cg_>0))/length(Cg_),mean(Cg_),median(Cg_),std(Cg_)),'Color',[0,0.55,0.69],'FontSize',14)
    subplot2n(4,1,4), histogram(Cw_,'EdgeColor','none','FaceColor',[0.65,0.11,0.19],'FaceAlpha',0.5,'Normalization','probability'); grid on,grid minor, xlim([-1 1]), hold on, xline(0,'Color','k','linewidth',1.5); hold off
       title(sprintf('WHITE MATTER: AFTER 17-HRF-deconvolution (all slices): %.1f%% of the correlations were > 0  [ mean=%.2f, median=%.2f, SD=%.2f ]',100*length(find(Cw_>0))/length(Cw_),mean(Cw_),median(Cw_),std(Cw_)),'Color',[0.65,0.11,0.19],'FontSize',14)
    saveas(fighndl,[QCpath 'hist_FC_compare_15filter_17deconv_allSlices' '.jpg'])
     close(fighndl)
end
clear svar1 spsc1 svar2 spsc2 sortorder1 sortorder2 F2g_ mF2g_ F2w_ mF2w_ Cg_ Cw_ F4_ mF4_ C2_ F6g_ F6w_ mF6g_ mF6w_ C3g_ C3w_

    catch
    end
end

%% tSNR maps
if (choices_preproc(5)==1)||(choices_preproc(6)==1)||(choices_preproc(8)==1)||(choices_preproc(9)==1)||(choices_preproc(11)==1)||(choices_preproc(12)==1)||(choices_preproc(13)==1)||(choices_preproc(14)==1)
clear QCpath save_videos fighndl stp
if ~(exist([base_dir_sub 'QC' '/tSNR_graphs'],'dir'))
    mkdir([base_dir_sub 'QC' '/tSNR_graphs'])
end
if Rns==0
    QCpath = [base_dir_sub 'QC' '/tSNR_graphs/'];
else
    if ~(exist([base_dir_sub 'QC' '/tSNR_graphs' '/run' num2str(Rns)],'dir'))
        mkdir([base_dir_sub 'QC' '/tSNR_graphs' '/run' num2str(Rns)])
    end
    QCpath = [base_dir_sub 'QC' '/tSNR_graphs' '/run' num2str(Rns) '/'];
end


try
if Rns==0
    load([base_dir_sub 'tSNR_across_preproc_steps.mat'])
else
    load([base_dir_sub 'tSNR_across_preproc_steps_run' num2str(Rns) '.mat'])
end

if ~(exist('F_tsnr','var'))
    scfMRItb_04_unzipFile(base_dir_sub, fname, '_mask_NS')
    eval(['load ' base_dir_sub fname '_mask_NS' '.mat;']); mask_NS = mask_NS.img;
    mask_CSF = load_untouch_nii([base_dir_sub fname_anat '_mask_CSF.nii']); mask_CSF = mask_CSF.img;
    mask_all = or(or(mask_GM,mask_WM),mask_CSF);
    if choices_preproc(5)==1 % raw data
        disp(sprintf('Calculating tSNR across pre-processing steps: raw data'))
        suffix = '';
        [tSNR_graph_SC(5,1), tSNR_graph_slicewise(:,1), F_tsnr{5,1}] = calc_tsnr(base_dir_sub, fname, suffix, ~mask_NS); clear suffix
    end
    if choices_preproc(6)==1 % after denoise1
        disp(sprintf('Calculating tSNR across pre-processing steps: step 6 (denoise1)'))
        suffix = '_denoised_before_MC80';
        [tSNR_graph_SC(6,1), tSNR_graph_slicewise(:,2), F_tsnr{6,1}] = calc_tsnr(base_dir_sub, fname, suffix, ~mask_NS); clear suffix
    end
    if choices_preproc(8)==1 % after motion correction
        disp(sprintf('Calculating tSNR across pre-processing steps: step 8 (motion correction)'))
        suffix = '_MC';
        [tSNR_graph_SC(8,1), tSNR_graph_slicewise(:,2), F_tsnr{8,1}] = calc_tsnr(base_dir_sub, fname, suffix, ~mask_NS); clear suffix
    end
    if choices_preproc(9)==1 % after denoise2 (RETROICOR)
        disp(sprintf('Calculating tSNR across pre-processing steps: step 9 (denoise2)'))
        suffix = '_MC_ricor';
        [tSNR_graph_SC(9,1), tSNR_graph_slicewise(:,2), F_tsnr{9,1}] = calc_tsnr(base_dir_sub, fname, suffix, ~mask_NS); clear suffix
    end
    if choices_preproc(11)==1 % after co-registration
        disp(sprintf('Calculating tSNR across pre-processing steps: step 11 (co-registration)'))
        suffix = '_warped';
        [tSNR_graph_SC(11,1), tSNR_graph_slicewise(:,2)] = calc_tsnr(base_dir_sub, fname, suffix, mask_all);
        [tSNR_graph_GM(11,1), tSNR_graph_slicewise(:,3), F_tsnr{11,1}] = calc_tsnr(base_dir_sub, fname, suffix, mask_GM);
        [tSNR_graph_WM(11,1), tSNR_graph_slicewise(:,4), F_tsnr{11,2}] = calc_tsnr(base_dir_sub, fname, suffix, mask_WM); clear suffix
    end
    if choices_preproc(12)==1 % after denoise3 (CSF regression)
        disp(sprintf('Calculating tSNR across pre-processing steps: step 12 (denoise3)'))
        suffix = '_denoised50';
        [tSNR_graph_SC(12,1), tSNR_graph_slicewise(:,2)] = calc_tsnr(base_dir_sub, fname, suffix, mask_all);
        [tSNR_graph_GM(12,1), tSNR_graph_slicewise(:,3), F_tsnr{12,1}] = calc_tsnr(base_dir_sub, fname, suffix, mask_GM);
        [tSNR_graph_WM(12,1), tSNR_graph_slicewise(:,4), F_tsnr{12,2}] = calc_tsnr(base_dir_sub, fname, suffix, mask_WM); clear suffix
    end
    if choices_preproc(13)==1 % after denoise4 (WM regression)
        disp(sprintf('Calculating tSNR across pre-processing steps: step 13 (denoise4)'))
        suffix = '_denoised50WM';
        [tSNR_graph_SC(13,1), tSNR_graph_slicewise(:,2)] = calc_tsnr(base_dir_sub, fname, suffix, mask_all);
        [tSNR_graph_GM(13,1), tSNR_graph_slicewise(:,3), F_tsnr{13,1}] = calc_tsnr(base_dir_sub, fname, suffix, mask_GM);
        [tSNR_graph_WM(13,1), tSNR_graph_slicewise(:,4), F_tsnr{13,2}] = calc_tsnr(base_dir_sub, fname, suffix, mask_WM); clear suffix
    end
    if choices_preproc(14)==1 % after denoise5 (additional covariates regression)
        disp(sprintf('Calculating tSNR across pre-processing steps: step 14 (denoise5)'))
        suffix = '_denoised50WMcov';
        [tSNR_graph_SC(14,1), tSNR_graph_slicewise(:,2)] = calc_tsnr(base_dir_sub, fname, suffix, mask_all);
        [tSNR_graph_GM(14,1), tSNR_graph_slicewise(:,3), F_tsnr{14,1}] = calc_tsnr(base_dir_sub, fname, suffix, mask_GM);
        [tSNR_graph_WM(14,1), tSNR_graph_slicewise(:,4), F_tsnr{14,2}] = calc_tsnr(base_dir_sub, fname, suffix, mask_WM); clear suffix
    end
end

%%% generate tSNR maps
temp=[];
for i=1:size(F_tsnr,1)
    for j=1:size(F_tsnr,2)
        temp2 = F_tsnr{i,j};
        temp = cat(1,temp,dezero(temp2(:)));
        clear temp2
    end
end; clear i j
climit = max(temp); clear temp
if choices_preproc(5)==1 % raw data
    disp(sprintf('Generating tSNR maps across pre-processing steps: raw data'))
    savename = [QCpath 'tSNR_maps_4_rawdata' '.jpg'];
    figtitle = sprintf('tSNR maps (slice 1 through %d) : raw data',size(mask_GM,3));
    gen_tsnr_maps(F_tsnr{5}, savename, figtitle, climit, tSNR_graph_SC(5)); clear savename figtitle
end
if choices_preproc(6)==1 % after denoise1
    disp(sprintf('Generating tSNR maps across pre-processing steps: step 6 (denoise1)'))
    savename = [QCpath 'tSNR_maps_6_denoise1' '.jpg'];
    figtitle = sprintf('tSNR maps (slice 1 through %d) after not-cord regression (denoise1)',size(mask_GM,3));
    gen_tsnr_maps(F_tsnr{6}, savename, figtitle, climit, tSNR_graph_SC(6)); clear savename figtitle
end
if choices_preproc(8)==1 % after motion correction
    disp(sprintf('Generating tSNR maps across pre-processing steps: step 8 (motion correction)'))
    savename = [QCpath 'tSNR_maps_8_motion_correction' '.jpg'];
    figtitle = sprintf('tSNR maps (slice 1 through %d) after motion correction',size(mask_GM,3));
    gen_tsnr_maps(F_tsnr{8}, savename, figtitle, climit, tSNR_graph_SC(8)); clear savename figtitle
end
if choices_preproc(9)==1 % after denoise2 (RETROICOR)
    disp(sprintf('Generating tSNR maps across pre-processing steps: step 9 (denoise2)'))
    savename = [QCpath 'tSNR_maps_9_denoise2' '.jpg'];
    figtitle = sprintf('tSNR maps (slice 1 through %d) after RETROICOR (denoise2)',size(mask_GM,3));
    gen_tsnr_maps(F_tsnr{9}, savename, figtitle, climit, tSNR_graph_SC(9)); clear savename figtitle
end
if choices_preproc(11)==1 % after co-registration
    disp(sprintf('Generating tSNR maps across pre-processing steps: step 11 (co-registration)'))
    savename = [QCpath 'tSNR_maps_11_coreg_GM' '.jpg'];
    figtitle = sprintf('tSNR maps (slice 1 through %d) in the gray matter after co-registration',size(mask_GM,3));
    gen_tsnr_maps(F_tsnr{11,1}, savename, figtitle, climit, tSNR_graph_GM(11)); clear savename figtitle
    savename = [QCpath 'tSNR_maps_11_coreg_WM' '.jpg'];
    figtitle = sprintf('tSNR maps (slice 1 through %d) in the white matter after co-registration',size(mask_GM,3));
    gen_tsnr_maps(F_tsnr{11,2}, savename, figtitle, climit, tSNR_graph_WM(11)); clear savename figtitle
end
if choices_preproc(12)==1 % after denoise3 (CSF regression)
    disp(sprintf('Generating tSNR maps across pre-processing steps: step 12 (denoise3)'))
    savename = [QCpath 'tSNR_maps_12_CSFregress_GM' '.jpg'];
    figtitle = sprintf('tSNR maps (slice 1 through %d) in the gray matter after CSF regression (denoise3)',size(mask_GM,3));
    gen_tsnr_maps(F_tsnr{12,1}, savename, figtitle, climit, tSNR_graph_GM(12)); clear savename figtitle
    savename = [QCpath 'tSNR_maps_12_CSFregress_WM' '.jpg'];
    figtitle = sprintf('tSNR maps (slice 1 through %d) in the white matter after CSF regression (denoise3)',size(mask_GM,3));
    gen_tsnr_maps(F_tsnr{12,2}, savename, figtitle, climit, tSNR_graph_WM(12)); clear savename figtitle
end
if choices_preproc(13)==1 % after denoise4 (WM regression)
    disp(sprintf('Generating tSNR maps across pre-processing steps: step 13 (denoise4)'))
    savename = [QCpath 'tSNR_maps_13_WMregress_GM' '.jpg'];
    figtitle = sprintf('tSNR maps (slice 1 through %d) in the gray matter after WM regression (denoise4)',size(mask_GM,3));
    gen_tsnr_maps(F_tsnr{13,1}, savename, figtitle, climit, tSNR_graph_GM(13)); clear savename figtitle
    savename = [QCpath 'tSNR_maps_13_WMregress_WM' '.jpg'];
    figtitle = sprintf('tSNR maps (slice 1 through %d) in the white matter after WM regression (denoise4)',size(mask_GM,3));
    gen_tsnr_maps(F_tsnr{13,2}, savename, figtitle, climit, tSNR_graph_WM(13)); clear savename figtitle
end
if choices_preproc(14)==1 % after denoise5 (additional covariates regression)
    disp(sprintf('Generating tSNR maps across pre-processing steps: step 14 (denoise5)'))
    savename = [QCpath 'tSNR_maps_14_COVregress_GM' '.jpg'];
    figtitle = sprintf('tSNR maps (slice 1 through %d) in the gray matter after additional covariates regression (denoise5)',size(mask_GM,3));
    gen_tsnr_maps(F_tsnr{14,1}, savename, figtitle, climit, tSNR_graph_GM(14)); clear savename figtitle
    savename = [QCpath 'tSNR_maps_14_COVregress_WM' '.jpg'];
    figtitle = sprintf('tSNR maps (slice 1 through %d) in the white matter after additional covariates regression (denoise5)',size(mask_GM,3));
    gen_tsnr_maps(F_tsnr{14,2}, savename, figtitle, climit, tSNR_graph_WM(14)); clear savename figtitle
end

%%% generate plots
fighndl = figure('Position',[1 1 1280 800],'Color',[0.85,0.90,0.85],'InvertHardcopy','off','Visible','off'); % red [0.95,0.9,0.85], green [0.85,0.90,0.85], blue [0.89,0.94,0.99]

labels = {[];[];[];[];'raw data';'not-cord regression';[];'motion correction';'RETROICOR';[];'co-registration';'CSF regression';'WM regression';'Other covariates regression'};
nzsteps = find(tSNR_graph_SC~=0);
tSNR_graph_GM(find((tSNR_graph_SC~=0)&(tSNR_graph_GM==0)))=NaN;
tSNR_graph_WM(find((tSNR_graph_SC~=0)&(tSNR_graph_WM==0)))=NaN;

for w=1:length(nzsteps)
    dispstring1{nzsteps(w)} = sprintf('%.2f',tSNR_graph_SC(nzsteps(w)));
end
for w=1:length(nzsteps)
    dispstring2{nzsteps(w)} = sprintf('%.2f',tSNR_graph_GM(nzsteps(w)));
    dispstring3{nzsteps(w)} = sprintf('%.2f',tSNR_graph_WM(nzsteps(w)));
end

subplot2n(2,1,1), plot(tSNR_graph_SC(nzsteps),'--p','color','b','linewidth',2.25, 'MarkerSize',18, 'MarkerEdgeColor',[0 0.37 0.22], 'MarkerFaceColor',[0.85,0.90,0.85]),
    grid on,grid minor, ylabel('tSNR'), xlim([0.5 length(nzsteps)+0.5]), ylim([0 max([tSNR_graph_SC;tSNR_graph_GM;tSNR_graph_GM])+6])
    xticks([1:length(nzsteps)]), xticklabels(labels(nzsteps)); a = get(gca,'xticklabels'); set(gca,'xticklabels',a,'fontsize',12); clear a
    text([1:length(nzsteps)]-0.1,tSNR_graph_SC(nzsteps)+4, dispstring1(nzsteps),'Color',[0 0.37 0.22],'FontSize',12)
    legend({'tSNR in GM+WM+CSF'},'FontSize',14,'TextColor','k','Location','best')
    title('BOLD temporal signal-to-noise ratio (tSNR) across pre-processing steps in the spinal cord (GM+WM+CSF)','Color',[0.65,0.11,0.19],'FontSize',16)
subplot2n(2,1,2), plot(tSNR_graph_GM(nzsteps),'--^','color',[0,0.55,0.69],'linewidth',2.25, 'MarkerSize',12, 'MarkerEdgeColor',[0,0.55,0.69], 'MarkerFaceColor',[0.89,0.94,0.99]),
    hold on, plot(tSNR_graph_WM(nzsteps),'--v','color',[0.65,0.11,0.19],'linewidth',2.25, 'MarkerSize',12, 'MarkerEdgeColor',[0.65,0.11,0.19], 'MarkerFaceColor',[0.95,0.9,0.85]),
    grid on,grid minor, ylabel('tSNR'), xlim([0.5 length(nzsteps)+0.5]), ylim([0 max([tSNR_graph_SC;tSNR_graph_GM;tSNR_graph_GM])+6])
    xticks([1:length(nzsteps)]), xticklabels(labels(nzsteps)); a = get(gca,'xticklabels'); set(gca,'xticklabels',a,'fontsize',12); clear a
    text([1:length(nzsteps)]-0.1,tSNR_graph_GM(nzsteps)+4, dispstring2(nzsteps),'Color',[0,0.55,0.69],'FontSize',12)
    text([1:length(nzsteps)]-0.1,tSNR_graph_WM(nzsteps)-4, dispstring3(nzsteps),'Color',[0.65,0.11,0.19],'FontSize',12)
    legend({'Gray matter tSNR';'White matter tSNR'},'FontSize',14,'TextColor','k','Location','best')
    title('BOLD tSNR across pre-processing steps in the gray and white matter only (co-registration step and beyond)','Color',[0.65,0.11,0.19],'FontSize',16)
saveas(fighndl,[QCpath 'tSNR_across_preproc_steps_avg' '.jpg'])
clf

for w=1:length(tSNR_graph_slicewise_rawdata)
    dispstring4{w} = sprintf('%.2f',tSNR_graph_slicewise_rawdata(w));
    dispstring5{w} = sprintf('%.2f',tSNR_graph_slicewise_final_SC(w));
    dispstring6{w} = sprintf('%.2f',tSNR_graph_slicewise_final_GM(w));
    dispstring7{w} = sprintf('%.2f',tSNR_graph_slicewise_final_WM(w));
end; clear w

subplot2n(2,1,1), plot(tSNR_graph_slicewise_rawdata,'--o','color',[0.25,0.25,0.25],'linewidth',2.25, 'MarkerSize',14, 'MarkerEdgeColor','k', 'MarkerFaceColor',[0.85,0.85,0.85]),
    hold on, plot(tSNR_graph_slicewise_final_SC,'--p','color','b','linewidth',2.25, 'MarkerSize',18, 'MarkerEdgeColor',[0 0.37 0.22], 'MarkerFaceColor',[0.85,0.90,0.85]),
    grid on,grid minor, ylabel('tSNR'), xlabel('SLICES'), xticks([1:length(tSNR_graph_slicewise_rawdata)]), 
    xlim([0.5 length(tSNR_graph_slicewise_rawdata)+0.5]), ylim([0 max([tSNR_graph_slicewise_rawdata;tSNR_graph_slicewise_final_SC;tSNR_graph_slicewise_final_GM;tSNR_graph_slicewise_final_WM])+8])
    text([1:length(tSNR_graph_slicewise_final_SC)]-0.1,tSNR_graph_slicewise_final_SC+6, dispstring5,'Color',[0 0.37 0.22],'FontSize',12)
    text([1:length(tSNR_graph_slicewise_rawdata)]-0.1,tSNR_graph_slicewise_rawdata+6, dispstring4,'Color',[0.05,0.05,0.05],'FontSize',12)
    legend({'Raw data tSNR';'Final tSNR in pre-processed GM+WM+CSF'},'FontSize',14,'TextColor','k','Location','best')
    title('BOLD temporal signal-to-noise ratio (tSNR) across SLICES before and after pre-processing in the spinal cord (GM+WM+CSF)','Color',[0.65,0.11,0.19],'FontSize',16)
subplot2n(2,1,2), plot(tSNR_graph_slicewise_final_GM,'--^','color',[0,0.55,0.69],'linewidth',2.25, 'MarkerSize',12, 'MarkerEdgeColor',[0,0.55,0.69], 'MarkerFaceColor',[0.89,0.94,0.99]),
    hold on, plot(tSNR_graph_slicewise_final_WM,'--v','color',[0.65,0.11,0.19],'linewidth',2.25, 'MarkerSize',12, 'MarkerEdgeColor',[0.65,0.11,0.19], 'MarkerFaceColor',[0.95,0.9,0.85]),
    grid on,grid minor, ylabel('tSNR'), xlabel('SLICES'), xticks([1:length(tSNR_graph_slicewise_rawdata)]), 
    xlim([0.5 length(tSNR_graph_slicewise_final_GM)+0.5]), ylim([0 max([tSNR_graph_slicewise_final_GM;tSNR_graph_slicewise_final_WM;tSNR_graph_slicewise_final_GM;tSNR_graph_slicewise_final_WM])+8])
    text([1:length(tSNR_graph_slicewise_final_GM)]-0.1,tSNR_graph_slicewise_final_GM+6, dispstring6,'Color',[0,0.55,0.69],'FontSize',12)
    text([1:length(tSNR_graph_slicewise_final_WM)]-0.1,tSNR_graph_slicewise_final_WM-6, dispstring7,'Color',[0.65,0.11,0.19],'FontSize',12)
    legend({'Final tSNR in pre-processed GRAY MATTER';'Final tSNR in pre-processed WHITE MATTER'},'FontSize',14,'TextColor','k','Location','best')
    title('BOLD tSNR across SLICES after pre-processing in the GRAY and WHITE matter','Color',[0.65,0.11,0.19],'FontSize',16)
saveas(fighndl,[QCpath 'tSNR_across_preproc_steps_slicewise' '.jpg'])

close(fighndl)
clear readme_tSNR_graph tSNR_graph_SC tSNR_graph_GM tSNR_graph_WM tSNR_graph_slicewise_rawdata tSNR_graph_slicewise_final_SC tSNR_graph_slicewise_final_GM tSNR_graph_slicewise_final_WM F_tsnr fighndl dispstring1 dispstring2 dispstring3 dispstring4 dispstring5 dispstring6 dispstring7 labels nzsteps 
catch
end
end

%% Save disk storage info and figures
if (k2==RUNS(k))
clear QCpath fighndl
if ~(exist([sub_dir{k} 'QC' '/19_disk_storage_info'],'dir'))
	mkdir([sub_dir{k} 'QC' '/19_disk_storage_info'])
end
QCpath = [sub_dir{k} 'QC' '/19_disk_storage_info/'];

try
    sdird = dir([sub_dir{k} 'disk_storage_space_info_*.mat']); sdird = sdird(end);
    load(sdird(1).name)
    clear sdird
catch
    sdirall = dir([sub_dir{k} '**/*.*']); sdirall = sdirall(3:end);
    sdirall_name = {sdirall.name}';
    excludefile = zeros(length(sdirall_name),1);
    for m=1:length(sdirall_name)
        if length(sdirall_name{m})>=4
            if strcmp(sdirall_name{m}(end-3:end),'.nii')==1
                for m2=1:length(sdirall_name)
                    if length(sdirall_name{m2})>=7
                        if (strcmp(sdirall_name{m}(1:end-4),sdirall_name{m2}(1:end-7))==1) && (strcmp(sdirall_name{m2}(end-6:end),'.nii.gz')==1) && (m~=m2)
                            excludefile(m) = 1; % if a .nii file is found AND if a corresponding .nii.gz file also exists, then do not count the .nii file for total storage computation
                        end
                    end
                end
            end
        end
    end
    sdirall_bytes = {sdirall.bytes}';
    sdirall_bytes = cell2mat(sdirall_bytes);
    sdirall_bytes = sdirall_bytes(find(excludefile==0));
    sdirall_bytes = sum(sdirall_bytes)/10^9; % size of subject folder in gigabytes
    folder_size_3_sub = sdirall_bytes;
    folder_size_2_sub = NaN;
    folder_size_1_sub = NaN;
    tmpc=clock; clk = sprintf('%.4d-%.2d-%.2d_%.2d-%.2d',tmpc(1),tmpc(2),tmpc(3),tmpc(4),tmpc(5)); clear tmpc
    dskfilename = sprintf('disk_storage_space_info_%s.mat',clk); clear clk
    save([sub_dir{k} dskfilename],'folder_size_*')
    clear sdirall sdirall_bytes sdirall_name m m2 excludefile dskfilename
end

fighndl = figure('Position',[1 1 1200 800],'Color',[0.85,0.90,0.85],'InvertHardcopy','off','Visible','off'); % [0.85,0.90,0.85] [0.8, 0.89, 0.94]
b = bar(1,folder_size_1_sub,0.5,'FaceColor',[0,0.55,0.69],'EdgeColor',[0.65,0.11,0.19],'LineWidth',1.5); hold on,
	xtips1 = b(1).XEndPoints; ytips1 = b(1).YEndPoints; labels1 = string(round(b(1).YData,2)); clear b
    labels1 = strcat(labels1,' GB');
    text(xtips1,ytips1,labels1,'HorizontalAlignment','center','VerticalAlignment','bottom','fontsize',17','color',[0.65,0.11,0.19]);
b = bar(2,folder_size_2_sub,0.5,'FaceColor',[0,0.65,0.69],'EdgeColor',[0.65,0.11,0.19],'LineWidth',1.5);
	xtips2 = b(1).XEndPoints; ytips2 = b(1).YEndPoints; labels2 = string(round(b(1).YData,2)); clear b
    labels2 = strcat(labels2,' GB');
    if folder_size_2_sub==0, labels2 = 'not performed'; end
    text(xtips2,ytips2,labels2,'HorizontalAlignment','center','VerticalAlignment','bottom','fontsize',17','color',[0.65,0.11,0.19]);
b = bar(3,folder_size_3_sub,0.5,'FaceColor',[0,0.75,0.69],'EdgeColor',[0.65,0.11,0.19],'LineWidth',1.5);
	xtips3 = b(1).XEndPoints; ytips3 = b(1).YEndPoints; labels3 = string(round(b(1).YData,2)); clear b
    labels3 = strcat(labels3,' GB');
    if folder_size_3_sub==0, labels3 = 'not performed'; end
    text(xtips3,ytips3,labels3,'HorizontalAlignment','center','VerticalAlignment','bottom','fontsize',17','color',[0.65,0.11,0.19]);
    grid on, grid minor, try ylim([0 max(folder_size_1_sub)+0.1.*max(folder_size_1_sub)]), catch, end, ylabel('Disk storage (in gigabytes)'),
    xticks([1,2,3]), xticklabels({'At the end of pre-processing','After removal of slice-wise files','After NIfTI file compression (gzip)'})
    a = get(gca,'xticklabels'); set(gca,'xticklabels',a,'fontsize',14); clear a
    legstr{1} = sprintf('At the end of pre-processing'); legstr{2} = sprintf('... AND after removal of duplicate slice-wise files'); legstr{3} = sprintf('... AND after NIfTI file compression (gzip)');
    legend(legstr,'Location','northeast','FontSize',18)
    title(sprintf('DISK STORAGE: size of subject folder at the end of pre-processing\n(%s)',sub_dir{k}),'Interpreter', 'none', 'Color',[0 0.37 0.22],'FontSize',18);
saveas(fighndl,[QCpath 'disk_storage_bargraph' '.jpg'])
close(fighndl)
clear folder_size_1_sub folder_size_2_sub folder_size_3_sub readme_disk_storage QCpath fighndl legstr b xtips1 ytips1 labels1 xtips2 ytips2 labels2 xtips3 ytips3 labels3 k dskfilename

end

%%
clear base_dir_sub fname fname3 fname_anat_orig fname_anat run
    end
end

%%
%% Save time taken for each step
if exist('time_taken_preproc','var')

ttaken_total = zeros(19,1); ttaken_step = zeros(19,1); ttaken_sub = zeros(length(sub_dir),1);
if choices_preproc(2)==1
    temp = time_taken_preproc.step02_slicetimingcorrection(find(time_taken_preproc.step02_slicetimingcorrection~=0));
    ttaken_total(2) = sum(temp); ttaken_step(2) = mean(temp); clear temp
    temp = sum(time_taken_preproc.step02_slicetimingcorrection,2); ttaken_sub = sum([ttaken_sub,temp],2); clear temp
end
if choices_preproc(3)==1
    temp = time_taken_preproc.step03_matchAnatFuncSlices(find(time_taken_preproc.step03_matchAnatFuncSlices~=0));
    ttaken_total(3) = sum(temp); ttaken_step(3) = mean(temp); clear temp
    temp = sum(time_taken_preproc.step03_matchAnatFuncSlices,2); ttaken_sub = sum([ttaken_sub,temp],2); clear temp
end
if choices_preproc(4)==1
    temp = time_taken_preproc.step04_splitData(find(time_taken_preproc.step04_splitData~=0));
    ttaken_total(4) = sum(temp); ttaken_step(4) = mean(temp); clear temp
    temp = sum(time_taken_preproc.step04_splitData,2); ttaken_sub = sum([ttaken_sub,temp],2); clear temp
end
if choices_preproc(5)==1
    temp = time_taken_preproc.step05_notCordMask(find(time_taken_preproc.step05_notCordMask~=0));
    ttaken_total(5) = sum(temp); ttaken_step(5) = mean(temp); clear temp
    temp = sum(time_taken_preproc.step05_notCordMask,2); ttaken_sub = sum([ttaken_sub,temp],2); clear temp
end
if choices_preproc(6)==1
    temp = time_taken_preproc.step06_denoise1(find(time_taken_preproc.step06_denoise1~=0));
    ttaken_total(6) = sum(temp); ttaken_step(6) = mean(temp); clear temp
    temp = sum(time_taken_preproc.step06_denoise1,2); ttaken_sub = sum([ttaken_sub,temp],2); clear temp
end
if choices_preproc(7)==1
    temp = time_taken_preproc.step07_gaussianMask(find(time_taken_preproc.step07_gaussianMask~=0));
    ttaken_total(7) = sum(temp); ttaken_step(7) = mean(temp); clear temp
    temp = sum(time_taken_preproc.step07_gaussianMask,2); ttaken_sub = sum([ttaken_sub,temp],2); clear temp
end
if choices_preproc(8)==1
    temp = time_taken_preproc.step08_motionCorrection(find(time_taken_preproc.step08_motionCorrection~=0));
    ttaken_total(8) = sum(temp); ttaken_step(8) = mean(temp); clear temp
    temp = sum(time_taken_preproc.step08_motionCorrection,2); ttaken_sub = sum([ttaken_sub,temp],2); clear temp
end
if choices_preproc(9)==1
    temp = time_taken_preproc.step09_retroicor(find(time_taken_preproc.step09_retroicor~=0));
    ttaken_total(9) = sum(temp); ttaken_step(9) = mean(temp); clear temp
    temp = sum(time_taken_preproc.step09_retroicor,2); ttaken_sub = sum([ttaken_sub,temp],2); clear temp
end
if choices_preproc(10)==1
    temp = time_taken_preproc.step10_GMWMCSFmasks(find(time_taken_preproc.step10_GMWMCSFmasks~=0));
    ttaken_total(10) = sum(temp); ttaken_step(10) = mean(temp); clear temp
    temp = sum(time_taken_preproc.step10_GMWMCSFmasks,2); ttaken_sub = sum([ttaken_sub,temp],2); clear temp
end
if choices_preproc(11)==1
    temp = time_taken_preproc.step11_FuncAnatRegistration(find(time_taken_preproc.step11_FuncAnatRegistration~=0));
    ttaken_total(11) = sum(temp); ttaken_step(11) = mean(temp); clear temp
    temp = sum(time_taken_preproc.step11_FuncAnatRegistration,2); ttaken_sub = sum([ttaken_sub,temp],2); clear temp
end
if choices_preproc(12)==1
    temp = time_taken_preproc.step12_CSFregress(find(time_taken_preproc.step12_CSFregress~=0));
    ttaken_total(12) = sum(temp); ttaken_step(12) = mean(temp); clear temp
    temp = sum(time_taken_preproc.step12_CSFregress,2); ttaken_sub = sum([ttaken_sub,temp],2); clear temp
end
if choices_preproc(13)==1
    temp = time_taken_preproc.step13_WMregress(find(time_taken_preproc.step13_WMregress~=0));
    ttaken_total(13) = sum(temp); ttaken_step(13) = mean(temp); clear temp
    temp = sum(time_taken_preproc.step13_WMregress,2); ttaken_sub = sum([ttaken_sub,temp],2); clear temp
end
if choices_preproc(14)==1
    temp = time_taken_preproc.step14_COVregress(find(time_taken_preproc.step14_COVregress~=0));
    ttaken_total(14) = sum(temp); ttaken_step(14) = mean(temp); clear temp
    temp = sum(time_taken_preproc.step14_COVregress,2); ttaken_sub = sum([ttaken_sub,temp],2); clear temp
end
if choices_preproc(15)==1
    temp = time_taken_preproc.step15_filterData(find(time_taken_preproc.step15_filterData~=0));
    ttaken_total(15) = sum(temp); ttaken_step(15) = mean(temp); clear temp
    temp = sum(time_taken_preproc.step15_filterData,2); ttaken_sub = sum([ttaken_sub,temp],2); clear temp
end
if choices_preproc(16)==1
    temp = time_taken_preproc.step16_cord_quadrants(find(time_taken_preproc.step16_cord_quadrants~=0));
    ttaken_total(16) = sum(temp); ttaken_step(16) = mean(temp); clear temp
    temp = sum(time_taken_preproc.step16_cord_quadrants,2); ttaken_sub = sum([ttaken_sub,temp],2); clear temp
end
if choices_preproc(17)==1
    temp = time_taken_preproc.step17_deconvolution(find(time_taken_preproc.step17_deconvolution~=0));
    ttaken_total(17) = sum(temp); ttaken_step(17) = mean(temp); clear temp
    temp = sum(time_taken_preproc.step17_deconvolution,2); ttaken_sub = sum([ttaken_sub,temp],2); clear temp
end
if choices_preproc(18)==1
    temp = time_taken_preproc.step18_standardspaceRegistration(find(time_taken_preproc.step18_standardspaceRegistration~=0));
    ttaken_total(18) = sum(temp); ttaken_step(18) = mean(temp); clear temp
    temp = sum(time_taken_preproc.step18_standardspaceRegistration,2); ttaken_sub = sum([ttaken_sub,temp],2); clear temp
end
try
    temp = time_taken_preproc.tSNR_computation(find(time_taken_preproc.tSNR_computation~=0));
    ttaken_total(19) = sum([ttaken_total(19),sum(temp)]); ttaken_step(19) = sum([ttaken_step(19),mean(temp)]); clear temp
    temp = sum(time_taken_preproc.tSNR_computation,2); ttaken_sub = sum([ttaken_sub,temp],2); clear temp
catch
end
try
    temp = time_taken_preproc.step19_remove_slicewise_files(find(time_taken_preproc.step19_remove_slicewise_files~=0));
    ttaken_total(19) = sum([ttaken_total(19),sum(temp)]); ttaken_step(19) = sum([ttaken_step(19),mean(temp)]); clear temp
    temp = sum(time_taken_preproc.step19_remove_slicewise_files,2); ttaken_sub = sum([ttaken_sub,temp],2); clear temp
catch
end
if choices_preproc(19)==1
    temp = time_taken_preproc.step19_gzip_delete_nii(find(time_taken_preproc.step19_gzip_delete_nii~=0));
    ttaken_total(19) = sum([ttaken_total(19),sum(temp)]); ttaken_step(19) = sum([ttaken_step(19),mean(temp)]); clear temp
    temp = sum(time_taken_preproc.step19_gzip_delete_nii,2); ttaken_sub = sum([ttaken_sub,temp],2); clear temp
end
ttaken_total = ttaken_total(find(choices_preproc==1));
ttaken_step = ttaken_step(find(choices_preproc==1));

fighndl = figure('Position',[1 1 1200 800],'Color',[0.85,0.90,0.85],'InvertHardcopy','off','Visible','off'); % [0.85,0.90,0.85] [0.8, 0.89, 0.94]
subplot2n(3,1,1), b = bar(ttaken_total,0.5,'FaceColor',[0,0.55,0.69],'EdgeColor',[0.65,0.11,0.19],'LineWidth',1.5);
    xticks([1:length(find(choices_preproc==1))]), xticklabels(mat2cell(find(choices_preproc==1),length(find(choices_preproc==1))))
	xtips1 = b(1).XEndPoints; ytips1 = b(1).YEndPoints;
    for w=1:length(ttaken_total)
        if round(ttaken_total(w)/3600)==0
            if round(mod(ttaken_total(w),3600)/60)==0
                if round(mod(ttaken_total(w),60))==0
                    labels1{w,1} = sprintf('%.0fms',round(mod(ttaken_total(w),1)*1000));
                else
                    labels1{w,1} = sprintf('%.0fs',round(mod(ttaken_total(w),60)));
                end
            else
                labels1{w,1} = sprintf('%.0fm:%.0fs',round(mod(ttaken_total(w),3600)/60),round(mod(ttaken_total(w),60)));
            end
        else
            labels1{w,1} = sprintf('%.0fh:%.0fm:%.0fs',round(ttaken_total(w)/3600),round(mod(ttaken_total(w),3600)/60),round(mod(ttaken_total(w),60)));
        end
    end
    labels1 = string(labels1); clear b
    text(xtips1,ytips1,labels1,'HorizontalAlignment','center','VerticalAlignment','bottom','fontsize',12','color',[0.65,0.11,0.19]);
    xlabel('Pre-processing steps'), ylabel('time taken (s)'), ylim([0 1.2*max(ttaken_total)])
    title(sprintf('(Total execution time, including all steps and subjects = %.0fh:%.0fm:%.0fs)\n\nTOTAL TIME TAKEN for each pre-processing step (%d subjects)',round(sum(ttaken_total)/3600),round(mod(sum(ttaken_total),3600)/60),round(mod(sum(ttaken_total),60)),length(sub_dir)),'Interpreter', 'none', 'Color',[0 0.37 0.22],'FontSize',16);
    clear labels1 xtips1 ytips1
subplot2n(3,1,2), b = bar(ttaken_step,0.5,'FaceColor',[0,0.65,0.69],'EdgeColor',[0.65,0.11,0.19],'LineWidth',1.5);
    xticks([1:length(ttaken_step)]), xticklabels(mat2cell(find(choices_preproc==1),length(find(choices_preproc==1))))
	xtips1 = b(1).XEndPoints; ytips1 = b(1).YEndPoints;
    for w=1:length(ttaken_step)
        if round(ttaken_step(w)/3600)==0
            if round(mod(ttaken_step(w),3600)/60)==0
                if round(mod(ttaken_step(w),60))==0
                    labels1{w,1} = sprintf('%.0fms',round(mod(ttaken_step(w),1)*1000));
                else
                    labels1{w,1} = sprintf('%.0fs',round(mod(ttaken_step(w),60)));
                end
            else
                labels1{w,1} = sprintf('%.0fm:%.0fs',round(mod(ttaken_step(w),3600)/60),round(mod(ttaken_step(w),60)));
            end
        else
            labels1{w,1} = sprintf('%.0fh:%.0fm:%.0fs',round(ttaken_step(w)/3600),round(mod(ttaken_step(w),3600)/60),round(mod(ttaken_step(w),60)));
        end
    end
    labels1 = string(labels1); clear b
    text(xtips1,ytips1,labels1,'HorizontalAlignment','center','VerticalAlignment','bottom','fontsize',12','color',[0.65,0.11,0.19]);
    xlabel('Pre-processing steps'), ylabel('time taken (s)'), ylim([0 1.2*max(ttaken_step)])
    title(sprintf('AVERAGE TIME TAKEN for each pre-processing step per subject'),'Interpreter', 'none', 'Color',[0 0.37 0.22],'FontSize',16);
    clear labels1 xtips1 ytips1
subplot2n(3,1,3), b = bar(ttaken_sub,0.5,'FaceColor',[0,0.75,0.69],'EdgeColor',[0.65,0.11,0.19],'LineWidth',1.5);
	xtips1 = b(1).XEndPoints; ytips1 = b(1).YEndPoints;
    for w=1:length(ttaken_sub)
        if round(ttaken_sub(w)/3600)==0
            if round(mod(ttaken_sub(w),3600)/60)==0
                if round(mod(ttaken_sub(w),60))==0
                    labels1{w,1} = sprintf('%.0fms',round(mod(ttaken_sub(w),1)*1000));
                else
                    labels1{w,1} = sprintf('%.0fs',round(mod(ttaken_sub(w),60)));
                end
            else
                labels1{w,1} = sprintf('%.0fm:%.0fs',round(mod(ttaken_sub(w),3600)/60),round(mod(ttaken_sub(w),60)));
            end
        else
            labels1{w,1} = sprintf('%.0fh:%.0fm:%.0fs',round(ttaken_sub(w)/3600),round(mod(ttaken_sub(w),3600)/60),round(mod(ttaken_sub(w),60)));
        end
    end
    labels1 = string(labels1); clear b
    text(xtips1,ytips1,labels1,'HorizontalAlignment','center','VerticalAlignment','bottom','fontsize',12','color',[0.65,0.11,0.19]);
    xlabel('Subjects'), ylabel('time taken (s)'), ylim([0 1.2*max(ttaken_sub)])
    title(sprintf('TOTAl TIME TAKEN for each SUBJECT'),'Interpreter', 'none', 'Color',[0 0.37 0.22],'FontSize',16);
```
