## Supplementary material for "Neptune: a toolbox for spinal cord functional MRI data processing and quality assurance": Neptune software: get_covariate_files.m

```
function cov_files = get_covariate_files(sub_dir,func_files,runs)

quest = sprintf('User-defined covariates for denoising - two options to choose files:\n\n1. Automatic: I have already named my covariates as\n<fMRI_data_filename>_covariates.txt\n(or file extensions .csv, .xls, .xlsx, .dat, .mat).\n\nOR\n\n2. Manual: I will choose the files in each subject separately.\n');
answer1 = questdlg(quest,'User-defined covariates','Automatic','Manual','Manual'); clear quest

switch answer1
    case 'Automatic'
        for k=1:size(func_files,1)
            for k2=1:size(func_files,2)
                sdirc = cat(1,dir([sub_dir{k} func_files{k,k2} '_covariates' '.txt']),dir([sub_dir{k} func_files{k,k2} '_covariates' '.csv']),dir([sub_dir{k} func_files{k,k2} '_covariates' '.xls']),dir([sub_dir{k} func_files{k,k2} '_covariates' '.xlsx']),dir([sub_dir{k} func_files{k,k2} '_covariates' '.dat']),dir([sub_dir{k} func_files{k,k2} '_covariates' '.mat']));
                if ~isempty('sdirc')
                    cov_files{k,k2} = [sdirc(1).folder '/' sdirc(1).name];
                else
                    cov_files{k,k2} = [];
                end
                numcovfiles(k,k2) = length(sdirc); clear sdirc
            end
        end
        
        [z1,z2] = find(numcovfiles==0); z12 = [z1,z2]; clear z1 z2
        if isempty('z12')
            dispstrng = sprintf('The following subject(s) do not have covariates.\nDo you want to manually select covariate files for these subjects, OR do not perform covariate regression for these subjects only?\n\n');
            for zz=1:size(z12,1)
                dispstrng = [dispstrng sprintf('Subject %d, run %d (%s)\n',z12(zz,1),z12(zz,2),[sub_dir{z12(zz,1)} func_files{z12(zz,1),z12(zz,2)}])]; %#ok<*AGROW>
            end; clear zz
            answer2 = questdlg(dispstrng,'User-defined covariates','Manually select covariate files','Do not perform covariate regression on these subjects','Manually select covariate files'); clear dispstrng
            switch answer2
                case 'Manually select covariate files'
                    for k=1:length(sub_dir)
                        cd(sub_dir{k})
                        sdir2 = dir(['**/' cov_string]); sdir2_folder = {sdir2.folder}; sdir2_name = {sdir2.name}; sdir2={sdir2.name};
                        for k2=1:length(sdir2_folder)
                            sdir2_name{k2} = [erase(sdir2_folder{k2},sub_dir{k}(1:end-1)) '/' sdir2{k2}];
                        end
                        for k2=1:runs(k)
                            if ~isempty(find(sum(abs(z12-[k,k2]),2)==0)) %#ok<EFIND>
                                prompt = sprintf('Select user-defined covariate file for SUBJECT %d, RUN %d  (.txt, .csv, .xls, .xlsx, .dat or .mat only)\n(%s)\n',k,k2,[sub_dir{k} func_files{k,k2}]);
                                indx = listdlg('PromptString',{prompt,''},...
                                    'SelectionMode','single','ListSize',[700 500],'ListString',sdir2_name);
                                cov_files{k,k2} = [sdir2_folder{indx} '/' sdir2{indx}];
                                clear prompt indx
                            end
                        end
                        clear sdir2* k2
                        cd ..
                    end; clear k
            end
        end; clear z12 answer2
        
        [y1,y2] = find(numcovfiles>1); y12 = [y1,y2]; clear y1 y2
        if isempty('y12')
            dispstrng = sprintf('The following subject(s) have MORE THAN ONE covariates file.\nDo you want to manually select a single covariate file for these subjects, OR perform covariate regression using the first identified file only?\n\n');
            for yy=1:size(y12,1)
                dispstrng = [dispstrng sprintf('Subject %d, run %d (%s)\n',y12(yy,1),y12(yy,2),[sub_dir{y12(yy,1)} func_files{y12(yy,1),y12(yy,2)}])];
            end; clear yy
            answer3 = questdlg(dispstrng,'User-defined covariates','Manually select covariate file','Perform covariate regression using the first identified file','Manually select covariate file'); clear dispstrng
            switch answer3
                case 'Manually select covariate file'
                    for k=1:length(sub_dir)
                        cd(sub_dir{k})
                        sdir2 = dir(['**/' cov_string]); sdir2_folder = {sdir2.folder}; sdir2_name = {sdir2.name}; sdir2={sdir2.name};
                        for k2=1:length(sdir2_folder)
                            sdir2_name{k2} = [erase(sdir2_folder{k2},sub_dir{k}(1:end-1)) '/' sdir2{k2}];
                        end
                        for k2=1:runs(k)
                            if ~isempty(find(sum(abs(y12-[k,k2]),2)==0)) %#ok<EFIND>
                                prompt = sprintf('Select user-defined covariate file for SUBJECT %d, RUN %d  (.txt, .csv, .xls, .xlsx, .dat or .mat only)\n(%s)\n',k,k2,[sub_dir{k} func_files{k,k2}]);
                                indx = listdlg('PromptString',{prompt,''},...
                                    'SelectionMode','single','ListSize',[700 500],'ListString',sdir2_name);
                                cov_files{k,k2} = [sdir2_folder{indx} '/' sdir2{indx}];
                                clear prompt indx
                            end
                        end
                        clear sdir2* k2
                        cd ..
                    end; clear k
            end
        end; clear y12 answer3

        
    case 'Manual'
        for k=1:length(sub_dir)
            cd(sub_dir{k})
            sdir2 = cat(1,dir('**/*.txt'),dir('**/*.csv'),dir('**/*.xls'),dir('**/*.xlsx'),dir('**/*.dat'),dir('**/*.mat'));
            sdir2_folder = {sdir2.folder}; sdir2_name = {sdir2.name}; sdir2={sdir2.name};
            for k2=1:length(sdir2_folder)
                sdir2_name{k2} = [erase(sdir2_folder{k2},sub_dir{k}(1:end-1)) '/' sdir2{k2}];
            end
            for k2=1:runs(k)
                prompt = sprintf('Select user-defined covariate file for SUBJECT %d, RUN %d  (.txt, .csv, .xls, .xlsx, .dat or .mat only)\n(%s)\n',k,k2,[sub_dir{k} func_files{k,k2}]);
                indx = listdlg('PromptString',{prompt,''},...
                    'SelectionMode','single','ListSize',[700 500],'ListString',sdir2_name);
                cov_files{k,k2} = [sdir2_folder{indx} '/' sdir2{indx}];
            end
            clear sdir2* prompt k2 indx
            cd ..
        end; clear k
        
end


end
```
