## Supplementary material for "Neptune: a toolbox for spinal cord functional MRI data processing and quality assurance": Neptune software: initdialog.m

```
function [selection1,selection2,selection3,selection4,selection5,selection6] = initdialog(bg1,bg2,bg3,bg4,bg5,bg6,flag,selection1,selection2,selection3,selection4,selection5,selection6)

rb1 = uiradiobutton(bg1,'Position',[10 15 540 15],'Value',strcmp(selection1,'Siemens scanner'));
rb2 = uiradiobutton(bg1,'Position',[180 15 540 15],'Value',strcmp(selection1,'Philips scanner'));
rb3 = uiradiobutton(bg1,'Position',[350 15 540 15],'Value',strcmp(selection1,'Other'));

rb4 = uiradiobutton(bg2,'Position',[10 15 540 15],'Value',strcmp(selection2,'7T scanner'));
rb5 = uiradiobutton(bg2,'Position',[180 15 540 15],'Value',strcmp(selection2,'3T scanner'));
rb6 = uiradiobutton(bg2,'Position',[350 15 540 15],'Value',strcmp(selection2,'Other'));

rb7 = uiradiobutton(bg3,'Position',[10 15 540 15],'Value',strcmp(selection3,'Cervical spine'));
rb8 = uiradiobutton(bg3,'Position',[180 15 540 15],'Value',strcmp(selection3,'Thoraco-lumbar spine'));

rb9 = uiradiobutton(bg4,'Position',[10 15 540 15],'Value',strcmp(selection4,'Resting-state fMRI'));
rb10 = uiradiobutton(bg4,'Position',[180 15 540 15],'Value',strcmp(selection4,'Task fMRI'));

rb11 = uiradiobutton(bg5,'Position',[10 15 540 15],'Value',strcmp(selection5,'Pre-processing only'));
rb12 = uiradiobutton(bg5,'Position',[180 15 540 15],'Value',strcmp(selection5,'Post-processing only'));
rb13 = uiradiobutton(bg5,'Position',[350 15 540 15],'Value',strcmp(selection5,'Both'));

rb14 = uiradiobutton(bg6,'Position',[450 15 540 15]);
rb15 = uiradiobutton(bg6,'Position',[225 15 540 15],'Value',strcmp(selection6,'DONE'));
rb16 = uiradiobutton(bg6,'Position',[450 15 540 15],'Value',strcmp(selection6,'Help'));

rb1.Text = 'Siemens scanner';
rb2.Text = 'Philips scanner';
rb3.Text = 'Other';
rb4.Text = '7T scanner';
rb5.Text = '3T scanner';
rb6.Text = 'Other';
rb7.Text = 'Cervical spine';
rb8.Text = 'Thoraco-lumbar spine';
rb9.Text = 'Resting-state fMRI';
rb10.Text = 'Task fMRI';
rb11.Text = 'Pre-processing only';
rb12.Text = 'Post-processing only';
rb13.Text = 'Both';
rb14.Text = '';
rb15.Text = 'DONE';
rb16.Text = 'Help';

pause(2)

selection1 = bg1.SelectedObject.Parent.SelectedObject.Text;
selection2 = bg2.SelectedObject.Parent.SelectedObject.Text;
selection3 = bg3.SelectedObject.Parent.SelectedObject.Text;
selection4 = bg4.SelectedObject.Parent.SelectedObject.Text;
selection5 = bg5.SelectedObject.Parent.SelectedObject.Text;
selection6 = bg6.SelectedObject.Parent.SelectedObject.Text;

if flag==1 && strcmp(selection6,'Help')==1
    selection6='';
end

end
```
