## Supplementary material for "Neptune: a toolbox for spinal cord functional MRI data processing and quality assurance": Neptune software: main_scfMRItb.m

```
% Neptune toolbox, created by Ranga Deshpande and Robert L Barry
% at the Athinoula A. Martinos Center for Biomedical Imaging, Massachusetts General Hospital, Harvard Medical School, Boston, MA, USA
% neptune_v1.YYMMDD was updated on DD-MM-20YY by Ranga Deshpande.

try
    purge
catch
    clear; close all; clc
end
first_dir = [pwd '/'];
tmp = which('neptune'); neptune_dir = tmp(1:end-9); clear tmp
tc00=tic;

eval('cd ~');
afni_directory = [pwd '/abin'];
setenv('PATH', [getenv('PATH') ':' afni_directory]); % path to AFNI's abin folder in your computer
cd(first_dir)
add_AFNI_to_path

try
    eval('cd ~');
    sdirSCT = dir('sct_5.4*');% use SCT 5.4 by default because Neptune was tested on it.
    if isempty(sdirSCT)
        sdirSCT = dir('sct_*');
        if ~isempty(sdirSCT)
            sdirSCT = sdirSCT(end); % if 5.4 is unavailable then use the latest available version.
        end
    else
        sdirSCT = sdirSCT(end);
    end
    if ~isempty(sdirSCT)
        SCT_directory = [sdirSCT(1).folder '/' sdirSCT(1).name '/bin'];
        setenv('PATH', [getenv('PATH') ':' SCT_directory]); % path to SCT's folder in your computer
        exist_SCT = 1;
        fprintf('Successfully added SCT to path\nLocation: %s\n',SCT_directory)
    else
        fprintf('Could not add SCT to path\n')
        exist_SCT = 0;
    end
    cd(first_dir)
    clear sdirSCT
catch
    fprintf('Could not add SCT to path\n')
    exist_SCT = 0;
end
warning('off')

month={'Jan','Feb','Mar','Apr','May','Jun','Jul','Aug','Sep','Oct','Nov','Dec'}';
L=0;
L=L+1; logg{L,1}=sprintf('------- Spinal cord fMRI toolbox (NEPTUNE) execution log -------');
clk=clock; L=L+2; logg{L,1}=sprintf('%d/%s/%d %dh:%dm:%ds - begin',clk(3),month{clk(2)},clk(1),clk(4),clk(5),round(clk(6))); clear clk

S = txt_update;  % Create the textbox.
set(S.tx,'string',logg)

tmp_init = clock; clk_init = sprintf('%.4d-%.2d-%.2d_%.2d-%.2d',tmp_init(1),tmp_init(2),tmp_init(3),tmp_init(4),tmp_init(5));

%% --- begin ---
welcome_fig = figure('Name','Welcome','NumberTitle','off','units','normalized','outerposition',[0 3/4 1/3 1/4]); % [position_along_x_from_bottomleft position_along_y_from_bottomleft size_x size_y] in normalized units (0,1)
subplot = @(m,n,p) subtightplot(m, n, p, [0.001 0.005], [0.01 0.001], [0.01 0.001]);
figure(welcome_fig), subplot(1,1,1), imshow('status_images/barrygp.jpg','InitialMagnification','fit')
neptune_fig = figure('Name','Neptune v1.0','NumberTitle','off','units','normalized','outerposition',[1/16 23.5/48 1/4 5/23]); % [position_along_x_from_bottomleft position_along_y_from_bottomleft size_x size_y] in normalized units (0,1)
subplot = @(m,n,p) subtightplot(m, n, p, [0.001 0.005], [0.01 0.001], [0.01 0.001]);
figure(neptune_fig), subplot(1,1,1), imshow('status_images/neptune_v1.0.jpg','InitialMagnification','fit')
msgicon = imread('status_images/neptune_v1.0.jpg'); msgicon=msgicon(1:10:end,1:10:end,:);
clear subplot

quest = ['\color[rgb]{0.65,0.11,0.19}' sprintf('Welcome to NEPTUNE v1.0!\n(The solar system nurtures our central nervous system (CNS).\nNeptune: last of the planets. Spinal cord: last of the CNS)\n\nThis spinal cord fMRI toolbox is offered by the \nBrain and Spinal Cord Laboratory\n(Creators: Ranga Deshpande and Robert Barry)\nAthinoula A. Martinos Center for Biomedical Imaging,\nMassachusetts General Hospital, Harvard Medical School and Harvard-MIT HST\n\nAre you ready to process spinal cord fMRI data?')];
opts.Interpreter = 'tex'; opts.Default = 'Yes! I am excited :)';
answer1 = questdlg(quest,'Welcome','Yes! I am excited :)','Not now','I am confused',opts); clear quest opts

switch answer1
    case 'Not now'
        return
    case 'I am confused'
        questdlg('To help you decide, I will redirect you to the toolbox documentation','Help / documentation','Awesome!','Awesome!');
        open('neptune_manual.pdf')
end; clear answer1


quest = sprintf('Load settings file?  (you can save input settings from a previous execution [.mat] and load in this step)');
answer2 = questdlg(quest,'Settings file','Yes, select .mat file','No, show all the settings','No, show all the settings'); clear quest

switch answer2
    case 'Yes, select .mat file'
        [f2,f1] = uigetfile(first_dir,'Choose settings file (.mat)...');
        settings_file = [f1 f2]; clear f1 f2
        load(settings_file)
        reselect_paths
end; clear answer2

wbar1 = waitbar(0,'Initializing...','Name','Initialization...','Units','normalized','Position',[4/5 0 1/5 1/17]);

%% --- Check unix and AFNI ---
try waitbar(0/8,wbar1,'Checking unix environment...'); catch, end
if ~exist('afni_test','var')
    afni_test=1; % if you do not want it to test for AFNI every time, create a variable called afni_test, set it to 0, and save it in the settings file to load next time.
end
if (afni_test==1)&&(~exist('notunix','var'))&&(~exist('noAFNI','var'))
notunix = unix('ls >output13.txt'); delete('output13.txt');
if notunix~=0
    msgbox(sprintf('The toolbox cannot be used without Unix support needed to run AFNI commands.\nWindows users could try setting up Linux virtual environment and installing AFNI on it (Google "AFNI on windows").\n\nTerminating execution...'),'Abort','error');
    try close(wbar1), catch, end
    return
end

try waitbar(0/8,wbar1,'Checking availability of AFNI...'); catch, end
% noAFNI = unix('afni_system_check.py -check_all >AFNI_error.txt'); if noAFNI==0, delete('AFNI_error.txt'); end
[noAFNI, ~] = unix('3dDespike');
if noAFNI~=0
    tmp=clock; clk = sprintf('%.4d-%.2d-%.2d_%.2d-%.2d',tmp(1),tmp(2),tmp(3),tmp(4),tmp(5));
    movefile('AFNI_error.txt',sprintf('AFNI_error_%s.txt',clk))
    msgbox(sprintf('AFNI is either not installed or is not in Matlab path (needed); cannot continue execution...\n- If you have not yet installed AFNI on your system, now is a good time.\n- If you have AFNI yet got this message, refer to line-6 (AFNI path) in the script main_scfMRItb.m\n\nAFNI error message has been saved to\n%s\n\nTerminating execution...',sprintf('AFNI_error_%s.txt',clk)),'Abort','error'); clear clk tmp
    try close(wbar1), catch, end
    return
end
end

%% --- initial choices ---
try waitbar(1/8,wbar1,'Initial choices...'); catch, end

if ~exist('rest','var') && ~exist('scanner','var') && ~exist('scan7T','var') && ~exist('C_spine','var')

fig1 = uifigure('Position',[680 680 600 380],'Color',[0.95,0.875,0.8]); drawnow; fig1.Visible='off'; fig1.Visible='on';
bg1 = uibuttongroup(fig1,'Position',[50 300 500 50],'BackgroundColor',[0.95,0.9,0.85],'BorderType','line');
bg2 = uibuttongroup(fig1,'Position',[50 250 500 50],'BackgroundColor',[0.95,0.9,0.85],'BorderType','line');
bg3 = uibuttongroup(fig1,'Position',[50 200 500 50],'BackgroundColor',[0.95,0.9,0.85],'BorderType','line');
bg4 = uibuttongroup(fig1,'Position',[50 150 500 50],'BackgroundColor',[0.95,0.9,0.85],'BorderType','line');
bg5 = uibuttongroup(fig1,'Position',[50 100 500 50],'BackgroundColor',[0.95,0.9,0.85],'BorderType','line');
bg6 = uibuttongroup(fig1,'Position',[50  30 500 50],'BackgroundColor',[0.95,0.9,0.85],'BorderType','line');

selection1='Siemens scanner'; selection2='7T scanner'; selection3='Cervical spine'; selection4='Resting-state fMRI'; selection5='Pre-processing only';
selection6=''; flag=0;
while strcmp(selection5,'DONE')~=1
    [selection1,selection2,selection3,selection4,selection5,selection6] = initdialog(bg1,bg2,bg3,bg4,bg5,bg6,flag,selection1,selection2,selection3,selection4,selection5,selection6);
    if strcmp(selection6,'DONE')==1
        closereq()
        break
    end
    if strcmp(selection6,'Help')==1
        fprintf('neptune_manual.pdf\n')
        open('neptune_manual.pdf')
        flag=1;
    end
end
try close(fig1); catch, end
clear fig1 bg1 bg2 bg3 bg4 bg5 bg6 flag selection6

% %% --- choose resting-state or task fMRI ---
switch selection4
    case 'Resting-state fMRI'
        rest=1;
    case 'Task fMRI'
        rest=0;
end; clear answer1
task=~rest;

switch selection1
    case 'Siemens scanner'
        scanner = 1;
    case 'Philips scanner'
        scanner = 2;
    case 'Other'
        quest = sprintf('Do you have reconstructed NIfTI files already?');
        answer2 = questdlg(quest,'NIfTI?','Yes'); clear quest
        switch answer2
            case 'Yes'
                scanner = 3;
            case 'No'
                msgbox('Sorry, only Siemens and Philips scanners are supported for now', 'Abort','error');
                tmp=clock; clk = sprintf('%.4d-%.2d-%.2d_%.2d-%.2d',tmp(1),tmp(2),tmp(3),tmp(4),tmp(5));
                save([home_dir sprintf('workspace_cancel_%s',clk)])
                msgbox(sprintf('Your choices so far have been saved to\n%s\nTerminating execution...',sprintf('workspace_cancel_%s.mat',clk)),'Choices saved','custom',msgicon); clear clk tmp
                return
            case 'Cancel'
                tmp=clock; clk = sprintf('%.4d-%.2d-%.2d_%.2d-%.2d',tmp(1),tmp(2),tmp(3),tmp(4),tmp(5));
                save([home_dir sprintf('workspace_cancel_%s',clk)])
                msgbox(sprintf('Your choices so far have been saved to\n%s\nTerminating execution...',sprintf('workspace_cancel_%s.mat',clk)),'Choices saved','custom',msgicon); clear clk tmp
                return
        end; clear answer2
end; clear answer1

switch selection2
    case '7T scanner'
        scan7T=1;
    case '3T scanner'
        scan7T=0;
    case 'Other'
        scan7T=0;
end; clear answer1
scan3T=~scan7T;

switch selection3
    case 'Cervical spine'
        C_spine=1;
    case 'Thoraco-lumbar spine'
        C_spine=0;
end; clear answer1
clear selection1 selection2 selection3 selection4
L_spine=~C_spine;

switch selection5
    case 'Pre-processing only'
        preproc=1; postproc=0;
    case 'Post-processing only'
        preproc=0; postproc=1;
    case 'Both'
        preproc=1; postproc=1;
end; clear answer1 selection5
nifti_convert=0;

end

%% --- choose home directory that contains all subject folders ---
try waitbar(2/8,wbar1,'Choose home directory...'); catch, end

if ~exist('home_dir','var')

answer1 = questdlg(sprintf('Choose home directory that contains all subject folders...'),'Home directory','Choose now','Not in the mood to continue today','Choose now');
switch answer1
    case 'Not in the mood to continue today'
        questdlg('Take a break, see you later!','Bye','Welcome, you too','Welcome, you too');
        tmp=clock; clk = sprintf('%.4d-%.2d-%.2d_%.2d-%.2d',tmp(1),tmp(2),tmp(3),tmp(4),tmp(5));
        save([first_dir sprintf('workspace_cancel_%s',clk)])
        msgbox(sprintf('Your choices so far have been saved to\n%s\nTerminating execution...',sprintf('workspace_cancel_%s.mat',clk)),'Choices saved','custom',msgicon); clear clk tmp
        return
end; clear answer1

home_dir = uigetdir(first_dir,'Choose home directory that contains all subject folders...');
if sum(isspace(home_dir))==1 % if home directory contains spaces
    prompt = sprintf('There cannot be spaces in any directory name.\nCheck for spaces: %s',home_dir);
    questdlg(prompt,'Error', 'I will try again after either renaming the affected directory or runing from a different path', 'I will try again after either renaming the affected directory or runing from a different path'); clear prompt
    tmp=clock; clk = sprintf('%.4d-%.2d-%.2d_%.2d-%.2d',tmp(1),tmp(2),tmp(3),tmp(4),tmp(5));
    save([home_dir sprintf('workspace_error_%s',clk)])
    msgbox(sprintf('Your choices so far have been saved to\n%s\nTerminating execution...',sprintf('workspace_error_%s.mat',clk)),'Choices saved','custom',msgicon); clear clk tmp
    return
end

home_dir = [home_dir '/'];
cd(home_dir)
temp = dir;
f=0;
for k=1:length(temp)
    if temp(k).isdir==1
        f=f+1;
        sdir1(f) = temp(k);
    end
end; clear k temp f
sdir1=sdir1(3:end);


else
if strcmp([pwd '/'],home_dir)~=1
quest = sprintf('Could you please confirm that this is your home directory that contains all subject folders?:\n   %s',home_dir);
answer_homedir = questdlg(quest,'Verifying home directory...', 'Yes, correct', 'No, reselect home directory', 'Yes, correct'); clear quest
switch answer_homedir
	case 'No, reselect home directory'
	redo_directories_flag = 1;
    home_dir = uigetdir(first_dir,'Choose home directory that contains all subject folders...');
    if sum(isspace(home_dir))==1 % if home directory contains spaces
        prompt = sprintf('There cannot be spaces in any directory name.\nCheck for spaces: %s',home_dir);
        questdlg(prompt,'Error', 'I will try again after either renaming the affected directory or runing from a different path', 'I will try again after either renaming the affected directory or runing from a different path'); clear prompt
        tmp=clock; clk = sprintf('%.4d-%.2d-%.2d_%.2d-%.2d',tmp(1),tmp(2),tmp(3),tmp(4),tmp(5));
        save([home_dir sprintf('workspace_error_%s',clk)])
        msgbox(sprintf('Your choices so far have been saved to\n%s\nTerminating execution...',sprintf('workspace_error_%s.mat',clk)),'Choices saved','custom',msgicon); clear clk tmp
        return
    end

    home_dir = [home_dir '/'];
    cd(home_dir)
    temp = dir;
    f=0;
    for k=1:length(temp)
        if temp(k).isdir==1
            f=f+1;
            sdir1(f) = temp(k);
        end
    end; clear k temp f
    sdir1=sdir1(3:end);
	case 'Yes, correct'
end
end
end

try
fileID = fopen([home_dir sprintf('log_neptune_%s.txt',clk_init)],'w'); clear tmp_init
fprintf(fileID,'%s\n',logg{1}); fprintf(fileID,'%s\n',logg{2}); 
catch
end

%% --- choose subjects ---
try waitbar(3/8,wbar1,'Choose subjects...'); catch, end

if ~exist('sub_dir','var') || exist('redo_directories_flag','var')

quest = sprintf('Please select subjects for processing');
answer1 = questdlg(quest,'Subjects','Alright, take me there :)','Nah, not today :(','Alright, take me there :)'); clear quest
switch answer1
    case 'Alright, take me there :)'
        cd(home_dir)
        sdir2 = dir; sdir2(find([sdir2.isdir]==0))=[]; sdir2=sdir2(3:end); sdir2={sdir2.name};
        prompt = sprintf('Select subjects...\n(hold Ctrl or Command button and click to choose multiple subjects)');
        indx = listdlg('PromptString',{prompt,''},...
            'SelectionMode','multiple','ListSize',[400 400],'ListString',sdir2);
        for k=1:length(indx)
            sub_dir{k,1} = [home_dir sdir2{indx(k)} '/'];
        end; clear k2 indx sdir2 prompt

    case 'Nah, not today :('
        tmp=clock; clk = sprintf('%.4d-%.2d-%.2d_%.2d-%.2d',tmp(1),tmp(2),tmp(3),tmp(4),tmp(5));
        save([home_dir sprintf('workspace_cancel_%s',clk)])
        msgbox(sprintf('Your choices so far have been saved to\n%s\nTerminating execution...',sprintf('workspace_cancel_%s.mat',clk)),'Choices saved','custom',msgicon); clear clk tmp
        return
end; clear answer1 sdir1

end

for k=1:length(sub_dir)
    subnames{k,1} = sub_dir{k}(max(strfind(sub_dir{k}(1:end-1),'/'))+1:end-1);
    if isempty(subnames{k,1})
        subnames{k,1} = sub_dir{k}(max(strfind(sub_dir{k}(1:end-1),'\'))+1:end-1);
    end
end; clear k

%% --- Subject-after-subject OR step-after-step ---
if ~exist('step_after_step','var')

step_after_step=1;
if length(sub_dir)~=1 % This step is inappropriate if only one subject is being processed
quest = sprintf('Do you want to perform all steps one subject after the other (subject after subject), OR one step at a time in all subjects (step after step)?');
answer1 = questdlg(quest,'A quick question','Subject after subject','Step after step','Whatever!','Step after step'); clear quest
switch answer1
    case 'Subject after subject'
        step_after_step=0;
    case 'Step after step'
        step_after_step=1;
    case 'Whatever!'
        quest = sprintf('I chose to run the pipeline for you step after step as the default option.');
        questdlg(quest,'Step after step','Fantastic!','Fantastic!'); clear quest
        step_after_step=1;
end; clear answer1
end

end

if step_after_step==0 % if doing subject after subject, ask which subject to start processing from (because some people run Neptune repeatedly on the same dataset and might want to restart from one of the middle subjects 'S' after processing has previously completed for the first 'S-1' subjects).
    prompt = sprintf('Which subject do you want to start processing from? (mention subject number)\n(Default value is 1 - start from the first subject)\n');
    answer3 = inputdlg(prompt,'Subject number to start from',1,{'1'}); clear prompt
    sub_aft_sub_num = str2num(cell2mat(answer3)); clear answer3
end

%% --- dicom (or PAR/REC) to NIfTI ---
if preproc==1 % no need to ask this if the user is only interested in post-processing
if exist('answer_NIfTIconvert','var')
    if strcmp(answer_NIfTIconvert,'Yes, convert')==1
        skip_NIfTIconvert_choice = 1; % it won't ask you to choose again if you want to perform NIfTI conversion
        scfMRItb_01_init_NIfTIconvert
    end
else
    skip_NIfTIconvert_choice = 0; % it will ask you if you want to perform NIfTI conversion, because the choice has not been made yet
    scfMRItb_01_init_NIfTIconvert
end
end

%% --- choose runs ---
try waitbar(5/8,wbar1,'Mention number of fMRI runs...'); catch, end

if ~exist('runs','var')

if length(sub_dir)~=1 % This step is inappropriate if only one subject is being processed
    quest = sprintf('Number of fMRI runs');
    answer1 = questdlg(quest,'Runs','Same for all subjects','Different across subjects','Same for all subjects'); clear quest
else
    answer1 = 'Same for all subjects';
end
switch answer1
    case 'Same for all subjects'
        prompt = sprintf('Number of fMRI runs in each subject');
        answer2 = inputdlg(prompt,'Runs',1,{'1'}); clear prompt
        runs = str2num(cell2mat(answer2)); clear answer2
        runs = repmat(runs,length(sub_dir),1);
    case 'Different across subjects'
        prompt = sprintf('Number of fMRI runs: enter number for each subject separated by a space');
        string1 = [];
        for k=1:length(sub_dir)
            string1 = strcat(string1, '1', {' '});
        end
        answer2 = inputdlg(prompt,'Runs',1,{cell2mat(string1)}); clear prompt string1
        runs = str2num(cell2mat(answer2)); clear answer2
        if length(runs)~=length(sub_dir)
            runs(end+1:length(sub_dir)) = runs(end);
        end
end; clear answer1
end

%% --- choose anatomical files ---
try waitbar(6/8,wbar1,'Choose anatomical files...'); catch, end

if ~exist('anat_files','var') || exist('redo_directories_flag','var')

quest = sprintf('Anatomical files: do you have a specific wildcard search string or will you choose the files in each subject?');
answer1 = questdlg(quest,'Anatomical files','Wildcard search string','Open subject folders','Open subject folders'); clear quest

switch answer1
    case 'Wildcard search string'
        prompt = sprintf('Enter wildcard search string (NIfTI anatomical files)');
        answer2 = inputdlg(prompt,'Anatomical files',1,{'sub-*_T1w.nii*'}); clear prompt
        anat_string = cell2mat(answer2); clear answer2
        
        for k=1:length(sub_dir)
            nf(k,1) = numel(dir([sub_dir{k} '**/' anat_string]));
        end; clear k
        S1=sub_dir(find(nf==0));
        for k=1:length(S1)
            s1 = strfind(S1{k},'/'); s1 = s1(end-1)+1;
            S1{k} = S1{k}(s1:end-1); clear s1
        end
        S2=sub_dir(find(nf>=2));
        for k=1:length(S2)
            s2 = strfind(S2{k},'/'); s2 = s2(end-1)+1;
            S2{k} = S2{k}(s2:end-1); clear s2
        end
        
        if (numel(S1)~=0)||(numel(S2)~=0)
        flg=1;
        quest = sprintf('Issues in locating anatomical files (see below). Do you want to try wildcard search again or choose files individually in each subject?\n\nNo files found in subject(s) - %s\n\nMore than one file in subject(s) - %s',strjoin(S1,', '),strjoin(S2,', ')); clear nf S1 S2
        answerb = questdlg(quest,'Anatomical files','Wildcard search string','Open subject folders','Open subject folders'); clear quest
        
        switch answerb
            case 'Wildcard search string'
                prompt = sprintf('Enter wildcard search string (NIfTI anatomical files)');
                answer2 = inputdlg(prompt,'Anatomical files',1,{'sub-*_T1w.nii*'}); clear prompt
                anat_string = cell2mat(answer2); clear answer2
                
                for k=1:length(sub_dir)
                    nf(k,1) = numel(dir([sub_dir{k} '**/' anat_string]));
                end; clear k
                S1=sub_dir(find(nf==0));
                for k=1:length(S1)
                    s1 = strfind(S1{k},'/'); s1 = s1(end-1)+1;
                    S1{k} = S1{k}(s1:end-1); clear s1
                end
                S2=sub_dir(find(nf>=2));
                for k=1:length(S2)
                    s2 = strfind(S2{k},'/'); s2 = s2(end-1)+1;
                    S2{k} = S2{k}(s2:end-1); clear s2
                end
                if (numel(S1)~=0)||(numel(S2)~=0)
                    quest = sprintf('Again, issues in locating anatomical files (see below). Please CHOOSE files individually in each subject.\n\nNo files found in subject(s) - %s\n\nMore than one file in subject(s) - %s',strjoin(S1,', '),strjoin(S2,', '));
                    answerc = questdlg(quest,'Anatomical files','OK, take me to subject folders','OK, take me to subject folders'); clear quest
                else
                    name=[];
                    for k=1:length(sub_dir)
                        cd(sub_dir{k})
                        sdir2 = dir(['**/' anat_string]); sdir2_folder = {sdir2.folder}; sdir2_name = {sdir2.name};
                        if strcmp(sdir2_name{1}(end-3:end),'.nii')==1
                            anat_files{k,1} = [sdir2_name{1}(1:end-4) erase(sdir2_name{1}(end-3:end),'.nii')];  % for .nii extension, this will also work here: sdir2{indx}(1:end-4);
                        else
                            anat_files{k,1} = [sdir2_name{1}(1:end-7) erase(sdir2_name{1}(end-6:end),'.nii.gz')];  % for .nii.gz extension, this will also work here: sdir2{indx}(1:end-7);
                        end
                        anat_home{k,1} = [sdir2_folder{1} '/'];
                        if strcmp(anat_home{k,1},sub_dir{k})==0
                            fprintf('Subject-%d''s anatomical is in a subfolder within the main subject directory (/%s). Moving it (and associated files) to the latter location.\n',k,erase(anat_home{k},sub_dir{k}))
                            clear sdir2_folder sdir2 sdir2_name
                            sdir2 = dir([anat_home{k,1} anat_files{k,1} '*']); sdir2_folder = {sdir2.folder}; sdir2_name = {sdir2.name};
                            for k2=1:length(sdir2_folder)
                                movefile([sdir2_folder{k2} '/' sdir2_name{k2}],[sub_dir{k} sdir2_name{k2}],'f')
                            end
                        end
                        name = strcat(name, sprintf('\n%s%s / %s',subnames{k},string(erase(sdir2_folder{1},sub_dir{k}(1:end-1))),anat_files{k,1}));
                        clear sdir2 sdir2_folder sdir2_name k2
                        cd ..
                    end; clear k
                    
                    quest = sprintf('Are these the NIfTI anatomical files?:\n%s',name);
                    answer2 = questdlg(quest,'Anatomical files','Yes','No, open subject folders','Yes'); clear quest
                    switch answer2
                        case 'No, open subject folders'
                            clear anat_files
                            for k=1:length(sub_dir)
                                cd(sub_dir{k})
                                sdir2 = cat(1,dir('**/*.nii'),dir('**/*.nii.gz'));
                                sdir2_folder = {sdir2.folder}; sdir2_name = {sdir2.name}; sdir2 = {sdir2.name};
                                for k2=1:length(sdir2_folder)
                                    sdir2_name{k2} = [erase(sdir2_folder{k2},sub_dir{k}(1:end-1)) '/' sdir2{k2}];
                                end
                                prompt = sprintf('Select NIfTI anatomical image for subject (%d) of (%d)...\n%s\n',k,length(sub_dir),sub_dir{k});
                                indx = listdlg('PromptString',{prompt,...
                                    'Only one file can be selected at a time.',''},...
                                    'SelectionMode','single','ListSize',[400 400],'ListString',sdir2_name);
                                cd ..
                                if strcmp(sdir2{indx}(end-3:end),'.nii')==1
                                    anat_files{k,1} = [sdir2{indx}(1:end-4) erase(sdir2{indx}(end-3:end),'.nii')];  % for .nii extension, this will also work here: sdir2{indx}(1:end-4);
                                else
                                    anat_files{k,1} = [sdir2{indx}(1:end-7) erase(sdir2{indx}(end-6:end),'.nii.gz')];  % for .nii.gz extension, this will also work here: sdir2{indx}(1:end-7);
                                end
                                anat_home{k,1} = [sdir2_folder{indx} '/'];
                                if strcmp(anat_home{k,1},sub_dir{k})==0
                                    fprintf('Subject-%d''s anatomical is in a subfolder within the main subject directory (/%s). Moving it (and associated files) to the latter location.\n',k,erase(anat_home{k},sub_dir{k}))
                                    clear sdir2_folder sdir2 sdir2_name
                                    sdir2 = dir([anat_home{k,1} anat_files{k,1} '*']); sdir2_folder = {sdir2.folder}; sdir2_name = {sdir2.name};
                                    for k2=1:length(sdir2_folder)
                                        movefile([sdir2_folder{k2} '/' sdir2_name{k2}],[sub_dir{k} sdir2_name{k2}],'f')
                                    end
                                end
                                clear sdir2 prompt sdir2_folder sdir2_name k2 indx f
                            end; clear k
                    end
                end
                clear nf S1 S2
            case 'No, open subject folders'
                clear anat_files
                for k=1:length(sub_dir)
                    cd(sub_dir{k})
                    sdir2 = cat(1,dir('**/*.nii'),dir('**/*.nii.gz'));
                    sdir2_folder = {sdir2.folder}; sdir2_name = {sdir2.name}; sdir2 = {sdir2.name};
                    for k2=1:length(sdir2_folder)
                        sdir2_name{k2} = [erase(sdir2_folder{k2},sub_dir{k}(1:end-1)) '/' sdir2{k2}];
                    end
                    prompt = sprintf('Select NIfTI anatomical image for subject (%d) of (%d)...\n%s\n',k,length(sub_dir),sub_dir{k});
                    indx = listdlg('PromptString',{prompt,...
                        'Only one file can be selected at a time.',''},...
                        'SelectionMode','single','ListSize',[400 400],'ListString',sdir2_name);
                    cd ..
                    if strcmp(sdir2{indx}(end-3:end),'.nii')==1
                        anat_files{k,1} = [sdir2{indx}(1:end-4) erase(sdir2{indx}(end-3:end),'.nii')];  % for .nii extension, this will also work here: sdir2{indx}(1:end-4);
                    else
                        anat_files{k,1} = [sdir2{indx}(1:end-7) erase(sdir2{indx}(end-6:end),'.nii.gz')];  % for .nii.gz extension, this will also work here: sdir2{indx}(1:end-7);
                    end
                    anat_home{k,1} = [sdir2_folder{indx} '/'];
                    if strcmp(anat_home{k,1},sub_dir{k})==0
                        fprintf('Subject-%d''s anatomical is in a subfolder within the main subject directory (/%s). Moving it (and associated files) to the latter location.\n',k,erase(anat_home{k},sub_dir{k}))
                        clear sdir2_folder sdir2 sdir2_name
                        sdir2 = dir([anat_home{k,1} anat_files{k,1} '*']); sdir2_folder = {sdir2.folder}; sdir2_name = {sdir2.name};
                        for k2=1:length(sdir2_folder)
                            movefile([sdir2_folder{k2} '/' sdir2_name{k2}],[sub_dir{k} sdir2_name{k2}],'f')
                        end
                    end
                    clear sdir2 prompt sdir2_folder sdir2_name k2 indx f
                end; clear k
        end
        if exist('answerc','var')
            clear anat_files
            for k=1:length(sub_dir)
                cd(sub_dir{k})
                sdir2 = cat(1,dir('**/*.nii'),dir('**/*.nii.gz'));
                sdir2_folder = {sdir2.folder}; sdir2_name = {sdir2.name}; sdir2 = {sdir2.name};
                for k2=1:length(sdir2_folder)
                    sdir2_name{k2} = [erase(sdir2_folder{k2},sub_dir{k}(1:end-1)) '/' sdir2{k2}];
                end
                prompt = sprintf('Select NIfTI anatomical image for subject (%d) of (%d)...\n%s\n',k,length(sub_dir),sub_dir{k});
                indx = listdlg('PromptString',{prompt,...
                    'Only one file can be selected at a time.',''},...
                    'SelectionMode','single','ListSize',[400 400],'ListString',sdir2_name);
                cd ..
                if strcmp(sdir2{indx}(end-3:end),'.nii')==1
                    anat_files{k,1} = [sdir2{indx}(1:end-4) erase(sdir2{indx}(end-3:end),'.nii')];  %#ok<*SAGROW> % for .nii extension, this will also work here: sdir2{indx}(1:end-4);
                else
                    anat_files{k,1} = [sdir2{indx}(1:end-7) erase(sdir2{indx}(end-6:end),'.nii.gz')];  % for .nii.gz extension, this will also work here: sdir2{indx}(1:end-7);
                end
                anat_home{k,1} = [sdir2_folder{indx} '/'];
                if strcmp(anat_home{k,1},sub_dir{k})==0
                    fprintf('Subject-%d''s anatomical is in a subfolder within the main subject directory (/%s). Moving it (and associated files) to the latter location.\n',k,erase(anat_home{k},sub_dir{k}))
                    clear sdir2_folder sdir2 sdir2_name
                    sdir2 = dir([anat_home{k,1} anat_files{k,1} '*']); sdir2_folder = {sdir2.folder}; sdir2_name = {sdir2.name};
                    for k2=1:length(sdir2_folder)
                        movefile([sdir2_folder{k2} '/' sdir2_name{k2}],[sub_dir{k} sdir2_name{k2}],'f')
                    end
                end
                clear sdir2 prompt sdir2_folder sdir2_name k2 indx f
            end; clear k
        end
        
        else
        name=[];
        for k=1:length(sub_dir)
            cd(sub_dir{k})
            sdir2 = dir(['**/' anat_string]); sdir2_folder = {sdir2.folder}; sdir2_name = {sdir2.name};
            if strcmp(sdir2_name{1}(end-3:end),'.nii')==1
                anat_files{k,1} = [sdir2_name{1}(1:end-4) erase(sdir2_name{1}(end-3:end),'.nii')];  % for .nii extension, this will also work here: sdir2{indx}(1:end-4);
            else
                anat_files{k,1} = [sdir2_name{1}(1:end-7) erase(sdir2_name{1}(end-6:end),'.nii.gz')];  % for .nii.gz extension, this will also work here: sdir2{indx}(1:end-7);
            end
            anat_home{k,1} = [sdir2_folder{1} '/'];
            if strcmp(anat_home{k,1},sub_dir{k})==0
                fprintf('Subject-%d''s anatomical is in a subfolder within the main subject directory (/%s). Moving it (and associated files) to the latter location.\n',k,erase(anat_home{k},sub_dir{k}))
                clear sdir2_folder sdir2 sdir2_name
                sdir2 = dir([anat_home{k,1} anat_files{k,1} '*']); sdir2_folder = {sdir2.folder}; sdir2_name = {sdir2.name};
                for k2=1:length(sdir2_folder)
                    movefile([sdir2_folder{k2} '/' sdir2_name{k2}],[sub_dir{k} sdir2_name{k2}],'f')
                end
            end
            name = strcat(name, sprintf('\n%s%s / %s',subnames{k},string(erase(sdir2_folder{1},sub_dir{k}(1:end-1))),anat_files{k,1}));
            clear sdir2 sdir2_folder sdir2_name k2
            cd ..
        end; clear k
        
        quest = sprintf('Are these the NIfTI anatomical files?:\n%s',name);
        answer2 = questdlg(quest,'Anatomical files','Yes','No, open subject folders','Yes'); clear quest
        switch answer2
            case 'No, open subject folders'
                clear anat_files
                for k=1:length(sub_dir)
                    cd(sub_dir{k})
                    sdir2 = cat(1,dir('**/*.nii'),dir('**/*.nii.gz'));
                    sdir2_folder = {sdir2.folder}; sdir2_name = {sdir2.name}; sdir2 = {sdir2.name};
                    for k2=1:length(sdir2_folder)
                        sdir2_name{k2} = [erase(sdir2_folder{k2},sub_dir{k}(1:end-1)) '/' sdir2{k2}];
                    end
                    prompt = sprintf('Select NIfTI anatomical image for subject (%d) of (%d)...\n%s\n',k,length(sub_dir),sub_dir{k});
                    indx = listdlg('PromptString',{prompt,...
                        'Only one file can be selected at a time.',''},...
                        'SelectionMode','single','ListSize',[400 400],'ListString',sdir2_name);
                    cd ..
                    if strcmp(sdir2{indx}(end-3:end),'.nii')==1
                        anat_files{k,1} = [sdir2{indx}(1:end-4) erase(sdir2{indx}(end-3:end),'.nii')];  % for .nii extension, this will also work here: sdir2{indx}(1:end-4);
                    else
                        anat_files{k,1} = [sdir2{indx}(1:end-7) erase(sdir2{indx}(end-6:end),'.nii.gz')];  % for .nii.gz extension, this will also work here: sdir2{indx}(1:end-7);
                    end
                    anat_home{k,1} = [sdir2_folder{indx} '/'];
                    if strcmp(anat_home{k,1},sub_dir{k})==0
                        fprintf('Subject-%d''s anatomical is in a subfolder within the main subject directory (/%s). Moving it (and associated files) to the latter location.\n',k,erase(anat_home{k},sub_dir{k}))
                        clear sdir2_folder sdir2 sdir2_name
                        sdir2 = dir([anat_home{k,1} anat_files{k,1} '*']); sdir2_folder = {sdir2.folder}; sdir2_name = {sdir2.name};
                        for k2=1:length(sdir2_folder)
                            movefile([sdir2_folder{k2} '/' sdir2_name{k2}],[sub_dir{k} sdir2_name{k2}],'f')
                        end
                    end
                    clear sdir2 prompt sdir2_folder sdir2_name k2 indx f
                end; clear k
        end
        end
        
    case 'Open subject folders'
        for k=1:length(sub_dir)
            cd(sub_dir{k})
            sdir2 = cat(1,dir('**/*.nii'),dir('**/*.nii.gz'));
            sdir2_folder = {sdir2.folder}; sdir2_name = {sdir2.name}; sdir2 = {sdir2.name};
            for k2=1:length(sdir2_folder)
                sdir2_name{k2} = [erase(sdir2_folder{k2},sub_dir{k}(1:end-1)) '/' sdir2{k2}];
            end
            prompt = sprintf('Select NIfTI anatomical image for subject (%d) of (%d)...\n%s\n',k,length(sub_dir),sub_dir{k});
            indx = listdlg('PromptString',{prompt,...
                'Only one file can be selected at a time.',''},...
                'SelectionMode','single','ListSize',[400 400],'ListString',sdir2_name);
            cd ..
            if strcmp(sdir2{indx}(end-3:end),'.nii')==1
                anat_files{k,1} = [sdir2{indx}(1:end-4) erase(sdir2{indx}(end-3:end),'.nii')];  % for .nii extension, this will also work here: sdir2{indx}(1:end-4);
            else
                anat_files{k,1} = [sdir2{indx}(1:end-7) erase(sdir2{indx}(end-6:end),'.nii.gz')];  % for .nii.gz extension, this will also work here: sdir2{indx}(1:end-7);
            end
            anat_home{k,1} = [sdir2_folder{indx} '/'];
            if strcmp(anat_home{k,1},sub_dir{k})==0
                fprintf('Subject-%d''s anatomical is in a subfolder within the main subject directory (/%s). Moving it (and associated files) to the latter location.\n',k,erase(anat_home{k},sub_dir{k}))
                clear sdir2_folder sdir2 sdir2_name
                sdir2 = dir([anat_home{k,1} anat_files{k,1} '*']); sdir2_folder = {sdir2.folder}; sdir2_name = {sdir2.name};
                for k2=1:length(sdir2_folder)
                    movefile([sdir2_folder{k2} '/' sdir2_name{k2}],[sub_dir{k} sdir2_name{k2}],'f')
                end
            end
            clear sdir2 prompt sdir2_folder sdir2_name k2 indx f
        end; clear k
end; clear answer1 f answer2 answerb answerc

end

%% --- choose functional files ---
try waitbar(7/8,wbar1,'Choose functional files...'); catch, end

if ~exist('func_files','var') || exist('redo_directories_flag','var')

quest = sprintf('Functional files: do you have a specific wildcard search string or will you choose the files in each subject?');
answer1 = questdlg(quest,'Functional files','Wildcard search string','Open subject folders','Open subject folders'); clear quest

switch answer1
    case 'Wildcard search string'
        prompt = sprintf('Enter wildcard search string (NIfTI functional files)');
        answer2 = inputdlg(prompt,'Functional files',1,{'sub-*_task-*_bold.nii*'}); clear prompt
        func_string = cell2mat(answer2); clear answer2
        
        for k=1:length(sub_dir)
            nf(k,1) = numel(dir([sub_dir{k} '**/' func_string]));
        end; clear k
        S1=sub_dir(find(nf==0));
        for k=1:length(S1)
            s1 = strfind(S1{k},'/'); s1 = s1(end-1)+1;
            S1{k} = S1{k}(s1:end-1); clear s1
        end
        S2=sub_dir(find(nf>runs));
        for k=1:length(S2)
            s2 = strfind(S2{k},'/'); s2 = s2(end-1)+1;
            S2{k} = S2{k}(s2:end-1); clear s2
        end
        
        if (numel(S1)~=0)||(numel(S2)~=0)
        flg=1;
        quest = sprintf('Issues in locating functional files (see below). Do you want to try wildcard search again or choose files individually in each subject?\n\nNo files found in subject(s) - %s\n\nMore files than #runs in subject(s) - %s',strjoin(S1,', '),strjoin(S2,', ')); clear nf S1 S2
        answerb = questdlg(quest,'Functional files','Wildcard search string','Open subject folders','Open subject folders'); clear quest
        
        switch answerb
            case 'Wildcard search string'
                prompt = sprintf('Enter wildcard search string (NIfTI functional files)');
                answer2 = inputdlg(prompt,'Functional files',1,{'sub-*_task-*_bold.nii*'}); clear prompt
                func_string = cell2mat(answer2); clear answer2
                
                for k=1:length(sub_dir)
                    nf(k,1) = numel(dir([sub_dir{k} '**/' func_string]));
                end; clear k
                S1=sub_dir(find(nf==0));
                for k=1:length(S1)
                    s1 = strfind(S1{k},'/'); s1 = s1(end-1)+1;
                    S1{k} = S1{k}(s1:end-1); clear s1
                end
                S2=sub_dir(find(nf>runs));
                for k=1:length(S2)
                    s2 = strfind(S2{k},'/'); s2 = s2(end-1)+1;
                    S2{k} = S2{k}(s2:end-1); clear s2
                end
                if (numel(S1)~=0)||(numel(S2)~=0)
                    quest = sprintf('Again, issues in locating functional files (see below). Please CHOOSE files individually in each subject.\n\nNo files found in subject(s) - %s\n\nMore files than #runs in subject(s) - %s',strjoin(S1,', '),strjoin(S2,', '));
                    answerc = questdlg(quest,'Functional files','OK, take me to subject folders','OK, take me to subject folders'); clear quest
                else
                    name=[];
                    for k=1:length(sub_dir)
                        cd(sub_dir{k})
                        sdir2 = dir(['**/' func_string]); sdir2_folder = {sdir2.folder}; sdir2_name = {sdir2.name};
                        for k2=1:runs(k)
                            if strcmp(sdir2_name{k2}(end-3:end),'.nii')==1
                                func_files{k,k2} = [sdir2_name{k2}(1:end-4) erase(sdir2_name{k2}(end-3:end),'.nii')];  % for .nii extension, this will also work here: sdir2{indx(k2)}(1:end-4);
                            else
                                func_files{k,k2} = [sdir2_name{k2}(1:end-7) erase(sdir2_name{k2}(end-6:end),'.nii.gz')];  % for .nii.gz extension, this will also work here: sdir2{indx(k2)}(1:end-7);
                            end
                            func_home{k,k2}  = [sdir2_folder{k2} '/'];
                            name = strcat(name, sprintf('\n%s%s / %s',subnames{k},string(erase(sdir2_folder{1},sub_dir{k}(1:end-1))),func_files{k,k2}));
                            if strcmp(func_home{k,k2},sub_dir{k})==0
                                fprintf('Subject-%d run-%d''s functional is in a subfolder within the main subject directory (/%s). Moving it (and associated files) to the latter location.\n',k,k2,erase(func_home{k,k2},sub_dir{k}))
                                sdir3 = cat(1,dir([func_home{k,k2} func_files{k,k2} '*']), dir([func_home{k,k2} sprintf('workspace_variables_run%d.mat',k2)]), dir([func_home{k,k2} sprintf('run%d_final_reg.mat',k2)]), dir([func_home{k,k2} 'cord_quadrants.mat']), dir([func_home{k,k2} 'cross_sectional_areas.mat']), dir([func_home{k,k2} 'motion_parameters.mat']), dir([func_home{k,k2} sprintf('tSNR_across_preproc_steps_run%d.mat',k2)]));
                                sdir3_folder = {sdir3.folder}; sdir3_name = {sdir3.name};
                                for k3=1:length(sdir3_folder)
                                    movefile([sdir3_folder{k3} '/' sdir3_name{k3}],[sub_dir{k} sdir3_name{k3}],'f')
                                end
                                clear sdir3_folder sdir3 sdir3_name
                            end
                        end
                        clear sdir2_folder sdir2 sdir2_name k2
                        cd ..
                    end; clear k
                    
                    quest = sprintf('Are these the NIfTI functional files?:\n%s',name); clear name
                    answer2 = questdlg(quest,'Functional files','Yes','No, open subject folders','Yes'); clear quest
                    switch answer2
                        case 'No, open subject folders'
                            clear func_files
                            for k=1:length(sub_dir)
                                cd(sub_dir{k})
                                sdir2 = cat(1,dir('**/*.nii'),dir('**/*.nii.gz'));
                                sdir2_folder = {sdir2.folder}; sdir2_name = {sdir2.name}; sdir2 = {sdir2.name};
                                for k2=1:length(sdir2_folder)
                                    sdir2_name{k2} = [erase(sdir2_folder{k2},sub_dir{k}(1:end-1)) '/' sdir2{k2}];
                                end
                                prompt = sprintf('Select NIfTI functional image(s) (all runs) for subject (%d) of (%d)...\n%s',k,length(sub_dir),sub_dir{k});
                                indx = listdlg('PromptString',{prompt,''},...
                                    'SelectionMode','multiple','ListSize',[400 400],'ListString',sdir2_name);
                                cd ..
                                for k2=1:runs(k)
                                    if strcmp(sdir2{indx(k2)}(end-3:end),'.nii')==1
                                        func_files{k,k2} = [sdir2{indx(k2)}(1:end-4) erase(sdir2{indx(k2)}(end-3:end),'.nii')];  % for .nii extension, this will also work here: sdir2{indx(k2)}(1:end-4);
                                    else
                                        func_files{k,k2} = [sdir2{indx(k2)}(1:end-7) erase(sdir2{indx(k2)}(end-6:end),'.nii.gz')];  % for .nii.gz extension, this will also work here: sdir2{indx(k2)}(1:end-7);
                                    end
                                    func_home{k,k2}  = [sdir2_folder{indx(k2)} '/'];
                                    if strcmp(func_home{k,k2},sub_dir{k})==0
                                        fprintf('Subject-%d run-%d''s functional is in a subfolder within the main subject directory (/%s). Moving it (and associated files) to the latter location.\n',k,k2,erase(func_home{k,k2},sub_dir{k}))
                                        sdir3 = cat(1,dir([func_home{k,k2} func_files{k,k2} '*']), dir([func_home{k,k2} sprintf('workspace_variables_run%d.mat',k2)]), dir([func_home{k,k2} sprintf('run%d_final_reg.mat',k2)]), dir([func_home{k,k2} 'cord_quadrants.mat']), dir([func_home{k,k2} 'cross_sectional_areas.mat']), dir([func_home{k,k2} 'motion_parameters.mat']), dir([func_home{k,k2} sprintf('tSNR_across_preproc_steps_run%d.mat',k2)]));
                                        sdir3_folder = {sdir3.folder}; sdir3_name = {sdir3.name};
                                        for k3=1:length(sdir3_folder)
                                            movefile([sdir3_folder{k3} '/' sdir3_name{k3}],[sub_dir{k} sdir3_name{k3}],'f')
                                        end
                                        clear sdir3_folder sdir3 sdir3_name
                                    end
                                end
                                clear sdir2 prompt sdir2_folder sdir2_name k2 k3 indx f
                            end; clear k f
                    end
                end
                clear nf S1 S2
            case 'Open subject folders'
                clear func_files
                for k=1:length(sub_dir)
                    cd(sub_dir{k})
                    sdir2 = cat(1,dir('**/*.nii'),dir('**/*.nii.gz'));
                    sdir2_folder = {sdir2.folder}; sdir2_name = {sdir2.name}; sdir2 = {sdir2.name};
                    for k2=1:length(sdir2_folder)
                        sdir2_name{k2} = [erase(sdir2_folder{k2},sub_dir{k}(1:end-1)) '/' sdir2{k2}];
                    end
                    prompt = sprintf('Select NIfTI functional image(s) (all runs) for subject (%d) of (%d)...\n%s',k,length(sub_dir),sub_dir{k});
                    indx = listdlg('PromptString',{prompt,''},...
                        'SelectionMode','multiple','ListSize',[400 400],'ListString',sdir2_name);
                    cd ..
                    for k2=1:runs(k)
                        if strcmp(sdir2{indx(k2)}(end-3:end),'.nii')==1
                            func_files{k,k2} = [sdir2{indx(k2)}(1:end-4) erase(sdir2{indx(k2)}(end-3:end),'.nii')];  % for .nii extension, this will also work here: sdir2{indx(k2)}(1:end-4);
                        else
                            func_files{k,k2} = [sdir2{indx(k2)}(1:end-7) erase(sdir2{indx(k2)}(end-6:end),'.nii.gz')];  % for .nii.gz extension, this will also work here: sdir2{indx(k2)}(1:end-7);
                        end
                        func_home{k,k2}  = [sdir2_folder{indx(k2)} '/'];
                        if strcmp(func_home{k,k2},sub_dir{k})==0
                            fprintf('Subject-%d run-%d''s functional is in a subfolder within the main subject directory (/%s). Moving it (and associated files) to the latter location.\n',k,k2,erase(func_home{k,k2},sub_dir{k}))
                            sdir3 = cat(1,dir([func_home{k,k2} func_files{k,k2} '*']), dir([func_home{k,k2} sprintf('workspace_variables_run%d.mat',k2)]), dir([func_home{k,k2} sprintf('run%d_final_reg.mat',k2)]), dir([func_home{k,k2} 'cord_quadrants.mat']), dir([func_home{k,k2} 'cross_sectional_areas.mat']), dir([func_home{k,k2} 'motion_parameters.mat']), dir([func_home{k,k2} sprintf('tSNR_across_preproc_steps_run%d.mat',k2)]));
                            sdir3_folder = {sdir3.folder}; sdir3_name = {sdir3.name};
                            for k3=1:length(sdir3_folder)
                                movefile([sdir3_folder{k3} '/' sdir3_name{k3}],[sub_dir{k} sdir3_name{k3}],'f')
                            end
                            clear sdir3_folder sdir3 sdir3_name
                        end
                    end
                    clear sdir2 prompt sdir2_folder sdir2_name k2 k3 indx f
                end; clear k f
        end
        if exist('answerc','var')
            clear func_files
            for k=1:length(sub_dir)
                cd(sub_dir{k})
                sdir2 = cat(1,dir('**/*.nii'),dir('**/*.nii.gz'));
                sdir2_folder = {sdir2.folder}; sdir2_name = {sdir2.name}; sdir2 = {sdir2.name};
                for k2=1:length(sdir2_folder)
                    sdir2_name{k2} = [erase(sdir2_folder{k2},sub_dir{k}(1:end-1)) '/' sdir2{k2}];
                end
                prompt = sprintf('Select NIfTI functional image(s) (all runs) for subject (%d) of (%d)...\n%s',k,length(sub_dir),sub_dir{k});
                indx = listdlg('PromptString',{prompt,''},...
                    'SelectionMode','multiple','ListSize',[400 400],'ListString',sdir2_name);
                cd ..
                for k2=1:runs(k)
                    if strcmp(sdir2{indx(k2)}(end-3:end),'.nii')==1
                        func_files{k,k2} = [sdir2{indx(k2)}(1:end-4) erase(sdir2{indx(k2)}(end-3:end),'.nii')];  % for .nii extension, this will also work here: sdir2{indx(k2)}(1:end-4);
                    else
                        func_files{k,k2} = [sdir2{indx(k2)}(1:end-7) erase(sdir2{indx(k2)}(end-6:end),'.nii.gz')];  % for .nii.gz extension, this will also work here: sdir2{indx(k2)}(1:end-7);
                    end
                    func_home{k,k2}  = [sdir2_folder{indx(k2)} '/'];
                    if strcmp(func_home{k,k2},sub_dir{k})==0
                        fprintf('Subject-%d run-%d''s functional is in a subfolder within the main subject directory (/%s). Moving it (and associated files) to the latter location.\n',k,k2,erase(func_home{k,k2},sub_dir{k}))
                        sdir3 = cat(1,dir([func_home{k,k2} func_files{k,k2} '*']), dir([func_home{k,k2} sprintf('workspace_variables_run%d.mat',k2)]), dir([func_home{k,k2} sprintf('run%d_final_reg.mat',k2)]), dir([func_home{k,k2} 'cord_quadrants.mat']), dir([func_home{k,k2} 'cross_sectional_areas.mat']), dir([func_home{k,k2} 'motion_parameters.mat']), dir([func_home{k,k2} sprintf('tSNR_across_preproc_steps_run%d.mat',k2)]));
                        sdir3_folder = {sdir3.folder}; sdir3_name = {sdir3.name};
                        for k3=1:length(sdir3_folder)
                            movefile([sdir3_folder{k3} '/' sdir3_name{k3}],[sub_dir{k} sdir3_name{k3}],'f')
                        end
                        clear sdir3_folder sdir3 sdir3_name
                    end
                end
                clear sdir2 prompt sdir2_folder sdir2_name k2 k3 indx f
            end; clear k f
        end
        
        else
        name=[];
        for k=1:length(sub_dir)
            cd(sub_dir{k})
            sdir2 = dir(['**/' func_string]); sdir2_folder = {sdir2.folder}; sdir2_name = {sdir2.name};
            for k2=1:runs(k)
                if strcmp(sdir2_name{k2}(end-3:end),'.nii')==1
                    func_files{k,k2} = [sdir2_name{k2}(1:end-4) erase(sdir2_name{k2}(end-3:end),'.nii')];  % for .nii extension, this will also work here: sdir2{indx(k2)}(1:end-4);
                else
                    func_files{k,k2} = [sdir2_name{k2}(1:end-7) erase(sdir2_name{k2}(end-6:end),'.nii.gz')];  % for .nii.gz extension, this will also work here: sdir2{indx(k2)}(1:end-7);
                end
                func_home{k,k2}  = [sdir2_folder{k2} '/'];
                name = strcat(name, sprintf('\n%s%s / %s',subnames{k},string(erase(sdir2_folder{1},sub_dir{k}(1:end-1))),func_files{k,k2}));
                if strcmp(func_home{k,k2},sub_dir{k})==0
                    fprintf('Subject-%d run-%d''s functional is in a subfolder within the main subject directory (/%s). Moving it (and associated files) to the latter location.\n',k,k2,erase(func_home{k,k2},sub_dir{k}))
                    sdir3 = cat(1,dir([func_home{k,k2} func_files{k,k2} '*']), dir([func_home{k,k2} sprintf('workspace_variables_run%d.mat',k2)]), dir([func_home{k,k2} sprintf('run%d_final_reg.mat',k2)]), dir([func_home{k,k2} 'cord_quadrants.mat']), dir([func_home{k,k2} 'cross_sectional_areas.mat']), dir([func_home{k,k2} 'motion_parameters.mat']), dir([func_home{k,k2} sprintf('tSNR_across_preproc_steps_run%d.mat',k2)]));
                    sdir3_folder = {sdir3.folder}; sdir3_name = {sdir3.name};
                    for k3=1:length(sdir3_folder)
                        movefile([sdir3_folder{k3} '/' sdir3_name{k3}],[sub_dir{k} sdir3_name{k3}],'f')
                    end
                    clear sdir3_folder sdir3 sdir3_name
                end
            end
            clear sdir2_folder sdir2 sdir2_name k2
            cd ..
        end; clear k
        
        quest = sprintf('Are these the NIfTI functional files?:\n%s',name); clear name
        answer2 = questdlg(quest,'Functional files','Yes','No, open subject folders','Yes'); clear quest
        switch answer2
            case 'No, open subject folders'
                clear func_files
                for k=1:length(sub_dir)
                    cd(sub_dir{k})
                    sdir2 = cat(1,dir('**/*.nii'),dir('**/*.nii.gz'));
                    sdir2_folder = {sdir2.folder}; sdir2_name = {sdir2.name}; sdir2 = {sdir2.name};
                    for k2=1:length(sdir2_folder)
                        sdir2_name{k2} = [erase(sdir2_folder{k2},sub_dir{k}(1:end-1)) '/' sdir2{k2}];
                    end
                    prompt = sprintf('Select NIfTI functional image(s) (all runs) for subject (%d) of (%d)...\n%s',k,length(sub_dir),sub_dir{k});
                    indx = listdlg('PromptString',{prompt,''},...
                        'SelectionMode','multiple','ListSize',[400 400],'ListString',sdir2_name);
                    cd ..
                    for k2=1:runs(k)
                        if strcmp(sdir2{indx(k2)}(end-3:end),'.nii')==1
                            func_files{k,k2} = [sdir2{indx(k2)}(1:end-4) erase(sdir2{indx(k2)}(end-3:end),'.nii')];  % for .nii extension, this will also work here: sdir2{indx(k2)}(1:end-4);
                        else
                            func_files{k,k2} = [sdir2{indx(k2)}(1:end-7) erase(sdir2{indx(k2)}(end-6:end),'.nii.gz')];  % for .nii.gz extension, this will also work here: sdir2{indx(k2)}(1:end-7);
                        end
                        func_home{k,k2}  = [sdir2_folder{indx(k2)} '/'];
                        if strcmp(func_home{k,k2},sub_dir{k})==0
                            fprintf('Subject-%d run-%d''s functional is in a subfolder within the main subject directory (/%s). Moving it (and associated files) to the latter location.\n',k,k2,erase(func_home{k,k2},sub_dir{k}))
                            sdir3 = cat(1,dir([func_home{k,k2} func_files{k,k2} '*']), dir([func_home{k,k2} sprintf('workspace_variables_run%d.mat',k2)]), dir([func_home{k,k2} sprintf('run%d_final_reg.mat',k2)]), dir([func_home{k,k2} 'cord_quadrants.mat']), dir([func_home{k,k2} 'cross_sectional_areas.mat']), dir([func_home{k,k2} 'motion_parameters.mat']), dir([func_home{k,k2} sprintf('tSNR_across_preproc_steps_run%d.mat',k2)]));
                            sdir3_folder = {sdir3.folder}; sdir3_name = {sdir3.name};
                            for k3=1:length(sdir3_folder)
                                movefile([sdir3_folder{k3} '/' sdir3_name{k3}],[sub_dir{k} sdir3_name{k3}],'f')
                            end
                            clear sdir3_folder sdir3 sdir3_name
                        end
                    end
                    clear sdir2 prompt sdir2_folder sdir2_name k2 k3 indx f
                end; clear k
        end
        end
        
    case 'Open subject folders'
        for k=1:length(sub_dir)
            cd(sub_dir{k})
            sdir2 = cat(1,dir('**/*.nii'),dir('**/*.nii.gz'));
            sdir2_folder = {sdir2.folder}; sdir2_name = {sdir2.name}; sdir2 = {sdir2.name};
            for k2=1:length(sdir2_folder)
                sdir2_name{k2} = [erase(sdir2_folder{k2},sub_dir{k}(1:end-1)) '/' sdir2{k2}];
            end
            prompt = sprintf('Select NIfTI functional image(s) (all runs) for subject (%d) of (%d)...\n%s',k,length(sub_dir),sub_dir{k});
            indx = listdlg('PromptString',{prompt,''},...
                'SelectionMode','multiple','ListSize',[400 400],'ListString',sdir2_name);
            cd ..
            for k2=1:runs(k)
                if strcmp(sdir2{indx(k2)}(end-3:end),'.nii')==1
                func_files{k,k2} = [sdir2{indx(k2)}(1:end-4) erase(sdir2{indx(k2)}(end-3:end),'.nii')];  % for .nii extension, this will also work here: sdir2{indx(k2)}(1:end-4);
                else
                func_files{k,k2} = [sdir2{indx(k2)}(1:end-7) erase(sdir2{indx(k2)}(end-6:end),'.nii.gz')];  % for .nii.gz extension, this will also work here: sdir2{indx(k2)}(1:end-7);
                end
                func_home{k,k2}  = [sdir2_folder{indx(k2)} '/'];
                if strcmp(func_home{k,k2},sub_dir{k})==0
                    fprintf('Subject-%d run-%d''s functional is in a subfolder within the main subject directory (/%s). Moving it (and associated files) to the latter location.\n',k,k2,erase(func_home{k,k2},sub_dir{k}))
                    sdir3 = cat(1,dir([func_home{k,k2} func_files{k,k2} '*']), dir([func_home{k,k2} sprintf('workspace_variables_run%d.mat',k2)]), dir([func_home{k,k2} sprintf('run%d_final_reg.mat',k2)]), dir([func_home{k,k2} 'cord_quadrants.mat']), dir([func_home{k,k2} 'cross_sectional_areas.mat']), dir([func_home{k,k2} 'motion_parameters.mat']), dir([func_home{k,k2} sprintf('tSNR_across_preproc_steps_run%d.mat',k2)]));
                    sdir3_folder = {sdir3.folder}; sdir3_name = {sdir3.name};
                    for k3=1:length(sdir3_folder)
                        movefile([sdir3_folder{k3} '/' sdir3_name{k3}],[sub_dir{k} sdir3_name{k3}],'f')
                    end
                    clear sdir3_folder sdir3 sdir3_name
                end
            end
            clear sdir2 prompt sdir2_folder sdir2_name k2 k3 indx f
        end; clear k
end; clear answer1 f answer2 answerb answerc

end

if ~exist('slices','var')||~exist('voxel_size_inplane_func','var')
try
    F = load_untouch_nii([sub_dir{1} func_files{1,1} '.nii']);
catch
    gunzip([sub_dir{1} func_files{1,1} '.nii.gz'])
    F = load_untouch_nii([sub_dir{1} func_files{1,1} '.nii']);
    delete([sub_dir{1} func_files{1,1} '.nii'])
end
slices = size(F.img,3); % This toolbox assumes that all subjects have the same number of slices. If not, you would have to treat differences as separate cohorts and process separately (and aggregate the data yourself in the end)
voxel_size_inplane_func = min([F.hdr.dime.pixdim(2),F.hdr.dime.pixdim(3)]); % assumes that the slices are oriented axially and the voxel dimensions are same along the x and y directions
clear F
end

%% --- Log the choices made so far ---
if preproc==1
    try waitbar(8/8,wbar1,'Providing pre-processing choices soon...'); catch, end
elseif postproc==1
    try waitbar(8/8,wbar1,'Providing post-processing choices soon...'); catch, end
end

try close(wbar1); catch, end
try close(welcome_fig); catch, end
try close(neptune_fig); catch, end
clear wbar1 welcome_fig neptune_fig

answer_singleop = 'Run pipeline';
try
L=L+2; logg{L,1}=sprintf('--- Initial choices: ---'); fprintf(fileID,'%s\n',logg{L,1}); try set(S.tx,'string',logg); catch, end; clear clk
L=L+1; logg{L,1}=sprintf('Home directory = %s',home_dir); fprintf(fileID,'%s\n',logg{L,1}); try set(S.tx,'string',logg); catch, end; clear clk
L=L+1; logg{L,1}=sprintf('Siemens scanner = %d; Philips scanner = %d',(scanner==1),(scanner==2)); fprintf(fileID,'%s\n',logg{L,1}); try set(S.tx,'string',logg); catch, end; clear clk
L=L+1; logg{L,1}=sprintf('7T scanner = %d; 3T scanner = %d',scan7T,~scan7T); fprintf(fileID,'%s\n',logg{L,1}); try set(S.tx,'string',logg); catch, end; clear clk
L=L+1; logg{L,1}=sprintf('Cervical spine = %d; thoracic/lumbar spine = %d',C_spine,~C_spine); fprintf(fileID,'%s\n',logg{L,1}); try set(S.tx,'string',logg); catch, end; clear clk
L=L+1; logg{L,1}=sprintf('Resting-state fMRI = %d; task fMRI = %d',rest,task); fprintf(fileID,'%s\n',logg{L,1}); try set(S.tx,'string',logg); catch, end; clear clk
L=L+1; logg{L,1}=sprintf('Pre-process step-after-step = %d; pre-process subject-after-subject = %d',step_after_step,~step_after_step); fprintf(fileID,'%s\n',logg{L,1}); try set(S.tx,'string',logg); catch, end; clear clk
L=L+1; logg{L,1}=sprintf('Pre-processing = %d; Post-processing = %d',preproc,postproc); fprintf(fileID,'%s\n',logg{L,1}); try set(S.tx,'string',logg); catch, end; clear clk
L=L+1; logg{L,1}=sprintf('Number of subjects = %d',length(sub_dir)); fprintf(fileID,'%s\n',logg{L,1}); try set(S.tx,'string',logg); catch, end; clear clk
L=L+1; logg{L,1}=sprintf('Runs per subject = %s',num2str(runs)); fprintf(fileID,'%s\n',logg{L,1}); try set(S.tx,'string',logg); catch, end; clear clk
L=L+2; logg{L,1}=sprintf('Subject folders:'); fprintf(fileID,'%s\n',logg{L,1}); try set(S.tx,'string',logg); catch, end; clear clk
for k=1:length(sub_dir)
    L=L+1; logg{L,1}=sprintf('%s',sub_dir{k}); fprintf(fileID,'%s\n',logg{L,1}); try set(S.tx,'string',logg); catch, end
end; clear k
L=L+2; logg{L,1}=sprintf('Anatomical files:'); fprintf(fileID,'%s\n',logg{L,1}); try set(S.tx,'string',logg); catch, end; clear clk
for k=1:length(sub_dir)
    temp=strfind(sub_dir{k},'/'); temp=temp(end-1)+1; temp=sub_dir{k}(temp:end);
    L=L+1; logg{L,1}=sprintf('%s%s',temp,anat_files{k}); fprintf(fileID,'%s\n',logg{L,1}); try set(S.tx,'string',logg); catch, end; clear temp
end; clear k
L=L+2; logg{L,1}=sprintf('Functional files:'); fprintf(fileID,'%s\n',logg{L,1}); try set(S.tx,'string',logg); catch, end; clear clk
for k=1:length(sub_dir)
    for k2=1:runs(k)
        temp=strfind(sub_dir{k},'/'); temp=temp(end-1)+1; temp=sub_dir{k}(temp:end);
        L=L+1; logg{L,1}=sprintf('%s%s',temp,func_files{k,k2}); fprintf(fileID,'%s\n',logg{L,1}); try set(S.tx,'string',logg); catch, end; clear temp
    end
end; clear k
catch
end

tmp=clock; clk_settings = sprintf('%.4d-%.2d-%.2d_%.2d-%.2d',tmp(1),tmp(2),tmp(3),tmp(4),tmp(5)); clear tmp
save([home_dir sprintf('neptune_settings_%s',clk_settings)]);

%% --- Branch out operations ---
if preproc==1
    refresh_html_log
    time_preproc_total = tic;
    preproc_scfMRItb
    time_taken_preproc.preproc_total = toc(time_preproc_total);
end
if postproc==1
    refresh_html_log
    time_postproc_total = tic;
    postproc_scfMRItb
    time_taken_postproc.postproc_total = toc(time_postproc_total);
end
clear answer_singleop time_preproc_total time_postproc_total

%% if anat and func files were in subfolders inside the subject folders, then move them (and associated files) back there
for k=1:length(sub_dir)
    if strcmp(anat_home{k},sub_dir{k})==0
        fprintf('Moving anatomical files from subject-%d''s main directory to the subfolder in which the anatomical originally resided (/%s)\n',k,erase(anat_home{k},sub_dir{k}))
        sdir2 = dir([sub_dir{k} anat_files{k} '*']); sdir2_folder = {sdir2.folder}; sdir2_name = {sdir2.name};
        for k2=1:length(sdir2_folder)
            movefile([sub_dir{k} sdir2_name{k2}],[anat_home{k} sdir2_name{k2}],'f')
        end
    end
    clear sdir2 sdir2_folder sdir2_name
end

for k=1:length(sub_dir)
    for k2=1:runs(k)
        if strcmp(func_home{k,k2},sub_dir{k})==0
            fprintf('Moving functional files from subject-%d run-%d''s main directory to the subfolder in which the functional originally resided (/%s)\n',k,k2,erase(func_home{k,k2},sub_dir{k}))
            sdir3 = cat(1,dir([func_home{k,k2} func_files{k,k2} '*']), dir([func_home{k,k2} sprintf('workspace_variables_run%d.mat',k2)]), dir([func_home{k,k2} sprintf('run%d_final_reg.mat',k2)]), dir([func_home{k,k2} 'cord_quadrants.mat']), dir([func_home{k,k2} 'cross_sectional_areas.mat']), dir([func_home{k,k2} 'motion_parameters.mat']), dir([func_home{k,k2} sprintf('tSNR_across_preproc_steps_run%d.mat',k2)]));
            sdir3_folder = {sdir3.folder}; sdir3_name = {sdir3.name};
            for k3=1:length(sdir3_folder)
                movefile([sub_dir{k} sdir3_name{k3}],[func_home{k,k2} sdir3_name{k3}],'f')
            end
            clear sdir3_folder sdir3 sdir3_name
        end
    end
end

%% final logging
try
L=L+2; logg{L,1}=sprintf('--- All Completed ---'); fprintf(fileID,'%s\n',logg{L,1}); try set(S.tx,'string',logg); catch, end; clear clk
tmp=clock; clk = sprintf('%.4d-%.2d-%.2d_%.2d-%.2d',tmp(1),tmp(2),tmp(3),tmp(4),tmp(5)); L=L+1; logg{L,1}=sprintf('The final workspace has been saved to:\n%s',[home_dir sprintf('workspace_final_%s.mat',clk)]); fprintf(fileID,'%s\n',logg{L,1}); try set(S.tx,'string',logg); catch, end; clear clk tmp
clk=clock; L=L+1; logg{L,1}=sprintf('%d/%s/%d %dh:%dm:%ds - everything COMPLETED as promised.',clk(3),month{clk(2)},clk(1),clk(4),clk(5),round(clk(6))); fprintf(fileID,'%s\n',logg{L,1}); try set(S.tx,'string',logg); catch, end; clear clk
L=L+2; logg{L,1}=sprintf('------- ------- -------'); fprintf(fileID,'%s\n',logg{L,1}); try set(S.tx,'string',logg); catch, end; clear clk
catch
end

clear A F
tmp=clock; clk = sprintf('%.4d-%.2d-%.2d_%.2d-%.2d',tmp(1),tmp(2),tmp(3),tmp(4),tmp(5)); clear tmp
save([home_dir sprintf('workspace_final_%s',clk)])
uiwait(msgbox(sprintf('The final workspace has been saved to\n%s\n\nExecution log with timestamps has been saved to\n%s\n',[home_dir sprintf('workspace_final_%s.mat',clk)],[home_dir sprintf('log_%s.txt',clk)]),'Choices saved','custom',msgicon)); clear clk tmp

if exist('tc00','var')
    try
    L=L+1; logg{L,1}=sprintf('Total time taken: %dh:%dm:%ds',floor(toc(tc00)/3600),floor(mod(toc(tc00),3600)/60),floor(mod(toc(tc00),60))); fprintf(fileID,'%s\n',logg{L,1}); try set(S.tx,'string',logg); catch, end; clear clk
    catch
    end
end

fclose(fileID);
clear fileID tmp clk row L tc

%% generate HTMLs

% within each subject's QC folder
for k=1:length(sub_dir)
    fprintf('Generate HTML files within each subject''s QC folder: subject (%d) of (%d)\n',k,length(sub_dir))
    html_dir = [sub_dir{k} 'QC' '/'];
    html_savefile = [html_dir 'report_neptune_sub.html'];
    refresh_html_sub
    movefile(html_savefile,[html_dir sprintf('report_neptune_%s.html',subnames{k})])
    clear html_dir html_savefile tmp_suffix
end

% master HTML
refresh_html_all
delete([home_dir 'report_neptune.html'])
movefile([home_dir 'report_neptune_all.html'],[home_dir 'report_neptune.html'],'f')

%% All DONE!
neptune_fig_alldone = figure('Name','All DONE!','NumberTitle','off','units','normalized','outerposition',[1/12 21/48 1/6 7/24]); % [position_along_x_from_bottomleft position_along_y_from_bottomleft size_x size_y] in normalized units (0,1)
subplot = @(m,n,p) subtightplot(m, n, p, [0.001 0.005], [0.01 0.001], [0.01 0.001]);
figure(neptune_fig_alldone), subplot(1,1,1), imshow('status_images/neptune_v1.0_all_done.jpg','InitialMagnification','fit')

fprintf('\n-------\n\n\n\n\nDear User,\n\nEverything has been COMPLETED as promised.\n\nRegards,\nSpinal cord fMRI toolbox (NEPTUNE)\n\n\n\n\n-------\n')
quest = ['\color[rgb]{0.65,0.11,0.19}' sprintf('Dear User,\n\nEverything COMPLETED as promised.\n\nRegards,\nSpinal cord fMRI toolbox (NEPTUNE)\n')]; %#ok<*NASGU>
opts.Interpreter = 'tex'; opts.WindowStyle = 'non-modal';
uiwait(msgbox(quest,'All done!','custom',msgicon,[1,1,1],opts)); clear quest opts

try close(neptune_fig_alldone), catch, end
web([home_dir 'report_neptune.html'],'-browser')

%%
```
