## Supplementary material for "Neptune: a toolbox for spinal cord functional MRI data processing and quality assurance": Neptune software: merge_mat_files.m

```
function merge_mat_files(C,A,B,varargin)

% Merges two .mat files A and B (i.e. its variables) into a single .mat file C
%
% If ignore=0 (4th argument)
%   In case A and B have common variable names, note that the common
%   variables in B will be renamed by appending a suffix
%   '_renamed_by_mergestructs' to the common variables, before saving as C
% 
% If ignore=1 (4th argument)
%   In case A and B have common variable names, the common variables in B will be ignored
%
% For completeness, A, B and C must preferrably contain the complete path

if isempty(varargin)
    ignore = 0;
else
    ignore = varargin{1};
end

x = load(A);
y = load(B);

z = mergestructs(x,y,ignore);

save(C,'-struct','z')

end
```
