## Supplementary material for "Neptune: a toolbox for spinal cord functional MRI data processing and quality assurance": Neptune software: mergestructs.m

```
function C = mergestructs(A,B,varargin)

% Merges two structures A and B into a single structure C
%
% If ignore=0 (3rd argument)
%   In case A and B have common variable names within their structure, 
%   note that the common variables in B will be renamed by appending a 
%   suffix '_renamed_by_mergestructs' to the common variables
% 
% If ignore=1 (3rd argument)
%   In case A and B have common variable names within their structure, 
%   note that the common variables in B will be ignored

if isempty(varargin)
    ignore = 0;
else
    ignore = varargin{1};
end

f1 = fieldnames(A);
f2 = fieldnames(B);
f = intersect(f1,f2); % identify variable names that are common between A and B

for k=1:length(f)
    if ignore==0
    eval(sprintf('B.%s_renamed_by_mergestructs = B.%s;',f{k},f{k})) % rename duplicates in B by appending '_renamed_by_mergestructs' to the end of the variable name
    end
    B = rmfield(B,f{k});
end

C = cell2struct([struct2cell(A);struct2cell(B)],[fieldnames(A);fieldnames(B)]); % merge structures

end
```
