## Supplementary material for "Neptune: a toolbox for spinal cord functional MRI data processing and quality assurance": Neptune software: postproc_choices.m

```
% Post-processing choices

%% --- choose post-processing steps ---
if ~exist('choices_postproc','var')

fig = uifigure('Position',[680 370 530 230],'Color',[0.95,0.875,0.8]); drawnow; fig.Visible='off'; fig.Visible='on';
pnl = uipanel(fig,'Position',[30 60 470 150],'BackgroundColor',[0.95,0.9,0.85],'BorderType','none');
pnl.Title = sprintf('Choose your post-processing steps and click DONE.\n''Help'' is available if required.\n');
pnl.TitlePosition = 'centertop'; pnl.ForegroundColor = [0.65,0.11,0.19];  % [0,0,0.8];
bg = uibuttongroup(fig,'Position',[30 30 470 30],'BackgroundColor',[0.95,0.875,0.8],'BorderType','none');

selection=''; flag=0;
choices_postproc=[1;1;1;0];
while strcmp(selection,'DONE')~=1
    [selection,choices_postproc] = postprocdialog(pnl,bg,choices_postproc,flag);
    if strcmp(selection,'DONE')==1
%         closereq()
        break
    end
    if strcmp(selection,'Help')==1
        fprintf('neptune_manual.pdf\n')
        open('neptune_manual.pdf')
        flag=1;
    end
end
close(fig)
clear fig bg pnl flag

end

%% --- user-defined settings not appearing in the GUI ---

if ~(exist('use_deconv_data_for_postproc','var'))
    % 1: try deconvolved data for post-processing; 0: try bandpass filtered data (before deconv) for post-processing
    use_deconv_data_for_postproc = 1;
end

if ~(exist('dont_do_pcor_betweenslice','var'))
    % 1: do not do partial correlation - between slice FC at any cost (master override). 0: ignore this flag and follow user-defined choices
    dont_do_pcor_betweenslice = 0;
end

%% --- GUI choices before proceeding ---
if ~exist('WM_flag','var') % quads
if (choices_postproc(1)==1)||(choices_postproc(2)==1)||(choices_postproc(3)==1)||(choices_postproc(4)==1)
    quest = sprintf('Do you wish to perform post-processing steps in the white matter as well?');
    answer_WM_flag = questdlg(quest,'White-matter flag','Yes','No','Yes'); clear quest
    if strcmp(answer_WM_flag,'Yes')
        WM_flag = 1;
    elseif strcmp(answer_WM_flag,'No')
        WM_flag = 0;
    end
    clear answer_WM_flag
end
end

if ~exist('assign_value','var') % quads
if choices_postproc(1)==1
    quest = sprintf('After computing SFC between all pairs of voxels between quadrants (and/or fALFF in all voxels), do you wish to:\n\n(1) assign the 95th percentile value as the final value\nOR\n(2) assign the 50th percentile value as the final value\nOR\n(3) instead obtain the mean time series for each quadrant first and compute SFC/fALFF using that (not recommended)');
    answer_assign_value = questdlg(quest,'Post-processing choices','(1) 95th percentile','(2) 50th percentile','(3) mean time series','(1) 95th percentile'); clear quest
    if strcmp(answer_assign_value,'(1) 95th percentile')
        assign_value = 1;
    elseif strcmp(answer_assign_value,'(2) 50th percentile')
        assign_value = 2;
    elseif strcmp(answer_assign_value,'(3) mean time series')
        assign_value = 3;
    end
    clear answer_assign_value
end
end

if ~exist('SFC_type','var') % SFC
if (choices_postproc(2)==1)||(choices_postproc(4)==1)
    quest = sprintf('Static functional connectivity: compute Pearson''s correlation, partial correlation or both?');
    answer_SFC_type = questdlg(quest,'SFC method','Pearson''s correlation','Partial correlation','Both','Pearson''s correlation'); clear quest
    if strcmp(answer_SFC_type,'Pearson''s correlation')
        SFC_type = 1;
    elseif strcmp(answer_SFC_type,'Partial correlation')
        SFC_type = 2;
    elseif strcmp(answer_SFC_type,'Both')
        SFC_type = 3;
    end
    clear answer_SFC_type

    quest = sprintf('Static functional connectivity:\nestimate (i) within-slice SFC, (ii) within- AND between-slices SFC, OR (iii) both?');
    answer_SFC_slices = questdlg(quest,'SFC slices','Within-slice SFC','Between-slices SFC','Both','Both'); clear quest
    if strcmp(answer_SFC_slices,'Within-slice')
        SFC_withinslice_flag = 1; SFC_betweenslice_flag = 0;
    elseif strcmp(answer_SFC_slices,'Between-slices')
        SFC_withinslice_flag = 0; SFC_betweenslice_flag = 1;
    elseif strcmp(answer_SFC_slices,'Both')
        SFC_withinslice_flag = 1; SFC_betweenslice_flag = 1;
    end
    clear answer_SFC_slices
end
end

if choices_postproc(4)==1
if ~exist('ttest_type','var')||~exist('indx_group1','var')||~exist('indx_group2','var')
    quest = sprintf('Stats - choose your test:\n\n(1) one-sample t-test (all subjects vs. null)\n(2) paired t-test (same subjects across two scans)\n(3) two-sample t-test (between two independent cohorts)');
    answer_stats = questdlg(quest,'Statistical test','(1) one-sample t-test','(2) paired t-test','(3) two-sample t-test','(1) one-sample t-test'); clear quest
    switch answer_stats
        
        case '(1) one-sample t-test'
            ttest_type = 1;
            quest = sprintf('For stats: please select subjects for the one-sample t-test');
            answer1 = questdlg(quest,'One-sample t-test','Sure, take me there','Not now','Sure, take me there'); clear quest
            switch answer1
                case 'Sure, take me there'
                    cd(home_dir)
                    sdir2 = dir; sdir2(find([sdir2.isdir]==0))=[]; sdir2=sdir2(3:end); sdir2={sdir2.name};
                    f2=0;
                    for u=1:length(sdir2)
                        for u2=1:length(subnames)
                            if strcmp(sdir2{u},subnames{u2})
                                f2=f2+1;
                                sdir3{f2} = sdir2{u};
                            end
                        end
                    end; clear f2 u u2
                    clear sdir2; sdir2=sdir3; clear sdir3
                    prompt = sprintf('Select subjects...\n(hold Ctrl or Command button and click to choose multiple subjects)');
                    indx_group1 = listdlg('PromptString',{prompt,''},...
                        'SelectionMode','multiple','ListSize',[400 400],'ListString',sdir2);
                    indx_group2 = [];
                    clear sdir2 prompt
                case 'Not now'
                    tmp=clock; clk = sprintf('%.4d-%.2d-%.2d_%.2d-%.2d',tmp(1),tmp(2),tmp(3),tmp(4),tmp(5));
                    save([home_dir sprintf('workspace_cancel_%s',clk)])
                    msgbox(sprintf('Your choices so far have been saved to\n%s\nTerminating execution...',sprintf('workspace_cancel_%s.mat',clk)),'Choices saved','custom',msgicon); clear clk tmp
                    return
            end; clear answer1
            
        case '(2) paired t-test'
            ttest_type = 2;
            if min(runs)==2 && max(runs)==2 % each subject has 2 runs for paired test
                quest = sprintf('For the paired t-test, do you want to use runs 1 and 2 in each subject as the groups for comparison?');
                answer_paired_test = questdlg(quest,'Paired t-test','Yes','No','Yes'); clear quest
                switch answer_paired_test
                    case 'Yes'
                        quest = sprintf('For stats: please select subjects for the paired t-test');
                        answer1 = questdlg(quest,'Paired t-test','Sure, take me there','Not now','Sure, take me there'); clear quest
                        switch answer1
                            case 'Sure, take me there'
                                cd(home_dir)
                                sdir2 = dir; sdir2(find([sdir2.isdir]==0))=[]; sdir2=sdir2(3:end); sdir2={sdir2.name};
                                f2=0;
                                for u=1:length(sdir2)
                                    for u2=1:length(subnames)
                                        if strcmp(sdir2{u},subnames{u2})
                                            f2=f2+1;
                                            sdir3{f2} = sdir2{u};
                                        end
                                    end
                                end; clear f2 u u2
                                clear sdir2; sdir2=sdir3; clear sdir3
                                prompt = sprintf('Select subjects...\n(hold Ctrl or Command button and click to choose multiple subjects)');
                                indx_group1 = listdlg('PromptString',{prompt,''},...
                                    'SelectionMode','multiple','ListSize',[400 400],'ListString',sdir2);
                                indx_group2 = indx_group1;
                                clear sdir2 prompt
                            case 'Not now'
                                tmp=clock; clk = sprintf('%.4d-%.2d-%.2d_%.2d-%.2d',tmp(1),tmp(2),tmp(3),tmp(4),tmp(5));
                                save([home_dir sprintf('workspace_cancel_%s',clk)])
                                msgbox(sprintf('Your choices so far have been saved to\n%s\nTerminating execution...',sprintf('workspace_cancel_%s.mat',clk)),'Choices saved','custom',msgicon); clear clk tmp
                                return
                        end; clear answer1
                        
                    case 'No'
                        quest = sprintf('For stats: please select subjects in GROUP-1 for the paired t-test');
                        answer1 = questdlg(quest,'Stats','Sure, take me there','Not now','Sure, take me there'); clear quest
                        switch answer1
                            case 'Sure, take me there'
                                cd(home_dir)
                                sdir2 = dir; sdir2(find([sdir2.isdir]==0))=[]; sdir2=sdir2(3:end); sdir2={sdir2.name};
                                f2=0;
                                for u=1:length(sdir2)
                                    for u2=1:length(subnames)
                                        if strcmp(sdir2{u},subnames{u2})
                                            f2=f2+1;
                                            sdir3{f2} = sdir2{u};
                                        end
                                    end
                                end; clear f2 u u2
                                clear sdir2; sdir2=sdir3; clear sdir3
                                prompt = sprintf('Select subjects in GROUP-1...\n(hold Ctrl or Command button and click to choose multiple subjects)');
                                indx_group1 = listdlg('PromptString',{prompt,''},...
                                    'SelectionMode','multiple','ListSize',[400 400],'ListString',sdir2);
                                clear sdir2 prompt
                            case 'Not now'
                                tmp=clock; clk = sprintf('%.4d-%.2d-%.2d_%.2d-%.2d',tmp(1),tmp(2),tmp(3),tmp(4),tmp(5));
                                save([home_dir sprintf('workspace_cancel_%s',clk)])
                                msgbox(sprintf('Your choices so far have been saved to\n%s\nTerminating execution...',sprintf('workspace_cancel_%s.mat',clk)),'Choices saved','custom',msgicon); clear clk tmp
                                return
                        end; clear answer1
                        
                        quest = sprintf('For stats: next, please select subjects in GROUP-2 for the paired t-test (must be the same number of subjects as GROUP-1)');
                        answer1 = questdlg(quest,'Stats','Sure, take me there','Not now','Sure, take me there'); clear quest
                        switch answer1
                            case 'Sure, take me there'
                                cd(home_dir)
                                sdir2 = dir; sdir2(find([sdir2.isdir]==0))=[]; sdir2=sdir2(3:end); sdir2={sdir2.name};
                                f2=0;
                                for u=1:length(sdir2)
                                    for u2=1:length(subnames)
                                        if strcmp(sdir2{u},subnames{u2})
                                            f2=f2+1;
                                            sdir3{f2} = sdir2{u};
                                        end
                                    end
                                end; clear f2 u u2
                                clear sdir2; sdir2=sdir3; clear sdir3
                                prompt = sprintf('Select subjects in GROUP-2...\n(hold Ctrl or Command button and click to choose multiple subjects)');
                                indx_group2 = listdlg('PromptString',{prompt,''},...
                                    'SelectionMode','multiple','ListSize',[400 400],'ListString',sdir2);
                                clear sdir2 prompt
                            case 'Not now'
                                tmp=clock; clk = sprintf('%.4d-%.2d-%.2d_%.2d-%.2d',tmp(1),tmp(2),tmp(3),tmp(4),tmp(5));
                                save([home_dir sprintf('workspace_cancel_%s',clk)])
                                msgbox(sprintf('Your choices so far have been saved to\n%s\nTerminating execution...',sprintf('workspace_cancel_%s.mat',clk)),'Choices saved','custom',msgicon); clear clk tmp
                                return
                        end; clear answer1
                        
                        if length(indx_group1)~=length(indx_group2)
                            clear indx_group1 indx_group1
                            quest = sprintf('Group-1 and Group-2 must have the same number of subjects for a paired t-test. Please choose the subjects correctly again.');
                            questdlg(quest,'Stats','Sure, take me there','Sure, take me there'); clear quest
                            cd(home_dir)
                            sdir2 = dir; sdir2(find([sdir2.isdir]==0))=[]; sdir2=sdir2(3:end); sdir2={sdir2.name};
                            f2=0;
                            for u=1:length(sdir2)
                                for u2=1:length(subnames)
                                    if strcmp(sdir2{u},subnames{u2})
                                        f2=f2+1;
                                        sdir3{f2} = sdir2{u};
                                    end
                                end
                            end; clear f2 u u2
                            clear sdir2; sdir2=sdir3; clear sdir3
                            prompt = sprintf('Select subjects in GROUP-1...\n(hold Ctrl or Command button and click to choose multiple subjects)');
                            indx_group1 = listdlg('PromptString',{prompt,''},...
                                'SelectionMode','multiple','ListSize',[400 400],'ListString',sdir2);
                            clear sdir2 prompt
                            
                            cd(home_dir)
                            sdir2 = dir; sdir2(find([sdir2.isdir]==0))=[]; sdir2=sdir2(3:end); sdir2={sdir2.name};
                            f2=0;
                            for u=1:length(sdir2)
                                for u2=1:length(subnames)
                                    if strcmp(sdir2{u},subnames{u2})
                                        f2=f2+1;
                                        sdir3{f2} = sdir2{u};
                                    end
                                end
                            end; clear f2 u u2
                            clear sdir2; sdir2=sdir3; clear sdir3
                            prompt = sprintf('Select subjects in GROUP-2...\n(hold Ctrl or Command button and click to choose multiple subjects)');
                            indx_group2 = listdlg('PromptString',{prompt,''},...
                                'SelectionMode','multiple','ListSize',[400 400],'ListString',sdir2);
                            clear sdir2 prompt
                            if length(indx_group1)~=length(indx_group2)
                                msgbox(sprintf('Error: again, the two groups do not have the same number of subjects for the paired t-test. Aborting.\n%s\n',overlap_str),'Abort','error');
                                return;
                            end
                        end
                        
                        indx_overlap = sort(unique(intersect(indx_group1,indx_group2)));
                        if ~isempty(indx_overlap)
                            overlap_str = '';
                            for w=1:length(indx_overlap)
                                tmp_suffix = sub_dir{indx_overlap(w)}(max(strfind(sub_dir{indx_overlap(w)}(1:end-1),'/'))+1:end-1);
                                if isempty(tmp_suffix)
                                    tmp_suffix = sub_dir{indx_overlap(w)}(max(strfind(sub_dir{indx_overlap(w)}(1:end-1),'\'))+1:end-1);
                                end
                                overlap_str = strcat(overlap_str,sprintf('\n%s',tmp_suffix));
                                clear tmp_suffix
                            end
                            answer2 = questdlg(sprintf('Error: some of the subjects (mentioned below) are in both the groups. Try choosing the subjects again without overlapping.\n%s\n',overlap_str),'Stats','Ok, take me there','Ignore this message and proceed as is','Ok, take me there');
                            switch answer2
                                case 'Ok, take me there'
                                    clear indx_group1 indx_group2 ttest_type indx_overlap overlap_str w
                                    cd(home_dir)
                                    sdir2 = dir; sdir2(find([sdir2.isdir]==0))=[]; sdir2=sdir2(3:end); sdir2={sdir2.name};
                                    f2=0;
                                    for u=1:length(sdir2)
                                        for u2=1:length(subnames)
                                            if strcmp(sdir2{u},subnames{u2})
                                                f2=f2+1;
                                                sdir3{f2} = sdir2{u};
                                            end
                                        end
                                    end; clear f2 u u2
                                    clear sdir2; sdir2=sdir3; clear sdir3
                                    prompt = sprintf('Select subjects for GROUP-1...\n(hold Ctrl or Command button and click to choose multiple subjects)');
                                    indx_group1 = listdlg('PromptString',{prompt,''},...
                                        'SelectionMode','multiple','ListSize',[400 400],'ListString',sdir2);
                                    clear sdir2 prompt
                                    
                                    cd(home_dir)
                                    sdir2 = dir; sdir2(find([sdir2.isdir]==0))=[]; sdir2=sdir2(3:end); sdir2={sdir2.name};
                                    f2=0;
                                    for u=1:length(sdir2)
                                        for u2=1:length(subnames)
                                            if strcmp(sdir2{u},subnames{u2})
                                                f2=f2+1;
                                                sdir3{f2} = sdir2{u};
                                            end
                                        end
                                    end; clear f2 u u2
                                    clear sdir2; sdir2=sdir3; clear sdir3
                                    prompt = sprintf('Select subjects for GROUP-2...\n(hold Ctrl or Command button and click to choose multiple subjects)');
                                    indx_group2 = listdlg('PromptString',{prompt,''},...
                                        'SelectionMode','multiple','ListSize',[400 400],'ListString',sdir2);
                                    clear sdir2 prompt
                                    
                                    indx_overlap = sort(unique(intersect(indx_group1,indx_group2)));
                                    if ~isempty(indx_overlap)
                                        overlap_str = '';
                                        for w=1:length(indx_overlap)
                                            tmp_suffix = sub_dir{indx_overlap(w)}(max(strfind(sub_dir{indx_overlap(w)}(1:end-1),'/'))+1:end-1);
                                            if isempty(tmp_suffix)
                                                tmp_suffix = sub_dir{indx_overlap(w)}(max(strfind(sub_dir{indx_overlap(w)}(1:end-1),'\'))+1:end-1);
                                            end
                                            overlap_str = strcat(overlap_str,sprintf('\n%s',tmp_suffix));
                                            clear tmp_suffix
                                        end
                                        msgbox(sprintf('Error: again, some of the subjects (mentioned below) are in both the groups. Aborting.\n%s\n',overlap_str),'Abort','error');
                                        return;
                                    end
                                    clear indx_overlap overlap_str w
                            end; clear answer2
                        end
                end
            else
                quest = sprintf('For stats: please select subjects in GROUP-1 for the paired t-test');
                answer1 = questdlg(quest,'Stats','Sure, take me there','Not now','Sure, take me there'); clear quest
                switch answer1
                    case 'Sure, take me there'
                        cd(home_dir)
                        sdir2 = dir; sdir2(find([sdir2.isdir]==0))=[]; sdir2=sdir2(3:end); sdir2={sdir2.name};
                        f2=0;
                        for u=1:length(sdir2)
                            for u2=1:length(subnames)
                                if strcmp(sdir2{u},subnames{u2})
                                    f2=f2+1;
                                    sdir3{f2} = sdir2{u};
                                end
                            end
                        end; clear f2 u u2
                        clear sdir2; sdir2=sdir3; clear sdir3
                        prompt = sprintf('Select subjects in GROUP-1...\n(hold Ctrl or Command button and click to choose multiple subjects)');
                        indx_group1 = listdlg('PromptString',{prompt,''},...
                            'SelectionMode','multiple','ListSize',[400 400],'ListString',sdir2);
                        clear sdir2 prompt
                    case 'Not now'
                        tmp=clock; clk = sprintf('%.4d-%.2d-%.2d_%.2d-%.2d',tmp(1),tmp(2),tmp(3),tmp(4),tmp(5));
                        save([home_dir sprintf('workspace_cancel_%s',clk)])
                        msgbox(sprintf('Your choices so far have been saved to\n%s\nTerminating execution...',sprintf('workspace_cancel_%s.mat',clk)),'Choices saved','custom',msgicon); clear clk tmp
                        return
                end; clear answer1
                
                quest = sprintf('For stats: next, please select subjects in GROUP-2 for the paired t-test (must be the same number of subjects as GROUP-1)');
                answer1 = questdlg(quest,'Stats','Sure, take me there','Not now','Sure, take me there'); clear quest
                switch answer1
                    case 'Sure, take me there'
                        cd(home_dir)
                        sdir2 = dir; sdir2(find([sdir2.isdir]==0))=[]; sdir2=sdir2(3:end); sdir2={sdir2.name};
                        f2=0;
                        for u=1:length(sdir2)
                            for u2=1:length(subnames)
                                if strcmp(sdir2{u},subnames{u2})
                                    f2=f2+1;
                                    sdir3{f2} = sdir2{u};
                                end
                            end
                        end; clear f2 u u2
                        clear sdir2; sdir2=sdir3; clear sdir3
                        prompt = sprintf('Select subjects in GROUP-2...\n(hold Ctrl or Command button and click to choose multiple subjects)');
                        indx_group2 = listdlg('PromptString',{prompt,''},...
                            'SelectionMode','multiple','ListSize',[400 400],'ListString',sdir2);
                        clear sdir2 prompt
                    case 'Not now'
                        tmp=clock; clk = sprintf('%.4d-%.2d-%.2d_%.2d-%.2d',tmp(1),tmp(2),tmp(3),tmp(4),tmp(5));
                        save([home_dir sprintf('workspace_cancel_%s',clk)])
                        msgbox(sprintf('Your choices so far have been saved to\n%s\nTerminating execution...',sprintf('workspace_cancel_%s.mat',clk)),'Choices saved','custom',msgicon); clear clk tmp
                        return
                end; clear answer1
                
                if length(indx_group1)~=length(indx_group2)
                    clear indx_group1 indx_group1
                    quest = sprintf('Group-1 and Group-2 must have the same number of subjects for a paired t-test. Please choose the subjects correctly again.');
                    questdlg(quest,'Stats','Sure, take me there','Sure, take me there'); clear quest
                    cd(home_dir)
                    sdir2 = dir; sdir2(find([sdir2.isdir]==0))=[]; sdir2=sdir2(3:end); sdir2={sdir2.name};
                    f2=0;
                    for u=1:length(sdir2)
                        for u2=1:length(subnames)
                            if strcmp(sdir2{u},subnames{u2})
                                f2=f2+1;
                                sdir3{f2} = sdir2{u};
                            end
                        end
                    end; clear f2 u u2
                    clear sdir2; sdir2=sdir3; clear sdir3
                    prompt = sprintf('Select subjects in GROUP-1...\n(hold Ctrl or Command button and click to choose multiple subjects)');
                    indx_group1 = listdlg('PromptString',{prompt,''},...
                        'SelectionMode','multiple','ListSize',[400 400],'ListString',sdir2);
                    clear sdir2 prompt
                    
                    cd(home_dir)
                    sdir2 = dir; sdir2(find([sdir2.isdir]==0))=[]; sdir2=sdir2(3:end); sdir2={sdir2.name};
                    f2=0;
                    for u=1:length(sdir2)
                        for u2=1:length(subnames)
                            if strcmp(sdir2{u},subnames{u2})
                                f2=f2+1;
                                sdir3{f2} = sdir2{u};
                            end
                        end
                    end; clear f2 u u2
                    clear sdir2; sdir2=sdir3; clear sdir3
                    prompt = sprintf('Select subjects in GROUP-2...\n(hold Ctrl or Command button and click to choose multiple subjects)');
                    indx_group2 = listdlg('PromptString',{prompt,''},...
                        'SelectionMode','multiple','ListSize',[400 400],'ListString',sdir2);
                    clear sdir2 prompt
                    if length(indx_group1)~=length(indx_group2)
                        msgbox(sprintf('Error: again, the two groups do not have the same number of subjects for the paired t-test. Aborting.\n%s\n',overlap_str),'Abort','error');
                        return;
                    end
                end
                
                indx_overlap = sort(unique(intersect(indx_group1,indx_group2)));
                if ~isempty(indx_overlap)
                    overlap_str = '';
                    for w=1:length(indx_overlap)
                        tmp_suffix = sub_dir{indx_overlap(w)}(max(strfind(sub_dir{indx_overlap(w)}(1:end-1),'/'))+1:end-1);
                        if isempty(tmp_suffix)
                            tmp_suffix = sub_dir{indx_overlap(w)}(max(strfind(sub_dir{indx_overlap(w)}(1:end-1),'\'))+1:end-1);
                        end
                        overlap_str = strcat(overlap_str,sprintf('\n%s',tmp_suffix));
                        clear tmp_suffix
                    end
                    answer2 = questdlg(sprintf('Error: some of the subjects (mentioned below) are in both the groups. Try choosing the subjects again without overlapping.\n%s\n',overlap_str),'Stats','Ok, take me there','Ignore this message and proceed as is','Ok, take me there');
                    switch answer2
                        case 'Ok, take me there'
                            clear indx_group1 indx_group2 ttest_type indx_overlap overlap_str w
                            cd(home_dir)
                            sdir2 = dir; sdir2(find([sdir2.isdir]==0))=[]; sdir2=sdir2(3:end); sdir2={sdir2.name};
                            f2=0;
                            for u=1:length(sdir2)
                                for u2=1:length(subnames)
                                    if strcmp(sdir2{u},subnames{u2})
                                        f2=f2+1;
                                        sdir3{f2} = sdir2{u};
                                    end
                                end
                            end; clear f2 u u2
                            clear sdir2; sdir2=sdir3; clear sdir3
                            prompt = sprintf('Select subjects for GROUP-1...\n(hold Ctrl or Command button and click to choose multiple subjects)');
                            indx_group1 = listdlg('PromptString',{prompt,''},...
                                'SelectionMode','multiple','ListSize',[400 400],'ListString',sdir2);
                            clear sdir2 prompt
                            
                            cd(home_dir)
                            sdir2 = dir; sdir2(find([sdir2.isdir]==0))=[]; sdir2=sdir2(3:end); sdir2={sdir2.name};
                            f2=0;
                            for u=1:length(sdir2)
                                for u2=1:length(subnames)
                                    if strcmp(sdir2{u},subnames{u2})
                                        f2=f2+1;
                                        sdir3{f2} = sdir2{u};
                                    end
                                end
                            end; clear f2 u u2
                            clear sdir2; sdir2=sdir3; clear sdir3
                            prompt = sprintf('Select subjects for GROUP-2...\n(hold Ctrl or Command button and click to choose multiple subjects)');
                            indx_group2 = listdlg('PromptString',{prompt,''},...
                                'SelectionMode','multiple','ListSize',[400 400],'ListString',sdir2);
                            clear sdir2 prompt
                            
                            indx_overlap = sort(unique(intersect(indx_group1,indx_group2)));
                            if ~isempty(indx_overlap)
                                overlap_str = '';
                                for w=1:length(indx_overlap)
                                    tmp_suffix = sub_dir{indx_overlap(w)}(max(strfind(sub_dir{indx_overlap(w)}(1:end-1),'/'))+1:end-1);
                                    if isempty(tmp_suffix)
                                        tmp_suffix = sub_dir{indx_overlap(w)}(max(strfind(sub_dir{indx_overlap(w)}(1:end-1),'\'))+1:end-1);
                                    end
                                    overlap_str = strcat(overlap_str,sprintf('\n%s',tmp_suffix));
                                    clear tmp_suffix
                                end
                                msgbox(sprintf('Error: again, some of the subjects (mentioned below) are in both the groups. Aborting.\n%s\n',overlap_str),'Abort','error');
                                return;
                            end
                            clear indx_overlap overlap_str w
                    end; clear answer2
                end
            end
            
        case '(3) two-sample t-test'
            ttest_type = 3;
            quest = sprintf('For stats: please select subjects in GROUP-1 for the two-sample t-test');
            answer1 = questdlg(quest,'Stats','Sure, take me there','Not now','Sure, take me there'); clear quest
            switch answer1
                case 'Sure, take me there'
                    cd(home_dir)
                    sdir2 = dir; sdir2(find([sdir2.isdir]==0))=[]; sdir2=sdir2(3:end); sdir2={sdir2.name};
                    f2=0;
                    for u=1:length(sdir2)
                        for u2=1:length(subnames)
                            if strcmp(sdir2{u},subnames{u2})
                                f2=f2+1;
                                sdir3{f2} = sdir2{u};
                            end
                        end
                    end; clear f2 u u2
                    clear sdir2; sdir2=sdir3; clear sdir3
                    prompt = sprintf('Select subjects in GROUP-1...\n(hold Ctrl or Command button and click to choose multiple subjects)');
                    indx_group1 = listdlg('PromptString',{prompt,''},...
                        'SelectionMode','multiple','ListSize',[400 400],'ListString',sdir2);
                    clear sdir2 prompt
                case 'Not now'
                    tmp=clock; clk = sprintf('%.4d-%.2d-%.2d_%.2d-%.2d',tmp(1),tmp(2),tmp(3),tmp(4),tmp(5));
                    save([home_dir sprintf('workspace_cancel_%s',clk)])
                    msgbox(sprintf('Your choices so far have been saved to\n%s\nTerminating execution...',sprintf('workspace_cancel_%s.mat',clk)),'Choices saved','custom',msgicon); clear clk tmp
                    return
            end; clear answer1
            
            quest = sprintf('For stats: next, please select subjects in GROUP-2 for the two-sample t-test');
            answer1 = questdlg(quest,'Stats','Sure, take me there','Not now','Sure, take me there'); clear quest
            switch answer1
                case 'Sure, take me there'
                    cd(home_dir)
                    sdir2 = dir; sdir2(find([sdir2.isdir]==0))=[]; sdir2=sdir2(3:end); sdir2={sdir2.name};
                    f2=0;
                    for u=1:length(sdir2)
                        for u2=1:length(subnames)
                            if strcmp(sdir2{u},subnames{u2})
                                f2=f2+1;
                                sdir3{f2} = sdir2{u};
                            end
                        end
                    end; clear f2 u u2
                    clear sdir2; sdir2=sdir3; clear sdir3
                    prompt = sprintf('Select subjects in GROUP-2...\n(hold Ctrl or Command button and click to choose multiple subjects)');
                    indx_group2 = listdlg('PromptString',{prompt,''},...
                        'SelectionMode','multiple','ListSize',[400 400],'ListString',sdir2);
                    clear sdir2 prompt
                case 'Not now'
                    tmp=clock; clk = sprintf('%.4d-%.2d-%.2d_%.2d-%.2d',tmp(1),tmp(2),tmp(3),tmp(4),tmp(5));
                    save([home_dir sprintf('workspace_cancel_%s',clk)])
                    msgbox(sprintf('Your choices so far have been saved to\n%s\nTerminating execution...',sprintf('workspace_cancel_%s.mat',clk)),'Choices saved','custom',msgicon); clear clk tmp
                    return
            end; clear answer1
            
            indx_overlap = sort(unique(intersect(indx_group1,indx_group2)));
            if ~isempty(indx_overlap)
                overlap_str = '';
                for w=1:length(indx_overlap)
                    tmp_suffix = sub_dir{indx_overlap(w)}(max(strfind(sub_dir{indx_overlap(w)}(1:end-1),'/'))+1:end-1);
                    if isempty(tmp_suffix)
                        tmp_suffix = sub_dir{indx_overlap(w)}(max(strfind(sub_dir{indx_overlap(w)}(1:end-1),'\'))+1:end-1);
                    end
                    overlap_str = strcat(overlap_str,sprintf('\n%s',tmp_suffix));
                    clear tmp_suffix
                end
                answer2 = questdlg(sprintf('Error: some of the subjects (mentioned below) are in both the groups. Try choosing the subjects again without overlapping.\n%s\n',overlap_str),'Stats','Ok, take me there','Ignore this message and proceed as is','Ok, take me there');
                switch answer2
                    case 'Ok, take me there'
                        clear indx_group1 indx_group2 ttest_type indx_overlap overlap_str w
                        cd(home_dir)
                        sdir2 = dir; sdir2(find([sdir2.isdir]==0))=[]; sdir2=sdir2(3:end); sdir2={sdir2.name};
                        f2=0;
                        for u=1:length(sdir2)
                            for u2=1:length(subnames)
                                if strcmp(sdir2{u},subnames{u2})
                                    f2=f2+1;
                                    sdir3{f2} = sdir2{u};
                                end
                            end
                        end; clear f2 u u2
                        clear sdir2; sdir2=sdir3; clear sdir3
                        prompt = sprintf('Select subjects for GROUP-1...\n(hold Ctrl or Command button and click to choose multiple subjects)');
                        indx_group1 = listdlg('PromptString',{prompt,''},...
                            'SelectionMode','multiple','ListSize',[400 400],'ListString',sdir2);
                        clear sdir2 prompt
                        
                        cd(home_dir)
                        sdir2 = dir; sdir2(find([sdir2.isdir]==0))=[]; sdir2=sdir2(3:end); sdir2={sdir2.name};
                        f2=0;
                        for u=1:length(sdir2)
                            for u2=1:length(subnames)
                                if strcmp(sdir2{u},subnames{u2})
                                    f2=f2+1;
                                    sdir3{f2} = sdir2{u};
                                end
                            end
                        end; clear f2 u u2
                        clear sdir2; sdir2=sdir3; clear sdir3
                        prompt = sprintf('Select subjects for GROUP-2...\n(hold Ctrl or Command button and click to choose multiple subjects)');
                        indx_group2 = listdlg('PromptString',{prompt,''},...
                            'SelectionMode','multiple','ListSize',[400 400],'ListString',sdir2);
                        clear sdir2 prompt
                        
                        indx_overlap = sort(unique(intersect(indx_group1,indx_group2)));
                        if ~isempty(indx_overlap)
                            overlap_str = '';
                            for w=1:length(indx_overlap)
                                tmp_suffix = sub_dir{indx_overlap(w)}(max(strfind(sub_dir{indx_overlap(w)}(1:end-1),'/'))+1:end-1);
                                if isempty(tmp_suffix)
                                    tmp_suffix = sub_dir{indx_overlap(w)}(max(strfind(sub_dir{indx_overlap(w)}(1:end-1),'\'))+1:end-1);
                                end
                                overlap_str = strcat(overlap_str,sprintf('\n%s',tmp_suffix));
                                clear tmp_suffix
                            end
                            msgbox(sprintf('Error: again, some of the subjects (mentioned below) are in both the groups. Aborting.\n%s\n',overlap_str),'Abort','error');
                            return;
                        end
                        clear indx_overlap overlap_str w
                end; clear answer2
            end
    end    
end

if ~exist('alpha_stats','var') % quads
    prompt = sprintf('Please provide p-value threshold for stats.\n\n(This is an uncorrected threshold; multiple comparisons correction is not automatically performed. If you want to perform correction, do stats with uncorrected threshold and use the saved p-values outside of Neptune.)\n');
    answer3 = inputdlg(prompt,'p-value threshold for stats',1,{'0.05'}); clear prompt
    alpha_stats = str2num(cell2mat(answer3)); clear answer3
end
end

%%
```
