## Supplementary material for "Neptune: a toolbox for spinal cord functional MRI data processing and quality assurance": Neptune software: postproc_scfMRItb.m

```
% POST-PROCESSING
L=L+2; logg{L,1}=sprintf('--- Post-processing ---'); fprintf(fileID,'%s\n',logg{L,1}); try set(S.tx,'string',logg); catch, end; clear clk
clk=clock; L=L+1; logg{L,1}=sprintf('%d/%s/%d %dh:%dm:%ds - begin making post-processing choices',clk(3),month{clk(2)},clk(1),clk(4),clk(5),round(clk(6))); fprintf(fileID,'%s\n',logg{L,1}); try set(S.tx,'string',logg); catch, end; clear clk
tc0=tic;

wbar3 = waitbar(0,'Choose post-processing steps...','Name','Post-processing status...','Units','normalized','Position',[4/5 0 1/5 1/17]);

%% --- Post-processing choices ---

postproc_choices


%% ---- Which of the pre-processed files to choose for post-processing? ----
try
    load([sub_dir{1} sprintf('workspace_variables_run%d.mat',1)], 'cutoff_str', 'f_BPF_max_str')
catch
    cutoff_str = '50'; f_BPF_max_str = '10';
end

% If file {1} is available then that will be used, else it will look for
% file {2}, and so on, until it tries to find file {8} in subject folders
if     exist([sub_dir{1} func_files{1,1} '_denoised' cutoff_str 'WMcov' '.nii'], 'file') || exist([sub_dir{1} func_files{1,1} '_denoised' cutoff_str 'WMcov' '.nii.gz'], 'file') || exist([sub_dir{1} func_files{1,1} '_denoised' cutoff_str 'WMcov' '.mat'], 'file') % if CSF, WM and COV regress steps were performed
    whichip=1;
elseif exist([sub_dir{1} func_files{1,1} '_denoised' cutoff_str 'cov' '.nii'], 'file') || exist([sub_dir{1} func_files{1,1} '_denoised' cutoff_str 'cov' '.nii.gz'], 'file') || exist([sub_dir{1} func_files{1,1} '_denoised' cutoff_str 'cov' '.mat'], 'file') % if both CSF and COV regress steps were performed
    whichip=2;
elseif exist([sub_dir{1} func_files{1,1} '_denoised' cutoff_str 'WM' '.nii'], 'file') || exist([sub_dir{1} func_files{1,1} '_denoised' cutoff_str 'WM' '.nii.gz'], 'file') || exist([sub_dir{1} func_files{1,1} '_denoised' cutoff_str 'WM' '.mat'], 'file') % if both CSF and WM regress steps were performed
    whichip=3;
elseif exist([sub_dir{1} func_files{1,1} '_WMcov' '.nii'], 'file') || exist([sub_dir{1} func_files{1,1} '_WMcov' '.nii.gz'], 'file') || exist([sub_dir{1} func_files{1,1} '_WMcov' '.mat'], 'file') % if both WM and COV regress steps were performed
    whichip=4;
elseif exist([sub_dir{1} func_files{1,1} '_cov' '.nii'], 'file') || exist([sub_dir{1} func_files{1,1} '_cov' '.nii.gz'], 'file') || exist([sub_dir{1} func_files{1,1} '_cov' '.mat'], 'file') % if only COV regress step was performed
    whichip=5;
elseif exist([sub_dir{1} func_files{1,1} '_denoised' cutoff_str '.nii'], 'file') || exist([sub_dir{1} func_files{1,1} '_denoised' cutoff_str '.nii.gz'], 'file') || exist([sub_dir{1} func_files{1,1} '_denoised' cutoff_str '.mat'], 'file') % if only CSF regress step was performed
    whichip=6;
elseif exist([sub_dir{1} func_files{1,1} '_WM' '.nii'], 'file') || exist([sub_dir{1} func_files{1,1} '_WM' '.nii.gz'], 'file') || exist([sub_dir{1} func_files{1,1} '_WM' '.mat'], 'file') % if only WM regress step was performed
    whichip=7;
elseif exist([sub_dir{1} func_files{1,1} '_warped' '.nii'], 'file') || exist([sub_dir{1} func_files{1,1} '_warped' '.nii.gz'], 'file') || exist([sub_dir{1} func_files{1,1} '_warped' '.nii.gz'], 'file') % if none of CSF, WM or COV regress steps were performed
    whichip=8;
end

if whichip==1
    if use_deconv_data_for_postproc == 1 % try deconvolved data for post-processing
        postproc_filesuffix{1} = ['_denoised' cutoff_str 'WMcov' '_deconv' '_BPF' f_BPF_max_str];
        postproc_filesuffix{2} = ['_denoised' cutoff_str 'WMcov' '_BPF' f_BPF_max_str '_deconv'];
        postproc_filesuffix{3} = ['_denoised' cutoff_str 'WMcov' '_HPF01' '_deconv'];
        postproc_filesuffix{4} = ['_denoised' cutoff_str 'WMcov' '_BPF' f_BPF_max_str];
        postproc_filesuffix{5} = ['_denoised' cutoff_str 'WMcov' '_HPF' '01'];
        postproc_filesuffix{6} = ['_denoised' cutoff_str 'WMcov'];
    elseif use_deconv_data_for_postproc == 0 % try bandpass filtered data (before deconv) for post-processing
        postproc_filesuffix{1} = ['_denoised' cutoff_str 'WMcov' '_BPF' f_BPF_max_str];
        postproc_filesuffix{2} = ['_denoised' cutoff_str 'WMcov' '_HPF' '01'];
        postproc_filesuffix{3} = ['_denoised' cutoff_str 'WMcov' '_deconv' '_BPF' f_BPF_max_str];
        postproc_filesuffix{4} = ['_denoised' cutoff_str 'WMcov' '_BPF' f_BPF_max_str '_deconv'];
        postproc_filesuffix{5} = ['_denoised' cutoff_str 'WMcov' '_HPF01' '_deconv'];
        postproc_filesuffix{6} = ['_denoised' cutoff_str 'WMcov'];
    end
elseif whichip==2
    if use_deconv_data_for_postproc == 1 % try deconvolved data for post-processing
        postproc_filesuffix{1} = ['_denoised' cutoff_str 'cov' '_deconv' '_BPF' f_BPF_max_str];
        postproc_filesuffix{2} = ['_denoised' cutoff_str 'cov' '_BPF' f_BPF_max_str '_deconv'];
        postproc_filesuffix{3} = ['_denoised' cutoff_str 'cov' '_HPF01' '_deconv'];
        postproc_filesuffix{4} = ['_denoised' cutoff_str 'cov' '_BPF' f_BPF_max_str];
        postproc_filesuffix{5} = ['_denoised' cutoff_str 'cov' '_HPF' '01'];
        postproc_filesuffix{6} = ['_denoised' cutoff_str 'cov'];
    elseif use_deconv_data_for_postproc == 0 % try bandpass filtered data (before deconv) for post-processing
        postproc_filesuffix{1} = ['_denoised' cutoff_str 'cov' '_BPF' f_BPF_max_str];
        postproc_filesuffix{2} = ['_denoised' cutoff_str 'cov' '_HPF' '01'];
        postproc_filesuffix{3} = ['_denoised' cutoff_str 'cov' '_deconv' '_BPF' f_BPF_max_str];
        postproc_filesuffix{4} = ['_denoised' cutoff_str 'cov' '_BPF' f_BPF_max_str '_deconv'];
        postproc_filesuffix{5} = ['_denoised' cutoff_str 'cov' '_HPF01' '_deconv'];
        postproc_filesuffix{6} = ['_denoised' cutoff_str 'cov'];
    end
elseif whichip==3
    if use_deconv_data_for_postproc == 1 % try deconvolved data for post-processing
        postproc_filesuffix{1} = ['_denoised' cutoff_str 'WM' '_deconv' '_BPF' f_BPF_max_str];
        postproc_filesuffix{2} = ['_denoised' cutoff_str 'WM' '_BPF' f_BPF_max_str '_deconv'];
        postproc_filesuffix{3} = ['_denoised' cutoff_str 'WM' '_HPF01' '_deconv'];
        postproc_filesuffix{4} = ['_denoised' cutoff_str 'WM' '_BPF' f_BPF_max_str];
        postproc_filesuffix{5} = ['_denoised' cutoff_str 'WM' '_HPF' '01'];
        postproc_filesuffix{6} = ['_denoised' cutoff_str 'WM'];
    elseif use_deconv_data_for_postproc == 0 % try bandpass filtered data (before deconv) for post-processing
        postproc_filesuffix{1} = ['_denoised' cutoff_str 'WM' '_BPF' f_BPF_max_str];
        postproc_filesuffix{2} = ['_denoised' cutoff_str 'WM' '_HPF' '01'];
        postproc_filesuffix{3} = ['_denoised' cutoff_str 'WM' '_deconv' '_BPF' f_BPF_max_str];
        postproc_filesuffix{4} = ['_denoised' cutoff_str 'WM' '_BPF' f_BPF_max_str '_deconv'];
        postproc_filesuffix{5} = ['_denoised' cutoff_str 'WM' '_HPF01' '_deconv'];
        postproc_filesuffix{6} = ['_denoised' cutoff_str 'WM'];
    end
elseif whichip==4
    if use_deconv_data_for_postproc == 1 % try deconvolved data for post-processing
        postproc_filesuffix{1} = ['_WMcov' '_deconv' '_BPF' f_BPF_max_str];
        postproc_filesuffix{2} = ['_WMcov' '_BPF' f_BPF_max_str '_deconv'];
        postproc_filesuffix{3} = ['_WMcov' '_HPF01' '_deconv'];
        postproc_filesuffix{4} = ['_WMcov' '_BPF' f_BPF_max_str];
        postproc_filesuffix{5} = ['_WMcov' '_HPF' '01'];
        postproc_filesuffix{6} = ['_WMcov']; %#ok<*NBRAK>
    elseif use_deconv_data_for_postproc == 0 % try bandpass filtered data (before deconv) for post-processing
        postproc_filesuffix{1} = ['_WMcov' '_BPF' f_BPF_max_str];
        postproc_filesuffix{2} = ['_WMcov' '_HPF' '01'];
        postproc_filesuffix{3} = ['_WMcov' '_deconv' '_BPF' f_BPF_max_str];
        postproc_filesuffix{4} = ['_WMcov' '_BPF' f_BPF_max_str '_deconv'];
        postproc_filesuffix{5} = ['_WMcov' '_HPF01' '_deconv'];
        postproc_filesuffix{6} = ['_WMcov'];
    end
elseif whichip==5
    if use_deconv_data_for_postproc == 1 % try deconvolved data for post-processing
        postproc_filesuffix{1} = ['_cov' '_deconv' '_BPF' f_BPF_max_str];
        postproc_filesuffix{2} = ['_cov' '_BPF' f_BPF_max_str '_deconv'];
        postproc_filesuffix{3} = ['_cov' '_HPF01' '_deconv'];
        postproc_filesuffix{4} = ['_cov' '_BPF' f_BPF_max_str];
        postproc_filesuffix{5} = ['_cov' '_HPF' '01'];
        postproc_filesuffix{6} = ['_cov'];
    elseif use_deconv_data_for_postproc == 0 % try bandpass filtered data (before deconv) for post-processing
        postproc_filesuffix{1} = ['_cov' '_BPF' f_BPF_max_str];
        postproc_filesuffix{2} = ['_cov' '_HPF' '01'];
        postproc_filesuffix{3} = ['_cov' '_deconv' '_BPF' f_BPF_max_str];
        postproc_filesuffix{4} = ['_cov' '_BPF' f_BPF_max_str '_deconv'];
        postproc_filesuffix{5} = ['_cov' '_HPF01' '_deconv'];
        postproc_filesuffix{6} = ['_cov'];
    end
elseif whichip==6
    if use_deconv_data_for_postproc == 1 % try deconvolved data for post-processing
        postproc_filesuffix{1} = ['_denoised' cutoff_str '_deconv' '_BPF' f_BPF_max_str];
        postproc_filesuffix{2} = ['_denoised' cutoff_str '_BPF' f_BPF_max_str '_deconv'];
        postproc_filesuffix{3} = ['_denoised' cutoff_str '_HPF01' '_deconv'];
        postproc_filesuffix{4} = ['_denoised' cutoff_str '_BPF' f_BPF_max_str];
        postproc_filesuffix{5} = ['_denoised' cutoff_str '_HPF' '01'];
        postproc_filesuffix{6} = ['_denoised' cutoff_str];
    elseif use_deconv_data_for_postproc == 0 % try bandpass filtered data (before deconv) for post-processing
        postproc_filesuffix{1} = ['_denoised' cutoff_str '_BPF' f_BPF_max_str];
        postproc_filesuffix{2} = ['_denoised' cutoff_str '_HPF' '01'];
        postproc_filesuffix{3} = ['_denoised' cutoff_str '_deconv' '_BPF' f_BPF_max_str];
        postproc_filesuffix{4} = ['_denoised' cutoff_str '_BPF' f_BPF_max_str '_deconv'];
        postproc_filesuffix{5} = ['_denoised' cutoff_str '_HPF01' '_deconv'];
        postproc_filesuffix{6} = ['_denoised' cutoff_str];
    end
elseif whichip==7
    if use_deconv_data_for_postproc == 1 % try deconvolved data for post-processing
        postproc_filesuffix{1} = ['_WM' '_deconv' '_BPF' f_BPF_max_str];
        postproc_filesuffix{2} = ['_WM' '_BPF' f_BPF_max_str '_deconv'];
        postproc_filesuffix{3} = ['_WM' '_HPF01' '_deconv'];
        postproc_filesuffix{4} = ['_WM' '_BPF' f_BPF_max_str];
        postproc_filesuffix{5} = ['_WM' '_HPF' '01'];
        postproc_filesuffix{6} = ['_WM'];
    elseif use_deconv_data_for_postproc == 0 % try bandpass filtered data (before deconv) for post-processing
        postproc_filesuffix{1} = ['_WM' '_BPF' f_BPF_max_str];
        postproc_filesuffix{2} = ['_WM' '_HPF' '01'];
        postproc_filesuffix{3} = ['_WM' '_deconv' '_BPF' f_BPF_max_str];
        postproc_filesuffix{4} = ['_WM' '_BPF' f_BPF_max_str '_deconv'];
        postproc_filesuffix{5} = ['_WM' '_HPF01' '_deconv'];
        postproc_filesuffix{6} = ['_WM'];
    end
elseif whichip==8
    if use_deconv_data_for_postproc == 1 % try deconvolved data for post-processing
        postproc_filesuffix{1} = ['_warped' '_deconv' '_BPF' f_BPF_max_str];
        postproc_filesuffix{2} = ['_warped' '_BPF' f_BPF_max_str '_deconv'];
        postproc_filesuffix{3} = ['_warped' '_HPF01' '_deconv'];
        postproc_filesuffix{4} = ['_warped' '_BPF' f_BPF_max_str];
        postproc_filesuffix{5} = ['_warped' '_HPF' '01'];
        postproc_filesuffix{6} = ['_warped'];
    elseif use_deconv_data_for_postproc == 0 % try bandpass filtered data (before deconv) for post-processing
        postproc_filesuffix{1} = ['_warped' '_BPF' f_BPF_max_str];
        postproc_filesuffix{2} = ['_warped' '_HPF' '01'];
        postproc_filesuffix{3} = ['_warped' '_deconv' '_BPF' f_BPF_max_str];
        postproc_filesuffix{4} = ['_warped' '_BPF' f_BPF_max_str '_deconv'];
        postproc_filesuffix{5} = ['_warped' '_HPF01' '_deconv'];
        postproc_filesuffix{6} = ['_warped'];
    end
end

%% update log:
postprocsteps = {'20. Extract quadrant time series''';'21. Estimate static functional connectivity (SFC)';'22. Estimate fractional amplitude of low frequency fluctuations (fALFF)';'04. Simple stats'};
L=L+2; logg{L,1}=sprintf('Post-processing steps CHOSEN:'); fprintf(fileID,'%s\n',logg{L,1}); try set(S.tx,'string',logg); catch, end
temp=postprocsteps(find(choices_postproc==1));
for k=1:length(temp)
    L=L+1; logg{L,1}=sprintf('%s',temp{k}); fprintf(fileID,'%s\n',logg{L,1}); try set(S.tx,'string',logg); catch, end
end; clear k temp
L=L+2; logg{L,1}=sprintf('Post-processing steps OMITTED:'); fprintf(fileID,'%s\n',logg{L,1}); try set(S.tx,'string',logg); catch, end
temp=postprocsteps(find(choices_postproc==0));
for k=1:length(temp)
    L=L+1; logg{L,1}=sprintf('%s',temp{k}); fprintf(fileID,'%s\n',logg{L,1}); try set(S.tx,'string',logg); catch, end
end; clear k temp

L=L+2; logg{L,1}=sprintf('Chosen post-processing parameters:'); fprintf(fileID,'%s\n',logg{L,1}); try set(S.tx,'string',logg); catch, end
if choices_postproc(1)==1
    L=L+1; logg{L,1}=sprintf('Perform computations in the white matter (WM_flag) = %d',WM_flag); fprintf(fileID,'%s\n',logg{L,1}); try set(S.tx,'string',logg); catch, end
    L=L+1; logg{L,1}=sprintf('Assign which value as the final SFC/fALFF value?: (1) 95th percentile, (2) 50th percentile, (3) use mean time series = %d',assign_value); fprintf(fileID,'%s\n',logg{L,1}); try set(S.tx,'string',logg); catch, end
end

if choices_postproc(2)==1
    L=L+1; logg{L,1}=sprintf('SFC - (1) Pearson''s correlation, or (2) partial correlation, or (3) both (SFC_type) = %d',SFC_type); fprintf(fileID,'%s\n',logg{L,1}); try set(S.tx,'string',logg); catch, end
    L=L+1; logg{L,1}=sprintf('SFC within each slice (SFC_withinslice_flag) = %d',SFC_withinslice_flag); fprintf(fileID,'%s\n',logg{L,1}); try set(S.tx,'string',logg); catch, end
    L=L+1; logg{L,1}=sprintf('SFC between slices (SFC_betweenslice_flag) = %d',SFC_betweenslice_flag); fprintf(fileID,'%s\n',logg{L,1}); try set(S.tx,'string',logg); catch, end
end
if choices_postproc(4)==1
    L=L+1; logg{L,1}=sprintf('Stats - which statistical test to perform: %s',answer_stats); fprintf(fileID,'%s\n',logg{L,1}); try set(S.tx,'string',logg); catch, end
    if exist('answer_paired_test','var')
    L=L+1; logg{L,1}=sprintf('Stats - for the paired t-test, use runs 1 and 2 in each subject as the groups for comparison?: %s',answer_paired_test); fprintf(fileID,'%s\n',logg{L,1}); try set(S.tx,'string',logg); catch, end
    end
end
L=L+1; logg{L,1}=sprintf('--- --- ---\n'); fprintf(fileID,'%s\n',logg{L,1}); try set(S.tx,'string',logg); catch, end; tc=tic;

clear fighndl
tmp=clock; clk = sprintf('%.4d-%.2d-%.2d_%.2d-%.2d',tmp(1),tmp(2),tmp(3),tmp(4),tmp(5));
save([home_dir sprintf('neptune_PostprocSettings_%s',clk)])
msgbox(sprintf('Your post-processing choices have been saved to\n%s',[home_dir sprintf('neptune_PostprocSettings_%s.mat',clk)]),'Choices saved','custom',msgicon); clear clk tmp
refresh_html_log


%% --- if step_after_step==1 ---
if step_after_step==1
    
%% '20. Quadrant time series extraction'
if choices_postproc(1)==1
L=L+2; logg{L,1}=sprintf('-------\n%s',postprocsteps{1}); fprintf(fileID,'%s\n',logg{L,1}); try set(S.tx,'string',logg); catch, end; tc=tic;
time_taken_postproc.step20_quadMask = zeros(size(func_files,1),size(func_files,2));

skip=0;
for k=1:length(sub_dir)
    for k2=1:runs(k)
        if (exist([sub_dir{k} func_files{k,k2} postproc_filesuffix{1} '.mat'], 'file'))
            ip_file = [func_files{k,k2} postproc_filesuffix{1}];
        elseif (exist([sub_dir{k} func_files{k,k2} postproc_filesuffix{2} '.mat'], 'file'))
            ip_file = [func_files{k,k2} postproc_filesuffix{2}];
        elseif (exist([sub_dir{k} func_files{k,k2} postproc_filesuffix{3} '.mat'], 'file'))
            ip_file = [func_files{k,k2} postproc_filesuffix{3}];
        elseif (exist([sub_dir{k} func_files{k,k2} postproc_filesuffix{4} '.mat'], 'file'))
            ip_file = [func_files{k,k2} postproc_filesuffix{4}];
        elseif (exist([sub_dir{k} func_files{k,k2} postproc_filesuffix{5} '.mat'], 'file'))
            ip_file = [func_files{k,k2} postproc_filesuffix{5}];
        elseif (exist([sub_dir{k} func_files{k,k2} postproc_filesuffix{6} '.mat'], 'file'))
            ip_file = [func_files{k,k2} postproc_filesuffix{6}];
        else
            questdlg('Unable to locate denoised, filtered or deconvolved 3D+time fMRI data','Error','Abort','Abort');
            break;
        end

        sdirz = dir([sub_dir{k} ip_file '_quads_timeseries' '.mat']);  %ip_file = sdirz(1).name(1:end-21);
        if isempty(sdirz)
            skip=skip+1;
            clk=clock; L=L+1; logg{L,1}=sprintf('%d/%s/%d %dh:%dm:%ds - begin (20) quadrant time series extraction for subject (%d) of (%d), run (%d) of (%d)',clk(3),month{clk(2)},clk(1),clk(4),clk(5),round(clk(6)),k,length(sub_dir),k2,runs(k)); fprintf(fileID,'%s\n',logg{L,1}); try set(S.tx,'string',logg); catch, end; clear clk
            clk=clock; L=L+1; logg{L,1}=sprintf('Input data for subject (%d) of (%d), run (%d) of (%d): %s',k,length(sub_dir),k2,runs(k),ip_file); fprintf(fileID,'%s\n',logg{L,1}); try set(S.tx,'string',logg); catch, end; clear clk
            try waitbar(((runs(k)*(k-1)+k2)/(length(sub_dir)*runs(k))),wbar3,sprintf('20. Quadrant time series extraction: Subject (%d) of (%d), Run (%d) of (%d)',k,length(sub_dir),k2,runs(k))); catch, end
            fprintf('20. Quadrant time series extraction: Subject (%d) of (%d), Run (%d) of (%d)\n',k,length(sub_dir),k2,runs(k))
            if runs(k)==1, Rns=0; else, Rns=k2; end
            time_20_quadMask = tic;
            scfMRItb_20_quadMask(sub_dir{k},ip_file, WM_flag,Rns);
            time_taken_postproc.step20_quadMask(k,k2) = toc(time_20_quadMask); clear time_20_quadMask
            clk=clock; L=L+1; logg{L,1}=sprintf('%d/%s/%d %dh:%dm:%ds - end (20) quadrant time series extraction for subject (%d) of (%d), run (%d) of (%d)',clk(3),month{clk(2)},clk(1),clk(4),clk(5),round(clk(6)),k,length(sub_dir),k2,runs(k)); fprintf(fileID,'%s\n',logg{L,1}); try set(S.tx,'string',logg); catch, end; clear clk
        else
            clk=clock; L=L+1; logg{L,1}=sprintf('%d/%s/%d %dh:%dm:%ds - skip (20) quadrant time series extraction for subject (%d) of (%d), run (%d) of (%d)',clk(3),month{clk(2)},clk(1),clk(4),clk(5),round(clk(6)),k,length(sub_dir),k2,runs(k)); fprintf(fileID,'%s\n',logg{L,1}); try set(S.tx,'string',logg); catch, end; clear clk
        end
        clear sdirz ip_file
    end
    clear k2
end

if skip==0
    clk=clock; L=L+1; logg{L,1}=sprintf('%d/%s/%d %dh:%dm:%ds - (20) quadrant time series extraction SKIPPED FOR ALL SUBJECTS',clk(3),month{clk(2)},clk(1),clk(4),clk(5),round(clk(6))); fprintf(fileID,'%s\n',logg{L,1}); try set(S.tx,'string',logg); catch, end; clear clk
else
    clk=clock; L=L+1; logg{L,1}=sprintf('%d/%s/%d %dh:%dm:%ds - (20) quadrant time series extraction COMPLETED FOR ALL SUBJECTS',clk(3),month{clk(2)},clk(1),clk(4),clk(5),round(clk(6))); fprintf(fileID,'%s\n',logg{L,1}); try set(S.tx,'string',logg); catch, end; clear clk
    L=L+1; logg{L,1}=sprintf('Total time taken so far: %dh:%dm:%ds\nTime taken for this step: %dh:%dm:%ds\nTime taken for this step in each run of each subject: %dh:%dm:%ds',floor(toc(tc0)/3600),floor(mod(toc(tc0),3600)/60),floor(mod(toc(tc0),60)),floor(toc(tc)/3600),floor(mod(toc(tc),3600)/60),floor(mod(toc(tc),60)),floor(toc(tc)/3600/skip),floor(mod(toc(tc),3600)/60/skip),floor(mod(toc(tc),60)/skip)); fprintf(fileID,'%s\n',logg{L,1}); try set(S.tx,'string',logg); catch, end; clear clk
    refresh_html
end
end

%% '21. Static functional connectivity'
if choices_postproc(2)==1
L=L+2; logg{L,1}=sprintf('-------\n%s',postprocsteps{2}); fprintf(fileID,'%s\n',logg{L,1}); try set(S.tx,'string',logg); catch, end; tc=tic;
time_taken_postproc.step21_SFC_withinslice = zeros(size(func_files,1),size(func_files,2));
time_taken_postproc.step21_SFC_betweenslice = zeros(size(func_files,1),size(func_files,2));

skip=0;
for k=1:length(sub_dir)
    for k2=1:runs(k)
        if (exist([sub_dir{k} func_files{k,k2} postproc_filesuffix{1} '_quads_timeseries' '.mat'], 'file'))
            ip_file = [func_files{k,k2} postproc_filesuffix{1} '_quads_timeseries'];
        elseif (exist([sub_dir{k} func_files{k,k2} postproc_filesuffix{2} '_quads_timeseries' '.mat'], 'file'))
            ip_file = [func_files{k,k2} postproc_filesuffix{2} '_quads_timeseries'];
        elseif (exist([sub_dir{k} func_files{k,k2} postproc_filesuffix{3} '_quads_timeseries' '.mat'], 'file'))
            ip_file = [func_files{k,k2} postproc_filesuffix{3} '_quads_timeseries'];
        elseif (exist([sub_dir{k} func_files{k,k2} postproc_filesuffix{4} '_quads_timeseries' '.mat'], 'file'))
            ip_file = [func_files{k,k2} postproc_filesuffix{4} '_quads_timeseries'];
        elseif (exist([sub_dir{k} func_files{k,k2} postproc_filesuffix{5} '_quads_timeseries' '.mat'], 'file'))
            ip_file = [func_files{k,k2} postproc_filesuffix{5} '_quads_timeseries'];
        elseif (exist([sub_dir{k} func_files{k,k2} postproc_filesuffix{6} '_quads_timeseries' '.mat'], 'file'))
            ip_file = [func_files{k,k2} postproc_filesuffix{6} '_quads_timeseries'];
        else
            questdlg('Unable to locate denoised, filtered or deconvolved 3D+time data','Error','Abort','Abort');
            break;
        end
        naming = ip_file(1:end-17);

        sdirz1 = dir([sub_dir{k} naming '_SFC_withinslice' '.mat']); sdirz2 = dir([sub_dir{k} naming '_SFC_withinslice_pcor' '.mat']);  sdirz3 = dir([sub_dir{k} naming '_SFC_betweenslice' '.mat']); sdirz4 = dir([sub_dir{k} naming '_SFC_betweenslice_pcor' '.mat']);  %ip_file = sdirz(1).name(1:end-8);
        if (SFC_withinslice_flag==1)&&(SFC_betweenslice_flag==1)&&(SFC_type==3)
            ifstring = (isempty(sdirz1))||(isempty(sdirz2))||(isempty(sdirz3))||(isempty(sdirz4));
        elseif (SFC_withinslice_flag==1)&&(SFC_betweenslice_flag==1)&&(SFC_type==2)
            ifstring = (isempty(sdirz2))||(isempty(sdirz4));
        elseif (SFC_withinslice_flag==1)&&(SFC_betweenslice_flag==1)&&(SFC_type==1)
            ifstring = (isempty(sdirz1))||(isempty(sdirz3));
        elseif (SFC_withinslice_flag==1)&&(SFC_betweenslice_flag==0)&&(SFC_type==3)
            ifstring = (isempty(sdirz1))||(isempty(sdirz2));
        elseif (SFC_withinslice_flag==1)&&(SFC_betweenslice_flag==0)&&(SFC_type==2)
            ifstring = (isempty(sdirz2));
        elseif (SFC_withinslice_flag==1)&&(SFC_betweenslice_flag==0)&&(SFC_type==1)
            ifstring = (isempty(sdirz1));
        elseif (SFC_withinslice_flag==0)&&(SFC_betweenslice_flag==1)&&(SFC_type==3)
            ifstring = (isempty(sdirz3))||(isempty(sdirz4));
        elseif (SFC_withinslice_flag==0)&&(SFC_betweenslice_flag==1)&&(SFC_type==2)
            ifstring = (isempty(sdirz4));
        elseif (SFC_withinslice_flag==0)&&(SFC_betweenslice_flag==1)&&(SFC_type==1)
            ifstring = (isempty(sdirz3));
        end

        if ifstring
            skip=skip+1;
            clk=clock; L=L+1; logg{L,1}=sprintf('%d/%s/%d %dh:%dm:%ds - begin (21) static functional connectivity for subject (%d) of (%d), run (%d) of (%d)',clk(3),month{clk(2)},clk(1),clk(4),clk(5),round(clk(6)),k,length(sub_dir),k2,runs(k)); fprintf(fileID,'%s\n',logg{L,1}); try set(S.tx,'string',logg); catch, end; clear clk
            try waitbar(((runs(k)*(k-1)+k2)/(length(sub_dir)*runs(k))),wbar3,sprintf('21. Static functional connectivity: Subject (%d) of (%d), Run (%d) of (%d)',k,length(sub_dir),k2,runs(k))); catch, end
            fprintf('21. Static functional connectivity: Subject (%d) of (%d), Run (%d) of (%d)\n',k,length(sub_dir),k2,runs(k))
            if runs(k)==1, Rns=0; else, Rns=k2; end

            SFC_type2 = SFC_type;
            if ~((isempty(sdirz1)) || (isempty(sdirz3)))
                if SFC_type==1, SFC_type2=0; elseif SFC_type==3, SFC_type2=2; end
            end
            if ~((isempty(sdirz2)) || (isempty(sdirz4)))
                if SFC_type==2, SFC_type2=0; elseif SFC_type==3, SFC_type2=1; end
            end
            
            if SFC_type2~=0
            if SFC_withinslice_flag==1
                time_21_SFC_withinslice = tic;
                scfMRItb_21_SFC_withinslice(sub_dir{k},ip_file,naming, SFC_type2,WM_flag,assign_value,Rns);
                time_taken_postproc.step21_SFC_withinslice(k,k2) = toc(time_21_SFC_withinslice); clear time_21_SFC_withinslice
            end
            if SFC_betweenslice_flag==1
                time_21_SFC_betweenslice = tic;
                scfMRItb_21_SFC_betweenslice(sub_dir{k},ip_file,naming, SFC_type2,WM_flag,assign_value,Rns, dont_do_pcor_betweenslice);
                time_taken_postproc.step21_SFC_betweenslice(k,k2) = toc(time_21_SFC_betweenslice); clear time_21_SFC_betweenslice
            end
            else
                fprintf('Did not perform SFC for subject %d run %d because a previously saved SFC file already exists. If you want to run, move that file from the subject''s folder, rename or delete it\n\n',k,k2)
                clk=clock; L=L+1; logg{L,1}=sprintf('%d/%s/%d %dh:%dm:%ds - Message: Did not perform SFC for subject %d run %d because a previously saved SFC file already exists. If you want to run, move that file from the subject''s folder or delete it\n\n',clk(3),month{clk(2)},clk(1),clk(4),clk(5),round(clk(6)),k,k2); fprintf(fileID,'%s\n',logg{L,1}); try set(S.tx,'string',logg); catch, end; clear clk
            end
            clk=clock; L=L+1; logg{L,1}=sprintf('%d/%s/%d %dh:%dm:%ds - end (21) static functional connectivity for subject (%d) of (%d), run (%d) of (%d)',clk(3),month{clk(2)},clk(1),clk(4),clk(5),round(clk(6)),k,length(sub_dir),k2,runs(k)); fprintf(fileID,'%s\n',logg{L,1}); try set(S.tx,'string',logg); catch, end; clear clk
        else
            clk=clock; L=L+1; logg{L,1}=sprintf('%d/%s/%d %dh:%dm:%ds - skip (21) static functional connectivity for subject (%d) of (%d), run (%d) of (%d)',clk(3),month{clk(2)},clk(1),clk(4),clk(5),round(clk(6)),k,length(sub_dir),k2,runs(k)); fprintf(fileID,'%s\n',logg{L,1}); try set(S.tx,'string',logg); catch, end; clear clk
        end
        clear sdirz1 sdirz2 sdirz3 sdirz4 ip_file naming SFC_type2 ifstring
    end
    clear k2
end

if skip==0
    clk=clock; L=L+1; logg{L,1}=sprintf('%d/%s/%d %dh:%dm:%ds - (21) static functional connectivity SKIPPED FOR ALL SUBJECTS',clk(3),month{clk(2)},clk(1),clk(4),clk(5),round(clk(6))); fprintf(fileID,'%s\n',logg{L,1}); try set(S.tx,'string',logg); catch, end; clear clk
else
    clk=clock; L=L+1; logg{L,1}=sprintf('%d/%s/%d %dh:%dm:%ds - (21) static functional connectivity COMPLETED FOR ALL SUBJECTS',clk(3),month{clk(2)},clk(1),clk(4),clk(5),round(clk(6))); fprintf(fileID,'%s\n',logg{L,1}); try set(S.tx,'string',logg); catch, end; clear clk
    L=L+1; logg{L,1}=sprintf('Total time taken so far: %dh:%dm:%ds\nTime taken for this step: %dh:%dm:%ds\nTime taken for this step in each run of each subject: %dh:%dm:%ds',floor(toc(tc0)/3600),floor(mod(toc(tc0),3600)/60),floor(mod(toc(tc0),60)),floor(toc(tc)/3600),floor(mod(toc(tc),3600)/60),floor(mod(toc(tc),60)),floor(toc(tc)/3600/skip),floor(mod(toc(tc),3600)/60/skip),floor(mod(toc(tc),60)/skip)); fprintf(fileID,'%s\n',logg{L,1}); try set(S.tx,'string',logg); catch, end; clear clk
    refresh_html
end
end

%% '22. fALFF'
if choices_postproc(3)==1
L=L+2; logg{L,1}=sprintf('-------\n%s',postprocsteps{3}); fprintf(fileID,'%s\n',logg{L,1}); try set(S.tx,'string',logg); catch, end; tc=tic;
time_taken_postproc.step22_fALFF = zeros(size(func_files,1),size(func_files,2));

skip=0;
for k=1:length(sub_dir)
    for k2=1:runs(k)
        if use_deconv_data_for_postproc == 1
            if (exist([sub_dir{k} func_files{k,k2} postproc_filesuffix{3} '.mat'], 'file'))
                ip_file = [func_files{k,k2} postproc_filesuffix{3}];
            elseif (exist([sub_dir{k} func_files{k,k2} postproc_filesuffix{5} '.mat'], 'file'))
                ip_file = [func_files{k,k2} postproc_filesuffix{5}];
            elseif (exist([sub_dir{k} func_files{k,k2} postproc_filesuffix{6} '.mat'], 'file'))
                ip_file = [func_files{k,k2} postproc_filesuffix{6}];
            else
                questdlg('Unable to locate unfiltered 3D+time fMRI data for fALFF','Error','Abort','Abort');
                break;
            end
        elseif use_deconv_data_for_postproc == 0
            if (exist([sub_dir{k} func_files{k,k2} postproc_filesuffix{2} '.mat'], 'file'))
                ip_file = [func_files{k,k2} postproc_filesuffix{2}];
            elseif (exist([sub_dir{k} func_files{k,k2} postproc_filesuffix{5} '.mat'], 'file'))
                ip_file = [func_files{k,k2} postproc_filesuffix{5}];
            elseif (exist([sub_dir{k} func_files{k,k2} postproc_filesuffix{6} '.mat'], 'file'))
                ip_file = [func_files{k,k2} postproc_filesuffix{6}];
            else
                questdlg('Unable to locate unfiltered 3D+time fMRI data for fALFF','Error','Abort','Abort');
                break;
            end
        end

        sdirz = dir([sub_dir{k} ip_file '_fALFF' '.mat']);
        if isempty(sdirz)
            skip=skip+1;
            clk=clock; L=L+1; logg{L,1}=sprintf('%d/%s/%d %dh:%dm:%ds - begin (22) fALFF for subject (%d) of (%d), run (%d) of (%d)',clk(3),month{clk(2)},clk(1),clk(4),clk(5),round(clk(6)),k,length(sub_dir),k2,runs(k)); fprintf(fileID,'%s\n',logg{L,1}); try set(S.tx,'string',logg); catch, end; clear clk
            try waitbar(((runs(k)*(k-1)+k2)/(length(sub_dir)*runs(k))),wbar3,sprintf('22. fALFF: Subject (%d) of (%d), Run (%d) of (%d)',k,length(sub_dir),k2,runs(k))); catch, end
            fprintf('22. fALFF: Subject (%d) of (%d), Run (%d) of (%d)\n',k,length(sub_dir),k2,runs(k))
            if runs(k)==1, Rns=0; else, Rns=k2; end
            time_22_fALFF = tic;
            scfMRItb_22_fALFF(sub_dir{k},ip_file, WM_flag,assign_value,Rns);
            time_taken_postproc.step22_fALFF(k,k2) = toc(time_22_fALFF); clear time_22_fALFF
            clk=clock; L=L+1; logg{L,1}=sprintf('%d/%s/%d %dh:%dm:%ds - end (22) fALFF for subject (%d) of (%d), run (%d) of (%d)',clk(3),month{clk(2)},clk(1),clk(4),clk(5),round(clk(6)),k,length(sub_dir),k2,runs(k)); fprintf(fileID,'%s\n',logg{L,1}); try set(S.tx,'string',logg); catch, end; clear clk
        else
            clk=clock; L=L+1; logg{L,1}=sprintf('%d/%s/%d %dh:%dm:%ds - skip (21) fALFF for subject (%d) of (%d), run (%d) of (%d)',clk(3),month{clk(2)},clk(1),clk(4),clk(5),round(clk(6)),k,length(sub_dir),k2,runs(k)); fprintf(fileID,'%s\n',logg{L,1}); try set(S.tx,'string',logg); catch, end; clear clk
        end
        clear sdirz ip_file naming
    end
    clear k2
end

if skip==0
    clk=clock; L=L+1; logg{L,1}=sprintf('%d/%s/%d %dh:%dm:%ds - (22) fALFF SKIPPED FOR ALL SUBJECTS',clk(3),month{clk(2)},clk(1),clk(4),clk(5),round(clk(6))); fprintf(fileID,'%s\n',logg{L,1}); try set(S.tx,'string',logg); catch, end; clear clk
else
    clk=clock; L=L+1; logg{L,1}=sprintf('%d/%s/%d %dh:%dm:%ds - (22) fALFF COMPLETED FOR ALL SUBJECTS',clk(3),month{clk(2)},clk(1),clk(4),clk(5),round(clk(6))); fprintf(fileID,'%s\n',logg{L,1}); try set(S.tx,'string',logg); catch, end; clear clk
    L=L+1; logg{L,1}=sprintf('Total time taken so far: %dh:%dm:%ds\nTime taken for this step: %dh:%dm:%ds\nTime taken for this step in each run of each subject: %dh:%dm:%ds',floor(toc(tc0)/3600),floor(mod(toc(tc0),3600)/60),floor(mod(toc(tc0),60)),floor(toc(tc)/3600),floor(mod(toc(tc),3600)/60),floor(mod(toc(tc),60)),floor(toc(tc)/3600/skip),floor(mod(toc(tc),3600)/60/skip),floor(mod(toc(tc),60)/skip)); fprintf(fileID,'%s\n',logg{L,1}); try set(S.tx,'string',logg); catch, end; clear clk
    refresh_html_log
end
end

end
%%
%%
%% --- if step_after_step==0 (i.e. do subject-after-subject) ---
if step_after_step==0

%%
for k=sub_aft_sub_num:length(sub_dir)
    for k2=1:runs(k)
        L=L+2; logg{L,1}=sprintf('-------\nPost-processing Subject (%d) of (%d), Run (%d) of (%d)',k,length(sub_dir),k2,runs(k)); fprintf(fileID,'%s\n',logg{L,1}); try set(S.tx,'string',logg); catch, end; tc=tic;

%% '20. Quadrant time series extraction'
if choices_postproc(2)==1
    if (exist([sub_dir{k} func_files{k,k2} postproc_filesuffix{1} '.mat'], 'file'))
        ip_file = [func_files{k,k2} postproc_filesuffix{1}];
    elseif (exist([sub_dir{k} func_files{k,k2} postproc_filesuffix{2} '.mat'], 'file'))
        ip_file = [func_files{k,k2} postproc_filesuffix{2}];
    elseif (exist([sub_dir{k} func_files{k,k2} postproc_filesuffix{3} '.mat'], 'file'))
        ip_file = [func_files{k,k2} postproc_filesuffix{3}];
    elseif (exist([sub_dir{k} func_files{k,k2} postproc_filesuffix{4} '.mat'], 'file'))
        ip_file = [func_files{k,k2} postproc_filesuffix{4}];
    elseif (exist([sub_dir{k} func_files{k,k2} postproc_filesuffix{5} '.mat'], 'file'))
        ip_file = [func_files{k,k2} postproc_filesuffix{5}];
    elseif (exist([sub_dir{k} func_files{k,k2} postproc_filesuffix{6} '.mat'], 'file'))
        ip_file = [func_files{k,k2} postproc_filesuffix{6}];
    else
        questdlg('Unable to locate denoised, filtered or deconvolved 3D+time data','Error','Abort','Abort');
        break;
    end

    sdirz = dir([sub_dir{k} ip_file '_quads_timeseries' '.mat']);  %ip_file = sdirz(1).name(1:end-21);
    if isempty(sdirz)
        clk=clock; L=L+1; logg{L,1}=sprintf('%d/%s/%d %dh:%dm:%ds - begin (20) quadrant time series extraction for subject (%d) of (%d), run (%d) of (%d)',clk(3),month{clk(2)},clk(1),clk(4),clk(5),round(clk(6)),k,length(sub_dir),k2,runs(k)); fprintf(fileID,'%s\n',logg{L,1}); try set(S.tx,'string',logg); catch, end; clear clk
        clk=clock; L=L+1; logg{L,1}=sprintf('Input data for subject (%d) of (%d), run (%d) of (%d): %s',k,length(sub_dir),k2,runs(k),ip_file); fprintf(fileID,'%s\n',logg{L,1}); try set(S.tx,'string',logg); catch, end; clear clk
        try waitbar(((runs(k)*(k-1)+k2)/(length(sub_dir)*runs(k))),wbar3,sprintf('20. Quadrant time series extraction: Subject (%d) of (%d), Run (%d) of (%d)',k,length(sub_dir),k2,runs(k))); catch, end
        fprintf('20. Quadrant time series extraction: Subject (%d) of (%d), Run (%d) of (%d)\n',k,length(sub_dir),k2,runs(k))
        if runs(k)==1, Rns=0; else, Rns=k2; end
        time_20_quadMask = tic;
        scfMRItb_20_quadMask(sub_dir{k},ip_file, WM_flag,Rns);
        time_taken_postproc.step20_quadMask(k,k2) = toc(time_20_quadMask); clear time_20_quadMask
        clk=clock; L=L+1; logg{L,1}=sprintf('%d/%s/%d %dh:%dm:%ds - end (20) quadrant time series extraction for subject (%d) of (%d), run (%d) of (%d)',clk(3),month{clk(2)},clk(1),clk(4),clk(5),round(clk(6)),k,length(sub_dir),k2,runs(k)); fprintf(fileID,'%s\n',logg{L,1}); try set(S.tx,'string',logg); catch, end; clear clk
    else
        clk=clock; L=L+1; logg{L,1}=sprintf('%d/%s/%d %dh:%dm:%ds - skip (20) quadrant time series extraction for subject (%d) of (%d), run (%d) of (%d)',clk(3),month{clk(2)},clk(1),clk(4),clk(5),round(clk(6)),k,length(sub_dir),k2,runs(k)); fprintf(fileID,'%s\n',logg{L,1}); try set(S.tx,'string',logg); catch, end; clear clk
    end
    clear sdirz ip_file
end

%% '21. Static functional connectivity'
if choices_postproc(2)==1
    if (exist([sub_dir{k} func_files{k,k2} postproc_filesuffix{1} '_quads_timeseries' '.mat'], 'file'))
        ip_file = [func_files{k,k2} postproc_filesuffix{1} '_quads_timeseries'];
    elseif (exist([sub_dir{k} func_files{k,k2} postproc_filesuffix{2} '_quads_timeseries' '.mat'], 'file'))
        ip_file = [func_files{k,k2} postproc_filesuffix{2} '_quads_timeseries'];
    elseif (exist([sub_dir{k} func_files{k,k2} postproc_filesuffix{3} '_quads_timeseries' '.mat'], 'file'))
        ip_file = [func_files{k,k2} postproc_filesuffix{3} '_quads_timeseries'];
    elseif (exist([sub_dir{k} func_files{k,k2} postproc_filesuffix{4} '_quads_timeseries' '.mat'], 'file'))
        ip_file = [func_files{k,k2} postproc_filesuffix{4} '_quads_timeseries'];
    elseif (exist([sub_dir{k} func_files{k,k2} postproc_filesuffix{5} '_quads_timeseries' '.mat'], 'file'))
        ip_file = [func_files{k,k2} postproc_filesuffix{5} '_quads_timeseries'];
    elseif (exist([sub_dir{k} func_files{k,k2} postproc_filesuffix{6} '_quads_timeseries' '.mat'], 'file'))
        ip_file = [func_files{k,k2} postproc_filesuffix{6} '_quads_timeseries'];
    else
        questdlg('Unable to locate denoised, filtered or deconvolved 3D+time data','Error','Abort','Abort');
        break;
    end
    naming = ip_file(1:end-17);
    
    sdirz1 = dir([sub_dir{k} naming '_SFC_withinslice' '.mat']); sdirz2 = dir([sub_dir{k} naming '_SFC_withinslice_pcor' '.mat']);  sdirz3 = dir([sub_dir{k} naming '_SFC_betweenslice' '.mat']); sdirz4 = dir([sub_dir{k} naming '_SFC_betweenslice_pcor' '.mat']);  %ip_file = sdirz(1).name(1:end-8);
    if (SFC_withinslice_flag==1)&&(SFC_betweenslice_flag==1)&&(SFC_type==3)
        ifstring = (isempty(sdirz1))||(isempty(sdirz2))||(isempty(sdirz3))||(isempty(sdirz4));
    elseif (SFC_withinslice_flag==1)&&(SFC_betweenslice_flag==1)&&(SFC_type==2)
        ifstring = (isempty(sdirz2))||(isempty(sdirz4));
    elseif (SFC_withinslice_flag==1)&&(SFC_betweenslice_flag==1)&&(SFC_type==1)
        ifstring = (isempty(sdirz1))||(isempty(sdirz3));
    elseif (SFC_withinslice_flag==1)&&(SFC_betweenslice_flag==0)&&(SFC_type==3)
        ifstring = (isempty(sdirz1))||(isempty(sdirz2));
    elseif (SFC_withinslice_flag==1)&&(SFC_betweenslice_flag==0)&&(SFC_type==2)
        ifstring = (isempty(sdirz2));
    elseif (SFC_withinslice_flag==1)&&(SFC_betweenslice_flag==0)&&(SFC_type==1)
        ifstring = (isempty(sdirz1));
    elseif (SFC_withinslice_flag==0)&&(SFC_betweenslice_flag==1)&&(SFC_type==3)
        ifstring = (isempty(sdirz3))||(isempty(sdirz4));
    elseif (SFC_withinslice_flag==0)&&(SFC_betweenslice_flag==1)&&(SFC_type==2)
        ifstring = (isempty(sdirz4));
    elseif (SFC_withinslice_flag==0)&&(SFC_betweenslice_flag==1)&&(SFC_type==1)
        ifstring = (isempty(sdirz3));
    end
    
    if ifstring
        clk=clock; L=L+1; logg{L,1}=sprintf('%d/%s/%d %dh:%dm:%ds - begin (21) static functional connectivity for subject (%d) of (%d), run (%d) of (%d)',clk(3),month{clk(2)},clk(1),clk(4),clk(5),round(clk(6)),k,length(sub_dir),k2,runs(k)); fprintf(fileID,'%s\n',logg{L,1}); try set(S.tx,'string',logg); catch, end; clear clk
        try waitbar(((runs(k)*(k-1)+k2)/(length(sub_dir)*runs(k))),wbar3,sprintf('21. Static functional connectivity: Subject (%d) of (%d), Run (%d) of (%d)',k,length(sub_dir),k2,runs(k))); catch, end
        fprintf('21. Static functional connectivity: Subject (%d) of (%d), Run (%d) of (%d)\n',k,length(sub_dir),k2,runs(k))
        if runs(k)==1, Rns=0; else, Rns=k2; end
        
        SFC_type2 = SFC_type;
        if ~((isempty(sdirz1)) || (isempty(sdirz3)))
            if SFC_type==1, SFC_type2=0; elseif SFC_type==3, SFC_type2=2; end
        end
        if ~((isempty(sdirz2)) || (isempty(sdirz4)))
            if SFC_type==2, SFC_type2=0; elseif SFC_type==3, SFC_type2=1; end
        end
        
        if SFC_type2~=0
            if SFC_withinslice_flag==1
                time_21_SFC_withinslice = tic;
                scfMRItb_21_SFC_withinslice(sub_dir{k},ip_file,naming, SFC_type2,WM_flag,assign_value,Rns);
                time_taken_postproc.step21_SFC_withinslice(k,k2) = toc(time_21_SFC_withinslice); clear time_21_SFC_withinslice
            end
            if SFC_betweenslice_flag==1
                time_21_SFC_betweenslice = tic;
                scfMRItb_21_SFC_betweenslice(sub_dir{k},ip_file,naming, SFC_type2,WM_flag,assign_value,Rns, dont_do_pcor_betweenslice);
                time_taken_postproc.step21_SFC_betweenslice(k,k2) = toc(time_21_SFC_betweenslice); clear time_21_SFC_betweenslice
            end
        else
            fprintf('Did not perform SFC for subject %d run %d because a previously saved SFC file already exists. If you want to run, move that file from the subject''s folder or delete it\n\n',k,k2)
            clk=clock; L=L+1; logg{L,1}=sprintf('%d/%s/%d %dh:%dm:%ds - Message: Did not perform SFC for subject %d run %d because a previously saved SFC file already exists. If you want to run, move that file from the subject''s folder or delete it\n\n',clk(3),month{clk(2)},clk(1),clk(4),clk(5),round(clk(6)),k,k2); fprintf(fileID,'%s\n',logg{L,1}); try set(S.tx,'string',logg); catch, end; clear clk
        end
        clk=clock; L=L+1; logg{L,1}=sprintf('%d/%s/%d %dh:%dm:%ds - end (21) static functional connectivity for subject (%d) of (%d), run (%d) of (%d)',clk(3),month{clk(2)},clk(1),clk(4),clk(5),round(clk(6)),k,length(sub_dir),k2,runs(k)); fprintf(fileID,'%s\n',logg{L,1}); try set(S.tx,'string',logg); catch, end; clear clk
    else
        clk=clock; L=L+1; logg{L,1}=sprintf('%d/%s/%d %dh:%dm:%ds - skip (21) static functional connectivity for subject (%d) of (%d), run (%d) of (%d)',clk(3),month{clk(2)},clk(1),clk(4),clk(5),round(clk(6)),k,length(sub_dir),k2,runs(k)); fprintf(fileID,'%s\n',logg{L,1}); try set(S.tx,'string',logg); catch, end; clear clk
    end
    clear sdirz1 sdirz2 sdirz3 sdirz4 ip_file naming SFC_type2 ifstring
end

%% '22. fALFF'
if choices_postproc(3)==1
    if use_deconv_data_for_postproc == 1
        if (exist([sub_dir{k} func_files{k,k2} postproc_filesuffix{3} '.mat'], 'file'))
            ip_file = [func_files{k,k2} postproc_filesuffix{3}];
        elseif (exist([sub_dir{k} func_files{k,k2} postproc_filesuffix{5} '.mat'], 'file'))
            ip_file = [func_files{k,k2} postproc_filesuffix{5}];
        elseif (exist([sub_dir{k} func_files{k,k2} postproc_filesuffix{6} '.mat'], 'file'))
            ip_file = [func_files{k,k2} postproc_filesuffix{6}];
        else
            questdlg('Unable to locate unfiltered 3D+time fMRI data for fALFF','Error','Abort','Abort');
            break;
        end
    elseif use_deconv_data_for_postproc == 0
        if (exist([sub_dir{k} func_files{k,k2} postproc_filesuffix{2} '.mat'], 'file'))
            ip_file = [func_files{k,k2} postproc_filesuffix{2}];
        elseif (exist([sub_dir{k} func_files{k,k2} postproc_filesuffix{5} '.mat'], 'file'))
            ip_file = [func_files{k,k2} postproc_filesuffix{5}];
        elseif (exist([sub_dir{k} func_files{k,k2} postproc_filesuffix{6} '.mat'], 'file'))
            ip_file = [func_files{k,k2} postproc_filesuffix{6}];
        else
            questdlg('Unable to locate unfiltered 3D+time fMRI data for fALFF','Error','Abort','Abort');
            break;
        end
    end
    
    sdirz = dir([sub_dir{k} ip_file '_fALFF' '.mat']);
    if isempty(sdirz)
        clk=clock; L=L+1; logg{L,1}=sprintf('%d/%s/%d %dh:%dm:%ds - begin (22) fALFF for subject (%d) of (%d), run (%d) of (%d)',clk(3),month{clk(2)},clk(1),clk(4),clk(5),round(clk(6)),k,length(sub_dir),k2,runs(k)); fprintf(fileID,'%s\n',logg{L,1}); try set(S.tx,'string',logg); catch, end; clear clk
        try waitbar(((runs(k)*(k-1)+k2)/(length(sub_dir)*runs(k))),wbar3,sprintf('22. fALFF: Subject (%d) of (%d), Run (%d) of (%d)',k,length(sub_dir),k2,runs(k))); catch, end
        fprintf('22. fALFF: Subject (%d) of (%d), Run (%d) of (%d)\n',k,length(sub_dir),k2,runs(k))
        if runs(k)==1, Rns=0; else, Rns=k2; end
        time_22_fALFF = tic;
        scfMRItb_22_fALFF(sub_dir{k},ip_file, WM_flag,assign_value,Rns);
        time_taken_postproc.step22_fALFF(k,k2) = toc(time_22_fALFF); clear time_22_fALFF
        clk=clock; L=L+1; logg{L,1}=sprintf('%d/%s/%d %dh:%dm:%ds - end (22) fALFF for subject (%d) of (%d), run (%d) of (%d)',clk(3),month{clk(2)},clk(1),clk(4),clk(5),round(clk(6)),k,length(sub_dir),k2,runs(k)); fprintf(fileID,'%s\n',logg{L,1}); try set(S.tx,'string',logg); catch, end; clear clk
    else
        clk=clock; L=L+1; logg{L,1}=sprintf('%d/%s/%d %dh:%dm:%ds - skip (21) fALFF for subject (%d) of (%d), run (%d) of (%d)',clk(3),month{clk(2)},clk(1),clk(4),clk(5),round(clk(6)),k,length(sub_dir),k2,runs(k)); fprintf(fileID,'%s\n',logg{L,1}); try set(S.tx,'string',logg); catch, end; clear clk
    end
    clear sdirz ip_file naming
end

%%
    clk=clock; L=L+1; logg{L,1}=sprintf('%d/%s/%d %dh:%dm:%ds - Subject (%d) of (%d), Run (%d) of (%d) COMPLETED',clk(3),month{clk(2)},clk(1),clk(4),clk(5),round(clk(6)),k,length(sub_dir),k2,runs(k)); fprintf(fileID,'%s\n',logg{L,1}); try set(S.tx,'string',logg); catch, end; clear clk
    L=L+1; logg{L,1}=sprintf('Total time taken so far: %dh:%dm:%ds\nTime taken for this subject: %dh:%dm:%ds',floor(toc(tc0)/3600),floor(mod(toc(tc0),3600)/60),floor(mod(toc(tc0),60)),floor(toc(tc)/3600),floor(mod(toc(tc),3600)/60),floor(mod(toc(tc),60))); fprintf(fileID,'%s\n',logg{L,1}); try set(S.tx,'string',logg); catch, end; clear clk
    end; clear k2
        
    refresh_html
    
end; clear k

end

%%
%% '23. Simple stats'
if choices_postproc(4)==1
L=L+2; logg{L,1}=sprintf('-------\n%s',postprocsteps{4}); fprintf(fileID,'%s\n',logg{L,1}); try set(S.tx,'string',logg); catch, end; tc=tic;
time_taken_postproc.step23_stats = zeros(size(func_files,1),size(func_files,2));

indx_group = sort(unique(union(indx_group1,indx_group2)));
f=0;
for k=1:length(sub_dir)
    if ismember(k,indx_group)
    for k2=1:runs(k)
        sdirStats = cat(1, dir([sub_dir{k} func_files{k,k2} '*SFC*' '.mat']), dir([sub_dir{k} func_files{k,k2} '*fALFF*' '.mat']));
        if isempty(sdirStats)
            f=f+1;
            missingdata(f,1) = [k,k2]; %#ok<*SAGROW>
        end
        clear sdirStats
    end
    end
end
if exist('missingdata','var')
    overlap_str = '';
    for w=1:size(missingdata,1)
        tmp_suffix = sub_dir{missingdata(w,1)}(max(strfind(sub_dir{missingdata(w,1)}(1:end-1),'/'))+1:end-1);
        if isempty(tmp_suffix)
            tmp_suffix = sub_dir{missingdata(w,1)}(max(strfind(sub_dir{missingdata(w,1)}(1:end-1),'\'))+1:end-1);
        end
        overlap_str = strcat(overlap_str,sprintf('\n%s / %s',tmp_suffix,func_files{missingdata(w,1),missingdata(w,2)}));
        clear tmp_suffix
    end
    msgbox(sprintf('Error: some of the subjects (mentioned below) do not have connectivity or fALFF data needed for stats. Aborting.\n%s\n',overlap_str),'Abort','error');
    clk=clock; L=L+1; logg{L,1}=sprintf('%d/%s/%d %dh:%dm:%ds - (23) simple stats SKIPPED. Aborting.',clk(3),month{clk(2)},clk(1),clk(4),clk(5),round(clk(6))); fprintf(fileID,'%s\n',logg{L,1}); try set(S.tx,'string',logg); catch, end; clear clk
    return;
end
clear f indx_group

clk=clock; L=L+1; logg{L,1}=sprintf('%d/%s/%d %dh:%dm:%ds - (23) simple stats STARTED',clk(3),month{clk(2)},clk(1),clk(4),clk(5),round(clk(6))); fprintf(fileID,'%s\n',logg{L,1}); try set(S.tx,'string',logg); catch, end; clear clk
try waitbar(0.5,wbar3,sprintf('23. Simple statistical analysis')); catch, end
fprintf('23. Performing simple statistical analysis\n')
if ~exist('SFC_type','var'), SFC_type=[]; end
if ~exist('SFC_withinslice_flag','var'), SFC_withinslice_flag=[]; end
if ~exist('SFC_betweenslice_flag','var'), SFC_betweenslice_flag=[]; end
time_23_stats = tic;
scfMRItb_23_stats1(sub_dir,func_files,postproc_filesuffix,choices_postproc, indx_group1,indx_group2,ttest_type,alpha_stats, WM_flag,slices,SFC_type,SFC_withinslice_flag,SFC_betweenslice_flag, use_deconv_data_for_postproc,dont_do_pcor_betweenslice);
time_taken_postproc.step23_stats = toc(time_23_stats); clear time_23_stats
clk=clock; L=L+1; logg{L,1}=sprintf('%d/%s/%d %dh:%dm:%ds - (23) simple stats COMPLETED',clk(3),month{clk(2)},clk(1),clk(4),clk(5),round(clk(6))); fprintf(fileID,'%s\n',logg{L,1}); try set(S.tx,'string',logg); catch, end; clear clk
L=L+1; logg{L,1}=sprintf('Total time taken so far: %dh:%dm:%ds\nTime taken for this step: %dh:%dm:%ds',floor(toc(tc0)/3600),floor(mod(toc(tc0),3600)/60),floor(mod(toc(tc0),60)),floor(toc(tc)/3600),floor(mod(toc(tc),3600)/60),floor(mod(toc(tc),60))); fprintf(fileID,'%s\n',logg{L,1}); try set(S.tx,'string',logg); catch, end; clear clk

end

%%
%%
try
if exist([home_dir filename_ExecutionTime],'file')
save([home_dir filename_ExecutionTime],'time_taken_postproc','-append')
end
catch
end

L=L+2; logg{L,1}=sprintf('--- Post-processing Completed ---'); fprintf(fileID,'%s\n',logg{L,1}); try set(S.tx,'string',logg); catch, end; clear clk
try close(wbar3), catch, end
refresh_html

quest = ['\color[rgb]{0.65,0.11,0.19}' sprintf('Dear User,\n\nAll post-processing steps were COMPLETED as promised.\n\nRegards,\nSpinal cord fMRI toolbox (NEPTUNE)\n')]; %#ok<*NASGU>
opts.Interpreter = 'tex'; opts.WindowStyle = 'non-modal';
msgbox(quest,'All post-processing done!','custom',msgicon,[1,1,1],opts); clear quest opts
clear wbar3

%%
```
