## Supplementary material for "Neptune: a toolbox for spinal cord functional MRI data processing and quality assurance": Neptune software: postprocdialog.m

```
function [selection,choices] = postprocdialog(pnl,bg,prevchoices,flag)

% cbx1 = uicheckbox(pnl,'Position',[10 434 410 15], 'Value',1);
cbx2 = uicheckbox(pnl,'Position',[10 82 410 15], 'Value',prevchoices(1));
cbx3 = uicheckbox(pnl,'Position',[10 60 410 15], 'Value',prevchoices(2));
cbx4 = uicheckbox(pnl,'Position',[10 38 410 15], 'Value',prevchoices(3));
cbx5 = uicheckbox(pnl,'Position',[10 16 410 15], 'Value',prevchoices(4));

rb1 = uiradiobutton(bg,'Position',[400 10 410 15]);
rb2 = uiradiobutton(bg,'Position',[200 10 410 15]);
rb3 = uiradiobutton(bg,'Position',[400 10 410 15]);

cbx2.Text = '01. Extract quadrant time series'' (mandatory)';
cbx3.Text = '02. Estimate static functional connectivity (SFC)';
cbx4.Text = '03. Estimate fractional amplitude of low frequency fluctuations (fALFF)';
cbx5.Text = '04. Simple stats';

rb1.Text = '';
rb2.Text = 'DONE';
rb3.Text = 'Help';

pause(2)
choices = [cbx2.Value;cbx3.Value;cbx4.Value;cbx5.Value];

selection = bg.SelectedObject.Parent.SelectedObject.Text;

if flag==1 && strcmp(selection,'Help')==1
    selection='';
end

end
```
