## Supplementary material for "Neptune: a toolbox for spinal cord functional MRI data processing and quality assurance": Neptune software: preproc_choices.m

```
% Pre-processing choices

%% --- non-GUI choices before proceeding ---

if choices_preproc(7)==1
if ~(exist('automatic_GaussianMask', 'var'))
    % by default, derive the Gaussian mask automatically from the not-cord mask. A value of 0 implies that the user will manually define the Gaussian mask.
    automatic_GaussianMask = 1; % default = 1
end
end

if (choices_preproc(10)==1)||(choices_preproc(18)==1)
if ~(exist('SCT_vert_auto', 'var'))
    % 1 = assume that we will perform automatic labeling of vertebrae during PAM50 registration
    SCT_vert_auto = 1;
end
if ~(exist('SCT_manualreg_labels', 'var'))
    % Vertebrae for manual labeling in SCT (if manual option was chosen), needed for registration
    SCT_manualreg_labels = '3,5';
end
end

if choices_preproc(17)==1
if ~(exist('run_on_HPF_data', 'var'))
    % by default, use high-pass filtered data to perform deconvolution. A value of 0 will perform deconvolution on band-pass filtered data
    run_on_HPF_data = 1; % default = 1
end
end

if ~exist('remove_slicewise_files','var')
    % Slice-wise files are combined to 3D+time data in step-14. After that it is safe to delete the slice-wise data (they can be regenerated later if needed). This is enabled by default. Set this to 0 if you don't want this be performed (unnecessary wastage of disk space, though).
    remove_slicewise_files = 1; % default = 1
end

if ~exist('run_allsubs_by_default','var')
    % By default, processing steps will only run on subjects in which they have not been executed before. However, if you want to run all processing steps on all subjects then set this to 1 (which will overwrite existing files).
    run_allsubs_by_default = 0; % default = 0
end

%% --- GUI-based choices before proceeding ---

if ~exist('ignore_extreme_slices','var')
    % 1 = removes first and last slice. 0 = uses all slices.
    quest = sprintf('Discard the first and last slice?\n(extreme slices are bad in some acquisitions)');
    answer_ignore_extreme_slices = questdlg(quest,'Ignore extreme slices','Yes, discard','No, use all slices','Yes, discard'); clear quest
    switch answer_ignore_extreme_slices
        case 'Yes, discard'
            ignore_extreme_slices = 1;
        case 'No, use all slices'
            ignore_extreme_slices = 0;
    end
end

if ~exist('discard_firstN_volumes','var')
    % By default, do not discard the first N volumes. If you want to discard the first N volumes then set the value of N to a non-zero value here
    prompt = sprintf('Do you want to discard the first N functional volumes?\n\n0 = do not discard.\nN = discard the first N volumes (N must be a non-zero positive integer) (typical value = 5)\n');
    answer3 = inputdlg(prompt,'Discard initial func volumes',1,{'0'}); clear prompt
    discard_firstN_volumes = str2num(cell2mat(answer3)); clear answer3
end

if choices_preproc(1)==1
if ~exist('NORDIC_type','var')
    quest = sprintf('Perform NORDIC using magnitude-only data OR magnitude+phase data?\n\nIf magnitude+phase, the phase data must be named as "<filename>_ph.nii" (or .nii.gz)');
    answer_NORDIC = questdlg(quest,'NORDIC','magnitude-only','magnitude+phase','magnitude-only'); clear quest
    switch answer_NORDIC
        case 'magnitude-only'
            NORDIC_type = 1;
        case 'magnitude+phase'
            NORDIC_type = 2;
    end
end
end

if choices_preproc(2)==1
if ~exist('sliceorder','var')
    prompt = sprintf('Please provide slice order for slice-timing correction (separated by spaces)');
    answer_sliceorder = inputdlg(prompt,'Slice-timing correction',1,{'1 3 5 7 9 11 2 4 6 8 10 12'}); clear prompt
    sliceorder = str2num(cell2mat(answer2));
end
end

if ~isempty(which('VideoWriter')) % check if Matlab's audio video toolbox exists
    if sum(choices_preproc([5,6,8,9,11,12,13,14]))~=0 % check if steps that do save videos have been chosen
        if ~(exist('save_videos', 'var'))
            prompt = sprintf('Quality Control (QC) videos take up considerable computational time and memory, but will provide useful QC data.\n\nDo you want to save QC videos? Three options:\n1. Yes, save for ALL steps\n2. Yes, save only at the end of all denoising (also shows raw data)\n3. No, do not save videos.\n');
            answer_videos = questdlg(prompt,'Save QC videos?','Yes, for all steps','Yes, only at the end','No','No'); clear prompt
            switch answer_videos
                case 'Yes, for all steps'
                    save_videos = 1; % save QC videos (generates sizeable .mp4 files and takes a few minutes to do it).
                    save_videos_steps = ones(8,1);
                case 'Yes, only at the end'
                    save_videos = 2; % save QC videos (generates sizeable .mp4 files and takes a few minutes to do it).
                    save_videos_steps = zeros(8,1);
                    if     choices_preproc(14)==1, save_videos_steps(8)=1;
                    elseif choices_preproc(13)==1, save_videos_steps(7)=1;
                    elseif choices_preproc(12)==1, save_videos_steps(6)=1;
                    elseif choices_preproc(11)==1, save_videos_steps(5)=1;
                    elseif choices_preproc(9)==1,  save_videos_steps(4)=1;
                    elseif choices_preproc(8)==1,  save_videos_steps(3)=1;
                    elseif choices_preproc(6)==1,  save_videos_steps(2)=1;
                    elseif choices_preproc(5)==1,  save_videos_steps(1)=1;
                    end
                case 'No'
                    save_videos = 0; % don't save QC videos.
                    save_videos_steps = zeros(8,1);
            end
        end
    else
        save_videos = 0;
    end
else
    save_videos = 0; % don't save QC videos if Matlab's audio video toolbox does not exist
end

if choices_preproc(8)==1
if ~(exist('motion_param_smoothing', 'var'))
    % smoothing of estimated motion parameters to suppress sporadic artifacts
    % 0 = do not smooth; any other positive integer N = smooth using an N-point median filter (typical value = 5)
    prompt = sprintf('Perform smoothing of estimated motion parameters to suppress spiky sporadic artifacts?\n\n0 = do not smooth.\nN = smooth using an N-point median filter (N must be a non-zero positive integer) (typical value = 5)\n');
    answer3 = inputdlg(prompt,'Motion parameter smoothing',1,{'0'}); clear prompt
    motion_param_smoothing = str2num(cell2mat(answer3)); clear answer3
end
end

if choices_preproc(10)==1
if ~exist('mask_GM_flag','var')
    if exist_SCT==1
    quest = sprintf('Obtaining anatomical gray matter mask:\n\nEstimate GM boundary using SCT deepseg?\nOR\nGet GM boundary by co-registration to PAM50 space and transforming the standard GM mask in SCT?\n');
    answer_masks = questdlg(quest,'GM mask','Use SCT deepseg','Do PAM50 co-registration','Use SCT deepseg'); clear quest
    switch answer_masks
        case 'Use SCT deepseg'
            mask_GM_flag = 1;
        case 'Do PAM50 co-registration'
            mask_GM_flag = 2;
    end
    else
        mask_GM_flag = 1;
    end
end
end


if choices_preproc(14)==1
if ~exist('covregress_options','var')

fig12 = uifigure('Position',[300 300 1000 300],'Color',[0.95,0.875,0.8]); drawnow; fig12.Visible='off'; fig12.Visible='on';
pnl = uipanel(fig12,'Position',[50 80 900 210],'BackgroundColor',[0.95,0.9,0.85],'BorderType','none');
pnl.Title = sprintf('\nDenoise-5 (additional covariates regression): Make one choice for each row and click DONE.\n\n');
pnl.TitlePosition = 'centertop'; pnl.ForegroundColor = [0.65,0.11,0.19]; pnl.FontSize = 13; pnl.FontWeight = 'bold';

bg1 = uibuttongroup(fig12,'Position',[50 200 900 50],'BackgroundColor',[0.95,0.9,0.85],'BorderType','line');
bg2 = uibuttongroup(fig12,'Position',[50 150 900 50],'BackgroundColor',[0.95,0.9,0.85],'BorderType','line');
bg3 = uibuttongroup(fig12,'Position',[50 100 900 50],'BackgroundColor',[0.95,0.9,0.85],'BorderType','line');
bg4 = uibuttongroup(fig12,'Position',[50  30 900 50],'BackgroundColor',[0.95,0.9,0.85],'BorderType','line');

selection1='Motion parameters (2 translation + 2 derivatives)'; selection2='NO scrubbing using motion'; selection3='NO user-defined covariates';
selection4=''; flag=0;
while strcmp(selection3,'DONE')~=1
    [selection1,selection2,selection3,selection4] = regresscovdialog(bg1,bg2,bg3,bg4,flag,selection1,selection2,selection3,selection4);
    if strcmp(selection4,'DONE')==1
        break
    end
    if strcmp(selection4,'Help')==1
        fprintf('neptune_manual.pdf\n')
        open('neptune_manual.pdf')
        flag=1;
    end
end
close(fig12)
clear fig12 bg1 bg2 bg3 bg4 flag selection4 pnl

switch selection1
    case 'Motion parameters (2 translation)'
        covregress_options(1,1) = 1;
    case 'Motion parameters (2 translation + 2 derivatives)'
        covregress_options(1,1) = 2;
    case 'NO motion parameters'
        covregress_options(1,1) = 3;
end
switch selection2
    case 'Scrubbing (volumes with motion > half voxel size)'
        covregress_options(2,1) = 1;
    case 'Scrubbing (user-defined motion threshold)'
        covregress_options(2,1) = 2;
    case 'NO scrubbing using motion'
        covregress_options(2,1) = 3;
end
switch selection3
    case 'Regression with user-defined covariate(s)'
        covregress_options(3,1) = 1;
    case 'NO user-defined covariates'
        covregress_options(3,1) = 2;
end

if covregress_options(2,1)==1
    covregress_options(2,2) = voxel_size_inplane_func/2;
elseif covregress_options(2,1)==2
    prompt = sprintf('Please provide motion threshold in millimeters for Scrubbing (user-defined motion threshold)');
    answer3 = inputdlg(prompt,'Scrubbing (motion threshold)',1,{'1.0'}); clear prompt
    covregress_options(2,2) = str2num(cell2mat(answer3)); clear answer3 %#ok<*ST2NM>
end
if covregress_options(3,1)==1
    cov_files = get_covariate_files(sub_dir,func_files,runs);
else
    cov_files = cell(size(func_files,1),size(func_files,2));
end

end % if ~exist('covregress_options','var')
end % if choices_preproc(14)==1


if (choices_preproc(15)==1)||(choices_preproc(17)==1)
if ~exist('bpf_hz','var')
    prompt = sprintf('Please provide upper cutoff frequency for temporal filtering (in Hertz) (commonly used value = 0.1 Hz)');
    answer3 = inputdlg(prompt,'Temporal filtering',1,{'0.1'}); clear prompt
    bpf_hz = str2num(cell2mat(answer3)); clear answer3
    if bpf_hz(1)<0.1
        bpf_hz_str = sprintf('0%.0f',bpf_hz(1)*100);
    else
        bpf_hz_str = sprintf('%.0f',bpf_hz(1)*100);
    end
end
if ~exist('detrend_flag','var')
    quest = sprintf('Do you wish to perform linear or quadratic detrending before filtering?\nIf you are unsure, choose ''No''.');
    answer_detrend = questdlg(quest,'Detrending','Linear detrending','Quadratic detrending','NO detrending','NO detrending'); clear quest
    switch answer_detrend
        case 'Linear detrending'
            detrend_flag = 1;
        case 'Quadratic detrending'
            detrend_flag = 2;
        case 'NO detrending'
            detrend_flag = 0;
    end
end
end

if ~exist('answer_gzip','var')
if choices_preproc(19)==1
    quest = sprintf('Do you wish to DELETE the .nii files AFTER saving the compressed .nii.gz files?');
    answer_gzip = questdlg(quest,'gzip NIfTI files','gzip only, don''t delete','delete only, I already gzipped','both gzip and delete','both gzip and delete'); clear quest
end
end

if step_after_step==1
if ~exist('UIstepsFirst','var')
    quest = sprintf('Do you wish to carry out all steps requiring user input first?\nThis will first do steps 1-5, 10 and 16 (if chosen) and post-processing choices (if selected), and then do the rest unto completion.');
    answer_UIstepsFirst = questdlg(quest,'Do UI steps first','Yes','No','Yes'); clear quest
    switch answer_UIstepsFirst
        case 'Yes'
            UIstepsFirst = 1;
        case 'No'
            UIstepsFirst = 0;
    end
end
else
    UIstepsFirst = -1; answer_UIstepsFirst = 'N/A';
end

%%
```
