## Supplementary material for "Neptune: a toolbox for spinal cord functional MRI data processing and quality assurance": Neptune software: preproc_scfMRItb.m

```
% PREPROCESSING
try
if ~exist('L','var'), L=0; end
L=L+2; logg{L,1}=sprintf('--- Pre-processing ---'); fprintf(fileID,'%s\n',logg{L,1}); try set(S.tx,'string',logg); catch, end; clear clk
clk=clock; L=L+1; logg{L,1}=sprintf('%d/%s/%d %dh:%dm:%ds - begin making pre-processing choices',clk(3),month{clk(2)},clk(1),clk(4),clk(5),round(clk(6))); fprintf(fileID,'%s\n',logg{L,1}); try set(S.tx,'string',logg); catch, end; clear clk
catch
end
tc0=tic;

%% --- Initialize pre-processing status figure ---
N=20;
status_fig = figure('Name','Status Bar','NumberTitle','off','units','normalized','outerposition',[2/3 0 1/3 1]); % [position_along_x_from_bottomleft position_along_y_from_bottomleft size_x size_y] in normalized units (0,1)
subplot = @(m,n,p) subtightplot(m, n, p, [0.001 0.005], [0.01 0.001], [0.01 0.001]);
figure(status_fig),
subplot(N,2,1),imshow(imread('status_images/step_01_white.jpeg')), subplot(N,2,2),imshow(imread('status_images/white.jpg'))
subplot(N,2,3),imshow(imread('status_images/step_02_white.jpeg')), subplot(N,2,4),imshow(imread('status_images/white.jpg')),
subplot(N,2,5),imshow(imread('status_images/step_03_white.jpeg')), subplot(N,2,6),imshow(imread('status_images/white.jpg')),
subplot(N,2,7),imshow(imread('status_images/step_04_white.jpeg')), subplot(N,2,8),imshow(imread('status_images/white.jpg')),
subplot(N,2,9),imshow(imread('status_images/step_05_white.jpeg')), subplot(N,2,10),imshow(imread('status_images/white.jpg')),
subplot(N,2,11),imshow(imread('status_images/step_06_white.jpeg')), subplot(N,2,12),imshow(imread('status_images/white.jpg')),
subplot(N,2,13),imshow(imread('status_images/step_07_white.jpeg')), subplot(N,2,14),imshow(imread('status_images/white.jpg')),
subplot(N,2,15),imshow(imread('status_images/step_08_white.jpeg')), subplot(N,2,16),imshow(imread('status_images/white.jpg')),
subplot(N,2,17),imshow(imread('status_images/step_09_white.jpeg')), subplot(N,2,18),imshow(imread('status_images/white.jpg')),
subplot(N,2,19),imshow(imread('status_images/step_10_white.jpeg')), subplot(N,2,20),imshow(imread('status_images/white.jpg')),
subplot(N,2,21),imshow(imread('status_images/step_11_white.jpeg')), subplot(N,2,22),imshow(imread('status_images/white.jpg')),
subplot(N,2,23),imshow(imread('status_images/step_12_white.jpeg')), subplot(N,2,24),imshow(imread('status_images/white.jpg')),
subplot(N,2,25),imshow(imread('status_images/step_13_white.jpeg')), subplot(N,2,26),imshow(imread('status_images/white.jpg')),
subplot(N,2,27),imshow(imread('status_images/step_14_white.jpeg')), subplot(N,2,28),imshow(imread('status_images/white.jpg')),
subplot(N,2,29),imshow(imread('status_images/step_15_white.jpeg')), subplot(N,2,30),imshow(imread('status_images/white.jpg')),
subplot(N,2,31),imshow(imread('status_images/step_16_white.jpeg')), subplot(N,2,32),imshow(imread('status_images/white.jpg')),
subplot(N,2,33),imshow(imread('status_images/step_17_white.jpeg')), subplot(N,2,34),imshow(imread('status_images/white.jpg')),
subplot(N,2,35),imshow(imread('status_images/step_18_white.jpeg')), subplot(N,2,36),imshow(imread('status_images/white.jpg')),
subplot(N,2,37),imshow(imread('status_images/step_19_white.jpeg')), subplot(N,2,38),imshow(imread('status_images/white.jpg')),
subplot(N,2,39),imshow(imread('status_images/white.jpg')), subplot(N,2,40),imshow(imread('status_images/white.jpg')),

wbar2 = waitbar(0,'Choose pre-processing steps...','Name','Pre-processing status...','Units','normalized','Position',[0 13/17 1/5 1/17]); % [4/5 0 1/5 1/17] [3/5 3/17 1/5 1/17]
wbar3 = waitbar(0,'Progress within each pre-processing step...','Name','Progress within each pre-processing step...','Units','normalized','Position',[0 11/17 1/5 1/17]); % [] [3/5 0 1/5 1/17]
if ~exist('clk_settings','var'), tmp=clock; clk_settings = sprintf('%.4d-%.2d-%.2d_%.2d-%.2d',tmp(1),tmp(2),tmp(3),tmp(4),tmp(5)); clear tmp; end

%% --- choose pre-processing steps ---
if ~exist('choices_preproc','var')

fig = uifigure('Position',[680 740 530 590],'Color',[0.95,0.875,0.8]); drawnow; fig.Visible='off'; fig.Visible='on';
pnl = uipanel(fig,'Position',[30 60 470 500],'BackgroundColor',[0.95,0.9,0.85],'BorderType','none');
pnl.Title = sprintf('Choose your pre-processing steps and click DONE.\n\nKindly note that steps [04, 05, 07, 08, 10, 11] are mandatory.\n''Help'' is available if required.\n');
pnl.TitlePosition = 'centertop'; pnl.ForegroundColor = [0.65,0.11,0.19];  % [0,0,0.8];
bg = uibuttongroup(fig,'Position',[30 30 470 30],'BackgroundColor',[0.95,0.875,0.8],'BorderType','none');

qSPM = which('spm_get_bf.m');
selection=''; flag=0;
if rest==1 && ~isempty(qSPM)
    choices_preproc=[1;0;1;1;1;1;1;1;1;1;1;1;1;1;1;1;1;0;1];
elseif task==1 || isempty(qSPM)
    choices_preproc=[1;0;1;1;1;1;1;1;1;1;1;1;1;1;1;1;0;0;1];
end
cbx4  = uicheckbox(pnl,'Position',[10 346 410 15], 'Enable','off', 'Value',choices_preproc(4));
cbx4.Text  = '04. Split data into slices (mandatory)';
if isempty(qSPM)
cbx17 = uicheckbox(pnl,'Position',[10  60 410 15], 'Enable','off', 'Value',choices_preproc(17));
cbx17.Text = '17. HRF deconvolution (requires SPM, which is not installed)';
end
if exist_SCT==0
cbx18 = uicheckbox(pnl,'Position',[10  38 410 15], 'Enable','off', 'Value',choices_preproc(18));
cbx18.Text = '18. Co-registration to PAM50 standard space (requires SCT)';
end

while strcmp(selection,'DONE')~=1
    [selection,choices_preproc] = preprocdialog(pnl,bg,choices_preproc,flag,exist_SCT);
    if strcmp(selection,'DONE')==1
%         closereq()
        break
    end
    if strcmp(selection,'Help')==1
        fprintf('neptune_manual.pdf\n')
        open('neptune_manual.pdf')
        flag=1;
    end
end
close(fig)
clear fig bg pnl flag answer1 cbx1 cbx4 cbx18 qSPM

end

%% --- update status figure if it is open---
if ishandle(status_fig)==1
try

subplot = @(m,n,p) subtightplot(m, n, p, [0.001 0.005], [0.01 0.001], [0.01 0.001]);
figure(status_fig)
if choices_preproc(1)==1
    subplot(N,2,1),imshow(imread('status_images/step_01_white.jpeg')), subplot(N,2,2),imshow(imread('status_images/status_white.jpg')),
else
    subplot(N,2,1),imshow(imread('status_images/step_01_gray.jpeg')), subplot(N,2,2),imshow(imread('status_images/status_gray.jpg')),
end
if choices_preproc(2)==1
    subplot(N,2,3),imshow(imread('status_images/step_02_white.jpeg')), subplot(N,2,4),imshow(imread('status_images/status_white_UI.jpg')),
else
    subplot(N,2,3),imshow(imread('status_images/step_02_gray.jpeg')), subplot(N,2,4),imshow(imread('status_images/status_gray.jpg')),
end
if choices_preproc(3)==1
    subplot(N,2,5),imshow(imread('status_images/step_03_white.jpeg')), subplot(N,2,6),imshow(imread('status_images/status_white.jpg')),
else
    subplot(N,2,5),imshow(imread('status_images/step_03_gray.jpeg')), subplot(N,2,6),imshow(imread('status_images/status_gray.jpg')),
end
if choices_preproc(4)==1
    subplot(N,2,7),imshow(imread('status_images/step_04_white.jpeg')), subplot(N,2,8),imshow(imread('status_images/status_white.jpg')),
else
    subplot(N,2,7),imshow(imread('status_images/step_04_gray.jpeg')), subplot(N,2,8),imshow(imread('status_images/status_gray.jpg')),
end
if choices_preproc(5)==1
    subplot(N,2,9),imshow(imread('status_images/step_05_white.jpeg')), subplot(N,2,10),imshow(imread('status_images/status_white_UI.jpg')),
else
    subplot(N,2,9),imshow(imread('status_images/step_05_gray.jpeg')), subplot(N,2,10),imshow(imread('status_images/status_gray.jpg')),
end
if choices_preproc(6)==1
    subplot(N,2,11),imshow(imread('status_images/step_06_white.jpeg')), subplot(N,2,12),imshow(imread('status_images/status_white.jpg')),
else
    subplot(N,2,11),imshow(imread('status_images/step_06_gray.jpeg')), subplot(N,2,12),imshow(imread('status_images/status_gray.jpg')),
end
if choices_preproc(7)==1
    subplot(N,2,13),imshow(imread('status_images/step_07_white.jpeg')), subplot(N,2,14),imshow(imread('status_images/status_white.jpg')),
else
    subplot(N,2,13),imshow(imread('status_images/step_07_gray.jpeg')), subplot(N,2,14),imshow(imread('status_images/status_gray.jpg')),
end
if choices_preproc(8)==1
    subplot(N,2,15),imshow(imread('status_images/step_08_white.jpeg')), subplot(N,2,16),imshow(imread('status_images/status_white.jpg')),
else
    subplot(N,2,15),imshow(imread('status_images/step_08_gray.jpeg')), subplot(N,2,16),imshow(imread('status_images/status_gray.jpg')),
end
if choices_preproc(9)==1
    subplot(N,2,17),imshow(imread('status_images/step_09_white.jpeg')), subplot(N,2,18),imshow(imread('status_images/status_white.jpg')),
else
    subplot(N,2,17),imshow(imread('status_images/step_09_gray.jpeg')), subplot(N,2,18),imshow(imread('status_images/status_gray.jpg')),
end
if choices_preproc(10)==1
    subplot(N,2,19),imshow(imread('status_images/step_10_white.jpeg')), subplot(N,2,20),imshow(imread('status_images/status_white_UI.jpg')),
else
    subplot(N,2,19),imshow(imread('status_images/step_10_gray.jpeg')), subplot(N,2,20),imshow(imread('status_images/status_gray.jpg')),
end
if choices_preproc(11)==1
    subplot(N,2,21),imshow(imread('status_images/step_11_white.jpeg')), subplot(N,2,22),imshow(imread('status_images/status_white.jpg')),
else
    subplot(N,2,21),imshow(imread('status_images/step_11_gray.jpeg')), subplot(N,2,22),imshow(imread('status_images/status_gray.jpg')),
end
if choices_preproc(12)==1
    subplot(N,2,23),imshow(imread('status_images/step_12_white.jpeg')), subplot(N,2,24),imshow(imread('status_images/status_white.jpg')),
else
    subplot(N,2,23),imshow(imread('status_images/step_12_gray.jpeg')), subplot(N,2,24),imshow(imread('status_images/status_gray.jpg')),
end
if choices_preproc(13)==1
    subplot(N,2,25),imshow(imread('status_images/step_13_white.jpeg')), subplot(N,2,26),imshow(imread('status_images/status_white.jpg')),
else
    subplot(N,2,25),imshow(imread('status_images/step_13_gray.jpeg')), subplot(N,2,26),imshow(imread('status_images/status_gray.jpg')),
end
if choices_preproc(14)==1
    subplot(N,2,27),imshow(imread('status_images/step_14_white.jpeg')), subplot(N,2,28),imshow(imread('status_images/status_white.jpg')),
else
    subplot(N,2,27),imshow(imread('status_images/step_14_gray.jpeg')), subplot(N,2,28),imshow(imread('status_images/status_gray.jpg')),
end
if choices_preproc(15)==1
    subplot(N,2,29),imshow(imread('status_images/step_15_white.jpeg')), subplot(N,2,30),imshow(imread('status_images/status_white.jpg')),
else
    subplot(N,2,29),imshow(imread('status_images/step_15_gray.jpeg')), subplot(N,2,30),imshow(imread('status_images/status_gray.jpg')),
end
if choices_preproc(16)==1
    subplot(N,2,31),imshow(imread('status_images/step_16_white.jpeg')), subplot(N,2,32),imshow(imread('status_images/status_white_UI.jpg')),
else
    subplot(N,2,31),imshow(imread('status_images/step_16_gray.jpeg')), subplot(N,2,32),imshow(imread('status_images/status_gray.jpg')),
end
if choices_preproc(17)==1
    subplot(N,2,33),imshow(imread('status_images/step_17_white.jpeg')), subplot(N,2,34),imshow(imread('status_images/status_white.jpg')),
else
    subplot(N,2,33),imshow(imread('status_images/step_17_gray.jpeg')), subplot(N,2,34),imshow(imread('status_images/status_gray.jpg')),
end
if choices_preproc(18)==1
    subplot(N,2,35),imshow(imread('status_images/step_18_white.jpeg')), subplot(N,2,36),imshow(imread('status_images/status_white.jpg')),
else
    subplot(N,2,35),imshow(imread('status_images/step_18_gray.jpeg')), subplot(N,2,36),imshow(imread('status_images/status_gray.jpg')),
end
if choices_preproc(19)==1
    subplot(N,2,37),imshow(imread('status_images/step_19_white.jpeg')), subplot(N,2,38),imshow(imread('status_images/status_white.jpg')),
else
    subplot(N,2,37),imshow(imread('status_images/step_19_gray.jpeg')), subplot(N,2,38),imshow(imread('status_images/status_gray.jpg')),
end

catch
end
end

preprocsteps = {'01. NORDIC denoising';'02. Slice-timing correction';'03. Match anatomical and functional slices';'04. Split data into slices';'05. Define "not cord" mask';'06. Denoise-1 (non-cord regression)';'07. Compute Gaussian mask';'08. Motion correction';'09. Denoise-2 (RETROICOR/3dDespike)';'10. Define GM, WM and CSF masks';'11. Co-registration (functional to anat.)';'12. Denoise-3 (CSF regression)';'13. Denoise-4 (white-matter signal regression from GM)';'14. Denoise-5 (additional covariates regression)';'15. Temporal filtering';'16. Define cord GM and WM quadrants';'17. HRF deconvolution';'18. Co-registration to PAM50 standard space';'19. gzip NIfTI files to save disk space'};

%% --- Pre-processing choices ---

preproc_choices


%% Update the log with newly made choices

try
L=L+2; logg{L,1}=sprintf('Pre-processing steps CHOSEN:'); fprintf(fileID,'%s\n',logg{L,1}); try set(S.tx,'string',logg); catch, end
temp=preprocsteps(find(choices_preproc==1));
for k=1:length(temp)
    L=L+1; logg{L,1}=sprintf('%s',temp{k}); fprintf(fileID,'%s\n',logg{L,1}); try set(S.tx,'string',logg); catch, end
end; clear k temp
L=L+2; logg{L,1}=sprintf('Pre-processing steps OMITTED:'); fprintf(fileID,'%s\n',logg{L,1}); try set(S.tx,'string',logg); catch, end
temp=preprocsteps(find(choices_preproc==0));
for k=1:length(temp)
    L=L+1; logg{L,1}=sprintf('%s',temp{k}); fprintf(fileID,'%s\n',logg{L,1}); try set(S.tx,'string',logg); catch, end
end; clear k temp

L=L+2; logg{L,1}=sprintf('Pre-processing control parameters:'); fprintf(fileID,'%s\n',logg{L,1}); try set(S.tx,'string',logg); catch, end
if choices_preproc(1)==1
    L=L+1; logg{L,1}=sprintf('NORDIC type = %s',answer_NORDIC); fprintf(fileID,'%s\n',logg{L,1}); try set(S.tx,'string',logg); catch, end
end
if choices_preproc(2)==1
    L=L+1; logg{L,1}=sprintf('Slice-timing correction - slice order = %s',answer_sliceorder); fprintf(fileID,'%s\n',logg{L,1}); try set(S.tx,'string',logg); catch, end
end
if sum(choices_preproc([5,6,8,9,11,12,13,14]))~=0
    L=L+1; logg{L,1}=sprintf('Save QC videos: %s',answer_videos); fprintf(fileID,'%s\n',logg{L,1}); try set(S.tx,'string',logg); catch, end
end
if choices_preproc(7)==1
    L=L+1; logg{L,1}=sprintf('Gaussian mask - automatic: %d',automatic_GaussianMask); fprintf(fileID,'%s\n',logg{L,1}); try set(S.tx,'string',logg); catch, end
end
if choices_preproc(8)==1
    L=L+1; logg{L,1}=sprintf('Smoothing of estimated motion parameters: %d',motion_param_smoothing); fprintf(fileID,'%s\n',logg{L,1}); try set(S.tx,'string',logg); catch, end
end
if choices_preproc(10)==1
    L=L+1; logg{L,1}=sprintf('GM mask: %s',answer_masks); fprintf(fileID,'%s\n',logg{L,1}); try set(S.tx,'string',logg); catch, end
end
if choices_preproc(14)==1
    L=L+1; logg{L,1}=sprintf('Additional covariates regression choices: (1) %s, (2) %s, (3) %s',selection1,selection2,selection3); fprintf(fileID,'%s\n',logg{L,1}); try set(S.tx,'string',logg); catch, end
    if covregress_options(2,1)==2
        L=L+1; logg{L,1}=sprintf('Additional covariates regression: user-defined motion threshold for scrubbing = %d millimeters',covregress_options(2,2)); fprintf(fileID,'%s\n',logg{L,1}); try set(S.tx,'string',logg); catch, end
    end
end
if choices_preproc(15)==1
    L=L+1; logg{L,1}=sprintf('Temporal band-pass filtering upper cut-off frequency = %.2f Hz (the lower cut-off is 0.01 Hz)',bpf_hz); fprintf(fileID,'%s\n',logg{L,1}); try set(S.tx,'string',logg); catch, end
end
if choices_preproc(17)==1
    L=L+1; logg{L,1}=sprintf('Deconvolution: run_on_HPF_data = %d',run_on_HPF_data); fprintf(fileID,'%s\n',logg{L,1}); try set(S.tx,'string',logg); catch, end
end
if choices_preproc(19)==1
    L=L+1; logg{L,1}=sprintf('Gzip Nifti files and/or delete Nifti files after that = %s',answer_gzip); fprintf(fileID,'%s\n',logg{L,1}); try set(S.tx,'string',logg); catch, end
end
L=L+1; logg{L,1}=sprintf('Ignore first and last slice = %d',ignore_extreme_slices); fprintf(fileID,'%s\n',logg{L,1}); try set(S.tx,'string',logg); catch, end
L=L+1; logg{L,1}=sprintf('Discard the first %d volumes',discard_firstN_volumes); fprintf(fileID,'%s\n',logg{L,1}); try set(S.tx,'string',logg); catch, end
L=L+1; logg{L,1}=sprintf('Remove slice-wise files after saving 3D+time data = %d',remove_slicewise_files); fprintf(fileID,'%s\n',logg{L,1}); try set(S.tx,'string',logg); catch, end
L=L+1; logg{L,1}=sprintf('Run all subjects by default = %d',run_allsubs_by_default); fprintf(fileID,'%s\n',logg{L,1}); try set(S.tx,'string',logg); catch, end
L=L+1; logg{L,1}=sprintf('Do UI steps first = %s',answer_UIstepsFirst); fprintf(fileID,'%s\n',logg{L,1}); try set(S.tx,'string',logg); catch, end
catch
end

try
    save([home_dir sprintf('neptune_settings_%s',clk_settings)],'-regexp','^(?!(cbx1|cbx4|cbx18|status_fig|wbar2|wbar3)$).') % exclude figures and status bars while saving the settings file, if possible
catch
    save([home_dir sprintf('neptune_settings_%s',clk_settings)]);
end
msgbox(sprintf('Your choices so far have been saved to\n%s',[home_dir sprintf('neptune_settings_%s',clk_settings)]),'Choices saved','custom',msgicon);
refresh_html_log


%% --- if step_after_step==1 ---
if step_after_step==1

%% initial prep
if ignore_extreme_slices~=0
    L=L+2; logg{L,1}=sprintf('-------\nIgnore first/last slices'); try fprintf(fileID,'%s\n',logg{L,1}); set(S.tx,'string',logg); catch, end; tc=tic;
    time_taken_preproc.step00_ignoreExtremeSlices = zeros(size(func_files,1),size(func_files,2));
    skip=0;
    if ( (exist([sub_dir{1} anat_files{1} '_origSlices.nii'], 'file')) || (exist([sub_dir{1} anat_files{1} '_origSlices.nii.gz'], 'file')) )
        scfMRItb_04_unzipFile(sub_dir{1},anat_files{1}, '')
        origdata = load_untouch_nii([sub_dir{1} anat_files{1} '.nii']);
        slices = size(origdata.img,3); clear origdata
    end
    for k=1:length(sub_dir)
        for k2=1:runs(k)
            if (k2==1) && ( ~( (exist([sub_dir{k} anat_files{k} '_origSlices.nii'], 'file')) || (exist([sub_dir{k} anat_files{k} '_origSlices.nii.gz'], 'file')) ) || (run_allsubs_by_default==1) )
                skip=skip+1;
                clk=clock; L=L+1; logg{L,1}=sprintf('%d/%s/%d %dh:%dm:%ds - begin ignoring first and last anatomical slice for subject (%d) of (%d)',clk(3),month{clk(2)},clk(1),clk(4),clk(5),round(clk(6)),k,length(sub_dir)); try fprintf(fileID,'%s\n',logg{L,1}); set(S.tx,'string',logg); catch, end; clear clk
                try waitbar((k/length(sub_dir)),wbar2,sprintf('Ignore first/last anatomical slices: Subject (%d) of (%d)',k,length(sub_dir))); catch, end
                fprintf('00. Ignore first and last anatomical slice: Subject (%d) of (%d)\n',k,length(sub_dir))
                time_ignoreExtremeSlices = tic;
                scfMRItb_04_unzipFile(sub_dir{k},anat_files{k}, '')
                origdata = load_untouch_nii([sub_dir{k} anat_files{k} '.nii']);
                origdata.img = origdata.img(:,:,2:end-1);
                origdata.hdr.dime.dim(4) = origdata.hdr.dime.dim(4)-2;
                slices = size(origdata.img,3);
                movefile([sub_dir{k} anat_files{k} '.nii'],[sub_dir{k} anat_files{k} '_origSlices.nii'])
                if exist([sub_dir{k} anat_files{k} '.nii.gz'],'file')
                    movefile([sub_dir{k} anat_files{k} '.nii.gz'],[sub_dir{k} anat_files{k} '_origSlices.nii.gz'])
                else
                    gzip([sub_dir{k} anat_files{k} '_origSlices.nii'])
                end
                save_untouch_nii(origdata,[sub_dir{k} anat_files{k} '.nii']);
                gzip([sub_dir{k} anat_files{k} '.nii'])
                clear origdata
                if exist('NORDIC_type','var')
                if NORDIC_type==2
                    try
                    scfMRItb_04_unzipFile(sub_dir{k},anat_files{k}, '_ph')
                    origdata = load_untouch_nii([sub_dir{k} anat_files{k} '_ph.nii']);
                    origdata.img = origdata.img(:,:,2:end-1);
                    origdata.hdr.dime.dim(4) = origdata.hdr.dime.dim(4)-2;
                    movefile([sub_dir{k} anat_files{k} '_ph.nii'],[sub_dir{k} anat_files{k} '_ph_origSlices.nii'])
                    if exist([sub_dir{k} anat_files{k} '_ph.nii.gz'],'file')
                        movefile([sub_dir{k} anat_files{k} '_ph.nii.gz'],[sub_dir{k} anat_files{k} '_ph_origSlices.nii.gz'])
                    else
                        gzip([sub_dir{k} anat_files{k} '_ph_origSlices.nii'])
                    end
                    save_untouch_nii(origdata,[sub_dir{k} anat_files{k} '_ph.nii']);
                    gzip([sub_dir{k} anat_files{k} '_ph.nii'])
                    clear origdata
                    catch
                    end
                end
                end
                clk=clock; L=L+1; logg{L,1}=sprintf('%d/%s/%d %dh:%dm:%ds - end ignoring first and last anatomical slice for subject (%d) of (%d)',clk(3),month{clk(2)},clk(1),clk(4),clk(5),round(clk(6)),k,length(sub_dir)); try fprintf(fileID,'%s\n',logg{L,1}); set(S.tx,'string',logg); catch, end; clear clk
                refresh_html_log
            else
                clk=clock; L=L+1; logg{L,1}=sprintf('%d/%s/%d %dh:%dm:%ds - did not ignore first and last anatomical slice for subject (%d) of (%d)',clk(3),month{clk(2)},clk(1),clk(4),clk(5),round(clk(6)),k,length(sub_dir)); try fprintf(fileID,'%s\n',logg{L,1}); set(S.tx,'string',logg); catch, end; clear clk
            end
            if ~( (exist([sub_dir{k} func_files{k,k2} '_origSlices.nii'], 'file')) || (exist([sub_dir{k} func_files{k,k2} '_origSlices.nii.gz'], 'file')) ) || (run_allsubs_by_default==1)
                skip=skip+1;
                clk=clock; L=L+1; logg{L,1}=sprintf('%d/%s/%d %dh:%dm:%ds - begin ignoring first and last functional slice for subject (%d) of (%d), run (%d) of (%d)',clk(3),month{clk(2)},clk(1),clk(4),clk(5),round(clk(6)),k,length(sub_dir),k2,runs(k)); try fprintf(fileID,'%s\n',logg{L,1}); set(S.tx,'string',logg); catch, end; clear clk
                try waitbar(((runs(k)*(k-1)+k2)/(length(sub_dir)*runs(k))),wbar2,sprintf('Ignore first/last functional slices: Subject (%d) of (%d), Run (%d) of (%d)',k,length(sub_dir),k2,runs(k))); catch, end
                fprintf('00. Ignore first and last functional slice: Subject (%d) of (%d), Run (%d) of (%d)\n',k,length(sub_dir),k2,runs(k))
                if ~exist('time_ignoreExtremeSlices','var')
                    time_ignoreExtremeSlices = tic;
                end
                scfMRItb_04_unzipFile(sub_dir{k},func_files{k,k2}, '')
                origdata = load_untouch_nii([sub_dir{k} func_files{k,k2} '.nii']);
                origdata.img = origdata.img(:,:,2:end-1,:);
                origdata.hdr.dime.dim(4) = origdata.hdr.dime.dim(4)-2;
                slices = size(origdata.img,3);
                movefile([sub_dir{k} func_files{k,k2} '.nii'],[sub_dir{k} func_files{k,k2} '_origSlices.nii'])
                if exist([sub_dir{k} func_files{k,k2} '.nii.gz'],'file')
                    movefile([sub_dir{k} func_files{k,k2} '.nii.gz'],[sub_dir{k} func_files{k,k2} '_origSlices.nii.gz'])
                else
                    gzip([sub_dir{k} func_files{k,k2} '_origSlices.nii'])
                end
                save_untouch_nii(origdata,[sub_dir{k} func_files{k,k2} '.nii']);
                gzip([sub_dir{k} func_files{k,k2} '.nii'])
                clear origdata
                if exist('NORDIC_type','var')
                if NORDIC_type==2
                    scfMRItb_04_unzipFile(sub_dir{k},func_files{k,k2}, '_ph')
                    origdata = load_untouch_nii([sub_dir{k} func_files{k,k2} '_ph.nii']);
                    origdata.img = origdata.img(:,:,2:end-1,:);
                    origdata.hdr.dime.dim(4) = origdata.hdr.dime.dim(4)-2;
                    movefile([sub_dir{k} func_files{k,k2} '_ph.nii'],[sub_dir{k} func_files{k,k2} '_ph_origSlices.nii'])
                    if exist([sub_dir{k} func_files{k,k2} '_ph.nii.gz'],'file')
                        movefile([sub_dir{k} func_files{k,k2} '_ph.nii.gz'],[sub_dir{k} func_files{k,k2} '_ph_origSlices.nii.gz'])
                    else
                        gzip([sub_dir{k} func_files{k,k2} '_ph_origSlices.nii'])
                    end
                    save_untouch_nii(origdata,[sub_dir{k} func_files{k,k2} '_ph.nii']);
                    gzip([sub_dir{k} func_files{k,k2} '_ph.nii'])
                    clear origdata
                end
                end
                time_taken_preproc.step00_ignoreExtremeSlices(k,k2) = toc(time_ignoreExtremeSlices); clear time_ignoreExtremeSlices Rns
                clk=clock; L=L+1; logg{L,1}=sprintf('%d/%s/%d %dh:%dm:%ds - end ignoring first and last functional slice for subject (%d) of (%d), run (%d) of (%d)',clk(3),month{clk(2)},clk(1),clk(4),clk(5),round(clk(6)),k,length(sub_dir),k2,runs(k)); try fprintf(fileID,'%s\n',logg{L,1}); set(S.tx,'string',logg); catch, end; clear clk
                refresh_html_log
            else
                clk=clock; L=L+1; logg{L,1}=sprintf('%d/%s/%d %dh:%dm:%ds - did not ignore first and last functional slice for subject (%d) of (%d), run (%d) of (%d)',clk(3),month{clk(2)},clk(1),clk(4),clk(5),round(clk(6)),k,length(sub_dir),k2,runs(k)); try fprintf(fileID,'%s\n',logg{L,1}); set(S.tx,'string',logg); catch, end; clear clk
            end
        end; clear k2
    end; clear k
    
    if skip==0
        clk=clock; L=L+1; logg{L,1}=sprintf('%d/%s/%d %dh:%dm:%ds - Ignoring first/last slices SKIPPED FOR ALL SUBJECTS',clk(3),month{clk(2)},clk(1),clk(4),clk(5),round(clk(6))); try fprintf(fileID,'%s\n',logg{L,1}); set(S.tx,'string',logg); catch, end; clear clk
    else
        clk=clock; L=L+1; logg{L,1}=sprintf('%d/%s/%d %dh:%dm:%ds - Ignoring first/last slices COMPLETED FOR ALL SUBJECTS',clk(3),month{clk(2)},clk(1),clk(4),clk(5),round(clk(6))); try fprintf(fileID,'%s\n',logg{L,1}); set(S.tx,'string',logg); catch, end; clear clk
        L=L+1; logg{L,1}=sprintf('Total time taken so far: %dh:%dm:%ds\nTime taken for this step: %dh:%dm:%ds\nTime taken for this step in each run of each subject: %dh:%dm:%ds',floor(toc(tc0)/3600),floor(mod(toc(tc0),3600)/60),floor(mod(toc(tc0),60)),floor(toc(tc)/3600),floor(mod(toc(tc),3600)/60),floor(mod(toc(tc),60)),floor(toc(tc)/3600/skip),floor(mod(toc(tc),3600)/60/skip),floor(mod(toc(tc),60)/skip)); try fprintf(fileID,'%s\n',logg{L,1}); set(S.tx,'string',logg); catch, end; clear clk
        refresh_html_log
    end
end

if discard_firstN_volumes~=0
    L=L+2; logg{L,1}=sprintf('-------\nDiscard first N functional volumes'); try fprintf(fileID,'%s\n',logg{L,1}); set(S.tx,'string',logg); catch, end; tc=tic;
    time_taken_preproc.step00_discardNvolumes = zeros(size(func_files,1),size(func_files,2));
    skip=0;
    for k=1:length(sub_dir)
        for k2=1:runs(k)
            if ~( (exist([sub_dir{k} func_files{k,k2} '_origLength.nii'], 'file')) || (exist([sub_dir{k} func_files{k,k2} '_origLength.nii.gz'], 'file')) ) || (run_allsubs_by_default==1)
                skip=skip+1;
                clk=clock; L=L+1; logg{L,1}=sprintf('%d/%s/%d %dh:%dm:%ds - begin discarding first %d volumes for subject (%d) of (%d), run (%d) of (%d)',clk(3),month{clk(2)},clk(1),clk(4),clk(5),round(clk(6)),discard_firstN_volumes,k,length(sub_dir),k2,runs(k)); try fprintf(fileID,'%s\n',logg{L,1}); set(S.tx,'string',logg); catch, end; clear clk
                try waitbar(((runs(k)*(k-1)+k2)/(length(sub_dir)*runs(k))),wbar2,sprintf('06. Discard first %d volumes: Subject (%d) of (%d), Run (%d) of (%d)',discard_firstN_volumes,k,length(sub_dir),k2,runs(k))); catch, end
                fprintf('00. Discard first %d volumes: Subject (%d) of (%d), Run (%d) of (%d)\n',discard_firstN_volumes,k,length(sub_dir),k2,runs(k))
                time_discardNvolumes = tic;
                scfMRItb_04_unzipFile(sub_dir{k},func_files{k,k2}, '')
                origdata = load_untouch_nii([sub_dir{k} func_files{k,k2} '.nii']);
                origdata.img = origdata.img(:,:,:,discard_firstN_volumes+1:end);
                origdata.hdr.dime.dim(5) = origdata.hdr.dime.dim(5)-discard_firstN_volumes;
                movefile([sub_dir{k} func_files{k,k2} '.nii'],[sub_dir{k} func_files{k,k2} '_origLength.nii'])
                if exist([sub_dir{k} func_files{k,k2} '.nii.gz'],'file')
                    movefile([sub_dir{k} func_files{k,k2} '.nii.gz'],[sub_dir{k} func_files{k,k2} '_origLength.nii.gz'])
                else
                    gzip([sub_dir{k} func_files{k,k2} '_origLength.nii'])
                end
                save_untouch_nii(origdata,[sub_dir{k} func_files{k,k2} '.nii']);
                gzip([sub_dir{k} func_files{k,k2} '.nii'])
                clear origdata
                if exist('NORDIC_type','var')
                if NORDIC_type==2
                    scfMRItb_04_unzipFile(sub_dir{k},func_files{k,k2}, '_ph')
                    origdata = load_untouch_nii([sub_dir{k} func_files{k,k2} '_ph.nii']);
                    origdata.img = origdata.img(:,:,:,discard_firstN_volumes+1:end);
                    origdata.hdr.dime.dim(5) = origdata.hdr.dime.dim(5)-discard_firstN_volumes;
                    movefile([sub_dir{k} func_files{k,k2} '_ph.nii'],[sub_dir{k} func_files{k,k2} '_ph_origLength.nii'])
                    if exist([sub_dir{k} func_files{k,k2} '_ph.nii.gz'],'file')
                        movefile([sub_dir{k} func_files{k,k2} '_ph.nii.gz'],[sub_dir{k} func_files{k,k2} '_ph_origLength.nii.gz'])
                    else
                        gzip([sub_dir{k} func_files{k,k2} '_ph_origLength.nii'])
                    end
                    save_untouch_nii(origdata,[sub_dir{k} func_files{k,k2} '_ph.nii']);
                    gzip([sub_dir{k} func_files{k,k2} '_ph.nii'])
                    clear origdata
                end
                end
                time_taken_preproc.step00_discardNvolumes(k,k2) = toc(time_discardNvolumes); clear time_discardNvolumes Rns
                clk=clock; L=L+1; logg{L,1}=sprintf('%d/%s/%d %dh:%dm:%ds - end discarding first %d volumes for subject (%d) of (%d), run (%d) of (%d)',clk(3),month{clk(2)},clk(1),clk(4),clk(5),round(clk(6)),discard_firstN_volumes,k,length(sub_dir),k2,runs(k)); try fprintf(fileID,'%s\n',logg{L,1}); set(S.tx,'string',logg); catch, end; clear clk
                refresh_html_log
            else
                clk=clock; L=L+1; logg{L,1}=sprintf('%d/%s/%d %dh:%dm:%ds - skip discarding first %d volumes for subject (%d) of (%d), run (%d) of (%d)',clk(3),month{clk(2)},clk(1),clk(4),clk(5),round(clk(6)),discard_firstN_volumes,k,length(sub_dir),k2,runs(k)); try fprintf(fileID,'%s\n',logg{L,1}); set(S.tx,'string',logg); catch, end; clear clk
            end
        end; clear k2
    end; clear k save_videos2
    
    if skip==0
        clk=clock; L=L+1; logg{L,1}=sprintf('%d/%s/%d %dh:%dm:%ds - Discard first %d volumes SKIPPED FOR ALL SUBJECTS',clk(3),month{clk(2)},clk(1),clk(4),clk(5),round(clk(6)),discard_firstN_volumes); try fprintf(fileID,'%s\n',logg{L,1}); set(S.tx,'string',logg); catch, end; clear clk
    else
        clk=clock; L=L+1; logg{L,1}=sprintf('%d/%s/%d %dh:%dm:%ds - Discard first %d volumes COMPLETED FOR ALL SUBJECTS',clk(3),month{clk(2)},clk(1),clk(4),clk(5),round(clk(6)),discard_firstN_volumes); try fprintf(fileID,'%s\n',logg{L,1}); set(S.tx,'string',logg); catch, end; clear clk
        L=L+1; logg{L,1}=sprintf('Total time taken so far: %dh:%dm:%ds\nTime taken for this step: %dh:%dm:%ds\nTime taken for this step in each run of each subject: %dh:%dm:%ds',floor(toc(tc0)/3600),floor(mod(toc(tc0),3600)/60),floor(mod(toc(tc0),60)),floor(toc(tc)/3600),floor(mod(toc(tc),3600)/60),floor(mod(toc(tc),60)),floor(toc(tc)/3600/skip),floor(mod(toc(tc),3600)/60/skip),floor(mod(toc(tc),60)/skip)); try fprintf(fileID,'%s\n',logg{L,1}); set(S.tx,'string',logg); catch, end; clear clk
        refresh_html_log
    end
end

%% '01. NORDIC denoising'
if choices_preproc(1)==1
try figure(status_fig),subplot(N,2,1),imshow(imread('status_images/step_01_blue.jpeg')), subplot(N,2,2),imshow(imread('status_images/status_blue.jpg')), catch, end
L=L+2; logg{L,1}=sprintf('-------\n%s',preprocsteps{1}); try fprintf(fileID,'%s\n',logg{L,1}); set(S.tx,'string',logg); catch, end; tc=tic;
time_taken_preproc.step01_NORDICdenoising = zeros(size(func_files,1),size(func_files,2));

skip=0;
for k=1:length(sub_dir)
    for k2=1:runs(k)
        if ~( (exist([sub_dir{k} func_files{k,k2} '_beforeNORDIC.nii'], 'file')) || (exist([sub_dir{k} func_files{k,k2} '_beforeNORDIC.nii.gz'], 'file')) ) || (run_allsubs_by_default==1)
            skip=skip+1;
            clk=clock; L=L+1; logg{L,1}=sprintf('%d/%s/%d %dh:%dm:%ds - begin (01) NORDIC denoising for subject (%d) of (%d), run (%d) of (%d)',clk(3),month{clk(2)},clk(1),clk(4),clk(5),round(clk(6)),k,length(sub_dir),k2,runs(k)); try fprintf(fileID,'%s\n',logg{L,1}); set(S.tx,'string',logg); catch, end; clear clk
            try waitbar(((runs(k)*(k-1)+k2)/(length(sub_dir)*runs(k))),wbar2,sprintf('01. NORDIC denoising: Subject (%d) of (%d), Run (%d) of (%d)',k,length(sub_dir),k2,runs(k))); catch, end
            fprintf('01. Perform NORDIC denoising: Subject (%d) of (%d), Run (%d) of (%d)\n',k,length(sub_dir),k2,runs(k))
            if runs(k)==1, Rns=0; else, Rns=k2; end
            time_01_NORDICdenoising = tic;
            scfMRItb_01_NORDIC_denoising(sub_dir{k},func_files{k,k2},anat_files{k,1},NORDIC_type,k2, wbar3,Rns);
            time_taken_preproc.step01_NORDICdenoising(k,k2) = toc(time_01_NORDICdenoising); clear time_01_NORDICdenoising Rns
            clk=clock; L=L+1; logg{L,1}=sprintf('%d/%s/%d %dh:%dm:%ds - end (01) NORDIC denoising for subject (%d) of (%d), run (%d) of (%d)',clk(3),month{clk(2)},clk(1),clk(4),clk(5),round(clk(6)),k,length(sub_dir),k2,runs(k)); try fprintf(fileID,'%s\n',logg{L,1}); set(S.tx,'string',logg); catch, end; clear clk
            refresh_html_log
        else
            clk=clock; L=L+1; logg{L,1}=sprintf('%d/%s/%d %dh:%dm:%ds - skip (01) NORDIC denoising for subject (%d) of (%d), run (%d) of (%d)',clk(3),month{clk(2)},clk(1),clk(4),clk(5),round(clk(6)),k,length(sub_dir),k2,runs(k)); try fprintf(fileID,'%s\n',logg{L,1}); set(S.tx,'string',logg); catch, end; clear clk
        end
    end; clear k2
end; clear k save_videos2

if skip==0
    clk=clock; L=L+1; logg{L,1}=sprintf('%d/%s/%d %dh:%dm:%ds - (01) NORDIC denoising SKIPPED FOR ALL SUBJECTS',clk(3),month{clk(2)},clk(1),clk(4),clk(5),round(clk(6))); try fprintf(fileID,'%s\n',logg{L,1}); set(S.tx,'string',logg); catch, end; clear clk
    try figure(status_fig),subplot(N,2,1),imshow(imread('status_images/step_01_green.jpeg')), subplot(N,2,2),imshow(imread('status_images/status_green_skip.jpg')), catch, end
else
    clk=clock; L=L+1; logg{L,1}=sprintf('%d/%s/%d %dh:%dm:%ds - (01) NORDIC denoising COMPLETED FOR ALL SUBJECTS',clk(3),month{clk(2)},clk(1),clk(4),clk(5),round(clk(6))); try fprintf(fileID,'%s\n',logg{L,1}); set(S.tx,'string',logg); catch, end; clear clk
    L=L+1; logg{L,1}=sprintf('Total time taken so far: %dh:%dm:%ds\nTime taken for this step: %dh:%dm:%ds\nTime taken for this step in each run of each subject: %dh:%dm:%ds',floor(toc(tc0)/3600),floor(mod(toc(tc0),3600)/60),floor(mod(toc(tc0),60)),floor(toc(tc)/3600),floor(mod(toc(tc),3600)/60),floor(mod(toc(tc),60)),floor(toc(tc)/3600/skip),floor(mod(toc(tc),3600)/60/skip),floor(mod(toc(tc),60)/skip)); try fprintf(fileID,'%s\n',logg{L,1}); set(S.tx,'string',logg); catch, end; clear clk
    try figure(status_fig),subplot(N,2,1),imshow(imread('status_images/step_01_green.jpeg')), subplot(N,2,2),imshow(imread('status_images/status_green.jpg')), catch, end
    refresh_html
end
end

%% '02. Slice-timing correction'
if choices_preproc(2)==1
try figure(status_fig),subplot(N,2,3),imshow(imread('status_images/step_02_blue.jpeg')), subplot(N,2,4),imshow(imread('status_images/status_blue_UI.jpg')), catch, end
L=L+2; logg{L,1}=sprintf('-------\n%s',preprocsteps{2}); try fprintf(fileID,'%s\n',logg{L,1}); set(S.tx,'string',logg); catch, end; tc=tic;
time_taken_preproc.step02_slicetimingcorrection = zeros(size(func_files,1),size(func_files,2));

skip=0;
for k=1:length(sub_dir)
    for k2=1:runs(k)
        if ~( (exist([sub_dir{k} func_files{k,k2} 'before_slicetiming' '.nii'], 'file')) || (exist([sub_dir{k} func_files{k,k2} 'before_slicetiming' '.nii.gz'], 'file')) ) || (run_allsubs_by_default==1)
            skip=skip+1;
            clk=clock; L=L+1; logg{L,1}=sprintf('%d/%s/%d %dh:%dm:%ds - begin (02) slice timing correction for subject (%d) of (%d), run (%d) of (%d)',clk(3),month{clk(2)},clk(1),clk(4),clk(5),round(clk(6)),k,length(sub_dir),k2,runs(k)); try fprintf(fileID,'%s\n',logg{L,1}); set(S.tx,'string',logg); catch, end; clear clk
            try waitbar(((runs(k)*(k-1)+k2)/(length(sub_dir)*runs(k))),wbar2,sprintf('02. Slice-timing correction: Subject (%d) of (%d), Run (%d) of (%d)',k,length(sub_dir),k2,runs(k))); catch, end
            fprintf('02. Slice-timing correction: Subject (%d) of (%d), Run (%d) of (%d)\n',k,length(sub_dir),k2,runs(k))
            time_02_slicetimingcorrection = tic;
            scfMRItb_02_slicetimingcorrection(sub_dir{k},func_files{k,k2},sliceorder');
            time_taken_preproc.step02_slicetimingcorrection(k,k2) = toc(time_02_slicetimingcorrection); clear time_02_slicetimingcorrection
            clk=clock; L=L+1; logg{L,1}=sprintf('%d/%s/%d %dh:%dm:%ds - end (02) slice timing correction for subject (%d) of (%d), run (%d) of (%d)',clk(3),month{clk(2)},clk(1),clk(4),clk(5),round(clk(6)),k,length(sub_dir),k2,runs(k)); try fprintf(fileID,'%s\n',logg{L,1}); set(S.tx,'string',logg); catch, end; clear clk time_02_slicetimingcorrection
            refresh_html_log
        else
            clk=clock; L=L+1; logg{L,1}=sprintf('%d/%s/%d %dh:%dm:%ds - skip (02) slice timing correction for subject (%d) of (%d), run (%d) of (%d)',clk(3),month{clk(2)},clk(1),clk(4),clk(5),round(clk(6)),k,length(sub_dir),k2,runs(k)); try fprintf(fileID,'%s\n',logg{L,1}); set(S.tx,'string',logg); catch, end; clear clk
        end
    end
    clear k2
end

if skip==0
    clk=clock; L=L+1; logg{L,1}=sprintf('%d/%s/%d %dh:%dm:%ds - (02) slice timing correction SKIPPED FOR ALL SUBJECTS',clk(3),month{clk(2)},clk(1),clk(4),clk(5),round(clk(6))); try fprintf(fileID,'%s\n',logg{L,1}); set(S.tx,'string',logg); catch, end; clear clk
    try figure(status_fig),subplot(N,2,3),imshow(imread('status_images/step_02_green.jpeg')), subplot(N,2,4),imshow(imread('status_images/status_green_skip.jpg')), catch, end
else
    clk=clock; L=L+1; logg{L,1}=sprintf('%d/%s/%d %dh:%dm:%ds - (02) slice timing correction COMPLETED FOR ALL SUBJECTS',clk(3),month{clk(2)},clk(1),clk(4),clk(5),round(clk(6))); try fprintf(fileID,'%s\n',logg{L,1}); set(S.tx,'string',logg); catch, end; clear clk
    L=L+1; logg{L,1}=sprintf('Total time taken so far: %dh:%dm:%ds\nTime taken for this step: %dh:%dm:%ds\nTime taken for this step in each run of each subject: %dh:%dm:%ds',floor(toc(tc0)/3600),floor(mod(toc(tc0),3600)/60),floor(mod(toc(tc0),60)),floor(toc(tc)/3600),floor(mod(toc(tc),3600)/60),floor(mod(toc(tc),60)),floor(toc(tc)/3600/skip),floor(mod(toc(tc),3600)/60/skip),floor(mod(toc(tc),60)/skip)); try fprintf(fileID,'%s\n',logg{L,1}); set(S.tx,'string',logg); catch, end; clear clk
    try figure(status_fig),subplot(N,2,3),imshow(imread('status_images/step_02_green.jpeg')), subplot(N,2,4),imshow(imread('status_images/status_green.jpg')), catch, end
end
refresh_html_log
end

%% '03. Match anatomical and functional slices'
if choices_preproc(3)==1
try figure(status_fig),subplot(N,2,5),imshow(imread('status_images/step_03_blue.jpeg')), subplot(N,2,6),imshow(imread('status_images/status_blue.jpg')), catch, end
L=L+2; logg{L,1}=sprintf('-------\n%s',preprocsteps{3}); try fprintf(fileID,'%s\n',logg{L,1}); set(S.tx,'string',logg); catch, end; tc=tic;
time_taken_preproc.step03_matchAnatFuncSlices = zeros(size(func_files,1),1);

skip=0;
for k=1:length(sub_dir)
    if ~( (exist([sub_dir{k} anat_files{k,1} '_reduced.nii'], 'file')) || (exist([sub_dir{k} anat_files{k,1} '_reduced.nii.gz'], 'file')) ) || (run_allsubs_by_default==1)
        skip=skip+1;
        clk=clock; L=L+1; logg{L,1}=sprintf('%d/%s/%d %dh:%dm:%ds - begin (03) match anatomical and functional slices for subject (%d) of (%d)',clk(3),month{clk(2)},clk(1),clk(4),clk(5),round(clk(6)),k,length(sub_dir)); try fprintf(fileID,'%s\n',logg{L,1}); set(S.tx,'string',logg); catch, end; clear clk
        try waitbar((k/length(sub_dir)),wbar2,sprintf('03. Match anatomical and functional slices: Subject (%d) of (%d)\n',k,length(sub_dir))); catch, end
        fprintf('03. Match anatomical and functional slices: Subject (%d) of (%d)\n',k,length(sub_dir))
        time_03_matchAnatFuncSlices = tic;
        [notunix, noAFNI] = scfMRItb_03_matchAnatFuncSlices(sub_dir{k},func_files{k,1},anat_files{k,1});
        time_taken_preproc.step03_matchAnatFuncSlices(k,1) = toc(time_03_matchAnatFuncSlices); clear time_03_matchAnatFuncSlices
        if notunix~=0
            tmp=clock; clk = sprintf('%.4d-%.2d-%.2d_%.2d-%.2d',tmp(1),tmp(2),tmp(3),tmp(4),tmp(5));
            save([home_dir sprintf('workspace_error_%s',clk)])
            msgbox(sprintf('The toolbox cannot be used without Unix support needed to run AFNI commands.\n\nYour choices so far have been saved to\n%s\n\nTerminating execution...',sprintf('workspace_error_%s.mat',clk)),'Abort','error'); clear clk tmp
            return
        end
        if noAFNI~=0
            tmp=clock; clk = sprintf('%.4d-%.2d-%.2d_%.2d-%.2d',tmp(1),tmp(2),tmp(3),tmp(4),tmp(5));
            save([home_dir sprintf('workspace_error_%s',clk)])
            msgbox(sprintf('AFNI is not in Matlab path (needed); cannot continue execution...\n- If you have not yet installed AFNI on your system, now is a good time.\n- If you have AFNI yet got this message, refer to line-9 (afni_directory = ...) in the script main_scfMRItb.m\n\nYour choices so far have been saved to\n%s\n\nTerminating execution...',sprintf('workspace_error_%s.mat',clk)),'Abort','error'); clear clk tmp
            return
        end
        clk=clock; L=L+1; logg{L,1}=sprintf('%d/%s/%d %dh:%dm:%ds - end (03) match anatomical and functional slices for subject (%d) of (%d)',clk(3),month{clk(2)},clk(1),clk(4),clk(5),round(clk(6)),k,length(sub_dir)); try fprintf(fileID,'%s\n',logg{L,1}); set(S.tx,'string',logg); catch, end; clear clk
        refresh_html_log
    else
        clk=clock; L=L+1; logg{L,1}=sprintf('%d/%s/%d %dh:%dm:%ds - skip (03) match anatomical and functional slices for subject (%d) of (%d)',clk(3),month{clk(2)},clk(1),clk(4),clk(5),round(clk(6)),k,length(sub_dir)); try fprintf(fileID,'%s\n',logg{L,1}); set(S.tx,'string',logg); catch, end; clear clk
    end
end

try % in case the functional was resliced along the z-axis to match the slice thickness of the anatomical, refresh the 'slices' variable that representes the number of slices
    scfMRItb_04_unzipFile(sub_dir{1}, func_files{1,1}, '_resliced')
    F = load_untouch_nii([sub_dir{1} func_files{1,1} '_resliced' '.nii']);
    slices = size(F.img,3); clear F % This toolbox assumes that all subjects have the same number of slices. If not, you would have to treat differences as separate cohorts and process separately (and aggregate yourself in the end)
catch
end

if skip==0
    clk=clock; L=L+1; logg{L,1}=sprintf('%d/%s/%d %dh:%dm:%ds - (03) match anatomical and functional slices SKIPPED FOR ALL SUBJECTS',clk(3),month{clk(2)},clk(1),clk(4),clk(5),round(clk(6))); try fprintf(fileID,'%s\n',logg{L,1}); set(S.tx,'string',logg); catch, end; clear clk
    try figure(status_fig),subplot(N,2,5),imshow(imread('status_images/step_03_green.jpeg')), subplot(N,2,6),imshow(imread('status_images/status_green_skip.jpg')), catch, end
else
    clk=clock; L=L+1; logg{L,1}=sprintf('%d/%s/%d %dh:%dm:%ds - (03) match anatomical and functional slices COMPLETED FOR ALL SUBJECTS',clk(3),month{clk(2)},clk(1),clk(4),clk(5),round(clk(6))); try fprintf(fileID,'%s\n',logg{L,1}); set(S.tx,'string',logg); catch, end; clear clk
    L=L+1; logg{L,1}=sprintf('Total time taken so far: %dh:%dm:%ds\nTime taken for this step: %dh:%dm:%ds\nTime taken for this step in each run of each subject: %dh:%dm:%ds',floor(toc(tc0)/3600),floor(mod(toc(tc0),3600)/60),floor(mod(toc(tc0),60)),floor(toc(tc)/3600),floor(mod(toc(tc),3600)/60),floor(mod(toc(tc),60)),floor(toc(tc)/3600/skip),floor(mod(toc(tc),3600)/60/skip),floor(mod(toc(tc),60)/skip)); try fprintf(fileID,'%s\n',logg{L,1}); set(S.tx,'string',logg); catch, end; clear clk
    try figure(status_fig),subplot(N,2,5),imshow(imread('status_images/step_03_green.jpeg')), subplot(N,2,6),imshow(imread('status_images/status_green.jpg')), catch, end
end
refresh_html_log
end
         
%% '04. Split data into slices'
if choices_preproc(4)==1
try figure(status_fig),subplot(N,2,7),imshow(imread('status_images/step_04_blue.jpeg')), subplot(N,2,8),imshow(imread('status_images/status_blue.jpg')), catch, end
L=L+2; logg{L,1}=sprintf('-------\n%s',preprocsteps{4}); try fprintf(fileID,'%s\n',logg{L,1}); set(S.tx,'string',logg); catch, end; tc=tic;
time_taken_preproc.step04_splitData = zeros(size(func_files,1),size(func_files,2));

skip=0;
for k=1:length(sub_dir)
    for k2=1:runs(k)
        if ~( (exist([sub_dir{k} func_files{k,k2} '_slice' num2str(slices) '.nii'], 'file')) || (exist([sub_dir{k} func_files{k,k2} '_slice' num2str(slices) '.nii.gz'], 'file')) || (exist([sub_dir{k} './' func_files{k,k2} '_slice' num2str(slices) '_MC_params.txt'], 'file')) ) || (run_allsubs_by_default==1)
            skip=skip+1;
            clk=clock; L=L+1; logg{L,1}=sprintf('%d/%s/%d %dh:%dm:%ds - begin (04) split data into slices for subject (%d) of (%d), run (%d) of (%d)',clk(3),month{clk(2)},clk(1),clk(4),clk(5),round(clk(6)),k,length(sub_dir),k2,runs(k)); try fprintf(fileID,'%s\n',logg{L,1}); set(S.tx,'string',logg); catch, end; clear clk
            try waitbar(((runs(k)*(k-1)+k2)/(length(sub_dir)*runs(k))),wbar2,sprintf('04. Split data into slices: Subject (%d) of (%d), Run (%d) of (%d)',k,length(sub_dir),k2,runs(k))); catch, end
            fprintf('04. Split data into slices: Subject (%d) of (%d), Run (%d) of (%d)\n',k,length(sub_dir),k2,runs(k))
            time_04_splitData = tic;
            % Splitting data need not be performed because it will be done within other subsequent functions as and when needed
            % [notunix, noAFNI] = scfMRItb_04_splitData(sub_dir{k},func_files{k,k2},anat_files{k,1},wbar3);
            time_taken_preproc.step04_splitData(k,k2) = toc(time_04_splitData); clear time_04_splitData
            clk=clock; L=L+1; logg{L,1}=sprintf('%d/%s/%d %dh:%dm:%ds - end (04) split data into slices for subject (%d) of (%d), run (%d) of (%d)',clk(3),month{clk(2)},clk(1),clk(4),clk(5),round(clk(6)),k,length(sub_dir),k2,runs(k)); try fprintf(fileID,'%s\n',logg{L,1}); set(S.tx,'string',logg); catch, end; clear clk
            refresh_html_log
        else
            clk=clock; L=L+1; logg{L,1}=sprintf('%d/%s/%d %dh:%dm:%ds - skip (04) split data into slices for subject (%d) of (%d), run (%d) of (%d)',clk(3),month{clk(2)},clk(1),clk(4),clk(5),round(clk(6)),k,length(sub_dir),k2,runs(k)); try fprintf(fileID,'%s\n',logg{L,1}); set(S.tx,'string',logg); catch, end; clear clk
        end
    end; clear k2
end

if skip==0
    clk=clock; L=L+1; logg{L,1}=sprintf('%d/%s/%d %dh:%dm:%ds - (04) split data into slices SKIPPED FOR ALL SUBJECTS',clk(3),month{clk(2)},clk(1),clk(4),clk(5),round(clk(6))); try fprintf(fileID,'%s\n',logg{L,1}); set(S.tx,'string',logg); catch, end; clear clk
    try figure(status_fig),subplot(N,2,7),imshow(imread('status_images/step_04_green.jpeg')), subplot(N,2,8),imshow(imread('status_images/status_green_skip.jpg')), catch, end
else
    clk=clock; L=L+1; logg{L,1}=sprintf('%d/%s/%d %dh:%dm:%ds - (04) split data into slices COMPLETED FOR ALL SUBJECTS',clk(3),month{clk(2)},clk(1),clk(4),clk(5),round(clk(6))); try fprintf(fileID,'%s\n',logg{L,1}); set(S.tx,'string',logg); catch, end; clear clk
    L=L+1; logg{L,1}=sprintf('Total time taken so far: %dh:%dm:%ds\nTime taken for this step: %dh:%dm:%ds\nTime taken for this step in each run of each subject: %dh:%dm:%ds',floor(toc(tc0)/3600),floor(mod(toc(tc0),3600)/60),floor(mod(toc(tc0),60)),floor(toc(tc)/3600),floor(mod(toc(tc),3600)/60),floor(mod(toc(tc),60)),floor(toc(tc)/3600/skip),floor(mod(toc(tc),3600)/60/skip),floor(mod(toc(tc),60)/skip)); try fprintf(fileID,'%s\n',logg{L,1}); set(S.tx,'string',logg); catch, end; clear clk
    try figure(status_fig),subplot(N,2,7),imshow(imread('status_images/step_04_green.jpeg')), subplot(N,2,8),imshow(imread('status_images/status_green.jpg')), catch, end
end
refresh_html_log
end

%% '05. Define "not cord" mask'
if choices_preproc(5)==1
try figure(status_fig),subplot(N,2,9),imshow(imread('status_images/step_05_blue.jpeg')), subplot(N,2,10),imshow(imread('status_images/status_blue_UI.jpg')), catch, end
L=L+2; logg{L,1}=sprintf('-------\n%s',preprocsteps{5}); try fprintf(fileID,'%s\n',logg{L,1}); set(S.tx,'string',logg); catch, end; tc=tic;
time_taken_preproc.step05_notCordMask = zeros(size(func_files,1),size(func_files,2));
if (save_videos==0)||(save_videos==1), save_videos2 = save_videos; elseif (save_videos==2)&&(sum([choices_preproc(6);choices_preproc(8:9);choices_preproc(11:14)])==0), save_videos2 = 1; else, save_videos2 = 0; end

skip=0;
for k=1:length(sub_dir)
    for k2=1:runs(k)
        if ~(exist([sub_dir{k} func_files{k,k2} '_mask_NS.mat'], 'file')) || (run_allsubs_by_default==1)
            skip=skip+1;
            clk=clock; L=L+1; logg{L,1}=sprintf('%d/%s/%d %dh:%dm:%ds - begin (05) define "not cord" mask for subject (%d) of (%d), run (%d) of (%d)',clk(3),month{clk(2)},clk(1),clk(4),clk(5),round(clk(6)),k,length(sub_dir),k2,runs(k)); try fprintf(fileID,'%s\n',logg{L,1}); set(S.tx,'string',logg); catch, end; clear clk
            try waitbar(((runs(k)*(k-1)+k2)/(length(sub_dir)*runs(k))),wbar2,sprintf('05. Define "not cord" mask: Subject (%d) of (%d), Run (%d) of (%d)',k,length(sub_dir),k2,runs(k))); catch, end
            fprintf('05. Define "not cord" mask: Subject (%d) of (%d), Run (%d) of (%d)\n',k,length(sub_dir),k2,runs(k))
            if runs(k)==1, Rns=0; else, Rns=k2; end
            time_05_notCordMask = tic;
            scfMRItb_05_notCordMask(sub_dir{k},func_files{k,k2}, wbar3,save_videos2,Rns,exist_SCT);
            time_taken_preproc.step05_notCordMask(k,k2) = toc(time_05_notCordMask); clear time_05_notCordMask Rns
            clk=clock; L=L+1; logg{L,1}=sprintf('%d/%s/%d %dh:%dm:%ds - end (05) define "not cord" mask for subject (%d) of (%d), run (%d) of (%d)',clk(3),month{clk(2)},clk(1),clk(4),clk(5),round(clk(6)),k,length(sub_dir),k2,runs(k)); try fprintf(fileID,'%s\n',logg{L,1}); set(S.tx,'string',logg); catch, end; clear clk
            refresh_html_log
        else
            if runs(k)==1, Rns=0; else, Rns=k2; end
            scfMRItb_05_notCordMask_onlyQC(sub_dir{k},func_files{k,k2}, wbar3,save_videos2,Rns,exist_SCT);
            clk=clock; L=L+1; logg{L,1}=sprintf('%d/%s/%d %dh:%dm:%ds - skip (05) define "not cord" mask for subject (%d) of (%d), run (%d) of (%d)',clk(3),month{clk(2)},clk(1),clk(4),clk(5),round(clk(6)),k,length(sub_dir),k2,runs(k)); try fprintf(fileID,'%s\n',logg{L,1}); set(S.tx,'string',logg); catch, end; clear clk
        end
    end; clear k2
end; clear k save_videos2

if skip==0
    clk=clock; L=L+1; logg{L,1}=sprintf('%d/%s/%d %dh:%dm:%ds - (05) define "not cord" mask SKIPPED FOR ALL SUBJECTS',clk(3),month{clk(2)},clk(1),clk(4),clk(5),round(clk(6))); try fprintf(fileID,'%s\n',logg{L,1}); set(S.tx,'string',logg); catch, end; clear clk
    try figure(status_fig),subplot(N,2,9),imshow(imread('status_images/step_05_green.jpeg')), subplot(N,2,10),imshow(imread('status_images/status_green_skip.jpg')), catch, end
else
    clk=clock; L=L+1; logg{L,1}=sprintf('%d/%s/%d %dh:%dm:%ds - (05) define "not cord" mask COMPLETED FOR ALL SUBJECTS',clk(3),month{clk(2)},clk(1),clk(4),clk(5),round(clk(6))); try fprintf(fileID,'%s\n',logg{L,1}); set(S.tx,'string',logg); catch, end; clear clk
    L=L+1; logg{L,1}=sprintf('Total time taken so far: %dh:%dm:%ds\nTime taken for this step: %dh:%dm:%ds\nTime taken for this step in each run of each subject: %dh:%dm:%ds',floor(toc(tc0)/3600),floor(mod(toc(tc0),3600)/60),floor(mod(toc(tc0),60)),floor(toc(tc)/3600),floor(mod(toc(tc),3600)/60),floor(mod(toc(tc),60)),floor(toc(tc)/3600/skip),floor(mod(toc(tc),3600)/60/skip),floor(mod(toc(tc),60)/skip)); try fprintf(fileID,'%s\n',logg{L,1}); set(S.tx,'string',logg); catch, end; clear clk
    try figure(status_fig),subplot(N,2,9),imshow(imread('status_images/step_05_green.jpeg')), subplot(N,2,10),imshow(imread('status_images/status_green.jpg')), catch, end
    refresh_html
end
end

%% 'Do "10. Define GM, WM and CSF masks" if UIstepsFirst=1'
if UIstepsFirst==1
if choices_preproc(10)==1
try figure(status_fig),subplot(N,2,19),imshow(imread('status_images/step_10_blue.jpeg')), subplot(N,2,20),imshow(imread('status_images/status_blue_UI.jpg')), catch, end
L=L+2; logg{L,1}=sprintf('-------\n%s',preprocsteps{10}); try fprintf(fileID,'%s\n',logg{L,1}); set(S.tx,'string',logg); catch, end; tc=tic;

skip=0;
time_taken_preproc.step10_GMWMCSFmasks_detailed = cell(length(sub_dir),1);
time_taken_preproc.step10_GMWMCSFmasks = zeros(size(func_files,1),size(func_files,2));
for k=1:length(sub_dir)
    if ~( exist([sub_dir{k} anat_files{k,1} '_reduced' '_mask_GM.nii'], 'file') || exist([sub_dir{k} anat_files{k,1} '_reduced' '_mask_GM.nii.gz'], 'file') ) || (run_allsubs_by_default==1)
        skip=skip+1;
        clk=clock; L=L+1; logg{L,1}=sprintf('%d/%s/%d %dh:%dm:%ds - begin (10) define GM, WM and CSF masks for subject (%d) of (%d), run (1) of (%d)',clk(3),month{clk(2)},clk(1),clk(4),clk(5),round(clk(6)),k,length(sub_dir),runs(k)); try fprintf(fileID,'%s\n',logg{L,1}); set(S.tx,'string',logg); catch, end; clear clk
        try waitbar((k*1/length(sub_dir)*runs(k)),wbar2,sprintf('10. Define GM, WM and CSF masks: Subject (%d) of (%d), Run (1) of (%d)',k,length(sub_dir),runs(k))); catch, end
        fprintf('10. Define GM, WM and CSF masks: Subject (%d) of (%d), Run (1) of (%d)\n',k,length(sub_dir),runs(k))
        time_10_GMWMCSFmasks = tic;
        time_taken_preproc.step10_GMWMCSFmasks_detailed{k,1} = scfMRItb_10_GMWMCSFmasks(sub_dir{k},func_files{k,1},anat_files{k,1},wbar3,exist_SCT,mask_GM_flag,choices_preproc,C_spine, SCT_vert_auto,SCT_manualreg_labels);
        time_taken_preproc.step10_GMWMCSFmasks(k,1) = toc(time_10_GMWMCSFmasks); clear time_10_GMWMCSFmasks
        clk=clock; L=L+1; logg{L,1}=sprintf('%d/%s/%d %dh:%dm:%ds - end (10) define GM, WM and CSF masks for subject (%d) of (%d), run (1) of (%d)',clk(3),month{clk(2)},clk(1),clk(4),clk(5),round(clk(6)),k,length(sub_dir),runs(k)); try fprintf(fileID,'%s\n',logg{L,1}); set(S.tx,'string',logg); catch, end; clear clk
        refresh_html_log
    else
        scfMRItb_10_GMWMCSFmasks_subset(sub_dir{k},func_files{k,1},anat_files{k,1},choices_preproc);
        clk=clock; L=L+1; logg{L,1}=sprintf('%d/%s/%d %dh:%dm:%ds - skip (10) define GM, WM and CSF masks for subject (%d) of (%d), run (1) of (%d)',clk(3),month{clk(2)},clk(1),clk(4),clk(5),round(clk(6)),k,length(sub_dir),runs(k)); try fprintf(fileID,'%s\n',logg{L,1}); set(S.tx,'string',logg); catch, end; clear clk
    end
end

if skip==0
    clk=clock; L=L+1; logg{L,1}=sprintf('%d/%s/%d %dh:%dm:%ds - (10) define GM, WM and CSF masks SKIPPED FOR ALL SUBJECTS',clk(3),month{clk(2)},clk(1),clk(4),clk(5),round(clk(6))); try fprintf(fileID,'%s\n',logg{L,1}); set(S.tx,'string',logg); catch, end; clear clk
    try figure(status_fig),subplot(N,2,19),imshow(imread('status_images/step_10_green.jpeg')), subplot(N,2,20),imshow(imread('status_images/status_green_skip.jpg')), catch, end
else
    clk=clock; L=L+1; logg{L,1}=sprintf('%d/%s/%d %dh:%dm:%ds - (10) define GM, WM and CSF masks COMPLETED FOR ALL SUBJECTS',clk(3),month{clk(2)},clk(1),clk(4),clk(5),round(clk(6))); try fprintf(fileID,'%s\n',logg{L,1}); set(S.tx,'string',logg); catch, end; clear clk
    L=L+1; logg{L,1}=sprintf('Total time taken so far: %dh:%dm:%ds\nTime taken for this step: %dh:%dm:%ds\nTime taken for this step in each run of each subject: %dh:%dm:%ds',floor(toc(tc0)/3600),floor(mod(toc(tc0),3600)/60),floor(mod(toc(tc0),60)),floor(toc(tc)/3600),floor(mod(toc(tc),3600)/60),floor(mod(toc(tc),60)),floor(toc(tc)/3600/skip),floor(mod(toc(tc),3600)/60/skip),floor(mod(toc(tc),60)/skip)); try fprintf(fileID,'%s\n',logg{L,1}); set(S.tx,'string',logg); catch, end; clear clk
    try figure(status_fig),subplot(N,2,19),imshow(imread('status_images/step_10_green.jpeg')), subplot(N,2,20),imshow(imread('status_images/status_green.jpg')), catch, end
    refresh_html_log
end
end
end

%% 'Do "16. Define cord quadrants" if UIstepsFirst=1'
if UIstepsFirst==1
if choices_preproc(16)==1
try figure(status_fig),subplot(N,2,31),imshow(imread('status_images/step_16_blue.jpeg')), subplot(N,2,32),imshow(imread('status_images/status_blue_UI.jpg')), catch, end
L=L+2; logg{L,1}=sprintf('-------\n%s',preprocsteps{16}); try fprintf(fileID,'%s\n',logg{L,1}); set(S.tx,'string',logg); catch, end; tc=tic;
time_taken_preproc.step16_cord_quadrants = zeros(size(func_files,1),1);

skip=0;
for k=1:length(sub_dir)
    if ~(exist([sub_dir{k} 'cord_quadrants.mat'], 'file')) || (run_allsubs_by_default==1)
        skip=skip+1;
        clk=clock; L=L+1; logg{L,1}=sprintf('%d/%s/%d %dh:%dm:%ds - begin (16) define cord quadrants for subject (%d) of (%d)',clk(3),month{clk(2)},clk(1),clk(4),clk(5),round(clk(6)),k,length(sub_dir)); try fprintf(fileID,'%s\n',logg{L,1}); set(S.tx,'string',logg); catch, end; clear clk
        try waitbar((k/length(sub_dir)),wbar2,sprintf('16. Define cord quadrants: Subject (%d) of (%d)',k,length(sub_dir))); catch, end
        fprintf('16. Define cord quadrants: Subject (%d) of (%d)\n',k,length(sub_dir))
        time_16_cord_quadrants = tic;
        scfMRItb_16_cord_quadrants_UIstepsFirst(sub_dir{k},anat_files{k,1},wbar3);
        time_taken_preproc.step16_cord_quadrants(k,1) = toc(time_16_cord_quadrants); clear time_16_cord_quadrants
        clk=clock; L=L+1; logg{L,1}=sprintf('%d/%s/%d %dh:%dm:%ds - end (16) define cord quadrants for subject (%d) of (%d)',clk(3),month{clk(2)},clk(1),clk(4),clk(5),round(clk(6)),k,length(sub_dir)); try fprintf(fileID,'%s\n',logg{L,1}); set(S.tx,'string',logg); catch, end; clear clk
        refresh_html_log
    else
        clk=clock; L=L+1; logg{L,1}=sprintf('%d/%s/%d %dh:%dm:%ds - skip (16) define cord quadrants for subject (%d) of (%d)',clk(3),month{clk(2)},clk(1),clk(4),clk(5),round(clk(6)),k,length(sub_dir)); try fprintf(fileID,'%s\n',logg{L,1}); set(S.tx,'string',logg); catch, end; clear clk
    end
end

if skip==0
    clk=clock; L=L+1; logg{L,1}=sprintf('%d/%s/%d %dh:%dm:%ds - (16) define cord quadrants SKIPPED FOR ALL SUBJECTS',clk(3),month{clk(2)},clk(1),clk(4),clk(5),round(clk(6))); try fprintf(fileID,'%s\n',logg{L,1}); set(S.tx,'string',logg); catch, end; clear clk
    try figure(status_fig),subplot(N,2,31),imshow(imread('status_images/step_16_green.jpeg')), subplot(N,2,32),imshow(imread('status_images/status_green_skip.jpg')), catch, end
else
    clk=clock; L=L+1; logg{L,1}=sprintf('%d/%s/%d %dh:%dm:%ds - (16) define cord quadrants COMPLETED FOR ALL SUBJECTS',clk(3),month{clk(2)},clk(1),clk(4),clk(5),round(clk(6))); try fprintf(fileID,'%s\n',logg{L,1}); set(S.tx,'string',logg); catch, end; clear clk
    L=L+1; logg{L,1}=sprintf('Total time taken so far: %dh:%dm:%ds\nTime taken for this step: %dh:%dm:%ds\nTime taken for this step in each run of each subject: %dh:%dm:%ds',floor(toc(tc0)/3600),floor(mod(toc(tc0),3600)/60),floor(mod(toc(tc0),60)),floor(toc(tc)/3600),floor(mod(toc(tc),3600)/60),floor(mod(toc(tc),60)),floor(toc(tc)/3600/skip),floor(mod(toc(tc),3600)/60/skip),floor(mod(toc(tc),60)/skip)); try fprintf(fileID,'%s\n',logg{L,1}); set(S.tx,'string',logg); catch, end; clear clk
    try figure(status_fig),subplot(N,2,31),imshow(imread('status_images/step_16_green.jpeg')), subplot(N,2,32),imshow(imread('status_images/status_green.jpg')), catch, end
    refresh_html_log
end
end
end

%% 'Do Post-processing choices (if in the pipeline) if UIstepsFirst=1'
if UIstepsFirst==1
if postproc==1
    postproc_choices

    try
        save([home_dir sprintf('neptune_settings_%s',clk_settings)],'-regexp','^(?!(cbx1|cbx4|cbx18|status_fig|wbar2|wbar3)$).') % exclude figures and status bars while saving the settings file, if possible
    catch
        save([home_dir sprintf('neptune_settings_%s',clk_settings)]);
    end
    msgbox(sprintf('Your post-processing choices have been updated in:\n%s',[home_dir sprintf('neptune_settings_%s',clk_settings)]),'Choices saved','custom',msgicon); clear clk_settings
    refresh_html_log
end
end

%% '06. Denoise-1 (non-cord regression)'
if choices_preproc(6)==1
try figure(status_fig),subplot(N,2,11),imshow(imread('status_images/step_06_blue.jpeg')), subplot(N,2,12),imshow(imread('status_images/status_blue.jpg')), catch, end
L=L+2; logg{L,1}=sprintf('-------\n%s',preprocsteps{6}); try fprintf(fileID,'%s\n',logg{L,1}); set(S.tx,'string',logg); catch, end; tc=tic;
time_taken_preproc.step06_denoise1 = zeros(size(func_files,1),size(func_files,2));
if (save_videos==0)||(save_videos==1), save_videos2 = save_videos; elseif (save_videos==2)&&(sum([choices_preproc(8:9);choices_preproc(11:14)])==0), save_videos2 = 1; else, save_videos2 = 0; end

skip=0;
for k=1:length(sub_dir)
    for k2=1:runs(k)
        if ~( (exist([sub_dir{k} func_files{k,k2} '_denoised_before_MC80.nii'], 'file')) || (exist([sub_dir{k} func_files{k,k2} '_slice' num2str(slices) '_denoised_before_MC80.nii'], 'file')) || (exist([sub_dir{k} func_files{k,k2} '_denoised_before_MC80.nii.gz'], 'file')) || (exist([sub_dir{k} func_files{k,k2} '_slice' num2str(slices) '_denoised_before_MC80.nii.gz'], 'file')) ) || (run_allsubs_by_default==1)
            skip=skip+1;
            clk=clock; L=L+1; logg{L,1}=sprintf('%d/%s/%d %dh:%dm:%ds - begin (06) denoise-1 (non-cord regression) for subject (%d) of (%d), run (%d) of (%d)',clk(3),month{clk(2)},clk(1),clk(4),clk(5),round(clk(6)),k,length(sub_dir),k2,runs(k)); try fprintf(fileID,'%s\n',logg{L,1}); set(S.tx,'string',logg); catch, end; clear clk
            try waitbar(((runs(k)*(k-1)+k2)/(length(sub_dir)*runs(k))),wbar2,sprintf('06. Denoise-1 (non-cord regression): Subject (%d) of (%d), Run (%d) of (%d)',k,length(sub_dir),k2,runs(k))); catch, end
            fprintf('06. Denoise-1 (non-cord regression): Subject (%d) of (%d), Run (%d) of (%d)\n',k,length(sub_dir),k2,runs(k))
            if runs(k)==1, Rns=0; else, Rns=k2; end
            time_06_denoise1 = tic;
            scfMRItb_06_denoise1(sub_dir{k},func_files{k,k2}, wbar3,save_videos2,Rns);
            time_taken_preproc.step06_denoise1(k,k2) = toc(time_06_denoise1); clear time_06_denoise1 Rns
            clk=clock; L=L+1; logg{L,1}=sprintf('%d/%s/%d %dh:%dm:%ds - end (06) denoise-1 (non-cord regression) for subject (%d) of (%d), run (%d) of (%d)',clk(3),month{clk(2)},clk(1),clk(4),clk(5),round(clk(6)),k,length(sub_dir),k2,runs(k)); try fprintf(fileID,'%s\n',logg{L,1}); set(S.tx,'string',logg); catch, end; clear clk
            refresh_html_log
        else
            scfMRItb_07_baseimage_if_no_06denoise1(sub_dir{k},func_files{k,k2}, wbar3);
            clk=clock; L=L+1; logg{L,1}=sprintf('%d/%s/%d %dh:%dm:%ds - skip (06) denoise-1 (non-cord regression) for subject (%d) of (%d), run (%d) of (%d)',clk(3),month{clk(2)},clk(1),clk(4),clk(5),round(clk(6)),k,length(sub_dir),k2,runs(k)); try fprintf(fileID,'%s\n',logg{L,1}); set(S.tx,'string',logg); catch, end; clear clk
        end
    end; clear k2
end; clear k save_videos2

if skip==0
    clk=clock; L=L+1; logg{L,1}=sprintf('%d/%s/%d %dh:%dm:%ds - (06) denoise-1 (non-cord regression) SKIPPED FOR ALL SUBJECTS',clk(3),month{clk(2)},clk(1),clk(4),clk(5),round(clk(6))); try fprintf(fileID,'%s\n',logg{L,1}); set(S.tx,'string',logg); catch, end; clear clk
    try figure(status_fig),subplot(N,2,11),imshow(imread('status_images/step_06_green.jpeg')), subplot(N,2,12),imshow(imread('status_images/status_green_skip.jpg')), catch, end
else
    clk=clock; L=L+1; logg{L,1}=sprintf('%d/%s/%d %dh:%dm:%ds - (06) denoise-1 (non-cord regression) COMPLETED FOR ALL SUBJECTS',clk(3),month{clk(2)},clk(1),clk(4),clk(5),round(clk(6))); try fprintf(fileID,'%s\n',logg{L,1}); set(S.tx,'string',logg); catch, end; clear clk
    L=L+1; logg{L,1}=sprintf('Total time taken so far: %dh:%dm:%ds\nTime taken for this step: %dh:%dm:%ds\nTime taken for this step in each run of each subject: %dh:%dm:%ds',floor(toc(tc0)/3600),floor(mod(toc(tc0),3600)/60),floor(mod(toc(tc0),60)),floor(toc(tc)/3600),floor(mod(toc(tc),3600)/60),floor(mod(toc(tc),60)),floor(toc(tc)/3600/skip),floor(mod(toc(tc),3600)/60/skip),floor(mod(toc(tc),60)/skip)); try fprintf(fileID,'%s\n',logg{L,1}); set(S.tx,'string',logg); catch, end; clear clk
    try figure(status_fig),subplot(N,2,11),imshow(imread('status_images/step_06_green.jpeg')), subplot(N,2,12),imshow(imread('status_images/status_green.jpg')), catch, end
    refresh_html
end
end

%% '07. Compute Gaussian mask'
if choices_preproc(7)==1
try figure(status_fig),subplot(N,2,13),imshow(imread('status_images/step_07_blue.jpeg')), subplot(N,2,14),
if automatic_GaussianMask==0, imshow(imread('status_images/status_blue_UI.jpg')), else, imshow(imread('status_images/status_blue.jpg')), end, catch, end
L=L+2; logg{L,1}=sprintf('-------\n%s',preprocsteps{7}); try fprintf(fileID,'%s\n',logg{L,1}); set(S.tx,'string',logg); catch, end; tc=tic;
time_taken_preproc.step07_gaussianMask = zeros(size(func_files,1),size(func_files,2));

skip=0;
for k=1:length(sub_dir)
    for k2=1:runs(k)
        if ~( (exist([sub_dir{k} func_files{k,k2} '_Gaussian_mask.nii'], 'file')) || (exist([sub_dir{k} func_files{k,k2} '_slice' num2str(slices) '_Gaussian_mask.nii'], 'file')) || (exist([sub_dir{k} func_files{k,k2} '_Gaussian_mask.nii.gz'], 'file')) || (exist([sub_dir{k} func_files{k,k2} '_slice' num2str(slices) '_Gaussian_mask.nii.gz'], 'file')) ) || (run_allsubs_by_default==1)
            skip=skip+1;
            clk=clock; L=L+1; logg{L,1}=sprintf('%d/%s/%d %dh:%dm:%ds - begin (07) compute Gaussian mask for subject (%d) of (%d), run (%d) of (%d)',clk(3),month{clk(2)},clk(1),clk(4),clk(5),round(clk(6)),k,length(sub_dir),k2,runs(k)); try fprintf(fileID,'%s\n',logg{L,1}); set(S.tx,'string',logg); catch, end; clear clk
            try waitbar(((runs(k)*(k-1)+k2)/(length(sub_dir)*runs(k))),wbar2,sprintf('07. Compute Gaussian mask: Subject (%d) of (%d), Run (%d) of (%d)',k,length(sub_dir),k2,runs(k))); catch, end
            fprintf('07. Compute Gaussian mask: Subject (%d) of (%d), Run (%d) of (%d)\n',k,length(sub_dir),k2,runs(k))
            if runs(k)==1, Rns=0; else, Rns=k2; end
            time_07_gaussianMask = tic;
            if choices_preproc(6)==0
                scfMRItb_07_baseimage_if_no_06denoise1(sub_dir{k},func_files{k,k2}, wbar3);
            end
            scfMRItb_07_gaussianMask(sub_dir{k},func_files{k,k2},automatic_GaussianMask, wbar3,Rns);
            time_taken_preproc.step07_gaussianMask(k,k2) = toc(time_07_gaussianMask); clear time_07_gaussianMask Rns
            clk=clock; L=L+1; logg{L,1}=sprintf('%d/%s/%d %dh:%dm:%ds - end (07) compute Gaussian mask for subject (%d) of (%d), run (%d) of (%d)',clk(3),month{clk(2)},clk(1),clk(4),clk(5),round(clk(6)),k,length(sub_dir),k2,runs(k)); try fprintf(fileID,'%s\n',logg{L,1}); set(S.tx,'string',logg); catch, end; clear clk
            refresh_html_log
        else
            clk=clock; L=L+1; logg{L,1}=sprintf('%d/%s/%d %dh:%dm:%ds - skip (07) compute Gaussian mask for subject (%d) of (%d), run (%d) of (%d)',clk(3),month{clk(2)},clk(1),clk(4),clk(5),round(clk(6)),k,length(sub_dir),k2,runs(k)); try fprintf(fileID,'%s\n',logg{L,1}); set(S.tx,'string',logg); catch, end; clear clk
        end
    end; clear k2
end

if skip==0
    clk=clock; L=L+1; logg{L,1}=sprintf('%d/%s/%d %dh:%dm:%ds - (07) compute Gaussian mask SKIPPED FOR ALL SUBJECTS',clk(3),month{clk(2)},clk(1),clk(4),clk(5),round(clk(6))); try fprintf(fileID,'%s\n',logg{L,1}); set(S.tx,'string',logg); catch, end; clear clk
    try figure(status_fig),subplot(N,2,13),imshow(imread('status_images/step_07_green.jpeg')), subplot(N,2,14),imshow(imread('status_images/status_green_skip.jpg')), catch, end
else
    clk=clock; L=L+1; logg{L,1}=sprintf('%d/%s/%d %dh:%dm:%ds - (07) compute Gaussian mask COMPLETED FOR ALL SUBJECTS',clk(3),month{clk(2)},clk(1),clk(4),clk(5),round(clk(6))); try fprintf(fileID,'%s\n',logg{L,1}); set(S.tx,'string',logg); catch, end; clear clk
    L=L+1; logg{L,1}=sprintf('Total time taken so far: %dh:%dm:%ds\nTime taken for this step: %dh:%dm:%ds\nTime taken for this step in each run of each subject: %dh:%dm:%ds',floor(toc(tc0)/3600),floor(mod(toc(tc0),3600)/60),floor(mod(toc(tc0),60)),floor(toc(tc)/3600),floor(mod(toc(tc),3600)/60),floor(mod(toc(tc),60)),floor(toc(tc)/3600/skip),floor(mod(toc(tc),3600)/60/skip),floor(mod(toc(tc),60)/skip)); try fprintf(fileID,'%s\n',logg{L,1}); set(S.tx,'string',logg); catch, end; clear clk
    try figure(status_fig),subplot(N,2,13),imshow(imread('status_images/step_07_green.jpeg')), subplot(N,2,14),imshow(imread('status_images/status_green.jpg')), catch, end
    refresh_html_log
end
end

%% '08. Motion correction'
if choices_preproc(8)==1
try figure(status_fig),subplot(N,2,15),imshow(imread('status_images/step_08_blue.jpeg')), subplot(N,2,16),imshow(imread('status_images/status_blue.jpg')), catch, end
L=L+2; logg{L,1}=sprintf('-------\n%s',preprocsteps{8}); try fprintf(fileID,'%s\n',logg{L,1}); set(S.tx,'string',logg); catch, end; tc=tic;
time_taken_preproc.step08_motionCorrection = zeros(size(func_files,1),size(func_files,2));
if (save_videos==0)||(save_videos==1), save_videos2 = save_videos; elseif (save_videos==2)&&(sum([choices_preproc(9);choices_preproc(11:14)])==0), save_videos2 = 1; else, save_videos2 = 0; end

skip=0;
for k=1:length(sub_dir)
    for k2=1:runs(k)
        if ~( (exist([sub_dir{k} func_files{k,k2} '_MC.nii'], 'file')) || (exist([sub_dir{k} func_files{k,k2} '_slice' num2str(slices) '_MC.nii'], 'file')) || (exist([sub_dir{k} func_files{k,k2} '_MC.nii.gz'], 'file')) || (exist([sub_dir{k} func_files{k,k2} '_slice' num2str(slices) '_MC.nii.gz'], 'file')) ) || (run_allsubs_by_default==1)
            skip=skip+1;
            clk=clock; L=L+1; logg{L,1}=sprintf('%d/%s/%d %dh:%dm:%ds - begin (08) motion correction for subject (%d) of (%d), run (%d) of (%d)',clk(3),month{clk(2)},clk(1),clk(4),clk(5),round(clk(6)),k,length(sub_dir),k2,runs(k)); try fprintf(fileID,'%s\n',logg{L,1}); set(S.tx,'string',logg); catch, end; clear clk
            try waitbar(((runs(k)*(k-1)+k2)/(length(sub_dir)*runs(k))),wbar2,sprintf('08. Motion correction: Subject (%d) of (%d), Run (%d) of (%d)',k,length(sub_dir),k2,runs(k))); catch, end
            fprintf('08. Motion correction: Subject (%d) of (%d), Run (%d) of (%d)\n',k,length(sub_dir),k2,runs(k))
            if runs(k)==1, Rns=0; else, Rns=k2; end
            time_08_motionCorrection = tic;
            scfMRItb_08_motionCorrection(sub_dir{k},func_files{k,k2}, motion_param_smoothing,wbar3,save_videos2,Rns);
            time_taken_preproc.step08_motionCorrection(k,k2) = toc(time_08_motionCorrection); clear time_08_motionCorrection Rns
            clk=clock; L=L+1; logg{L,1}=sprintf('%d/%s/%d %dh:%dm:%ds - end (08) motion correction for subject (%d) of (%d), run (%d) of (%d)',clk(3),month{clk(2)},clk(1),clk(4),clk(5),round(clk(6)),k,length(sub_dir),k2,runs(k)); try fprintf(fileID,'%s\n',logg{L,1}); set(S.tx,'string',logg); catch, end; clear clk
            refresh_html_log
        else
            clk=clock; L=L+1; logg{L,1}=sprintf('%d/%s/%d %dh:%dm:%ds - skip (08) motion correction for subject (%d) of (%d), run (%d) of (%d)',clk(3),month{clk(2)},clk(1),clk(4),clk(5),round(clk(6)),k,length(sub_dir),k2,runs(k)); try fprintf(fileID,'%s\n',logg{L,1}); set(S.tx,'string',logg); catch, end; clear clk
        end
    end; clear k2
end; clear k save_videos2

if skip==0
    clk=clock; L=L+1; logg{L,1}=sprintf('%d/%s/%d %dh:%dm:%ds - (08) motion correction SKIPPED FOR ALL SUBJECTS',clk(3),month{clk(2)},clk(1),clk(4),clk(5),round(clk(6))); try fprintf(fileID,'%s\n',logg{L,1}); set(S.tx,'string',logg); catch, end; clear clk
    try figure(status_fig),subplot(N,2,15),imshow(imread('status_images/step_08_green.jpeg')), subplot(N,2,16),imshow(imread('status_images/status_green_skip.jpg')), catch, end
else
    clk=clock; L=L+1; logg{L,1}=sprintf('%d/%s/%d %dh:%dm:%ds - (08) motion correction COMPLETED FOR ALL SUBJECTS',clk(3),month{clk(2)},clk(1),clk(4),clk(5),round(clk(6))); try fprintf(fileID,'%s\n',logg{L,1}); set(S.tx,'string',logg); catch, end; clear clk
    L=L+1; logg{L,1}=sprintf('Total time taken so far: %dh:%dm:%ds\nTime taken for this step: %dh:%dm:%ds\nTime taken for this step in each run of each subject: %dh:%dm:%ds',floor(toc(tc0)/3600),floor(mod(toc(tc0),3600)/60),floor(mod(toc(tc0),60)),floor(toc(tc)/3600),floor(mod(toc(tc),3600)/60),floor(mod(toc(tc),60)),floor(toc(tc)/3600/skip),floor(mod(toc(tc),3600)/60/skip),floor(mod(toc(tc),60)/skip)); try fprintf(fileID,'%s\n',logg{L,1}); set(S.tx,'string',logg); catch, end; clear clk
    try figure(status_fig),subplot(N,2,15),imshow(imread('status_images/step_08_green.jpeg')), subplot(N,2,16),imshow(imread('status_images/status_green.jpg')), catch, end
    refresh_html_log
end
end

%% '09. Denoise-2 (RETROICOR/3dDespike)'
if choices_preproc(9)==1
try figure(status_fig),subplot(N,2,17),imshow(imread('status_images/step_09_blue.jpeg')), subplot(N,2,18),imshow(imread('status_images/status_blue.jpg')), catch, end
L=L+2; logg{L,1}=sprintf('-------\n%s',preprocsteps{9}); try fprintf(fileID,'%s\n',logg{L,1}); set(S.tx,'string',logg); catch, end; tc=tic;
time_taken_preproc.step09_retroicor = zeros(size(func_files,1),size(func_files,2));
if (save_videos==0)||(save_videos==1), save_videos2 = save_videos; elseif (save_videos==2)&&(sum(choices_preproc(11:14))==0), save_videos2 = 1; else, save_videos2 = 0; end

skip=0;
for k=1:length(sub_dir)
    for k2=1:runs(k)
        if ~( (exist([sub_dir{k} func_files{k,k2} '_MC_ricor.nii'], 'file')) || (exist([sub_dir{k} func_files{k,k2} '_slice' num2str(slices) '_MC_ricor.nii'], 'file')) || (exist([sub_dir{k} func_files{k,k2} '_MC_ricor.nii.gz'], 'file')) || (exist([sub_dir{k} func_files{k,k2} '_slice' num2str(slices) '_MC_ricor.nii.gz'], 'file')) ) || (run_allsubs_by_default==1)
            skip=skip+1;
            clk=clock; L=L+1; logg{L,1}=sprintf('%d/%s/%d %dh:%dm:%ds - begin (09) Denoise-2 (RETROICOR/3dDespike) for subject (%d) of (%d), run (%d) of (%d)',clk(3),month{clk(2)},clk(1),clk(4),clk(5),round(clk(6)),k,length(sub_dir),k2,runs(k)); try fprintf(fileID,'%s\n',logg{L,1}); set(S.tx,'string',logg); catch, end; clear clk
            try waitbar(((runs(k)*(k-1)+k2)/(length(sub_dir)*runs(k))),wbar2,sprintf('09. Denoise-2 (RETROICOR/3dDespike): Subject (%d) of (%d), Run (%d) of (%d)',k,length(sub_dir),k2,runs(k))); catch, end
            fprintf('09. Denoise-2 (RETROICOR/3dDespike): Subject (%d) of (%d), Run (%d) of (%d)\n',k,length(sub_dir),k2,runs(k))
            if runs(k)==1, Rns=0; else, Rns=k2; end
            time_09_retroicor = tic;
            noRETROICOR = scfMRItb_09_retroicor(sub_dir{k},func_files{k,k2},scanner, wbar3,save_videos2,Rns);
            time_taken_preproc.step09_retroicor(k,k2) = toc(time_09_retroicor); clear time_09_retroicor Rns
            if noRETROICOR==1
                clk=clock; L=L+1; logg{L,1}=sprintf('%d/%s/%d %dh:%dm:%ds - .log file (physiological data) unavailable or in incorrect format, and hence RETROICOR was not performed for subject (%d) of (%d), run (%d) of (%d). 3dDespike was performed instead.',clk(3),month{clk(2)},clk(1),clk(4),clk(5),round(clk(6)),k,length(sub_dir),k2,runs(k)); try fprintf(fileID,'%s\n',logg{L,1}); set(S.tx,'string',logg); catch, end; clear clk
                disp(sprintf('Denoise-2: RETROICOR was not performed for subject (%d) run (%d), because the .log file with physiological data was not found or was in incorrect format. 3dDespike was performed instead.',k,k2)) %#ok<DSPS>
                if skip==1
                    try figure(status_fig),subplot(N,2,17),imshow(imread('status_images/step_09_blue_2.jpeg')), subplot(N,2,18),imshow(imread('status_images/status_blue.jpg')), catch, end
                end
            clk=clock; L=L+1; logg{L,1}=sprintf('%d/%s/%d %dh:%dm:%ds - end (09) Denoise-2 (3dDespike) for subject (%d) of (%d), run (%d) of (%d)',clk(3),month{clk(2)},clk(1),clk(4),clk(5),round(clk(6)),k,length(sub_dir),k2,runs(k)); try fprintf(fileID,'%s\n',logg{L,1}); set(S.tx,'string',logg); catch, end; clear clk
            else
            clk=clock; L=L+1; logg{L,1}=sprintf('%d/%s/%d %dh:%dm:%ds - end (09) Denoise-2 (RETROICOR) for subject (%d) of (%d), run (%d) of (%d)',clk(3),month{clk(2)},clk(1),clk(4),clk(5),round(clk(6)),k,length(sub_dir),k2,runs(k)); try fprintf(fileID,'%s\n',logg{L,1}); set(S.tx,'string',logg); catch, end; clear clk
            end
            refresh_html_log
        else
            clk=clock; L=L+1; logg{L,1}=sprintf('%d/%s/%d %dh:%dm:%ds - skip (09) Denoise-2 (RETROICOR/3dDespike) for subject (%d) of (%d), run (%d) of (%d)',clk(3),month{clk(2)},clk(1),clk(4),clk(5),round(clk(6)),k,length(sub_dir),k2,runs(k)); try fprintf(fileID,'%s\n',logg{L,1}); set(S.tx,'string',logg); catch, end; clear clk
        end
    end; clear k2
end; clear k save_videos2

if skip==0
    clk=clock; L=L+1; logg{L,1}=sprintf('%d/%s/%d %dh:%dm:%ds - (09) Denoise-2 (RETROICOR/3dDespike) SKIPPED FOR ALL SUBJECTS',clk(3),month{clk(2)},clk(1),clk(4),clk(5),round(clk(6))); try fprintf(fileID,'%s\n',logg{L,1}); set(S.tx,'string',logg); catch, end; clear clk
    try figure(status_fig),subplot(N,2,17),imshow(imread('status_images/step_09_green.jpeg')), subplot(N,2,18),imshow(imread('status_images/status_green_skip.jpg')), catch, end
else
    if noRETROICOR==0
    clk=clock; L=L+1; logg{L,1}=sprintf('%d/%s/%d %dh:%dm:%ds - (09) Denoise-2 (RETROICOR/3dDespike) COMPLETED FOR ALL SUBJECTS',clk(3),month{clk(2)},clk(1),clk(4),clk(5),round(clk(6))); try fprintf(fileID,'%s\n',logg{L,1}); set(S.tx,'string',logg); catch, end; clear clk
    L=L+1; logg{L,1}=sprintf('Total time taken so far: %dh:%dm:%ds\nTime taken for this step: %dh:%dm:%ds\nTime taken for this step in each run of each subject: %dh:%dm:%ds',floor(toc(tc0)/3600),floor(mod(toc(tc0),3600)/60),floor(mod(toc(tc0),60)),floor(toc(tc)/3600),floor(mod(toc(tc),3600)/60),floor(mod(toc(tc),60)),floor(toc(tc)/3600/skip),floor(mod(toc(tc),3600)/60/skip),floor(mod(toc(tc),60)/skip)); try fprintf(fileID,'%s\n',logg{L,1}); set(S.tx,'string',logg); catch, end; clear clk
    try figure(status_fig),subplot(N,2,17),imshow(imread('status_images/step_09_green.jpeg')), subplot(N,2,18),imshow(imread('status_images/status_green.jpg')), catch, end
    else
    clk=clock; L=L+1; logg{L,1}=sprintf('%d/%s/%d %dh:%dm:%ds - (09) Denoise-2 (3dDespike) COMPLETED FOR ALL SUBJECTS',clk(3),month{clk(2)},clk(1),clk(4),clk(5),round(clk(6))); try fprintf(fileID,'%s\n',logg{L,1}); set(S.tx,'string',logg); catch, end; clear clk
    L=L+1; logg{L,1}=sprintf('Total time taken so far: %dh:%dm:%ds\nTime taken for this step: %dh:%dm:%ds\nTime taken for this step in each run of each subject: %dh:%dm:%ds',floor(toc(tc0)/3600),floor(mod(toc(tc0),3600)/60),floor(mod(toc(tc0),60)),floor(toc(tc)/3600),floor(mod(toc(tc),3600)/60),floor(mod(toc(tc),60)),floor(toc(tc)/3600/skip),floor(mod(toc(tc),3600)/60/skip),floor(mod(toc(tc),60)/skip)); try fprintf(fileID,'%s\n',logg{L,1}); set(S.tx,'string',logg); catch, end; clear clk
    try figure(status_fig),subplot(N,2,17),imshow(imread('status_images/step_09_green_2.jpeg')), subplot(N,2,18),imshow(imread('status_images/status_green.jpg')), catch, end
    end
    refresh_html_log
end
end

%% '10. Define GM, WM and CSF masks'
if choices_preproc(10)==1
try figure(status_fig),subplot(N,2,19),imshow(imread('status_images/step_10_blue.jpeg')), subplot(N,2,20),imshow(imread('status_images/status_blue_UI.jpg')), catch, end
L=L+2; logg{L,1}=sprintf('-------\n%s',preprocsteps{10}); try fprintf(fileID,'%s\n',logg{L,1}); set(S.tx,'string',logg); catch, end; tc=tic;

skip=0;
if UIstepsFirst~=1
time_taken_preproc.step10_GMWMCSFmasks_detailed = cell(length(sub_dir),1);
time_taken_preproc.step10_GMWMCSFmasks = zeros(size(func_files,1),1);
end
for k=1:length(sub_dir)
    if ~( exist([sub_dir{k} anat_files{k,1} '_reduced' '_mask_GM.nii'], 'file') || exist([sub_dir{k} anat_files{k,1} '_reduced' '_mask_GM.nii.gz'], 'file') ) || (run_allsubs_by_default==1)
        skip=skip+1;
        clk=clock; L=L+1; logg{L,1}=sprintf('%d/%s/%d %dh:%dm:%ds - begin (10) define GM, WM and CSF masks for subject (%d) of (%d), run (1) of (%d)',clk(3),month{clk(2)},clk(1),clk(4),clk(5),round(clk(6)),k,length(sub_dir),runs(k)); try fprintf(fileID,'%s\n',logg{L,1}); set(S.tx,'string',logg); catch, end; clear clk
        try waitbar((k*1/length(sub_dir)*runs(k)),wbar2,sprintf('10. Define GM, WM and CSF masks: Subject (%d) of (%d), Run (1) of (%d)',k,length(sub_dir),runs(k))); catch, end
        fprintf('10. Define GM, WM and CSF masks: Subject (%d) of (%d), Run (1) of (%d)\n',k,length(sub_dir),runs(k))
        time_10_GMWMCSFmasks = tic;
        time_taken_preproc.step10_GMWMCSFmasks_detailed{k,1} = scfMRItb_10_GMWMCSFmasks(sub_dir{k},func_files{k,1},anat_files{k,1},wbar3,exist_SCT,mask_GM_flag,choices_preproc,C_spine, SCT_vert_auto,SCT_manualreg_labels);
        time_taken_preproc.step10_GMWMCSFmasks(k,1) = toc(time_10_GMWMCSFmasks); clear time_10_GMWMCSFmasks
        clk=clock; L=L+1; logg{L,1}=sprintf('%d/%s/%d %dh:%dm:%ds - end (10) define GM, WM and CSF masks for subject (%d) of (%d), run (1) of (%d)',clk(3),month{clk(2)},clk(1),clk(4),clk(5),round(clk(6)),k,length(sub_dir),runs(k)); try fprintf(fileID,'%s\n',logg{L,1}); set(S.tx,'string',logg); catch, end; clear clk
        refresh_html_log
    else
        if UIstepsFirst==1
            clk=clock; L=L+1; logg{L,1}=sprintf('%d/%s/%d %dh:%dm:%ds - begin remnants of step-10 (GM, WM and CSF masks) for subject (%d) of (%d), run (1) of (%d)',clk(3),month{clk(2)},clk(1),clk(4),clk(5),round(clk(6)),k,length(sub_dir),runs(k)); try fprintf(fileID,'%s\n',logg{L,1}); set(S.tx,'string',logg); catch, end; clear clk
            try waitbar((k*1/length(sub_dir)*runs(k)),wbar2,sprintf('10. Remnants of step-10 (GM, WM and CSF masks): Subject (%d) of (%d), Run (1) of (%d)',k,length(sub_dir),runs(k))); catch, end
            fprintf('10. Remnants of step-10 (GM, WM and CSF masks): Subject (%d) of (%d), Run (1) of (%d)\n',k,length(sub_dir),runs(k))
        else
            clk=clock; L=L+1; logg{L,1}=sprintf('%d/%s/%d %dh:%dm:%ds - skip remnants of step-10 (GM, WM and CSF masks) for subject (%d) of (%d), run (1) of (%d)',clk(3),month{clk(2)},clk(1),clk(4),clk(5),round(clk(6)),k,length(sub_dir),runs(k)); try fprintf(fileID,'%s\n',logg{L,1}); set(S.tx,'string',logg); catch, end; clear clk
        end
        scfMRItb_10_GMWMCSFmasks_subset(sub_dir{k},func_files{k,1},anat_files{k,1},choices_preproc);
    end
    for k2=2:runs(k)
        if ~( (exist([sub_dir{k} func_files{k,k2} '_base_masked.nii'], 'file')) || (exist([sub_dir{k} func_files{k,k2} '_slice' num2str(slices) '_base_masked.nii'], 'file')) || (exist([sub_dir{k} func_files{k,k2} '_base_masked.nii.gz'], 'file')) || (exist([sub_dir{k} func_files{k,k2} '_slice' num2str(slices) '_base_masked.nii.gz'], 'file')) ) || (run_allsubs_by_default==1)
            skip=skip+1;
            if UIstepsFirst~=1
                clk=clock; L=L+1; logg{L,1}=sprintf('%d/%s/%d %dh:%dm:%ds - begin (10) define GM, WM and CSF masks for subject (%d) of (%d), run (%d) of (%d)',clk(3),month{clk(2)},clk(1),clk(4),clk(5),round(clk(6)),k,length(sub_dir),k2,runs(k)); try fprintf(fileID,'%s\n',logg{L,1}); set(S.tx,'string',logg); catch, end; clear clk
                try waitbar(((runs(k)*(k-1)+k2)/(length(sub_dir)*runs(k))),wbar2,sprintf('10. Define GM, WM and CSF masks: Subject (%d) of (%d), Run (%d) of (%d)',k,length(sub_dir),k2,runs(k))); catch, end
                fprintf('10. Define GM, WM and CSF masks: Subject (%d) of (%d), Run (%d) of (%d)\n',k,length(sub_dir),k2,runs(k))
            else
                clk=clock; L=L+1; logg{L,1}=sprintf('%d/%s/%d %dh:%dm:%ds - begin remnants of step-10 (GM, WM and CSF masks) for subject (%d) of (%d), run (%d) of (%d)',clk(3),month{clk(2)},clk(1),clk(4),clk(5),round(clk(6)),k,length(sub_dir),k2,runs(k)); try fprintf(fileID,'%s\n',logg{L,1}); set(S.tx,'string',logg); catch, end; clear clk
                try waitbar((k*1/length(sub_dir)*runs(k)),wbar2,sprintf('10. Remnants of step-10 (GM, WM and CSF masks): Subject (%d) of (%d), Run (%d) of (%d)',k,length(sub_dir),k2,runs(k))); catch, end
                fprintf('10. Remnants of step-10 (GM, WM and CSF masks): Subject (%d) of (%d), Run (%d) of (%d)\n',k,length(sub_dir),k2,runs(k))
            end
            scfMRItb_10_GMWMCSFmasks_run2(sub_dir{k},func_files{k,k2},wbar3);
            if UIstepsFirst~=1
                clk=clock; L=L+1; logg{L,1}=sprintf('%d/%s/%d %dh:%dm:%ds - end (10) define GM, WM and CSF masks for subject (%d) of (%d), run (%d) of (%d)',clk(3),month{clk(2)},clk(1),clk(4),clk(5),round(clk(6)),k,length(sub_dir),k2,runs(k)); try fprintf(fileID,'%s\n',logg{L,1}); set(S.tx,'string',logg); catch, end; clear clk
            else
                clk=clock; L=L+1; logg{L,1}=sprintf('%d/%s/%d %dh:%dm:%ds - end remnants of step-10 (GM, WM and CSF masks) for subject (%d) of (%d), run (%d) of (%d)',clk(3),month{clk(2)},clk(1),clk(4),clk(5),round(clk(6)),k,length(sub_dir),k2,runs(k)); try fprintf(fileID,'%s\n',logg{L,1}); set(S.tx,'string',logg); catch, end; clear clk
            end
        else
            clk=clock; L=L+1; logg{L,1}=sprintf('%d/%s/%d %dh:%dm:%ds - skip (10) define GM, WM and CSF masks for subject (%d) of (%d), run (%d) of (%d)',clk(3),month{clk(2)},clk(1),clk(4),clk(5),round(clk(6)),k,length(sub_dir),k2,runs(k)); try fprintf(fileID,'%s\n',logg{L,1}); set(S.tx,'string',logg); catch, end; clear clk
        end
    end; clear k2
end

if skip==0
    if UIstepsFirst~=1
        clk=clock; L=L+1; logg{L,1}=sprintf('%d/%s/%d %dh:%dm:%ds - (10) define GM, WM and CSF masks SKIPPED FOR ALL SUBJECTS',clk(3),month{clk(2)},clk(1),clk(4),clk(5),round(clk(6))); try fprintf(fileID,'%s\n',logg{L,1}); set(S.tx,'string',logg); catch, end; clear clk
    else
        clk=clock; L=L+1; logg{L,1}=sprintf('%d/%s/%d %dh:%dm:%ds - remnants of step-10 (GM, WM and CSF masks) SKIPPED FOR ALL SUBJECTS',clk(3),month{clk(2)},clk(1),clk(4),clk(5),round(clk(6))); try fprintf(fileID,'%s\n',logg{L,1}); set(S.tx,'string',logg); catch, end; clear clk
    end
    try figure(status_fig),subplot(N,2,19),imshow(imread('status_images/step_10_green.jpeg')), subplot(N,2,20),imshow(imread('status_images/status_green_skip.jpg')), catch, end
else
    if UIstepsFirst~=1
        clk=clock; L=L+1; logg{L,1}=sprintf('%d/%s/%d %dh:%dm:%ds - (10) define GM, WM and CSF masks COMPLETED FOR ALL SUBJECTS',clk(3),month{clk(2)},clk(1),clk(4),clk(5),round(clk(6))); try fprintf(fileID,'%s\n',logg{L,1}); set(S.tx,'string',logg); catch, end; clear clk
    else
        clk=clock; L=L+1; logg{L,1}=sprintf('%d/%s/%d %dh:%dm:%ds - remnants of step-10 (GM, WM and CSF masks) COMPLETED FOR ALL SUBJECTS',clk(3),month{clk(2)},clk(1),clk(4),clk(5),round(clk(6))); try fprintf(fileID,'%s\n',logg{L,1}); set(S.tx,'string',logg); catch, end; clear clk
    end
    L=L+1; logg{L,1}=sprintf('Total time taken so far: %dh:%dm:%ds\nTime taken for this step: %dh:%dm:%ds\nTime taken for this step in each run of each subject: %dh:%dm:%ds',floor(toc(tc0)/3600),floor(mod(toc(tc0),3600)/60),floor(mod(toc(tc0),60)),floor(toc(tc)/3600),floor(mod(toc(tc),3600)/60),floor(mod(toc(tc),60)),floor(toc(tc)/3600/skip),floor(mod(toc(tc),3600)/60/skip),floor(mod(toc(tc),60)/skip)); try fprintf(fileID,'%s\n',logg{L,1}); set(S.tx,'string',logg); catch, end; clear clk
    try figure(status_fig),subplot(N,2,19),imshow(imread('status_images/step_10_green.jpeg')), subplot(N,2,20),imshow(imread('status_images/status_green.jpg')), catch, end
    refresh_html_log
end
end

%% '11. Co-registration (functional to anat.)'
if choices_preproc(11)==1
try figure(status_fig),subplot(N,2,21),imshow(imread('status_images/step_11_blue.jpeg')), subplot(N,2,22),imshow(imread('status_images/status_blue.jpg')), catch, end
L=L+2; logg{L,1}=sprintf('-------\n%s',preprocsteps{11}); try fprintf(fileID,'%s\n',logg{L,1}); set(S.tx,'string',logg); catch, end; tc=tic;
time_taken_preproc.step11_FuncAnatRegistration = zeros(size(func_files,1),size(func_files,2));
if (save_videos==0)||(save_videos==1), save_videos2 = save_videos; elseif (save_videos==2)&&(sum(choices_preproc(12:14))==0), save_videos2 = 1; else, save_videos2 = 0; end

skip=0;
for k=1:length(sub_dir)
    for k2=1:runs(k)
        if ~( (exist([sub_dir{k} func_files{k,k2} '_warped.nii'], 'file')) || (exist([sub_dir{k} func_files{k,k2} '_slice' num2str(slices) '_warped.nii'], 'file')) || (exist([sub_dir{k} func_files{k,k2} '_warped.nii.gz'], 'file')) || (exist([sub_dir{k} func_files{k,k2} '_slice' num2str(slices) '_warped.nii.gz'], 'file')) ) || (run_allsubs_by_default==1)
            skip=skip+1;
            clk=clock; L=L+1; logg{L,1}=sprintf('%d/%s/%d %dh:%dm:%ds - begin (11) co-registration (functional to anat.) for subject (%d) of (%d), run (%d) of (%d)',clk(3),month{clk(2)},clk(1),clk(4),clk(5),round(clk(6)),k,length(sub_dir),k2,runs(k)); try fprintf(fileID,'%s\n',logg{L,1}); set(S.tx,'string',logg); catch, end; clear clk
            try waitbar(((runs(k)*(k-1)+k2)/(length(sub_dir)*runs(k))),wbar2,sprintf('11. Co-registration (functional to anat.): Subject (%d) of (%d), Run (%d) of (%d)',k,length(sub_dir),k2,runs(k))); catch, end
            fprintf('11. Co-registration (functional to anat.): Subject (%d) of (%d), Run (%d) of (%d)\n',k,length(sub_dir),k2,runs(k))
            if runs(k)==1, Rns=0; else, Rns=k2; end
            time_11_FuncAnatRegistration = tic;
            scfMRItb_11_FuncAnatRegistration(sub_dir{k},func_files{k,k2},anat_files{k,1},k2,wbar3,save_videos2,Rns);
            time_taken_preproc.step11_FuncAnatRegistration(k,k2) = toc(time_11_FuncAnatRegistration); clear time_11_FuncAnatRegistration Rns
            clk=clock; L=L+1; logg{L,1}=sprintf('%d/%s/%d %dh:%dm:%ds - end (11) co-registration (functional to anat.) for subject (%d) of (%d), run (%d) of (%d)',clk(3),month{clk(2)},clk(1),clk(4),clk(5),round(clk(6)),k,length(sub_dir),k2,runs(k)); try fprintf(fileID,'%s\n',logg{L,1}); set(S.tx,'string',logg); catch, end; clear clk
            refresh_html_log
        else
            clk=clock; L=L+1; logg{L,1}=sprintf('%d/%s/%d %dh:%dm:%ds - skip (11) co-registration (functional to anat.) for subject (%d) of (%d), run (%d) of (%d)',clk(3),month{clk(2)},clk(1),clk(4),clk(5),round(clk(6)),k,length(sub_dir),k2,runs(k)); try fprintf(fileID,'%s\n',logg{L,1}); set(S.tx,'string',logg); catch, end; clear clk
        end
    end; clear k2
end; clear k save_videos2

if skip==0
    clk=clock; L=L+1; logg{L,1}=sprintf('%d/%s/%d %dh:%dm:%ds - (11) co-registration (functional to anat.) SKIPPED FOR ALL SUBJECTS',clk(3),month{clk(2)},clk(1),clk(4),clk(5),round(clk(6))); try fprintf(fileID,'%s\n',logg{L,1}); set(S.tx,'string',logg); catch, end; clear clk
    try figure(status_fig),subplot(N,2,21),imshow(imread('status_images/step_11_green.jpeg')), subplot(N,2,22),imshow(imread('status_images/status_green_skip.jpg')), catch, end
else
    clk=clock; L=L+1; logg{L,1}=sprintf('%d/%s/%d %dh:%dm:%ds - (11) co-registration (functional to anat.) COMPLETED FOR ALL SUBJECTS',clk(3),month{clk(2)},clk(1),clk(4),clk(5),round(clk(6))); try fprintf(fileID,'%s\n',logg{L,1}); set(S.tx,'string',logg); catch, end; clear clk
    L=L+1; logg{L,1}=sprintf('Total time taken so far: %dh:%dm:%ds\nTime taken for this step: %dh:%dm:%ds\nTime taken for this step in each run of each subject: %dh:%dm:%ds',floor(toc(tc0)/3600),floor(mod(toc(tc0),3600)/60),floor(mod(toc(tc0),60)),floor(toc(tc)/3600),floor(mod(toc(tc),3600)/60),floor(mod(toc(tc),60)),floor(toc(tc)/3600/skip),floor(mod(toc(tc),3600)/60/skip),floor(mod(toc(tc),60)/skip)); try fprintf(fileID,'%s\n',logg{L,1}); set(S.tx,'string',logg); catch, end; clear clk
    try figure(status_fig),subplot(N,2,21),imshow(imread('status_images/step_11_green.jpeg')), subplot(N,2,22),imshow(imread('status_images/status_green.jpg')), catch, end
    refresh_html
end
end

%% '12. Denoise-3 (CSF regression)'
if choices_preproc(12)==1
try figure(status_fig),subplot(N,2,23),imshow(imread('status_images/step_12_blue.jpeg')), subplot(N,2,24),imshow(imread('status_images/status_blue.jpg')), catch, end
L=L+2; logg{L,1}=sprintf('-------\n%s',preprocsteps{12}); try fprintf(fileID,'%s\n',logg{L,1}); set(S.tx,'string',logg); catch, end; tc=tic;
time_taken_preproc.step12_CSFregress = zeros(size(func_files,1),size(func_files,2));
if (save_videos==0)||(save_videos==1), save_videos2 = save_videos; elseif (save_videos==2)&&(sum(choices_preproc(13:14))==0), save_videos2 = 2; else, save_videos2 = 0; end

skip=0;
for k=1:length(sub_dir)
    for k2=1:runs(k)
        try load([sub_dir{k} sprintf('workspace_variables_run%d.mat',k2)], 'cutoff_str'); catch, cutoff_str = '50'; end
        if ~( (exist([sub_dir{k} func_files{k,k2} '_denoised' cutoff_str '.nii'], 'file')) || (exist([sub_dir{k} func_files{k,k2} '_slice' num2str(slices) '_denoised' cutoff_str '.nii'], 'file')) || (exist([sub_dir{k} func_files{k,k2} '_denoised' cutoff_str '.nii.gz'], 'file')) || (exist([sub_dir{k} func_files{k,k2} '_slice' num2str(slices) '_denoised' cutoff_str '.nii.gz'], 'file')) ) || (run_allsubs_by_default==1)
            skip=skip+1;
            clk=clock; L=L+1; logg{L,1}=sprintf('%d/%s/%d %dh:%dm:%ds - begin (12) denoise-3 (CSF regression) for subject (%d) of (%d), run (%d) of (%d)',clk(3),month{clk(2)},clk(1),clk(4),clk(5),round(clk(6)),k,length(sub_dir),k2,runs(k)); try fprintf(fileID,'%s\n',logg{L,1}); set(S.tx,'string',logg); catch, end; clear clk
            try waitbar(((runs(k)*(k-1)+k2)/(length(sub_dir)*runs(k))),wbar2,sprintf('12. Denoise-3 (CSF regression): Subject (%d) of (%d), Run (%d) of (%d)',k,length(sub_dir),k2,runs(k))); catch, end
            fprintf('12. Denoise-3 (CSF regression): Subject (%d) of (%d), Run (%d) of (%d)\n',k,length(sub_dir),k2,runs(k))
            if runs(k)==1, Rns=0; else, Rns=k2; end
            time_12_CSFregress = tic;
            scfMRItb_12_CSFregress(sub_dir{k},func_files{k,k2},anat_files{k,1}, wbar3,save_videos2,Rns);
            time_taken_preproc.step12_CSFregress(k,k2) = toc(time_12_CSFregress); clear time_12_CSFregress Rns
            clk=clock; L=L+1; logg{L,1}=sprintf('%d/%s/%d %dh:%dm:%ds - end (12) denoise-3 (CSF regression) for subject (%d) of (%d), run (%d) of (%d)',clk(3),month{clk(2)},clk(1),clk(4),clk(5),round(clk(6)),k,length(sub_dir),k2,runs(k)); try fprintf(fileID,'%s\n',logg{L,1}); set(S.tx,'string',logg); catch, end; clear clk
            refresh_html_log
        else
            clk=clock; L=L+1; logg{L,1}=sprintf('%d/%s/%d %dh:%dm:%ds - skip (12) denoise-3 (CSF regression) for subject (%d) of (%d), run (%d) of (%d)',clk(3),month{clk(2)},clk(1),clk(4),clk(5),round(clk(6)),k,length(sub_dir),k2,runs(k)); try fprintf(fileID,'%s\n',logg{L,1}); set(S.tx,'string',logg); catch, end; clear clk
        end
    end; clear k2
end; clear k save_videos2

if skip==0
    clk=clock; L=L+1; logg{L,1}=sprintf('%d/%s/%d %dh:%dm:%ds - (12) denoise-3 (CSF regression) SKIPPED FOR ALL SUBJECTS',clk(3),month{clk(2)},clk(1),clk(4),clk(5),round(clk(6))); try fprintf(fileID,'%s\n',logg{L,1}); set(S.tx,'string',logg); catch, end; clear clk
    try figure(status_fig),subplot(N,2,23),imshow(imread('status_images/step_12_green.jpeg')), subplot(N,2,24),imshow(imread('status_images/status_green_skip.jpg')), catch, end
else
    clk=clock; L=L+1; logg{L,1}=sprintf('%d/%s/%d %dh:%dm:%ds - (12) denoise-3 (CSF regression) COMPLETED FOR ALL SUBJECTS',clk(3),month{clk(2)},clk(1),clk(4),clk(5),round(clk(6))); try fprintf(fileID,'%s\n',logg{L,1}); set(S.tx,'string',logg); catch, end; clear clk
    L=L+1; logg{L,1}=sprintf('Total time taken so far: %dh:%dm:%ds\nTime taken for this step: %dh:%dm:%ds\nTime taken for this step in each run of each subject: %dh:%dm:%ds',floor(toc(tc0)/3600),floor(mod(toc(tc0),3600)/60),floor(mod(toc(tc0),60)),floor(toc(tc)/3600),floor(mod(toc(tc),3600)/60),floor(mod(toc(tc),60)),floor(toc(tc)/3600/skip),floor(mod(toc(tc),3600)/60/skip),floor(mod(toc(tc),60)/skip)); try fprintf(fileID,'%s\n',logg{L,1}); set(S.tx,'string',logg); catch, end; clear clk
    try figure(status_fig),subplot(N,2,23),imshow(imread('status_images/step_12_green.jpeg')), subplot(N,2,24),imshow(imread('status_images/status_green.jpg')), catch, end
    refresh_html
end
end

%% '13. Denoise-4 (WM regression)'
if choices_preproc(13)==1
try figure(status_fig),subplot(N,2,25),imshow(imread('status_images/step_13_blue.jpeg')), subplot(N,2,26),imshow(imread('status_images/status_blue.jpg')), catch, end
L=L+2; logg{L,1}=sprintf('-------\n%s',preprocsteps{13}); try fprintf(fileID,'%s\n',logg{L,1}); set(S.tx,'string',logg); catch, end; tc=tic;
time_taken_preproc.step13_WMregress = zeros(size(func_files,1),size(func_files,2));
if (save_videos==0)||(save_videos==1), save_videos2 = save_videos; elseif (save_videos==2)&&(sum(choices_preproc(14:14))==0), save_videos2 = 2; else, save_videos2 = 0; end

skip=0;
for k=1:length(sub_dir)
    for k2=1:runs(k)
        try load([sub_dir{k} sprintf('workspace_variables_run%d.mat',k2)], 'cutoff_str'); catch, cutoff_str = '50'; end
        if ~( (exist([sub_dir{k} func_files{k,k2} '_denoised' cutoff_str 'WM.nii'], 'file')) || (exist([sub_dir{k} func_files{k,k2} '_slice' num2str(slices) '_denoised' cutoff_str 'WM.nii'], 'file')) || (exist([sub_dir{k} func_files{k,k2} '_denoised' cutoff_str 'WM.nii.gz'], 'file')) || (exist([sub_dir{k} func_files{k,k2} '_slice' num2str(slices) '_denoised' cutoff_str 'WM.nii.gz'], 'file'))  ||  (exist([sub_dir{k} func_files{k,k2} '_WM.nii'], 'file')) || (exist([sub_dir{k} func_files{k,k2} '_slice' num2str(slices) '_WM.nii'], 'file')) || (exist([sub_dir{k} func_files{k,k2} '_WM.nii.gz'], 'file')) || (exist([sub_dir{k} func_files{k,k2} '_slice' num2str(slices) '_WM.nii.gz'], 'file')) ) || (run_allsubs_by_default==1)
            skip=skip+1;
            clk=clock; L=L+1; logg{L,1}=sprintf('%d/%s/%d %dh:%dm:%ds - begin (13) Regress-out white-matter signal from GM for subject (%d) of (%d), run (%d) of (%d)',clk(3),month{clk(2)},clk(1),clk(4),clk(5),round(clk(6)),k,length(sub_dir),k2,runs(k)); try fprintf(fileID,'%s\n',logg{L,1}); set(S.tx,'string',logg); catch, end; clear clk
            try waitbar(((runs(k)*(k-1)+k2)/(length(sub_dir)*runs(k))),wbar2,sprintf('13. Regress-out white-matter signal from GM: Subject (%d) of (%d), Run (%d) of (%d)',k,length(sub_dir),k2,runs(k))); catch, end
            fprintf('13. Regress-out white-matter signal from GM: Subject (%d) of (%d), Run (%d) of (%d)\n',k,length(sub_dir),k2,runs(k))
            if runs(k)==1, Rns=0; else, Rns=k2; end
            time_13_WMregress = tic;
            scfMRItb_13_WMregress(sub_dir{k},func_files{k,k2},anat_files{k,1}, wbar3,save_videos2,Rns);
            time_taken_preproc.step13_WMregress(k,k2) = toc(time_13_WMregress); clear time_13_WMregress Rns
            clk=clock; L=L+1; logg{L,1}=sprintf('%d/%s/%d %dh:%dm:%ds - end (13) regress-out white-matter signal from GM for subject (%d) of (%d), run (%d) of (%d)',clk(3),month{clk(2)},clk(1),clk(4),clk(5),round(clk(6)),k,length(sub_dir),k2,runs(k)); try fprintf(fileID,'%s\n',logg{L,1}); set(S.tx,'string',logg); catch, end; clear clk
            refresh_html_log
        else
            clk=clock; L=L+1; logg{L,1}=sprintf('%d/%s/%d %dh:%dm:%ds - skip (13) regress-out white-matter signal from GM for subject (%d) of (%d), run (%d) of (%d)',clk(3),month{clk(2)},clk(1),clk(4),clk(5),round(clk(6)),k,length(sub_dir),k2,runs(k)); try fprintf(fileID,'%s\n',logg{L,1}); set(S.tx,'string',logg); catch, end; clear clk
        end
    end; clear k2
end; clear k save_videos2

if skip==0
    clk=clock; L=L+1; logg{L,1}=sprintf('%d/%s/%d %dh:%dm:%ds - (13) regress-out white-matter signal from GM SKIPPED FOR ALL SUBJECTS',clk(3),month{clk(2)},clk(1),clk(4),clk(5),round(clk(6))); try fprintf(fileID,'%s\n',logg{L,1}); set(S.tx,'string',logg); catch, end; clear clk
    try figure(status_fig),subplot(N,2,25),imshow(imread('status_images/step_13_green.jpeg')), subplot(N,2,26),imshow(imread('status_images/status_green_skip.jpg')), catch, end
else
    clk=clock; L=L+1; logg{L,1}=sprintf('%d/%s/%d %dh:%dm:%ds - (13) regress-out white-matter signal from GM COMPLETED FOR ALL SUBJECTS',clk(3),month{clk(2)},clk(1),clk(4),clk(5),round(clk(6))); try fprintf(fileID,'%s\n',logg{L,1}); set(S.tx,'string',logg); catch, end; clear clk
    L=L+1; logg{L,1}=sprintf('Total time taken so far: %dh:%dm:%ds\nTime taken for this step: %dh:%dm:%ds\nTime taken for this step in each run of each subject: %dh:%dm:%ds',floor(toc(tc0)/3600),floor(mod(toc(tc0),3600)/60),floor(mod(toc(tc0),60)),floor(toc(tc)/3600),floor(mod(toc(tc),3600)/60),floor(mod(toc(tc),60)),floor(toc(tc)/3600/skip),floor(mod(toc(tc),3600)/60/skip),floor(mod(toc(tc),60)/skip)); try fprintf(fileID,'%s\n',logg{L,1}); set(S.tx,'string',logg); catch, end; clear clk
    try figure(status_fig),subplot(N,2,25),imshow(imread('status_images/step_13_green.jpeg')), subplot(N,2,26),imshow(imread('status_images/status_green.jpg')), catch, end
    refresh_html
end
end

%% '14. Denoise-5 (additional covariates regression)'
if (choices_preproc(14)==1) && ~( covregress_options(1,1)==3 && covregress_options(2,1)==3 && covregress_options(3,1)==2 )
try figure(status_fig),subplot(N,2,27),imshow(imread('status_images/step_14_blue.jpeg')), subplot(N,2,28),imshow(imread('status_images/status_blue.jpg')), catch, end
L=L+2; logg{L,1}=sprintf('-------\n%s',preprocsteps{14}); try fprintf(fileID,'%s\n',logg{L,1}); set(S.tx,'string',logg); catch, end; tc=tic;
time_taken_preproc.step14_COVregress = zeros(size(func_files,1),size(func_files,2));
save_videos2 = save_videos;

skip=0;
for k=1:length(sub_dir)
    for k2=1:runs(k)
        try load([sub_dir{k} sprintf('workspace_variables_run%d.mat',k2)], 'cutoff_str'); catch, cutoff_str = '50'; end
        if ~( (exist([sub_dir{k} func_files{k,k2} '_denoised' cutoff_str 'WMcov.nii'], 'file')) || (exist([sub_dir{k} func_files{k,k2} '_slice' num2str(slices) '_denoised' cutoff_str 'WMcov.nii'], 'file')) || (exist([sub_dir{k} func_files{k,k2} '_denoised' cutoff_str 'WMcov.nii.gz'], 'file')) || (exist([sub_dir{k} func_files{k,k2} '_slice' num2str(slices) '_denoised' cutoff_str 'WMcov.nii.gz'], 'file'))  ||  (exist([sub_dir{k} func_files{k,k2} '_denoised' cutoff_str 'cov.nii'], 'file')) || (exist([sub_dir{k} func_files{k,k2} '_slice' num2str(slices) '_denoised' cutoff_str 'cov.nii'], 'file')) || (exist([sub_dir{k} func_files{k,k2} '_denoised' cutoff_str 'cov.nii.gz'], 'file')) || (exist([sub_dir{k} func_files{k,k2} '_slice' num2str(slices) '_denoised' cutoff_str 'cov.nii.gz'], 'file'))  ||  (exist([sub_dir{k} func_files{k,k2} '_WMcov.nii'], 'file')) || (exist([sub_dir{k} func_files{k,k2} '_slice' num2str(slices) '_WMcov.nii'], 'file')) || (exist([sub_dir{k} func_files{k,k2} '_WMcov.nii.gz'], 'file')) || (exist([sub_dir{k} func_files{k,k2} '_slice' num2str(slices) '_WMcov.nii.gz'], 'file'))  ||  (exist([sub_dir{k} func_files{k,k2} '_cov.nii'], 'file')) || (exist([sub_dir{k} func_files{k,k2} '_slice' num2str(slices) '_cov.nii'], 'file')) || (exist([sub_dir{k} func_files{k,k2} '_cov.nii.gz'], 'file')) || (exist([sub_dir{k} func_files{k,k2} '_slice' num2str(slices) '_cov.nii.gz'], 'file')) ) || (run_allsubs_by_default==1)
            skip=skip+1;
            clk=clock; L=L+1; logg{L,1}=sprintf('%d/%s/%d %dh:%dm:%ds - begin (14) Regress-out additional covariates for subject (%d) of (%d), run (%d) of (%d)',clk(3),month{clk(2)},clk(1),clk(4),clk(5),round(clk(6)),k,length(sub_dir),k2,runs(k)); try fprintf(fileID,'%s\n',logg{L,1}); set(S.tx,'string',logg); catch, end; clear clk
            try waitbar(((runs(k)*(k-1)+k2)/(length(sub_dir)*runs(k))),wbar2,sprintf('14. Regress-out additional covariates: Subject (%d) of (%d), Run (%d) of (%d)',k,length(sub_dir),k2,runs(k))); catch, end
            fprintf('14. Regress-out additional covariates: Subject (%d) of (%d), Run (%d) of (%d)\n',k,length(sub_dir),k2,runs(k))
            if runs(k)==1, Rns=0; else, Rns=k2; end
            time_14_COVregress = tic;
            scfMRItb_14_COVregress(sub_dir{k},func_files{k,k2},anat_files{k,1},covregress_options,cov_files{k,k2},slices, wbar3,save_videos2,Rns);
            time_taken_preproc.step14_COVregress(k,k2) = toc(time_14_COVregress); clear time_14_COVregress Rns
            clk=clock; L=L+1; logg{L,1}=sprintf('%d/%s/%d %dh:%dm:%ds - end (14) regress-out additional covariates for subject (%d) of (%d), run (%d) of (%d)',clk(3),month{clk(2)},clk(1),clk(4),clk(5),round(clk(6)),k,length(sub_dir),k2,runs(k)); try fprintf(fileID,'%s\n',logg{L,1}); set(S.tx,'string',logg); catch, end; clear clk
            refresh_html_log
        else
            clk=clock; L=L+1; logg{L,1}=sprintf('%d/%s/%d %dh:%dm:%ds - skip (14) regress-out additional covariates for subject (%d) of (%d), run (%d) of (%d)',clk(3),month{clk(2)},clk(1),clk(4),clk(5),round(clk(6)),k,length(sub_dir),k2,runs(k)); try fprintf(fileID,'%s\n',logg{L,1}); set(S.tx,'string',logg); catch, end; clear clk
        end
    end; clear k2
end; clear k save_videos2

if skip==0
    clk=clock; L=L+1; logg{L,1}=sprintf('%d/%s/%d %dh:%dm:%ds - (14) regress-out additional covariates SKIPPED FOR ALL SUBJECTS',clk(3),month{clk(2)},clk(1),clk(4),clk(5),round(clk(6))); try fprintf(fileID,'%s\n',logg{L,1}); set(S.tx,'string',logg); catch, end; clear clk
    try figure(status_fig),subplot(N,2,27),imshow(imread('status_images/step_14_green.jpeg')), subplot(N,2,28),imshow(imread('status_images/status_green_skip.jpg')), catch, end
else
    clk=clock; L=L+1; logg{L,1}=sprintf('%d/%s/%d %dh:%dm:%ds - (14) regress-out additional covariates COMPLETED FOR ALL SUBJECTS',clk(3),month{clk(2)},clk(1),clk(4),clk(5),round(clk(6))); try fprintf(fileID,'%s\n',logg{L,1}); set(S.tx,'string',logg); catch, end; clear clk
    L=L+1; logg{L,1}=sprintf('Total time taken so far: %dh:%dm:%ds\nTime taken for this step: %dh:%dm:%ds\nTime taken for this step in each run of each subject: %dh:%dm:%ds',floor(toc(tc0)/3600),floor(mod(toc(tc0),3600)/60),floor(mod(toc(tc0),60)),floor(toc(tc)/3600),floor(mod(toc(tc),3600)/60),floor(mod(toc(tc),60)),floor(toc(tc)/3600/skip),floor(mod(toc(tc),3600)/60/skip),floor(mod(toc(tc),60)/skip)); try fprintf(fileID,'%s\n',logg{L,1}); set(S.tx,'string',logg); catch, end; clear clk
    try figure(status_fig),subplot(N,2,27),imshow(imread('status_images/step_14_green.jpeg')), subplot(N,2,28),imshow(imread('status_images/status_green.jpg')), catch, end
    refresh_html
end
end

%% 'Save denoised 3D+time fMRI data'
if sum([choices_preproc(6);choices_preproc(8);choices_preproc(10:14)])~=0
    try L=L+2; logg{L,1}=sprintf('-------\nSave denoised 3D+time fMRI data'); catch, end; try fprintf(fileID,'%s\n',logg{L,1}); set(S.tx,'string',logg); catch, end; tc=tic;
time_taken_preproc.save4Ddata_denoised = zeros(size(func_files,1),size(func_files,2));

skip=0;
for k=1:length(sub_dir)
    for k2=1:runs(k)
        try load([sub_dir{k} sprintf('workspace_variables_run%d.mat',k2)], 'cutoff_str'); catch, cutoff_str = '50'; end
        if exist([sub_dir{k} func_files{k,k2} '_WM' '.nii'], 'file') || exist([sub_dir{k} func_files{k,k2} '_WM' '.nii.gz'], 'file') % if only WM regress step was performed
            existname = [sub_dir{k} func_files{k,k2} '_WM' '.mat'];
        end
        if exist([sub_dir{k} func_files{k,k2} '_denoised' cutoff_str '.nii'], 'file') || exist([sub_dir{k} func_files{k,k2} '_denoised' cutoff_str '.nii.gz'], 'file') % if only CSF regress step was performed
            existname = [sub_dir{k} func_files{k,k2} '_denoised' cutoff_str '.mat'];
        end
        if exist([sub_dir{k} func_files{k,k2} '_cov' '.nii'], 'file') || exist([sub_dir{k} func_files{k,k2} '_cov' '.nii.gz'], 'file') % if only cov regress step was performed
            existname = [sub_dir{k} func_files{k,k2} '_cov' '.mat'];
        end
        if exist([sub_dir{k} func_files{k,k2} '_WMcov' '.nii'], 'file') || exist([sub_dir{k} func_files{k,k2} '_WMcov' '.nii.gz'], 'file') % if both WM and cov regress steps were performed
            existname = [sub_dir{k} func_files{k,k2} '_WMcov' '.mat'];
        end
        if exist([sub_dir{k} func_files{k,k2} '_denoised' cutoff_str 'WM' '.nii'], 'file') || exist([sub_dir{k} func_files{k,k2} '_denoised' cutoff_str 'WM' '.nii.gz'], 'file') % if both CSF and WM regress steps were performed
            existname = [sub_dir{k} func_files{k,k2} '_denoised' cutoff_str 'WM' '.mat'];
        end
        if exist([sub_dir{k} func_files{k,k2} '_denoised' cutoff_str 'cov' '.nii'], 'file') || exist([sub_dir{k} func_files{k,k2} '_denoised' cutoff_str 'cov' '.nii.gz'], 'file') % if both CSF and cov regress steps were performed
            existname = [sub_dir{k} func_files{k,k2} '_denoised' cutoff_str 'cov' '.mat'];
        end
        if exist([sub_dir{k} func_files{k,k2} '_denoised' cutoff_str 'WMcov' '.nii'], 'file') || exist([sub_dir{k} func_files{k,k2} '_denoised' cutoff_str 'WMcov' '.nii.gz'], 'file') % if both CSF, WM and cov regress steps were performed
            existname = [sub_dir{k} func_files{k,k2} '_denoised' cutoff_str 'WMcov' '.mat'];
        end
        if ~(exist('existname', 'var'))
            existname = [];
        end
        if ~(exist(existname, 'file')) || (run_allsubs_by_default==1)
            skip=skip+1;
            try clk=clock; L=L+1; logg{L,1}=sprintf('%d/%s/%d %dh:%dm:%ds - begin save denoised 3D+time fMRI data for subject (%d) of (%d), run (%d) of (%d)',clk(3),month{clk(2)},clk(1),clk(4),clk(5),round(clk(6)),k,length(sub_dir),k2,runs(k)); catch, end; try fprintf(fileID,'%s\n',logg{L,1}); set(S.tx,'string',logg); catch, end; clear clk
            try waitbar(((runs(k)*(k-1)+k2)/(length(sub_dir)*runs(k))),wbar2,sprintf('Save denoised 3D+time fMRI data: Subject (%d) of (%d), Run (%d) of (%d)',k,length(sub_dir),k2,runs(k))); catch, end
            if ~exist('wbar3','var'), wbar3 = waitbar(0,'14. Save denoised 3D+time fMRI data...','Name','Progress(14): Save denoised 3D+time fMRI data...','Units','normalized','Position',[3/5 0 1/5 1/17]); end
            fprintf('Save denoised 3D+time fMRI data: Subject (%d) of (%d), Run (%d) of (%d)\n',k,length(sub_dir),k2,runs(k))
            time_save4Ddata_denoised = tic;
            scfMRItb_14_save4Ddata_denoised(sub_dir{k},func_files{k,k2},anat_files{k,1},wbar3);
            time_taken_preproc.save4Ddata_denoised(k,k2) = toc(time_save4Ddata_denoised); clear time_save4Ddata_denoised
            try clk=clock; L=L+1; logg{L,1}=sprintf('%d/%s/%d %dh:%dm:%ds - end save denoised 3D+time fMRI data for subject (%d) of (%d), run (%d) of (%d)',clk(3),month{clk(2)},clk(1),clk(4),clk(5),round(clk(6)),k,length(sub_dir),k2,runs(k)); catch, end; try fprintf(fileID,'%s\n',logg{L,1}); set(S.tx,'string',logg); catch, end; clear clk
        else
            try clk=clock; L=L+1; logg{L,1}=sprintf('%d/%s/%d %dh:%dm:%ds - skip save denoised 3D+time fMRI data for subject (%d) of (%d), run (%d) of (%d)',clk(3),month{clk(2)},clk(1),clk(4),clk(5),round(clk(6)),k,length(sub_dir),k2,runs(k)); catch, end; try fprintf(fileID,'%s\n',logg{L,1}); set(S.tx,'string',logg); catch, end; clear clk
        end
        clear existname
        refresh_html_log
    end; clear k2
end

if skip==0
    try clk=clock; L=L+1; logg{L,1}=sprintf('%d/%s/%d %dh:%dm:%ds - save denoised 3D+time fMRI data SKIPPED FOR ALL SUBJECTS',clk(3),month{clk(2)},clk(1),clk(4),clk(5),round(clk(6))); catch, end; try fprintf(fileID,'%s\n',logg{L,1}); set(S.tx,'string',logg); catch, end; clear clk
else
    try clk=clock; L=L+1; logg{L,1}=sprintf('%d/%s/%d %dh:%dm:%ds - save denoised 3D+time fMRI data COMPLETED FOR ALL SUBJECTS',clk(3),month{clk(2)},clk(1),clk(4),clk(5),round(clk(6))); catch, end; try fprintf(fileID,'%s\n',logg{L,1}); set(S.tx,'string',logg); catch, end; clear clk
    try L=L+1; logg{L,1}=sprintf('Total time taken so far: %dh:%dm:%ds\nTime taken for this step: %dh:%dm:%ds\nTime taken for this step in each run of each subject: %dh:%dm:%ds',floor(toc(tc0)/3600),floor(mod(toc(tc0),3600)/60),floor(mod(toc(tc0),60)),floor(toc(tc)/3600),floor(mod(toc(tc),3600)/60),floor(mod(toc(tc),60)),floor(toc(tc)/3600/skip),floor(mod(toc(tc),3600)/60/skip),floor(mod(toc(tc),60)/skip)); catch, end; try fprintf(fileID,'%s\n',logg{L,1}); set(S.tx,'string',logg); catch, end; clear clk
end
refresh_html
end

%% '15. Temporal band-pass filtering'
if choices_preproc(15)==1
try figure(status_fig),subplot(N,2,29),imshow(imread('status_images/step_15_blue.jpeg')), subplot(N,2,30),imshow(imread('status_images/status_blue.jpg')), catch, end
L=L+2; logg{L,1}=sprintf('-------\n%s',preprocsteps{15}); try fprintf(fileID,'%s\n',logg{L,1}); set(S.tx,'string',logg); catch, end; tc=tic;
time_taken_preproc.step15_filterData = zeros(size(func_files,1),size(func_files,2));

skip=0;
for k=1:length(sub_dir)
    for k2=1:runs(k)
        try load([sub_dir{k} sprintf('workspace_variables_run%d.mat',k2)], 'cutoff_str'); catch, cutoff_str = '50'; end
        if exist([sub_dir{k} func_files{k,k2} '_WM' '.nii'], 'file') || exist([sub_dir{k} func_files{k,k2} '_WM' '.nii.gz'], 'file') % if only WM regress step was performed
            existname = [sub_dir{k} func_files{k,k2} '_WM' '_HPF01.mat'];
        end
        if exist([sub_dir{k} func_files{k,k2} '_denoised' cutoff_str '.nii'], 'file') || exist([sub_dir{k} func_files{k,k2} '_denoised' cutoff_str '.nii.gz'], 'file') % if only CSF regress step was performed
            existname = [sub_dir{k} func_files{k,k2} '_denoised' cutoff_str '_HPF01.mat'];
        end
        if exist([sub_dir{k} func_files{k,k2} '_cov' '.nii'], 'file') || exist([sub_dir{k} func_files{k,k2} '_cov' '.nii.gz'], 'file') % if only cov regress step was performed
            existname = [sub_dir{k} func_files{k,k2} '_cov' '_HPF01.mat'];
        end
        if exist([sub_dir{k} func_files{k,k2} '_WMcov' '.nii'], 'file') || exist([sub_dir{k} func_files{k,k2} '_WMcov' '.nii.gz'], 'file') % if both WM and cov regress steps were performed
            existname = [sub_dir{k} func_files{k,k2} '_WMcov' '_HPF01.mat'];
        end
        if exist([sub_dir{k} func_files{k,k2} '_denoised' cutoff_str 'WM' '.nii'], 'file') || exist([sub_dir{k} func_files{k,k2} '_denoised' cutoff_str 'WM' '.nii.gz'], 'file') % if both CSF and WM regress steps were performed
            existname = [sub_dir{k} func_files{k,k2} '_denoised' cutoff_str 'WM' '_HPF01.mat'];
        end
        if exist([sub_dir{k} func_files{k,k2} '_denoised' cutoff_str 'cov' '.nii'], 'file') || exist([sub_dir{k} func_files{k,k2} '_denoised' cutoff_str 'cov' '.nii.gz'], 'file') % if both CSF and cov regress steps were performed
            existname = [sub_dir{k} func_files{k,k2} '_denoised' cutoff_str 'cov' '_HPF01.mat'];
        end
        if exist([sub_dir{k} func_files{k,k2} '_denoised' cutoff_str 'WMcov' '.nii'], 'file') || exist([sub_dir{k} func_files{k,k2} '_denoised' cutoff_str 'WMcov' '.nii.gz'], 'file') % if both CSF, WM and cov regress steps were performed
            existname = [sub_dir{k} func_files{k,k2} '_denoised' cutoff_str 'WMcov' '_HPF01.mat'];
        end
        if ~(exist('existname', 'var'))
            existname = [];
        end
        if ~(exist(existname, 'file')) || (run_allsubs_by_default==1)
            skip=skip+1;
            clk=clock; L=L+1; logg{L,1}=sprintf('%d/%s/%d %dh:%dm:%ds - begin (15) temporal band-pass filtering for subject (%d) of (%d), run (%d) of (%d)',clk(3),month{clk(2)},clk(1),clk(4),clk(5),round(clk(6)),k,length(sub_dir),k2,runs(k)); try fprintf(fileID,'%s\n',logg{L,1}); set(S.tx,'string',logg); catch, end; clear clk
            try waitbar(((runs(k)*(k-1)+k2)/(length(sub_dir)*runs(k))),wbar2,sprintf('15. Temporal band-pass filtering: Subject (%d) of (%d), Run (%d) of (%d)',k,length(sub_dir),k2,runs(k))); catch, end
            fprintf('15. Temporal band-pass filtering: Subject (%d) of (%d), Run (%d) of (%d)\n',k,length(sub_dir),k2,runs(k))
            if runs(k)==1, Rns=0; else, Rns=k2; end
            time_15_filterData = tic;
            scfMRItb_15_filterData(sub_dir{k},func_files{k,k2},anat_files{k,1},bpf_hz,detrend_flag, wbar3,save_videos,Rns);
            time_taken_preproc.step15_filterData(k,k2) = toc(time_15_filterData); clear time_15_filterData Rns
            clk=clock; L=L+1; logg{L,1}=sprintf('%d/%s/%d %dh:%dm:%ds - end (15) temporal band-pass filtering for subject (%d) of (%d), run (%d) of (%d)',clk(3),month{clk(2)},clk(1),clk(4),clk(5),round(clk(6)),k,length(sub_dir),k2,runs(k)); try fprintf(fileID,'%s\n',logg{L,1}); set(S.tx,'string',logg); catch, end; clear clk
            refresh_html_log
        else
            if ~(exist([sub_dir{k} 'workspace_variables_temp.mat'], 'file'))
                base_dir_sub = sub_dir{k}; fname = func_files{k,k2}; cutoff = 0.50; cutoff_str = '50'; f_BPF_max_vec = 0.10; f_BPF_max_str_vec = {'10'};
                scfMRItb_04_unzipFile(sub_dir{k}, func_files{k,k2}, '')
                F = load_untouch_nii([sub_dir{k} func_files{k,k2} '.nii']); TR = F.hdr.dime.pixdim(1,5);
                f_BPF_max = f_BPF_max_vec(1); f_BPF_max_str = f_BPF_max_str_vec{1};
                save([sub_dir{k} 'workspace_variables_temp.mat'], 'base_dir_sub', 'fname', 'cutoff', 'cutoff_str', 'f_BPF_max_vec', 'f_BPF_max_str_vec', 'f_BPF_max', 'f_BPF_max_str', 'TR', 'bpf_hz')
                clear base_dir_sub fname F TR
            end
            clk=clock; L=L+1; logg{L,1}=sprintf('%d/%s/%d %dh:%dm:%ds - skip (15) temporal band-pass filtering for subject (%d) of (%d), run (%d) of (%d)',clk(3),month{clk(2)},clk(1),clk(4),clk(5),round(clk(6)),k,length(sub_dir),k2,runs(k)); try fprintf(fileID,'%s\n',logg{L,1}); set(S.tx,'string',logg); catch, end; clear clk
        end
        clear existname
    end; clear k2
end

if skip==0
    clk=clock; L=L+1; logg{L,1}=sprintf('%d/%s/%d %dh:%dm:%ds - (15) temporal band-pass filtering SKIPPED FOR ALL SUBJECTS',clk(3),month{clk(2)},clk(1),clk(4),clk(5),round(clk(6))); try fprintf(fileID,'%s\n',logg{L,1}); set(S.tx,'string',logg); catch, end; clear clk
    try figure(status_fig),subplot(N,2,29),imshow(imread('status_images/step_15_green.jpeg')), subplot(N,2,30),imshow(imread('status_images/status_green_skip.jpg')), catch, end
else
    clk=clock; L=L+1; logg{L,1}=sprintf('%d/%s/%d %dh:%dm:%ds - (15) temporal band-pass filtering COMPLETED FOR ALL SUBJECTS',clk(3),month{clk(2)},clk(1),clk(4),clk(5),round(clk(6))); try fprintf(fileID,'%s\n',logg{L,1}); set(S.tx,'string',logg); catch, end; clear clk
    L=L+1; logg{L,1}=sprintf('Total time taken so far: %dh:%dm:%ds\nTime taken for this step: %dh:%dm:%ds\nTime taken for this step in each run of each subject: %dh:%dm:%ds',floor(toc(tc0)/3600),floor(mod(toc(tc0),3600)/60),floor(mod(toc(tc0),60)),floor(toc(tc)/3600),floor(mod(toc(tc),3600)/60),floor(mod(toc(tc),60)),floor(toc(tc)/3600/skip),floor(mod(toc(tc),3600)/60/skip),floor(mod(toc(tc),60)/skip)); try fprintf(fileID,'%s\n',logg{L,1}); set(S.tx,'string',logg); catch, end; clear clk
    try figure(status_fig),subplot(N,2,29),imshow(imread('status_images/step_15_green.jpeg')), subplot(N,2,30),imshow(imread('status_images/status_green.jpg')), catch, end
    refresh_html_log
end
end

%% '16. Define cord quadrants'
if choices_preproc(16)==1
if UIstepsFirst~=1
try figure(status_fig),subplot(N,2,31),imshow(imread('status_images/step_16_blue.jpeg')), subplot(N,2,32),imshow(imread('status_images/status_blue_UI.jpg')), catch, end
L=L+2; logg{L,1}=sprintf('-------\n%s',preprocsteps{16}); try fprintf(fileID,'%s\n',logg{L,1}); set(S.tx,'string',logg); catch, end; tc=tic;
time_taken_preproc.step16_cord_quadrants = zeros(size(func_files,1),1);
end

skip=0;
for k=1:length(sub_dir)
    if ~(exist([sub_dir{k} 'workspace_variables_temp.mat'], 'file'))
        try
        base_dir_sub = sub_dir{k}; fname = func_files{k,1}; cutoff = 0.50; cutoff_str = '50'; f_BPF_max_vec = 0.10; f_BPF_max_str_vec = {'10'}; if ~exist('bpf_hz','var'), bpf_hz = f_BPF_max_vec; end
        scfMRItb_04_unzipFile(sub_dir{k}, func_files{k,1}, ''); F = load_untouch_nii([sub_dir{k} func_files{k,1} '.nii']); TR = F.hdr.dime.pixdim(1,5);
        if TR>3.65, wz=2; else, wz=3; end; f_BPF_max = f_BPF_max_vec(1,1); f_BPF_max_str = f_BPF_max_str_vec{1,1}; clear wz
        save([sub_dir{k} 'workspace_variables_temp.mat'], 'base_dir_sub', 'fname', 'cutoff', 'cutoff_str', 'f_BPF_max_vec', 'f_BPF_max_str_vec', 'f_BPF_max', 'f_BPF_max_str', 'TR', 'bpf_hz')
        clear base_dir_sub fname F TR
        catch
        end
    end
    
    if (run_allsubs_by_default==1) && (UIstepsFirst~=1)
        skip=skip+1;
        clk=clock; L=L+1; logg{L,1}=sprintf('%d/%s/%d %dh:%dm:%ds - begin (16) define cord quadrants for subject (%d) of (%d)',clk(3),month{clk(2)},clk(1),clk(4),clk(5),round(clk(6)),k,length(sub_dir)); try fprintf(fileID,'%s\n',logg{L,1}); set(S.tx,'string',logg); catch, end; clear clk
        try waitbar((k/length(sub_dir)),wbar2,sprintf('16. Define cord quadrants: Subject (%d) of (%d)',k,length(sub_dir))); catch, end
        fprintf('16. Define cord quadrants: Subject (%d) of (%d)\n',k,length(sub_dir))
        time_16_cord_quadrants = tic;
        scfMRItb_16_cord_quadrants(sub_dir{k},func_files{k,1},anat_files{k,1}, scan7T,C_spine,wbar3);
        time_taken_preproc.step16_cord_quadrants(k,1) = toc(time_16_cord_quadrants); clear time_16_cord_quadrants
        merge_mat_files([sub_dir{k} sprintf('workspace_variables_run%d.mat',1)],[sub_dir{k} sprintf('workspace_variables_run%d.mat',1)],[sub_dir{k} 'workspace_variables_temp.mat'],1)
        clk=clock; L=L+1; logg{L,1}=sprintf('%d/%s/%d %dh:%dm:%ds - end (16) define cord quadrants for subject (%d) of (%d)',clk(3),month{clk(2)},clk(1),clk(4),clk(5),round(clk(6)),k,length(sub_dir)); try fprintf(fileID,'%s\n',logg{L,1}); set(S.tx,'string',logg); catch, end; clear clk
        refresh_html_log
    elseif (run_allsubs_by_default~=1) && (UIstepsFirst~=1)
    try
        load([sub_dir{k} 'cord_quadrants.mat'], 'FC_mask_quads')
        clk=clock; L=L+1; logg{L,1}=sprintf('%d/%s/%d %dh:%dm:%ds - skip (16) define cord quadrants for subject (%d) of (%d)',clk(3),month{clk(2)},clk(1),clk(4),clk(5),round(clk(6)),k,length(sub_dir)); try fprintf(fileID,'%s\n',logg{L,1}); set(S.tx,'string',logg); catch, end; clear clk
        if ~exist([sub_dir{k} sprintf('workspace_variables_run%d.mat',1)],'file')
            try
                load([base_dir_sub 'workspace_variables_temp.mat'])
            catch
                f_BPF_max_vec = 0.10; f_BPF_max_str_vec = {'10'};
                cutoff = 0.50; cutoff_str = '50';
                f_BPF_max = f_BPF_max_vec(1,1); f_BPF_max_str = f_BPF_max_str_vec{1,1};
                if ~exist('TR','var')
                    scfMRItb_04_unzipFile(sub_dir{k}, func_files{k,1}, '')
                    F = load_untouch_nii([sub_dir{k} func_files{k,1} '.nii']); TR = F.hdr.dime.pixdim(1,5); clear F
                end
            end
            base_dir_sub = sub_dir{k}; fname = func_files{k,1};
            try
            save([base_dir_sub 'workspace_variables_run1.mat'],'base_dir_sub','fname*','cutoff*','mask_*','f_BPF_max*','*FC_mask_*','*spine','TR','scan3T','new_size')
            catch
            save([base_dir_sub 'workspace_variables_run1.mat'],'base_dir_sub','fname*','cutoff*','f_BPF_max*','*FC_mask_*','*spine','TR','scan3T')
            end
        end
    catch
        skip=skip+1;
        clk=clock; L=L+1; logg{L,1}=sprintf('%d/%s/%d %dh:%dm:%ds - begin (16) define cord quadrants for subject (%d) of (%d)',clk(3),month{clk(2)},clk(1),clk(4),clk(5),round(clk(6)),k,length(sub_dir)); try fprintf(fileID,'%s\n',logg{L,1}); set(S.tx,'string',logg); catch, end; clear clk
        try waitbar((k/length(sub_dir)),wbar2,sprintf('16. Define cord quadrants: Subject (%d) of (%d)',k,length(sub_dir))); catch, end
        fprintf('16. Define cord quadrants: Subject (%d) of (%d)\n',k,length(sub_dir))
        time_16_cord_quadrants = tic;
        scfMRItb_16_cord_quadrants(sub_dir{k},func_files{k,1},anat_files{k,1}, scan7T,C_spine,wbar3);
        time_taken_preproc.step16_cord_quadrants(k,1) = toc(time_16_cord_quadrants); clear time_16_cord_quadrants
        clk=clock; L=L+1; logg{L,1}=sprintf('%d/%s/%d %dh:%dm:%ds - end (16) define cord quadrants for subject (%d) of (%d)',clk(3),month{clk(2)},clk(1),clk(4),clk(5),round(clk(6)),k,length(sub_dir)); try fprintf(fileID,'%s\n',logg{L,1}); set(S.tx,'string',logg); catch, end; clear clk
        refresh_html_log
    end
    elseif UIstepsFirst==1
        scfMRItb_16_cord_quadrants_remnants(sub_dir{k},func_files{k,1},anat_files{k,1}, scan7T,C_spine);
    end
    try
        merge_mat_files([sub_dir{k} sprintf('workspace_variables_run%d.mat',1)],[sub_dir{k} 'workspace_variables_temp.mat'],[sub_dir{k} sprintf('workspace_variables_run%d.mat',1)],1)
    catch
    end
    if exist([sub_dir{k} 'workspace_variables_temp.mat'],'file') && exist([sub_dir{k} sprintf('workspace_variables_run%d.mat',1)],'file')
        delete([sub_dir{k} 'workspace_variables_temp.mat'])
    end
    
    for k2=2:runs(k)
        try
            load([sub_dir{k} sprintf('workspace_variables_run%d.mat',k2)], 'FC_mask_quads')
        catch
            skip=skip+1;
            copyfile([sub_dir{k} sprintf('workspace_variables_run%d.mat',1)],[sub_dir{k} sprintf('workspace_variables_run%d.mat',k2)],'f')
            clear fname TR
            fname = func_files{k,k2};
            if ~exist('TR','var')
                scfMRItb_04_unzipFile(sub_dir{k}, func_files{k,k2}, '')
                F = load_untouch_nii([sub_dir{k} func_files{k,k2} '.nii']); TR = F.hdr.dime.pixdim(1,5); clear F
            end
            save([sub_dir{k} sprintf('workspace_variables_run%d.mat',k2)], 'fname','TR', '-append')
        end
    end; clear k2
end

if (skip==0) && (UIstepsFirst~=1)
    clk=clock; L=L+1; logg{L,1}=sprintf('%d/%s/%d %dh:%dm:%ds - (16) define cord quadrants SKIPPED FOR ALL SUBJECTS',clk(3),month{clk(2)},clk(1),clk(4),clk(5),round(clk(6))); try fprintf(fileID,'%s\n',logg{L,1}); set(S.tx,'string',logg); catch, end; clear clk
    try figure(status_fig),subplot(N,2,31),imshow(imread('status_images/step_16_green.jpeg')), subplot(N,2,32),imshow(imread('status_images/status_green_skip.jpg')), catch, end
elseif (skip~=0) && (UIstepsFirst~=1)
    clk=clock; L=L+1; logg{L,1}=sprintf('%d/%s/%d %dh:%dm:%ds - (16) define cord quadrants COMPLETED FOR ALL SUBJECTS',clk(3),month{clk(2)},clk(1),clk(4),clk(5),round(clk(6))); try fprintf(fileID,'%s\n',logg{L,1}); set(S.tx,'string',logg); catch, end; clear clk
    L=L+1; logg{L,1}=sprintf('Total time taken so far: %dh:%dm:%ds\nTime taken for this step: %dh:%dm:%ds\nTime taken for this step in each run of each subject: %dh:%dm:%ds',floor(toc(tc0)/3600),floor(mod(toc(tc0),3600)/60),floor(mod(toc(tc0),60)),floor(toc(tc)/3600),floor(mod(toc(tc),3600)/60),floor(mod(toc(tc),60)),floor(toc(tc)/3600/skip),floor(mod(toc(tc),3600)/60/skip),floor(mod(toc(tc),60)/skip)); try fprintf(fileID,'%s\n',logg{L,1}); set(S.tx,'string',logg); catch, end; clear clk
    try figure(status_fig),subplot(N,2,31),imshow(imread('status_images/step_16_green.jpeg')), subplot(N,2,32),imshow(imread('status_images/status_green.jpg')), catch, end
    refresh_html_log
end
end

%% '17. HRF deconvolution'
if choices_preproc(17)==1
try figure(status_fig),subplot(N,2,33),imshow(imread('status_images/step_17_blue.jpeg')), subplot(N,2,34),imshow(imread('status_images/status_blue.jpg')), catch, end
L=L+2; logg{L,1}=sprintf('-------\n%s',preprocsteps{17}); try fprintf(fileID,'%s\n',logg{L,1}); set(S.tx,'string',logg); catch, end; tc=tic;
time_taken_preproc.step17_deconvolution = zeros(size(func_files,1),size(func_files,2));

skip=0;
if ~(exist('bpf_hz', 'var')), bpf_hz = 0.1; end
for k=1:length(sub_dir)
    for k2=1:runs(k)
        try load([sub_dir{k} sprintf('workspace_variables_run%d.mat',k2)], 'cutoff_str'); catch, cutoff_str = '50'; end
        if run_on_HPF_data==1
            if exist([sub_dir{k} func_files{k,k2} '_WM' '.nii'], 'file') || exist([sub_dir{k} func_files{k,k2} '_WM' '.nii.gz'], 'file') % if only WM regress step was performed
                if exist([sub_dir{k} func_files{k,k2} '_WM' '_HPF' '01' '.mat'], 'file')
                    namefile = [sub_dir{k} func_files{k,k2} '_WM' '_deconv' '_BPF' bpf_hz_str '.mat'];
                else
                    namefile = [sub_dir{k} func_files{k,k2} '_WM' '_deconv' '.mat'];
                end
            end
            if exist([sub_dir{k} func_files{k,k2} '_denoised' cutoff_str '.nii'], 'file') || exist([sub_dir{k} func_files{k,k2} '_denoised' cutoff_str '.nii.gz'], 'file') % if only CSF regress step was performed
                if exist([sub_dir{k} func_files{k,k2} '_denoised' cutoff_str '_HPF' '01' '.mat'], 'file')
                    namefile = [sub_dir{k} func_files{k,k2} '_denoised' cutoff_str '_deconv' '_BPF' bpf_hz_str '.mat'];
                else
                    namefile = [sub_dir{k} func_files{k,k2} '_denoised' cutoff_str '_deconv' '.mat'];
                end
            end
            if exist([sub_dir{k} func_files{k,k2} '_cov' '.nii'], 'file') || exist([sub_dir{k} func_files{k,k2} '_cov' '.nii.gz'], 'file') % if only cov regress step was performed
                if exist([sub_dir{k} func_files{k,k2} '_cov' '_HPF' '01' '.mat'], 'file')
                    namefile = [sub_dir{k} func_files{k,k2} '_cov' '_deconv' '_BPF' bpf_hz_str '.mat'];
                else
                    namefile = [sub_dir{k} func_files{k,k2} '_cov' '_deconv' '.mat'];
                end
            end
            if exist([sub_dir{k} func_files{k,k2} '_WMcov' '.nii'], 'file') || exist([sub_dir{k} func_files{k,k2} '_WMcov' '.nii.gz'], 'file') % if both WM and cov regress steps were performed
                if exist([sub_dir{k} func_files{k,k2} '_WMcov' '_HPF' '01' '.mat'], 'file')
                    namefile = [sub_dir{k} func_files{k,k2} '_WMcov' '_deconv' '_BPF' bpf_hz_str '.mat'];
                else
                    namefile = [sub_dir{k} func_files{k,k2} '_WMcov' '_deconv' '.mat'];
                end
            end
            if exist([sub_dir{k} func_files{k,k2} '_denoised' cutoff_str 'WM' '.nii'], 'file') || exist([sub_dir{k} func_files{k,k2} '_denoised' cutoff_str 'WM' '.nii.gz'], 'file') % if both CSF and WM regress steps were performed
                if exist([sub_dir{k} func_files{k,k2} '_denoised' cutoff_str 'WM' '_HPF' '01' '.mat'], 'file')
                    namefile = [sub_dir{k} func_files{k,k2} '_denoised' cutoff_str 'WM' '_deconv' '_BPF' bpf_hz_str '.mat'];
                else
                    namefile = [sub_dir{k} func_files{k,k2} '_denoised' cutoff_str 'WM' '_deconv' '.mat'];
                end
            end
            if exist([sub_dir{k} func_files{k,k2} '_denoised' cutoff_str 'cov' '.nii'], 'file') || exist([sub_dir{k} func_files{k,k2} '_denoised' cutoff_str 'cov' '.nii.gz'], 'file') % if both CSF and cov regress steps were performed
                if exist([sub_dir{k} func_files{k,k2} '_denoised' cutoff_str 'cov' '_HPF' '01' '.mat'], 'file')
                    namefile = [sub_dir{k} func_files{k,k2} '_denoised' cutoff_str 'cov' '_deconv' '_BPF' bpf_hz_str '.mat'];
                else
                    namefile = [sub_dir{k} func_files{k,k2} '_denoised' cutoff_str 'cov' '_deconv' '.mat'];
                end
            end
            if exist([sub_dir{k} func_files{k,k2} '_denoised' cutoff_str 'WMcov' '.nii'], 'file') || exist([sub_dir{k} func_files{k,k2} '_denoised' cutoff_str 'WMcov' '.nii.gz'], 'file') % if CSF, WM and cov regress steps were performed
                if exist([sub_dir{k} func_files{k,k2} '_denoised' cutoff_str 'WMcov' '_HPF' '01' '.mat'], 'file')
                    namefile = [sub_dir{k} func_files{k,k2} '_denoised' cutoff_str 'WMcov' '_deconv' '_BPF' bpf_hz_str '.mat'];
                else
                    namefile = [sub_dir{k} func_files{k,k2} '_denoised' cutoff_str 'WMcov' '_deconv' '.mat'];
                end
            end
        elseif run_on_HPF_data==0
            if exist([sub_dir{k} func_files{k,k2} '_WM' '.nii'], 'file') || exist([sub_dir{k} func_files{k,k2} '_WM' '.nii.gz'], 'file') % if only WM regress step was performed
                if exist([sub_dir{k} func_files{k,k2} '_WM' '_BPF' bpf_hz_str '_deconv' '.mat'], 'file')
                    namefile = [sub_dir{k} func_files{k,k2} '_WM' '_BPF' bpf_hz_str '_deconv' '.mat'];
                else
                    namefile = [sub_dir{k} func_files{k,k2} '_WM' '_BPF' '10' '_deconv' '.mat'];
                end
            end
            if exist([sub_dir{k} func_files{k,k2} '_denoised' cutoff_str '.nii'], 'file') || exist([sub_dir{k} func_files{k,k2} '_denoised' cutoff_str '.nii.gz'], 'file') % if only CSF regress step was performed
                if exist([sub_dir{k} func_files{k,k2} '_denoised' cutoff_str '_BPF' bpf_hz_str '_deconv' '.mat'], 'file')
                    namefile = [sub_dir{k} func_files{k,k2} '_denoised' cutoff_str '_BPF' bpf_hz_str '_deconv' '.mat'];
                else
                    namefile = [sub_dir{k} func_files{k,k2} '_denoised' cutoff_str '_BPF' '10' '_deconv' '.mat'];
                end
            end
            if exist([sub_dir{k} func_files{k,k2} '_cov' '.nii'], 'file') || exist([sub_dir{k} func_files{k,k2} '_cov' '.nii.gz'], 'file') % if only cov regress step was performed
                if exist([sub_dir{k} func_files{k,k2} '_cov' '_BPF' bpf_hz_str '_deconv' '.mat'], 'file')
                    namefile = [sub_dir{k} func_files{k,k2} '_cov' '_BPF' bpf_hz_str '_deconv' '.mat'];
                else
                    namefile = [sub_dir{k} func_files{k,k2} '_cov' '_BPF' '10' '_deconv' '.mat'];
                end
            end
            if exist([sub_dir{k} func_files{k,k2} '_denoised' cutoff_str 'WM' '.nii'], 'file') || exist([sub_dir{k} func_files{k,k2} '_denoised' cutoff_str 'WM' '.nii.gz'], 'file') % if both CSF and WM regress steps were performed
                if exist([sub_dir{k} func_files{k,k2} '_denoised' cutoff_str 'WM' '_BPF' bpf_hz_str '_deconv' '.mat'], 'file')
                    namefile = [sub_dir{k} func_files{k,k2} '_denoised' cutoff_str 'WM' '_BPF' bpf_hz_str '_deconv' '.mat'];
                else
                    namefile = [sub_dir{k} func_files{k,k2} '_denoised' cutoff_str 'WM' '_BPF' '10' '_deconv' '.mat'];
                end
            end
            if exist([sub_dir{k} func_files{k,k2} '_denoised' cutoff_str 'cov' '.nii'], 'file') || exist([sub_dir{k} func_files{k,k2} '_denoised' cutoff_str 'cov' '.nii.gz'], 'file') % if both CSF and cov regress steps were performed
                if exist([sub_dir{k} func_files{k,k2} '_denoised' cutoff_str 'cov' '_BPF' bpf_hz_str '_deconv' '.mat'], 'file')
                    namefile = [sub_dir{k} func_files{k,k2} '_denoised' cutoff_str 'cov' '_BPF' bpf_hz_str '_deconv' '.mat'];
                else
                    namefile = [sub_dir{k} func_files{k,k2} '_denoised' cutoff_str 'cov' '_BPF' '10' '_deconv' '.mat'];
                end
            end
            if exist([sub_dir{k} func_files{k,k2} '_denoised' cutoff_str 'WMcov' '.nii'], 'file') || exist([sub_dir{k} func_files{k,k2} '_denoised' cutoff_str 'WMcov' '.nii.gz'], 'file') % if CSF, WM and cov regress steps were performed
                if exist([sub_dir{k} func_files{k,k2} '_denoised' cutoff_str 'WMcov' '_BPF' bpf_hz_str '_deconv' '.mat'], 'file')
                    namefile = [sub_dir{k} func_files{k,k2} '_denoised' cutoff_str 'WMcov' '_BPF' bpf_hz_str '_deconv' '.mat'];
                else
                    namefile = [sub_dir{k} func_files{k,k2} '_denoised' cutoff_str 'WMcov' '_BPF' '10' '_deconv' '.mat'];
                end
            end
        end
        if ~(exist('namefile', 'var'))
            namefile = [];
        end
        if ~(exist('bpf_hz', 'var'))
            bpf_hz = 0.1;
        end
        if ~(exist(namefile, 'file')) || (run_allsubs_by_default==1)
            skip=skip+1;
            clk=clock; L=L+1; logg{L,1}=sprintf('%d/%s/%d %dh:%dm:%ds - begin (17) HRF deconvolution for subject (%d) of (%d), run (%d) of (%d)',clk(3),month{clk(2)},clk(1),clk(4),clk(5),round(clk(6)),k,length(sub_dir),k2,runs(k)); try fprintf(fileID,'%s\n',logg{L,1}); set(S.tx,'string',logg); catch, end; clear clk
            try waitbar(((runs(k)*(k-1)+k2)/(length(sub_dir)*runs(k))),wbar2,sprintf('17. HRF deconvolution: Subject (%d) of (%d), Run (%d) of (%d)',k,length(sub_dir),k2,runs(k))); catch, end
            fprintf('17. HRF deconvolution: Subject (%d) of (%d), Run (%d) of (%d)\n',k,length(sub_dir),k2,runs(k))
            if runs(k)==1, Rns=0; else, Rns=k2; end
            time_17_deconvolution = tic;
            if rest==1
                scfMRItb_17_deconvolution_rest(sub_dir{k},func_files{k,k2},anat_files{k,1}, run_on_HPF_data,bpf_hz, wbar3,save_videos,Rns);
            elseif task==1
                scfMRItb_17_deconvolution_rest(sub_dir{k},func_files{k,k2},anat_files{k,1}, run_on_HPF_data,bpf_hz, wbar3,save_videos,Rns);
            end
            time_taken_preproc.step17_deconvolution(k,k2) = toc(time_17_deconvolution); clear time_17_deconvolution Rns
            clk=clock; L=L+1; logg{L,1}=sprintf('%d/%s/%d %dh:%dm:%ds - end (17) HRF deconvolution for subject (%d) of (%d), run (%d) of (%d)',clk(3),month{clk(2)},clk(1),clk(4),clk(5),round(clk(6)),k,length(sub_dir),k2,runs(k)); try fprintf(fileID,'%s\n',logg{L,1}); set(S.tx,'string',logg); catch, end; clear clk
            refresh_html_log
        else
            clk=clock; L=L+1; logg{L,1}=sprintf('%d/%s/%d %dh:%dm:%ds - skip (17) HRF deconvolution for subject (%d) of (%d), run (%d) of (%d)',clk(3),month{clk(2)},clk(1),clk(4),clk(5),round(clk(6)),k,length(sub_dir),k2,runs(k)); try fprintf(fileID,'%s\n',logg{L,1}); set(S.tx,'string',logg); catch, end; clear clk
        end
        clear namefile
    end; clear k2
end

if skip==0
    clk=clock; L=L+1; logg{L,1}=sprintf('%d/%s/%d %dh:%dm:%ds - (17) HRF deconvolution SKIPPED FOR ALL SUBJECTS',clk(3),month{clk(2)},clk(1),clk(4),clk(5),round(clk(6))); try fprintf(fileID,'%s\n',logg{L,1}); set(S.tx,'string',logg); catch, end; clear clk
    try figure(status_fig),subplot(N,2,33),imshow(imread('status_images/step_17_green.jpeg')), subplot(N,2,34),imshow(imread('status_images/status_green_skip.jpg')), catch, end
else
    clk=clock; L=L+1; logg{L,1}=sprintf('%d/%s/%d %dh:%dm:%ds - (17) HRF deconvolution COMPLETED FOR ALL SUBJECTS',clk(3),month{clk(2)},clk(1),clk(4),clk(5),round(clk(6))); try fprintf(fileID,'%s\n',logg{L,1}); set(S.tx,'string',logg); catch, end; clear clk
    L=L+1; logg{L,1}=sprintf('Total time taken so far: %dh:%dm:%ds\nTime taken for this step: %dh:%dm:%ds\nTime taken for this step in each run of each subject: %dh:%dm:%ds',floor(toc(tc0)/3600),floor(mod(toc(tc0),3600)/60),floor(mod(toc(tc0),60)),floor(toc(tc)/3600),floor(mod(toc(tc),3600)/60),floor(mod(toc(tc),60)),floor(toc(tc)/3600/skip),floor(mod(toc(tc),3600)/60/skip),floor(mod(toc(tc),60)/skip)); try fprintf(fileID,'%s\n',logg{L,1}); set(S.tx,'string',logg); catch, end; clear clk
    try figure(status_fig),subplot(N,2,33),imshow(imread('status_images/step_17_green.jpeg')), subplot(N,2,34),imshow(imread('status_images/status_green.jpg')), catch, end
    refresh_html_log
end
end

%% '18. Co-registration to PAM50 standard space'
if choices_preproc(18)==1
try figure(status_fig),subplot(N,2,35),imshow(imread('status_images/step_18_blue.jpeg')), subplot(N,2,36),imshow(imread('status_images/status_blue.jpg')), catch, end
L=L+2; logg{L,1}=sprintf('-------\n%s',preprocsteps{18}); try fprintf(fileID,'%s\n',logg{L,1}); set(S.tx,'string',logg); catch, end; tc=tic;
clk=clock; L=L+1; logg{L,1}=sprintf('%d/%s/%d %dh:%dm:%ds - co-registration to PAM50 standard space',clk(3),month{clk(2)},clk(1),clk(4),clk(5),round(clk(6))); try fprintf(fileID,'%s\n',logg{L,1}); set(S.tx,'string',logg); catch, end; clear clk
time_taken_preproc.step18_PAM50registration = zeros(size(func_files,1),size(func_files,2));

skip=0;
for k=1:length(sub_dir)
    if ((exist_SCT==1) && ~(exist([sub_dir{k} 'SCT_PAM50registration_step18/' 'warp_template2anat.nii.gz'],'file'))) || (run_allsubs_by_default==1)
        skip=skip+1;
        clk=clock; L=L+1; logg{L,1}=sprintf('%d/%s/%d %dh:%dm:%ds - begin (18) co-registration to PAM50 standard space for subject (%d) of (%d)',clk(3),month{clk(2)},clk(1),clk(4),clk(5),round(clk(6)),k,length(sub_dir)); try fprintf(fileID,'%s\n',logg{L,1}); set(S.tx,'string',logg); catch, end; clear clk
        try waitbar(((k)/(length(sub_dir))),wbar2,sprintf('18. Co-registration to PAM50 standard space: Subject (%d) of (%d)',k,length(sub_dir))); catch, end
        fprintf('18. Co-registration to PAM50 standard space: Subject (%d) of (%d)\n',k,length(sub_dir))
        time_18_PAM50registration = tic;
        try
        scfMRItb_18_PAM50registration(sub_dir{k},anat_files{k}, wbar3,C_spine,SCT_vert_auto,SCT_manualreg_labels);
        catch
        msgbox(sprintf('PAM50 registration using SCT did not run successfully for subject %d of %d',k,length(sub_dir)),'PAM50 reg failed','error')
        end
        time_taken_preproc.step18_PAM50registration(k,1) = toc(time_18_PAM50registration); clear time_18_PAM50registration Rns
        clk=clock; L=L+1; logg{L,1}=sprintf('%d/%s/%d %dh:%dm:%ds - end (18) co-registration to PAM50 standard space for subject (%d) of (%d)',clk(3),month{clk(2)},clk(1),clk(4),clk(5),round(clk(6)),k,length(sub_dir)); try fprintf(fileID,'%s\n',logg{L,1}); set(S.tx,'string',logg); catch, end; clear clk
        refresh_html_log
    else
        clk=clock; L=L+1; logg{L,1}=sprintf('%d/%s/%d %dh:%dm:%ds - skip (18) co-registration to PAM50 standard space for subject (%d) of (%d)',clk(3),month{clk(2)},clk(1),clk(4),clk(5),round(clk(6)),k,length(sub_dir)); try fprintf(fileID,'%s\n',logg{L,1}); set(S.tx,'string',logg); catch, end; clear clk
    end
end

if skip==0
    clk=clock; L=L+1; logg{L,1}=sprintf('%d/%s/%d %dh:%dm:%ds - (18) co-registration to PAM50 standard space SKIPPED FOR ALL SUBJECTS',clk(3),month{clk(2)},clk(1),clk(4),clk(5),round(clk(6))); try fprintf(fileID,'%s\n',logg{L,1}); set(S.tx,'string',logg); catch, end; clear clk
    try figure(status_fig),subplot(N,2,35),imshow(imread('status_images/step_18_green.jpeg')), subplot(N,2,36),imshow(imread('status_images/status_green_skip.jpg')), catch, end
else
    clk=clock; L=L+1; logg{L,1}=sprintf('%d/%s/%d %dh:%dm:%ds - (18) co-registration to PAM50 standard space COMPLETED FOR ALL SUBJECTS',clk(3),month{clk(2)},clk(1),clk(4),clk(5),round(clk(6))); try fprintf(fileID,'%s\n',logg{L,1}); set(S.tx,'string',logg); catch, end; clear clk
    L=L+1; logg{L,1}=sprintf('Total time taken so far: %dh:%dm:%ds\nTime taken for this step: %dh:%dm:%ds\nTime taken for this step in each run of each subject: %dh:%dm:%ds',floor(toc(tc0)/3600),floor(mod(toc(tc0),3600)/60),floor(mod(toc(tc0),60)),floor(toc(tc)/3600),floor(mod(toc(tc),3600)/60),floor(mod(toc(tc),60)),floor(toc(tc)/3600/skip),floor(mod(toc(tc),3600)/60/skip),floor(mod(toc(tc),60)/skip)); try fprintf(fileID,'%s\n',logg{L,1}); set(S.tx,'string',logg); catch, end; clear clk
    try figure(status_fig),subplot(N,2,35),imshow(imread('status_images/step_18_green.jpeg')), subplot(N,2,36),imshow(imread('status_images/status_green.jpg')), catch, end
    refresh_html_log
end
end

%% Calculate tSNR across pre-processing steps
if (choices_preproc(5)==1)||(choices_preproc(6)==1)||(choices_preproc(8)==1)||(choices_preproc(9)==1)||(choices_preproc(11)==1)||(choices_preproc(12)==1)||(choices_preproc(13)==1)||(choices_preproc(14)==1)
L=L+2; logg{L,1}=sprintf('-------\nCalculate tSNR across pre-processing steps'); try fprintf(fileID,'%s\n',logg{L,1}); set(S.tx,'string',logg); catch, end; tc=tic;
time_taken_preproc.tSNR_computation = zeros(size(func_files,1),size(func_files,2));

skip=0;
for k=1:length(sub_dir)
    for k2=1:runs(k)
        if ~( (exist([sub_dir{k} 'tSNR_across_preproc_steps.mat'], 'file'))||(exist([sub_dir{k} 'tSNR_across_preproc_steps_run' num2str(k2) '.mat'], 'file')) ) || (run_allsubs_by_default==1)
            skip=skip+1;
            clk=clock; L=L+1; logg{L,1}=sprintf('%d/%s/%d %dh:%dm:%ds - begin calculating tSNR across pre-processing steps for subject (%d) of (%d), run (%d) of (%d)',clk(3),month{clk(2)},clk(1),clk(4),clk(5),round(clk(6)),k,length(sub_dir),k2,runs(k)); try fprintf(fileID,'%s\n',logg{L,1}); set(S.tx,'string',logg); catch, end; clear clk
            try waitbar(((runs(k)*(k-1)+k2)/(length(sub_dir)*runs(k))),wbar2,sprintf('Calculate tSNR across pre-processing steps: Subject (%d) of (%d), Run (%d) of (%d)',k,length(sub_dir),k2,runs(k))); catch, end
            fprintf('Calculate tSNR across pre-processing steps: Subject (%d) of (%d), Run (%d) of (%d)\n',k,length(sub_dir),k2,runs(k))
            if runs(k)==1, Rns=0; else, Rns=k2; end
            time_tSNR_graphs = tic;
            scfMRItb_tSNR_graphs(sub_dir{k},func_files{k,k2},anat_files{k,1},choices_preproc, wbar3,Rns);
            time_taken_preproc.tSNR_computation(k,k2) = toc(time_tSNR_graphs); clear time_tSNR_graphs Rns
            clk=clock; L=L+1; logg{L,1}=sprintf('%d/%s/%d %dh:%dm:%ds - end calculating tSNR across pre-processing steps for subject (%d) of (%d), run (%d) of (%d)',clk(3),month{clk(2)},clk(1),clk(4),clk(5),round(clk(6)),k,length(sub_dir),k2,runs(k)); try fprintf(fileID,'%s\n',logg{L,1}); set(S.tx,'string',logg); catch, end; clear clk
        else
            clk=clock; L=L+1; logg{L,1}=sprintf('%d/%s/%d %dh:%dm:%ds - skip calculating tSNR across pre-processing steps for subject (%d) of (%d), run (%d) of (%d)',clk(3),month{clk(2)},clk(1),clk(4),clk(5),round(clk(6)),k,length(sub_dir),k2,runs(k)); try fprintf(fileID,'%s\n',logg{L,1}); set(S.tx,'string',logg); catch, end; clear clk
        end
    end; clear k2
end

if skip==0
    clk=clock; L=L+1; logg{L,1}=sprintf('%d/%s/%d %dh:%dm:%ds - Calculate tSNR across pre-processing steps SKIPPED FOR ALL SUBJECTS',clk(3),month{clk(2)},clk(1),clk(4),clk(5),round(clk(6))); try fprintf(fileID,'%s\n',logg{L,1}); set(S.tx,'string',logg); catch, end; clear clk
else
    clk=clock; L=L+1; logg{L,1}=sprintf('%d/%s/%d %dh:%dm:%ds - Calculate tSNR across pre-processing steps COMPLETED FOR ALL SUBJECTS',clk(3),month{clk(2)},clk(1),clk(4),clk(5),round(clk(6))); try fprintf(fileID,'%s\n',logg{L,1}); set(S.tx,'string',logg); catch, end; clear clk
    L=L+1; logg{L,1}=sprintf('Total time taken so far: %dh:%dm:%ds\nTime taken for this step: %dh:%dm:%ds\nTime taken for this step in each run of each subject: %dh:%dm:%ds',floor(toc(tc0)/3600),floor(mod(toc(tc0),3600)/60),floor(mod(toc(tc0),60)),floor(toc(tc)/3600),floor(mod(toc(tc),3600)/60),floor(mod(toc(tc),60)),floor(toc(tc)/3600/skip),floor(mod(toc(tc),3600)/60/skip),floor(mod(toc(tc),60)/skip)); try fprintf(fileID,'%s\n',logg{L,1}); set(S.tx,'string',logg); catch, end; clear clk
    refresh_html
end
end

%% '19. gzip NIfTI files to save disk space'
for k=1:length(sub_dir) % size of subject folder before removal of slicewise files and gzipping
    sdirall = dir([sub_dir{k} '**/*.*']); sdirall = sdirall(3:end);
    sdirall_bytes = {sdirall.bytes}';
    sdirall_bytes = cell2mat(sdirall_bytes);
    sdirall_bytes = sum(sdirall_bytes)/10^9; % size of subject folder in gigabytes
    folder_size_1(k,1) = sdirall_bytes;
    clear sdirall sdirall_bytes
end

if remove_slicewise_files==1
    if ~exist('wbar3','var')
        wbar3 = waitbar(0,'14. Save denoised 3D+time fMRI data...','Name','Progress(14): Save denoised 3D+time fMRI data...','Units','normalized','Position',[3/5 0 1/5 1/17]);
    end
    for k=1:length(sub_dir)
        fprintf('19. Remove slice-wise files: Subject (%d) of (%d)\n',k,length(sub_dir))
        time_19_remove_slicewise = tic;
        scfMRItb_19_remove_slicewise_files(sub_dir{k},func_files(k,:),anat_files{k,1},wbar3)
        time_taken_preproc.step19_remove_slicewise_files(k,1) = toc(time_19_remove_slicewise); clear time_19_remove_slicewise %#ok<*SAGROW>
    end
    for k=1:length(sub_dir) % size of subject folder after removal of slicewise files but before gzipping
        sdirall = dir([sub_dir{k} '**/*.*']); sdirall = sdirall(3:end);
        sdirall_bytes = {sdirall.bytes}';
        sdirall_bytes = cell2mat(sdirall_bytes);
        sdirall_bytes = sum(sdirall_bytes)/10^9; % size of subject folder in gigabytes
        folder_size_2(k,1) = sdirall_bytes;
        clear sdirall sdirall_bytes
    end
end

if choices_preproc(19)==1
try figure(status_fig),subplot(N,2,37),imshow(imread('status_images/step_19_blue.jpeg')), subplot(N,2,38),imshow(imread('status_images/status_blue.jpg')), catch, end
try L=L+2; logg{L,1}=sprintf('-------\n%s',preprocsteps{19}); catch, end; try fprintf(fileID,'%s\n',logg{L,1}); set(S.tx,'string',logg); catch, end; tc=tic;

if ~exist('answer_gzip','var')
    answer_gzip = 'both gzip and delete';
end
skip=0;
for k=1:length(sub_dir)
    switch answer_gzip
        case 'gzip only, don''t delete'
            try clk=clock; L=L+1; logg{L,1}=sprintf('%d/%s/%d %dh:%dm:%ds - gzip NIfTI files for subject (%d) of (%d)',clk(3),month{clk(2)},clk(1),clk(4),clk(5),round(clk(6)),k,length(sub_dir)); catch, end; try fprintf(fileID,'%s\n',logg{L,1}); set(S.tx,'string',logg); catch, end; clear clk
            try waitbar((k/length(sub_dir)),wbar2,sprintf('19. gzip NIfTI files: Subject (%d) of (%d)',k,length(sub_dir))); catch, end
            fprintf('19. gzip NIfTI files: Subject (%d) of (%d)\n',k,length(sub_dir))
            time_19_gzip_nii = tic;
            skp = scfMRItb_19_gzip_nii(sub_dir{k},wbar3); skip=skip+single(skp>0); clear skp
            time_taken_preproc.step19_gzip_nii(k,1) = toc(time_19_gzip_nii); clear time_19_gzip_nii %#ok<*SAGROW>
        case 'delete only, I already gzipped'
            try clk=clock; L=L+1; logg{L,1}=sprintf('%d/%s/%d %dh:%dm:%ds - delete .nii files for subject (%d) of (%d)',clk(3),month{clk(2)},clk(1),clk(4),clk(5),round(clk(6)),k,length(sub_dir)); catch, end; try fprintf(fileID,'%s\n',logg{L,1}); set(S.tx,'string',logg); catch, end; clear clk
            try waitbar((k/length(sub_dir)),wbar2,sprintf('19. Delete .nii files to save disk space (.nii.gz preserved): Subject (%d) of (%d)',k,length(sub_dir))); catch, end
            fprintf('19. Delete .nii files to save disk space (.nii.gz preserved): Subject (%d) of (%d)\n',k,length(sub_dir))
            time_19_delete_nii = tic;
            skp = scfMRItb_19_delete_nii(sub_dir{k},wbar3); skip=skip+single(skp>0); clear skp
            time_taken_preproc.step19_delete_nii(k,1) = toc(time_19_delete_nii); clear time_19_delete_nii
        case 'both gzip and delete'
            try clk=clock; L=L+1; logg{L,1}=sprintf('%d/%s/%d %dh:%dm:%ds - gzip NIfTI files for subject (%d) of (%d)',clk(3),month{clk(2)},clk(1),clk(4),clk(5),round(clk(6)),k,length(sub_dir)); catch, end; try fprintf(fileID,'%s\n',logg{L,1}); set(S.tx,'string',logg); catch, end; clear clk
            try waitbar((k/length(sub_dir)),wbar2,sprintf('19. gzip NIfTI files: Subject (%d) of (%d)',k,length(sub_dir))); catch, end
            fprintf('19. gzip NIfTI files: Subject (%d) of (%d)\n',k,length(sub_dir))
            time_19_gzip_delete_nii = tic;
            skp = scfMRItb_19_gzip_nii(sub_dir{k},wbar3); skip=skip+single(skp>0); clear skp
            try clk=clock; L=L+1; logg{L,1}=sprintf('%d/%s/%d %dh:%dm:%ds - delete .nii files for subject (%d) of (%d)',clk(3),month{clk(2)},clk(1),clk(4),clk(5),round(clk(6)),k,length(sub_dir)); catch, end; try fprintf(fileID,'%s\n',logg{L,1}); set(S.tx,'string',logg); catch, end; clear clk
            try waitbar((k/length(sub_dir)),wbar2,sprintf('19. Delete .nii files to save disk space (.nii.gz preserved): Subject (%d) of (%d)',k,length(sub_dir))); catch, end
            fprintf('19. Delete .nii files to save disk space (.nii.gz preserved): Subject (%d) of (%d)\n',k,length(sub_dir))
            skp = scfMRItb_19_delete_nii(sub_dir{k},wbar3); skip=skip+single(skp>0); clear skp
            time_taken_preproc.step19_gzip_delete_nii(k,1) = toc(time_19_gzip_delete_nii); clear time_19_gzip_delete_nii
    end
    refresh_html_log
end; clear k

for k=1:length(sub_dir) % size of subject folder after removal of slicewise files and gzipping
    sdirall = dir([sub_dir{k} '**/*.*']); sdirall = sdirall(3:end);
    sdirall_name = {sdirall.name}';
    excludefile = zeros(length(sdirall_name),1);
    for m=1:length(sdirall_name)
        if length(sdirall_name{m})>=4
            if strcmp(sdirall_name{m}(end-3:end),'.nii')==1
                for m2=1:length(sdirall_name)
                    if length(sdirall_name{m2})>=7
                        if (strcmp(sdirall_name{m}(1:end-4),sdirall_name{m2}(1:end-7))==1) && (strcmp(sdirall_name{m2}(end-6:end),'.nii.gz')==1) && (m~=m2)
                            excludefile(m) = 1; % if a .nii file is found AND if a corresponding .nii.gz file also exists, then do not count the .nii file for total storage computation
                        end
                    end
                end
            end
        end
    end
    sdirall_bytes = {sdirall.bytes}';
    sdirall_bytes = cell2mat(sdirall_bytes);
    sdirall_bytes = sdirall_bytes(find(excludefile==0));
    sdirall_bytes = sum(sdirall_bytes)/10^9; % size of subject folder in gigabytes
    folder_size_3(k,1) = sdirall_bytes;
    clear sdirall sdirall_bytes sdirall_name m m2 excludefile
end

if skip==0
    try clk=clock; L=L+1; logg{L,1}=sprintf('%d/%s/%d %dh:%dm:%ds - gzip NIfTI files SKIPPED FOR ALL SUBJECTS',clk(3),month{clk(2)},clk(1),clk(4),clk(5),round(clk(6))); catch, end; try fprintf(fileID,'%s\n',logg{L,1}); set(S.tx,'string',logg); catch, end; clear clk
    try figure(status_fig),subplot(N,2,37),imshow(imread('status_images/step_19_green.jpeg')), subplot(N,2,38),imshow(imread('status_images/status_green_skip.jpg')), catch, end
else
    try clk=clock; L=L+1; logg{L,1}=sprintf('%d/%s/%d %dh:%dm:%ds - gzip NIfTI files COMPLETED FOR ALL SUBJECTS',clk(3),month{clk(2)},clk(1),clk(4),clk(5),round(clk(6))); catch, end; try fprintf(fileID,'%s\n',logg{L,1}); set(S.tx,'string',logg); catch, end; clear clk
    try L=L+1; logg{L,1}=sprintf('Total time taken so far: %dh:%dm:%ds\nTime taken for this step: %dh:%dm:%ds\nTime taken for this step in each subject: %dh:%dm:%ds',floor(toc(tc0)/3600),floor(mod(toc(tc0),3600)/60),floor(mod(toc(tc0),60)),floor(toc(tc)/3600),floor(mod(toc(tc),3600)/60),floor(mod(toc(tc),60)),floor(toc(tc)/3600/skip),floor(mod(toc(tc),3600)/60/skip),floor(mod(toc(tc),60)/skip)); catch, end; try fprintf(fileID,'%s\n',logg{L,1}); set(S.tx,'string',logg); catch, end; clear clk
    try figure(status_fig),subplot(N,2,37),imshow(imread('status_images/step_19_green.jpeg')), subplot(N,2,38),imshow(imread('status_images/status_green.jpg')), catch, end
    refresh_html_log
end
end

%% Save disk storage info and figures
tmpc=clock; clk = sprintf('%.4d-%.2d-%.2d_%.2d-%.2d',tmpc(1),tmpc(2),tmpc(3),tmpc(4),tmpc(5)); clear tmpc
dskfilename = sprintf('disk_storage_space_info_%s.mat',clk); clear clk

% save disk storage info and figures
for k=1:length(sub_dir)
    % save disk storage info into .mat file
    if exist('folder_size_2','var') && exist('folder_size_3','var')
        folder_size_1_sub = folder_size_1(k); folder_size_2_sub = folder_size_2(k); folder_size_3_sub = folder_size_3(k);
        readme_disk_storage = sprintf('These numbers are the total storage space (in gigabytes) taken by all files in this subject folder (%s)\nfolder_size_1_sub = %.2f (size of subject folder after pre-processing before removal of slicewise files and gzipping)\nfolder_size_2_sub = %.2f (size of subject folder after pre-processing and removal of slicewise files but before gzipping)\nfolder_size_3_sub = %.2f (size of subject folder after pre-processing, removal of slicewise files and gzipping)\n',sub_dir{k},folder_size_1_sub,folder_size_2_sub,folder_size_3_sub);
        save([sub_dir{k} dskfilename],'folder_size_1_sub','folder_size_2_sub','folder_size_3_sub','readme_disk_storage')
    elseif exist('folder_size_2','var') && ~exist('folder_size_3','var')
        folder_size_1_sub = folder_size_1(k); folder_size_2_sub = folder_size_2(k);
        readme_disk_storage = sprintf('These numbers are the total storage space (in gigabytes) taken by all files in this subject folder (%s)\nfolder_size_1_sub = %.2f (size of subject folder after pre-processing before removal of slicewise files and gzipping)\nfolder_size_2_sub = %.2f (size of subject folder after pre-processing and removal of slicewise files but before gzipping)\nfolder_size_3_sub = <"gzipping" not performed> (size of subject folder after pre-processing, removal of slicewise files and gzipping)\n',sub_dir{k},folder_size_1_sub,folder_size_2_sub);
        save([sub_dir{k} dskfilename],'folder_size_1_sub','folder_size_2_sub','readme_disk_storage')
    elseif ~exist('folder_size_2','var') && exist('folder_size_3','var')
        folder_size_1_sub = folder_size_1(k); folder_size_3_sub = folder_size_3(k);
        readme_disk_storage = sprintf('These numbers are the total storage space (in gigabytes) taken by all files in this subject folder (%s)\nfolder_size_1_sub = %.2f (size of subject folder after pre-processing before removal of slicewise files and gzipping)\nfolder_size_2_sub = <"removal of slicewise files" not performed> (size of subject folder after pre-processing and removal of slicewise files but before gzipping)\nfolder_size_3_sub = %.2f (size of subject folder after pre-processing, removal of slicewise files and gzipping)\n',sub_dir{k},folder_size_1_sub,folder_size_3_sub);
        save([sub_dir{k} dskfilename],'folder_size_1_sub','folder_size_3_sub','readme_disk_storage')
    elseif ~exist('folder_size_2','var') && ~exist('folder_size_3','var')
        folder_size_1_sub = folder_size_1(k);
        readme_disk_storage = sprintf('These numbers are the total storage space (in gigabytes) taken by all files in this subject folder (%s)\nfolder_size_1_sub = %.2f (size of subject folder after pre-processing before removal of slicewise files and gzipping)\nfolder_size_2_sub = <"removal of slicewise files" not performed> (size of subject folder after pre-processing and removal of slicewise files but before gzipping)\nfolder_size_3_sub = <"gzipping" not performed> (size of subject folder after pre-processing, removal of slicewise files and gzipping)\n',sub_dir{k},folder_size_1_sub);
        save([sub_dir{k} dskfilename],'folder_size_1_sub','readme_disk_storage')
    end

    % save bar graphs
    if ~(exist([sub_dir{k} 'QC' '/19_disk_storage_info'],'dir'))
        mkdir([sub_dir{k} 'QC' '/19_disk_storage_info'])
    end
    QCpath = [sub_dir{k} 'QC' '/19_disk_storage_info/'];
    if ~exist('folder_size_2_sub','var'), folder_size_2_sub = 0; end
    if ~exist('folder_size_3_sub','var'), folder_size_3_sub = 0; end
    
    fighndl = figure('Position',[1 1 1200 800],'Color',[0.85,0.90,0.85],'InvertHardcopy','off','Visible','off'); % [0.85,0.90,0.85] [0.8, 0.89, 0.94]
    b = bar(1,folder_size_1_sub,0.5,'FaceColor',[0,0.55,0.69],'EdgeColor',[0.65,0.11,0.19],'LineWidth',1.5); hold on,
        xtips1 = b(1).XEndPoints; ytips1 = b(1).YEndPoints; labels1 = string(round(b(1).YData,2)); clear b
        labels1 = strcat(labels1,' GB');
        text(xtips1,ytips1,labels1,'HorizontalAlignment','center','VerticalAlignment','bottom','fontsize',17','color',[0.65,0.11,0.19]);
    b = bar(2,folder_size_2_sub,0.5,'FaceColor',[0,0.65,0.69],'EdgeColor',[0.65,0.11,0.19],'LineWidth',1.5);
        xtips2 = b(1).XEndPoints; ytips2 = b(1).YEndPoints; labels2 = string(round(b(1).YData,2)); clear b
        labels2 = strcat(labels2,' GB');
        if folder_size_2_sub==0, labels2 = 'not performed'; end
        text(xtips2,ytips2,labels2,'HorizontalAlignment','center','VerticalAlignment','bottom','fontsize',17','color',[0.65,0.11,0.19]);
    b = bar(3,folder_size_3_sub,0.5,'FaceColor',[0,0.75,0.69],'EdgeColor',[0.65,0.11,0.19],'LineWidth',1.5);
        xtips3 = b(1).XEndPoints; ytips3 = b(1).YEndPoints; labels3 = string(round(b(1).YData,2)); clear b
        labels3 = strcat(labels3,' GB');
        if folder_size_3_sub==0, labels3 = 'not performed'; end
        text(xtips3,ytips3,labels3,'HorizontalAlignment','center','VerticalAlignment','bottom','fontsize',17','color',[0.65,0.11,0.19]);
        grid on, grid minor, ylim([0 max(folder_size_1_sub)+0.1.*max(folder_size_1_sub)]), ylabel('Disk storage (in gigabytes)'),
        xticks([1,2,3]), xticklabels({'At the end of pre-processing','After removal of slice-wise files','After NIfTI file compression (gzip)'})
        a = get(gca,'xticklabels'); set(gca,'xticklabels',a,'fontsize',14); clear a
        legstr{1} = sprintf('At the end of pre-processing'); legstr{2} = sprintf('... AND after removal of duplicate slice-wise files'); legstr{3} = sprintf('... AND after NIfTI file compression (gzip)');
        legend(legstr,'Location','northeast','FontSize',18)
        title(sprintf('DISK STORAGE: size of subject folder at the end of pre-processing\n(%s)',sub_dir{k}),'Interpreter', 'none', 'Color',[0 0.37 0.22],'FontSize',18);
    saveas(fighndl,[QCpath 'disk_storage_bargraph' '.jpg'])
    tmp=clock; clk_settings = sprintf('%.4d-%.2d-%.2d_%.2d-%.2d',tmp(1),tmp(2),tmp(3),tmp(4),tmp(5)); clear tmp
    saveas(fighndl,[QCpath sprintf('disk_storage_bargraph_%s',clk_settings) '.jpg'])
    close(fighndl)
    clear folder_size_1_sub folder_size_2_sub folder_size_3_sub readme_disk_storage QCpath fighndl legstr b xtips1 ytips1 labels1 xtips2 ytips2 labels2 xtips3 ytips3 labels3 clk_settings
end
clear k dskfilename


end
%%
%%
%% --- if step_after_step==0 (i.e. do subject-after-subject) ---
if step_after_step==0
    if ~exist('sub_aft_sub_num','var'), sub_aft_sub_num=1; end

%%
for k=sub_aft_sub_num:length(sub_dir)
    for k2=1:runs(k)
        L=L+2; logg{L,1}=sprintf('-------\nPre-processing Subject (%d) of (%d), Run (%d) of (%d)',k,length(sub_dir),k2,runs(k)); try fprintf(fileID,'%s\n',logg{L,1}); set(S.tx,'string',logg); catch, end; tc=tic;

%% initial prep
if ignore_extreme_slices~=0
    if ( (exist([sub_dir{1} anat_files{1} '_origSlices.nii'], 'file')) || (exist([sub_dir{1} anat_files{1} '_origSlices.nii.gz'], 'file')) )
        scfMRItb_04_unzipFile(sub_dir{1},anat_files{1}, '')
        origdata = load_untouch_nii([sub_dir{1} anat_files{1} '.nii']);
        slices = size(origdata.img,3); clear origdata
    end
    if (k==1) && ( ~( (exist([sub_dir{k} anat_files{k} '_origSlices.nii'], 'file')) || (exist([sub_dir{k} anat_files{k} '_origSlices.nii.gz'], 'file')) ) || (run_allsubs_by_default==1) )
        clk=clock; L=L+1; logg{L,1}=sprintf('%d/%s/%d %dh:%dm:%ds - begin ignoring first and last anatomical slice for subject (%d) of (%d)',clk(3),month{clk(2)},clk(1),clk(4),clk(5),round(clk(6)),k,length(sub_dir)); try fprintf(fileID,'%s\n',logg{L,1}); set(S.tx,'string',logg); catch, end; clear clk
        try waitbar((k/length(sub_dir)),wbar2,sprintf('Ignore first/last anatomical slices: Subject (%d) of (%d)',k,length(sub_dir))); catch, end
        fprintf('00. Ignore first and last anatomical slice: Subject (%d) of (%d), Run (%d) of (%d)\n',k,length(sub_dir),k2,runs(k))
        time_ignoreExtremeSlices = tic;
        scfMRItb_04_unzipFile(sub_dir{k},anat_files{k}, '')
        origdata = load_untouch_nii([sub_dir{k} anat_files{k} '.nii']);
        origdata.img = origdata.img(:,:,2:end-1,:);
        origdata.hdr.dime.dim(4) = origdata.hdr.dime.dim(4)-2;
        movefile([sub_dir{k} anat_files{k} '.nii'],[sub_dir{k} anat_files{k} '_origSlices.nii'])
        if exist([sub_dir{k} anat_files{k} '.nii.gz'],'file')
            movefile([sub_dir{k} anat_files{k} '.nii.gz'],[sub_dir{k} anat_files{k} '_origSlices.nii.gz'])
        else
            gzip([sub_dir{k} anat_files{k} '_origSlices.nii'])
        end
        save_untouch_nii(origdata,[sub_dir{k} anat_files{k} '.nii']);
        gzip([sub_dir{k} anat_files{k} '.nii'])
        clear origdata
        if exist('NORDIC_type','var') && (NORDIC_type==2)
            try
                scfMRItb_04_unzipFile(sub_dir{k},anat_files{k}, '_ph')
                origdata = load_untouch_nii([sub_dir{k} anat_files{k} '_ph.nii']);
                origdata.img = origdata.img(:,:,2:end-1);
                origdata.hdr.dime.dim(4) = origdata.hdr.dime.dim(4)-2;
                movefile([sub_dir{k} anat_files{k} '_ph.nii'],[sub_dir{k} anat_files{k} '_ph_origSlices.nii'])
                if exist([sub_dir{k} anat_files{k} '_ph.nii.gz'],'file')
                    movefile([sub_dir{k} anat_files{k} '_ph.nii.gz'],[sub_dir{k} anat_files{k} '_ph_origSlices.nii.gz'])
                else
                    gzip([sub_dir{k} anat_files{k} '_ph_origSlices.nii'])
                end
                save_untouch_nii(origdata,[sub_dir{k} anat_files{k} '_ph.nii']);
                gzip([sub_dir{k} anat_files{k} '_ph.nii'])
                clear origdata
            catch
            end
        end
        time_taken_preproc.step00_ignoreExtremeSlices(k,k2) = toc(time_ignoreExtremeSlices); clear time_ignoreExtremeSlices Rns
        clk=clock; L=L+1; logg{L,1}=sprintf('%d/%s/%d %dh:%dm:%ds - end ignoring first and last anatomical slice for subject (%d) of (%d)',clk(3),month{clk(2)},clk(1),clk(4),clk(5),round(clk(6)),k,length(sub_dir)); try fprintf(fileID,'%s\n',logg{L,1}); set(S.tx,'string',logg); catch, end; clear clk
        refresh_html_log
    else
        clk=clock; L=L+1; logg{L,1}=sprintf('%d/%s/%d %dh:%dm:%ds - did not ignore first and last anatomical slice for subject (%d) of (%d)',clk(3),month{clk(2)},clk(1),clk(4),clk(5),round(clk(6)),k,length(sub_dir)); try fprintf(fileID,'%s\n',logg{L,1}); set(S.tx,'string',logg); catch, end; clear clk
    end
    if ~( (exist([sub_dir{k} func_files{k,k2} '_origSlices.nii'], 'file')) || (exist([sub_dir{k} func_files{k,k2} '_origSlices.nii.gz'], 'file')) ) || (run_allsubs_by_default==1)
        clk=clock; L=L+1; logg{L,1}=sprintf('%d/%s/%d %dh:%dm:%ds - begin ignoring first and last functional slice for subject (%d) of (%d), run (%d) of (%d)',clk(3),month{clk(2)},clk(1),clk(4),clk(5),round(clk(6)),k,length(sub_dir),k2,runs(k)); try fprintf(fileID,'%s\n',logg{L,1}); set(S.tx,'string',logg); catch, end; clear clk
        try waitbar(((runs(k)*(k-1)+k2)/(length(sub_dir)*runs(k))),wbar2,sprintf('Ignore first/last functional slices: Subject (%d) of (%d), Run (%d) of (%d)',k,length(sub_dir),k2,runs(k))); catch, end
        fprintf('00. Ignore first and last functional slice: Subject (%d) of (%d), Run (%d) of (%d)\n',k,length(sub_dir),k2,runs(k))
        time_ignoreExtremeSlices = tic;
        scfMRItb_04_unzipFile(sub_dir{k},func_files{k,k2}, '')
        origdata = load_untouch_nii([sub_dir{k} func_files{k,k2} '.nii']);
        origdata.img = origdata.img(:,:,2:end-1,:);
        origdata.hdr.dime.dim(4) = origdata.hdr.dime.dim(4)-2;
        movefile([sub_dir{k} func_files{k,k2} '.nii'],[sub_dir{k} func_files{k,k2} '_origSlices.nii'])
        if exist([sub_dir{k} func_files{k,k2} '.nii.gz'],'file')
            movefile([sub_dir{k} func_files{k,k2} '.nii.gz'],[sub_dir{k} func_files{k,k2} '_origSlices.nii.gz'])
        else
            gzip([sub_dir{k} func_files{k,k2} '_origSlices.nii'])
        end
        save_untouch_nii(origdata,[sub_dir{k} func_files{k,k2} '.nii']);
        gzip([sub_dir{k} func_files{k,k2} '.nii'])
        clear origdata
        if exist('NORDIC_type','var')
        if NORDIC_type==2
            scfMRItb_04_unzipFile(sub_dir{k},func_files{k,k2}, '_ph')
            origdata = load_untouch_nii([sub_dir{k} func_files{k,k2} '_ph.nii']);
            origdata.img = origdata.img(:,:,2:end-1,:);
            origdata.hdr.dime.dim(4) = origdata.hdr.dime.dim(4)-2;
            movefile([sub_dir{k} func_files{k,k2} '_ph.nii'],[sub_dir{k} func_files{k,k2} '_ph_origSlices.nii'])
            if exist([sub_dir{k} func_files{k,k2} '_ph.nii.gz'],'file')
                movefile([sub_dir{k} func_files{k,k2} '_ph.nii.gz'],[sub_dir{k} func_files{k,k2} '_ph_origSlices.nii.gz'])
            else
                gzip([sub_dir{k} func_files{k,k2} '_ph_origSlices.nii'])
            end
            save_untouch_nii(origdata,[sub_dir{k} func_files{k,k2} '_ph.nii']);
            gzip([sub_dir{k} func_files{k,k2} '_ph.nii'])
            clear origdata
        end
        end
        time_taken_preproc.step00_ignoreExtremeSlices(k,k2) = toc(time_ignoreExtremeSlices); clear time_ignoreExtremeSlices Rns
        clk=clock; L=L+1; logg{L,1}=sprintf('%d/%s/%d %dh:%dm:%ds - end ignoring first and last functional slice for subject (%d) of (%d), run (%d) of (%d)',clk(3),month{clk(2)},clk(1),clk(4),clk(5),round(clk(6)),k,length(sub_dir),k2,runs(k)); try fprintf(fileID,'%s\n',logg{L,1}); set(S.tx,'string',logg); catch, end; clear clk
        refresh_html_log
    else
        clk=clock; L=L+1; logg{L,1}=sprintf('%d/%s/%d %dh:%dm:%ds - did not ignore first and last functional slice for subject (%d) of (%d), run (%d) of (%d)',clk(3),month{clk(2)},clk(1),clk(4),clk(5),round(clk(6)),k,length(sub_dir),k2,runs(k)); try fprintf(fileID,'%s\n',logg{L,1}); set(S.tx,'string',logg); catch, end; clear clk
    end
end

if discard_firstN_volumes~=0
    if ~( (exist([sub_dir{k} func_files{k,k2} '_origLength.nii'], 'file')) || (exist([sub_dir{k} func_files{k,k2} '_origLength.nii.gz'], 'file')) ) || (run_allsubs_by_default==1)
        clk=clock; L=L+1; logg{L,1}=sprintf('%d/%s/%d %dh:%dm:%ds - begin discarding first %d volumes for subject (%d) of (%d), run (%d) of (%d)',clk(3),month{clk(2)},clk(1),clk(4),clk(5),round(clk(6)),discard_firstN_volumes,k,length(sub_dir),k2,runs(k)); try fprintf(fileID,'%s\n',logg{L,1}); set(S.tx,'string',logg); catch, end; clear clk
        try waitbar(((runs(k)*(k-1)+k2)/(length(sub_dir)*runs(k))),wbar2,sprintf('06. Discard first %d volumes: Subject (%d) of (%d), Run (%d) of (%d)',discard_firstN_volumes,k,length(sub_dir),k2,runs(k))); catch, end
        fprintf('00. Discard first %d volumes: Subject (%d) of (%d), Run (%d) of (%d)\n',discard_firstN_volumes,k,length(sub_dir),k2,runs(k))
        time_discardNvolumes = tic;
        scfMRItb_04_unzipFile(sub_dir{k},func_files{k,k2}, '')
        origdata = load_untouch_nii([sub_dir{k} func_files{k,k2} '.nii']);
        origdata.img = origdata.img(:,:,:,discard_firstN_volumes+1:end);
        origdata.hdr.dime.dim(5) = origdata.hdr.dime.dim(5)-discard_firstN_volumes;
        movefile([sub_dir{k} func_files{k,k2} '.nii'],[sub_dir{k} func_files{k,k2} '_origLength.nii'])
        if exist([sub_dir{k} func_files{k,k2} '.nii.gz'],'file')
            movefile([sub_dir{k} func_files{k,k2} '.nii.gz'],[sub_dir{k} func_files{k,k2} '_origLength.nii.gz'])
        else
            gzip([sub_dir{k} func_files{k,k2} '_origLength.nii'])
        end
        save_untouch_nii(origdata,[sub_dir{k} func_files{k,k2} '.nii']);
        gzip([sub_dir{k} func_files{k,k2} '.nii'])
        clear origdata
        if exist('NORDIC_type','var')
        if NORDIC_type==2
            scfMRItb_04_unzipFile(sub_dir{k},func_files{k,k2}, '_ph')
            origdata = load_untouch_nii([sub_dir{k} func_files{k,k2} '_ph.nii']);
            origdata.img = origdata.img(:,:,:,discard_firstN_volumes+1:end);
            origdata.hdr.dime.dim(5) = origdata.hdr.dime.dim(5)-discard_firstN_volumes;
            movefile([sub_dir{k} func_files{k,k2} '_ph.nii'],[sub_dir{k} func_files{k,k2} '_ph_origLength.nii'])
            if exist([sub_dir{k} func_files{k,k2} '_ph.nii.gz'],'file')
                movefile([sub_dir{k} func_files{k,k2} '_ph.nii.gz'],[sub_dir{k} func_files{k,k2} '_ph_origLength.nii.gz'])
            else
                gzip([sub_dir{k} func_files{k,k2} '_ph_origLength.nii'])
            end
            save_untouch_nii(origdata,[sub_dir{k} func_files{k,k2} '_ph.nii']);
            gzip([sub_dir{k} func_files{k,k2} '_ph.nii'])
            clear origdata
        end
        end
        time_taken_preproc.step00_discardNvolumes(k,k2) = toc(time_discardNvolumes); clear time_discardNvolumes Rns
        clk=clock; L=L+1; logg{L,1}=sprintf('%d/%s/%d %dh:%dm:%ds - end discarding first %d volumes for subject (%d) of (%d), run (%d) of (%d)',clk(3),month{clk(2)},clk(1),clk(4),clk(5),round(clk(6)),discard_firstN_volumes,k,length(sub_dir),k2,runs(k)); try fprintf(fileID,'%s\n',logg{L,1}); set(S.tx,'string',logg); catch, end; clear clk
        refresh_html_log
    else
        clk=clock; L=L+1; logg{L,1}=sprintf('%d/%s/%d %dh:%dm:%ds - skip discarding first %d volumes for subject (%d) of (%d), run (%d) of (%d)',clk(3),month{clk(2)},clk(1),clk(4),clk(5),round(clk(6)),discard_firstN_volumes,k,length(sub_dir),k2,runs(k)); try fprintf(fileID,'%s\n',logg{L,1}); set(S.tx,'string',logg); catch, end; clear clk
    end
end

%% '01. NORDIC denoising'
if choices_preproc(1)==1
    if ~( (exist([sub_dir{k} func_files{k,k2} '_beforeNORDIC.nii'], 'file')) || (exist([sub_dir{k} func_files{k,k2} '_beforeNORDIC.nii.gz'], 'file')) ) || (run_allsubs_by_default==1)
        try figure(status_fig),subplot(N,2,1),imshow(imread('status_images/step_01_blue.jpeg')), subplot(N,2,2),imshow(imread('status_images/status_blue.jpg')), catch, end
        clk=clock; L=L+1; logg{L,1}=sprintf('%d/%s/%d %dh:%dm:%ds - begin (01) NORDIC denoising for subject (%d) of (%d), run (%d) of (%d)',clk(3),month{clk(2)},clk(1),clk(4),clk(5),round(clk(6)),k,length(sub_dir),k2,runs(k)); try fprintf(fileID,'%s\n',logg{L,1}); set(S.tx,'string',logg); catch, end; clear clk
        try waitbar(((runs(k)*(k-1)+k2)/(length(sub_dir)*runs(k))),wbar2,sprintf('01. NORDIC denoising: Subject (%d) of (%d), Run (%d) of (%d)',k,length(sub_dir),k2,runs(k))); catch, end
        fprintf('01. Perform NORDIC denoising: Subject (%d) of (%d), Run (%d) of (%d)\n',k,length(sub_dir),k2,runs(k))
        if runs(k)==1, Rns=0; else, Rns=k2; end
        time_01_NORDICdenoising = tic;
        scfMRItb_01_NORDIC_denoising(sub_dir{k},func_files{k,k2},anat_files{k,1},NORDIC_type,k2, wbar3,Rns);
        time_taken_preproc.step01_NORDICdenoising(k,k2) = toc(time_01_NORDICdenoising); clear time_01_NORDICdenoising Rns
        try figure(status_fig),subplot(N,2,1),imshow(imread('status_images/step_01_green.jpeg')), subplot(N,2,2),imshow(imread('status_images/status_green.jpg')), catch, end
        clk=clock; L=L+1; logg{L,1}=sprintf('%d/%s/%d %dh:%dm:%ds - end (01) NORDIC denoising for subject (%d) of (%d), run (%d) of (%d)',clk(3),month{clk(2)},clk(1),clk(4),clk(5),round(clk(6)),k,length(sub_dir),k2,runs(k)); try fprintf(fileID,'%s\n',logg{L,1}); set(S.tx,'string',logg); catch, end; clear clk
        refresh_html
    else
        clk=clock; L=L+1; logg{L,1}=sprintf('%d/%s/%d %dh:%dm:%ds - skip (01) NORDIC denoising for subject (%d) of (%d), run (%d) of (%d)',clk(3),month{clk(2)},clk(1),clk(4),clk(5),round(clk(6)),k,length(sub_dir),k2,runs(k)); try fprintf(fileID,'%s\n',logg{L,1}); set(S.tx,'string',logg); catch, end; clear clk
        try figure(status_fig),subplot(N,2,1),imshow(imread('status_images/step_01_green.jpeg')), subplot(N,2,2),imshow(imread('status_images/status_green_skip.jpg')), catch, end
    end
end

%% '02. Slice-timing correction'
if choices_preproc(2)==1
    if ~(exist([sub_dir{k} func_files{k,k2} 'before_slicetiming' '.nii'], 'file')) || (run_allsubs_by_default==1)
        try figure(status_fig),subplot(N,2,3),imshow(imread('status_images/step_02_blue.jpeg')), subplot(N,2,4),imshow(imread('status_images/status_blue_UI.jpg')), catch, end
        clk=clock; L=L+1; logg{L,1}=sprintf('%d/%s/%d %dh:%dm:%ds - begin (02) slice timing correction for subject (%d) of (%d), run (%d) of (%d)',clk(3),month{clk(2)},clk(1),clk(4),clk(5),round(clk(6)),k,length(sub_dir),k2,runs(k)); try fprintf(fileID,'%s\n',logg{L,1}); set(S.tx,'string',logg); catch, end; clear clk
        try waitbar(((runs(k)*(k-1)+k2)/(length(sub_dir)*runs(k))),wbar2,sprintf('02. Slice-timing correction: Subject (%d) of (%d), Run (%d) of (%d)',k,length(sub_dir),k2,runs(k))); catch, end
        fprintf('02. Slice-timing correction: Subject (%d) of (%d), Run (%d) of (%d)\n',k,length(sub_dir),k2,runs(k))
        time_02_slicetimingcorrection = tic;
        scfMRItb_02_slicetimingcorrection(sub_dir{k},func_files{k,k2},sliceorder');
        time_taken_preproc.step02_slicetimingcorrection(k,k2) = toc(time_02_slicetimingcorrection); clear time_02_slicetimingcorrection
        try figure(status_fig),subplot(N,2,3),imshow(imread('status_images/step_02_green.jpeg')), subplot(N,2,4),imshow(imread('status_images/status_green.jpg')), catch, end
        clk=clock; L=L+1; logg{L,1}=sprintf('%d/%s/%d %dh:%dm:%ds - end (02) slice timing correction for subject (%d) of (%d), run (%d) of (%d)',clk(3),month{clk(2)},clk(1),clk(4),clk(5),round(clk(6)),k,length(sub_dir),k2,runs(k)); try fprintf(fileID,'%s\n',logg{L,1}); set(S.tx,'string',logg); catch, end; clear clk
        refresh_html_log
    else
        clk=clock; L=L+1; logg{L,1}=sprintf('%d/%s/%d %dh:%dm:%ds - skip (02) slice timing correction for subject (%d) of (%d), run (%d) of (%d)',clk(3),month{clk(2)},clk(1),clk(4),clk(5),round(clk(6)),k,length(sub_dir),k2,runs(k)); try fprintf(fileID,'%s\n',logg{L,1}); set(S.tx,'string',logg); catch, end; clear clk
        try figure(status_fig),subplot(N,2,3),imshow(imread('status_images/step_02_green.jpeg')), subplot(N,2,4),imshow(imread('status_images/status_green_skip.jpg')), catch, end
    end
end

%% '03. Match anatomical and functional slices'
if choices_preproc(3)==1
    if ~(exist([sub_dir{k} anat_files{k,1} '_reduced.nii'], 'file')) || (run_allsubs_by_default==1)
        try figure(status_fig),subplot(N,2,5),imshow(imread('status_images/step_03_blue.jpeg')), subplot(N,2,6),imshow(imread('status_images/status_blue.jpg')), catch, end
        clk=clock; L=L+1; logg{L,1}=sprintf('%d/%s/%d %dh:%dm:%ds - begin (03) match anatomical and functional slices for subject (%d) of (%d), run (%d) of (%d)',clk(3),month{clk(2)},clk(1),clk(4),clk(5),round(clk(6)),k,length(sub_dir),k2,runs(k)); try fprintf(fileID,'%s\n',logg{L,1}); set(S.tx,'string',logg); catch, end; clear clk
        try waitbar((k/length(sub_dir)),wbar2,sprintf('03. Match anatomical and functional slices: Subject (%d) of (%d)\n',k,length(sub_dir))); catch, end
        fprintf('03. Match anatomical and functional slices: Subject (%d) of (%d)\n',k,length(sub_dir))
        time_03_matchAnatFuncSlices = tic;
        [notunix, noAFNI] = scfMRItb_03_matchAnatFuncSlices(sub_dir{k},func_files{k,1},anat_files{k,1});
        time_taken_preproc.step03_matchAnatFuncSlices(k,k2) = toc(time_03_matchAnatFuncSlices); clear time_03_matchAnatFuncSlices
        if notunix~=0
            tmp=clock; clk = sprintf('%.4d-%.2d-%.2d_%.2d-%.2d',tmp(1),tmp(2),tmp(3),tmp(4),tmp(5));
            save([home_dir sprintf('workspace_error_%s',clk)])
            msgbox(sprintf('The toolbox cannot be used without Unix support needed to run AFNI commands.\n\nYour choices so far have been saved to\n%s\n\nTerminating execution...',sprintf('workspace_error_%s.mat',clk)),'Abort','error'); clear clk tmp
            return
        end
        if noAFNI~=0
            tmp=clock; clk = sprintf('%.4d-%.2d-%.2d_%.2d-%.2d',tmp(1),tmp(2),tmp(3),tmp(4),tmp(5));
            save([home_dir sprintf('workspace_error_%s',clk)])
            msgbox(sprintf('AFNI is not in Matlab path (needed); cannot continue execution...\n- If you have not yet installed AFNI on your system, now is a good time.\n- If you have AFNI yet got this message, refer to line-9 (afni_directory = ...) in the script main_scfMRItb.m\n\nYour choices so far have been saved to\n%s\n\nTerminating execution...',sprintf('workspace_error_%s.mat',clk)),'Abort','error'); clear clk tmp
            return
        end
        try figure(status_fig),subplot(N,2,5),imshow(imread('status_images/step_03_green.jpeg')), subplot(N,2,6),imshow(imread('status_images/status_green.jpg')), catch, end
        clk=clock; L=L+1; logg{L,1}=sprintf('%d/%s/%d %dh:%dm:%ds - end (03) match anatomical and functional slices for subject (%d) of (%d), run (%d) of (%d)',clk(3),month{clk(2)},clk(1),clk(4),clk(5),round(clk(6)),k,length(sub_dir),k2,runs(k)); try fprintf(fileID,'%s\n',logg{L,1}); set(S.tx,'string',logg); catch, end; clear clk
        refresh_html_log
    else
        clk=clock; L=L+1; logg{L,1}=sprintf('%d/%s/%d %dh:%dm:%ds - skip (03) match anatomical and functional slices for subject (%d) of (%d), run (%d) of (%d)',clk(3),month{clk(2)},clk(1),clk(4),clk(5),round(clk(6)),k,length(sub_dir),k2,runs(k)); try fprintf(fileID,'%s\n',logg{L,1}); set(S.tx,'string',logg); catch, end; clear clk
        try figure(status_fig),subplot(N,2,5),imshow(imread('status_images/step_03_green.jpeg')), subplot(N,2,6),imshow(imread('status_images/status_green_skip.jpg')), catch, end
    end
end

%% '04. Split data into slices'
if choices_preproc(4)==1
    scfMRItb_04_resplitData(sub_dir{k}, func_files{k,k2}, '', slices) % if slice-wise data doesn't exist but combined 4D data exists, then split the data to generate the needed slice-wise data
    scfMRItb_04_resplitData(sub_dir{k}, func_files{k,k2}, '_base', slices) % if slice-wise data doesn't exist but combined 4D data exists, then split the data to generate the needed slice-wise data
    if ~(exist([sub_dir{k} func_files{k,k2} '_slice' num2str(slices) '_base.nii'], 'file')) || (run_allsubs_by_default==1)
        try figure(status_fig),subplot(N,2,7),imshow(imread('status_images/step_04_blue.jpeg')), subplot(N,2,8),imshow(imread('status_images/status_blue.jpg')), catch, end
        clk=clock; L=L+1; logg{L,1}=sprintf('%d/%s/%d %dh:%dm:%ds - begin (04) split data into slices for subject (%d) of (%d), run (%d) of (%d)',clk(3),month{clk(2)},clk(1),clk(4),clk(5),round(clk(6)),k,length(sub_dir),k2,runs(k)); try fprintf(fileID,'%s\n',logg{L,1}); set(S.tx,'string',logg); catch, end; clear clk
        try waitbar(((runs(k)*(k-1)+k2)/(length(sub_dir)*runs(k))),wbar2,sprintf('04. Split data into slices: Subject (%d) of (%d), Run (%d) of (%d)',k,length(sub_dir),k2,runs(k))); catch, end
        fprintf('04. Split data into slices: Subject (%d) of (%d), Run (%d) of (%d)\n',k,length(sub_dir),k2,runs(k))
        time_04_splitData = tic;
        % Splitting data need not be performed because it will be done within other subsequent functions as and when needed
        % [notunix, noAFNI] = scfMRItb_04_splitData(sub_dir{k},func_files{k,k2},anat_files{k,1},wbar3);
        time_taken_preproc.step04_splitData(k,k2) = toc(time_04_splitData); clear time_04_splitData
        try figure(status_fig),subplot(N,2,7),imshow(imread('status_images/step_04_green.jpeg')), subplot(N,2,8),imshow(imread('status_images/status_green.jpg')), catch, end
        clk=clock; L=L+1; logg{L,1}=sprintf('%d/%s/%d %dh:%dm:%ds - end (04) split data into slices for subject (%d) of (%d), run (%d) of (%d)',clk(3),month{clk(2)},clk(1),clk(4),clk(5),round(clk(6)),k,length(sub_dir),k2,runs(k)); try fprintf(fileID,'%s\n',logg{L,1}); set(S.tx,'string',logg); catch, end; clear clk
        refresh_html_log
    else
        clk=clock; L=L+1; logg{L,1}=sprintf('%d/%s/%d %dh:%dm:%ds - skip (04) split data into slices for subject (%d) of (%d), run (%d) of (%d)',clk(3),month{clk(2)},clk(1),clk(4),clk(5),round(clk(6)),k,length(sub_dir),k2,runs(k)); try fprintf(fileID,'%s\n',logg{L,1}); set(S.tx,'string',logg); catch, end; clear clk
        try figure(status_fig),subplot(N,2,7),imshow(imread('status_images/step_04_green.jpeg')), subplot(N,2,8),imshow(imread('status_images/status_green_skip.jpg')), catch, end
    end
end

%% '05. Define "not cord" mask'
if choices_preproc(5)==1
    if ~(exist([sub_dir{k} func_files{k,k2} '_mask_NS.mat'], 'file')) || (run_allsubs_by_default==1)
        if (save_videos==0)||(save_videos==1), save_videos2 = save_videos; elseif (save_videos==2)&&(sum([choices_preproc(6);choices_preproc(8:9);choices_preproc(11:14)])==0), save_videos2 = 1; else, save_videos2 = 0; end
        try figure(status_fig),subplot(N,2,9),imshow(imread('status_images/step_05_blue.jpeg')), subplot(N,2,10),imshow(imread('status_images/status_blue_UI.jpg')), catch, end
        clk=clock; L=L+1; logg{L,1}=sprintf('%d/%s/%d %dh:%dm:%ds - begin (05) define "not cord" mask for subject (%d) of (%d), run (%d) of (%d)',clk(3),month{clk(2)},clk(1),clk(4),clk(5),round(clk(6)),k,length(sub_dir),k2,runs(k)); try fprintf(fileID,'%s\n',logg{L,1}); set(S.tx,'string',logg); catch, end; clear clk
        try waitbar(((runs(k)*(k-1)+k2)/(length(sub_dir)*runs(k))),wbar2,sprintf('05. Define "not cord" mask: Subject (%d) of (%d), Run (%d) of (%d)',k,length(sub_dir),k2,runs(k))); catch, end
        fprintf('05. Define "not cord" mask: Subject (%d) of (%d), Run (%d) of (%d)\n',k,length(sub_dir),k2,runs(k))
        if runs(k)==1, Rns=0; else, Rns=k2; end
        time_05_notCordMask = tic;
        scfMRItb_05_notCordMask(sub_dir{k},func_files{k,k2}, wbar3,save_videos2,Rns,exist_SCT);
        time_taken_preproc.step05_notCordMask(k,k2) = toc(time_05_notCordMask); clear time_05_notCordMask Rns save_videos2
        try figure(status_fig),subplot(N,2,9),imshow(imread('status_images/step_05_green.jpeg')), subplot(N,2,10),imshow(imread('status_images/status_green.jpg')), catch, end
        clk=clock; L=L+1; logg{L,1}=sprintf('%d/%s/%d %dh:%dm:%ds - end (05) define "not cord" mask for subject (%d) of (%d), run (%d) of (%d)',clk(3),month{clk(2)},clk(1),clk(4),clk(5),round(clk(6)),k,length(sub_dir),k2,runs(k)); try fprintf(fileID,'%s\n',logg{L,1}); set(S.tx,'string',logg); catch, end; clear clk
        refresh_html_log
    else
        if runs(k)==1, Rns=0; else, Rns=k2; end
        if (save_videos==0)||(save_videos==1), save_videos2 = save_videos; elseif (save_videos==2)&&(sum([choices_preproc(6);choices_preproc(8:9);choices_preproc(11:14)])==0), save_videos2 = 1; else, save_videos2 = 0; end
        try figure(status_fig),subplot(N,2,9),imshow(imread('status_images/step_05_blue.jpeg')), subplot(N,2,10),imshow(imread('status_images/status_blue_UI.jpg')), catch, end
        scfMRItb_05_notCordMask_onlyQC(sub_dir{k},func_files{k,k2}, wbar3,save_videos2,Rns,exist_SCT);
        clk=clock; L=L+1; logg{L,1}=sprintf('%d/%s/%d %dh:%dm:%ds - skip (05) define "not cord" mask for subject (%d) of (%d), run (%d) of (%d)',clk(3),month{clk(2)},clk(1),clk(4),clk(5),round(clk(6)),k,length(sub_dir),k2,runs(k)); try fprintf(fileID,'%s\n',logg{L,1}); set(S.tx,'string',logg); catch, end; clear clk
        try figure(status_fig),subplot(N,2,9),imshow(imread('status_images/step_05_green.jpeg')), subplot(N,2,10),imshow(imread('status_images/status_green_skip.jpg')), catch, end
    end
end

%% '06. Denoise-1 (non-cord regression)'
if choices_preproc(6)==1
    if ~( (exist([sub_dir{k} func_files{k,k2} '_denoised_before_MC80.nii'], 'file')) || (exist([sub_dir{k} func_files{k,k2} '_slice' num2str(slices) '_denoised_before_MC80.nii'], 'file')) || (exist([sub_dir{k} func_files{k,k2} '_denoised_before_MC80.nii.gz'], 'file')) || (exist([sub_dir{k} func_files{k,k2} '_slice' num2str(slices) '_denoised_before_MC80.nii.gz'], 'file')) ) || (run_allsubs_by_default==1)
        if (save_videos==0)||(save_videos==1), save_videos2 = save_videos; elseif (save_videos==2)&&(sum([choices_preproc(8:9);choices_preproc(11:14)])==0), save_videos2 = 1; else, save_videos2 = 0; end
        try figure(status_fig),subplot(N,2,11),imshow(imread('status_images/step_06_blue.jpeg')), subplot(N,2,12),imshow(imread('status_images/status_blue.jpg')), catch, end
        clk=clock; L=L+1; logg{L,1}=sprintf('%d/%s/%d %dh:%dm:%ds - begin (06) denoise-1 (non-cord regression) for subject (%d) of (%d), run (%d) of (%d)',clk(3),month{clk(2)},clk(1),clk(4),clk(5),round(clk(6)),k,length(sub_dir),k2,runs(k)); try fprintf(fileID,'%s\n',logg{L,1}); set(S.tx,'string',logg); catch, end; clear clk
        try waitbar(((runs(k)*(k-1)+k2)/(length(sub_dir)*runs(k))),wbar2,sprintf('06. Denoise-1 (non-cord regression): Subject (%d) of (%d), Run (%d) of (%d)',k,length(sub_dir),k2,runs(k))); catch, end
        fprintf('06. Denoise-1 (non-cord regression): Subject (%d) of (%d), Run (%d) of (%d)\n',k,length(sub_dir),k2,runs(k))
        if runs(k)==1, Rns=0; else, Rns=k2; end
        time_06_denoise1 = tic;
        scfMRItb_06_denoise1(sub_dir{k},func_files{k,k2}, wbar3,save_videos2,Rns,exist_SCT);
        time_taken_preproc.step06_denoise1(k,k2) = toc(time_06_denoise1); clear time_06_denoise1 Rns save_videos2
        try figure(status_fig),subplot(N,2,11),imshow(imread('status_images/step_06_green.jpeg')), subplot(N,2,12),imshow(imread('status_images/status_green.jpg')), catch, end
        clk=clock; L=L+1; logg{L,1}=sprintf('%d/%s/%d %dh:%dm:%ds - end (06) denoise-1 (non-cord regression) for subject (%d) of (%d), run (%d) of (%d)',clk(3),month{clk(2)},clk(1),clk(4),clk(5),round(clk(6)),k,length(sub_dir),k2,runs(k)); try fprintf(fileID,'%s\n',logg{L,1}); set(S.tx,'string',logg); catch, end; clear clk
        refresh_html
    else
        scfMRItb_07_baseimage_if_no_06denoise1(sub_dir{k},func_files{k,k2}, wbar3);
        clk=clock; L=L+1; logg{L,1}=sprintf('%d/%s/%d %dh:%dm:%ds - skip (06) denoise-1 (non-cord regression) for subject (%d) of (%d), run (%d) of (%d)',clk(3),month{clk(2)},clk(1),clk(4),clk(5),round(clk(6)),k,length(sub_dir),k2,runs(k)); try fprintf(fileID,'%s\n',logg{L,1}); set(S.tx,'string',logg); catch, end; clear clk
        try figure(status_fig),subplot(N,2,11),imshow(imread('status_images/step_06_green.jpeg')), subplot(N,2,12),imshow(imread('status_images/status_green_skip.jpg')), catch, end
    end
end

%% '07. Compute Gaussian mask'
if choices_preproc(7)==1
    if ~( (exist([sub_dir{k} func_files{k,k2} '_Gaussian_mask.nii'], 'file')) || (exist([sub_dir{k} func_files{k,k2} '_slice' num2str(slices) '_Gaussian_mask.nii'], 'file')) || (exist([sub_dir{k} func_files{k,k2} '_Gaussian_mask.nii.gz'], 'file')) || (exist([sub_dir{k} func_files{k,k2} '_slice' num2str(slices) '_Gaussian_mask.nii.gz'], 'file')) ) || (run_allsubs_by_default==1)
        try figure(status_fig),subplot(N,2,13),imshow(imread('status_images/step_07_blue.jpeg')), subplot(N,2,14),
        if automatic_GaussianMask==0, imshow(imread('status_images/status_blue_UI.jpg')), else, imshow(imread('status_images/status_blue.jpg')), end, catch, end
        clk=clock; L=L+1; logg{L,1}=sprintf('%d/%s/%d %dh:%dm:%ds - begin (07) compute Gaussian mask for subject (%d) of (%d), run (%d) of (%d)',clk(3),month{clk(2)},clk(1),clk(4),clk(5),round(clk(6)),k,length(sub_dir),k2,runs(k)); try fprintf(fileID,'%s\n',logg{L,1}); set(S.tx,'string',logg); catch, end; clear clk
        try waitbar(((runs(k)*(k-1)+k2)/(length(sub_dir)*runs(k))),wbar2,sprintf('07. Compute Gaussian mask: Subject (%d) of (%d), Run (%d) of (%d)',k,length(sub_dir),k2,runs(k))); catch, end
        fprintf('07. Compute Gaussian mask: Subject (%d) of (%d), Run (%d) of (%d)\n',k,length(sub_dir),k2,runs(k))
        if runs(k)==1, Rns=0; else, Rns=k2; end
        time_07_gaussianMask = tic;
        if choices_preproc(6)==0
            scfMRItb_07_baseimage_if_no_06denoise1(sub_dir{k},func_files{k,k2}, wbar3);
        end
        scfMRItb_07_gaussianMask(sub_dir{k},func_files{k,k2},automatic_GaussianMask, wbar3,Rns);
        time_taken_preproc.step07_gaussianMask(k,k2) = toc(time_07_gaussianMask); clear time_07_gaussianMask Rns
        try figure(status_fig),subplot(N,2,13),imshow(imread('status_images/step_07_green.jpeg')), subplot(N,2,14),imshow(imread('status_images/status_green.jpg')), catch, end
        clk=clock; L=L+1; logg{L,1}=sprintf('%d/%s/%d %dh:%dm:%ds - end (07) compute Gaussian mask for subject (%d) of (%d), run (%d) of (%d)',clk(3),month{clk(2)},clk(1),clk(4),clk(5),round(clk(6)),k,length(sub_dir),k2,runs(k)); try fprintf(fileID,'%s\n',logg{L,1}); set(S.tx,'string',logg); catch, end; clear clk
        refresh_html_log
    else
        clk=clock; L=L+1; logg{L,1}=sprintf('%d/%s/%d %dh:%dm:%ds - skip (07) compute Gaussian mask for subject (%d) of (%d), run (%d) of (%d)',clk(3),month{clk(2)},clk(1),clk(4),clk(5),round(clk(6)),k,length(sub_dir),k2,runs(k)); try fprintf(fileID,'%s\n',logg{L,1}); set(S.tx,'string',logg); catch, end; clear clk
        try figure(status_fig),subplot(N,2,13),imshow(imread('status_images/step_07_green.jpeg')), subplot(N,2,14),imshow(imread('status_images/status_green_skip.jpg')), catch, end
    end
end

%% '08. Motion correction'
if choices_preproc(8)==1
    if ~( (exist([sub_dir{k} func_files{k,k2} '_slice' num2str(slices) '_MC.nii'], 'file')) || (exist([sub_dir{k} func_files{k,k2} '_MC.nii'], 'file')) || (exist([sub_dir{k} func_files{k,k2} '_slice' num2str(slices) '_MC.nii.gz'], 'file')) || (exist([sub_dir{k} func_files{k,k2} '_MC.nii.gz'], 'file')) ) || (run_allsubs_by_default==1)
        if (save_videos==0)||(save_videos==1), save_videos2 = save_videos; elseif (save_videos==2)&&(sum([choices_preproc(9);choices_preproc(11:14)])==0), save_videos2 = 1; else, save_videos2 = 0; end
        try figure(status_fig),subplot(N,2,15),imshow(imread('status_images/step_08_blue.jpeg')), subplot(N,2,16),imshow(imread('status_images/status_blue.jpg')), catch, end
        clk=clock; L=L+1; logg{L,1}=sprintf('%d/%s/%d %dh:%dm:%ds - begin (08) motion correction for subject (%d) of (%d), run (%d) of (%d)',clk(3),month{clk(2)},clk(1),clk(4),clk(5),round(clk(6)),k,length(sub_dir),k2,runs(k)); try fprintf(fileID,'%s\n',logg{L,1}); set(S.tx,'string',logg); catch, end; clear clk
        try waitbar(((runs(k)*(k-1)+k2)/(length(sub_dir)*runs(k))),wbar2,sprintf('08. Motion correction: Subject (%d) of (%d), Run (%d) of (%d)',k,length(sub_dir),k2,runs(k))); catch, end
        fprintf('08. Motion correction: Subject (%d) of (%d), Run (%d) of (%d)\n',k,length(sub_dir),k2,runs(k))
        if runs(k)==1, Rns=0; else, Rns=k2; end
        time_08_motionCorrection = tic;
        scfMRItb_08_motionCorrection(sub_dir{k},func_files{k,k2}, motion_param_smoothing,wbar3,save_videos2,Rns);
        time_taken_preproc.step08_motionCorrection(k,k2) = toc(time_08_motionCorrection); clear time_08_motionCorrection Rns save_videos2
        try figure(status_fig),subplot(N,2,15),imshow(imread('status_images/step_08_green.jpeg')), subplot(N,2,16),imshow(imread('status_images/status_green.jpg')), catch, end
        clk=clock; L=L+1; logg{L,1}=sprintf('%d/%s/%d %dh:%dm:%ds - end (08) motion correction for subject (%d) of (%d), run (%d) of (%d)',clk(3),month{clk(2)},clk(1),clk(4),clk(5),round(clk(6)),k,length(sub_dir),k2,runs(k)); try fprintf(fileID,'%s\n',logg{L,1}); set(S.tx,'string',logg); catch, end; clear clk
        refresh_html_log
    else
        clk=clock; L=L+1; logg{L,1}=sprintf('%d/%s/%d %dh:%dm:%ds - skip (08) motion correction for subject (%d) of (%d), run (%d) of (%d)',clk(3),month{clk(2)},clk(1),clk(4),clk(5),round(clk(6)),k,length(sub_dir),k2,runs(k)); try fprintf(fileID,'%s\n',logg{L,1}); set(S.tx,'string',logg); catch, end; clear clk
        try figure(status_fig),subplot(N,2,15),imshow(imread('status_images/step_08_green.jpeg')), subplot(N,2,16),imshow(imread('status_images/status_green_skip.jpg')), catch, end
    end
end

%% '09. Denoise-2 (RETROICOR/3dDespike)'
if choices_preproc(9)==1
    if ~( (exist([sub_dir{k} func_files{k,k2} '_MC_ricor.nii'], 'file')) || (exist([sub_dir{k} func_files{k,k2} '_slice' num2str(slices) '_MC_ricor.nii'], 'file')) || (exist([sub_dir{k} func_files{k,k2} '_MC_ricor.nii.gz'], 'file')) || (exist([sub_dir{k} func_files{k,k2} '_slice' num2str(slices) '_MC_ricor.nii.gz'], 'file')) ) || (run_allsubs_by_default==1)
        if (save_videos==0)||(save_videos==1), save_videos2 = save_videos; elseif (save_videos==2)&&(sum(choices_preproc(11:14))==0), save_videos2 = 1; else, save_videos2 = 0; end
        try figure(status_fig),subplot(N,2,17),imshow(imread('status_images/step_09_blue.jpeg')), subplot(N,2,18),imshow(imread('status_images/status_blue.jpg')), catch, end
        clk=clock; L=L+1; logg{L,1}=sprintf('%d/%s/%d %dh:%dm:%ds - begin (09) Denoise-2 (RETROICOR/3dDespike) for subject (%d) of (%d), run (%d) of (%d)',clk(3),month{clk(2)},clk(1),clk(4),clk(5),round(clk(6)),k,length(sub_dir),k2,runs(k)); try fprintf(fileID,'%s\n',logg{L,1}); set(S.tx,'string',logg); catch, end; clear clk
        try waitbar(((runs(k)*(k-1)+k2)/(length(sub_dir)*runs(k))),wbar2,sprintf('09. Denoise-2 (RETROICOR/3dDespike): Subject (%d) of (%d), Run (%d) of (%d)',k,length(sub_dir),k2,runs(k))); catch, end
        fprintf('09. Denoise-2 (RETROICOR/3dDespike): Subject (%d) of (%d), Run (%d) of (%d)\n',k,length(sub_dir),k2,runs(k))
        if runs(k)==1, Rns=0; else, Rns=k2; end
        time_09_retroicor = tic;
        noRETROICOR = scfMRItb_09_retroicor(sub_dir{k},func_files{k,k2},scanner, wbar3,save_videos2,Rns);
        time_taken_preproc.step09_retroicor(k,k2) = toc(time_09_retroicor); clear time_09_retroicor Rns save_videos2
        if noRETROICOR==1
            clk=clock; L=L+1; logg{L,1}=sprintf('%d/%s/%d %dh:%dm:%ds - .log file (physiological data) unavailable or in incorrect format, and hence RETROICOR was not performed for subject (%d) of (%d), run (%d) of (%d). 3dDespike was performed instead.',clk(3),month{clk(2)},clk(1),clk(4),clk(5),round(clk(6)),k,length(sub_dir),k2,runs(k)); try fprintf(fileID,'%s\n',logg{L,1}); set(S.tx,'string',logg); catch, end; clear clk
            disp(sprintf('Denoise-2: RETROICOR was not performed for subject (%d) run (%d), because the .log file with physiological data was not found or was in incorrect format. 3dDespike was performed instead.',k,k2)) %#ok<DSPS>
        try figure(status_fig),subplot(N,2,17),imshow(imread('status_images/step_09_green_2.jpeg')), subplot(N,2,18),imshow(imread('status_images/status_green.jpg')), catch, end
        clk=clock; L=L+1; logg{L,1}=sprintf('%d/%s/%d %dh:%dm:%ds - end (09) Denoise-2 (3dDespike) for subject (%d) of (%d), run (%d) of (%d)',clk(3),month{clk(2)},clk(1),clk(4),clk(5),round(clk(6)),k,length(sub_dir),k2,runs(k)); try fprintf(fileID,'%s\n',logg{L,1}); set(S.tx,'string',logg); catch, end; clear clk
        else
        try figure(status_fig),subplot(N,2,17),imshow(imread('status_images/step_09_green.jpeg')), subplot(N,2,18),imshow(imread('status_images/status_green.jpg')), catch, end
        clk=clock; L=L+1; logg{L,1}=sprintf('%d/%s/%d %dh:%dm:%ds - end (09) Denoise-2 (RETROICOR/3dDespike) for subject (%d) of (%d), run (%d) of (%d)',clk(3),month{clk(2)},clk(1),clk(4),clk(5),round(clk(6)),k,length(sub_dir),k2,runs(k)); try fprintf(fileID,'%s\n',logg{L,1}); set(S.tx,'string',logg); catch, end; clear clk
        end
        refresh_html_log
    else
        clk=clock; L=L+1; logg{L,1}=sprintf('%d/%s/%d %dh:%dm:%ds - skip (09) Denoise-2 (RETROICOR/3dDespike) for subject (%d) of (%d), run (%d) of (%d)',clk(3),month{clk(2)},clk(1),clk(4),clk(5),round(clk(6)),k,length(sub_dir),k2,runs(k)); try fprintf(fileID,'%s\n',logg{L,1}); set(S.tx,'string',logg); catch, end; clear clk
        try figure(status_fig),subplot(N,2,17),imshow(imread('status_images/step_09_green.jpeg')), subplot(N,2,18),imshow(imread('status_images/status_green_skip.jpg')), catch, end
    end
end

%% '10. Define GM, WM and CSF masks'
if choices_preproc(10)==1
    try figure(status_fig),subplot(N,2,19),imshow(imread('status_images/step_10_blue.jpeg')), subplot(N,2,20),imshow(imread('status_images/status_blue_UI.jpg')), catch, end
    skip=0;
    if k2==1
        if ~( exist([sub_dir{k} anat_files{k,1} '_reduced' '_mask_GM.nii'], 'file') || exist([sub_dir{k} anat_files{k,1} '_reduced' '_mask_GM.nii.gz'], 'file') ) || (run_allsubs_by_default==1)
            skip=skip+1;
            clk=clock; L=L+1; logg{L,1}=sprintf('%d/%s/%d %dh:%dm:%ds - begin (10) define GM, WM and CSF masks for subject (%d) of (%d), run (1) of (%d)',clk(3),month{clk(2)},clk(1),clk(4),clk(5),round(clk(6)),k,length(sub_dir),runs(k)); try fprintf(fileID,'%s\n',logg{L,1}); set(S.tx,'string',logg); catch, end; clear clk
            try waitbar((k*1/length(sub_dir)*runs(k)),wbar2,sprintf('10. Define GM, WM and CSF masks: Subject (%d) of (%d), Run (1) of (%d)',k,length(sub_dir),runs(k))); catch, end
            fprintf('10. Define GM, WM and CSF masks: Subject (%d) of (%d), Run (1) of (%d)\n',k,length(sub_dir),runs(k))
            time_10_GMWMCSFmasks = tic;
            time_taken_preproc.step10_GMWMCSFmasks_detailed{k,1} = scfMRItb_10_GMWMCSFmasks(sub_dir{k},func_files{k,1},anat_files{k,1}, wbar3,exist_SCT,mask_GM_flag,choices_preproc,C_spine, SCT_vert_auto,SCT_manualreg_labels);
            time_taken_preproc.step10_GMWMCSFmasks(k,1) = toc(time_10_GMWMCSFmasks); clear time_10_GMWMCSFmasks
            clk=clock; L=L+1; logg{L,1}=sprintf('%d/%s/%d %dh:%dm:%ds - end (10) define GM, WM and CSF masks for subject (%d) of (%d), run (1) of (%d)',clk(3),month{clk(2)},clk(1),clk(4),clk(5),round(clk(6)),k,length(sub_dir),runs(k)); try fprintf(fileID,'%s\n',logg{L,1}); set(S.tx,'string',logg); catch, end; clear clk
        else
            clk=clock; L=L+1; logg{L,1}=sprintf('%d/%s/%d %dh:%dm:%ds - skip (10) define GM, WM and CSF masks for subject (%d) of (%d), run (1) of (%d)',clk(3),month{clk(2)},clk(1),clk(4),clk(5),round(clk(6)),k,length(sub_dir),runs(k)); try fprintf(fileID,'%s\n',logg{L,1}); set(S.tx,'string',logg); catch, end; clear clk
        end
    else
        if ~( (exist([sub_dir{k} func_files{k,k2} '_base_masked.nii'], 'file')) || (exist([sub_dir{k} func_files{k,k2} '_slice' num2str(slices) '_base_masked.nii'], 'file')) || (exist([sub_dir{k} func_files{k,k2} '_base_masked.nii.gz'], 'file')) || (exist([sub_dir{k} func_files{k,k2} '_slice' num2str(slices) '_base_masked.nii.gz'], 'file')) ) || (run_allsubs_by_default==1)
            skip=skip+1;
            clk=clock; L=L+1; logg{L,1}=sprintf('%d/%s/%d %dh:%dm:%ds - begin (10) define GM, WM and CSF masks for subject (%d) of (%d), run (%d) of (%d)',clk(3),month{clk(2)},clk(1),clk(4),clk(5),round(clk(6)),k,length(sub_dir),k2,runs(k)); try fprintf(fileID,'%s\n',logg{L,1}); set(S.tx,'string',logg); catch, end; clear clk
            try waitbar(((runs(k)*(k-1)+k2)/(length(sub_dir)*runs(k))),wbar2,sprintf('10. Define GM, WM and CSF masks: Subject (%d) of (%d), Run (%d) of (%d)',k,length(sub_dir),k2,runs(k))); catch, end
            fprintf('10. Define GM, WM and CSF masks: Subject (%d) of (%d), Run (%d) of (%d)\n',k,length(sub_dir),k2,runs(k))
            time_10_GMWMCSFmasks = tic;
            scfMRItb_10_GMWMCSFmasks_run2(sub_dir{k},func_files{k,k2},wbar3);
            time_taken_preproc.step10_GMWMCSFmasks(k,k2) = toc(time_10_GMWMCSFmasks); clear time_10_GMWMCSFmasks
            clk=clock; L=L+1; logg{L,1}=sprintf('%d/%s/%d %dh:%dm:%ds - end (10) define GM, WM and CSF masks for subject (%d) of (%d), run (%d) of (%d)',clk(3),month{clk(2)},clk(1),clk(4),clk(5),round(clk(6)),k,length(sub_dir),k2,runs(k)); try fprintf(fileID,'%s\n',logg{L,1}); set(S.tx,'string',logg); catch, end; clear clk
        else
            clk=clock; L=L+1; logg{L,1}=sprintf('%d/%s/%d %dh:%dm:%ds - skip (10) define GM, WM and CSF masks for subject (%d) of (%d), run (%d) of (%d)',clk(3),month{clk(2)},clk(1),clk(4),clk(5),round(clk(6)),k,length(sub_dir),k2,runs(k)); try fprintf(fileID,'%s\n',logg{L,1}); set(S.tx,'string',logg); catch, end; clear clk
        end
    end
    if skip==0
        try figure(status_fig),subplot(N,2,19),imshow(imread('status_images/step_10_green.jpeg')), subplot(N,2,20),imshow(imread('status_images/status_green_skip.jpg')), catch, end
    else
        try figure(status_fig),subplot(N,2,19),imshow(imread('status_images/step_10_green.jpeg')), subplot(N,2,20),imshow(imread('status_images/status_green.jpg')), catch, end
    end
    clear skip
    refresh_html_log
end

%% '11. Co-registration (functional to anat.)'
if choices_preproc(11)==1
    if ~( (exist([sub_dir{k} func_files{k,k2} '_warped.nii'], 'file')) || (exist([sub_dir{k} func_files{k,k2} '_slice' num2str(slices) '_warped.nii'], 'file')) || (exist([sub_dir{k} func_files{k,k2} '_warped.nii.gz'], 'file')) || (exist([sub_dir{k} func_files{k,k2} '_slice' num2str(slices) '_warped.nii.gz'], 'file')) ) || (run_allsubs_by_default==1)
        if (save_videos==0)||(save_videos==1), save_videos2 = save_videos; elseif (save_videos==2)&&(sum(choices_preproc(12:14))==0), save_videos2 = 1; else, save_videos2 = 0; end
        try figure(status_fig),subplot(N,2,21),imshow(imread('status_images/step_11_blue.jpeg')), subplot(N,2,22),imshow(imread('status_images/status_blue.jpg')), catch, end
        clk=clock; L=L+1; logg{L,1}=sprintf('%d/%s/%d %dh:%dm:%ds - begin (11) co-registration (functional to anat.) for subject (%d) of (%d), run (%d) of (%d)',clk(3),month{clk(2)},clk(1),clk(4),clk(5),round(clk(6)),k,length(sub_dir),k2,runs(k)); try fprintf(fileID,'%s\n',logg{L,1}); set(S.tx,'string',logg); catch, end; clear clk
        try waitbar(((runs(k)*(k-1)+k2)/(length(sub_dir)*runs(k))),wbar2,sprintf('11. Co-registration (functional to anat.): Subject (%d) of (%d), Run (%d) of (%d)',k,length(sub_dir),k2,runs(k))); catch, end
        fprintf('11. Co-registration (functional to anat.): Subject (%d) of (%d), Run (%d) of (%d)\n',k,length(sub_dir),k2,runs(k))
        if runs(k)==1, Rns=0; else, Rns=k2; end
        time_11_FuncAnatRegistration = tic;
        scfMRItb_11_FuncAnatRegistration(sub_dir{k},func_files{k,k2},anat_files{k,1},k2, wbar3,save_videos2,Rns);
        time_taken_preproc.step11_FuncAnatRegistration(k,k2) = toc(time_11_FuncAnatRegistration); clear time_11_FuncAnatRegistration Rns save_videos2
        try figure(status_fig),subplot(N,2,21),imshow(imread('status_images/step_11_green.jpeg')), subplot(N,2,22),imshow(imread('status_images/status_green.jpg')), catch, end
        clk=clock; L=L+1; logg{L,1}=sprintf('%d/%s/%d %dh:%dm:%ds - end (11) co-registration (functional to anat.) for subject (%d) of (%d), run (%d) of (%d)',clk(3),month{clk(2)},clk(1),clk(4),clk(5),round(clk(6)),k,length(sub_dir),k2,runs(k)); try fprintf(fileID,'%s\n',logg{L,1}); set(S.tx,'string',logg); catch, end; clear clk
        refresh_html
    else
        clk=clock; L=L+1; logg{L,1}=sprintf('%d/%s/%d %dh:%dm:%ds - skip (11) co-registration (functional to anat.) for subject (%d) of (%d), run (%d) of (%d)',clk(3),month{clk(2)},clk(1),clk(4),clk(5),round(clk(6)),k,length(sub_dir),k2,runs(k)); try fprintf(fileID,'%s\n',logg{L,1}); set(S.tx,'string',logg); catch, end; clear clk
        try figure(status_fig),subplot(N,2,21),imshow(imread('status_images/step_11_green.jpeg')), subplot(N,2,22),imshow(imread('status_images/status_green_skip.jpg')), catch, end
    end
end

%% '12. Denoise-3 (CSF regression)'
if choices_preproc(12)==1
    try load([sub_dir{k} sprintf('workspace_variables_run%d.mat',k2)], 'cutoff_str'); catch, cutoff_str = '50'; end
    if ~( (exist([sub_dir{k} func_files{k,k2} '_denoised' cutoff_str '.nii'], 'file')) || (exist([sub_dir{k} func_files{k,k2} '_slice' num2str(slices) '_denoised' cutoff_str '.nii'], 'file')) || (exist([sub_dir{k} func_files{k,k2} '_denoised' cutoff_str '.nii.gz'], 'file')) || (exist([sub_dir{k} func_files{k,k2} '_slice' num2str(slices) '_denoised' cutoff_str '.nii.gz'], 'file')) ) || (run_allsubs_by_default==1)
        if (save_videos==0)||(save_videos==1), save_videos2 = save_videos; elseif (save_videos==2)&&(sum(choices_preproc(13:14))==0), save_videos2 = 2; else, save_videos2 = 0; end
        try figure(status_fig),subplot(N,2,23),imshow(imread('status_images/step_12_blue.jpeg')), subplot(N,2,24),imshow(imread('status_images/status_blue.jpg')), catch, end
        clk=clock; L=L+1; logg{L,1}=sprintf('%d/%s/%d %dh:%dm:%ds - begin (12) denoise-3 (CSF regression) for subject (%d) of (%d), run (%d) of (%d)',clk(3),month{clk(2)},clk(1),clk(4),clk(5),round(clk(6)),k,length(sub_dir),k2,runs(k)); try fprintf(fileID,'%s\n',logg{L,1}); set(S.tx,'string',logg); catch, end; clear clk
        try waitbar(((runs(k)*(k-1)+k2)/(length(sub_dir)*runs(k))),wbar2,sprintf('12. Denoise-3 (CSF regression): Subject (%d) of (%d), Run (%d) of (%d)',k,length(sub_dir),k2,runs(k))); catch, end
        fprintf('12. Denoise-3 (CSF regression): Subject (%d) of (%d), Run (%d) of (%d)\n',k,length(sub_dir),k2,runs(k))
        if runs(k)==1, Rns=0; else, Rns=k2; end
        time_12_CSFregress = tic;
        scfMRItb_12_CSFregress(sub_dir{k},func_files{k,k2},anat_files{k,1}, wbar3,save_videos2,Rns);
        time_taken_preproc.step12_CSFregress(k,k2) = toc(time_12_CSFregress); clear time_12_CSFregress Rns save_videos2
        try figure(status_fig),subplot(N,2,23),imshow(imread('status_images/step_12_green.jpeg')), subplot(N,2,24),imshow(imread('status_images/status_green.jpg')), catch, end
        clk=clock; L=L+1; logg{L,1}=sprintf('%d/%s/%d %dh:%dm:%ds - end (12) denoise-3 (CSF regression) for subject (%d) of (%d), run (%d) of (%d)',clk(3),month{clk(2)},clk(1),clk(4),clk(5),round(clk(6)),k,length(sub_dir),k2,runs(k)); try fprintf(fileID,'%s\n',logg{L,1}); set(S.tx,'string',logg); catch, end; clear clk
        refresh_html
    else
        clk=clock; L=L+1; logg{L,1}=sprintf('%d/%s/%d %dh:%dm:%ds - skip (12) denoise-3 (CSF regression) for subject (%d) of (%d), run (%d) of (%d)',clk(3),month{clk(2)},clk(1),clk(4),clk(5),round(clk(6)),k,length(sub_dir),k2,runs(k)); try fprintf(fileID,'%s\n',logg{L,1}); set(S.tx,'string',logg); catch, end; clear clk
        try figure(status_fig),subplot(N,2,23),imshow(imread('status_images/step_12_green.jpeg')), subplot(N,2,24),imshow(imread('status_images/status_green_skip.jpg')), catch, end
    end
end

%% '13. Denoise-4 (WM regression)'
if choices_preproc(13)==1
    try load([sub_dir{k} sprintf('workspace_variables_run%d.mat',k2)], 'cutoff_str'); catch, cutoff_str = '50'; end
    if ~( (exist([sub_dir{k} func_files{k,k2} '_denoised' cutoff_str 'WM.nii'], 'file')) || (exist([sub_dir{k} func_files{k,k2} '_slice' num2str(slices) '_denoised' cutoff_str 'WM.nii'], 'file')) || (exist([sub_dir{k} func_files{k,k2} '_denoised' cutoff_str 'WM.nii.gz'], 'file')) || (exist([sub_dir{k} func_files{k,k2} '_slice' num2str(slices) '_denoised' cutoff_str 'WM.nii.gz'], 'file'))  ||  (exist([sub_dir{k} func_files{k,k2} '_WM.nii'], 'file')) || (exist([sub_dir{k} func_files{k,k2} '_slice' num2str(slices) '_WM.nii'], 'file')) || (exist([sub_dir{k} func_files{k,k2} '_WM.nii.gz'], 'file')) || (exist([sub_dir{k} func_files{k,k2} '_slice' num2str(slices) '_WM.nii.gz'], 'file')) ) || (run_allsubs_by_default==1)
        if (save_videos==0)||(save_videos==1), save_videos2 = save_videos; elseif (save_videos==2)&&(sum(choices_preproc(14:14))==0), save_videos2 = 2; else, save_videos2 = 0; end
        try figure(status_fig),subplot(N,2,25),imshow(imread('status_images/step_13_blue.jpeg')), subplot(N,2,26),imshow(imread('status_images/status_blue.jpg')), catch, end
        clk=clock; L=L+1; logg{L,1}=sprintf('%d/%s/%d %dh:%dm:%ds - begin (13) regress-out white-matter signal from GM for subject (%d) of (%d), run (%d) of (%d)',clk(3),month{clk(2)},clk(1),clk(4),clk(5),round(clk(6)),k,length(sub_dir),k2,runs(k)); try fprintf(fileID,'%s\n',logg{L,1}); set(S.tx,'string',logg); catch, end; clear clk
        try waitbar(((runs(k)*(k-1)+k2)/(length(sub_dir)*runs(k))),wbar2,sprintf('13. Regress-out white-matter signal from GM: Subject (%d) of (%d), Run (%d) of (%d)',k,length(sub_dir),k2,runs(k))); catch, end
        fprintf('13. Regress-out white-matter signal from GM: Subject (%d) of (%d), Run (%d) of (%d)\n',k,length(sub_dir),k2,runs(k))
        if runs(k)==1, Rns=0; else, Rns=k2; end
        time_13_WMregress = tic;
        scfMRItb_13_WMregress(sub_dir{k},func_files{k,k2},anat_files{k,1}, wbar3,save_videos2,Rns);
        time_taken_preproc.step13_WMregress(k,k2) = toc(time_13_WMregress); clear time_13_WMregress Rns save_videos2
        try figure(status_fig),subplot(N,2,25),imshow(imread('status_images/step_13_green.jpeg')), subplot(N,2,26),imshow(imread('status_images/status_green.jpg')), catch, end
        clk=clock; L=L+1; logg{L,1}=sprintf('%d/%s/%d %dh:%dm:%ds - end (13) regress-out white-matter signal from GM for subject (%d) of (%d), run (%d) of (%d)',clk(3),month{clk(2)},clk(1),clk(4),clk(5),round(clk(6)),k,length(sub_dir),k2,runs(k)); try fprintf(fileID,'%s\n',logg{L,1}); set(S.tx,'string',logg); catch, end; clear clk
        refresh_html
    else
        clk=clock; L=L+1; logg{L,1}=sprintf('%d/%s/%d %dh:%dm:%ds - skip (13) regress-out white-matter signal from GM for subject (%d) of (%d), run (%d) of (%d)',clk(3),month{clk(2)},clk(1),clk(4),clk(5),round(clk(6)),k,length(sub_dir),k2,runs(k)); try fprintf(fileID,'%s\n',logg{L,1}); set(S.tx,'string',logg); catch, end; clear clk
        try figure(status_fig),subplot(N,2,25),imshow(imread('status_images/step_13_green.jpeg')), subplot(N,2,26),imshow(imread('status_images/status_green_skip.jpg')), catch, end
    end
end

%% '14. Denoise-5 (additional covariates regression)'
if (choices_preproc(14)==1) && ~( covregress_options(1,1)==3 && covregress_options(2,1)==3 && covregress_options(3,1)==2 )
    try load([sub_dir{k} sprintf('workspace_variables_run%d.mat',k2)], 'cutoff_str'); catch, cutoff_str = '50'; end
    if ~( (exist([sub_dir{k} func_files{k,k2} '_denoised' cutoff_str 'WMcov.nii'], 'file')) || (exist([sub_dir{k} func_files{k,k2} '_slice' num2str(slices) '_denoised' cutoff_str 'WMcov.nii'], 'file')) || (exist([sub_dir{k} func_files{k,k2} '_denoised' cutoff_str 'WMcov.nii.gz'], 'file')) || (exist([sub_dir{k} func_files{k,k2} '_slice' num2str(slices) '_denoised' cutoff_str 'WMcov.nii.gz'], 'file'))  ||  (exist([sub_dir{k} func_files{k,k2} '_denoised' cutoff_str 'cov.nii'], 'file')) || (exist([sub_dir{k} func_files{k,k2} '_slice' num2str(slices) '_denoised' cutoff_str 'cov.nii'], 'file')) || (exist([sub_dir{k} func_files{k,k2} '_denoised' cutoff_str 'cov.nii.gz'], 'file')) || (exist([sub_dir{k} func_files{k,k2} '_slice' num2str(slices) '_denoised' cutoff_str 'cov.nii.gz'], 'file'))  ||  (exist([sub_dir{k} func_files{k,k2} '_WMcov.nii'], 'file')) || (exist([sub_dir{k} func_files{k,k2} '_slice' num2str(slices) '_WMcov.nii'], 'file')) || (exist([sub_dir{k} func_files{k,k2} '_WMcov.nii.gz'], 'file')) || (exist([sub_dir{k} func_files{k,k2} '_slice' num2str(slices) '_WMcov.nii.gz'], 'file'))  ||  (exist([sub_dir{k} func_files{k,k2} '_cov.nii'], 'file')) || (exist([sub_dir{k} func_files{k,k2} '_slice' num2str(slices) '_cov.nii'], 'file')) || (exist([sub_dir{k} func_files{k,k2} '_cov.nii.gz'], 'file')) || (exist([sub_dir{k} func_files{k,k2} '_slice' num2str(slices) '_cov.nii.gz'], 'file')) ) || (run_allsubs_by_default==1)
        save_videos2 = save_videos;
        try figure(status_fig),subplot(N,2,27),imshow(imread('status_images/step_14_blue.jpeg')), subplot(N,2,28),imshow(imread('status_images/status_blue.jpg')), catch, end
        clk=clock; L=L+1; logg{L,1}=sprintf('%d/%s/%d %dh:%dm:%ds - begin (14) Regress-out additional covariates for subject (%d) of (%d), run (%d) of (%d)',clk(3),month{clk(2)},clk(1),clk(4),clk(5),round(clk(6)),k,length(sub_dir),k2,runs(k)); try fprintf(fileID,'%s\n',logg{L,1}); set(S.tx,'string',logg); catch, end; clear clk
        try waitbar(((runs(k)*(k-1)+k2)/(length(sub_dir)*runs(k))),wbar2,sprintf('14. Regress-out additional covariates: Subject (%d) of (%d), Run (%d) of (%d)',k,length(sub_dir),k2,runs(k))); catch, end
        fprintf('14. Regress-out additional covariates: Subject (%d) of (%d), Run (%d) of (%d)\n',k,length(sub_dir),k2,runs(k))
        if runs(k)==1, Rns=0; else, Rns=k2; end
        time_14_COVregress = tic;
        scfMRItb_14_COVregress(sub_dir{k},func_files{k,k2},anat_files{k,1},covregress_options,cov_files{k,k2},slices, wbar3,save_videos2,Rns);
        time_taken_preproc.step14_COVregress(k,k2) = toc(time_14_COVregress); clear time_14_COVregress Rns save_videos2
        try figure(status_fig),subplot(N,2,27),imshow(imread('status_images/step_14_green.jpeg')), subplot(N,2,28),imshow(imread('status_images/status_green.jpg')), catch, end
        clk=clock; L=L+1; logg{L,1}=sprintf('%d/%s/%d %dh:%dm:%ds - end (14) regress-out additional covariates for subject (%d) of (%d), run (%d) of (%d)',clk(3),month{clk(2)},clk(1),clk(4),clk(5),round(clk(6)),k,length(sub_dir),k2,runs(k)); try fprintf(fileID,'%s\n',logg{L,1}); set(S.tx,'string',logg); catch, end; clear clk
        refresh_html
    else
        clk=clock; L=L+1; logg{L,1}=sprintf('%d/%s/%d %dh:%dm:%ds - skip (14) regress-out additional covariates for subject (%d) of (%d), run (%d) of (%d)',clk(3),month{clk(2)},clk(1),clk(4),clk(5),round(clk(6)),k,length(sub_dir),k2,runs(k)); try fprintf(fileID,'%s\n',logg{L,1}); set(S.tx,'string',logg); catch, end; clear clk
        try figure(status_fig),subplot(N,2,27),imshow(imread('status_images/step_14_green.jpeg')), subplot(N,2,28),imshow(imread('status_images/status_green_skip.jpg')), catch, end
    end
end

%% 'Save denoised 3D+time fMRI data'
if sum([choices_preproc(6);choices_preproc(8);choices_preproc(10:14)])~=0
    try load([sub_dir{k} sprintf('workspace_variables_run%d.mat',k2)], 'cutoff_str'); catch, cutoff_str = '50'; end
    if exist([sub_dir{k} func_files{k,k2} '_WM' '.nii'], 'file') || exist([sub_dir{k} func_files{k,k2} '_WM' '.nii.gz'], 'file') % if only WM regress step was performed
        existname = [sub_dir{k} func_files{k,k2} '_WM' '.mat'];
    end
    if exist([sub_dir{k} func_files{k,k2} '_denoised' cutoff_str '.nii'], 'file') || exist([sub_dir{k} func_files{k,k2} '_denoised' cutoff_str '.nii.gz'], 'file') % if only CSF regress step was performed
        existname = [sub_dir{k} func_files{k,k2} '_denoised' cutoff_str '.mat'];
    end
    if exist([sub_dir{k} func_files{k,k2} '_cov' '.nii'], 'file') || exist([sub_dir{k} func_files{k,k2} '_cov' '.nii.gz'], 'file') % if only cov regress step was performed
        existname = [sub_dir{k} func_files{k,k2} '_cov' '.mat'];
    end
    if exist([sub_dir{k} func_files{k,k2} '_WMcov' '.nii'], 'file') || exist([sub_dir{k} func_files{k,k2} '_WMcov' '.nii.gz'], 'file') % if both WM and cov regress steps were performed
        existname = [sub_dir{k} func_files{k,k2} '_WMcov' '.mat'];
    end
    if exist([sub_dir{k} func_files{k,k2} '_denoised' cutoff_str 'WM' '.nii'], 'file') || exist([sub_dir{k} func_files{k,k2} '_denoised' cutoff_str 'WM' '.nii.gz'], 'file') % if both CSF and WM regress steps were performed
        existname = [sub_dir{k} func_files{k,k2} '_denoised' cutoff_str 'WM' '.mat'];
    end
    if exist([sub_dir{k} func_files{k,k2} '_denoised' cutoff_str 'cov' '.nii'], 'file') || exist([sub_dir{k} func_files{k,k2} '_denoised' cutoff_str 'cov' '.nii.gz'], 'file') % if both CSF and cov regress steps were performed
        existname = [sub_dir{k} func_files{k,k2} '_denoised' cutoff_str 'cov' '.mat'];
    end
    if exist([sub_dir{k} func_files{k,k2} '_denoised' cutoff_str 'WMcov' '.nii'], 'file') || exist([sub_dir{k} func_files{k,k2} '_denoised' cutoff_str 'WMcov' '.nii.gz'], 'file') % if both CSF, WM and cov regress steps were performed
        existname = [sub_dir{k} func_files{k,k2} '_denoised' cutoff_str 'WMcov' '.mat'];
    end
    if ~(exist('existname', 'var'))
        existname = [];
    end
    if ~(exist(existname, 'file')) || (run_allsubs_by_default==1)
        clk=clock; L=L+1; logg{L,1}=sprintf('%d/%s/%d %dh:%dm:%ds - begin save denoised 3D+time fMRI data for subject (%d) of (%d), run (%d) of (%d)',clk(3),month{clk(2)},clk(1),clk(4),clk(5),round(clk(6)),k,length(sub_dir),k2,runs(k)); try fprintf(fileID,'%s\n',logg{L,1}); set(S.tx,'string',logg); catch, end; clear clk
        try waitbar(((runs(k)*(k-1)+k2)/(length(sub_dir)*runs(k))),wbar2,sprintf('Save denoised 3D+time fMRI data: Subject (%d) of (%d), Run (%d) of (%d)',k,length(sub_dir),k2,runs(k))); catch, end
        fprintf('Save denoised 3D+time fMRI data: Subject (%d) of (%d), Run (%d) of (%d)\n',k,length(sub_dir),k2,runs(k))
        time_save4Ddata_denoised = tic;
        scfMRItb_14_save4Ddata_denoised(sub_dir{k},func_files{k,k2},anat_files{k,1},wbar3);
        time_taken_preproc.save4Ddata_denoised(k,k2) = toc(time_save4Ddata_denoised); clear time_save4Ddata_denoised
        clk=clock; L=L+1; logg{L,1}=sprintf('%d/%s/%d %dh:%dm:%ds - end save denoised 3D+time fMRI data for subject (%d) of (%d), run (%d) of (%d)',clk(3),month{clk(2)},clk(1),clk(4),clk(5),round(clk(6)),k,length(sub_dir),k2,runs(k)); try fprintf(fileID,'%s\n',logg{L,1}); set(S.tx,'string',logg); catch, end; clear clk
        refresh_html_log
    else
        clk=clock; L=L+1; logg{L,1}=sprintf('%d/%s/%d %dh:%dm:%ds - skip save denoised 3D+time fMRI data for subject (%d) of (%d), run (%d) of (%d)',clk(3),month{clk(2)},clk(1),clk(4),clk(5),round(clk(6)),k,length(sub_dir),k2,runs(k)); try fprintf(fileID,'%s\n',logg{L,1}); set(S.tx,'string',logg); catch, end; clear clk
    end
    clear existname
end

%% '15. Temporal band-pass filtering'
if choices_preproc(15)==1
    try load([sub_dir{k} sprintf('workspace_variables_run%d.mat',k2)], 'cutoff_str'); catch, cutoff_str = '50'; end
    if exist([sub_dir{k} func_files{k,k2} '_WM' '.nii'], 'file') || exist([sub_dir{k} func_files{k,k2} '_WM' '.nii.gz'], 'file') % if only WM regress step was performed
        existname = [sub_dir{k} func_files{k,k2} '_WM' '_HPF01.mat'];
    end
    if exist([sub_dir{k} func_files{k,k2} '_denoised' cutoff_str '.nii'], 'file') || exist([sub_dir{k} func_files{k,k2} '_denoised' cutoff_str '.nii.gz'], 'file') % if only CSF regress step was performed
        existname = [sub_dir{k} func_files{k,k2} '_denoised' cutoff_str '_HPF01.mat'];
    end
    if exist([sub_dir{k} func_files{k,k2} '_cov' '.nii'], 'file') || exist([sub_dir{k} func_files{k,k2} '_cov' '.nii.gz'], 'file') % if only cov regress step was performed
        existname = [sub_dir{k} func_files{k,k2} '_cov' '_HPF01.mat'];
    end
    if exist([sub_dir{k} func_files{k,k2} '_WMcov' '.nii'], 'file') || exist([sub_dir{k} func_files{k,k2} '_WMcov' '.nii.gz'], 'file') % if both WM and cov regress steps were performed
        existname = [sub_dir{k} func_files{k,k2} '_WMcov' '_HPF01.mat'];
    end
    if exist([sub_dir{k} func_files{k,k2} '_denoised' cutoff_str 'WM' '.nii'], 'file') || exist([sub_dir{k} func_files{k,k2} '_denoised' cutoff_str 'WM' '.nii.gz'], 'file') % if both CSF and WM regress steps were performed
        existname = [sub_dir{k} func_files{k,k2} '_denoised' cutoff_str 'WM' '_HPF01.mat'];
    end
    if exist([sub_dir{k} func_files{k,k2} '_denoised' cutoff_str 'cov' '.nii'], 'file') || exist([sub_dir{k} func_files{k,k2} '_denoised' cutoff_str 'cov' '.nii.gz'], 'file') % if both CSF and cov regress steps were performed
        existname = [sub_dir{k} func_files{k,k2} '_denoised' cutoff_str 'cov' '_HPF01.mat'];
    end
    if exist([sub_dir{k} func_files{k,k2} '_denoised' cutoff_str 'WMcov' '.nii'], 'file') || exist([sub_dir{k} func_files{k,k2} '_denoised' cutoff_str 'WMcov' '.nii.gz'], 'file') % if both CSF, WM and cov regress steps were performed
        existname = [sub_dir{k} func_files{k,k2} '_denoised' cutoff_str 'WMcov' '_HPF01.mat'];
    end
    if ~(exist('existname', 'var'))
        existname = [];
    end
    if ~(exist(existname, 'file')) || (run_allsubs_by_default==1)
        try figure(status_fig),subplot(N,2,29),imshow(imread('status_images/step_15_blue.jpeg')), subplot(N,2,30),imshow(imread('status_images/status_blue.jpg')), catch, end
        clk=clock; L=L+1; logg{L,1}=sprintf('%d/%s/%d %dh:%dm:%ds - begin (15) temporal band-pass filtering for subject (%d) of (%d), run (%d) of (%d)',clk(3),month{clk(2)},clk(1),clk(4),clk(5),round(clk(6)),k,length(sub_dir),k2,runs(k)); try fprintf(fileID,'%s\n',logg{L,1}); set(S.tx,'string',logg); catch, end; clear clk
        try waitbar(((runs(k)*(k-1)+k2)/(length(sub_dir)*runs(k))),wbar2,sprintf('15. Temporal band-pass filtering: Subject (%d) of (%d), Run (%d) of (%d)',k,length(sub_dir),k2,runs(k))); catch, end
        fprintf('15. Temporal band-pass filtering: Subject (%d) of (%d), Run (%d) of (%d)\n',k,length(sub_dir),k2,runs(k))
        if runs(k)==1, Rns=0; else, Rns=k2; end
        time_15_filterData = tic;
        scfMRItb_15_filterData(sub_dir{k},func_files{k,k2},anat_files{k,1},bpf_hz,detrend_flag, wbar3,save_videos,Rns);
        time_taken_preproc.step15_filterData(k,k2) = toc(time_15_filterData); clear time_15_filterData Rns
        try figure(status_fig),subplot(N,2,29),imshow(imread('status_images/step_15_green.jpeg')), subplot(N,2,30),imshow(imread('status_images/status_green.jpg')), catch, end
        clk=clock; L=L+1; logg{L,1}=sprintf('%d/%s/%d %dh:%dm:%ds - end (15) temporal band-pass filtering for subject (%d) of (%d), run (%d) of (%d)',clk(3),month{clk(2)},clk(1),clk(4),clk(5),round(clk(6)),k,length(sub_dir),k2,runs(k)); try fprintf(fileID,'%s\n',logg{L,1}); set(S.tx,'string',logg); catch, end; clear clk
        refresh_html_log
    else
        if ~(exist([sub_dir{k} 'workspace_variables_temp.mat'], 'file'))
            base_dir_sub = sub_dir{k}; fname = func_files{k,k2}; cutoff = 0.50; cutoff_str = '50'; f_BPF_max_vec = 0.10; f_BPF_max_str_vec = {'10'};
            scfMRItb_04_unzipFile(sub_dir{k}, func_files{k,k2}, '')
            F = load_untouch_nii([sub_dir{k} func_files{k,k2} '.nii']); TR = F.hdr.dime.pixdim(1,5);
            f_BPF_max = f_BPF_max_vec(1); f_BPF_max_str = f_BPF_max_str_vec{1};
            save([sub_dir{k} 'workspace_variables_temp.mat'], 'base_dir_sub', 'fname', 'cutoff', 'cutoff_str', 'f_BPF_max_vec', 'f_BPF_max_str_vec', 'f_BPF_max', 'f_BPF_max_str', 'TR', 'bpf_hz')
            clear base_dir_sub fname F TR
        end
        clk=clock; L=L+1; logg{L,1}=sprintf('%d/%s/%d %dh:%dm:%ds - skip (15) temporal band-pass filtering for subject (%d) of (%d), run (%d) of (%d)',clk(3),month{clk(2)},clk(1),clk(4),clk(5),round(clk(6)),k,length(sub_dir),k2,runs(k)); try fprintf(fileID,'%s\n',logg{L,1}); set(S.tx,'string',logg); catch, end; clear clk
        try figure(status_fig),subplot(N,2,29),imshow(imread('status_images/step_15_green.jpeg')), subplot(N,2,30),imshow(imread('status_images/status_green_skip.jpg')), catch, end
    end
    clear existname
end

%% '16. Define cord quadrants'
if choices_preproc(16)==1
    try figure(status_fig),subplot(N,2,31),imshow(imread('status_images/step_16_blue.jpeg')), subplot(N,2,32),imshow(imread('status_images/status_blue_UI.jpg')), catch, end
    skip=0;
    if ~(exist([sub_dir{k} 'workspace_variables_temp.mat'], 'file'))
        try
        base_dir_sub = sub_dir{k}; fname = func_files{k,1}; cutoff = 0.50; cutoff_str = '50'; f_BPF_max_vec = 0.10; f_BPF_max_str_vec = {'10'}; if ~exist('bpf_hz','var'), bpf_hz = f_BPF_max_vec; end
        scfMRItb_04_unzipFile(sub_dir{k}, func_files{k,1}, ''); F = load_untouch_nii([sub_dir{k} func_files{k,1} '.nii']); TR = F.hdr.dime.pixdim(1,5);
        if TR>3.65, wz=2; else, wz=3; end; f_BPF_max = f_BPF_max_vec(1,1); f_BPF_max_str = f_BPF_max_str_vec{1,1}; clear wz
        save([sub_dir{k} 'workspace_variables_temp.mat'], 'base_dir_sub', 'fname', 'cutoff', 'cutoff_str', 'f_BPF_max_vec', 'f_BPF_max_str_vec', 'f_BPF_max', 'f_BPF_max_str', 'TR', 'bpf_hz')
        clear base_dir_sub fname F TR
        catch
        end
    end
    if k2==1
        if (run_allsubs_by_default==1)
            skip=skip+1;
            clk=clock; L=L+1; logg{L,1}=sprintf('%d/%s/%d %dh:%dm:%ds - begin (16) define cord quadrants for subject (%d) of (%d)',clk(3),month{clk(2)},clk(1),clk(4),clk(5),round(clk(6)),k,length(sub_dir)); try fprintf(fileID,'%s\n',logg{L,1}); set(S.tx,'string',logg); catch, end; clear clk
            try waitbar((k/length(sub_dir)),wbar2,sprintf('16. Define cord quadrants: Subject (%d) of (%d)',k,length(sub_dir))); catch, end
            fprintf('16. Define cord quadrants: Subject (%d) of (%d)\n',k,length(sub_dir))
            time_16_cord_quadrants = tic;
            scfMRItb_16_cord_quadrants(sub_dir{k},func_files{k,1},anat_files{k,1}, scan7T,C_spine,wbar3);
            time_taken_preproc.step16_cord_quadrants(k,1) = toc(time_16_cord_quadrants); clear time_16_cord_quadrants
            merge_mat_files([sub_dir{k} sprintf('workspace_variables_run%d.mat',1)],[sub_dir{k} sprintf('workspace_variables_run%d.mat',1)],[sub_dir{k} 'workspace_variables_temp.mat'],1)
            clk=clock; L=L+1; logg{L,1}=sprintf('%d/%s/%d %dh:%dm:%ds - end (16) define cord quadrants for subject (%d) of (%d)',clk(3),month{clk(2)},clk(1),clk(4),clk(5),round(clk(6)),k,length(sub_dir)); try fprintf(fileID,'%s\n',logg{L,1}); set(S.tx,'string',logg); catch, end; clear clk
            refresh_html_log
        else
        try
            load([sub_dir{k} 'cord_quadrants.mat'], 'FC_mask_quads')
            clk=clock; L=L+1; logg{L,1}=sprintf('%d/%s/%d %dh:%dm:%ds - skip (16) define cord quadrants for subject (%d) of (%d), run (1) of (%d)',clk(3),month{clk(2)},clk(1),clk(4),clk(5),round(clk(6)),k,length(sub_dir),runs(k)); try fprintf(fileID,'%s\n',logg{L,1}); set(S.tx,'string',logg); catch, end; clear clk
            if ~exist([sub_dir{k} sprintf('workspace_variables_run%d.mat',1)],'file')
                try
                    load([base_dir_sub 'workspace_variables_temp.mat'])
                catch
                    f_BPF_max_vec = 0.10; f_BPF_max_str_vec = {'10'};
                    cutoff = 0.50; cutoff_str = '50';
                    f_BPF_max = f_BPF_max_vec(1,1); f_BPF_max_str = f_BPF_max_str_vec{1,1};
                    if ~exist('TR','var')
                        scfMRItb_04_unzipFile(sub_dir{k}, func_files{k,1}, '')
                        F = load_untouch_nii([sub_dir{k} func_files{k,1} '.nii']); TR = F.hdr.dime.pixdim(1,5); clear F
                    end
                end
                base_dir_sub = sub_dir{k}; fname = func_files{k,1};
                try
                save([base_dir_sub 'workspace_variables_run1.mat'],'base_dir_sub','fname*','cutoff*','mask_*','f_BPF_max*','*FC_mask_*','*spine','TR','scan3T','new_size')
                catch
                save([base_dir_sub 'workspace_variables_run1.mat'],'base_dir_sub','fname*','cutoff*','f_BPF_max*','*FC_mask_*','*spine','TR','scan3T')
                end
            end
        catch
            skip=skip+1;
            clk=clock; L=L+1; logg{L,1}=sprintf('%d/%s/%d %dh:%dm:%ds - begin (16) define cord quadrants for subject (%d) of (%d)',clk(3),month{clk(2)},clk(1),clk(4),clk(5),round(clk(6)),k,length(sub_dir)); try fprintf(fileID,'%s\n',logg{L,1}); set(S.tx,'string',logg); catch, end; clear clk
            try waitbar((k/length(sub_dir)),wbar2,sprintf('16. Define cord quadrants: Subject (%d) of (%d)',k,length(sub_dir))); catch, end
            fprintf('16. Define cord quadrants: Subject (%d) of (%d)\n',k,length(sub_dir))
            time_16_cord_quadrants = tic;
            scfMRItb_16_cord_quadrants(sub_dir{k},func_files{k,1},anat_files{k,1}, scan7T,C_spine,wbar3);
            time_taken_preproc.step16_cord_quadrants(k,1) = toc(time_16_cord_quadrants); clear time_16_cord_quadrants
            merge_mat_files([sub_dir{k} sprintf('workspace_variables_run%d.mat',1)],[sub_dir{k} sprintf('workspace_variables_run%d.mat',1)],[sub_dir{k} 'workspace_variables_temp.mat'],1)
            clk=clock; L=L+1; logg{L,1}=sprintf('%d/%s/%d %dh:%dm:%ds - end (16) define cord quadrants for subject (%d) of (%d)',clk(3),month{clk(2)},clk(1),clk(4),clk(5),round(clk(6)),k,length(sub_dir)); try fprintf(fileID,'%s\n',logg{L,1}); set(S.tx,'string',logg); catch, end; clear clk
            refresh_html_log
        end
        end
        
    else
        try
            load([sub_dir{k} sprintf('workspace_variables_run%d.mat',k2)], 'FC_mask_quads')
        catch
            skip=skip+1;
            copyfile([sub_dir{k} sprintf('workspace_variables_run%d.mat',1)],[sub_dir{k} sprintf('workspace_variables_run%d.mat',k2)],'f')
            clear fname TR
            fname = func_files{k,k2};
            if ~exist('TR','var')
                scfMRItb_04_unzipFile(sub_dir{k}, func_files{k,k2}, '')
                F = load_untouch_nii([sub_dir{k} func_files{k,k2} '.nii']); TR = F.hdr.dime.pixdim(1,5); clear F
            end
            save([sub_dir{k} sprintf('workspace_variables_run%d.mat',k2)], 'fname','TR', '-append')
        end
    end
    try
        merge_mat_files([sub_dir{k} sprintf('workspace_variables_run%d.mat',1)],[sub_dir{k} 'workspace_variables_temp.mat'],[sub_dir{k} sprintf('workspace_variables_run%d.mat',1)],1)
    catch
    end
    if exist(sprintf('%sworkspace_variables_temp.mat',sub_dir{k}),'file') && exist([sub_dir{k} sprintf('workspace_variables_run%d.mat',1)],'file')
        delete(sprintf('%sworkspace_variables_temp.mat',sub_dir{k}))
    end
    if skip==0
        try figure(status_fig),subplot(N,2,31),imshow(imread('status_images/step_16_green.jpeg')), subplot(N,2,32),imshow(imread('status_images/status_green_skip.jpg')), catch, end
    else
        try figure(status_fig),subplot(N,2,31),imshow(imread('status_images/step_16_green.jpeg')), subplot(N,2,32),imshow(imread('status_images/status_green.jpg')), catch, end
    end
    clear skip
end

%% '17. HRF deconvolution'
if choices_preproc(17)==1
    try load([sub_dir{k} sprintf('workspace_variables_run%d.mat',k2)], 'cutoff_str'); catch, cutoff_str = '50'; end
    if run_on_HPF_data==1
        if exist([sub_dir{k} func_files{k,k2} '_WM' '.nii'], 'file') || exist([sub_dir{k} func_files{k,k2} '_WM' '.nii.gz'], 'file') % if only WM regress step was performed
            if exist([sub_dir{k} func_files{k,k2} '_WM' '_HPF' '01' '.mat'], 'file')
                namefile = [sub_dir{k} func_files{k,k2} '_WM' '_deconv' '_BPF' bpf_hz_str '.mat'];
            else
                namefile = [sub_dir{k} func_files{k,k2} '_WM' '_deconv' '.mat'];
            end
        end
        if exist([sub_dir{k} func_files{k,k2} '_denoised' cutoff_str '.nii'], 'file') || exist([sub_dir{k} func_files{k,k2} '_denoised' cutoff_str '.nii.gz'], 'file') % if only CSF regress step was performed
            if exist([sub_dir{k} func_files{k,k2} '_denoised' cutoff_str '_HPF' '01' '.mat'], 'file')
                namefile = [sub_dir{k} func_files{k,k2} '_denoised' cutoff_str '_deconv' '_BPF' bpf_hz_str '.mat'];
            else
                namefile = [sub_dir{k} func_files{k,k2} '_denoised' cutoff_str '_deconv' '.mat'];
            end
        end
        if exist([sub_dir{k} func_files{k,k2} '_cov' '.nii'], 'file') || exist([sub_dir{k} func_files{k,k2} '_cov' '.nii.gz'], 'file') % if only cov regress step was performed
            if exist([sub_dir{k} func_files{k,k2} '_cov' '_HPF' '01' '.mat'], 'file')
                namefile = [sub_dir{k} func_files{k,k2} '_cov' '_deconv' '_BPF' bpf_hz_str '.mat'];
            else
                namefile = [sub_dir{k} func_files{k,k2} '_cov' '_deconv' '.mat'];
            end
        end
        if exist([sub_dir{k} func_files{k,k2} '_WMcov' '.nii'], 'file') || exist([sub_dir{k} func_files{k,k2} '_WMcov' '.nii.gz'], 'file') % if both WM and cov regress steps were performed
            if exist([sub_dir{k} func_files{k,k2} '_WMcov' '_HPF' '01' '.mat'], 'file')
                namefile = [sub_dir{k} func_files{k,k2} '_WMcov' '_deconv' '_BPF' bpf_hz_str '.mat'];
            else
                namefile = [sub_dir{k} func_files{k,k2} '_WMcov' '_deconv' '.mat'];
            end
        end
        if exist([sub_dir{k} func_files{k,k2} '_denoised' cutoff_str 'WM' '.nii'], 'file') || exist([sub_dir{k} func_files{k,k2} '_denoised' cutoff_str 'WM' '.nii.gz'], 'file') % if both CSF and WM regress steps were performed
            if exist([sub_dir{k} func_files{k,k2} '_denoised' cutoff_str 'WM' '_HPF' '01' '.mat'], 'file')
                namefile = [sub_dir{k} func_files{k,k2} '_denoised' cutoff_str 'WM' '_deconv' '_BPF' bpf_hz_str '.mat'];
            else
                namefile = [sub_dir{k} func_files{k,k2} '_denoised' cutoff_str 'WM' '_deconv' '.mat'];
            end
        end
        if exist([sub_dir{k} func_files{k,k2} '_denoised' cutoff_str 'cov' '.nii'], 'file') || exist([sub_dir{k} func_files{k,k2} '_denoised' cutoff_str 'cov' '.nii.gz'], 'file') % if both CSF and cov regress steps were performed
            if exist([sub_dir{k} func_files{k,k2} '_denoised' cutoff_str 'cov' '_HPF' '01' '.mat'], 'file')
                namefile = [sub_dir{k} func_files{k,k2} '_denoised' cutoff_str 'cov' '_deconv' '_BPF' bpf_hz_str '.mat'];
            else
                namefile = [sub_dir{k} func_files{k,k2} '_denoised' cutoff_str 'cov' '_deconv' '.mat'];
            end
        end
        if exist([sub_dir{k} func_files{k,k2} '_denoised' cutoff_str 'WMcov' '.nii'], 'file') || exist([sub_dir{k} func_files{k,k2} '_denoised' cutoff_str 'WMcov' '.nii.gz'], 'file') % if CSF, WM and cov regress steps were performed
            if exist([sub_dir{k} func_files{k,k2} '_denoised' cutoff_str 'WMcov' '_HPF' '01' '.mat'], 'file')
                namefile = [sub_dir{k} func_files{k,k2} '_denoised' cutoff_str 'WMcov' '_deconv' '_BPF' bpf_hz_str '.mat'];
            else
                namefile = [sub_dir{k} func_files{k,k2} '_denoised' cutoff_str 'WMcov' '_deconv' '.mat'];
            end
        end
    elseif run_on_HPF_data==0
        if exist([sub_dir{k} func_files{k,k2} '_WM' '.nii'], 'file') || exist([sub_dir{k} func_files{k,k2} '_WM' '.nii.gz'], 'file') % if only WM regress step was performed
            if exist([sub_dir{k} func_files{k,k2} '_WM' '_BPF' bpf_hz_str '_deconv' '.mat'], 'file')
                namefile = [sub_dir{k} func_files{k,k2} '_WM' '_BPF' bpf_hz_str '_deconv' '.mat'];
            else
                namefile = [sub_dir{k} func_files{k,k2} '_WM' '_BPF' '10' '_deconv' '.mat'];
            end
        end
        if exist([sub_dir{k} func_files{k,k2} '_denoised' cutoff_str '.nii'], 'file') || exist([sub_dir{k} func_files{k,k2} '_denoised' cutoff_str '.nii.gz'], 'file') % if only CSF regress step was performed
            if exist([sub_dir{k} func_files{k,k2} '_denoised' cutoff_str '_BPF' bpf_hz_str '_deconv' '.mat'], 'file')
                namefile = [sub_dir{k} func_files{k,k2} '_denoised' cutoff_str '_BPF' bpf_hz_str '_deconv' '.mat'];
            else
                namefile = [sub_dir{k} func_files{k,k2} '_denoised' cutoff_str '_BPF' '10' '_deconv' '.mat'];
            end
        end
        if exist([sub_dir{k} func_files{k,k2} '_cov' '.nii'], 'file') || exist([sub_dir{k} func_files{k,k2} '_cov' '.nii.gz'], 'file') % if only cov regress step was performed
            if exist([sub_dir{k} func_files{k,k2} '_cov' '_BPF' bpf_hz_str '_deconv' '.mat'], 'file')
                namefile = [sub_dir{k} func_files{k,k2} '_cov' '_BPF' bpf_hz_str '_deconv' '.mat'];
            else
                namefile = [sub_dir{k} func_files{k,k2} '_cov' '_BPF' '10' '_deconv' '.mat'];
            end
        end
        if exist([sub_dir{k} func_files{k,k2} '_denoised' cutoff_str 'WM' '.nii'], 'file') || exist([sub_dir{k} func_files{k,k2} '_denoised' cutoff_str 'WM' '.nii.gz'], 'file') % if both CSF and WM regress steps were performed
            if exist([sub_dir{k} func_files{k,k2} '_denoised' cutoff_str 'WM' '_BPF' bpf_hz_str '_deconv' '.mat'], 'file')
                namefile = [sub_dir{k} func_files{k,k2} '_denoised' cutoff_str 'WM' '_BPF' bpf_hz_str '_deconv' '.mat'];
            else
                namefile = [sub_dir{k} func_files{k,k2} '_denoised' cutoff_str 'WM' '_BPF' '10' '_deconv' '.mat'];
            end
        end
        if exist([sub_dir{k} func_files{k,k2} '_denoised' cutoff_str 'cov' '.nii'], 'file') || exist([sub_dir{k} func_files{k,k2} '_denoised' cutoff_str 'cov' '.nii.gz'], 'file') % if both CSF and cov regress steps were performed
            if exist([sub_dir{k} func_files{k,k2} '_denoised' cutoff_str 'cov' '_BPF' bpf_hz_str '_deconv' '.mat'], 'file')
                namefile = [sub_dir{k} func_files{k,k2} '_denoised' cutoff_str 'cov' '_BPF' bpf_hz_str '_deconv' '.mat'];
            else
                namefile = [sub_dir{k} func_files{k,k2} '_denoised' cutoff_str 'cov' '_BPF' '10' '_deconv' '.mat'];
            end
        end
        if exist([sub_dir{k} func_files{k,k2} '_denoised' cutoff_str 'WMcov' '.nii'], 'file') || exist([sub_dir{k} func_files{k,k2} '_denoised' cutoff_str 'WMcov' '.nii.gz'], 'file') % if CSF, WM and cov regress steps were performed
            if exist([sub_dir{k} func_files{k,k2} '_denoised' cutoff_str 'WMcov' '_BPF' bpf_hz_str '_deconv' '.mat'], 'file')
                namefile = [sub_dir{k} func_files{k,k2} '_denoised' cutoff_str 'WMcov' '_BPF' bpf_hz_str '_deconv' '.mat'];
            else
                namefile = [sub_dir{k} func_files{k,k2} '_denoised' cutoff_str 'WMcov' '_BPF' '10' '_deconv' '.mat'];
            end
        end
    end
    if ~(exist('namefile', 'var'))
        namefile = [];
    end
    if ~(exist('bpf_hz', 'var'))
        bpf_hz = 0.1;
    end
    if ~(exist(namefile, 'file')) || (run_allsubs_by_default==1)
        try figure(status_fig),subplot(N,2,33),imshow(imread('status_images/step_17_blue.jpeg')), subplot(N,2,34),imshow(imread('status_images/status_blue.jpg')), catch, end
        clk=clock; L=L+1; logg{L,1}=sprintf('%d/%s/%d %dh:%dm:%ds - begin (17) HRF deconvolution for subject (%d) of (%d), run (%d) of (%d)',clk(3),month{clk(2)},clk(1),clk(4),clk(5),round(clk(6)),k,length(sub_dir),k2,runs(k)); try fprintf(fileID,'%s\n',logg{L,1}); set(S.tx,'string',logg); catch, end; clear clk
        try waitbar(((runs(k)*(k-1)+k2)/(length(sub_dir)*runs(k))),wbar2,sprintf('17. HRF deconvolution: Subject (%d) of (%d), Run (%d) of (%d)',k,length(sub_dir),k2,runs(k))); catch, end
        fprintf('17. HRF deconvolution: Subject (%d) of (%d), Run (%d) of (%d)\n',k,length(sub_dir),k2,runs(k))
        if runs(k)==1, Rns=0; else, Rns=k2; end
        time_17_deconvolution = tic;
        if rest==1
            scfMRItb_17_deconvolution_rest(sub_dir{k},func_files{k,k2},anat_files{k,1}, run_on_HPF_data,bpf_hz, wbar3,save_videos,Rns);
        elseif task==1
            scfMRItb_17_deconvolution_rest(sub_dir{k},func_files{k,k2},anat_files{k,1}, run_on_HPF_data,bpf_hz, wbar3,save_videos,Rns);
        end
        time_taken_preproc.step17_deconvolution(k,k2) = toc(time_17_deconvolution); clear time_17_deconvolution Rns
        try figure(status_fig),subplot(N,2,33),imshow(imread('status_images/step_17_green.jpeg')), subplot(N,2,34),imshow(imread('status_images/status_green.jpg')), catch, end
        clk=clock; L=L+1; logg{L,1}=sprintf('%d/%s/%d %dh:%dm:%ds - end (17) HRF deconvolution for subject (%d) of (%d), run (%d) of (%d)',clk(3),month{clk(2)},clk(1),clk(4),clk(5),round(clk(6)),k,length(sub_dir),k2,runs(k)); try fprintf(fileID,'%s\n',logg{L,1}); set(S.tx,'string',logg); catch, end; clear clk
        refresh_html_log
    else
        clk=clock; L=L+1; logg{L,1}=sprintf('%d/%s/%d %dh:%dm:%ds - skip (17) HRF deconvolution for subject (%d) of (%d), run (%d) of (%d)',clk(3),month{clk(2)},clk(1),clk(4),clk(5),round(clk(6)),k,length(sub_dir),k2,runs(k)); try fprintf(fileID,'%s\n',logg{L,1}); set(S.tx,'string',logg); catch, end; clear clk
        try figure(status_fig),subplot(N,2,33),imshow(imread('status_images/step_17_green.jpeg')), subplot(N,2,34),imshow(imread('status_images/status_green_skip.jpg')), catch, end
    end
end
% if task==1
%     clk=clock; L=L+1; logg{L,1}=sprintf('%d/%s/%d %dh:%dm:%ds - deconvolution for task fMRI data is yet to be adopted in this toolbox; not performing this step',clk(3),month{clk(2)},clk(1),clk(4),clk(5),round(clk(6))); try fprintf(fileID,'%s\n',logg{L,1}); set(S.tx,'string',logg); catch, end; clear clk
%     try figure(status_fig),subplot(N,2,33),imshow(imread('status_images/step_17_gray.jpeg')), subplot(N,2,34),imshow(imread('status_images/status_gray.jpg')), catch, end
%     msgbox('Deconvolution for task fMRI data is yet to be adopted in this toolbox; not performing this step.','Deconvolution not available for task fMRI','error')
% end

%% '18. Co-registration to PAM50 standard space'
if choices_preproc(18)==1
    if ((exist_SCT==1) && ~(exist([sub_dir{k} 'SCT_PAM50registration_step18/' 'warp_template2anat.nii.gz'],'file'))) || (run_allsubs_by_default==1)
        try figure(status_fig),subplot(N,2,35),imshow(imread('status_images/step_18_blue.jpeg')), subplot(N,2,36),imshow(imread('status_images/status_blue.jpg')), catch, end
        clk=clock; L=L+1; logg{L,1}=sprintf('%d/%s/%d %dh:%dm:%ds - begin (18) co-registration to PAM50 standard space for subject (%d) of (%d)',clk(3),month{clk(2)},clk(1),clk(4),clk(5),round(clk(6)),k,length(sub_dir)); try fprintf(fileID,'%s\n',logg{L,1}); set(S.tx,'string',logg); catch, end; clear clk
        try waitbar(((k)/(length(sub_dir))),wbar2,sprintf('18. Co-registration to PAM50 standard space: Subject (%d) of (%d)',k,length(sub_dir))); catch, end
        fprintf('18. Co-registration to PAM50 standard space: Subject (%d) of (%d)\n',k,length(sub_dir))
        time_18_PAM50registration = tic;
        scfMRItb_18_PAM50registration(sub_dir{k},anat_files{k}, wbar3,C_spine,SCT_vert_auto,SCT_manualreg_labels);
        time_taken_preproc.step18_PAM50registration(k,1) = toc(time_18_PAM50registration); clear time_18_PAM50registration Rns
        try figure(status_fig),subplot(N,2,35),imshow(imread('status_images/step_18_green.jpeg')), subplot(N,2,36),imshow(imread('status_images/status_green.jpg')), catch, end
        clk=clock; L=L+1; logg{L,1}=sprintf('%d/%s/%d %dh:%dm:%ds - end (18) co-registration to PAM50 standard space for subject (%d) of (%d)',clk(3),month{clk(2)},clk(1),clk(4),clk(5),round(clk(6)),k,length(sub_dir)); try fprintf(fileID,'%s\n',logg{L,1}); set(S.tx,'string',logg); catch, end; clear clk
        refresh_html_log
    else
        clk=clock; L=L+1; logg{L,1}=sprintf('%d/%s/%d %dh:%dm:%ds - skip (18) co-registration to PAM50 standard space for subject (%d) of (%d)',clk(3),month{clk(2)},clk(1),clk(4),clk(5),round(clk(6)),k,length(sub_dir)); try fprintf(fileID,'%s\n',logg{L,1}); set(S.tx,'string',logg); catch, end; clear clk
    end
end

%% Calculate tSNR across pre-processing steps
if (choices_preproc(5)==1)||(choices_preproc(6)==1)||(choices_preproc(8)==1)||(choices_preproc(9)==1)||(choices_preproc(11)==1)||(choices_preproc(12)==1)||(choices_preproc(13)==1)||(choices_preproc(14)==1)
    if ~( (exist([sub_dir{k} 'tSNR_across_preproc_steps.mat'], 'file'))||(exist([sub_dir{k} 'tSNR_across_preproc_steps_run' num2str(k2) '.mat'], 'file')) ) || (run_allsubs_by_default==1)
        clk=clock; L=L+1; logg{L,1}=sprintf('%d/%s/%d %dh:%dm:%ds - begin calculating tSNR across pre-processing steps for subject (%d) of (%d), run (%d) of (%d)',clk(3),month{clk(2)},clk(1),clk(4),clk(5),round(clk(6)),k,length(sub_dir),k2,runs(k)); try fprintf(fileID,'%s\n',logg{L,1}); set(S.tx,'string',logg); catch, end; clear clk
        try waitbar(((runs(k)*(k-1)+k2)/(length(sub_dir)*runs(k))),wbar2,sprintf('Calculate tSNR across pre-processing steps: Subject (%d) of (%d), Run (%d) of (%d)',k,length(sub_dir),k2,runs(k))); catch, end
        fprintf('Calculate tSNR across pre-processing steps: Subject (%d) of (%d), Run (%d) of (%d)\n',k,length(sub_dir),k2,runs(k))
        if runs(k)==1, Rns=0; else, Rns=k2; end
        time_tSNR_graphs = tic;
        scfMRItb_tSNR_graphs(sub_dir{k},func_files{k,k2},anat_files{k,1},choices_preproc, wbar3,Rns);
        time_taken_preproc.tSNR_computation(k,k2) = toc(time_tSNR_graphs); clear time_tSNR_graphs Rns
        clk=clock; L=L+1; logg{L,1}=sprintf('%d/%s/%d %dh:%dm:%ds - end calculating tSNR across pre-processing steps for subject (%d) of (%d), run (%d) of (%d)',clk(3),month{clk(2)},clk(1),clk(4),clk(5),round(clk(6)),k,length(sub_dir),k2,runs(k)); try fprintf(fileID,'%s\n',logg{L,1}); set(S.tx,'string',logg); catch, end; clear clk
        refresh_html
    else
        clk=clock; L=L+1; logg{L,1}=sprintf('%d/%s/%d %dh:%dm:%ds - skip calculating tSNR across pre-processing steps for subject (%d) of (%d), run (%d) of (%d)',clk(3),month{clk(2)},clk(1),clk(4),clk(5),round(clk(6)),k,length(sub_dir),k2,runs(k)); try fprintf(fileID,'%s\n',logg{L,1}); set(S.tx,'string',logg); catch, end; clear clk
    end
end

%% '19. gzip NIfTI files to save disk space'
sdirall = dir([sub_dir{k} '**/*.*']); sdirall = sdirall(3:end);
sdirall_bytes = {sdirall.bytes}';
sdirall_bytes = cell2mat(sdirall_bytes);
sdirall_bytes = sum(sdirall_bytes)/10^9; % size of subject folder in gigabytes
folder_size_1(k,1) = sdirall_bytes;
clear sdirall sdirall_bytes

if remove_slicewise_files==1
    fprintf('19. Remove slice-wise files: Subject (%d) of (%d), Run (%d) of (%d)\n',k,length(sub_dir),k2,runs(k))
    time_19_remove_slicewise = tic;
    scfMRItb_19_remove_slicewise_files(sub_dir{k},func_files(k,k2),anat_files{k,1},wbar3)
    time_taken_preproc.step19_remove_slicewise_files(k,1) = toc(time_19_remove_slicewise); clear time_19_remove_slicewise %#ok<*SAGROW>
    
    sdirall = dir([sub_dir{k} '**/*.*']); sdirall = sdirall(3:end);
    sdirall_bytes = {sdirall.bytes}';
    sdirall_bytes = cell2mat(sdirall_bytes);
    sdirall_bytes = sum(sdirall_bytes)/10^9; % size of subject folder in gigabytes
    folder_size_2(k,1) = sdirall_bytes;
    clear sdirall sdirall_bytes
end

if choices_preproc(19)==1
    try figure(status_fig),subplot(N,2,37),imshow(imread('status_images/step_19_blue.jpeg')), subplot(N,2,38),imshow(imread('status_images/status_blue.jpg')), catch, end
    skip=0;
    switch answer_gzip
        case 'gzip only, don''t delete'
            clk=clock; L=L+1; logg{L,1}=sprintf('%d/%s/%d %dh:%dm:%ds - gzip NIfTI files for Subject (%d) of (%d), Run (%d) of (%d)',clk(3),month{clk(2)},clk(1),clk(4),clk(5),round(clk(6)),k,length(sub_dir),k2,runs(k)); try fprintf(fileID,'%s\n',logg{L,1}); set(S.tx,'string',logg); catch, end; clear clk
            try waitbar((k/length(sub_dir)),wbar2,sprintf('19. gzip NIfTI files: Subject (%d) of (%d), Run (%d) of (%d)',k,length(sub_dir),k2,runs(k))); catch, end
            fprintf('19. gzip NIfTI files: Subject (%d) of (%d), Run (%d) of (%d)\n',k,length(sub_dir),k2,runs(k))
            time_19_gzip_nii = tic;
            skp = scfMRItb_19_gzip_nii(sub_dir{k},wbar3); skip=skip+single(skp>0); clear skp
            time_taken_preproc.step19_gzip_nii(k,1) = toc(time_19_gzip_nii); clear time_19_gzip_nii
        case 'delete only, I already gzipped'
            clk=clock; L=L+1; logg{L,1}=sprintf('%d/%s/%d %dh:%dm:%ds - delete .nii files for Subject (%d) of (%d), Run (%d) of (%d)',clk(3),month{clk(2)},clk(1),clk(4),clk(5),round(clk(6)),k,length(sub_dir),k2,runs(k)); try fprintf(fileID,'%s\n',logg{L,1}); set(S.tx,'string',logg); catch, end; clear clk
            try waitbar((k/length(sub_dir)),wbar2,sprintf('19. Delete .nii files to save disk space (.nii.gz preserved): Subject (%d) of (%d), Run (%d) of (%d)',k,length(sub_dir),k2,runs(k))); catch, end
            fprintf('19. Delete .nii files to save disk space (.nii.gz preserved): Subject (%d) of (%d), Run (%d) of (%d)\n',k,length(sub_dir),k2,runs(k))
            time_19_delete_nii = tic;
            skp = scfMRItb_19_delete_nii(sub_dir{k},wbar3); skip=skip+single(skp>0); clear skp
            time_taken_preproc.step19_delete_nii(k,1) = toc(time_19_delete_nii); clear time_19_delete_nii
        case 'both gzip and delete'
            clk=clock; L=L+1; logg{L,1}=sprintf('%d/%s/%d %dh:%dm:%ds - gzip NIfTI files for Subject (%d) of (%d), Run (%d) of (%d)',clk(3),month{clk(2)},clk(1),clk(4),clk(5),round(clk(6)),k,length(sub_dir),k2,runs(k)); try fprintf(fileID,'%s\n',logg{L,1}); set(S.tx,'string',logg); catch, end; clear clk
            try waitbar((k/length(sub_dir)),wbar2,sprintf('19. gzip NIfTI files: Subject (%d) of (%d), Run (%d) of (%d)',k,length(sub_dir),k2,runs(k))); catch, end
            fprintf('19. gzip NIfTI files: Subject (%d) of (%d), Run (%d) of (%d)\n',k,length(sub_dir),k2,runs(k))
            time_19_gzip_delete_nii = tic;
            skp = scfMRItb_19_gzip_nii(sub_dir{k},wbar3); skip=skip+single(skp>0); clear skp
            clk=clock; L=L+1; logg{L,1}=sprintf('%d/%s/%d %dh:%dm:%ds - delete .nii files for Subject (%d) of (%d), Run (%d) of (%d)',clk(3),month{clk(2)},clk(1),clk(4),clk(5),round(clk(6)),k,length(sub_dir),k2,runs(k)); try fprintf(fileID,'%s\n',logg{L,1}); set(S.tx,'string',logg); catch, end; clear clk
            try waitbar((k/length(sub_dir)),wbar2,sprintf('19. Delete .nii files to save disk space (.nii.gz preserved): Subject (%d) of (%d), Run (%d) of (%d)',k,length(sub_dir),k2,runs(k))); catch, end
            fprintf('19. Delete .nii files to save disk space (.nii.gz preserved): Subject (%d) of (%d), Run (%d) of (%d)\n',k,length(sub_dir),k2,runs(k))
            skp = scfMRItb_19_delete_nii(sub_dir{k},wbar3); skip=skip+single(skp>0); clear skp
            time_taken_preproc.step19_gzip_delete_nii(k,1) = toc(time_19_gzip_delete_nii); clear time_19_gzip_delete_nii
    end
    
    % size of subject folder after removal of slicewise files and gzipping
    sdirall = dir([sub_dir{k} '**/*.*']); sdirall = sdirall(3:end);
    sdirall_name = {sdirall.name}';
    excludefile = zeros(length(sdirall_name),1);
    for m=1:length(sdirall_name)
        if length(sdirall_name{m})>=4
            if strcmp(sdirall_name{m}(end-3:end),'.nii')==1
                for m2=1:length(sdirall_name)
                    if length(sdirall_name{m2})>=7
                        if (strcmp(sdirall_name{m}(1:end-4),sdirall_name{m2}(1:end-7))==1) && (strcmp(sdirall_name{m2}(end-6:end),'.nii.gz')==1) && (m~=m2)
                            excludefile(m) = 1; % if a .nii file is found AND if a corresponding .nii.gz file also exists, then do not count the .nii file for total storage computation
                        end
                    end
                end
            end
        end
    end
    sdirall_bytes = {sdirall.bytes}';
    sdirall_bytes = cell2mat(sdirall_bytes);
    sdirall_bytes = sdirall_bytes(find(excludefile==0));
    sdirall_bytes = sum(sdirall_bytes)/10^9; % size of subject folder in gigabytes
    folder_size_3(k,1) = sdirall_bytes;
    clear sdirall sdirall_bytes sdirall_name m m2 excludefile

    if skip==0
        try figure(status_fig),subplot(N,2,37),imshow(imread('status_images/step_19_green.jpeg')), subplot(N,2,38),imshow(imread('status_images/status_green_skip.jpg')), catch, end
    else
        try figure(status_fig),subplot(N,2,37),imshow(imread('status_images/step_19_green.jpeg')), subplot(N,2,38),imshow(imread('status_images/status_green.jpg')), catch, end
    end
    clear skip
    refresh_html_log
end

tmpc=clock; clk = sprintf('%.4d-%.2d-%.2d_%.2d-%.2d',tmpc(1),tmpc(2),tmpc(3),tmpc(4),tmpc(5)); clear tmpc
dskfilename = sprintf('disk_storage_space_info_%s.mat',clk); clear clk
% save disk storage info and figures
% save disk storage info into .mat file
if exist('folder_size_2','var') && exist('folder_size_3','var')
    folder_size_1_sub = folder_size_1(k); folder_size_2_sub = folder_size_2(k); folder_size_3_sub = folder_size_3(k);
    readme_disk_storage = sprintf('These numbers are the total storage space (in gigabytes) taken by all files in this subject folder (%s)\nfolder_size_1_sub = %.2f (size of subject folder after pre-processing before removal of slicewise files and gzipping)\nfolder_size_2_sub = %.2f (size of subject folder after pre-processing and removal of slicewise files but before gzipping)\nfolder_size_3_sub = %.2f (size of subject folder after pre-processing, removal of slicewise files and gzipping)\n',sub_dir{k},folder_size_1_sub,folder_size_2_sub,folder_size_3_sub);
    save([sub_dir{k} dskfilename],'folder_size_1_sub','folder_size_2_sub','folder_size_3_sub','readme_disk_storage')
elseif exist('folder_size_2','var') && ~exist('folder_size_3','var')
    folder_size_1_sub = folder_size_1(k); folder_size_2_sub = folder_size_2(k);
    readme_disk_storage = sprintf('These numbers are the total storage space (in gigabytes) taken by all files in this subject folder (%s)\nfolder_size_1_sub = %.2f (size of subject folder after pre-processing before removal of slicewise files and gzipping)\nfolder_size_2_sub = %.2f (size of subject folder after pre-processing and removal of slicewise files but before gzipping)\nfolder_size_3_sub = <"gzipping" not performed> (size of subject folder after pre-processing, removal of slicewise files and gzipping)\n',sub_dir{k},folder_size_1_sub,folder_size_2_sub);
    save([sub_dir{k} dskfilename],'folder_size_1_sub','folder_size_2_sub','readme_disk_storage')
elseif ~exist('folder_size_2','var') && exist('folder_size_3','var')
    folder_size_1_sub = folder_size_1(k); folder_size_3_sub = folder_size_3(k);
    readme_disk_storage = sprintf('These numbers are the total storage space (in gigabytes) taken by all files in this subject folder (%s)\nfolder_size_1_sub = %.2f (size of subject folder after pre-processing before removal of slicewise files and gzipping)\nfolder_size_2_sub = <"removal of slicewise files" not performed> (size of subject folder after pre-processing and removal of slicewise files but before gzipping)\nfolder_size_3_sub = %.2f (size of subject folder after pre-processing, removal of slicewise files and gzipping)\n',sub_dir{k},folder_size_1_sub,folder_size_3_sub);
    save([sub_dir{k} dskfilename],'folder_size_1_sub','folder_size_3_sub','readme_disk_storage')
elseif ~exist('folder_size_2','var') && ~exist('folder_size_3','var')
    folder_size_1_sub = folder_size_1(k);
    readme_disk_storage = sprintf('These numbers are the total storage space (in gigabytes) taken by all files in this subject folder (%s)\nfolder_size_1_sub = %.2f (size of subject folder after pre-processing before removal of slicewise files and gzipping)\nfolder_size_2_sub = <"removal of slicewise files" not performed> (size of subject folder after pre-processing and removal of slicewise files but before gzipping)\nfolder_size_3_sub = <"gzipping" not performed> (size of subject folder after pre-processing, removal of slicewise files and gzipping)\n',sub_dir{k},folder_size_1_sub);
    save([sub_dir{k} dskfilename],'folder_size_1_sub','readme_disk_storage')
end

% save bar graphs
if ~(exist([sub_dir{k} 'QC' '/19_disk_storage_info'],'dir'))
    mkdir([sub_dir{k} 'QC' '/19_disk_storage_info'])
end
QCpath = [sub_dir{k} 'QC' '/19_disk_storage_info/'];
if ~exist('folder_size_2_sub','var'), folder_size_2_sub = 0; end
if ~exist('folder_size_3_sub','var'), folder_size_3_sub = 0; end

fighndl = figure('Position',[1 1 1200 800],'Color',[0.85,0.90,0.85],'InvertHardcopy','off','Visible','off'); % [0.85,0.90,0.85] [0.8, 0.89, 0.94]
b = bar(1,folder_size_1_sub,0.5,'FaceColor',[0,0.55,0.69],'EdgeColor',[0.65,0.11,0.19],'LineWidth',1.5); hold on,
    xtips1 = b(1).XEndPoints; ytips1 = b(1).YEndPoints; labels1 = string(round(b(1).YData,2)); clear b
    labels1 = strcat(labels1,' GB');
    text(xtips1,ytips1,labels1,'HorizontalAlignment','center','VerticalAlignment','bottom','fontsize',17','color',[0.65,0.11,0.19]);
b = bar(2,folder_size_2_sub,0.5,'FaceColor',[0,0.65,0.69],'EdgeColor',[0.65,0.11,0.19],'LineWidth',1.5);
    xtips2 = b(1).XEndPoints; ytips2 = b(1).YEndPoints; labels2 = string(round(b(1).YData,2)); clear b
    labels2 = strcat(labels2,' GB');
    if folder_size_2_sub==0, labels2 = 'not performed'; end
    text(xtips2,ytips2,labels2,'HorizontalAlignment','center','VerticalAlignment','bottom','fontsize',17','color',[0.65,0.11,0.19]);
b = bar(3,folder_size_3_sub,0.5,'FaceColor',[0,0.75,0.69],'EdgeColor',[0.65,0.11,0.19],'LineWidth',1.5);
	xtips3 = b(1).XEndPoints; ytips3 = b(1).YEndPoints; labels3 = string(round(b(1).YData,2)); clear b
    labels3 = strcat(labels3,' GB');
    if folder_size_3_sub==0, labels3 = 'not performed'; end
    text(xtips3,ytips3,labels3,'HorizontalAlignment','center','VerticalAlignment','bottom','fontsize',17','color',[0.65,0.11,0.19]);
    grid on, grid minor, ylim([0 max(folder_size_1_sub)+0.1.*max(folder_size_1_sub)]), ylabel('Disk storage (in gigabytes)'),
    xticks([1,2,3]), xticklabels({'At the end of pre-processing','After removal of slice-wise files','After NIfTI file compression (gzip)'})
    a = get(gca,'xticklabels'); set(gca,'xticklabels',a,'fontsize',14); clear a
    legstr{1} = sprintf('At the end of pre-processing'); legstr{2} = sprintf('... AND after removal of duplicate slice-wise files'); legstr{3} = sprintf('... AND after NIfTI file compression (gzip)');
    legend(legstr,'Location','northeast','FontSize',18)
    title(sprintf('DISK STORAGE: size of subject folder at the end of pre-processing\n(%s)',sub_dir{k}),'Interpreter', 'none', 'Color',[0 0.37 0.22],'FontSize',18);
saveas(fighndl,[QCpath 'disk_storage_bargraph' '.jpg'])
tmp=clock; clk_settings = sprintf('%.4d-%.2d-%.2d_%.2d-%.2d',tmp(1),tmp(2),tmp(3),tmp(4),tmp(5)); clear tmp
saveas(fighndl,[QCpath sprintf('disk_storage_bargraph_%s',clk_settings) '.jpg'])
close(fighndl)
clear folder_size_1_sub folder_size_2_sub folder_size_3_sub readme_disk_storage QCpath fighndl legstr b xtips1 ytips1 labels1 xtips2 ytips2 labels2 xtips3 ytips3 labels3 clk_settings

clk=clock; L=L+1; logg{L,1}=sprintf('%d/%s/%d %dh:%dm:%ds - Subject (%d) of (%d), Run (%d) of (%d) COMPLETED',clk(3),month{clk(2)},clk(1),clk(4),clk(5),round(clk(6)),k,length(sub_dir),k2,runs(k)); try fprintf(fileID,'%s\n',logg{L,1}); set(S.tx,'string',logg); catch, end; clear clk
L=L+1; logg{L,1}=sprintf('Total time taken so far: %dh:%dm:%ds\nTime taken for this subject: %dh:%dm:%ds',floor(toc(tc0)/3600),floor(mod(toc(tc0),3600)/60),floor(mod(toc(tc0),60)),floor(toc(tc)/3600),floor(mod(toc(tc),3600)/60),floor(mod(toc(tc),60))); try fprintf(fileID,'%s\n',logg{L,1}); set(S.tx,'string',logg); catch, end; clear clk

%% Refresh status figures to the original state for the next subject / run
if ishandle(status_fig)==1
try
figure(status_fig)
if choices_preproc(1)==1
    subplot(N,2,1),imshow(imread('status_images/step_01_green.jpeg')), subplot(N,2,2),imshow(imread('status_images/status_green.jpg')),
else
    subplot(N,2,1),imshow(imread('status_images/step_01_gray.jpeg')), subplot(N,2,2),imshow(imread('status_images/status_gray.jpg')),
end
if choices_preproc(2)==1
    subplot(N,2,3),imshow(imread('status_images/step_02_white.jpeg')), subplot(N,2,4),imshow(imread('status_images/status_white_UI.jpg')),
else
    subplot(N,2,3),imshow(imread('status_images/step_02_gray.jpeg')), subplot(N,2,4),imshow(imread('status_images/status_gray.jpg')),
end
if choices_preproc(3)==1
    subplot(N,2,5),imshow(imread('status_images/step_03_white.jpeg')), subplot(N,2,6),imshow(imread('status_images/status_white.jpg')),
else
    subplot(N,2,5),imshow(imread('status_images/step_03_gray.jpeg')), subplot(N,2,6),imshow(imread('status_images/status_gray.jpg')),
end
if choices_preproc(4)==1
    subplot(N,2,7),imshow(imread('status_images/step_04_white.jpeg')), subplot(N,2,8),imshow(imread('status_images/status_white.jpg')),
else
    subplot(N,2,7),imshow(imread('status_images/step_04_gray.jpeg')), subplot(N,2,8),imshow(imread('status_images/status_gray.jpg')),
end
if choices_preproc(5)==1
    subplot(N,2,9),imshow(imread('status_images/step_05_white.jpeg')), subplot(N,2,10),imshow(imread('status_images/status_white_UI.jpg')),
else
    subplot(N,2,9),imshow(imread('status_images/step_05_gray.jpeg')), subplot(N,2,10),imshow(imread('status_images/status_gray.jpg')),
end
if choices_preproc(6)==1
    subplot(N,2,11),imshow(imread('status_images/step_06_white.jpeg')), subplot(N,2,12),imshow(imread('status_images/status_white.jpg')),
else
    subplot(N,2,11),imshow(imread('status_images/step_06_gray.jpeg')), subplot(N,2,12),imshow(imread('status_images/status_gray.jpg')),
end
if choices_preproc(7)==1
    subplot(N,2,13),imshow(imread('status_images/step_07_white.jpeg')), subplot(N,2,14),imshow(imread('status_images/status_white_UI.jpg')),
else
    subplot(N,2,13),imshow(imread('status_images/step_07_gray.jpeg')), subplot(N,2,14),imshow(imread('status_images/status_gray.jpg')),
end
if choices_preproc(8)==1
    subplot(N,2,15),imshow(imread('status_images/step_08_white.jpeg')), subplot(N,2,16),imshow(imread('status_images/status_white.jpg')),
else
    subplot(N,2,15),imshow(imread('status_images/step_08_gray.jpeg')), subplot(N,2,16),imshow(imread('status_images/status_gray.jpg')),
end
if choices_preproc(9)==1
    subplot(N,2,17),imshow(imread('status_images/step_09_white.jpeg')), subplot(N,2,18),imshow(imread('status_images/status_white.jpg')),
else
    subplot(N,2,17),imshow(imread('status_images/step_09_gray.jpeg')), subplot(N,2,18),imshow(imread('status_images/status_gray.jpg')),
end
if choices_preproc(10)==1
    subplot(N,2,19),imshow(imread('status_images/step_10_white.jpeg')), subplot(N,2,20),imshow(imread('status_images/status_white_UI.jpg')),
else
    subplot(N,2,19),imshow(imread('status_images/step_10_gray.jpeg')), subplot(N,2,20),imshow(imread('status_images/status_gray.jpg')),
end
if choices_preproc(11)==1
    subplot(N,2,21),imshow(imread('status_images/step_11_white.jpeg')), subplot(N,2,22),imshow(imread('status_images/status_white.jpg')),
else
    subplot(N,2,21),imshow(imread('status_images/step_11_gray.jpeg')), subplot(N,2,22),imshow(imread('status_images/status_gray.jpg')),
end
if choices_preproc(12)==1
    subplot(N,2,23),imshow(imread('status_images/step_12_white.jpeg')), subplot(N,2,24),imshow(imread('status_images/status_white.jpg')),
else
    subplot(N,2,23),imshow(imread('status_images/step_12_gray.jpeg')), subplot(N,2,24),imshow(imread('status_images/status_gray.jpg')),
end
if choices_preproc(13)==1
    subplot(N,2,25),imshow(imread('status_images/step_13_white.jpeg')), subplot(N,2,26),imshow(imread('status_images/status_white.jpg')),
else
    subplot(N,2,25),imshow(imread('status_images/step_13_gray.jpeg')), subplot(N,2,26),imshow(imread('status_images/status_gray.jpg')),
end
if choices_preproc(14)==1
    subplot(N,2,27),imshow(imread('status_images/step_14_white.jpeg')), subplot(N,2,28),imshow(imread('status_images/status_white.jpg')),
else
    subplot(N,2,27),imshow(imread('status_images/step_14_gray.jpeg')), subplot(N,2,28),imshow(imread('status_images/status_gray.jpg')),
end
if choices_preproc(15)==1
    subplot(N,2,29),imshow(imread('status_images/step_15_white.jpeg')), subplot(N,2,30),imshow(imread('status_images/status_white.jpg')),
else
    subplot(N,2,29),imshow(imread('status_images/step_15_gray.jpeg')), subplot(N,2,30),imshow(imread('status_images/status_gray.jpg')),
end
if choices_preproc(16)==1
    subplot(N,2,31),imshow(imread('status_images/step_16_white.jpeg')), subplot(N,2,32),imshow(imread('status_images/status_white_UI.jpg')),
else
    subplot(N,2,31),imshow(imread('status_images/step_16_gray.jpeg')), subplot(N,2,32),imshow(imread('status_images/status_gray.jpg')),
end
if choices_preproc(17)==1
    subplot(N,2,33),imshow(imread('status_images/step_17_white.jpeg')), subplot(N,2,34),imshow(imread('status_images/status_white.jpg')),
else
    subplot(N,2,33),imshow(imread('status_images/step_17_gray.jpeg')), subplot(N,2,34),imshow(imread('status_images/status_gray.jpg')),
end
if choices_preproc(18)==1
    subplot(N,2,35),imshow(imread('status_images/step_18_white.jpeg')), subplot(N,2,36),imshow(imread('status_images/status_white.jpg')),
else
    subplot(N,2,35),imshow(imread('status_images/step_18_gray.jpeg')), subplot(N,2,36),imshow(imread('status_images/status_gray.jpg')),
end
if choices_preproc(19)==1
    subplot(N,2,37),imshow(imread('status_images/step_19_white.jpeg')), subplot(N,2,38),imshow(imread('status_images/status_white.jpg')),
else
    subplot(N,2,37),imshow(imread('status_images/step_19_gray.jpeg')), subplot(N,2,38),imshow(imread('status_images/status_gray.jpg')),
    
end
catch
end
end

    end; clear k2
    
    refresh_html
    
end; clear k

end

%%
%% Save time taken for each step
ttaken_total = zeros(19,1); ttaken_step = zeros(19,1); ttaken_sub = zeros(length(sub_dir),1);
try
if choices_preproc(1)==1
    temp = time_taken_preproc.step01_NORDICdenoising(find(time_taken_preproc.step01_NORDICdenoising~=0));
    ttaken_total(1) = sum(temp); ttaken_step(1) = mean(temp); clear temp
    temp = sum(time_taken_preproc.step01_NORDICdenoising,2); ttaken_sub = sum([ttaken_sub,temp],2); clear temp
end
if choices_preproc(2)==1
    temp = time_taken_preproc.step02_slicetimingcorrection(find(time_taken_preproc.step02_slicetimingcorrection~=0));
    ttaken_total(2) = sum(temp); ttaken_step(2) = mean(temp); clear temp
    temp = sum(time_taken_preproc.step02_slicetimingcorrection,2); ttaken_sub = sum([ttaken_sub,temp],2); clear temp
end
if choices_preproc(3)==1
    temp = time_taken_preproc.step03_matchAnatFuncSlices(find(time_taken_preproc.step03_matchAnatFuncSlices~=0));
    ttaken_total(3) = sum(temp); ttaken_step(3) = mean(temp); clear temp
    temp = sum(time_taken_preproc.step03_matchAnatFuncSlices,2); ttaken_sub = sum([ttaken_sub,temp],2); clear temp
end
if choices_preproc(4)==1
    temp = time_taken_preproc.step04_splitData(find(time_taken_preproc.step04_splitData~=0));
    ttaken_total(4) = sum(temp); ttaken_step(4) = mean(temp); clear temp
    temp = sum(time_taken_preproc.step04_splitData,2); ttaken_sub = sum([ttaken_sub,temp],2); clear temp
end
if choices_preproc(5)==1
    temp = time_taken_preproc.step05_notCordMask(find(time_taken_preproc.step05_notCordMask~=0));
    ttaken_total(5) = sum(temp); ttaken_step(5) = mean(temp); clear temp
    temp = sum(time_taken_preproc.step05_notCordMask,2); ttaken_sub = sum([ttaken_sub,temp],2); clear temp
end
if choices_preproc(6)==1
    temp = time_taken_preproc.step06_denoise1(find(time_taken_preproc.step06_denoise1~=0));
    ttaken_total(6) = sum(temp); ttaken_step(6) = mean(temp); clear temp
    temp = sum(time_taken_preproc.step06_denoise1,2); ttaken_sub = sum([ttaken_sub,temp],2); clear temp
end
if choices_preproc(7)==1
    temp = time_taken_preproc.step07_gaussianMask(find(time_taken_preproc.step07_gaussianMask~=0));
    ttaken_total(7) = sum(temp); ttaken_step(7) = mean(temp); clear temp
    temp = sum(time_taken_preproc.step07_gaussianMask,2); ttaken_sub = sum([ttaken_sub,temp],2); clear temp
end
if choices_preproc(8)==1
    temp = time_taken_preproc.step08_motionCorrection(find(time_taken_preproc.step08_motionCorrection~=0));
    ttaken_total(8) = sum(temp); ttaken_step(8) = mean(temp); clear temp
    temp = sum(time_taken_preproc.step08_motionCorrection,2); ttaken_sub = sum([ttaken_sub,temp],2); clear temp
end
if choices_preproc(9)==1
    temp = time_taken_preproc.step09_retroicor(find(time_taken_preproc.step09_retroicor~=0));
    ttaken_total(9) = sum(temp); ttaken_step(9) = mean(temp); clear temp
    temp = sum(time_taken_preproc.step09_retroicor,2); ttaken_sub = sum([ttaken_sub,temp],2); clear temp
end
if choices_preproc(10)==1
    temp = time_taken_preproc.step10_GMWMCSFmasks(find(time_taken_preproc.step10_GMWMCSFmasks~=0));
    ttaken_total(10) = sum(temp); ttaken_step(10) = mean(temp); clear temp
    temp = sum(time_taken_preproc.step10_GMWMCSFmasks,2); ttaken_sub = sum([ttaken_sub,temp],2); clear temp
end
if choices_preproc(11)==1
    temp = time_taken_preproc.step11_FuncAnatRegistration(find(time_taken_preproc.step11_FuncAnatRegistration~=0));
    ttaken_total(11) = sum(temp); ttaken_step(11) = mean(temp); clear temp
    temp = sum(time_taken_preproc.step11_FuncAnatRegistration,2); ttaken_sub = sum([ttaken_sub,temp],2); clear temp
end
if choices_preproc(12)==1
    temp = time_taken_preproc.step12_CSFregress(find(time_taken_preproc.step12_CSFregress~=0));
    ttaken_total(12) = sum(temp); ttaken_step(12) = mean(temp); clear temp
    temp = sum(time_taken_preproc.step12_CSFregress,2); ttaken_sub = sum([ttaken_sub,temp],2); clear temp
end
if choices_preproc(13)==1
    temp = time_taken_preproc.step13_WMregress(find(time_taken_preproc.step13_WMregress~=0));
    ttaken_total(13) = sum(temp); ttaken_step(13) = mean(temp); clear temp
    temp = sum(time_taken_preproc.step13_WMregress,2); ttaken_sub = sum([ttaken_sub,temp],2); clear temp
end
if choices_preproc(14)==1
    temp = time_taken_preproc.step14_COVregress(find(time_taken_preproc.step14_COVregress~=0));
    ttaken_total(14) = sum(temp); ttaken_step(14) = mean(temp); clear temp
    temp = sum(time_taken_preproc.step14_COVregress,2); ttaken_sub = sum([ttaken_sub,temp],2); clear temp
end
if choices_preproc(15)==1
    temp = time_taken_preproc.step15_filterData(find(time_taken_preproc.step15_filterData~=0));
    ttaken_total(15) = sum(temp); ttaken_step(15) = mean(temp); clear temp
    temp = sum(time_taken_preproc.step15_filterData,2); ttaken_sub = sum([ttaken_sub,temp],2); clear temp
end
if choices_preproc(16)==1
    temp = time_taken_preproc.step16_cord_quadrants(find(time_taken_preproc.step16_cord_quadrants~=0));
    ttaken_total(16) = sum(temp); ttaken_step(16) = mean(temp); clear temp
    temp = sum(time_taken_preproc.step16_cord_quadrants,2); ttaken_sub = sum([ttaken_sub,temp],2); clear temp
end
if choices_preproc(17)==1
    temp = time_taken_preproc.step17_deconvolution(find(time_taken_preproc.step17_deconvolution~=0));
    ttaken_total(17) = sum(temp); ttaken_step(17) = mean(temp); clear temp
    temp = sum(time_taken_preproc.step17_deconvolution,2); ttaken_sub = sum([ttaken_sub,temp],2); clear temp
end
if choices_preproc(18)==1
    temp = time_taken_preproc.step18_PAM50registration(find(time_taken_preproc.step18_PAM50registration~=0));
    ttaken_total(18) = sum(temp); ttaken_step(18) = mean(temp); clear temp
    temp = sum(time_taken_preproc.step18_PAM50registration,2); ttaken_sub = sum([ttaken_sub,temp],2); clear temp
end
try
    temp = time_taken_preproc.tSNR_computation(find(time_taken_preproc.tSNR_computation~=0));
    ttaken_total(19) = sum([ttaken_total(19),sum(temp)]); ttaken_step(19) = sum([ttaken_step(19),mean(temp)]); clear temp
    temp = sum(time_taken_preproc.tSNR_computation,2); ttaken_sub = sum([ttaken_sub,temp],2); clear temp
catch
end
try
    temp = time_taken_preproc.step19_remove_slicewise_files(find(time_taken_preproc.step19_remove_slicewise_files~=0));
    ttaken_total(19) = sum([ttaken_total(19),sum(temp)]); ttaken_step(19) = sum([ttaken_step(19),mean(temp)]); clear temp
    temp = sum(time_taken_preproc.step19_remove_slicewise_files,2); ttaken_sub = sum([ttaken_sub,temp],2); clear temp
catch
end
if choices_preproc(19)==1
    temp = time_taken_preproc.step19_gzip_delete_nii(find(time_taken_preproc.step19_gzip_delete_nii~=0));
    ttaken_total(19) = sum([ttaken_total(19),sum(temp)]); ttaken_step(19) = sum([ttaken_step(19),mean(temp)]); clear temp
    temp = sum(time_taken_preproc.step19_gzip_delete_nii,2); ttaken_sub = sum([ttaken_sub,temp],2); clear temp
end
catch
end

try
ttaken_total = ttaken_total(find(choices_preproc==1));
ttaken_step = ttaken_step(find(choices_preproc==1));

fighndl = figure('Position',[1 1 1200 800],'Color',[0.85,0.90,0.85],'InvertHardcopy','off','Visible','off'); % [0.85,0.90,0.85] [0.8, 0.89, 0.94]
subplot2n(3,1,1), b = bar(ttaken_total,0.5,'FaceColor',[0,0.55,0.69],'EdgeColor',[0.65,0.11,0.19],'LineWidth',1.5);
    xticks([1:length(find(choices_preproc==1))]), xticklabels(mat2cell(find(choices_preproc==1),length(find(choices_preproc==1))))
	xtips1 = b(1).XEndPoints; ytips1 = b(1).YEndPoints;
    for w=1:length(ttaken_total)
        if floor(ttaken_total(w)/3600)==0
            if floor(mod(ttaken_total(w),3600)/60)==0
                if floor(mod(ttaken_total(w),60))==0
                    labels1{w,1} = sprintf('%.0fms',floor(mod(ttaken_total(w),1)*1000));
                else
                    labels1{w,1} = sprintf('%.0fs',floor(mod(ttaken_total(w),60)));
                end
            else
                labels1{w,1} = sprintf('%.0fm:%.0fs',floor(mod(ttaken_total(w),3600)/60),floor(mod(ttaken_total(w),60)));
            end
        else
            labels1{w,1} = sprintf('%.0fh:%.0fm:%.0fs',floor(ttaken_total(w)/3600),floor(mod(ttaken_total(w),3600)/60),floor(mod(ttaken_total(w),60)));
        end
    end
    labels1 = string(labels1); clear b
    text(xtips1,ytips1,labels1,'HorizontalAlignment','center','VerticalAlignment','bottom','fontsize',12','color',[0.65,0.11,0.19]);
    try xlabel('Pre-processing steps'), ylabel('time taken (s)'), ylim([0 1.2*max(ttaken_total)]), catch, end
    title(sprintf('(Total execution time, including all steps and subjects = %.0fh:%.0fm:%.0fs)\n\nTOTAL TIME TAKEN for each pre-processing step (%d subjects)',floor(sum(ttaken_total)/3600),floor(mod(sum(ttaken_total),3600)/60),floor(mod(sum(ttaken_total),60)),length(sub_dir)),'Interpreter', 'none', 'Color',[0 0.37 0.22],'FontSize',16);
    clear labels1 xtips1 ytips1
subplot2n(3,1,2), b = bar(ttaken_step,0.5,'FaceColor',[0,0.65,0.69],'EdgeColor',[0.65,0.11,0.19],'LineWidth',1.5);
    xticks([1:length(ttaken_step)]), xticklabels(mat2cell(find(choices_preproc==1),length(find(choices_preproc==1))))
	xtips1 = b(1).XEndPoints; ytips1 = b(1).YEndPoints;
    for w=1:length(ttaken_step)
        if floor(ttaken_step(w)/3600)==0
            if floor(mod(ttaken_step(w),3600)/60)==0
                if floor(mod(ttaken_step(w),60))==0
                    labels1{w,1} = sprintf('%.0fms',floor(mod(ttaken_step(w),1)*1000));
                else
                    labels1{w,1} = sprintf('%.0fs',floor(mod(ttaken_step(w),60)));
                end
            else
                labels1{w,1} = sprintf('%.0fm:%.0fs',floor(mod(ttaken_step(w),3600)/60),floor(mod(ttaken_step(w),60)));
            end
        else
            labels1{w,1} = sprintf('%.0fh:%.0fm:%.0fs',floor(ttaken_step(w)/3600),floor(mod(ttaken_step(w),3600)/60),floor(mod(ttaken_step(w),60)));
        end
    end
    labels1 = string(labels1); clear b
    text(xtips1,ytips1,labels1,'HorizontalAlignment','center','VerticalAlignment','bottom','fontsize',12','color',[0.65,0.11,0.19]);
    try xlabel('Pre-processing steps'), ylabel('time taken (s)'), ylim([0 1.2*max(ttaken_step)]), catch, end
    title(sprintf('AVERAGE TIME TAKEN for each pre-processing step per subject'),'Interpreter', 'none', 'Color',[0 0.37 0.22],'FontSize',16);
    clear labels1 xtips1 ytips1
subplot2n(3,1,3), b = bar(ttaken_sub,0.5,'FaceColor',[0,0.75,0.69],'EdgeColor',[0.65,0.11,0.19],'LineWidth',1.5);
	xtips1 = b(1).XEndPoints; ytips1 = b(1).YEndPoints;
    for w=1:length(ttaken_sub)
        if floor(ttaken_sub(w)/3600)==0
            if floor(mod(ttaken_sub(w),3600)/60)==0
                if floor(mod(ttaken_sub(w),60))==0
                    labels1{w,1} = sprintf('%.0fms',floor(mod(ttaken_sub(w),1)*1000));
                else
                    labels1{w,1} = sprintf('%.0fs',floor(mod(ttaken_sub(w),60)));
                end
            else
                labels1{w,1} = sprintf('%.0fm:%.0fs',floor(mod(ttaken_sub(w),3600)/60),floor(mod(ttaken_sub(w),60)));
            end
        else
            labels1{w,1} = sprintf('%.0fh:%.0fm:%.0fs',floor(ttaken_sub(w)/3600),floor(mod(ttaken_sub(w),3600)/60),floor(mod(ttaken_sub(w),60)));
        end
    end
    labels1 = string(labels1); clear b
    text(xtips1,ytips1,labels1,'HorizontalAlignment','center','VerticalAlignment','bottom','fontsize',12','color',[0.65,0.11,0.19]);
    try xlabel('Subjects'), ylabel('time taken (s)'), ylim([0 1.2*max(ttaken_sub)]), catch, end
    title(sprintf('TOTAL TIME TAKEN for each SUBJECT'),'Interpreter', 'none', 'Color',[0 0.37 0.22],'FontSize',16);
saveas(fighndl,[home_dir 'ExecutionTime_graph' '.jpg'])
tmp=clock; clk_settings = sprintf('%.4d-%.2d-%.2d_%.2d-%.2d',tmp(1),tmp(2),tmp(3),tmp(4),tmp(5)); clear tmp
saveas(fighndl,[home_dir sprintf('ExecutionTime_graph_%s',clk_settings) '.jpg'])
close(fighndl)
catch
end

try
filename_ExecutionTime = sprintf('ExecutionTime_%s.mat',clk_settings);
readme_time_taken = sprintf('Time taken for different processing steps:\ntime_taken_preproc = raw measurements of time taken for pre-processing\nttaken_total = total execution time, including all steps and subjects\nttaken_step = average time taken for each pre-processing step per subject\nttaken_sub = total time taken for each subject\n');
save([home_dir filename_ExecutionTime],'ttaken_total','ttaken_step','ttaken_sub','time_taken_preproc','readme_time_taken')
clear labels1 xtips1 ytips1 clk_settings readme_time_taken
catch
end

%%
L=L+2; logg{L,1}=sprintf('--- Pre-processing Completed ---'); try fprintf(fileID,'%s\n',logg{L,1}); set(S.tx,'string',logg); catch, end; clear clk
try close(wbar2), catch, end
try close(wbar3), catch, end
refresh_html_log

quest = ['\color[rgb]{0.65,0.11,0.19}' sprintf('Dear User,\n\nAll pre-processing steps were COMPLETED as promised.\n\nRegards,\nSpinal cord fMRI toolbox (NEPTUNE)\n')]; %#ok<*NASGU>
opts.Interpreter = 'tex'; opts.WindowStyle = 'non-modal';
msgbox(quest,'All pre-processing done!','custom',msgicon,[1,1,1],opts); clear quest opts

try close(status_fig), catch, end
clear status_fig wbar2 wbar3

%%
```
