## Supplementary material for "Neptune: a toolbox for spinal cord functional MRI data processing and quality assurance": Neptune software: preprocdialog.m

```
function [selection,choices] = preprocdialog(pnl,bg,prevchoices,flag,exist_SCT)

qSPM = which('spm_get_bf.m');

cbx1  = uicheckbox(pnl,'Position',[10 412 410 15], 'Value',prevchoices(1));
cbx2  = uicheckbox(pnl,'Position',[10 390 410 15], 'Value',prevchoices(2));
cbx3  = uicheckbox(pnl,'Position',[10 368 410 15], 'Value',prevchoices(3));
% cbx4  = uicheckbox(pnl,'Position',[10 346 410 15], 'Enable','off', 'Value',prevchoices(4));
cbx5  = uicheckbox(pnl,'Position',[10 324 410 15], 'Value',prevchoices(5));
cbx6  = uicheckbox(pnl,'Position',[10 302 410 15], 'Value',prevchoices(6));
cbx7  = uicheckbox(pnl,'Position',[10 280 410 15], 'Value',prevchoices(7));
cbx8  = uicheckbox(pnl,'Position',[10 258 410 15], 'Value',prevchoices(8));
cbx9  = uicheckbox(pnl,'Position',[10 236 410 15], 'Value',prevchoices(9));
cbx10 = uicheckbox(pnl,'Position',[10 214 410 15], 'Value',prevchoices(10));
cbx11 = uicheckbox(pnl,'Position',[10 192 410 15], 'Value',prevchoices(11));
cbx12 = uicheckbox(pnl,'Position',[10 170 410 15], 'Value',prevchoices(12));
cbx13 = uicheckbox(pnl,'Position',[10 148 410 15], 'Value',prevchoices(13));
cbx14 = uicheckbox(pnl,'Position',[10 126 410 15], 'Value',prevchoices(14));
cbx15 = uicheckbox(pnl,'Position',[10 104 410 15], 'Value',prevchoices(15));
cbx16 = uicheckbox(pnl,'Position',[10  82 410 15], 'Value',prevchoices(16));
if ~isempty(qSPM)
cbx17 = uicheckbox(pnl,'Position',[10  60 410 15], 'Value',prevchoices(17));
else
% cbx17 = uicheckbox(pnl,'Position',[10  60 410 15], 'Enable','off', 'Value',prevchoices(17));
end
if exist_SCT==1
cbx18 = uicheckbox(pnl,'Position',[10  38 410 15], 'Value',prevchoices(18));
else
% cbx18 = uicheckbox(pnl,'Position',[10  38 410 15], 'Enable','off', 'Value',prevchoices(18));
end
cbx19 = uicheckbox(pnl,'Position',[10  16 410 15], 'Value',prevchoices(19));

rb1 = uiradiobutton(bg,'Position',[400 10 410 15]);
rb2 = uiradiobutton(bg,'Position',[200 10 410 15]);
rb3 = uiradiobutton(bg,'Position',[400 10 410 15]);

cbx1.Text  = '01. NORDIC denoising';
cbx2.Text  = '02. Slice-timing correction';
cbx3.Text  = '03. Match anatomical and functional slices';
% cbx4.Text  = '04. Split data into slices (mandatory)';
cbx5.Text  = '05. Define "not cord" mask (mandatory)';
cbx6.Text  = '06. Denoise-1 (non-cord signal regression)';
cbx7.Text  = '07. Compute Gaussian mask (mandatory)';
cbx8.Text  = '08. Motion correction (mandatory)';
cbx9.Text  = '09. Denoise-2 (RETROICOR/3dDespike)';
cbx10.Text = '10. Define GM, WM and CSF masks (mandatory)';
cbx11.Text = '11. Co-registration (functional to anatomical) (mandatory)';
cbx12.Text = '12. Denoise-3 (cerebrospinal fluid signal regression)';
cbx13.Text = '13. Denoise-4 (white-matter signal regression)';
cbx14.Text = '14. Denoise-5 (additional covariates regression)';
cbx15.Text = '15. Temporal filtering';
cbx16.Text = '16. Define spinal cord GM and WM quadrants';
if ~isempty(qSPM)
cbx17.Text = '17. HRF deconvolution (preferred for connectivity studies only)';
else
% cbx17.Text = '17. HRF deconvolution (requires SPM, which is not installed)';
end
if exist_SCT==1
cbx18.Text = '18. Co-registration to PAM50 standard space';
else
% cbx18.Text = '18. Co-registration to PAM50 standard space (requires SCT)';
end
cbx19.Text = '19. gzip NIfTI files to save disk space';

rb1.Text = '';
rb2.Text = 'DONE';
rb3.Text = 'Help';

pause(2)
if ~isempty(qSPM) && exist_SCT==1
choices = [cbx1.Value;cbx2.Value;cbx3.Value;1;cbx5.Value;cbx6.Value;cbx7.Value;cbx8.Value;cbx9.Value;cbx10.Value;cbx11.Value;cbx12.Value;cbx13.Value;cbx14.Value;cbx15.Value;cbx16.Value;cbx17.Value;cbx18.Value;cbx19.Value];
elseif isempty(qSPM) && exist_SCT==1
choices = [cbx1.Value;cbx2.Value;cbx3.Value;1;cbx5.Value;cbx6.Value;cbx7.Value;cbx8.Value;cbx9.Value;cbx10.Value;cbx11.Value;cbx12.Value;cbx13.Value;cbx14.Value;cbx15.Value;cbx16.Value;0;cbx18.Value;cbx19.Value];
elseif ~isempty(qSPM) && exist_SCT==0
choices = [cbx1.Value;cbx2.Value;cbx3.Value;1;cbx5.Value;cbx6.Value;cbx7.Value;cbx8.Value;cbx9.Value;cbx10.Value;cbx11.Value;cbx12.Value;cbx13.Value;cbx14.Value;cbx15.Value;cbx16.Value;cbx17.Value;0;cbx19.Value];
elseif isempty(qSPM) && exist_SCT==0
choices = [cbx1.Value;cbx2.Value;cbx3.Value;1;cbx5.Value;cbx6.Value;cbx7.Value;cbx8.Value;cbx9.Value;cbx10.Value;cbx11.Value;cbx12.Value;cbx13.Value;cbx14.Value;cbx15.Value;cbx16.Value;0;0;cbx19.Value];
end

selection = bg.SelectedObject.Parent.SelectedObject.Text;

if flag==1 && strcmp(selection,'Help')==1
    selection='';
end

end
```
