## Supplementary material for "Neptune: a toolbox for spinal cord functional MRI data processing and quality assurance": Neptune software: refresh_html.m

```
if ~(exist('home_dir','var'))
    home_dir = erase(base_dir_sub,base_dir_sub(max(strfind(base_dir_sub(1:end-1),'/'))+1:end));
    if isempty(home_dir)
        home_dir = erase(base_dir_sub,base_dir_sub(max(strfind(base_dir_sub(1:end-1),'\'))+1:end));
    end
end
if ~(exist('neptune_dir','var'))
    tmp = which('neptune'); neptune_dir = tmp(1:end-9); clear tmp
end

%% report_neptune.html

html0.outdir = home_dir;
html0.savefile = [neptune_dir 'report_neptune.m'];
set(0,'DefaultFigureVisible','off')
html0.htmlFile = publish(html0.savefile,'format','html','outputDir',html0.outdir,'figureSnapMethod','entireFigureWindow','showCode',false);
set(0,'DefaultFigureVisible','on')

% Read and manipulate text
html0.fileIdRead  = fopen([home_dir 'report_neptune.html'], 'r');
html0.fileTextNew = strrep(fscanf(html0.fileIdRead,'%c'), '&lt;', '<');
html0.fileTextNew = strrep(html0.fileTextNew, '&gt;', '>');
html0.fileTextNew2 = html0.fileTextNew(1:strfind(html0.fileTextNew,'##### SOURCE BEGIN #####'));
if isempty(strfind(html0.fileTextNew2,'pending execution'))
    html0.headingstyle = sprintf('</style></head><body><br><center><img src="%sstatus_images/barrygp.jpg" style="width:360px;height:126px;">&nbsp;&nbsp;&nbsp;&nbsp;<img src="%sstatus_images/neptune_v1.0.jpg" style="width:350px;height:126px;"><p style="font-size:20px ; color:#A61C30"><b>NEPTUNE execution outputs and quality control</p><p style="font-size:17px ; color:#00A000">COMPLETED!</b></p></center>',neptune_dir,neptune_dir);
else
    html0.headingstyle = sprintf('</style></head><body><br><center><img src="%sstatus_images/barrygp.jpg" style="width:360px;height:126px;">&nbsp;&nbsp;&nbsp;&nbsp;<img src="%sstatus_images/neptune_v1.0.jpg" style="width:350px;height:126px;"><p style="font-size:20px ; color:#A61C30"><b>NEPTUNE execution outputs and quality control</p><p style="font-size:17px ; color:#FF0000">INCOMPLETE...</b></p></center>',neptune_dir,neptune_dir);
end
html0.fileTextNew = strrep(html0.fileTextNew, '</style></head><body>', html0.headingstyle);
html0.fileTextNew = strrep(html0.fileTextNew, '<h1></h1><!--introduction--><!--/introduction-->', '');
html0.fileTextNew = strrep(html0.fileTextNew, 'font-size:10px', 'font-size:12px');
html0.fileTextNew = strrep(html0.fileTextNew, 'line-height:140%', 'line-height:150%');
html0.fileTextNew = strrep(html0.fileTextNew, '<a href="https://www.mathworks.com/products/matlab/">Published with', 'Generated by Neptune (creators: Ranga Deshpande and Robert L Barry @A.A.Martinos Center for Biomedical Imaging, Massachusetts General Hospital, Harvard Medical School and Harvard-MIT HST). <a>Generated using');
html0.fileTextNew = strrep(html0.fileTextNew, 'background:#fff none', 'background:#F2E6D9');
html0.fileTextNew = strrep(html0.fileTextNew, '<h2 ', '<h2 style="color:#008CB0;" ');
html0.fileTextNew = strrep(html0.fileTextNew, '#d6d4d4', '#A0A0A0');
html0.fileTextNew = strrep(html0.fileTextNew, 'line-height:150%; background:#F2E6D9', 'line-height:100%; background:#F2E6D9');
html0.fileTextNew = strrep(html0.fileTextNew, '.content { font-size:1.2em; line-height:150%; padding: 20px; }', '.content { font-size:1.2em; line-height:140%; padding: 20px; }');
fclose(html0.fileIdRead);
% Write Text
html0.fileIdWrite = fopen([home_dir 'report_neptune.html'], 'w');
fprintf(html0.fileIdWrite, '%c', html0.fileTextNew);
fclose(html0.fileIdWrite);
clear html0*

%% detailed log (report_neptune_log.html)
html0.outdir = home_dir;
html0.savefile = [neptune_dir 'report_neptune_log.m'];
set(0,'DefaultFigureVisible','off')
html0.htmlFile = publish(html0.savefile,'format','html','outputDir',html0.outdir,'figureSnapMethod','entireFigureWindow','showCode',false);
set(0,'DefaultFigureVisible','on')

% Read and manipulate text
html0.fileIdRead  = fopen([home_dir 'report_neptune_log.html'], 'r');
html0.fileTextNew = strrep(fscanf(html0.fileIdRead,'%c'), '&lt;', '<');
html0.fileTextNew = strrep(html0.fileTextNew, '&gt;', '>');
html0.fileTextNew2 = html0.fileTextNew(1:strfind(html0.fileTextNew,'##### SOURCE BEGIN #####'));
if isempty(strfind(html0.fileTextNew2,'everything COMPLETED as promised'))
    html0.headingstyle = sprintf('</style></head><body><br><center><img src="%sstatus_images/barrygp.jpg" style="width:360px;height:126px;">&nbsp;&nbsp;&nbsp;&nbsp;<img src="%sstatus_images/neptune_v1.0.jpg" style="width:146px;height:126px;"><p style="font-size:20px ; color:#A61C30"><b>NEPTUNE detailed execution log</p><p style="font-size:17px ; color:#00A000">COMPLETED!</b></p></center>',neptune_dir,neptune_dir);
else
    html0.headingstyle = sprintf('</style></head><body><br><center><img src="%sstatus_images/barrygp.jpg" style="width:360px;height:126px;">&nbsp;&nbsp;&nbsp;&nbsp;<img src="%sstatus_images/neptune_v1.0.jpg" style="width:146px;height:126px;"><p style="font-size:20px ; color:#A61C30"><b>NEPTUNE detailed execution log</p><p style="font-size:17px ; color:#FF0000">INCOMPLETE...</b></p></center>',neptune_dir,neptune_dir);
end
html0.fileTextNew = strrep(html0.fileTextNew, '</style></head><body>', html0.headingstyle);
html0.fileTextNew = strrep(html0.fileTextNew, '<h1></h1><!--introduction--><!--/introduction-->', '');
html0.fileTextNew = strrep(html0.fileTextNew, 'font-size:10px', 'font-size:12px');
html0.fileTextNew = strrep(html0.fileTextNew, 'line-height:140%', 'line-height:150%');
html0.fileTextNew = strrep(html0.fileTextNew, '<a href="https://www.mathworks.com/products/matlab/">Published with', 'Generated by Neptune (creators: Ranga Deshpande and Robert L Barry @A.A.Martinos Center for Biomedical Imaging, Massachusetts General Hospital, Harvard Medical School and Harvard-MIT HST). <a>Generated using');
html0.fileTextNew = strrep(html0.fileTextNew, 'background:#fff none', 'background:#F2E6D9');
html0.fileTextNew = strrep(html0.fileTextNew, '<h2 ', '<h2 style="color:#008CB0;" ');
html0.fileTextNew = strrep(html0.fileTextNew, '#d6d4d4', '#A0A0A0');
html0.fileTextNew = strrep(html0.fileTextNew, 'line-height:150%; background:#F2E6D9', 'line-height:100%; background:#F2E6D9');
html0.fileTextNew = strrep(html0.fileTextNew, '.content { font-size:1.2em; line-height:150%; padding: 20px; }', '.content { font-size:1.2em; line-height:140%; padding: 20px; }');
fclose(html0.fileIdRead);
% Write Text
html0.fileIdWrite = fopen([home_dir 'report_neptune_log.html'], 'w');
fprintf(html0.fileIdWrite, '%c', html0.fileTextNew);
fclose(html0.fileIdWrite);
clear html0*
```
