## Supplementary material for "Neptune: a toolbox for spinal cord functional MRI data processing and quality assurance": Neptune software: refresh_html_manual.m

```
% Use this file to generate the webpage (report_neptune.html)
% using manually entered/chosen subject folders and choices


% Step-1: define the following variables or load it onto the workspace manually
% (the fields populated below are for reference only
home_dir = '/home_dir'; % full path to the directory where your subject folders exist
sub_dir = {'subject_dir'};  % full path to all subjects' directories that you are interested in generating the HTML
runs = [1;1]; % number of functional runs in each subject
slices = 12; % number of slices (assumes same value for all subjects) (if different subs have different slices, enter the maximum value)
choices_preproc = [0;0;1;1;1;1;1;1;1;1;1;1;1;1;1;1;1;0;0]; % 19x1 vector of pre-processing choices made by you
save_videos_steps = [0;0;0;0;0;0;0;0]; % optional input (defaults to all zeros if missing). Set to all zeros if QC videos were not saved

% If you want to load the above variables directly from a previous NEPTUNE run, 
% OR if you want a starting point to generate these variables, then
% load one of your saved neptune_settings_* files. Example below:
load('neptune_settings_2021-01-01_12-00.mat', 'home_dir', 'sub_dir', 'runs', 'slices', 'choices_preproc', 'save_videos_steps')
tmp=clock; clk_init = sprintf('%.4d-%.2d-%.2d_%.2d-%.2d',tmp(1),tmp(2),tmp(3),tmp(4),tmp(5)); clear tmp
save(sprintf('settings_for_report_html_manual_%s.mat',clk_init), 'sub_dir', 'choices_preproc', 'home_dir', 'runs', 'save_videos_steps', 'slices', 'clk_init')


% Step-2: once the above variables are ready, execute the following lines
refresh_html_all
delete([home_dir 'report_neptune.html'])
movefile([home_dir 'report_neptune_all.html'],[home_dir 'report_neptune.html'],'f')

movefile('report_neptune.html',sprintf('report_neptune_%s.html',clk_init))
web(sprintf('report_neptune_%s.html',clk_init),'-browser')


%%
```
