## Supplementary material for "Neptune: a toolbox for spinal cord functional MRI data processing and quality assurance": Neptune software: regresscovdialog.m

```
function [selection1,selection2,selection3,selection4] = regresscovdialog(bg1,bg2,bg3,bg4,flag,selection1,selection2,selection3,selection4)

rb1 = uiradiobutton(bg1,'Position',[10 15 940 15],'Value',strcmp(selection1,'Motion parameters (2 translation)'));
rb2 = uiradiobutton(bg1,'Position',[310 15 940 15],'Value',strcmp(selection1,'Motion parameters (2 translation + 2 derivatives)'));
rb3 = uiradiobutton(bg1,'Position',[620 15 940 15],'Value',strcmp(selection1,'NO motion parameters'));

rb4 = uiradiobutton(bg2,'Position',[10 15 940 15],'Value',strcmp(selection2,'Scrubbing (volumes with motion > half voxel size)'));
rb5 = uiradiobutton(bg2,'Position',[310 15 940 15],'Value',strcmp(selection2,'Scrubbing (user-defined motion threshold)'));
rb6 = uiradiobutton(bg2,'Position',[620 15 940 15],'Value',strcmp(selection2,'NO scrubbing using motion'));

rb7 = uiradiobutton(bg3,'Position',[10 15 940 15],'Value',strcmp(selection3,'Regression with user-defined covariate(s)'));
rb8 = uiradiobutton(bg3,'Position',[310 15 940 15],'Value',strcmp(selection3,'NO user-defined covariates'));

rb10 = uiradiobutton(bg4,'Position',[825 15 940 15]);
rb11 = uiradiobutton(bg4,'Position',[425 15 940 15],'Value',strcmp(selection4,'DONE'));
rb12 = uiradiobutton(bg4,'Position',[825 15 940 15],'Value',strcmp(selection4,'Help'));

rb1.Text = 'Motion parameters (2 translation)';
rb2.Text = 'Motion parameters (2 translation + 2 derivatives)';
rb3.Text = 'NO motion parameters';
rb4.Text = 'Scrubbing (volumes with motion > half voxel size)';
rb5.Text = 'Scrubbing (user-defined motion threshold)';
rb6.Text = 'NO scrubbing using motion';
rb7.Text = 'Regression with user-defined covariate(s)';
rb8.Text = 'NO user-defined covariates';
rb10.Text = '';
rb11.Text = 'DONE';
rb12.Text = 'Help';

pause(2)

selection1 = bg1.SelectedObject.Parent.SelectedObject.Text;
selection2 = bg2.SelectedObject.Parent.SelectedObject.Text;
selection3 = bg3.SelectedObject.Parent.SelectedObject.Text;
selection4 = bg4.SelectedObject.Parent.SelectedObject.Text;

if flag==1 && strcmp(selection4,'Help')==1
    selection4='';
end

end
```
