## Supplementary material for "Neptune: a toolbox for spinal cord functional MRI data processing and quality assurance": Neptune software: report_neptune_all.m

```
clk=clock;
month={'Jan','Feb','Mar','Apr','May','Jun','Jul','Aug','Sep','Oct','Nov','Dec'}';
fprintf('<p style="font-size:14px ; color:#000000">')
disp(sprintf('Home directory: <i>%s</i>',home_dir(1:end-1)))
disp(sprintf('Page generation date/time: %.2d/%s/%.4d %.2dh:%.2dm:%.2ds',clk(3),month{clk(2)},clk(1),clk(4),clk(5),round(clk(6))))
fprintf('</p>')
clear clk
if ~exist('save_videos_steps','var'), save_videos_steps=zeros(8,1); html1.saveVIDflag=1; else, html1.saveVIDflag=0; end
if ~exist('WM_flag','var'), WM_flag=1; html1.saveWMflag=1; else, html1.saveWMflag=0; end
if ~exist('SFC_type','var'), SFC_type=3; html1.saveSFC1flag=1; else, html1.saveSFC1flag=0; end
if ~exist('SFC_withinslice_flag','var'), SFC_withinslice_flag=1; html1.saveSFC2flag=1; else, html1.saveSFC2flag=0; end
if ~exist('SFC_betweenslice_flag','var'), SFC_betweenslice_flag=1; html1.saveSFC3flag=1; else, html1.saveSFC3flag=0; end
if ~exist('dont_do_pcor_betweenslice','var'), dont_do_pcor_betweenslice=0; html1.saveSFC4flag=1; else, html1.saveSFC4flag=0; end

%% Shortlist of QC outputs
fprintf('<p style="font-size:14px ; color:#000000">')
fprintf('<b>00.1. Time series within the spinal cord after:</b>\n')
for html1k=1:length(sub_dir)
    for html1k2=1:runs(html1k)
        if runs(html1k)==1
            html1.QCdir = ['.' erase(sub_dir{html1k},home_dir(1:end-1)) 'QC/' 'QC_shortlist/'];
        else
            html1.QCdir = ['.' erase(sub_dir{html1k},home_dir(1:end-1)) 'QC/' 'QC_shortlist/' 'run' num2str(html1k2) '/'];
        end
        html1.slicesstr = '';
        if exist([home_dir html1.QCdir(3:end) 'fMRI_ts_01nordic_allSlices.jpg'],'file')
            html1.slicesstr = strcat(html1.slicesstr,sprintf(' <a href="%s">01-NORDIC</a>,',[html1.QCdir 'fMRI_ts_01nordic_allSlices.jpg']));
        else
            html1.slicesstr = strcat(html1.slicesstr,sprintf(' 01-NORDIC,'));
        end
        if exist([home_dir html1.QCdir(3:end) 'fMRI_ts_04rawdata_allSlices.jpg'],'file')
            html1.slicesstr = strcat(html1.slicesstr,sprintf(' <a href="%s">04-RawData</a>,',[html1.QCdir 'fMRI_ts_04rawdata_allSlices.jpg']));
        else
            html1.slicesstr = strcat(html1.slicesstr,sprintf(' 04-RawData,'));
        end
        if exist([home_dir html1.QCdir(3:end) 'fMRI_ts_06denoise1_allSlices.jpg'],'file')
            html1.slicesstr = strcat(html1.slicesstr,sprintf(' <a href="%s">06-denoise-1-notCord</a>,',[html1.QCdir 'fMRI_ts_06denoise1_allSlices.jpg']));
        else
            html1.slicesstr = strcat(html1.slicesstr,sprintf(' 06-denoise-1-notCord,'));
        end
        if exist([home_dir html1.QCdir(3:end) 'fMRI_ts_08moco_allSlices.jpg'],'file')
            html1.slicesstr = strcat(html1.slicesstr,sprintf(' <a href="%s">08-MoCo</a>,',[html1.QCdir 'fMRI_ts_08moco_allSlices.jpg']));
        else
            html1.slicesstr = strcat(html1.slicesstr,sprintf(' 08-MoCo,'));
        end
        if exist([home_dir html1.QCdir(3:end) 'fMRI_ts_09ricor_allSlices.jpg'],'file')
            html1.slicesstr = strcat(html1.slicesstr,sprintf(' <a href="%s">09-denoise-2-RETROICOR</a>,',[html1.QCdir 'fMRI_ts_09ricor_allSlices.jpg']));
        else
            html1.slicesstr = strcat(html1.slicesstr,sprintf(' 09-denoise-2-RETROICOR,'));
        end
        if exist([home_dir html1.QCdir(3:end) 'fMRI_ts_11coreg_allSlices.jpg'],'file')
            html1.slicesstr = strcat(html1.slicesstr,sprintf(' <a href="%s">11-coregistration</a>,',[html1.QCdir 'fMRI_ts_11coreg_allSlices.jpg']));
        else
            html1.slicesstr = strcat(html1.slicesstr,sprintf(' 11-coregistration,'));
        end
        if exist([home_dir html1.QCdir(3:end) 'fMRI_ts_12denoise3_allSlices.jpg'],'file')
            html1.slicesstr = strcat(html1.slicesstr,sprintf(' <a href="%s">12-denoise-3-CSF</a>,',[html1.QCdir 'fMRI_ts_12denoise3_allSlices.jpg']));
        else
            html1.slicesstr = strcat(html1.slicesstr,sprintf(' 12-denoise-3-CSF,'));
        end
        if exist([home_dir html1.QCdir(3:end) 'fMRI_ts_13denoise4_allSlices.jpg'],'file')
            html1.slicesstr = strcat(html1.slicesstr,sprintf(' <a href="%s">13-denoise-4-WM</a>,',[html1.QCdir 'fMRI_ts_13denoise4_allSlices.jpg']));
        else
            html1.slicesstr = strcat(html1.slicesstr,sprintf(' 13-denoise-4-WM,'));
        end
        if exist([home_dir html1.QCdir(3:end) 'fMRI_ts_14denoise5_allSlices.jpg'],'file')
            html1.slicesstr = strcat(html1.slicesstr,sprintf(' <a href="%s">14-denoise-5-covariates</a>,',[html1.QCdir 'fMRI_ts_14denoise5_allSlices.jpg']));
        else
            html1.slicesstr = strcat(html1.slicesstr,sprintf(' 14-denoise-5-covariates,'));
        end
        if exist([home_dir html1.QCdir(3:end) 'fMRI_ts_15filter_allSlices.jpg'],'file')
            html1.slicesstr = strcat(html1.slicesstr,sprintf(' <a href="%s">15-filtering</a>,',[html1.QCdir 'fMRI_ts_15filter_allSlices.jpg']));
        else
            html1.slicesstr = strcat(html1.slicesstr,sprintf(' 15-filtering,'));
        end
        if exist([home_dir html1.QCdir(3:end) 'fMRI_ts_17deconv_allSlices.jpg'],'file')
            html1.slicesstr = strcat(html1.slicesstr,sprintf(' <a href="%s">17-deconvolution</a>,',[html1.QCdir 'fMRI_ts_17deconv_allSlices.jpg']));
        else
            html1.slicesstr = strcat(html1.slicesstr,sprintf(' 17-deconvolution,'));
        end
        if exist([home_dir html1.QCdir(3:end) 'fMRI_ts_20quadTS_allSlices.jpg'],'file')
            html1.slicesstr = strcat(html1.slicesstr,sprintf(' <a href="%s">20-quadTimeseries</a>,',[html1.QCdir 'fMRI_ts_20quadTS_allSlices.jpg']));
        else
            html1.slicesstr = strcat(html1.slicesstr,sprintf(' 20-quadTimeseries,'));
        end
        
        if (html1k2==1) && (runs(html1k)==1)
            disp(sprintf('      Subject %d of %d - %s  %s',html1k,length(sub_dir),subnames{html1k},html1.slicesstr(2:end-1)))
        elseif (html1k2==1) && (runs(html1k)~=1)
            disp(sprintf('      Subject %d of %d, run %d of %d - %s  %s',html1k,length(sub_dir),html1k2,runs(html1k),subnames{html1k},html1.slicesstr(2:end-1)))
        else
            disp(sprintf('                      run %d of %d - %s  %s',html1k2,runs(html1k),subnames{html1k},html1.slicesstr(2:end-1)))
        end
        clear html1.QCdir html1.slicesstr html1k3
    end
end

fprintf('\n')
fprintf('<b>00.2. Global mean signal within the spinal cord after:</b>\n')
for html1k=1:length(sub_dir)
    for html1k2=1:runs(html1k)
        if runs(html1k)==1
            html1.QCdir = ['.' erase(sub_dir{html1k},home_dir(1:end-1)) 'QC/' 'QC_shortlist/'];
        else
            html1.QCdir = ['.' erase(sub_dir{html1k},home_dir(1:end-1)) 'QC/' 'QC_shortlist/' 'run' num2str(html1k2) '/'];
        end
        html1.slicesstr = '';
        if exist([home_dir html1.QCdir(3:end) 'global_signal_01nordic_allSlices.jpg'],'file')
            html1.slicesstr = strcat(html1.slicesstr,sprintf(' <a href="%s">01-NORDIC</a>,',[html1.QCdir 'global_signal_01nordic_allSlices.jpg']));
        else
            html1.slicesstr = strcat(html1.slicesstr,sprintf(' 01-NORDIC,'));
        end
        if exist([home_dir html1.QCdir(3:end) 'global_signal_04rawdata_allSlices.jpg'],'file')
            html1.slicesstr = strcat(html1.slicesstr,sprintf(' <a href="%s">04-RawData</a>,',[html1.QCdir 'global_signal_04rawdata_allSlices.jpg']));
        else
            html1.slicesstr = strcat(html1.slicesstr,sprintf(' 04-RawData,'));
        end
        if exist([home_dir html1.QCdir(3:end) 'global_signal_06denoise1_allSlices.jpg'],'file')
            html1.slicesstr = strcat(html1.slicesstr,sprintf(' <a href="%s">06-denoise-1-notCord</a>,',[html1.QCdir 'global_signal_06denoise1_allSlices.jpg']));
        else
            html1.slicesstr = strcat(html1.slicesstr,sprintf(' 06-denoise-1-notCord,'));
        end
        if exist([home_dir html1.QCdir(3:end) 'global_signal_06denoise1_vs_08moco_allSlices.jpg'],'file')
            html1.slicesstr = strcat(html1.slicesstr,sprintf(' <a href="%s">08-MoCo</a>,',[html1.QCdir 'global_signal_06denoise1_vs_08moco_allSlices.jpg']));
        else
            html1.slicesstr = strcat(html1.slicesstr,sprintf(' 08-MoCo,'));
        end
        if exist([home_dir html1.QCdir(3:end) 'global_signal_08moco_vs_09ricor_allSlices.jpg'],'file')
            html1.slicesstr = strcat(html1.slicesstr,sprintf(' <a href="%s">09-denoise-2-RETROICOR</a>,',[html1.QCdir 'global_signal_08moco_vs_09ricor_allSlices.jpg']));
        else
            html1.slicesstr = strcat(html1.slicesstr,sprintf(' 09-denoise-2-RETROICOR,'));
        end
        if exist([home_dir html1.QCdir(3:end) 'global_signal_08moco_vs_11coreg_allSlices.jpg'],'file')
            html1.slicesstr = strcat(html1.slicesstr,sprintf(' <a href="%s">11-coregistration</a>,',[html1.QCdir 'global_signal_08moco_vs_11coreg_allSlices.jpg']));
        else
            html1.slicesstr = strcat(html1.slicesstr,sprintf(' 11-coregistration,'));
        end
        if exist([home_dir html1.QCdir(3:end) 'global_signal_11coreg_vs_12denoise3_allSlices.jpg'],'file')
            html1.slicesstr = strcat(html1.slicesstr,sprintf(' <a href="%s">12-denoise-3-CSF</a>,',[html1.QCdir 'global_signal_11coreg_vs_12denoise3_allSlices.jpg']));
        else
            html1.slicesstr = strcat(html1.slicesstr,sprintf(' 12-denoise-3-CSF,'));
        end
        if exist([home_dir html1.QCdir(3:end) 'global_signal_12denoise3_vs_13denoise4_allSlices.jpg'],'file')
            html1.slicesstr = strcat(html1.slicesstr,sprintf(' <a href="%s">13-denoise-4-WM</a>,',[html1.QCdir 'global_signal_12denoise3_vs_13denoise4_allSlices.jpg']));
        else
            html1.slicesstr = strcat(html1.slicesstr,sprintf(' 13-denoise-4-WM,'));
        end
        if exist([home_dir html1.QCdir(3:end) 'global_signal_13denoise4_vs_14denoise5_allSlices.jpg'],'file')
            html1.slicesstr = strcat(html1.slicesstr,sprintf(' <a href="%s">14-denoise-5-covariates</a>,',[html1.QCdir 'global_signal_13denoise4_vs_14denoise5_allSlices.jpg']));
        else
            html1.slicesstr = strcat(html1.slicesstr,sprintf(' 14-denoise-5-covariates,'));
        end
        if exist([home_dir html1.QCdir(3:end) 'global_signal_14denoise5_vs_15filter_allSlices.jpg'],'file')
            html1.slicesstr = strcat(html1.slicesstr,sprintf(' <a href="%s">15-filtering</a>,',[html1.QCdir 'global_signal_14denoise5_vs_15filter_allSlices.jpg']));
        else
            html1.slicesstr = strcat(html1.slicesstr,sprintf(' 15-filtering,'));
        end
        if exist([home_dir html1.QCdir(3:end) 'global_signal_15filter_vs_17deconv_allSlices.jpg'],'file')
            html1.slicesstr = strcat(html1.slicesstr,sprintf(' <a href="%s">17-deconvolution</a>,',[html1.QCdir 'global_signal_15filter_vs_17deconv_allSlices.jpg']));
        else
            html1.slicesstr = strcat(html1.slicesstr,sprintf(' 17-deconvolution,'));
        end
        if exist([home_dir html1.QCdir(3:end) 'global_signal_20quadTS_allSlices_GM.jpg'],'file')
            html1.slicesstr = strcat(html1.slicesstr,sprintf(' <a href="%s">20-quadTimeseries</a>,',[html1.QCdir 'global_signal_20quadTS_allSlices_GM.jpg']));
        else
            html1.slicesstr = strcat(html1.slicesstr,sprintf(' 20-quadTimeseries,'));
        end
        
        if (html1k2==1) && (runs(html1k)==1)
            disp(sprintf('      Subject %d of %d - %s  %s',html1k,length(sub_dir),subnames{html1k},html1.slicesstr(2:end-1)))
        elseif (html1k2==1) && (runs(html1k)~=1)
            disp(sprintf('      Subject %d of %d, run %d of %d - %s  %s',html1k,length(sub_dir),html1k2,runs(html1k),subnames{html1k},html1.slicesstr(2:end-1)))
        else
            disp(sprintf('                      run %d of %d - %s  %s',html1k2,runs(html1k),subnames{html1k},html1.slicesstr(2:end-1)))
        end
        clear html1.QCdir html1.slicesstr html1k3
    end
end

fprintf('\n')
fprintf('<b>00.3. Histogram of voxel-to-voxel correlations within the spinal cord after:</b>\n')
for html1k=1:length(sub_dir)
    for html1k2=1:runs(html1k)
        if runs(html1k)==1
            html1.QCdir = ['.' erase(sub_dir{html1k},home_dir(1:end-1)) 'QC/' 'QC_shortlist/'];
        else
            html1.QCdir = ['.' erase(sub_dir{html1k},home_dir(1:end-1)) 'QC/' 'QC_shortlist/' 'run' num2str(html1k2) '/'];
        end
        html1.slicesstr = '';
        if exist([home_dir html1.QCdir(3:end) 'hist_FC_compare_01nordic_allSlices.jpg'],'file')
            html1.slicesstr = strcat(html1.slicesstr,sprintf(' <a href="%s">01-NORDIC</a>,',[html1.QCdir 'hist_FC_compare_01nordic_allSlices.jpg']));
        else
            html1.slicesstr = strcat(html1.slicesstr,sprintf(' 01-NORDIC,'));
        end
        if exist([home_dir html1.QCdir(3:end) 'hist_FC_compare_04rawdata_allSlices.jpg'],'file')
            html1.slicesstr = strcat(html1.slicesstr,sprintf(' <a href="%s">04-RawData</a>,',[html1.QCdir 'hist_FC_compare_04rawdata_allSlices.jpg']));
        else
            html1.slicesstr = strcat(html1.slicesstr,sprintf(' 04-RawData,'));
        end
        if exist([home_dir html1.QCdir(3:end) 'hist_FC_compare_06denoise1_allSlices.jpg'],'file')
            html1.slicesstr = strcat(html1.slicesstr,sprintf(' <a href="%s">06-denoise-1-notCord</a>,',[html1.QCdir 'hist_FC_compare_06denoise1_allSlices.jpg']));
        else
            html1.slicesstr = strcat(html1.slicesstr,sprintf(' 06-denoise-1-notCord,'));
        end
        if exist([home_dir html1.QCdir(3:end) 'hist_FC_compare_06denoise1_08moco_allSlices.jpg'],'file')
            html1.slicesstr = strcat(html1.slicesstr,sprintf(' <a href="%s">08-MoCo</a>,',[html1.QCdir 'hist_FC_compare_06denoise1_08moco_allSlices.jpg']));
        else
            html1.slicesstr = strcat(html1.slicesstr,sprintf(' 08-MoCo,'));
        end
        if exist([home_dir html1.QCdir(3:end) 'hist_FC_compare_08moco_09ricor_allSlices.jpg'],'file')
            html1.slicesstr = strcat(html1.slicesstr,sprintf(' <a href="%s">09-denoise-2-RETROICOR</a>,',[html1.QCdir 'hist_FC_compare_08moco_09ricor_allSlices.jpg']));
        else
            html1.slicesstr = strcat(html1.slicesstr,sprintf(' 09-denoise-2-RETROICOR,'));
        end
        if exist([home_dir html1.QCdir(3:end) 'hist_FC_compare_08moco_11coreg_allSlices.jpg'],'file')
            html1.slicesstr = strcat(html1.slicesstr,sprintf(' <a href="%s">11-coregistration</a>,',[html1.QCdir 'hist_FC_compare_08moco_11coreg_allSlices.jpg']));
        else
            html1.slicesstr = strcat(html1.slicesstr,sprintf(' 11-coregistration,'));
        end
        if exist([home_dir html1.QCdir(3:end) 'hist_FC_compare_11coreg_12denoise3_allSlices.jpg'],'file')
            html1.slicesstr = strcat(html1.slicesstr,sprintf(' <a href="%s">12-denoise-3-CSF</a>,',[html1.QCdir 'hist_FC_compare_11coreg_12denoise3_allSlices.jpg']));
        else
            html1.slicesstr = strcat(html1.slicesstr,sprintf(' 12-denoise-3-CSF,'));
        end
        if exist([home_dir html1.QCdir(3:end) 'hist_FC_compare_12denoise3_13denoise4_allSlices.jpg'],'file')
            html1.slicesstr = strcat(html1.slicesstr,sprintf(' <a href="%s">13-denoise-4-WM</a>,',[html1.QCdir 'hist_FC_compare_12denoise3_13denoise4_allSlices.jpg']));
        else
            html1.slicesstr = strcat(html1.slicesstr,sprintf(' 13-denoise-4-WM,'));
        end
        if exist([home_dir html1.QCdir(3:end) 'hist_FC_compare_13denoise4_14denoise5_allSlices.jpg'],'file')
            html1.slicesstr = strcat(html1.slicesstr,sprintf(' <a href="%s">14-denoise-5-covariates</a>,',[html1.QCdir 'hist_FC_compare_13denoise4_14denoise5_allSlices.jpg']));
        else
            html1.slicesstr = strcat(html1.slicesstr,sprintf(' 14-denoise-5-covariates,'));
        end
        if exist([home_dir html1.QCdir(3:end) 'hist_FC_compare_14denoise5_15filter_allSlices.jpg'],'file')
            html1.slicesstr = strcat(html1.slicesstr,sprintf(' <a href="%s">15-filtering</a>,',[html1.QCdir 'hist_FC_compare_14denoise5_15filter_allSlices.jpg']));
        else
            html1.slicesstr = strcat(html1.slicesstr,sprintf(' 15-filtering,'));
        end
        if exist([home_dir html1.QCdir(3:end) 'hist_FC_compare_15filter_17deconv_allSlices.jpg'],'file')
            html1.slicesstr = strcat(html1.slicesstr,sprintf(' <a href="%s">17-deconvolution</a>,',[html1.QCdir 'hist_FC_compare_15filter_17deconv_allSlices.jpg']));
        else
            html1.slicesstr = strcat(html1.slicesstr,sprintf(' 17-deconvolution,'));
        end
        
        if (html1k2==1) && (runs(html1k)==1)
            disp(sprintf('      Subject %d of %d - %s  %s',html1k,length(sub_dir),subnames{html1k},html1.slicesstr(2:end-1)))
        elseif (html1k2==1) && (runs(html1k)~=1)
            disp(sprintf('      Subject %d of %d, run %d of %d - %s  %s',html1k,length(sub_dir),html1k2,runs(html1k),subnames{html1k},html1.slicesstr(2:end-1)))
        else
            disp(sprintf('                      run %d of %d - %s  %s',html1k2,runs(html1k),subnames{html1k},html1.slicesstr(2:end-1)))
        end
        clear html1.QCdir html1.slicesstr html1k3
    end
end

fprintf('\n')
fprintf('<b>00.4. Motion parameters:</b>\n')
for html1k=1:length(sub_dir)
    for html1k2=1:runs(html1k)
        if runs(html1k)==1
            html1.QCdir = ['.' erase(sub_dir{html1k},home_dir(1:end-1)) 'QC/' 'QC_shortlist/'];
        else
            html1.QCdir = ['.' erase(sub_dir{html1k},home_dir(1:end-1)) 'QC/' 'QC_shortlist/' 'run' num2str(html1k2) '/'];
        end
        html1.slicesstr = '';
        if exist([home_dir html1.QCdir(3:end) 'motion_params.jpg'],'file')
            html1.slicesstr = strcat(html1.slicesstr,sprintf(' <a href="%s">motion parameters</a>,',[html1.QCdir 'motion_params.jpg']));
        else
            html1.slicesstr = strcat(html1.slicesstr,sprintf(' motion parameters,'));
        end
        if exist([home_dir html1.QCdir(3:end) 'relative_displacement.jpg'],'file')
            html1.slicesstr = strcat(html1.slicesstr,sprintf(' <a href="%s">relative displacement</a>,',[html1.QCdir 'relative_displacement.jpg']));
        else
            html1.slicesstr = strcat(html1.slicesstr,sprintf(' relative displacement,'));
        end
        if exist([home_dir html1.QCdir(3:end) 'mean_max_FD.jpg'],'file')
            html1.slicesstr = strcat(html1.slicesstr,sprintf(' <a href="%s">framewise displacement across slices</a>,',[html1.QCdir 'mean_max_FD.jpg']));
        else
            html1.slicesstr = strcat(html1.slicesstr,sprintf(' framewise displacement across slices,'));
        end
        
        if (html1k2==1) && (runs(html1k)==1)
            disp(sprintf('      Subject %d of %d - %s  %s',html1k,length(sub_dir),subnames{html1k},html1.slicesstr(2:end-1)))
        elseif (html1k2==1) && (runs(html1k)~=1)
            disp(sprintf('      Subject %d of %d, run %d of %d - %s  %s',html1k,length(sub_dir),html1k2,runs(html1k),subnames{html1k},html1.slicesstr(2:end-1)))
        else
            disp(sprintf('                      run %d of %d - %s  %s',html1k2,runs(html1k),subnames{html1k},html1.slicesstr(2:end-1)))
        end
        clear html1.QCdir html1.slicesstr html1k3
    end
end

fprintf('\n')
fprintf('<b>00.5. HRF parameters and/or fALFF (if estimated):</b>\n')
for html1k=1:length(sub_dir)
    for html1k2=1:runs(html1k)
        if runs(html1k)==1
            html1.QCdir = ['.' erase(sub_dir{html1k},home_dir(1:end-1)) 'QC/' 'QC_shortlist/'];
        else
            html1.QCdir = ['.' erase(sub_dir{html1k},home_dir(1:end-1)) 'QC/' 'QC_shortlist/' 'run' num2str(html1k2) '/'];
        end
        html1.slicesstr = '';
        if exist([home_dir html1.QCdir(3:end) 'HRF_plots_17deconv_allSlices.jpg'],'file')
            html1.slicesstr = strcat(html1.slicesstr,sprintf(' <a href="%s">HRF plots</a>,',[html1.QCdir 'HRF_plots_17deconv_allSlices.jpg']));
        else
            html1.slicesstr = strcat(html1.slicesstr,sprintf(' HRF plots,'));
        end
        if exist([home_dir html1.QCdir(3:end) 'HRF_parameters_17deconv_allSlices.jpg'],'file')
            html1.slicesstr = strcat(html1.slicesstr,sprintf(' <a href="%s">HRF parameters</a>,',[html1.QCdir 'HRF_parameters_17deconv_allSlices.jpg']));
        else
            html1.slicesstr = strcat(html1.slicesstr,sprintf(' HRF parameters,'));
        end
        if exist([home_dir html1.QCdir(3:end) 'fALFF_across_slices.jpg'],'file')
            html1.slicesstr = strcat(html1.slicesstr,sprintf(' <a href="%s">fALFF</a>,',[html1.QCdir 'fALFF_across_slices.jpg']));
        else
            html1.slicesstr = strcat(html1.slicesstr,sprintf(' fALFF,'));
        end
        
        if (html1k2==1) && (runs(html1k)==1)
            disp(sprintf('      Subject %d of %d - %s  %s',html1k,length(sub_dir),subnames{html1k},html1.slicesstr(2:end-1)))
        elseif (html1k2==1) && (runs(html1k)~=1)
            disp(sprintf('      Subject %d of %d, run %d of %d - %s  %s',html1k,length(sub_dir),html1k2,runs(html1k),subnames{html1k},html1.slicesstr(2:end-1)))
        else
            disp(sprintf('                      run %d of %d - %s  %s',html1k2,runs(html1k),subnames{html1k},html1.slicesstr(2:end-1)))
        end
        clear html1.QCdir html1.slicesstr html1k3
    end
end

fprintf('\n')
fprintf('<b>00.6. tSNR:</b>\n')
for html1k=1:length(sub_dir)
    for html1k2=1:runs(html1k)
        if runs(html1k)==1
            html1.QCdir = ['.' erase(sub_dir{html1k},home_dir(1:end-1)) 'QC/' 'QC_shortlist/'];
        else
            html1.QCdir = ['.' erase(sub_dir{html1k},home_dir(1:end-1)) 'QC/' 'QC_shortlist/' 'run' num2str(html1k2) '/'];
        end
        html1.slicesstr = '';
        if exist([home_dir html1.QCdir(3:end) 'tSNR_across_preproc_steps_avg.jpg'],'file')
            html1.slicesstr = strcat(html1.slicesstr,sprintf(' <a href="%s">tSNR across preproc steps</a>,',[html1.QCdir 'tSNR_across_preproc_steps_avg.jpg']));
        else
            html1.slicesstr = strcat(html1.slicesstr,sprintf(' tSNR across preproc steps,'));
        end
        if exist([home_dir html1.QCdir(3:end) 'tSNR_across_preproc_steps_slicewise.jpg'],'file')
            html1.slicesstr = strcat(html1.slicesstr,sprintf(' <a href="%s">tSNR across slices</a>,',[html1.QCdir 'tSNR_across_preproc_steps_slicewise.jpg']));
        else
            html1.slicesstr = strcat(html1.slicesstr,sprintf(' tSNR across slices,'));
        end
        if exist([home_dir html1.QCdir(3:end) 'tSNR_maps_1_rawdata_beforeNORDIC.jpg'],'file')
            html1.slicesstr = strcat(html1.slicesstr,sprintf(' <a href="%s">tSNR-maps-before-NORDIC</a>,',[html1.QCdir 'tSNR_maps_1_rawdata_beforeNORDIC.jpg']));
        else
            html1.slicesstr = strcat(html1.slicesstr,sprintf(' tSNR-maps-before-NORDIC,'));
        end
        if exist([home_dir html1.QCdir(3:end) 'tSNR_maps_4_rawdata.jpg'],'file')
            html1.slicesstr = strcat(html1.slicesstr,sprintf(' <a href="%s">tSNR-maps-rawData</a>,',[html1.QCdir 'tSNR_maps_4_rawdata.jpg']));
        else
            html1.slicesstr = strcat(html1.slicesstr,sprintf(' tSNR-maps-rawData,'));
        end
        if exist([home_dir html1.QCdir(3:end) 'tSNR_maps_6_denoise1.jpg'],'file')
            html1.slicesstr = strcat(html1.slicesstr,sprintf(' <a href="%s">tSNR-maps-after-denoise1</a>,',[html1.QCdir 'tSNR_maps_6_denoise1.jpg']));
        else
            html1.slicesstr = strcat(html1.slicesstr,sprintf(' tSNR-maps-after-denoise1,'));
        end
        if exist([home_dir html1.QCdir(3:end) 'tSNR_maps_8_motion_correction.jpg'],'file')
            html1.slicesstr = strcat(html1.slicesstr,sprintf(' <a href="%s">tSNR-maps-after-MoCo</a>,',[html1.QCdir 'tSNR_maps_8_motion_correction.jpg']));
        else
            html1.slicesstr = strcat(html1.slicesstr,sprintf(' tSNR-maps-after-MoCo,'));
        end
        if exist([home_dir html1.QCdir(3:end) 'tSNR_maps_9_denoise2.jpg'],'file')
            html1.slicesstr = strcat(html1.slicesstr,sprintf(' <a href="%s">tSNR-maps-after-RETROICOR</a>,',[html1.QCdir 'tSNR_maps_9_denoise2.jpg']));
        else
            html1.slicesstr = strcat(html1.slicesstr,sprintf(' tSNR-maps-after-RETROICOR,'));
        end
        if exist([home_dir html1.QCdir(3:end) 'tSNR_maps_GM_11_coreg.jpg'],'file')
            html1.slicesstr = strcat(html1.slicesstr,sprintf(' <a href="%s">tSNR-maps-after-coreg</a>,',[html1.QCdir 'tSNR_maps_GM_11_coreg.jpg']));
        else
            html1.slicesstr = strcat(html1.slicesstr,sprintf(' tSNR-maps-after-coreg,'));
        end
        if exist([home_dir html1.QCdir(3:end) 'tSNR_maps_GM_12_CSFregress.jpg'],'file')
            html1.slicesstr = strcat(html1.slicesstr,sprintf(' <a href="%s">tSNR-maps-after-CSFregress</a>,',[html1.QCdir 'tSNR_maps_GM_12_CSFregress.jpg']));
        else
            html1.slicesstr = strcat(html1.slicesstr,sprintf(' tSNR-maps-after-CSFregress,'));
        end
        if exist([home_dir html1.QCdir(3:end) 'tSNR_maps_GM_13_WMregress.jpg'],'file')
            html1.slicesstr = strcat(html1.slicesstr,sprintf(' <a href="%s">tSNR-maps-after-WMregress</a>,',[html1.QCdir 'tSNR_maps_GM_13_WMregress.jpg']));
        else
            html1.slicesstr = strcat(html1.slicesstr,sprintf(' tSNR-maps-after-WMregress,'));
        end
        if exist([home_dir html1.QCdir(3:end) 'tSNR_maps_GM_14_COVregress.jpg'],'file')
            html1.slicesstr = strcat(html1.slicesstr,sprintf(' <a href="%s">tSNR-maps-after-COVregress</a>,',[html1.QCdir 'tSNR_maps_GM_14_COVregress.jpg']));
        else
            html1.slicesstr = strcat(html1.slicesstr,sprintf(' tSNR-maps-after-COVregress,'));
        end
        
        if (html1k2==1) && (runs(html1k)==1)
            disp(sprintf('      Subject %d of %d - %s  %s',html1k,length(sub_dir),subnames{html1k},html1.slicesstr(2:end-1)))
        elseif (html1k2==1) && (runs(html1k)~=1)
            disp(sprintf('      Subject %d of %d, run %d of %d - %s  %s',html1k,length(sub_dir),html1k2,runs(html1k),subnames{html1k},html1.slicesstr(2:end-1)))
        else
            disp(sprintf('                      run %d of %d - %s  %s',html1k2,runs(html1k),subnames{html1k},html1.slicesstr(2:end-1)))
        end
        clear html1.QCdir html1.slicesstr html1k3
    end
end

fprintf('\n')
fprintf('<b>00.7. Functional connectivity (if estimated):</b>\n')
for html1k=1:length(sub_dir)
    for html1k2=1:runs(html1k)
        if runs(html1k)==1
            html1.QCdir = ['.' erase(sub_dir{html1k},home_dir(1:end-1)) 'QC/' 'QC_shortlist/'];
        else
            html1.QCdir = ['.' erase(sub_dir{html1k},home_dir(1:end-1)) 'QC/' 'QC_shortlist/' 'run' num2str(html1k2) '/'];
        end
        html1.slicesstr = '';
        if exist([home_dir html1.QCdir(3:end) 'FC_compare_boxplot_20quadTS_allSlices.jpg'],'file')
            html1.slicesstr = strcat(html1.slicesstr,sprintf(' <a href="%s">boxplot-quadFC</a>,',[html1.QCdir 'FC_compare_boxplot_20quadTS_allSlices.jpg']));
        else
            html1.slicesstr = strcat(html1.slicesstr,sprintf(' boxplot-quadFC,'));
        end
        if exist([home_dir html1.QCdir(3:end) 'FCmatrix_WithinSlice_allSlices.jpg'],'file')
            html1.slicesstr = strcat(html1.slicesstr,sprintf(' <a href="%s">FC-withinSlice</a>,',[html1.QCdir 'FCmatrix_WithinSlice_allSlices.jpg']));
        else
            html1.slicesstr = strcat(html1.slicesstr,sprintf(' FC-withinSlice,'));
        end
        if exist([home_dir html1.QCdir(3:end) 'FCmatrix_partialcorr_WithinSlice_allSlices.jpg'],'file')
            html1.slicesstr = strcat(html1.slicesstr,sprintf(' <a href="%s">FC-withinSlice-pcor</a>,',[html1.QCdir 'FCmatrix_partialcorr_WithinSlice_allSlices.jpg']));
        else
            html1.slicesstr = strcat(html1.slicesstr,sprintf(' FC-withinSlice-pcor,'));
        end
        if exist([home_dir html1.QCdir(3:end) 'FCmatrix_Within_and_BetweenSlices_GM.jpg'],'file')
            html1.slicesstr = strcat(html1.slicesstr,sprintf(' <a href="%s">FC-GM-wholeCord</a>,',[html1.QCdir 'FCmatrix_Within_and_BetweenSlices_GM.jpg']));
        else
            html1.slicesstr = strcat(html1.slicesstr,sprintf(' FC-GM-wholeCord,'));
        end
        if exist([home_dir html1.QCdir(3:end) 'FCmatrix_Within_and_BetweenSlices_WM.jpg'],'file')
            html1.slicesstr = strcat(html1.slicesstr,sprintf(' <a href="%s">FC-WM-wholeCord</a>,',[html1.QCdir 'FCmatrix_Within_and_BetweenSlices_WM.jpg']));
        else
            html1.slicesstr = strcat(html1.slicesstr,sprintf(' FC-WM-wholeCord,'));
        end
        if exist([home_dir html1.QCdir(3:end) 'FCmatrix_partialcorr_Within_and_BetweenSlices_GM.jpg'],'file')
            html1.slicesstr = strcat(html1.slicesstr,sprintf(' <a href="%s">FC-GM-wholeCord-pcor</a>,',[html1.QCdir 'FCmatrix_partialcorr_Within_and_BetweenSlices_GM.jpg']));
        else
            html1.slicesstr = strcat(html1.slicesstr,sprintf(' FC-GM-wholeCord-pcor,'));
        end
        if exist([home_dir html1.QCdir(3:end) 'FCmatrix_partialcorr_Within_and_BetweenSlices_WM.jpg'],'file')
            html1.slicesstr = strcat(html1.slicesstr,sprintf(' <a href="%s">FC-WM-wholeCord-pcor</a>,',[html1.QCdir 'FCmatrix_partialcorr_Within_and_BetweenSlices_WM.jpg']));
        else
            html1.slicesstr = strcat(html1.slicesstr,sprintf(' FC-WM-wholeCord-pcor,'));
        end
        
        if (html1k2==1) && (runs(html1k)==1)
            disp(sprintf('      Subject %d of %d - %s  %s',html1k,length(sub_dir),subnames{html1k},html1.slicesstr(2:end-1)))
        elseif (html1k2==1) && (runs(html1k)~=1)
            disp(sprintf('      Subject %d of %d, run %d of %d - %s  %s',html1k,length(sub_dir),html1k2,runs(html1k),subnames{html1k},html1.slicesstr(2:end-1)))
        else
            disp(sprintf('                      run %d of %d - %s  %s',html1k2,runs(html1k),subnames{html1k},html1.slicesstr(2:end-1)))
        end
        clear html1.QCdir html1.slicesstr html1k3
    end
end

fprintf('</p>')

%% Unprocessed raw data (BEFORE NORDIC denoising)
if exist('choices_preproc','var')
if choices_preproc(1)==1
fprintf('<p style="font-size:14px ; color:#000000">')

fprintf('<b>01.1. Time series within the spinal cord in unprocessed raw data BEFORE NORDIC denoising:</b>\n')
for html1k=1:length(sub_dir)
    for html1k2=1:runs(html1k)
        if runs(html1k)==1
            html1.QCdir = ['.' erase(sub_dir{html1k},home_dir(1:end-1)) 'QC/' '01_NORDIC_denoising/'];
        else
            html1.QCdir = ['.' erase(sub_dir{html1k},home_dir(1:end-1)) 'QC/' '01_NORDIC_denoising/' 'run' num2str(html1k2) '/'];
        end
        if exist([home_dir html1.QCdir(3:end) 'fMRI_ts_01nordic_allSlices.jpg'],'file')
            if (html1k2==1) && (runs(html1k)==1)
                disp(sprintf('      Subject %d of %d - %s<a href="%s"> time series plot (all slices combined)</a>',html1k,length(sub_dir),subnames{html1k},[html1.QCdir 'fMRI_ts_01nordic_allSlices.jpg']))
            elseif (html1k2==1) && (runs(html1k)~=1)
                disp(sprintf('      Subject %d of %d, run %d of %d - %s<a href="%s"> time series plot (all slices combined)</a>',html1k,length(sub_dir),html1k2,runs(html1k),subnames{html1k},[html1.QCdir 'fMRI_ts_01nordic_allSlices.jpg']))
            else
                disp(sprintf('                      run %d of %d - %s<a href="%s"> time series plot (all slices combined)</a>',html1k2,runs(html1k),subnames{html1k},[html1.QCdir 'fMRI_ts_01nordic_allSlices.jpg']))
            end
        else
            if (html1k2==1) && (runs(html1k)==1)
                disp(sprintf('      Subject %d of %d - (pending execution)',html1k,length(sub_dir)))
            elseif (html1k2==1) && (runs(html1k)~=1)
                disp(sprintf('      Subject %d of %d, run %d of %d - (pending execution)',html1k,length(sub_dir),html1k2,runs(html1k)))
            else
                disp(sprintf('                      run %d of %d - (pending execution)',html1k2,runs(html1k)))
            end
        end
        html1.slicesstr = '';
        for html1k3=1:slices
            if exist([home_dir html1.QCdir(3:end) 'fMRI_ts_01nordic_slice' sprintf('%.2d',html1k3) '.jpg'],'file')
                html1.slicesstr = strcat(html1.slicesstr,sprintf(' <a href="%s">slice-%d</a>,',[html1.QCdir 'fMRI_ts_01nordic_slice' sprintf('%.2d',html1k3) '.jpg'],html1k3));
            else
                html1.slicesstr = strcat(html1.slicesstr,sprintf(' slice-%d,',html1k3));
            end
        end
        if exist([home_dir html1.QCdir(3:end) 'fMRI_ts_01nordic_slice' sprintf('%.2d',1) '.jpg'],'file')
            disp(sprintf('                                   (%s)',html1.slicesstr(2:end-1)))
        end
        clear html1.QCdir html1.slicesstr html1k3
    end
end

fprintf('\n')
fprintf('<b>01.2. Global mean signal within the spinal cord in unprocessed raw data BEFORE NORDIC denoising:</b>\n')
for html1k=1:length(sub_dir)
    for html1k2=1:runs(html1k)
        if runs(html1k)==1
            html1.QCdir = ['.' erase(sub_dir{html1k},home_dir(1:end-1)) 'QC/' '01_NORDIC_denoising/'];
        else
            html1.QCdir = ['.' erase(sub_dir{html1k},home_dir(1:end-1)) 'QC/' '01_NORDIC_denoising/' 'run' num2str(html1k2) '/'];
        end
        if exist([home_dir html1.QCdir(3:end) 'global_signal_01nordic_allSlices.jpg'],'file')
            if (html1k2==1) && (runs(html1k)==1)
                disp(sprintf('      Subject %d of %d - %s<a href="%s"> global mean signal (all slices combined)</a>',html1k,length(sub_dir),subnames{html1k},[html1.QCdir 'global_signal_01nordic_allSlices.jpg']))
            elseif (html1k2==1) && (runs(html1k)~=1)
                disp(sprintf('      Subject %d of %d, run %d of %d - %s<a href="%s"> global mean signal (all slices combined)</a>',html1k,length(sub_dir),html1k2,runs(html1k),subnames{html1k},[html1.QCdir 'global_signal_01nordic_allSlices.jpg']))
            else
                disp(sprintf('                      run %d of %d - %s<a href="%s"> global mean signal (all slices combined)</a>',html1k2,runs(html1k),subnames{html1k},[html1.QCdir 'global_signal_01nordic_allSlices.jpg']))
            end
        else
            if (html1k2==1) && (runs(html1k)==1)
                disp(sprintf('      Subject %d of %d - (pending execution)',html1k,length(sub_dir)))
            elseif (html1k2==1) && (runs(html1k)~=1)
                disp(sprintf('      Subject %d of %d, run %d of %d - (pending execution)',html1k,length(sub_dir),html1k2,runs(html1k)))
            else
                disp(sprintf('                      run %d of %d - (pending execution)',html1k2,runs(html1k)))
            end
        end
        html1.slicesstr = '';
        for html1k3=1:slices
            if exist([home_dir html1.QCdir(3:end) 'global_signal_01nordic_slice' sprintf('%.2d',html1k3) '.jpg'],'file')
                html1.slicesstr = strcat(html1.slicesstr,sprintf(' <a href="%s">slice-%d</a>,',[html1.QCdir 'global_signal_01nordic_slice' sprintf('%.2d',html1k3) '.jpg'],html1k3));
            else
                html1.slicesstr = strcat(html1.slicesstr,sprintf(' slice-%d,',html1k3));
            end
        end
        if exist([home_dir html1.QCdir(3:end) 'global_signal_01nordic_slice' sprintf('%.2d',1) '.jpg'],'file')
            disp(sprintf('                                   (%s)',html1.slicesstr(2:end-1)))
        end
        clear html1.QCdir html1.slicesstr html1k3
    end
end

fprintf('\n')
fprintf('<b>01.3. Histogram of voxel-to-voxel correlations within the spinal cord in unprocessed raw data BEFORE NORDIC denoising:</b>\n')
for html1k=1:length(sub_dir)
    for html1k2=1:runs(html1k)
        if runs(html1k)==1
            html1.QCdir = ['.' erase(sub_dir{html1k},home_dir(1:end-1)) 'QC/' '01_NORDIC_denoising/'];
        else
            html1.QCdir = ['.' erase(sub_dir{html1k},home_dir(1:end-1)) 'QC/' '01_NORDIC_denoising/' 'run' num2str(html1k2) '/'];
        end
        if exist([home_dir html1.QCdir(3:end) 'hist_FC_compare_01nordic_allSlices.jpg'],'file')
            if (html1k2==1) && (runs(html1k)==1)
                disp(sprintf('      Subject %d of %d - %s<a href="%s"> histogram of functional connectivity (all slices combined)</a>',html1k,length(sub_dir),subnames{html1k},[html1.QCdir 'hist_FC_compare_01nordic_allSlices.jpg']))
            elseif (html1k2==1) && (runs(html1k)~=1)
                disp(sprintf('      Subject %d of %d, run %d of %d - %s<a href="%s"> histogram of functional connectivity (all slices combined)</a>',html1k,length(sub_dir),html1k2,runs(html1k),subnames{html1k},[html1.QCdir 'hist_FC_compare_01nordic_allSlices.jpg']))
            else
                disp(sprintf('                      run %d of %d - %s<a href="%s"> histogram of functional connectivity (all slices combined)</a>',html1k2,runs(html1k),subnames{html1k},[html1.QCdir 'hist_FC_compare_01nordic_allSlices.jpg']))
            end
        else
            if (html1k2==1) && (runs(html1k)==1)
                disp(sprintf('      Subject %d of %d - (pending execution)',html1k,length(sub_dir)))
            elseif (html1k2==1) && (runs(html1k)~=1)
                disp(sprintf('      Subject %d of %d, run %d of %d - (pending execution)',html1k,length(sub_dir),html1k2,runs(html1k)))
            else
                disp(sprintf('                      run %d of %d - (pending execution)',html1k2,runs(html1k)))
            end
        end
        html1.slicesstr = '';
        for html1k3=1:slices
            if exist([home_dir html1.QCdir(3:end) 'hist_FC_compare_01nordic_slice' sprintf('%.2d',html1k3) '.jpg'],'file')
                html1.slicesstr = strcat(html1.slicesstr,sprintf(' <a href="%s">slice-%d</a>,',[html1.QCdir 'hist_FC_compare_01nordic_slice' sprintf('%.2d',html1k3) '.jpg'],html1k3));
            else
                html1.slicesstr = strcat(html1.slicesstr,sprintf(' slice-%d,',html1k3));
            end
        end
        if exist([home_dir html1.QCdir(3:end) 'hist_FC_compare_01nordic_slice' sprintf('%.2d',1) '.jpg'],'file')
            disp(sprintf('                                   (%s)',html1.slicesstr(2:end-1)))
        end
        clear html1.QCdir html1.slicesstr html1k3
    end
end

if ~isempty(which('VideoWriter')) && (save_videos_steps(1)==1) % check if Matlab's audio video toolbox exists
fprintf('\n')
fprintf('<b>01.4. FMRI video of unprocessed raw data BEFORE NORDIC denoising:</b>\n')
for html1k=1:length(sub_dir)
    for html1k2=1:runs(html1k)
        if runs(html1k)==1
            html1.QCdir = ['.' erase(sub_dir{html1k},home_dir(1:end-1)) 'QC/' '01_NORDIC_denoising/'];
        else
            html1.QCdir = ['.' erase(sub_dir{html1k},home_dir(1:end-1)) 'QC/' '01_NORDIC_denoising/' 'run' num2str(html1k2) '/'];
        end
        html1.slicesstr = '';
        for html1k3=1:slices
            if exist([home_dir html1.QCdir(3:end) 'fMRIvideo_01nordic_slice' sprintf('%.2d',html1k3) '.mp4'],'file')
                html1.slicesstr = strcat(html1.slicesstr,sprintf(' <a href="%s">slice-%d</a>,',[html1.QCdir 'fMRIvideo_01nordic_slice' sprintf('%.2d',html1k3) '.mp4'],html1k3));
            else
                html1.slicesstr = strcat(html1.slicesstr,sprintf(' slice-%d,',html1k3));
            end
        end
        if exist([home_dir html1.QCdir(3:end) 'fMRIvideo_01nordic_slice' sprintf('%.2d',1) '.mp4'],'file')
            if (html1k2==1) && (runs(html1k)==1)
                disp(sprintf('      Subject %d of %d - %s %s',html1k,length(sub_dir),subnames{html1k},html1.slicesstr(2:end-1)))
            elseif (html1k2==1) && (runs(html1k)~=1)
                disp(sprintf('      Subject %d of %d, run %d of %d - %s %s',html1k,length(sub_dir),html1k2,runs(html1k),subnames{html1k},html1.slicesstr(2:end-1)))
            else
                disp(sprintf('                      run %d of %d - %s %s',html1k2,runs(html1k),subnames{html1k},html1.slicesstr(2:end-1)))
            end
        else
            if (html1k2==1) && (runs(html1k)==1)
                disp(sprintf('      Subject %d of %d - (pending execution)',html1k,length(sub_dir)))
            elseif (html1k2==1) && (runs(html1k)~=1)
                disp(sprintf('      Subject %d of %d, run %d of %d - (pending execution)',html1k,length(sub_dir),html1k2,runs(html1k)))
            else
                disp(sprintf('                      run %d of %d - (pending execution)',html1k2,runs(html1k)))
            end
        end
    end
    clear html1.QCdir html1.slicesstr html1k3
end
end

else
fprintf('<b>Not in pipeline</b>\n')
fprintf('</p>')
end
end

%% Unprocessed raw data (AFTER NORDIC denoising, if NORDIC was performed) (this will be the default raw data moving forward)
fprintf('<p style="font-size:14px ; color:#000000">')
fprintf('<b>01.1. Time series within the spinal cord in UNPROCESSED RAW DATA:</b>\n')
for html1k=1:length(sub_dir)
    for html1k2=1:runs(html1k)
        if runs(html1k)==1
            html1.QCdir = ['.' erase(sub_dir{html1k},home_dir(1:end-1)) 'QC/' '04_splitData_rawQC/'];
        else
            html1.QCdir = ['.' erase(sub_dir{html1k},home_dir(1:end-1)) 'QC/' '04_splitData_rawQC/' 'run' num2str(html1k2) '/'];
        end
        if exist([home_dir html1.QCdir(3:end) 'fMRI_ts_rawdata_allSlices.jpg'],'file')
            if (html1k2==1) && (runs(html1k)==1)
                disp(sprintf('      Subject %d of %d - %s<a href="%s"> time series plot (all slices combined)</a>',html1k,length(sub_dir),subnames{html1k},[html1.QCdir 'fMRI_ts_rawdata_allSlices.jpg']))
            elseif (html1k2==1) && (runs(html1k)~=1)
                disp(sprintf('      Subject %d of %d, run %d of %d - %s<a href="%s"> time series plot (all slices combined)</a>',html1k,length(sub_dir),html1k2,runs(html1k),subnames{html1k},[html1.QCdir 'fMRI_ts_rawdata_allSlices.jpg']))
            else
                disp(sprintf('                      run %d of %d - %s<a href="%s"> time series plot (all slices combined)</a>',html1k2,runs(html1k),subnames{html1k},[html1.QCdir 'fMRI_ts_rawdata_allSlices.jpg']))
            end
        else
            if (html1k2==1) && (runs(html1k)==1)
                disp(sprintf('      Subject %d of %d - (pending execution)',html1k,length(sub_dir)))
            elseif (html1k2==1) && (runs(html1k)~=1)
                disp(sprintf('      Subject %d of %d, run %d of %d - (pending execution)',html1k,length(sub_dir),html1k2,runs(html1k)))
            else
                disp(sprintf('                      run %d of %d - (pending execution)',html1k2,runs(html1k)))
            end
        end
        html1.slicesstr = '';
        for html1k3=1:slices
            if exist([home_dir html1.QCdir(3:end) 'fMRI_ts_rawdata_slice' sprintf('%.2d',html1k3) '.jpg'],'file')
                html1.slicesstr = strcat(html1.slicesstr,sprintf(' <a href="%s">slice-%d</a>,',[html1.QCdir 'fMRI_ts_rawdata_slice' sprintf('%.2d',html1k3) '.jpg'],html1k3));
            else
                html1.slicesstr = strcat(html1.slicesstr,sprintf(' slice-%d,',html1k3));
            end
        end
        if exist([home_dir html1.QCdir(3:end) 'fMRI_ts_rawdata_slice' sprintf('%.2d',1) '.jpg'],'file')
            disp(sprintf('                                   (%s)',html1.slicesstr(2:end-1)))
        end
        clear html1.QCdir html1.slicesstr html1k3
    end
end

fprintf('\n')
fprintf('<b>01.2. Global mean signal within the spinal cord in UNPROCESSED RAW DATA:</b>\n')
for html1k=1:length(sub_dir)
    for html1k2=1:runs(html1k)
        if runs(html1k)==1
            html1.QCdir = ['.' erase(sub_dir{html1k},home_dir(1:end-1)) 'QC/' '04_splitData_rawQC/'];
        else
            html1.QCdir = ['.' erase(sub_dir{html1k},home_dir(1:end-1)) 'QC/' '04_splitData_rawQC/' 'run' num2str(html1k2) '/'];
        end
        if exist([home_dir html1.QCdir(3:end) 'global_signal_rawdata_allSlices.jpg'],'file')
            if (html1k2==1) && (runs(html1k)==1)
                disp(sprintf('      Subject %d of %d - %s<a href="%s"> global mean signal (all slices combined)</a>',html1k,length(sub_dir),subnames{html1k},[html1.QCdir 'global_signal_rawdata_allSlices.jpg']))
            elseif (html1k2==1) && (runs(html1k)~=1)
                disp(sprintf('      Subject %d of %d, run %d of %d - %s<a href="%s"> global mean signal (all slices combined)</a>',html1k,length(sub_dir),html1k2,runs(html1k),subnames{html1k},[html1.QCdir 'global_signal_rawdata_allSlices.jpg']))
            else
                disp(sprintf('                      run %d of %d - %s<a href="%s"> global mean signal (all slices combined)</a>',html1k2,runs(html1k),subnames{html1k},[html1.QCdir 'global_signal_rawdata_allSlices.jpg']))
            end
        else
            if (html1k2==1) && (runs(html1k)==1)
                disp(sprintf('      Subject %d of %d - (pending execution)',html1k,length(sub_dir)))
            elseif (html1k2==1) && (runs(html1k)~=1)
                disp(sprintf('      Subject %d of %d, run %d of %d - (pending execution)',html1k,length(sub_dir),html1k2,runs(html1k)))
            else
                disp(sprintf('                      run %d of %d - (pending execution)',html1k2,runs(html1k)))
            end
        end
        html1.slicesstr = '';
        for html1k3=1:slices
            if exist([home_dir html1.QCdir(3:end) 'global_signal_rawdata_slice' sprintf('%.2d',html1k3) '.jpg'],'file')
                html1.slicesstr = strcat(html1.slicesstr,sprintf(' <a href="%s">slice-%d</a>,',[html1.QCdir 'global_signal_rawdata_slice' sprintf('%.2d',html1k3) '.jpg'],html1k3));
            else
                html1.slicesstr = strcat(html1.slicesstr,sprintf(' slice-%d,',html1k3));
            end
        end
        if exist([home_dir html1.QCdir(3:end) 'global_signal_rawdata_slice' sprintf('%.2d',1) '.jpg'],'file')
            disp(sprintf('                                   (%s)',html1.slicesstr(2:end-1)))
        end
        clear html1.QCdir html1.slicesstr html1k3
    end
end

fprintf('\n')
fprintf('<b>01.3. Histogram of voxel-to-voxel correlations within the spinal cord in UNPROCESSED RAW DATA:</b>\n')
for html1k=1:length(sub_dir)
    for html1k2=1:runs(html1k)
        if runs(html1k)==1
            html1.QCdir = ['.' erase(sub_dir{html1k},home_dir(1:end-1)) 'QC/' '04_splitData_rawQC/'];
        else
            html1.QCdir = ['.' erase(sub_dir{html1k},home_dir(1:end-1)) 'QC/' '04_splitData_rawQC/' 'run' num2str(html1k2) '/'];
        end
        if exist([home_dir html1.QCdir(3:end) 'hist_FC_rawdata_allSlices.jpg'],'file')
            if (html1k2==1) && (runs(html1k)==1)
                disp(sprintf('      Subject %d of %d - %s<a href="%s"> histogram of functional connectivity (all slices combined)</a>',html1k,length(sub_dir),subnames{html1k},[html1.QCdir 'hist_FC_rawdata_allSlices.jpg']))
            elseif (html1k2==1) && (runs(html1k)~=1)
                disp(sprintf('      Subject %d of %d, run %d of %d - %s<a href="%s"> histogram of functional connectivity (all slices combined)</a>',html1k,length(sub_dir),html1k2,runs(html1k),subnames{html1k},[html1.QCdir 'hist_FC_rawdata_allSlices.jpg']))
            else
                disp(sprintf('                      run %d of %d - %s<a href="%s"> histogram of functional connectivity (all slices combined)</a>',html1k2,runs(html1k),subnames{html1k},[html1.QCdir 'hist_FC_rawdata_allSlices.jpg']))
            end
        else
            if (html1k2==1) && (runs(html1k)==1)
                disp(sprintf('      Subject %d of %d - (pending execution)',html1k,length(sub_dir)))
            elseif (html1k2==1) && (runs(html1k)~=1)
                disp(sprintf('      Subject %d of %d, run %d of %d - (pending execution)',html1k,length(sub_dir),html1k2,runs(html1k)))
            else
                disp(sprintf('                      run %d of %d - (pending execution)',html1k2,runs(html1k)))
            end
        end
        html1.slicesstr = '';
        for html1k3=1:slices
            if exist([home_dir html1.QCdir(3:end) 'hist_FC_rawdata_slice' sprintf('%.2d',html1k3) '.jpg'],'file')
                html1.slicesstr = strcat(html1.slicesstr,sprintf(' <a href="%s">slice-%d</a>,',[html1.QCdir 'hist_FC_rawdata_slice' sprintf('%.2d',html1k3) '.jpg'],html1k3));
            else
                html1.slicesstr = strcat(html1.slicesstr,sprintf(' slice-%d,',html1k3));
            end
        end
        if exist([home_dir html1.QCdir(3:end) 'hist_FC_rawdata_slice' sprintf('%.2d',1) '.jpg'],'file')
            disp(sprintf('                                   (%s)',html1.slicesstr(2:end-1)))
        end
        clear html1.QCdir html1.slicesstr html1k3
    end
end

if ~isempty(which('VideoWriter')) && (save_videos_steps(1)==1) % check if Matlab's audio video toolbox exists
fprintf('\n')
fprintf('<b>01.4. FMRI video of UNPROCESSED RAW DATA:</b>\n')
for html1k=1:length(sub_dir)
    for html1k2=1:runs(html1k)
        if runs(html1k)==1
            html1.QCdir = ['.' erase(sub_dir{html1k},home_dir(1:end-1)) 'QC/' '04_splitData_rawQC/'];
        else
            html1.QCdir = ['.' erase(sub_dir{html1k},home_dir(1:end-1)) 'QC/' '04_splitData_rawQC/' 'run' num2str(html1k2) '/'];
        end
        html1.slicesstr = '';
        for html1k3=1:slices
            if exist([home_dir html1.QCdir(3:end) 'fMRIvideo_rawdata_slice' sprintf('%.2d',html1k3) '.mp4'],'file')
                html1.slicesstr = strcat(html1.slicesstr,sprintf(' <a href="%s">slice-%d</a>,',[html1.QCdir 'fMRIvideo_rawdata_slice' sprintf('%.2d',html1k3) '.mp4'],html1k3));
            else
                html1.slicesstr = strcat(html1.slicesstr,sprintf(' slice-%d,',html1k3));
            end
        end
        if exist([home_dir html1.QCdir(3:end) 'fMRIvideo_rawdata_slice' sprintf('%.2d',1) '.mp4'],'file')
            if (html1k2==1) && (runs(html1k)==1)
                disp(sprintf('      Subject %d of %d - %s %s',html1k,length(sub_dir),subnames{html1k},html1.slicesstr(2:end-1)))
            elseif (html1k2==1) && (runs(html1k)~=1)
                disp(sprintf('      Subject %d of %d, run %d of %d - %s %s',html1k,length(sub_dir),html1k2,runs(html1k),subnames{html1k},html1.slicesstr(2:end-1)))
            else
                disp(sprintf('                      run %d of %d - %s %s',html1k2,runs(html1k),subnames{html1k},html1.slicesstr(2:end-1)))
            end
        else
            if (html1k2==1) && (runs(html1k)==1)
                disp(sprintf('      Subject %d of %d - (pending execution)',html1k,length(sub_dir)))
            elseif (html1k2==1) && (runs(html1k)~=1)
                disp(sprintf('      Subject %d of %d, run %d of %d - (pending execution)',html1k,length(sub_dir),html1k2,runs(html1k)))
            else
                disp(sprintf('                      run %d of %d - (pending execution)',html1k2,runs(html1k)))
            end
        end
    end
    clear html1.QCdir html1.slicesstr html1k3
end
end

fprintf('</p>')

%% 05. Define "not cord" mask
if exist('choices_preproc','var')
if choices_preproc(5)==1
fprintf('<p style="font-size:14px ; color:#000000">')

fprintf('<b>05.1. User-defined boundary for the "not cord" mask, overlaid on the mean functional image:</b>\n')
for html1k=1:length(sub_dir)
    for html1k2=1:runs(html1k)
        if runs(html1k)==1
            html1.QCdir = ['.' erase(sub_dir{html1k},home_dir(1:end-1)) 'QC/' '05_notCordMask/'];
        else
            html1.QCdir = ['.' erase(sub_dir{html1k},home_dir(1:end-1)) 'QC/' '05_notCordMask/' 'run' num2str(html1k2) '/'];
        end
        html1.slicesstr = '';
        for html1k3=1:slices
            if exist([home_dir html1.QCdir(3:end) 'notCordMask_slice' sprintf('%.2d',html1k3) '.jpg'],'file')
                html1.slicesstr = strcat(html1.slicesstr,sprintf(' <a href="%s">slice-%d</a>,',[html1.QCdir 'notCordMask_slice' sprintf('%.2d',html1k3) '.jpg'],html1k3));
            else
                html1.slicesstr = strcat(html1.slicesstr,sprintf(' slice-%d,',html1k3));
            end
        end
        if exist([home_dir html1.QCdir(3:end) 'notCordMask_slice' sprintf('%.2d',1) '.jpg'],'file')
            if (html1k2==1) && (runs(html1k)==1)
                disp(sprintf('      Subject %d of %d - %s %s',html1k,length(sub_dir),subnames{html1k},html1.slicesstr(2:end-1)))
            elseif (html1k2==1) && (runs(html1k)~=1)
                disp(sprintf('      Subject %d of %d, run %d of %d - %s %s',html1k,length(sub_dir),html1k2,runs(html1k),subnames{html1k},html1.slicesstr(2:end-1)))
            else
                disp(sprintf('                      run %d of %d - %s %s',html1k2,runs(html1k),subnames{html1k},html1.slicesstr(2:end-1)))
            end
        else
            if (html1k2==1) && (runs(html1k)==1)
                disp(sprintf('      Subject %d of %d - (pending execution)',html1k,length(sub_dir)))
            elseif (html1k2==1) && (runs(html1k)~=1)
                disp(sprintf('      Subject %d of %d, run %d of %d - (pending execution)',html1k,length(sub_dir),html1k2,runs(html1k)))
            else
                disp(sprintf('                      run %d of %d - (pending execution)',html1k2,runs(html1k)))
            end
        end
    end
    clear html1.QCdir html1.slicesstr html1k3
end
else
fprintf('<b>Not in pipeline</b>\n')
fprintf('</p>')
end
end

%% 06. Denoise-1 (non-cord signal regression)
if exist('choices_preproc','var')
if choices_preproc(6)==1
fprintf('<p style="font-size:14px ; color:#000000">')

fprintf('<b>06.1. Time series within the spinal cord after "06-denoise1":</b>\n')
for html1k=1:length(sub_dir)
    for html1k2=1:runs(html1k)
        if runs(html1k)==1
            html1.QCdir = ['.' erase(sub_dir{html1k},home_dir(1:end-1)) 'QC/' '06_denoise1/'];
        else
            html1.QCdir = ['.' erase(sub_dir{html1k},home_dir(1:end-1)) 'QC/' '06_denoise1/' 'run' num2str(html1k2) '/'];
        end
        if exist([home_dir html1.QCdir(3:end) 'fMRI_ts_06denoise1_allSlices.jpg'],'file')
            if (html1k2==1) && (runs(html1k)==1)
                disp(sprintf('      Subject %d of %d - %s<a href="%s"> time series plot (all slices combined)</a>',html1k,length(sub_dir),subnames{html1k},[html1.QCdir 'fMRI_ts_06denoise1_allSlices.jpg']))
            elseif (html1k2==1) && (runs(html1k)~=1)
                disp(sprintf('      Subject %d of %d, run %d of %d - %s<a href="%s"> time series plot (all slices combined)</a>',html1k,length(sub_dir),html1k2,runs(html1k),subnames{html1k},[html1.QCdir 'fMRI_ts_06denoise1_allSlices.jpg']))
            else
                disp(sprintf('                      run %d of %d - %s<a href="%s"> time series plot (all slices combined)</a>',html1k2,runs(html1k),subnames{html1k},[html1.QCdir 'fMRI_ts_06denoise1_allSlices.jpg']))
            end
        else
            if (html1k2==1) && (runs(html1k)==1)
                disp(sprintf('      Subject %d of %d - (pending execution)',html1k,length(sub_dir)))
            elseif (html1k2==1) && (runs(html1k)~=1)
                disp(sprintf('      Subject %d of %d, run %d of %d - (pending execution)',html1k,length(sub_dir),html1k2,runs(html1k)))
            else
                disp(sprintf('                      run %d of %d - (pending execution)',html1k2,runs(html1k)))
            end
        end
        html1.slicesstr = '';
        for html1k3=1:slices
            if exist([home_dir html1.QCdir(3:end) 'fMRI_ts_06denoise1_slice' sprintf('%.2d',html1k3) '.jpg'],'file')
                html1.slicesstr = strcat(html1.slicesstr,sprintf(' <a href="%s">slice-%d</a>,',[html1.QCdir 'fMRI_ts_06denoise1_slice' sprintf('%.2d',html1k3) '.jpg'],html1k3));
            else
                html1.slicesstr = strcat(html1.slicesstr,sprintf(' slice-%d,',html1k3));
            end
        end
        if exist([home_dir html1.QCdir(3:end) 'fMRI_ts_06denoise1_slice' sprintf('%.2d',1) '.jpg'],'file')
            disp(sprintf('                                   (%s)',html1.slicesstr(2:end-1)))
        end
        clear html1.QCdir html1.slicesstr html1k3
    end
end

fprintf('\n')
fprintf('<b>06.2. Global mean signal within the spinal cord after "06-denoise1":</b>\n')
for html1k=1:length(sub_dir)
    for html1k2=1:runs(html1k)
        if runs(html1k)==1
            html1.QCdir = ['.' erase(sub_dir{html1k},home_dir(1:end-1)) 'QC/' '06_denoise1/'];
        else
            html1.QCdir = ['.' erase(sub_dir{html1k},home_dir(1:end-1)) 'QC/' '06_denoise1/' 'run' num2str(html1k2) '/'];
        end
        if exist([home_dir html1.QCdir(3:end) 'global_signal_06denoise1_allSlices.jpg'],'file')
            if (html1k2==1) && (runs(html1k)==1)
                disp(sprintf('      Subject %d of %d - %s<a href="%s"> global mean signal (all slices combined)</a>',html1k,length(sub_dir),subnames{html1k},[html1.QCdir 'global_signal_06denoise1_allSlices.jpg']))
            elseif (html1k2==1) && (runs(html1k)~=1)
                disp(sprintf('      Subject %d of %d, run %d of %d - %s<a href="%s"> global mean signal (all slices combined)</a>',html1k,length(sub_dir),html1k2,runs(html1k),subnames{html1k},[html1.QCdir 'global_signal_06denoise1_allSlices.jpg']))
            else
                disp(sprintf('                      run %d of %d - %s<a href="%s"> global mean signal (all slices combined)</a>',html1k2,runs(html1k),subnames{html1k},[html1.QCdir 'global_signal_06denoise1_allSlices.jpg']))
            end
        else
            if (html1k2==1) && (runs(html1k)==1)
                disp(sprintf('      Subject %d of %d - (pending execution)',html1k,length(sub_dir)))
            elseif (html1k2==1) && (runs(html1k)~=1)
                disp(sprintf('      Subject %d of %d, run %d of %d - (pending execution)',html1k,length(sub_dir),html1k2,runs(html1k)))
            else
                disp(sprintf('                      run %d of %d - (pending execution)',html1k2,runs(html1k)))
            end
        end
        html1.slicesstr = '';
        for html1k3=1:slices
            if exist([home_dir html1.QCdir(3:end) 'global_signal_06denoise1_slice' sprintf('%.2d',html1k3) '.jpg'],'file')
                html1.slicesstr = strcat(html1.slicesstr,sprintf(' <a href="%s">slice-%d</a>,',[html1.QCdir 'global_signal_06denoise1_slice' sprintf('%.2d',html1k3) '.jpg'],html1k3));
            else
                html1.slicesstr = strcat(html1.slicesstr,sprintf(' slice-%d,',html1k3));
            end
        end
        if exist([home_dir html1.QCdir(3:end) 'global_signal_06denoise1_slice' sprintf('%.2d',1) '.jpg'],'file')
            disp(sprintf('                                   (%s)',html1.slicesstr(2:end-1)))
        end
        clear html1.QCdir html1.slicesstr html1k3
    end
end

fprintf('\n')
fprintf('<b>06.3. Histogram of voxel-to-voxel correlations within the spinal cord after "06-denoise1":</b>\n')
for html1k=1:length(sub_dir)
    for html1k2=1:runs(html1k)
        if runs(html1k)==1
            html1.QCdir = ['.' erase(sub_dir{html1k},home_dir(1:end-1)) 'QC/' '06_denoise1/'];
        else
            html1.QCdir = ['.' erase(sub_dir{html1k},home_dir(1:end-1)) 'QC/' '06_denoise1/' 'run' num2str(html1k2) '/'];
        end
        if exist([home_dir html1.QCdir(3:end) 'hist_FC_compare_06denoise1_allSlices.jpg'],'file')
            if (html1k2==1) && (runs(html1k)==1)
                disp(sprintf('      Subject %d of %d - %s<a href="%s"> histogram of functional connectivity (all slices combined)</a>',html1k,length(sub_dir),subnames{html1k},[html1.QCdir 'hist_FC_compare_06denoise1_allSlices.jpg']))
            elseif (html1k2==1) && (runs(html1k)~=1)
                disp(sprintf('      Subject %d of %d, run %d of %d - %s<a href="%s"> histogram of functional connectivity (all slices combined)</a>',html1k,length(sub_dir),html1k2,runs(html1k),subnames{html1k},[html1.QCdir 'hist_FC_compare_06denoise1_allSlices.jpg']))
            else
                disp(sprintf('                      run %d of %d - %s<a href="%s"> histogram of functional connectivity (all slices combined)</a>',html1k2,runs(html1k),subnames{html1k},[html1.QCdir 'hist_FC_compare_06denoise1_allSlices.jpg']))
            end
        else
            if (html1k2==1) && (runs(html1k)==1)
                disp(sprintf('      Subject %d of %d - (pending execution)',html1k,length(sub_dir)))
            elseif (html1k2==1) && (runs(html1k)~=1)
                disp(sprintf('      Subject %d of %d, run %d of %d - (pending execution)',html1k,length(sub_dir),html1k2,runs(html1k)))
            else
                disp(sprintf('                      run %d of %d - (pending execution)',html1k2,runs(html1k)))
            end
        end
        html1.slicesstr = '';
        for html1k3=1:slices
            if exist([home_dir html1.QCdir(3:end) 'hist_FC_compare_06denoise1_slice' sprintf('%.2d',html1k3) '.jpg'],'file')
                html1.slicesstr = strcat(html1.slicesstr,sprintf(' <a href="%s">slice-%d</a>,',[html1.QCdir 'hist_FC_compare_06denoise1_slice' sprintf('%.2d',html1k3) '.jpg'],html1k3));
            else
                html1.slicesstr = strcat(html1.slicesstr,sprintf(' slice-%d,',html1k3));
            end
        end
        if exist([home_dir html1.QCdir(3:end) 'hist_FC_compare_06denoise1_slice' sprintf('%.2d',1) '.jpg'],'file')
            disp(sprintf('                                   (%s)',html1.slicesstr(2:end-1)))
        end
        clear html1.QCdir html1.slicesstr html1k3
    end
end

if ~isempty(which('VideoWriter')) && (save_videos_steps(2)==1) % check if Matlab's audio video toolbox exists
fprintf('\n')
fprintf('<b>06.4. FMRI video of data after "06-denoise1":</b>\n')
for html1k=1:length(sub_dir)
    for html1k2=1:runs(html1k)
        if runs(html1k)==1
            html1.QCdir = ['.' erase(sub_dir{html1k},home_dir(1:end-1)) 'QC/' '06_denoise1/'];
        else
            html1.QCdir = ['.' erase(sub_dir{html1k},home_dir(1:end-1)) 'QC/' '06_denoise1/' 'run' num2str(html1k2) '/'];
        end
        html1.slicesstr = '';
        for html1k3=1:slices
            if exist([home_dir html1.QCdir(3:end) 'fMRIvideo_06denoise1_slice' sprintf('%.2d',html1k3) '.mp4'],'file')
                html1.slicesstr = strcat(html1.slicesstr,sprintf(' <a href="%s">slice-%d</a>,',[html1.QCdir 'fMRIvideo_06denoise1_slice' sprintf('%.2d',html1k3) '.mp4'],html1k3));
            else
                html1.slicesstr = strcat(html1.slicesstr,sprintf(' slice-%d,',html1k3));
            end
        end
        if exist([home_dir html1.QCdir(3:end) 'fMRIvideo_06denoise1_slice' sprintf('%.2d',1) '.mp4'],'file')
            if (html1k2==1) && (runs(html1k)==1)
                disp(sprintf('      Subject %d of %d - %s %s',html1k,length(sub_dir),subnames{html1k},html1.slicesstr(2:end-1)))
            elseif (html1k2==1) && (runs(html1k)~=1)
                disp(sprintf('      Subject %d of %d, run %d of %d - %s %s',html1k,length(sub_dir),html1k2,runs(html1k),subnames{html1k},html1.slicesstr(2:end-1)))
            else
                disp(sprintf('                      run %d of %d - %s %s',html1k2,runs(html1k),subnames{html1k},html1.slicesstr(2:end-1)))
            end
        else
            if (html1k2==1) && (runs(html1k)==1)
                disp(sprintf('      Subject %d of %d - (pending execution)',html1k,length(sub_dir)))
            elseif (html1k2==1) && (runs(html1k)~=1)
                disp(sprintf('      Subject %d of %d, run %d of %d - (pending execution)',html1k,length(sub_dir),html1k2,runs(html1k)))
            else
                disp(sprintf('                      run %d of %d - (pending execution)',html1k2,runs(html1k)))
            end
        end
    end
    clear html1.QCdir html1.slicesstr html1k3
end
end

else
fprintf('<b>Not in pipeline</b>\n')
fprintf('</p>')
end
end

%% 07. Define Gaussian mask
if exist('choices_preproc','var')
if choices_preproc(7)==1
fprintf('<p style="font-size:14px ; color:#000000">')
if ~exist('automatic_GaussianMask','var'), automatic_GaussianMask=1; end
if automatic_GaussianMask==0
    fprintf('<b>07.1. Mean functional image masked by the user-defined Gaussian mask:</b>\n')
else
    fprintf('<b>07.1. Mean functional image masked by the auto-generated Gaussian mask:</b>\n')
end
for html1k=1:length(sub_dir)
    for html1k2=1:runs(html1k)
        if runs(html1k)==1
            html1.QCdir = ['.' erase(sub_dir{html1k},home_dir(1:end-1)) 'QC/' '07_gaussianMask/'];
        else
            html1.QCdir = ['.' erase(sub_dir{html1k},home_dir(1:end-1)) 'QC/' '07_gaussianMask/' 'run' num2str(html1k2) '/'];
        end
        html1.slicesstr = '';
        for html1k3=1:slices
            if exist([home_dir html1.QCdir(3:end) 'MeanFunc_GaussianMasked_slice' sprintf('%.2d',html1k3) '.jpg'],'file')
                html1.slicesstr = strcat(html1.slicesstr,sprintf(' <a href="%s">slice-%d</a>,',[html1.QCdir 'MeanFunc_GaussianMasked_slice' sprintf('%.2d',html1k3) '.jpg'],html1k3));
            else
                html1.slicesstr = strcat(html1.slicesstr,sprintf(' slice-%d,',html1k3));
            end
        end
        if exist([home_dir html1.QCdir(3:end) 'MeanFunc_GaussianMasked_slice' sprintf('%.2d',1) '.jpg'],'file')
            if (html1k2==1) && (runs(html1k)==1)
                disp(sprintf('      Subject %d of %d - %s %s',html1k,length(sub_dir),subnames{html1k},html1.slicesstr(2:end-1)))
            elseif (html1k2==1) && (runs(html1k)~=1)
                disp(sprintf('      Subject %d of %d, run %d of %d - %s %s',html1k,length(sub_dir),html1k2,runs(html1k),subnames{html1k},html1.slicesstr(2:end-1)))
            else
                disp(sprintf('                      run %d of %d - %s %s',html1k2,runs(html1k),subnames{html1k},html1.slicesstr(2:end-1)))
            end
        else
            if (html1k2==1) && (runs(html1k)==1)
                disp(sprintf('      Subject %d of %d - (pending execution)',html1k,length(sub_dir)))
            elseif (html1k2==1) && (runs(html1k)~=1)
                disp(sprintf('      Subject %d of %d, run %d of %d - (pending execution)',html1k,length(sub_dir),html1k2,runs(html1k)))
            else
                disp(sprintf('                      run %d of %d - (pending execution)',html1k2,runs(html1k)))
            end
        end
    end
    clear html1.QCdir html1.slicesstr html1k3
end
else
fprintf('<b>Not in pipeline</b>\n')
fprintf('</p>')
end
end

%% 08. Motion correction
if exist('choices_preproc','var')
if choices_preproc(8)==1
fprintf('<p style="font-size:14px ; color:#000000">')

fprintf('<b>08.1. Motion parameters:</b>\n')
for html1k=1:length(sub_dir)
    for html1k2=1:runs(html1k)
        if runs(html1k)==1
            html1.QCdir = ['.' erase(sub_dir{html1k},home_dir(1:end-1)) 'QC/' '08_motionCorrection/'];
        else
            html1.QCdir = ['.' erase(sub_dir{html1k},home_dir(1:end-1)) 'QC/' '08_motionCorrection/' 'run' num2str(html1k2) '/'];
        end
        if exist([home_dir html1.QCdir(3:end) 'motion_params.jpg'],'file')
            if (html1k2==1) && (runs(html1k)==1)
                disp(sprintf('      Subject %d of %d - %s<a href="%s"> motion parameters (all slices)</a>',html1k,length(sub_dir),subnames{html1k},[html1.QCdir 'motion_params.jpg']))
            elseif (html1k2==1) && (runs(html1k)~=1)
                disp(sprintf('      Subject %d of %d, run %d of %d - %s<a href="%s"> motion parameters (all slices)</a>',html1k,length(sub_dir),html1k2,runs(html1k),subnames{html1k},[html1.QCdir 'motion_params.jpg']))
            else
                disp(sprintf('                      run %d of %d - %s<a href="%s"> motion parameters (all slices)</a>',html1k2,runs(html1k),subnames{html1k},[html1.QCdir 'motion_params.jpg']))
            end
        else
            if (html1k2==1) && (runs(html1k)==1)
                disp(sprintf('      Subject %d of %d - (pending execution)',html1k,length(sub_dir)))
            elseif (html1k2==1) && (runs(html1k)~=1)
                disp(sprintf('      Subject %d of %d, run %d of %d - (pending execution)',html1k,length(sub_dir),html1k2,runs(html1k)))
            else
                disp(sprintf('                      run %d of %d - (pending execution)',html1k2,runs(html1k)))
            end
        end
        if exist([home_dir html1.QCdir(3:end) 'relative_displacement.jpg'],'file')
            if (html1k2==1) && (runs(html1k)==1)
                disp(sprintf('                     - %s<a href="%s"> framewise displacement (all slices)</a>',subnames{html1k},[html1.QCdir 'relative_displacement.jpg']))
            elseif (html1k2==1) && (runs(html1k)~=1)
                disp(sprintf('                                 - %s<a href="%s"> framewise displacement (all slices)</a>',subnames{html1k},[html1.QCdir 'relative_displacement.jpg']))
            else
                disp(sprintf('                                 - %s<a href="%s"> framewise displacement (all slices)</a>',subnames{html1k},[html1.QCdir 'relative_displacement.jpg']))
            end
        end
        if exist([home_dir html1.QCdir(3:end) 'mean_max_FD.jpg'],'file')
            if (html1k2==1) && (runs(html1k)==1)
                disp(sprintf('                     - %s<a href="%s"> mean and max framewise displacement per slice</a>',subnames{html1k},[html1.QCdir 'mean_max_FD.jpg']))
            elseif (html1k2==1) && (runs(html1k)~=1)
                disp(sprintf('                                 - %s<a href="%s"> mean and max framewise displacement per slice</a>',subnames{html1k},[html1.QCdir 'mean_max_FD.jpg']))
            else
                disp(sprintf('                                 - %s<a href="%s"> mean and max framewise displacement per slice</a>',subnames{html1k},[html1.QCdir 'mean_max_FD.jpg']))
            end
        end
        clear html1.QCdir html1k3
    end
end

fprintf('\n')
fprintf('<b>08.2. Time series within the spinal cord after "08-motion-correction":</b>\n')
for html1k=1:length(sub_dir)
    for html1k2=1:runs(html1k)
        if runs(html1k)==1
            html1.QCdir = ['.' erase(sub_dir{html1k},home_dir(1:end-1)) 'QC/' '08_motionCorrection/'];
        else
            html1.QCdir = ['.' erase(sub_dir{html1k},home_dir(1:end-1)) 'QC/' '08_motionCorrection/' 'run' num2str(html1k2) '/'];
        end
        if exist([home_dir html1.QCdir(3:end) 'fMRI_ts_08moco_allSlices.jpg'],'file')
            if (html1k2==1) && (runs(html1k)==1)
                disp(sprintf('      Subject %d of %d - %s<a href="%s"> time series plot (all slices combined)</a>',html1k,length(sub_dir),subnames{html1k},[html1.QCdir 'fMRI_ts_08moco_allSlices.jpg']))
            elseif (html1k2==1) && (runs(html1k)~=1)
                disp(sprintf('      Subject %d of %d, run %d of %d - %s<a href="%s"> time series plot (all slices combined)</a>',html1k,length(sub_dir),html1k2,runs(html1k),subnames{html1k},[html1.QCdir 'fMRI_ts_08moco_allSlices.jpg']))
            else
                disp(sprintf('                      run %d of %d - %s<a href="%s"> time series plot (all slices combined)</a>',html1k2,runs(html1k),subnames{html1k},[html1.QCdir 'fMRI_ts_08moco_allSlices.jpg']))
            end
        else
            if (html1k2==1) && (runs(html1k)==1)
                disp(sprintf('      Subject %d of %d - (pending execution)',html1k,length(sub_dir)))
            elseif (html1k2==1) && (runs(html1k)~=1)
                disp(sprintf('      Subject %d of %d, run %d of %d - (pending execution)',html1k,length(sub_dir),html1k2,runs(html1k)))
            else
                disp(sprintf('                      run %d of %d - (pending execution)',html1k2,runs(html1k)))
            end
        end
        html1.slicesstr = '';
        for html1k3=1:slices
            if exist([home_dir html1.QCdir(3:end) 'fMRI_ts_08moco_slice' sprintf('%.2d',html1k3) '.jpg'],'file')
                html1.slicesstr = strcat(html1.slicesstr,sprintf(' <a href="%s">slice-%d</a>,',[html1.QCdir 'fMRI_ts_08moco_slice' sprintf('%.2d',html1k3) '.jpg'],html1k3));
            else
                html1.slicesstr = strcat(html1.slicesstr,sprintf(' slice-%d,',html1k3));
            end
        end
        if exist([home_dir html1.QCdir(3:end) 'fMRI_ts_08moco_slice' sprintf('%.2d',1) '.jpg'],'file')
            disp(sprintf('                                   (%s)',html1.slicesstr(2:end-1)))
        end
        clear html1.QCdir html1.slicesstr html1k3
    end
end

fprintf('\n')
fprintf('<b>08.3. Global mean signal within the spinal cord after "08-motion-correction":</b>\n')
for html1k=1:length(sub_dir)
    for html1k2=1:runs(html1k)
        if runs(html1k)==1
            html1.QCdir = ['.' erase(sub_dir{html1k},home_dir(1:end-1)) 'QC/' '08_motionCorrection/'];
        else
            html1.QCdir = ['.' erase(sub_dir{html1k},home_dir(1:end-1)) 'QC/' '08_motionCorrection/' 'run' num2str(html1k2) '/'];
        end
        if exist([home_dir html1.QCdir(3:end) 'global_signal_08moco_allSlices.jpg'],'file')
            if (html1k2==1) && (runs(html1k)==1)
                disp(sprintf('      Subject %d of %d - %s<a href="%s"> global mean signal (all slices combined)</a>',html1k,length(sub_dir),subnames{html1k},[html1.QCdir 'global_signal_08moco_allSlices.jpg']))
            elseif (html1k2==1) && (runs(html1k)~=1)
                disp(sprintf('      Subject %d of %d, run %d of %d - %s<a href="%s"> global mean signal (all slices combined)</a>',html1k,length(sub_dir),html1k2,runs(html1k),subnames{html1k},[html1.QCdir 'global_signal_08moco_allSlices.jpg']))
            else
                disp(sprintf('                      run %d of %d - %s<a href="%s"> global mean signal (all slices combined)</a>',html1k2,runs(html1k),subnames{html1k},[html1.QCdir 'global_signal_08moco_allSlices.jpg']))
            end
        else
            if (html1k2==1) && (runs(html1k)==1)
                disp(sprintf('      Subject %d of %d - (pending execution)',html1k,length(sub_dir)))
            elseif (html1k2==1) && (runs(html1k)~=1)
                disp(sprintf('      Subject %d of %d, run %d of %d - (pending execution)',html1k,length(sub_dir),html1k2,runs(html1k)))
            else
                disp(sprintf('                      run %d of %d - (pending execution)',html1k2,runs(html1k)))
            end
        end
        html1.slicesstr = '';
        for html1k3=1:slices
            if exist([home_dir html1.QCdir(3:end) 'global_signal_08moco_slice' sprintf('%.2d',html1k3) '.jpg'],'file')
                html1.slicesstr = strcat(html1.slicesstr,sprintf(' <a href="%s">slice-%d</a>,',[html1.QCdir 'global_signal_08moco_slice' sprintf('%.2d',html1k3) '.jpg'],html1k3));
            else
                html1.slicesstr = strcat(html1.slicesstr,sprintf(' slice-%d,',html1k3));
            end
        end
        if exist([home_dir html1.QCdir(3:end) 'global_signal_08moco_slice' sprintf('%.2d',1) '.jpg'],'file')
            disp(sprintf('                                   (%s)',html1.slicesstr(2:end-1)))
        end
        clear html1.QCdir html1.slicesstr html1k3
    end
end

fprintf('\n')
fprintf('<b>08.4. Global mean signal: "08-motion-correction" vs. "06-denoise1":</b>\n')
for html1k=1:length(sub_dir)
    for html1k2=1:runs(html1k)
        if runs(html1k)==1
            html1.QCdir = ['.' erase(sub_dir{html1k},home_dir(1:end-1)) 'QC/' '08_motionCorrection/'];
        else
            html1.QCdir = ['.' erase(sub_dir{html1k},home_dir(1:end-1)) 'QC/' '08_motionCorrection/' 'run' num2str(html1k2) '/'];
        end
        if exist([home_dir html1.QCdir(3:end) 'global_signal_06denoise1_vs_08moco_allSlices.jpg'],'file')
            if (html1k2==1) && (runs(html1k)==1)
                disp(sprintf('      Subject %d of %d - %s<a href="%s"> global mean signal (all slices combined)</a>',html1k,length(sub_dir),subnames{html1k},[html1.QCdir 'global_signal_06denoise1_vs_08moco_allSlices.jpg']))
            elseif (html1k2==1) && (runs(html1k)~=1)
                disp(sprintf('      Subject %d of %d, run %d of %d - %s<a href="%s"> global mean signal (all slices combined)</a>',html1k,length(sub_dir),html1k2,runs(html1k),subnames{html1k},[html1.QCdir 'global_signal_06denoise1_vs_08moco_allSlices.jpg']))
            else
                disp(sprintf('                      run %d of %d - %s<a href="%s"> global mean signal (all slices combined)</a>',html1k2,runs(html1k),subnames{html1k},[html1.QCdir 'global_signal_06denoise1_vs_08moco_allSlices.jpg']))
            end
        else
            if (html1k2==1) && (runs(html1k)==1)
                disp(sprintf('      Subject %d of %d - (pending execution)',html1k,length(sub_dir)))
            elseif (html1k2==1) && (runs(html1k)~=1)
                disp(sprintf('      Subject %d of %d, run %d of %d - (pending execution)',html1k,length(sub_dir),html1k2,runs(html1k)))
            else
                disp(sprintf('                      run %d of %d - (pending execution)',html1k2,runs(html1k)))
            end
        end
        html1.slicesstr = '';
        for html1k3=1:slices
            if exist([home_dir html1.QCdir(3:end) 'global_signal_06denoise1_vs_08moco_slice' sprintf('%.2d',html1k3) '.jpg'],'file')
                html1.slicesstr = strcat(html1.slicesstr,sprintf(' <a href="%s">slice-%d</a>,',[html1.QCdir 'global_signal_06denoise1_vs_08moco_slice' sprintf('%.2d',html1k3) '.jpg'],html1k3));
            else
                html1.slicesstr = strcat(html1.slicesstr,sprintf(' slice-%d,',html1k3));
            end
        end
        if exist([home_dir html1.QCdir(3:end) 'global_signal_06denoise1_vs_08moco_slice' sprintf('%.2d',1) '.jpg'],'file')
            disp(sprintf('                                   (%s)',html1.slicesstr(2:end-1)))
        end
        clear html1.QCdir html1.slicesstr html1k3
    end
end

fprintf('\n')
fprintf('<b>08.5. Histogram of voxel-to-voxel correlations within the spinal cord after "08-motion-correction":</b>\n')
for html1k=1:length(sub_dir)
    for html1k2=1:runs(html1k)
        if runs(html1k)==1
            html1.QCdir = ['.' erase(sub_dir{html1k},home_dir(1:end-1)) 'QC/' '08_motionCorrection/'];
        else
            html1.QCdir = ['.' erase(sub_dir{html1k},home_dir(1:end-1)) 'QC/' '08_motionCorrection/' 'run' num2str(html1k2) '/'];
        end
        if exist([home_dir html1.QCdir(3:end) 'hist_FC_compare_08moco_allSlices.jpg'],'file')
            if (html1k2==1) && (runs(html1k)==1)
                disp(sprintf('      Subject %d of %d - %s<a href="%s"> histogram of functional connectivity (all slices combined)</a>',html1k,length(sub_dir),subnames{html1k},[html1.QCdir 'hist_FC_compare_08moco_allSlices.jpg']))
            elseif (html1k2==1) && (runs(html1k)~=1)
                disp(sprintf('      Subject %d of %d, run %d of %d - %s<a href="%s"> histogram of functional connectivity (all slices combined)</a>',html1k,length(sub_dir),html1k2,runs(html1k),subnames{html1k},[html1.QCdir 'hist_FC_compare_08moco_allSlices.jpg']))
            else
                disp(sprintf('                      run %d of %d - %s<a href="%s"> histogram of functional connectivity (all slices combined)</a>',html1k2,runs(html1k),subnames{html1k},[html1.QCdir 'hist_FC_compare_08moco_allSlices.jpg']))
            end
        else
            if (html1k2==1) && (runs(html1k)==1)
                disp(sprintf('      Subject %d of %d - (pending execution)',html1k,length(sub_dir)))
            elseif (html1k2==1) && (runs(html1k)~=1)
                disp(sprintf('      Subject %d of %d, run %d of %d - (pending execution)',html1k,length(sub_dir),html1k2,runs(html1k)))
            else
                disp(sprintf('                      run %d of %d - (pending execution)',html1k2,runs(html1k)))
            end
        end
        html1.slicesstr = '';
        for html1k3=1:slices
            if exist([home_dir html1.QCdir(3:end) 'hist_FC_compare_08moco_slice' sprintf('%.2d',html1k3) '.jpg'],'file')
                html1.slicesstr = strcat(html1.slicesstr,sprintf(' <a href="%s">slice-%d</a>,',[html1.QCdir 'hist_FC_compare_08moco_slice' sprintf('%.2d',html1k3) '.jpg'],html1k3));
            else
                html1.slicesstr = strcat(html1.slicesstr,sprintf(' slice-%d,',html1k3));
            end
        end
        if exist([home_dir html1.QCdir(3:end) 'hist_FC_compare_08moco_slice' sprintf('%.2d',1) '.jpg'],'file')
            disp(sprintf('                                   (%s)',html1.slicesstr(2:end-1)))
        end
        clear html1.QCdir html1.slicesstr html1k3
    end
end

fprintf('\n')
fprintf('<b>08.6. Histogram of voxel-to-voxel correlations: "08-motion-correction" vs. "06-denoise1":</b>\n')
for html1k=1:length(sub_dir)
    for html1k2=1:runs(html1k)
        if runs(html1k)==1
            html1.QCdir = ['.' erase(sub_dir{html1k},home_dir(1:end-1)) 'QC/' '08_motionCorrection/'];
        else
            html1.QCdir = ['.' erase(sub_dir{html1k},home_dir(1:end-1)) 'QC/' '08_motionCorrection/' 'run' num2str(html1k2) '/'];
        end
        if exist([home_dir html1.QCdir(3:end) 'hist_FC_compare_06denoise1_08moco_allSlices.jpg'],'file')
            if (html1k2==1) && (runs(html1k)==1)
                disp(sprintf('      Subject %d of %d - %s<a href="%s"> histogram of functional connectivity (all slices combined)</a>',html1k,length(sub_dir),subnames{html1k},[html1.QCdir 'hist_FC_compare_06denoise1_08moco_allSlices.jpg']))
            elseif (html1k2==1) && (runs(html1k)~=1)
                disp(sprintf('      Subject %d of %d, run %d of %d - %s<a href="%s"> histogram of functional connectivity (all slices combined)</a>',html1k,length(sub_dir),html1k2,runs(html1k),subnames{html1k},[html1.QCdir 'hist_FC_compare_06denoise1_08moco_allSlices.jpg']))
            else
                disp(sprintf('                      run %d of %d - %s<a href="%s"> histogram of functional connectivity (all slices combined)</a>',html1k2,runs(html1k),subnames{html1k},[html1.QCdir 'hist_FC_compare_06denoise1_08moco_allSlices.jpg']))
            end
        else
            if (html1k2==1) && (runs(html1k)==1)
                disp(sprintf('      Subject %d of %d - (pending execution)',html1k,length(sub_dir)))
            elseif (html1k2==1) && (runs(html1k)~=1)
                disp(sprintf('      Subject %d of %d, run %d of %d - (pending execution)',html1k,length(sub_dir),html1k2,runs(html1k)))
            else
                disp(sprintf('                      run %d of %d - (pending execution)',html1k2,runs(html1k)))
            end
        end
        html1.slicesstr = '';
        for html1k3=1:slices
            if exist([home_dir html1.QCdir(3:end) 'hist_FC_compare_06denoise1_08moco_slice' sprintf('%.2d',html1k3) '.jpg'],'file')
                html1.slicesstr = strcat(html1.slicesstr,sprintf(' <a href="%s">slice-%d</a>,',[html1.QCdir 'hist_FC_compare_06denoise1_08moco_slice' sprintf('%.2d',html1k3) '.jpg'],html1k3));
            else
                html1.slicesstr = strcat(html1.slicesstr,sprintf(' slice-%d,',html1k3));
            end
        end
        if exist([home_dir html1.QCdir(3:end) 'hist_FC_compare_06denoise1_08moco_slice' sprintf('%.2d',1) '.jpg'],'file')
            disp(sprintf('                                   (%s)',html1.slicesstr(2:end-1)))
        end
        clear html1.QCdir html1.slicesstr html1k3
    end
end

if ~isempty(which('VideoWriter')) && (save_videos_steps(3)==1) % check if Matlab's audio video toolbox exists
fprintf('\n')
fprintf('<b>08.7. FMRI video of data after "08-motion-correction":</b>\n')
for html1k=1:length(sub_dir)
    for html1k2=1:runs(html1k)
        if runs(html1k)==1
            html1.QCdir = ['.' erase(sub_dir{html1k},home_dir(1:end-1)) 'QC/' '08_motionCorrection/'];
        else
            html1.QCdir = ['.' erase(sub_dir{html1k},home_dir(1:end-1)) 'QC/' '08_motionCorrection/' 'run' num2str(html1k2) '/'];
        end
        html1.slicesstr = '';
        for html1k3=1:slices
            if exist([home_dir html1.QCdir(3:end) 'fMRIvideo_08moco_slice' sprintf('%.2d',html1k3) '.mp4'],'file')
                html1.slicesstr = strcat(html1.slicesstr,sprintf(' <a href="%s">slice-%d</a>,',[html1.QCdir 'fMRIvideo_08moco_slice' sprintf('%.2d',html1k3) '.mp4'],html1k3));
            else
                html1.slicesstr = strcat(html1.slicesstr,sprintf(' slice-%d,',html1k3));
            end
        end
        if exist([home_dir html1.QCdir(3:end) 'fMRIvideo_08moco_slice' sprintf('%.2d',1) '.mp4'],'file')
            if (html1k2==1) && (runs(html1k)==1)
                disp(sprintf('      Subject %d of %d - %s %s',html1k,length(sub_dir),subnames{html1k},html1.slicesstr(2:end-1)))
            elseif (html1k2==1) && (runs(html1k)~=1)
                disp(sprintf('      Subject %d of %d, run %d of %d - %s %s',html1k,length(sub_dir),html1k2,runs(html1k),subnames{html1k},html1.slicesstr(2:end-1)))
            else
                disp(sprintf('                      run %d of %d - %s %s',html1k2,runs(html1k),subnames{html1k},html1.slicesstr(2:end-1)))
            end
        else
            if (html1k2==1) && (runs(html1k)==1)
                disp(sprintf('      Subject %d of %d - (pending execution)',html1k,length(sub_dir)))
            elseif (html1k2==1) && (runs(html1k)~=1)
                disp(sprintf('      Subject %d of %d, run %d of %d - (pending execution)',html1k,length(sub_dir),html1k2,runs(html1k)))
            else
                disp(sprintf('                      run %d of %d - (pending execution)',html1k2,runs(html1k)))
            end
        end
    end
    clear html1.QCdir html1.slicesstr html1k3
end
end

else
fprintf('<b>Not in pipeline</b>\n')
fprintf('</p>')
end
end

%% 09. Denoise-2 (RETROICOR)
if exist('choices_preproc','var')
if choices_preproc(9)==1
fprintf('<p style="font-size:14px ; color:#000000">')

fprintf('<b>09.1. Time series within the spinal cord after "09-denoise2-RETROICOR":</b>\n')
for html1k=1:length(sub_dir)
    for html1k2=1:runs(html1k)
        if runs(html1k)==1
            html1.QCdir = ['.' erase(sub_dir{html1k},home_dir(1:end-1)) 'QC/' '09_denoise2-RETROICOR/'];
        else
            html1.QCdir = ['.' erase(sub_dir{html1k},home_dir(1:end-1)) 'QC/' '09_denoise2-RETROICOR/' 'run' num2str(html1k2) '/'];
        end
        if exist([home_dir html1.QCdir(3:end) 'fMRI_ts_09ricor_allSlices.jpg'],'file')
            if (html1k2==1) && (runs(html1k)==1)
                disp(sprintf('      Subject %d of %d - %s<a href="%s"> time series plot (all slices combined)</a>',html1k,length(sub_dir),subnames{html1k},[html1.QCdir 'fMRI_ts_09ricor_allSlices.jpg']))
            elseif (html1k2==1) && (runs(html1k)~=1)
                disp(sprintf('      Subject %d of %d, run %d of %d - %s<a href="%s"> time series plot (all slices combined)</a>',html1k,length(sub_dir),html1k2,runs(html1k),subnames{html1k},[html1.QCdir 'fMRI_ts_09ricor_allSlices.jpg']))
            else
                disp(sprintf('                      run %d of %d - %s<a href="%s"> time series plot (all slices combined)</a>',html1k2,runs(html1k),subnames{html1k},[html1.QCdir 'fMRI_ts_09ricor_allSlices.jpg']))
            end
        else
            if (html1k2==1) && (runs(html1k)==1)
                disp(sprintf('      Subject %d of %d - (pending execution)',html1k,length(sub_dir)))
            elseif (html1k2==1) && (runs(html1k)~=1)
                disp(sprintf('      Subject %d of %d, run %d of %d - (pending execution)',html1k,length(sub_dir),html1k2,runs(html1k)))
            else
                disp(sprintf('                      run %d of %d - (pending execution)',html1k2,runs(html1k)))
            end
        end
        html1.slicesstr = '';
        for html1k3=1:slices
            if exist([home_dir html1.QCdir(3:end) 'fMRI_ts_09ricor_slice' sprintf('%.2d',html1k3) '.jpg'],'file')
                html1.slicesstr = strcat(html1.slicesstr,sprintf(' <a href="%s">slice-%d</a>,',[html1.QCdir 'fMRI_ts_09ricor_slice' sprintf('%.2d',html1k3) '.jpg'],html1k3));
            else
                html1.slicesstr = strcat(html1.slicesstr,sprintf(' slice-%d,',html1k3));
            end
        end
        if exist([home_dir html1.QCdir(3:end) 'fMRI_ts_09ricor_slice' sprintf('%.2d',1) '.jpg'],'file')
            disp(sprintf('                                   (%s)',html1.slicesstr(2:end-1)))
        end
        clear html1.QCdir html1.slicesstr html1k3
    end
end

fprintf('\n')
fprintf('<b>09.2. Global mean signal within the spinal cord after "09-denoise2-RETROICOR":</b>\n')
for html1k=1:length(sub_dir)
    for html1k2=1:runs(html1k)
        if runs(html1k)==1
            html1.QCdir = ['.' erase(sub_dir{html1k},home_dir(1:end-1)) 'QC/' '09_denoise2-RETROICOR/'];
        else
            html1.QCdir = ['.' erase(sub_dir{html1k},home_dir(1:end-1)) 'QC/' '09_denoise2-RETROICOR/' 'run' num2str(html1k2) '/'];
        end
        if exist([home_dir html1.QCdir(3:end) 'global_signal_09ricor_allSlices.jpg'],'file')
            if (html1k2==1) && (runs(html1k)==1)
                disp(sprintf('      Subject %d of %d - %s<a href="%s"> global mean signal (all slices combined)</a>',html1k,length(sub_dir),subnames{html1k},[html1.QCdir 'global_signal_09ricor_allSlices.jpg']))
            elseif (html1k2==1) && (runs(html1k)~=1)
                disp(sprintf('      Subject %d of %d, run %d of %d - %s<a href="%s"> global mean signal (all slices combined)</a>',html1k,length(sub_dir),html1k2,runs(html1k),subnames{html1k},[html1.QCdir 'global_signal_09ricor_allSlices.jpg']))
            else
                disp(sprintf('                      run %d of %d - %s<a href="%s"> global mean signal (all slices combined)</a>',html1k2,runs(html1k),subnames{html1k},[html1.QCdir 'global_signal_09ricor_allSlices.jpg']))
            end
        else
            if (html1k2==1) && (runs(html1k)==1)
                disp(sprintf('      Subject %d of %d - (pending execution)',html1k,length(sub_dir)))
            elseif (html1k2==1) && (runs(html1k)~=1)
                disp(sprintf('      Subject %d of %d, run %d of %d - (pending execution)',html1k,length(sub_dir),html1k2,runs(html1k)))
            else
                disp(sprintf('                      run %d of %d - (pending execution)',html1k2,runs(html1k)))
            end
        end
        html1.slicesstr = '';
        for html1k3=1:slices
            if exist([home_dir html1.QCdir(3:end) 'global_signal_09ricor_slice' sprintf('%.2d',html1k3) '.jpg'],'file')
                html1.slicesstr = strcat(html1.slicesstr,sprintf(' <a href="%s">slice-%d</a>,',[html1.QCdir 'global_signal_09ricor_slice' sprintf('%.2d',html1k3) '.jpg'],html1k3));
            else
                html1.slicesstr = strcat(html1.slicesstr,sprintf(' slice-%d,',html1k3));
            end
        end
        if exist([home_dir html1.QCdir(3:end) 'global_signal_09ricor_slice' sprintf('%.2d',1) '.jpg'],'file')
            disp(sprintf('                                   (%s)',html1.slicesstr(2:end-1)))
        end
        clear html1.QCdir html1.slicesstr html1k3
    end
end

fprintf('\n')
fprintf('<b>09.3. Global mean signal: "09-denoise2-RETROICOR" vs. "08-motion-correction":</b>\n')
for html1k=1:length(sub_dir)
    for html1k2=1:runs(html1k)
        if runs(html1k)==1
            html1.QCdir = ['.' erase(sub_dir{html1k},home_dir(1:end-1)) 'QC/' '09_denoise2-RETROICOR/'];
        else
            html1.QCdir = ['.' erase(sub_dir{html1k},home_dir(1:end-1)) 'QC/' '09_denoise2-RETROICOR/' 'run' num2str(html1k2) '/'];
        end
        if exist([home_dir html1.QCdir(3:end) 'global_signal_08moco_vs_09ricor_allSlices.jpg'],'file')
            if (html1k2==1) && (runs(html1k)==1)
                disp(sprintf('      Subject %d of %d - %s<a href="%s"> global mean signal (all slices combined)</a>',html1k,length(sub_dir),subnames{html1k},[html1.QCdir 'global_signal_08moco_vs_09ricor_allSlices.jpg']))
            elseif (html1k2==1) && (runs(html1k)~=1)
                disp(sprintf('      Subject %d of %d, run %d of %d - %s<a href="%s"> global mean signal (all slices combined)</a>',html1k,length(sub_dir),html1k2,runs(html1k),subnames{html1k},[html1.QCdir 'global_signal_08moco_vs_09ricor_allSlices.jpg']))
            else
                disp(sprintf('                      run %d of %d - %s<a href="%s"> global mean signal (all slices combined)</a>',html1k2,runs(html1k),subnames{html1k},[html1.QCdir 'global_signal_08moco_vs_09ricor_allSlices.jpg']))
            end
        else
            if (html1k2==1) && (runs(html1k)==1)
                disp(sprintf('      Subject %d of %d - (pending execution)',html1k,length(sub_dir)))
            elseif (html1k2==1) && (runs(html1k)~=1)
                disp(sprintf('      Subject %d of %d, run %d of %d - (pending execution)',html1k,length(sub_dir),html1k2,runs(html1k)))
            else
                disp(sprintf('                      run %d of %d - (pending execution)',html1k2,runs(html1k)))
            end
        end
        html1.slicesstr = '';
        for html1k3=1:slices
            if exist([home_dir html1.QCdir(3:end) 'global_signal_08moco_vs_09ricor_slice' sprintf('%.2d',html1k3) '.jpg'],'file')
                html1.slicesstr = strcat(html1.slicesstr,sprintf(' <a href="%s">slice-%d</a>,',[html1.QCdir 'global_signal_08moco_vs_09ricor_slice' sprintf('%.2d',html1k3) '.jpg'],html1k3));
            else
                html1.slicesstr = strcat(html1.slicesstr,sprintf(' slice-%d,',html1k3));
            end
        end
        if exist([home_dir html1.QCdir(3:end) 'global_signal_08moco_vs_09ricor_slice' sprintf('%.2d',1) '.jpg'],'file')
            disp(sprintf('                                   (%s)',html1.slicesstr(2:end-1)))
        end
        clear html1.QCdir html1.slicesstr html1k3
    end
end

fprintf('\n')
fprintf('<b>09.4. Histogram of voxel-to-voxel correlations within the spinal cord after "09-denoise2-RETROICOR":</b>\n')
for html1k=1:length(sub_dir)
    for html1k2=1:runs(html1k)
        if runs(html1k)==1
            html1.QCdir = ['.' erase(sub_dir{html1k},home_dir(1:end-1)) 'QC/' '09_denoise2-RETROICOR/'];
        else
            html1.QCdir = ['.' erase(sub_dir{html1k},home_dir(1:end-1)) 'QC/' '09_denoise2-RETROICOR/' 'run' num2str(html1k2) '/'];
        end
        if exist([home_dir html1.QCdir(3:end) 'hist_FC_compare_09ricor_allSlices.jpg'],'file')
            if (html1k2==1) && (runs(html1k)==1)
                disp(sprintf('      Subject %d of %d - %s<a href="%s"> histogram of functional connectivity (all slices combined)</a>',html1k,length(sub_dir),subnames{html1k},[html1.QCdir 'hist_FC_compare_09ricor_allSlices.jpg']))
            elseif (html1k2==1) && (runs(html1k)~=1)
                disp(sprintf('      Subject %d of %d, run %d of %d - %s<a href="%s"> histogram of functional connectivity (all slices combined)</a>',html1k,length(sub_dir),html1k2,runs(html1k),subnames{html1k},[html1.QCdir 'hist_FC_compare_09ricor_allSlices.jpg']))
            else
                disp(sprintf('                      run %d of %d - %s<a href="%s"> histogram of functional connectivity (all slices combined)</a>',html1k2,runs(html1k),subnames{html1k},[html1.QCdir 'hist_FC_compare_09ricor_allSlices.jpg']))
            end
        else
            if (html1k2==1) && (runs(html1k)==1)
                disp(sprintf('      Subject %d of %d - (pending execution)',html1k,length(sub_dir)))
            elseif (html1k2==1) && (runs(html1k)~=1)
                disp(sprintf('      Subject %d of %d, run %d of %d - (pending execution)',html1k,length(sub_dir),html1k2,runs(html1k)))
            else
                disp(sprintf('                      run %d of %d - (pending execution)',html1k2,runs(html1k)))
            end
        end
        html1.slicesstr = '';
        for html1k3=1:slices
            if exist([home_dir html1.QCdir(3:end) 'hist_FC_compare_09ricor_slice' sprintf('%.2d',html1k3) '.jpg'],'file')
                html1.slicesstr = strcat(html1.slicesstr,sprintf(' <a href="%s">slice-%d</a>,',[html1.QCdir 'hist_FC_compare_09ricor_slice' sprintf('%.2d',html1k3) '.jpg'],html1k3));
            else
                html1.slicesstr = strcat(html1.slicesstr,sprintf(' slice-%d,',html1k3));
            end
        end
        if exist([home_dir html1.QCdir(3:end) 'hist_FC_compare_09ricor_slice' sprintf('%.2d',1) '.jpg'],'file')
            disp(sprintf('                                   (%s)',html1.slicesstr(2:end-1)))
        end
        clear html1.QCdir html1.slicesstr html1k3
    end
end

fprintf('\n')
fprintf('<b>09.5. Histogram of voxel-to-voxel correlations: "09-denoise2-RETROICOR" vs. "08-motion-correction":</b>\n')
for html1k=1:length(sub_dir)
    for html1k2=1:runs(html1k)
        if runs(html1k)==1
            html1.QCdir = ['.' erase(sub_dir{html1k},home_dir(1:end-1)) 'QC/' '09_denoise2-RETROICOR/'];
        else
            html1.QCdir = ['.' erase(sub_dir{html1k},home_dir(1:end-1)) 'QC/' '09_denoise2-RETROICOR/' 'run' num2str(html1k2) '/'];
        end
        if exist([home_dir html1.QCdir(3:end) 'hist_FC_compare_08moco_09ricor_allSlices.jpg'],'file')
            if (html1k2==1) && (runs(html1k)==1)
                disp(sprintf('      Subject %d of %d - %s<a href="%s"> histogram of functional connectivity (all slices combined)</a>',html1k,length(sub_dir),subnames{html1k},[html1.QCdir 'hist_FC_compare_08moco_09ricor_allSlices.jpg']))
            elseif (html1k2==1) && (runs(html1k)~=1)
                disp(sprintf('      Subject %d of %d, run %d of %d - %s<a href="%s"> histogram of functional connectivity (all slices combined)</a>',html1k,length(sub_dir),html1k2,runs(html1k),subnames{html1k},[html1.QCdir 'hist_FC_compare_08moco_09ricor_allSlices.jpg']))
            else
                disp(sprintf('                      run %d of %d - %s<a href="%s"> histogram of functional connectivity (all slices combined)</a>',html1k2,runs(html1k),subnames{html1k},[html1.QCdir 'hist_FC_compare_08moco_09ricor_allSlices.jpg']))
            end
        else
            if (html1k2==1) && (runs(html1k)==1)
                disp(sprintf('      Subject %d of %d - (pending execution)',html1k,length(sub_dir)))
            elseif (html1k2==1) && (runs(html1k)~=1)
                disp(sprintf('      Subject %d of %d, run %d of %d - (pending execution)',html1k,length(sub_dir),html1k2,runs(html1k)))
            else
                disp(sprintf('                      run %d of %d - (pending execution)',html1k2,runs(html1k)))
            end
        end
        html1.slicesstr = '';
        for html1k3=1:slices
            if exist([home_dir html1.QCdir(3:end) 'hist_FC_compare_08moco_09ricor_slice' sprintf('%.2d',html1k3) '.jpg'],'file')
                html1.slicesstr = strcat(html1.slicesstr,sprintf(' <a href="%s">slice-%d</a>,',[html1.QCdir 'hist_FC_compare_08moco_09ricor_slice' sprintf('%.2d',html1k3) '.jpg'],html1k3));
            else
                html1.slicesstr = strcat(html1.slicesstr,sprintf(' slice-%d,',html1k3));
            end
        end
        if exist([home_dir html1.QCdir(3:end) 'hist_FC_compare_08moco_09ricor_slice' sprintf('%.2d',1) '.jpg'],'file')
            disp(sprintf('                                   (%s)',html1.slicesstr(2:end-1)))
        end
        clear html1.QCdir html1.slicesstr html1k3
    end
end

if ~isempty(which('VideoWriter')) && (save_videos_steps(4)==1) % check if Matlab's audio video toolbox exists
fprintf('\n')
fprintf('<b>09.6. FMRI video of data after "09-denoise2-RETROICOR":</b>\n')
for html1k=1:length(sub_dir)
    for html1k2=1:runs(html1k)
        if runs(html1k)==1
            html1.QCdir = ['.' erase(sub_dir{html1k},home_dir(1:end-1)) 'QC/' '09_denoise2-RETROICOR/'];
        else
            html1.QCdir = ['.' erase(sub_dir{html1k},home_dir(1:end-1)) 'QC/' '09_denoise2-RETROICOR/' 'run' num2str(html1k2) '/'];
        end
        html1.slicesstr = '';
        for html1k3=1:slices
            if exist([home_dir html1.QCdir(3:end) 'fMRIvideo_09ricor_slice' sprintf('%.2d',html1k3) '.mp4'],'file')
                html1.slicesstr = strcat(html1.slicesstr,sprintf(' <a href="%s">slice-%d</a>,',[html1.QCdir 'fMRIvideo_09ricor_slice' sprintf('%.2d',html1k3) '.mp4'],html1k3));
            else
                html1.slicesstr = strcat(html1.slicesstr,sprintf(' slice-%d,',html1k3));
            end
        end
        if exist([home_dir html1.QCdir(3:end) 'fMRIvideo_09ricor_slice' sprintf('%.2d',1) '.mp4'],'file')
            if (html1k2==1) && (runs(html1k)==1)
                disp(sprintf('      Subject %d of %d - %s %s',html1k,length(sub_dir),subnames{html1k},html1.slicesstr(2:end-1)))
            elseif (html1k2==1) && (runs(html1k)~=1)
                disp(sprintf('      Subject %d of %d, run %d of %d - %s %s',html1k,length(sub_dir),html1k2,runs(html1k),subnames{html1k},html1.slicesstr(2:end-1)))
            else
                disp(sprintf('                      run %d of %d - %s %s',html1k2,runs(html1k),subnames{html1k},html1.slicesstr(2:end-1)))
            end
        else
            if (html1k2==1) && (runs(html1k)==1)
                disp(sprintf('      Subject %d of %d - (pending execution)',html1k,length(sub_dir)))
            elseif (html1k2==1) && (runs(html1k)~=1)
                disp(sprintf('      Subject %d of %d, run %d of %d - (pending execution)',html1k,length(sub_dir),html1k2,runs(html1k)))
            else
                disp(sprintf('                      run %d of %d - (pending execution)',html1k2,runs(html1k)))
            end
        end
    end
    clear html1.QCdir html1.slicesstr html1k3
end
end

else
fprintf('<b>Not in pipeline</b>\n')
fprintf('</p>')
end
end

%% 10. Define GM, WM and CSF masks
if exist('choices_preproc','var')
if choices_preproc(10)==1
fprintf('<p style="font-size:14px ; color:#000000">')

fprintf('<b>10.1. Plot of cross-sectional areas across slices:</b>\n')
for html1k=1:length(sub_dir)
    html1.QCdir = ['.' erase(sub_dir{html1k},home_dir(1:end-1)) 'QC/' '10_GM-WM-CSF-masks/'];
    if exist([home_dir html1.QCdir(3:end) 'cross_sectional_areas_across_slices.jpg'],'file')
        disp(sprintf('      Subject %d of %d - %s<a href="%s"> Cross-sectional areas</a>',html1k,length(sub_dir),subnames{html1k},[html1.QCdir 'cross_sectional_areas_across_slices.jpg']))
    else
        disp(sprintf('      Subject %d of %d - (pending execution)',html1k,length(sub_dir)))
    end
    clear html1.QCdir html1k3
end

fprintf('\n')
fprintf('<b>10.2. User-defined cerebrospinal fluid masks:</b>\n')
for html1k=1:length(sub_dir)
    html1.QCdir = ['.' erase(sub_dir{html1k},home_dir(1:end-1)) 'QC/' '10_GM-WM-CSF-masks/'];
    html1.slicesstr = '';
    for html1k3=1:slices
        if exist([home_dir html1.QCdir(3:end) 'CSFmask_slice' sprintf('%.2d',html1k3) '.jpg'],'file')
            html1.slicesstr = strcat(html1.slicesstr,sprintf(' <a href="%s">slice-%d</a>,',[html1.QCdir 'CSFmask_slice' sprintf('%.2d',html1k3) '.jpg'],html1k3));
        else
            html1.slicesstr = strcat(html1.slicesstr,sprintf(' slice-%d,',html1k3));
        end
    end
    if exist([home_dir html1.QCdir(3:end) 'CSFmask_slice' sprintf('%.2d',1) '.jpg'],'file')
        disp(sprintf('      Subject %d of %d - %s %s',html1k,length(sub_dir),subnames{html1k},html1.slicesstr(2:end-1)))
    else
        disp(sprintf('      Subject %d of %d - (pending execution)',html1k,length(sub_dir)))
    end
    clear html1.QCdir html1.slicesstr html1k3
end

fprintf('\n')
fprintf('<b>10.3. User-defined white-matter masks:</b>\n')
for html1k=1:length(sub_dir)
    html1.QCdir = ['.' erase(sub_dir{html1k},home_dir(1:end-1)) 'QC/' '10_GM-WM-CSF-masks/'];
    html1.slicesstr = '';
    for html1k3=1:slices
        if exist([home_dir html1.QCdir(3:end) 'WMmask_slice' sprintf('%.2d',html1k3) '.jpg'],'file')
            html1.slicesstr = strcat(html1.slicesstr,sprintf(' <a href="%s">slice-%d</a>,',[html1.QCdir 'WMmask_slice' sprintf('%.2d',html1k3) '.jpg'],html1k3));
        else
            html1.slicesstr = strcat(html1.slicesstr,sprintf(' slice-%d,',html1k3));
        end
    end
    if exist([home_dir html1.QCdir(3:end) 'WMmask_slice' sprintf('%.2d',1) '.jpg'],'file')
        disp(sprintf('      Subject %d of %d - %s %s',html1k,length(sub_dir),subnames{html1k},html1.slicesstr(2:end-1)))
    else
        disp(sprintf('      Subject %d of %d - (pending execution)',html1k,length(sub_dir)))
    end
    clear html1.QCdir html1.slicesstr html1k3
end

fprintf('\n')
fprintf('<b>10.4. User-defined gray-matter masks:</b>\n')
for html1k=1:length(sub_dir)
    html1.QCdir = ['.' erase(sub_dir{html1k},home_dir(1:end-1)) 'QC/' '10_GM-WM-CSF-masks/'];
    html1.slicesstr = '';
    for html1k3=1:slices
        if exist([home_dir html1.QCdir(3:end) 'GMmask_slice' sprintf('%.2d',html1k3) '.jpg'],'file')
            html1.slicesstr = strcat(html1.slicesstr,sprintf(' <a href="%s">slice-%d</a>,',[html1.QCdir 'GMmask_slice' sprintf('%.2d',html1k3) '.jpg'],html1k3));
        else
            html1.slicesstr = strcat(html1.slicesstr,sprintf(' slice-%d,',html1k3));
        end
    end
    if exist([home_dir html1.QCdir(3:end) 'GMmask_slice' sprintf('%.2d',1) '.jpg'],'file')
        disp(sprintf('      Subject %d of %d - %s %s',html1k,length(sub_dir),subnames{html1k},html1.slicesstr(2:end-1)))
    else
        disp(sprintf('      Subject %d of %d - (pending execution)',html1k,length(sub_dir)))
    end
    clear html1.QCdir html1.slicesstr html1k3
end

fprintf('\n')
fprintf('<b>10.5. All masks final - full field of view:</b>\n')
for html1k=1:length(sub_dir)
    html1.QCdir = ['.' erase(sub_dir{html1k},home_dir(1:end-1)) 'QC/' '10_GM-WM-CSF-masks/'];
    html1.slicesstr = '';
    for html1k3=1:slices
        if exist([home_dir html1.QCdir(3:end) 'final_masks_slice' sprintf('%.2d',html1k3) '_fullFOV' '.jpg'],'file')
            html1.slicesstr = strcat(html1.slicesstr,sprintf(' <a href="%s">slice-%d</a>,',[html1.QCdir 'final_masks_slice' sprintf('%.2d',html1k3) '_fullFOV' '.jpg'],html1k3));
        else
            html1.slicesstr = strcat(html1.slicesstr,sprintf(' slice-%d,',html1k3));
        end
    end
    if exist([home_dir html1.QCdir(3:end) 'final_masks_slice' sprintf('%.2d',1) '_fullFOV' '.jpg'],'file')
        disp(sprintf('      Subject %d of %d - %s %s',html1k,length(sub_dir),subnames{html1k},html1.slicesstr(2:end-1)))
    else
        disp(sprintf('      Subject %d of %d - (pending execution)',html1k,length(sub_dir)))
    end
    clear html1.QCdir html1.slicesstr html1k3
end

fprintf('\n')
fprintf('<b>10.6. All masks final - zoomed view:</b>\n')
for html1k=1:length(sub_dir)
    html1.QCdir = ['.' erase(sub_dir{html1k},home_dir(1:end-1)) 'QC/' '10_GM-WM-CSF-masks/'];
    html1.slicesstr = '';
    for html1k3=1:slices
        if exist([home_dir html1.QCdir(3:end) 'final_masks_slice' sprintf('%.2d',html1k3) '_zoomed' '.jpg'],'file')
            html1.slicesstr = strcat(html1.slicesstr,sprintf(' <a href="%s">slice-%d</a>,',[html1.QCdir 'final_masks_slice' sprintf('%.2d',html1k3) '_zoomed' '.jpg'],html1k3));
        else
            html1.slicesstr = strcat(html1.slicesstr,sprintf(' slice-%d,',html1k3));
        end
    end
    if exist([home_dir html1.QCdir(3:end) 'final_masks_slice' sprintf('%.2d',1) '_zoomed' '.jpg'],'file')
        disp(sprintf('      Subject %d of %d - %s %s',html1k,length(sub_dir),subnames{html1k},html1.slicesstr(2:end-1)))
    else
        disp(sprintf('      Subject %d of %d - (pending execution)',html1k,length(sub_dir)))
    end
    clear html1.QCdir html1.slicesstr html1k3
end

else
fprintf('<b>Not in pipeline</b>\n')
fprintf('</p>')
end
end

%% 11. Co-registration (functional to anatomical)
if exist('choices_preproc','var')
if choices_preproc(11)==1
fprintf('<p style="font-size:14px ; color:#000000">')

fprintf('<b>11.1. Comparing co-registered mean functional images with the corresponding anatomical images:</b>\n')
for html1k=1:length(sub_dir)
    for html1k2=1:runs(html1k)
        if runs(html1k)==1
            html1.QCdir = ['.' erase(sub_dir{html1k},home_dir(1:end-1)) 'QC/' '11_func-anat-registration/'];
        else
            html1.QCdir = ['.' erase(sub_dir{html1k},home_dir(1:end-1)) 'QC/' '11_func-anat-registration/' 'run' num2str(html1k2) '/'];
        end
        html1.slicesstr = '';
        for html1k3=1:slices
            if exist([home_dir html1.QCdir(3:end) 'func_anat_reg_slice' sprintf('%.2d',html1k3) '.jpg'],'file')
                html1.slicesstr = strcat(html1.slicesstr,sprintf(' <a href="%s">slice-%d</a>,',[html1.QCdir 'func_anat_reg_slice' sprintf('%.2d',html1k3) '.jpg'],html1k3));
            else
                html1.slicesstr = strcat(html1.slicesstr,sprintf(' slice-%d,',html1k3));
            end
        end
        if exist([home_dir html1.QCdir(3:end) 'func_anat_reg_slice' sprintf('%.2d',1) '.jpg'],'file')
            if (html1k2==1) && (runs(html1k)==1)
                disp(sprintf('      Subject %d of %d - %s %s',html1k,length(sub_dir),subnames{html1k},html1.slicesstr(2:end-1)))
            elseif (html1k2==1) && (runs(html1k)~=1)
                disp(sprintf('      Subject %d of %d, run %d of %d - %s %s',html1k,length(sub_dir),html1k2,runs(html1k),subnames{html1k},html1.slicesstr(2:end-1)))
            else
                disp(sprintf('                      run %d of %d - %s %s',html1k2,runs(html1k),subnames{html1k},html1.slicesstr(2:end-1)))
            end
        else
            if (html1k2==1) && (runs(html1k)==1)
                disp(sprintf('      Subject %d of %d - (pending execution)',html1k,length(sub_dir)))
            elseif (html1k2==1) && (runs(html1k)~=1)
                disp(sprintf('      Subject %d of %d, run %d of %d - (pending execution)',html1k,length(sub_dir),html1k2,runs(html1k)))
            else
                disp(sprintf('                      run %d of %d - (pending execution)',html1k2,runs(html1k)))
            end
        end
    end
    clear html1.QCdir html1.slicesstr html1k3
end

fprintf('\n')
fprintf('<b>11.2. Time series within the gray and white matter after "11-functional-anatomical-registration":</b>\n')
for html1k=1:length(sub_dir)
    for html1k2=1:runs(html1k)
        if runs(html1k)==1
            html1.QCdir = ['.' erase(sub_dir{html1k},home_dir(1:end-1)) 'QC/' '11_func-anat-registration/'];
        else
            html1.QCdir = ['.' erase(sub_dir{html1k},home_dir(1:end-1)) 'QC/' '11_func-anat-registration/' 'run' num2str(html1k2) '/'];
        end
        if exist([home_dir html1.QCdir(3:end) 'fMRI_ts_11coreg_allSlices.jpg'],'file')
            if (html1k2==1) && (runs(html1k)==1)
                disp(sprintf('      Subject %d of %d - %s<a href="%s"> time series plot (all slices combined)</a>',html1k,length(sub_dir),subnames{html1k},[html1.QCdir 'fMRI_ts_11coreg_allSlices.jpg']))
            elseif (html1k2==1) && (runs(html1k)~=1)
                disp(sprintf('      Subject %d of %d, run %d of %d - %s<a href="%s"> time series plot (all slices combined)</a>',html1k,length(sub_dir),html1k2,runs(html1k),subnames{html1k},[html1.QCdir 'fMRI_ts_11coreg_allSlices.jpg']))
            else
                disp(sprintf('                      run %d of %d - %s<a href="%s"> time series plot (all slices combined)</a>',html1k2,runs(html1k),subnames{html1k},[html1.QCdir 'fMRI_ts_11coreg_allSlices.jpg']))
            end
        else
            if (html1k2==1) && (runs(html1k)==1)
                disp(sprintf('      Subject %d of %d - (pending execution)',html1k,length(sub_dir)))
            elseif (html1k2==1) && (runs(html1k)~=1)
                disp(sprintf('      Subject %d of %d, run %d of %d - (pending execution)',html1k,length(sub_dir),html1k2,runs(html1k)))
            else
                disp(sprintf('                      run %d of %d - (pending execution)',html1k2,runs(html1k)))
            end
        end
        html1.slicesstr = '';
        for html1k3=1:slices
            if exist([home_dir html1.QCdir(3:end) 'fMRI_ts_11coreg_slice' sprintf('%.2d',html1k3) '.jpg'],'file')
                html1.slicesstr = strcat(html1.slicesstr,sprintf(' <a href="%s">slice-%d</a>,',[html1.QCdir 'fMRI_ts_11coreg_slice' sprintf('%.2d',html1k3) '.jpg'],html1k3));
            else
                html1.slicesstr = strcat(html1.slicesstr,sprintf(' slice-%d,',html1k3));
            end
        end
        if exist([home_dir html1.QCdir(3:end) 'fMRI_ts_11coreg_slice' sprintf('%.2d',1) '.jpg'],'file')
            disp(sprintf('                                   (%s)',html1.slicesstr(2:end-1)))
        end
        clear html1.QCdir html1.slicesstr html1k3
    end
end

fprintf('\n')
fprintf('<b>11.3. Global mean signal within the gray and white matter after "11-functional-anatomical-registration":</b>\n')
for html1k=1:length(sub_dir)
    for html1k2=1:runs(html1k)
        if runs(html1k)==1
            html1.QCdir = ['.' erase(sub_dir{html1k},home_dir(1:end-1)) 'QC/' '11_func-anat-registration/'];
        else
            html1.QCdir = ['.' erase(sub_dir{html1k},home_dir(1:end-1)) 'QC/' '11_func-anat-registration/' 'run' num2str(html1k2) '/'];
        end
        if exist([home_dir html1.QCdir(3:end) 'global_signal_11coreg_allSlices.jpg'],'file')
            if (html1k2==1) && (runs(html1k)==1)
                disp(sprintf('      Subject %d of %d - %s<a href="%s"> global mean signal (all slices combined)</a>',html1k,length(sub_dir),subnames{html1k},[html1.QCdir 'global_signal_11coreg_allSlices.jpg']))
            elseif (html1k2==1) && (runs(html1k)~=1)
                disp(sprintf('      Subject %d of %d, run %d of %d - %s<a href="%s"> global mean signal (all slices combined)</a>',html1k,length(sub_dir),html1k2,runs(html1k),subnames{html1k},[html1.QCdir 'global_signal_11coreg_allSlices.jpg']))
            else
                disp(sprintf('                      run %d of %d - %s<a href="%s"> global mean signal (all slices combined)</a>',html1k2,runs(html1k),subnames{html1k},[html1.QCdir 'global_signal_11coreg_allSlices.jpg']))
            end
        else
            if (html1k2==1) && (runs(html1k)==1)
                disp(sprintf('      Subject %d of %d - (pending execution)',html1k,length(sub_dir)))
            elseif (html1k2==1) && (runs(html1k)~=1)
                disp(sprintf('      Subject %d of %d, run %d of %d - (pending execution)',html1k,length(sub_dir),html1k2,runs(html1k)))
            else
                disp(sprintf('                      run %d of %d - (pending execution)',html1k2,runs(html1k)))
            end
        end
        html1.slicesstr = '';
        for html1k3=1:slices
            if exist([home_dir html1.QCdir(3:end) 'global_signal_11coreg_slice' sprintf('%.2d',html1k3) '.jpg'],'file')
                html1.slicesstr = strcat(html1.slicesstr,sprintf(' <a href="%s">slice-%d</a>,',[html1.QCdir 'global_signal_11coreg_slice' sprintf('%.2d',html1k3) '.jpg'],html1k3));
            else
                html1.slicesstr = strcat(html1.slicesstr,sprintf(' slice-%d,',html1k3));
            end
        end
        if exist([home_dir html1.QCdir(3:end) 'global_signal_11coreg_slice' sprintf('%.2d',1) '.jpg'],'file')
            disp(sprintf('                                   (%s)',html1.slicesstr(2:end-1)))
        end
        clear html1.QCdir html1.slicesstr html1k3
    end
end

fprintf('\n')
fprintf('<b>11.4. Global mean signal: "11-functional-anatomical-registration" vs. "08-motion-correction":</b>\n')
for html1k=1:length(sub_dir)
    for html1k2=1:runs(html1k)
        if runs(html1k)==1
            html1.QCdir = ['.' erase(sub_dir{html1k},home_dir(1:end-1)) 'QC/' '11_func-anat-registration/'];
        else
            html1.QCdir = ['.' erase(sub_dir{html1k},home_dir(1:end-1)) 'QC/' '11_func-anat-registration/' 'run' num2str(html1k2) '/'];
        end
        if exist([home_dir html1.QCdir(3:end) 'global_signal_08moco_vs_11coreg_allSlices.jpg'],'file')
            if (html1k2==1) && (runs(html1k)==1)
                disp(sprintf('      Subject %d of %d - %s<a href="%s"> global mean signal (all slices combined)</a>',html1k,length(sub_dir),subnames{html1k},[html1.QCdir 'global_signal_08moco_vs_11coreg_allSlices.jpg']))
            elseif (html1k2==1) && (runs(html1k)~=1)
                disp(sprintf('      Subject %d of %d, run %d of %d - %s<a href="%s"> global mean signal (all slices combined)</a>',html1k,length(sub_dir),html1k2,runs(html1k),subnames{html1k},[html1.QCdir 'global_signal_08moco_vs_11coreg_allSlices.jpg']))
            else
                disp(sprintf('                      run %d of %d - %s<a href="%s"> global mean signal (all slices combined)</a>',html1k2,runs(html1k),subnames{html1k},[html1.QCdir 'global_signal_08moco_vs_11coreg_allSlices.jpg']))
            end
        else
            if (html1k2==1) && (runs(html1k)==1)
                disp(sprintf('      Subject %d of %d - (pending execution)',html1k,length(sub_dir)))
            elseif (html1k2==1) && (runs(html1k)~=1)
                disp(sprintf('      Subject %d of %d, run %d of %d - (pending execution)',html1k,length(sub_dir),html1k2,runs(html1k)))
            else
                disp(sprintf('                      run %d of %d - (pending execution)',html1k2,runs(html1k)))
            end
        end
        html1.slicesstr = '';
        for html1k3=1:slices
            if exist([home_dir html1.QCdir(3:end) 'global_signal_08moco_vs_11coreg_slice' sprintf('%.2d',html1k3) '.jpg'],'file')
                html1.slicesstr = strcat(html1.slicesstr,sprintf(' <a href="%s">slice-%d</a>,',[html1.QCdir 'global_signal_08moco_vs_11coreg_slice' sprintf('%.2d',html1k3) '.jpg'],html1k3));
            else
                html1.slicesstr = strcat(html1.slicesstr,sprintf(' slice-%d,',html1k3));
            end
        end
        if exist([home_dir html1.QCdir(3:end) 'global_signal_08moco_vs_11coreg_slice' sprintf('%.2d',1) '.jpg'],'file')
            disp(sprintf('                                   (%s)',html1.slicesstr(2:end-1)))
        end
        clear html1.QCdir html1.slicesstr html1k3
    end
end

fprintf('\n')
fprintf('<b>11.5. Histogram of voxel-to-voxel correlations within the gray and white matter after "11-functional-anatomical-registration":</b>\n')
for html1k=1:length(sub_dir)
    for html1k2=1:runs(html1k)
        if runs(html1k)==1
            html1.QCdir = ['.' erase(sub_dir{html1k},home_dir(1:end-1)) 'QC/' '11_func-anat-registration/'];
        else
            html1.QCdir = ['.' erase(sub_dir{html1k},home_dir(1:end-1)) 'QC/' '11_func-anat-registration/' 'run' num2str(html1k2) '/'];
        end
        if exist([home_dir html1.QCdir(3:end) 'hist_FC_compare_11coreg_allSlices.jpg'],'file')
            if (html1k2==1) && (runs(html1k)==1)
                disp(sprintf('      Subject %d of %d - %s<a href="%s"> histogram of functional connectivity (all slices combined)</a>',html1k,length(sub_dir),subnames{html1k},[html1.QCdir 'hist_FC_compare_11coreg_allSlices.jpg']))
            elseif (html1k2==1) && (runs(html1k)~=1)
                disp(sprintf('      Subject %d of %d, run %d of %d - %s<a href="%s"> histogram of functional connectivity (all slices combined)</a>',html1k,length(sub_dir),html1k2,runs(html1k),subnames{html1k},[html1.QCdir 'hist_FC_compare_11coreg_allSlices.jpg']))
            else
                disp(sprintf('                      run %d of %d - %s<a href="%s"> histogram of functional connectivity (all slices combined)</a>',html1k2,runs(html1k),subnames{html1k},[html1.QCdir 'hist_FC_compare_11coreg_allSlices.jpg']))
            end
        else
            if (html1k2==1) && (runs(html1k)==1)
                disp(sprintf('      Subject %d of %d - (pending execution)',html1k,length(sub_dir)))
            elseif (html1k2==1) && (runs(html1k)~=1)
                disp(sprintf('      Subject %d of %d, run %d of %d - (pending execution)',html1k,length(sub_dir),html1k2,runs(html1k)))
            else
                disp(sprintf('                      run %d of %d - (pending execution)',html1k2,runs(html1k)))
            end
        end
        html1.slicesstr = '';
        for html1k3=1:slices
            if exist([home_dir html1.QCdir(3:end) 'hist_FC_compare_11coreg_slice' sprintf('%.2d',html1k3) '.jpg'],'file')
                html1.slicesstr = strcat(html1.slicesstr,sprintf(' <a href="%s">slice-%d</a>,',[html1.QCdir 'hist_FC_compare_11coreg_slice' sprintf('%.2d',html1k3) '.jpg'],html1k3));
            else
                html1.slicesstr = strcat(html1.slicesstr,sprintf(' slice-%d,',html1k3));
            end
        end
        if exist([home_dir html1.QCdir(3:end) 'hist_FC_compare_11coreg_slice' sprintf('%.2d',1) '.jpg'],'file')
            disp(sprintf('                                   (%s)',html1.slicesstr(2:end-1)))
        end
        clear html1.QCdir html1.slicesstr html1k3
    end
end

fprintf('\n')
fprintf('<b>11.6. Histogram of voxel-to-voxel correlations: "11-functional-anatomical-registration" vs. "08-motion-correction":</b>\n')
for html1k=1:length(sub_dir)
    for html1k2=1:runs(html1k)
        if runs(html1k)==1
            html1.QCdir = ['.' erase(sub_dir{html1k},home_dir(1:end-1)) 'QC/' '11_func-anat-registration/'];
        else
            html1.QCdir = ['.' erase(sub_dir{html1k},home_dir(1:end-1)) 'QC/' '11_func-anat-registration/' 'run' num2str(html1k2) '/'];
        end
        if exist([home_dir html1.QCdir(3:end) 'hist_FC_compare_08moco_11coreg_allSlices.jpg'],'file')
            if (html1k2==1) && (runs(html1k)==1)
                disp(sprintf('      Subject %d of %d - %s<a href="%s"> histogram of functional connectivity (all slices combined)</a>',html1k,length(sub_dir),subnames{html1k},[html1.QCdir 'hist_FC_compare_08moco_11coreg_allSlices.jpg']))
            elseif (html1k2==1) && (runs(html1k)~=1)
                disp(sprintf('      Subject %d of %d, run %d of %d - %s<a href="%s"> histogram of functional connectivity (all slices combined)</a>',html1k,length(sub_dir),html1k2,runs(html1k),subnames{html1k},[html1.QCdir 'hist_FC_compare_08moco_11coreg_allSlices.jpg']))
            else
                disp(sprintf('                      run %d of %d - %s<a href="%s"> histogram of functional connectivity (all slices combined)</a>',html1k2,runs(html1k),subnames{html1k},[html1.QCdir 'hist_FC_compare_08moco_11coreg_allSlices.jpg']))
            end
        else
            if (html1k2==1) && (runs(html1k)==1)
                disp(sprintf('      Subject %d of %d - (pending execution)',html1k,length(sub_dir)))
            elseif (html1k2==1) && (runs(html1k)~=1)
                disp(sprintf('      Subject %d of %d, run %d of %d - (pending execution)',html1k,length(sub_dir),html1k2,runs(html1k)))
            else
                disp(sprintf('                      run %d of %d - (pending execution)',html1k2,runs(html1k)))
            end
        end
        html1.slicesstr = '';
        for html1k3=1:slices
            if exist([home_dir html1.QCdir(3:end) 'hist_FC_compare_08moco_11coreg_slice' sprintf('%.2d',html1k3) '.jpg'],'file')
                html1.slicesstr = strcat(html1.slicesstr,sprintf(' <a href="%s">slice-%d</a>,',[html1.QCdir 'hist_FC_compare_08moco_11coreg_slice' sprintf('%.2d',html1k3) '.jpg'],html1k3));
            else
                html1.slicesstr = strcat(html1.slicesstr,sprintf(' slice-%d,',html1k3));
            end
        end
        if exist([home_dir html1.QCdir(3:end) 'hist_FC_compare_08moco_11coreg_slice' sprintf('%.2d',1) '.jpg'],'file')
            disp(sprintf('                                   (%s)',html1.slicesstr(2:end-1)))
        end
        clear html1.QCdir html1.slicesstr html1k3
    end
end

if ~isempty(which('VideoWriter')) && (save_videos_steps(5)==1) % check if Matlab's audio video toolbox exists
fprintf('\n')
fprintf('<b>11.7. FMRI video of data after "11-functional-anatomical-registration":</b>\n')
for html1k=1:length(sub_dir)
    for html1k2=1:runs(html1k)
        if runs(html1k)==1
            html1.QCdir = ['.' erase(sub_dir{html1k},home_dir(1:end-1)) 'QC/' '11_func-anat-registration/'];
        else
            html1.QCdir = ['.' erase(sub_dir{html1k},home_dir(1:end-1)) 'QC/' '11_func-anat-registration/' 'run' num2str(html1k2) '/'];
        end
        html1.slicesstr = '';
        for html1k3=1:slices
            if exist([home_dir html1.QCdir(3:end) 'fMRIvideo_11coreg_slice' sprintf('%.2d',html1k3) '.mp4'],'file')
                html1.slicesstr = strcat(html1.slicesstr,sprintf(' <a href="%s">slice-%d</a>,',[html1.QCdir 'fMRIvideo_11coreg_slice' sprintf('%.2d',html1k3) '.mp4'],html1k3));
            else
                html1.slicesstr = strcat(html1.slicesstr,sprintf(' slice-%d,',html1k3));
            end
        end
        if exist([home_dir html1.QCdir(3:end) 'fMRIvideo_11coreg_slice' sprintf('%.2d',1) '.mp4'],'file')
            if (html1k2==1) && (runs(html1k)==1)
                disp(sprintf('      Subject %d of %d - %s %s',html1k,length(sub_dir),subnames{html1k},html1.slicesstr(2:end-1)))
            elseif (html1k2==1) && (runs(html1k)~=1)
                disp(sprintf('      Subject %d of %d, run %d of %d - %s %s',html1k,length(sub_dir),html1k2,runs(html1k),subnames{html1k},html1.slicesstr(2:end-1)))
            else
                disp(sprintf('                      run %d of %d - %s %s',html1k2,runs(html1k),subnames{html1k},html1.slicesstr(2:end-1)))
            end
        else
            if (html1k2==1) && (runs(html1k)==1)
                disp(sprintf('      Subject %d of %d - (pending execution)',html1k,length(sub_dir)))
            elseif (html1k2==1) && (runs(html1k)~=1)
                disp(sprintf('      Subject %d of %d, run %d of %d - (pending execution)',html1k,length(sub_dir),html1k2,runs(html1k)))
            else
                disp(sprintf('                      run %d of %d - (pending execution)',html1k2,runs(html1k)))
            end
        end
    end
    clear html1.QCdir html1.slicesstr html1k3
end
end

else
fprintf('<b>Not in pipeline</b>\n')
fprintf('</p>')
end
end

%% 12. Denoise-3 (cerebrospinal fluid signal regression)
if exist('choices_preproc','var')
if choices_preproc(12)==1
fprintf('<p style="font-size:14px ; color:#000000">')

fprintf('<b>12.1. Time series within the gray and white matter after "12-denoise3-CSFregress":</b>\n')
for html1k=1:length(sub_dir)
    for html1k2=1:runs(html1k)
        if runs(html1k)==1
            html1.QCdir = ['.' erase(sub_dir{html1k},home_dir(1:end-1)) 'QC/' '12_denoise3-CSFregress/'];
        else
            html1.QCdir = ['.' erase(sub_dir{html1k},home_dir(1:end-1)) 'QC/' '12_denoise3-CSFregress/' 'run' num2str(html1k2) '/'];
        end
        if exist([home_dir html1.QCdir(3:end) 'fMRI_ts_12denoise3_allSlices.jpg'],'file')
            if (html1k2==1) && (runs(html1k)==1)
                disp(sprintf('      Subject %d of %d - %s<a href="%s"> time series plot (all slices combined)</a>',html1k,length(sub_dir),subnames{html1k},[html1.QCdir 'fMRI_ts_12denoise3_allSlices.jpg']))
            elseif (html1k2==1) && (runs(html1k)~=1)
                disp(sprintf('      Subject %d of %d, run %d of %d - %s<a href="%s"> time series plot (all slices combined)</a>',html1k,length(sub_dir),html1k2,runs(html1k),subnames{html1k},[html1.QCdir 'fMRI_ts_12denoise3_allSlices.jpg']))
            else
                disp(sprintf('                      run %d of %d - %s<a href="%s"> time series plot (all slices combined)</a>',html1k2,runs(html1k),subnames{html1k},[html1.QCdir 'fMRI_ts_12denoise3_allSlices.jpg']))
            end
        else
            if (html1k2==1) && (runs(html1k)==1)
                disp(sprintf('      Subject %d of %d - (pending execution)',html1k,length(sub_dir)))
            elseif (html1k2==1) && (runs(html1k)~=1)
                disp(sprintf('      Subject %d of %d, run %d of %d - (pending execution)',html1k,length(sub_dir),html1k2,runs(html1k)))
            else
                disp(sprintf('                      run %d of %d - (pending execution)',html1k2,runs(html1k)))
            end
        end
        html1.slicesstr = '';
        for html1k3=1:slices
            if exist([home_dir html1.QCdir(3:end) 'fMRI_ts_12denoise3_slice' sprintf('%.2d',html1k3) '.jpg'],'file')
                html1.slicesstr = strcat(html1.slicesstr,sprintf(' <a href="%s">slice-%d</a>,',[html1.QCdir 'fMRI_ts_12denoise3_slice' sprintf('%.2d',html1k3) '.jpg'],html1k3));
            else
                html1.slicesstr = strcat(html1.slicesstr,sprintf(' slice-%d,',html1k3));
            end
        end
        if exist([home_dir html1.QCdir(3:end) 'fMRI_ts_12denoise3_slice' sprintf('%.2d',1) '.jpg'],'file')
            disp(sprintf('                                   (%s)',html1.slicesstr(2:end-1)))
        end
        clear html1.QCdir html1.slicesstr html1k3
    end
end

fprintf('\n')
fprintf('<b>12.2. Global mean signal within the gray and white matter after "12-denoise3-CSFregress":</b>\n')
for html1k=1:length(sub_dir)
    for html1k2=1:runs(html1k)
        if runs(html1k)==1
            html1.QCdir = ['.' erase(sub_dir{html1k},home_dir(1:end-1)) 'QC/' '12_denoise3-CSFregress/'];
        else
            html1.QCdir = ['.' erase(sub_dir{html1k},home_dir(1:end-1)) 'QC/' '12_denoise3-CSFregress/' 'run' num2str(html1k2) '/'];
        end
        if exist([home_dir html1.QCdir(3:end) 'global_signal_12denoise3_allSlices.jpg'],'file')
            if (html1k2==1) && (runs(html1k)==1)
                disp(sprintf('      Subject %d of %d - %s<a href="%s"> global mean signal (all slices combined)</a>',html1k,length(sub_dir),subnames{html1k},[html1.QCdir 'global_signal_12denoise3_allSlices.jpg']))
            elseif (html1k2==1) && (runs(html1k)~=1)
                disp(sprintf('      Subject %d of %d, run %d of %d - %s<a href="%s"> global mean signal (all slices combined)</a>',html1k,length(sub_dir),html1k2,runs(html1k),subnames{html1k},[html1.QCdir 'global_signal_12denoise3_allSlices.jpg']))
            else
                disp(sprintf('                      run %d of %d - %s<a href="%s"> global mean signal (all slices combined)</a>',html1k2,runs(html1k),subnames{html1k},[html1.QCdir 'global_signal_12denoise3_allSlices.jpg']))
            end
        else
            if (html1k2==1) && (runs(html1k)==1)
                disp(sprintf('      Subject %d of %d - (pending execution)',html1k,length(sub_dir)))
            elseif (html1k2==1) && (runs(html1k)~=1)
                disp(sprintf('      Subject %d of %d, run %d of %d - (pending execution)',html1k,length(sub_dir),html1k2,runs(html1k)))
            else
                disp(sprintf('                      run %d of %d - (pending execution)',html1k2,runs(html1k)))
            end
        end
        html1.slicesstr = '';
        for html1k3=1:slices
            if exist([home_dir html1.QCdir(3:end) 'global_signal_12denoise3_slice' sprintf('%.2d',html1k3) '.jpg'],'file')
                html1.slicesstr = strcat(html1.slicesstr,sprintf(' <a href="%s">slice-%d</a>,',[html1.QCdir 'global_signal_12denoise3_slice' sprintf('%.2d',html1k3) '.jpg'],html1k3));
            else
                html1.slicesstr = strcat(html1.slicesstr,sprintf(' slice-%d,',html1k3));
            end
        end
        if exist([home_dir html1.QCdir(3:end) 'global_signal_12denoise3_slice' sprintf('%.2d',1) '.jpg'],'file')
            disp(sprintf('                                   (%s)',html1.slicesstr(2:end-1)))
        end
        clear html1.QCdir html1.slicesstr html1k3
    end
end

fprintf('\n')
fprintf('<b>12.3. Global mean signal: "12-denoise3-CSFregress" vs. "11-func-anat-registration":</b>\n')
for html1k=1:length(sub_dir)
    for html1k2=1:runs(html1k)
        if runs(html1k)==1
            html1.QCdir = ['.' erase(sub_dir{html1k},home_dir(1:end-1)) 'QC/' '12_denoise3-CSFregress/'];
        else
            html1.QCdir = ['.' erase(sub_dir{html1k},home_dir(1:end-1)) 'QC/' '12_denoise3-CSFregress/' 'run' num2str(html1k2) '/'];
        end
        if exist([home_dir html1.QCdir(3:end) 'global_signal_11coreg_vs_12denoise3_allSlices.jpg'],'file')
            if (html1k2==1) && (runs(html1k)==1)
                disp(sprintf('      Subject %d of %d - %s<a href="%s"> global mean signal (all slices combined)</a>',html1k,length(sub_dir),subnames{html1k},[html1.QCdir 'global_signal_11coreg_vs_12denoise3_allSlices.jpg']))
            elseif (html1k2==1) && (runs(html1k)~=1)
                disp(sprintf('      Subject %d of %d, run %d of %d - %s<a href="%s"> global mean signal (all slices combined)</a>',html1k,length(sub_dir),html1k2,runs(html1k),subnames{html1k},[html1.QCdir 'global_signal_11coreg_vs_12denoise3_allSlices.jpg']))
            else
                disp(sprintf('                      run %d of %d - %s<a href="%s"> global mean signal (all slices combined)</a>',html1k2,runs(html1k),subnames{html1k},[html1.QCdir 'global_signal_11coreg_vs_12denoise3_allSlices.jpg']))
            end
        else
            if (html1k2==1) && (runs(html1k)==1)
                disp(sprintf('      Subject %d of %d - (pending execution)',html1k,length(sub_dir)))
            elseif (html1k2==1) && (runs(html1k)~=1)
                disp(sprintf('      Subject %d of %d, run %d of %d - (pending execution)',html1k,length(sub_dir),html1k2,runs(html1k)))
            else
                disp(sprintf('                      run %d of %d - (pending execution)',html1k2,runs(html1k)))
            end
        end
        html1.slicesstr = '';
        for html1k3=1:slices
            if exist([home_dir html1.QCdir(3:end) 'global_signal_11coreg_vs_12denoise3_slice' sprintf('%.2d',html1k3) '.jpg'],'file')
                html1.slicesstr = strcat(html1.slicesstr,sprintf(' <a href="%s">slice-%d</a>,',[html1.QCdir 'global_signal_11coreg_vs_12denoise3_slice' sprintf('%.2d',html1k3) '.jpg'],html1k3));
            else
                html1.slicesstr = strcat(html1.slicesstr,sprintf(' slice-%d,',html1k3));
            end
        end
        if exist([home_dir html1.QCdir(3:end) 'global_signal_11coreg_vs_12denoise3_slice' sprintf('%.2d',1) '.jpg'],'file')
            disp(sprintf('                                   (%s)',html1.slicesstr(2:end-1)))
        end
        clear html1.QCdir html1.slicesstr html1k3
    end
end

fprintf('\n')
fprintf('<b>12.4. Histogram of voxel-to-voxel correlations within the gray and white matter after "12-denoise3-CSFregress":</b>\n')
for html1k=1:length(sub_dir)
    for html1k2=1:runs(html1k)
        if runs(html1k)==1
            html1.QCdir = ['.' erase(sub_dir{html1k},home_dir(1:end-1)) 'QC/' '12_denoise3-CSFregress/'];
        else
            html1.QCdir = ['.' erase(sub_dir{html1k},home_dir(1:end-1)) 'QC/' '12_denoise3-CSFregress/' 'run' num2str(html1k2) '/'];
        end
        if exist([home_dir html1.QCdir(3:end) 'hist_FC_compare_12denoise3_allSlices.jpg'],'file')
            if (html1k2==1) && (runs(html1k)==1)
                disp(sprintf('      Subject %d of %d - %s<a href="%s"> histogram of functional connectivity (all slices combined)</a>',html1k,length(sub_dir),subnames{html1k},[html1.QCdir 'hist_FC_compare_12denoise3_allSlices.jpg']))
            elseif (html1k2==1) && (runs(html1k)~=1)
                disp(sprintf('      Subject %d of %d, run %d of %d - %s<a href="%s"> histogram of functional connectivity (all slices combined)</a>',html1k,length(sub_dir),html1k2,runs(html1k),subnames{html1k},[html1.QCdir 'hist_FC_compare_12denoise3_allSlices.jpg']))
            else
                disp(sprintf('                      run %d of %d - %s<a href="%s"> histogram of functional connectivity (all slices combined)</a>',html1k2,runs(html1k),subnames{html1k},[html1.QCdir 'hist_FC_compare_12denoise3_allSlices.jpg']))
            end
        else
            if (html1k2==1) && (runs(html1k)==1)
                disp(sprintf('      Subject %d of %d - (pending execution)',html1k,length(sub_dir)))
            elseif (html1k2==1) && (runs(html1k)~=1)
                disp(sprintf('      Subject %d of %d, run %d of %d - (pending execution)',html1k,length(sub_dir),html1k2,runs(html1k)))
            else
                disp(sprintf('                      run %d of %d - (pending execution)',html1k2,runs(html1k)))
            end
        end
        html1.slicesstr = '';
        for html1k3=1:slices
            if exist([home_dir html1.QCdir(3:end) 'hist_FC_compare_12denoise3_slice' sprintf('%.2d',html1k3) '.jpg'],'file')
                html1.slicesstr = strcat(html1.slicesstr,sprintf(' <a href="%s">slice-%d</a>,',[html1.QCdir 'hist_FC_compare_12denoise3_slice' sprintf('%.2d',html1k3) '.jpg'],html1k3));
            else
                html1.slicesstr = strcat(html1.slicesstr,sprintf(' slice-%d,',html1k3));
            end
        end
        if exist([home_dir html1.QCdir(3:end) 'hist_FC_compare_12denoise3_slice' sprintf('%.2d',1) '.jpg'],'file')
            disp(sprintf('                                   (%s)',html1.slicesstr(2:end-1)))
        end
        clear html1.QCdir html1.slicesstr html1k3
    end
end

fprintf('\n')
fprintf('<b>12.5. Histogram of voxel-to-voxel correlations: "12-denoise3-CSFregress" vs. "11-func-anat-registration":</b>\n')
for html1k=1:length(sub_dir)
    for html1k2=1:runs(html1k)
        if runs(html1k)==1
            html1.QCdir = ['.' erase(sub_dir{html1k},home_dir(1:end-1)) 'QC/' '12_denoise3-CSFregress/'];
        else
            html1.QCdir = ['.' erase(sub_dir{html1k},home_dir(1:end-1)) 'QC/' '12_denoise3-CSFregress/' 'run' num2str(html1k2) '/'];
        end
        if exist([home_dir html1.QCdir(3:end) 'hist_FC_compare_11coreg_12denoise3_allSlices.jpg'],'file')
            if (html1k2==1) && (runs(html1k)==1)
                disp(sprintf('      Subject %d of %d - %s<a href="%s"> histogram of functional connectivity (all slices combined)</a>',html1k,length(sub_dir),subnames{html1k},[html1.QCdir 'hist_FC_compare_11coreg_12denoise3_allSlices.jpg']))
            elseif (html1k2==1) && (runs(html1k)~=1)
                disp(sprintf('      Subject %d of %d, run %d of %d - %s<a href="%s"> histogram of functional connectivity (all slices combined)</a>',html1k,length(sub_dir),html1k2,runs(html1k),subnames{html1k},[html1.QCdir 'hist_FC_compare_11coreg_12denoise3_allSlices.jpg']))
            else
                disp(sprintf('                      run %d of %d - %s<a href="%s"> histogram of functional connectivity (all slices combined)</a>',html1k2,runs(html1k),subnames{html1k},[html1.QCdir 'hist_FC_compare_11coreg_12denoise3_allSlices.jpg']))
            end
        else
            if (html1k2==1) && (runs(html1k)==1)
                disp(sprintf('      Subject %d of %d - (pending execution)',html1k,length(sub_dir)))
            elseif (html1k2==1) && (runs(html1k)~=1)
                disp(sprintf('      Subject %d of %d, run %d of %d - (pending execution)',html1k,length(sub_dir),html1k2,runs(html1k)))
            else
                disp(sprintf('                      run %d of %d - (pending execution)',html1k2,runs(html1k)))
            end
        end
        html1.slicesstr = '';
        for html1k3=1:slices
            if exist([home_dir html1.QCdir(3:end) 'hist_FC_compare_11coreg_12denoise3_slice' sprintf('%.2d',html1k3) '.jpg'],'file')
                html1.slicesstr = strcat(html1.slicesstr,sprintf(' <a href="%s">slice-%d</a>,',[html1.QCdir 'hist_FC_compare_11coreg_12denoise3_slice' sprintf('%.2d',html1k3) '.jpg'],html1k3));
            else
                html1.slicesstr = strcat(html1.slicesstr,sprintf(' slice-%d,',html1k3));
            end
        end
        if exist([home_dir html1.QCdir(3:end) 'hist_FC_compare_11coreg_12denoise3_slice' sprintf('%.2d',1) '.jpg'],'file')
            disp(sprintf('                                   (%s)',html1.slicesstr(2:end-1)))
        end
        clear html1.QCdir html1.slicesstr html1k3
    end
end

if ~isempty(which('VideoWriter')) && (save_videos_steps(6)==1) % check if Matlab's audio video toolbox exists
fprintf('\n')
fprintf('<b>12.6. FMRI video of data after "12-denoise3-CSFregress":</b>\n')
for html1k=1:length(sub_dir)
    for html1k2=1:runs(html1k)
        if runs(html1k)==1
            html1.QCdir = ['.' erase(sub_dir{html1k},home_dir(1:end-1)) 'QC/' '12_denoise3-CSFregress/'];
        else
            html1.QCdir = ['.' erase(sub_dir{html1k},home_dir(1:end-1)) 'QC/' '12_denoise3-CSFregress/' 'run' num2str(html1k2) '/'];
        end
        html1.slicesstr = '';
        for html1k3=1:slices
            if exist([home_dir html1.QCdir(3:end) 'fMRIvideo_12denoise3_slice' sprintf('%.2d',html1k3) '.mp4'],'file')
                html1.slicesstr = strcat(html1.slicesstr,sprintf(' <a href="%s">slice-%d</a>,',[html1.QCdir 'fMRIvideo_12denoise3_slice' sprintf('%.2d',html1k3) '.mp4'],html1k3));
            else
                html1.slicesstr = strcat(html1.slicesstr,sprintf(' slice-%d,',html1k3));
            end
        end
        if exist([home_dir html1.QCdir(3:end) 'fMRIvideo_12denoise3_slice' sprintf('%.2d',1) '.mp4'],'file')
            if (html1k2==1) && (runs(html1k)==1)
                disp(sprintf('      Subject %d of %d - %s %s',html1k,length(sub_dir),subnames{html1k},html1.slicesstr(2:end-1)))
            elseif (html1k2==1) && (runs(html1k)~=1)
                disp(sprintf('      Subject %d of %d, run %d of %d - %s %s',html1k,length(sub_dir),html1k2,runs(html1k),subnames{html1k},html1.slicesstr(2:end-1)))
            else
                disp(sprintf('                      run %d of %d - %s %s',html1k2,runs(html1k),subnames{html1k},html1.slicesstr(2:end-1)))
            end
        else
            if (html1k2==1) && (runs(html1k)==1)
                disp(sprintf('      Subject %d of %d - (pending execution)',html1k,length(sub_dir)))
            elseif (html1k2==1) && (runs(html1k)~=1)
                disp(sprintf('      Subject %d of %d, run %d of %d - (pending execution)',html1k,length(sub_dir),html1k2,runs(html1k)))
            else
                disp(sprintf('                      run %d of %d - (pending execution)',html1k2,runs(html1k)))
            end
        end
    end
    clear html1.QCdir html1.slicesstr html1k3
end
end

else
fprintf('<b>Not in pipeline</b>\n')
fprintf('</p>')
end
end

%% 13. Denoise-4 (white-matter signal regression)
if exist('choices_preproc','var')
if choices_preproc(13)==1
fprintf('<p style="font-size:14px ; color:#000000">')

fprintf('<b>13.1. Time series within the gray and white matter after "13-denoise4-WMregress":</b>\n')
for html1k=1:length(sub_dir)
    for html1k2=1:runs(html1k)
        if runs(html1k)==1
            html1.QCdir = ['.' erase(sub_dir{html1k},home_dir(1:end-1)) 'QC/' '13_denoise4-WMregress/'];
        else
            html1.QCdir = ['.' erase(sub_dir{html1k},home_dir(1:end-1)) 'QC/' '13_denoise4-WMregress/' 'run' num2str(html1k2) '/'];
        end
        if exist([home_dir html1.QCdir(3:end) 'fMRI_ts_13denoise4_allSlices.jpg'],'file')
            if (html1k2==1) && (runs(html1k)==1)
                disp(sprintf('      Subject %d of %d - %s<a href="%s"> time series plot (all slices combined)</a>',html1k,length(sub_dir),subnames{html1k},[html1.QCdir 'fMRI_ts_13denoise4_allSlices.jpg']))
            elseif (html1k2==1) && (runs(html1k)~=1)
                disp(sprintf('      Subject %d of %d, run %d of %d - %s<a href="%s"> time series plot (all slices combined)</a>',html1k,length(sub_dir),html1k2,runs(html1k),subnames{html1k},[html1.QCdir 'fMRI_ts_13denoise4_allSlices.jpg']))
            else
                disp(sprintf('                      run %d of %d - %s<a href="%s"> time series plot (all slices combined)</a>',html1k2,runs(html1k),subnames{html1k},[html1.QCdir 'fMRI_ts_13denoise4_allSlices.jpg']))
            end
        else
            if (html1k2==1) && (runs(html1k)==1)
                disp(sprintf('      Subject %d of %d - (pending execution)',html1k,length(sub_dir)))
            elseif (html1k2==1) && (runs(html1k)~=1)
                disp(sprintf('      Subject %d of %d, run %d of %d - (pending execution)',html1k,length(sub_dir),html1k2,runs(html1k)))
            else
                disp(sprintf('                      run %d of %d - (pending execution)',html1k2,runs(html1k)))
            end
        end
        html1.slicesstr = '';
        for html1k3=1:slices
            if exist([home_dir html1.QCdir(3:end) 'fMRI_ts_13denoise4_slice' sprintf('%.2d',html1k3) '.jpg'],'file')
                html1.slicesstr = strcat(html1.slicesstr,sprintf(' <a href="%s">slice-%d</a>,',[html1.QCdir 'fMRI_ts_13denoise4_slice' sprintf('%.2d',html1k3) '.jpg'],html1k3));
            else
                html1.slicesstr = strcat(html1.slicesstr,sprintf(' slice-%d,',html1k3));
            end
        end
        if exist([home_dir html1.QCdir(3:end) 'fMRI_ts_13denoise4_slice' sprintf('%.2d',1) '.jpg'],'file')
            disp(sprintf('                                   (%s)',html1.slicesstr(2:end-1)))
        end
        clear html1.QCdir html1.slicesstr html1k3
    end
end

fprintf('\n')
fprintf('<b>13.2. Global mean signal within the gray and white matter after "13-denoise4-WMregress":</b>\n')
for html1k=1:length(sub_dir)
    for html1k2=1:runs(html1k)
        if runs(html1k)==1
            html1.QCdir = ['.' erase(sub_dir{html1k},home_dir(1:end-1)) 'QC/' '13_denoise4-WMregress/'];
        else
            html1.QCdir = ['.' erase(sub_dir{html1k},home_dir(1:end-1)) 'QC/' '13_denoise4-WMregress/' 'run' num2str(html1k2) '/'];
        end
        if exist([home_dir html1.QCdir(3:end) 'global_signal_13denoise4_allSlices.jpg'],'file')
            if (html1k2==1) && (runs(html1k)==1)
                disp(sprintf('      Subject %d of %d - %s<a href="%s"> global mean signal (all slices combined)</a>',html1k,length(sub_dir),subnames{html1k},[html1.QCdir 'global_signal_13denoise4_allSlices.jpg']))
            elseif (html1k2==1) && (runs(html1k)~=1)
                disp(sprintf('      Subject %d of %d, run %d of %d - %s<a href="%s"> global mean signal (all slices combined)</a>',html1k,length(sub_dir),html1k2,runs(html1k),subnames{html1k},[html1.QCdir 'global_signal_13denoise4_allSlices.jpg']))
            else
                disp(sprintf('                      run %d of %d - %s<a href="%s"> global mean signal (all slices combined)</a>',html1k2,runs(html1k),subnames{html1k},[html1.QCdir 'global_signal_13denoise4_allSlices.jpg']))
            end
        else
            if (html1k2==1) && (runs(html1k)==1)
                disp(sprintf('      Subject %d of %d - (pending execution)',html1k,length(sub_dir)))
            elseif (html1k2==1) && (runs(html1k)~=1)
                disp(sprintf('      Subject %d of %d, run %d of %d - (pending execution)',html1k,length(sub_dir),html1k2,runs(html1k)))
            else
                disp(sprintf('                      run %d of %d - (pending execution)',html1k2,runs(html1k)))
            end
        end
        html1.slicesstr = '';
        for html1k3=1:slices
            if exist([home_dir html1.QCdir(3:end) 'global_signal_13denoise4_slice' sprintf('%.2d',html1k3) '.jpg'],'file')
                html1.slicesstr = strcat(html1.slicesstr,sprintf(' <a href="%s">slice-%d</a>,',[html1.QCdir 'global_signal_13denoise4_slice' sprintf('%.2d',html1k3) '.jpg'],html1k3));
            else
                html1.slicesstr = strcat(html1.slicesstr,sprintf(' slice-%d,',html1k3));
            end
        end
        if exist([home_dir html1.QCdir(3:end) 'global_signal_13denoise4_slice' sprintf('%.2d',1) '.jpg'],'file')
            disp(sprintf('                                   (%s)',html1.slicesstr(2:end-1)))
        end
        clear html1.QCdir html1.slicesstr html1k3
    end
end

fprintf('\n')
fprintf('<b>13.3. Global mean signal: "13-denoise4-WMregress" vs. "12-denoise3-CSFregress":</b>\n')
for html1k=1:length(sub_dir)
    for html1k2=1:runs(html1k)
        if runs(html1k)==1
            html1.QCdir = ['.' erase(sub_dir{html1k},home_dir(1:end-1)) 'QC/' '13_denoise4-WMregress/'];
        else
            html1.QCdir = ['.' erase(sub_dir{html1k},home_dir(1:end-1)) 'QC/' '13_denoise4-WMregress/' 'run' num2str(html1k2) '/'];
        end
        if exist([home_dir html1.QCdir(3:end) 'global_signal_12denoise3_vs_13denoise4_allSlices.jpg'],'file')
            if (html1k2==1) && (runs(html1k)==1)
                disp(sprintf('      Subject %d of %d - %s<a href="%s"> global mean signal (all slices combined)</a>',html1k,length(sub_dir),subnames{html1k},[html1.QCdir 'global_signal_12denoise3_vs_13denoise4_allSlices.jpg']))
            elseif (html1k2==1) && (runs(html1k)~=1)
                disp(sprintf('      Subject %d of %d, run %d of %d - %s<a href="%s"> global mean signal (all slices combined)</a>',html1k,length(sub_dir),html1k2,runs(html1k),subnames{html1k},[html1.QCdir 'global_signal_12denoise3_vs_13denoise4_allSlices.jpg']))
            else
                disp(sprintf('                      run %d of %d - %s<a href="%s"> global mean signal (all slices combined)</a>',html1k2,runs(html1k),subnames{html1k},[html1.QCdir 'global_signal_12denoise3_vs_13denoise4_allSlices.jpg']))
            end
        else
            if (html1k2==1) && (runs(html1k)==1)
                disp(sprintf('      Subject %d of %d - (pending execution)',html1k,length(sub_dir)))
            elseif (html1k2==1) && (runs(html1k)~=1)
                disp(sprintf('      Subject %d of %d, run %d of %d - (pending execution)',html1k,length(sub_dir),html1k2,runs(html1k)))
            else
                disp(sprintf('                      run %d of %d - (pending execution)',html1k2,runs(html1k)))
            end
        end
        html1.slicesstr = '';
        for html1k3=1:slices
            if exist([home_dir html1.QCdir(3:end) 'global_signal_12denoise3_vs_13denoise4_slice' sprintf('%.2d',html1k3) '.jpg'],'file')
                html1.slicesstr = strcat(html1.slicesstr,sprintf(' <a href="%s">slice-%d</a>,',[html1.QCdir 'global_signal_12denoise3_vs_13denoise4_slice' sprintf('%.2d',html1k3) '.jpg'],html1k3));
            else
                html1.slicesstr = strcat(html1.slicesstr,sprintf(' slice-%d,',html1k3));
            end
        end
        if exist([home_dir html1.QCdir(3:end) 'global_signal_12denoise3_vs_13denoise4_slice' sprintf('%.2d',1) '.jpg'],'file')
            disp(sprintf('                                   (%s)',html1.slicesstr(2:end-1)))
        end
        clear html1.QCdir html1.slicesstr html1k3
    end
end

fprintf('\n')
fprintf('<b>13.4. Histogram of voxel-to-voxel correlations within the gray and white matter after "13-denoise4-WMregress":</b>\n')
for html1k=1:length(sub_dir)
    for html1k2=1:runs(html1k)
        if runs(html1k)==1
            html1.QCdir = ['.' erase(sub_dir{html1k},home_dir(1:end-1)) 'QC/' '13_denoise4-WMregress/'];
        else
            html1.QCdir = ['.' erase(sub_dir{html1k},home_dir(1:end-1)) 'QC/' '13_denoise4-WMregress/' 'run' num2str(html1k2) '/'];
        end
        if exist([home_dir html1.QCdir(3:end) 'hist_FC_compare_13denoise4_allSlices.jpg'],'file')
            if (html1k2==1) && (runs(html1k)==1)
                disp(sprintf('      Subject %d of %d - %s<a href="%s"> histogram of functional connectivity (all slices combined)</a>',html1k,length(sub_dir),subnames{html1k},[html1.QCdir 'hist_FC_compare_13denoise4_allSlices.jpg']))
            elseif (html1k2==1) && (runs(html1k)~=1)
                disp(sprintf('      Subject %d of %d, run %d of %d - %s<a href="%s"> histogram of functional connectivity (all slices combined)</a>',html1k,length(sub_dir),html1k2,runs(html1k),subnames{html1k},[html1.QCdir 'hist_FC_compare_13denoise4_allSlices.jpg']))
            else
                disp(sprintf('                      run %d of %d - %s<a href="%s"> histogram of functional connectivity (all slices combined)</a>',html1k2,runs(html1k),subnames{html1k},[html1.QCdir 'hist_FC_compare_13denoise4_allSlices.jpg']))
            end
        else
            if (html1k2==1) && (runs(html1k)==1)
                disp(sprintf('      Subject %d of %d - (pending execution)',html1k,length(sub_dir)))
            elseif (html1k2==1) && (runs(html1k)~=1)
                disp(sprintf('      Subject %d of %d, run %d of %d - (pending execution)',html1k,length(sub_dir),html1k2,runs(html1k)))
            else
                disp(sprintf('                      run %d of %d - (pending execution)',html1k2,runs(html1k)))
            end
        end
        html1.slicesstr = '';
        for html1k3=1:slices
            if exist([home_dir html1.QCdir(3:end) 'hist_FC_compare_13denoise4_slice' sprintf('%.2d',html1k3) '.jpg'],'file')
                html1.slicesstr = strcat(html1.slicesstr,sprintf(' <a href="%s">slice-%d</a>,',[html1.QCdir 'hist_FC_compare_13denoise4_slice' sprintf('%.2d',html1k3) '.jpg'],html1k3));
            else
                html1.slicesstr = strcat(html1.slicesstr,sprintf(' slice-%d,',html1k3));
            end
        end
        if exist([home_dir html1.QCdir(3:end) 'hist_FC_compare_13denoise4_slice' sprintf('%.2d',1) '.jpg'],'file')
            disp(sprintf('                                   (%s)',html1.slicesstr(2:end-1)))
        end
        clear html1.QCdir html1.slicesstr html1k3
    end
end

fprintf('\n')
fprintf('<b>13.5. Histogram of voxel-to-voxel correlations: "13-denoise4-WMregress" vs. "12-denoise3-CSFregress":</b>\n')
for html1k=1:length(sub_dir)
    for html1k2=1:runs(html1k)
        if runs(html1k)==1
            html1.QCdir = ['.' erase(sub_dir{html1k},home_dir(1:end-1)) 'QC/' '13_denoise4-WMregress/'];
        else
            html1.QCdir = ['.' erase(sub_dir{html1k},home_dir(1:end-1)) 'QC/' '13_denoise4-WMregress/' 'run' num2str(html1k2) '/'];
        end
        if exist([home_dir html1.QCdir(3:end) 'hist_FC_compare_12denoise3_13denoise4_allSlices.jpg'],'file')
            if (html1k2==1) && (runs(html1k)==1)
                disp(sprintf('      Subject %d of %d - %s<a href="%s"> histogram of functional connectivity (all slices combined)</a>',html1k,length(sub_dir),subnames{html1k},[html1.QCdir 'hist_FC_compare_12denoise3_13denoise4_allSlices.jpg']))
            elseif (html1k2==1) && (runs(html1k)~=1)
                disp(sprintf('      Subject %d of %d, run %d of %d - %s<a href="%s"> histogram of functional connectivity (all slices combined)</a>',html1k,length(sub_dir),html1k2,runs(html1k),subnames{html1k},[html1.QCdir 'hist_FC_compare_12denoise3_13denoise4_allSlices.jpg']))
            else
                disp(sprintf('                      run %d of %d - %s<a href="%s"> histogram of functional connectivity (all slices combined)</a>',html1k2,runs(html1k),subnames{html1k},[html1.QCdir 'hist_FC_compare_12denoise3_13denoise4_allSlices.jpg']))
            end
        else
            if (html1k2==1) && (runs(html1k)==1)
                disp(sprintf('      Subject %d of %d - (pending execution)',html1k,length(sub_dir)))
            elseif (html1k2==1) && (runs(html1k)~=1)
                disp(sprintf('      Subject %d of %d, run %d of %d - (pending execution)',html1k,length(sub_dir),html1k2,runs(html1k)))
            else
                disp(sprintf('                      run %d of %d - (pending execution)',html1k2,runs(html1k)))
            end
        end
        html1.slicesstr = '';
        for html1k3=1:slices
            if exist([home_dir html1.QCdir(3:end) 'hist_FC_compare_12denoise3_13denoise4_slice' sprintf('%.2d',html1k3) '.jpg'],'file')
                html1.slicesstr = strcat(html1.slicesstr,sprintf(' <a href="%s">slice-%d</a>,',[html1.QCdir 'hist_FC_compare_12denoise3_13denoise4_slice' sprintf('%.2d',html1k3) '.jpg'],html1k3));
            else
                html1.slicesstr = strcat(html1.slicesstr,sprintf(' slice-%d,',html1k3));
            end
        end
        if exist([home_dir html1.QCdir(3:end) 'hist_FC_compare_12denoise3_13denoise4_slice' sprintf('%.2d',1) '.jpg'],'file')
            disp(sprintf('                                   (%s)',html1.slicesstr(2:end-1)))
        end
        clear html1.QCdir html1.slicesstr html1k3
    end
end

if ~isempty(which('VideoWriter')) && (save_videos_steps(7)==1) % check if Matlab's audio video toolbox exists
fprintf('\n')
fprintf('<b>13.6. FMRI video of data after "13-denoise4-WMregress":</b>\n')
for html1k=1:length(sub_dir)
    for html1k2=1:runs(html1k)
        if runs(html1k)==1
            html1.QCdir = ['.' erase(sub_dir{html1k},home_dir(1:end-1)) 'QC/' '13_denoise4-WMregress/'];
        else
            html1.QCdir = ['.' erase(sub_dir{html1k},home_dir(1:end-1)) 'QC/' '13_denoise4-WMregress/' 'run' num2str(html1k2) '/'];
        end
        html1.slicesstr = '';
        for html1k3=1:slices
            if exist([home_dir html1.QCdir(3:end) 'fMRIvideo_13denoise4_slice' sprintf('%.2d',html1k3) '.mp4'],'file')
                html1.slicesstr = strcat(html1.slicesstr,sprintf(' <a href="%s">slice-%d</a>,',[html1.QCdir 'fMRIvideo_13denoise4_slice' sprintf('%.2d',html1k3) '.mp4'],html1k3));
            else
                html1.slicesstr = strcat(html1.slicesstr,sprintf(' slice-%d,',html1k3));
            end
        end
        if exist([home_dir html1.QCdir(3:end) 'fMRIvideo_13denoise4_slice' sprintf('%.2d',1) '.mp4'],'file')
            if (html1k2==1) && (runs(html1k)==1)
                disp(sprintf('      Subject %d of %d - %s %s',html1k,length(sub_dir),subnames{html1k},html1.slicesstr(2:end-1)))
            elseif (html1k2==1) && (runs(html1k)~=1)
                disp(sprintf('      Subject %d of %d, run %d of %d - %s %s',html1k,length(sub_dir),html1k2,runs(html1k),subnames{html1k},html1.slicesstr(2:end-1)))
            else
                disp(sprintf('                      run %d of %d - %s %s',html1k2,runs(html1k),subnames{html1k},html1.slicesstr(2:end-1)))
            end
        else
            if (html1k2==1) && (runs(html1k)==1)
                disp(sprintf('      Subject %d of %d - (pending execution)',html1k,length(sub_dir)))
            elseif (html1k2==1) && (runs(html1k)~=1)
                disp(sprintf('      Subject %d of %d, run %d of %d - (pending execution)',html1k,length(sub_dir),html1k2,runs(html1k)))
            else
                disp(sprintf('                      run %d of %d - (pending execution)',html1k2,runs(html1k)))
            end
        end
    end
    clear html1.QCdir html1.slicesstr html1k3
end
end

else
fprintf('<b>Not in pipeline</b>\n')
fprintf('</p>')
end
end

%% 14. Denoise-5 (additional covariates regression)
if exist('choices_preproc','var')
if choices_preproc(14)==1
fprintf('<p style="font-size:14px ; color:#000000">')

fprintf('<b>14.1. Time series within the gray and white matter after "14-denoise5-COVregress":</b>\n')
for html1k=1:length(sub_dir)
    for html1k2=1:runs(html1k)
        if runs(html1k)==1
            html1.QCdir = ['.' erase(sub_dir{html1k},home_dir(1:end-1)) 'QC/' '14_denoise5-COVregress/'];
        else
            html1.QCdir = ['.' erase(sub_dir{html1k},home_dir(1:end-1)) 'QC/' '14_denoise5-COVregress/' 'run' num2str(html1k2) '/'];
        end
        if exist([home_dir html1.QCdir(3:end) 'fMRI_ts_14denoise5_allSlices.jpg'],'file')
            if (html1k2==1) && (runs(html1k)==1)
                disp(sprintf('      Subject %d of %d - %s<a href="%s"> time series plot (all slices combined)</a>',html1k,length(sub_dir),subnames{html1k},[html1.QCdir 'fMRI_ts_14denoise5_allSlices.jpg']))
            elseif (html1k2==1) && (runs(html1k)~=1)
                disp(sprintf('      Subject %d of %d, run %d of %d - %s<a href="%s"> time series plot (all slices combined)</a>',html1k,length(sub_dir),html1k2,runs(html1k),subnames{html1k},[html1.QCdir 'fMRI_ts_14denoise5_allSlices.jpg']))
            else
                disp(sprintf('                      run %d of %d - %s<a href="%s"> time series plot (all slices combined)</a>',html1k2,runs(html1k),subnames{html1k},[html1.QCdir 'fMRI_ts_14denoise5_allSlices.jpg']))
            end
        else
            if (html1k2==1) && (runs(html1k)==1)
                disp(sprintf('      Subject %d of %d - (pending execution)',html1k,length(sub_dir)))
            elseif (html1k2==1) && (runs(html1k)~=1)
                disp(sprintf('      Subject %d of %d, run %d of %d - (pending execution)',html1k,length(sub_dir),html1k2,runs(html1k)))
            else
                disp(sprintf('                      run %d of %d - (pending execution)',html1k2,runs(html1k)))
            end
        end
        html1.slicesstr = '';
        for html1k3=1:slices
            if exist([home_dir html1.QCdir(3:end) 'fMRI_ts_14denoise5_slice' sprintf('%.2d',html1k3) '.jpg'],'file')
                html1.slicesstr = strcat(html1.slicesstr,sprintf(' <a href="%s">slice-%d</a>,',[html1.QCdir 'fMRI_ts_14denoise5_slice' sprintf('%.2d',html1k3) '.jpg'],html1k3));
            else
                html1.slicesstr = strcat(html1.slicesstr,sprintf(' slice-%d,',html1k3));
            end
        end
        if exist([home_dir html1.QCdir(3:end) 'fMRI_ts_14denoise5_slice' sprintf('%.2d',1) '.jpg'],'file')
            disp(sprintf('                                   (%s)',html1.slicesstr(2:end-1)))
        end
        clear html1.QCdir html1.slicesstr html1k3
    end
end

fprintf('\n')
fprintf('<b>14.2. Global mean signal within the gray and white matter after "14-denoise5-COVregress":</b>\n')
for html1k=1:length(sub_dir)
    for html1k2=1:runs(html1k)
        if runs(html1k)==1
            html1.QCdir = ['.' erase(sub_dir{html1k},home_dir(1:end-1)) 'QC/' '14_denoise5-COVregress/'];
        else
            html1.QCdir = ['.' erase(sub_dir{html1k},home_dir(1:end-1)) 'QC/' '14_denoise5-COVregress/' 'run' num2str(html1k2) '/'];
        end
        if exist([home_dir html1.QCdir(3:end) 'global_signal_14denoise5_allSlices.jpg'],'file')
            if (html1k2==1) && (runs(html1k)==1)
                disp(sprintf('      Subject %d of %d - %s<a href="%s"> global mean signal (all slices combined)</a>',html1k,length(sub_dir),subnames{html1k},[html1.QCdir 'global_signal_14denoise5_allSlices.jpg']))
            elseif (html1k2==1) && (runs(html1k)~=1)
                disp(sprintf('      Subject %d of %d, run %d of %d - %s<a href="%s"> global mean signal (all slices combined)</a>',html1k,length(sub_dir),html1k2,runs(html1k),subnames{html1k},[html1.QCdir 'global_signal_14denoise5_allSlices.jpg']))
            else
                disp(sprintf('                      run %d of %d - %s<a href="%s"> global mean signal (all slices combined)</a>',html1k2,runs(html1k),subnames{html1k},[html1.QCdir 'global_signal_14denoise5_allSlices.jpg']))
            end
        else
            if (html1k2==1) && (runs(html1k)==1)
                disp(sprintf('      Subject %d of %d - (pending execution)',html1k,length(sub_dir)))
            elseif (html1k2==1) && (runs(html1k)~=1)
                disp(sprintf('      Subject %d of %d, run %d of %d - (pending execution)',html1k,length(sub_dir),html1k2,runs(html1k)))
            else
                disp(sprintf('                      run %d of %d - (pending execution)',html1k2,runs(html1k)))
            end
        end
        html1.slicesstr = '';
        for html1k3=1:slices
            if exist([home_dir html1.QCdir(3:end) 'global_signal_14denoise5_slice' sprintf('%.2d',html1k3) '.jpg'],'file')
                html1.slicesstr = strcat(html1.slicesstr,sprintf(' <a href="%s">slice-%d</a>,',[html1.QCdir 'global_signal_14denoise5_slice' sprintf('%.2d',html1k3) '.jpg'],html1k3));
            else
                html1.slicesstr = strcat(html1.slicesstr,sprintf(' slice-%d,',html1k3));
            end
        end
        if exist([home_dir html1.QCdir(3:end) 'global_signal_14denoise5_slice' sprintf('%.2d',1) '.jpg'],'file')
            disp(sprintf('                                   (%s)',html1.slicesstr(2:end-1)))
        end
        clear html1.QCdir html1.slicesstr html1k3
    end
end

fprintf('\n')
fprintf('<b>14.3. Global mean signal: "14-denoise5-COVregress" vs. "13-denoise4-WMregress":</b>\n')
for html1k=1:length(sub_dir)
    for html1k2=1:runs(html1k)
        if runs(html1k)==1
            html1.QCdir = ['.' erase(sub_dir{html1k},home_dir(1:end-1)) 'QC/' '14_denoise5-COVregress/'];
        else
            html1.QCdir = ['.' erase(sub_dir{html1k},home_dir(1:end-1)) 'QC/' '14_denoise5-COVregress/' 'run' num2str(html1k2) '/'];
        end
        if exist([home_dir html1.QCdir(3:end) 'global_signal_13denoise4_vs_14denoise5_allSlices.jpg'],'file')
            if (html1k2==1) && (runs(html1k)==1)
                disp(sprintf('      Subject %d of %d - %s<a href="%s"> global mean signal (all slices combined)</a>',html1k,length(sub_dir),subnames{html1k},[html1.QCdir 'global_signal_13denoise4_vs_14denoise5_allSlices.jpg']))
            elseif (html1k2==1) && (runs(html1k)~=1)
                disp(sprintf('      Subject %d of %d, run %d of %d - %s<a href="%s"> global mean signal (all slices combined)</a>',html1k,length(sub_dir),html1k2,runs(html1k),subnames{html1k},[html1.QCdir 'global_signal_13denoise4_vs_14denoise5_allSlices.jpg']))
            else
                disp(sprintf('                      run %d of %d - %s<a href="%s"> global mean signal (all slices combined)</a>',html1k2,runs(html1k),subnames{html1k},[html1.QCdir 'global_signal_13denoise4_vs_14denoise5_allSlices.jpg']))
            end
        else
            if (html1k2==1) && (runs(html1k)==1)
                disp(sprintf('      Subject %d of %d - (pending execution)',html1k,length(sub_dir)))
            elseif (html1k2==1) && (runs(html1k)~=1)
                disp(sprintf('      Subject %d of %d, run %d of %d - (pending execution)',html1k,length(sub_dir),html1k2,runs(html1k)))
            else
                disp(sprintf('                      run %d of %d - (pending execution)',html1k2,runs(html1k)))
            end
        end
        html1.slicesstr = '';
        for html1k3=1:slices
            if exist([home_dir html1.QCdir(3:end) 'global_signal_13denoise4_vs_14denoise5_slice' sprintf('%.2d',html1k3) '.jpg'],'file')
                html1.slicesstr = strcat(html1.slicesstr,sprintf(' <a href="%s">slice-%d</a>,',[html1.QCdir 'global_signal_13denoise4_vs_14denoise5_slice' sprintf('%.2d',html1k3) '.jpg'],html1k3));
            else
                html1.slicesstr = strcat(html1.slicesstr,sprintf(' slice-%d,',html1k3));
            end
        end
        if exist([home_dir html1.QCdir(3:end) 'global_signal_13denoise4_vs_14denoise5_slice' sprintf('%.2d',1) '.jpg'],'file')
            disp(sprintf('                                   (%s)',html1.slicesstr(2:end-1)))
        end
        clear html1.QCdir html1.slicesstr html1k3
    end
end

fprintf('\n')
fprintf('<b>14.4. Histogram of voxel-to-voxel correlations within the gray and white matter after "14-denoise5-COVregress":</b>\n')
for html1k=1:length(sub_dir)
    for html1k2=1:runs(html1k)
        if runs(html1k)==1
            html1.QCdir = ['.' erase(sub_dir{html1k},home_dir(1:end-1)) 'QC/' '14_denoise5-COVregress/'];
        else
            html1.QCdir = ['.' erase(sub_dir{html1k},home_dir(1:end-1)) 'QC/' '14_denoise5-COVregress/' 'run' num2str(html1k2) '/'];
        end
        if exist([home_dir html1.QCdir(3:end) 'hist_FC_compare_14denoise5_allSlices.jpg'],'file')
            if (html1k2==1) && (runs(html1k)==1)
                disp(sprintf('      Subject %d of %d - %s<a href="%s"> histogram of functional connectivity (all slices combined)</a>',html1k,length(sub_dir),subnames{html1k},[html1.QCdir 'hist_FC_compare_14denoise5_allSlices.jpg']))
            elseif (html1k2==1) && (runs(html1k)~=1)
                disp(sprintf('      Subject %d of %d, run %d of %d - %s<a href="%s"> histogram of functional connectivity (all slices combined)</a>',html1k,length(sub_dir),html1k2,runs(html1k),subnames{html1k},[html1.QCdir 'hist_FC_compare_14denoise5_allSlices.jpg']))
            else
                disp(sprintf('                      run %d of %d - %s<a href="%s"> histogram of functional connectivity (all slices combined)</a>',html1k2,runs(html1k),subnames{html1k},[html1.QCdir 'hist_FC_compare_14denoise5_allSlices.jpg']))
            end
        else
            if (html1k2==1) && (runs(html1k)==1)
                disp(sprintf('      Subject %d of %d - (pending execution)',html1k,length(sub_dir)))
            elseif (html1k2==1) && (runs(html1k)~=1)
                disp(sprintf('      Subject %d of %d, run %d of %d - (pending execution)',html1k,length(sub_dir),html1k2,runs(html1k)))
            else
                disp(sprintf('                      run %d of %d - (pending execution)',html1k2,runs(html1k)))
            end
        end
        html1.slicesstr = '';
        for html1k3=1:slices
            if exist([home_dir html1.QCdir(3:end) 'hist_FC_compare_14denoise5_slice' sprintf('%.2d',html1k3) '.jpg'],'file')
                html1.slicesstr = strcat(html1.slicesstr,sprintf(' <a href="%s">slice-%d</a>,',[html1.QCdir 'hist_FC_compare_14denoise5_slice' sprintf('%.2d',html1k3) '.jpg'],html1k3));
            else
                html1.slicesstr = strcat(html1.slicesstr,sprintf(' slice-%d,',html1k3));
            end
        end
        if exist([home_dir html1.QCdir(3:end) 'hist_FC_compare_14denoise5_slice' sprintf('%.2d',1) '.jpg'],'file')
            disp(sprintf('                                   (%s)',html1.slicesstr(2:end-1)))
        end
        clear html1.QCdir html1.slicesstr html1k3
    end
end

fprintf('\n')
fprintf('<b>14.5. Histogram of voxel-to-voxel correlations: "14-denoise5-COVregress" vs. "13-denoise4-WMregress":</b>\n')
for html1k=1:length(sub_dir)
    for html1k2=1:runs(html1k)
        if runs(html1k)==1
            html1.QCdir = ['.' erase(sub_dir{html1k},home_dir(1:end-1)) 'QC/' '14_denoise5-COVregress/'];
        else
            html1.QCdir = ['.' erase(sub_dir{html1k},home_dir(1:end-1)) 'QC/' '14_denoise5-COVregress/' 'run' num2str(html1k2) '/'];
        end
        if exist([home_dir html1.QCdir(3:end) 'hist_FC_compare_13denoise4_14denoise5_allSlices.jpg'],'file')
            if (html1k2==1) && (runs(html1k)==1)
                disp(sprintf('      Subject %d of %d - %s<a href="%s"> histogram of functional connectivity (all slices combined)</a>',html1k,length(sub_dir),subnames{html1k},[html1.QCdir 'hist_FC_compare_13denoise4_14denoise5_allSlices.jpg']))
            elseif (html1k2==1) && (runs(html1k)~=1)
                disp(sprintf('      Subject %d of %d, run %d of %d - %s<a href="%s"> histogram of functional connectivity (all slices combined)</a>',html1k,length(sub_dir),html1k2,runs(html1k),subnames{html1k},[html1.QCdir 'hist_FC_compare_13denoise4_14denoise5_allSlices.jpg']))
            else
                disp(sprintf('                      run %d of %d - %s<a href="%s"> histogram of functional connectivity (all slices combined)</a>',html1k2,runs(html1k),subnames{html1k},[html1.QCdir 'hist_FC_compare_13denoise4_14denoise5_allSlices.jpg']))
            end
        else
            if (html1k2==1) && (runs(html1k)==1)
                disp(sprintf('      Subject %d of %d - (pending execution)',html1k,length(sub_dir)))
            elseif (html1k2==1) && (runs(html1k)~=1)
                disp(sprintf('      Subject %d of %d, run %d of %d - (pending execution)',html1k,length(sub_dir),html1k2,runs(html1k)))
            else
                disp(sprintf('                      run %d of %d - (pending execution)',html1k2,runs(html1k)))
            end
        end
        html1.slicesstr = '';
        for html1k3=1:slices
            if exist([home_dir html1.QCdir(3:end) 'hist_FC_compare_13denoise4_14denoise5_slice' sprintf('%.2d',html1k3) '.jpg'],'file')
                html1.slicesstr = strcat(html1.slicesstr,sprintf(' <a href="%s">slice-%d</a>,',[html1.QCdir 'hist_FC_compare_13denoise4_14denoise5_slice' sprintf('%.2d',html1k3) '.jpg'],html1k3));
            else
                html1.slicesstr = strcat(html1.slicesstr,sprintf(' slice-%d,',html1k3));
            end
        end
        if exist([home_dir html1.QCdir(3:end) 'hist_FC_compare_13denoise4_14denoise5_slice' sprintf('%.2d',1) '.jpg'],'file')
            disp(sprintf('                                   (%s)',html1.slicesstr(2:end-1)))
        end
        clear html1.QCdir html1.slicesstr html1k3
    end
end

if ~isempty(which('VideoWriter')) && (save_videos_steps(8)==1) % check if Matlab's audio video toolbox exists
fprintf('\n')
fprintf('<b>14.6. FMRI video of data after "14-denoise5-COVregress":</b>\n')
for html1k=1:length(sub_dir)
    for html1k2=1:runs(html1k)
        if runs(html1k)==1
            html1.QCdir = ['.' erase(sub_dir{html1k},home_dir(1:end-1)) 'QC/' '14_denoise5-COVregress/'];
        else
            html1.QCdir = ['.' erase(sub_dir{html1k},home_dir(1:end-1)) 'QC/' '14_denoise5-COVregress/' 'run' num2str(html1k2) '/'];
        end
        html1.slicesstr = '';
        for html1k3=1:slices
            if exist([home_dir html1.QCdir(3:end) 'fMRIvideo_14denoise5_slice' sprintf('%.2d',html1k3) '.mp4'],'file')
                html1.slicesstr = strcat(html1.slicesstr,sprintf(' <a href="%s">slice-%d</a>,',[html1.QCdir 'fMRIvideo_14denoise5_slice' sprintf('%.2d',html1k3) '.mp4'],html1k3));
            else
                html1.slicesstr = strcat(html1.slicesstr,sprintf(' slice-%d,',html1k3));
            end
        end
        if exist([home_dir html1.QCdir(3:end) 'fMRIvideo_14denoise5_slice' sprintf('%.2d',1) '.mp4'],'file')
            if (html1k2==1) && (runs(html1k)==1)
                disp(sprintf('      Subject %d of %d - %s %s',html1k,length(sub_dir),subnames{html1k},html1.slicesstr(2:end-1)))
            elseif (html1k2==1) && (runs(html1k)~=1)
                disp(sprintf('      Subject %d of %d, run %d of %d - %s %s',html1k,length(sub_dir),html1k2,runs(html1k),subnames{html1k},html1.slicesstr(2:end-1)))
            else
                disp(sprintf('                      run %d of %d - %s %s',html1k2,runs(html1k),subnames{html1k},html1.slicesstr(2:end-1)))
            end
        else
            if (html1k2==1) && (runs(html1k)==1)
                disp(sprintf('      Subject %d of %d - (pending execution)',html1k,length(sub_dir)))
            elseif (html1k2==1) && (runs(html1k)~=1)
                disp(sprintf('      Subject %d of %d, run %d of %d - (pending execution)',html1k,length(sub_dir),html1k2,runs(html1k)))
            else
                disp(sprintf('                      run %d of %d - (pending execution)',html1k2,runs(html1k)))
            end
        end
    end
    clear html1.QCdir html1.slicesstr html1k3
end
end

else
fprintf('<b>Not in pipeline</b>\n')
fprintf('</p>')
end
end

%% 15. Temporal band-pass filtering
if exist('choices_preproc','var')
if choices_preproc(15)==1
fprintf('<p style="font-size:14px ; color:#000000">')

fprintf('<b>15.1. Time series within the gray and white matter after "15-temporal-filtering":</b>\n')
for html1k=1:length(sub_dir)
    for html1k2=1:runs(html1k)
        if runs(html1k)==1
            html1.QCdir = ['.' erase(sub_dir{html1k},home_dir(1:end-1)) 'QC/' '15_temporal-filtering/'];
        else
            html1.QCdir = ['.' erase(sub_dir{html1k},home_dir(1:end-1)) 'QC/' '15_temporal-filtering/' 'run' num2str(html1k2) '/'];
        end
        if exist([home_dir html1.QCdir(3:end) 'fMRI_ts_15filter_allSlices.jpg'],'file')
            if (html1k2==1) && (runs(html1k)==1)
                disp(sprintf('      Subject %d of %d - %s<a href="%s"> time series plot (all slices combined)</a>',html1k,length(sub_dir),subnames{html1k},[html1.QCdir 'fMRI_ts_15filter_allSlices.jpg']))
            elseif (html1k2==1) && (runs(html1k)~=1)
                disp(sprintf('      Subject %d of %d, run %d of %d - %s<a href="%s"> time series plot (all slices combined)</a>',html1k,length(sub_dir),html1k2,runs(html1k),subnames{html1k},[html1.QCdir 'fMRI_ts_15filter_allSlices.jpg']))
            else
                disp(sprintf('                      run %d of %d - %s<a href="%s"> time series plot (all slices combined)</a>',html1k2,runs(html1k),subnames{html1k},[html1.QCdir 'fMRI_ts_15filter_allSlices.jpg']))
            end
        else
            if (html1k2==1) && (runs(html1k)==1)
                disp(sprintf('      Subject %d of %d - (pending execution)',html1k,length(sub_dir)))
            elseif (html1k2==1) && (runs(html1k)~=1)
                disp(sprintf('      Subject %d of %d, run %d of %d - (pending execution)',html1k,length(sub_dir),html1k2,runs(html1k)))
            else
                disp(sprintf('                      run %d of %d - (pending execution)',html1k2,runs(html1k)))
            end
        end
        html1.slicesstr = '';
        for html1k3=1:slices
            if exist([home_dir html1.QCdir(3:end) 'fMRI_ts_15filter_slice' sprintf('%.2d',html1k3) '.jpg'],'file')
                html1.slicesstr = strcat(html1.slicesstr,sprintf(' <a href="%s">slice-%d</a>,',[html1.QCdir 'fMRI_ts_15filter_slice' sprintf('%.2d',html1k3) '.jpg'],html1k3));
            else
                html1.slicesstr = strcat(html1.slicesstr,sprintf(' slice-%d,',html1k3));
            end
        end
        if exist([home_dir html1.QCdir(3:end) 'fMRI_ts_15filter_slice' sprintf('%.2d',1) '.jpg'],'file')
            disp(sprintf('                                   (%s)',html1.slicesstr(2:end-1)))
        end
        clear html1.QCdir html1.slicesstr html1k3
    end
end

fprintf('\n')
fprintf('<b>15.2. Normalized mean gray and white matter signals: "15-temporal-filtering" vs. "13-denoise4-WMregress":</b>\n')
for html1k=1:length(sub_dir)
    for html1k2=1:runs(html1k)
        if runs(html1k)==1
            html1.QCdir = ['.' erase(sub_dir{html1k},home_dir(1:end-1)) 'QC/' '15_temporal-filtering/'];
        else
            html1.QCdir = ['.' erase(sub_dir{html1k},home_dir(1:end-1)) 'QC/' '15_temporal-filtering/' 'run' num2str(html1k2) '/'];
        end
        if exist([home_dir html1.QCdir(3:end) 'mean_timeseries_14denoise5_vs_15filter_allSlices.jpg'],'file')
            if (html1k2==1) && (runs(html1k)==1)
                disp(sprintf('      Subject %d of %d - %s<a href="%s"> global mean signal (all slices combined)</a>',html1k,length(sub_dir),subnames{html1k},[html1.QCdir 'mean_timeseries_14denoise5_vs_15filter_allSlices.jpg']))
            elseif (html1k2==1) && (runs(html1k)~=1)
                disp(sprintf('      Subject %d of %d, run %d of %d - %s<a href="%s"> global mean signal (all slices combined)</a>',html1k,length(sub_dir),html1k2,runs(html1k),subnames{html1k},[html1.QCdir 'mean_timeseries_14denoise5_vs_15filter_allSlices.jpg']))
            else
                disp(sprintf('                      run %d of %d - %s<a href="%s"> global mean signal (all slices combined)</a>',html1k2,runs(html1k),subnames{html1k},[html1.QCdir 'mean_timeseries_14denoise5_vs_15filter_allSlices.jpg']))
            end
        else
            if (html1k2==1) && (runs(html1k)==1)
                disp(sprintf('      Subject %d of %d - (pending execution)',html1k,length(sub_dir)))
            elseif (html1k2==1) && (runs(html1k)~=1)
                disp(sprintf('      Subject %d of %d, run %d of %d - (pending execution)',html1k,length(sub_dir),html1k2,runs(html1k)))
            else
                disp(sprintf('                      run %d of %d - (pending execution)',html1k2,runs(html1k)))
            end
        end
        html1.slicesstr = '';
        for html1k3=1:slices
            if exist([home_dir html1.QCdir(3:end) 'mean_timeseries_14denoise5_vs_15filter_slice' sprintf('%.2d',html1k3) '.jpg'],'file')
                html1.slicesstr = strcat(html1.slicesstr,sprintf(' <a href="%s">slice-%d</a>,',[html1.QCdir 'mean_timeseries_14denoise5_vs_15filter_slice' sprintf('%.2d',html1k3) '.jpg'],html1k3));
            else
                html1.slicesstr = strcat(html1.slicesstr,sprintf(' slice-%d,',html1k3));
            end
        end
        if exist([home_dir html1.QCdir(3:end) 'mean_timeseries_14denoise5_vs_15filter_slice' sprintf('%.2d',1) '.jpg'],'file')
            disp(sprintf('                                   (%s)',html1.slicesstr(2:end-1)))
        end
        clear html1.QCdir html1.slicesstr html1k3
    end
end

fprintf('\n')
fprintf('<b>15.3. Histogram of voxel-to-voxel correlations within the gray and white matter after "15-temporal-filtering":</b>\n')
for html1k=1:length(sub_dir)
    for html1k2=1:runs(html1k)
        if runs(html1k)==1
            html1.QCdir = ['.' erase(sub_dir{html1k},home_dir(1:end-1)) 'QC/' '15_temporal-filtering/'];
        else
            html1.QCdir = ['.' erase(sub_dir{html1k},home_dir(1:end-1)) 'QC/' '15_temporal-filtering/' 'run' num2str(html1k2) '/'];
        end
        if exist([home_dir html1.QCdir(3:end) 'hist_FC_compare_15filter_allSlices.jpg'],'file')
            if (html1k2==1) && (runs(html1k)==1)
                disp(sprintf('      Subject %d of %d - %s<a href="%s"> histogram of functional connectivity (all slices combined)</a>',html1k,length(sub_dir),subnames{html1k},[html1.QCdir 'hist_FC_compare_15filter_allSlices.jpg']))
            elseif (html1k2==1) && (runs(html1k)~=1)
                disp(sprintf('      Subject %d of %d, run %d of %d - %s<a href="%s"> histogram of functional connectivity (all slices combined)</a>',html1k,length(sub_dir),html1k2,runs(html1k),subnames{html1k},[html1.QCdir 'hist_FC_compare_15filter_allSlices.jpg']))
            else
                disp(sprintf('                      run %d of %d - %s<a href="%s"> histogram of functional connectivity (all slices combined)</a>',html1k2,runs(html1k),subnames{html1k},[html1.QCdir 'hist_FC_compare_15filter_allSlices.jpg']))
            end
        else
            if (html1k2==1) && (runs(html1k)==1)
                disp(sprintf('      Subject %d of %d - (pending execution)',html1k,length(sub_dir)))
            elseif (html1k2==1) && (runs(html1k)~=1)
                disp(sprintf('      Subject %d of %d, run %d of %d - (pending execution)',html1k,length(sub_dir),html1k2,runs(html1k)))
            else
                disp(sprintf('                      run %d of %d - (pending execution)',html1k2,runs(html1k)))
            end
        end
        html1.slicesstr = '';
        for html1k3=1:slices
            if exist([home_dir html1.QCdir(3:end) 'hist_FC_compare_15filter_slice' sprintf('%.2d',html1k3) '.jpg'],'file')
                html1.slicesstr = strcat(html1.slicesstr,sprintf(' <a href="%s">slice-%d</a>,',[html1.QCdir 'hist_FC_compare_15filter_slice' sprintf('%.2d',html1k3) '.jpg'],html1k3));
            else
                html1.slicesstr = strcat(html1.slicesstr,sprintf(' slice-%d,',html1k3));
            end
        end
        if exist([home_dir html1.QCdir(3:end) 'hist_FC_compare_15filter_slice' sprintf('%.2d',1) '.jpg'],'file')
            disp(sprintf('                                   (%s)',html1.slicesstr(2:end-1)))
        end
        clear html1.QCdir html1.slicesstr html1k3
    end
end

fprintf('\n')
fprintf('<b>15.4. Histogram of voxel-to-voxel correlations: "15-temporal-filtering" vs. "13-denoise4-WMregress":</b>\n')
for html1k=1:length(sub_dir)
    for html1k2=1:runs(html1k)
        if runs(html1k)==1
            html1.QCdir = ['.' erase(sub_dir{html1k},home_dir(1:end-1)) 'QC/' '15_temporal-filtering/'];
        else
            html1.QCdir = ['.' erase(sub_dir{html1k},home_dir(1:end-1)) 'QC/' '15_temporal-filtering/' 'run' num2str(html1k2) '/'];
        end
        if exist([home_dir html1.QCdir(3:end) 'hist_FC_compare_14denoise5_15filter_allSlices.jpg'],'file')
            if (html1k2==1) && (runs(html1k)==1)
                disp(sprintf('      Subject %d of %d - %s<a href="%s"> histogram of functional connectivity (all slices combined)</a>',html1k,length(sub_dir),subnames{html1k},[html1.QCdir 'hist_FC_compare_14denoise5_15filter_allSlices.jpg']))
            elseif (html1k2==1) && (runs(html1k)~=1)
                disp(sprintf('      Subject %d of %d, run %d of %d - %s<a href="%s"> histogram of functional connectivity (all slices combined)</a>',html1k,length(sub_dir),html1k2,runs(html1k),subnames{html1k},[html1.QCdir 'hist_FC_compare_14denoise5_15filter_allSlices.jpg']))
            else
                disp(sprintf('                      run %d of %d - %s<a href="%s"> histogram of functional connectivity (all slices combined)</a>',html1k2,runs(html1k),subnames{html1k},[html1.QCdir 'hist_FC_compare_14denoise5_15filter_allSlices.jpg']))
            end
        else
            if (html1k2==1) && (runs(html1k)==1)
                disp(sprintf('      Subject %d of %d - (pending execution)',html1k,length(sub_dir)))
            elseif (html1k2==1) && (runs(html1k)~=1)
                disp(sprintf('      Subject %d of %d, run %d of %d - (pending execution)',html1k,length(sub_dir),html1k2,runs(html1k)))
            else
                disp(sprintf('                      run %d of %d - (pending execution)',html1k2,runs(html1k)))
            end
        end
        html1.slicesstr = '';
        for html1k3=1:slices
            if exist([home_dir html1.QCdir(3:end) 'hist_FC_compare_14denoise5_15filter_slice' sprintf('%.2d',html1k3) '.jpg'],'file')
                html1.slicesstr = strcat(html1.slicesstr,sprintf(' <a href="%s">slice-%d</a>,',[html1.QCdir 'hist_FC_compare_14denoise5_15filter_slice' sprintf('%.2d',html1k3) '.jpg'],html1k3));
            else
                html1.slicesstr = strcat(html1.slicesstr,sprintf(' slice-%d,',html1k3));
            end
        end
        if exist([home_dir html1.QCdir(3:end) 'hist_FC_compare_14denoise5_15filter_slice' sprintf('%.2d',1) '.jpg'],'file')
            disp(sprintf('                                   (%s)',html1.slicesstr(2:end-1)))
        end
        clear html1.QCdir html1.slicesstr html1k3
    end
end

else
fprintf('<b>Not in pipeline</b>\n')
fprintf('</p>')
end
end

%% 16. Define cord quadrants
if exist('choices_preproc','var')
if choices_preproc(16)==1
fprintf('<p style="font-size:14px ; color:#000000">')

fprintf('<b>Gray and white matter quadrants in the cord (manually defined):</b>\n')
for html1k=1:length(sub_dir)
    html1.QCdir = ['.' erase(sub_dir{html1k},home_dir(1:end-1)) 'QC/' '16_cord_quadrants/'];
    html1.slicesstr = '';
    for html1k3=1:slices
        if exist([home_dir html1.QCdir(3:end) 'cord_quadrants_slice' sprintf('%.2d',html1k3) '.jpg'],'file')
            html1.slicesstr = strcat(html1.slicesstr,sprintf(' <a href="%s">slice-%d</a>,',[html1.QCdir 'cord_quadrants_slice' sprintf('%.2d',html1k3) '.jpg'],html1k3));
        else
            html1.slicesstr = strcat(html1.slicesstr,sprintf(' slice-%d,',html1k3));
        end
    end
    if exist([home_dir html1.QCdir(3:end) 'cord_quadrants_slice' sprintf('%.2d',1) '.jpg'],'file')
        disp(sprintf('      Subject %d of %d - %s %s',html1k,length(sub_dir),subnames{html1k},html1.slicesstr(2:end-1)))
    else
        disp(sprintf('      Subject %d of %d - (pending execution)',html1k,length(sub_dir)))
    end
    clear html1.QCdir html1.slicesstr html1k3
end

else
fprintf('<b>Not in pipeline</b>\n')
fprintf('</p>')
end
end

%% 17. HRF deconvolution
if exist('choices_preproc','var')
if choices_preproc(17)==1
fprintf('<p style="font-size:14px ; color:#000000">')

fprintf('<b>17.1. The hemodynamic response function (HRF) and HRF parameters across slices and quadrants:</b>\n')
for html1k=1:length(sub_dir)
    for html1k2=1:runs(html1k)
        if runs(html1k)==1
            html1.QCdir = ['.' erase(sub_dir{html1k},home_dir(1:end-1)) 'QC/' '17_deconvolution/'];
        else
            html1.QCdir = ['.' erase(sub_dir{html1k},home_dir(1:end-1)) 'QC/' '17_deconvolution/' 'run' num2str(html1k2) '/'];
        end
        if exist([home_dir html1.QCdir(3:end) 'HRF_plots_17deconv_allSlices.jpg'],'file')
            if (html1k2==1) && (runs(html1k)==1)
                disp(sprintf('      Subject %d of %d - %s<a href="%s"> Plots of the HRF across slices and quadrants</a>',html1k,length(sub_dir),subnames{html1k},[html1.QCdir 'HRF_plots_17deconv_allSlices.jpg']))
            elseif (html1k2==1) && (runs(html1k)~=1)
                disp(sprintf('      Subject %d of %d, run %d of %d - %s<a href="%s"> Plots of the HRF across slices and quadrants</a>',html1k,length(sub_dir),html1k2,runs(html1k),subnames{html1k},[html1.QCdir 'HRF_plots_17deconv_allSlices.jpg']))
            else
                disp(sprintf('                      run %d of %d - %s<a href="%s"> Plots of the HRF across slices and quadrants</a>',html1k2,runs(html1k),subnames{html1k},[html1.QCdir 'HRF_plots_17deconv_allSlices.jpg']))
            end
        else
            if (html1k2==1) && (runs(html1k)==1)
                disp(sprintf('      Subject %d of %d - (pending execution)',html1k,length(sub_dir)))
            elseif (html1k2==1) && (runs(html1k)~=1)
                disp(sprintf('      Subject %d of %d, run %d of %d - (pending execution)',html1k,length(sub_dir),html1k2,runs(html1k)))
            else
                disp(sprintf('                      run %d of %d - (pending execution)',html1k2,runs(html1k)))
            end
        end
        if exist([home_dir html1.QCdir(3:end) 'HRF_parameters_17deconv_allSlices.jpg'],'file')
            if (html1k2==1) && (runs(html1k)==1)
                disp(sprintf('                     - %s<a href="%s"> HRF shape parameters across slices and quadrants</a>',subnames{html1k},[html1.QCdir 'HRF_parameters_17deconv_allSlices.jpg']))
            elseif (html1k2==1) && (runs(html1k)~=1)
                disp(sprintf('                                 - %s<a href="%s"> HRF shape parameters across slices and quadrants</a>',subnames{html1k},[html1.QCdir 'HRF_parameters_17deconv_allSlices.jpg']))
            else
                disp(sprintf('                                 - %s<a href="%s"> HRF shape parameters across slices and quadrants</a>',subnames{html1k},[html1.QCdir 'HRF_parameters_17deconv_allSlices.jpg']))
            end
        end
        clear html1.QCdir html1k3
    end
end

fprintf('\n')
fprintf('<b>17.2. Time series within the gray and white matter after "17-HRF-deconvolution":</b>\n')
for html1k=1:length(sub_dir)
    for html1k2=1:runs(html1k)
        if runs(html1k)==1
            html1.QCdir = ['.' erase(sub_dir{html1k},home_dir(1:end-1)) 'QC/' '17_deconvolution/'];
        else
            html1.QCdir = ['.' erase(sub_dir{html1k},home_dir(1:end-1)) 'QC/' '17_deconvolution/' 'run' num2str(html1k2) '/'];
        end
        if exist([home_dir html1.QCdir(3:end) 'fMRI_ts_17deconv_allSlices.jpg'],'file')
            if (html1k2==1) && (runs(html1k)==1)
                disp(sprintf('      Subject %d of %d - %s<a href="%s"> time series plot (all slices combined)</a>',html1k,length(sub_dir),subnames{html1k},[html1.QCdir 'fMRI_ts_17deconv_allSlices.jpg']))
            elseif (html1k2==1) && (runs(html1k)~=1)
                disp(sprintf('      Subject %d of %d, run %d of %d - %s<a href="%s"> time series plot (all slices combined)</a>',html1k,length(sub_dir),html1k2,runs(html1k),subnames{html1k},[html1.QCdir 'fMRI_ts_17deconv_allSlices.jpg']))
            else
                disp(sprintf('                      run %d of %d - %s<a href="%s"> time series plot (all slices combined)</a>',html1k2,runs(html1k),subnames{html1k},[html1.QCdir 'fMRI_ts_17deconv_allSlices.jpg']))
            end
        else
            if (html1k2==1) && (runs(html1k)==1)
                disp(sprintf('      Subject %d of %d - (pending execution)',html1k,length(sub_dir)))
            elseif (html1k2==1) && (runs(html1k)~=1)
                disp(sprintf('      Subject %d of %d, run %d of %d - (pending execution)',html1k,length(sub_dir),html1k2,runs(html1k)))
            else
                disp(sprintf('                      run %d of %d - (pending execution)',html1k2,runs(html1k)))
            end
        end
        html1.slicesstr = '';
        for html1k3=1:slices
            if exist([home_dir html1.QCdir(3:end) 'fMRI_ts_17deconv_slice' sprintf('%.2d',html1k3) '.jpg'],'file')
                html1.slicesstr = strcat(html1.slicesstr,sprintf(' <a href="%s">slice-%d</a>,',[html1.QCdir 'fMRI_ts_17deconv_slice' sprintf('%.2d',html1k3) '.jpg'],html1k3));
            else
                html1.slicesstr = strcat(html1.slicesstr,sprintf(' slice-%d,',html1k3));
            end
        end
        if exist([home_dir html1.QCdir(3:end) 'fMRI_ts_17deconv_slice' sprintf('%.2d',1) '.jpg'],'file')
            disp(sprintf('                                   (%s)',html1.slicesstr(2:end-1)))
        end
        clear html1.QCdir html1.slicesstr html1k3
    end
end

fprintf('\n')
fprintf('<b>17.3. Normalized mean gray and white matter signals: "17-HRF-deconvolution" vs. "15-temporal-filtering":</b>\n')
for html1k=1:length(sub_dir)
    for html1k2=1:runs(html1k)
        if runs(html1k)==1
            html1.QCdir = ['.' erase(sub_dir{html1k},home_dir(1:end-1)) 'QC/' '17_deconvolution/'];
        else
            html1.QCdir = ['.' erase(sub_dir{html1k},home_dir(1:end-1)) 'QC/' '17_deconvolution/' 'run' num2str(html1k2) '/'];
        end
        if exist([home_dir html1.QCdir(3:end) 'mean_timeseries_15filter_vs_17deconv_allSlices.jpg'],'file')
            if (html1k2==1) && (runs(html1k)==1)
                disp(sprintf('      Subject %d of %d - %s<a href="%s"> global mean signal (all slices combined)</a>',html1k,length(sub_dir),subnames{html1k},[html1.QCdir 'mean_timeseries_15filter_vs_17deconv_allSlices.jpg']))
            elseif (html1k2==1) && (runs(html1k)~=1)
                disp(sprintf('      Subject %d of %d, run %d of %d - %s<a href="%s"> global mean signal (all slices combined)</a>',html1k,length(sub_dir),html1k2,runs(html1k),subnames{html1k},[html1.QCdir 'mean_timeseries_15filter_vs_17deconv_allSlices.jpg']))
            else
                disp(sprintf('                      run %d of %d - %s<a href="%s"> global mean signal (all slices combined)</a>',html1k2,runs(html1k),subnames{html1k},[html1.QCdir 'mean_timeseries_15filter_vs_17deconv_allSlices.jpg']))
            end
        else
            if (html1k2==1) && (runs(html1k)==1)
                disp(sprintf('      Subject %d of %d - (pending execution)',html1k,length(sub_dir)))
            elseif (html1k2==1) && (runs(html1k)~=1)
                disp(sprintf('      Subject %d of %d, run %d of %d - (pending execution)',html1k,length(sub_dir),html1k2,runs(html1k)))
            else
                disp(sprintf('                      run %d of %d - (pending execution)',html1k2,runs(html1k)))
            end
        end
        html1.slicesstr = '';
        for html1k3=1:slices
            if exist([home_dir html1.QCdir(3:end) 'mean_timeseries_15filter_vs_17deconv_slice' sprintf('%.2d',html1k3) '.jpg'],'file')
                html1.slicesstr = strcat(html1.slicesstr,sprintf(' <a href="%s">slice-%d</a>,',[html1.QCdir 'mean_timeseries_15filter_vs_17deconv_slice' sprintf('%.2d',html1k3) '.jpg'],html1k3));
            else
                html1.slicesstr = strcat(html1.slicesstr,sprintf(' slice-%d,',html1k3));
            end
        end
        if exist([home_dir html1.QCdir(3:end) 'mean_timeseries_15filter_vs_17deconv_slice' sprintf('%.2d',1) '.jpg'],'file')
            disp(sprintf('                                   (%s)',html1.slicesstr(2:end-1)))
        end
        clear html1.QCdir html1.slicesstr html1k3
    end
end

fprintf('\n')
fprintf('<b>17.4. Histogram of voxel-to-voxel correlations within the gray and white matter after "17-HRF-deconvolution":</b>\n')
for html1k=1:length(sub_dir)
    for html1k2=1:runs(html1k)
        if runs(html1k)==1
            html1.QCdir = ['.' erase(sub_dir{html1k},home_dir(1:end-1)) 'QC/' '17_deconvolution/'];
        else
            html1.QCdir = ['.' erase(sub_dir{html1k},home_dir(1:end-1)) 'QC/' '17_deconvolution/' 'run' num2str(html1k2) '/'];
        end
        if exist([home_dir html1.QCdir(3:end) 'hist_FC_compare_17deconv_allSlices.jpg'],'file')
            if (html1k2==1) && (runs(html1k)==1)
                disp(sprintf('      Subject %d of %d - %s<a href="%s"> histogram of functional connectivity (all slices combined)</a>',html1k,length(sub_dir),subnames{html1k},[html1.QCdir 'hist_FC_compare_17deconv_allSlices.jpg']))
            elseif (html1k2==1) && (runs(html1k)~=1)
                disp(sprintf('      Subject %d of %d, run %d of %d - %s<a href="%s"> histogram of functional connectivity (all slices combined)</a>',html1k,length(sub_dir),html1k2,runs(html1k),subnames{html1k},[html1.QCdir 'hist_FC_compare_17deconv_allSlices.jpg']))
            else
                disp(sprintf('                      run %d of %d - %s<a href="%s"> histogram of functional connectivity (all slices combined)</a>',html1k2,runs(html1k),subnames{html1k},[html1.QCdir 'hist_FC_compare_17deconv_allSlices.jpg']))
            end
        else
            if (html1k2==1) && (runs(html1k)==1)
                disp(sprintf('      Subject %d of %d - (pending execution)',html1k,length(sub_dir)))
            elseif (html1k2==1) && (runs(html1k)~=1)
                disp(sprintf('      Subject %d of %d, run %d of %d - (pending execution)',html1k,length(sub_dir),html1k2,runs(html1k)))
            else
                disp(sprintf('                      run %d of %d - (pending execution)',html1k2,runs(html1k)))
            end
        end
        html1.slicesstr = '';
        for html1k3=1:slices
            if exist([home_dir html1.QCdir(3:end) 'hist_FC_compare_17deconv_slice' sprintf('%.2d',html1k3) '.jpg'],'file')
                html1.slicesstr = strcat(html1.slicesstr,sprintf(' <a href="%s">slice-%d</a>,',[html1.QCdir 'hist_FC_compare_17deconv_slice' sprintf('%.2d',html1k3) '.jpg'],html1k3));
            else
                html1.slicesstr = strcat(html1.slicesstr,sprintf(' slice-%d,',html1k3));
            end
        end
        if exist([home_dir html1.QCdir(3:end) 'hist_FC_compare_17deconv_slice' sprintf('%.2d',1) '.jpg'],'file')
            disp(sprintf('                                   (%s)',html1.slicesstr(2:end-1)))
        end
        clear html1.QCdir html1.slicesstr html1k3
    end
end

fprintf('\n')
fprintf('<b>17.5. Histogram of voxel-to-voxel correlations: "17-HRF-deconvolution" vs. "15-temporal-filtering":</b>\n')
for html1k=1:length(sub_dir)
    for html1k2=1:runs(html1k)
        if runs(html1k)==1
            html1.QCdir = ['.' erase(sub_dir{html1k},home_dir(1:end-1)) 'QC/' '17_deconvolution/'];
        else
            html1.QCdir = ['.' erase(sub_dir{html1k},home_dir(1:end-1)) 'QC/' '17_deconvolution/' 'run' num2str(html1k2) '/'];
        end
        if exist([home_dir html1.QCdir(3:end) 'hist_FC_compare_15filter_17deconv_allSlices.jpg'],'file')
            if (html1k2==1) && (runs(html1k)==1)
                disp(sprintf('      Subject %d of %d - %s<a href="%s"> histogram of functional connectivity (all slices combined)</a>',html1k,length(sub_dir),subnames{html1k},[html1.QCdir 'hist_FC_compare_15filter_17deconv_allSlices.jpg']))
            elseif (html1k2==1) && (runs(html1k)~=1)
                disp(sprintf('      Subject %d of %d, run %d of %d - %s<a href="%s"> histogram of functional connectivity (all slices combined)</a>',html1k,length(sub_dir),html1k2,runs(html1k),subnames{html1k},[html1.QCdir 'hist_FC_compare_15filter_17deconv_allSlices.jpg']))
            else
                disp(sprintf('                      run %d of %d - %s<a href="%s"> histogram of functional connectivity (all slices combined)</a>',html1k2,runs(html1k),subnames{html1k},[html1.QCdir 'hist_FC_compare_15filter_17deconv_allSlices.jpg']))
            end
        else
            if (html1k2==1) && (runs(html1k)==1)
                disp(sprintf('      Subject %d of %d - (pending execution)',html1k,length(sub_dir)))
            elseif (html1k2==1) && (runs(html1k)~=1)
                disp(sprintf('      Subject %d of %d, run %d of %d - (pending execution)',html1k,length(sub_dir),html1k2,runs(html1k)))
            else
                disp(sprintf('                      run %d of %d - (pending execution)',html1k2,runs(html1k)))
            end
        end
        html1.slicesstr = '';
        for html1k3=1:slices
            if exist([home_dir html1.QCdir(3:end) 'hist_FC_compare_15filter_17deconv_slice' sprintf('%.2d',html1k3) '.jpg'],'file')
                html1.slicesstr = strcat(html1.slicesstr,sprintf(' <a href="%s">slice-%d</a>,',[html1.QCdir 'hist_FC_compare_15filter_17deconv_slice' sprintf('%.2d',html1k3) '.jpg'],html1k3));
            else
                html1.slicesstr = strcat(html1.slicesstr,sprintf(' slice-%d,',html1k3));
            end
        end
        if exist([home_dir html1.QCdir(3:end) 'hist_FC_compare_15filter_17deconv_slice' sprintf('%.2d',1) '.jpg'],'file')
            disp(sprintf('                                   (%s)',html1.slicesstr(2:end-1)))
        end
        clear html1.QCdir html1.slicesstr html1k3
    end
end

else
fprintf('<b>Not in pipeline</b>\n')
fprintf('</p>')
end
end

%% Temporal signal-to-noise ratio (tSNR) graphs and maps (at the end of all pre-processing steps)
if exist('choices_preproc','var')
if (choices_preproc(5)==1)||(choices_preproc(6)==1)||(choices_preproc(8)==1)||(choices_preproc(9)==1)||(choices_preproc(11)==1)||(choices_preproc(12)==1)||(choices_preproc(13)==1)||(choices_preproc(14)==1)
fprintf('<p style="font-size:14px ; color:#000000">')

fprintf('<b>Graphs showing tSNR across pre-processing steps:</b>\n')
for html1k=1:length(sub_dir)
    for html1k2=1:runs(html1k)
        if runs(html1k)==1
            html1.QCdir = ['.' erase(sub_dir{html1k},home_dir(1:end-1)) 'QC/' 'tSNR_graphs/'];
        else
            html1.QCdir = ['.' erase(sub_dir{html1k},home_dir(1:end-1)) 'QC/' 'tSNR_graphs/' 'run' num2str(html1k2) '/'];
        end
        if exist([home_dir html1.QCdir(3:end) 'tSNR_across_preproc_steps_avg.jpg'],'file')
            if (html1k2==1) && (runs(html1k)==1)
                disp(sprintf('      Subject %d of %d - %s<a href="%s"> Graph of median tSNR across pre-processing steps</a>',html1k,length(sub_dir),subnames{html1k},[html1.QCdir 'tSNR_across_preproc_steps_avg.jpg']))
            elseif (html1k2==1) && (runs(html1k)~=1)
                disp(sprintf('      Subject %d of %d, run %d of %d - %s<a href="%s"> Graph of median tSNR across pre-processing steps</a>',html1k,length(sub_dir),html1k2,runs(html1k),subnames{html1k},[html1.QCdir 'tSNR_across_preproc_steps_avg.jpg']))
            else
                disp(sprintf('                      run %d of %d - %s<a href="%s"> Graph of median tSNR across pre-processing steps</a>',html1k2,runs(html1k),subnames{html1k},[html1.QCdir 'tSNR_across_preproc_steps_avg.jpg']))
            end
        else
            if (html1k2==1) && (runs(html1k)==1)
                disp(sprintf('      Subject %d of %d - (pending execution)',html1k,length(sub_dir)))
            elseif (html1k2==1) && (runs(html1k)~=1)
                disp(sprintf('      Subject %d of %d, run %d of %d - (pending execution)',html1k,length(sub_dir),html1k2,runs(html1k)))
            else
                disp(sprintf('                      run %d of %d - (pending execution)',html1k2,runs(html1k)))
            end
        end
        if exist([home_dir html1.QCdir(3:end) 'tSNR_across_preproc_steps_slicewise.jpg'],'file')
            if (html1k2==1) && (runs(html1k)==1)
                disp(sprintf('                     - %s<a href="%s"> Graph of slice-wise median tSNR before and after pre-processing</a>',subnames{html1k},[html1.QCdir 'tSNR_across_preproc_steps_slicewise.jpg']))
            elseif (html1k2==1) && (runs(html1k)~=1)
                disp(sprintf('                                 - %s<a href="%s"> Graph of slice-wise median tSNR before and after pre-processing</a>',subnames{html1k},[html1.QCdir 'tSNR_across_preproc_steps_slicewise.jpg']))
            else
                disp(sprintf('                                 - %s<a href="%s"> Graph of slice-wise median tSNR before and after pre-processing</a>',subnames{html1k},[html1.QCdir 'tSNR_across_preproc_steps_slicewise.jpg']))
            end
        end
        clear html1.QCdir html1k3
    end
end

fprintf('\n')
fprintf('<b>tSNR maps across pre-processing steps:</b>\n')
for html1k=1:length(sub_dir)
    for html1k2=1:runs(html1k)
        if runs(html1k)==1
            html1.QCdir = ['.' erase(sub_dir{html1k},home_dir(1:end-1)) 'QC/' 'tSNR_graphs/'];
        else
            html1.QCdir = ['.' erase(sub_dir{html1k},home_dir(1:end-1)) 'QC/' 'tSNR_graphs/' 'run' num2str(html1k2) '/'];
        end
        if (choices_preproc(1)==1)
            if exist([home_dir html1.QCdir(3:end) 'tSNR_maps_1_rawdata_beforeNORDIC.jpg'],'file') || exist([home_dir html1.QCdir(3:end) 'tSNR_maps_4_rawdata.jpg'],'file')            
                html1.slicesstr = '';
                if (runs(html1k)==1)
                    html1.slicesstr = strcat(html1.slicesstr,sprintf('                     - %s Slice-wise tSNR maps of raw data (GM+WM+CSF): ',subnames{html1k}));
                else
                    html1.slicesstr = strcat(html1.slicesstr,sprintf('                                 - %s Slice-wise tSNR maps of raw data (GM+WM+CSF): ',subnames{html1k}));
                end
                if exist([home_dir html1.QCdir(3:end) 'tSNR_maps_1_rawdata_beforeNORDIC.jpg'],'file') && choices_preproc(1)==1
                    html1.slicesstr = strcat(html1.slicesstr,sprintf(' <a href="%s"> (1) before NORDIC denoising,</a>',[html1.QCdir 'tSNR_maps_1_rawdata_beforeNORDIC.jpg']));
                else
                    html1.slicesstr = strcat(html1.slicesstr,sprintf('  (1) before NORDIC denoising,'));
                end
                if exist([home_dir html1.QCdir(3:end) 'tSNR_maps_4_rawdata.jpg'],'file') && choices_preproc(4)==1
                    html1.slicesstr = strcat(html1.slicesstr,sprintf(' <a href="%s">(4) after NORDIC denoising,</a>',[html1.QCdir 'tSNR_maps_4_rawdata.jpg']));
                else
                    html1.slicesstr = strcat(html1.slicesstr,sprintf(' (4) after NORDIC denoising,'));
                end
                disp(sprintf(' %s ',html1.slicesstr(2:end-1)))
                clear html1.slicesstr
            else
                if (runs(html1k)==1)
                    disp(sprintf('                     - (pending execution)'))
                else
                    disp(sprintf('                                 - (pending execution)'))
                end
            end
        else
            if exist([home_dir html1.QCdir(3:end) 'tSNR_maps_4_rawdata.jpg'],'file')
                if (html1k2==1) && (runs(html1k)==1)
                    disp(sprintf('      Subject %d of %d - %s<a href="%s"> Slice-wise tSNR maps of raw data (GM+WM+CSF)</a>',html1k,length(sub_dir),subnames{html1k},[html1.QCdir 'tSNR_maps_4_rawdata.jpg']))
                elseif (html1k2==1) && (runs(html1k)~=1)
                    disp(sprintf('      Subject %d of %d, run %d of %d - %s<a href="%s"> Slice-wise tSNR maps of raw data</a>',html1k,length(sub_dir),html1k2,runs(html1k),subnames{html1k},[html1.QCdir 'tSNR_maps_4_rawdata.jpg']))
                else
                    disp(sprintf('                      run %d of %d - %s<a href="%s"> Slice-wise tSNR maps of raw data</a>',html1k2,runs(html1k),subnames{html1k},[html1.QCdir 'tSNR_maps_4_rawdata.jpg']))
                end
            else
                if (html1k2==1) && (runs(html1k)==1)
                    disp(sprintf('      Subject %d of %d - (pending execution)',html1k,length(sub_dir)))
                elseif (html1k2==1) && (runs(html1k)~=1)
                    disp(sprintf('      Subject %d of %d, run %d of %d - (pending execution)',html1k,length(sub_dir),html1k2,runs(html1k)))
                else
                    disp(sprintf('                      run %d of %d - (pending execution)',html1k2,runs(html1k)))
                end
            end
        end
        
        if (choices_preproc(6)==1)||(choices_preproc(8)==1)||(choices_preproc(9)==1)
        if exist([home_dir html1.QCdir(3:end) 'tSNR_maps_6_denoise1.jpg'],'file') || exist([home_dir html1.QCdir(3:end) 'tSNR_maps_8_motion_correction.jpg'],'file') || exist([home_dir html1.QCdir(3:end) 'tSNR_maps_9_denoise2.jpg'],'file')
        html1.slicesstr = '';
        if (runs(html1k)==1)
            html1.slicesstr = strcat(html1.slicesstr,sprintf('                     - %s tSNR maps (GM+WM+CSF) after: ',subnames{html1k}));
        else
            html1.slicesstr = strcat(html1.slicesstr,sprintf('                                 - %s tSNR maps (GM+WM+CSF) after: ',subnames{html1k}));
        end
        if exist([home_dir html1.QCdir(3:end) 'tSNR_maps_6_denoise1.jpg'],'file') && choices_preproc(6)==1
            html1.slicesstr = strcat(html1.slicesstr,sprintf(' <a href="%s"> (6) not-cord regression (denoise1),</a>',[html1.QCdir 'tSNR_maps_6_denoise1.jpg']));
        else
            html1.slicesstr = strcat(html1.slicesstr,sprintf('  (6) not-cord regression (denoise1),'));
        end
        if exist([home_dir html1.QCdir(3:end) 'tSNR_maps_8_motion_correction.jpg'],'file') && choices_preproc(8)==1
            html1.slicesstr = strcat(html1.slicesstr,sprintf(' <a href="%s">(8) motion correction,</a>',[html1.QCdir 'tSNR_maps_8_motion_correction.jpg']));
        else
            html1.slicesstr = strcat(html1.slicesstr,sprintf(' (8) motion correction,'));
        end
        if exist([home_dir html1.QCdir(3:end) 'tSNR_maps_9_denoise2.jpg'],'file') && choices_preproc(9)==1
            html1.slicesstr = strcat(html1.slicesstr,sprintf(' <a href="%s">(9) RETROICOR (denoise2)</a>,',[html1.QCdir 'tSNR_maps_9_denoise2.jpg']));
        else
            html1.slicesstr = strcat(html1.slicesstr,sprintf(' (9) RETROICOR (denoise2),'));
        end
        disp(sprintf(' %s ',html1.slicesstr(2:end-1)))
        clear html1.slicesstr
        else
        if (runs(html1k)==1)
            disp(sprintf('                     - (pending execution)'))
        else
            disp(sprintf('                                 - (pending execution)'))
        end
        end
        end

        if (choices_preproc(11)==1)||(choices_preproc(12)==1)||(choices_preproc(13)==1)||(choices_preproc(14)==1)
        if exist([home_dir html1.QCdir(3:end) 'tSNR_maps_GM_11_coreg.jpg'],'file') || exist([home_dir html1.QCdir(3:end) 'tSNR_maps_GM_12_CSFregress.jpg'],'file') || exist([home_dir html1.QCdir(3:end) 'tSNR_maps_GM_13_WMregress.jpg'],'file') || exist([home_dir html1.QCdir(3:end) 'tSNR_maps_GM_14_COVregress.jpg'],'file')
        html1.slicesstr = '';
        if (runs(html1k)==1)
            html1.slicesstr = strcat(html1.slicesstr,sprintf('                     - %s GRAY MATTER tSNR maps after: ',subnames{html1k}));
        else
            html1.slicesstr = strcat(html1.slicesstr,sprintf('                                 - %s GRAY MATTER tSNR maps after: ',subnames{html1k}));
        end
        if exist([home_dir html1.QCdir(3:end) 'tSNR_maps_GM_11_coreg.jpg'],'file') && choices_preproc(11)==1
            html1.slicesstr = strcat(html1.slicesstr,sprintf(' <a href="%s"> (11) co-registration,</a>',[html1.QCdir 'tSNR_maps_GM_11_coreg.jpg']));
        else
            html1.slicesstr = strcat(html1.slicesstr,sprintf('  (11) co-registration,'));
        end
        if exist([home_dir html1.QCdir(3:end) 'tSNR_maps_GM_12_CSFregress.jpg'],'file') && choices_preproc(12)==1
            html1.slicesstr = strcat(html1.slicesstr,sprintf(' <a href="%s">(12) CSF regression (denoise3),</a>',[html1.QCdir 'tSNR_maps_GM_12_CSFregress.jpg']));
        else
            html1.slicesstr = strcat(html1.slicesstr,sprintf(' (12) CSF regression (denoise3),'));
        end
        if exist([home_dir html1.QCdir(3:end) 'tSNR_maps_GM_13_WMregress.jpg'],'file') && choices_preproc(13)==1
            html1.slicesstr = strcat(html1.slicesstr,sprintf(' <a href="%s">(13) WM regression (denoise4)</a>,',[html1.QCdir 'tSNR_maps_GM_13_WMregress.jpg']));
        else
            html1.slicesstr = strcat(html1.slicesstr,sprintf(' (13) WM regression (denoise4),'));
        end
        if exist([home_dir html1.QCdir(3:end) 'tSNR_maps_GM_14_COVregress.jpg'],'file') && choices_preproc(14)==1
            html1.slicesstr = strcat(html1.slicesstr,sprintf(' <a href="%s">(14) COV regression (denoise5)</a>,',[html1.QCdir 'tSNR_maps_GM_14_COVregress.jpg']));
        else
            html1.slicesstr = strcat(html1.slicesstr,sprintf(' (14) COV regression (denoise5),'));
        end
        disp(sprintf(' %s ',html1.slicesstr(2:end-1)))
        clear html1.slicesstr
        else
        if (runs(html1k)==1)
            disp(sprintf('                     - (pending execution)'))
        else
            disp(sprintf('                                 - (pending execution)'))
        end
        end
        end
        
        if (choices_preproc(11)==1)||(choices_preproc(12)==1)||(choices_preproc(13)==1)||(choices_preproc(14)==1)
        if exist([home_dir html1.QCdir(3:end) 'tSNR_maps_WM_11_coreg.jpg'],'file') || exist([home_dir html1.QCdir(3:end) 'tSNR_maps_WM_12_CSFregress.jpg'],'file') || exist([home_dir html1.QCdir(3:end) 'tSNR_maps_WM_13_WMregress.jpg'],'file') || exist([home_dir html1.QCdir(3:end) 'tSNR_maps_WM_14_COVregress.jpg'],'file')
        html1.slicesstr = '';
        if (runs(html1k)==1)
            html1.slicesstr = strcat(html1.slicesstr,sprintf('                     - %s WHITE MATTER tSNR maps after: ',subnames{html1k}));
        else
            html1.slicesstr = strcat(html1.slicesstr,sprintf('                                 - %s WHITE MATTER tSNR maps after: ',subnames{html1k}));
        end
        if exist([home_dir html1.QCdir(3:end) 'tSNR_maps_WM_11_coreg.jpg'],'file') && choices_preproc(11)==1
            html1.slicesstr = strcat(html1.slicesstr,sprintf(' <a href="%s">(11) co-registration,</a>',[html1.QCdir 'tSNR_maps_WM_11_coreg.jpg']));
        else
            html1.slicesstr = strcat(html1.slicesstr,sprintf(' (11) co-registration,'));
        end
        if exist([home_dir html1.QCdir(3:end) 'tSNR_maps_WM_12_CSFregress.jpg'],'file') && choices_preproc(12)==1
            html1.slicesstr = strcat(html1.slicesstr,sprintf(' <a href="%s">(12) CSF regression (denoise3),</a>',[html1.QCdir 'tSNR_maps_WM_12_CSFregress.jpg']));
        else
            html1.slicesstr = strcat(html1.slicesstr,sprintf(' (12) CSF regression (denoise3),'));
        end
        if exist([home_dir html1.QCdir(3:end) 'tSNR_maps_WM_13_WMregress.jpg'],'file') && choices_preproc(13)==1
            html1.slicesstr = strcat(html1.slicesstr,sprintf(' <a href="%s">(13) WM regression (denoise4)</a>,',[html1.QCdir 'tSNR_maps_WM_13_WMregress.jpg']));
        else
            html1.slicesstr = strcat(html1.slicesstr,sprintf(' (13) WM regression (denoise4),'));
        end
        if exist([home_dir html1.QCdir(3:end) 'tSNR_maps_WM_14_COVregress.jpg'],'file') && choices_preproc(14)==1
            html1.slicesstr = strcat(html1.slicesstr,sprintf(' <a href="%s">(14) COV regression (denoise5)</a>,',[html1.QCdir 'tSNR_maps_WM_14_COVregress.jpg']));
        else
            html1.slicesstr = strcat(html1.slicesstr,sprintf(' (14) COV regression (denoise5),'));
        end
        disp(sprintf(' %s ',html1.slicesstr(2:end-1)))
        clear html1.slicesstr
        else
        if (runs(html1k)==1)
            disp(sprintf('                     - (pending execution)'))
        else
            disp(sprintf('                                 - (pending execution)'))
        end
        end
        end
        clear html1.QCdir html1k3
    end
end

else
fprintf('<b>Not performed because none of the following steps were in the pipeline: 5, 6, 8, 9, 11, 12, 13, 14</b>\n')
fprintf('</p>')
end
end

%% 19. Disk storage information
fprintf('<p style="font-size:14px ; color:#000000">')

fprintf('<b>Disk storage: size of subject folder at the end of pre-processing:</b>\n')
for html1k=1:length(sub_dir)
    html1.QCdir = ['.' erase(sub_dir{html1k},home_dir(1:end-1)) 'QC/' '19_disk_storage_info/'];
    if exist([home_dir html1.QCdir(3:end) 'disk_storage_bargraph.jpg'],'file')
        disp(sprintf('      Subject %d of %d - %s<a href="%s"> Disk storage bar graph</a>',html1k,length(sub_dir),subnames{html1k},[html1.QCdir 'disk_storage_bargraph.jpg']))
    else
        disp(sprintf('      Subject %d of %d - (pending execution)',html1k,length(sub_dir)))
    end
    clear html1.QCdir
end

fprintf('\n')
fprintf('<b>Execution time: time taken for each processing step as well as for each subject:</b>\n')
if exist([home_dir 'ExecutionTime_graph.jpg'],'file')
    disp(sprintf('      <a href="%s"> Execution time bar graphs</a>',[home_dir 'ExecutionTime_graph.jpg']))
else
    disp(sprintf('      Execution time bar graphs (processing is yet to complete OR file is missing)'))
end


%%
if exist('choices_postproc','var')
%% Post-processing (if chosen):
fprintf('\n')

%% 20. Quadrant time series extraction
if choices_postproc(1)==1
fprintf('<p style="font-size:14px ; color:#000000">')

fprintf('<b>20.1. Eroded gray and white matter quadrant masks for post-processing:</b>\n')
for html1k=1:length(sub_dir)
    if runs(html1k)==1
        html1.QCdir = ['.' erase(sub_dir{html1k},home_dir(1:end-1)) 'QC/' '20_quadrant_timeseries/'];
    else
        html1.QCdir = ['.' erase(sub_dir{html1k},home_dir(1:end-1)) 'QC/' '20_quadrant_timeseries/' 'run1' '/'];
    end    
    html1.slicesstr = '';
    for html1k3=1:slices
        if exist([home_dir html1.QCdir(3:end) 'QuadMask_for_postproc_slice' sprintf('%.2d',html1k3) '.jpg'],'file')
            html1.slicesstr = strcat(html1.slicesstr,sprintf(' <a href="%s">slice-%d</a>,',[html1.QCdir 'QuadMask_for_postproc_slice' sprintf('%.2d',html1k3) '.jpg'],html1k3));
        else
            html1.slicesstr = strcat(html1.slicesstr,sprintf(' slice-%d,',html1k3));
        end
    end
    if exist([home_dir html1.QCdir(3:end) 'QuadMask_for_postproc_slice' sprintf('%.2d',1) '.jpg'],'file')
        disp(sprintf('      Subject %d of %d - %s %s',html1k,length(sub_dir),subnames{html1k},html1.slicesstr(2:end-1)))
    else
        disp(sprintf('      Subject %d of %d - (pending execution)',html1k,length(sub_dir)))
    end
    clear html1.QCdir html1.slicesstr html1k3
end

fprintf('\n')
fprintf('<b>20.2. Time series within the four quadrants (left/right ventral/dorsal horns):</b>\n')
for html1k=1:length(sub_dir)
    for html1k2=1:runs(html1k)
        if runs(html1k)==1
            html1.QCdir = ['.' erase(sub_dir{html1k},home_dir(1:end-1)) 'QC/' '20_quadrant_timeseries/'];
        else
            html1.QCdir = ['.' erase(sub_dir{html1k},home_dir(1:end-1)) 'QC/' '20_quadrant_timeseries/' 'run' num2str(html1k2) '/'];
        end
        if exist([home_dir html1.QCdir(3:end) 'fMRI_ts_20quadTS_allSlices.jpg'],'file')
            if (html1k2==1) && (runs(html1k)==1)
                disp(sprintf('      Subject %d of %d - %s<a href="%s"> time series plot (all slices combined)</a>',html1k,length(sub_dir),subnames{html1k},[html1.QCdir 'fMRI_ts_20quadTS_allSlices.jpg']))
            elseif (html1k2==1) && (runs(html1k)~=1)
                disp(sprintf('      Subject %d of %d, run %d of %d - %s<a href="%s"> time series plot (all slices combined)</a>',html1k,length(sub_dir),html1k2,runs(html1k),subnames{html1k},[html1.QCdir 'fMRI_ts_20quadTS_allSlices.jpg']))
            else
                disp(sprintf('                      run %d of %d - %s<a href="%s"> time series plot (all slices combined)</a>',html1k2,runs(html1k),subnames{html1k},[html1.QCdir 'fMRI_ts_20quadTS_allSlices.jpg']))
            end
        else
            if (html1k2==1) && (runs(html1k)==1)
                disp(sprintf('      Subject %d of %d - (pending execution)',html1k,length(sub_dir)))
            elseif (html1k2==1) && (runs(html1k)~=1)
                disp(sprintf('      Subject %d of %d, run %d of %d - (pending execution)',html1k,length(sub_dir),html1k2,runs(html1k)))
            else
                disp(sprintf('                      run %d of %d - (pending execution)',html1k2,runs(html1k)))
            end
        end
        html1.slicesstr = '';
        for html1k3=1:slices
            if exist([home_dir html1.QCdir(3:end) 'fMRI_ts_20quadTS_slice' sprintf('%.2d',html1k3) '.jpg'],'file')
                html1.slicesstr = strcat(html1.slicesstr,sprintf(' <a href="%s">slice-%d</a>,',[html1.QCdir 'fMRI_ts_20quadTS_slice' sprintf('%.2d',html1k3) '.jpg'],html1k3));
            else
                html1.slicesstr = strcat(html1.slicesstr,sprintf(' slice-%d,',html1k3));
            end
        end
        if exist([home_dir html1.QCdir(3:end) 'fMRI_ts_20quadTS_slice' sprintf('%.2d',1) '.jpg'],'file')
            disp(sprintf('                                   (%s)',html1.slicesstr(2:end-1)))
        end
        clear html1.QCdir html1.slicesstr html1k3
    end
end

fprintf('\n')
fprintf('<b>20.3. Mean GRAY matter time series within each quadrant:</b>\n')
for html1k=1:length(sub_dir)
    for html1k2=1:runs(html1k)
        if runs(html1k)==1
            html1.QCdir = ['.' erase(sub_dir{html1k},home_dir(1:end-1)) 'QC/' '20_quadrant_timeseries/'];
        else
            html1.QCdir = ['.' erase(sub_dir{html1k},home_dir(1:end-1)) 'QC/' '20_quadrant_timeseries/' 'run' num2str(html1k2) '/'];
        end
        if exist([home_dir html1.QCdir(3:end) 'mean_timeseries_GM_20quadTS_allSlices.jpg'],'file')
            if (html1k2==1) && (runs(html1k)==1)
                disp(sprintf('      Subject %d of %d - %s<a href="%s"> mean time series GM (all slices combined)</a>',html1k,length(sub_dir),subnames{html1k},[html1.QCdir 'mean_timeseries_GM_20quadTS_allSlices.jpg']))
            elseif (html1k2==1) && (runs(html1k)~=1)
                disp(sprintf('      Subject %d of %d, run %d of %d - %s<a href="%s"> mean time series GM (all slices combined)</a>',html1k,length(sub_dir),html1k2,runs(html1k),subnames{html1k},[html1.QCdir 'mean_timeseries_GM_20quadTS_allSlices.jpg']))
            else
                disp(sprintf('                      run %d of %d - %s<a href="%s"> mean time series GM (all slices combined)</a>',html1k2,runs(html1k),subnames{html1k},[html1.QCdir 'mean_timeseries_GM_20quadTS_allSlices.jpg']))
            end
        else
            if (html1k2==1) && (runs(html1k)==1)
                disp(sprintf('      Subject %d of %d - (pending execution)',html1k,length(sub_dir)))
            elseif (html1k2==1) && (runs(html1k)~=1)
                disp(sprintf('      Subject %d of %d, run %d of %d - (pending execution)',html1k,length(sub_dir),html1k2,runs(html1k)))
            else
                disp(sprintf('                      run %d of %d - (pending execution)',html1k2,runs(html1k)))
            end
        end
        html1.slicesstr = '';
        for html1k3=1:slices
            if exist([home_dir html1.QCdir(3:end) 'mean_timeseries_GM_20quadTS_slice' sprintf('%.2d',html1k3) '.jpg'],'file')
                html1.slicesstr = strcat(html1.slicesstr,sprintf(' <a href="%s">slice-%d</a>,',[html1.QCdir 'mean_timeseries_GM_20quadTS_slice' sprintf('%.2d',html1k3) '.jpg'],html1k3));
            else
                html1.slicesstr = strcat(html1.slicesstr,sprintf(' slice-%d,',html1k3));
            end
        end
        if exist([home_dir html1.QCdir(3:end) 'mean_timeseries_GM_20quadTS_slice' sprintf('%.2d',1) '.jpg'],'file')
            disp(sprintf('                                   (%s)',html1.slicesstr(2:end-1)))
        end
        clear html1.QCdir html1.slicesstr html1k3
    end
end

if WM_flag==1
fprintf('\n')
fprintf('<b>20.4. Mean WHITE matter time series within each quadrant:</b>\n')
for html1k=1:length(sub_dir)
    for html1k2=1:runs(html1k)
        if runs(html1k)==1
            html1.QCdir = ['.' erase(sub_dir{html1k},home_dir(1:end-1)) 'QC/' '20_quadrant_timeseries/'];
        else
            html1.QCdir = ['.' erase(sub_dir{html1k},home_dir(1:end-1)) 'QC/' '20_quadrant_timeseries/' 'run' num2str(html1k2) '/'];
        end
        if exist([home_dir html1.QCdir(3:end) 'mean_timeseries_WM_20quadTS_allSlices.jpg'],'file')
            if (html1k2==1) && (runs(html1k)==1)
                disp(sprintf('      Subject %d of %d - %s<a href="%s"> mean time series WM (all slices combined)</a>',html1k,length(sub_dir),subnames{html1k},[html1.QCdir 'mean_timeseries_WM_20quadTS_allSlices.jpg']))
            elseif (html1k2==1) && (runs(html1k)~=1)
                disp(sprintf('      Subject %d of %d, run %d of %d - %s<a href="%s"> mean time series WM (all slices combined)</a>',html1k,length(sub_dir),html1k2,runs(html1k),subnames{html1k},[html1.QCdir 'mean_timeseries_WM_20quadTS_allSlices.jpg']))
            else
                disp(sprintf('                      run %d of %d - %s<a href="%s"> mean time series WM (all slices combined)</a>',html1k2,runs(html1k),subnames{html1k},[html1.QCdir 'mean_timeseries_WM_20quadTS_allSlices.jpg']))
            end
        else
            if (html1k2==1) && (runs(html1k)==1)
                disp(sprintf('      Subject %d of %d - (pending execution)',html1k,length(sub_dir)))
            elseif (html1k2==1) && (runs(html1k)~=1)
                disp(sprintf('      Subject %d of %d, run %d of %d - (pending execution)',html1k,length(sub_dir),html1k2,runs(html1k)))
            else
                disp(sprintf('                      run %d of %d - (pending execution)',html1k2,runs(html1k)))
            end
        end
        html1.slicesstr = '';
        for html1k3=1:slices
            if exist([home_dir html1.QCdir(3:end) 'mean_timeseries_WM_20quadTS_slice' sprintf('%.2d',html1k3) '.jpg'],'file')
                html1.slicesstr = strcat(html1.slicesstr,sprintf(' <a href="%s">slice-%d</a>,',[html1.QCdir 'mean_timeseries_WM_20quadTS_slice' sprintf('%.2d',html1k3) '.jpg'],html1k3));
            else
                html1.slicesstr = strcat(html1.slicesstr,sprintf(' slice-%d,',html1k3));
            end
        end
        if exist([home_dir html1.QCdir(3:end) 'mean_timeseries_WM_20quadTS_slice' sprintf('%.2d',1) '.jpg'],'file')
            disp(sprintf('                                   (%s)',html1.slicesstr(2:end-1)))
        end
        clear html1.QCdir html1.slicesstr html1k3
    end
end
else
fprintf('\n')
fprintf('<b>20.4. Mean WHITE matter time series within each quadrant:</b> not available because the user chose not to perform post-processing in the white matter.\n')
end

fprintf('\n')
fprintf('<b>20.5. Boxplot of voxel-to-voxel correlations within and between quadrants:</b>\n')
for html1k=1:length(sub_dir)
    for html1k2=1:runs(html1k)
        if runs(html1k)==1
            html1.QCdir = ['.' erase(sub_dir{html1k},home_dir(1:end-1)) 'QC/' '20_quadrant_timeseries/'];
        else
            html1.QCdir = ['.' erase(sub_dir{html1k},home_dir(1:end-1)) 'QC/' '20_quadrant_timeseries/' 'run' num2str(html1k2) '/'];
        end
        if exist([home_dir html1.QCdir(3:end) 'boxplot_FC_compare_20quadTS_allSlices.jpg'],'file')
            if (html1k2==1) && (runs(html1k)==1)
                disp(sprintf('      Subject %d of %d - %s<a href="%s"> boxplot of functional connectivity (pooled across all slices)</a>',html1k,length(sub_dir),subnames{html1k},[html1.QCdir 'boxplot_FC_compare_20quadTS_allSlices.jpg']))
            elseif (html1k2==1) && (runs(html1k)~=1)
                disp(sprintf('      Subject %d of %d, run %d of %d - %s<a href="%s"> boxplot of functional connectivity (pooled across all slices)</a>',html1k,length(sub_dir),html1k2,runs(html1k),subnames{html1k},[html1.QCdir 'boxplot_FC_compare_20quadTS_allSlices.jpg']))
            else
                disp(sprintf('                      run %d of %d - %s<a href="%s"> boxplot of functional connectivity (pooled across all slices)</a>',html1k2,runs(html1k),subnames{html1k},[html1.QCdir 'boxplot_FC_compare_20quadTS_allSlices.jpg']))
            end
        else
            if (html1k2==1) && (runs(html1k)==1)
                disp(sprintf('      Subject %d of %d - (pending execution)',html1k,length(sub_dir)))
            elseif (html1k2==1) && (runs(html1k)~=1)
                disp(sprintf('      Subject %d of %d, run %d of %d - (pending execution)',html1k,length(sub_dir),html1k2,runs(html1k)))
            else
                disp(sprintf('                      run %d of %d - (pending execution)',html1k2,runs(html1k)))
            end
        end
        html1.slicesstr = '';
        for html1k3=1:slices
            if exist([home_dir html1.QCdir(3:end) 'boxplot_FC_compare_20quadTS_slice' sprintf('%.2d',html1k3) '.jpg'],'file')
                html1.slicesstr = strcat(html1.slicesstr,sprintf(' <a href="%s">slice-%d</a>,',[html1.QCdir 'boxplot_FC_compare_20quadTS_slice' sprintf('%.2d',html1k3) '.jpg'],html1k3));
            else
                html1.slicesstr = strcat(html1.slicesstr,sprintf(' slice-%d,',html1k3));
            end
        end
        if exist([home_dir html1.QCdir(3:end) 'boxplot_FC_compare_20quadTS_slice' sprintf('%.2d',1) '.jpg'],'file')
            disp(sprintf('                                   (%s)',html1.slicesstr(2:end-1)))
        end
        clear html1.QCdir html1.slicesstr html1k3
    end
end

else
fprintf('<b>Not in pipeline</b>\n')
fprintf('</p>')
end

%% 21. Static functional connectivity
if choices_postproc(2)==1
fprintf('<p style="font-size:14px ; color:#000000">')

if ((SFC_type==1)||(SFC_type==3))&&(SFC_withinslice_flag==1)
fprintf('<b>21.1. Within-slice functional connectivity - Pearson''s correlation based FC:</b>\n')
for html1k=1:length(sub_dir)
    for html1k2=1:runs(html1k)
        if runs(html1k)==1
            html1.QCdir = ['.' erase(sub_dir{html1k},home_dir(1:end-1)) 'QC/' '21_static_functional_connectivity/'];
        else
            html1.QCdir = ['.' erase(sub_dir{html1k},home_dir(1:end-1)) 'QC/' '21_static_functional_connectivity/' 'run' num2str(html1k2) '/'];
        end
        if exist([home_dir html1.QCdir(3:end) 'FCmatrix_WithinSlice_allSlices.jpg'],'file')
            if (html1k2==1) && (runs(html1k)==1)
                disp(sprintf('      Subject %d of %d - %s<a href="%s"> within-slice FC matrix (median across all slices)</a>',html1k,length(sub_dir),subnames{html1k},[html1.QCdir 'FCmatrix_WithinSlice_allSlices.jpg']))
            elseif (html1k2==1) && (runs(html1k)~=1)
                disp(sprintf('      Subject %d of %d, run %d of %d - %s<a href="%s"> within-slice FC matrix (median across all slices)</a>',html1k,length(sub_dir),html1k2,runs(html1k),subnames{html1k},[html1.QCdir 'FCmatrix_WithinSlice_allSlices.jpg']))
            else
                disp(sprintf('                      run %d of %d - %s<a href="%s"> within-slice FC matrix (median across all slices)</a>',html1k2,runs(html1k),subnames{html1k},[html1.QCdir 'FCmatrix_WithinSlice_allSlices.jpg']))
            end
        else
            if (html1k2==1) && (runs(html1k)==1)
                disp(sprintf('      Subject %d of %d - (pending execution)',html1k,length(sub_dir)))
            elseif (html1k2==1) && (runs(html1k)~=1)
                disp(sprintf('      Subject %d of %d, run %d of %d - (pending execution)',html1k,length(sub_dir),html1k2,runs(html1k)))
            else
                disp(sprintf('                      run %d of %d - (pending execution)',html1k2,runs(html1k)))
            end
        end
        html1.slicesstr = '';
        for html1k3=1:slices
            if exist([home_dir html1.QCdir(3:end) 'FCmatrix_WithinSlice_slice' sprintf('%.2d',html1k3) '.jpg'],'file')
                html1.slicesstr = strcat(html1.slicesstr,sprintf(' <a href="%s">slice-%d</a>,',[html1.QCdir 'FCmatrix_WithinSlice_slice' sprintf('%.2d',html1k3) '.jpg'],html1k3));
            else
                html1.slicesstr = strcat(html1.slicesstr,sprintf(' slice-%d,',html1k3));
            end
        end
        if exist([home_dir html1.QCdir(3:end) 'FCmatrix_WithinSlice_slice' sprintf('%.2d',1) '.jpg'],'file')
            disp(sprintf('                                   (%s)',html1.slicesstr(2:end-1)))
        end
        clear html1.QCdir html1.slicesstr html1k3
    end
end
else
fprintf('\n')
if ~((SFC_type==1)||(SFC_type==3))&&(SFC_withinslice_flag==1)
fprintf('<b>21.1. Within-slice functional connectivity - Pearson''s correlation based FC:</b> not available because the user chose not to perform Pearson''s correlation based FC.\n')
elseif ((SFC_type==1)||(SFC_type==3))&&~(SFC_withinslice_flag==1)
fprintf('<b>21.1. Within-slice functional connectivity - Pearson''s correlation based FC:</b> not available because the user chose not to perform within-slice FC.\n')
elseif ~((SFC_type==1)||(SFC_type==3))&&~(SFC_withinslice_flag==1)
fprintf('<b>21.1. Within-slice functional connectivity - Pearson''s correlation based FC:</b> not available because the user chose not to perform within-slice FC and Pearson''s correlation based FC.\n')
end
end

fprintf('\n')
if ((SFC_type==2)||(SFC_type==3))&&(SFC_withinslice_flag==1)
fprintf('<b>21.2. Within-slice functional connectivity - partial correlation based FC:</b>\n')
for html1k=1:length(sub_dir)
    for html1k2=1:runs(html1k)
        if runs(html1k)==1
            html1.QCdir = ['.' erase(sub_dir{html1k},home_dir(1:end-1)) 'QC/' '21_static_functional_connectivity/'];
        else
            html1.QCdir = ['.' erase(sub_dir{html1k},home_dir(1:end-1)) 'QC/' '21_static_functional_connectivity/' 'run' num2str(html1k2) '/'];
        end
        if exist([home_dir html1.QCdir(3:end) 'FCmatrix_partialcorr_WithinSlice_allSlices.jpg'],'file')
            if (html1k2==1) && (runs(html1k)==1)
                disp(sprintf('      Subject %d of %d - %s<a href="%s"> within-slice FC matrix (median partial correlation across all slices)</a>',html1k,length(sub_dir),subnames{html1k},[html1.QCdir 'FCmatrix_partialcorr_WithinSlice_allSlices.jpg']))
            elseif (html1k2==1) && (runs(html1k)~=1)
                disp(sprintf('      Subject %d of %d, run %d of %d - %s<a href="%s"> within-slice FC matrix (median partial correlation across all slices)</a>',html1k,length(sub_dir),html1k2,runs(html1k),subnames{html1k},[html1.QCdir 'FCmatrix_partialcorr_WithinSlice_allSlices.jpg']))
            else
                disp(sprintf('                      run %d of %d - %s<a href="%s"> within-slice FC matrix (median partial correlation across all slices)</a>',html1k2,runs(html1k),subnames{html1k},[html1.QCdir 'FCmatrix_partialcorr_WithinSlice_allSlices.jpg']))
            end
        else
            if (html1k2==1) && (runs(html1k)==1)
                disp(sprintf('      Subject %d of %d - (pending execution)',html1k,length(sub_dir)))
            elseif (html1k2==1) && (runs(html1k)~=1)
                disp(sprintf('      Subject %d of %d, run %d of %d - (pending execution)',html1k,length(sub_dir),html1k2,runs(html1k)))
            else
                disp(sprintf('                      run %d of %d - (pending execution)',html1k2,runs(html1k)))
            end
        end
        html1.slicesstr = '';
        for html1k3=1:slices
            if exist([home_dir html1.QCdir(3:end) 'FCmatrix_partialcorr_WithinSlice_slice' sprintf('%.2d',html1k3) '.jpg'],'file')
                html1.slicesstr = strcat(html1.slicesstr,sprintf(' <a href="%s">slice-%d</a>,',[html1.QCdir 'FCmatrix_partialcorr_WithinSlice_slice' sprintf('%.2d',html1k3) '.jpg'],html1k3));
            else
                html1.slicesstr = strcat(html1.slicesstr,sprintf(' slice-%d,',html1k3));
            end
        end
        if exist([home_dir html1.QCdir(3:end) 'FCmatrix_partialcorr_WithinSlice_slice' sprintf('%.2d',1) '.jpg'],'file')
            disp(sprintf('                                   (%s)',html1.slicesstr(2:end-1)))
        end
        clear html1.QCdir html1.slicesstr html1k3
    end
end
else
fprintf('\n')
if ~((SFC_type==2)||(SFC_type==3))&&(SFC_withinslice_flag==1)
fprintf('<b>21.2. Within-slice functional connectivity - partial correlation based FC:</b> not available because the user chose not to perform partial correlation based FC.\n')
elseif ((SFC_type==2)||(SFC_type==3))&&~(SFC_withinslice_flag==1)
fprintf('<b>21.2. Within-slice functional connectivity - partial correlation based FC:</b> not available because the user chose not to perform within-slice FC.\n')
elseif ~((SFC_type==2)||(SFC_type==3))&&~(SFC_withinslice_flag==1)
fprintf('<b>21.2. Within-slice functional connectivity - partial correlation based FC:</b> not available because the user chose not to perform within-slice FC and partial correlation based FC.\n')
end
end

fprintf('\n')
if ((SFC_type==1)||(SFC_type==3))&&(SFC_betweenslice_flag==1)
fprintf('<b>21.3. Within- and between-slice functional connectivity - Pearson''s correlation based FC:</b>\n')
for html1k=1:length(sub_dir)
    for html1k2=1:runs(html1k)
        if runs(html1k)==1
            html1.QCdir = ['.' erase(sub_dir{html1k},home_dir(1:end-1)) 'QC/' '21_static_functional_connectivity/'];
        else
            html1.QCdir = ['.' erase(sub_dir{html1k},home_dir(1:end-1)) 'QC/' '21_static_functional_connectivity/' 'run' num2str(html1k2) '/'];
        end
        if exist([home_dir html1.QCdir(3:end) 'FCmatrix_Within_and_BetweenSlices_GM.jpg'],'file')
            if (html1k2==1) && (runs(html1k)==1)
                disp(sprintf('      Subject %d of %d - %s<a href="%s"> matrix of within- and between-slice GRAY MATTER FC across all slices</a>',html1k,length(sub_dir),subnames{html1k},[html1.QCdir 'FCmatrix_Within_and_BetweenSlices_GM.jpg']))
            elseif (html1k2==1) && (runs(html1k)~=1)
                disp(sprintf('      Subject %d of %d, run %d of %d - %s<a href="%s"> matrix of within- and between-slice GRAY MATTER FC across all slices</a>',html1k,length(sub_dir),html1k2,runs(html1k),subnames{html1k},[html1.QCdir 'FCmatrix_Within_and_BetweenSlices_GM.jpg']))
            else
                disp(sprintf('                      run %d of %d - %s<a href="%s"> matrix of within- and between-slice GRAY MATTER FC across all slices</a>',html1k2,runs(html1k),subnames{html1k},[html1.QCdir 'FCmatrix_Within_and_BetweenSlices_GM.jpg']))
            end
        else
            if (html1k2==1) && (runs(html1k)==1)
                disp(sprintf('      Subject %d of %d - (pending execution)',html1k,length(sub_dir)))
            elseif (html1k2==1) && (runs(html1k)~=1)
                disp(sprintf('      Subject %d of %d, run %d of %d - (pending execution)',html1k,length(sub_dir),html1k2,runs(html1k)))
            else
                disp(sprintf('                      run %d of %d - (pending execution)',html1k2,runs(html1k)))
            end
        end
        if WM_flag==1
        if exist([home_dir html1.QCdir(3:end) 'FCmatrix_Within_and_BetweenSlices_WM.jpg'],'file')
            if (html1k2==1) && (runs(html1k)==1)
                disp(sprintf('                     - %s<a href="%s"> matrix of within- and between-slice WHITE MATTER FC across all slices</a>',subnames{html1k},[html1.QCdir 'FCmatrix_Within_and_BetweenSlices_WM.jpg']))
            elseif (html1k2==1) && (runs(html1k)~=1)
                disp(sprintf('                                 - %s<a href="%s"> matrix of within- and between-slice WHITE MATTER FC across all slices</a>',subnames{html1k},[html1.QCdir 'FCmatrix_Within_and_BetweenSlices_WM.jpg']))
            else
                disp(sprintf('                                 - %s<a href="%s"> matrix of within- and between-slice WHITE MATTER FC across all slices</a>',subnames{html1k},[html1.QCdir 'FCmatrix_Within_and_BetweenSlices_WM.jpg']))
            end
        end
        end
        clear html1.QCdir html1.slicesstr html1k3
    end
end
else
fprintf('\n')
if ~((SFC_type==1)||(SFC_type==3))&&(SFC_betweenslice_flag==1)
fprintf('<b>21.3. Within- and between-slice functional connectivity - Pearson''s correlation based FC:</b> not available because the user chose not to perform Pearson''s correlation based FC.\n')
elseif ((SFC_type==1)||(SFC_type==3))&&~(SFC_betweenslice_flag==1)
fprintf('<b>21.3. Within- and between-slice functional connectivity - Pearson''s correlation based FC:</b> not available because the user chose not to perform between-slice FC.\n')
elseif ~((SFC_type==1)||(SFC_type==3))&&~(SFC_betweenslice_flag==1)
fprintf('<b>21.3. Within- and between-slice functional connectivity - Pearson''s correlation based FC:</b> not available because the user chose not to perform between-slice FC and Pearson''s correlation based FC.\n')
end
end

fprintf('\n')
if ((SFC_type==2)||(SFC_type==3))&&(SFC_betweenslice_flag==1)&&(dont_do_pcor_betweenslice==0)
fprintf('<b>21.4. Within- and between-slice functional connectivity - partial correlation based FC:</b>\n')
for html1k=1:length(sub_dir)
    for html1k2=1:runs(html1k)
        if runs(html1k)==1
            html1.QCdir = ['.' erase(sub_dir{html1k},home_dir(1:end-1)) 'QC/' '21_static_functional_connectivity/'];
        else
            html1.QCdir = ['.' erase(sub_dir{html1k},home_dir(1:end-1)) 'QC/' '21_static_functional_connectivity/' 'run' num2str(html1k2) '/'];
        end
        if exist([home_dir html1.QCdir(3:end) 'FCmatrix_partialcorr_Within_and_BetweenSlices_GM.jpg'],'file')
            if (html1k2==1) && (runs(html1k)==1)
                disp(sprintf('      Subject %d of %d - %s<a href="%s"> matrix of within- and between-slice GRAY MATTER FC across all slices (partial correlation)</a>',html1k,length(sub_dir),subnames{html1k},[html1.QCdir 'FCmatrix_partialcorr_Within_and_BetweenSlices_GM.jpg']))
            elseif (html1k2==1) && (runs(html1k)~=1)
                disp(sprintf('      Subject %d of %d, run %d of %d - %s<a href="%s"> matrix of within- and between-slice GRAY MATTER FC across all slices (partial correlation)</a>',html1k,length(sub_dir),html1k2,runs(html1k),subnames{html1k},[html1.QCdir 'FCmatrix_partialcorr_Within_and_BetweenSlices_GM.jpg']))
            else
                disp(sprintf('                      run %d of %d - %s<a href="%s"> matrix of within- and between-slice GRAY MATTER FC across all slices (partial correlation)</a>',html1k2,runs(html1k),subnames{html1k},[html1.QCdir 'FCmatrix_partialcorr_Within_and_BetweenSlices_GM.jpg']))
            end
        else
            if (html1k2==1) && (runs(html1k)==1)
                disp(sprintf('      Subject %d of %d - (pending execution)',html1k,length(sub_dir)))
            elseif (html1k2==1) && (runs(html1k)~=1)
                disp(sprintf('      Subject %d of %d, run %d of %d - (pending execution)',html1k,length(sub_dir),html1k2,runs(html1k)))
            else
                disp(sprintf('                      run %d of %d - (pending execution)',html1k2,runs(html1k)))
            end
        end
        if WM_flag==1
        if exist([home_dir html1.QCdir(3:end) 'FCmatrix_partialcorr_Within_and_BetweenSlices_WM.jpg'],'file')
            if (html1k2==1) && (runs(html1k)==1)
                disp(sprintf('                     - %s<a href="%s"> matrix of within- and between-slice WHITE MATTER FC across all slices (partial correlation)</a>',subnames{html1k},[html1.QCdir 'FCmatrix_partialcorr_Within_and_BetweenSlices_WM.jpg']))
            elseif (html1k2==1) && (runs(html1k)~=1)
                disp(sprintf('                                 - %s<a href="%s"> matrix of within- and between-slice WHITE MATTER FC across all slices (partial correlation)</a>',subnames{html1k},[html1.QCdir 'FCmatrix_partialcorr_Within_and_BetweenSlices_WM.jpg']))
            else
                disp(sprintf('                                 - %s<a href="%s"> matrix of within- and between-slice WHITE MATTER FC across all slices (partial correlation)</a>',subnames{html1k},[html1.QCdir 'FCmatrix_partialcorr_Within_and_BetweenSlices_WM.jpg']))
            end
        end
        end
        clear html1.QCdir html1.slicesstr html1k3
    end
end
else
fprintf('\n')
if ~((SFC_type==2)||(SFC_type==3))&&(SFC_betweenslice_flag==1)&&(dont_do_pcor_betweenslice==0)
fprintf('<b>21.4. Within- and between-slice functional connectivity - partial correlation based FC:</b> not available because the user chose not to perform partial correlation based FC.\n')
elseif ((SFC_type==2)||(SFC_type==3))&&~(SFC_betweenslice_flag==1)&&(dont_do_pcor_betweenslice==0)
fprintf('<b>21.4. Within- and between-slice functional connectivity - partial correlation based FC:</b> not available because the user chose not to perform between-slice FC.\n')
elseif ~((SFC_type==2)||(SFC_type==3))&&~(SFC_betweenslice_flag==1)&&(dont_do_pcor_betweenslice==0)
fprintf('<b>21.4. Within- and between-slice functional connectivity - partial correlation based FC:</b> not available because the user chose not to perform between-slice FC and partial correlation based FC.\n')
elseif (dont_do_pcor_betweenslice==1)
fprintf('<b>21.4. Within- and between-slice functional connectivity - partial correlation based FC:</b> not available because there is a default setting to not perform this computationally intensive step. To manually override the setting, set the value of the variable "dont_do_pcor_betweenslice" to 0 in the code "postproc_scfMRItb".\n')
end
end

else
fprintf('<b>Not in pipeline</b>\n')
fprintf('</p>')
end

%% 22. Fractional amplitude of low-frequency BOLD fluctuations (fALFF)
if choices_postproc(3)==1
fprintf('<p style="font-size:14px ; color:#000000">')

fprintf('<b>fALFF across slices and quadrants:</b>\n')
for html1k=1:length(sub_dir)
    for html1k2=1:runs(html1k)
        if runs(html1k)==1
            html1.QCdir = ['.' erase(sub_dir{html1k},home_dir(1:end-1)) 'QC/' '22_fALFF/'];
        else
            html1.QCdir = ['.' erase(sub_dir{html1k},home_dir(1:end-1)) 'QC/' '22_fALFF/' 'run' num2str(html1k2) '/'];
        end
        if exist([home_dir html1.QCdir(3:end) 'fALFF_across_slices.jpg'],'file')
            if (html1k2==1) && (runs(html1k)==1)
                disp(sprintf('      Subject %d of %d - %s<a href="%s"> fALFF across slices</a>',html1k,length(sub_dir),subnames{html1k},[html1.QCdir 'fALFF_across_slices.jpg']))
            elseif (html1k2==1) && (runs(html1k)~=1)
                disp(sprintf('      Subject %d of %d, run %d of %d - %s<a href="%s"> fALFF across slices</a>',html1k,length(sub_dir),html1k2,runs(html1k),subnames{html1k},[html1.QCdir 'fALFF_across_slices.jpg']))
            else
                disp(sprintf('                      run %d of %d - %s<a href="%s"> fALFF across slices</a>',html1k2,runs(html1k),subnames{html1k},[html1.QCdir 'fALFF_across_slices.jpg']))
            end
        else
            if (html1k2==1) && (runs(html1k)==1)
                disp(sprintf('      Subject %d of %d - (pending execution)',html1k,length(sub_dir)))
            elseif (html1k2==1) && (runs(html1k)~=1)
                disp(sprintf('      Subject %d of %d, run %d of %d - (pending execution)',html1k,length(sub_dir),html1k2,runs(html1k)))
            else
                disp(sprintf('                      run %d of %d - (pending execution)',html1k2,runs(html1k)))
            end
        end
        clear html1.QCdir
    end
end

else
fprintf('<b>Not in pipeline</b>\n')
fprintf('</p>')
end

%% 23. Simple stats
if choices_postproc(4)==1
fprintf('<p style="font-size:14px ; color:#000000">')

if ((SFC_type==1)||(SFC_type==3))&&(SFC_withinslice_flag==1)
fprintf('<b>23.1. Stats: within-slice functional connectivity (Pearson''s correlation):</b>\n')
    html1.QCdir = './stats/figures/';
    if exist([home_dir html1.QCdir(3:end) 'stats_SFC_WithinSlice_SlicesPooled.jpg'],'file')
        disp(sprintf('      <a href="%s"> significant group-level T-statistics for within-slice FC (FC pooled across all slices for each connection)</a>',[html1.QCdir 'stats_SFC_WithinSlice_SlicesPooled.jpg']))
    else
        disp(sprintf('      (pending execution)'))
    end
    html1.slicesstr = '';
    for html1k3=1:slices
        if exist([home_dir html1.QCdir(3:end) 'stats_SFC_WithinSlice_slice' sprintf('%.2d',html1k3) '.jpg'],'file')
            html1.slicesstr = strcat(html1.slicesstr,sprintf(' <a href="%s">slice-%d</a>,',[html1.QCdir 'stats_SFC_WithinSlice_slice' sprintf('%.2d',html1k3) '.jpg'],html1k3));
        else
            html1.slicesstr = strcat(html1.slicesstr,sprintf(' slice-%d,',html1k3));
        end
    end
    if exist([home_dir html1.QCdir(3:end) 'stats_SFC_WithinSlice_slice' sprintf('%.2d',1) '.jpg'],'file')
        disp(sprintf('          (%s)',html1.slicesstr(2:end-1)))
    end
    clear html1.QCdir html1.slicesstr html1k3
else
    fprintf('\n')
    if ~((SFC_type==1)||(SFC_type==3))&&(SFC_withinslice_flag==1)
        fprintf('<b>23.1. Stats: within-slice functional connectivity (Pearson''s correlation):</b> not available because the user chose not to perform Pearson''s correlation based FC.\n')
    elseif ((SFC_type==1)||(SFC_type==3))&&~(SFC_withinslice_flag==1)
        fprintf('<b>23.1. Stats: within-slice functional connectivity (Pearson''s correlation):</b> not available because the user chose not to perform within-slice FC.\n')
    elseif ~((SFC_type==1)||(SFC_type==3))&&~(SFC_withinslice_flag==1)
        fprintf('<b>23.1. Stats: within-slice functional connectivity (Pearson''s correlation):</b> not available because the user chose not to perform within-slice FC and Pearson''s correlation based FC.\n')
    end
end

fprintf('\n')
if ((SFC_type==2)||(SFC_type==3))&&(SFC_withinslice_flag==1)
fprintf('<b>23.2. Stats: within-slice functional connectivity (partial correlation):</b>\n')
    html1.QCdir = './stats/figures/';
    if exist([home_dir html1.QCdir(3:end) 'stats_SFC_pcor_WithinSlice_SlicesPooled.jpg'],'file')
        disp(sprintf('      <a href="%s"> significant group-level T-statistics for within-slice FC (partial correlation values pooled across all slices for each connection)</a>',[html1.QCdir 'stats_SFC_pcor_WithinSlice_SlicesPooled.jpg']))
    else
        disp(sprintf('      (pending execution)'))
    end
    html1.slicesstr = '';
    for html1k3=1:slices
        if exist([home_dir html1.QCdir(3:end) 'stats_SFC_pcor_WithinSlice_slice' sprintf('%.2d',html1k3) '.jpg'],'file')
            html1.slicesstr = strcat(html1.slicesstr,sprintf(' <a href="%s">slice-%d</a>,',[html1.QCdir 'stats_SFC_pcor_WithinSlice_slice' sprintf('%.2d',html1k3) '.jpg'],html1k3));
        else
            html1.slicesstr = strcat(html1.slicesstr,sprintf(' slice-%d,',html1k3));
        end
    end
    if exist([home_dir html1.QCdir(3:end) 'stats_SFC_pcor_WithinSlice_slice' sprintf('%.2d',1) '.jpg'],'file')
        disp(sprintf('          (%s)',html1.slicesstr(2:end-1)))
    end
    clear html1.QCdir html1.slicesstr html1k3
else
    fprintf('\n')
    if ~((SFC_type==2)||(SFC_type==3))&&(SFC_withinslice_flag==1)
        fprintf('<b>23.2. Stats: within-slice functional connectivity (partial correlation):</b> not available because the user chose not to perform partial correlation based FC.\n')
    elseif ((SFC_type==2)||(SFC_type==3))&&~(SFC_withinslice_flag==1)
        fprintf('<b>23.2. Stats: within-slice functional connectivity (partial correlation):</b> not available because the user chose not to perform within-slice FC.\n')
    elseif ~((SFC_type==2)||(SFC_type==3))&&~(SFC_withinslice_flag==1)
        fprintf('<b>23.2. Stats: within-slice functional connectivity (partial correlation):</b> not available because the user chose not to perform within-slice FC and partial correlation based FC.\n')
    end
end

fprintf('\n')
if ((SFC_type==1)||(SFC_type==3))&&(SFC_betweenslice_flag==1)
fprintf('<b>23.3. Stats: within- and between-slices functional connectivity (Pearson''s correlation):</b>\n')
    html1.QCdir = './stats/figures/';
    if exist([home_dir html1.QCdir(3:end) 'stats_SFC_Within_and_BetweenSlices_GM.jpg'],'file')
        disp(sprintf('      <a href="%s"> significant group-level T-statistics for within- and between-slice GRAY MATTER FC across all slices</a>',[html1.QCdir 'stats_SFC_Within_and_BetweenSlices_GM.jpg']))
    else
        disp(sprintf('      (pending execution)'))
    end
    if WM_flag==1
        if exist([home_dir html1.QCdir(3:end) 'stats_SFC_Within_and_BetweenSlices_WM.jpg'],'file')
        disp(sprintf('      <a href="%s"> significant group-level T-statistics for within- and between-slice WHITE MATTER FC across all slices</a>',[html1.QCdir 'stats_SFC_Within_and_BetweenSlices_WM.jpg']))
        end
    end
    if exist([home_dir html1.QCdir(3:end) 'stats_SFC_Within_and_BetweenSlices_GM_unthresholded.jpg'],'file')
        fprintf('\n')
        disp(sprintf('      <a href="%s"> unthresholded group-level T-statistics for within- and between-slice GRAY MATTER FC across all slices</a>',[html1.QCdir 'stats_SFC_Within_and_BetweenSlices_GM_unthresholded.jpg']))
    end
    if WM_flag==1
        if exist([home_dir html1.QCdir(3:end) 'stats_SFC_Within_and_BetweenSlices_WM_unthresholded.jpg'],'file')
        disp(sprintf('      <a href="%s"> unthresholded group-level T-statistics for within- and between-slice WHITE MATTER FC across all slices</a>',[html1.QCdir 'stats_SFC_Within_and_BetweenSlices_WM_unthresholded.jpg']))
        end
    end
    clear html1.QCdir html1.slicesstr html1k3
else
    fprintf('\n')
    if ~((SFC_type==1)||(SFC_type==3))&&(SFC_betweenslice_flag==1)
        fprintf('<b>23.3. Stats: within- and between-slice functional connectivity (Pearson''s correlation):</b> not available because the user chose not to perform Pearson''s correlation based FC.\n')
    elseif ((SFC_type==1)||(SFC_type==3))&&~(SFC_betweenslice_flag==1)
        fprintf('<b>23.3. Stats: within- and between-slice functional connectivity (Pearson''s correlation):</b> not available because the user chose not to perform between-slice FC.\n')
    elseif ~((SFC_type==1)||(SFC_type==3))&&~(SFC_betweenslice_flag==1)
        fprintf('<b>23.3. Stats: within- and between-slice functional connectivity (Pearson''s correlation):</b> not available because the user chose not to perform between-slice FC and Pearson''s correlation based FC.\n')
    end
end

fprintf('\n')
if ((SFC_type==2)||(SFC_type==3))&&(SFC_betweenslice_flag==1)&&(dont_do_pcor_betweenslice==0)
fprintf('<b>23.4. Stats: within- and between-slices functional connectivity (partial correlation):</b>\n')
    html1.QCdir = './stats/figures/';
    if exist([home_dir html1.QCdir(3:end) 'stats_SFC_pcor_Within_and_BetweenSlices_GM.jpg'],'file')
        disp(sprintf('      <a href="%s"> significant group-level T-statistics for within- and between-slice GRAY MATTER FC across all slices (partial correlation)</a>',[html1.QCdir 'stats_SFC_pcor_Within_and_BetweenSlices_GM.jpg']))
    else
        disp(sprintf('      (pending execution)'))
    end
    if WM_flag==1
        if exist([home_dir html1.QCdir(3:end) 'stats_SFC_pcor_Within_and_BetweenSlices_WM.jpg'],'file')
        disp(sprintf('      <a href="%s"> significant group-level T-statistics for within- and between-slice WHITE MATTER FC across all slices (partial correlation)</a>',[html1.QCdir 'stats_SFC_pcor_Within_and_BetweenSlices_WM.jpg']))
        end
    end
    if exist([home_dir html1.QCdir(3:end) 'stats_SFC_pcor_Within_and_BetweenSlices_GM_unthresholded.jpg'],'file')
        fprintf('\n')
        disp(sprintf('      <a href="%s"> unthresholded group-level T-statistics for within- and between-slice GRAY MATTER FC across all slices</a>',[html1.QCdir 'stats_SFC_pcor_Within_and_BetweenSlices_GM_unthresholded.jpg']))
    end
    if WM_flag==1
        if exist([home_dir html1.QCdir(3:end) 'stats_SFC_pcor_Within_and_BetweenSlices_WM_unthresholded.jpg'],'file')
        disp(sprintf('      <a href="%s"> unthresholded group-level T-statistics for within- and between-slice WHITE MATTER FC across all slices</a>',[html1.QCdir 'stats_SFC_pcor_Within_and_BetweenSlices_WM_unthresholded.jpg']))
        end
    end
    clear html1.QCdir html1.slicesstr html1k3
else
    fprintf('\n')
    if ~((SFC_type==2)||(SFC_type==3))&&(SFC_betweenslice_flag==1)&&(dont_do_pcor_betweenslice==0)
        fprintf('<b>23.4. Stats: within- and between-slice functional connectivity (partial correlation):</b> not available because the user chose not to perform partial correlation based FC.\n')
    elseif ((SFC_type==2)||(SFC_type==3))&&~(SFC_betweenslice_flag==1)&&(dont_do_pcor_betweenslice==0)
        fprintf('<b>23.4. Stats: within- and between-slice functional connectivity (partial correlation):</b> not available because the user chose not to perform between-slice FC.\n')
    elseif ~((SFC_type==2)||(SFC_type==3))&&~(SFC_betweenslice_flag==1)&&(dont_do_pcor_betweenslice==0)
        fprintf('<b>23.4. Stats: within- and between-slice functional connectivity (partial correlation):</b> not available because the user chose not to perform between-slice FC and partial correlation based FC.\n')
    elseif (dont_do_pcor_betweenslice==1)
        fprintf('<b>23.4. Within- and between-slice functional connectivity - partial correlation based FC:</b> not available because there is a default setting to not perform this computationally intensive step. To manually override the setting, set the value of the variable "dont_do_pcor_betweenslice" to 0 in the code "postproc_scfMRItb".\n')
    end
end

fprintf('\n')
fprintf('<b>23.5. Stats: fALFF across slices and quadrants:</b>\n')
html1.QCdir = './stats/figures/';
if exist([home_dir html1.QCdir(3:end) 'stats_fALFF.jpg'],'file')
    disp(sprintf('      <a href="%s"> significant group-level T-statistics for fALFF</a>',[html1.QCdir 'stats_fALFF.jpg']))
else
    disp(sprintf('      (pending execution)'))
end
clear html1.QCdir

else
fprintf('<b>Not in pipeline</b>\n')
fprintf('</p>')
end

end

%% Detailed execution log
fprintf('<p style="font-size:14px ; color:#000000">')
disp(sprintf('<a href="%s">Click here to see the detailed execution log</a>',[home_dir 'report_neptune_log.html']))
fprintf('</p>')
if html1.saveVIDflag==1, clear save_videos_steps; end
if html1.saveWMflag==1, clear WM_flag; end
if html1.saveSFC1flag==1, clear SFC_type; end
if html1.saveSFC2flag==1, clear SFC_withinslice_flag; end
if html1.saveSFC3flag==1, clear SFC_betweenslice_flag; end
if html1.saveSFC4flag==1, clear dont_do_pcor_betweenslice; end
clear html1*

%%
```
