## Supplementary material for "Neptune: a toolbox for spinal cord functional MRI data processing and quality assurance": Neptune software: report_neptune_log.m

```
fprintf('<p style="font-size:14px ; color:#000000">')
clk=clock;
month={'Jan','Feb','Mar','Apr','May','Jun','Jul','Aug','Sep','Oct','Nov','Dec'}';
disp(sprintf('Home directory: <i>%s</i>',home_dir(1:end-1)))
disp(sprintf('Page generation date/time: %.2d/%s/%.4d %.2dh:%.2dm:%.2ds',clk(3),month{clk(2)},clk(1),clk(4),clk(5),round(clk(6))))
disp(sprintf('<a href="%s">Click here to return to the execution outputs and quality control page</a>',[home_dir 'report_neptune.html']))
clear clk
fprintf('</p>')

%% DETAILED EXECUTION LOG
fprintf('<p style="font-size:14px ; color:#202020">')
fprintf('<b>The detailed log was saved to: <i>%s</i></b>\n',[home_dir sprintf('log_neptune_%s.txt',clk_init)])
fprintf('<b>The log is reproduced below as is:</b>\n\n')
try
for html1k=1:length(logg)
    fprintf('%s\n',logg{html1k})
end
catch
end
fprintf('\n')
fprintf('<b>__________________________________________________________________________________________</b>\n')
fprintf('</p>')
clear html1*
```
