## Supplementary material for "Neptune: a toolbox for spinal cord functional MRI data processing and quality assurance": Neptune software: add_AFNI_to_path.m

```
% is the path to AFNI already set?
[afninotfound, unixoutput] = unix('3dcopy');
% if not, then we will look for it...
if afninotfound
    % is AFNI in /abin?
    [afninotfound, unixoutput] = unix('/abin/3dcopy');
    if ~afninotfound % found it!
        setenv('PATH', [getenv('PATH') ':/abin']);
        fprintf('Adding /abin to Matlab''s path (for AFNI).\n');
    end
end
if afninotfound
    % is AFNI in ~/abin?
    [afninotfound, unixoutput] = unix('~/abin/3dcopy');
    if ~afninotfound % found it!
%         current_directory = robustdirname(pwd);
        current_directory = pwd;
        eval('cd ~');
        afni_directory = [pwd '/abin'];
        setenv('PATH', [getenv('PATH') ':' afni_directory]);
        disp(sprintf(['Adding ' afni_directory ' to Matlab''s path (for AFNI).']));
        eval(['cd(''' current_directory ''')']);
        clear current_directory
    end
end
if afninotfound
    % if AFNI is not in /abin nor ~/abin, then the user will have to add
    % the correct path to AFNI before this function will work.
    fprintf('*****\n');
    fprintf('The required AFNI functions cannot be found.\n');
    fprintf('AFNI needs to be added to Matlab''s path.\n');
    fprintf('You can do this with the following command:\n');
    fprintf(' \n');
    fprintf('   setenv(''PATH'', [getenv(''PATH'') '':/abin'']);\n');
    fprintf(' \n');
    fprintf('where (as an example) ''/abin'' is the AFNI directory.\n');
    fprintf('Exiting...\n');
    fprintf('*****\n');
    return;
end
if afninotfound==0
    fprintf('Successfully added AFNI to path\nLocation: %s\n',afni_directory)
end

clear afninotfound unixoutput afni_directory
```
