## Supplementary material for "Neptune: a toolbox for spinal cord functional MRI data processing and quality assurance": Neptune software: add_SCT_to_path.m

```
% 2019-04-11 : Robert L. Barry
% edited by Ranga Deshpande on 21/Nov/2021

% Is the path to SCT already set?
[sctnotfound, unixoutput] = unix('sct_propseg');
if sctnotfound
    current_directory = pwd;
    eval('cd ~');
    sdirSCT = dir('sct_*'); sdirSCT = sdirSCT(end);
    if ~isempty(sdirSCT)
        SCT_directory = [sdirSCT(1).folder '/' sdirSCT(1).name '/bin'];
        setenv('PATH', [getenv('PATH') ':' SCT_directory]); % path to SCT's folder in your computer
        fprintf('Successfully added SCT to path\nLocation: %s\n',SCT_directory)
    else
        fprintf('Could not add SCT to path\n')
    end
    cd(current_directory)
%     setenv('PATH', [getenv('PATH') ':/Applications/SCT/ver_curr/bin']);
%     disp(sprintf(['Adding SCT to Matlab''s path.']));
end
clear sctnotfound unixoutput current_directory sdirSCT
```
