## Supplementary material for "Neptune: a toolbox for spinal cord functional MRI data processing and quality assurance": Neptune software: calc_tsnr.m

```
function [tSNR, tSNR_slicewise, F_tsnr] = calc_tsnr(base_dir_sub, fname, suffix, mask)

%% CALCULATE MEDIAN tSNR VALUE

[S1,S2,S3] = size(mask);

if exist([base_dir_sub fname '_slice' num2str(1) suffix '.nii'],'file') || exist([base_dir_sub fname '_slice' num2str(1) suffix '.nii.gz'],'file')
    scfMRItb_04_unzipFile(base_dir_sub, fname, '_mean')
    F = load_untouch_nii([base_dir_sub fname '_mean' '.nii']);
    F_tsnr = F; F_tsnr.img = zeros(S2,S1,S3);
    siz3 = size(F.img,3); clear F
    for i3 = 1 : siz3
        scfMRItb_04_unzipFile(base_dir_sub, fname, ['_slice' num2str(i3) suffix])
        F = load_untouch_nii([base_dir_sub fname '_slice' num2str(i3) suffix '.nii']);
        F = single(squeeze(F.img));
        F = F .* repmat(single(mask(:,:,i3)), [1 1 size(F,3)]);
        F_tsnr.img(:,:,i3) = rot90(mean(F,3) ./ (std(F, [], 3)+1E-20));
        tSNR_slicewise(i3,1) = median(dezero(F_tsnr.img(:,:,i3)));
        clear F;
    end

elseif exist([base_dir_sub fname suffix '.nii'],'file') || exist([base_dir_sub fname suffix '.nii.gz'],'file')
    scfMRItb_04_unzipFile(base_dir_sub, fname, suffix)
    F = load_untouch_nii([base_dir_sub fname suffix '.nii']);
    F.hdr.dime.dim(1) = 3; F.hdr.dime.dim(5) = 1;
    F_tsnr = F; F_tsnr.img = zeros(S2,S1,S3);
    siz3 = size(F.img,3);
    for i3 = 1 : siz3
        F1 = single(squeeze(F.img(:,:,i3,:)));
        F1 = F1 .* repmat(single(mask(:,:,i3)), [1 1 size(F1,3)]);
        F_tsnr.img(:,:,i3) = rot90(mean(F1,3) ./ (std(F1, [], 3)+1E-20));
        tSNR_slicewise(i3,1) = median(dezero(F_tsnr.img(:,:,i3)));
        clear F1;
    end
end

tSNR = median(tSNR_slicewise);

end
```
