## Supplementary material for "Neptune: a toolbox for spinal cord functional MRI data processing and quality assurance": Neptune software: Dicom_Protocol_Rename_noip.m

```
% Dicom_Protocol_Rename_noip.m
%
% Utility program to open user-selected Seimens-format DICOM MRI image files and
% rename them by their individual "Protocol Name Strings" and copying them
% to a user-selectable directory.
%
% Original by Dr. Pat Helm
% Significantly modified by Ron Beyers
% Last update: Oct 29, 2009
% 
% Modified to *_noip by Ranga Deshpande @MGH (started modifications on
% 11-Mar-2020 and ended on 15-Sep-2021).
%
function Dicom_Protocol_Rename_noip(varargin)
%
if nargin==2
    directory = varargin{1};
    out_directory = varargin{2};
end
if nargin==1
    directory = varargin{1};
    if ispc
        out_directory = [directory '\' 'RenamedFiles' '\'];
    elseif isunix
        out_directory = [directory '/' 'RenamedFiles' '/'];
    end
end
if nargin==0
    directory = pwd;
    if ispc
        out_directory = [directory '\' 'RenamedFiles' '\'];
    elseif isunix
        out_directory = [directory '/' 'RenamedFiles' '/'];
    end
end


Start_Dir = directory;
% Proceed   = 1;
%
% while Proceed,
    fc = 0; % Count of valid DICOM files found
    fp = 1; % New Directory first pass flag
    %
%     if ~exist( 'File_Search_Path', 'var' ), % Initialize search path to Root Directory
%         File_Search_Path = 'C:\';
%     end
%     %
%     CmdTxt = 'Select desired DICOM Image Directory for File Renaming: ';
%     fprintf( '\n%s\n', CmdTxt );
%     %
%     Selected_Directory_Name = uigetdir( File_Search_Path, CmdTxt );
    Selected_Directory_Name = directory;
    %
        if ispc
            File_Search_Path  = sprintf( '%s\\', Selected_Directory_Name ); % Append a slash '\' on the end of Name to make it a Path
        elseif isunix
            File_Search_Path  = sprintf( '%s/', Selected_Directory_Name ); % Append a slash '/' on the end of Name to make it a Path
        end
        fprintf( 'DICOM Image Source Directory set as: %s\n', File_Search_Path );
        %
        cd( File_Search_Path );
        File_Search_Dir_Struct = dir;
        File_Search_Dir_Struct = File_Search_Dir_Struct(3:end);
        %
%         CmdTxt = 'Select REQUIRED Dicom file search suffix: ';
%         fprintf( '\n%s\n', CmdTxt );
%         Button_Names_Cell_Array = { 'None', '.IMA', '.DCM' };
%         %
%         Suffix_Select = menu( CmdTxt, Button_Names_Cell_Array );
%         %
%         if Suffix_Select == 0 || Suffix_Select == 1,
%             fprintf( 'Suffix "None" selected.\n' );
%             Suffix_String = '';
%         elseif Suffix_Select == 2,
%             fprintf( 'Suffix ".IMA" selected.\n' );
%             Suffix_String = '.ima';
%         elseif Suffix_Select == 3,
%             fprintf( 'Suffix ".DCM" selected.\n' );
%             Suffix_String = '.dcm';
%         end
        %
        New_Dicom_Filename_CellArray   = {}; % Initialize an empty array of cells for the new filenames
        New_Dicom_Filename_Version_vec = 0;  % Initialize a zero vector of filename version number
        %
        for k = 1:length( File_Search_Dir_Struct )
            skiploop=0;
            fprintf('Converting file (%d) of (%d)\n',k,length( File_Search_Dir_Struct ))
            Dir_Filename = lower( File_Search_Dir_Struct( k ).name );
            %
            if File_Search_Dir_Struct( k ).isdir == 0 && length( Dir_Filename ) > 2
%                 if isempty( Suffix_String ) || strcmp( Dir_Filename( end-3:end ), Suffix_String ),
% %                 if ( strcmp( Dir_Filename( end-3:end ), '.IMA' ) ||...
% %                      strcmp( Dir_Filename( end-3:end ), '.ima' ) ), % Found a valid DICOM filename
                    %
                    if fp % On the first found Dicom file, create the RenamedFiles directory
                        [ success, mess, messid ] = mkdir( out_directory );
                        if ~success
                            fprintf( 'WARNING: Failed to create output directory: %s, %s\n', mess, messid );
                            % k = inf;
                            break ; % Break/Terminate the For k = loop
                        elseif success && ( ~isempty( mess ) || ~isempty( messid ) )
                            fprintf( 'NOTICE: Regarding output directory: %s, %s\n', mess, messid );
                        elseif success
                            fprintf( 'RenamedFiles directory created.\n' );
                        end
                        fp = 0;
                    end         
                    %
                    fc = fc + 1;
                    fprintf( 'Found Dicomfile #%2d: %s,  ', fc, Dir_Filename );
                    try
                        Input_Dicom_MetaData = dicominfo( Dir_Filename );
                    catch
                        skiploop=1;
                    end
                    
                    if skiploop~=1
                    [ Input_Dicom_Image, Input_Dicom_Map ] = dicomread( Dir_Filename );
                    %
%                     Dicom_File_Search_Orig_Path = pwd;
%                     cd( 'RenamedFiles' );
                    %
                    if isfield( Input_Dicom_MetaData, 'ProtocolName' ) && isfield( Input_Dicom_MetaData, 'InstanceNumber' ),
                        New_Dicom_Filename = sprintf( '%s-%d', Input_Dicom_MetaData.ProtocolName, Input_Dicom_MetaData.InstanceNumber );
                        % New_Dicom_Filename = sprintf( '%s-%d.dcm', Input_Dicom_MetaData.ProtocolName, Input_Dicom_MetaData.InstanceNumber );
                    else
                        fprintf( 'NOTICE: No Rename Info,  ' );
                        New_Dicom_Filename = Dir_Filename;                
                        % New_Dicom_Filename = sprintf( '%s.dcm', Dir_Filename );
                    end
                    %
                    New_Dicom_Filename( New_Dicom_Filename == ' ' ) = '-';  % Find any "blank" spaces and Replace with a "-" (dash)
                    %
                    if ~isempty( New_Dicom_Filename_CellArray ) % At least one previous New Filename exists
                        Filename_Cell_Match_Index = strmatch( New_Dicom_Filename, New_Dicom_Filename_CellArray, 'exact' ); % Test for and find any previous Filename match
                        if ~isempty( Filename_Cell_Match_Index ) % Previous Filename found
                            New_Dicom_Filename_Version_vec( Filename_Cell_Match_Index ) = New_Dicom_Filename_Version_vec( Filename_Cell_Match_Index ) + 1; % Increment the version number
                            New_Dicom_Filename = sprintf( '%s-%d', New_Dicom_Filename, New_Dicom_Filename_Version_vec( Filename_Cell_Match_Index ) );            % Append the new version number
                        else % Not previously used, so append to end of CellArray
                            New_Dicom_Filename_CellArray( end + 1 )   = { New_Dicom_Filename };                % Append unfound New Filename to Cells
                            New_Dicom_Filename_Version_vec( end + 1 ) = 1;                                     % Set New Filename Version number to "1" (one)
                            New_Dicom_Filename                        = sprintf( '%s-1', New_Dicom_Filename ); % Append a "1" (one) as the initial version number
                        end
                    else
                        New_Dicom_Filename_CellArray(1)   = { New_Dicom_Filename };                % Store in first cell
                        New_Dicom_Filename_Version_vec(1) = 1;                                     % Set New Filename Version number to "1" (one)
                        New_Dicom_Filename                = sprintf( '%s-1', New_Dicom_Filename ); % Append a "1" (one) as the initial version number
                    end
                    %
                    New_Dicom_Filename = sprintf( '%s.dcm', New_Dicom_Filename ); % Append .dcm to end of the New Filename
                    fprintf( 'Renamed to: %s\n', New_Dicom_Filename );
                    %
                    New_Dicom_MetaData           = Input_Dicom_MetaData;
                    New_Dicom_MetaData.Filename  = New_Dicom_Filename;
                    if ispc
                        New_Dicom_Folder = [out_directory Input_Dicom_MetaData.ProtocolName '\'];
                        if ~exist(New_Dicom_Folder,'dir')
                            mkdir(New_Dicom_Folder);
                        end
                        Copy_Cmnd_String = sprintf( 'copy "%s" "%s%s" /V /Y', Dir_Filename, New_Dicom_Folder, New_Dicom_Filename );
                    elseif isunix
                        New_Dicom_Folder = [out_directory Input_Dicom_MetaData.ProtocolName '/'];
                        if ~exist(New_Dicom_Folder,'dir')
                            mkdir(New_Dicom_Folder);
                        end
                        Copy_Cmnd_String = sprintf( 'cp %s %s%s', Dir_Filename, New_Dicom_Folder, New_Dicom_Filename );
                    end
                    [ Copy_Error, ErrorMessage ] = system( Copy_Cmnd_String );
                    if Copy_Error
                        fprintf( 'ERROR: Copy Command Error: %s for Filenames, In: %s and Out: %s%s\n',...
                                 ErrorMessage, Dir_Filename, out_directory, New_Dicom_Filename );
                    end
%                     Dicom_Write_Status = dicomwrite( Input_Dicom_Image, Input_Dicom_Map, New_Dicom_Filename, New_Dicom_MetaData,...
%                                                      'Endian', 'ieee-le', 'VR', 'explicit', 'CreateMode', 'Copy',...
%                                                      'CompressionMode', 'None', 'ObjectType', 'MR Image Storage' );
%                     Dicom_Write_Status = dicomwrite( Input_Dicom_Image, New_Dicom_Filename, Input_Dicom_MetaData );
%                     %
%                     if ~isempty( Dicom_Write_Status ),
%                         Status_FieldList_cell = fieldnames( Dicom_Write_Status );
%                         for cn = 1:length( Status_FieldList_cell ),
%                             Error_Struct_string = sprintf( 'Dicom_Write_Status.%s', Status_FieldList_cell{ cn } );
%                             if ~isempty( eval( Error_Struct_string ) ),
%                                 fprintf( 'WARNING: DICOM Write Status message: %s = %s for Filename: %s\n',...
%                                          Error_Struct_string, eval( sprintf( '%s{:}', Error_Struct_string ) ), New_Dicom_Filename );
%                             end
%                         end
%                     end
%                     %
%                     cd( Dicom_File_Search_Orig_Path );
%                 end
                    end
            end
        end
        %
        fprintf( 'Found a total of %d Dicom Files to Rename.\n', fc );

%
%     CmdTxt = 'Select to Rename another Directory or End Program: ';
%     fprintf( '\n%s\n', CmdTxt );
%     Button_Names_Cell_Array = { 'Rename another Directory', 'End Program' };
%     %
%     Proceed = menu( CmdTxt, Button_Names_Cell_Array );
%     %
%     if Proceed == 1,
%         fprintf( 'Rename another Directory selected.\n' );
%     else
%         fprintf( 'End Program selected.\n' );
%         Proceed = 0; % Set flag to exit the While loop
%     end
% end % End of While loop
%
cd( Start_Dir ); % Return to the original program start directory
fprintf( 'Dicom Protocol Rename Program completed.\n' );
%
% clear( 'CmdTxt', 'Dir_Filename', 'File_Search_Path',...
%        'Input_Dicom_Image', 'Input_Dicom_MetaData', 'Jb', 'New_Dicom_Filename', 'File_Search_Dir_Struct', 'Proceed',...
%        'Dicom_File_Search_Orig_Path', 'Selected_Directory_Name', 'Start_Dir',...
%        'k', 'fc', 'Filename_Cell_Match_Index', 'New_Dicom_Filename_CellArray',...
%        'New_Dicom_Filename_Version_vec', 'fp', 'mess', 'messid', 'success' );
%
% End of Dicom_Protocol_Rename.m code
```
