## Supplementary material for "Neptune: a toolbox for spinal cord functional MRI data processing and quality assurance": Neptune software: exec_deconv.m

```
function [op, HRF, adjust_global, PARA]=exec_deconv(ip,TR)

% ip is the input time series of dimensions (time points X ROIs)
% TR is in seconds

% A blind deconvolution approach to recover effective connectivity brain networks from resting state fMRI data
% https://pubmed.ncbi.nlm.nih.gov/23422254/

Threshold = 1; % default value taken from the paper
event_lag_max = round(8/TR); % default value taken from the paper

HRF = zeros(round(30/TR),size(ip,2));
adjust_global = zeros(1,size(ip,2));
PARA = zeros(3,size(ip,2)); % the three parameters are (column-1): RH, TTP, FWHM

y = 0;
na = zscore(ip);
z1 = 1;
for i=1:size(ip,2)
    if na(:,i)~=0
        y=1;
        subip(:,z1) = na(:,i);
        nzero1(z1,1) = i;
        z1=z1+1;
    end
end
clear i z1

if y==1 % do deconvolution if time series is non-zero (not flat)
    [data_deconv, ~, HRF1, adjust_global1, PARA1] = wgr_deconv_canonhrf_par(subip,Threshold,event_lag_max,TR);
else
    op = zeros(size(ip,1),size(ip,2));
    return
end

op = zeros(size(ip,1),size(ip,2));
for i=1:size(subip,2)
    op(:,nzero1(i)) = data_deconv(:,i);
    HRF(:,nzero1(i)) = HRF1(:,i);
    adjust_global(nzero1(i)) = adjust_global1(i);
    PARA(:,nzero1(i)) = PARA1(:,i);
end; clear i y

end
```
