## Supplementary material for "Neptune: a toolbox for spinal cord functional MRI data processing and quality assurance": Neptune software: falff2.m

```
function [fALFF] = falff2(data, TR, HighCutoff, LowCutoff, varargin)

% Use ALFF method to compute the brain and return a ALFF brain map which reflects the "energy" of the voxels' BOLD signal
% Ref: Zang, Y.F., He, Y., Zhu, C.Z., Cao, Q.J., Sui, M.Q., Liang, M., Tian, L.X., Jiang, T.Z., Wang, Y.F., 2007. Altered baseline brain activity in children with ADHD revealed by resting-state functional MRI. Brain Dev 29, 83-91.
% 
% Input:
% 	data            -	Nxixjxk data as in "time series x 3 spatial	dimensions"
% 	TR              -   repetition time
% 	LowCutoff		-	the low edge of the pass band
% 	HighCutoff		-	the High edge of the pass band
%   mask (optional) -   gray matter whole-brain mask
% 
% Output:
%	ALFFBrain       -   The ALFF results
% 
%-----------------------------------------------------------
%   Algorithm originally Written by Xiao-Wei Song.
%   Algorithm Re-Written by YAN Chao-Gan on 120328.
%   Note: the ALFF generated by the new version is sqrt(2/N) times of the
%   original version. (new version used: 2*abs(fft(x))/N; original version
%   used:  sqrt(2*abs(fft(x))^2/N)) 
%	http://restfmri.net 
% 
% Code modified @Northwestern by Ranga Deshpande (7-May-2019)
% Code modified @MGH by Ranga Deshpande (16-Mar-2020)
% 

if nargin>=5
    mask = varargin{1};
else
    mask = ones(size(data,2),size(data,3),size(data,4));
end
if nargin>=6 % used to calculate percentage complete
    s1 = varargin{2};
    ss1 = varargin{3};
    k1 = varargin{4};
end

% Get the frequency index
sampleFreq 	 = 1/TR;
sampleLength = size(data,1);
paddedLength = 2^nextpow2(sampleLength); %jiaxize 150808
if (LowCutoff >= sampleFreq/2) % All high included
    idx_LowCutoff = paddedLength/2 + 1;
else % high cut off, such as freq > 0.01 Hz
    idx_LowCutoff = ceil(LowCutoff * paddedLength * TR + 1);
    % Change from round to ceil: idx_LowCutoff = round(LowCutoff *paddedLength *TR + 1);
end
if (HighCutoff>=sampleFreq/2)||(HighCutoff==0) % All low pass
    idx_HighCutoff = paddedLength/2 + 1;
else % Low pass, such as freq < 0.08 Hz
    idx_HighCutoff = fix(HighCutoff *paddedLength *TR + 1);
    % Change from round to fix: idx_HighCutoff	=round(HighCutoff *paddedLength *TR + 1);
end

[~,L1,L2,L3] = size(data);

for j=1:L2
    if nargin>=6, sm = sum(sum(s1(1:j-1,1:size(data,4)))); end
    for k=1:L3
        if nargin>=6 % used to calculate percentage complete
            pcent=100*(sm+sum(s1(j,1:k)))/ss1; %calculation of percentage complete
            fprintf('fALFF subject (%d), voxel (%d,%d) done, percent complete = %.2f\n', k1,j,k,pcent) %display progress
        end
        for i=1:L1
            if mask(i,j,k)~=0
                data1 = data(:,i,j,k);
                % Zero Padding
                data1 = [data1;zeros(paddedLength -sampleLength,size(data1,2))]; %padded with zero
                
                data1 = 2*abs(fft(data1))/sampleLength;
                
                temp = sum(data1(idx_LowCutoff:idx_HighCutoff)) ./ sum(data1(2:(paddedLength/2 + 1)));
                temp(~isfinite(temp))=0;
                fALFF(i,j,k) = temp;
                clear temp data1
            end
        end
    end
end; clear i j k

end
```
