## Supplementary material for "Neptune: a toolbox for spinal cord functional MRI data processing and quality assurance": Neptune software: FD_metric2.m

```
function [FD,mean_FD,AD] = FD_metric2(motion_parameters,radius)
%%%%%%%%%%%%%%%%%%%%%%%%%%%%%%%%%%%%%%%%%%%%%%%%%%%%%%%%%%%%%%%%%%%%%%%%%
% This program is used to calculate the framewise displacement to express
% head motion as a scalar quantity.
%
%%%%%%%%%%%%%%%%%
% Inputs
% motion_parameters : This is an m x 6 matrix with m being the number of
%                     timepoints and a total of six motion parameters. The
%                     first 3 columns being x,y and z translation(in mm) and
%                     the next three columns x,y and z rotaions(in degrees).
% radius            : This is the radius of the sphere. Based on 
%                     Power et al.,2012 the radius value is 50(in mm).
%
% Outputs
% FD      : This is the framewise displacement value over time. This is a
%           column vector of size (m-1) x 1 where m is the number of
%           timepoints.
% mean_FD : This is the mean framewise displacement value over all
%           timepoints.
% AD      : This is the absolute displacement in mm (added by Ranga
%           Deshpande, MGH; March 2021)
%%%%%%%%%%%%%%%%%
% 
%
% Reference : Jonathan D. Power, Kelly A. Barnes, Abraham Z. Snyder,
%             Bradley L. Schlaggar, Steven E. Petersen, Spurious but systematic
%             correlations in functional connectivity MRI networks arise from subject
%             motion, NeuroImage, Volume 59, Issue 3, 1 February 2012, Pages 2142-2154, 
%             ISSN 1053-8119, http://dx.doi.org/10.1016/j.neuroimage.2011.10.018.
%
%%%%%%%%%%%%%%%%%%%%%%%%%%%%%%%%%%%%%%%%%%%%%%%%%%%%%%%%%%%%%%%%%%%%%%%%%

x_rotation_mm=conversion_parameter(motion_parameters(:,4),radius);
y_rotation_mm=conversion_parameter(motion_parameters(:,5),radius);
z_rotation_mm=conversion_parameter(motion_parameters(:,6),radius);

motion_all_in_mm(:,1:3)=motion_parameters(:,1:3);
motion_all_in_mm(:,4)=x_rotation_mm;
motion_all_in_mm(:,5)=y_rotation_mm;
motion_all_in_mm(:,6)=z_rotation_mm;

for j=1:size(motion_all_in_mm,1)
    AD(j,1)=sum(abs(motion_all_in_mm(j,:))); % absolute displacement in mm
end

for j=2:size(motion_all_in_mm,1)
    delta_motion_parameters((j-1),:)=motion_all_in_mm((j-1),:)-motion_all_in_mm(j,:);
end

for j=1:size(delta_motion_parameters)
    FD(j,1)=sum(abs(delta_motion_parameters(j,:))); % relative displacement in mm
end

mean_FD=mean(FD(:,1));  % Taking mean over time to get one value of FD

end

%%%%%%%%%%%%%%%%%%%%%%%%%%%%%%%%%%%%%%%%%%%%%%%%%%%%%%%%%%%%%%%%%%%%%%%%%%
function arc_length=conversion_parameter(degree,rad)

% This function is used to convert the angular displacement from degrees to mm.
%
% input terms
% degree : input parameter for the fuction. This is the vector (size 1*n)
%          of angular displacements in degree
% rad    : radius of the circle (in mm)
% 
% output terms
% arc_length : the output where the angle is converted to the arc lenth
%              with units same as the units of the radius (in mm).

for i=1:size(degree,1)
    radians(i,1)=degree(i,1)*(pi/180);      % converts the angle from degree to radians
    arc_length(i,1)=rad*radians(i,1);    % gives the length of arc (units are same as the units of the radius i.e in mm)
end

end
```
