## Supplementary material for "Neptune: a toolbox for spinal cord functional MRI data processing and quality assurance": Neptune software: figmriF.m

```
function fighndl2 = figmriF(img)

img = squeeze(img);

if size(img,3) > 1
    img = img(:,:,1);
    disp('Only showing first slice of 3D volume.');
end
fighndl2 = imshow(single(abs(img)), [], 'InitialMagnification', 'fit');
colorbar;
```
