## Supplementary material for "Neptune: a toolbox for spinal cord functional MRI data processing and quality assurance": Neptune software: fitT2star.m

```
function params = fitT2star(TEs, Obs)

options = optimset('Display','none');

if size(Obs,1) > size(Obs,2)
    Obs = Obs';
end
%fit = fminsearch(@(x) objfcn(x,TEs,Obs),[1.15*Obs(1,1) 20/1000 Obs(1,end)/2],options);
fit = fminsearch(@(x) objfcn(x,TEs,Obs),[1.15*Obs(1,1) 20/1000],options);
params = fit;

end

function err = objfcn(x,TE,Obs)
%  S = x(1,1) .* (exp(-TE ./ x(1,2))) + x(1,3);
  S = x(1,1) .* (exp(-TE ./ x(1,2)));
  err = sum((S-Obs).*(S-Obs),2);
end
```
