## Supplementary material for "Neptune: a toolbox for spinal cord functional MRI data processing and quality assurance": Neptune software: fitT2star_err.m

```
function [params, sos_err] = fitT2star_err(TEs, Obs)

options = optimset('Display','none');

if size(Obs,1) > size(Obs,2)
    Obs = Obs';
end
params = fminsearch(@(x) objfcn(x,TEs,Obs),[1.15*Obs(1,1) 20/1000],options);

% calculate error in fit (sum of squares)
%sos_err = sum((Obs - (params(1,1) .* exp(-TEs ./ params(1,2)))) .^ 2, 2);

% calculate goodness of fit (chi-squared)
expected = (params(1,1) .* exp(-TEs ./ params(1,2)));
weighting = size(Obs,2):-1:1;
weighting = weighting ./ sum(weighting,2);
sos_err = sum((((Obs - expected) .^ 2) ./ expected) .* weighting, 2);

end

function err = objfcn(x,TE,Obs)
  S = x(1,1) .* (exp(-TE ./ x(1,2)));
  err = sum((S-Obs).*(S-Obs),2);
end
```
