## Supplementary material for "Neptune: a toolbox for spinal cord functional MRI data processing and quality assurance": Neptune software: gauss2D_R.m

```
function I = gauss2D_R(X, Y, dx, dy, theta, order)
%GAUSS2: Calculates a rotated 2D gaussian (cylindrical)
%
%	Usage:
%		y = gauss2D_R(X, Y, FWHM_x, FWHM_y, theta, order);
%
%		X		= Matrix of x indices
%		Y		= Matrix of y indices
%		FWHM_x	= FWHM in x dimension
%		FWHM_y	= FWHM in y dimension
%		theta	= Rotation angle in degress (+ve = anticlockwise)
%		order	= Gaussian order
%
%	Set:
%		X = ones(Ny, 1) * ((0:Nx-1)*dx - x0)
%		Y = ((0:Ny-1)*dy - y0) * ones(1, Nx);
%
%	v1.0

order = abs(floor(order));
theta = theta*pi/180;

X_ = cos(theta)*X - sin(theta)*Y;
Y_ = sin(theta)*X + cos(theta)*Y;
R_ = (X_/dx).^2 + (Y_/dy).^2;
I = exp(-log(2)*(2^(2*order))*(R_.^order));
```
