## Supplementary material for "Neptune: a toolbox for spinal cord functional MRI data processing and quality assurance": Neptune software: gen_tsnr_maps.m

```
function gen_tsnr_maps(F_tsnr, savename, figtitle, climit, tSNR)

%% GENERATE tSNR MAPS

if isstruct(F_tsnr), F_tsnr = F_tsnr.img; end

siz3 = size(F_tsnr,3);
for i3 = 1 : siz3
    a2 = F_tsnr(:,:,i3);
    [r, c] = find(a2 > 0);
    space_above = min(r) - 1;
    space_below = size(a2,1) - max(r);
    space_on_left = min(c) - 1;
    space_on_right = size(a2,2) - max(c);
    F_tsnr(:,:,i3) = circshift(a2, [(floor((space_below-space_above)/2)) (floor((space_on_right-space_on_left)/2))]);
    clear a2 r c space_*;
end

a2 = sum(F_tsnr,3);
[r, c] = find(a2 > 0);
a1_tsnr = F_tsnr(min(r):max(r),min(c):max(c),:);
a1_tsnr_flipped = a1_tsnr(:,:,fliplr(1:1:size(F_tsnr,3)));
a1_tsnr_flipped = a1_tsnr_flipped(:,:,end:-1:1);

fighndl = figure('Position',[1 1 960 600],'Color',[0.85,0.90,0.85],'InvertHardcopy','off','Visible','off'); % red [0.95,0.9,0.85], green [0.85,0.90,0.85], blue [0.89,0.94,0.99]
figmriF(mosaic(a1_tsnr_flipped)); colormap(jet), caxis([0 climit])
title(sprintf('\n%s\nMedian tSNR = %.2f\n',figtitle,tSNR),'Color',[0,0.55,0.69],'FontSize',16)
saveas(fighndl,savename)
close(fighndl)

end
```
