## Supplementary material for "Neptune: a toolbox for spinal cord functional MRI data processing and quality assurance": Neptune software: ICC.m

```
function out = ICC(cse,typ,dat)
%function to work out ICCs according to shrout & fleiss' schema (Shrout PE,
%Fleiss JL. Intraclass correlations: uses in assessing rater reliability.
%Psychol Bull. 1979;86:420-428).
%
% 'dat' is data whose rows represent different ratings or raters & whose
% columns represent different cases or targets being measured. Each target
% is assumed to be a random sample from a population of targets.
%
% 'cse' is either 1,2,3. 'cse' is: 1 if each target is measured by a
% different set of raters from a population of raters, 2 if each target is
% measured by the same raters, but that these raters are sampled from a
% population of raters, 3 if each target is measured by the same raters and
% these raters are the only raters of interest.
%
% 'typ' is either 'single' or 'k' & denotes whether the ICC is based on a
% single measurement or on an average of k measurements, where k = the
% number of ratings/raters.
%
% This has been tested using the example data in the paper by shrout & fleiss.
% 
% Example: out = ICC(3,'k',S_Fdata)
% returns ICC(3,k) of data 'S_Fdata' to double 'out'.
%
% Kevin Brownhill, Imaging Sciences, KCL, London
%~~~~~~~~~~~~~~~~~~~~~~~~~~~~~~~~~~~~~~~~~~~~~~~~~~~~~~~~~~~~~~~~~~~~~~~~~~

%number of raters/ratings
k = size(dat,2);
%number of targets
n = size(dat,1);
%mean per target
mpt = mean(dat,2);
%mean per rater/rating
mpr = mean(dat);
%get total mean
tm = mean(mpt);
%within target sum sqrs
WSS = sum(sum(bsxfun(@minus,dat,mpt).^2));
%within target mean sqrs
WMS = WSS / (n * (k - 1));
%between rater sum sqrs
RSS = sum((mpr - tm).^2) * n;
%between rater mean sqrs
RMS = RSS / (k - 1);
% %get total sum sqrs
% TSS = sum(sum((dat - tm).^2));
%between target sum sqrs
BSS = sum((mpt - tm).^2) * k;
%between targets mean squares
BMS = BSS / (n - 1);
%residual sum of squares
ESS = WSS - RSS;
%residual mean sqrs
EMS = ESS / ((k - 1) * (n - 1));
switch cse
    case 1
        switch typ
            case 'single'
                out = (BMS - WMS) / (BMS + (k - 1) * WMS);
            case 'k'
                out = (BMS - WMS) / BMS;
            otherwise
               error('Wrong value for input typ') 
        end
    case 2
        switch typ
            case 'single'
                out = (BMS - EMS) / (BMS + (k - 1) * EMS + k * (RMS - EMS) / n);
            case 'k'
                out = (BMS - EMS) / (BMS + (RMS - EMS) / n);
            otherwise
               error('Wrong value for input typ') 
        end
    case 3
        switch typ
            case 'single'
                out = (BMS - EMS) / (BMS + (k - 1) * EMS);
            case 'k'
                out = (BMS - EMS) / BMS;
            otherwise
               error('Wrong value for input typ') 
        end
    otherwise
        error('Wrong value for input cse')
end
```
