## Supplementary material for "Neptune: a toolbox for spinal cord functional MRI data processing and quality assurance": Neptune software: ICC_RLB.m

```
function [r, LB, UB, F, df1, df2, p] = ICC_RLB(M1, M2)

% 2015-03-25 Modified ICC - so it works for bootstrap!
% 
% Intraclass correlation
%   [r, LB, UB, F, df1, df2, p] = ICC(M, type, alpha, r0)
%
%   M is matrix of observations. Each row is an object of measurement and
%   each column is a judge or measurement.
%
%   'type' is a string that can be one of the six possible codes for the desired
%   type of ICC:
%       '1-1': The degree of absolute agreement among measurements made on
%         randomly seleted objects. It estimates the correlation of any two
%         measurements.
%       '1-k': The degree of absolute agreement of measurements that are
%         averages of k independent measurements on randomly selected
%         objects.
%       'C-1': case 2: The degree of consistency among measurements. Also known
%         as norm-referenced reliability and as Winer's adjustment for
%         anchor points. case 3: The degree of consistency among measurements maded under
%         the fixed levels of the column factor. This ICC estimates the
%         corrlation of any two measurements, but when interaction is
%         present, it underestimates reliability.
%       'C-k': case 2: The degree of consistency for measurements that are
%         averages of k independent measurements on randomly selected
%         onbjectgs. Known as Cronbach's alpha in psychometrics. case 3:  
%         The degree of consistency for averages of k independent
%         measures made under the fixed levels of column factor.
%       'A-1': case 2: The degree of absolute agreement among measurements. Also
%         known as criterion-referenced reliability. case 3: The absolute 
%         agreement of measurements made under the fixed levels of the column factor.
%       'A-k': case 2: The degree of absolute agreement for measurements that are
%         averages of k independent measurements on randomly selected objects.
%         case 3: he degree of absolute agreement for measurements that are
%         based on k independent measurements maded under the fixed levels
%         of the column factor.
%
%       ICC is the estimated intraclass correlation. LB and UB are upper
%       and lower bounds of the ICC with alpha level of significance. 
%
%       In addition to estimation of ICC, a hypothesis test is performed
%       with the null hypothesis that ICC = r0. The F value, degrees of
%       freedom and the corresponding p-value of the this test are
%       reported.
%
%       (c) Arash Salarian, 2008
%
%       Reference: McGraw, K. O., Wong, S. P., "Forming Inferences About
%       Some Intraclass Correlation Coefficients", Psychological Methods,
%       Vol. 1, No. 1, pp. 30-46, 1996
%

M = cat(2,M1,M2); % this is what the function needs

type = 'C-1';
alpha = .05;
r0 = 0;

[n, k] = size(M);
[p, table] = anova_rm(M, 'off');

SSR = table{3,2};
SSE = table{4,2};
SSC = table{2,2};
SSW = SSE + SSC;

MSR = SSR / (n-1);
MSE = SSE / ((n-1)*(k-1));
MSC = SSC / (k-1);
MSW = SSW / (n*(k-1));

switch type
    case '1-1'
        [r, LB, UB, F, df1, df2, p] = ICC_case_1_1(MSR, MSE, MSC, MSW, alpha, r0, n, k);
    case '1-k'
        [r, LB, UB, F, df1, df2, p] = ICC_case_1_k(MSR, MSE, MSC, MSW, alpha, r0, n, k);
    case 'C-1'
        [r, LB, UB, F, df1, df2, p] = ICC_case_C_1(MSR, MSE, MSC, MSW, alpha, r0, n, k);
    case 'C-k'
        [r, LB, UB, F, df1, df2, p] = ICC_case_C_k(MSR, MSE, MSC, MSW, alpha, r0, n, k);
    case 'A-1'
        [r, LB, UB, F, df1, df2, p] = ICC_case_A_1(MSR, MSE, MSC, MSW, alpha, r0, n, k);
    case 'A-k'
        [r, LB, UB, F, df1, df2, p] = ICC_case_A_k(MSR, MSE, MSC, MSW, alpha, r0, n, k);
end


%----------------------------------------
function [r, LB, UB, F, df1, df2, p] = ICC_case_1_1(MSR, MSE, MSC, MSW, alpha, r0, n, k)
r = (MSR - MSW) / (MSR + (k-1)*MSW);

F = (MSR/MSW) * (1-r0)/(1+(k-1)*r0);
df1 = n-1;
df2 = n*(k-1);
p = 1-fcdf(F, df1, df2);

FL = F / finv(1-alpha/2, n-1, n*(k-1));
FU = F * finv(1-alpha/2, n*(k-1), n-1);

LB = (FL - 1) / (FL + (k-1));
UB = (FU - 1) / (FU + (k-1));

%----------------------------------------
function [r, LB, UB, F, df1, df2, p] = ICC_case_1_k(MSR, MSE, MSC, MSW, alpha, r0, n, k)
r = (MSR - MSW) / MSR;

F = (MSR/MSW) * (1-r0);
df1 = n-1;
df2 = n*(k-1);
p = 1-fcdf(F, df1, df2);

FL = F / finv(1-alpha/2, n-1, n*(k-1));
FU = F * finv(1-alpha/2, n*(k-1), n-1);

LB = 1 - 1 / FL;
UB = 1 - 1 / FU;

%----------------------------------------
function [r, LB, UB, F, df1, df2, p] = ICC_case_C_1(MSR, MSE, MSC, MSW, alpha, r0, n, k)
r = (MSR - MSE) / (MSR + (k-1)*MSE);

F = (MSR/MSE) * (1-r0)/(1+(k-1)*r0);
df1 = n - 1;
df2 = (n-1)*(k-1);
p = 1-fcdf(F, df1, df2);

FL = F / finv(1-alpha/2, n-1, (n-1)*(k-1));
FU = F * finv(1-alpha/2, (n-1)*(k-1), n-1);

LB = (FL - 1) / (FL + (k-1));
UB = (FU - 1) / (FU + (k-1));

%----------------------------------------
function [r, LB, UB, F, df1, df2, p] = ICC_case_C_k(MSR, MSE, MSC, MSW, alpha, r0, n, k)
r = (MSR - MSE) / MSR;

F = (MSR/MSE) * (1-r0);
df1 = n - 1;
df2 = (n-1)*(k-1);
p = 1-fcdf(F, df1, df2);

FL = F / finv(1-alpha/2, n-1, (n-1)*(k-1));
FU = F * finv(1-alpha/2, (n-1)*(k-1), n-1);

LB = 1 - 1 / FL;
UB = 1 - 1 / FU;

%----------------------------------------
function [r, LB, UB, F, df1, df2, p] = ICC_case_A_1(MSR, MSE, MSC, MSW, alpha, r0, n, k)
r = (MSR - MSE) / (MSR + (k-1)*MSE + k*(MSC-MSE)/n);

a = (k*r0) / (n*(1-r0));
b = 1 + (k*r0*(n-1))/(n*(1-r0));
F = MSR / (a*MSC + b*MSE);
df1 = n - 1;
df2 = (a*MSC + b*MSE)^2/((a*MSC)^2/(k-1) + (b*MSE)^2/((n-1)*(k-1)));
p = 1-fcdf(F, df1, df2);

a = k*r/(n*(1-r));
b = 1+k*r*(n-1)/(n*(1-r));
v = (a*MSC + b*MSE)^2/((a*MSC)^2/(k-1) + (b*MSE)^2/((n-1)*(k-1)));

Fs = finv(1-alpha/2, n-1, v);
LB = n*(MSR - Fs*MSE)/(Fs*(k*MSC + (k*n - k - n)*MSE) + n*MSR);

Fs = finv(1-alpha/2, v, n-1);
UB = n*(Fs*MSR-MSE)/(k*MSC + (k*n - k - n)*MSE + n*Fs*MSR);

%----------------------------------------
function [r, LB, UB, F, df1, df2, p] = ICC_case_A_k(MSR, MSE, MSC, MSW, alpha, r0, n, k)
r = (MSR - MSE) / (MSR + (MSC-MSE)/n);

c = r0/(n*(1-r0));
d = 1 + (r0*(n-1))/(n*(1-r0));
F = MSR / (c*MSC + d*MSE);
df1 = n - 1;
df2 = (c*MSC + d*MSE)^2/((c*MSC)^2/(k-1) + (d*MSE)^2/((n-1)*(k-1)));
p = 1-fcdf(F, df1, df2);

a = r/(n*(1-r));
b = 1+r*(n-1)/(n*(1-r));
v = (a*MSC + b*MSE)^2/((a*MSC)^2/(k-1) + (b*MSE)^2/((n-1)*(k-1)));

Fs = finv(1-alpha/2, n-1, v);
LB = n*(MSR - Fs*MSE)/(Fs*(MSC-MSE) + n*MSR);

Fs = finv(1-alpha/2, v, n-1);
UB = n*(Fs*MSR - MSE)/(MSC - MSE + n*Fs*MSR);
```
