## Supplementary material for "Neptune: a toolbox for spinal cord functional MRI data processing and quality assurance": Neptune software: meanPSC.m

```
function op = meanPSC(ip)

% Input is an ROI x timepoints matrix of raw data
% Output is an ROI x 1 matrix of mean percentage signal change values
% Author: Ranga Deshpande (March 2021)

op = zeros(size(ip,1),1);
for k = 1:size(ip,1)
    temp = ip(k,:)';
    temp = 100*(temp-mean(temp))./mean(temp);
    op(k,1) = mean(abs(temp));
    clear temp
end
```
