## Supplementary material for "Neptune: a toolbox for spinal cord functional MRI data processing and quality assurance": Neptune software: mosaic.m

```
function M = mosaic (stack)
%%MOSAIC Create a 2-D mosaic of images from a 3-D stack.
%
%  M=MOSAIC(STACK) takes a 3-D STACK of images and attempts to stitch them
%  together in a 2-D image with dimensions as close to square as possible.
%
%  Written by David S. Smith, 3 Apr 2011.
%
%  History
%    v1.0 - Initial revision.

[n1 n2 n3] = size(stack);

if n3 == 1, error('MOSAIC> Input must be 3D.'); end

n = round(sqrt(n3)); % starting point: integer nearest to the square root

% find largest integer less than or equal to sqrt that evenly divides the
% number of images in the stack
for k = n:-1:1
       if ~mod(n3,k), break; end
end

% figure out the best dimensions
j = n3 / k;
ny = min(j,k);
nx = max(j,k);

% stick them together
if isa(stack,'logical')
 M = false(ny*n1,nx*n2);
else
       M = zeros(ny*n1,nx*n2,class(stack));
end
for k = 1:ny
       for j = 1:nx
   M(1+n1*(k-1):n1*k,1+n2*(j-1):n2*j) = stack(:,:,j+(k-1)*nx);
       end
end
```
