## Supplementary material for "Neptune: a toolbox for spinal cord functional MRI data processing and quality assurance": Neptune software: NIFTI_NORDIC.m

```
function  NIFTI_NORDIC(fn_magn_in,fn_phase_in,fn_out,ARG)
% fMRI
%  fn_magn_in='name.nii.gz';
%  fn_phase_in='name2.nii.gz';
%  fn_out=['NORDIC_' fn_magn_in(1:end-7)];
%  ARG.temporal_phase=1;
%  ARG.phase_filter_width=10;
%  NIFTI_NORDIC(fn_magn_in,fn_phase_in,fn_out,ARG)
%
% dMRI
%  fn_magn_in='name.nii.gz';
%  fn_phase_in='name2.nii.gz';
%  fn_out=['NORDIC_' fn_magn_in(1:end-7)];
%  ARG.temporal_phase=3;
%  ARG.phase_filter_width=3;
%  NIFTI_NORDIC(fn_magn_in,fn_phase_in,fn_out,ARG)
%
%
%  file_input assumes 4D data
%
%OPTIONS
%   ARG.DIROUT    VAL=      string        Default is empty
%   ARG.noise_volume_last   VAL  = num  specifiec volume from the end of the series
%                                          0 default
%
%   ARG.factor_error        val  = num    >1 use higher noisefloor <1 use lower noisefloor 
%                                          1 default 
%
%   ARG.full_dynamic_range  val = [ 0 1]   0 keep the input scale, output maximizes range. 
%                                            Default 0
%   ARG.temporal_phase      val = [1 2 3]  1 was default, 3 now in dMRI due tophase errors in some data
%   ARG.NORDIC              val = [0 1]    1 Default
%   ARG.MP                  val = [0 1 2]  1 NORDIC gfactor with MP estimation. 
%                                          2 MP without gfactor correction
%                                          0 default
%   ARG.kernel_size_gfactor val = [val1 val2 val], defautl is [14 14 1]
%   ARG.kernel_size_PCA     val = [val1 val2 val], default is val1=val2=val3; 
%                                                  ratio of 11:1 between spatial and temproal voxels
%   ARG.magnitude_only      val =[] or 1.  Using complex or magntiude only. Default is []
%                                          Function still needs two inputs but will ignore the second
%
%   ARG.save_add_info       val =[0 1];  If it is 1, then an additonal matlab file is being saved with degress removed etc.
%                                         default is 0
%   ARG.make_complex_nii    if the field exist, then the phase is being saved in a similar format as the input phase
%
%   ARG.phase_slice_average_for_kspace_centering     val = [0 1]   
%                                         if val =0, not used, if val=1 the series average pr slice is first removed
%                                         default is now 0
%   ARG.phase_filter_width  val = [1... 10]  Specifiec the width of the smoothing filter for the phase
%                                         default is now 3
%   
%   ARG.save_gfactor_map   val = [1 2].  1, saves the RELATIVE gfactor, 2 saves the
%                                            gfactor and does not complete the NORDIC processing


%  TODO
%  Scaling relative to the width of the MP spectrum, if one wants to be
%  conservative
%  
%  4/15/21 swapped the uint16 and in16 for the phase
%
%  VERSION 4/22/2021


if ~exist('ARG')  % initialize ARG structure
    ARG.DIROUT=[pwd '/'];
elseif ~isfield(ARG,'DIROUT') % Specify where to save data
    ARG.DIROUT=[pwd '/'];
end

if ~isfield(ARG,'noise_volume_last')
    ARG.noise_volume_last=0;  % there is no noise volume   {0 1 2 ...}
end

if ~isfield(ARG,'factor_error')
    ARG.factor_error=1.0;  % error in gfactor estimatetion. >1 use higher noisefloor <1 use lower noisefloor
end

if ~isfield(ARG,'full_dynamic_range')
    ARG.full_dynamic_range=0;  % Format o
end

if ~isfield(ARG,'temporal_phase')
    ARG.temporal_phase=1;  % Correction for slice and time-specific phase
end

if ~isfield(ARG,'NORDIC') & ~isfield(ARG,'MP')
    ARG.NORDIC=1;  %  threshold based on Noise
    ARG.MP=0;  % threshold based on Marchencko-Pastur
elseif ~isfield(ARG,'NORDIC') %  MP selected
    if ARG.MP==1
        ARG.NORDIC=0;
    else
        ARG.NORDIC=1;
    end
    
elseif  ~isfield(ARG,'MP')   %  NORDIC selected
    if ARG.NORDIC==1
        ARG.MP=0;
    else
        ARG.MP=1;
    end
end

if ~isfield(ARG,'phase_filter_width')
    ARG.phase_filter_width=3;  %  default is [14 14 90]
end


if ~isfield(ARG,'NORDIC_patch_overlap')
    ARG.NORDIC_patch_overlap=2;  %  default is [14 14 90]
end

if ~isfield(ARG,'gfactor_patch_overlap')
    ARG.gfactor_patch_overlap=2;  %  default is [14 14 90]
end


if ~isfield(ARG,'kernel_size_gfactor')
    ARG.kernel_size_gfactor=[];  %  default is [14 14 90]
end

if ~isfield(ARG,'kernel_size_PCA')
    ARG.kernel_size_PCA=[]; % default is 11:1 ratio
end

if ~isfield(ARG,'phase_slice_average_for_kspace_centering');
ARG.phase_slice_average_for_kspace_centering=0;
end

if ~isfield(ARG,'magnitude_only') % if legacy data
    ARG.magnitude_only=0; %
end

if isfield(ARG,'save_add_info'); end   %  additional information is saved in matlab file
if isfield(ARG,'make_complex_nii'); end   %  two output NII files are saved

if ~isfield(ARG,'save_gfactor_map') % save out a map of a relative gfactor
    ARG.save_gfactor_map=[]; %
end


if isfield(ARG,'use_generic_NII_read') % save out a map of a relative gfactor
    if ARG.use_generic_NII_read==1
    path(path,'/home/range6-raid1/moeller/matlab/ADD/NIFTI/');
    end
else
    ARG.use_generic_NII_read=0;
end

if ~isfield(ARG,'data_has_zero_elements') %
    ARG.data_has_zero_elements=0; %  % If there are pixels that are constant zero
end


ARG;
%%%%%%%%%%%%%%%%%%%%%%%%%%%%%%%%%%%%%%%%%%%%%%%%%%%%%%%%%%%%%%%%%%%

if ARG.magnitude_only~=1

    try
    info_phase=niftiinfo(fn_phase_in);
    info=niftiinfo(fn_magn_in);
    catch;  disp('The niftiinfo fails at reading the header')  ;end
    
    
    if ARG.use_generic_NII_read~=1
        I_M=abs(single(niftiread(fn_magn_in)));
        I_P=single(niftiread(fn_phase_in));
    else
        try
        tmp=load_nii(fn_magn_in);
        I_M=abs(single(tmp.img));
        tmp=load_nii(fn_phase_in);
        I_P=single(tmp.img);
        catch
           disp('Missing nfiti tool. Serach mathworks for load_nii  fileexchange 8797') 
        end
        
    end
    
    phase_range=single(max(I_P(:)));
    phase_range_min=single(min(I_P(:)));
if ~exist('info_phase')    
    info_phase.Datatype=class(I_P);
    info.Datatype=class(I_M);
end
		% Here, we combine magnitude and phase data into complex form 
        fprintf('Phase should be -pi to pi...\n')
	
	% convert to single and then scale the phase
I_P = single(I_P);
            range_norm=phase_range-phase_range_min;
            range_center=(phase_range+phase_range_min)/range_norm*1/2;
            I_P = (single(I_P)./range_norm -range_center)*2*pi;
            II=single(I_M)  .* exp(1i*I_P);       

	
	if 0
        if strmatch(info_phase.Datatype,'uint16')
            I_P = single(I_P)/phase_range*2*pi;
            II=single(I_M)  .* exp(1i*I_P);
        elseif strmatch(info_phase.Datatype,'int16')
            I_P = (single(I_P)+1-(phase_range+1)/2)/(phase_range+1)*2*pi;
            II=single(I_M)  .* exp(1i*I_P);
        elseif strmatch(info_phase.Datatype,'single')
            phase_range_min=min(I_P(:));
            range_norm=phase_range-phase_range_min;
            range_center=(phase_range+phase_range_min)/range_norm*1/2;
            I_P = (single(I_P)./range_norm -range_center)*2*pi;
            II=single(I_M)  .* exp(1i*I_P);                 
              
        end
	end
		
        fprintf('Phase data range is %.2f to %.2f\n', min(I_P(:)), max(I_P(:)))
else
    
     try
     info=niftiinfo(fn_magn_in);
    catch;  disp('The niftiinfo fails at reading the header')  ;end
 
    
    if ARG.use_generic_NII_read~=1
        I_M=abs(single(niftiread(fn_magn_in)));
    else
        tmp=load_nii(fn_magn_in);
        I_M=abs(single(tmp.img));
    end
    
    
if ~exist('info_phase')    
     info.Datatype=class(I_M);
end

end 


if ~isempty(ARG.magnitude_only)
    if ARG.magnitude_only==1
        II=single(I_M);
        ARG.temporal_phase=0;
    end
end


TEMPVOL=abs(II(:,:,:,1));
ARG.ABSOLUTE_SCALE=min(TEMPVOL(TEMPVOL~=0));
II=II./ARG.ABSOLUTE_SCALE;


%   load test_DATA
%  II=II(:,:,:,1:95);

if size(II,4)<6
    disp('Too few volumes')
    % return
end

KSP2=II;
matdim=size(KSP2);

tt=mean(reshape(abs(KSP2),[],size(KSP2,4)));
[idx]=find(tt>0.95*max(tt));
meanphase=mean(KSP2(:,:,:,idx(1)),4);

if 1
    disp('estimating slice-dependent phases ...')
    meanphase=mean(KSP2(:,:,:,[1:end-ARG.noise_volume_last]),4);
    for nsl=1:size(meanphase,3);
        %meanphase2(:,:,nsl)=complex(  medfilt2(squeeze(real(meanphase(:,:,nsl))),[7 7]), medfilt2(squeeze(imag(meanphase(:,:,nsl))),[7 7]) );
    end
    meanphase=meanphase*ARG.phase_slice_average_for_kspace_centering;
end


for slice=matdim(3):-1:1
    for n=1:size(KSP2,4); % include the noise
        KKSP2(:,:,slice,n)=KSP2(:,:,slice,n).*exp(-i*angle(meanphase(:,:,slice)));
    end
end
DD_phase=0*KSP2;

if           ARG.temporal_phase>0; % Standarad low-pass filtered map
    for slice=matdim(3):-1:1
        for n=1:size(KSP2,4);
            tmp=KSP2(:,:,slice,n);
            for ndim=[1:2]; tmp=ifftshift(ifft(ifftshift( tmp ,ndim),[],ndim),ndim+0); end
            [nx, ny, nc, nb] = size(tmp(:,:,:,:,1,1));
            tmp = bsxfun(@times,tmp,reshape(tukeywin(ny,1).^ARG.phase_filter_width,[1 ny]));
            tmp = bsxfun(@times,tmp,reshape(tukeywin(nx,1).^ARG.phase_filter_width,[nx 1]));
            for ndim=[1:2]; tmp=fftshift(fft(fftshift( tmp ,ndim),[],ndim),ndim+0); end
            DD_phase(:,:,slice,n)=tmp;
        end
    end
end


if           ARG.temporal_phase==2; % Secondary step for filtered phase with residual spikes
    for slice=matdim(3):-1:1
        for n=1:size(KSP2,4);
            
            phase_diff=angle(KSP2(:,:,slice,n)./DD_phase(:,:,slice,n));
            mask=abs(phase_diff)>1;
            DD_phase2=DD_phase(:,:,slice,n);
            tmp=(KSP2(:,:,slice,n));
            DD_phase2(mask)= tmp(mask);
            DD_phase(:,:,slice,n)=DD_phase2;
        end
    end
end


for slice=matdim(3):-1:1
    for n=1:size(KSP2,4);
        KSP2(:,:,slice,n)= KSP2(:,:,slice,n).*exp(-i*angle( DD_phase(:,:,slice,n)   ));
    end
end


disp('Completed estimating slice-dependent phases ...')
if isfield(ARG,'use_magn_for_gfactor')
    
    if isempty(ARG.kernel_size_gfactor) | size(ARG.kernel_size_gfactor,2)<3
        KSP2=abs(KSP2(:,:,1:end,1:min(90,end),1));  % should be at least 30 volumes
    else
        KSP2=abs(KSP2(:,:,1:end,1:min(ARG.kernel_size_gfactor(3),end),1));
    end
    
else
    
    if (isempty(ARG.kernel_size_gfactor) | size(ARG.kernel_size_gfactor,2)<3)
        KSP2=(KSP2(:,:,1:end,1:min(90,end),1));  % should be at least 30 volumes
    else
       % KSP2=(KSP2(:,:,1:end,1:min(ARG.kernel_size_gfactor(3),end),1));
        KSP2=(KSP2(:,:,1:end,1:min(ARG.kernel_size_gfactor(4),end),1));        
        
    end
    
end


KSP2(isnan(KSP2))=0;
KSP2(isinf(KSP2))=0;
master_fast=1;
KSP_recon=0*KSP2;

if isempty(ARG.kernel_size_gfactor)
    ARG.kernel_size=[14 14 1];
else
    ARG.kernel_size=[ARG.kernel_size_gfactor(1) ARG.kernel_size_gfactor(2) 1];
    ARG.kernel_size=[ARG.kernel_size_gfactor(1) ARG.kernel_size_gfactor(2) ARG.kernel_size_gfactor(3)];
end

QQ.KSP_processed=zeros(1,size(KSP2,1)-ARG.kernel_size(1));
ARG.patch_average=0;
ARG.patch_average_sub= ARG.gfactor_patch_overlap;

ARG.LLR_scale=0;
ARG.NVR_threshold=1;
ARG.soft_thrs=10;  % MPPCa   (When Noise varies)
%ARG.soft_thrs=[];  % NORDIC  (When noise is flat)
KSP_weight=KSP2(:,:,:,1)*0;
NOISE=KSP_weight;
Component_threshold=KSP_weight;
energy_removed=KSP_weight;
SNR_weight=KSP_weight;
QQ.KSP_processed=zeros(1,size(KSP2,1)-ARG.kernel_size(1));

if ARG.patch_average==0
    KSP_processed=QQ.KSP_processed*0;
    for nw1=2:max(1,floor(ARG.kernel_size(1)/ARG.patch_average_sub));
        KSP_processed(1,nw1 : max(1,floor(ARG.kernel_size(1)/ARG.patch_average_sub)):end)=2;
    end
    KSP_processed(end)=0; % disp
    QQ.KSP_processed=KSP_processed;
end
KSP_processed;

disp('estimating g-factor ...')
QQ.ARG=ARG;
for n1=1:size(QQ.KSP_processed,2)
    % fprintf( [num2str(n1) ' '])
    [KSP_recon,~,KSP_weight,NOISE,Component_threshold,energy_removed,SNR_weight]=sub_LLR_Processing(KSP_recon,KSP2,ARG,n1,QQ,master_fast,KSP_weight,NOISE,Component_threshold,energy_removed,SNR_weight)   ;    % save all files
    % fprintf( [num2str(n1) ' '])
    % if mod(n1,20)==0; disp(' ');end
end
KSP_recon=KSP_recon./repmat((KSP_weight),[1 1 1 size(KSP2,4)]);
ARG.NOISE=  sqrt(NOISE./KSP_weight);
ARG.Component_threshold = Component_threshold./KSP_weight;
ARG.energy_removed = energy_removed./KSP_weight;
ARG.SNR_weight  = SNR_weight./KSP_weight;
IMG2=KSP_recon;
ARG2=ARG;
disp('completed estimating g-factor')
gfactor=ARG.NOISE;

if size(KSP2,4)<6;  % gfactor stimation most likely failed, replace with median estimated
    gfactor(isnan(gfactor))=0;
    gfactor(gfactor==0)=median(gfactor(gfactor~=0));
end


if sum(gfactor(:)==0)>0;  % gfactor stimation most likely failed since it is zero
    gfactor(isnan(gfactor))=0;
    gfactor(gfactor<1)=median(gfactor(gfactor~=0));
    ARG.data_has_zero_elements=1;
end


%%%%%%%%%%%%%%%%%%%%%%%%%%%%%%%%%%%%%%%%%%%%%%%%%%%%%%%%%%%%%%%%%%%%%%%%%%%%%%%
if ARG.MP==2;
    gfactor=ones(size(gfactor));
end

% gfactor=ones(size(gfactor));


if ( ARG.save_gfactor_map==2 )  | ( ARG.save_gfactor_map==1 )
    
    g_IMG=abs(gfactor(:,:,:,1:end)); % remove g-factor and noise for DUAL 1
    g_IMG(isnan(g_IMG))=0;
    tmp=sort(abs(g_IMG(:)));  sn_scale=2*tmp(round(0.99*end));%sn_scale=max();
    gain_level=floor(log2(32000/sn_scale));
    
    if  ARG.full_dynamic_range==0; gain_level=0;end
    
    if strmatch(info.Datatype,'uint16')
        g_IMG= uint16(abs(g_IMG)*2^gain_level);
    elseif strmatch(info.Datatype,'int16')
        g_IMG= int16(abs(g_IMG)*2^gain_level);
    else
        g_IMG= single(abs(g_IMG)*2^gain_level);
    end
     
    niftiwrite((g_IMG),[ARG.DIROUT fn_out(1:end) '_gfactor' '.nii'])
    if ARG.save_gfactor_map==2
     return
    end
    
end


KSP2=II;
matdim=size(KSP2);


for slice=matdim(3):-1:1
    for n=1:size(KSP2,4); % include the noise
        KSP2(:,:,slice,n)=KSP2(:,:,slice,n).*exp(-i*angle(meanphase(:,:,slice)));
    end
end

for n=1:size(KSP2,4);
    KSP2(:,:,:,n)= KSP2(:,:,:,n)./ gfactor;
end

if ARG.noise_volume_last>0
    KSP2_NOISE =KSP2(:,:,:,end+1-ARG.noise_volume_last);
end


if           ARG.temporal_phase==3; % Secondary step for filtered phase with residual spikes
    for slice=matdim(3):-1:1
        for n=1:size(KSP2,4);
            
            phase_diff=angle(KSP2(:,:,slice,n)./DD_phase(:,:,slice,n));
            mask  = abs(phase_diff)>1;
            mask2 = abs(KSP2(:,:,slice,n))>sqrt(2);
            DD_phase2=DD_phase(:,:,slice,n);
            tmp=(KSP2(:,:,slice,n));
            DD_phase2(mask.*mask2==1)= tmp(mask.*mask2==1);
            DD_phase(:,:,slice,n)=DD_phase2;
            
        end
    end
end

for slice=matdim(3):-1:1
    for n=1:size(KSP2,4);
        KSP2(:,:,slice,n)= KSP2(:,:,slice,n).*exp(-i*angle( DD_phase(:,:,slice,n)   ));
    end
end

KSP2(isnan(KSP2))=0;
KSP2(isinf(KSP2))=0;
%%%%%%%%%%%%%%%%%%%%%%%%%%%%%%%%%%%%%%%%%%%%%%%%%%%%%%%%%%%%%%%%%%%%%%%%%%%%%%%
if ARG.noise_volume_last>0
    %tmp_noise=KSP2(:,:,:,end+1-ARG.noise_volume_last);
    tmp_noise=KSP2_NOISE;
    
    tmp_noise(isnan(tmp_noise))=0;
    tmp_noise(isinf(tmp_noise))=0;
    ARG.measured_noise=std(tmp_noise(tmp_noise~=0));  % sqrt(2) for real and complex
else
    ARG.measured_noise=1;  % IF COMPLEX DATA
end


if  ~isfield(ARG,'use_magn_for_gfactor') & (isempty(ARG.magnitude_only) | ARG.magnitude_only==0)  %% WOULD THIS BE THE ISSUE  & replaced by |
    ARG.measured_noise =  ARG.measured_noise/sqrt(2)
end


%%%%%%%%%%%%%%%%%%%%%%%%%%%%%

if    ARG.data_has_zero_elements==1
    MASK=(sum(abs(KSP2),4)==0);
    Num_zero_elements=sum(MASK(:));
    for nvol=1:size(KSP2,4)
        tmp=KSP2(:,:,:,nvol);
        tmp(MASK)=(randn(Num_zero_elements,1)+1i*randn(Num_zero_elements,1))/sqrt(2);
        KSP2(:,:,:,nvol)=tmp;
    end
end


%%%%%%%%%%%%%%%%%%%%%%%%%%%%%
for readout=1:size(KSP2,1)
    TT(readout)= std(reshape(KSP2(readout,:,:,:),[],1));
    TT1(readout)= mean(reshape(KSP2(readout,:,:,:),[],1));
end

% ARG.measured_noise=median(TT([2:8 end-8:end-1]))

%%%%%%%%%%%%%%%%%%%%%%%%%%%%%%%%%%%%%%%%%%%%%%%%%%%%%%%%%%%%%%%%%%%%%%%%%%%%%%%
master_fast=1;
KSP_recon=0*KSP2;
ARG.kernel_size=repmat([ round((size(KSP2,4)*11)^(1/3))   ],1,3);

if  isempty(ARG.kernel_size_PCA)
    ARG.kernel_size=repmat([ round((size(KSP2,4)*11)^(1/3))   ],1,3);
else
    ARG.kernel_size = ARG.kernel_size_PCA ;
end


if matdim(3) <= ARG.kernel_size(3)  % Number of slices is less than cubic kernel    
    ARG.kernel_size   = repmat([ round((size(KSP2,4)*11/matdim(3) )^(1/2))   ],1,2);
    ARG.kernel_size(3)= matdim(3);    
end


QQ.KSP_processed=zeros(1,size(KSP2,1)-ARG.kernel_size(1));
ARG.patch_average=0;
ARG.patch_average_sub= ARG.NORDIC_patch_overlap ;
% ARG.kernel_size=[7 7 7]; ARG.patch_average_sub=7;  MPPCA
% ARG.soft_thrs=10;  % MPPCa   (When Noise varies)
ARG.LLR_scale=1;
ARG.NVR_threshold=0;

ARG.soft_thrs=[];  % NORDIC  (When noise is flat)
%%%%%%%%%%%%%%%%%%%%%%%%%%%%%%%%%%%%%%%%%%%%%%%%%%%%%%%%%%%%%%%%%%%%%%%%%%%%%%%
ARG.NVR_threshold=0;
for ntmp=1:10
    [~,S,~]=svd(randn(prod(ARG.kernel_size),size(KSP2,4)));
    ARG.NVR_threshold=ARG.NVR_threshold+S(1,1);
end

if ARG.magnitude_only~=1  % 4/29/2021
ARG.NVR_threshold= ARG.NVR_threshold/10*sqrt(2)* ARG.measured_noise*ARG.factor_error;  % sqrt(2) due to complex  1.20 due to understimate of g-factor
else
ARG.NVR_threshold= ARG.NVR_threshold/10*ARG.measured_noise*ARG.factor_error;  % sqrt(2) due to complex  1.20 due to understimate of g-factor    
end

%%%%%%%%%%%%%%%%%%%%%%%%%%%%%%%%%%%%%%%%%%%%%%%%%%%%%%%%%%%%%%%%%%%%%%%%%%%%%

if ARG.MP>0
    ARG.soft_thrs=10;
end


if isfield(ARG,'soft_thrs_in')
    ARG.soft_thrs=ARG.soft_thrs_in;
end


%%%%%%%%%%%%%%%%%%%%%%%%%%%%%%%%%%%%%%%%%%%%%%%%%%%%%%%%%%%%%%%%%%%%%%%%%%%%%%%


KSP_weight=KSP2(:,:,:,1)*0;
NOISE=KSP_weight;
Component_threshold=KSP_weight;
energy_removed=KSP_weight;
SNR_weight=KSP_weight;
QQ.KSP_processed=zeros(1,size(KSP2,1)-ARG.kernel_size(1));

if ARG.patch_average==0
    KSP_processed=QQ.KSP_processed*0;
    for nw1=2:max(1,floor(ARG.kernel_size(1)/ARG.patch_average_sub));
        KSP_processed(1,nw1 : max(1,floor(ARG.kernel_size(1)/ARG.patch_average_sub)):end)=2;
    end
    KSP_processed(end)=0; % disp
    QQ.KSP_processed=KSP_processed;
end
disp('starting NORDIC ...')
KSP_processed;
QQ.ARG=ARG;
for n1=1:size(QQ.KSP_processed,2)
    [KSP_recon,~,KSP_weight,NOISE,Component_threshold,energy_removed,SNR_weight]=sub_LLR_Processing(KSP_recon,KSP2,ARG,n1,QQ,master_fast,KSP_weight,NOISE,Component_threshold,energy_removed,SNR_weight)   ;    % save all files
end
KSP_recon=KSP_recon./repmat((KSP_weight),[1 1 1 size(KSP2,4)]);  % Assumes that the combination is with N instead of sqrt(N). Works for NVR not MPPCA
ARG.NOISE=  sqrt(NOISE./KSP_weight);
ARG.Component_threshold = Component_threshold./KSP_weight;
ARG.energy_removed = energy_removed./KSP_weight;
ARG.SNR_weight  = SNR_weight./KSP_weight;
IMG2=KSP_recon;
disp('completing NORDIC ...')


if isfield(ARG,'save_residual_matlab')
if ARG.save_residual_matlab==1;
Residual=KSP2-KSP_recon;
  save([ARG.DIROUT 'RESIDUAL'  fn_out '.mat'   ],'Residual','-v7.3')
end
end


for n=1:size(IMG2,4);
    IMG2(:,:,:,n)= IMG2(:,:,:,n).* gfactor;
end


for slice=matdim(3):-1:1
    for n=1:size(IMG2,4); % include the noise
        IMG2(:,:,slice,n)=IMG2(:,:,slice,n).*exp(i*angle(meanphase(:,:,slice)));
    end
end

for slice=matdim(3):-1:1
    for n=1:size(IMG2,4);
        IMG2(:,:,slice,n)= IMG2(:,:,slice,n).*exp(i*angle( DD_phase(:,:,slice,n)   ));
    end
end


IMG2=IMG2.*ARG.ABSOLUTE_SCALE;

IMG2(isnan(IMG2))=0;

if isfield(ARG,'make_complex_nii')
    IMG2_tmp=abs(IMG2(:,:,:,1:end)); % remove g-factor and noise for DUAL 1
    IMG2_tmp(isnan(IMG2_tmp))=0;
    tmp=sort(abs(IMG2_tmp(:)));  sn_scale=2*tmp(round(0.99*end));%sn_scale=max();
    gain_level=floor(log2(32000/sn_scale));
    %IMG2_tmp= int16(abs(IMG2_tmp)*2^gain_level);
    
    if  ARG.full_dynamic_range==0; gain_level=0;end
    
    if strmatch(info.Datatype,'uint16')
        IMG2_tmp= uint16(abs(IMG2_tmp)*2^gain_level);
    elseif strmatch(info.Datatype,'int16')
        IMG2_tmp= int16(abs(IMG2_tmp)*2^gain_level);
    else
        IMG2_tmp= single(abs(IMG2_tmp)*2^gain_level);
    end
    
    niftiwrite((IMG2_tmp),[ARG.DIROUT fn_out '_magnitudeIMG.nii'],info)
    
    
    
    IMG2_tmp=angle(IMG2(:,:,:,1:end));
    if strmatch(info_phase.Datatype,'int16')
    %    IMG2_tmp=IMG2_tmp+pi;
    end
    
     IMG2_tmp=    (IMG2_tmp/(2*pi)+range_center)*range_norm;
    
     if strmatch(info_phase.Datatype,'uint16')      
        IMG2_tmp= uint16(IMG2_tmp);
     elseif strmatch(info_phase.Datatype,'int16')       
        IMG2_tmp= int16(IMG2_tmp);
     else
        IMG2_tmp= single((IMG2_tmp));
     end
    
    
    if 0
    if strmatch(info_phase.Datatype,'uint16')
        IMG2_tmp=IMG2_tmp/(2*pi)*phase_range;
        IMG2_tmp= uint16(abs(IMG2_tmp)*2^gain_level);
    elseif strmatch(info_phase.Datatype,'int16')
        IMG2_tmp=IMG2_tmp/(2*pi)*phase_range;
        IMG2_tmp= int16((IMG2_tmp)*2^gain_level);
    else
        IMG2_tmp= single(abs(IMG2_tmp)*2^gain_level);
    end
    end
    
    
    niftiwrite((IMG2_tmp),[ARG.DIROUT fn_out '_phaseIMG.nii'],info_phase)
    
else
    IMG2=abs(IMG2(:,:,:,1:end)); % remove g-factor and noise for DUAL 1
    IMG2(isnan(IMG2))=0;
    tmp=sort(abs(IMG2(:)));  sn_scale=2*tmp(round(0.99*end));%sn_scale=max();
    gain_level=floor(log2(32000/sn_scale));
    
    if  ARG.full_dynamic_range==0; gain_level=0;end
    
    if strmatch(info.Datatype,'uint16')
        IMG2= uint16(abs(IMG2)*2^gain_level);
    elseif strmatch(info.Datatype,'int16')
        IMG2= int16(abs(IMG2)*2^gain_level);
    else
        IMG2= single(abs(IMG2)*2^gain_level);
    end
    if ARG.use_generic_NII_read==0;
    niftiwrite((IMG2),[ARG.DIROUT fn_out(1:end) '.nii'],info)
    else
     nii=make_nii(IMG2);   
     save_nii(nii, [ARG.DIROUT fn_out(1:end) '.nii'])
    end
end


if isfield(ARG,'save_add_info')
    if  ARG.save_add_info==1
        disp('saving additional info')
        save([ARG.DIROUT fn_out '.mat'   ],'ARG2','ARG','-v7.3')
    end
end


return


function  [KSP_recon,KSP2,KSP2_weight,NOISE, Component_threshold,energy_removed,SNR_weight]=sub_LLR_Processing(KSP_recon,KSP2,ARG,n1,QQ,master,KSP2_weight,NOISE,Component_threshold,energy_removed,SNR_weight)

if ~exist('NOISE'); NOISE=[];  end
if ~exist('Component_threshold');Component_threshold=[];  end
if ~exist('energy_removed');  energy_removed=[]; end
if ~exist('SNR_weight'); SNR_weight=[]; end

%  QQ.KSP_processed  0 nothing done, 1 running, 2 saved 3 completed and  averaged

if master==0 && ARG.patch_average==0
    OPTION='NO_master_NO_PA';
    
elseif master==0 && ARG.patch_average==1
    OPTION='NO_master_PA'    ;
    
elseif master==1 && ARG.patch_average==0
    OPTION='master_NO_PA'    ;
    
elseif master==1 && ARG.patch_average==1
    OPTION='master_PA'   ;
    
end

switch OPTION
    
    case 'NO_master_NO_PA'
        
    case  'NO_master_PA'
        
    case  'master_NO_PA'
        
    case  'master_PA'
        
end


if     QQ.KSP_processed(1,n1)~=1  && QQ.KSP_processed(1,n1)~=3 % not being processed also not completed yet
    
    if     QQ.KSP_processed(1,n1)==2 && master==1%  processed but not added.
        % loading instead of processing
        % load file as soon as save, if more than 10 sec, just do the recon
        % instead.
        try  % try to load otherwise go to next slice
            load([ARG.filename  'slice' num2str(n1)  '.mat'],'DATA_full2')
        catch;
            QQ.KSP_processed(1,n1)=0;  % identified as bad file and being identified for reprocessing
            return  ;end
    end
    
    if QQ.KSP_processed(1,n1)~=2
        QQ.KSP_processed(1,n1)=1; % block for other processes
        if ~exist('DATA_full2')
            ARG2=QQ.ARG;
            if master==0
                QQ.KSP_processed(1,n1)=1    ;  % STARTING
                KSP2a=QQ.KSP2([1:ARG.kernel_size(1)]+(n1-1),:,:,:); lambda=ARG2.LLR_scale*ARG.NVR_threshold;
            else
                QQ.KSP_processed(1,n1)=1    ;  % STARTING
                KSP2a=KSP2([1:ARG.kernel_size(1)]+(n1-1),:,:,:); lambda=ARG2.LLR_scale*ARG.NVR_threshold;
                
            end
            
            if    ARG.patch_average==1
                %  [DATA_full2, ~,NOISE, Component_threshold] =subfunction_loop_for_NVR_avg(KSP2a,ARG.kernel_size(3),ARG.kernel_size(2),ARG.kernel_size(1),lambda,1,ARG.soft_thrs);
                [DATA_full2, KSP2_weight] =subfunction_loop_for_NVR_avg(KSP2a,ARG.kernel_size(3),ARG.kernel_size(2),ARG.kernel_size(1),lambda,1,ARG.soft_thrs,KSP2_weight);
            else
                
                KSP2_weight_tmp         =KSP2_weight([1:ARG.kernel_size(1)]+(n1-1),:,:,:);
                NOISE_tmp               =NOISE([1:ARG.kernel_size(1)]+(n1-1),:,:,:);
                Component_threshold_tmp =Component_threshold([1:ARG.kernel_size(1)]+(n1-1),:,:,:);
                energy_removed_tmp      =energy_removed([1:ARG.kernel_size(1)]+(n1-1),:,:,:);
                SNR_weight_tmp          =SNR_weight([1:ARG.kernel_size(1)]+(n1-1),:,:,:);
                
                [DATA_full2,KSP2_weight_tmp,NOISE_tmp, Component_threshold_tmp,energy_removed_tmp,SNR_weight_tmp] =...
                    subfunction_loop_for_NVR_avg_update(KSP2a,ARG.kernel_size(3),ARG.kernel_size(2),ARG.kernel_size(1),lambda,1,ARG.soft_thrs,KSP2_weight_tmp,ARG,NOISE_tmp,Component_threshold_tmp,energy_removed_tmp,SNR_weight_tmp);
                
                KSP2_weight([1:ARG.kernel_size(1)]+(n1-1),:,:,:)=KSP2_weight_tmp;
                
                try;     NOISE([1:ARG.kernel_size(1)]+(n1-1),:,:,:) =NOISE_tmp;  catch;end
                Component_threshold([1:ARG.kernel_size(1)]+(n1-1),:,:,:) = Component_threshold_tmp;
                energy_removed([1:ARG.kernel_size(1)]+(n1-1),:,:,:)  = energy_removed_tmp;
                SNR_weight([1:ARG.kernel_size(1)]+(n1-1),:,:,:)  = SNR_weight_tmp;
                %DATA_full=subfunction_loop_for_NVR(KSP2a,ARG.kernel_size(3),ARG.kernel_size(2),ARG.kernel_size(1),lambda);
                %DATA_full2(1, round(w2/2)+[1:size(DATA_full,1)],:,:  )=DATA_full;  % center plane only
            end
            
        end
        
    end
    
    
    
    if master==0
        if QQ.KSP_processed(1,n1)~=2
            save([ARG.filename  'slice' num2str(n1)  '.mat'],'DATA_full2', '-v7.3'        )
            QQ.KSP_processed(1,n1)=2    ;  % COMPLETED
        end
    else
        
        if    ARG.patch_average==1
            tmp=KSP_recon([1:ARG.kernel_size(1)]+(n1-1),:,:,:) ;
            KSP_recon([1:ARG.kernel_size(1)]+(n1-1),:,:,:)= tmp + DATA_full2;
        else
            KSP_recon([1:ARG.kernel_size(1)]+(n1-1) ,1:size(DATA_full2,2),:,:)=  KSP_recon([1:ARG.kernel_size(1)]+(n1-1) ,1:size(DATA_full2,2),:,:) +  DATA_full2;
        end
        QQ.KSP_processed(1,n1)=3     ;
    end
    
    
end


return


function [KSP2_tmp_update, KSP2_weight]=subfunction_loop_for_NVR_avg(KSP2a,w3,w2,w1,lambda2,patch_avg, soft_thrs,KSP2_weight,ARG)
if ~exist('patch_avg'); patch_avg=1;end
if ~exist('soft_thrs'); soft_thrs=[]; end

if ~exist('KSP2_weight')
    KSP2_weight=zeros(size(KSP2a(:,:,:,1)));
elseif isempty(KSP2_weight)
    KSP2_weight=zeros(size(KSP2a(:,:,:,1)));
end


if ~exist('KSP2_tmp_update')
    KSP2_tmp_update=zeros(size(KSP2a(:,:,:,:)));
elseif isempty(KSP2_tmp_update)
    KSP2_tmp_update=zeros(size(KSP2a(:,:,:,:)));
end


%   KSP2_weight=zeros(size(KSP2a(:,:,:,1)));
%   KSP2_tmp_update=zeros(size(KSP2a));

%   for n2=1:size(KSP2a,2)-w2+1;
%        for n3=1:size(KSP2a,3)-w3+1;
for n2=[1: max(1,floor(w2/ARG.patch_average_sub)):size(KSP2a,2)*1-w2+1  size(KSP2a,2)-w2+1];
    for n3=[1: max(1,floor(w3/ARG.patch_average_sub)):size(KSP2a,3)*1-w3+1  size(KSP2a,3)-w3+1  ];
        
        KSP2_tmp=KSP2a(:,[1:w2]+(n2-1),[1:w3]+(n3-1),:);
        tmp1=reshape(KSP2_tmp,[],size(KSP2_tmp,4));
        
        [U,S,V]=svd([(tmp1) ],'econ');
        S=diag(S);
        
        
        
        
        [idx]=sum(S<lambda2);
        if isempty(soft_thrs)
            S(S<lambda2)=0;
        elseif soft_thrs==10  % USING MPPCA
            
            
          %  disp('test for zero entries')
            Test_mat=sum(tmp1,2);
            sum(Test_mat==0)
            
            centering=0;
            MM=size(tmp1,1);
            NNN=size(tmp1,2);
            R = min(MM, NNN);
            scaling = (max(MM, NNN) - (0:R-centering-1)) / NNN;
            scaling = scaling(:);
            vals=S;
            vals = (vals).^2 / NNN;
            % First estimation of Sigma^2;  Eq 1 from ISMRM presentation
            csum = cumsum(vals(R-centering:-1:1)); cmean = csum(R-centering:-1:1)./(R-centering:-1:1)'; sigmasq_1 = cmean./scaling;
            % Second estimation of Sigma^2; Eq 2 from ISMRM presentation
            gamma = (MM - (0:R-centering-1)) / NNN;
            rangeMP = 4*sqrt(gamma(:));
            rangeData = vals(1:R-centering) - vals(R-centering);
            sigmasq_2 = rangeData./rangeMP;
            t = find(sigmasq_2 < sigmasq_1, 1);
            S(t:end)=0;
            
            
        else
            S(max(1,end-floor(idx*soft_thrs)):end)=0;
        end
        
        tmp1=U*diag(S)*V';
        
        tmp1=reshape(tmp1,size(KSP2_tmp));
        if patch_avg==1
            
            KSP2_tmp_update(:,[1:w2]+(n2-1),[1:w3]+(n3-1),:) =...
                KSP2_tmp_update(:,[1:w2]+(n2-1),[1:w3]+(n3-1),:) +tmp1;
            KSP2_weight(:,[1:w2]+(n2-1),[1:w3]+(n3-1),:) =...
                KSP2_weight(:,[1:w2]+(n2-1),[1:w3]+(n3-1),:) +1;
        else
            KSP2_tmp_update(:,round(w2/2)+(n2-1),round(w3/2)+(n3-1),:) =...
                KSP2_tmp_update(:,round(w2/2)+(n2-1),round(w3/2)+(n3-1),:) +tmp1(1,round(end/2),round(end/2),:);
            KSP2_weight(:,round(w2/2)+(n2-1),round(w3/2)+(n3-1),:) =...
                KSP2_weight(:,round(w2/2)+(n2-1),round(w3/2)+(n3-1),:) +1;
            
        end
        
        
    end
end


return


function [KSP2_tmp_update, KSP2_weight,NOISE,KSP2_tmp_update_threshold,energy_removed,SNR_weight]=subfunction_loop_for_NVR_avg_update(KSP2a,w3,w2,w1,lambda2,patch_avg, soft_thrs,KSP2_weight,ARG,NOISE,KSP2_tmp_update_threshold,energy_removed,SNR_weight)

if ~isfield(ARG,'patch_scale'); patch_scale=1; else; patch_scale=ARG.patch_scale;end
if ~exist('patch_avg'); patch_avg=1;end    %   patch_avg=0; means zero only
if ~exist('soft_thrs'); soft_thrs=[]; end

if ~exist('KSP2_weight')
    KSP2_weight=zeros(size(KSP2a(:,:,:,1)));
elseif isempty(KSP2_weight)
    KSP2_weight=zeros(size(KSP2a(:,:,:,1)));
end

if ~exist('NOISE_tmp')'%  ~exist('NOISE_tmpKSP2_tmp_update')
    NOISE_tmp=zeros(size(KSP2a(:,:,:,1)));
elseif isempty(KSP2_tmp_update)
    NOISE_tmp=zeros(size(KSP2a(:,:,:,1)));
end

if ~exist('KSP2_tmp_update_threshold')
    KSP2_tmp_update_threshold=zeros(size(KSP2a(:,:,:,1)));
elseif isempty(KSP2_tmp_update_threshold)
    KSP2_tmp_update_threshold=zeros(size(KSP2a(:,:,:,1)));
end

if ~exist('energy_removed')
    energy_removed=zeros(size(KSP2a(:,:,:,1)));
elseif isempty(energy_removed)
    energy_removed=zeros(size(KSP2a(:,:,:,1)));
end


if ~exist('SNR_weight')
    SNR_weight=zeros(size(KSP2a(:,:,:,1)));
elseif isempty(SNR_weight)
    SNR_weight=zeros(size(KSP2a(:,:,:,1)));
end


KSP2_tmp_update=0*KSP2a;
%NOISE=[];


for n2=[1: max(1,floor(w2/ARG.patch_average_sub)):size(KSP2a,2)*1-w2+1  size(KSP2a,2)-w2+1];
    for n3=[1: max(1,floor(w3/ARG.patch_average_sub)):size(KSP2a,3)*1-w3+1  size(KSP2a,3)-w3+1  ];
        
        KSP2_tmp=KSP2a(:,[1:w2]+(n2-1),[1:w3]+(n3-1),:);
        tmp1=reshape(KSP2_tmp,[],size(KSP2_tmp,4));
        
        [U,S,V]=svd([(tmp1) ],'econ');
        S=diag(S);
        
        
        
        [idx]=sum(S<lambda2);
        if isempty(soft_thrs)
            energy_scrub=sqrt(sum(S.^1)).\sqrt(sum(S(S<lambda2).^1));
            S(S<lambda2)=0;
            t=idx;
        elseif soft_thrs~=10;
            
            S=S-lambda2*soft_thrs;
            S(S<0)=0;
            energy_scrub=0;
            t=1;
            
        elseif soft_thrs==10  % USING MPPCA
            
         %  disp('test for zero entries')
            Test_mat=sum(tmp1,2);
            MM0=sum(Test_mat==0);
            
            
            if MM0>1  & MM0<100
             %  2 
            end
            
            
            centering=0;
            MM=size(tmp1,1)-MM0;  % Correction for some zero entries
            
            if MM>0
            NNN=size(tmp1,2);
            R = min(MM, NNN);
            scaling = (max(MM, NNN) - (0:R-centering-1)) / NNN;
            scaling = scaling(:);
            vals=S;
            vals = (vals).^2 / NNN;
            % First estimation of Sigma^2;  Eq 1 from ISMRM presentation
            csum = cumsum(vals(R-centering:-1:1)); cmean = csum(R-centering:-1:1)./(R-centering:-1:1)'; sigmasq_1 = cmean./scaling;
            % Second estimation of Sigma^2; Eq 2 from ISMRM presentation
            gamma = (MM - (0:R-centering-1)) / NNN;
            rangeMP = 4*sqrt(gamma(:));
            rangeData = vals(1:R-centering) - vals(R-centering);
            sigmasq_2 = rangeData./rangeMP;
            t = find(sigmasq_2 < sigmasq_1, 1);
            % NOISE(1:size(KSP2a,1),[1:w2]+(n2-1),[1:w3]+(n3-1),1) = sigmasq_2(t);
            idx=size(S(t:end),1)  ;
            energy_scrub=sqrt(sum(S.^1)).\sqrt(sum(S(t:end).^1));
            S(t:end)=0;
            else  % all zero entries
                t=1;
                energy_scrub=0;
                sigmasq_2=0;
            end
            
        else
            S(max(1,end-floor(idx*soft_thrs)):end)=0;
        end
        
        tmp1=U*diag(S)*V';
        
        tmp1=reshape(tmp1,size(KSP2_tmp));
        
        if patch_scale==1; else; patch_scale=size(S,1)-idx; end
        
        if isempty(t);  t=1; end  % threshold removed all.
        
        if patch_avg==1
            
            KSP2_tmp_update(:,[1:w2]+(n2-1),[1:w3]+(n3-1),:) =...
                KSP2_tmp_update(:,[1:w2]+(n2-1),[1:w3]+(n3-1),:) +patch_scale*tmp1;
            KSP2_weight(:,[1:w2]+(n2-1),[1:w3]+(n3-1),:) =...
                KSP2_weight(:,[1:w2]+(n2-1),[1:w3]+(n3-1),:) + patch_scale;
            KSP2_tmp_update_threshold(:,[1:w2]+(n2-1),[1:w3]+(n3-1),:) =...
                KSP2_tmp_update_threshold(:,[1:w2]+(n2-1),[1:w3]+(n3-1),:) +idx;
            energy_removed(:,[1:w2]+(n2-1),[1:w3]+(n3-1),:) =...
                energy_removed(:,[1:w2]+(n2-1),[1:w3]+(n3-1),:) +energy_scrub;
            
            SNR_weight(:,[1:w2]+(n2-1),[1:w3]+(n3-1),1) =...
                SNR_weight(:,[1:w2]+(n2-1),[1:w3]+(n3-1),1) + S(1)./S(max(1,t-1));
            
            try
                NOISE(1:size(KSP2a,1),[1:w2]+(n2-1),[1:w3]+(n3-1),1) = ...
                    NOISE(1:size(KSP2a,1),[1:w2]+(n2-1),[1:w3]+(n3-1),1) +   sigmasq_2(t);; catch; end
        else
            KSP2_tmp_update(:,round(w2/2)+(n2-1),round(w3/2)+(n3-1),:) =...
                KSP2_tmp_update(:,round(w2/2)+(n2-1),round(w3/2)+(n3-1),:) +patch_scale*tmp1(1,round(end/2),round(end/2),:);
            KSP2_weight(:,round(w2/2)+(n2-1),round(w3/2)+(n3-1),:) =...
                KSP2_weight(:,round(w2/2)+(n2-1),round(w3/2)+(n3-1),:) +patch_scale;
            KSP2_tmp_update_threshold(:,round(w2/2)+(n2-1),round(w3/2)+(n3-1),:) =...
                KSP2_tmp_update_threshold(:,round(w2/2)+(n2-1),round(w3/2)+(n3-1),:) +idx;
            energy_removed(:,round(w2/2)+(n2-1),round(w3/2)+(n3-1),:)  =...
                energy_removed(:,round(w2/2)+(n2-1),round(w3/2)+(n3-1),:) +energy_scrub;
            
            SNR_weight(:,round(w2/2)+(n2-1),round(w3/2)+(n3-1),1) =...
                SNR_weight(:,round(w2/2)+(n2-1),round(w3/2)+(n3-1),1) + S(1)./S(max(1,t-1));
            try
                NOISE(:,round(w2/2)+(n2-1),round(w3/2)+(n3-1),:) = ...
                    NOISE(:,round(w2/2)+(n2-1),round(w3/2)+(n3-1),:) +   sigmasq_2(t);; catch; end
            
        end
        
%            if MM0>1  & MM0<196
            
            %   [  sigmasq_2(t) MM0] %  2 
%            end
        
        
    end
end


return
```
