## Supplementary material for "Neptune: a toolbox for spinal cord functional MRI data processing and quality assurance": Neptune software: pcorr_zscore.m

```
function [z, r] = pcorr_zscore(a, b, dof_correction, junk, siz4)

% Created 2015-Feb-20 (RLB)
% Revised 2015-Feb-23 - updated calculation for degrees of freedom
% Revised 2015-Apr-07 - permit truncation of time series (via siz4)
% Revised 2015-Apr-10 - bug fix (junk needs to be truncated too)

if nargin < 5
    siz4 = size(a,2);
    if nargin < 3
        dof_correction = 1;
    end
end

% only truncate time series if user specifies length
if exist('siz4','var')
    if (size(a,2)~=siz4)||(size(b,2)~=siz4)
        a = a(:,1:siz4);
        b = b(:,1:siz4);
        junk = junk(:,1:siz4);
    end
end

% zero mean
a = a - mean(a,2);
b = b - mean(b,2);
junk = junk - repmat(mean(junk,2), [1 size(junk,2)]);

% partial correlation
x = [a; b]';
[r,~] = partialcorr(x,junk');
r = r(1,2);

z_from_r = atanh(r);

if dof_correction
    
    % regress the nuisance time series from the time series of interest to
    % improve the estimate of the degrees of freedom
    a = reshape(((a') - (junk' * lscov(junk',a'))), [1 siz4]);
    b = reshape(((b') - (junk' * lscov(junk',b'))), [1 siz4]);
    
    % first order autocorrelation
    p1 = corrcoef(a(2:end),a(1:end-1));
    p1 = p1(1,2);
    p2 = corrcoef(b(2:end),b(1:end-1));
    p2 = p2(1,2);
    
    % actual degrees of freedom estimates
    dof1 = (1-p1^2)/(1+p1^2);
    dof2 = (1-p2^2)/(1+p2^2);
    
    % corrected z-score
    z1 = z_from_r .* sqrt((siz4.*dof1)-3);
    z2 = z_from_r .* sqrt((siz4.*dof2)-3);
    z = (z1 + z2) / 2;
    
else % do not correct for inflated degrees of freedom
    
    z = z_from_r .* sqrt(siz4-3);
end
```
