## Supplementary material for "Neptune: a toolbox for spinal cord functional MRI data processing and quality assurance": Neptune software: pcorr_zscore_vec.m

```
function [z, temp1f, temp2f, r] = pcorr_zscore_vec(a, b, dof_correction, junk, JLTM_cutoff, siz4)

% Created 2015-Mar-16 (RLB)
% Revised 2015-Apr-07 - permit truncation of time series (via siz4)
% Updated on 2023-10-03 by Ranga Deshpande to discard NaN values

if nargin < 6
    siz4 = size(a,2);
    if nargin < 5
        JLTM_cutoff = 0.95;
        if nargin < 3
            dof_correction = 1;
        end
    end
end

% perform all possible correlations
temp1 = zeros((size(a,1).*size(b,1)), 1); temp2=temp1;
ii = 0;
for i1 = 1 : size(a,1)
    for i2 = 1 : size(b,1)
        ii = ii + 1;
        [temp1(ii,1), temp2(ii,1)] = pcorr_zscore(a(i1,:), b(i2,:), dof_correction, junk, siz4);
    end % i2
end % i1
temps1 = sort(temp1(:));
temps2 = sort(temp2(:));

% Discard NaN values (this option was added in October 2023 by Ranga Deshpande)
temps1(find(isnan(temps1))) = [];
temps2(find(isnan(temps2))) = [];

% the correlation that we want :)
z = temps1(ceil(size(temps1,1)*JLTM_cutoff),1);
r = temps2(ceil(size(temps2,1)*JLTM_cutoff),1);

% find the corresponding time series
ii = 0;
for i1 = 1 : size(a,1)
    for i2 = 1 : size(b,1)
        ii = ii + 1;
        if temp1(ii,1) == z
            temp1f = a(i1,:);
            temp2f = b(i2,:);
        end
    end % i2
end % i1

end
```
