## Supplementary material for "Neptune: a toolbox for spinal cord functional MRI data processing and quality assurance": Neptune software: phys_filt_sync_RLB.m

```
function   [scan_sync, ...
            phys_out, ...
            phys_out_allchan] = phys_filt_sync(current_file_phys, TR, parms, channel, nDrop)

%**************************************************************************
% This function displays the frequency components of the physiological
% log file, revealing frequency components of heartrate and respiration.
%**************************************************************************

%**************************************************************************
% Read the prescribed channels of the physlog
%**************************************************************************

channels = 1:10;            % Total channels = 1:10; Cardiac = 5; Resp = 6; 

phys_out_allchan = read_physlog_file(current_file_phys, channels);

%**************************************************************************
% Filter the physlog with a gaussian to remove spikes (no need to detrend)
%**************************************************************************
   
% gaussian_filter_1d = fspecial('gaussian',[1 450],250);

% phys_out = filtfilt(gaussian_filter_1d, 1, phys_out_allchan(:, channel));

phys_out = phys_out_allchan(:,channel)';       % Transpose to row vector

%**************************************************************************
% Here we indentify the portion of the dynamic image data from the 
% scanner that lines up with the phys log data.  The physlog starts
% recording at the beginning of the preparation pulses and continues until
% the scan is done.  Because some scans are started automatically and
% others are started manually (with a variable amount of time between the
% preparaton pulses and the actual start of the scan), it is necessary to
% start at the end of the phys log and work backwards to find the
% correct timing.
%
% The physlog data is sampled at 500 Hz, or 500 times per second.  Thus,
% the time between each physlog sample is 0.002 seconds, or 2 miliseconds
%**************************************************************************

fs_phys = 500;

%**************************************************************************
% Find the end scan marker(s) in the physlog.  The marker is '20' in the
% physlog in hexidecimal format.  The physlog is read in as decimal, so we
% look for the marker 32.  (20 = hex(32)).
%
% The Mark channel in the phys log is channel 10, so we search there:
%**************************************************************************

scan_end_indices = find(phys_out_allchan(:,10) > 31);

%**************************************************************************
% For reasons unknown, sometimes there is more than one stop marker.
% We only care about the last one:
%**************************************************************************

scan_end = scan_end_indices(end);

%**************************************************************************
% Now that we have the end of the physio data we can determine how long, in
% seconds, the physiological data was being accumulated:
%**************************************************************************

phys_run_time = scan_end * (1/fs_phys);

%**************************************************************************
% Compare this to the scan duration information (also in seconds) from the 
% scan parameter file (read in by the VUIIS par file reader) and 
% Calculate the time offset between the scan and the physlog data:
%**************************************************************************

phys_time_offset = phys_run_time - parms.scan_dur;

%**************************************************************************
% Get the difference in calculated dynamics and actual dynamics
% This difference is PROBABLY the result of the dummy dynamic scans and
% other things.
%**************************************************************************

calculated_dyns = parms.scan_dur / TR;

dyn_scan_offset = calculated_dyns - parms.max_dynamics;

%**************************************************************************
% We know the length of one repetition time and the end of the physlog
% scan.  So now we can step back any number of dynamic scans to find where
% in the physlog corresponds to the beginning of the collection of MRI
% data.  ONE TR Corresponds to ONE Dynamic Scan
%**************************************************************************

phys_samples_TR = TR / (1/fs_phys);

%**************************************************************************
% Now that we know the number of physlog samples in one TR, and how many
% TRs in the scan, we can determine the number of physlog samples in the
% scan:
%**************************************************************************

phys_samples_total_scan = parms.max_dynamics * phys_samples_TR;

phys_sync_dur = phys_samples_total_scan * (1/fs_phys);

%**************************************************************************
% Now we can start at the scan end marker in the physlog and backtrack by
% the amount of physlog samples in the total scan to find where the physlog
% data syncs up with the MRI scan.
%**************************************************************************

scan_start = scan_end - phys_samples_total_scan;

phys_sync_duration_scans = scan_end - scan_start;

%**************************************************************************
% Find the synchronized data
%**************************************************************************

scan_sync = phys_out(scan_start+1:scan_end);                               

%**************************************************************************
% Downsample the physlog, one per dynamic scan.
%**************************************************************************

% 2009-05-27 do not downsample

%one_per_TR = round(length(scan_sync)/parms.max_dynamics);

%scan_sync = downsample(scan_sync, one_per_TR);

%**************************************************************************
% Make the scan_sync the length as the number of dynamic scans used from
% the image set.  We may discard scans at the beginning of the acquisition
% period to allow the signal to stabilize, so start at end, work backwards.
%**************************************************************************

%scan_sync = scan_sync(nDrop+1:parms.max_dynamics);
```
