## Supplementary material for "Neptune: a toolbox for spinal cord functional MRI data processing and quality assurance": Neptune software: read_1D_xform_file.m

```
function xform_out = read_1D_xform_file(xform_file)

% Robert L. Barry
% 2014-04-04
% 
% Reads in an AFNI transformation matrix.

fid = fopen(xform_file);
xform_out = textscan(fid,'%n %n %n %n %n %n %n %n %n %n %n %n ','commentStyle','#');
fclose(fid);
```
