## Supplementary material for "Neptune: a toolbox for spinal cord functional MRI data processing and quality assurance": Neptune software: robustdirname.m

```
function rdname = robustdirname(dname)

% Fix a directory path when it contains one or more spaces.

% 2017-09-18
% 2020-02-23 - removed special handling for Rob's computer

if ~strcmp(dname(end), '/')
    dname = [dname '/'];
end

idx_sp = any(ismember(dname, ' ')); % spaces?
rdname = dname; % trivial case - no spaces exist

if idx_sp % if any spaces exist...
    idx_fs = find(ismember(rdname, '/')); % forward slash
    for i1 = 1 : size(idx_fs,2)-1
        i1a = idx_fs(1,i1);
        i1b = idx_fs(1,i1+1);
        temp = rdname(1,i1a:i1b);
        idx_temp = find(ismember(temp, ' '));
        if ~isempty(idx_temp) && ~strcmp(temp(1,2), '''') && ~strcmp(temp(1,end-1), '''') 
            rdname = [rdname(1,1:i1a) '''' rdname(1,i1a+1:i1b-1) '''' rdname(1,i1b:end)];
            idx_fs = find(ismember(rdname, '/')); % forward slash
        end
    end
end
```
