## Supplementary material for "Neptune: a toolbox for spinal cord functional MRI data processing and quality assurance": Neptune software: spinal_cord_registration_9b.m

```
function [final_reg, adj_final_trans, use_cmass] = spinal_cord_registration_9b(base_dir, fname, fname_anat, siz3, siz4, varargin)

% Last updated: 2021-04-12 - Added functionality to use motion corrected data if RETROICOR was not performed or skipped (*_MC* must be used and *_MC_mean* must be computed and used instead of *_MC_ricor* and *_MC_ricor_mean* respectively) (Ranga Deshpande)
%               2015-01-19 - Added boundary (outside CSF) constraint
%               2015-09-04 - Updated -ls and -sp weightings
%               2015-09-09 - Modified to use mean functional image
%               2015-11-23 - Normalized intensities of images before
%                            estimation of transform parameters

% no rotation, no shearing
affine_parms = '-parfix 3 0 -parfix 4 0 -parfix 5 0 -parfix 6 0 -parfix 9 1 -parfix 10 0 -parfix 11 0 -parfix 12 0 -onepass -nmatch 100% -parang 7 1/1.01 1.01 -parang 8 1/1.02 1.02 ';
%affine_parms = '-parfix 3 0 -parfix 4 0 -parfix 5 0 -parfix 6 0 -parfix 9 1 -parfix 11 0 -parfix 12 0 -onepass -nmatch 100% -parang 7 1/1.01 1.01 -parang 8 1/1.05 1.05 -maxshr 0.05 ';
%affine_parms = '-parfix 3 0 -parfix 4 0 -parfix 5 0 -parfix 6 0 -parfix 9 1 -parfix 11 0 -parfix 12 0 -onepass -nmatch 100% -parang 7 1/1.001 1.001 -parang 8 1/1.001 1.001 -maxshr 0.001 ';

% maximum translations for coarse and fine passes
%smax_vec = [2.5 1.0];
%smax_vec = [2.0 1.0];
%smax_vec = [1.5 0.75];
smax_vec = [5 1]; % don't need to be too specific here
smax1 = smax_vec(1,1);
smax2 = smax_vec(1,2);

siz4max = min(siz4, 10); % max number of dynamics used to estimate initial translations
siz4vec = 1:floor(siz4/siz4max):siz4;
siz4vec = siz4vec(1,1:siz4max);

%cost_fcns = {'-lpa '}; % fixed hel/nmi/mi problem of wild translations
%cost_fcns = {'-hel ' '-nmi ' '-mi '};
%cost_fcns = {'-hel ' '-nmi ' '-mi ' '-ls ' '-crM ' '-crA ' '-crU '};
%cost_fcns = {'-hel ' '-nmi ' '-mi ' '-ls ' '-crM ' '-crA ' '-crU ' '-sp ' '-je '};
cost_fcns = {'-hel ' '-nmi ' '-mi ' '-ls ' '-crM ' '-crA ' '-crU ' '-sp ' '-je ' '-lpa '};
%cost_fcns = {'-hel ' '-nmi ' '-mi ' '-ls ' '-crM ' '-crA ' '-crU ' '-sp ' '-lpa '};
%cost_fcns = {'-hel ' '-nmi ' '-mi ' '-ls ' '-crM ' '-crA ' '-crU ' '-sp ' '-je ' '-lss ' '-lpc ' '-lpa '};
%cost_fcns = {'-ls ' '-crM ' '-crA ' '-crU ' '-lpa '};
%cost_fcns = {'-crM ' '-crA '};
%cost_fcns = {'-hel '};
%cost_fcns = {'-ls ' '-crM ' '-crA ' '-crU ' '-sp '};
%cost_fcns = {'-ls '};

%cost_fcns = {'-ls ' '-sp ' '-hel ' '-nmi ' '-mi '};

%cost_fcns = {'-ls ' '-sp '}; % This version of the function requires these
                             % values to remain fixed and in this order.

% From 3dAllineate documentation:
% 
% "standard":
%   hel :: - Hellinger distance(base,source)
%   nmi :: 1/Normalized MI = H(base,source)/[H(base)+H(source)]
%   mi  :: - Mutual Information = H(base,source)-H(base)-H(source)
%   ls  :: 1 - abs(Pearson correlation coefficient)
%   crM :: 1 - abs[ CR(base,source) * CR(source,base) ]
%   crA :: 1 - abs[ CR(base,source) + CR(source,base) ]
%   crU :: CR(source,base) = Var(source|base) / Var(source)
% 
% "extra":
%   sp  :: 1 - abs(Spearman correlation coefficient)
%   je  :: H(base,source) = joint entropy of image pair
% 
% not sure about these:
%   lss :: Pearson correlation coefficient between image pair
%   lpc :: nonlinear average of Pearson cc over local neighborhoods
%   lpa :: 1 - abs(lpc)


if isempty(varargin)
    wbar3 = waitbar(0,'11. Co-registration (functional to anatomical)...','Name','Progress(11): Co-registration (functional to anatomical)...','Units','normalized','Position',[3/5 0 1/5 1/17]);
else
    wbar3 = varargin{1};
end

final_reg = zeros(siz3, size(cost_fcns,2), 2);

try % if the function "scfMRItb_04_resplitData.m" is not available then 'try' will ensure that this code will run smoothly nonetheless
    scfMRItb_04_resplitData(base_dir, fname, '_Smask', siz3); % if slice-wise data doesn't exist but combined 4D data exists, then split the data to generate the needed slice-wise data
    if ~(exist([base_dir fname '_slice' num2str(siz3) '_Smask.nii'], 'file'))
        try scfMRItb_04_resplitData(base_dir, fname, '_Gaussian_mask', siz3); catch, end % if slice-wise data doesn't exist but combined 4D data exists, then split the data to generate the needed slice-wise data
    end
    scfMRItb_04_resplitData(base_dir, fname_anat, '_affine_reg_mask', siz3); % if slice-wise data doesn't exist but combined 4D data exists, then split the data to generate the needed slice-wise data
    scfMRItb_04_resplitData(base_dir, fname_anat, '', siz3); % if slice-wise data doesn't exist but combined 4D data exists, then split the data to generate the needed slice-wise data
    scfMRItb_04_resplitData(base_dir, fname, '_MC_ricor', siz3); % if slice-wise data doesn't exist but combined 4D data exists, then split the data to generate the needed slice-wise data
    scfMRItb_04_resplitData(base_dir, fname, '_MC', siz3); % if slice-wise data doesn't exist but combined 4D data exists, then split the data to generate the needed slice-wise datacatch
    scfMRItb_04_resplitData(base_dir, fname, '_MC_ricor_mean', siz3); % if slice-wise data doesn't exist but combined 4D data exists, then split the data to generate the needed slice-wise data
    scfMRItb_04_resplitData(base_dir, fname, '_MC_mean', siz3); % if slice-wise data doesn't exist but combined 4D data exists, then split the data to generate the needed slice-wise data
    scfMRItb_04_resplitData(base_dir, fname, '_base_masked', siz3); % if slice-wise data doesn't exist but combined 4D data exists, then split the data to generate the needed slice-wise data
catch
end

%% Check if anat affine_reg_mask and func base_masked exist. If not, generate them

try scfMRItb_04_resplitData(base_dir, fname_anat, '_affine_reg_mask', siz3); catch, end % if slice-wise data doesn't exist but combined 4D data exists, then split the data to generate the needed slice-wise data

if ~exist([base_dir fname_anat '_slice' num2str(siz3) '_affine_reg_mask.nii'], 'file')

try
scfMRItb_04_unzipFile(base_dir, fname_anat, '_mask_GM')
scfMRItb_04_unzipFile(base_dir, fname_anat, '_mask_WM')
scfMRItb_04_unzipFile(base_dir, fname_anat, '_mask_CSF')
catch
end
mask_GM = load_untouch_nii([base_dir fname_anat '_mask_GM.nii']);
mask_WM = load_untouch_nii([base_dir fname_anat '_mask_WM.nii']);
mask_CSF = load_untouch_nii([base_dir fname_anat '_mask_CSF.nii']);

mask_all = or(or(mask_GM.img, mask_WM.img), mask_CSF.img) > 0;
mask_all_d = imdilate(mask_all, strel('disk', 20));

% consider only GM, WM, and a boundary condition (a ring around CSF)
affine_reg_mask = 5.00 .* single(mask_GM.img) + 5.00 .* single(mask_WM.img) + ...
    + 5.00 .* single(mask_CSF.img) + 5.00 .* xor(mask_all, mask_all_d);

% save affine_reg_mask
try scfMRItb_04_resplitData(base_dir, fname_anat, '_affine_reg_mask', siz3); catch, end % if slice-wise data doesn't exist but combined 4D data exists, then split the data to generate the needed slice-wise data
try
scfMRItb_04_unzipFile(base_dir, fname_anat, '')
scfMRItb_04_resplitData(base_dir, fname_anat, '', siz3) % if slice-wise data doesn't exist but combined 4D data exists, then split the data to generate the needed slice-wise data
catch
end
for i3 = 1 : siz3
    unix(['rm -f ' base_dir fname_anat '_slice' num2str(i3) '_affine_reg_mask.nii']);
    try scfMRItb_04_unzipFile(base_dir, fname_anat, ['_slice' num2str(i3)]); catch, end
    AM = load_untouch_nii([base_dir fname_anat '_slice' num2str(i3) '.nii']);
    AM.img = affine_reg_mask(:,:,i3);
    save_untouch_nii(AM, [base_dir fname_anat '_slice' num2str(i3) '_affine_reg_mask.nii']);
    clear AM
end
end


try scfMRItb_04_resplitData(base_dir, fname, '_base_masked', siz3); catch, end % if slice-wise data doesn't exist but combined 4D data exists, then split the data to generate the needed slice-wise data
if ~exist([base_dir fname '_slice' num2str(siz3) '_base_masked.nii'], 'file')
    
fname3 = [fname '_mean'];
try scfMRItb_04_unzipFile(base_dir, fname3, ''); catch, end
try scfMRItb_04_unzipFile(base_dir, fname, '_Gaussian_mask'); catch, end
F = load_untouch_nii([base_dir fname3 '.nii']);
F_Gmask = load_untouch_nii([base_dir fname '_Gaussian_mask.nii']);
funct_mask = single(F_Gmask.img);

try scfMRItb_04_resplitData(base_dir, fname, '_base_masked', siz3); catch, end % if slice-wise data doesn't exist but combined 4D data exists, then split the data to generate the needed slice-wise data
try
scfMRItb_04_unzipFile(base_dir, fname, '_base')
scfMRItb_04_resplitData(base_dir, fname, '_base', siz3) % if slice-wise data doesn't exist but combined 4D data exists, then split the data to generate the needed slice-wise data
catch
end
for i3 = 1 : size(F.img,3)
    waitbar((i3/size(F.img,3)),wbar3,sprintf('10. Save masked functional: slice (%d) of (%d)',i3,size(F.img,3)));
    unix(['rm -f ' base_dir fname '_slice' num2str(i3) '_base_masked.nii']);
    try scfMRItb_04_unzipFile(base_dir, fname, ['_slice' num2str(i3) '_base']); catch, end
    FM = load_untouch_nii([base_dir fname '_slice' num2str(i3) '_base.nii']);
    FM.img = single(FM.img) .* funct_mask(:,:,i3); % functional * mask
    save_untouch_nii(FM, [base_dir fname '_slice' num2str(i3) '_base_masked.nii']);
    clear FM;
end
end

%% ----- CENTER OF MASS ALIGNMENT FROM FUNCTIONAL TO ANATOMICAL ------------

eval(['load ' base_dir fname '_mask_NS.mat']); % load mask_NS
CoM_shifts = zeros(2,siz3);

try
scfMRItb_04_unzipFile(base_dir, fname, '_Gaussian_mask')
scfMRItb_04_resplitData(base_dir, fname, '_Gaussian_mask', siz3) % if slice-wise data doesn't exist but combined 4D data exists, then split the data to generate the needed slice-wise data
catch
end
try
scfMRItb_04_unzipFile(base_dir, fname, '_MC_ricor')
scfMRItb_04_resplitData(base_dir, fname, '_MC_ricor', siz3) % if slice-wise data doesn't exist but combined 4D data exists, then split the data to generate the needed slice-wise data
catch
end
try
scfMRItb_04_unzipFile(base_dir, fname, '_MC')
scfMRItb_04_resplitData(base_dir, fname, '_MC', siz3) % if slice-wise data doesn't exist but combined 4D data exists, then split the data to generate the needed slice-wise data
catch
end
try
scfMRItb_04_unzipFile(base_dir, fname_anat, '')
scfMRItb_04_resplitData(base_dir, fname_anat, '', siz3) % if slice-wise data doesn't exist but combined 4D data exists, then split the data to generate the needed slice-wise data
catch
end
try
scfMRItb_04_unzipFile(base_dir, fname_anat, '_affine_reg_mask')
scfMRItb_04_resplitData(base_dir, fname_anat, '_affine_reg_mask', siz3) % if slice-wise data doesn't exist but combined 4D data exists, then split the data to generate the needed slice-wise data
catch
end

for i3 = 1 : siz3
    waitbar(0/4+(i3/siz3)/4,wbar3,sprintf('11. Co-registration (1/4) center of mass alignment: slice (%d) of (%d)',i3,siz3));
    
    if ~(exist([base_dir fname '_slice' num2str(i3) '_Smask.nii'], 'file'))
        try scfMRItb_04_unzipFile(base_dir, fname, ['_slice' num2str(i3) '_Gaussian_mask']); catch, end
        F_weights = load_untouch_nii([base_dir fname '_slice' num2str(i3) '_Gaussian_mask.nii']);
        unix(['rm -f ' base_dir fname '_slice' num2str(i3) '_Smask.nii']);
        F_weights.img = ~mask_NS.img(:,:,i3);
        save_untouch_nii(F_weights, [base_dir fname '_slice' num2str(i3) '_Smask.nii']);
        clear F_weights;
    end
    
    % A_base = anatomical
    try
        scfMRItb_04_unzipFile(base_dir, fname_anat, ['_slice' num2str(i3) '_affine_reg_mask'])
        scfMRItb_04_unzipFile(base_dir, fname_anat, ['_slice' num2str(i3)])
    catch
    end
    A_base_mask = load_untouch_nii([base_dir fname_anat '_slice' num2str(i3) '_affine_reg_mask.nii']);
    A_base = load_untouch_nii([base_dir fname_anat '_slice' num2str(i3) '.nii']);
    A_base.img = single(single(A_base.img) .* (A_base_mask.img > 0));
    
    % A_input = functional
    try
        try scfMRItb_04_unzipFile(base_dir, fname, ['_slice' num2str(i3) '_MC_ricor']); catch, end
        A_input_all_dyn = load_untouch_nii([base_dir fname '_slice' num2str(i3) '_MC_ricor.nii']); % use data output from RETROICOR, if available
    catch
        try scfMRItb_04_unzipFile(base_dir, fname, ['_slice' num2str(i3) '_MC']); catch, end
        A_input_all_dyn = load_untouch_nii([base_dir fname '_slice' num2str(i3) '_MC.nii']);       % if RETROICOR was skipped or not performed, then use pre-RETROICOR motion-corrected data
    end
    
    % resize anatomical
    A_target = imresize(A_base.img, [size(A_input_all_dyn.img,1) size(A_input_all_dyn.img,2)], 'bilinear');

    % Estimate translation from several functional images
    CoM_shifts_i4 = zeros(size(siz4vec,2), 2);
    i4i = 0;
    for i4 = siz4vec
        i4i = i4i + 1;
        A_input = single(A_input_all_dyn.img(:,:,1,i4)) .* ~mask_NS.img(:,:,i3);
        
        A_target_sum_rows = sum(A_target,2);
        CoM_target_r = (sum(A_target_sum_rows .* ((1:1:size(A_target_sum_rows,1))'),1) ./ sum(A_target_sum_rows,1));
        A_target_sum_cols = sum(A_target,1);
        CoM_target_c = (sum(A_target_sum_cols .* (1:1:size(A_target_sum_cols,2)),2) ./ sum(A_target_sum_cols,2));
        
        A_input_sum_rows = sum(A_input,2);
        CoM_input_r = (sum(A_input_sum_rows .* ((1:1:size(A_input_sum_rows,1))'),1) ./ sum(A_input_sum_rows,1));
        A_input_sum_cols = sum(A_input,1);
        CoM_input_c = (sum(A_input_sum_cols .* (1:1:size(A_input_sum_cols,2)),2) ./ sum(A_input_sum_cols,2));
    
        % row = A-P translation (+ve value = shift anterior)
    %    -(-CoM_input_r+CoM_target_r)
        % column = L-R translation (+ve value = shift left)
    %    -CoM_input_c+CoM_target_c
        
        CoM_shifts_i4(i4i,:) = [-(-CoM_input_r+CoM_target_r) -CoM_input_c+CoM_target_c];
    end
    
    CoM_shifts(:,i3) = median(CoM_shifts_i4,1);
    clear A_base A_input* A_target* CoM_input* CoM_target* CoM_shifts_i4;
end
% CoM_shifts
CoM_shifts = CoM_shifts ./ 6; % ballpark estimate is okay


%% ----- SLICE-BY-SLICE AFFINE REGISTRATION OF BASE ------------------------

try
scfMRItb_04_unzipFile(base_dir, fname, '_base_masked')
scfMRItb_04_resplitData(base_dir, fname, '_base_masked', siz3) % if slice-wise data doesn't exist but combined 4D data exists, then split the data to generate the needed slice-wise data
catch
end
try
scfMRItb_04_unzipFile(base_dir, fname, '_MC_ricor_mean')
scfMRItb_04_resplitData(base_dir, fname, '_MC_ricor_mean', siz3) % if slice-wise data doesn't exist but combined 4D data exists, then split the data to generate the needed slice-wise data
catch
end
try
scfMRItb_04_unzipFile(base_dir, fname, '_MC_mean')
scfMRItb_04_resplitData(base_dir, fname, '_MC_mean', siz3) % if slice-wise data doesn't exist but combined 4D data exists, then split the data to generate the needed slice-wise data
catch
end
try
scfMRItb_04_unzipFile(base_dir, fname, '_Smask')
scfMRItb_04_resplitData(base_dir, fname, '_Smask', siz3) % if slice-wise data doesn't exist but combined 4D data exists, then split the data to generate the needed slice-wise data
catch
end


% was RETROICOR performed?
% if RETROICOR was skipped or not performed, then compute mean functional from pre-RETROICOR motion-corrected data as follows (next 12 lines)
if ~(exist([base_dir fname '_MC_ricor_mean.nii'], 'file'))    
    unix(['rm -f ' base_dir fname '_MC_mean.nii']);
    unix(['3dTstat -mean -prefix ' base_dir fname '_MC_mean.nii ' base_dir fname '_MC.nii']);
end
for i3 = 1 : siz3
    if ~(exist([base_dir fname '_slice' num2str(i3) '_MC_ricor_mean.nii'], 'file'))
    unix(['rm -f ' base_dir fname '_slice' num2str(i3) '_MC_mean.nii']);
    unix(['3dZcutup -keep ' num2str(i3-1) ' ' num2str(i3-1) ' ' ...
        '-prefix ' base_dir fname '_slice' num2str(i3) '_MC_mean.nii' ' ' ...
        base_dir fname '_MC_mean.nii']);
    unix(['3dTstat -mean -prefix ' base_dir fname '_slice' num2str(i3) '_MC_mean.nii ' base_dir fname '_slice' num2str(i3) '_MC.nii']);
    end
end

use_cmass = 0; % by default, aligning centers of mass is not required

for i3 = 1 : siz3
    waitbar(1/4+(i3/siz3)/4,wbar3,sprintf('11. Co-registration (2/4) affine reg of base image: slice (%d) of (%d)',i3,siz3));
    fprintf('func-to-anat registration: slice (%d) of (%d) -> subject %s\n',i3,siz3,fname)
    
    try scfMRItb_04_unzipFile(base_dir, fname, ['_slice' num2str(i3) '_Smask']); catch, end
    F_weights = load_untouch_nii([base_dir fname '_slice' num2str(i3) '_Smask.nii']);
    F_weights = single(F_weights.img);
    
    try % use data output from RETROICOR, if available
        try scfMRItb_04_unzipFile(base_dir, fname, ['_slice' num2str(i3) '_MC_ricor_mean']); catch, end
        F_oneslice = load_untouch_nii([base_dir fname '_slice' num2str(i3) '_MC_ricor_mean.nii']);
    catch % if RETROICOR was skipped or not performed, then use mean functional from pre-RETROICOR motion-corrected data
        try scfMRItb_04_unzipFile(base_dir, fname, ['_slice' num2str(i3) '_MC_mean']); catch, end
        F_oneslice = load_untouch_nii([base_dir fname '_slice' num2str(i3) '_MC_mean.nii']);
    end
    F_oneslice = single(F_oneslice.img);

    % prepare functional image
    try scfMRItb_04_unzipFile(base_dir, fname, ['_slice' num2str(i3) '_base_masked']); catch, end
    F = load_untouch_nii([base_dir fname '_slice' num2str(i3) '_base_masked.nii']); % junk load
    i4 = 0; % mean image
    i4val = i4;
    unix(['rm -f ' base_dir fname '_slice' num2str(i3) '_one_masked_' num2str(i4) '.nii']);
    F.img = F_oneslice .* F_weights;
    junk = sort(dezero(F.img));
    junk_index = floor(size(junk,1)*.95);
    norm_factor_funct = junk(junk_index,1);
    disp(sprintf(['Normalizing functional image by factor of: ' num2str((norm_factor_funct ./ 100),4)]));
    F.img = F.img ./ norm_factor_funct .* 100;
    save_untouch_nii(F, [base_dir fname '_slice' num2str(i3) '_one_masked_' num2str(i4) '.nii']);
    clear F
    
    % prepare anatomical image
    try scfMRItb_04_unzipFile(base_dir, fname_anat, ['_slice' num2str(i3)]); catch, end
    A = load_untouch_nii([base_dir fname_anat '_slice' num2str(i3) '.nii']); % junk load
    unix(['rm -f ' base_dir fname_anat '_slice' num2str(i3) '_' num2str(i4) '.nii']);
    try scfMRItb_04_unzipFile(base_dir, fname_anat, ['_slice' num2str(i3) '_affine_reg_mask']); catch, end
    A_base_mask = load_untouch_nii([base_dir fname_anat '_slice' num2str(i3) '_affine_reg_mask.nii']);
    A.img = single(A.img) .* (A_base_mask.img > 0);
    junk = sort(dezero(A.img));
    junk_index = floor(size(junk,1)*.95);
    norm_factor_anat = junk(junk_index,1);
    disp(sprintf(['Normalizing anatomical image by factor of: ' num2str((norm_factor_anat ./ 100),4)]));
    A.img = A.img ./ norm_factor_anat .* 100;
    save_untouch_nii(A, [base_dir fname_anat '_slice' num2str(i3) '_' num2str(i4) '.nii']);
    clear A

    median_translations = zeros(size(cost_fcns,2), 2);
    for i5 = 1 : size(cost_fcns,2)
    cost_fcn = cost_fcns{1,i5};
    
    all_translations = zeros(siz4max, 2);

    % (1) coarse search: max additional shift of {coarse} mm
    smax = smax1;
    unix(['rm -f ' base_dir fname '_slice' num2str(i3) '_base_warped_' num2str(i4val) '.nii']);
    % parfix 1 --> +ve = shift left
    % parfix 2 --> +ve = shift anterior
    % 
    % constrain each translation to be equal to the CoM shifts +/- smax mm
    % no rotation is allowed (because the spinal cord is kinda circular)
    % constrain L-R scaling to be +/- 1%
    % constrain A-P scaling to be +/- 2% (because this is the phase-encode
    %      direction and more stretching is expected in this direction)
    if use_cmass==0
    str = ['3dAllineate -VERB ' ...
        '-base ' base_dir fname_anat '_slice' num2str(i3) '_' num2str(i4) '.nii ' ...
        '-input ' base_dir fname '_slice' num2str(i3) '_one_masked_' num2str(i4val) '.nii ' ...
        '-prefix ' base_dir fname '_slice' num2str(i3) '_base_warped_' num2str(i4val) '.nii ' ...
        '-weight ' base_dir fname_anat '_slice' num2str(i3) '_affine_reg_mask.nii ' '-nomask ' ...
        '-interp quintic -final NN ' ...
        cost_fcn ...
        '-warp affine_general ' ...
        affine_parms ...
        '-parang 1 ' num2str(CoM_shifts(1,i3)-smax*0.75) ' ' num2str(CoM_shifts(1,i3)+smax*0.75) ' ' ...
        '-parang 2 ' num2str(CoM_shifts(2,i3)-smax) ' ' num2str(CoM_shifts(2,i3)+smax)];
    T = evalc_parfor(str);
    findme = '++ image warp: parameters = ';
    findme_loc = strfind(T, '++ image warp: parameters =');
    findme_loc = findme_loc(1,end); % in case there is more than one
    temp = T(1,findme_loc+size(findme,2):findme_loc+size(findme,2)+17);
    findspaces = strfind(temp, ' ');
    space1 = findspaces(1,1);
    space2 = findspaces(1,2);
    newstartingshift1 = str2num(temp(1,1:space1-1));
    newstartingshift2 = str2num(temp(1,space1+1:space2-1));
    if newstartingshift1==0 && newstartingshift2==0 % happens when the centers of mass are misaligned
        use_cmass = 1;
        clear newstartingshift* space* findspaces temp findme_loc findme T str
    end
    end

    if use_cmass==1
    str = ['3dAllineate -VERB ' ...
        '-base ' base_dir fname_anat '_slice' num2str(i3) '_' num2str(i4) '.nii ' ...
        '-input ' base_dir fname '_slice' num2str(i3) '_one_masked_' num2str(i4val) '.nii ' ...
        '-prefix ' base_dir fname '_slice' num2str(i3) '_base_warped_' num2str(i4val) '.nii ' ...
        '-weight ' base_dir fname_anat '_slice' num2str(i3) '_affine_reg_mask.nii ' '-nomask ' ...
        '-cmass z' ...
        '-interp quintic -final NN ' ...
        cost_fcn ...
        '-warp affine_general ' ...
        affine_parms ...
        '-parang 1 ' num2str(CoM_shifts(1,i3)-smax*0.75) ' ' num2str(CoM_shifts(1,i3)+smax*0.75) ' ' ...
        '-parang 2 ' num2str(CoM_shifts(2,i3)-smax) ' ' num2str(CoM_shifts(2,i3)+smax)];
    T = evalc_parfor(str);
    findme = '++ image warp: parameters = ';
    findme_loc = strfind(T, '++ image warp: parameters =');
    findme_loc = findme_loc(1,end); % in case there is more than one
    temp = T(1,findme_loc+size(findme,2):findme_loc+size(findme,2)+17);
    findspaces = strfind(temp, ' ');
    space1 = findspaces(1,1);
    space2 = findspaces(1,2);
    newstartingshift1 = str2num(temp(1,1:space1-1));
    newstartingshift2 = str2num(temp(1,space1+1:space2-1));        
    end
    
    % (2) refined search: max additional shift of {fine} mm
    smax = smax2;
    if (smax > 0) % skip fine search if smax <= 0 (may not be advisable?)
    unix(['rm -f ' base_dir fname '_slice' num2str(i3) '_base_warped_' num2str(i4val) '.nii']);

    if use_cmass==0
    str = ['3dAllineate -VERB ' ...
        '-base ' base_dir fname_anat '_slice' num2str(i3) '_' num2str(i4) '.nii ' ...
        '-input ' base_dir fname '_slice' num2str(i3) '_one_masked_' num2str(i4val) '.nii ' ...
        '-prefix ' base_dir fname '_slice' num2str(i3) '_base_warped_' num2str(i4val) '.nii ' ...
        '-weight ' base_dir fname_anat '_slice' num2str(i3) '_affine_reg_mask.nii ' '-nomask ' ...
        '-interp quintic -final NN ' ...
        cost_fcn ...
        '-warp affine_general ' ...
        affine_parms ...
        '-parang 1 ' num2str(newstartingshift1-smax*0.50) ' ' num2str(newstartingshift1+smax*0.50) ' ' ...
        '-parang 2 ' num2str(newstartingshift2-smax) ' ' num2str(newstartingshift2+smax)];
    else
    str = ['3dAllineate -VERB ' ...
        '-base ' base_dir fname_anat '_slice' num2str(i3) '_' num2str(i4) '.nii ' ...
        '-input ' base_dir fname '_slice' num2str(i3) '_one_masked_' num2str(i4val) '.nii ' ...
        '-prefix ' base_dir fname '_slice' num2str(i3) '_base_warped_' num2str(i4val) '.nii ' ...
        '-weight ' base_dir fname_anat '_slice' num2str(i3) '_affine_reg_mask.nii ' '-nomask ' ...
        '-cmass z' ...
        '-interp quintic -final NN ' ...
        cost_fcn ...
        '-warp affine_general ' ...
        affine_parms ...
        '-parang 1 ' num2str(newstartingshift1-smax*0.50) ' ' num2str(newstartingshift1+smax*0.50) ' ' ...
        '-parang 2 ' num2str(newstartingshift2-smax) ' ' num2str(newstartingshift2+smax)];
    end

    T = evalc_parfor(str);
    findme = '++ image warp: parameters = ';
    findme_loc = strfind(T, '++ image warp: parameters =');
    findme_loc = findme_loc(1,end); % in case there is more than one
    temp = T(1,findme_loc+size(findme,2):findme_loc+size(findme,2)+17);
    findspaces = strfind(temp, ' ');
    space1 = findspaces(1,1);
    space2 = findspaces(1,2);
    newstartingshift1 = str2num(temp(1,1:space1-1));
    newstartingshift2 = str2num(temp(1,space1+1:space2-1));
    end % smax2 (fine search)
    
    all_translations(1,:) = [newstartingshift1 newstartingshift2];
    all_translations = all_translations(1,:);
    
    % clean up
    unix(['rm -f ' base_dir fname '_slice' num2str(i3) '_base_warped_*.nii']);
    
    median_translations(i5,:) = median(all_translations,1);
%     median_translations(1:i5,:)
    end % i5 - all cost functions
    
%     median_translations
    final_reg(i3,:,:) = median_translations;
    
    % Save warping parameters for upcoming affine registrations
    unix(['rm -f ' base_dir fname '_slice' num2str(i3) '_base_warped.nii']);
    final_translations = median(median_translations,1);
    
    % display translations that will be used
    adjusted_final_translations = final_translations;
    adj_final_trans = adjusted_final_translations;
    clear abs_diff adjusted_final_translations;
    
    g1 = final_translations(1,1);
    g2 = final_translations(1,2);
    
    if use_cmass==0
    str = ['3dAllineate -VERB ' ...
        '-base ' base_dir fname_anat '_slice' num2str(i3) '_' num2str(i4) '.nii ' ...
        '-input ' base_dir fname '_slice' num2str(i3) '_base_masked.nii ' ...
        '-prefix ' base_dir fname '_slice' num2str(i3) '_base_warped.nii ' ...
        '-1Dmatrix_save ' base_dir fname '_slice' num2str(i3) '_xform.aff12.1D ' ...
        '-weight ' base_dir fname_anat '_slice' num2str(i3) '_affine_reg_mask.nii ' '-nomask ' ...
        '-nmi ' ...
        '-interp cubic -final NN ' ...
        '-warp affine_general ' ...
        affine_parms ...
        '-parfix 1 ' num2str(g1) ' ' ...
        '-parfix 2 ' num2str(g2)];
    else
    str = ['3dAllineate -VERB ' ...
        '-base ' base_dir fname_anat '_slice' num2str(i3) '_' num2str(i4) '.nii ' ...
        '-input ' base_dir fname '_slice' num2str(i3) '_base_masked.nii ' ...
        '-prefix ' base_dir fname '_slice' num2str(i3) '_base_warped.nii ' ...
        '-1Dmatrix_save ' base_dir fname '_slice' num2str(i3) '_xform.aff12.1D ' ...
        '-weight ' base_dir fname_anat '_slice' num2str(i3) '_affine_reg_mask.nii ' '-nomask ' ...
        '-cmass z' ...
        '-nmi ' ...
        '-interp cubic -final NN ' ...
        '-warp affine_general ' ...
        affine_parms ...
        '-parfix 1 ' num2str(g1) ' ' ...
        '-parfix 2 ' num2str(g2)];
    end
    T = evalc('unix(str)');
    clear T findme* temp findspaces space1 space2;
    
    clear newstartingshift* nss1 nss2 smax r_min c_min cost_fcn;
    
    % clean up
    unix(['rm -f ' base_dir fname '_slice' num2str(i3) '_one_masked_*.nii']);
    
end % i3

end
```
