## Supplementary material for "Neptune: a toolbox for spinal cord functional MRI data processing and quality assurance": Neptune software: spinal_cord_registration_T2s.m

```
function final_reg = spinal_cord_registration_T2s(base_dir, fname_anat, fname_T2s, i3T2s, g)

% Updated:      2015-07-12 - Modified for T2* mapping
% Previous:     2015-01-19 - Added boundary (outside CSF) constraint

siz3 = 12; % but only 1 slice is being registered
siz4 = 1; % 1 dynamic (echo #2)

% no rotation, no shearing
affine_parms = '-parfix 3 0 -parfix 4 0 -parfix 5 0 -parfix 6 0 -parfix 9 1 -parfix 10 0 -parfix 11 0 -parfix 12 0 -onepass -nmatch 100% -parang 7 1/1.01 1.01 -parang 8 1/1.02 1.02 ';

% maximum translations for coarse and fine passes
smax_vec = [2.5 1.0];
smax1 = smax_vec(1,1); % define separately for parfor loop
smax2 = smax_vec(1,2);

siz4max = min(siz4, 50); % max number of dynamics used to estimate translations
siz4vec = 1:floor(siz4/siz4max):siz4;
siz4vec = siz4vec(1,1:siz4max);

cost_fcns = {'-ls ' '-sp '}; % This version of the function requires these
                             % values to remain fixed and in this order.

final_reg = zeros(siz3, size(cost_fcns,2), 2);
CoM_shifts = zeros(2,siz3);

% ----- SLICE-BY-SLICE AFFINE REGISTRATION OF BASE ------------------------

for i3 = i3T2s : i3T2s

    median_translations = zeros(size(cost_fcns,2), 2);
    for i5 = 1 : size(cost_fcns,2)
    cost_fcn = cost_fcns{1,i5};
    
    all_translations = zeros(siz4max, 2);
    for i4 = 1 : siz4max
    i4val = siz4vec(1,i4);

    % (1) coarse search: max additional shift of {coarse} mm
    smax = smax1;
    unix(['rm -f ' base_dir fname_T2s '_warped' '.nii']);
    % parfix 1 --> +ve = shift left
    % parfix 2 --> +ve = shift anterior
    % 
    % constrain each translation to be equal to the CoM shifts +/- smax mm
    % no rotation is allowed (because the spinal cord is kinda circular)
    % constrain L-R scaling to be +/- 1%
    % constrain A-P scaling to be +/- 2% (because this is the phase-encode
    %      direction and more stretching is expected in this direction)
    str = ['3dAllineate -VERB ' ...
        '-base ' base_dir fname_anat '_slice' num2str(i3) '.nii ' ...
        '-input ' base_dir fname_T2s '_masked' '.nii ' ...
        '-prefix ' base_dir fname_T2s '_warped' '.nii ' ...
        '-weight ' base_dir fname_anat '_slice' num2str(i3) '_affine_reg_mask.nii ' '-nomask ' ...
        '-interp quintic -final NN ' ...
        cost_fcn ...
        '-warp affine_general ' ...
        affine_parms ...
        '-parang 1 ' num2str(CoM_shifts(1,i3)-smax*0.75) ' ' num2str(CoM_shifts(1,i3)*0.75) ' ' ...
        '-parang 2 ' num2str(CoM_shifts(2,i3)-smax) ' ' num2str(CoM_shifts(2,i3)+smax)];
    T = evalc_parfor(str);
    findme = '++ image warp: parameters = ';
    findme_loc = strfind(T, '++ image warp: parameters =');
    findme_loc = findme_loc(1,end); % in case there is more than one
    temp = T(1,findme_loc+size(findme,2):findme_loc+size(findme,2)+17);
    findspaces = strfind(temp, ' ');
    space1 = findspaces(1,1);
    space2 = findspaces(1,2);
    newstartingshift1 = str2num(temp(1,1:space1-1));
    newstartingshift2 = str2num(temp(1,space1+1:space2-1));
    
    % (2) refined search: max additional shift of {fine} mm
    smax = smax2;
    if (smax > 0) % skip fine search if smax <= 0 (may not be advisable?)
    unix(['rm -f ' base_dir fname_T2s '_warped' '.nii']);
    str = ['3dAllineate -VERB ' ...
        '-base ' base_dir fname_anat '_slice' num2str(i3) '.nii ' ...
        '-input ' base_dir fname_T2s '_masked' '.nii ' ...
        '-prefix ' base_dir fname_T2s '_warped' '.nii ' ...
        '-weight ' base_dir fname_anat '_slice' num2str(i3) '_affine_reg_mask.nii ' '-nomask ' ...
        '-interp quintic -final NN ' ...
        cost_fcn ...
        '-warp affine_general ' ...
        affine_parms ...
        '-parang 1 ' num2str(newstartingshift1-smax*0.50) ' ' num2str(newstartingshift1+smax*0.50) ' ' ...
        '-parang 2 ' num2str(newstartingshift2-smax) ' ' num2str(newstartingshift2+smax)];
    T = evalc_parfor(str);
    findme = '++ image warp: parameters = ';
    findme_loc = strfind(T, '++ image warp: parameters =');
    findme_loc = findme_loc(1,end); % in case there is more than one
    temp = T(1,findme_loc+size(findme,2):findme_loc+size(findme,2)+17);
    findspaces = strfind(temp, ' ');
    space1 = findspaces(1,1);
    space2 = findspaces(1,2);
    newstartingshift1 = str2num(temp(1,1:space1-1));
    newstartingshift2 = str2num(temp(1,space1+1:space2-1));
    end % smax2 (fine search)
    
    all_translations(i4,:) = [newstartingshift1 newstartingshift2];
    
    end % for i4 - all dynamics
    
    median_translations(i5,:) = median(all_translations,1);
%     median_translations(1:i5,:)
    end % i5 - all cost functions
    
%     median_translations
    final_reg(i3,:,:) = median_translations;
    
    % Save warping parameters for upcoming affine registrations
    unix(['rm -f ' base_dir fname_T2s '_warped' '.nii']);
    final_translations = median(median_translations,1)
    
    % The previous version of this function only had the following line:
    % 
    % final_translations = median(median_translations,1)
    % 
    % But -sp sometimes give unreliable translations if the fMRI data
    % quality are poor. -ls seems to work well most of the time, but it
    % would be nice to protect against the possibility that -ls produces
    % less-than-optimal results.
    % 
    % Therefore, let us propose an empirical compromise.
    % 
    % If abs(difference) (= abs_diff) between -ls and -sp is:
    %      abs_diff < 0.5 mm,        use the median of -ls and -sp
    %      0.5 <= abs_diff < 1.0 mm, use 2/3*ls + 1/3*sp
    %      1.0 <= abs_diff < 1.5 mm, use 3/4*ls + 1/4*sp
    %      abs_diff >= 1.5 mm,       use 9/10*ls + 1/10*sp
    
    % translation left-right
    abs_diff = abs(median_translations(1,1)-median_translations(2,1));
    if     (0.5 <= abs_diff) && (abs_diff < 1.0)
        final_translations(1,1) = 2/3*median_translations(1,1) + 1/3*median_translations(2,1);
    elseif (1.0 <= abs_diff) && (abs_diff < 1.5)
        final_translations(1,1) = 3/4*median_translations(1,1) + 1/4*median_translations(2,1);
    elseif (1.5 <= abs_diff)
        final_translations(1,1) = 9/10*median_translations(1,1) + 1/10*median_translations(2,1);
    end
        
    % translation anterior-posterior
    abs_diff = abs(median_translations(1,2)-median_translations(2,2));
    if     (0.5 <= abs_diff) && (abs_diff < 1.0)
        final_translations(1,2) = 2/3*median_translations(1,2) + 1/3*median_translations(2,2);
    elseif (1.0 <= abs_diff) && (abs_diff < 1.5)
        final_translations(1,2) = 3/4*median_translations(1,2) + 1/4*median_translations(2,2);
    elseif (1.5 <= abs_diff)
        final_translations(1,2) = 9/10*median_translations(1,2) + 1/10*median_translations(2,2);
    end
    % display translations that will be used
    adjusted_final_translations = final_translations;
    clear abs_diff adjusted_final_translations;
    
    % override registration if necessary
    if nargin == 5
        g1 = g(1,1);
        g2 = g(1,2);
        overridden_final_translations = g;
    else % nargin == 4
        g1 = final_translations(1,1);
        g2 = final_translations(1,2);
    end
    
    str = ['3dAllineate -VERB ' ...
        '-base ' base_dir fname_anat '_slice' num2str(i3) '.nii ' ...
        '-input ' base_dir fname_T2s '_masked' '.nii ' ...
        '-prefix ' base_dir fname_T2s '_warped' '.nii ' ...
        '-1Dmatrix_save ' base_dir fname_T2s '_xform.aff12.1D ' ...
        '-weight ' base_dir fname_anat '_slice' num2str(i3) '_affine_reg_mask.nii ' '-nomask ' ...
        '-nmi ' ...
        '-interp cubic -final NN ' ...
        '-warp affine_general ' ...
        affine_parms ...
        '-parfix 1 ' num2str(g1) ' ' ...
        '-parfix 2 ' num2str(g2)];
    T = evalc('unix(str)');
    clear T findme* temp findspaces space1 space2;
    
    clear newstartingshift* nss1 nss2 smax r_min c_min cost_fcn;
    
end % i3
```
