## Supplementary material for "Neptune: a toolbox for spinal cord functional MRI data processing and quality assurance": Neptune software: subplot2n.m

```
function theAxis = subplot2n(varargin)
    %SUBPLOT Create axes in tiled positions.
    %   H = SUBPLOT(m,n,p), or SUBPLOT(mnp), breaks the Figure window
    %   into an m-by-n matrix of small axes, selects the p-th axes for
    %   the current plot, and returns the axes handle.  The axes are
    %   counted along the top row of the Figure window, then the second
    %   row, etc.  For example,
    %
    %       SUBPLOT(2,1,1), PLOT(income)
    %       SUBPLOT(2,1,2), PLOT(outgo)
    %
    %   plots income on the top half of the window and outgo on the
    %   bottom half. If the CurrentAxes is nested in a uipanel the
    %   panel is used as the parent for the subplot instead of the
    %   current figure.
    %
    %   SUBPLOT(m,n,p), if the axes already exists, makes it current.
    %   SUBPLOT(m,n,p,'replace'), if the axes already exists, deletes it and
    %   creates a new axes.
    %   SUBPLOT(m,n,p,'align') places the axes so that the plot boxes
    %   are aligned, but does not prevent the labels and ticks from
    %   overlapping.
    %   SUBPLOT(m,n,P), where P is a vector, specifies an axes position
    %   that covers all the subplot positions listed in P.
    %   SUBPLOT(H), where H is an axes handle, is another way of making
    %   an axes current for subsequent plotting commands.
    %
    %   SUBPLOT('position',[left bottom width height]) creates an
    %   axes at the specified position in normalized coordinates (in
    %   in the range from 0.0 to 1.0).
    %
    %   SUBPLOT(..., PROP1, VALUE1, PROP2, VALUE2, ...) sets the
    %   specified property-value pairs on the subplot axes. To add the
    %   subplot to a specific figure pass the figure handle as the
    %   value for the 'Parent' property.
    %
    %   If a SUBPLOT specification causes a new axes to overlap an
    %   existing axes, the existing axes is deleted - unless the position
    %   of the new and existing axes are identical.  For example,
    %   the statement SUBPLOT(1,2,1) deletes all existing axes overlapping
    %   the left side of the Figure window and creates a new axes on that
    %   side - unless there is an axes there with a position that exactly
    %   matches the position of the new axes (and 'replace' was not specified),
    %   in which case all other overlapping axes will be deleted and the
    %   matching axes will become the current axes.
    %
    %   SUBPLOT(111) is an exception to the rules above, and is not
    %   identical in behavior to SUBPLOT(1,1,1).  For reasons of backwards
    %   compatibility, it is a special case of subplot which does not
    %   immediately create an axes, but instead sets up the figure so that
    %   the next graphics command executes CLF RESET in the figure
    %   (deleting all children of the figure), and creates a new axes in
    %   the default position.  This syntax does not return a handle, so it
    %   is an error to specify a return argument.  The delayed CLF RESET
    %   is accomplished by setting the figure's NextPlot to 'replace'.
    %
    %   Be aware when creating subplots from scripts that the Position
    %   property of subplots is not finalized until either a drawnow
    %   command is issued, or MATLAB returns to await a user command.
    %   That is, the value obtained for subplot i by the command
    %   h(i).Position will not be correct until the script
    %   refreshes the plot or exits.
    %
    %   See also  GCA, GCF, AXES, FIGURE, UIPANEL    

    %   SUBPLOT(m,n,p,H) when H is an axes will move H to the specified
    %   position.
    %   SUBPLOT(m,n,p,H,PROP1,VALUE1,...) will move H and apply the
    %   specified property-value pairs
    %
    %   SUBPLOT(m,n,p) for non-integer p places the subplot at the
    %   fraction p-floor(p) between the positions floor(p) and ceil(p)
    
    %   Copyright 1984-2019 The MathWorks, Inc.
    
    % Separate out name/value pairs and string arguments from other arguments.
    % This will also convert a '222' first input into [2,2,2].
    [args,pvpairs,narg] = subplot2n_parseargs(varargin);
    
    % Check whether we should ignore a possible 'v6' argument.
    if ~isempty(pvpairs) && strcmpi(pvpairs{1}, 'v6')
        filename = 'subplot';
        warning(['MATLAB:', filename, ':IgnoringV6Argument'],...
            getString(message('MATLAB:usev6plotapi:IgnoringV6ArgumentForFilename', upper(filename))));
        pvpairs(1) = [];
    end
    
    
    % we will kill all overlapping axes siblings if we encounter the mnp
    % or m,n,p specifier (excluding '111').
    % But if we get the 'position' or H specifier, we won't check for and
    % delete overlapping siblings:
    killSiblings = 0;
    createAxis = true;
    moveAxis = false;
    delayDestroy = false;
    useAutoLayout = true;
    tol = sqrt(eps);
    parent = handle(get(0, 'CurrentFigure'));
    ancestorFigure = parent;
    if ~isempty(parent) && ~isempty(parent.CurrentAxes)
        parent = parent.CurrentAxes.Parent;
        ancestorFigure = parent;
        if ~strcmp(ancestorFigure.Type, 'figure')
            ancestorFigure = ancestor(parent, 'figure');
        end
    end
    preventMove = false;

    % This is the percent offset from the subplot grid of the plotbox.
    inset = [.2, .18, .04, .1]; % [left bottom right top]

    %check for encoded format
    h = [];
    position = [];
    explicitParent = false;
    explicitPosition = false;
    nRows = [];
    
    if narg == 0
        % The argument could be either:
        % 1) subplot()
        % 2) subplot('Position',positionVector)
        if isempty(pvpairs)
            % subplot()
            % make compatible with 3.5, i.e. subplot == subplot(111)
            args{1} = 111;
            narg = 1;
        elseif strcmpi(pvpairs{1}, 'position')
            % subplot('Position',positionVector)
            if numel(pvpairs)>=2
                pos_size = size(pvpairs{2});
                if (pos_size(1) * pos_size(2) == 4)
                    position = pvpairs{2};
                    explicitPosition = true;
                else
                    error(message('MATLAB:subplot:InvalidPositionParameter'))
                end
            else
                error(message('MATLAB:subplot:InvalidPositionParameter'))
            end
            killSiblings = 1; % Kill overlaps here also.
            useAutoLayout = false;
            pvpairs(1:2) = [];
        else
            error(message('MATLAB:subplot:UnknownOption'))
        end
    end
    
    if narg == 1
        % The argument could be one of 2 things:
        % 1) a 3-digit number 100 < num < 1000, of the format mnp
        % 2) an axes handle
        arg = args{1};
        
        % Check whether arg is a handle to an Axes/Chart.
        if isempty(arg)
            error(message('MATLAB:subplot:UnknownOption'))
        elseif isSubplotCandidate(arg)
            h = handle(arg);
            if isa(h, 'matlab.graphics.chart.Chart') ...
                    && ~isprop(h,'InnerPosition') ...
                    && ~isappdata(h,'SubplotPosition')
                error(message('MATLAB:subplot:UnsupportedSyntaxWithChart',h.Type));
            end
            
            createAxis = false;
        elseif isa(arg,'matlab.ui.control.UIAxes')
            % Error if given uiaxes
            try
                restrictedAccess = feature('WebGraphicsRestriction');
            catch
                restrictedAccess = 1;
            end
            if restrictedAccess
                error(message('MATLAB:ui:uiaxes:general'));
            end
        elseif isnumeric(arg)
            % Check for NaN and Inf.
            if (~isfinite(arg))
                error(message('MATLAB:subplot:SubplotIndexNonFinite'))
            end
            
            % Check for input out of range
            if (arg <= 100 || arg >= 1000)
                error(message('MATLAB:subplot:SubplotIndexOutOfRange'))
            end
            
            plotId = rem(arg, 10);
            nCols = rem(fix(arg - plotId) / 10, 10);
            nRows = fix(arg / 100);
            if nRows * nCols < plotId
                error(message('MATLAB:subplot:SubplotIndexTooLarge'));
            end
            killSiblings = 1;
            if (arg == 111)
                createAxis = false;
                delayDestroy = true;
                if nargout > 0
                    error(message('MATLAB:subplot:TooManyOutputs'))
                end
            else
                createAxis = true;
                delayDestroy = false;
            end
        else
            error(message('MATLAB:subplot:InvalidAxesHandle'))           
        end
        
    elseif narg == 2
        % passed in subplot(a,b) where a/b are not a name/value pair
        error(message('MATLAB:subplot:InvalidSyntax'))
    
    elseif narg == 3
        % passed in subplot(m,n,p)
        nRows = args{1};
        nCols = args{2};
        plotId = args{3};
        
        % we should kill overlaps here too:
        killSiblings = 1;
        
    elseif narg == 4
        % passed in subplot(m,n,p,ax)
        
        nRows = args{1};
        nCols = args{2};
        plotId = args{3};

        arg = args{4};
        if isempty(arg)
            error(message('MATLAB:subplot:InvalidAxesHandle'))
        else
            h = handle(arg);
            if isa(h,'matlab.ui.control.UIAxes')
                % Error if given uiaxes
                error(message('MATLAB:ui:uiaxes:general'));
            elseif isa(h, 'matlab.graphics.chart.Chart') ...
                    && ~isObjectAlreadyInCell(args{1},args{2},args{3},h)...                    
                    && ~(isa(h,'matlab.graphics.chart.internal.SubplotPositionableChart')...
                    || isa(h,'matlab.graphics.chart.internal.PositionableChartWithAxes'))
                error(message('MATLAB:subplot:UnsupportedSyntaxWithChart',h.Type));
            elseif ~isSubplotCandidate(h)
                error(message('MATLAB:subplot:InvalidAxesHandle'))
            end
            
            parent = h.Parent;
            ancestorFigure = ancestor(h, 'figure');
            
            % If the parent is passed in explicitly, don't create a new figure
            % when the "NextPlot" property is set to "new" in the figure.
            explicitParent = true;
            ancestorFigure.CurrentAxes = h;
            moveAxis = true;
            createAxis = false;
            
            if ~isempty(pvpairs) && strcmpi(pvpairs{1}, 'PreventMove')
                preventMove = true;
                pvpairs{1} = [];
            end
        end
        
    elseif narg > 4
        % String inputs have already been removed
        % so any other number of non-string inputs is invalid syntax.
        error(message('MATLAB:subplot:UnknownOption'))
        
    end
    
    % Check for 'replace' or 'align' in the properties.
    if ~isempty(pvpairs)
        arg = pvpairs{1};
        if strncmpi(arg, 'replace', 1)
            % passed in subplot(m,n,p,'replace')
            killSiblings = 2; % kill nomatter what
            pvpairs(1) = [];
        elseif strcmpi(arg, 'align')
            % passed in subplot(m,n,p,'align')
            % since obeying position will remove the axes from the grid just set
            % useAutoLayout to false to skip adding it to the grid to start with
            useAutoLayout = false;
            killSiblings = 1; % kill if it overlaps stuff
            pvpairs(1) = [];
        end
    end
    
    % Find and remove any 'Parent' property passed in as a name/value pair.
    par = 2*find(strncmpi('Parent', pvpairs(1 : 2 : end), 6));
    if any(par)
        % If the parent is passed in explicitly, don't create a new figure
        % when the "NextPlot" property is set to "new" in the figure.
        explicitParent = true;
        parent = handle(pvpairs{par(end)});
        ancestorFigure = ancestor(parent, 'figure');
        pvpairs([par-1 par]) = [];
    end
    
    % Warn for n-v pairs that shouldn't be set on Axes in a subplot.
    % Also warns on substrings of these properties
    unsupportedArgs = {'Position','OuterPosition','ActivePositionProperty'};
    for prop = pvpairs(1 : 2 : end)
        argmatches = strncmpi(unsupportedArgs, prop{:}, length(prop{:}));
        if(argmatches(1))
            warning(message('MATLAB:subplot:InvalidPositionSyntax'));
        elseif any(argmatches(2:end))            
            warning(message('MATLAB:subplot:InvalidNVPair', ...
                unsupportedArgs{find(argmatches,1)}));
        end
    end
    
    % if we recovered an identifier earlier, use it:
    if ~isempty(h) && ~moveAxis
        % Syntax: subplot(ax)
        parent = h.Parent;
        ancestorFigure = ancestor(h, 'figure');
        ancestorFigure.CurrentAxes = h;
    else  % if we haven't recovered position yet, generate it from mnp info:
        if isempty(parent)
            parent = gcf;
            ancestorFigure = parent;
        end
        
        % Error if AutoResizeChildren is 'on'
        if isprop(parent,'AutoResizeChildren') && strcmp(parent.AutoResizeChildren,'on')
            error(message('MATLAB:subplot:AutoResizeChildren'))
        end

        if isempty(position)
            if min(plotId) < 1
                error(message('MATLAB:subplot:SubplotIndexTooSmall'))
            elseif max(plotId) > nCols * nRows
                error(message('MATLAB:subplot:SubplotIndexTooLarge'));
            else
                
                row = (nRows - 1) - fix((plotId - 1) / nCols);
                col = rem(plotId - 1, nCols);
                
                % get default axes position in normalized units
                % If we have checked this quantity once, cache it.
                if ~isappdata(ancestorFigure, 'SubplotDefaultAxesLocation')
                    if ~strcmp(get(ancestorFigure, 'DefaultAxesUnits'), 'normalized')
                        tmp = axes('Parent',ancestorFigure);
                        tmp.Units = 'normalized';
                        def_pos = tmp.InnerPosition;
                        delete(tmp)
                    else
                        def_pos = get(ancestorFigure, 'DefaultAxesPosition');
                    end
                    setappdata(ancestorFigure, 'SubplotDefaultAxesLocation', def_pos);
                    if(parent ~= ancestorFigure)
                        setappdata(parent, 'SubplotDefaultAxesLocation', def_pos);
                    end
                else
                    def_pos = getappdata(ancestorFigure, 'SubplotDefaultAxesLocation');
                end
                
                % compute outerposition and insets relative to figure bounds
                rw = max(row) - min(row) + 1;
                cw = max(col) - min(col) + 1;
                width = def_pos(3) / (nCols - inset(1) - inset(3));
                height = def_pos(4) / (nRows - inset(2) - inset(4));
                inset = inset .* [width, height, width, height];
                outerpos = [def_pos(1) + min(col) * width - inset(1), ...
                    def_pos(2) + min(row) * height - inset(2), ...
                    width * cw, height * rw];
                
                % adjust outerpos and insets for axes around the outside edges
                if min(col) == 0
                    inset(1) = def_pos(1);
                    outerpos(3) = outerpos(1) + outerpos(3);
                    outerpos(1) = 0;
                end
                if min(row) == 0
                    inset(2) = def_pos(2);
                    outerpos(4) = outerpos(2) + outerpos(4);
                    outerpos(2) = 0;
                end
                if max(col) == nCols - 1
                    inset(3) = max(0, 1 - def_pos(1) - def_pos(3));
                    outerpos(3) = 1 - outerpos(1);
                end
                if max(row) == nRows - 1
                    inset(4) = max(0, 1 - def_pos(2) - def_pos(4));
                    outerpos(4) = 1 - outerpos(2);
                end
                
                % compute inner position
                position = [outerpos(1 : 2) + inset(1 : 2), ...
                    outerpos(3 : 4) - inset(1 : 2) - inset(3 : 4)];
                
            end
        end
    end
    
    % kill overlapping siblings if mnp specifier was used:
    nextstate = ancestorFigure.NextPlot;
    
    if strncmp(nextstate, 'replace', 7)
        nextstate = 'add';
    elseif strncmp(nextstate, 'new', 3)
        killSiblings = 0;
    end
    
    if killSiblings
        if delayDestroy
            ancestorFigure.NextPlot = 'replace';
            return
        end
        hasDeleted = false;
        sibs = datasiblings(parent);
        newcurrent = [];
        for i = 1 : length(sibs)
            % Be aware that handles in this list might be destroyed before
            % we get to them, because of other objects' DeleteFcn callbacks...
            if isSubplotCandidate(sibs(i))
                units = sibs(i).Units;
                
                if isa(sibs(i),'matlab.graphics.chart.Chart') ...
                        && ~(isa(sibs(i),'matlab.graphics.chart.internal.SubplotPositionableChart') || ...
                             isa(sibs(i),'matlab.graphics.chart.internal.PositionableChartWithAxes'))
                    % when existing chart replaced a subplot, use
                    % inner position of the axes it replaced when
                    % checking for matching/overlapping.
                    if isappdata(sibs(i),'SubplotPosition')
                        sibpos = getappdata(sibs(i),'SubplotPosition');
                        if length(sibpos) < 4
                            sibpos = sibs(i).OuterPosition;
                        end
                    else                       
                        if all(sibs(i).OuterPosition == [0,0,1,1]) && ...
                                ~explicitPosition && all(nRows == 1) && all(nCols == 1)
                            % special case to make subplot(1,1,1) select 
                            % (and not clobber) a [0,0,1,1] full-container chart. 
                            % Even though the requested subplot's position
                            % doesn't match the existing Chart's 
                            % OuterPosition, pretend the existing
                            % chart was positioned at the exact location 
                            % of the requested subplot
                            sibpos = position; 
                        else
                            % otherwise, compare against the chart's
                            % OuterPosition when deciding if we should
                            % clobber existing chart with the incoming subplot
                            sibpos = sibs(i).OuterPosition;

                        end
                    end
                                        
                else
                    sibpos = sibs(i).InnerPosition;
                end
                % If a legend or colorbar has resized the axes, use the original axes
                % position as the "Position" property:
                if ~explicitPosition
                    if isappdata(sibs(i), 'LegendColorbarExpectedPosition') && ...
                            isequal(getappdata(sibs(i), 'LegendColorbarExpectedPosition'), get(sibs(i), 'Position'))
                        sibinset = getappdata(sibs(i), 'LegendColorbarOriginalInset');
                        if isempty(sibinset)
                            % during load the appdata might not be present
                            sibinset = get(sibs(i).Parent, 'DefaultAxesLooseInset');
                        end
                        sibinset = offsetsInUnits(sibs(i), sibinset, 'normalized', get(sibs(i), 'Units'));
                        if strcmpi(sibs(i).ActivePositionProperty, 'position')
                            pos = sibs(i).Position;
                            loose = sibs(i).LooseInset;
                            opos = getOuterFromPosAndLoose(pos, loose, get(sibs(i), 'Units'));
                            if strcmp(sibs(i).Units, 'normalized')
                                sibinset = [opos(3 : 4), opos(3 : 4)] .* sibinset;
                            end
                            sibpos = [opos(1 : 2) + sibinset(1 : 2), opos(3 : 4) - sibinset(1 : 2) - sibinset(3 : 4)];
                        end
                    end
                end
                if ~strcmp(units, 'normalized')
                    sibpos = hgconvertunits(ancestorFigure, sibpos, units, 'normalized', parent);
                end
                intersect = 1;
                if ((position(1) >= sibpos(1) + sibpos(3) - tol) || ...
                        (sibpos(1) >= position(1) + position(3) - tol) || ...
                        (position(2) >= sibpos(2) + sibpos(4) - tol) || ...
                        (sibpos(2) >= position(2) + position(4) - tol))
                    intersect = 0;
                end
                if intersect
                    % position is the proposed position of an axes, and
                    % sibpos is the current position of an existing axes.
                    % Since the bounding boxes of position and sibpos overlap,
                    % we must determine whether to delete the sibling sibs(i)
                    % whose normalized position is sibpos.
                    
                    % First of all, we check whether we must kill the sibling
                    % "no matter what."
                    if (killSiblings == 2)
                        if ~hasDeleted
                            hasDeleted = true;
                            % Notify the editor that an axes is being deleted
                            matlab.graphics.internal.clearNotify(ancestorFigure, 'delete');
                        end
                            
                        delete(sibs(i));
                        
                        % If the proposed and existing axes overlap exactly, we do
                        % not kill the sibling.  Rather we shall ensure later that
                        % this sibling axes is set as the 'CurrentAxes' of its
                        % ancestorFigure.
                        
                        % Next we check for a partial overlap.
                    elseif (any(abs(sibpos - position) > tol))
                        % The proposed and existing axes partially overlap.
                        % Since the proposed and existing axes could each be
                        % "grid-generated" or "explicitly-specified", we must
                        % consider four possibilities for the overlap of
                        % "proposed" vs. "existing", i.e.
                        % (1) "grid-generated" vs. "grid-generated"
                        % (2) "grid-generated" vs. "explicitly-specified"
                        % (3) "explicitly-specified" vs. "grid-generated"
                        % (4) "explicitly-specified" vs. "explicitly-specified"
                        
                        % If the position of the proposed axes is
                        % "explicitly-specified", then the only condition that
                        % avoids killing the sibling is an exact overlap.
                        % However, we know that the overlap is partial.
                        if (explicitPosition)
                            if ~hasDeleted
                                hasDeleted = true;
                                % Notify the editor that an axes is being deleted
                                matlab.graphics.internal.clearNotify(ancestorFigure, 'delete');
                            end
                            delete(sibs(i));
                        else
                            % We know that the position of the proposed axes is
                            % "grid-generated".
                            
                            grid = getappdata(parent, 'SubplotGrid');
                            % The SubplotGrid maintains an array of axes
                            % handles, one per grid location.  Axes that span
                            % multiple grid locations do not store handles in
                            % the SubplotGrid.
                            
                            if isempty(grid) || ~any(grid(:) == sibs(i)) || ...
                                    size(grid, 1) ~= nRows || size(grid, 2) ~= nCols || ...
                                    ~isscalar(row) || ~isscalar(col)
                                % If the sibling cannot be found in the grid, we
                                % kill the sibling.  Otherwise, the proposed and
                                % existing axes are "grid-generated".  If we
                                % are changing the size of the grid, we kill
                                % the sibling.  Otherwise, "plotId" may be a
                                % vector of multiple grid locations, which
                                % causes a partial overlap between the proposed
                                % and existing axes, so we kill the sibling.
                                
                                % This check recognizes that there may be
                                % labels, colorbars, legends, etc. attached to
                                % the existing axes that have affected its
                                % position.  In such a case, we do not kill the
                                % sibling.
                                if ~hasDeleted
                                    hasDeleted = true;
                                    % Notify the editor that an axes is being deleted
                                    matlab.graphics.internal.clearNotify(ancestorFigure, 'delete');
                                end
                                delete(sibs(i));
                            end
                        end
                    end
                    % if this axes overlaps the other one exactly then
                    if ~isempty(newcurrent) && isgraphics(newcurrent)
                        delete(newcurrent);
                    end
                    newcurrent = sibs(i);
                    if ~isempty(pvpairs) && isvalid(newcurrent)
                        set(newcurrent, pvpairs{:});
                    end
                end
            end
        end
        if ~isempty(newcurrent) && isgraphics(newcurrent)
            ancestorFigure.CurrentAxes = newcurrent;
            createAxis = false;
            matlab.graphics.internal.markFigure(newcurrent);
        end
        ancestorFigure.NextPlot = nextstate;
    end
    
    if isa(parent,'matlab.graphics.layout.Layout') 
        lay=parent;
        parent=lay.Parent;
        delete(lay)
    elseif isa(parent,'matlab.graphics.Graphics') && ...
                isa(parent,'matlab.ui.container.CanvasContainer') && ...
                isvalid(parent)
        objs = findall(parent, '-depth', 1, {'-isa','matlab.graphics.layout.Layout'});
        delete(objs);
    end
    
    % create the axes:
    if createAxis
        if strcmp(nextstate, 'new') && ~explicitParent
            parent = figure;
            ancestorFigure = parent;
        end
        ax = axes('Units', 'normalized', 'Position', position, ...
            'LooseInset', inset, 'Parent', parent);
        % TODO: Get axes to accept position args on command line
        ax.Units = get(ancestorFigure, 'DefaultAxesUnits');
        if useAutoLayout
            addAxesToGrid(ax, nRows, nCols, row, col, position, plotId);
        else
            addNonGridAxes(ax,position);
        end
        if ~isempty(pvpairs)
            set(ax, pvpairs{:});
        end
    elseif moveAxis && ~preventMove
        
        % moving an existing axes/chart into a subplot layout.
        
        %first, remove h from any pre-existing subplot
        matlab.graphics.internal.removeAxesFromGrid(h.Parent, h);
        ax = h;
        units = h.Units;

        % Some Charts position by outer position only & don't support
        % insets. Don't modify position, just use the position of the
        % outgoing subplot, as set by swapaxes.
        if ~isa(h,'matlab.graphics.chart.Chart') %case for axes/polaraxes         
            set(h, 'Units', 'normalized', 'InnerPosition', position, ...
                'LooseInset', inset, 'Parent', parent);
        elseif isa(h,'matlab.graphics.chart.internal.SubplotPositionableChart') || ...
               isa(h,'matlab.graphics.chart.internal.PositionableChartWithAxes') 
            outerposition = [position(1:2) - inset(1:2), position(3:4) + inset(1:2) + inset(3:4)];
            set(h, 'Units', 'normalized', 'InnerPosition', position, ...
                'MaxInsetForSubplotCell', inset, ...
            'SubplotCellOuterPosition', outerposition, 'Parent', parent);        
        else %orange chart that has been positioned in grid by swapaxes
            set(h,'Parent',parent);
        end
        
            
        
        h.Units = units;
        if ~isempty(pvpairs)
            set(h, pvpairs{:});
        end
        if useAutoLayout
            addAxesToGrid(ax, nRows, nCols, row, col, position, plotId);
        else
            addNonGridAxes(ax,position);
        end
            
    else
        % this should only happen with subplot(H)
        ax = ancestorFigure.CurrentAxes;
    end
    % return identifier, if requested:
    if(nargout > 0)
        theAxis = ax;
    end
    
end


% Add ax to a matrix of handles in the specified location.
% The grid is stored on the parent appdata.
% Also store the insets in ax appdata.
% Only stores the axes if it is in a 1-by-1 cell and
% the grid size matches any existing grid.
function addAxesToGrid(ax, nRows, nCols, row, col, position, plotId)
    p = ax.Parent;
    grid = getappdata(p, 'SubplotGrid');
    if isempty(grid)
        grid = gobjects(nRows,nCols);
    end
    
    % add SubplotListenersManager to p
    if ~isappdata(p,'SubplotListenersManager')
        lm =  matlab.graphics.internal.SubplotListenersManager(nRows*nCols);
        % create an empty filed so that other tests wont complain
        setappdata(p,'SubplotListeners',[]);
    else
        lm=getappdata(p,'SubplotListenersManager');
    end
    lm.addToListeners(ax,[]);
    setappdata(p,'SubplotListenersManager',lm);
    
    % add SubplotDeleteListenersManager to axes
    if ~isappdata(ax,'SubplotDeleteListenersManager')
        dlm =  matlab.graphics.internal.SubplotDeleteListenersManager();
        dlm.addToListeners(ax);
        setappdata(ax,'SubplotDeleteListenersManager',dlm); 
    end
    
    setappdata(ax,'SubplotGridLocation',{nRows,nCols,plotId})
    setappdata(ax, 'SubplotPosition', position); % normalized
    subplotlayoutInvalid(handle(ax), [], p);
    
    %when subplot is not in a single grid cell for the current grid,
    %don't add it to the auto-layout
    if any(size(grid) ~= [nRows, nCols]) ... %active grid shape does not match n,m
            || length(row) ~= 1 || length(col) ~= 1 ... %multi-cell subplot
            || round(row) ~= row || round(col) ~= col ... %non-integer cell specified
            || grid(row + 1, col + 1) == ax %axes is already in cell m,n,p
        addAxesToSpanGrid(ax, nRows, nCols, row, col);
        return
    end
    
    % only add axes to grid if it doesn't span multiple columns or rows
    grid(row + 1, col + 1) = ax;
    setappdata(p,  'SubplotGrid', grid)
end

%remember that this was a subplot, even though it was positioned explicitly
function addNonGridAxes(ax, position)    
    setappdata(ax,  'SubplotGridLocation', [])
    setappdata(ax, 'SubplotPosition', position); % normalized
end

% Remember that this was a 'span' subplot.  We don't adjust these
% for overlap but we do adjust for subplot titles.
function addAxesToSpanGrid(ax, nRows, nCols, row, col)    
    p = ax.Parent;
    spanList = getappdata(p, 'SubplotSpanGrid');
    if isempty(spanList)
        spanList = gobjects(nRows,nCols);
    end   
    for i=1:length(row)
        for j =1:length(col)
            spanList(floor(row(i)) + 1, floor(col(j)) + 1) = ax;
        end
    end
    setappdata(p,'SubplotSpanGrid',spanList);
end


%----------------------------------------------------------------%
% Convert units of offsets like LooseInset or TightInset
% Note: Copied from legendcolorbarlayout.m
function out = offsetsInUnits(ax, in, from, to)
    fig = ancestor(ax, 'figure');
    par = ax.Parent;
    p1 = hgconvertunits(fig, [0, 0, in(1 : 2)], from, to, par);
    p2 = hgconvertunits(fig, [0, 0, in(3 : 4)], from, to, par);
    out = [p1(3 : 4), p2(3 : 4)];
end

%----------------------------------------------------------------%
% Compute reference OuterPos from pos and loose. Note that
% loose insets are relative to outerposition
% Note: Copied from legendcolorbarlayout.m
function outer = getOuterFromPosAndLoose(pos, loose, units)
    if strcmp(units, 'normalized')
        % compute outer width and height and normalize loose to them
        w = pos(3) / (1 - loose(1) - loose(3));
        h = pos(4) / (1 - loose(2) - loose(4));
        loose = [w, h, w, h] .* loose;
    end
    outer = [pos(1 : 2) - loose(1 : 2), pos(3 : 4) + loose(1 : 2) + loose(3 : 4)];
end

function sibs = datasiblings(parent) 
% Returns any siblings which do not have the appdata 'NonDataObject'. In 
% other words, returns "real" axes, not things like legend or scribe objects.
%
    sibs = parent.Children;
    nondatachild = logical([]);
    for i=length(sibs):-1:1
        nondatachild(i) = isappdata(sibs(i),'NonDataObject');
    end
    sibs(nondatachild) = [];
end

function out = isSubplotCandidate(obj)
    out = false;
    if isgraphics(obj)
        h = handle(obj);
        out = isa(h,'matlab.graphics.axis.AbstractAxes') ...
            || isa(h, 'matlab.graphics.chart.ChartGroup') ...
            || isa(h, 'matlab.graphics.chart.Chart');
    end
end

function match = isObjectAlreadyInCell(m,n,p,h)
% Check if an axes/chart was already designated for a specified subplot 
% grid location by examining appdata. This check is performed to 
% allow special case of swapaxes using m,n,p,h syntax for charts. Subplot
% for charts doesn't yet know how to compute OuterPosition; we only want 
% to support m,n,p syntax for sealed charts only when the chart
% has already been initialized with an OuterPosition and grid location 
% ahead of time by swapaxes.
    match = false;
    if(isappdata(h,'SubplotGridLocation'))
        ad = getappdata(h,'SubplotGridLocation');
        match = iscell(ad) && length(ad) == 3 && ...
            ad{1} == m && ad{2} == n && ad{3} == p;
            
    end
end
```
