## Supplementary material for "Neptune: a toolbox for spinal cord functional MRI data processing and quality assurance": Neptune software: subplot2n_parseargs.m

```
function [args,strargs,narg] = subplot2n_parseargs(args)
% Helper function used by subplot for separating out name/value pairs and
% string arguments from other arguments. entries out of the pvpair list.
% Note that the arguments may include a three digit string as the first
% input argument, which represents 'mnp'. That will be separated out into
% three separate numeric input arguments for [m, n, p].

% Copyright 2015-2017 The MathWorks, Inc.

% Check if the first input argument is a string representing 'mnp'
% If so, convert it to three separate numeric inputs.

import matlab.graphics.internal.*;
if numel(args)>=1 && isCharOrString(args{1}) ... % it is a string
        && numel(args{1})==3 ... % it has three characters
        && all(args{1} >= '0' & args{1} <= '9') % characters are between 0-9

    args = [{str2double(args{1}(1)) ,... % first character
             str2double(args{1}(2)) ,... % second character
             str2double(args{1}(3))},... % third character
             args(2:end)];
end

args = convertStringToCharArgs(args);

% Find the first string input.
% (append true so that firststr is never empty)
firststr = find([cellfun(@ischar,args), true],1);

% Split the arguments at the first string input.
strargs = args(firststr:end);
args = args(1:firststr-1);
narg = numel(args);

end
```
