## Supplementary material for "Neptune: a toolbox for spinal cord functional MRI data processing and quality assurance": Neptune software: suptitle2.m

```
function hout=suptitle2(str)
%SUPTITLE puts a title above all subplots.
%
%	SUPTITLE('text') adds text to the top of the figure
%	above all subplots (a "super title"). Use this function
%	after all subplot commands.
%
%   SUPTITLE is a helper function for yeastdemo.

%   Copyright 2003-2014 The MathWorks, Inc.

% Warning: If the figure or axis units are non-default, this
% function will temporarily change the units.

% Parameters used to position the supertitle.

% Amount of the figure window devoted to subplots
plotregion = .92;

% Y position of title in normalized coordinates
titleypos  = .95;

% Fontsize for supertitle
fs = get(gcf,'defaultaxesfontsize')+4;

% Fudge factor to adjust y spacing between subplots
fudge=1;

haold = gca;
figunits = get(gcf,'units');

% Get the (approximate) difference between full height (plot + title
% + xlabel) and bounding rectangle.

if ~strcmp(figunits,'pixels')
    set(gcf,'units','pixels');
    pos = get(gcf,'position');
    set(gcf,'units',figunits);
else
    pos = get(gcf,'position');
end
ff = (fs-4)*1.27*5/pos(4)*fudge;

% The 5 here reflects about 3 characters of height below
% an axis and 2 above. 1.27 is pixels per point.

% Determine the bounding rectangle for all the plots

h = findobj(gcf,'Type','axes');

oldUnits = get(h, {'Units'});
if ~all(strcmp(oldUnits, 'normalized'))
    % This code is based on normalized units, so we need to temporarily
    % change the axes to normalized units.
    set(h, 'Units', 'normalized');
    cleanup = onCleanup(@()resetUnits(h, oldUnits));
end

max_y=0;
min_y=1;
oldtitle = [];
numAxes = length(h);
thePositions = zeros(numAxes,4);
for i=1:numAxes
    pos=get(h(i),'pos');
    thePositions(i,:) = pos;
    if ~strcmp(get(h(i),'Tag'),'suptitle')
        if pos(2) < min_y
            min_y=pos(2)-ff/5*3;
        end
        if pos(4)+pos(2) > max_y
            max_y=pos(4)+pos(2)+ff/5*2;
        end
    else
        oldtitle = h(i);
    end
end

if max_y > plotregion
    scale = (plotregion-min_y)/(max_y-min_y);
    for i=1:numAxes
        pos = thePositions(i,:);
        pos(2) = (pos(2)-min_y)*scale+min_y;
        pos(4) = pos(4)*scale-(1-scale)*ff/5*3;
        set(h(i),'position',pos);
    end
end

np = get(gcf,'nextplot');
set(gcf,'nextplot','add');
if ~isempty(oldtitle)
    delete(oldtitle);
end
axes('pos',[0 1 1 1],'visible','off','Tag','suptitle');
ht=text(.5,titleypos-1,str);set(ht,'horizontalalignment','center','fontsize',fs);
set(gcf,'nextplot',np);
% axes(haold);
if nargout
    hout=ht;
end
end

function resetUnits(h, oldUnits)
    % Reset units on axes object. Note that one of these objects could have
    % been an old supertitle that has since been deleted.
    valid = isgraphics(h);
    set(h(valid), {'Units'}, oldUnits(valid));
end
```
