## Supplementary material for "Neptune: a toolbox for spinal cord functional MRI data processing and quality assurance": Neptune software: wgr_deconv_canonhrf_par.m

```
function [data_deconv, event, HRF, adjust_global, PARA] = wgr_deconv_canonhrf_par(data,thr,event_lag_max,TR)

%%% this function implements the method described in 
%%% Wu et al, 
%%% A blind deconvolution approach to recover effective connectivity brain networks from resting state fMRI data,
%%% Med Image Anal. 2013 Jan 29. pii: S1361-8415(13)00004-2. doi: 10.1016/j.media.2013.01.003

%%% input
%%% data, dimensions time points x number of voxels, normalized
%%% threshold, assuming data are normalized
%%% event_lag_max: maximum time from neural event to BOLD event in bins, not time 
%%% (e.g. if we assume 10seconds, and TR=2s,  set the value to 10/2=5)
%%% TR is the TR parameter, in seconds.

%%% Some parts of the code (subfunction: Fit_Canonical_HRF, CanonicalBasisSet, get_parameters2) were modified from the hemodynamic response estimation toolbox(http://www.stat.columbia.edu/~martin/HRF_Est_Toolbox.zip).
%%% 
%%% the code uses the parallel for loop ¡°parfor¡±. In case of older matlab versions, parfor can be changed to standard for. 
%%% 
%%% The default is using canonical hrf and two derivatives, as described in the paper.
%%% The function can be modified to use instead point processes, FIR model, etc. 


[N, nvar] = size(data);
[even_new]=wgr_trigger_onset(data,thr);
p_m=3;
% Options: p_m=1 - only canonical HRF
%          p_m=2 - canonical + temporal derivative
%          p_m=3 - canonical + time and dispersion derivative
T = round(30/TR);%% assume HRF effect lasts 30s.
data_deconv = zeros(N,nvar);
HRF = zeros(T,nvar);
PARA = zeros(3,nvar);
warning off
parfor i=1:nvar
    [data_deconv(:,i), HRF(:,i), event{i}, adjust_global(i),  PARA(:,i)] = wgr_adjust_onset(data(:,i),even_new{i},event_lag_max,TR,p_m,T,N);
end
warning on


function [dat_deconv, hrf, even_new, ad_global, param] = wgr_adjust_onset(dat,even_new,event_lag_max,TR,p_m,T,N)
%% global adjust.
kk=1;
hrf = zeros(T,event_lag_max+1);
for event_lag=0:event_lag_max
    RR = even_new-event_lag; RR(RR<=0)=[];
    design = zeros(N,1);
    design(RR) = 1;
    [hrf(:,kk), e3, param(:,kk)] = Fit_Canonical_HRF(dat,TR,design,T,p_m);
    Cov_E(kk) = cov(e3);
    kk = kk+1;
end
[C, ind] = min(Cov_E); ad_global=ind-1;%begin with 0.
even_new = even_new-ad_global; even_new(even_new<=0)=[]; 
hrf = hrf(:,ind);
param = param(:,ind);
%% linear deconvolution.
H=fft([hrf; zeros(N-T,1)]);
M=fft(dat);
dat_deconv = ifft(conj(H).*M./(H.*conj(H)+C));

return

function [oneset] = wgr_trigger_onset(matrix,thr)
[N, nvar] = size(matrix);
matrix = zscore(matrix);
% Computes pseudo event.
for i = 1:nvar
    oneset_temp = [];
    for t  = 2:N-1
        if matrix(t,i) > thr && matrix(t-1,i)<matrix(t,i) && matrix(t,i)>matrix(t+1,i)% detects threshold
            oneset_temp = [oneset_temp t] ;
        end
    end
    oneset{i} = oneset_temp;      
end

return

function [hrf, e, param] = Fit_Canonical_HRF(tc,TR,Run,T,p)
%
% Fits GLM using canonical hrf (with option of using time and dispersion derivatives)';
%
% INPUTS:
% 
% tc    - time course
% TR    - time resolution
% Runs  - expermental design
% T     - length of estimated HRF
% p     - Model type
%
% Options: p=1 - only canonical HRF
%          p=2 - canonical + temporal derivative
%          p=3 - canonical + time and dispersion derivative
% 
% OUTPUTS:
%
% hrf   - estimated hemodynamic response function
% fit   - estimated time course
% e     - residual time course
% param - estimated amplitude, height and width

len = length(Run);

X = zeros(len,p);

[h, dh, dh2] = CanonicalBasisSet(TR);
v = conv(Run,h);
X(:,1) = v(1:len);

if (p>1)
    v = conv(Run,dh);
    X(:,2) = v(1:len);
end

if (p>2)
    v = conv(Run,dh2);
    X(:,3) = v(1:len);
end

X = [(zeros(len,1)+1) X];
b = pinv(X)*tc;
e = tc-X*b;
fit = X*b;

b = b(2:end);

if (p == 2)
    bc = sign(b(1))*sqrt(b(1)^2 + b(2)^2); 
    H = [h dh];    
elseif (p==1)
    bc = b(1);
    H = h;
elseif (p>2)
    bc = sign(b(1))*sqrt(b(1)^2 + b(2)^2 + b(3)^2);
    H = [h dh dh2];
end    

hrf = H*b;

param = get_parameters2(hrf,T);

return
% END MAIN FUNCTION
%%%%%%%%%%%%%%%%%%%%%%%%%%%%%%%%%%%%%%%%%%%%%%%%%%%%%%%%%%%%%%%%%%%%%%%%%%%%%%%%

function [h, dh, dh2] = CanonicalBasisSet(TR)

len = round(30/TR);
xBF.dt = TR;
xBF.length= len;
xBF.name = 'hrf (with time and dispersion derivatives)';
xBF = spm_get_bf(xBF);

v1 = xBF.bf(1:len,1);
v2 = xBF.bf(1:len,2);
v3 = xBF.bf(1:len,3);

h = v1;
dh =  v2 - (v2'*v1/norm(v1)^2).*v1;
dh2 =  v3 - (v3'*v1/norm(v1)^2).*v1 - (v3'*dh/norm(dh)^2).*dh;

h = h./max(h);
dh = dh./max(dh);
dh2 = dh2./max(dh2);

return


function [param] = get_parameters2(hdrf,t)

% Find model parameters
%
% Height - h
% Time to peak - p (in time units of TR seconds)
% Width (at half peak) - w  


% Calculate Heights and Time to peak:

% n = t(end)*0.6;
n = round(t*0.8);

[h,p] = max(abs(hdrf(1:n)));
h = hdrf(p);

%if (p > t(end)*0.6), warning('Late time to peak'), end;

if (h >0)
    v = (hdrf >= h/2);    
else
    v = (hdrf <= h/2);
end;
    
[a,b] = min(diff(v));
v(b+1:end) = 0;
w = sum(v);

cnt = p-1;
g =hdrf(2:end) - hdrf(1:(end-1));
while((cnt > 0) & (abs(g(cnt)) <0.001)),
    h = hdrf(cnt);
    p = cnt;
    cnt = cnt-1;
end;


param = zeros(3,1);
param(1) = h;
param(2) = p;
param(3) = w;

return;
```
