## Supplementary material for "Neptune: a toolbox for spinal cord functional MRI data processing and quality assurance": Neptune software: xcorr_zscore.m

```
function [z, r] = xcorr_zscore(a, b, dof_correction, siz4)

% Created 2014-Aug-26 (RLB)
% Revised 2014-Nov-24 - permit truncation of time series (via siz4)

if nargin < 4
    siz4 = size(a,2);
    if nargin < 3
        dof_correction = 1;
    end
end

% only truncate time series if user specifies length
if exist('siz4','var')
    if (size(a,2)~=siz4)||(size(b,2)~=siz4)
        a = a(:,1:siz4);
        b = b(:,1:siz4);
    end
end

% % zero mean
% a = a - mean(a,2);
% b = b - mean(b,2);

% cross correlation
r = corrcoef(a,b);
r = r(1,2);
z_from_r = atanh(r);

if dof_correction
    
    % first order autocorrelation
    p1 = corrcoef(a(2:end),a(1:end-1));
    p1 = p1(1,2);
    p2 = corrcoef(b(2:end),b(1:end-1));
    p2 = p2(1,2);
    
    % actual degrees of freedom estimates
    dof1 = (1-p1^2)/(1+p1^2);
    dof2 = (1-p2^2)/(1+p2^2);
    
    % corrected z-score
    z1 = z_from_r .* sqrt((siz4.*dof1)-3);
    z2 = z_from_r .* sqrt((siz4.*dof2)-3);
    z = (z1 + z2) / 2;
    
else % do not correct for inflated degrees of freedom
    
    z = z_from_r .* sqrt(siz4-3);
end
```
