## Supplementary material for "Neptune: a toolbox for spinal cord functional MRI data processing and quality assurance": Neptune software: subtightplot.m

```
function h=subtightplot(m,n,p,gap,marg_h,marg_w,varargin)
%
% Functional purpose: A wrapper function for Matlab function subplot. Adds the ability to define the gap between
% neighbouring subplots. Unfotrtunately Matlab subplot function lacks this functionality, and the gap between
% subplots can reach 40% of figure area, which is pretty lavish.  
%
% Input arguments (defaults exist):
%   gap- two elements vector [vertical,horizontal] defining the gap between neighbouring axes. Default value
%            is 0.01. Note this vale will cause titles legends and labels to collide with the subplots, while presenting
%            relatively large axis. 
%   marg_h  margins in height in normalized units (0...1)
%            or [lower uppper] for different lower and upper margins 
%   marg_w  margins in width in normalized units (0...1)
%            or [left right] for different left and right margins 
%
% Output arguments: same as subplot- none, or axes handle according to function call.
%
% Issues & Comments: Note that if additional elements are used in order to be passed to subplot, gap parameter must
%       be defined. For default gap value use empty element- [].      
%
% Usage example: h=subtightplot((2,3,1:2,[0.5,0.2])

if (nargin<4) || isempty(gap),    gap=0.01;  end
if (nargin<5) || isempty(marg_h),  marg_h=0.05;  end
if (nargin<5) || isempty(marg_w),  marg_w=marg_h;  end
if isscalar(gap),   gap(2)=gap;  end
if isscalar(marg_h),  marg_h(2)=marg_h;  end
if isscalar(marg_w),  marg_w(2)=marg_w;  end
gap_vert   = gap(1);
gap_horz   = gap(2);
marg_lower = marg_h(1);
marg_upper = marg_h(2);
marg_left  = marg_w(1);
marg_right = marg_w(2);

%note n and m are switched as Matlab indexing is column-wise, while subplot indexing is row-wise :(
[subplot_col,subplot_row]=ind2sub([n,m],p);  

% note subplot suppors vector p inputs- so a merged subplot of higher dimentions will be created
subplot_cols=1+max(subplot_col)-min(subplot_col); % number of column elements in merged subplot 
subplot_rows=1+max(subplot_row)-min(subplot_row); % number of row elements in merged subplot   

% single subplot dimensions:
height=(1-(marg_lower+marg_upper)-(m-1)*gap_vert)/m;
width =(1-(marg_left+marg_right)-(n-1)*gap_horz)/n;

% merged subplot dimensions:
merged_height=subplot_rows*( height+gap_vert )- gap_vert;
merged_width= subplot_cols*( width +gap_horz )- gap_horz;

% merged subplot position:
merged_bottom=(m-max(subplot_row))*(height+gap_vert) +marg_lower;
merged_left=(min(subplot_col)-1)*(width+gap_horz) +marg_left;
pos_vec=[merged_left merged_bottom merged_width merged_height];

h=subplot('Position',pos_vec,varargin{:});

if (nargout < 1),  clear h;  end

end
```
