## Supplementary material for "Neptune: a toolbox for spinal cord functional MRI data processing and quality assurance": Neptune software: txt_update.m

```
function [S] = txt_update
% Creates a textbox.
S.fh = figure('units','normalized',...
              'position',[0 1/11 9/40 5/12],...
              'menubar','none',...
              'resize','on',...
              'numbertitle','off',...
              'name','Execution log');
S.tx = uicontrol('style','edit',...
                 'units','normalized',...
                 'Position',[0 0 1 1],...
                 'backgroundcolor','w',...
                 'HorizontalAlign','left',...
                 'min',0,'max',10,...
                 'enable','inactive');
end
```
