## Supplementary material for "Neptune: a toolbox for spinal cord functional MRI data processing and quality assurance": Neptune software: setup_neptune.m

```
% NEPTUNE SETUP
% 
% Run this script from its current directory to set up Neptune
% 
% It removes older versions of NEPTUNE from MATLAB path, 
% moves NEPTUNE toolbox directory to MATLAB's toolboxes home directory, and
% adds NEPTUNE to MATLAB path and pathdef file, and
% saves installation notes into the current directory and NEPTUNE's installed directory
% 
% Ranga Deshpande | November 2022
% 
% ------- ------- -------

remove_existing_installation = 1; % 1 = remove; 0 = do not remove existing Neptune installation if it exists

if remove_existing_installation==1
    existing_version = which('neptune.m');
    if ~isempty(existing_version) % remove existing Neptune installation if it exists
        existing_version = existing_version(1:end-10);
        rmpath(genpath(existing_version))
        movefile(existing_version,[existing_version '_uninstalled'])
        % The older version will not be deleted (in case any of your projects depended on it),
        % but it will be removed from MATLAB path and renamed with the '_uninstalled' suffix.
    end
end

current_dir = [pwd '/'];
sdir = dir('neptune_v*');
if isempty(sdir)
    clear current_dir sdir
    cd ..
    current_dir = [pwd '/'];
    sdir = dir('neptune_v*');
end
for k = length(sdir):-1:1
    if sdir(k).isdir==0
        sdir(k) = [];
    end
end; clear k

destination_dir = which('spm.m'); % if SPM is installed, choose destination directory as the one in which SPM is installed. This is just a pointer; Neptune does not require SPM.
destination_dir = destination_dir(1:end-5);
tmp = destination_dir(1:max(strfind(destination_dir(1:end-1),'/')));
if isempty(tmp)
    tmp = destination_dir(1:max(strfind(destination_dir(1:end-1),'\')));
end
clear destination_dir
destination_dir = tmp; clear tmp

if isempty(destination_dir) % if SPM is not installed, choose destination directory as the MATLAB toolbox directory within the MATLAB installation folder
    clear destination_dir
    destination_dir = [fullfile(matlabroot,'toolbox') '/'];
end

if ~exist([destination_dir sdir(1).name],'dir')
    mkdir([destination_dir sdir(1).name])
end
copyfile([current_dir sdir(1).name],[destination_dir sdir(1).name])

addpath(genpath([destination_dir sdir(1).name]))
savepath(which('pathdef.m'))

clk = clock;
month={'Jan','Feb','Mar','Apr','May','Jun','Jul','Aug','Sep','Oct','Nov','Dec'}';
clk_filename = sprintf('%.4d-%.2d-%.2d_%.2d-%.2d',clk(1),clk(2),clk(3),clk(4),clk(5));
if (remove_existing_installation==1)&&(~isempty(existing_version))
    message_done = sprintf('\nDate and time: %d/%s/%d %dh:%dm:%ds\n\nNEPTUNE has been installed into the following directory:\n%s%s/\n\nNEPTUNE has been added to MATLAB path, and will remain in path every time MATLAB is opened.\nYou could choose to delete the current folder with setup file if you wish.\n\nAn older version of Neptune in the following location has been removed from MATLAB path and renamed with the suffix ''_uninstalled'': %s',clk(3),month{clk(2)},clk(1),clk(4),clk(5),round(clk(6)), destination_dir,sdir(1).name,existing_version);
else
    message_done = sprintf('\nDate and time: %d/%s/%d %dh:%dm:%ds\n\nNEPTUNE has been installed into the following directory:\n%s%s/\n\nNEPTUNE has been added to MATLAB path, and will remain in path every time MATLAB is opened.\nYou could choose to delete the current folder with setup file if you wish.\n',clk(3),month{clk(2)},clk(1),clk(4),clk(5),round(clk(6)), destination_dir,sdir(1).name);
end

fileID = fopen([current_dir sprintf('neptune_installation_notes_%s.txt',clk_filename)],'w');
fprintf(fileID,'%s\n',message_done);

copyfile( [current_dir sprintf('neptune_installation_notes_%s.txt',clk_filename)], [destination_dir sdir(1).name '/' sprintf('neptune_installation_notes_%s.txt',clk_filename)] )

disp(message_done)

clear current_dir sdir destination_dir clk clk_filename message_done fileID remove_existing_installation existing_version

%%
```
