## Supplementary material for "Neptune: a toolbox for spinal cord functional MRI data processing and quality assurance": Neptune software: updates_info.txt

neptune_v1.211110
   Release date: 10-November-2021
   Initial release

neptune_v1.211116
   Release date: 16-November-2021
   Minor bug fixes

neptune_v1.211118
   Release date: 18-November-2021
   Minor bug fixes

-------
neptune_v1.211227
   Release date: 27-December-2021
   SCT-guided manual segmentation masks (for steps 5 and 10):
   - Step 5 (not-cord mask): Spinal Cord Toolbox (SCT) propseg is used to generate the cord mask boundary, which is displayed to the user. The user can either accept it as is, or modify the points and accept.
   - Step 10 (anatomical masks): Likewise, SCT propseg is used to generate WM and CSF boundaries while deepseg is used to generate GM boundary. The boundaries are displayed to the user, as and before they can either accept it as is or modify the points as required.
   In both cases, if SCT is unavailable then the user is given the opportunity to manually draw the boundaries themselves. SCT-guided manual masks' generation saves approximately 90% time on average compared to earlier Neptune versions.

-------
neptune_v1.220505
   Release date: 05-May-2022
   - Step 1 (NORDIC denoising): GUI option to perform NORDIC denoising of NIfTI fMRI data (magnitude-only data, OR, if available magnitude+phase data).
   - Step 10 (anatomical masks): the previous version obtained tissue boundaries using SCT deepseg; now the user also has the option of using PAM50 GM masks warped to the anatomical space upon co-registration in SCT. The updated interface allows the user to choose either the PAM50-based boundary, or the deepseg segmentation boundary, or if none of those are satisfactory then draw one's own boundary.
   - Step 18 (standard space co-registration using SCT): perform co-registration to PAM50 space and obtain templates, masks, atlases and labels in anatomical space. These masks are currently not used during post-processing, but the user could make use of them outside of Neptune.
   - Option to discard first and last slices (because with some sequences they are bad).
   - Option to discard the first N functional volumes.
   - Option to smooth motion parameters to suppress sporadic artifacts.
   - Neptune now provides a small set of shortlisted QC outputs for a quick assessment, which has been integrated into the HTML report.
   - Neptune setup now first uninstalls the older version of Neptune (removes it from MATLAB path and deletes Neptune directory), and then proceeds with the installation of the current version.
   - Bug fixes and performance improvements.

-------
neptune_v1.231111
   Release date: 11-November-2023
   - Neptune setup now does not delete the older version (if it exists), but renames it and removes it from the MATLAB path. This way, for records, researchers can keep a copy of Neptune used to process their older data.
   - Step-9 (added 3dDespike option with modifications): if RETROICOR is unable to run due to unavailability of physiological data or incompatibility, Neptune now performs AFNI's 3dDespike denoising to minimize spikes in fMRI time series caused by high motion or swallowing volumes. Hence, Step-9 can now be chosen even when physiological data are not recorded.
   - Fixed bugs that occurred when running Neptune subject after subject.
   - A new GUI option has been added to choose which subject number to start processing from when running subject after subject. This helps those running Neptune repeatedly on the same dataset who might want to restart from one of the middle subjects 'S' after previously completing the first 'S-1' subjects.
   - Significantly, innumerable minor enhancements, bug fixes and performance improvements have been made.

-------
neptune_v1.260301
   Release date: 1-March-2026
   - Step-11: During functional-to-anatomical co-registration, if the centers of mass are misaligned, then it will now perform co-registration after aligning the centers of mass, thus avoiding co-registration failures in several circumstances.
   - Fixed minor bugs and improved the performance for a slight increase in the speed of processing 

-------
