## Supplementary figures and images for "Neptune: a toolbox for spinal cord functional MRI data processing and quality assurance"

### barrygp.jpg

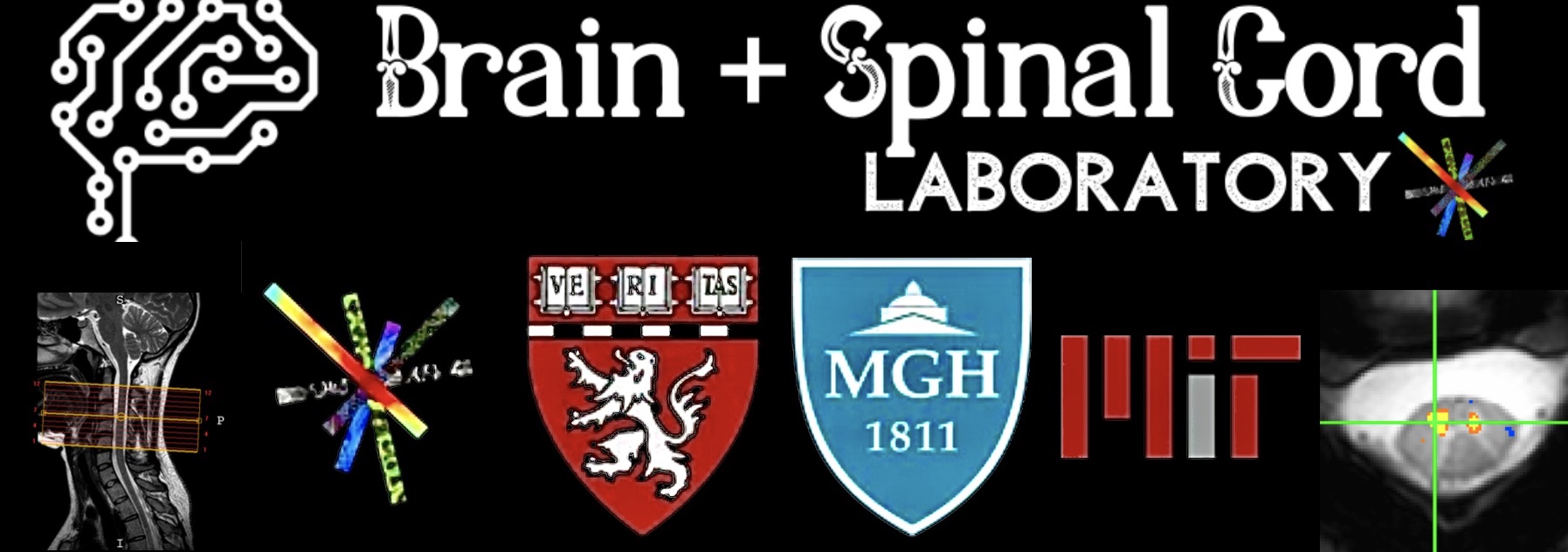

### completed.jpg

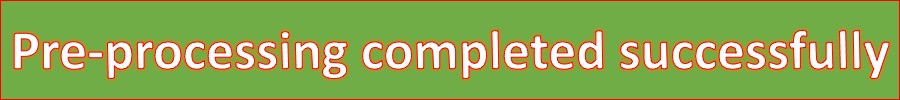

### green.jpg

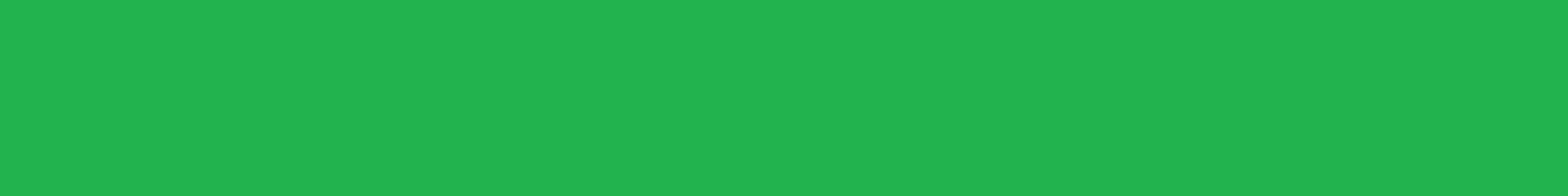

### neptune_v1.0.jpg

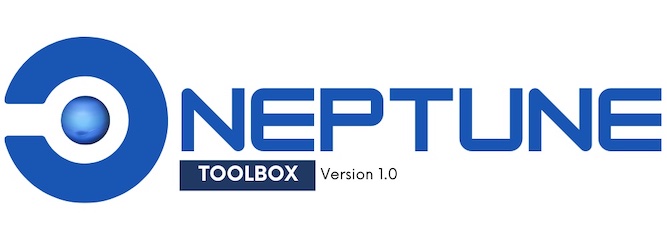

### neptune_v1.0_all_done.jpg

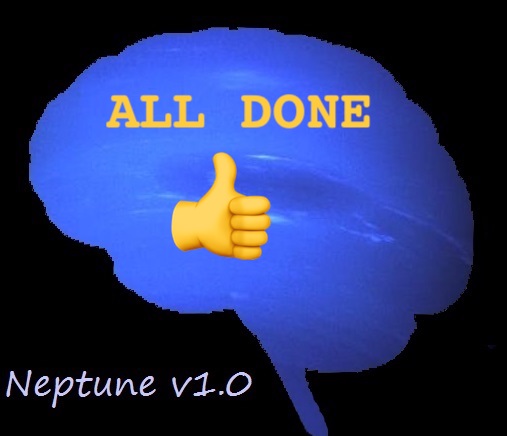

### neptune_v1.0_old.jpg

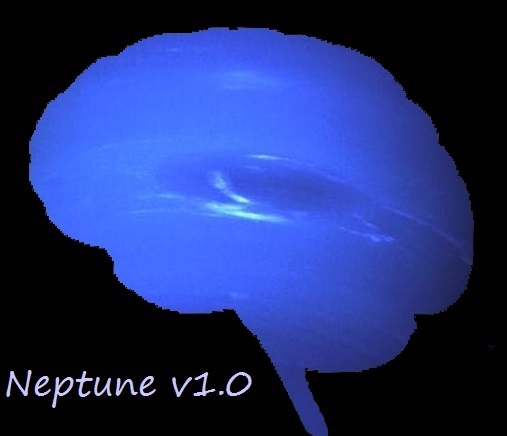

### red.jpg

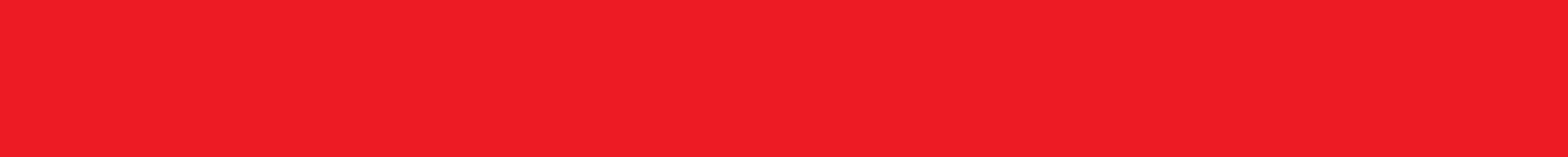

### status_blue.jpg

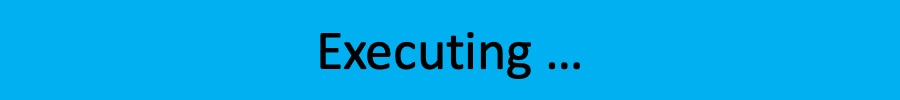

### status_blue_UI.jpg

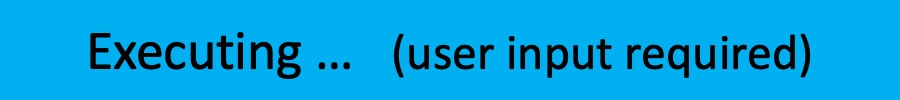

### status_gray.jpg

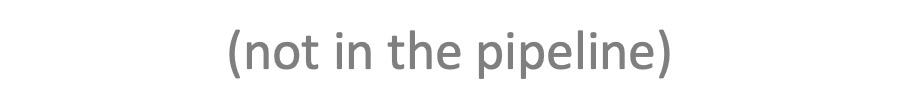

### status_green.jpg

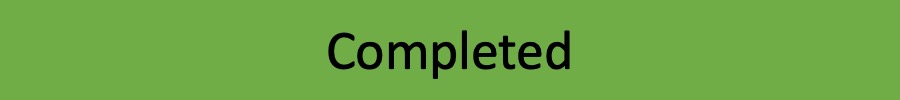

### status_green_skip.jpg

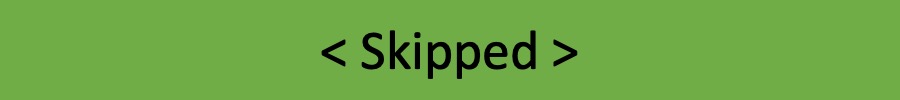

### status_white.jpg

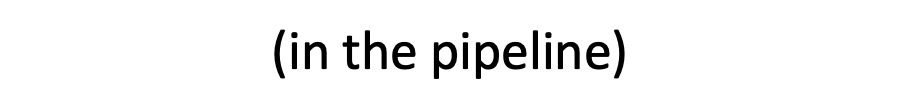

### status_white_UI.jpg

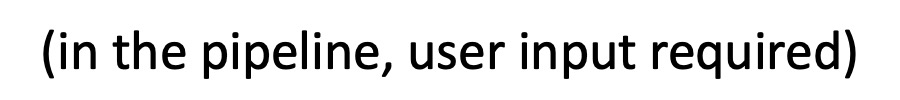

### step_01_blue.jpeg

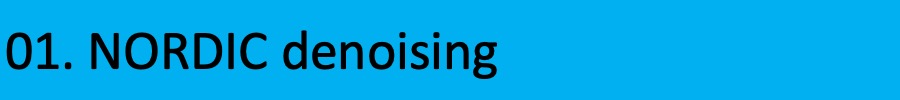

### step_01_gray.jpeg

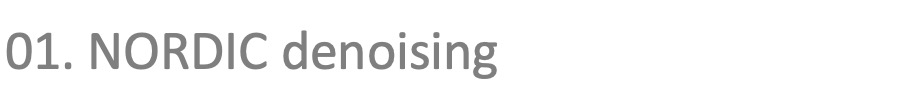

### step_01_green.jpeg

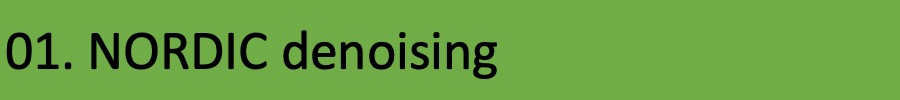

### step_01_white.jpeg

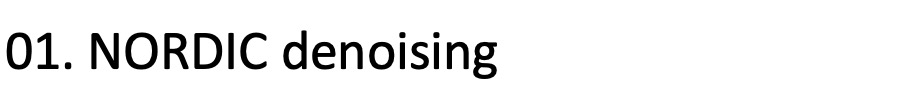

### step_02_blue.jpeg

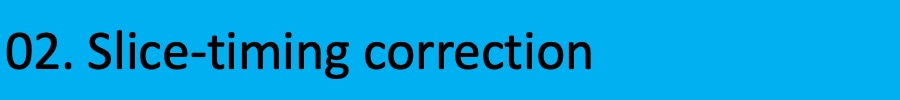

### step_02_gray.jpeg

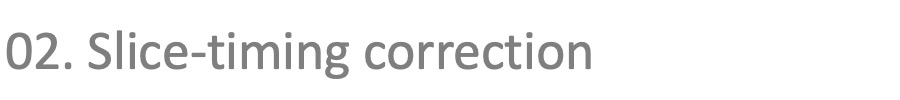

### step_02_green.jpeg

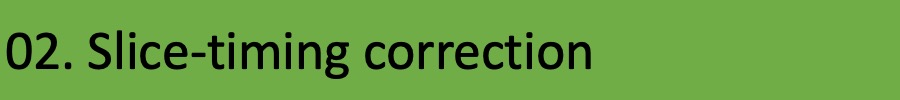

### step_02_white.jpeg

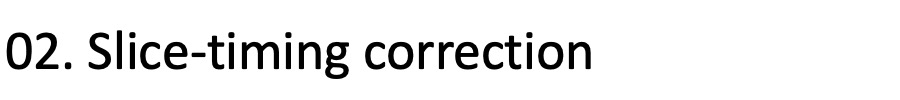

### step_03_blue.jpeg

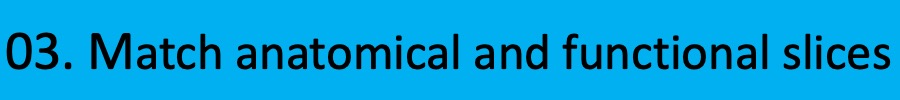

### step_03_gray.jpeg

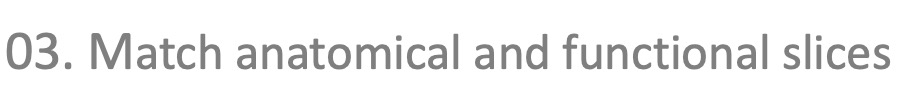

### step_03_green.jpeg

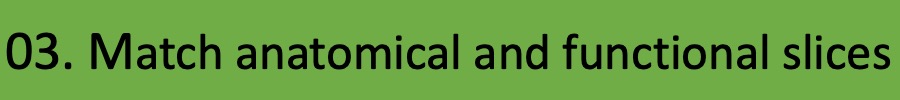

### step_03_white.jpeg

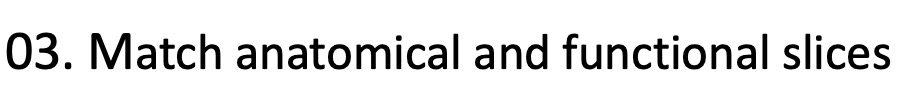

### step_04_blue.jpeg

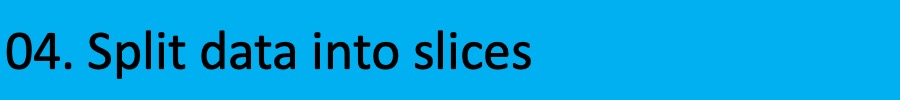

### step_04_gray.jpeg

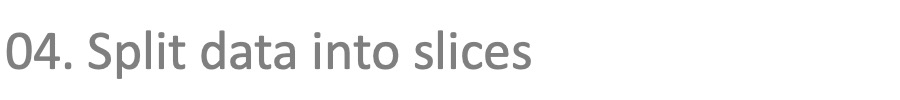

### step_04_green.jpeg

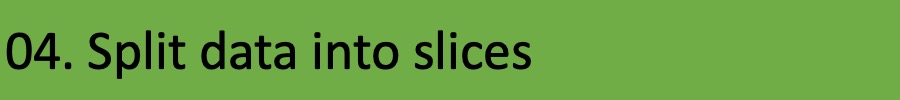

### step_04_white.jpeg

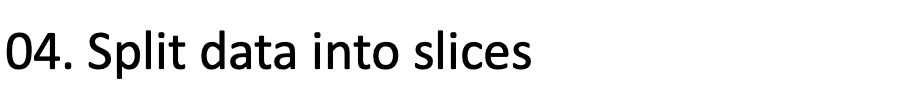

### step_05_blue.jpeg

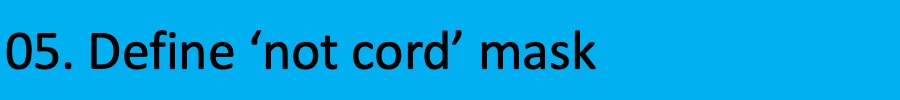

### step_05_gray.jpeg

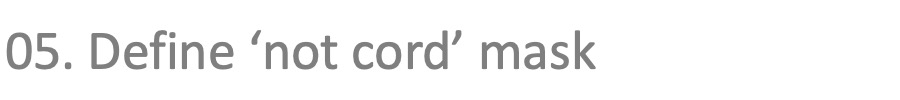

### step_05_green.jpeg

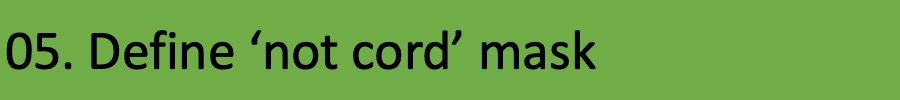

### step_05_white.jpeg

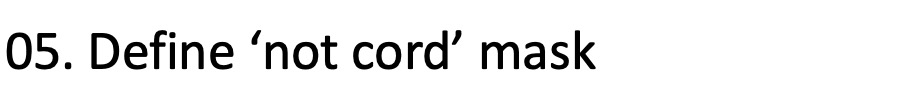

### step_06_blue.jpeg

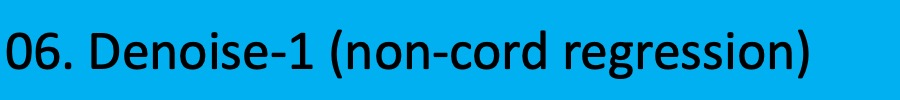

### step_06_gray.jpeg

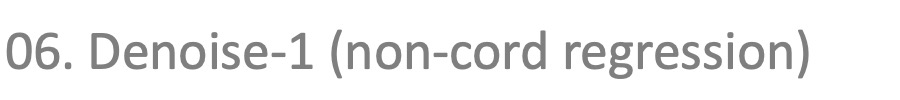

### step_06_green.jpeg

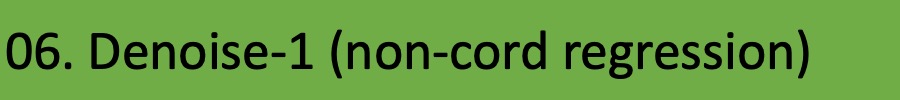

### step_06_white.jpeg

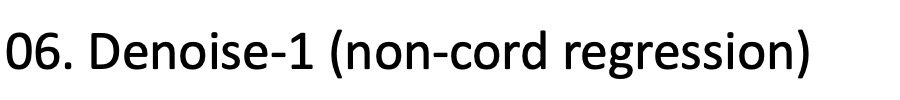

### step_07_blue.jpeg

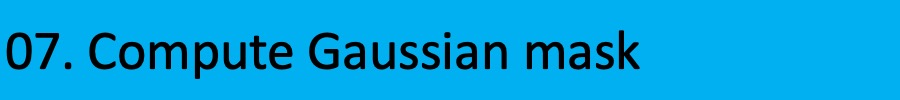

### step_07_gray.jpeg

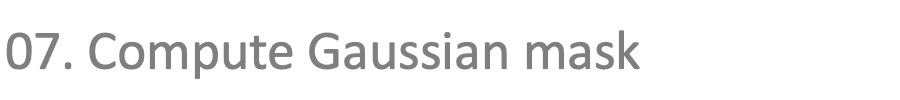

### step_07_green.jpeg

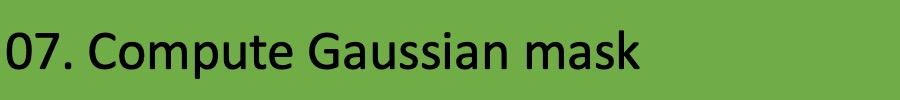

### step_07_white.jpeg

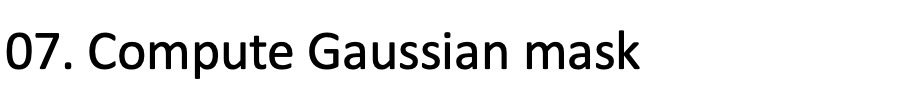

### step_08_blue.jpeg

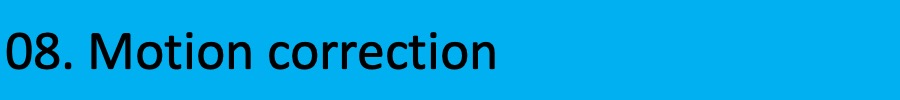

### step_08_gray.jpeg

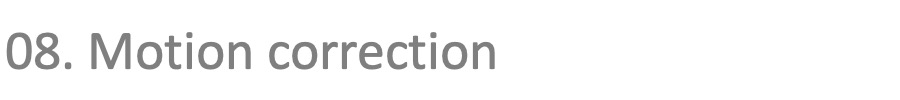

### step_08_green.jpeg

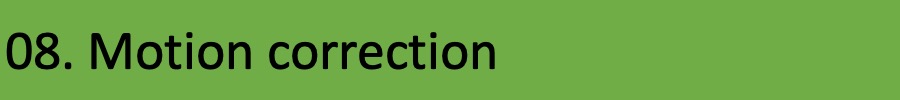

### step_08_white.jpeg

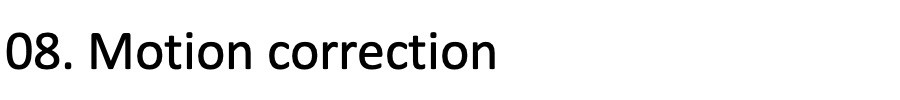

### step_09_blue.jpeg

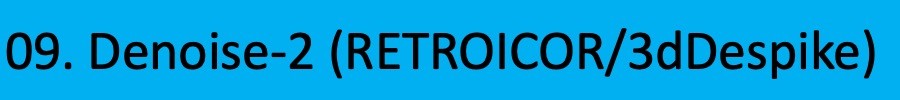

### step_09_blue_2.jpeg

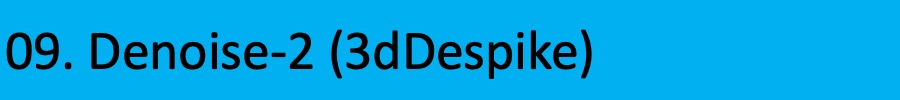

### step_09_gray.jpeg

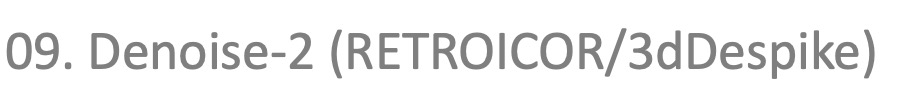

### step_09_green.jpeg

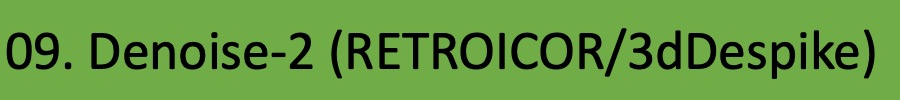

### step_09_green_2.jpeg

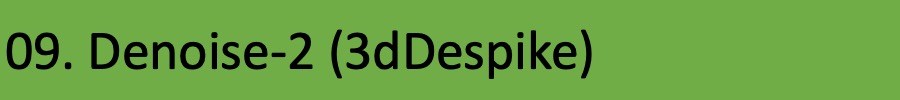

### step_09_white.jpeg

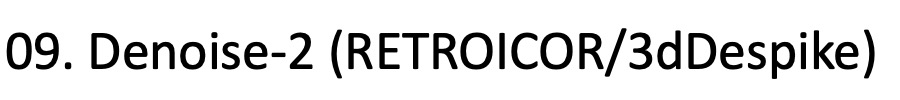

### step_10_blue.jpeg

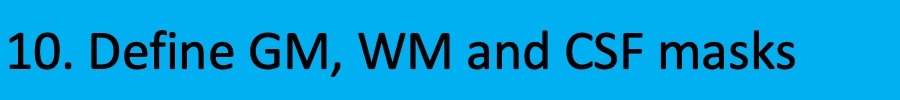

### step_10_gray.jpeg

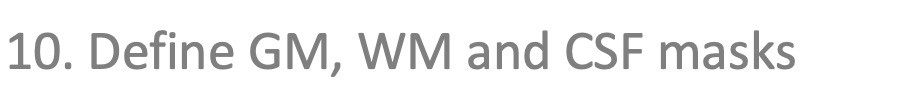

### step_10_green.jpeg

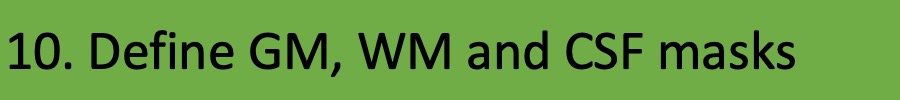

### step_10_white.jpeg

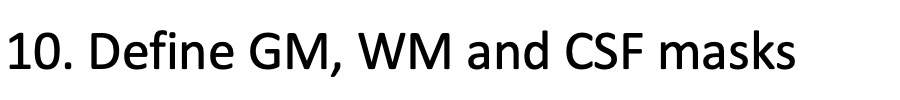

### step_11_blue.jpeg

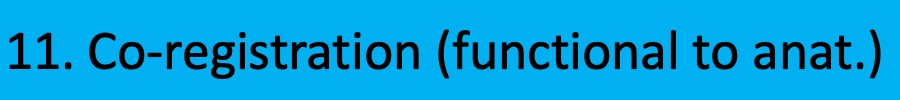

### step_11_gray.jpeg

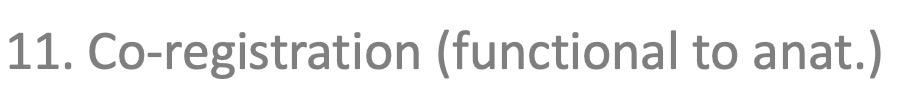

### step_11_green.jpeg

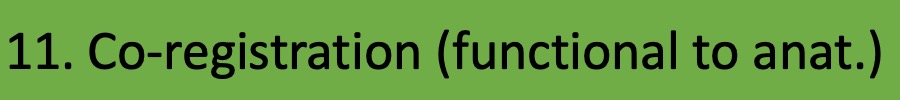

### step_11_white.jpeg

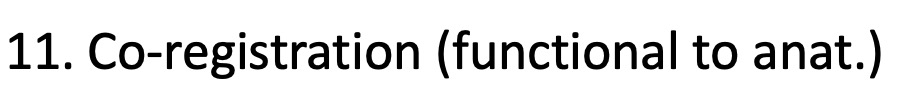

### step_12_blue.jpeg

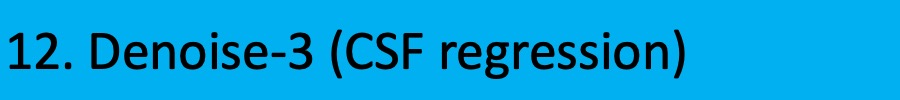

### step_12_gray.jpeg

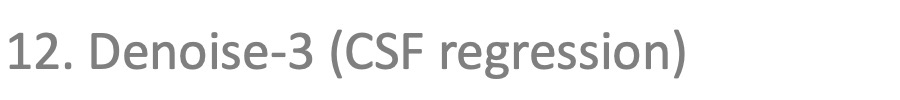

### step_12_green.jpeg

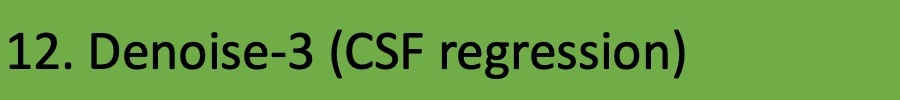

### step_12_white.jpeg

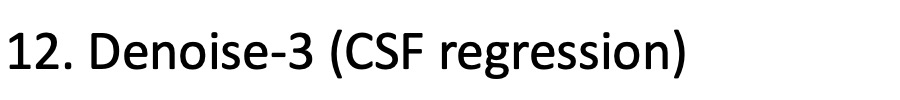

### step_13_blue.jpeg

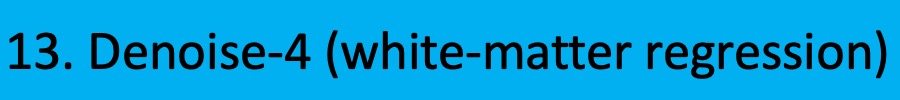

### step_13_gray.jpeg

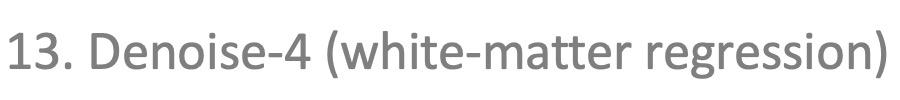

### step_13_green.jpeg

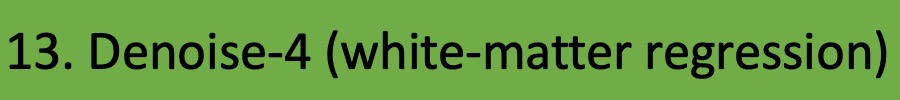

### step_13_white.jpeg

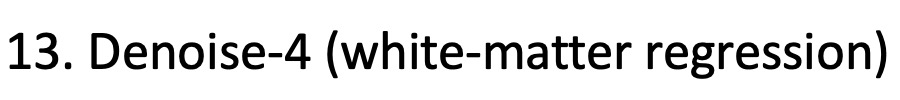

### step_14_blue.jpeg

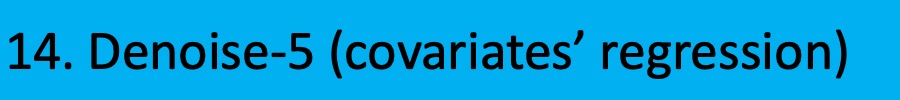

### step_14_gray.jpeg

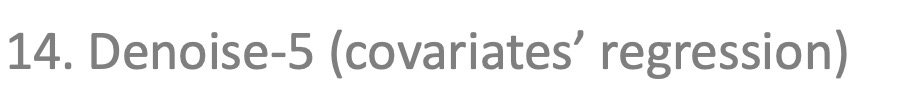

### step_14_green.jpeg

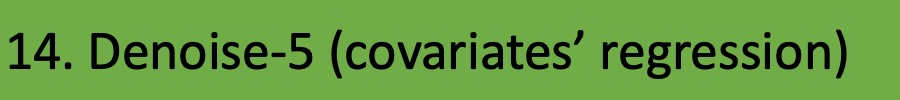

### step_14_white.jpeg

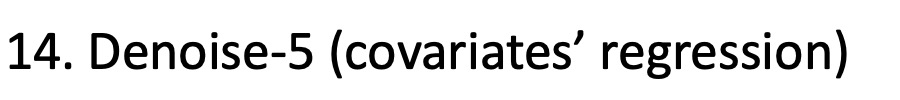

### step_15_blue.jpeg

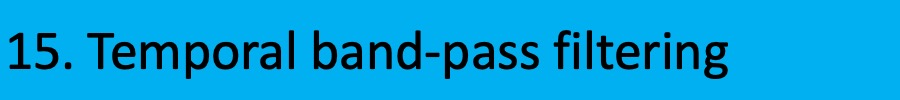

### step_15_gray.jpeg

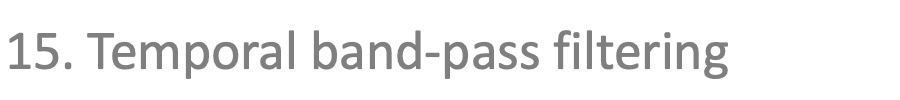

### step_15_green.jpeg

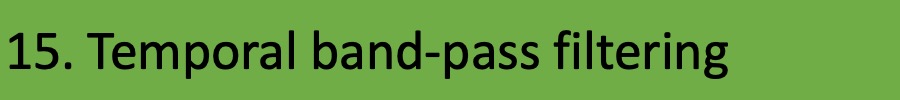

### step_15_white.jpeg

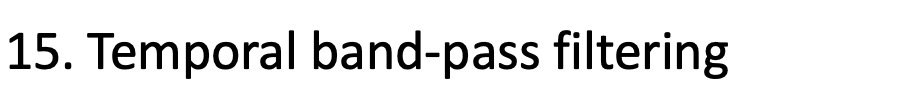

### step_16_blue.jpeg

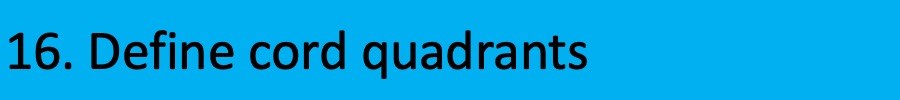

### step_16_gray.jpeg

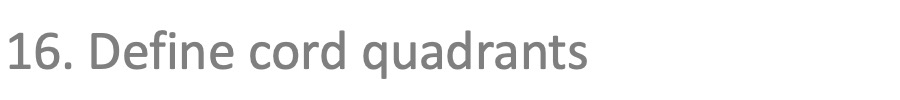

### step_16_green.jpeg

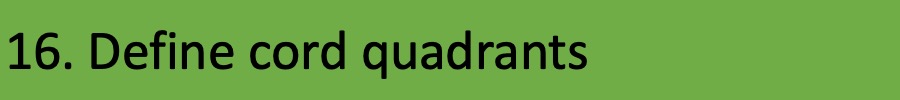

### step_16_white.jpeg

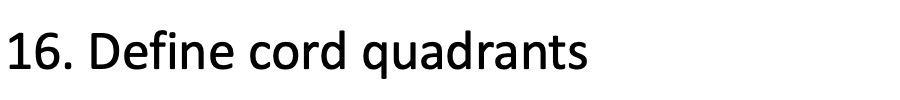

### step_17_blue.jpeg

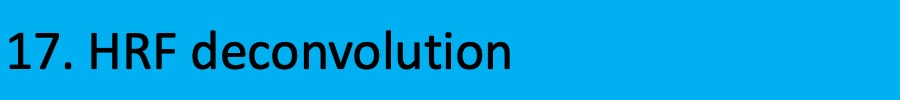

### step_17_gray.jpeg

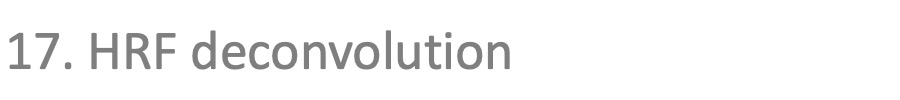

### step_17_green.jpeg

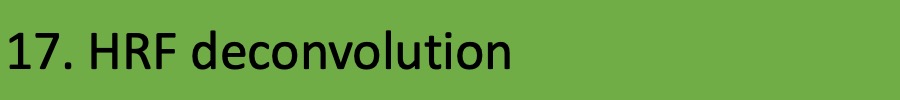

### step_17_white.jpeg

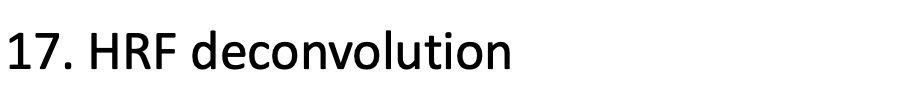

### step_18_blue.jpeg

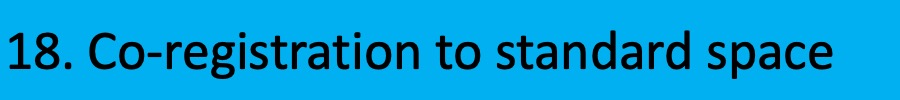

### step_18_gray.jpeg

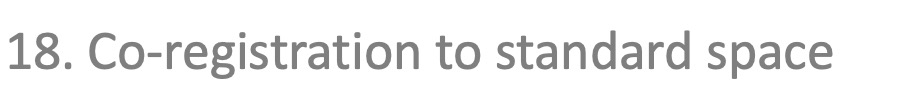

### step_18_green.jpeg

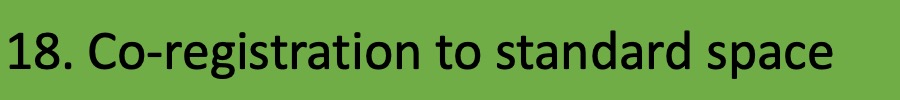

### step_18_white.jpeg

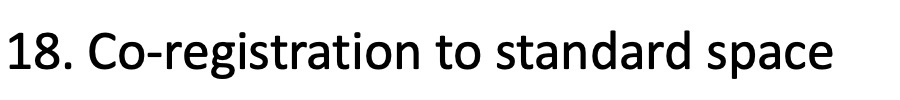

### step_19_blue.jpeg

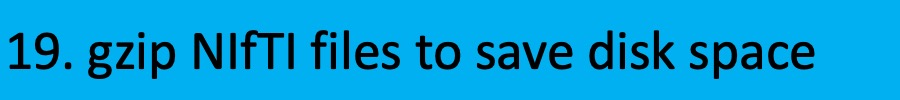

### step_19_gray.jpeg

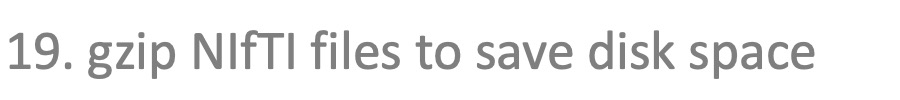

### step_19_green.jpeg

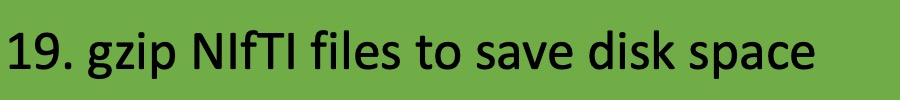

### step_19_white.jpeg

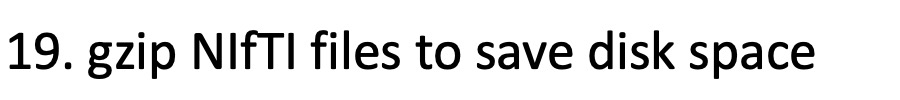

### white.jpg

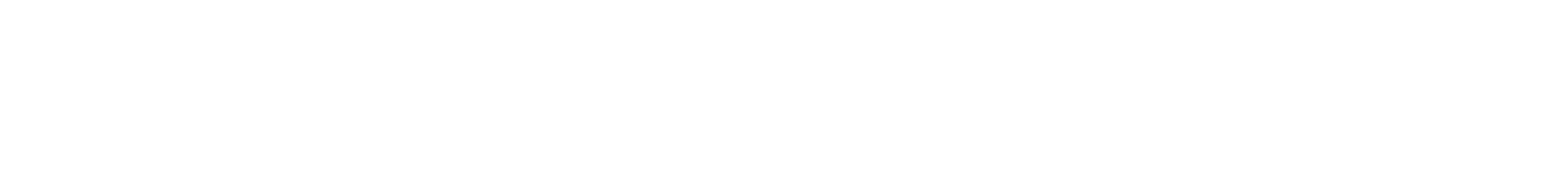
